## Supplemental Table 5 for "SETDB1 Fuels the Lung Cancer Phenotype by Modulating Epigenome, 3D Genome Organization and Chromatin Mechanical Properties"

**Supplementary Table 5, Annotation of compartment switches, Related to Figure 5B, C**

| chrom | start | end | Control_PC1 | SETDB1_LOF_PC1 | dynamics |
|---|---|---|---|---|---|
| chr1 | 0 | 100000 | 0.00 | 0.00 | NO |
| chr1 | 100000 | 200000 | 0.00 | 0.00 | NO |
| chr1 | 200000 | 300000 | 0.00 | 0.00 | NO |
| chr1 | 300000 | 400000 | 0.00 | 0.00 | NO |
| chr1 | 400000 | 500000 | 0.00 | 0.00 | NO |
| chr1 | 500000 | 600000 | 0.00 | 0.00 | NO |
| chr1 | 600000 | 700000 | 0.00 | 0.00 | NO |
| chr1 | 700000 | 800000 | 0.00 | 0.00 | NO |
| chr1 | 800000 | 900000 | 0.00 | 0.00 | NO |
| chr1 | 900000 | 1000000 | 1.34 | 1.08 | A>A |
| chr1 | 1000000 | 1100000 | 0.00 | 1.34 | NO |
| chr1 | 1100000 | 1200000 | 1.45 | 1.31 | A>A |
| chr1 | 1200000 | 1300000 | 1.52 | 1.11 | A>A |
| chr1 | 1300000 | 1400000 | 0.00 | 0.00 | NO |
| chr1 | 1400000 | 1500000 | 0.00 | 0.00 | NO |
| chr1 | 1500000 | 1600000 | 1.51 | 1.29 | A>A |
| chr1 | 1600000 | 1700000 | 0.00 | 0.00 | NO |
| chr1 | 1700000 | 1800000 | 1.48 | 1.56 | A>A |
| chr1 | 1800000 | 1900000 | 1.37 | 1.30 | A>A |
| chr1 | 1900000 | 2000000 | 1.43 | 1.40 | A>A |
| chr1 | 2000000 | 2100000 | 1.26 | 1.14 | A>A |
| chr1 | 2100000 | 2200000 | 1.24 | 1.11 | A>A |
| chr1 | 2200000 | 2300000 | 1.14 | 1.08 | A>A |
| chr1 | 2300000 | 2400000 | 1.08 | 0.91 | A>A |
| chr1 | 2400000 | 2500000 | 1.11 | 1.15 | A>A |
| chr1 | 2500000 | 2600000 | 0.77 | 0.94 | A>A |
| chr1 | 2600000 | 2700000 | 0.00 | 0.00 | NO |
| chr1 | 2700000 | 2800000 | 0.56 | 0.53 | A>A |
| chr1 | 2800000 | 2900000 | 0.63 | 0.57 | A>A |
| chr1 | 2900000 | 3000000 | 0.56 | 0.51 | A>A |
| chr1 | 3000000 | 3100000 | 0.71 | 0.73 | A>A |
| chr1 | 3100000 | 3200000 | 0.70 | 0.63 | A>A |
| chr1 | 3200000 | 3300000 | 0.70 | 0.60 | A>A |
| chr1 | 3300000 | 3400000 | 0.59 | 0.59 | A>A |
| chr1 | 3400000 | 3500000 | 1.13 | 1.02 | A>A |
| chr1 | 3500000 | 3600000 | 1.07 | 1.08 | A>A |
| chr1 | 3600000 | 3700000 | 1.15 | 1.02 | A>A |
| chr1 | 3700000 | 3800000 | 0.78 | 0.82 | A>A |
| chr1 | 3800000 | 3900000 | 0.00 | 0.14 | NO |
| chr1 | 3900000 | 4000000 | 0.00 | 0.00 | NO |
| chr1 | 4000000 | 4100000 | 0.18 | -0.06 | A>A |
| chr1 | 4100000 | 4200000 | 0.10 | -0.06 | A>A |
| chr1 | 4200000 | 4300000 | -0.06 | -0.11 | B>B |
| chr1 | 4300000 | 4400000 | -0.12 | -0.11 | B>B |
| chr1 | 4400000 | 4500000 | -0.05 | -0.04 | B>B |

| | | | | | |
|---|---|---|---|---|---|
| chr1 | 4500000 | 4600000 | -0.08 | -0.13 | B>B |
| chr1 | 4600000 | 4700000 | -0.03 | -0.08 | B>B |
| chr1 | 4700000 | 4800000 | -0.08 | -0.11 | B>B |
| chr1 | 4800000 | 4900000 | -0.36 | -0.38 | B>B |
| chr1 | 4900000 | 5000000 | -0.55 | -0.58 | B>B |
| chr1 | 5000000 | 5100000 | -0.66 | -0.64 | B>B |
| chr1 | 5100000 | 5200000 | -0.67 | -0.70 | B>B |
| chr1 | 5200000 | 5300000 | -0.72 | -0.72 | B>B |
| chr1 | 5300000 | 5400000 | -0.68 | -0.65 | B>B |
| chr1 | 5400000 | 5500000 | -0.61 | -0.49 | B>B |
| chr1 | 5500000 | 5600000 | -0.40 | -0.37 | B>B |
| chr1 | 5600000 | 5700000 | -0.45 | -0.42 | B>B |
| chr1 | 5700000 | 5800000 | -0.25 | -0.30 | B>B |
| chr1 | 5800000 | 5900000 | -0.01 | -0.16 | B>B |
| chr1 | 5900000 | 6000000 | 0.18 | 0.13 | A>A |
| chr1 | 6000000 | 6100000 | 0.58 | 0.63 | A>A |
| chr1 | 6100000 | 6200000 | 0.95 | 0.88 | A>A |
| chr1 | 6200000 | 6300000 | 1.05 | 1.13 | A>A |
| chr1 | 6300000 | 6400000 | 1.21 | 1.09 | A>A |
| chr1 | 6400000 | 6500000 | 1.17 | 1.19 | A>A |
| chr1 | 6500000 | 6600000 | 1.22 | 1.11 | A>A |
| chr1 | 6600000 | 6700000 | 1.24 | 1.17 | A>A |
| chr1 | 6700000 | 6800000 | 0.88 | 0.93 | A>A |
| chr1 | 6800000 | 6900000 | 0.89 | 0.95 | A>A |
| chr1 | 6900000 | 7000000 | 0.86 | 0.79 | A>A |
| chr1 | 7000000 | 7100000 | 0.59 | 0.68 | A>A |
| chr1 | 7100000 | 7200000 | 0.57 | 0.61 | A>A |
| chr1 | 7200000 | 7300000 | 0.65 | 0.62 | A>A |
| chr1 | 7300000 | 7400000 | 0.81 | 0.97 | A>A |
| chr1 | 7400000 | 7500000 | 0.82 | 0.84 | A>A |
| chr1 | 7500000 | 7600000 | 0.69 | 0.72 | A>A |
| chr1 | 7600000 | 7700000 | 0.54 | 0.66 | A>A |
| chr1 | 7700000 | 7800000 | 0.69 | 0.64 | A>A |
| chr1 | 7800000 | 7900000 | 0.83 | 0.84 | A>A |
| chr1 | 7900000 | 8000000 | 0.83 | 0.87 | A>A |
| chr1 | 8000000 | 8100000 | 0.68 | 0.76 | A>A |
| chr1 | 8100000 | 8200000 | 1.18 | 1.14 | A>A |
| chr1 | 8200000 | 8300000 | 1.20 | 1.23 | A>A |
| chr1 | 8300000 | 8400000 | 0.89 | 0.92 | A>A |
| chr1 | 8400000 | 8500000 | 0.77 | 0.85 | A>A |
| chr1 | 8500000 | 8600000 | 1.01 | 0.98 | A>A |
| chr1 | 8600000 | 8700000 | 0.91 | 0.94 | A>A |
| chr1 | 8700000 | 8800000 | 0.98 | 1.06 | A>A |
| chr1 | 8800000 | 8900000 | 1.04 | 0.91 | A>A |
| chr1 | 8900000 | 9000000 | 0.96 | 1.04 | A>A |
| chr1 | 9000000 | 9100000 | 1.00 | 1.09 | A>A |
| chr1 | 9100000 | 9200000 | 1.01 | 1.07 | A>A |
| chr1 | 9200000 | 9300000 | 1.23 | 1.09 | A>A |
| chr1 | 9300000 | 9400000 | 0.89 | 0.99 | A>A |
| chr1 | 9400000 | 9500000 | 1.31 | 1.19 | A>A |
| chr1 | 9500000 | 9600000 | 0.82 | 0.95 | A>A |

| | | | | | |
|---|---|---|---|---|---|
| chr1 | 9600000 | 9700000 | 0.97 | 1.21 | A>A |
| chr1 | 9700000 | 9800000 | 1.08 | 1.15 | A>A |
| chr1 | 9800000 | 9900000 | 1.17 | 1.23 | A>A |
| chr1 | 9900000 | 10000000 | 0.92 | 1.04 | A>A |
| chr1 | 10000000 | 10100000 | 1.15 | 1.09 | A>A |
| chr1 | 10100000 | 10200000 | 1.13 | 1.22 | A>A |
| chr1 | 10200000 | 10300000 | 1.05 | 1.06 | A>A |
| chr1 | 10300000 | 10400000 | 1.14 | 1.12 | A>A |
| chr1 | 10400000 | 10500000 | 1.15 | 1.23 | A>A |
| chr1 | 10500000 | 10600000 | 0.75 | 0.94 | A>A |
| chr1 | 10600000 | 10700000 | 0.99 | 1.00 | A>A |
| chr1 | 10700000 | 10800000 | 0.99 | 1.11 | A>A |
| chr1 | 10800000 | 10900000 | 1.04 | 1.13 | A>A |
| chr1 | 10900000 | 11000000 | 0.79 | 0.88 | A>A |
| chr1 | 11000000 | 11100000 | 0.89 | 0.90 | A>A |
| chr1 | 11100000 | 11200000 | 1.10 | 1.09 | A>A |
| chr1 | 11200000 | 11300000 | 1.07 | 1.07 | A>A |
| chr1 | 11300000 | 11400000 | 0.82 | 0.77 | A>A |
| chr1 | 11400000 | 11500000 | 0.84 | 0.82 | A>A |
| chr1 | 11500000 | 11600000 | 0.92 | 0.78 | A>A |
| chr1 | 11600000 | 11700000 | 0.77 | 0.77 | A>A |
| chr1 | 11700000 | 11800000 | 0.97 | 0.91 | A>A |
| chr1 | 11800000 | 11900000 | 1.14 | 1.12 | A>A |
| chr1 | 11900000 | 12000000 | 1.16 | 1.20 | A>A |
| chr1 | 12000000 | 12100000 | 1.12 | 1.07 | A>A |
| chr1 | 12100000 | 12200000 | 0.79 | 0.71 | A>A |
| chr1 | 12200000 | 12300000 | 0.68 | 0.50 | A>A |
| chr1 | 12300000 | 12400000 | 0.40 | 0.11 | A>A |
| chr1 | 12400000 | 12500000 | 0.62 | 0.41 | A>A |
| chr1 | 12500000 | 12600000 | 0.69 | 0.51 | A>A |
| chr1 | 12600000 | 12700000 | 0.45 | 0.28 | A>A |
| chr1 | 12700000 | 12800000 | 0.17 | -0.31 | B>B |
| chr1 | 12800000 | 12900000 | 0.00 | -0.61 | NO |
| chr1 | 12900000 | 13000000 | 0.00 | 0.00 | NO |
| chr1 | 13000000 | 13100000 | 0.00 | 0.00 | NO |
| chr1 | 13100000 | 13200000 | 0.00 | 0.00 | NO |
| chr1 | 13200000 | 13300000 | 0.00 | 0.00 | NO |
| chr1 | 13300000 | 13400000 | 0.00 | 0.00 | NO |
| chr1 | 13400000 | 13500000 | 0.00 | 0.00 | NO |
| chr1 | 13500000 | 13600000 | 0.00 | 0.00 | NO |
| chr1 | 13600000 | 13700000 | 0.00 | 0.00 | NO |
| chr1 | 13700000 | 13800000 | 0.00 | 0.00 | NO |
| chr1 | 13800000 | 13900000 | -0.23 | -0.97 | A>B |
| chr1 | 13900000 | 14000000 | -0.28 | -0.91 | A>B |
| chr1 | 14000000 | 14100000 | 0.04 | -0.35 | B>B |
| chr1 | 14100000 | 14200000 | -0.37 | -0.67 | B>B |
| chr1 | 14200000 | 14300000 | -0.54 | -0.89 | B>B |
| chr1 | 14300000 | 14400000 | -0.68 | -0.99 | B>B |
| chr1 | 14400000 | 14500000 | -0.68 | -1.01 | B>B |
| chr1 | 14500000 | 14600000 | -0.75 | -1.02 | B>B |
| chr1 | 14600000 | 14700000 | -0.67 | -0.94 | B>B |
| chr1 | 14700000 | 14800000 | -0.53 | -0.86 | B>B |
| chr1 | 14800000 | 14900000 | -0.35 | -0.81 | B>B |
| chr1 | 14900000 | 15000000 | -0.16 | -0.69 | B>B |
| chr1 | 15000000 | 15100000 | 0.03 | -0.44 | B>B |
| chr1 | 15100000 | 15200000 | 0.00 | -0.42 | B>B |

| | | | | | |
|---|---|---|---|---|---|
| chr1 | 15200000 | 15300000 | 0.11 | -0.25 | B>B |
| chr1 | 15300000 | 15400000 | 0.04 | -0.13 | B>B |
| chr1 | 15400000 | 15500000 | 0.35 | 0.32 | A>A |
| chr1 | 15500000 | 15600000 | 0.61 | 0.59 | A>A |
| chr1 | 15600000 | 15700000 | 0.69 | 0.75 | A>A |
| chr1 | 15700000 | 15800000 | 0.85 | 0.99 | A>A |
| chr1 | 15800000 | 15900000 | 1.04 | 1.11 | A>A |
| chr1 | 15900000 | 16000000 | 1.20 | 1.41 | A>A |
| chr1 | 16000000 | 16100000 | 1.16 | 1.33 | A>A |
| chr1 | 16100000 | 16200000 | 1.31 | 1.37 | A>A |
| chr1 | 16200000 | 16300000 | 1.33 | 1.50 | A>A |
| chr1 | 16300000 | 16400000 | 1.02 | 1.27 | A>A |
| chr1 | 16400000 | 16500000 | 1.44 | 1.53 | A>A |
| chr1 | 16500000 | 16600000 | 1.30 | 1.38 | A>A |
| chr1 | 16600000 | 16700000 | 1.31 | 1.42 | A>A |
| chr1 | 16700000 | 16800000 | 1.34 | 1.51 | A>A |
| chr1 | 16800000 | 16900000 | 0.00 | 0.00 | NO |
| chr1 | 16900000 | 17000000 | 1.49 | 1.44 | A>A |
| chr1 | 17000000 | 17100000 | 0.00 | 0.00 | NO |
| chr1 | 17100000 | 17200000 | 0.00 | 0.00 | NO |
| chr1 | 17200000 | 17300000 | 0.00 | 0.00 | NO |
| chr1 | 17300000 | 17400000 | 1.01 | 1.02 | A>A |
| chr1 | 17400000 | 17500000 | 1.09 | 1.08 | A>A |
| chr1 | 17500000 | 17600000 | 1.31 | 1.27 | A>A |
| chr1 | 17600000 | 17700000 | 1.00 | 0.96 | A>A |
| chr1 | 17700000 | 17800000 | 0.69 | 0.78 | A>A |
| chr1 | 17800000 | 17900000 | 0.88 | 0.87 | A>A |
| chr1 | 17900000 | 18000000 | 0.65 | 0.74 | A>A |
| chr1 | 18000000 | 18100000 | 0.57 | 0.57 | A>A |
| chr1 | 18100000 | 18200000 | 0.50 | 0.51 | A>A |
| chr1 | 18200000 | 18300000 | 0.40 | 0.42 | A>A |
| chr1 | 18300000 | 18400000 | 0.36 | 0.39 | A>A |
| chr1 | 18400000 | 18500000 | 0.36 | 0.37 | A>A |
| chr1 | 18500000 | 18600000 | 0.34 | 0.32 | A>A |
| chr1 | 18600000 | 18700000 | 0.35 | 0.42 | A>A |
| chr1 | 18700000 | 18800000 | 0.50 | 0.64 | A>A |
| chr1 | 18800000 | 18900000 | 0.35 | 0.47 | A>A |
| chr1 | 18900000 | 19000000 | 0.54 | 0.57 | A>A |
| chr1 | 19000000 | 19100000 | 0.79 | 0.60 | A>A |
| chr1 | 19100000 | 19200000 | 0.45 | 0.44 | A>A |
| chr1 | 19200000 | 19300000 | 0.68 | 0.71 | A>A |
| chr1 | 19300000 | 19400000 | 1.12 | 1.01 | A>A |
| chr1 | 19400000 | 19500000 | 0.73 | 0.82 | A>A |
| chr1 | 19500000 | 19600000 | 0.86 | 0.97 | A>A |
| chr1 | 19600000 | 19700000 | 0.80 | 0.75 | A>A |
| chr1 | 19700000 | 19800000 | 0.87 | 0.90 | A>A |
| chr1 | 19800000 | 19900000 | 0.93 | 0.90 | A>A |
| chr1 | 19900000 | 20000000 | 0.59 | 0.72 | A>A |
| chr1 | 20000000 | 20100000 | 0.68 | 0.71 | A>A |
| chr1 | 20100000 | 20200000 | 0.65 | 0.77 | A>A |
| chr1 | 20200000 | 20300000 | 0.43 | 0.53 | A>A |
| chr1 | 20300000 | 20400000 | 0.40 | 0.50 | A>A |
| chr1 | 20400000 | 20500000 | 0.49 | 0.49 | A>A |
| chr1 | 20500000 | 20600000 | 0.49 | 0.40 | A>A |
| chr1 | 20600000 | 20700000 | 0.68 | 0.68 | A>A |
| chr1 | 20700000 | 20800000 | 0.65 | 0.59 | A>A |

| | | | | | |
|---|---|---|---|---|---|
| chr1 | 20800000 | 20900000 | 0.62 | 0.64 | A>A |
| chr1 | 20900000 | 21000000 | 0.72 | 0.81 | A>A |
| chr1 | 21000000 | 21100000 | 0.70 | 0.81 | A>A |
| chr1 | 21100000 | 21200000 | 1.09 | 0.91 | A>A |
| chr1 | 21200000 | 21300000 | 0.75 | 0.75 | A>A |
| chr1 | 21300000 | 21400000 | 0.95 | 0.89 | A>A |
| chr1 | 21400000 | 21500000 | 0.79 | 0.80 | A>A |
| chr1 | 21500000 | 21600000 | 0.52 | 0.65 | A>A |
| chr1 | 21600000 | 21700000 | 1.01 | 0.87 | A>A |
| chr1 | 21700000 | 21800000 | 0.00 | 0.00 | NO |
| chr1 | 21800000 | 21900000 | 0.68 | 0.62 | A>A |
| chr1 | 21900000 | 22000000 | 0.82 | 0.85 | A>A |
| chr1 | 22000000 | 22100000 | 0.86 | 0.72 | A>A |
| chr1 | 22100000 | 22200000 | 1.02 | 1.01 | A>A |
| chr1 | 22200000 | 22300000 | 0.96 | 0.92 | A>A |
| chr1 | 22300000 | 22400000 | 0.00 | 0.00 | NO |
| chr1 | 22400000 | 22500000 | 0.59 | 0.61 | A>A |
| chr1 | 22500000 | 22600000 | 0.83 | 0.76 | A>A |
| chr1 | 22600000 | 22700000 | 0.70 | 0.79 | A>A |
| chr1 | 22700000 | 22800000 | 0.57 | 0.60 | A>A |
| chr1 | 22800000 | 22900000 | 0.74 | 0.79 | A>A |
| chr1 | 22900000 | 23000000 | 0.50 | 0.53 | A>A |
| chr1 | 23000000 | 23100000 | 0.71 | 0.83 | A>A |
| chr1 | 23100000 | 23200000 | 0.88 | 0.87 | A>A |
| chr1 | 23200000 | 23300000 | 0.49 | 0.52 | A>A |
| chr1 | 23300000 | 23400000 | 0.71 | 0.77 | A>A |
| chr1 | 23400000 | 23500000 | 1.00 | 0.89 | A>A |
| chr1 | 23500000 | 23600000 | 0.86 | 0.94 | A>A |
| chr1 | 23600000 | 23700000 | 0.93 | 0.93 | A>A |
| chr1 | 23700000 | 23800000 | 0.90 | 0.90 | A>A |
| chr1 | 23800000 | 23900000 | 0.84 | 0.88 | A>A |
| chr1 | 23900000 | 24000000 | 0.93 | 0.92 | A>A |
| chr1 | 24000000 | 24100000 | 0.93 | 1.05 | A>A |
| chr1 | 24100000 | 24200000 | 1.04 | 1.07 | A>A |
| chr1 | 24200000 | 24300000 | 0.00 | 0.00 | NO |
| chr1 | 24300000 | 24400000 | 0.00 | 0.00 | NO |
| chr1 | 24400000 | 24500000 | 0.80 | 0.87 | A>A |
| chr1 | 24500000 | 24600000 | 0.89 | 0.86 | A>A |
| chr1 | 24600000 | 24700000 | 1.14 | 1.12 | A>A |
| chr1 | 24700000 | 24800000 | 0.72 | 0.78 | A>A |
| chr1 | 24800000 | 24900000 | 0.90 | 0.94 | A>A |
| chr1 | 24900000 | 25000000 | 0.66 | 0.71 | A>A |
| chr1 | 25000000 | 25100000 | 0.84 | 0.87 | A>A |
| chr1 | 25100000 | 25200000 | 0.68 | 0.70 | A>A |
| chr1 | 25200000 | 25300000 | 0.83 | 0.90 | A>A |
| chr1 | 25300000 | 25400000 | 0.80 | 0.97 | A>A |
| chr1 | 25400000 | 25500000 | 0.80 | 0.86 | A>A |
| chr1 | 25500000 | 25600000 | 0.66 | 0.75 | A>A |
| chr1 | 25600000 | 25700000 | 0.00 | 0.00 | NO |
| chr1 | 25700000 | 25800000 | 0.00 | 0.00 | NO |
| chr1 | 25800000 | 25900000 | 0.77 | 0.86 | A>A |
| chr1 | 25900000 | 26000000 | 0.93 | 0.93 | A>A |
| chr1 | 26000000 | 26100000 | 0.96 | 0.95 | A>A |
| chr1 | 26100000 | 26200000 | 0.72 | 0.80 | A>A |
| chr1 | 26200000 | 26300000 | 0.96 | 1.04 | A>A |
| chr1 | 26300000 | 26400000 | 0.78 | 0.89 | A>A |

| | | | | | |
|---|---|---|---|---|---|
| chr1 | 26400000 | 26500000 | 0.89 | 0.98 | A>A |
| chr1 | 26500000 | 26600000 | 0.96 | 0.97 | A>A |
| chr1 | 26600000 | 26700000 | 1.08 | 1.03 | A>A |
| chr1 | 26700000 | 26800000 | 0.91 | 0.95 | A>A |
| chr1 | 26800000 | 26900000 | 1.13 | 1.15 | A>A |
| chr1 | 26900000 | 27000000 | 0.97 | 1.00 | A>A |
| chr1 | 27000000 | 27100000 | 1.18 | 1.25 | A>A |
| chr1 | 27100000 | 27200000 | 0.90 | 1.04 | A>A |
| chr1 | 27200000 | 27300000 | 1.15 | 1.02 | A>A |
| chr1 | 27300000 | 27400000 | 0.93 | 1.07 | A>A |
| chr1 | 27400000 | 27500000 | 0.90 | 1.00 | A>A |
| chr1 | 27500000 | 27600000 | 1.05 | 1.01 | A>A |
| chr1 | 27600000 | 27700000 | 0.94 | 0.92 | A>A |
| chr1 | 27700000 | 27800000 | 0.97 | 1.05 | A>A |
| chr1 | 27800000 | 27900000 | 1.25 | 1.22 | A>A |
| chr1 | 27900000 | 28000000 | 0.84 | 0.90 | A>A |
| chr1 | 28000000 | 28100000 | 1.02 | 1.03 | A>A |
| chr1 | 28100000 | 28200000 | 0.89 | 1.11 | A>A |
| chr1 | 28200000 | 28300000 | 0.99 | 1.15 | A>A |
| chr1 | 28300000 | 28400000 | 1.07 | 1.06 | A>A |
| chr1 | 28400000 | 28500000 | 1.06 | 1.13 | A>A |
| chr1 | 28500000 | 28600000 | 1.01 | 1.05 | A>A |
| chr1 | 28600000 | 28700000 | 1.11 | 1.18 | A>A |
| chr1 | 28700000 | 28800000 | 1.11 | 1.06 | A>A |
| chr1 | 28800000 | 28900000 | 1.01 | 1.16 | A>A |
| chr1 | 28900000 | 29000000 | 0.90 | 1.09 | A>A |
| chr1 | 29000000 | 29100000 | 0.95 | 0.95 | A>A |
| chr1 | 29100000 | 29200000 | 0.83 | 0.88 | A>A |
| chr1 | 29200000 | 29300000 | 0.86 | 0.84 | A>A |
| chr1 | 29300000 | 29400000 | 0.80 | 0.74 | A>A |
| chr1 | 29400000 | 29500000 | 0.79 | 0.74 | A>A |
| chr1 | 29500000 | 29600000 | 0.55 | 0.48 | A>A |
| chr1 | 29600000 | 29700000 | 0.54 | 0.45 | A>A |
| chr1 | 29700000 | 29800000 | 0.28 | 0.10 | A>A |
| chr1 | 29800000 | 29900000 | 0.06 | 0.07 | A>A |
| chr1 | 29900000 | 30000000 | 0.00 | 0.00 | NO |
| chr1 | 30000000 | 30100000 | 0.22 | 0.09 | A>A |
| chr1 | 30100000 | 30200000 | 0.28 | 0.08 | A>A |
| chr1 | 30200000 | 30300000 | 0.23 | 0.07 | A>A |
| chr1 | 30300000 | 30400000 | 0.23 | 0.09 | A>A |
| chr1 | 30400000 | 30500000 | 0.35 | 0.13 | A>A |
| chr1 | 30500000 | 30600000 | 0.22 | 0.09 | A>A |
| chr1 | 30600000 | 30700000 | 0.11 | 0.03 | A>A |
| chr1 | 30700000 | 30800000 | 0.14 | -0.01 | A>A |
| chr1 | 30800000 | 30900000 | 0.14 | 0.03 | A>A |
| chr1 | 30900000 | 31000000 | 0.16 | 0.07 | A>A |
| chr1 | 31000000 | 31100000 | 0.23 | 0.17 | A>A |
| chr1 | 31100000 | 31200000 | 0.23 | 0.32 | A>A |
| chr1 | 31200000 | 31300000 | 0.68 | 0.61 | A>A |
| chr1 | 31300000 | 31400000 | 0.78 | 0.79 | A>A |
| chr1 | 31400000 | 31500000 | 0.87 | 0.91 | A>A |
| chr1 | 31500000 | 31600000 | 0.98 | 1.04 | A>A |
| chr1 | 31600000 | 31700000 | 0.83 | 0.90 | A>A |
| chr1 | 31700000 | 31800000 | 1.04 | 1.02 | A>A |
| chr1 | 31800000 | 31900000 | 0.90 | 0.94 | A>A |
| chr1 | 31900000 | 32000000 | 0.98 | 1.13 | A>A |

| | | | | | |
|---|---|---|---|---|---|
| chr1 | 32000000 | 32100000 | 0.98 | 1.00 | A>A |
| chr1 | 32100000 | 32200000 | 1.15 | 1.24 | A>A |
| chr1 | 32200000 | 32300000 | 1.03 | 1.15 | A>A |
| chr1 | 32300000 | 32400000 | 1.40 | 1.35 | A>A |
| chr1 | 32400000 | 32500000 | 1.09 | 1.23 | A>A |
| chr1 | 32500000 | 32600000 | 1.16 | 1.28 | A>A |
| chr1 | 32600000 | 32700000 | 1.37 | 1.42 | A>A |
| chr1 | 32700000 | 32800000 | 1.36 | 1.42 | A>A |
| chr1 | 32800000 | 32900000 | 1.08 | 1.25 | A>A |
| chr1 | 32900000 | 33000000 | 0.99 | 1.20 | A>A |
| chr1 | 33000000 | 33100000 | 0.99 | 1.29 | A>A |
| chr1 | 33100000 | 33200000 | 1.18 | 1.17 | A>A |
| chr1 | 33200000 | 33300000 | 1.19 | 1.31 | A>A |
| chr1 | 33300000 | 33400000 | 0.91 | 1.03 | A>A |
| chr1 | 33400000 | 33500000 | 1.03 | 1.11 | A>A |
| chr1 | 33500000 | 33600000 | 0.73 | 0.71 | A>A |
| chr1 | 33600000 | 33700000 | 0.48 | 0.44 | A>A |
| chr1 | 33700000 | 33800000 | 0.53 | 0.37 | A>A |
| chr1 | 33800000 | 33900000 | 0.32 | 0.31 | A>A |
| chr1 | 33900000 | 34000000 | -0.01 | -0.23 | B>B |
| chr1 | 34000000 | 34100000 | -0.33 | -0.70 | B>B |
| chr1 | 34100000 | 34200000 | -0.53 | -0.91 | B>B |
| chr1 | 34200000 | 34300000 | -0.57 | -1.05 | B>B |
| chr1 | 34300000 | 34400000 | -0.59 | -1.06 | B>B |
| chr1 | 34400000 | 34500000 | -0.58 | -0.99 | B>B |
| chr1 | 34500000 | 34600000 | -0.57 | -0.99 | B>B |
| chr1 | 34600000 | 34700000 | -0.52 | -0.95 | B>B |
| chr1 | 34700000 | 34800000 | -0.62 | -1.01 | B>B |
| chr1 | 34800000 | 34900000 | -0.55 | -1.02 | B>B |
| chr1 | 34900000 | 35000000 | -0.61 | -0.98 | B>B |
| chr1 | 35000000 | 35100000 | -0.52 | -0.98 | B>B |
| chr1 | 35100000 | 35200000 | -0.31 | -0.72 | B>B |
| chr1 | 35200000 | 35300000 | 0.07 | -0.24 | B>B |
| chr1 | 35300000 | 35400000 | 0.36 | 0.09 | A>A |
| chr1 | 35400000 | 35500000 | 0.42 | 0.38 | A>A |
| chr1 | 35500000 | 35600000 | 0.71 | 0.55 | A>A |
| chr1 | 35600000 | 35700000 | 0.77 | 0.76 | A>A |
| chr1 | 35700000 | 35800000 | 0.92 | 0.94 | A>A |
| chr1 | 35800000 | 35900000 | 0.82 | 0.85 | A>A |
| chr1 | 35900000 | 36000000 | 0.87 | 0.94 | A>A |
| chr1 | 36000000 | 36100000 | 0.94 | 1.00 | A>A |
| chr1 | 36100000 | 36200000 | 1.10 | 1.07 | A>A |
| chr1 | 36200000 | 36300000 | 0.92 | 0.93 | A>A |
| chr1 | 36300000 | 36400000 | 0.97 | 1.04 | A>A |
| chr1 | 36400000 | 36500000 | 1.11 | 0.97 | A>A |
| chr1 | 36500000 | 36600000 | 0.81 | 0.85 | A>A |
| chr1 | 36600000 | 36700000 | 1.06 | 1.00 | A>A |
| chr1 | 36700000 | 36800000 | 1.08 | 0.93 | A>A |
| chr1 | 36800000 | 36900000 | 0.83 | 0.72 | A>A |
| chr1 | 36900000 | 37000000 | 0.64 | 0.18 | A>A |
| chr1 | 37000000 | 37100000 | 0.17 | -0.22 | B>B |
| chr1 | 37100000 | 37200000 | -0.03 | -0.38 | B>B |
| chr1 | 37200000 | 37300000 | 0.06 | -0.42 | B>B |
| chr1 | 37300000 | 37400000 | -0.08 | -0.51 | B>B |
| chr1 | 37400000 | 37500000 | -0.05 | -0.51 | B>B |
| chr1 | 37500000 | 37600000 | -0.06 | -0.58 | B>B |

| | | | | | |
|---|---|---|---|---|---|
| chr1 | 37600000 | 37700000 | -0.16 | -0.52 | B>B |
| chr1 | 37700000 | 37800000 | -0.10 | -0.47 | B>B |
| chr1 | 37800000 | 37900000 | 0.06 | -0.32 | B>B |
| chr1 | 37900000 | 38000000 | 0.54 | 0.23 | A>A |
| chr1 | 38000000 | 38100000 | 0.77 | 0.43 | A>A |
| chr1 | 38100000 | 38200000 | 0.78 | 0.39 | A>A |
| chr1 | 38200000 | 38300000 | 0.79 | 0.57 | A>A |
| chr1 | 38300000 | 38400000 | 0.74 | 0.66 | A>A |
| chr1 | 38400000 | 38500000 | 0.60 | 0.34 | A>A |
| chr1 | 38500000 | 38600000 | 0.08 | -0.37 | B>B |
| chr1 | 38600000 | 38700000 | -0.15 | -0.58 | B>B |
| chr1 | 38700000 | 38800000 | -0.15 | -0.62 | B>B |
| chr1 | 38800000 | 38900000 | -0.32 | -0.71 | B>B |
| chr1 | 38900000 | 39000000 | -0.25 | -0.65 | B>B |
| chr1 | 39000000 | 39100000 | -0.10 | -0.57 | B>B |
| chr1 | 39100000 | 39200000 | -0.19 | -0.69 | B>B |
| chr1 | 39200000 | 39300000 | -0.07 | -0.39 | B>B |
| chr1 | 39300000 | 39400000 | 0.48 | 0.32 | A>A |
| chr1 | 39400000 | 39500000 | 0.70 | 0.67 | A>A |
| chr1 | 39500000 | 39600000 | 0.64 | 0.59 | A>A |
| chr1 | 39600000 | 39700000 | 0.88 | 1.04 | A>A |
| chr1 | 39700000 | 39800000 | 0.89 | 0.87 | A>A |
| chr1 | 39800000 | 39900000 | 0.88 | 1.00 | A>A |
| chr1 | 39900000 | 40000000 | 0.99 | 0.88 | A>A |
| chr1 | 40000000 | 40100000 | 1.02 | 1.01 | A>A |
| chr1 | 40100000 | 40200000 | 0.98 | 1.09 | A>A |
| chr1 | 40200000 | 40300000 | 1.05 | 0.97 | A>A |
| chr1 | 40300000 | 40400000 | 1.17 | 1.25 | A>A |
| chr1 | 40400000 | 40500000 | 1.12 | 1.15 | A>A |
| chr1 | 40500000 | 40600000 | 0.95 | 1.23 | A>A |
| chr1 | 40600000 | 40700000 | 0.93 | 1.29 | A>A |
| chr1 | 40700000 | 40800000 | 0.89 | 1.27 | A>A |
| chr1 | 40800000 | 40900000 | 0.79 | 1.08 | A>A |
| chr1 | 40900000 | 41000000 | 0.77 | 1.26 | A>A |
| chr1 | 41000000 | 41100000 | 0.90 | 1.06 | A>A |
| chr1 | 41100000 | 41200000 | 1.17 | 1.31 | A>A |
| chr1 | 41200000 | 41300000 | 1.07 | 1.12 | A>A |
| chr1 | 41300000 | 41400000 | 1.28 | 1.42 | A>A |
| chr1 | 41400000 | 41500000 | 1.09 | 1.20 | A>A |
| chr1 | 41500000 | 41600000 | 0.86 | 0.98 | A>A |
| chr1 | 41600000 | 41700000 | 1.01 | 1.06 | A>A |
| chr1 | 41700000 | 41800000 | 0.86 | 0.92 | A>A |
| chr1 | 41800000 | 41900000 | 0.98 | 1.23 | A>A |
| chr1 | 41900000 | 42000000 | 0.70 | 0.88 | A>A |
| chr1 | 42000000 | 42100000 | 0.60 | 0.76 | A>A |
| chr1 | 42100000 | 42200000 | 0.79 | 0.85 | A>A |
| chr1 | 42200000 | 42300000 | 0.61 | 0.67 | A>A |
| chr1 | 42300000 | 42400000 | 0.79 | 0.95 | A>A |
| chr1 | 42400000 | 42500000 | 0.83 | 0.82 | A>A |
| chr1 | 42500000 | 42600000 | 0.55 | 0.67 | A>A |
| chr1 | 42600000 | 42700000 | 0.64 | 0.78 | A>A |
| chr1 | 42700000 | 42800000 | 0.89 | 0.96 | A>A |
| chr1 | 42800000 | 42900000 | 0.94 | 0.99 | A>A |
| chr1 | 42900000 | 43000000 | 0.91 | 0.91 | A>A |
| chr1 | 43000000 | 43100000 | 0.80 | 0.78 | A>A |
| chr1 | 43100000 | 43200000 | 0.95 | 1.01 | A>A |

| | | | | | |
|---|---|---|---|---|---|
| chr1 | 43200000 | 43300000 | 0.89 | 0.90 | A>A |
| chr1 | 43300000 | 43400000 | 0.89 | 0.93 | A>A |
| chr1 | 43400000 | 43500000 | 1.36 | 1.36 | A>A |
| chr1 | 43500000 | 43600000 | 0.84 | 0.93 | A>A |
| chr1 | 43600000 | 43700000 | 0.95 | 0.94 | A>A |
| chr1 | 43700000 | 43800000 | 1.07 | 1.21 | A>A |
| chr1 | 43800000 | 43900000 | 1.06 | 1.16 | A>A |
| chr1 | 43900000 | 44000000 | 1.06 | 1.06 | A>A |
| chr1 | 44000000 | 44100000 | 1.21 | 1.20 | A>A |
| chr1 | 44100000 | 44200000 | 1.12 | 1.18 | A>A |
| chr1 | 44200000 | 44300000 | 1.08 | 1.12 | A>A |
| chr1 | 44300000 | 44400000 | 1.20 | 1.17 | A>A |
| chr1 | 44400000 | 44500000 | 1.03 | 1.10 | A>A |
| chr1 | 44500000 | 44600000 | 1.11 | 1.02 | A>A |
| chr1 | 44600000 | 44700000 | 0.99 | 1.08 | A>A |
| chr1 | 44700000 | 44800000 | 1.05 | 1.08 | A>A |
| chr1 | 44800000 | 44900000 | 1.15 | 1.00 | A>A |
| chr1 | 44900000 | 45000000 | 1.25 | 1.03 | A>A |
| chr1 | 45000000 | 45100000 | 1.03 | 1.05 | A>A |
| chr1 | 45100000 | 45200000 | 1.12 | 1.15 | A>A |
| chr1 | 45200000 | 45300000 | 1.42 | 1.34 | A>A |
| chr1 | 45300000 | 45400000 | 1.36 | 1.29 | A>A |
| chr1 | 45400000 | 45500000 | 0.98 | 1.18 | A>A |
| chr1 | 45500000 | 45600000 | 1.21 | 1.25 | A>A |
| chr1 | 45600000 | 45700000 | 1.18 | 1.32 | A>A |
| chr1 | 45700000 | 45800000 | 0.98 | 1.13 | A>A |
| chr1 | 45800000 | 45900000 | 1.00 | 1.15 | A>A |
| chr1 | 45900000 | 46000000 | 1.28 | 1.33 | A>A |
| chr1 | 46000000 | 46100000 | 1.07 | 1.39 | A>A |
| chr1 | 46100000 | 46200000 | 1.26 | 1.30 | A>A |
| chr1 | 46200000 | 46300000 | 1.24 | 1.36 | A>A |
| chr1 | 46300000 | 46400000 | 1.20 | 1.24 | A>A |
| chr1 | 46400000 | 46500000 | 1.15 | 1.21 | A>A |
| chr1 | 46500000 | 46600000 | 1.32 | 1.38 | A>A |
| chr1 | 46600000 | 46700000 | 1.20 | 1.21 | A>A |
| chr1 | 46700000 | 46800000 | 1.30 | 1.42 | A>A |
| chr1 | 46800000 | 46900000 | 1.15 | 1.09 | A>A |
| chr1 | 46900000 | 47000000 | 1.00 | 1.02 | A>A |
| chr1 | 47000000 | 47100000 | 0.84 | 0.66 | A>A |
| chr1 | 47100000 | 47200000 | 0.41 | 0.47 | A>A |
| chr1 | 47200000 | 47300000 | -0.23 | -0.31 | B>B |
| chr1 | 47300000 | 47400000 | 0.00 | 0.00 | NO |
| chr1 | 47400000 | 47500000 | -0.47 | -0.48 | B>B |
| chr1 | 47500000 | 47600000 | 0.00 | 0.00 | NO |
| chr1 | 47600000 | 47700000 | 0.22 | 0.07 | A>A |
| chr1 | 47700000 | 47800000 | 0.55 | 0.41 | A>A |
| chr1 | 47800000 | 47900000 | 0.41 | 0.25 | A>A |
| chr1 | 47900000 | 48000000 | 0.45 | 0.29 | A>A |
| chr1 | 48000000 | 48100000 | 0.50 | 0.38 | A>A |
| chr1 | 48100000 | 48200000 | 0.20 | 0.20 | A>A |
| chr1 | 48200000 | 48300000 | -0.02 | -0.23 | B>B |
| chr1 | 48300000 | 48400000 | -0.04 | -0.30 | B>B |
| chr1 | 48400000 | 48500000 | -0.15 | -0.48 | B>B |
| chr1 | 48500000 | 48600000 | -0.39 | -0.63 | B>B |
| chr1 | 48600000 | 48700000 | -0.56 | -0.72 | B>B |
| chr1 | 48700000 | 48800000 | -0.66 | -0.80 | B>B |

| | | | | | |
|---|---|---|---|---|---|
| chr1 | 48800000 | 48900000 | -0.63 | -0.73 | B>B |
| chr1 | 48900000 | 49000000 | -0.48 | -0.56 | B>B |
| chr1 | 49000000 | 49100000 | -0.46 | -0.76 | B>B |
| chr1 | 49100000 | 49200000 | -0.41 | -0.73 | B>B |
| chr1 | 49200000 | 49300000 | -0.40 | -0.76 | B>B |
| chr1 | 49300000 | 49400000 | -0.28 | -0.64 | B>B |
| chr1 | 49400000 | 49500000 | -0.38 | -0.72 | B>B |
| chr1 | 49500000 | 49600000 | -0.54 | -0.85 | B>B |
| chr1 | 49600000 | 49700000 | -0.62 | -0.88 | B>B |
| chr1 | 49700000 | 49800000 | -0.52 | -0.86 | B>B |
| chr1 | 49800000 | 49900000 | -0.66 | -0.92 | B>B |
| chr1 | 49900000 | 50000000 | -0.84 | -0.90 | B>B |
| chr1 | 50000000 | 50100000 | -0.81 | -0.89 | B>B |
| chr1 | 50100000 | 50200000 | -0.85 | -0.91 | B>B |
| chr1 | 50200000 | 50300000 | -0.86 | -0.95 | B>B |
| chr1 | 50300000 | 50400000 | -0.90 | -0.97 | B>B |
| chr1 | 50400000 | 50500000 | -0.78 | -0.93 | B>B |
| chr1 | 50500000 | 50600000 | -0.76 | -1.03 | B>B |
| chr1 | 50600000 | 50700000 | -0.77 | -0.96 | B>B |
| chr1 | 50700000 | 50800000 | -0.59 | -0.80 | B>B |
| chr1 | 50800000 | 50900000 | -0.32 | -0.37 | B>B |
| chr1 | 50900000 | 51000000 | -0.19 | -0.20 | B>B |
| chr1 | 51000000 | 51100000 | -0.13 | -0.05 | B>B |
| chr1 | 51100000 | 51200000 | 0.02 | 0.08 | A>A |
| chr1 | 51200000 | 51300000 | 0.08 | 0.27 | A>A |
| chr1 | 51300000 | 51400000 | 0.14 | 0.38 | A>A |
| chr1 | 51400000 | 51500000 | 0.34 | 0.52 | A>A |
| chr1 | 51500000 | 51600000 | 0.25 | 0.32 | A>A |
| chr1 | 51600000 | 51700000 | 0.31 | 0.40 | A>A |
| chr1 | 51700000 | 51800000 | 0.73 | 0.77 | A>A |
| chr1 | 51800000 | 51900000 | 0.81 | 0.94 | A>A |
| chr1 | 51900000 | 52000000 | 0.86 | 0.94 | A>A |
| chr1 | 52000000 | 52100000 | 0.96 | 1.10 | A>A |
| chr1 | 52100000 | 52200000 | 1.07 | 1.22 | A>A |
| chr1 | 52200000 | 52300000 | 0.93 | 1.09 | A>A |
| chr1 | 52300000 | 52400000 | 1.08 | 1.15 | A>A |
| chr1 | 52400000 | 52500000 | 1.11 | 1.32 | A>A |
| chr1 | 52500000 | 52600000 | 1.10 | 1.23 | A>A |
| chr1 | 52600000 | 52700000 | 1.34 | 1.36 | A>A |
| chr1 | 52700000 | 52800000 | 1.23 | 1.30 | A>A |
| chr1 | 52800000 | 52900000 | 1.17 | 1.32 | A>A |
| chr1 | 52900000 | 53000000 | 1.40 | 1.45 | A>A |
| chr1 | 53000000 | 53100000 | 1.41 | 1.38 | A>A |
| chr1 | 53100000 | 53200000 | 1.08 | 1.17 | A>A |
| chr1 | 53200000 | 53300000 | 1.30 | 1.22 | A>A |
| chr1 | 53300000 | 53400000 | 1.19 | 1.22 | A>A |
| chr1 | 53400000 | 53500000 | 1.34 | 1.26 | A>A |
| chr1 | 53500000 | 53600000 | 1.41 | 1.53 | A>A |
| chr1 | 53600000 | 53700000 | 1.10 | 1.28 | A>A |
| chr1 | 53700000 | 53800000 | 1.26 | 1.29 | A>A |
| chr1 | 53800000 | 53900000 | 1.24 | 1.20 | A>A |
| chr1 | 53900000 | 54000000 | 0.91 | 0.93 | A>A |
| chr1 | 54000000 | 54100000 | 1.14 | 1.14 | A>A |
| chr1 | 54100000 | 54200000 | 1.00 | 0.96 | A>A |
| chr1 | 54200000 | 54300000 | 1.09 | 0.98 | A>A |
| chr1 | 54300000 | 54400000 | 1.25 | 1.17 | A>A |

| | | | | | |
|---|---|---|---|---|---|
| chr1 | 54400000 | 54500000 | 1.24 | 1.15 | A>A |
| chr1 | 54500000 | 54600000 | 0.91 | 0.95 | A>A |
| chr1 | 54600000 | 54700000 | 1.01 | 1.02 | A>A |
| chr1 | 54700000 | 54800000 | 1.10 | 1.16 | A>A |
| chr1 | 54800000 | 54900000 | 1.02 | 1.00 | A>A |
| chr1 | 54900000 | 55000000 | 0.82 | 0.85 | A>A |
| chr1 | 55000000 | 55100000 | 1.01 | 0.97 | A>A |
| chr1 | 55100000 | 55200000 | 0.94 | 0.95 | A>A |
| chr1 | 55200000 | 55300000 | 0.87 | 0.87 | A>A |
| chr1 | 55300000 | 55400000 | 1.03 | 0.90 | A>A |
| chr1 | 55400000 | 55500000 | 0.79 | 0.79 | A>A |
| chr1 | 55500000 | 55600000 | 0.54 | 0.56 | A>A |
| chr1 | 55600000 | 55700000 | 0.55 | 0.33 | A>A |
| chr1 | 55700000 | 55800000 | 0.27 | 0.03 | A>A |
| chr1 | 55800000 | 55900000 | 0.37 | -0.13 | A>A |
| chr1 | 55900000 | 56000000 | 0.39 | -0.14 | A>A |
| chr1 | 56000000 | 56100000 | 0.46 | -0.32 | A>B |
| chr1 | 56100000 | 56200000 | 0.34 | -0.36 | A>B |
| chr1 | 56200000 | 56300000 | 0.24 | -0.45 | A>B |
| chr1 | 56300000 | 56400000 | 0.20 | -0.48 | A>B |
| chr1 | 56400000 | 56500000 | 0.17 | -0.60 | A>B |
| chr1 | 56500000 | 56600000 | 0.14 | -0.64 | A>B |
| chr1 | 56600000 | 56700000 | 0.05 | -0.56 | A>B |
| chr1 | 56700000 | 56800000 | 0.14 | -0.48 | A>B |
| chr1 | 56800000 | 56900000 | 0.15 | -0.32 | B>B |
| chr1 | 56900000 | 57000000 | 0.19 | -0.16 | A>A |
| chr1 | 57000000 | 57100000 | 0.10 | -0.22 | B>B |
| chr1 | 57100000 | 57200000 | 0.14 | -0.24 | B>B |
| chr1 | 57200000 | 57300000 | -0.23 | -0.56 | B>B |
| chr1 | 57300000 | 57400000 | -0.54 | -0.76 | B>B |
| chr1 | 57400000 | 57500000 | -0.61 | -0.88 | B>B |
| chr1 | 57500000 | 57600000 | -0.66 | -0.91 | B>B |
| chr1 | 57600000 | 57700000 | -0.68 | -0.90 | B>B |
| chr1 | 57700000 | 57800000 | -0.68 | -0.86 | B>B |
| chr1 | 57800000 | 57900000 | -0.66 | -0.91 | B>B |
| chr1 | 57900000 | 58000000 | -0.74 | -0.87 | B>B |
| chr1 | 58000000 | 58100000 | -0.76 | -0.89 | B>B |
| chr1 | 58100000 | 58200000 | -0.76 | -0.89 | B>B |
| chr1 | 58200000 | 58300000 | -0.80 | -0.88 | B>B |
| chr1 | 58300000 | 58400000 | -0.78 | -0.90 | B>B |
| chr1 | 58400000 | 58500000 | -0.73 | -0.86 | B>B |
| chr1 | 58500000 | 58600000 | -0.75 | -0.87 | B>B |
| chr1 | 58600000 | 58700000 | -0.75 | -0.86 | B>B |
| chr1 | 58700000 | 58800000 | -0.71 | -0.89 | B>B |
| chr1 | 58800000 | 58900000 | -0.65 | -0.87 | B>B |
| chr1 | 58900000 | 59000000 | -0.44 | -0.62 | B>B |
| chr1 | 59000000 | 59100000 | -0.21 | -0.29 | B>B |
| chr1 | 59100000 | 59200000 | -0.06 | -0.13 | B>B |
| chr1 | 59200000 | 59300000 | 0.08 | 0.14 | A>A |
| chr1 | 59300000 | 59400000 | 0.32 | 0.42 | A>A |
| chr1 | 59400000 | 59500000 | 0.26 | 0.39 | A>A |
| chr1 | 59500000 | 59600000 | 0.22 | 0.29 | A>A |
| chr1 | 59600000 | 59700000 | 0.10 | 0.28 | A>A |
| chr1 | 59700000 | 59800000 | 0.01 | 0.08 | A>A |
| chr1 | 59800000 | 59900000 | -0.11 | 0.02 | B>B |
| chr1 | 59900000 | 60000000 | -0.13 | 0.05 | B>B |

| | | | | | |
|---|---|---|---|---|---|
| chr1 | 60000000 | 60100000 | -0.32 | -0.03 | B>B |
| chr1 | 60100000 | 60200000 | -0.25 | -0.05 | B>B |
| chr1 | 60200000 | 60300000 | -0.44 | -0.24 | B>B |
| chr1 | 60300000 | 60400000 | -0.58 | -0.49 | B>B |
| chr1 | 60400000 | 60500000 | -0.66 | -0.70 | B>B |
| chr1 | 60500000 | 60600000 | -0.77 | -0.96 | B>B |
| chr1 | 60600000 | 60700000 | -0.80 | -1.02 | B>B |
| chr1 | 60700000 | 60800000 | -0.75 | -1.00 | B>B |
| chr1 | 60800000 | 60900000 | -0.87 | -1.02 | B>B |
| chr1 | 60900000 | 61000000 | -0.83 | -0.94 | B>B |
| chr1 | 61000000 | 61100000 | -0.83 | -1.01 | B>B |
| chr1 | 61100000 | 61200000 | -0.83 | -0.97 | B>B |
| chr1 | 61200000 | 61300000 | -0.71 | -0.90 | B>B |
| chr1 | 61300000 | 61400000 | -0.56 | -0.67 | B>B |
| chr1 | 61400000 | 61500000 | -0.23 | -0.41 | B>B |
| chr1 | 61500000 | 61600000 | 0.05 | 0.04 | A>A |
| chr1 | 61600000 | 61700000 | 0.10 | 0.29 | A>A |
| chr1 | 61700000 | 61800000 | 0.00 | 0.20 | A>A |
| chr1 | 61800000 | 61900000 | -0.06 | 0.03 | B>B |
| chr1 | 61900000 | 62000000 | 0.04 | 0.02 | A>A |
| chr1 | 62000000 | 62100000 | -0.08 | -0.01 | B>B |
| chr1 | 62100000 | 62200000 | 0.21 | 0.16 | A>A |
| chr1 | 62200000 | 62300000 | 0.37 | 0.26 | A>A |
| chr1 | 62300000 | 62400000 | 0.01 | 0.10 | A>A |
| chr1 | 62400000 | 62500000 | 0.03 | 0.28 | A>A |
| chr1 | 62500000 | 62600000 | 0.12 | 0.27 | A>A |
| chr1 | 62600000 | 62700000 | 0.34 | 0.38 | A>A |
| chr1 | 62700000 | 62800000 | 0.55 | 0.67 | A>A |
| chr1 | 62800000 | 62900000 | 0.39 | 0.49 | A>A |
| chr1 | 62900000 | 63000000 | 0.41 | 0.37 | A>A |
| chr1 | 63000000 | 63100000 | 0.62 | 0.48 | A>A |
| chr1 | 63100000 | 63200000 | 0.61 | 0.39 | A>A |
| chr1 | 63200000 | 63300000 | 0.15 | 0.04 | A>A |
| chr1 | 63300000 | 63400000 | 0.00 | -0.28 | B>B |
| chr1 | 63400000 | 63500000 | -0.12 | -0.48 | B>B |
| chr1 | 63500000 | 63600000 | -0.27 | -0.52 | B>B |
| chr1 | 63600000 | 63700000 | -0.19 | -0.52 | B>B |
| chr1 | 63700000 | 63800000 | 0.09 | -0.19 | B>B |
| chr1 | 63800000 | 63900000 | 0.30 | 0.06 | A>A |
| chr1 | 63900000 | 64000000 | 0.27 | 0.07 | A>A |
| chr1 | 64000000 | 64100000 | 0.38 | 0.36 | A>A |
| chr1 | 64100000 | 64200000 | 0.62 | 0.74 | A>A |
| chr1 | 64200000 | 64300000 | 0.87 | 0.90 | A>A |
| chr1 | 64300000 | 64400000 | 0.73 | 0.91 | A>A |
| chr1 | 64400000 | 64500000 | 0.64 | 0.84 | A>A |
| chr1 | 64500000 | 64600000 | 0.63 | 0.76 | A>A |
| chr1 | 64600000 | 64700000 | 0.35 | 0.54 | A>A |
| chr1 | 64700000 | 64800000 | 0.42 | 0.45 | A>A |
| chr1 | 64800000 | 64900000 | 0.71 | 0.55 | A>A |
| chr1 | 64900000 | 65000000 | 0.71 | 0.54 | A>A |
| chr1 | 65000000 | 65100000 | 0.70 | 0.56 | A>A |
| chr1 | 65100000 | 65200000 | 0.53 | 0.54 | A>A |
| chr1 | 65200000 | 65300000 | 0.63 | 0.60 | A>A |
| chr1 | 65300000 | 65400000 | 0.79 | 0.69 | A>A |
| chr1 | 65400000 | 65500000 | 0.76 | 0.72 | A>A |
| chr1 | 65500000 | 65600000 | 0.52 | 0.25 | A>A |

| | | | | | |
|---|---|---|---|---|---|
| chr1 | 65600000 | 65700000 | 0.35 | 0.11 | A>A |
| chr1 | 65700000 | 65800000 | 0.25 | 0.12 | A>A |
| chr1 | 65800000 | 65900000 | 0.31 | 0.16 | A>A |
| chr1 | 65900000 | 66000000 | 0.20 | -0.04 | A>A |
| chr1 | 66000000 | 66100000 | 0.00 | -0.08 | B>B |
| chr1 | 66100000 | 66200000 | -0.15 | -0.27 | B>B |
| chr1 | 66200000 | 66300000 | -0.22 | -0.39 | B>B |
| chr1 | 66300000 | 66400000 | -0.15 | -0.28 | B>B |
| chr1 | 66400000 | 66500000 | 0.07 | 0.12 | A>A |
| chr1 | 66500000 | 66600000 | 0.33 | 0.40 | A>A |
| chr1 | 66600000 | 66700000 | 0.34 | 0.45 | A>A |
| chr1 | 66700000 | 66800000 | 0.39 | 0.46 | A>A |
| chr1 | 66800000 | 66900000 | 0.42 | 0.24 | A>A |
| chr1 | 66900000 | 67000000 | 0.01 | -0.18 | B>B |
| chr1 | 67000000 | 67100000 | 0.07 | -0.45 | B>B |
| chr1 | 67100000 | 67200000 | -0.03 | -0.37 | B>B |
| chr1 | 67200000 | 67300000 | -0.06 | -0.26 | B>B |
| chr1 | 67300000 | 67400000 | 0.11 | -0.07 | A>A |
| chr1 | 67400000 | 67500000 | 0.22 | 0.27 | A>A |
| chr1 | 67500000 | 67600000 | 0.18 | 0.06 | A>A |
| chr1 | 67600000 | 67700000 | 0.06 | 0.02 | A>A |
| chr1 | 67700000 | 67800000 | 0.16 | 0.07 | A>A |
| chr1 | 67800000 | 67900000 | 0.55 | 0.42 | A>A |
| chr1 | 67900000 | 68000000 | 0.31 | 0.19 | A>A |
| chr1 | 68000000 | 68100000 | 0.48 | 0.49 | A>A |
| chr1 | 68100000 | 68200000 | 0.50 | 0.60 | A>A |
| chr1 | 68200000 | 68300000 | 0.29 | 0.40 | A>A |
| chr1 | 68300000 | 68400000 | 0.30 | 0.38 | A>A |
| chr1 | 68400000 | 68500000 | 0.39 | 0.53 | A>A |
| chr1 | 68500000 | 68600000 | 0.11 | 0.11 | A>A |
| chr1 | 68600000 | 68700000 | -0.01 | 0.09 | A>A |
| chr1 | 68700000 | 68800000 | -0.07 | 0.00 | B>B |
| chr1 | 68800000 | 68900000 | -0.16 | -0.16 | B>B |
| chr1 | 68900000 | 69000000 | -0.38 | -0.37 | B>B |
| chr1 | 69000000 | 69100000 | -0.65 | -0.48 | B>B |
| chr1 | 69100000 | 69200000 | -0.68 | -0.36 | B>B |
| chr1 | 69200000 | 69300000 | -0.75 | -0.53 | B>B |
| chr1 | 69300000 | 69400000 | -0.77 | -0.63 | B>B |
| chr1 | 69400000 | 69500000 | -0.74 | -0.77 | B>B |
| chr1 | 69500000 | 69600000 | -0.77 | -0.77 | B>B |
| chr1 | 69600000 | 69700000 | -0.79 | -0.82 | B>B |
| chr1 | 69700000 | 69800000 | -0.72 | -0.82 | B>B |
| chr1 | 69800000 | 69900000 | -0.77 | -0.82 | B>B |
| chr1 | 69900000 | 70000000 | -0.76 | -0.85 | B>B |
| chr1 | 70000000 | 70100000 | -0.70 | -0.86 | B>B |
| chr1 | 70100000 | 70200000 | -0.72 | -0.78 | B>B |
| chr1 | 70200000 | 70300000 | -0.68 | -0.84 | B>B |
| chr1 | 70300000 | 70400000 | -0.78 | -0.89 | B>B |
| chr1 | 70400000 | 70500000 | -0.76 | -0.90 | B>B |
| chr1 | 70500000 | 70600000 | -0.65 | -0.77 | B>B |
| chr1 | 70600000 | 70700000 | -0.06 | -0.36 | B>B |
| chr1 | 70700000 | 70800000 | 0.13 | 0.00 | A>A |
| chr1 | 70800000 | 70900000 | 0.08 | -0.18 | B>B |
| chr1 | 70900000 | 71000000 | -0.33 | -0.51 | B>B |
| chr1 | 71000000 | 71100000 | -0.60 | -0.80 | B>B |
| chr1 | 71100000 | 71200000 | -0.72 | -0.90 | B>B |

| | | | | | |
|---|---|---|---|---|---|
| chr1 | 71200000 | 71300000 | -0.74 | -0.89 | B>B |
| chr1 | 71300000 | 71400000 | -0.79 | -0.94 | B>B |
| chr1 | 71400000 | 71500000 | -0.68 | -0.81 | B>B |
| chr1 | 71500000 | 71600000 | -0.52 | -0.56 | B>B |
| chr1 | 71600000 | 71700000 | -0.77 | -0.75 | B>B |
| chr1 | 71700000 | 71800000 | -0.78 | -0.81 | B>B |
| chr1 | 71800000 | 71900000 | -0.79 | -0.81 | B>B |
| chr1 | 71900000 | 72000000 | -0.76 | -0.86 | B>B |
| chr1 | 72000000 | 72100000 | -0.75 | -0.77 | B>B |
| chr1 | 72100000 | 72200000 | -0.74 | -0.79 | B>B |
| chr1 | 72200000 | 72300000 | -0.75 | -0.80 | B>B |
| chr1 | 72300000 | 72400000 | -0.73 | -0.77 | B>B |
| chr1 | 72400000 | 72500000 | -0.70 | -0.75 | B>B |
| chr1 | 72500000 | 72600000 | -0.68 | -0.77 | B>B |
| chr1 | 72600000 | 72700000 | -0.70 | -0.73 | B>B |
| chr1 | 72700000 | 72800000 | -0.69 | -0.69 | B>B |
| chr1 | 72800000 | 72900000 | -0.65 | -0.72 | B>B |
| chr1 | 72900000 | 73000000 | -0.68 | -0.73 | B>B |
| chr1 | 73000000 | 73100000 | -0.69 | -0.73 | B>B |
| chr1 | 73100000 | 73200000 | -0.69 | -0.72 | B>B |
| chr1 | 73200000 | 73300000 | -0.71 | -0.72 | B>B |
| chr1 | 73300000 | 73400000 | -0.72 | -0.76 | B>B |
| chr1 | 73400000 | 73500000 | -0.68 | -0.75 | B>B |
| chr1 | 73500000 | 73600000 | -0.66 | -0.72 | B>B |
| chr1 | 73600000 | 73700000 | -0.69 | -0.74 | B>B |
| chr1 | 73700000 | 73800000 | -0.73 | -0.73 | B>B |
| chr1 | 73800000 | 73900000 | -0.69 | -0.76 | B>B |
| chr1 | 73900000 | 74000000 | -0.70 | -0.74 | B>B |
| chr1 | 74000000 | 74100000 | -0.67 | -0.71 | B>B |
| chr1 | 74100000 | 74200000 | -0.65 | -0.74 | B>B |
| chr1 | 74200000 | 74300000 | -0.62 | -0.72 | B>B |
| chr1 | 74300000 | 74400000 | -0.60 | -0.75 | B>B |
| chr1 | 74400000 | 74500000 | -0.63 | -0.70 | B>B |
| chr1 | 74500000 | 74600000 | -0.67 | -0.75 | B>B |
| chr1 | 74600000 | 74700000 | -0.54 | -0.71 | B>B |
| chr1 | 74700000 | 74800000 | -0.58 | -0.76 | B>B |
| chr1 | 74800000 | 74900000 | -0.61 | -0.75 | B>B |
| chr1 | 74900000 | 75000000 | -0.55 | -0.71 | B>B |
| chr1 | 75000000 | 75100000 | -0.56 | -0.69 | B>B |
| chr1 | 75100000 | 75200000 | -0.47 | -0.57 | B>B |
| chr1 | 75200000 | 75300000 | -0.54 | -0.62 | B>B |
| chr1 | 75300000 | 75400000 | -0.75 | -0.76 | B>B |
| chr1 | 75400000 | 75500000 | -0.78 | -0.78 | B>B |
| chr1 | 75500000 | 75600000 | -0.64 | -0.55 | B>B |
| chr1 | 75600000 | 75700000 | -0.56 | -0.47 | B>B |
| chr1 | 75700000 | 75800000 | -0.69 | -0.33 | B>B |
| chr1 | 75800000 | 75900000 | -0.76 | -0.28 | B>B |
| chr1 | 75900000 | 76000000 | -0.74 | -0.35 | B>B |
| chr1 | 76000000 | 76100000 | -0.69 | -0.32 | B>B |
| chr1 | 76100000 | 76200000 | -0.54 | -0.34 | B>B |
| chr1 | 76200000 | 76300000 | -0.13 | 0.00 | B>B |
| chr1 | 76300000 | 76400000 | -0.41 | -0.31 | B>B |
| chr1 | 76400000 | 76500000 | -0.34 | -0.29 | B>B |
| chr1 | 76500000 | 76600000 | -0.38 | -0.38 | B>B |
| chr1 | 76600000 | 76700000 | -0.57 | -0.64 | B>B |
| chr1 | 76700000 | 76800000 | -0.67 | -0.83 | B>B |

| | | | | | |
|---|---|---|---|---|---|
| chr1 | 76800000 | 76900000 | -0.70 | -0.83 | B>B |
| chr1 | 76900000 | 77000000 | -0.67 | -0.83 | B>B |
| chr1 | 77000000 | 77100000 | -0.62 | -0.84 | B>B |
| chr1 | 77100000 | 77200000 | -0.67 | -0.83 | B>B |
| chr1 | 77200000 | 77300000 | -0.61 | -0.75 | B>B |
| chr1 | 77300000 | 77400000 | -0.43 | -0.56 | B>B |
| chr1 | 77400000 | 77500000 | -0.25 | -0.41 | B>B |
| chr1 | 77500000 | 77600000 | -0.25 | -0.38 | B>B |
| chr1 | 77600000 | 77700000 | 0.15 | 0.14 | A>A |
| chr1 | 77700000 | 77800000 | 0.42 | 0.47 | A>A |
| chr1 | 77800000 | 77900000 | 0.51 | 0.55 | A>A |
| chr1 | 77900000 | 78000000 | 0.49 | 0.49 | A>A |
| chr1 | 78000000 | 78100000 | 0.46 | 0.44 | A>A |
| chr1 | 78100000 | 78200000 | 0.57 | 0.48 | A>A |
| chr1 | 78200000 | 78300000 | 0.69 | 0.60 | A>A |
| chr1 | 78300000 | 78400000 | 0.41 | 0.49 | A>A |
| chr1 | 78400000 | 78500000 | 0.46 | 0.26 | A>A |
| chr1 | 78500000 | 78600000 | -0.03 | -0.09 | B>B |
| chr1 | 78600000 | 78700000 | -0.56 | -0.62 | B>B |
| chr1 | 78700000 | 78800000 | -0.69 | -0.75 | B>B |
| chr1 | 78800000 | 78900000 | -0.69 | -0.81 | B>B |
| chr1 | 78900000 | 79000000 | -0.79 | -0.75 | B>B |
| chr1 | 79000000 | 79100000 | -0.61 | -0.58 | B>B |
| chr1 | 79100000 | 79200000 | -0.47 | -0.31 | B>B |
| chr1 | 79200000 | 79300000 | -0.39 | -0.30 | B>B |
| chr1 | 79300000 | 79400000 | -0.21 | -0.30 | B>B |
| chr1 | 79400000 | 79500000 | -0.03 | -0.28 | B>B |
| chr1 | 79500000 | 79600000 | -0.03 | -0.15 | B>B |
| chr1 | 79600000 | 79700000 | -0.20 | -0.16 | B>B |
| chr1 | 79700000 | 79800000 | -0.32 | -0.03 | B>B |
| chr1 | 79800000 | 79900000 | -0.43 | 0.14 | B>A |
| chr1 | 79900000 | 80000000 | -0.48 | 0.03 | B>B |
| chr1 | 80000000 | 80100000 | -0.59 | 0.19 | B>A |
| chr1 | 80100000 | 80200000 | -0.58 | 0.31 | B>A |
| chr1 | 80200000 | 80300000 | -0.57 | 0.39 | B>A |
| chr1 | 80300000 | 80400000 | -0.61 | 0.34 | B>A |
| chr1 | 80400000 | 80500000 | -0.57 | 0.35 | B>A |
| chr1 | 80500000 | 80600000 | -0.56 | 0.29 | B>A |
| chr1 | 80600000 | 80700000 | -0.56 | 0.14 | B>A |
| chr1 | 80700000 | 80800000 | -0.58 | 0.25 | B>A |
| chr1 | 80800000 | 80900000 | -0.56 | 0.27 | B>A |
| chr1 | 80900000 | 81000000 | -0.53 | 0.33 | B>A |
| chr1 | 81000000 | 81100000 | -0.56 | 0.30 | B>A |
| chr1 | 81100000 | 81200000 | -0.58 | 0.33 | B>A |
| chr1 | 81200000 | 81300000 | -0.59 | 0.32 | B>A |
| chr1 | 81300000 | 81400000 | -0.61 | 0.25 | B>A |
| chr1 | 81400000 | 81500000 | -0.60 | 0.24 | B>A |
| chr1 | 81500000 | 81600000 | -0.60 | 0.24 | B>A |
| chr1 | 81600000 | 81700000 | -0.63 | 0.21 | B>A |
| chr1 | 81700000 | 81800000 | -0.62 | 0.08 | B>A |
| chr1 | 81800000 | 81900000 | -0.62 | 0.03 | B>A |
| chr1 | 81900000 | 82000000 | -0.56 | 0.00 | B>A |
| chr1 | 82000000 | 82100000 | -0.51 | -0.01 | B>B |
| chr1 | 82100000 | 82200000 | -0.38 | 0.13 | B>B |
| chr1 | 82200000 | 82300000 | -0.41 | -0.09 | B>B |
| chr1 | 82300000 | 82400000 | -0.39 | -0.18 | B>B |

| | | | | | |
|---|---|---|---|---|---|
| chr1 | 82400000 | 82500000 | -0.47 | -0.29 | B>B |
| chr1 | 82500000 | 82600000 | -0.59 | -0.39 | B>B |
| chr1 | 82600000 | 82700000 | -0.66 | -0.28 | B>B |
| chr1 | 82700000 | 82800000 | -0.68 | -0.36 | B>B |
| chr1 | 82800000 | 82900000 | -0.70 | -0.40 | B>B |
| chr1 | 82900000 | 83000000 | -0.74 | -0.46 | B>B |
| chr1 | 83000000 | 83100000 | -0.71 | -0.56 | B>B |
| chr1 | 83100000 | 83200000 | -0.70 | -0.68 | B>B |
| chr1 | 83200000 | 83300000 | -0.75 | -0.68 | B>B |
| chr1 | 83300000 | 83400000 | -0.75 | -0.77 | B>B |
| chr1 | 83400000 | 83500000 | -0.77 | -0.77 | B>B |
| chr1 | 83500000 | 83600000 | -0.77 | -0.79 | B>B |
| chr1 | 83600000 | 83700000 | 0.00 | 0.00 | NO |
| chr1 | 83700000 | 83800000 | 0.00 | 0.00 | NO |
| chr1 | 83800000 | 83900000 | 0.00 | 0.00 | NO |
| chr1 | 83900000 | 84000000 | 0.00 | 0.00 | NO |
| chr1 | 84000000 | 84100000 | -0.75 | -0.78 | B>B |
| chr1 | 84100000 | 84200000 | -0.68 | -0.87 | B>B |
| chr1 | 84200000 | 84300000 | -0.64 | -0.77 | B>B |
| chr1 | 84300000 | 84400000 | -0.34 | -0.36 | B>B |
| chr1 | 84400000 | 84500000 | 0.04 | -0.04 | B>B |
| chr1 | 84500000 | 84600000 | 0.08 | 0.04 | A>A |
| chr1 | 84600000 | 84700000 | 0.27 | 0.39 | A>A |
| chr1 | 84700000 | 84800000 | 0.43 | 0.63 | A>A |
| chr1 | 84800000 | 84900000 | 0.33 | 0.45 | A>A |
| chr1 | 84900000 | 85000000 | 0.22 | 0.33 | A>A |
| chr1 | 85000000 | 85100000 | 0.29 | 0.21 | A>A |
| chr1 | 85100000 | 85200000 | 0.31 | 0.21 | A>A |
| chr1 | 85200000 | 85300000 | -0.03 | 0.13 | A>A |
| chr1 | 85300000 | 85400000 | 0.07 | 0.28 | A>A |
| chr1 | 85400000 | 85500000 | 0.32 | 0.32 | A>A |
| chr1 | 85500000 | 85600000 | 0.53 | 0.48 | A>A |
| chr1 | 85600000 | 85700000 | 0.60 | 0.58 | A>A |
| chr1 | 85700000 | 85800000 | 0.29 | 0.34 | A>A |
| chr1 | 85800000 | 85900000 | 0.47 | 0.45 | A>A |
| chr1 | 85900000 | 86000000 | 0.97 | 0.82 | A>A |
| chr1 | 86000000 | 86100000 | 0.58 | 0.46 | A>A |
| chr1 | 86100000 | 86200000 | 0.38 | 0.17 | A>A |
| chr1 | 86200000 | 86300000 | 0.35 | 0.09 | A>A |
| chr1 | 86300000 | 86400000 | 0.05 | -0.09 | B>B |
| chr1 | 86400000 | 86500000 | -0.17 | -0.30 | B>B |
| chr1 | 86500000 | 86600000 | -0.39 | -0.57 | B>B |
| chr1 | 86600000 | 86700000 | -0.42 | -0.64 | B>B |
| chr1 | 86700000 | 86800000 | -0.46 | -0.69 | B>B |
| chr1 | 86800000 | 86900000 | -0.35 | -0.53 | B>B |
| chr1 | 86900000 | 87000000 | -0.41 | -0.56 | B>B |
| chr1 | 87000000 | 87100000 | -0.55 | -0.66 | B>B |
| chr1 | 87100000 | 87200000 | -0.35 | -0.38 | B>B |
| chr1 | 87200000 | 87300000 | -0.10 | -0.04 | B>B |
| chr1 | 87300000 | 87400000 | 0.21 | 0.19 | A>A |
| chr1 | 87400000 | 87500000 | 0.17 | 0.01 | A>A |
| chr1 | 87500000 | 87600000 | -0.13 | -0.19 | B>B |
| chr1 | 87600000 | 87700000 | -0.28 | -0.45 | B>B |
| chr1 | 87700000 | 87800000 | -0.27 | -0.33 | B>B |
| chr1 | 87800000 | 87900000 | 0.09 | 0.09 | A>A |
| chr1 | 87900000 | 88000000 | 0.09 | 0.20 | A>A |

| | | | | | |
|---|---|---|---|---|---|
| chr1 | 88000000 | 88100000 | 0.25 | 0.29 | A>A |
| chr1 | 88100000 | 88200000 | 0.21 | 0.33 | A>A |
| chr1 | 88200000 | 88300000 | 0.29 | 0.41 | A>A |
| chr1 | 88300000 | 88400000 | 0.27 | 0.41 | A>A |
| chr1 | 88400000 | 88500000 | 0.05 | 0.14 | A>A |
| chr1 | 88500000 | 88600000 | -0.13 | -0.04 | B>B |
| chr1 | 88600000 | 88700000 | -0.30 | -0.27 | B>B |
| chr1 | 88700000 | 88800000 | -0.44 | -0.39 | B>B |
| chr1 | 88800000 | 88900000 | -0.67 | -0.61 | B>B |
| chr1 | 88900000 | 89000000 | -0.60 | -0.60 | B>B |
| chr1 | 89000000 | 89100000 | -0.52 | -0.51 | B>B |
| chr1 | 89100000 | 89200000 | -0.05 | -0.11 | B>B |
| chr1 | 89200000 | 89300000 | 0.18 | 0.23 | A>A |
| chr1 | 89300000 | 89400000 | -0.03 | 0.03 | B>B |
| chr1 | 89400000 | 89500000 | -0.29 | -0.24 | B>B |
| chr1 | 89500000 | 89600000 | -0.69 | -0.29 | B>B |
| chr1 | 89600000 | 89700000 | -0.81 | -0.50 | B>B |
| chr1 | 89700000 | 89800000 | -0.66 | -0.66 | B>B |
| chr1 | 89800000 | 89900000 | -0.56 | -0.58 | B>B |
| chr1 | 89900000 | 90000000 | -0.31 | -0.35 | B>B |
| chr1 | 90000000 | 90100000 | -0.01 | -0.11 | B>B |
| chr1 | 90100000 | 90200000 | 0.01 | -0.11 | B>B |
| chr1 | 90200000 | 90300000 | 0.04 | -0.03 | A>A |
| chr1 | 90300000 | 90400000 | 0.46 | 0.42 | A>A |
| chr1 | 90400000 | 90500000 | 0.20 | 0.02 | A>A |
| chr1 | 90500000 | 90600000 | -0.38 | -0.52 | B>B |
| chr1 | 90600000 | 90700000 | -0.61 | -0.75 | B>B |
| chr1 | 90700000 | 90800000 | -0.73 | -0.86 | B>B |
| chr1 | 90800000 | 90900000 | -0.68 | -0.80 | B>B |
| chr1 | 90900000 | 91000000 | -0.73 | -0.81 | B>B |
| chr1 | 91000000 | 91100000 | -0.62 | -0.81 | B>B |
| chr1 | 91100000 | 91200000 | -0.61 | -0.72 | B>B |
| chr1 | 91200000 | 91300000 | -0.46 | -0.74 | B>B |
| chr1 | 91300000 | 91400000 | -0.30 | -0.45 | B>B |
| chr1 | 91400000 | 91500000 | -0.05 | -0.31 | B>B |
| chr1 | 91500000 | 91600000 | -0.22 | -0.56 | B>B |
| chr1 | 91600000 | 91700000 | -0.39 | -0.60 | B>B |
| chr1 | 91700000 | 91800000 | -0.35 | -0.62 | B>B |
| chr1 | 91800000 | 91900000 | -0.34 | -0.53 | B>B |
| chr1 | 91900000 | 92000000 | 0.10 | -0.25 | B>B |
| chr1 | 92000000 | 92100000 | 0.63 | 0.43 | A>A |
| chr1 | 92100000 | 92200000 | 0.60 | 0.48 | A>A |
| chr1 | 92200000 | 92300000 | 0.70 | 0.49 | A>A |
| chr1 | 92300000 | 92400000 | 0.56 | 0.31 | A>A |
| chr1 | 92400000 | 92500000 | 0.28 | 0.11 | A>A |
| chr1 | 92500000 | 92600000 | 0.35 | 0.18 | A>A |
| chr1 | 92600000 | 92700000 | 0.36 | 0.08 | A>A |
| chr1 | 92700000 | 92800000 | 0.29 | 0.30 | A>A |
| chr1 | 92800000 | 92900000 | 0.31 | 0.42 | A>A |
| chr1 | 92900000 | 93000000 | 0.53 | 0.59 | A>A |
| chr1 | 93000000 | 93100000 | 0.47 | 0.53 | A>A |
| chr1 | 93100000 | 93200000 | 0.78 | 0.84 | A>A |
| chr1 | 93200000 | 93300000 | 1.05 | 1.06 | A>A |
| chr1 | 93300000 | 93400000 | 1.04 | 1.25 | A>A |
| chr1 | 93400000 | 93500000 | 1.23 | 1.29 | A>A |
| chr1 | 93500000 | 93600000 | 1.17 | 1.33 | A>A |

| | | | | | |
|---|---|---|---|---|---|
| chr1 | 93600000 | 93700000 | 1.27 | 1.59 | A>A |
| chr1 | 93700000 | 93800000 | 1.05 | 1.39 | A>A |
| chr1 | 93800000 | 93900000 | 1.20 | 1.44 | A>A |
| chr1 | 93900000 | 94000000 | 1.04 | 1.51 | A>A |
| chr1 | 94000000 | 94100000 | 0.97 | 1.39 | A>A |
| chr1 | 94100000 | 94200000 | 1.46 | 1.82 | A>A |
| chr1 | 94200000 | 94300000 | 1.32 | 1.59 | A>A |
| chr1 | 94300000 | 94400000 | 0.86 | 1.22 | A>A |
| chr1 | 94400000 | 94500000 | 1.18 | 1.14 | A>A |
| chr1 | 94500000 | 94600000 | 1.39 | 1.48 | A>A |
| chr1 | 94600000 | 94700000 | 1.58 | 1.66 | A>A |
| chr1 | 94700000 | 94800000 | 1.35 | 1.46 | A>A |
| chr1 | 94800000 | 94900000 | 1.02 | 1.25 | A>A |
| chr1 | 94900000 | 95000000 | 0.61 | 0.75 | A>A |
| chr1 | 95000000 | 95100000 | 0.94 | 0.99 | A>A |
| chr1 | 95100000 | 95200000 | 0.88 | 1.01 | A>A |
| chr1 | 95200000 | 95300000 | 0.61 | 0.69 | A>A |
| chr1 | 95300000 | 95400000 | 0.57 | 0.65 | A>A |
| chr1 | 95400000 | 95500000 | 0.71 | 0.79 | A>A |
| chr1 | 95500000 | 95600000 | 0.29 | 0.25 | A>A |
| chr1 | 95600000 | 95700000 | 0.28 | 0.23 | A>A |
| chr1 | 95700000 | 95800000 | -0.35 | -0.35 | B>B |
| chr1 | 95800000 | 95900000 | -0.63 | -0.76 | B>B |
| chr1 | 95900000 | 96000000 | -0.69 | -0.82 | B>B |
| chr1 | 96000000 | 96100000 | -0.69 | -0.87 | B>B |
| chr1 | 96100000 | 96200000 | -0.73 | -0.87 | B>B |
| chr1 | 96200000 | 96300000 | -0.71 | -0.81 | B>B |
| chr1 | 96300000 | 96400000 | -0.72 | -0.79 | B>B |
| chr1 | 96400000 | 96500000 | -0.72 | -0.78 | B>B |
| chr1 | 96500000 | 96600000 | -0.70 | -0.73 | B>B |
| chr1 | 96600000 | 96700000 | -0.61 | -0.61 | B>B |
| chr1 | 96700000 | 96800000 | -0.51 | -0.34 | B>B |
| chr1 | 96800000 | 96900000 | -0.42 | -0.30 | B>B |
| chr1 | 96900000 | 97000000 | -0.46 | -0.29 | B>B |
| chr1 | 97000000 | 97100000 | -0.39 | -0.29 | B>B |
| chr1 | 97100000 | 97200000 | -0.35 | -0.20 | B>B |
| chr1 | 97200000 | 97300000 | -0.17 | -0.06 | B>B |
| chr1 | 97300000 | 97400000 | -0.43 | -0.11 | B>B |
| chr1 | 97400000 | 97500000 | -0.40 | -0.09 | B>B |
| chr1 | 97500000 | 97600000 | -0.34 | -0.02 | B>B |
| chr1 | 97600000 | 97700000 | -0.24 | 0.01 | B>B |
| chr1 | 97700000 | 97800000 | -0.24 | 0.14 | B>B |
| chr1 | 97800000 | 97900000 | -0.28 | 0.10 | B>B |
| chr1 | 97900000 | 98000000 | -0.23 | 0.08 | B>B |
| chr1 | 98000000 | 98100000 | -0.21 | 0.05 | B>B |
| chr1 | 98100000 | 98200000 | -0.16 | 0.12 | B>B |
| chr1 | 98200000 | 98300000 | -0.22 | 0.05 | B>B |
| chr1 | 98300000 | 98400000 | -0.04 | 0.22 | A>A |
| chr1 | 98400000 | 98500000 | -0.02 | 0.25 | A>A |
| chr1 | 98500000 | 98600000 | 0.10 | 0.18 | A>A |
| chr1 | 98600000 | 98700000 | 0.25 | 0.29 | A>A |
| chr1 | 98700000 | 98800000 | 0.29 | 0.40 | A>A |
| chr1 | 98800000 | 98900000 | 0.32 | 0.29 | A>A |
| chr1 | 98900000 | 99000000 | -0.21 | -0.08 | B>B |
| chr1 | 99000000 | 99100000 | -0.39 | -0.18 | B>B |
| chr1 | 99100000 | 99200000 | -0.26 | -0.14 | B>B |

| | | | | | |
|---|---|---|---|---|---|
| chr1 | 99200000 | 99300000 | -0.49 | -0.43 | B>B |
| chr1 | 99300000 | 99400000 | -0.61 | -0.48 | B>B |
| chr1 | 99400000 | 99500000 | -0.68 | -0.53 | B>B |
| chr1 | 99500000 | 99600000 | -0.71 | -0.51 | B>B |
| chr1 | 99600000 | 99700000 | -0.56 | -0.51 | B>B |
| chr1 | 99700000 | 99800000 | -0.70 | -0.62 | B>B |
| chr1 | 99800000 | 99900000 | -0.73 | -0.67 | B>B |
| chr1 | 99900000 | 100000000 | -0.46 | -0.39 | B>B |
| chr1 | 1E+08 | 100100000 | -0.35 | -0.19 | B>B |
| chr1 | 1E+08 | 100200000 | -0.35 | -0.20 | B>B |
| chr1 | 1E+08 | 100300000 | -0.07 | -0.03 | B>B |
| chr1 | 1E+08 | 100400000 | 0.37 | 0.38 | A>A |
| chr1 | 1E+08 | 100500000 | 0.50 | 0.58 | A>A |
| chr1 | 1.01E+08 | 100600000 | 0.53 | 0.56 | A>A |
| chr1 | 1.01E+08 | 100700000 | 0.49 | 0.58 | A>A |
| chr1 | 1.01E+08 | 100800000 | 0.64 | 0.62 | A>A |
| chr1 | 1.01E+08 | 100900000 | 0.45 | 0.37 | A>A |
| chr1 | 1.01E+08 | 101000000 | 0.02 | 0.14 | A>A |
| chr1 | 1.01E+08 | 101100000 | -0.05 | 0.17 | A>A |
| chr1 | 1.01E+08 | 101200000 | -0.03 | 0.38 | A>A |
| chr1 | 1.01E+08 | 101300000 | -0.07 | 0.31 | A>A |
| chr1 | 1.01E+08 | 101400000 | -0.01 | -0.05 | B>B |
| chr1 | 1.01E+08 | 101500000 | -0.01 | -0.16 | B>B |
| chr1 | 1.02E+08 | 101600000 | -0.22 | -0.50 | B>B |
| chr1 | 1.02E+08 | 101700000 | -0.33 | -0.61 | B>B |
| chr1 | 1.02E+08 | 101800000 | -0.31 | -0.46 | B>B |
| chr1 | 1.02E+08 | 101900000 | -0.63 | -0.73 | B>B |
| chr1 | 1.02E+08 | 102000000 | -0.70 | -0.79 | B>B |
| chr1 | 1.02E+08 | 102100000 | -0.74 | -0.83 | B>B |
| chr1 | 1.02E+08 | 102200000 | -0.72 | -0.81 | B>B |
| chr1 | 1.02E+08 | 102300000 | -0.70 | -0.81 | B>B |
| chr1 | 1.02E+08 | 102400000 | -0.71 | -0.81 | B>B |
| chr1 | 1.02E+08 | 102500000 | -0.69 | -0.73 | B>B |
| chr1 | 1.03E+08 | 102600000 | -0.71 | -0.77 | B>B |
| chr1 | 1.03E+08 | 102700000 | -0.65 | -0.77 | B>B |
| chr1 | 1.03E+08 | 102800000 | -0.67 | -0.76 | B>B |
| chr1 | 1.03E+08 | 102900000 | -0.66 | -0.79 | B>B |
| chr1 | 1.03E+08 | 103000000 | -0.67 | -0.74 | B>B |
| chr1 | 1.03E+08 | 103100000 | -0.67 | -0.75 | B>B |
| chr1 | 1.03E+08 | 103200000 | -0.68 | -0.75 | B>B |
| chr1 | 1.03E+08 | 103300000 | -0.59 | -0.76 | B>B |
| chr1 | 1.03E+08 | 103400000 | -0.56 | -0.72 | B>B |
| chr1 | 1.03E+08 | 103500000 | -0.62 | -0.70 | B>B |
| chr1 | 1.04E+08 | 103600000 | -0.63 | -0.69 | B>B |
| chr1 | 1.04E+08 | 103700000 | -0.70 | -0.70 | B>B |
| chr1 | 1.04E+08 | 103800000 | -0.71 | -0.74 | B>B |
| chr1 | 1.04E+08 | 103900000 | 0.00 | 0.00 | NO |
| chr1 | 1.04E+08 | 104000000 | 0.00 | 0.00 | NO |
| chr1 | 1.04E+08 | 104100000 | -0.55 | -0.60 | B>B |
| chr1 | 1.04E+08 | 104200000 | 0.00 | 0.00 | NO |
| chr1 | 1.04E+08 | 104300000 | 0.00 | 0.00 | NO |
| chr1 | 1.04E+08 | 104400000 | -0.67 | -0.59 | B>B |
| chr1 | 1.04E+08 | 104500000 | -0.69 | -0.62 | B>B |
| chr1 | 1.05E+08 | 104600000 | -0.65 | -0.49 | B>B |
| chr1 | 1.05E+08 | 104700000 | -0.60 | -0.54 | B>B |
| chr1 | 1.05E+08 | 104800000 | -0.67 | -0.51 | B>B |

| | | | | | |
|---|---|---|---|---|---|
| chr1 | 1.05E+08 | 104900000 | -0.63 | -0.46 | B>B |
| chr1 | 1.05E+08 | 105000000 | -0.65 | -0.53 | B>B |
| chr1 | 1.05E+08 | 105100000 | -0.65 | -0.52 | B>B |
| chr1 | 1.05E+08 | 105200000 | -0.61 | -0.53 | B>B |
| chr1 | 1.05E+08 | 105300000 | -0.62 | -0.41 | B>B |
| chr1 | 1.05E+08 | 105400000 | -0.62 | -0.38 | B>B |
| chr1 | 1.05E+08 | 105500000 | -0.65 | -0.29 | B>B |
| chr1 | 1.06E+08 | 105600000 | -0.59 | -0.31 | B>B |
| chr1 | 1.06E+08 | 105700000 | -0.64 | -0.28 | B>B |
| chr1 | 1.06E+08 | 105800000 | -0.62 | -0.25 | B>B |
| chr1 | 1.06E+08 | 105900000 | -0.66 | -0.33 | B>B |
| chr1 | 1.06E+08 | 106000000 | -0.64 | -0.23 | B>B |
| chr1 | 1.06E+08 | 106100000 | -0.64 | -0.33 | B>B |
| chr1 | 1.06E+08 | 106200000 | -0.60 | -0.31 | B>B |
| chr1 | 1.06E+08 | 106300000 | -0.61 | -0.36 | B>B |
| chr1 | 1.06E+08 | 106400000 | -0.63 | -0.41 | B>B |
| chr1 | 1.06E+08 | 106500000 | -0.63 | -0.49 | B>B |
| chr1 | 1.07E+08 | 106600000 | -0.61 | -0.51 | B>B |
| chr1 | 1.07E+08 | 106700000 | -0.63 | -0.63 | B>B |
| chr1 | 1.07E+08 | 106800000 | -0.62 | -0.68 | B>B |
| chr1 | 1.07E+08 | 106900000 | -0.62 | -0.66 | B>B |
| chr1 | 1.07E+08 | 107000000 | -0.67 | -0.70 | B>B |
| chr1 | 1.07E+08 | 107100000 | -0.66 | -0.70 | B>B |
| chr1 | 1.07E+08 | 107200000 | -0.67 | -0.74 | B>B |
| chr1 | 1.07E+08 | 107300000 | -0.68 | -0.71 | B>B |
| chr1 | 1.07E+08 | 107400000 | -0.68 | -0.74 | B>B |
| chr1 | 1.07E+08 | 107500000 | -0.70 | -0.72 | B>B |
| chr1 | 1.08E+08 | 107600000 | -0.64 | -0.73 | B>B |
| chr1 | 1.08E+08 | 107700000 | -0.62 | -0.67 | B>B |
| chr1 | 1.08E+08 | 107800000 | -0.69 | -0.72 | B>B |
| chr1 | 1.08E+08 | 107900000 | -0.61 | -0.77 | B>B |
| chr1 | 1.08E+08 | 108000000 | -0.54 | -0.67 | B>B |
| chr1 | 1.08E+08 | 108100000 | -0.40 | -0.59 | B>B |
| chr1 | 1.08E+08 | 108200000 | -0.16 | -0.28 | B>B |
| chr1 | 1.08E+08 | 108300000 | -0.22 | -0.26 | B>B |
| chr1 | 1.08E+08 | 108400000 | -0.46 | -0.54 | B>B |
| chr1 | 1.08E+08 | 108500000 | -0.54 | -0.77 | B>B |
| chr1 | 1.09E+08 | 108600000 | -0.47 | -0.67 | B>B |
| chr1 | 1.09E+08 | 108700000 | -0.39 | -0.54 | B>B |
| chr1 | 1.09E+08 | 108800000 | 0.00 | 0.00 | NO |
| chr1 | 1.09E+08 | 108900000 | 0.00 | 0.00 | NO |
| chr1 | 1.09E+08 | 109000000 | 0.00 | 0.00 | NO |
| chr1 | 1.09E+08 | 109100000 | 0.00 | 0.00 | NO |
| chr1 | 1.09E+08 | 109200000 | 0.52 | 0.59 | A>A |
| chr1 | 1.09E+08 | 109300000 | 1.14 | 1.01 | A>A |
| chr1 | 1.09E+08 | 109400000 | 1.15 | 1.18 | A>A |
| chr1 | 1.09E+08 | 109500000 | 1.25 | 1.34 | A>A |
| chr1 | 1.1E+08 | 109600000 | 1.48 | 1.59 | A>A |
| chr1 | 1.1E+08 | 109700000 | 1.64 | 1.92 | A>A |
| chr1 | 1.1E+08 | 109800000 | 1.75 | 1.98 | A>A |
| chr1 | 1.1E+08 | 109900000 | 1.84 | 1.82 | A>A |
| chr1 | 1.1E+08 | 110000000 | 1.59 | 1.73 | A>A |
| chr1 | 1.1E+08 | 110100000 | 1.62 | 1.68 | A>A |
| chr1 | 1.1E+08 | 110200000 | 1.56 | 1.64 | A>A |
| chr1 | 1.1E+08 | 110300000 | 0.00 | 0.00 | NO |
| chr1 | 1.1E+08 | 110400000 | 1.65 | 1.64 | A>A |

| | | | | | |
|---|---|---|---|---|---|
| chr1 | 1.1E+08 | 110500000 | 2.30 | 2.07 | A>A |
| chr1 | 1.11E+08 | 110600000 | 1.50 | 1.40 | A>A |
| chr1 | 1.11E+08 | 110700000 | 1.35 | 0.94 | A>A |
| chr1 | 1.11E+08 | 110800000 | 0.98 | 0.68 | A>A |
| chr1 | 1.11E+08 | 110900000 | 1.11 | 0.74 | A>A |
| chr1 | 1.11E+08 | 111000000 | 0.99 | 0.61 | A>A |
| chr1 | 1.11E+08 | 111100000 | 0.49 | 0.03 | A>A |
| chr1 | 1.11E+08 | 111200000 | 0.29 | -0.33 | A>B |
| chr1 | 1.11E+08 | 111300000 | 0.00 | -0.48 | B>B |
| chr1 | 1.11E+08 | 111400000 | -0.11 | -0.41 | B>B |
| chr1 | 1.11E+08 | 111500000 | 0.06 | -0.20 | B>B |
| chr1 | 1.12E+08 | 111600000 | -0.11 | -0.26 | B>B |
| chr1 | 1.12E+08 | 111700000 | 0.16 | -0.16 | A>A |
| chr1 | 1.12E+08 | 111800000 | 0.18 | -0.18 | B>B |
| chr1 | 1.12E+08 | 111900000 | -0.04 | -0.35 | B>B |
| chr1 | 1.12E+08 | 112000000 | 0.21 | -0.20 | A>A |
| chr1 | 1.12E+08 | 112100000 | 0.30 | -0.05 | A>A |
| chr1 | 1.12E+08 | 112200000 | 0.53 | -0.06 | A>A |
| chr1 | 1.12E+08 | 112300000 | 0.34 | -0.10 | A>A |
| chr1 | 1.12E+08 | 112400000 | 0.05 | -0.48 | B>B |
| chr1 | 1.12E+08 | 112500000 | 0.04 | -0.63 | A>B |
| chr1 | 1.13E+08 | 112600000 | -0.06 | -0.71 | A>B |
| chr1 | 1.13E+08 | 112700000 | -0.24 | -0.74 | B>B |
| chr1 | 1.13E+08 | 112800000 | -0.08 | -0.72 | A>B |
| chr1 | 1.13E+08 | 112900000 | 0.04 | -0.49 | B>B |
| chr1 | 1.13E+08 | 113000000 | 0.50 | 0.16 | A>A |
| chr1 | 1.13E+08 | 113100000 | 0.71 | 0.37 | A>A |
| chr1 | 1.13E+08 | 113200000 | 0.84 | 0.63 | A>A |
| chr1 | 1.13E+08 | 113300000 | 0.81 | 0.73 | A>A |
| chr1 | 1.13E+08 | 113400000 | 0.00 | 0.78 | NO |
| chr1 | 1.13E+08 | 113500000 | 1.16 | 0.99 | A>A |
| chr1 | 1.14E+08 | 113600000 | 1.16 | 0.94 | A>A |
| chr1 | 1.14E+08 | 113700000 | 0.65 | 0.74 | A>A |
| chr1 | 1.14E+08 | 113800000 | 0.83 | 0.98 | A>A |
| chr1 | 1.14E+08 | 113900000 | 1.05 | 1.17 | A>A |
| chr1 | 1.14E+08 | 114000000 | 0.84 | 0.91 | A>A |
| chr1 | 1.14E+08 | 114100000 | 0.68 | 0.81 | A>A |
| chr1 | 1.14E+08 | 114200000 | 0.65 | 0.80 | A>A |
| chr1 | 1.14E+08 | 114300000 | 0.61 | 0.75 | A>A |
| chr1 | 1.14E+08 | 114400000 | 0.94 | 0.90 | A>A |
| chr1 | 1.14E+08 | 114500000 | 0.77 | 0.78 | A>A |
| chr1 | 1.15E+08 | 114600000 | 0.27 | 0.27 | A>A |
| chr1 | 1.15E+08 | 114700000 | 0.38 | 0.20 | A>A |
| chr1 | 1.15E+08 | 114800000 | 0.22 | 0.18 | A>A |
| chr1 | 1.15E+08 | 114900000 | 0.21 | 0.09 | A>A |
| chr1 | 1.15E+08 | 115000000 | 0.49 | 0.47 | A>A |
| chr1 | 1.15E+08 | 115100000 | 0.68 | 0.59 | A>A |
| chr1 | 1.15E+08 | 115200000 | 0.35 | 0.15 | A>A |
| chr1 | 1.15E+08 | 115300000 | 0.32 | 0.20 | A>A |
| chr1 | 1.15E+08 | 115400000 | -0.09 | -0.24 | B>B |
| chr1 | 1.15E+08 | 115500000 | -0.26 | -0.34 | B>B |
| chr1 | 1.16E+08 | 115600000 | -0.21 | -0.26 | B>B |
| chr1 | 1.16E+08 | 115700000 | -0.07 | -0.17 | B>B |
| chr1 | 1.16E+08 | 115800000 | 0.26 | 0.20 | A>A |
| chr1 | 1.16E+08 | 115900000 | 0.21 | 0.17 | A>A |
| chr1 | 1.16E+08 | 116000000 | 0.17 | 0.09 | A>A |

| | | | | | |
|---|---|---|---|---|---|
| chr1 | 1.16E+08 | 116100000 | -0.04 | -0.01 | B>B |
| chr1 | 1.16E+08 | 116200000 | -0.03 | -0.03 | B>B |
| chr1 | 1.16E+08 | 116300000 | 0.02 | -0.08 | B>B |
| chr1 | 1.16E+08 | 116400000 | -0.17 | -0.30 | B>B |
| chr1 | 1.16E+08 | 116500000 | -0.17 | -0.34 | B>B |
| chr1 | 1.17E+08 | 116600000 | -0.23 | -0.18 | B>B |
| chr1 | 1.17E+08 | 116700000 | -0.14 | -0.16 | B>B |
| chr1 | 1.17E+08 | 116800000 | -0.01 | -0.19 | B>B |
| chr1 | 1.17E+08 | 116900000 | 0.11 | -0.21 | B>B |
| chr1 | 1.17E+08 | 117000000 | 0.23 | 0.00 | A>A |
| chr1 | 1.17E+08 | 117100000 | 0.17 | 0.06 | A>A |
| chr1 | 1.17E+08 | 117200000 | 0.00 | 0.00 | NO |
| chr1 | 1.17E+08 | 117300000 | 0.57 | 0.37 | A>A |
| chr1 | 1.17E+08 | 117400000 | 0.53 | 0.46 | A>A |
| chr1 | 1.17E+08 | 117500000 | 0.60 | 0.59 | A>A |
| chr1 | 1.18E+08 | 117600000 | 0.51 | 0.43 | A>A |
| chr1 | 1.18E+08 | 117700000 | 0.52 | 0.60 | A>A |
| chr1 | 1.18E+08 | 117800000 | 0.78 | 1.00 | A>A |
| chr1 | 1.18E+08 | 117900000 | 0.36 | 0.54 | A>A |
| chr1 | 1.18E+08 | 118000000 | 0.44 | 0.46 | A>A |
| chr1 | 1.18E+08 | 118100000 | 0.10 | 0.15 | A>A |
| chr1 | 1.18E+08 | 118200000 | 0.36 | 0.24 | A>A |
| chr1 | 1.18E+08 | 118300000 | 0.14 | 0.12 | A>A |
| chr1 | 1.18E+08 | 118400000 | 0.14 | 0.05 | A>A |
| chr1 | 1.18E+08 | 118500000 | 0.02 | -0.03 | B>B |
| chr1 | 1.19E+08 | 118600000 | -0.41 | -0.66 | B>B |
| chr1 | 1.19E+08 | 118700000 | -0.52 | -0.68 | B>B |
| chr1 | 1.19E+08 | 118800000 | -0.53 | -0.74 | B>B |
| chr1 | 1.19E+08 | 118900000 | -0.62 | -0.83 | B>B |
| chr1 | 1.19E+08 | 119000000 | -0.71 | -0.86 | B>B |
| chr1 | 1.19E+08 | 119100000 | -0.74 | -0.79 | B>B |
| chr1 | 1.19E+08 | 119200000 | -0.67 | -0.83 | B>B |
| chr1 | 1.19E+08 | 119300000 | -0.68 | -0.74 | B>B |
| chr1 | 1.19E+08 | 119400000 | -0.65 | -0.76 | B>B |
| chr1 | 1.19E+08 | 119500000 | -0.52 | -0.70 | B>B |
| chr1 | 1.2E+08 | 119600000 | -0.46 | -0.57 | B>B |
| chr1 | 1.2E+08 | 119700000 | -0.18 | -0.34 | B>B |
| chr1 | 1.2E+08 | 119800000 | -0.17 | -0.37 | B>B |
| chr1 | 1.2E+08 | 119900000 | -0.25 | -0.58 | B>B |
| chr1 | 1.2E+08 | 120000000 | -0.19 | -0.76 | B>B |
| chr1 | 1.2E+08 | 120100000 | 0.00 | -0.75 | NO |
| chr1 | 1.2E+08 | 120200000 | 0.00 | -0.45 | NO |
| chr1 | 1.2E+08 | 120300000 | 0.50 | -0.20 | A>B |
| chr1 | 1.2E+08 | 120400000 | 0.72 | 0.04 | A>B |
| chr1 | 1.2E+08 | 120500000 | 0.00 | 0.39 | NO |
| chr1 | 1.21E+08 | 120600000 | 0.00 | 0.00 | NO |
| chr1 | 1.21E+08 | 120700000 | 0.00 | 0.00 | NO |
| chr1 | 1.21E+08 | 120800000 | 0.00 | 0.00 | NO |
| chr1 | 1.21E+08 | 120900000 | 0.00 | 0.00 | NO |
| chr1 | 1.21E+08 | 121000000 | 0.00 | 0.00 | NO |
| chr1 | 1.21E+08 | 121100000 | 0.00 | 0.00 | NO |
| chr1 | 1.21E+08 | 121200000 | 0.00 | 0.00 | NO |
| chr1 | 1.21E+08 | 121300000 | 0.00 | 0.00 | NO |
| chr1 | 1.21E+08 | 121400000 | 0.00 | 0.00 | NO |
| chr1 | 1.21E+08 | 121500000 | 0.00 | 0.00 | NO |
| chr1 | 1.22E+08 | 121600000 | 0.00 | 0.00 | NO |

| | | | | | |
|---|---|---|---|---|---|
| chr1 | 1.22E+08 | 121700000 | 0.00 | 0.00 | NO |
| chr1 | 1.22E+08 | 121800000 | 0.00 | 0.00 | NO |
| chr1 | 1.22E+08 | 121900000 | 0.00 | 0.00 | NO |
| chr1 | 1.22E+08 | 122000000 | 0.00 | 0.00 | NO |
| chr1 | 1.22E+08 | 122100000 | 0.00 | 0.00 | NO |
| chr1 | 1.22E+08 | 122200000 | 0.00 | 0.00 | NO |
| chr1 | 1.22E+08 | 122300000 | 0.00 | 0.00 | NO |
| chr1 | 1.22E+08 | 122400000 | 0.00 | 0.00 | NO |
| chr1 | 1.22E+08 | 122500000 | 0.00 | 0.00 | NO |
| chr1 | 1.23E+08 | 122600000 | 0.00 | 0.00 | NO |
| chr1 | 1.23E+08 | 122700000 | 0.00 | 0.00 | NO |
| chr1 | 1.23E+08 | 122800000 | 0.00 | 0.00 | NO |
| chr1 | 1.23E+08 | 122900000 | 0.00 | 0.00 | NO |
| chr1 | 1.23E+08 | 123000000 | 0.00 | 0.00 | NO |
| chr1 | 1.23E+08 | 123100000 | 0.00 | 0.00 | NO |
| chr1 | 1.23E+08 | 123200000 | 0.00 | 0.00 | NO |
| chr1 | 1.23E+08 | 123300000 | 0.00 | 0.00 | NO |
| chr1 | 1.23E+08 | 123400000 | 0.00 | 0.00 | NO |
| chr1 | 1.23E+08 | 123500000 | 0.00 | 0.00 | NO |
| chr1 | 1.24E+08 | 123600000 | 0.00 | 0.00 | NO |
| chr1 | 1.24E+08 | 123700000 | 0.00 | 0.00 | NO |
| chr1 | 1.24E+08 | 123800000 | 0.00 | 0.00 | NO |
| chr1 | 1.24E+08 | 123900000 | 0.00 | 0.00 | NO |
| chr1 | 1.24E+08 | 124000000 | 0.00 | 0.00 | NO |
| chr1 | 1.24E+08 | 124100000 | 0.00 | 0.00 | NO |
| chr1 | 1.24E+08 | 124200000 | 0.00 | 0.00 | NO |
| chr1 | 1.24E+08 | 124300000 | 0.00 | 0.00 | NO |
| chr1 | 1.24E+08 | 124400000 | 0.00 | 0.00 | NO |
| chr1 | 1.24E+08 | 124500000 | 0.00 | 0.00 | NO |
| chr1 | 1.25E+08 | 124600000 | 0.00 | 0.00 | NO |
| chr1 | 1.25E+08 | 124700000 | 0.00 | 0.00 | NO |
| chr1 | 1.25E+08 | 124800000 | 0.00 | 0.00 | NO |
| chr1 | 1.25E+08 | 124900000 | 0.00 | 0.00 | NO |
| chr1 | 1.25E+08 | 125000000 | 0.00 | 0.00 | NO |
| chr1 | 1.25E+08 | 125100000 | 0.00 | 0.00 | NO |
| chr1 | 1.25E+08 | 125200000 | 0.00 | 0.00 | NO |
| chr1 | 1.25E+08 | 125300000 | 0.00 | 0.00 | NO |
| chr1 | 1.25E+08 | 125400000 | 0.00 | 0.00 | NO |
| chr1 | 1.25E+08 | 125500000 | 0.00 | 0.00 | NO |
| chr1 | 1.26E+08 | 125600000 | 0.00 | 0.00 | NO |
| chr1 | 1.26E+08 | 125700000 | 0.00 | 0.00 | NO |
| chr1 | 1.26E+08 | 125800000 | 0.00 | 0.00 | NO |
| chr1 | 1.26E+08 | 125900000 | 0.00 | 0.00 | NO |
| chr1 | 1.26E+08 | 126000000 | 0.00 | 0.00 | NO |
| chr1 | 1.26E+08 | 126100000 | 0.00 | 0.00 | NO |
| chr1 | 1.26E+08 | 126200000 | 0.00 | 0.00 | NO |
| chr1 | 1.26E+08 | 126300000 | 0.00 | 0.00 | NO |
| chr1 | 1.26E+08 | 126400000 | 0.00 | 0.00 | NO |
| chr1 | 1.26E+08 | 126500000 | 0.00 | 0.00 | NO |
| chr1 | 1.27E+08 | 126600000 | 0.00 | 0.00 | NO |
| chr1 | 1.27E+08 | 126700000 | 0.00 | 0.00 | NO |
| chr1 | 1.27E+08 | 126800000 | 0.00 | 0.00 | NO |
| chr1 | 1.27E+08 | 126900000 | 0.00 | 0.00 | NO |
| chr1 | 1.27E+08 | 127000000 | 0.00 | 0.00 | NO |
| chr1 | 1.27E+08 | 127100000 | 0.00 | 0.00 | NO |
| chr1 | 1.27E+08 | 127200000 | 0.00 | 0.00 | NO |

| | | | | | |
|---|---|---|---|---|---|
| chr1 | 1.27E+08 | 127300000 | 0.00 | 0.00 | NO |
| chr1 | 1.27E+08 | 127400000 | 0.00 | 0.00 | NO |
| chr1 | 1.27E+08 | 127500000 | 0.00 | 0.00 | NO |
| chr1 | 1.28E+08 | 127600000 | 0.00 | 0.00 | NO |
| chr1 | 1.28E+08 | 127700000 | 0.00 | 0.00 | NO |
| chr1 | 1.28E+08 | 127800000 | 0.00 | 0.00 | NO |
| chr1 | 1.28E+08 | 127900000 | 0.00 | 0.00 | NO |
| chr1 | 1.28E+08 | 128000000 | 0.00 | 0.00 | NO |
| chr1 | 1.28E+08 | 128100000 | 0.00 | 0.00 | NO |
| chr1 | 1.28E+08 | 128200000 | 0.00 | 0.00 | NO |
| chr1 | 1.28E+08 | 128300000 | 0.00 | 0.00 | NO |
| chr1 | 1.28E+08 | 128400000 | 0.00 | 0.00 | NO |
| chr1 | 1.28E+08 | 128500000 | 0.00 | 0.00 | NO |
| chr1 | 1.29E+08 | 128600000 | 0.00 | 0.00 | NO |
| chr1 | 1.29E+08 | 128700000 | 0.00 | 0.00 | NO |
| chr1 | 1.29E+08 | 128800000 | 0.00 | 0.00 | NO |
| chr1 | 1.29E+08 | 128900000 | 0.00 | 0.00 | NO |
| chr1 | 1.29E+08 | 129000000 | 0.00 | 0.00 | NO |
| chr1 | 1.29E+08 | 129100000 | 0.00 | 0.00 | NO |
| chr1 | 1.29E+08 | 129200000 | 0.00 | 0.00 | NO |
| chr1 | 1.29E+08 | 129300000 | 0.00 | 0.00 | NO |
| chr1 | 1.29E+08 | 129400000 | 0.00 | 0.00 | NO |
| chr1 | 1.29E+08 | 129500000 | 0.00 | 0.00 | NO |
| chr1 | 1.3E+08 | 129600000 | 0.00 | 0.00 | NO |
| chr1 | 1.3E+08 | 129700000 | 0.00 | 0.00 | NO |
| chr1 | 1.3E+08 | 129800000 | 0.00 | 0.00 | NO |
| chr1 | 1.3E+08 | 129900000 | 0.00 | 0.00 | NO |
| chr1 | 1.3E+08 | 130000000 | 0.00 | 0.00 | NO |
| chr1 | 1.3E+08 | 130100000 | 0.00 | 0.00 | NO |
| chr1 | 1.3E+08 | 130200000 | 0.00 | 0.00 | NO |
| chr1 | 1.3E+08 | 130300000 | 0.00 | 0.00 | NO |
| chr1 | 1.3E+08 | 130400000 | 0.00 | 0.00 | NO |
| chr1 | 1.3E+08 | 130500000 | 0.00 | 0.00 | NO |
| chr1 | 1.31E+08 | 130600000 | 0.00 | 0.00 | NO |
| chr1 | 1.31E+08 | 130700000 | 0.00 | 0.00 | NO |
| chr1 | 1.31E+08 | 130800000 | 0.00 | 0.00 | NO |
| chr1 | 1.31E+08 | 130900000 | 0.00 | 0.00 | NO |
| chr1 | 1.31E+08 | 131000000 | 0.00 | 0.00 | NO |
| chr1 | 1.31E+08 | 131100000 | 0.00 | 0.00 | NO |
| chr1 | 1.31E+08 | 131200000 | 0.00 | 0.00 | NO |
| chr1 | 1.31E+08 | 131300000 | 0.00 | 0.00 | NO |
| chr1 | 1.31E+08 | 131400000 | 0.00 | 0.00 | NO |
| chr1 | 1.31E+08 | 131500000 | 0.00 | 0.00 | NO |
| chr1 | 1.32E+08 | 131600000 | 0.00 | 0.00 | NO |
| chr1 | 1.32E+08 | 131700000 | 0.00 | 0.00 | NO |
| chr1 | 1.32E+08 | 131800000 | 0.00 | 0.00 | NO |
| chr1 | 1.32E+08 | 131900000 | 0.00 | 0.00 | NO |
| chr1 | 1.32E+08 | 132000000 | 0.00 | 0.00 | NO |
| chr1 | 1.32E+08 | 132100000 | 0.00 | 0.00 | NO |
| chr1 | 1.32E+08 | 132200000 | 0.00 | 0.00 | NO |
| chr1 | 1.32E+08 | 132300000 | 0.00 | 0.00 | NO |
| chr1 | 1.32E+08 | 132400000 | 0.00 | 0.00 | NO |
| chr1 | 1.32E+08 | 132500000 | 0.00 | 0.00 | NO |
| chr1 | 1.33E+08 | 132600000 | 0.00 | 0.00 | NO |
| chr1 | 1.33E+08 | 132700000 | 0.00 | 0.00 | NO |
| chr1 | 1.33E+08 | 132800000 | 0.00 | 0.00 | NO |

| | | | | | |
|---|---|---|---|---|---|
| chr1 | 1.33E+08 | 132900000 | 0.00 | 0.00 | NO |
| chr1 | 1.33E+08 | 133000000 | 0.00 | 0.00 | NO |
| chr1 | 1.33E+08 | 133100000 | 0.00 | 0.00 | NO |
| chr1 | 1.33E+08 | 133200000 | 0.00 | 0.00 | NO |
| chr1 | 1.33E+08 | 133300000 | 0.00 | 0.00 | NO |
| chr1 | 1.33E+08 | 133400000 | 0.00 | 0.00 | NO |
| chr1 | 1.33E+08 | 133500000 | 0.00 | 0.00 | NO |
| chr1 | 1.34E+08 | 133600000 | 0.00 | 0.00 | NO |
| chr1 | 1.34E+08 | 133700000 | 0.00 | 0.00 | NO |
| chr1 | 1.34E+08 | 133800000 | 0.00 | 0.00 | NO |
| chr1 | 1.34E+08 | 133900000 | 0.00 | 0.00 | NO |
| chr1 | 1.34E+08 | 134000000 | 0.00 | 0.00 | NO |
| chr1 | 1.34E+08 | 134100000 | 0.00 | 0.00 | NO |
| chr1 | 1.34E+08 | 134200000 | 0.00 | 0.00 | NO |
| chr1 | 1.34E+08 | 134300000 | 0.00 | 0.00 | NO |
| chr1 | 1.34E+08 | 134400000 | 0.00 | 0.00 | NO |
| chr1 | 1.34E+08 | 134500000 | 0.00 | 0.00 | NO |
| chr1 | 1.35E+08 | 134600000 | 0.00 | 0.00 | NO |
| chr1 | 1.35E+08 | 134700000 | 0.00 | 0.00 | NO |
| chr1 | 1.35E+08 | 134800000 | 0.00 | 0.00 | NO |
| chr1 | 1.35E+08 | 134900000 | 0.00 | 0.00 | NO |
| chr1 | 1.35E+08 | 135000000 | 0.00 | 0.00 | NO |
| chr1 | 1.35E+08 | 135100000 | 0.00 | 0.00 | NO |
| chr1 | 1.35E+08 | 135200000 | 0.00 | 0.00 | NO |
| chr1 | 1.35E+08 | 135300000 | 0.00 | 0.00 | NO |
| chr1 | 1.35E+08 | 135400000 | 0.00 | 0.00 | NO |
| chr1 | 1.35E+08 | 135500000 | 0.00 | 0.00 | NO |
| chr1 | 1.36E+08 | 135600000 | 0.00 | 0.00 | NO |
| chr1 | 1.36E+08 | 135700000 | 0.00 | 0.00 | NO |
| chr1 | 1.36E+08 | 135800000 | 0.00 | 0.00 | NO |
| chr1 | 1.36E+08 | 135900000 | 0.00 | 0.00 | NO |
| chr1 | 1.36E+08 | 136000000 | 0.00 | 0.00 | NO |
| chr1 | 1.36E+08 | 136100000 | 0.00 | 0.00 | NO |
| chr1 | 1.36E+08 | 136200000 | 0.00 | 0.00 | NO |
| chr1 | 1.36E+08 | 136300000 | 0.00 | 0.00 | NO |
| chr1 | 1.36E+08 | 136400000 | 0.00 | 0.00 | NO |
| chr1 | 1.36E+08 | 136500000 | 0.00 | 0.00 | NO |
| chr1 | 1.37E+08 | 136600000 | 0.00 | 0.00 | NO |
| chr1 | 1.37E+08 | 136700000 | 0.00 | 0.00 | NO |
| chr1 | 1.37E+08 | 136800000 | 0.00 | 0.00 | NO |
| chr1 | 1.37E+08 | 136900000 | 0.00 | 0.00 | NO |
| chr1 | 1.37E+08 | 137000000 | 0.00 | 0.00 | NO |
| chr1 | 1.37E+08 | 137100000 | 0.00 | 0.00 | NO |
| chr1 | 1.37E+08 | 137200000 | 0.00 | 0.00 | NO |
| chr1 | 1.37E+08 | 137300000 | 0.00 | 0.00 | NO |
| chr1 | 1.37E+08 | 137400000 | 0.00 | 0.00 | NO |
| chr1 | 1.37E+08 | 137500000 | 0.00 | 0.00 | NO |
| chr1 | 1.38E+08 | 137600000 | 0.00 | 0.00 | NO |
| chr1 | 1.38E+08 | 137700000 | 0.00 | 0.00 | NO |
| chr1 | 1.38E+08 | 137800000 | 0.00 | 0.00 | NO |
| chr1 | 1.38E+08 | 137900000 | 0.00 | 0.00 | NO |
| chr1 | 1.38E+08 | 138000000 | 0.00 | 0.00 | NO |
| chr1 | 1.38E+08 | 138100000 | 0.00 | 0.00 | NO |
| chr1 | 1.38E+08 | 138200000 | 0.00 | 0.00 | NO |
| chr1 | 1.38E+08 | 138300000 | 0.00 | 0.00 | NO |
| chr1 | 1.38E+08 | 138400000 | 0.00 | 0.00 | NO |

| | | | | | |
|---|---|---|---|---|---|
| chr1 | 1.38E+08 | 138500000 | 0.00 | 0.00 | NO |
| chr1 | 1.39E+08 | 138600000 | 0.00 | 0.00 | NO |
| chr1 | 1.39E+08 | 138700000 | 0.00 | 0.00 | NO |
| chr1 | 1.39E+08 | 138800000 | 0.00 | 0.00 | NO |
| chr1 | 1.39E+08 | 138900000 | 0.00 | 0.00 | NO |
| chr1 | 1.39E+08 | 139000000 | 0.00 | 0.00 | NO |
| chr1 | 1.39E+08 | 139100000 | 0.00 | 0.00 | NO |
| chr1 | 1.39E+08 | 139200000 | 0.00 | 0.00 | NO |
| chr1 | 1.39E+08 | 139300000 | 0.00 | 0.00 | NO |
| chr1 | 1.39E+08 | 139400000 | 0.00 | 0.00 | NO |
| chr1 | 1.39E+08 | 139500000 | 0.00 | 0.00 | NO |
| chr1 | 1.4E+08 | 139600000 | 0.00 | 0.00 | NO |
| chr1 | 1.4E+08 | 139700000 | 0.00 | 0.00 | NO |
| chr1 | 1.4E+08 | 139800000 | 0.00 | 0.00 | NO |
| chr1 | 1.4E+08 | 139900000 | 0.00 | 0.00 | NO |
| chr1 | 1.4E+08 | 140000000 | 0.00 | 0.00 | NO |
| chr1 | 1.4E+08 | 140100000 | 0.00 | 0.00 | NO |
| chr1 | 1.4E+08 | 140200000 | 0.00 | 0.00 | NO |
| chr1 | 1.4E+08 | 140300000 | 0.00 | 0.00 | NO |
| chr1 | 1.4E+08 | 140400000 | 0.00 | 0.00 | NO |
| chr1 | 1.4E+08 | 140500000 | 0.00 | 0.00 | NO |
| chr1 | 1.41E+08 | 140600000 | 0.00 | 0.00 | NO |
| chr1 | 1.41E+08 | 140700000 | 0.00 | 0.00 | NO |
| chr1 | 1.41E+08 | 140800000 | 0.00 | 0.00 | NO |
| chr1 | 1.41E+08 | 140900000 | 0.00 | 0.00 | NO |
| chr1 | 1.41E+08 | 141000000 | 0.00 | 0.00 | NO |
| chr1 | 1.41E+08 | 141100000 | 0.00 | 0.00 | NO |
| chr1 | 1.41E+08 | 141200000 | 0.00 | 0.00 | NO |
| chr1 | 1.41E+08 | 141300000 | 0.00 | 0.00 | NO |
| chr1 | 1.41E+08 | 141400000 | 0.00 | 0.00 | NO |
| chr1 | 1.41E+08 | 141500000 | 0.00 | 0.00 | NO |
| chr1 | 1.42E+08 | 141600000 | 0.00 | 0.00 | NO |
| chr1 | 1.42E+08 | 141700000 | 0.00 | 0.00 | NO |
| chr1 | 1.42E+08 | 141800000 | 0.00 | 0.00 | NO |
| chr1 | 1.42E+08 | 141900000 | 0.00 | 0.00 | NO |
| chr1 | 1.42E+08 | 142000000 | 0.00 | 0.00 | NO |
| chr1 | 1.42E+08 | 142100000 | 0.00 | 0.00 | NO |
| chr1 | 1.42E+08 | 142200000 | 0.00 | 0.00 | NO |
| chr1 | 1.42E+08 | 142300000 | 0.00 | 0.00 | NO |
| chr1 | 1.42E+08 | 142400000 | 0.00 | 0.00 | NO |
| chr1 | 1.42E+08 | 142500000 | 0.00 | 0.00 | NO |
| chr1 | 1.43E+08 | 142600000 | 0.00 | 0.00 | NO |
| chr1 | 1.43E+08 | 142700000 | 0.00 | 0.00 | NO |
| chr1 | 1.43E+08 | 142800000 | 0.00 | 0.00 | NO |
| chr1 | 1.43E+08 | 142900000 | 0.00 | 0.00 | NO |
| chr1 | 1.43E+08 | 143000000 | 0.00 | 0.00 | NO |
| chr1 | 1.43E+08 | 143100000 | 0.00 | 0.00 | NO |
| chr1 | 1.43E+08 | 143200000 | 0.00 | 0.00 | NO |
| chr1 | 1.43E+08 | 143300000 | 0.00 | 0.00 | NO |
| chr1 | 1.43E+08 | 143400000 | 0.00 | 0.00 | NO |
| chr1 | 1.43E+08 | 143500000 | 0.00 | 0.00 | NO |
| chr1 | 1.44E+08 | 143600000 | 0.00 | 0.00 | NO |
| chr1 | 1.44E+08 | 143700000 | 0.00 | 0.00 | NO |
| chr1 | 1.44E+08 | 143800000 | 0.00 | 0.00 | NO |
| chr1 | 1.44E+08 | 143900000 | 0.00 | 0.00 | NO |
| chr1 | 1.44E+08 | 144000000 | 0.00 | 0.00 | NO |

| | | | | | |
|---|---|---|---|---|---|
| chr1 | 1.44E+08 | 144100000 | 0.00 | 0.00 | NO |
| chr1 | 1.44E+08 | 144200000 | 0.00 | 0.00 | NO |
| chr1 | 1.44E+08 | 144300000 | 0.00 | 0.00 | NO |
| chr1 | 1.44E+08 | 144400000 | 0.00 | 0.00 | NO |
| chr1 | 1.44E+08 | 144500000 | 0.00 | 0.00 | NO |
| chr1 | 1.45E+08 | 144600000 | 0.00 | 0.00 | NO |
| chr1 | 1.45E+08 | 144700000 | 0.00 | 0.00 | NO |
| chr1 | 1.45E+08 | 144800000 | 0.00 | 0.00 | NO |
| chr1 | 1.45E+08 | 144900000 | 0.00 | 0.00 | NO |
| chr1 | 1.45E+08 | 145000000 | 1.09 | 1.00 | A>A |
| chr1 | 1.45E+08 | 145100000 | 1.22 | 0.98 | A>A |
| chr1 | 1.45E+08 | 145200000 | 0.00 | 0.00 | NO |
| chr1 | 1.45E+08 | 145300000 | 0.00 | 0.00 | NO |
| chr1 | 1.45E+08 | 145400000 | 0.00 | 0.00 | NO |
| chr1 | 1.45E+08 | 145500000 | 1.92 | 1.74 | A>A |
| chr1 | 1.46E+08 | 145600000 | 2.21 | 2.05 | A>A |
| chr1 | 1.46E+08 | 145700000 | 1.86 | 1.69 | A>A |
| chr1 | 1.46E+08 | 145800000 | 0.00 | 0.00 | NO |
| chr1 | 1.46E+08 | 145900000 | 0.00 | 0.00 | NO |
| chr1 | 1.46E+08 | 146000000 | 0.00 | 0.00 | NO |
| chr1 | 1.46E+08 | 146100000 | 0.00 | 0.00 | NO |
| chr1 | 1.46E+08 | 146200000 | 0.00 | 0.00 | NO |
| chr1 | 1.46E+08 | 146300000 | 0.00 | 0.00 | NO |
| chr1 | 1.46E+08 | 146400000 | 0.00 | 0.00 | NO |
| chr1 | 1.46E+08 | 146500000 | 0.00 | 0.00 | NO |
| chr1 | 1.47E+08 | 146600000 | 0.00 | -0.36 | NO |
| chr1 | 1.47E+08 | 146700000 | 0.33 | 0.01 | A>A |
| chr1 | 1.47E+08 | 146800000 | 0.20 | -0.16 | A>A |
| chr1 | 1.47E+08 | 146900000 | -0.30 | -0.36 | B>B |
| chr1 | 1.47E+08 | 147000000 | -0.15 | -0.26 | B>B |
| chr1 | 1.47E+08 | 147100000 | 0.24 | 0.11 | A>A |
| chr1 | 1.47E+08 | 147200000 | 0.16 | 0.10 | A>A |
| chr1 | 1.47E+08 | 147300000 | 0.07 | -0.22 | B>B |
| chr1 | 1.47E+08 | 147400000 | 0.22 | 0.01 | A>A |
| chr1 | 1.47E+08 | 147500000 | 0.00 | 0.00 | NO |
| chr1 | 1.48E+08 | 147600000 | 0.00 | 0.00 | NO |
| chr1 | 1.48E+08 | 147700000 | 0.00 | 0.00 | NO |
| chr1 | 1.48E+08 | 147800000 | 0.00 | 0.00 | NO |
| chr1 | 1.48E+08 | 147900000 | 0.00 | 0.00 | NO |
| chr1 | 1.48E+08 | 148000000 | 0.00 | 0.00 | NO |
| chr1 | 1.48E+08 | 148100000 | 0.00 | 0.00 | NO |
| chr1 | 1.48E+08 | 148200000 | 0.00 | 0.00 | NO |
| chr1 | 1.48E+08 | 148300000 | 0.00 | 0.00 | NO |
| chr1 | 1.48E+08 | 148400000 | 0.00 | 0.00 | NO |
| chr1 | 1.48E+08 | 148500000 | 0.00 | 0.00 | NO |
| chr1 | 1.49E+08 | 148600000 | 0.00 | 0.00 | NO |
| chr1 | 1.49E+08 | 148700000 | 0.00 | 0.00 | NO |
| chr1 | 1.49E+08 | 148800000 | 0.00 | 0.00 | NO |
| chr1 | 1.49E+08 | 148900000 | 0.00 | 0.00 | NO |
| chr1 | 1.49E+08 | 149000000 | 0.00 | 0.00 | NO |
| chr1 | 1.49E+08 | 149100000 | 0.00 | 0.00 | NO |
| chr1 | 1.49E+08 | 149200000 | 0.00 | 0.00 | NO |
| chr1 | 1.49E+08 | 149300000 | 0.00 | 0.00 | NO |
| chr1 | 1.49E+08 | 149400000 | 0.00 | 0.00 | NO |
| chr1 | 1.49E+08 | 149500000 | 0.00 | 0.00 | NO |
| chr1 | 1.5E+08 | 149600000 | 0.00 | 0.00 | NO |

| | | | | | |
|---|---|---|---|---|---|
| chr1 | 1.5E+08 | 149700000 | 0.00 | 0.00 | NO |
| chr1 | 1.5E+08 | 149800000 | 0.00 | 0.00 | NO |
| chr1 | 1.5E+08 | 149900000 | 1.31 | 1.19 | A>A |
| chr1 | 1.5E+08 | 150000000 | 1.57 | 1.27 | A>A |
| chr1 | 1.5E+08 | 150100000 | 1.69 | 1.43 | A>A |
| chr1 | 1.5E+08 | 150200000 | 1.55 | 1.35 | A>A |
| chr1 | 1.5E+08 | 150300000 | 1.68 | 1.55 | A>A |
| chr1 | 1.5E+08 | 150400000 | 1.81 | 1.67 | A>A |
| chr1 | 1.5E+08 | 150500000 | 1.75 | 1.66 | A>A |
| chr1 | 1.51E+08 | 150600000 | 1.63 | 1.59 | A>A |
| chr1 | 1.51E+08 | 150700000 | 1.77 | 1.69 | A>A |
| chr1 | 1.51E+08 | 150800000 | 1.78 | 1.75 | A>A |
| chr1 | 1.51E+08 | 150900000 | 1.67 | 1.63 | A>A |
| chr1 | 1.51E+08 | 151000000 | 1.60 | 1.57 | A>A |
| chr1 | 1.51E+08 | 151100000 | 1.66 | 1.55 | A>A |
| chr1 | 1.51E+08 | 151200000 | 1.68 | 1.47 | A>A |
| chr1 | 1.51E+08 | 151300000 | 1.61 | 1.44 | A>A |
| chr1 | 1.51E+08 | 151400000 | 1.35 | 1.30 | A>A |
| chr1 | 1.51E+08 | 151500000 | 1.22 | 1.19 | A>A |
| chr1 | 1.52E+08 | 151600000 | 1.30 | 1.13 | A>A |
| chr1 | 1.52E+08 | 151700000 | 1.15 | 0.95 | A>A |
| chr1 | 1.52E+08 | 151800000 | 0.97 | 0.91 | A>A |
| chr1 | 1.52E+08 | 151900000 | 0.75 | 0.57 | A>A |
| chr1 | 1.52E+08 | 152000000 | 0.69 | 0.45 | A>A |
| chr1 | 1.52E+08 | 152100000 | 0.20 | -0.21 | B>B |
| chr1 | 1.52E+08 | 152200000 | -0.09 | -0.41 | B>B |
| chr1 | 1.52E+08 | 152300000 | -0.48 | -0.65 | B>B |
| chr1 | 1.52E+08 | 152400000 | -0.29 | -0.64 | B>B |
| chr1 | 1.52E+08 | 152500000 | -0.08 | -0.35 | B>B |
| chr1 | 1.53E+08 | 152600000 | -0.33 | -0.61 | B>B |
| chr1 | 1.53E+08 | 152700000 | -0.41 | -0.76 | B>B |
| chr1 | 1.53E+08 | 152800000 | -0.40 | -0.86 | B>B |
| chr1 | 1.53E+08 | 152900000 | -0.39 | -0.81 | B>B |
| chr1 | 1.53E+08 | 153000000 | -0.48 | -0.79 | B>B |
| chr1 | 1.53E+08 | 153100000 | -0.50 | -0.84 | B>B |
| chr1 | 1.53E+08 | 153200000 | -0.45 | -0.76 | B>B |
| chr1 | 1.53E+08 | 153300000 | -0.22 | -0.63 | B>B |
| chr1 | 1.53E+08 | 153400000 | 0.10 | -0.38 | B>B |
| chr1 | 1.53E+08 | 153500000 | 0.42 | -0.05 | A>A |
| chr1 | 1.54E+08 | 153600000 | 0.88 | 0.66 | A>A |
| chr1 | 1.54E+08 | 153700000 | 0.96 | 0.73 | A>A |
| chr1 | 1.54E+08 | 153800000 | 0.96 | 0.75 | A>A |
| chr1 | 1.54E+08 | 153900000 | 1.34 | 1.13 | A>A |
| chr1 | 1.54E+08 | 154000000 | 1.16 | 1.08 | A>A |
| chr1 | 1.54E+08 | 154100000 | 1.17 | 1.16 | A>A |
| chr1 | 1.54E+08 | 154200000 | 1.41 | 1.23 | A>A |
| chr1 | 1.54E+08 | 154300000 | 1.37 | 1.29 | A>A |
| chr1 | 1.54E+08 | 154400000 | 1.30 | 1.45 | A>A |
| chr1 | 1.54E+08 | 154500000 | 1.43 | 1.24 | A>A |
| chr1 | 1.55E+08 | 154600000 | 1.36 | 1.20 | A>A |
| chr1 | 1.55E+08 | 154700000 | 1.50 | 1.36 | A>A |
| chr1 | 1.55E+08 | 154800000 | 1.68 | 1.57 | A>A |
| chr1 | 1.55E+08 | 154900000 | 1.53 | 1.42 | A>A |
| chr1 | 1.55E+08 | 155000000 | 1.59 | 1.69 | A>A |
| chr1 | 1.55E+08 | 155100000 | 1.61 | 1.54 | A>A |
| chr1 | 1.55E+08 | 155200000 | 1.63 | 1.71 | A>A |

| | | | | | |
|---|---|---|---|---|---|
| chr1 | 1.55E+08 | 155300000 | 1.72 | 1.83 | A>A |
| chr1 | 1.55E+08 | 155400000 | 1.84 | 1.85 | A>A |
| chr1 | 1.55E+08 | 155500000 | 2.05 | 2.10 | A>A |
| chr1 | 1.56E+08 | 155600000 | 1.68 | 1.82 | A>A |
| chr1 | 1.56E+08 | 155700000 | 1.69 | 1.75 | A>A |
| chr1 | 1.56E+08 | 155800000 | 1.93 | 1.86 | A>A |
| chr1 | 1.56E+08 | 155900000 | 2.00 | 2.00 | A>A |
| chr1 | 1.56E+08 | 156000000 | 1.92 | 2.03 | A>A |
| chr1 | 1.56E+08 | 156100000 | 1.85 | 2.02 | A>A |
| chr1 | 1.56E+08 | 156200000 | 2.00 | 2.05 | A>A |
| chr1 | 1.56E+08 | 156300000 | 1.90 | 1.84 | A>A |
| chr1 | 1.56E+08 | 156400000 | 1.83 | 1.84 | A>A |
| chr1 | 1.56E+08 | 156500000 | 1.84 | 1.86 | A>A |
| chr1 | 1.57E+08 | 156600000 | 1.56 | 1.65 | A>A |
| chr1 | 1.57E+08 | 156700000 | 1.40 | 1.28 | A>A |
| chr1 | 1.57E+08 | 156800000 | 1.50 | 1.29 | A>A |
| chr1 | 1.57E+08 | 156900000 | 1.17 | 1.07 | A>A |
| chr1 | 1.57E+08 | 157000000 | 0.90 | 0.73 | A>A |
| chr1 | 1.57E+08 | 157100000 | 0.50 | 0.17 | A>A |
| chr1 | 1.57E+08 | 157200000 | -0.04 | -0.39 | B>B |
| chr1 | 1.57E+08 | 157300000 | -0.43 | -0.74 | B>B |
| chr1 | 1.57E+08 | 157400000 | -0.50 | -0.85 | B>B |
| chr1 | 1.57E+08 | 157500000 | -0.47 | -0.76 | B>B |
| chr1 | 1.58E+08 | 157600000 | -0.61 | -0.79 | B>B |
| chr1 | 1.58E+08 | 157700000 | -0.64 | -0.85 | B>B |
| chr1 | 1.58E+08 | 157800000 | -0.62 | -0.84 | B>B |
| chr1 | 1.58E+08 | 157900000 | -0.58 | -0.77 | B>B |
| chr1 | 1.58E+08 | 158000000 | -0.11 | -0.19 | B>B |
| chr1 | 1.58E+08 | 158100000 | -0.03 | -0.09 | B>B |
| chr1 | 1.58E+08 | 158200000 | -0.45 | -0.45 | B>B |
| chr1 | 1.58E+08 | 158300000 | -0.71 | -0.76 | B>B |
| chr1 | 1.58E+08 | 158400000 | -0.74 | -0.84 | B>B |
| chr1 | 1.58E+08 | 158500000 | -0.73 | -0.83 | B>B |
| chr1 | 1.59E+08 | 158600000 | -0.73 | -0.89 | B>B |
| chr1 | 1.59E+08 | 158700000 | -0.72 | -0.94 | B>B |
| chr1 | 1.59E+08 | 158800000 | -0.74 | -0.87 | B>B |
| chr1 | 1.59E+08 | 158900000 | -0.72 | -0.83 | B>B |
| chr1 | 1.59E+08 | 159000000 | -0.60 | -0.72 | B>B |
| chr1 | 1.59E+08 | 159100000 | -0.39 | -0.62 | B>B |
| chr1 | 1.59E+08 | 159200000 | -0.06 | -0.55 | B>B |
| chr1 | 1.59E+08 | 159300000 | 0.09 | -0.72 | A>B |
| chr1 | 1.59E+08 | 159400000 | 0.05 | -0.77 | A>B |
| chr1 | 1.59E+08 | 159500000 | -0.05 | -0.77 | A>B |
| chr1 | 1.6E+08 | 159600000 | -0.06 | -0.79 | A>B |
| chr1 | 1.6E+08 | 159700000 | 0.06 | -0.74 | A>B |
| chr1 | 1.6E+08 | 159800000 | 0.20 | -0.40 | A>B |
| chr1 | 1.6E+08 | 159900000 | 0.43 | -0.10 | A>A |
| chr1 | 1.6E+08 | 160000000 | 0.61 | 0.01 | A>B |
| chr1 | 1.6E+08 | 160100000 | 0.76 | 0.42 | A>A |
| chr1 | 1.6E+08 | 160200000 | 1.06 | 0.67 | A>A |
| chr1 | 1.6E+08 | 160300000 | 1.11 | 0.82 | A>A |
| chr1 | 1.6E+08 | 160400000 | 0.91 | 0.78 | A>A |
| chr1 | 1.6E+08 | 160500000 | 1.04 | 0.78 | A>A |
| chr1 | 1.61E+08 | 160600000 | 0.97 | 0.73 | A>A |
| chr1 | 1.61E+08 | 160700000 | 0.79 | 0.64 | A>A |
| chr1 | 1.61E+08 | 160800000 | 1.21 | 0.94 | A>A |

| | | | | | |
|---|---|---|---|---|---|
| chr1 | 1.61E+08 | 160900000 | 1.33 | 1.08 | A>A |
| chr1 | 1.61E+08 | 161000000 | 1.35 | 1.23 | A>A |
| chr1 | 1.61E+08 | 161100000 | 1.56 | 1.46 | A>A |
| chr1 | 1.61E+08 | 161200000 | 1.72 | 1.50 | A>A |
| chr1 | 1.61E+08 | 161300000 | 1.81 | 1.62 | A>A |
| chr1 | 1.61E+08 | 161400000 | 1.63 | 1.40 | A>A |
| chr1 | 1.61E+08 | 161500000 | 0.00 | 0.00 | NO |
| chr1 | 1.62E+08 | 161600000 | 0.00 | 0.00 | NO |
| chr1 | 1.62E+08 | 161700000 | 0.00 | 0.00 | NO |
| chr1 | 1.62E+08 | 161800000 | 0.69 | 0.39 | A>A |
| chr1 | 1.62E+08 | 161900000 | 0.71 | 0.40 | A>A |
| chr1 | 1.62E+08 | 162000000 | 0.59 | 0.25 | A>A |
| chr1 | 1.62E+08 | 162100000 | 0.17 | -0.03 | A>A |
| chr1 | 1.62E+08 | 162200000 | 0.35 | -0.10 | A>A |
| chr1 | 1.62E+08 | 162300000 | 0.27 | -0.10 | A>A |
| chr1 | 1.62E+08 | 162400000 | 0.18 | -0.04 | A>A |
| chr1 | 1.62E+08 | 162500000 | 0.23 | 0.01 | A>A |
| chr1 | 1.63E+08 | 162600000 | 0.11 | -0.12 | B>B |
| chr1 | 1.63E+08 | 162700000 | -0.09 | -0.47 | B>B |
| chr1 | 1.63E+08 | 162800000 | -0.15 | -0.49 | B>B |
| chr1 | 1.63E+08 | 162900000 | -0.57 | -0.73 | B>B |
| chr1 | 1.63E+08 | 163000000 | -0.69 | -0.91 | B>B |
| chr1 | 1.63E+08 | 163100000 | -0.73 | -0.89 | B>B |
| chr1 | 1.63E+08 | 163200000 | -0.65 | -0.87 | B>B |
| chr1 | 1.63E+08 | 163300000 | -0.45 | -0.59 | B>B |
| chr1 | 1.63E+08 | 163400000 | -0.51 | -0.63 | B>B |
| chr1 | 1.63E+08 | 163500000 | -0.64 | -0.76 | B>B |
| chr1 | 1.64E+08 | 163600000 | -0.65 | -0.75 | B>B |
| chr1 | 1.64E+08 | 163700000 | -0.74 | -0.82 | B>B |
| chr1 | 1.64E+08 | 163800000 | -0.70 | -0.79 | B>B |
| chr1 | 1.64E+08 | 163900000 | -0.66 | -0.80 | B>B |
| chr1 | 1.64E+08 | 164000000 | -0.66 | -0.78 | B>B |
| chr1 | 1.64E+08 | 164100000 | -0.65 | -0.75 | B>B |
| chr1 | 1.64E+08 | 164200000 | -0.55 | -0.76 | B>B |
| chr1 | 1.64E+08 | 164300000 | -0.46 | -0.75 | B>B |
| chr1 | 1.64E+08 | 164400000 | -0.36 | -0.69 | B>B |
| chr1 | 1.64E+08 | 164500000 | -0.31 | -0.56 | B>B |
| chr1 | 1.65E+08 | 164600000 | -0.16 | -0.27 | B>B |
| chr1 | 1.65E+08 | 164700000 | -0.13 | -0.33 | B>B |
| chr1 | 1.65E+08 | 164800000 | -0.24 | -0.37 | B>B |
| chr1 | 1.65E+08 | 164900000 | -0.44 | -0.58 | B>B |
| chr1 | 1.65E+08 | 165000000 | -0.55 | -0.74 | B>B |
| chr1 | 1.65E+08 | 165100000 | -0.55 | -0.66 | B>B |
| chr1 | 1.65E+08 | 165200000 | -0.55 | -0.76 | B>B |
| chr1 | 1.65E+08 | 165300000 | -0.46 | -0.81 | B>B |
| chr1 | 1.65E+08 | 165400000 | -0.41 | -0.72 | B>B |
| chr1 | 1.65E+08 | 165500000 | -0.39 | -0.81 | B>B |
| chr1 | 1.66E+08 | 165600000 | -0.22 | -0.53 | B>B |
| chr1 | 1.66E+08 | 165700000 | -0.01 | -0.07 | B>B |
| chr1 | 1.66E+08 | 165800000 | 0.16 | 0.04 | A>A |
| chr1 | 1.66E+08 | 165900000 | 0.02 | -0.16 | B>B |
| chr1 | 1.66E+08 | 166000000 | -0.58 | -0.73 | B>B |
| chr1 | 1.66E+08 | 166100000 | -0.68 | -0.86 | B>B |
| chr1 | 1.66E+08 | 166200000 | -0.67 | -0.86 | B>B |
| chr1 | 1.66E+08 | 166300000 | -0.77 | -0.85 | B>B |
| chr1 | 1.66E+08 | 166400000 | -0.74 | -0.84 | B>B |

| | | | | | |
|---|---|---|---|---|---|
| chr1 | 1.66E+08 | 166500000 | -0.68 | -0.76 | B>B |
| chr1 | 1.67E+08 | 166600000 | -0.78 | -0.95 | B>B |
| chr1 | 1.67E+08 | 166700000 | -0.82 | -0.94 | B>B |
| chr1 | 1.67E+08 | 166800000 | -0.70 | -0.87 | B>B |
| chr1 | 1.67E+08 | 166900000 | -0.28 | -0.45 | B>B |
| chr1 | 1.67E+08 | 167000000 | -0.24 | -0.53 | B>B |
| chr1 | 1.67E+08 | 167100000 | -0.05 | -0.26 | B>B |
| chr1 | 1.67E+08 | 167200000 | 0.18 | -0.16 | A>A |
| chr1 | 1.67E+08 | 167300000 | 0.42 | 0.27 | A>A |
| chr1 | 1.67E+08 | 167400000 | 0.39 | 0.24 | A>A |
| chr1 | 1.67E+08 | 167500000 | 0.41 | 0.27 | A>A |
| chr1 | 1.68E+08 | 167600000 | 0.64 | 0.46 | A>A |
| chr1 | 1.68E+08 | 167700000 | 0.93 | 0.77 | A>A |
| chr1 | 1.68E+08 | 167800000 | 0.95 | 0.93 | A>A |
| chr1 | 1.68E+08 | 167900000 | 1.15 | 1.07 | A>A |
| chr1 | 1.68E+08 | 168000000 | 1.08 | 1.16 | A>A |
| chr1 | 1.68E+08 | 168100000 | 0.86 | 0.95 | A>A |
| chr1 | 1.68E+08 | 168200000 | 0.84 | 1.09 | A>A |
| chr1 | 1.68E+08 | 168300000 | 0.69 | 1.02 | A>A |
| chr1 | 1.68E+08 | 168400000 | 0.52 | 0.76 | A>A |
| chr1 | 1.68E+08 | 168500000 | 0.34 | 0.59 | A>A |
| chr1 | 1.69E+08 | 168600000 | 0.23 | 0.50 | A>A |
| chr1 | 1.69E+08 | 168700000 | 0.43 | 0.66 | A>A |
| chr1 | 1.69E+08 | 168800000 | 0.40 | 0.79 | A>A |
| chr1 | 1.69E+08 | 168900000 | 0.50 | 0.78 | A>A |
| chr1 | 1.69E+08 | 169000000 | 0.60 | 0.80 | A>A |
| chr1 | 1.69E+08 | 169100000 | 0.50 | 0.73 | A>A |
| chr1 | 1.69E+08 | 169200000 | 0.43 | 0.67 | A>A |
| chr1 | 1.69E+08 | 169300000 | 0.40 | 0.60 | A>A |
| chr1 | 1.69E+08 | 169400000 | 0.13 | 0.27 | A>A |
| chr1 | 1.69E+08 | 169500000 | -0.01 | 0.15 | A>A |
| chr1 | 1.7E+08 | 169600000 | -0.32 | -0.13 | B>B |
| chr1 | 1.7E+08 | 169700000 | -0.42 | -0.27 | B>B |
| chr1 | 1.7E+08 | 169800000 | -0.25 | -0.23 | B>B |
| chr1 | 1.7E+08 | 169900000 | -0.29 | -0.27 | B>B |
| chr1 | 1.7E+08 | 170000000 | -0.47 | -0.42 | B>B |
| chr1 | 1.7E+08 | 170100000 | -0.53 | -0.60 | B>B |
| chr1 | 1.7E+08 | 170200000 | -0.75 | -0.89 | B>B |
| chr1 | 1.7E+08 | 170300000 | -0.80 | -0.91 | B>B |
| chr1 | 1.7E+08 | 170400000 | -0.71 | -0.81 | B>B |
| chr1 | 1.7E+08 | 170500000 | -0.71 | -0.76 | B>B |
| chr1 | 1.71E+08 | 170600000 | -0.59 | -0.57 | B>B |
| chr1 | 1.71E+08 | 170700000 | -0.56 | -0.67 | B>B |
| chr1 | 1.71E+08 | 170800000 | -0.78 | -0.81 | B>B |
| chr1 | 1.71E+08 | 170900000 | -0.69 | -0.75 | B>B |
| chr1 | 1.71E+08 | 171000000 | -0.71 | -0.83 | B>B |
| chr1 | 1.71E+08 | 171100000 | -0.75 | -0.92 | B>B |
| chr1 | 1.71E+08 | 171200000 | -0.70 | -0.93 | B>B |
| chr1 | 1.71E+08 | 171300000 | -0.50 | -0.68 | B>B |
| chr1 | 1.71E+08 | 171400000 | -0.24 | -0.28 | B>B |
| chr1 | 1.71E+08 | 171500000 | 0.12 | 0.19 | A>A |
| chr1 | 1.72E+08 | 171600000 | 0.34 | 0.46 | A>A |
| chr1 | 1.72E+08 | 171700000 | 0.42 | 0.46 | A>A |
| chr1 | 1.72E+08 | 171800000 | 0.57 | 0.67 | A>A |
| chr1 | 1.72E+08 | 171900000 | 0.64 | 0.84 | A>A |
| chr1 | 1.72E+08 | 172000000 | 0.40 | 0.64 | A>A |

| | | | | | |
|---|---|---|---|---|---|
| chr1 | 1.72E+08 | 172100000 | 0.59 | 0.84 | A>A |
| chr1 | 1.72E+08 | 172200000 | 0.76 | 1.11 | A>A |
| chr1 | 1.72E+08 | 172300000 | 0.80 | 1.16 | A>A |
| chr1 | 1.72E+08 | 172400000 | 0.59 | 0.92 | A>A |
| chr1 | 1.72E+08 | 172500000 | 0.71 | 0.89 | A>A |
| chr1 | 1.73E+08 | 172600000 | 0.65 | 0.79 | A>A |
| chr1 | 1.73E+08 | 172700000 | 0.56 | 0.72 | A>A |
| chr1 | 1.73E+08 | 172800000 | 0.19 | 0.31 | A>A |
| chr1 | 1.73E+08 | 172900000 | 0.68 | 0.77 | A>A |
| chr1 | 1.73E+08 | 173000000 | 0.63 | 0.87 | A>A |
| chr1 | 1.73E+08 | 173100000 | 0.73 | 0.86 | A>A |
| chr1 | 1.73E+08 | 173200000 | 0.39 | 0.55 | A>A |
| chr1 | 1.73E+08 | 173300000 | 0.30 | 0.31 | A>A |
| chr1 | 1.73E+08 | 173400000 | 0.24 | 0.35 | A>A |
| chr1 | 1.73E+08 | 173500000 | 0.33 | 0.37 | A>A |
| chr1 | 1.74E+08 | 173600000 | 0.39 | 0.35 | A>A |
| chr1 | 1.74E+08 | 173700000 | 0.40 | 0.29 | A>A |
| chr1 | 1.74E+08 | 173800000 | 0.41 | 0.50 | A>A |
| chr1 | 1.74E+08 | 173900000 | 0.65 | 0.73 | A>A |
| chr1 | 1.74E+08 | 174000000 | 0.53 | 0.69 | A>A |
| chr1 | 1.74E+08 | 174100000 | 0.41 | 0.47 | A>A |
| chr1 | 1.74E+08 | 174200000 | 0.08 | 0.27 | A>A |
| chr1 | 1.74E+08 | 174300000 | -0.25 | -0.09 | B>B |
| chr1 | 1.74E+08 | 174400000 | -0.33 | -0.18 | B>B |
| chr1 | 1.74E+08 | 174500000 | -0.47 | -0.27 | B>B |
| chr1 | 1.75E+08 | 174600000 | -0.63 | -0.33 | B>B |
| chr1 | 1.75E+08 | 174700000 | -0.46 | -0.26 | B>B |
| chr1 | 1.75E+08 | 174800000 | -0.25 | -0.11 | B>B |
| chr1 | 1.75E+08 | 174900000 | -0.29 | 0.04 | B>B |
| chr1 | 1.75E+08 | 175000000 | -0.13 | -0.07 | B>B |
| chr1 | 1.75E+08 | 175100000 | -0.31 | -0.60 | B>B |
| chr1 | 1.75E+08 | 175200000 | -0.54 | -0.61 | B>B |
| chr1 | 1.75E+08 | 175300000 | -0.63 | -0.75 | B>B |
| chr1 | 1.75E+08 | 175400000 | -0.64 | -0.83 | B>B |
| chr1 | 1.75E+08 | 175500000 | -0.61 | -0.87 | B>B |
| chr1 | 1.76E+08 | 175600000 | -0.68 | -0.87 | B>B |
| chr1 | 1.76E+08 | 175700000 | -0.57 | -0.80 | B>B |
| chr1 | 1.76E+08 | 175800000 | -0.55 | -0.74 | B>B |
| chr1 | 1.76E+08 | 175900000 | -0.36 | -0.57 | B>B |
| chr1 | 1.76E+08 | 176000000 | -0.17 | -0.26 | B>B |
| chr1 | 1.76E+08 | 176100000 | -0.13 | -0.28 | B>B |
| chr1 | 1.76E+08 | 176200000 | -0.18 | -0.35 | B>B |
| chr1 | 1.76E+08 | 176300000 | -0.66 | -0.74 | B>B |
| chr1 | 1.76E+08 | 176400000 | -0.66 | -0.76 | B>B |
| chr1 | 1.76E+08 | 176500000 | -0.67 | -0.89 | B>B |
| chr1 | 1.77E+08 | 176600000 | -0.66 | -0.86 | B>B |
| chr1 | 1.77E+08 | 176700000 | -0.76 | -0.84 | B>B |
| chr1 | 1.77E+08 | 176800000 | -0.74 | -0.81 | B>B |
| chr1 | 1.77E+08 | 176900000 | -0.58 | -0.64 | B>B |
| chr1 | 1.77E+08 | 177000000 | -0.59 | -0.79 | B>B |
| chr1 | 1.77E+08 | 177100000 | -0.68 | -0.83 | B>B |
| chr1 | 1.77E+08 | 177200000 | -0.50 | -0.62 | B>B |
| chr1 | 1.77E+08 | 177300000 | -0.56 | -0.70 | B>B |
| chr1 | 1.77E+08 | 177400000 | -0.62 | -0.75 | B>B |
| chr1 | 1.77E+08 | 177500000 | -0.70 | -0.78 | B>B |
| chr1 | 1.78E+08 | 177600000 | -0.70 | -0.80 | B>B |

| | | | | | |
|---|---|---|---|---|---|
| chr1 | 1.78E+08 | 177700000 | -0.77 | -0.88 | B>B |
| chr1 | 1.78E+08 | 177800000 | -0.73 | -0.84 | B>B |
| chr1 | 1.78E+08 | 177900000 | -0.58 | -0.73 | B>B |
| chr1 | 1.78E+08 | 178000000 | -0.34 | -0.28 | B>B |
| chr1 | 1.78E+08 | 178100000 | 0.19 | 0.17 | A>A |
| chr1 | 1.78E+08 | 178200000 | 0.29 | 0.25 | A>A |
| chr1 | 1.78E+08 | 178300000 | 0.12 | 0.12 | A>A |
| chr1 | 1.78E+08 | 178400000 | 0.21 | 0.21 | A>A |
| chr1 | 1.78E+08 | 178500000 | 0.18 | 0.23 | A>A |
| chr1 | 1.79E+08 | 178600000 | 0.21 | 0.25 | A>A |
| chr1 | 1.79E+08 | 178700000 | 0.34 | 0.34 | A>A |
| chr1 | 1.79E+08 | 178800000 | 0.32 | 0.27 | A>A |
| chr1 | 1.79E+08 | 178900000 | 0.19 | 0.36 | A>A |
| chr1 | 1.79E+08 | 179000000 | 0.27 | 0.33 | A>A |
| chr1 | 1.79E+08 | 179100000 | 0.38 | 0.50 | A>A |
| chr1 | 1.79E+08 | 179200000 | 0.65 | 0.68 | A>A |
| chr1 | 1.79E+08 | 179300000 | 0.49 | 0.49 | A>A |
| chr1 | 1.79E+08 | 179400000 | 0.06 | 0.02 | A>A |
| chr1 | 1.79E+08 | 179500000 | -0.28 | -0.29 | B>B |
| chr1 | 1.8E+08 | 179600000 | -0.28 | -0.33 | B>B |
| chr1 | 1.8E+08 | 179700000 | -0.18 | -0.20 | B>B |
| chr1 | 1.8E+08 | 179800000 | -0.05 | -0.13 | B>B |
| chr1 | 1.8E+08 | 179900000 | 0.53 | 0.56 | A>A |
| chr1 | 1.8E+08 | 180000000 | 0.76 | 0.73 | A>A |
| chr1 | 1.8E+08 | 180100000 | 0.76 | 0.76 | A>A |
| chr1 | 1.8E+08 | 180200000 | 0.54 | 0.57 | A>A |
| chr1 | 1.8E+08 | 180300000 | 0.58 | 0.75 | A>A |
| chr1 | 1.8E+08 | 180400000 | 0.61 | 0.69 | A>A |
| chr1 | 1.8E+08 | 180500000 | 0.46 | 0.41 | A>A |
| chr1 | 1.81E+08 | 180600000 | 0.64 | 0.63 | A>A |
| chr1 | 1.81E+08 | 180700000 | 0.68 | 0.60 | A>A |
| chr1 | 1.81E+08 | 180800000 | 0.55 | 0.42 | A>A |
| chr1 | 1.81E+08 | 180900000 | 0.25 | 0.15 | A>A |
| chr1 | 1.81E+08 | 181000000 | 0.46 | 0.29 | A>A |
| chr1 | 1.81E+08 | 181100000 | 0.72 | 0.41 | A>A |
| chr1 | 1.81E+08 | 181200000 | -0.04 | -0.15 | B>B |
| chr1 | 1.81E+08 | 181300000 | -0.18 | -0.42 | B>B |
| chr1 | 1.81E+08 | 181400000 | -0.26 | -0.48 | B>B |
| chr1 | 1.81E+08 | 181500000 | -0.41 | -0.62 | B>B |
| chr1 | 1.82E+08 | 181600000 | -0.61 | -0.91 | B>B |
| chr1 | 1.82E+08 | 181700000 | -0.66 | -0.98 | B>B |
| chr1 | 1.82E+08 | 181800000 | -0.67 | -0.96 | B>B |
| chr1 | 1.82E+08 | 181900000 | -0.70 | -0.91 | B>B |
| chr1 | 1.82E+08 | 182000000 | -0.59 | -0.82 | B>B |
| chr1 | 1.82E+08 | 182100000 | -0.28 | -0.50 | B>B |
| chr1 | 1.82E+08 | 182200000 | -0.08 | -0.55 | B>B |
| chr1 | 1.82E+08 | 182300000 | -0.03 | -0.57 | B>B |
| chr1 | 1.82E+08 | 182400000 | -0.17 | -0.56 | B>B |
| chr1 | 1.82E+08 | 182500000 | -0.37 | -0.80 | B>B |
| chr1 | 1.83E+08 | 182600000 | -0.20 | -0.68 | B>B |
| chr1 | 1.83E+08 | 182700000 | -0.11 | -0.88 | A>B |
| chr1 | 1.83E+08 | 182800000 | 0.08 | -0.52 | A>B |
| chr1 | 1.83E+08 | 182900000 | 0.45 | 0.12 | A>A |
| chr1 | 1.83E+08 | 183000000 | 0.70 | 0.33 | A>A |
| chr1 | 1.83E+08 | 183100000 | 1.06 | 0.73 | A>A |
| chr1 | 1.83E+08 | 183200000 | 1.00 | 0.70 | A>A |

| | | | | | |
|---|---|---|---|---|---|
| chr1 | 1.83E+08 | 183300000 | 0.62 | 0.24 | A>A |
| chr1 | 1.83E+08 | 183400000 | 0.57 | 0.26 | A>A |
| chr1 | 1.83E+08 | 183500000 | 0.67 | 0.33 | A>A |
| chr1 | 1.84E+08 | 183600000 | 0.47 | 0.09 | A>A |
| chr1 | 1.84E+08 | 183700000 | 0.18 | -0.32 | B>B |
| chr1 | 1.84E+08 | 183800000 | 0.30 | -0.32 | A>B |
| chr1 | 1.84E+08 | 183900000 | 0.15 | -0.37 | B>B |
| chr1 | 1.84E+08 | 184000000 | -0.09 | -0.83 | A>B |
| chr1 | 1.84E+08 | 184100000 | -0.17 | -0.74 | B>B |
| chr1 | 1.84E+08 | 184200000 | -0.42 | -0.90 | B>B |
| chr1 | 1.84E+08 | 184300000 | -0.41 | -0.90 | B>B |
| chr1 | 1.84E+08 | 184400000 | -0.36 | -0.89 | B>B |
| chr1 | 1.84E+08 | 184500000 | -0.40 | -0.93 | B>B |
| chr1 | 1.85E+08 | 184600000 | -0.42 | -0.92 | B>B |
| chr1 | 1.85E+08 | 184700000 | -0.22 | -0.67 | B>B |
| chr1 | 1.85E+08 | 184800000 | -0.11 | -0.41 | B>B |
| chr1 | 1.85E+08 | 184900000 | -0.35 | -0.63 | B>B |
| chr1 | 1.85E+08 | 185000000 | -0.21 | -0.59 | B>B |
| chr1 | 1.85E+08 | 185100000 | 0.00 | -0.21 | B>B |
| chr1 | 1.85E+08 | 185200000 | 0.01 | -0.04 | B>B |
| chr1 | 1.85E+08 | 185300000 | 0.05 | 0.20 | A>A |
| chr1 | 1.85E+08 | 185400000 | 0.23 | 0.50 | A>A |
| chr1 | 1.85E+08 | 185500000 | 0.24 | 0.53 | A>A |
| chr1 | 1.86E+08 | 185600000 | 0.29 | 0.47 | A>A |
| chr1 | 1.86E+08 | 185700000 | -0.05 | 0.05 | A>A |
| chr1 | 1.86E+08 | 185800000 | -0.39 | -0.15 | B>B |
| chr1 | 1.86E+08 | 185900000 | -0.34 | -0.04 | B>B |
| chr1 | 1.86E+08 | 186000000 | -0.40 | -0.08 | B>B |
| chr1 | 1.86E+08 | 186100000 | -0.45 | -0.03 | B>B |
| chr1 | 1.86E+08 | 186200000 | -0.28 | -0.03 | B>B |
| chr1 | 1.86E+08 | 186300000 | -0.05 | 0.19 | A>A |
| chr1 | 1.86E+08 | 186400000 | 0.24 | 0.48 | A>A |
| chr1 | 1.86E+08 | 186500000 | 0.23 | 0.46 | A>A |
| chr1 | 1.87E+08 | 186600000 | 0.31 | 0.51 | A>A |
| chr1 | 1.87E+08 | 186700000 | 0.36 | 0.54 | A>A |
| chr1 | 1.87E+08 | 186800000 | 0.44 | 0.71 | A>A |
| chr1 | 1.87E+08 | 186900000 | 0.25 | 0.50 | A>A |
| chr1 | 1.87E+08 | 187000000 | -0.05 | 0.17 | A>A |
| chr1 | 1.87E+08 | 187100000 | -0.38 | -0.17 | B>B |
| chr1 | 1.87E+08 | 187200000 | -0.47 | -0.19 | B>B |
| chr1 | 1.87E+08 | 187300000 | -0.46 | -0.24 | B>B |
| chr1 | 1.87E+08 | 187400000 | -0.49 | -0.21 | B>B |
| chr1 | 1.87E+08 | 187500000 | -0.48 | -0.15 | B>B |
| chr1 | 1.88E+08 | 187600000 | -0.51 | -0.08 | B>B |
| chr1 | 1.88E+08 | 187700000 | -0.45 | -0.10 | B>B |
| chr1 | 1.88E+08 | 187800000 | -0.50 | -0.12 | B>B |
| chr1 | 1.88E+08 | 187900000 | -0.50 | -0.09 | B>B |
| chr1 | 1.88E+08 | 188000000 | -0.40 | 0.02 | B>B |
| chr1 | 1.88E+08 | 188100000 | -0.43 | -0.07 | B>B |
| chr1 | 1.88E+08 | 188200000 | -0.42 | -0.11 | B>B |
| chr1 | 1.88E+08 | 188300000 | -0.48 | -0.07 | B>B |
| chr1 | 1.88E+08 | 188400000 | -0.44 | -0.02 | B>B |
| chr1 | 1.88E+08 | 188500000 | -0.43 | -0.07 | B>B |
| chr1 | 1.89E+08 | 188600000 | -0.49 | -0.02 | B>B |
| chr1 | 1.89E+08 | 188700000 | -0.45 | 0.05 | B>B |
| chr1 | 1.89E+08 | 188800000 | -0.48 | -0.05 | B>B |

| | | | | | |
|---|---|---|---|---|---|
| chr1 | 1.89E+08 | 188900000 | -0.45 | -0.01 | B>B |
| chr1 | 1.89E+08 | 189000000 | -0.39 | 0.00 | B>B |
| chr1 | 1.89E+08 | 189100000 | -0.37 | -0.04 | B>B |
| chr1 | 1.89E+08 | 189200000 | -0.43 | -0.03 | B>B |
| chr1 | 1.89E+08 | 189300000 | -0.35 | 0.06 | B>B |
| chr1 | 1.89E+08 | 189400000 | -0.26 | 0.12 | B>B |
| chr1 | 1.89E+08 | 189500000 | -0.13 | 0.18 | A>A |
| chr1 | 1.9E+08 | 189600000 | -0.10 | 0.22 | A>A |
| chr1 | 1.9E+08 | 189700000 | -0.12 | 0.22 | A>A |
| chr1 | 1.9E+08 | 189800000 | -0.08 | 0.24 | A>A |
| chr1 | 1.9E+08 | 189900000 | -0.17 | 0.19 | A>A |
| chr1 | 1.9E+08 | 190000000 | -0.18 | 0.24 | A>A |
| chr1 | 1.9E+08 | 190100000 | -0.19 | 0.22 | A>A |
| chr1 | 1.9E+08 | 190200000 | -0.21 | 0.26 | A>A |
| chr1 | 1.9E+08 | 190300000 | -0.35 | 0.02 | B>B |
| chr1 | 1.9E+08 | 190400000 | -0.45 | -0.15 | B>B |
| chr1 | 1.9E+08 | 190500000 | -0.57 | -0.36 | B>B |
| chr1 | 1.91E+08 | 190600000 | -0.70 | -0.51 | B>B |
| chr1 | 1.91E+08 | 190700000 | -0.70 | -0.49 | B>B |
| chr1 | 1.91E+08 | 190800000 | -0.75 | -0.46 | B>B |
| chr1 | 1.91E+08 | 190900000 | -0.70 | -0.50 | B>B |
| chr1 | 1.91E+08 | 191000000 | -0.72 | -0.39 | B>B |
| chr1 | 1.91E+08 | 191100000 | -0.65 | -0.29 | B>B |
| chr1 | 1.91E+08 | 191200000 | -0.66 | -0.21 | B>B |
| chr1 | 1.91E+08 | 191300000 | -0.64 | -0.15 | B>B |
| chr1 | 1.91E+08 | 191400000 | -0.66 | -0.13 | B>A |
| chr1 | 1.91E+08 | 191500000 | -0.65 | -0.09 | B>A |
| chr1 | 1.92E+08 | 191600000 | -0.65 | -0.17 | B>B |
| chr1 | 1.92E+08 | 191700000 | -0.67 | -0.20 | B>B |
| chr1 | 1.92E+08 | 191800000 | -0.70 | -0.31 | B>B |
| chr1 | 1.92E+08 | 191900000 | -0.67 | -0.41 | B>B |
| chr1 | 1.92E+08 | 192000000 | -0.66 | -0.52 | B>B |
| chr1 | 1.92E+08 | 192100000 | -0.71 | -0.47 | B>B |
| chr1 | 1.92E+08 | 192200000 | -0.74 | -0.46 | B>B |
| chr1 | 1.92E+08 | 192300000 | -0.69 | -0.48 | B>B |
| chr1 | 1.92E+08 | 192400000 | -0.58 | -0.42 | B>B |
| chr1 | 1.92E+08 | 192500000 | -0.34 | -0.06 | B>B |
| chr1 | 1.93E+08 | 192600000 | -0.16 | 0.10 | B>B |
| chr1 | 1.93E+08 | 192700000 | -0.04 | 0.30 | A>A |
| chr1 | 1.93E+08 | 192800000 | -0.03 | 0.20 | A>A |
| chr1 | 1.93E+08 | 192900000 | 0.01 | 0.22 | A>A |
| chr1 | 1.93E+08 | 193000000 | 0.00 | 0.23 | A>A |
| chr1 | 1.93E+08 | 193100000 | 0.10 | 0.14 | A>A |
| chr1 | 1.93E+08 | 193200000 | -0.10 | 0.02 | B>B |
| chr1 | 1.93E+08 | 193300000 | -0.16 | 0.08 | B>B |
| chr1 | 1.93E+08 | 193400000 | -0.18 | 0.01 | B>B |
| chr1 | 1.93E+08 | 193500000 | -0.15 | 0.17 | A>A |
| chr1 | 1.94E+08 | 193600000 | -0.15 | 0.14 | B>B |
| chr1 | 1.94E+08 | 193700000 | -0.13 | 0.18 | A>A |
| chr1 | 1.94E+08 | 193800000 | -0.14 | 0.02 | B>B |
| chr1 | 1.94E+08 | 193900000 | -0.21 | -0.06 | B>B |
| chr1 | 1.94E+08 | 194000000 | -0.27 | -0.03 | B>B |
| chr1 | 1.94E+08 | 194100000 | -0.29 | -0.14 | B>B |
| chr1 | 1.94E+08 | 194200000 | -0.38 | -0.15 | B>B |
| chr1 | 1.94E+08 | 194300000 | -0.46 | -0.19 | B>B |
| chr1 | 1.94E+08 | 194400000 | -0.55 | -0.35 | B>B |

| | | | | | |
|---|---|---|---|---|---|
| chr1 | 1.94E+08 | 194500000 | -0.66 | -0.47 | B>B |
| chr1 | 1.95E+08 | 194600000 | -0.66 | -0.60 | B>B |
| chr1 | 1.95E+08 | 194700000 | -0.68 | -0.63 | B>B |
| chr1 | 1.95E+08 | 194800000 | -0.71 | -0.66 | B>B |
| chr1 | 1.95E+08 | 194900000 | -0.74 | -0.67 | B>B |
| chr1 | 1.95E+08 | 195000000 | -0.70 | -0.68 | B>B |
| chr1 | 1.95E+08 | 195100000 | -0.68 | -0.67 | B>B |
| chr1 | 1.95E+08 | 195200000 | -0.71 | -0.69 | B>B |
| chr1 | 1.95E+08 | 195300000 | -0.72 | -0.69 | B>B |
| chr1 | 1.95E+08 | 195400000 | -0.73 | -0.68 | B>B |
| chr1 | 1.95E+08 | 195500000 | -0.72 | -0.65 | B>B |
| chr1 | 1.96E+08 | 195600000 | -0.73 | -0.66 | B>B |
| chr1 | 1.96E+08 | 195700000 | -0.77 | -0.65 | B>B |
| chr1 | 1.96E+08 | 195800000 | -0.69 | -0.58 | B>B |
| chr1 | 1.96E+08 | 195900000 | -0.72 | -0.72 | B>B |
| chr1 | 1.96E+08 | 196000000 | -0.79 | -0.70 | B>B |
| chr1 | 1.96E+08 | 196100000 | -0.71 | -0.50 | B>B |
| chr1 | 1.96E+08 | 196200000 | -0.55 | -0.32 | B>B |
| chr1 | 1.96E+08 | 196300000 | -0.60 | -0.27 | B>B |
| chr1 | 1.96E+08 | 196400000 | -0.43 | 0.04 | B>B |
| chr1 | 1.96E+08 | 196500000 | -0.44 | 0.16 | B>A |
| chr1 | 1.97E+08 | 196600000 | -0.38 | 0.19 | B>A |
| chr1 | 1.97E+08 | 196700000 | -0.46 | 0.01 | B>B |
| chr1 | 1.97E+08 | 196800000 | 0.00 | 0.00 | NO |
| chr1 | 1.97E+08 | 196900000 | -0.53 | 0.13 | B>A |
| chr1 | 1.97E+08 | 197000000 | -0.40 | 0.16 | B>A |
| chr1 | 1.97E+08 | 197100000 | -0.19 | 0.27 | A>A |
| chr1 | 1.97E+08 | 197200000 | -0.19 | 0.28 | A>A |
| chr1 | 1.97E+08 | 197300000 | -0.43 | 0.26 | B>A |
| chr1 | 1.97E+08 | 197400000 | -0.52 | 0.00 | B>A |
| chr1 | 1.97E+08 | 197500000 | -0.62 | -0.02 | B>A |
| chr1 | 1.98E+08 | 197600000 | -0.42 | 0.14 | B>A |
| chr1 | 1.98E+08 | 197700000 | -0.26 | 0.19 | B>B |
| chr1 | 1.98E+08 | 197800000 | -0.29 | 0.27 | B>A |
| chr1 | 1.98E+08 | 197900000 | -0.36 | 0.09 | B>B |
| chr1 | 1.98E+08 | 198000000 | -0.34 | 0.17 | B>B |
| chr1 | 1.98E+08 | 198100000 | -0.36 | 0.14 | B>B |
| chr1 | 1.98E+08 | 198200000 | -0.18 | 0.34 | B>A |
| chr1 | 1.98E+08 | 198300000 | -0.29 | 0.16 | B>B |
| chr1 | 1.98E+08 | 198400000 | -0.49 | -0.01 | B>B |
| chr1 | 1.98E+08 | 198500000 | -0.37 | 0.08 | B>B |
| chr1 | 1.99E+08 | 198600000 | -0.38 | 0.21 | B>A |
| chr1 | 1.99E+08 | 198700000 | -0.42 | 0.11 | B>A |
| chr1 | 1.99E+08 | 198800000 | -0.53 | -0.05 | B>B |
| chr1 | 1.99E+08 | 198900000 | -0.55 | 0.19 | B>A |
| chr1 | 1.99E+08 | 199000000 | -0.55 | 0.16 | B>A |
| chr1 | 1.99E+08 | 199100000 | -0.68 | -0.09 | B>A |
| chr1 | 1.99E+08 | 199200000 | -0.62 | -0.05 | B>A |
| chr1 | 1.99E+08 | 199300000 | -0.70 | -0.04 | B>A |
| chr1 | 1.99E+08 | 199400000 | -0.76 | -0.07 | B>A |
| chr1 | 1.99E+08 | 199500000 | -0.60 | -0.04 | B>A |
| chr1 | 2E+08 | 199600000 | -0.35 | 0.18 | B>A |
| chr1 | 2E+08 | 199700000 | 0.01 | 0.55 | B>A |
| chr1 | 2E+08 | 199800000 | 0.30 | 0.86 | B>A |
| chr1 | 2E+08 | 199900000 | 0.38 | 0.86 | A>A |
| chr1 | 2E+08 | 200000000 | 0.37 | 0.87 | A>A |

| | | | | | |
|---|---|---|---|---|---|
| chr1 | 2E+08 | 200100000 | 0.35 | 0.68 | A>A |
| chr1 | 2E+08 | 200200000 | 0.27 | 0.64 | A>A |
| chr1 | 2E+08 | 200300000 | 0.36 | 0.71 | A>A |
| chr1 | 2E+08 | 200400000 | 0.54 | 0.98 | A>A |
| chr1 | 2E+08 | 200500000 | 0.26 | 0.76 | A>A |
| chr1 | 2.01E+08 | 200600000 | 0.53 | 0.85 | A>A |
| chr1 | 2.01E+08 | 200700000 | 1.00 | 1.31 | A>A |
| chr1 | 2.01E+08 | 200800000 | 0.98 | 1.17 | A>A |
| chr1 | 2.01E+08 | 200900000 | 1.04 | 1.28 | A>A |
| chr1 | 2.01E+08 | 201000000 | 1.17 | 1.29 | A>A |
| chr1 | 2.01E+08 | 201100000 | 1.03 | 1.19 | A>A |
| chr1 | 2.01E+08 | 201200000 | 1.28 | 1.30 | A>A |
| chr1 | 2.01E+08 | 201300000 | 1.44 | 1.37 | A>A |
| chr1 | 2.01E+08 | 201400000 | 1.37 | 1.28 | A>A |
| chr1 | 2.01E+08 | 201500000 | 1.34 | 1.41 | A>A |
| chr1 | 2.02E+08 | 201600000 | 1.36 | 1.43 | A>A |
| chr1 | 2.02E+08 | 201700000 | 1.50 | 1.46 | A>A |
| chr1 | 2.02E+08 | 201800000 | 1.48 | 1.30 | A>A |
| chr1 | 2.02E+08 | 201900000 | 1.47 | 1.39 | A>A |
| chr1 | 2.02E+08 | 202000000 | 1.75 | 1.65 | A>A |
| chr1 | 2.02E+08 | 202100000 | 1.94 | 1.94 | A>A |
| chr1 | 2.02E+08 | 202200000 | 1.54 | 1.37 | A>A |
| chr1 | 2.02E+08 | 202300000 | 1.52 | 1.51 | A>A |
| chr1 | 2.02E+08 | 202400000 | 1.38 | 1.26 | A>A |
| chr1 | 2.02E+08 | 202500000 | 0.00 | 0.00 | NO |
| chr1 | 2.03E+08 | 202600000 | 1.19 | 1.16 | A>A |
| chr1 | 2.03E+08 | 202700000 | 1.21 | 1.31 | A>A |
| chr1 | 2.03E+08 | 202800000 | 1.40 | 1.39 | A>A |
| chr1 | 2.03E+08 | 202900000 | 1.46 | 1.52 | A>A |
| chr1 | 2.03E+08 | 203000000 | 1.40 | 1.17 | A>A |
| chr1 | 2.03E+08 | 203100000 | 1.16 | 0.98 | A>A |
| chr1 | 2.03E+08 | 203200000 | 1.34 | 1.09 | A>A |
| chr1 | 2.03E+08 | 203300000 | 1.09 | 0.89 | A>A |
| chr1 | 2.03E+08 | 203400000 | 1.11 | 0.88 | A>A |
| chr1 | 2.03E+08 | 203500000 | 1.02 | 0.85 | A>A |
| chr1 | 2.04E+08 | 203600000 | 1.18 | 1.04 | A>A |
| chr1 | 2.04E+08 | 203700000 | 1.12 | 0.92 | A>A |
| chr1 | 2.04E+08 | 203800000 | 1.13 | 0.97 | A>A |
| chr1 | 2.04E+08 | 203900000 | 1.07 | 1.09 | A>A |
| chr1 | 2.04E+08 | 204000000 | 1.39 | 1.29 | A>A |
| chr1 | 2.04E+08 | 204100000 | 1.30 | 1.19 | A>A |
| chr1 | 2.04E+08 | 204200000 | 1.00 | 0.99 | A>A |
| chr1 | 2.04E+08 | 204300000 | 1.38 | 1.42 | A>A |
| chr1 | 2.04E+08 | 204400000 | 1.26 | 1.26 | A>A |
| chr1 | 2.04E+08 | 204500000 | 1.34 | 1.45 | A>A |
| chr1 | 2.05E+08 | 204600000 | 1.05 | 1.07 | A>A |
| chr1 | 2.05E+08 | 204700000 | 1.36 | 1.37 | A>A |
| chr1 | 2.05E+08 | 204800000 | 1.01 | 1.00 | A>A |
| chr1 | 2.05E+08 | 204900000 | 1.16 | 1.07 | A>A |
| chr1 | 2.05E+08 | 205000000 | 1.10 | 0.99 | A>A |
| chr1 | 2.05E+08 | 205100000 | 1.27 | 1.17 | A>A |
| chr1 | 2.05E+08 | 205200000 | 1.39 | 1.34 | A>A |
| chr1 | 2.05E+08 | 205300000 | 1.42 | 1.33 | A>A |
| chr1 | 2.05E+08 | 205400000 | 1.34 | 1.32 | A>A |
| chr1 | 2.05E+08 | 205500000 | 1.33 | 1.36 | A>A |
| chr1 | 2.06E+08 | 205600000 | 1.49 | 1.42 | A>A |

| | | | | | |
|---|---|---|---|---|---|
| chr1 | 2.06E+08 | 205700000 | 1.39 | 1.40 | A>A |
| chr1 | 2.06E+08 | 205800000 | 1.45 | 1.31 | A>A |
| chr1 | 2.06E+08 | 205900000 | 1.44 | 1.28 | A>A |
| chr1 | 2.06E+08 | 206000000 | 0.00 | 0.00 | NO |
| chr1 | 2.06E+08 | 206100000 | 0.00 | 0.00 | NO |
| chr1 | 2.06E+08 | 206200000 | 0.00 | 0.00 | NO |
| chr1 | 2.06E+08 | 206300000 | 1.14 | 1.10 | A>A |
| chr1 | 2.06E+08 | 206400000 | 0.00 | 0.00 | NO |
| chr1 | 2.06E+08 | 206500000 | 0.00 | 0.00 | NO |
| chr1 | 2.07E+08 | 206600000 | 0.00 | 0.00 | NO |
| chr1 | 2.07E+08 | 206700000 | 1.24 | 1.24 | A>A |
| chr1 | 2.07E+08 | 206800000 | 1.25 | 1.17 | A>A |
| chr1 | 2.07E+08 | 206900000 | 1.03 | 1.04 | A>A |
| chr1 | 2.07E+08 | 207000000 | 0.86 | 0.78 | A>A |
| chr1 | 2.07E+08 | 207100000 | 0.95 | 0.75 | A>A |
| chr1 | 2.07E+08 | 207200000 | 0.82 | 0.86 | A>A |
| chr1 | 2.07E+08 | 207300000 | 0.52 | 0.72 | A>A |
| chr1 | 2.07E+08 | 207400000 | 0.44 | 0.67 | A>A |
| chr1 | 2.07E+08 | 207500000 | 0.59 | 0.88 | A>A |
| chr1 | 2.08E+08 | 207600000 | 0.46 | 0.50 | A>A |
| chr1 | 2.08E+08 | 207700000 | 0.31 | 0.29 | A>A |
| chr1 | 2.08E+08 | 207800000 | 0.00 | 0.00 | NO |
| chr1 | 2.08E+08 | 207900000 | 0.38 | 0.20 | A>A |
| chr1 | 2.08E+08 | 208000000 | 0.49 | 0.41 | A>A |
| chr1 | 2.08E+08 | 208100000 | 0.38 | 0.07 | A>A |
| chr1 | 2.08E+08 | 208200000 | 0.14 | -0.23 | B>B |
| chr1 | 2.08E+08 | 208300000 | -0.04 | -0.27 | B>B |
| chr1 | 2.08E+08 | 208400000 | 0.16 | 0.02 | A>A |
| chr1 | 2.08E+08 | 208500000 | -0.30 | -0.66 | B>B |
| chr1 | 2.09E+08 | 208600000 | -0.56 | -0.96 | B>B |
| chr1 | 2.09E+08 | 208700000 | -0.65 | -1.01 | B>B |
| chr1 | 2.09E+08 | 208800000 | -0.58 | -0.93 | B>B |
| chr1 | 2.09E+08 | 208900000 | -0.56 | -0.85 | B>B |
| chr1 | 2.09E+08 | 209000000 | -0.53 | -0.85 | B>B |
| chr1 | 2.09E+08 | 209100000 | -0.41 | -0.76 | B>B |
| chr1 | 2.09E+08 | 209200000 | -0.45 | -0.62 | B>B |
| chr1 | 2.09E+08 | 209300000 | -0.33 | -0.50 | B>B |
| chr1 | 2.09E+08 | 209400000 | -0.32 | -0.48 | B>B |
| chr1 | 2.09E+08 | 209500000 | -0.24 | -0.62 | B>B |
| chr1 | 2.1E+08 | 209600000 | -0.15 | -0.65 | B>B |
| chr1 | 2.1E+08 | 209700000 | -0.22 | -0.75 | B>B |
| chr1 | 2.1E+08 | 209800000 | -0.03 | -0.47 | B>B |
| chr1 | 2.1E+08 | 209900000 | 0.04 | -0.39 | B>B |
| chr1 | 2.1E+08 | 210000000 | -0.05 | -0.54 | B>B |
| chr1 | 2.1E+08 | 210100000 | -0.28 | -0.70 | B>B |
| chr1 | 2.1E+08 | 210200000 | -0.53 | -0.80 | B>B |
| chr1 | 2.1E+08 | 210300000 | -0.51 | -0.87 | B>B |
| chr1 | 2.1E+08 | 210400000 | -0.30 | -0.63 | B>B |
| chr1 | 2.1E+08 | 210500000 | -0.13 | -0.49 | B>B |
| chr1 | 2.11E+08 | 210600000 | 0.00 | -0.33 | B>B |
| chr1 | 2.11E+08 | 210700000 | -0.02 | -0.56 | B>B |
| chr1 | 2.11E+08 | 210800000 | 0.02 | -0.55 | B>B |
| chr1 | 2.11E+08 | 210900000 | 0.19 | -0.61 | A>B |
| chr1 | 2.11E+08 | 211000000 | 0.09 | -0.65 | A>B |
| chr1 | 2.11E+08 | 211100000 | 0.04 | -0.63 | A>B |
| chr1 | 2.11E+08 | 211200000 | 0.04 | -0.58 | A>B |

| | | | | | |
|---|---|---|---|---|---|
| chr1 | 2.11E+08 | 211300000 | -0.01 | -0.55 | B>B |
| chr1 | 2.11E+08 | 211400000 | 0.08 | -0.45 | B>B |
| chr1 | 2.11E+08 | 211500000 | 0.54 | 0.12 | A>A |
| chr1 | 2.12E+08 | 211600000 | 0.72 | 0.37 | A>A |
| chr1 | 2.12E+08 | 211700000 | 0.62 | 0.44 | A>A |
| chr1 | 2.12E+08 | 211800000 | 0.86 | 0.71 | A>A |
| chr1 | 2.12E+08 | 211900000 | 0.68 | 0.56 | A>A |
| chr1 | 2.12E+08 | 212000000 | 1.05 | 0.97 | A>A |
| chr1 | 2.12E+08 | 212100000 | 1.14 | 1.08 | A>A |
| chr1 | 2.12E+08 | 212200000 | 0.89 | 0.81 | A>A |
| chr1 | 2.12E+08 | 212300000 | 0.60 | 0.66 | A>A |
| chr1 | 2.12E+08 | 212400000 | 0.81 | 0.97 | A>A |
| chr1 | 2.12E+08 | 212500000 | 0.87 | 0.94 | A>A |
| chr1 | 2.13E+08 | 212600000 | 0.78 | 0.79 | A>A |
| chr1 | 2.13E+08 | 212700000 | 0.86 | 0.84 | A>A |
| chr1 | 2.13E+08 | 212800000 | 1.01 | 1.01 | A>A |
| chr1 | 2.13E+08 | 212900000 | 0.72 | 0.78 | A>A |
| chr1 | 2.13E+08 | 213000000 | 0.71 | 0.74 | A>A |
| chr1 | 2.13E+08 | 213100000 | 0.80 | 0.81 | A>A |
| chr1 | 2.13E+08 | 213200000 | 0.64 | 0.55 | A>A |
| chr1 | 2.13E+08 | 213300000 | 0.16 | 0.03 | A>A |
| chr1 | 2.13E+08 | 213400000 | -0.21 | -0.30 | B>B |
| chr1 | 2.13E+08 | 213500000 | -0.47 | -0.54 | B>B |
| chr1 | 2.14E+08 | 213600000 | -0.64 | -0.76 | B>B |
| chr1 | 2.14E+08 | 213700000 | -0.74 | -0.90 | B>B |
| chr1 | 2.14E+08 | 213800000 | -0.84 | -0.89 | B>B |
| chr1 | 2.14E+08 | 213900000 | -0.78 | -0.85 | B>B |
| chr1 | 2.14E+08 | 214000000 | -0.77 | -0.85 | B>B |
| chr1 | 2.14E+08 | 214100000 | -0.75 | -0.86 | B>B |
| chr1 | 2.14E+08 | 214200000 | -0.64 | -0.61 | B>B |
| chr1 | 2.14E+08 | 214300000 | -0.61 | -0.52 | B>B |
| chr1 | 2.14E+08 | 214400000 | -0.42 | -0.55 | B>B |
| chr1 | 2.14E+08 | 214500000 | -0.18 | -0.35 | B>B |
| chr1 | 2.15E+08 | 214600000 | -0.01 | -0.06 | B>B |
| chr1 | 2.15E+08 | 214700000 | 0.29 | 0.18 | A>A |
| chr1 | 2.15E+08 | 214800000 | 0.06 | -0.06 | B>B |
| chr1 | 2.15E+08 | 214900000 | -0.43 | -0.50 | B>B |
| chr1 | 2.15E+08 | 215000000 | -0.59 | -0.67 | B>B |
| chr1 | 2.15E+08 | 215100000 | -0.64 | -0.62 | B>B |
| chr1 | 2.15E+08 | 215200000 | -0.60 | -0.64 | B>B |
| chr1 | 2.15E+08 | 215300000 | -0.67 | -0.48 | B>B |
| chr1 | 2.15E+08 | 215400000 | -0.70 | -0.62 | B>B |
| chr1 | 2.15E+08 | 215500000 | -0.78 | -0.80 | B>B |
| chr1 | 2.16E+08 | 215600000 | -0.76 | -0.85 | B>B |
| chr1 | 2.16E+08 | 215700000 | -0.69 | -0.71 | B>B |
| chr1 | 2.16E+08 | 215800000 | -0.39 | -0.41 | B>B |
| chr1 | 2.16E+08 | 215900000 | -0.64 | -0.68 | B>B |
| chr1 | 2.16E+08 | 216000000 | -0.79 | -0.79 | B>B |
| chr1 | 2.16E+08 | 216100000 | -0.79 | -0.79 | B>B |
| chr1 | 2.16E+08 | 216200000 | -0.78 | -0.81 | B>B |
| chr1 | 2.16E+08 | 216300000 | -0.79 | -0.79 | B>B |
| chr1 | 2.16E+08 | 216400000 | -0.74 | -0.75 | B>B |
| chr1 | 2.16E+08 | 216500000 | -0.71 | -0.73 | B>B |
| chr1 | 2.17E+08 | 216600000 | -0.77 | -0.75 | B>B |
| chr1 | 2.17E+08 | 216700000 | -0.75 | -0.73 | B>B |
| chr1 | 2.17E+08 | 216800000 | -0.66 | -0.70 | B>B |

| | | | | | |
|---|---|---|---|---|---|
| chr1 | 2.17E+08 | 216900000 | -0.61 | -0.66 | B>B |
| chr1 | 2.17E+08 | 217000000 | -0.62 | -0.68 | B>B |
| chr1 | 2.17E+08 | 217100000 | -0.70 | -0.70 | B>B |
| chr1 | 2.17E+08 | 217200000 | -0.67 | -0.60 | B>B |
| chr1 | 2.17E+08 | 217300000 | -0.61 | -0.48 | B>B |
| chr1 | 2.17E+08 | 217400000 | -0.59 | -0.50 | B>B |
| chr1 | 2.17E+08 | 217500000 | -0.58 | -0.47 | B>B |
| chr1 | 2.18E+08 | 217600000 | -0.49 | -0.39 | B>B |
| chr1 | 2.18E+08 | 217700000 | -0.36 | -0.31 | B>B |
| chr1 | 2.18E+08 | 217800000 | -0.24 | -0.19 | B>B |
| chr1 | 2.18E+08 | 217900000 | -0.29 | -0.24 | B>B |
| chr1 | 2.18E+08 | 218000000 | -0.50 | -0.42 | B>B |
| chr1 | 2.18E+08 | 218100000 | -0.63 | -0.57 | B>B |
| chr1 | 2.18E+08 | 218200000 | -0.67 | -0.69 | B>B |
| chr1 | 2.18E+08 | 218300000 | -0.71 | -0.70 | B>B |
| chr1 | 2.18E+08 | 218400000 | -0.50 | -0.56 | B>B |
| chr1 | 2.18E+08 | 218500000 | -0.30 | -0.47 | B>B |
| chr1 | 2.19E+08 | 218600000 | 0.12 | 0.08 | A>A |
| chr1 | 2.19E+08 | 218700000 | 0.32 | 0.26 | A>A |
| chr1 | 2.19E+08 | 218800000 | 0.36 | 0.28 | A>A |
| chr1 | 2.19E+08 | 218900000 | 0.32 | 0.22 | A>A |
| chr1 | 2.19E+08 | 219000000 | 0.28 | 0.28 | A>A |
| chr1 | 2.19E+08 | 219100000 | 0.28 | 0.24 | A>A |
| chr1 | 2.19E+08 | 219200000 | 0.24 | 0.19 | A>A |
| chr1 | 2.19E+08 | 219300000 | 0.14 | -0.04 | A>A |
| chr1 | 2.19E+08 | 219400000 | 0.04 | -0.15 | B>B |
| chr1 | 2.19E+08 | 219500000 | -0.04 | -0.23 | B>B |
| chr1 | 2.2E+08 | 219600000 | -0.09 | -0.47 | B>B |
| chr1 | 2.2E+08 | 219700000 | -0.20 | -0.78 | B>B |
| chr1 | 2.2E+08 | 219800000 | -0.31 | -0.76 | B>B |
| chr1 | 2.2E+08 | 219900000 | -0.44 | -0.76 | B>B |
| chr1 | 2.2E+08 | 220000000 | -0.46 | -0.69 | B>B |
| chr1 | 2.2E+08 | 220100000 | -0.38 | -0.42 | B>B |
| chr1 | 2.2E+08 | 220200000 | -0.06 | -0.22 | B>B |
| chr1 | 2.2E+08 | 220300000 | 0.22 | 0.08 | A>A |
| chr1 | 2.2E+08 | 220400000 | 0.01 | -0.11 | B>B |
| chr1 | 2.2E+08 | 220500000 | -0.23 | -0.47 | B>B |
| chr1 | 2.21E+08 | 220600000 | -0.48 | -0.76 | B>B |
| chr1 | 2.21E+08 | 220700000 | -0.55 | -0.79 | B>B |
| chr1 | 2.21E+08 | 220800000 | -0.59 | -0.77 | B>B |
| chr1 | 2.21E+08 | 220900000 | -0.35 | -0.55 | B>B |
| chr1 | 2.21E+08 | 221000000 | -0.23 | -0.31 | B>B |
| chr1 | 2.21E+08 | 221100000 | -0.30 | -0.25 | B>B |
| chr1 | 2.21E+08 | 221200000 | -0.34 | -0.08 | B>B |
| chr1 | 2.21E+08 | 221300000 | -0.19 | -0.05 | B>B |
| chr1 | 2.21E+08 | 221400000 | -0.28 | -0.11 | B>B |
| chr1 | 2.21E+08 | 221500000 | -0.32 | -0.22 | B>B |
| chr1 | 2.22E+08 | 221600000 | -0.30 | -0.22 | B>B |
| chr1 | 2.22E+08 | 221700000 | -0.20 | -0.13 | B>B |
| chr1 | 2.22E+08 | 221800000 | -0.03 | 0.01 | B>B |
| chr1 | 2.22E+08 | 221900000 | -0.03 | 0.07 | A>A |
| chr1 | 2.22E+08 | 222000000 | -0.13 | -0.02 | B>B |
| chr1 | 2.22E+08 | 222100000 | -0.04 | -0.05 | B>B |
| chr1 | 2.22E+08 | 222200000 | -0.19 | -0.17 | B>B |
| chr1 | 2.22E+08 | 222300000 | -0.40 | -0.42 | B>B |
| chr1 | 2.22E+08 | 222400000 | -0.52 | -0.62 | B>B |

| | | | | | |
|---|---|---|---|---|---|
| chr1 | 2.22E+08 | 222500000 | -0.58 | -0.82 | B>B |
| chr1 | 2.23E+08 | 222600000 | -0.57 | -0.80 | B>B |
| chr1 | 2.23E+08 | 222700000 | -0.33 | -0.47 | B>B |
| chr1 | 2.23E+08 | 222800000 | 0.18 | -0.03 | A>A |
| chr1 | 2.23E+08 | 222900000 | 0.16 | 0.07 | A>A |
| chr1 | 2.23E+08 | 223000000 | 0.08 | -0.05 | A>A |
| chr1 | 2.23E+08 | 223100000 | -0.03 | -0.03 | B>B |
| chr1 | 2.23E+08 | 223200000 | -0.02 | -0.16 | B>B |
| chr1 | 2.23E+08 | 223300000 | -0.03 | -0.18 | B>B |
| chr1 | 2.23E+08 | 223400000 | 0.01 | -0.28 | B>B |
| chr1 | 2.23E+08 | 223500000 | -0.01 | -0.34 | B>B |
| chr1 | 2.24E+08 | 223600000 | -0.09 | -0.29 | B>B |
| chr1 | 2.24E+08 | 223700000 | 0.19 | -0.29 | B>B |
| chr1 | 2.24E+08 | 223800000 | 0.00 | 0.00 | NO |
| chr1 | 2.24E+08 | 223900000 | 0.26 | 0.09 | A>A |
| chr1 | 2.24E+08 | 224000000 | 0.73 | 0.66 | A>A |
| chr1 | 2.24E+08 | 224100000 | 0.69 | 0.65 | A>A |
| chr1 | 2.24E+08 | 224200000 | 0.00 | 0.00 | NO |
| chr1 | 2.24E+08 | 224300000 | 0.49 | 0.50 | A>A |
| chr1 | 2.24E+08 | 224400000 | 0.79 | 0.94 | A>A |
| chr1 | 2.24E+08 | 224500000 | 0.83 | 1.06 | A>A |
| chr1 | 2.25E+08 | 224600000 | 0.71 | 0.83 | A>A |
| chr1 | 2.25E+08 | 224700000 | 0.26 | 0.56 | A>A |
| chr1 | 2.25E+08 | 224800000 | -0.09 | 0.06 | B>B |
| chr1 | 2.25E+08 | 224900000 | -0.19 | 0.05 | B>B |
| chr1 | 2.25E+08 | 225000000 | -0.26 | 0.07 | B>B |
| chr1 | 2.25E+08 | 225100000 | -0.25 | 0.14 | B>B |
| chr1 | 2.25E+08 | 225200000 | -0.32 | 0.13 | B>B |
| chr1 | 2.25E+08 | 225300000 | -0.50 | 0.02 | B>A |
| chr1 | 2.25E+08 | 225400000 | -0.49 | -0.02 | B>B |
| chr1 | 2.25E+08 | 225500000 | -0.43 | 0.11 | B>A |
| chr1 | 2.26E+08 | 225600000 | -0.01 | 0.54 | B>A |
| chr1 | 2.26E+08 | 225700000 | 0.48 | 0.90 | A>A |
| chr1 | 2.26E+08 | 225800000 | 0.84 | 1.18 | A>A |
| chr1 | 2.26E+08 | 225900000 | 1.21 | 1.29 | A>A |
| chr1 | 2.26E+08 | 226000000 | 0.97 | 1.21 | A>A |
| chr1 | 2.26E+08 | 226100000 | 0.89 | 1.24 | A>A |
| chr1 | 2.26E+08 | 226200000 | 0.80 | 1.06 | A>A |
| chr1 | 2.26E+08 | 226300000 | 1.23 | 1.38 | A>A |
| chr1 | 2.26E+08 | 226400000 | 1.31 | 1.35 | A>A |
| chr1 | 2.26E+08 | 226500000 | 1.29 | 1.31 | A>A |
| chr1 | 2.27E+08 | 226600000 | 1.15 | 1.18 | A>A |
| chr1 | 2.27E+08 | 226700000 | 0.94 | 1.01 | A>A |
| chr1 | 2.27E+08 | 226800000 | 0.88 | 0.95 | A>A |
| chr1 | 2.27E+08 | 226900000 | 0.86 | 0.93 | A>A |
| chr1 | 2.27E+08 | 227000000 | 0.97 | 1.08 | A>A |
| chr1 | 2.27E+08 | 227100000 | 0.88 | 0.97 | A>A |
| chr1 | 2.27E+08 | 227200000 | 0.81 | 0.94 | A>A |
| chr1 | 2.27E+08 | 227300000 | 0.83 | 0.91 | A>A |
| chr1 | 2.27E+08 | 227400000 | 0.76 | 0.96 | A>A |
| chr1 | 2.27E+08 | 227500000 | 1.05 | 1.11 | A>A |
| chr1 | 2.28E+08 | 227600000 | 1.08 | 1.19 | A>A |
| chr1 | 2.28E+08 | 227700000 | 0.57 | 0.76 | A>A |
| chr1 | 2.28E+08 | 227800000 | 0.62 | 1.07 | A>A |
| chr1 | 2.28E+08 | 227900000 | 0.53 | 1.13 | B>A |
| chr1 | 2.28E+08 | 228000000 | 0.76 | 1.21 | A>A |

| | | | | | |
|---|---|---|---|---|---|
| chr1 | 2.28E+08 | 228100000 | 0.88 | 1.35 | A>A |
| chr1 | 2.28E+08 | 228200000 | 0.85 | 1.10 | A>A |
| chr1 | 2.28E+08 | 228300000 | 1.35 | 1.58 | A>A |
| chr1 | 2.28E+08 | 228400000 | 1.45 | 1.69 | A>A |
| chr1 | 2.28E+08 | 228500000 | 1.55 | 1.70 | A>A |
| chr1 | 2.29E+08 | 228600000 | 1.66 | 1.77 | A>A |
| chr1 | 2.29E+08 | 228700000 | 1.53 | 1.62 | A>A |
| chr1 | 2.29E+08 | 228800000 | 1.29 | 1.36 | A>A |
| chr1 | 2.29E+08 | 228900000 | 0.83 | 0.79 | A>A |
| chr1 | 2.29E+08 | 229000000 | 1.18 | 1.07 | A>A |
| chr1 | 2.29E+08 | 229100000 | 1.07 | 0.97 | A>A |
| chr1 | 2.29E+08 | 229200000 | 1.01 | 0.97 | A>A |
| chr1 | 2.29E+08 | 229300000 | 1.07 | 1.00 | A>A |
| chr1 | 2.29E+08 | 229400000 | 0.84 | 0.81 | A>A |
| chr1 | 2.29E+08 | 229500000 | 1.08 | 0.96 | A>A |
| chr1 | 2.3E+08 | 229600000 | 1.09 | 0.95 | A>A |
| chr1 | 2.3E+08 | 229700000 | 0.89 | 0.83 | A>A |
| chr1 | 2.3E+08 | 229800000 | 0.73 | 0.57 | A>A |
| chr1 | 2.3E+08 | 229900000 | 0.32 | 0.25 | A>A |
| chr1 | 2.3E+08 | 230000000 | 0.34 | 0.19 | A>A |
| chr1 | 2.3E+08 | 230100000 | 0.44 | 0.29 | A>A |
| chr1 | 2.3E+08 | 230200000 | 0.56 | 0.37 | A>A |
| chr1 | 2.3E+08 | 230300000 | 0.73 | 0.55 | A>A |
| chr1 | 2.3E+08 | 230400000 | 0.56 | 0.41 | A>A |
| chr1 | 2.3E+08 | 230500000 | 0.31 | 0.18 | A>A |
| chr1 | 2.31E+08 | 230600000 | 0.30 | 0.01 | A>A |
| chr1 | 2.31E+08 | 230700000 | 0.24 | -0.05 | A>A |
| chr1 | 2.31E+08 | 230800000 | 0.33 | 0.11 | A>A |
| chr1 | 2.31E+08 | 230900000 | 0.57 | 0.28 | A>A |
| chr1 | 2.31E+08 | 231000000 | 0.59 | 0.23 | A>A |
| chr1 | 2.31E+08 | 231100000 | 0.68 | 0.44 | A>A |
| chr1 | 2.31E+08 | 231200000 | 0.76 | 0.55 | A>A |
| chr1 | 2.31E+08 | 231300000 | 0.97 | 0.50 | A>A |
| chr1 | 2.31E+08 | 231400000 | 0.71 | 0.53 | A>A |
| chr1 | 2.31E+08 | 231500000 | 0.69 | 0.56 | A>A |
| chr1 | 2.32E+08 | 231600000 | 0.96 | 0.81 | A>A |
| chr1 | 2.32E+08 | 231700000 | 1.00 | 0.78 | A>A |
| chr1 | 2.32E+08 | 231800000 | 0.47 | 0.31 | A>A |
| chr1 | 2.32E+08 | 231900000 | 0.35 | 0.15 | A>A |
| chr1 | 2.32E+08 | 232000000 | 0.49 | 0.21 | A>A |
| chr1 | 2.32E+08 | 232100000 | 0.31 | 0.05 | A>A |
| chr1 | 2.32E+08 | 232200000 | 0.37 | 0.08 | A>A |
| chr1 | 2.32E+08 | 232300000 | 0.44 | 0.10 | A>A |
| chr1 | 2.32E+08 | 232400000 | 0.36 | 0.08 | A>A |
| chr1 | 2.32E+08 | 232500000 | 0.36 | 0.07 | A>A |
| chr1 | 2.33E+08 | 232600000 | 0.32 | 0.09 | A>A |
| chr1 | 2.33E+08 | 232700000 | 0.35 | 0.16 | A>A |
| chr1 | 2.33E+08 | 232800000 | 0.28 | 0.21 | A>A |
| chr1 | 2.33E+08 | 232900000 | 0.52 | 0.61 | A>A |
| chr1 | 2.33E+08 | 233000000 | 0.45 | 0.59 | A>A |
| chr1 | 2.33E+08 | 233100000 | 0.54 | 0.47 | A>A |
| chr1 | 2.33E+08 | 233200000 | 0.52 | 0.50 | A>A |
| chr1 | 2.33E+08 | 233300000 | 0.55 | 0.55 | A>A |
| chr1 | 2.33E+08 | 233400000 | 0.42 | 0.39 | A>A |
| chr1 | 2.33E+08 | 233500000 | 0.51 | 0.57 | A>A |
| chr1 | 2.34E+08 | 233600000 | 0.34 | 0.42 | A>A |

| | | | | | |
|---|---|---|---|---|---|
| chr1 | 2.34E+08 | 233700000 | 0.50 | 0.51 | A>A |
| chr1 | 2.34E+08 | 233800000 | 0.64 | 0.70 | A>A |
| chr1 | 2.34E+08 | 233900000 | 0.56 | 0.65 | A>A |
| chr1 | 2.34E+08 | 234000000 | 0.35 | 0.42 | A>A |
| chr1 | 2.34E+08 | 234100000 | 0.53 | 0.61 | A>A |
| chr1 | 2.34E+08 | 234200000 | 0.53 | 0.59 | A>A |
| chr1 | 2.34E+08 | 234300000 | 0.31 | 0.37 | A>A |
| chr1 | 2.34E+08 | 234400000 | 0.43 | 0.60 | A>A |
| chr1 | 2.34E+08 | 234500000 | 0.38 | 0.60 | A>A |
| chr1 | 2.35E+08 | 234600000 | 0.70 | 0.88 | A>A |
| chr1 | 2.35E+08 | 234700000 | 1.00 | 1.09 | A>A |
| chr1 | 2.35E+08 | 234800000 | 1.15 | 1.28 | A>A |
| chr1 | 2.35E+08 | 234900000 | 1.39 | 1.43 | A>A |
| chr1 | 2.35E+08 | 235000000 | 1.82 | 1.74 | A>A |
| chr1 | 2.35E+08 | 235100000 | 1.28 | 1.34 | A>A |
| chr1 | 2.35E+08 | 235200000 | 1.40 | 1.46 | A>A |
| chr1 | 2.35E+08 | 235300000 | 0.74 | 0.91 | A>A |
| chr1 | 2.35E+08 | 235400000 | 1.09 | 1.17 | A>A |
| chr1 | 2.35E+08 | 235500000 | 1.13 | 1.23 | A>A |
| chr1 | 2.36E+08 | 235600000 | 1.11 | 1.18 | A>A |
| chr1 | 2.36E+08 | 235700000 | 0.74 | 0.78 | A>A |
| chr1 | 2.36E+08 | 235800000 | 0.72 | 0.59 | A>A |
| chr1 | 2.36E+08 | 235900000 | 0.74 | 0.65 | A>A |
| chr1 | 2.36E+08 | 236000000 | 0.62 | 0.56 | A>A |
| chr1 | 2.36E+08 | 236100000 | 0.85 | 0.63 | A>A |
| chr1 | 2.36E+08 | 236200000 | 0.48 | 0.47 | A>A |
| chr1 | 2.36E+08 | 236300000 | 0.35 | 0.21 | A>A |
| chr1 | 2.36E+08 | 236400000 | 0.34 | 0.14 | A>A |
| chr1 | 2.36E+08 | 236500000 | 0.38 | 0.13 | A>A |
| chr1 | 2.37E+08 | 236600000 | 0.42 | 0.17 | A>A |
| chr1 | 2.37E+08 | 236700000 | 0.36 | 0.21 | A>A |
| chr1 | 2.37E+08 | 236800000 | 0.00 | -0.21 | B>B |
| chr1 | 2.37E+08 | 236900000 | -0.31 | -0.68 | B>B |
| chr1 | 2.37E+08 | 237000000 | -0.25 | -0.52 | B>B |
| chr1 | 2.37E+08 | 237100000 | -0.32 | -0.45 | B>B |
| chr1 | 2.37E+08 | 237200000 | -0.45 | -0.61 | B>B |
| chr1 | 2.37E+08 | 237300000 | -0.62 | -0.76 | B>B |
| chr1 | 2.37E+08 | 237400000 | -0.71 | -0.91 | B>B |
| chr1 | 2.37E+08 | 237500000 | -0.87 | -0.93 | B>B |
| chr1 | 2.38E+08 | 237600000 | -0.75 | -0.81 | B>B |
| chr1 | 2.38E+08 | 237700000 | -0.78 | -0.87 | B>B |
| chr1 | 2.38E+08 | 237800000 | -0.67 | -0.83 | B>B |
| chr1 | 2.38E+08 | 237900000 | -0.67 | -0.81 | B>B |
| chr1 | 2.38E+08 | 238000000 | -0.56 | -0.61 | B>B |
| chr1 | 2.38E+08 | 238100000 | -0.65 | -0.63 | B>B |
| chr1 | 2.38E+08 | 238200000 | -0.71 | -0.77 | B>B |
| chr1 | 2.38E+08 | 238300000 | -0.84 | -0.82 | B>B |
| chr1 | 2.38E+08 | 238400000 | -0.74 | -0.76 | B>B |
| chr1 | 2.38E+08 | 238500000 | -0.66 | -0.73 | B>B |
| chr1 | 2.39E+08 | 238600000 | -0.71 | -0.78 | B>B |
| chr1 | 2.39E+08 | 238700000 | -0.73 | -0.75 | B>B |
| chr1 | 2.39E+08 | 238800000 | -0.69 | -0.73 | B>B |
| chr1 | 2.39E+08 | 238900000 | -0.68 | -0.73 | B>B |
| chr1 | 2.39E+08 | 239000000 | -0.68 | -0.74 | B>B |
| chr1 | 2.39E+08 | 239100000 | -0.70 | -0.68 | B>B |
| chr1 | 2.39E+08 | 239200000 | -0.70 | -0.60 | B>B |

| | | | | | |
|---|---|---|---|---|---|
| chr1 | 2.39E+08 | 239300000 | -0.69 | -0.48 | B>B |
| chr1 | 2.39E+08 | 239400000 | -0.68 | -0.38 | B>B |
| chr1 | 2.39E+08 | 239500000 | -0.62 | -0.42 | B>B |
| chr1 | 2.4E+08 | 239600000 | -0.53 | -0.45 | B>B |
| chr1 | 2.4E+08 | 239700000 | -0.51 | -0.59 | B>B |
| chr1 | 2.4E+08 | 239800000 | -0.46 | -0.65 | B>B |
| chr1 | 2.4E+08 | 239900000 | -0.39 | -0.66 | B>B |
| chr1 | 2.4E+08 | 240000000 | -0.28 | -0.48 | B>B |
| chr1 | 2.4E+08 | 240100000 | -0.30 | -0.39 | B>B |
| chr1 | 2.4E+08 | 240200000 | -0.34 | -0.41 | B>B |
| chr1 | 2.4E+08 | 240300000 | -0.07 | -0.34 | B>B |
| chr1 | 2.4E+08 | 240400000 | -0.16 | -0.18 | B>B |
| chr1 | 2.4E+08 | 240500000 | -0.03 | 0.04 | A>A |
| chr1 | 2.41E+08 | 240600000 | 0.02 | 0.08 | A>A |
| chr1 | 2.41E+08 | 240700000 | -0.09 | -0.10 | B>B |
| chr1 | 2.41E+08 | 240800000 | -0.17 | -0.16 | B>B |
| chr1 | 2.41E+08 | 240900000 | -0.26 | -0.25 | B>B |
| chr1 | 2.41E+08 | 241000000 | -0.20 | -0.02 | B>B |
| chr1 | 2.41E+08 | 241100000 | -0.39 | -0.23 | B>B |
| chr1 | 2.41E+08 | 241200000 | -0.45 | -0.57 | B>B |
| chr1 | 2.41E+08 | 241300000 | -0.41 | -0.81 | B>B |
| chr1 | 2.41E+08 | 241400000 | -0.41 | -0.72 | B>B |
| chr1 | 2.41E+08 | 241500000 | -0.48 | -0.73 | B>B |
| chr1 | 2.42E+08 | 241600000 | -0.40 | -0.71 | B>B |
| chr1 | 2.42E+08 | 241700000 | -0.11 | -0.42 | B>B |
| chr1 | 2.42E+08 | 241800000 | 0.23 | 0.08 | A>A |
| chr1 | 2.42E+08 | 241900000 | 0.62 | 0.46 | A>A |
| chr1 | 2.42E+08 | 242000000 | 0.53 | 0.32 | A>A |
| chr1 | 2.42E+08 | 242100000 | 0.30 | 0.05 | A>A |
| chr1 | 2.42E+08 | 242200000 | 0.43 | -0.01 | A>A |
| chr1 | 2.42E+08 | 242300000 | 0.43 | -0.01 | A>A |
| chr1 | 2.42E+08 | 242400000 | 0.54 | 0.04 | A>A |
| chr1 | 2.42E+08 | 242500000 | 0.00 | 0.00 | NO |
| chr1 | 2.43E+08 | 242600000 | 0.38 | -0.12 | A>A |
| chr1 | 2.43E+08 | 242700000 | 0.32 | -0.43 | A>B |
| chr1 | 2.43E+08 | 242800000 | 0.43 | -0.65 | A>B |
| chr1 | 2.43E+08 | 242900000 | 0.26 | -0.76 | A>B |
| chr1 | 2.43E+08 | 243000000 | 0.27 | -0.72 | A>B |
| chr1 | 2.43E+08 | 243100000 | 0.24 | -0.66 | A>B |
| chr1 | 2.43E+08 | 243200000 | 0.00 | 0.00 | NO |
| chr1 | 2.43E+08 | 243300000 | 0.00 | 0.00 | NO |
| chr1 | 2.43E+08 | 243400000 | 0.00 | 0.00 | NO |
| chr1 | 2.43E+08 | 243500000 | 0.54 | 0.26 | A>A |
| chr1 | 2.44E+08 | 243600000 | 0.51 | 0.24 | A>A |
| chr1 | 2.44E+08 | 243700000 | 0.58 | 0.33 | A>A |
| chr1 | 2.44E+08 | 243800000 | 0.66 | 0.44 | A>A |
| chr1 | 2.44E+08 | 243900000 | 0.57 | 0.33 | A>A |
| chr1 | 2.44E+08 | 244000000 | 0.77 | 0.47 | A>A |
| chr1 | 2.44E+08 | 244100000 | 0.80 | 0.41 | A>A |
| chr1 | 2.44E+08 | 244200000 | 0.75 | 0.39 | A>A |
| chr1 | 2.44E+08 | 244300000 | 0.89 | 0.46 | A>A |
| chr1 | 2.44E+08 | 244400000 | 0.91 | 0.45 | A>A |
| chr1 | 2.44E+08 | 244500000 | 0.51 | 0.43 | A>A |
| chr1 | 2.45E+08 | 244600000 | 0.80 | 0.75 | A>A |
| chr1 | 2.45E+08 | 244700000 | 0.53 | 0.37 | A>A |
| chr1 | 2.45E+08 | 244800000 | 0.42 | 0.36 | A>A |

| | | | | | |
|---|---|---|---|---|---|
| chr1 | 2.45E+08 | 244900000 | 0.63 | 0.45 | A>A |
| chr1 | 2.45E+08 | 245000000 | 0.52 | 0.29 | A>A |
| chr1 | 2.45E+08 | 245100000 | 1.06 | 0.75 | A>A |
| chr1 | 2.45E+08 | 245200000 | 1.06 | 0.60 | A>A |
| chr1 | 2.45E+08 | 245300000 | 0.17 | -0.05 | A>A |
| chr1 | 2.45E+08 | 245400000 | -0.14 | -0.41 | B>B |
| chr1 | 2.45E+08 | 245500000 | -0.17 | -0.57 | B>B |
| chr1 | 2.46E+08 | 245600000 | -0.25 | -0.67 | B>B |
| chr1 | 2.46E+08 | 245700000 | -0.28 | -0.67 | B>B |
| chr1 | 2.46E+08 | 245800000 | -0.31 | -0.70 | B>B |
| chr1 | 2.46E+08 | 245900000 | -0.36 | -0.64 | B>B |
| chr1 | 2.46E+08 | 246000000 | -0.26 | -0.53 | B>B |
| chr1 | 2.46E+08 | 246100000 | -0.23 | -0.48 | B>B |
| chr1 | 2.46E+08 | 246200000 | -0.20 | -0.34 | B>B |
| chr1 | 2.46E+08 | 246300000 | -0.09 | -0.34 | B>B |
| chr1 | 2.46E+08 | 246400000 | 0.00 | -0.29 | B>B |
| chr1 | 2.46E+08 | 246500000 | 0.13 | -0.14 | B>B |
| chr1 | 2.47E+08 | 246600000 | 0.14 | -0.11 | A>A |
| chr1 | 2.47E+08 | 246700000 | 0.12 | 0.08 | A>A |
| chr1 | 2.47E+08 | 246800000 | 0.49 | 0.60 | A>A |
| chr1 | 2.47E+08 | 246900000 | 0.31 | 0.65 | A>A |
| chr1 | 2.47E+08 | 247000000 | 0.15 | 0.80 | B>A |
| chr1 | 2.47E+08 | 247100000 | 0.14 | 0.91 | B>A |
| chr1 | 2.47E+08 | 247200000 | -0.40 | 0.73 | B>A |
| chr1 | 2.47E+08 | 247300000 | -0.38 | 0.59 | B>A |
| chr1 | 2.47E+08 | 247400000 | -0.52 | 0.16 | B>A |
| chr1 | 2.47E+08 | 247500000 | -0.53 | -0.03 | B>B |
| chr1 | 2.48E+08 | 247600000 | -0.62 | -0.41 | B>B |
| chr1 | 2.48E+08 | 247700000 | -0.86 | -0.83 | B>B |
| chr1 | 2.48E+08 | 247800000 | -0.97 | -1.06 | B>B |
| chr1 | 2.48E+08 | 247900000 | -0.92 | -1.08 | B>B |
| chr1 | 2.48E+08 | 248000000 | -0.99 | -1.12 | B>B |
| chr1 | 2.48E+08 | 248100000 | -0.90 | -1.03 | B>B |
| chr1 | 2.48E+08 | 248200000 | -0.98 | -1.06 | B>B |
| chr1 | 2.48E+08 | 248300000 | -0.98 | -1.10 | B>B |
| chr1 | 2.48E+08 | 248400000 | -0.94 | -1.09 | B>B |
| chr1 | 2.48E+08 | 248500000 | -1.01 | -1.16 | B>B |
| chr1 | 2.49E+08 | 248600000 | -0.95 | -1.11 | B>B |
| chr1 | 2.49E+08 | 248700000 | 0.00 | 0.00 | NO |
| chr1 | 2.49E+08 | 248800000 | -1.06 | -1.35 | B>B |
| chr1 | 2.49E+08 | 248900000 | -0.96 | -1.18 | B>B |
| chr1 | 2.49E+08 | 249000000 | 0.00 | 0.00 | NO |
| chr1 | 2.49E+08 | 249100000 | 0.00 | 0.00 | NO |
| chr1 | 2.49E+08 | 249200000 | -0.19 | -0.25 | B>B |
| chr1 | 2.49E+08 | 249250621 | 0.00 | 0.00 | NO |
| chr2 | 0 | 100000 | -0.35 | 0.38 | B>A |
| chr2 | 100000 | 200000 | -0.50 | -0.08 | B>B |
| chr2 | 200000 | 300000 | -0.22 | 0.10 | B>B |
| chr2 | 300000 | 400000 | -0.53 | -0.21 | B>B |
| chr2 | 400000 | 500000 | -0.61 | -0.18 | B>B |
| chr2 | 500000 | 600000 | -0.50 | -0.21 | B>B |
| chr2 | 600000 | 700000 | -0.05 | 0.05 | B>B |
| chr2 | 700000 | 800000 | -0.02 | -0.06 | B>B |
| chr2 | 800000 | 900000 | -0.22 | -0.42 | B>B |
| chr2 | 900000 | 1000000 | -0.53 | -0.50 | B>B |
| chr2 | 1000000 | 1100000 | -0.71 | -0.48 | B>B |

| | | | | | |
|---|---|---|---|---|---|
| chr2 | 1100000 | 1200000 | -0.72 | -0.56 | B>B |
| chr2 | 1200000 | 1300000 | -0.72 | -0.57 | B>B |
| chr2 | 1300000 | 1400000 | -0.73 | -0.66 | B>B |
| chr2 | 1400000 | 1500000 | -0.54 | -0.49 | B>B |
| chr2 | 1500000 | 1600000 | -0.26 | 0.00 | B>B |
| chr2 | 1600000 | 1700000 | -0.18 | 0.52 | B>A |
| chr2 | 1700000 | 1800000 | -0.14 | 0.24 | A>A |
| chr2 | 1800000 | 1900000 | -0.27 | -0.30 | B>B |
| chr2 | 1900000 | 2000000 | -0.47 | -0.60 | B>B |
| chr2 | 2000000 | 2100000 | -0.64 | -0.83 | B>B |
| chr2 | 2100000 | 2200000 | -0.68 | -0.81 | B>B |
| chr2 | 2200000 | 2300000 | -0.76 | -0.99 | B>B |
| chr2 | 2300000 | 2400000 | -0.87 | -1.01 | B>B |
| chr2 | 2400000 | 2500000 | -0.71 | -0.93 | B>B |
| chr2 | 2500000 | 2600000 | -0.33 | -0.74 | B>B |
| chr2 | 2600000 | 2700000 | 0.05 | -0.40 | B>B |
| chr2 | 2700000 | 2800000 | -0.01 | -0.51 | B>B |
| chr2 | 2800000 | 2900000 | 0.01 | -0.50 | B>B |
| chr2 | 2900000 | 3000000 | 0.02 | -0.43 | B>B |
| chr2 | 3000000 | 3100000 | 0.14 | -0.22 | B>B |
| chr2 | 3100000 | 3200000 | 0.34 | 0.02 | A>A |
| chr2 | 3200000 | 3300000 | 0.35 | 0.28 | A>A |
| chr2 | 3300000 | 3400000 | 0.79 | 0.61 | A>A |
| chr2 | 3400000 | 3500000 | 0.59 | 0.58 | A>A |
| chr2 | 3500000 | 3600000 | 0.82 | 0.85 | A>A |
| chr2 | 3600000 | 3700000 | 0.94 | 0.76 | A>A |
| chr2 | 3700000 | 3800000 | 0.11 | -0.14 | B>B |
| chr2 | 3800000 | 3900000 | -0.13 | -0.39 | B>B |
| chr2 | 3900000 | 4000000 | -0.42 | -0.61 | B>B |
| chr2 | 4000000 | 4100000 | -0.47 | -0.69 | B>B |
| chr2 | 4100000 | 4200000 | -0.70 | -0.84 | B>B |
| chr2 | 4200000 | 4300000 | -0.87 | -0.88 | B>B |
| chr2 | 4300000 | 4400000 | -0.86 | -0.82 | B>B |
| chr2 | 4400000 | 4500000 | -0.83 | -0.79 | B>B |
| chr2 | 4500000 | 4600000 | -0.87 | -0.75 | B>B |
| chr2 | 4600000 | 4700000 | -0.80 | -0.72 | B>B |
| chr2 | 4700000 | 4800000 | -0.79 | -0.54 | B>B |
| chr2 | 4800000 | 4900000 | -0.76 | -0.59 | B>B |
| chr2 | 4900000 | 5000000 | -0.61 | -0.29 | B>B |
| chr2 | 5000000 | 5100000 | 0.00 | 0.00 | NO |
| chr2 | 5100000 | 5200000 | -0.60 | -0.29 | B>B |
| chr2 | 5200000 | 5300000 | -0.64 | -0.32 | B>B |
| chr2 | 5300000 | 5400000 | -0.74 | -0.29 | B>B |
| chr2 | 5400000 | 5500000 | -0.70 | -0.41 | B>B |
| chr2 | 5500000 | 5600000 | -0.64 | -0.51 | B>B |
| chr2 | 5600000 | 5700000 | -0.56 | -0.55 | B>B |
| chr2 | 5700000 | 5800000 | -0.71 | -0.69 | B>B |
| chr2 | 5800000 | 5900000 | -0.62 | -0.68 | B>B |
| chr2 | 5900000 | 6000000 | -0.65 | -0.73 | B>B |
| chr2 | 6000000 | 6100000 | -0.61 | -0.71 | B>B |
| chr2 | 6100000 | 6200000 | -0.55 | -0.62 | B>B |
| chr2 | 6200000 | 6300000 | -0.31 | -0.40 | B>B |
| chr2 | 6300000 | 6400000 | -0.33 | -0.33 | B>B |
| chr2 | 6400000 | 6500000 | -0.52 | -0.60 | B>B |
| chr2 | 6500000 | 6600000 | -0.62 | -0.70 | B>B |
| chr2 | 6600000 | 6700000 | -0.66 | -0.68 | B>B |

| | | | | | |
|---|---|---|---|---|---|
| chr2 | 6700000 | 6800000 | -0.68 | -0.76 | B>B |
| chr2 | 6800000 | 6900000 | -0.63 | -0.69 | B>B |
| chr2 | 6900000 | 7000000 | -0.32 | -0.41 | B>B |
| chr2 | 7000000 | 7100000 | 0.09 | -0.01 | A>A |
| chr2 | 7100000 | 7200000 | 0.07 | 0.06 | A>A |
| chr2 | 7200000 | 7300000 | -0.10 | -0.13 | B>B |
| chr2 | 7300000 | 7400000 | -0.22 | -0.32 | B>B |
| chr2 | 7400000 | 7500000 | -0.32 | -0.49 | B>B |
| chr2 | 7500000 | 7600000 | -0.28 | -0.27 | B>B |
| chr2 | 7600000 | 7700000 | -0.44 | -0.43 | B>B |
| chr2 | 7700000 | 7800000 | -0.33 | -0.33 | B>B |
| chr2 | 7800000 | 7900000 | -0.11 | 0.05 | B>B |
| chr2 | 7900000 | 8000000 | 0.08 | 0.36 | A>A |
| chr2 | 8000000 | 8100000 | 0.03 | 0.27 | A>A |
| chr2 | 8100000 | 8200000 | 0.09 | 0.24 | A>A |
| chr2 | 8200000 | 8300000 | 0.15 | 0.32 | A>A |
| chr2 | 8300000 | 8400000 | 0.27 | 0.39 | A>A |
| chr2 | 8400000 | 8500000 | 0.38 | 0.50 | A>A |
| chr2 | 8500000 | 8600000 | 0.49 | 0.59 | A>A |
| chr2 | 8600000 | 8700000 | 0.59 | 0.59 | A>A |
| chr2 | 8700000 | 8800000 | 0.62 | 0.59 | A>A |
| chr2 | 8800000 | 8900000 | 0.36 | 0.31 | A>A |
| chr2 | 8900000 | 9000000 | 0.55 | 0.51 | A>A |
| chr2 | 9000000 | 9100000 | 0.58 | 0.63 | A>A |
| chr2 | 9100000 | 9200000 | 0.79 | 0.76 | A>A |
| chr2 | 9200000 | 9300000 | 0.77 | 0.75 | A>A |
| chr2 | 9300000 | 9400000 | 1.14 | 1.18 | A>A |
| chr2 | 9400000 | 9500000 | 1.15 | 1.20 | A>A |
| chr2 | 9500000 | 9600000 | 0.97 | 1.08 | A>A |
| chr2 | 9600000 | 9700000 | 1.24 | 1.21 | A>A |
| chr2 | 9700000 | 9800000 | 1.38 | 1.50 | A>A |
| chr2 | 9800000 | 9900000 | 1.46 | 1.58 | A>A |
| chr2 | 9900000 | 10000000 | 1.13 | 1.29 | A>A |
| chr2 | 10000000 | 10100000 | 1.42 | 1.58 | A>A |
| chr2 | 10100000 | 10200000 | 1.37 | 1.51 | A>A |
| chr2 | 10200000 | 10300000 | 1.17 | 1.39 | A>A |
| chr2 | 10300000 | 10400000 | 1.36 | 1.56 | A>A |
| chr2 | 10400000 | 10500000 | 1.62 | 1.74 | A>A |
| chr2 | 10500000 | 10600000 | 1.30 | 1.43 | A>A |
| chr2 | 10600000 | 10700000 | 1.35 | 1.60 | A>A |
| chr2 | 10700000 | 10800000 | 1.39 | 1.48 | A>A |
| chr2 | 10800000 | 10900000 | 1.20 | 1.31 | A>A |
| chr2 | 10900000 | 11000000 | 1.04 | 1.15 | A>A |
| chr2 | 11000000 | 11100000 | 1.28 | 1.36 | A>A |
| chr2 | 11100000 | 11200000 | 1.28 | 1.33 | A>A |
| chr2 | 11200000 | 11300000 | 0.89 | 1.04 | A>A |
| chr2 | 11300000 | 11400000 | 0.99 | 1.18 | A>A |
| chr2 | 11400000 | 11500000 | 1.23 | 1.44 | A>A |
| chr2 | 11500000 | 11600000 | 0.99 | 1.08 | A>A |
| chr2 | 11600000 | 11700000 | 0.95 | 0.83 | A>A |
| chr2 | 11700000 | 11800000 | 0.74 | 0.74 | A>A |
| chr2 | 11800000 | 11900000 | 0.63 | 0.56 | A>A |
| chr2 | 11900000 | 12000000 | 0.11 | 0.04 | A>A |
| chr2 | 12000000 | 12100000 | -0.05 | -0.17 | B>B |
| chr2 | 12100000 | 12200000 | 0.00 | -0.14 | B>B |
| chr2 | 12200000 | 12300000 | -0.04 | -0.12 | B>B |

| | | | | | |
|---|---|---|---|---|---|
| chr2 | 12300000 | 12400000 | -0.23 | -0.27 | B>B |
| chr2 | 12400000 | 12500000 | -0.39 | -0.34 | B>B |
| chr2 | 12500000 | 12600000 | -0.51 | -0.54 | B>B |
| chr2 | 12600000 | 12700000 | -0.57 | -0.76 | B>B |
| chr2 | 12700000 | 12800000 | -0.59 | -0.82 | B>B |
| chr2 | 12800000 | 12900000 | -0.52 | -0.71 | B>B |
| chr2 | 12900000 | 13000000 | -0.74 | -0.87 | B>B |
| chr2 | 13000000 | 13100000 | -0.79 | -0.79 | B>B |
| chr2 | 13100000 | 13200000 | -0.59 | -0.65 | B>B |
| chr2 | 13200000 | 13300000 | -0.88 | -0.78 | B>B |
| chr2 | 13300000 | 13400000 | -0.85 | -0.79 | B>B |
| chr2 | 13400000 | 13500000 | -0.88 | -0.81 | B>B |
| chr2 | 13500000 | 13600000 | -0.81 | -0.77 | B>B |
| chr2 | 13600000 | 13700000 | -0.83 | -0.57 | B>B |
| chr2 | 13700000 | 13800000 | -0.82 | -0.46 | B>B |
| chr2 | 13800000 | 13900000 | -0.85 | -0.34 | B>B |
| chr2 | 13900000 | 14000000 | -0.84 | -0.30 | B>A |
| chr2 | 14000000 | 14100000 | -0.82 | -0.31 | B>B |
| chr2 | 14100000 | 14200000 | -0.85 | -0.36 | B>B |
| chr2 | 14200000 | 14300000 | -0.81 | -0.54 | B>B |
| chr2 | 14300000 | 14400000 | -0.81 | -0.74 | B>B |
| chr2 | 14400000 | 14500000 | -0.84 | -0.81 | B>B |
| chr2 | 14500000 | 14600000 | -0.79 | -0.79 | B>B |
| chr2 | 14600000 | 14700000 | -0.74 | -0.76 | B>B |
| chr2 | 14700000 | 14800000 | -0.64 | -0.75 | B>B |
| chr2 | 14800000 | 14900000 | -0.74 | -0.79 | B>B |
| chr2 | 14900000 | 15000000 | -0.71 | -0.76 | B>B |
| chr2 | 15000000 | 15100000 | -0.60 | -0.80 | B>B |
| chr2 | 15100000 | 15200000 | -0.51 | -0.81 | B>B |
| chr2 | 15200000 | 15300000 | -0.38 | -0.70 | B>B |
| chr2 | 15300000 | 15400000 | -0.10 | -0.28 | B>B |
| chr2 | 15400000 | 15500000 | 0.14 | 0.00 | A>A |
| chr2 | 15500000 | 15600000 | 0.28 | 0.16 | A>A |
| chr2 | 15600000 | 15700000 | 0.29 | 0.08 | A>A |
| chr2 | 15700000 | 15800000 | 0.05 | -0.06 | B>B |
| chr2 | 15800000 | 15900000 | 0.15 | -0.11 | A>A |
| chr2 | 15900000 | 16000000 | 0.13 | -0.50 | A>B |
| chr2 | 16000000 | 16100000 | -0.11 | -0.55 | B>B |
| chr2 | 16100000 | 16200000 | -0.13 | -0.40 | B>B |
| chr2 | 16200000 | 16300000 | -0.08 | -0.31 | B>B |
| chr2 | 16300000 | 16400000 | -0.19 | -0.40 | B>B |
| chr2 | 16400000 | 16500000 | -0.31 | -0.66 | B>B |
| chr2 | 16500000 | 16600000 | -0.51 | -0.78 | B>B |
| chr2 | 16600000 | 16700000 | -0.48 | -0.72 | B>B |
| chr2 | 16700000 | 16800000 | -0.52 | -0.73 | B>B |
| chr2 | 16800000 | 16900000 | -0.42 | -0.73 | B>B |
| chr2 | 16900000 | 17000000 | -0.53 | -0.79 | B>B |
| chr2 | 17000000 | 17100000 | -0.77 | -0.87 | B>B |
| chr2 | 17100000 | 17200000 | -0.83 | -0.85 | B>B |
| chr2 | 17200000 | 17300000 | -0.90 | -0.86 | B>B |
| chr2 | 17300000 | 17400000 | -0.64 | -0.84 | B>B |
| chr2 | 17400000 | 17500000 | -0.51 | -0.74 | B>B |
| chr2 | 17500000 | 17600000 | -0.42 | -0.61 | B>B |
| chr2 | 17600000 | 17700000 | -0.48 | -0.66 | B>B |
| chr2 | 17700000 | 17800000 | -0.30 | -0.69 | B>B |
| chr2 | 17800000 | 17900000 | 0.08 | -0.32 | B>B |

| | | | | | |
|---|---|---|---|---|---|
| chr2 | 17900000 | 18000000 | 0.45 | -0.15 | A>B |
| chr2 | 18000000 | 18100000 | 0.04 | -0.58 | A>B |
| chr2 | 18100000 | 18200000 | -0.12 | -0.82 | A>B |
| chr2 | 18200000 | 18300000 | -0.17 | -0.90 | A>B |
| chr2 | 18300000 | 18400000 | 0.00 | -0.85 | A>B |
| chr2 | 18400000 | 18500000 | 0.18 | -0.79 | A>B |
| chr2 | 18500000 | 18600000 | 0.12 | -0.58 | A>B |
| chr2 | 18600000 | 18700000 | 0.14 | -0.63 | A>B |
| chr2 | 18700000 | 18800000 | 0.04 | -0.58 | A>B |
| chr2 | 18800000 | 18900000 | 0.05 | -0.77 | A>B |
| chr2 | 18900000 | 19000000 | 0.25 | -0.22 | A>A |
| chr2 | 19000000 | 19100000 | 0.45 | 0.05 | A>A |
| chr2 | 19100000 | 19200000 | 0.54 | 0.18 | A>A |
| chr2 | 19200000 | 19300000 | 0.54 | 0.34 | A>A |
| chr2 | 19300000 | 19400000 | 0.56 | 0.32 | A>A |
| chr2 | 19400000 | 19500000 | 0.60 | 0.31 | A>A |
| chr2 | 19500000 | 19600000 | 0.32 | 0.05 | A>A |
| chr2 | 19600000 | 19700000 | 0.61 | 0.30 | A>A |
| chr2 | 19700000 | 19800000 | 0.70 | 0.47 | A>A |
| chr2 | 19800000 | 19900000 | 0.68 | 0.23 | A>A |
| chr2 | 19900000 | 20000000 | 0.62 | 0.20 | A>A |
| chr2 | 20000000 | 20100000 | 0.52 | 0.15 | A>A |
| chr2 | 20100000 | 20200000 | 0.90 | 0.68 | A>A |
| chr2 | 20200000 | 20300000 | 1.06 | 0.99 | A>A |
| chr2 | 20300000 | 20400000 | 1.07 | 1.01 | A>A |
| chr2 | 20400000 | 20500000 | 0.94 | 0.91 | A>A |
| chr2 | 20500000 | 20600000 | 0.72 | 0.58 | A>A |
| chr2 | 20600000 | 20700000 | 0.68 | 0.51 | A>A |
| chr2 | 20700000 | 20800000 | 0.85 | 0.68 | A>A |
| chr2 | 20800000 | 20900000 | 0.30 | 0.06 | A>A |
| chr2 | 20900000 | 21000000 | 0.02 | -0.18 | B>B |
| chr2 | 21000000 | 21100000 | -0.10 | -0.52 | B>B |
| chr2 | 21100000 | 21200000 | -0.14 | -0.61 | B>B |
| chr2 | 21200000 | 21300000 | -0.33 | -0.64 | B>B |
| chr2 | 21300000 | 21400000 | -0.57 | -0.79 | B>B |
| chr2 | 21400000 | 21500000 | -0.63 | -0.85 | B>B |
| chr2 | 21500000 | 21600000 | -0.70 | -0.84 | B>B |
| chr2 | 21600000 | 21700000 | -0.79 | -0.87 | B>B |
| chr2 | 21700000 | 21800000 | -0.85 | -0.85 | B>B |
| chr2 | 21800000 | 21900000 | -0.81 | -0.85 | B>B |
| chr2 | 21900000 | 22000000 | -0.84 | -0.85 | B>B |
| chr2 | 22000000 | 22100000 | -0.86 | -0.85 | B>B |
| chr2 | 22100000 | 22200000 | -0.81 | -0.87 | B>B |
| chr2 | 22200000 | 22300000 | -0.85 | -0.82 | B>B |
| chr2 | 22300000 | 22400000 | -0.85 | -0.82 | B>B |
| chr2 | 22400000 | 22500000 | -0.85 | -0.87 | B>B |
| chr2 | 22500000 | 22600000 | -0.87 | -0.86 | B>B |
| chr2 | 22600000 | 22700000 | -0.89 | -0.87 | B>B |
| chr2 | 22700000 | 22800000 | -0.86 | -0.87 | B>B |
| chr2 | 22800000 | 22900000 | -0.82 | -0.86 | B>B |
| chr2 | 22900000 | 23000000 | -0.87 | -0.89 | B>B |
| chr2 | 23000000 | 23100000 | -0.79 | -0.87 | B>B |
| chr2 | 23100000 | 23200000 | -0.61 | -0.85 | B>B |
| chr2 | 23200000 | 23300000 | -0.48 | -0.84 | B>B |
| chr2 | 23300000 | 23400000 | -0.36 | -0.76 | B>B |
| chr2 | 23400000 | 23500000 | -0.16 | -0.59 | B>B |

| | | | | | |
|---|---|---|---|---|---|
| chr2 | 23500000 | 23600000 | 0.02 | -0.35 | B>B |
| chr2 | 23600000 | 23700000 | 0.19 | -0.05 | A>A |
| chr2 | 23700000 | 23800000 | 0.22 | 0.05 | A>A |
| chr2 | 23800000 | 23900000 | 0.31 | 0.17 | A>A |
| chr2 | 23900000 | 24000000 | 0.34 | 0.32 | A>A |
| chr2 | 24000000 | 24100000 | 0.72 | 0.72 | A>A |
| chr2 | 24100000 | 24200000 | 0.91 | 0.94 | A>A |
| chr2 | 24200000 | 24300000 | 1.04 | 0.94 | A>A |
| chr2 | 24300000 | 24400000 | 0.89 | 1.00 | A>A |
| chr2 | 24400000 | 24500000 | 0.91 | 1.00 | A>A |
| chr2 | 24500000 | 24600000 | 0.99 | 1.05 | A>A |
| chr2 | 24600000 | 24700000 | 0.75 | 0.94 | A>A |
| chr2 | 24700000 | 24800000 | 0.97 | 1.16 | A>A |
| chr2 | 24800000 | 24900000 | 1.03 | 1.14 | A>A |
| chr2 | 24900000 | 25000000 | 0.90 | 1.19 | A>A |
| chr2 | 25000000 | 25100000 | 1.00 | 1.01 | A>A |
| chr2 | 25100000 | 25200000 | 1.05 | 1.21 | A>A |
| chr2 | 25200000 | 25300000 | 1.22 | 1.27 | A>A |
| chr2 | 25300000 | 25400000 | 1.13 | 1.24 | A>A |
| chr2 | 25400000 | 25500000 | 1.20 | 1.13 | A>A |
| chr2 | 25500000 | 25600000 | 1.22 | 1.32 | A>A |
| chr2 | 25600000 | 25700000 | 1.38 | 1.47 | A>A |
| chr2 | 25700000 | 25800000 | 1.15 | 1.27 | A>A |
| chr2 | 25800000 | 25900000 | 1.27 | 1.36 | A>A |
| chr2 | 25900000 | 26000000 | 1.19 | 1.33 | A>A |
| chr2 | 26000000 | 26100000 | 1.28 | 1.30 | A>A |
| chr2 | 26100000 | 26200000 | 1.22 | 1.31 | A>A |
| chr2 | 26200000 | 26300000 | 1.35 | 1.62 | A>A |
| chr2 | 26300000 | 26400000 | 1.26 | 1.50 | A>A |
| chr2 | 26400000 | 26500000 | 1.20 | 1.29 | A>A |
| chr2 | 26500000 | 26600000 | 1.25 | 1.28 | A>A |
| chr2 | 26600000 | 26700000 | 1.18 | 1.27 | A>A |
| chr2 | 26700000 | 26800000 | 1.02 | 1.19 | A>A |
| chr2 | 26800000 | 26900000 | 1.12 | 1.20 | A>A |
| chr2 | 26900000 | 27000000 | 1.14 | 1.20 | A>A |
| chr2 | 27000000 | 27100000 | 1.04 | 1.17 | A>A |
| chr2 | 27100000 | 27200000 | 1.01 | 1.20 | A>A |
| chr2 | 27200000 | 27300000 | 1.09 | 1.13 | A>A |
| chr2 | 27300000 | 27400000 | 1.05 | 1.14 | A>A |
| chr2 | 27400000 | 27500000 | 1.07 | 1.22 | A>A |
| chr2 | 27500000 | 27600000 | 1.10 | 1.15 | A>A |
| chr2 | 27600000 | 27700000 | 1.05 | 1.29 | A>A |
| chr2 | 27700000 | 27800000 | 1.21 | 1.40 | A>A |
| chr2 | 27800000 | 27900000 | 0.93 | 1.31 | A>A |
| chr2 | 27900000 | 28000000 | 0.68 | 0.96 | A>A |
| chr2 | 28000000 | 28100000 | 1.14 | 1.21 | A>A |
| chr2 | 28100000 | 28200000 | 0.99 | 1.16 | A>A |
| chr2 | 28200000 | 28300000 | 1.05 | 1.30 | A>A |
| chr2 | 28300000 | 28400000 | 1.02 | 1.02 | A>A |
| chr2 | 28400000 | 28500000 | 1.12 | 1.27 | A>A |
| chr2 | 28500000 | 28600000 | 1.03 | 1.23 | A>A |
| chr2 | 28600000 | 28700000 | 0.79 | 0.96 | A>A |
| chr2 | 28700000 | 28800000 | 0.88 | 1.07 | A>A |
| chr2 | 28800000 | 28900000 | 1.07 | 1.04 | A>A |
| chr2 | 28900000 | 29000000 | 1.06 | 1.11 | A>A |
| chr2 | 29000000 | 29100000 | 0.82 | 0.81 | A>A |

| | | | | | |
|---|---|---|---|---|---|
| chr2 | 29100000 | 29200000 | 0.70 | 0.44 | A>A |
| chr2 | 29200000 | 29300000 | 0.85 | 0.40 | A>A |
| chr2 | 29300000 | 29400000 | 0.92 | 0.53 | A>A |
| chr2 | 29400000 | 29500000 | 0.37 | -0.16 | A>A |
| chr2 | 29500000 | 29600000 | 0.18 | -0.58 | A>B |
| chr2 | 29600000 | 29700000 | -0.05 | -0.86 | A>B |
| chr2 | 29700000 | 29800000 | -0.03 | -0.85 | A>B |
| chr2 | 29800000 | 29900000 | -0.03 | -0.89 | A>B |
| chr2 | 29900000 | 30000000 | 0.01 | -0.83 | A>B |
| chr2 | 30000000 | 30100000 | 0.02 | -0.84 | A>B |
| chr2 | 30100000 | 30200000 | 0.05 | -0.85 | A>B |
| chr2 | 30200000 | 30300000 | 0.07 | -0.63 | A>B |
| chr2 | 30300000 | 30400000 | 0.25 | -0.20 | A>A |
| chr2 | 30400000 | 30500000 | 0.54 | 0.30 | A>A |
| chr2 | 30500000 | 30600000 | 0.80 | 0.42 | A>A |
| chr2 | 30600000 | 30700000 | 0.73 | 0.39 | A>A |
| chr2 | 30700000 | 30800000 | 0.67 | 0.37 | A>A |
| chr2 | 30800000 | 30900000 | 0.50 | 0.26 | A>A |
| chr2 | 30900000 | 31000000 | 0.19 | -0.27 | B>B |
| chr2 | 31000000 | 31100000 | 0.20 | -0.52 | A>B |
| chr2 | 31100000 | 31200000 | 0.35 | -0.49 | A>B |
| chr2 | 31200000 | 31300000 | 0.48 | -0.41 | A>B |
| chr2 | 31300000 | 31400000 | 0.39 | -0.36 | A>B |
| chr2 | 31400000 | 31500000 | 0.34 | -0.26 | A>B |
| chr2 | 31500000 | 31600000 | 0.30 | -0.23 | A>A |
| chr2 | 31600000 | 31700000 | 0.32 | -0.18 | A>A |
| chr2 | 31700000 | 31800000 | 0.21 | -0.40 | A>B |
| chr2 | 31800000 | 31900000 | 0.14 | -0.43 | B>B |
| chr2 | 31900000 | 32000000 | 0.23 | -0.16 | A>A |
| chr2 | 32000000 | 32100000 | 0.46 | 0.07 | A>A |
| chr2 | 32100000 | 32200000 | 0.78 | 0.59 | A>A |
| chr2 | 32200000 | 32300000 | 0.91 | 0.70 | A>A |
| chr2 | 32300000 | 32400000 | 0.90 | 0.67 | A>A |
| chr2 | 32400000 | 32500000 | 0.80 | 0.70 | A>A |
| chr2 | 32500000 | 32600000 | 0.98 | 0.84 | A>A |
| chr2 | 32600000 | 32700000 | 1.03 | 1.04 | A>A |
| chr2 | 32700000 | 32800000 | 0.82 | 0.90 | A>A |
| chr2 | 32800000 | 32900000 | 0.74 | 0.74 | A>A |
| chr2 | 32900000 | 33000000 | 0.51 | 0.47 | A>A |
| chr2 | 33000000 | 33100000 | 0.44 | 0.45 | A>A |
| chr2 | 33100000 | 33200000 | 0.48 | 0.44 | A>A |
| chr2 | 33200000 | 33300000 | 0.47 | 0.42 | A>A |
| chr2 | 33300000 | 33400000 | 0.50 | 0.50 | A>A |
| chr2 | 33400000 | 33500000 | 0.48 | 0.50 | A>A |
| chr2 | 33500000 | 33600000 | 0.46 | 0.44 | A>A |
| chr2 | 33600000 | 33700000 | 0.31 | 0.30 | A>A |
| chr2 | 33700000 | 33800000 | 0.25 | 0.19 | A>A |
| chr2 | 33800000 | 33900000 | -0.08 | -0.17 | B>B |
| chr2 | 33900000 | 34000000 | -0.43 | -0.60 | B>B |
| chr2 | 34000000 | 34100000 | -0.65 | -0.70 | B>B |
| chr2 | 34100000 | 34200000 | -0.66 | -0.70 | B>B |
| chr2 | 34200000 | 34300000 | -0.65 | -0.73 | B>B |
| chr2 | 34300000 | 34400000 | -0.68 | -0.60 | B>B |
| chr2 | 34400000 | 34500000 | -0.78 | -0.76 | B>B |
| chr2 | 34500000 | 34600000 | -0.85 | -0.72 | B>B |
| chr2 | 34600000 | 34700000 | -0.79 | -0.60 | B>B |

| | | | | | |
|---|---|---|---|---|---|
| chr2 | 34700000 | 34800000 | -0.77 | -0.55 | B>B |
| chr2 | 34800000 | 34900000 | -0.78 | -0.29 | B>B |
| chr2 | 34900000 | 35000000 | -0.74 | -0.36 | B>B |
| chr2 | 35000000 | 35100000 | -0.80 | -0.40 | B>B |
| chr2 | 35100000 | 35200000 | -0.75 | -0.46 | B>B |
| chr2 | 35200000 | 35300000 | -0.71 | -0.44 | B>B |
| chr2 | 35300000 | 35400000 | -0.69 | -0.43 | B>B |
| chr2 | 35400000 | 35500000 | -0.73 | -0.61 | B>B |
| chr2 | 35500000 | 35600000 | -0.74 | -0.63 | B>B |
| chr2 | 35600000 | 35700000 | -0.70 | -0.60 | B>B |
| chr2 | 35700000 | 35800000 | -0.73 | -0.68 | B>B |
| chr2 | 35800000 | 35900000 | -0.76 | -0.63 | B>B |
| chr2 | 35900000 | 36000000 | -0.62 | -0.56 | B>B |
| chr2 | 36000000 | 36100000 | -0.57 | -0.54 | B>B |
| chr2 | 36100000 | 36200000 | -0.47 | -0.44 | B>B |
| chr2 | 36200000 | 36300000 | -0.47 | -0.40 | B>B |
| chr2 | 36300000 | 36400000 | -0.50 | -0.42 | B>B |
| chr2 | 36400000 | 36500000 | -0.31 | -0.32 | B>B |
| chr2 | 36500000 | 36600000 | -0.02 | -0.09 | B>B |
| chr2 | 36600000 | 36700000 | 0.27 | 0.43 | A>A |
| chr2 | 36700000 | 36800000 | 0.14 | 0.24 | A>A |
| chr2 | 36800000 | 36900000 | 0.18 | 0.19 | A>A |
| chr2 | 36900000 | 37000000 | 0.11 | 0.12 | A>A |
| chr2 | 37000000 | 37100000 | 0.34 | 0.47 | A>A |
| chr2 | 37100000 | 37200000 | 0.63 | 0.63 | A>A |
| chr2 | 37200000 | 37300000 | 0.71 | 0.81 | A>A |
| chr2 | 37300000 | 37400000 | 0.83 | 1.03 | A>A |
| chr2 | 37400000 | 37500000 | 0.99 | 1.05 | A>A |
| chr2 | 37500000 | 37600000 | 0.85 | 1.01 | A>A |
| chr2 | 37600000 | 37700000 | 0.72 | 0.77 | A>A |
| chr2 | 37700000 | 37800000 | 0.93 | 1.22 | A>A |
| chr2 | 37800000 | 37900000 | 1.01 | 1.13 | A>A |
| chr2 | 37900000 | 38000000 | 0.99 | 1.28 | A>A |
| chr2 | 38000000 | 38100000 | 0.79 | 0.87 | A>A |
| chr2 | 38100000 | 38200000 | 0.67 | 0.75 | A>A |
| chr2 | 38200000 | 38300000 | 0.91 | 0.94 | A>A |
| chr2 | 38300000 | 38400000 | 0.98 | 1.12 | A>A |
| chr2 | 38400000 | 38500000 | 0.94 | 1.00 | A>A |
| chr2 | 38500000 | 38600000 | 0.75 | 0.89 | A>A |
| chr2 | 38600000 | 38700000 | 1.13 | 1.18 | A>A |
| chr2 | 38700000 | 38800000 | 0.84 | 0.90 | A>A |
| chr2 | 38800000 | 38900000 | 1.14 | 1.17 | A>A |
| chr2 | 38900000 | 39000000 | 1.03 | 1.04 | A>A |
| chr2 | 39000000 | 39100000 | 0.90 | 0.96 | A>A |
| chr2 | 39100000 | 39200000 | 0.75 | 0.87 | A>A |
| chr2 | 39200000 | 39300000 | 0.70 | 0.74 | A>A |
| chr2 | 39300000 | 39400000 | 0.70 | 0.67 | A>A |
| chr2 | 39400000 | 39500000 | 0.57 | 0.63 | A>A |
| chr2 | 39500000 | 39600000 | 0.52 | 0.52 | A>A |
| chr2 | 39600000 | 39700000 | 0.50 | 0.59 | A>A |
| chr2 | 39700000 | 39800000 | 0.51 | 0.45 | A>A |
| chr2 | 39800000 | 39900000 | 0.41 | 0.31 | A>A |
| chr2 | 39900000 | 40000000 | 0.23 | 0.14 | A>A |
| chr2 | 40000000 | 40100000 | -0.07 | -0.43 | B>B |
| chr2 | 40100000 | 40200000 | -0.42 | -0.65 | B>B |
| chr2 | 40200000 | 40300000 | -0.56 | -0.86 | B>B |

| | | | | | |
|---|---|---|---|---|---|
| chr2 | 40300000 | 40400000 | -0.36 | -0.78 | B>B |
| chr2 | 40400000 | 40500000 | -0.30 | -0.49 | B>B |
| chr2 | 40500000 | 40600000 | -0.15 | -0.18 | B>B |
| chr2 | 40600000 | 40700000 | -0.04 | -0.24 | B>B |
| chr2 | 40700000 | 40800000 | -0.20 | -0.32 | B>B |
| chr2 | 40800000 | 40900000 | -0.58 | -0.71 | B>B |
| chr2 | 40900000 | 41000000 | -0.61 | -0.81 | B>B |
| chr2 | 41000000 | 41100000 | -0.60 | -0.80 | B>B |
| chr2 | 41100000 | 41200000 | -0.66 | -0.89 | B>B |
| chr2 | 41200000 | 41300000 | -0.70 | -0.85 | B>B |
| chr2 | 41300000 | 41400000 | -0.63 | -0.91 | B>B |
| chr2 | 41400000 | 41500000 | -0.64 | -0.87 | B>B |
| chr2 | 41500000 | 41600000 | -0.58 | -0.86 | B>B |
| chr2 | 41600000 | 41700000 | -0.45 | -0.87 | B>B |
| chr2 | 41700000 | 41800000 | -0.46 | -0.79 | B>B |
| chr2 | 41800000 | 41900000 | -0.47 | -0.78 | B>B |
| chr2 | 41900000 | 42000000 | -0.20 | -0.47 | B>B |
| chr2 | 42000000 | 42100000 | 0.13 | 0.03 | A>A |
| chr2 | 42100000 | 42200000 | 0.61 | 0.39 | A>A |
| chr2 | 42200000 | 42300000 | 0.63 | 0.47 | A>A |
| chr2 | 42300000 | 42400000 | 0.57 | 0.50 | A>A |
| chr2 | 42400000 | 42500000 | 0.69 | 0.60 | A>A |
| chr2 | 42500000 | 42600000 | 0.64 | 0.53 | A>A |
| chr2 | 42600000 | 42700000 | 0.88 | 0.78 | A>A |
| chr2 | 42700000 | 42800000 | 1.01 | 0.89 | A>A |
| chr2 | 42800000 | 42900000 | 1.02 | 0.95 | A>A |
| chr2 | 42900000 | 43000000 | 0.93 | 0.94 | A>A |
| chr2 | 43000000 | 43100000 | 0.87 | 0.88 | A>A |
| chr2 | 43100000 | 43200000 | 0.88 | 0.96 | A>A |
| chr2 | 43200000 | 43300000 | 1.05 | 1.10 | A>A |
| chr2 | 43300000 | 43400000 | 1.03 | 1.10 | A>A |
| chr2 | 43400000 | 43500000 | 0.96 | 1.08 | A>A |
| chr2 | 43500000 | 43600000 | 1.13 | 1.22 | A>A |
| chr2 | 43600000 | 43700000 | 1.07 | 1.27 | A>A |
| chr2 | 43700000 | 43800000 | 0.78 | 0.82 | A>A |
| chr2 | 43800000 | 43900000 | 0.87 | 0.95 | A>A |
| chr2 | 43900000 | 44000000 | 1.23 | 1.24 | A>A |
| chr2 | 44000000 | 44100000 | 1.15 | 1.13 | A>A |
| chr2 | 44100000 | 44200000 | 1.00 | 1.06 | A>A |
| chr2 | 44200000 | 44300000 | 0.83 | 0.85 | A>A |
| chr2 | 44300000 | 44400000 | 1.09 | 1.15 | A>A |
| chr2 | 44400000 | 44500000 | 1.15 | 1.09 | A>A |
| chr2 | 44500000 | 44600000 | 0.97 | 0.95 | A>A |
| chr2 | 44600000 | 44700000 | 0.76 | 0.74 | A>A |
| chr2 | 44700000 | 44800000 | 0.74 | 0.90 | A>A |
| chr2 | 44800000 | 44900000 | 0.84 | 0.92 | A>A |
| chr2 | 44900000 | 45000000 | 0.68 | 0.77 | A>A |
| chr2 | 45000000 | 45100000 | 0.75 | 0.83 | A>A |
| chr2 | 45100000 | 45200000 | 0.73 | 0.75 | A>A |
| chr2 | 45200000 | 45300000 | 0.65 | 0.75 | A>A |
| chr2 | 45300000 | 45400000 | 0.82 | 0.86 | A>A |
| chr2 | 45400000 | 45500000 | 0.80 | 0.85 | A>A |
| chr2 | 45500000 | 45600000 | 0.72 | 0.97 | A>A |
| chr2 | 45600000 | 45700000 | 0.78 | 0.95 | A>A |
| chr2 | 45700000 | 45800000 | 0.72 | 0.81 | A>A |
| chr2 | 45800000 | 45900000 | 0.85 | 0.86 | A>A |

| | | | | | |
|---|---|---|---|---|---|
| chr2 | 45900000 | 46000000 | 1.03 | 1.03 | A>A |
| chr2 | 46000000 | 46100000 | 1.05 | 1.07 | A>A |
| chr2 | 46100000 | 46200000 | 0.84 | 0.94 | A>A |
| chr2 | 46200000 | 46300000 | 0.71 | 0.77 | A>A |
| chr2 | 46300000 | 46400000 | 1.04 | 1.18 | A>A |
| chr2 | 46400000 | 46500000 | 1.04 | 1.20 | A>A |
| chr2 | 46500000 | 46600000 | 0.93 | 0.92 | A>A |
| chr2 | 46600000 | 46700000 | 0.76 | 0.80 | A>A |
| chr2 | 46700000 | 46800000 | 1.01 | 1.02 | A>A |
| chr2 | 46800000 | 46900000 | 0.95 | 0.89 | A>A |
| chr2 | 46900000 | 47000000 | 1.17 | 1.20 | A>A |
| chr2 | 47000000 | 47100000 | 1.14 | 1.15 | A>A |
| chr2 | 47100000 | 47200000 | 0.90 | 0.86 | A>A |
| chr2 | 47200000 | 47300000 | 1.01 | 1.03 | A>A |
| chr2 | 47300000 | 47400000 | 1.20 | 1.09 | A>A |
| chr2 | 47400000 | 47500000 | 1.11 | 1.16 | A>A |
| chr2 | 47500000 | 47600000 | 0.85 | 0.82 | A>A |
| chr2 | 47600000 | 47700000 | 0.87 | 0.76 | A>A |
| chr2 | 47700000 | 47800000 | 0.76 | 0.58 | A>A |
| chr2 | 47800000 | 47900000 | 0.79 | 0.47 | A>A |
| chr2 | 47900000 | 48000000 | 0.67 | 0.40 | A>A |
| chr2 | 48000000 | 48100000 | 0.50 | 0.42 | A>A |
| chr2 | 48100000 | 48200000 | 0.47 | 0.08 | A>A |
| chr2 | 48200000 | 48300000 | 0.20 | -0.16 | A>A |
| chr2 | 48300000 | 48400000 | 0.24 | -0.07 | A>A |
| chr2 | 48400000 | 48500000 | 0.34 | 0.09 | A>A |
| chr2 | 48500000 | 48600000 | 0.53 | 0.37 | A>A |
| chr2 | 48600000 | 48700000 | 0.47 | 0.21 | A>A |
| chr2 | 48700000 | 48800000 | 0.25 | -0.08 | A>A |
| chr2 | 48800000 | 48900000 | -0.27 | -0.57 | B>B |
| chr2 | 48900000 | 49000000 | -0.58 | -0.80 | B>B |
| chr2 | 49000000 | 49100000 | -0.69 | -0.85 | B>B |
| chr2 | 49100000 | 49200000 | -0.60 | -0.85 | B>B |
| chr2 | 49200000 | 49300000 | -0.74 | -0.94 | B>B |
| chr2 | 49300000 | 49400000 | -0.88 | -0.93 | B>B |
| chr2 | 49400000 | 49500000 | -0.86 | -0.91 | B>B |
| chr2 | 49500000 | 49600000 | -0.86 | -0.93 | B>B |
| chr2 | 49600000 | 49700000 | -0.88 | -0.87 | B>B |
| chr2 | 49700000 | 49800000 | -0.84 | -0.94 | B>B |
| chr2 | 49800000 | 49900000 | -0.84 | -0.91 | B>B |
| chr2 | 49900000 | 50000000 | -0.86 | -0.92 | B>B |
| chr2 | 50000000 | 50100000 | -0.89 | -0.92 | B>B |
| chr2 | 50100000 | 50200000 | -0.91 | -0.94 | B>B |
| chr2 | 50200000 | 50300000 | -0.87 | -0.93 | B>B |
| chr2 | 50300000 | 50400000 | -0.73 | -0.90 | B>B |
| chr2 | 50400000 | 50500000 | -0.90 | -0.93 | B>B |
| chr2 | 50500000 | 50600000 | -0.85 | -0.88 | B>B |
| chr2 | 50600000 | 50700000 | -0.72 | -0.86 | B>B |
| chr2 | 50700000 | 50800000 | -0.67 | -0.70 | B>B |
| chr2 | 50800000 | 50900000 | -0.83 | -0.82 | B>B |
| chr2 | 50900000 | 51000000 | -0.86 | -0.87 | B>B |
| chr2 | 51000000 | 51100000 | -0.85 | -0.87 | B>B |
| chr2 | 51100000 | 51200000 | -0.90 | -0.86 | B>B |
| chr2 | 51200000 | 51300000 | -0.81 | -0.91 | B>B |
| chr2 | 51300000 | 51400000 | -0.83 | -0.87 | B>B |
| chr2 | 51400000 | 51500000 | -0.84 | -0.82 | B>B |

| | | | | | |
|---|---|---|---|---|---|
| chr2 | 51500000 | 51600000 | -0.87 | -0.84 | B>B |
| chr2 | 51600000 | 51700000 | -0.81 | -0.82 | B>B |
| chr2 | 51700000 | 51800000 | -0.82 | -0.81 | B>B |
| chr2 | 51800000 | 51900000 | -0.77 | -0.82 | B>B |
| chr2 | 51900000 | 52000000 | -0.85 | -0.77 | B>B |
| chr2 | 52000000 | 52100000 | -0.78 | -0.82 | B>B |
| chr2 | 52100000 | 52200000 | -0.80 | -0.81 | B>B |
| chr2 | 52200000 | 52300000 | -0.80 | -0.82 | B>B |
| chr2 | 52300000 | 52400000 | -0.84 | -0.76 | B>B |
| chr2 | 52400000 | 52500000 | -0.77 | -0.77 | B>B |
| chr2 | 52500000 | 52600000 | -0.81 | -0.70 | B>B |
| chr2 | 52600000 | 52700000 | -0.81 | -0.68 | B>B |
| chr2 | 52700000 | 52800000 | -0.85 | -0.66 | B>B |
| chr2 | 52800000 | 52900000 | -0.79 | -0.67 | B>B |
| chr2 | 52900000 | 53000000 | -0.76 | -0.66 | B>B |
| chr2 | 53000000 | 53100000 | -0.73 | -0.48 | B>B |
| chr2 | 53100000 | 53200000 | -0.73 | -0.27 | B>B |
| chr2 | 53200000 | 53300000 | -0.71 | -0.31 | B>B |
| chr2 | 53300000 | 53400000 | -0.71 | -0.27 | B>B |
| chr2 | 53400000 | 53500000 | -0.80 | -0.44 | B>B |
| chr2 | 53500000 | 53600000 | -0.75 | -0.44 | B>B |
| chr2 | 53600000 | 53700000 | -0.69 | -0.43 | B>B |
| chr2 | 53700000 | 53800000 | -0.55 | -0.44 | B>B |
| chr2 | 53800000 | 53900000 | -0.37 | -0.40 | B>B |
| chr2 | 53900000 | 54000000 | -0.04 | -0.11 | B>B |
| chr2 | 54000000 | 54100000 | 0.31 | 0.44 | A>A |
| chr2 | 54100000 | 54200000 | 0.60 | 0.67 | A>A |
| chr2 | 54200000 | 54300000 | 0.50 | 0.77 | A>A |
| chr2 | 54300000 | 54400000 | 0.28 | 0.46 | A>A |
| chr2 | 54400000 | 54500000 | 0.30 | 0.45 | A>A |
| chr2 | 54500000 | 54600000 | 0.39 | 0.51 | A>A |
| chr2 | 54600000 | 54700000 | 0.53 | 0.63 | A>A |
| chr2 | 54700000 | 54800000 | 0.55 | 0.75 | A>A |
| chr2 | 54800000 | 54900000 | 0.70 | 0.81 | A>A |
| chr2 | 54900000 | 55000000 | 0.46 | 0.55 | A>A |
| chr2 | 55000000 | 55100000 | 0.55 | 0.55 | A>A |
| chr2 | 55100000 | 55200000 | 0.70 | 0.78 | A>A |
| chr2 | 55200000 | 55300000 | 0.85 | 0.82 | A>A |
| chr2 | 55300000 | 55400000 | 0.96 | 1.01 | A>A |
| chr2 | 55400000 | 55500000 | 0.67 | 0.75 | A>A |
| chr2 | 55500000 | 55600000 | 0.90 | 0.86 | A>A |
| chr2 | 55600000 | 55700000 | 0.92 | 0.90 | A>A |
| chr2 | 55700000 | 55800000 | 0.79 | 0.80 | A>A |
| chr2 | 55800000 | 55900000 | 0.68 | 0.72 | A>A |
| chr2 | 55900000 | 56000000 | 0.22 | 0.27 | A>A |
| chr2 | 56000000 | 56100000 | 0.51 | 0.58 | A>A |
| chr2 | 56100000 | 56200000 | 0.42 | 0.58 | A>A |
| chr2 | 56200000 | 56300000 | 0.07 | 0.17 | A>A |
| chr2 | 56300000 | 56400000 | -0.25 | -0.12 | B>B |
| chr2 | 56400000 | 56500000 | -0.49 | -0.23 | B>B |
| chr2 | 56500000 | 56600000 | -0.52 | -0.24 | B>B |
| chr2 | 56600000 | 56700000 | -0.51 | -0.17 | B>B |
| chr2 | 56700000 | 56800000 | -0.47 | -0.21 | B>B |
| chr2 | 56800000 | 56900000 | -0.57 | -0.16 | B>B |
| chr2 | 56900000 | 57000000 | -0.31 | -0.15 | B>B |
| chr2 | 57000000 | 57100000 | -0.47 | -0.14 | B>B |

| | | | | | |
|---|---|---|---|---|---|
| chr2 | 57100000 | 57200000 | -0.58 | -0.26 | B>B |
| chr2 | 57200000 | 57300000 | -0.64 | -0.32 | B>B |
| chr2 | 57300000 | 57400000 | -0.61 | -0.23 | B>B |
| chr2 | 57400000 | 57500000 | -0.73 | -0.21 | B>A |
| chr2 | 57500000 | 57600000 | -0.75 | -0.27 | B>B |
| chr2 | 57600000 | 57700000 | -0.62 | -0.28 | B>B |
| chr2 | 57700000 | 57800000 | -0.47 | -0.29 | B>B |
| chr2 | 57800000 | 57900000 | -0.65 | -0.35 | B>B |
| chr2 | 57900000 | 58000000 | -0.36 | -0.25 | B>B |
| chr2 | 58000000 | 58100000 | 0.02 | -0.05 | B>B |
| chr2 | 58100000 | 58200000 | 0.10 | 0.05 | A>A |
| chr2 | 58200000 | 58300000 | 0.29 | 0.11 | A>A |
| chr2 | 58300000 | 58400000 | 0.19 | 0.16 | A>A |
| chr2 | 58400000 | 58500000 | 0.18 | -0.02 | A>A |
| chr2 | 58500000 | 58600000 | -0.08 | -0.43 | B>B |
| chr2 | 58600000 | 58700000 | -0.16 | -0.41 | B>B |
| chr2 | 58700000 | 58800000 | -0.14 | -0.32 | B>B |
| chr2 | 58800000 | 58900000 | -0.23 | -0.24 | B>B |
| chr2 | 58900000 | 59000000 | -0.40 | -0.44 | B>B |
| chr2 | 59000000 | 59100000 | -0.59 | -0.67 | B>B |
| chr2 | 59100000 | 59200000 | -0.70 | -0.74 | B>B |
| chr2 | 59200000 | 59300000 | -0.71 | -0.76 | B>B |
| chr2 | 59300000 | 59400000 | -0.72 | -0.76 | B>B |
| chr2 | 59400000 | 59500000 | -0.78 | -0.76 | B>B |
| chr2 | 59500000 | 59600000 | -0.75 | -0.74 | B>B |
| chr2 | 59600000 | 59700000 | -0.82 | -0.73 | B>B |
| chr2 | 59700000 | 59800000 | -0.78 | -0.76 | B>B |
| chr2 | 59800000 | 59900000 | -0.80 | -0.79 | B>B |
| chr2 | 59900000 | 60000000 | -0.73 | -0.76 | B>B |
| chr2 | 60000000 | 60100000 | -0.72 | -0.80 | B>B |
| chr2 | 60100000 | 60200000 | -0.69 | -0.80 | B>B |
| chr2 | 60200000 | 60300000 | -0.69 | -0.79 | B>B |
| chr2 | 60300000 | 60400000 | -0.63 | -0.77 | B>B |
| chr2 | 60400000 | 60500000 | -0.55 | -0.80 | B>B |
| chr2 | 60500000 | 60600000 | -0.35 | -0.73 | B>B |
| chr2 | 60600000 | 60700000 | -0.27 | -0.65 | B>B |
| chr2 | 60700000 | 60800000 | -0.17 | -0.44 | B>B |
| chr2 | 60800000 | 60900000 | -0.04 | -0.23 | B>B |
| chr2 | 60900000 | 61000000 | 0.40 | 0.15 | A>A |
| chr2 | 61000000 | 61100000 | 0.72 | 0.56 | A>A |
| chr2 | 61100000 | 61200000 | 0.73 | 0.68 | A>A |
| chr2 | 61200000 | 61300000 | 0.64 | 0.62 | A>A |
| chr2 | 61300000 | 61400000 | 0.83 | 0.83 | A>A |
| chr2 | 61400000 | 61500000 | 1.02 | 1.02 | A>A |
| chr2 | 61500000 | 61600000 | 1.09 | 1.11 | A>A |
| chr2 | 61600000 | 61700000 | 1.12 | 1.05 | A>A |
| chr2 | 61700000 | 61800000 | 0.97 | 1.05 | A>A |
| chr2 | 61800000 | 61900000 | 1.00 | 1.06 | A>A |
| chr2 | 61900000 | 62000000 | 1.02 | 1.03 | A>A |
| chr2 | 62000000 | 62100000 | 0.88 | 0.94 | A>A |
| chr2 | 62100000 | 62200000 | 0.57 | 0.74 | A>A |
| chr2 | 62200000 | 62300000 | 0.50 | 0.80 | A>A |
| chr2 | 62300000 | 62400000 | 0.60 | 0.85 | A>A |
| chr2 | 62400000 | 62500000 | 0.58 | 0.81 | A>A |
| chr2 | 62500000 | 62600000 | 0.54 | 0.72 | A>A |
| chr2 | 62600000 | 62700000 | 0.39 | 0.57 | A>A |

| | | | | | |
|---|---|---|---|---|---|
| chr2 | 62700000 | 62800000 | 0.80 | 0.92 | A>A |
| chr2 | 62800000 | 62900000 | 0.85 | 0.98 | A>A |
| chr2 | 62900000 | 63000000 | 0.72 | 0.92 | A>A |
| chr2 | 63000000 | 63100000 | 0.36 | 0.69 | A>A |
| chr2 | 63100000 | 63200000 | 0.27 | 0.66 | A>A |
| chr2 | 63200000 | 63300000 | 0.24 | 0.61 | A>A |
| chr2 | 63300000 | 63400000 | 0.09 | 0.62 | B>A |
| chr2 | 63400000 | 63500000 | -0.04 | 0.53 | B>A |
| chr2 | 63500000 | 63600000 | -0.02 | 0.66 | B>A |
| chr2 | 63600000 | 63700000 | 0.11 | 0.61 | A>A |
| chr2 | 63700000 | 63800000 | 0.20 | 0.53 | A>A |
| chr2 | 63800000 | 63900000 | 0.36 | 0.64 | A>A |
| chr2 | 63900000 | 64000000 | 0.42 | 0.70 | A>A |
| chr2 | 64000000 | 64100000 | 0.55 | 0.73 | A>A |
| chr2 | 64100000 | 64200000 | 0.49 | 0.71 | A>A |
| chr2 | 64200000 | 64300000 | 0.50 | 0.52 | A>A |
| chr2 | 64300000 | 64400000 | 0.63 | 0.79 | A>A |
| chr2 | 64400000 | 64500000 | 0.59 | 0.72 | A>A |
| chr2 | 64500000 | 64600000 | 0.46 | 0.64 | A>A |
| chr2 | 64600000 | 64700000 | 0.63 | 0.77 | A>A |
| chr2 | 64700000 | 64800000 | 0.60 | 0.67 | A>A |
| chr2 | 64800000 | 64900000 | 0.70 | 0.79 | A>A |
| chr2 | 64900000 | 65000000 | 1.06 | 1.00 | A>A |
| chr2 | 65000000 | 65100000 | 0.92 | 0.89 | A>A |
| chr2 | 65100000 | 65200000 | 0.66 | 0.66 | A>A |
| chr2 | 65200000 | 65300000 | 0.76 | 0.72 | A>A |
| chr2 | 65300000 | 65400000 | 0.81 | 0.74 | A>A |
| chr2 | 65400000 | 65500000 | 0.55 | 0.46 | A>A |
| chr2 | 65500000 | 65600000 | 0.50 | 0.44 | A>A |
| chr2 | 65600000 | 65700000 | 0.59 | 0.46 | A>A |
| chr2 | 65700000 | 65800000 | 0.45 | 0.26 | A>A |
| chr2 | 65800000 | 65900000 | 0.06 | -0.21 | B>B |
| chr2 | 65900000 | 66000000 | -0.34 | -0.66 | B>B |
| chr2 | 66000000 | 66100000 | -0.46 | -0.75 | B>B |
| chr2 | 66100000 | 66200000 | -0.52 | -0.78 | B>B |
| chr2 | 66200000 | 66300000 | -0.51 | -0.80 | B>B |
| chr2 | 66300000 | 66400000 | -0.60 | -0.79 | B>B |
| chr2 | 66400000 | 66500000 | -0.55 | -0.71 | B>B |
| chr2 | 66500000 | 66600000 | -0.51 | -0.69 | B>B |
| chr2 | 66600000 | 66700000 | -0.25 | -0.40 | B>B |
| chr2 | 66700000 | 66800000 | -0.13 | -0.25 | B>B |
| chr2 | 66800000 | 66900000 | -0.47 | -0.47 | B>B |
| chr2 | 66900000 | 67000000 | -0.56 | -0.60 | B>B |
| chr2 | 67000000 | 67100000 | -0.58 | -0.57 | B>B |
| chr2 | 67100000 | 67200000 | -0.54 | -0.49 | B>B |
| chr2 | 67200000 | 67300000 | -0.55 | -0.34 | B>B |
| chr2 | 67300000 | 67400000 | -0.50 | -0.17 | B>B |
| chr2 | 67400000 | 67500000 | -0.36 | -0.13 | B>B |
| chr2 | 67500000 | 67600000 | -0.20 | -0.03 | B>B |
| chr2 | 67600000 | 67700000 | 0.01 | 0.20 | A>A |
| chr2 | 67700000 | 67800000 | -0.10 | 0.07 | B>B |
| chr2 | 67800000 | 67900000 | -0.13 | -0.10 | B>B |
| chr2 | 67900000 | 68000000 | -0.36 | -0.42 | B>B |
| chr2 | 68000000 | 68100000 | -0.30 | -0.41 | B>B |
| chr2 | 68100000 | 68200000 | -0.23 | -0.55 | B>B |
| chr2 | 68200000 | 68300000 | -0.12 | -0.34 | B>B |

| | | | | | |
|---|---|---|---|---|---|
| chr2 | 68300000 | 68400000 | 0.39 | 0.07 | A>A |
| chr2 | 68400000 | 68500000 | 0.60 | 0.26 | A>A |
| chr2 | 68500000 | 68600000 | 0.35 | -0.03 | A>A |
| chr2 | 68600000 | 68700000 | 0.32 | -0.08 | A>A |
| chr2 | 68700000 | 68800000 | 0.29 | 0.11 | A>A |
| chr2 | 68800000 | 68900000 | 0.24 | 0.11 | A>A |
| chr2 | 68900000 | 69000000 | 0.29 | 0.10 | A>A |
| chr2 | 69000000 | 69100000 | 0.56 | 0.11 | A>A |
| chr2 | 69100000 | 69200000 | 0.53 | 0.11 | A>A |
| chr2 | 69200000 | 69300000 | 0.66 | 0.44 | A>A |
| chr2 | 69300000 | 69400000 | 0.68 | 0.65 | A>A |
| chr2 | 69400000 | 69500000 | 0.73 | 0.61 | A>A |
| chr2 | 69500000 | 69600000 | 0.72 | 0.55 | A>A |
| chr2 | 69600000 | 69700000 | 0.98 | 0.88 | A>A |
| chr2 | 69700000 | 69800000 | 0.98 | 0.99 | A>A |
| chr2 | 69800000 | 69900000 | 1.14 | 1.01 | A>A |
| chr2 | 69900000 | 70000000 | 0.99 | 1.01 | A>A |
| chr2 | 70000000 | 70100000 | 1.10 | 1.08 | A>A |
| chr2 | 70100000 | 70200000 | 1.03 | 1.06 | A>A |
| chr2 | 70200000 | 70300000 | 1.09 | 1.08 | A>A |
| chr2 | 70300000 | 70400000 | 1.09 | 1.19 | A>A |
| chr2 | 70400000 | 70500000 | 1.09 | 1.11 | A>A |
| chr2 | 70500000 | 70600000 | 0.83 | 0.86 | A>A |
| chr2 | 70600000 | 70700000 | 1.14 | 1.07 | A>A |
| chr2 | 70700000 | 70800000 | 1.09 | 1.20 | A>A |
| chr2 | 70800000 | 70900000 | 0.76 | 0.86 | A>A |
| chr2 | 70900000 | 71000000 | 1.10 | 1.11 | A>A |
| chr2 | 71000000 | 71100000 | 0.94 | 1.00 | A>A |
| chr2 | 71100000 | 71200000 | 1.13 | 1.03 | A>A |
| chr2 | 71200000 | 71300000 | 1.06 | 0.97 | A>A |
| chr2 | 71300000 | 71400000 | 0.98 | 1.03 | A>A |
| chr2 | 71400000 | 71500000 | 0.96 | 0.95 | A>A |
| chr2 | 71500000 | 71600000 | 0.98 | 1.10 | A>A |
| chr2 | 71600000 | 71700000 | 1.09 | 1.06 | A>A |
| chr2 | 71700000 | 71800000 | 0.57 | 0.60 | A>A |
| chr2 | 71800000 | 71900000 | 0.58 | 0.74 | A>A |
| chr2 | 71900000 | 72000000 | 0.75 | 0.94 | A>A |
| chr2 | 72000000 | 72100000 | 0.59 | 0.75 | A>A |
| chr2 | 72100000 | 72200000 | 0.56 | 0.76 | A>A |
| chr2 | 72200000 | 72300000 | 0.61 | 0.97 | A>A |
| chr2 | 72300000 | 72400000 | 0.63 | 0.78 | A>A |
| chr2 | 72400000 | 72500000 | 0.71 | 0.89 | A>A |
| chr2 | 72500000 | 72600000 | 0.73 | 0.98 | A>A |
| chr2 | 72600000 | 72700000 | 0.59 | 0.88 | A>A |
| chr2 | 72700000 | 72800000 | 0.65 | 0.98 | A>A |
| chr2 | 72800000 | 72900000 | 0.63 | 0.90 | A>A |
| chr2 | 72900000 | 73000000 | 0.72 | 0.90 | A>A |
| chr2 | 73000000 | 73100000 | 0.83 | 0.93 | A>A |
| chr2 | 73100000 | 73200000 | 0.88 | 0.88 | A>A |
| chr2 | 73200000 | 73300000 | 1.11 | 1.08 | A>A |
| chr2 | 73300000 | 73400000 | 0.95 | 1.00 | A>A |
| chr2 | 73400000 | 73500000 | 0.91 | 0.93 | A>A |
| chr2 | 73500000 | 73600000 | 0.83 | 0.85 | A>A |
| chr2 | 73600000 | 73700000 | 0.80 | 0.84 | A>A |
| chr2 | 73700000 | 73800000 | 0.71 | 0.82 | A>A |
| chr2 | 73800000 | 73900000 | 0.69 | 0.75 | A>A |

| | | | | | |
|---|---|---|---|---|---|
| chr2 | 73900000 | 74000000 | 0.85 | 0.90 | A>A |
| chr2 | 74000000 | 74100000 | 0.87 | 0.90 | A>A |
| chr2 | 74100000 | 74200000 | 0.98 | 1.05 | A>A |
| chr2 | 74200000 | 74300000 | 1.03 | 1.12 | A>A |
| chr2 | 74300000 | 74400000 | 0.89 | 1.01 | A>A |
| chr2 | 74400000 | 74500000 | 1.01 | 1.00 | A>A |
| chr2 | 74500000 | 74600000 | 1.01 | 0.98 | A>A |
| chr2 | 74600000 | 74700000 | 1.04 | 1.06 | A>A |
| chr2 | 74700000 | 74800000 | 1.00 | 0.88 | A>A |
| chr2 | 74800000 | 74900000 | 0.78 | 0.66 | A>A |
| chr2 | 74900000 | 75000000 | 0.57 | 0.47 | A>A |
| chr2 | 75000000 | 75100000 | 0.59 | 0.51 | A>A |
| chr2 | 75100000 | 75200000 | 0.39 | 0.24 | A>A |
| chr2 | 75200000 | 75300000 | 0.34 | 0.11 | A>A |
| chr2 | 75300000 | 75400000 | 0.09 | -0.12 | B>B |
| chr2 | 75400000 | 75500000 | 0.24 | -0.08 | A>A |
| chr2 | 75500000 | 75600000 | 0.11 | -0.22 | B>B |
| chr2 | 75600000 | 75700000 | -0.03 | -0.15 | B>B |
| chr2 | 75700000 | 75800000 | -0.03 | -0.11 | B>B |
| chr2 | 75800000 | 75900000 | 0.07 | -0.08 | B>B |
| chr2 | 75900000 | 76000000 | -0.24 | -0.42 | B>B |
| chr2 | 76000000 | 76100000 | -0.67 | -0.68 | B>B |
| chr2 | 76100000 | 76200000 | -0.81 | -0.71 | B>B |
| chr2 | 76200000 | 76300000 | -0.85 | -0.61 | B>B |
| chr2 | 76300000 | 76400000 | -0.80 | -0.52 | B>B |
| chr2 | 76400000 | 76500000 | -0.78 | -0.47 | B>B |
| chr2 | 76500000 | 76600000 | -0.76 | -0.45 | B>B |
| chr2 | 76600000 | 76700000 | -0.76 | -0.46 | B>B |
| chr2 | 76700000 | 76800000 | -0.77 | -0.52 | B>B |
| chr2 | 76800000 | 76900000 | -0.73 | -0.62 | B>B |
| chr2 | 76900000 | 77000000 | -0.77 | -0.68 | B>B |
| chr2 | 77000000 | 77100000 | -0.77 | -0.78 | B>B |
| chr2 | 77100000 | 77200000 | -0.72 | -0.72 | B>B |
| chr2 | 77200000 | 77300000 | -0.76 | -0.77 | B>B |
| chr2 | 77300000 | 77400000 | -0.75 | -0.80 | B>B |
| chr2 | 77400000 | 77500000 | -0.74 | -0.77 | B>B |
| chr2 | 77500000 | 77600000 | -0.74 | -0.78 | B>B |
| chr2 | 77600000 | 77700000 | -0.77 | -0.76 | B>B |
| chr2 | 77700000 | 77800000 | -0.77 | -0.72 | B>B |
| chr2 | 77800000 | 77900000 | -0.74 | -0.78 | B>B |
| chr2 | 77900000 | 78000000 | -0.66 | -0.76 | B>B |
| chr2 | 78000000 | 78100000 | -0.72 | -0.73 | B>B |
| chr2 | 78100000 | 78200000 | -0.74 | -0.77 | B>B |
| chr2 | 78200000 | 78300000 | -0.71 | -0.77 | B>B |
| chr2 | 78300000 | 78400000 | -0.72 | -0.77 | B>B |
| chr2 | 78400000 | 78500000 | -0.72 | -0.78 | B>B |
| chr2 | 78500000 | 78600000 | -0.70 | -0.76 | B>B |
| chr2 | 78600000 | 78700000 | -0.74 | -0.73 | B>B |
| chr2 | 78700000 | 78800000 | -0.71 | -0.79 | B>B |
| chr2 | 78800000 | 78900000 | -0.72 | -0.75 | B>B |
| chr2 | 78900000 | 79000000 | -0.68 | -0.77 | B>B |
| chr2 | 79000000 | 79100000 | -0.68 | -0.77 | B>B |
| chr2 | 79100000 | 79200000 | -0.67 | -0.75 | B>B |
| chr2 | 79200000 | 79300000 | -0.63 | -0.73 | B>B |
| chr2 | 79300000 | 79400000 | -0.64 | -0.76 | B>B |
| chr2 | 79400000 | 79500000 | -0.62 | -0.74 | B>B |

| | | | | | |
|---|---|---|---|---|---|
| chr2 | 79500000 | 79600000 | -0.65 | -0.78 | B>B |
| chr2 | 79600000 | 79700000 | -0.72 | -0.84 | B>B |
| chr2 | 79700000 | 79800000 | -0.66 | -0.77 | B>B |
| chr2 | 79800000 | 79900000 | -0.54 | -0.82 | B>B |
| chr2 | 79900000 | 80000000 | -0.62 | -0.80 | B>B |
| chr2 | 80000000 | 80100000 | -0.59 | -0.72 | B>B |
| chr2 | 80100000 | 80200000 | -0.59 | -0.69 | B>B |
| chr2 | 80200000 | 80300000 | -0.66 | -0.75 | B>B |
| chr2 | 80300000 | 80400000 | -0.63 | -0.76 | B>B |
| chr2 | 80400000 | 80500000 | -0.68 | -0.78 | B>B |
| chr2 | 80500000 | 80600000 | -0.64 | -0.78 | B>B |
| chr2 | 80600000 | 80700000 | -0.69 | -0.81 | B>B |
| chr2 | 80700000 | 80800000 | -0.70 | -0.81 | B>B |
| chr2 | 80800000 | 80900000 | -0.65 | -0.79 | B>B |
| chr2 | 80900000 | 81000000 | -0.66 | -0.76 | B>B |
| chr2 | 81000000 | 81100000 | -0.68 | -0.83 | B>B |
| chr2 | 81100000 | 81200000 | -0.68 | -0.70 | B>B |
| chr2 | 81200000 | 81300000 | -0.65 | -0.63 | B>B |
| chr2 | 81300000 | 81400000 | -0.66 | -0.58 | B>B |
| chr2 | 81400000 | 81500000 | -0.71 | -0.46 | B>B |
| chr2 | 81500000 | 81600000 | -0.69 | -0.24 | B>B |
| chr2 | 81600000 | 81700000 | -0.69 | -0.11 | B>A |
| chr2 | 81700000 | 81800000 | -0.70 | 0.09 | B>A |
| chr2 | 81800000 | 81900000 | -0.63 | 0.03 | B>A |
| chr2 | 81900000 | 82000000 | -0.66 | 0.01 | B>A |
| chr2 | 82000000 | 82100000 | -0.65 | 0.03 | B>A |
| chr2 | 82100000 | 82200000 | -0.64 | -0.08 | B>A |
| chr2 | 82200000 | 82300000 | -0.61 | -0.07 | B>A |
| chr2 | 82300000 | 82400000 | -0.65 | -0.07 | B>A |
| chr2 | 82400000 | 82500000 | -0.60 | 0.00 | B>A |
| chr2 | 82500000 | 82600000 | -0.56 | 0.13 | B>A |
| chr2 | 82600000 | 82700000 | -0.60 | 0.17 | B>A |
| chr2 | 82700000 | 82800000 | -0.60 | 0.19 | B>A |
| chr2 | 82800000 | 82900000 | -0.58 | 0.14 | B>A |
| chr2 | 82900000 | 83000000 | -0.54 | 0.26 | B>A |
| chr2 | 83000000 | 83100000 | -0.50 | 0.32 | B>A |
| chr2 | 83100000 | 83200000 | -0.59 | 0.22 | B>A |
| chr2 | 83200000 | 83300000 | -0.54 | 0.27 | B>A |
| chr2 | 83300000 | 83400000 | -0.55 | 0.26 | B>A |
| chr2 | 83400000 | 83500000 | -0.58 | 0.10 | B>A |
| chr2 | 83500000 | 83600000 | -0.60 | 0.17 | B>A |
| chr2 | 83600000 | 83700000 | -0.64 | 0.13 | B>A |
| chr2 | 83700000 | 83800000 | -0.63 | 0.08 | B>A |
| chr2 | 83800000 | 83900000 | -0.67 | 0.08 | B>A |
| chr2 | 83900000 | 84000000 | -0.66 | 0.07 | B>A |
| chr2 | 84000000 | 84100000 | -0.65 | -0.05 | B>A |
| chr2 | 84100000 | 84200000 | -0.64 | -0.16 | B>B |
| chr2 | 84200000 | 84300000 | -0.58 | -0.10 | B>B |
| chr2 | 84300000 | 84400000 | -0.59 | -0.07 | B>A |
| chr2 | 84400000 | 84500000 | -0.55 | -0.10 | B>B |
| chr2 | 84500000 | 84600000 | -0.46 | -0.25 | B>B |
| chr2 | 84600000 | 84700000 | -0.22 | -0.25 | B>B |
| chr2 | 84700000 | 84800000 | -0.12 | -0.27 | B>B |
| chr2 | 84800000 | 84900000 | -0.23 | -0.41 | B>B |
| chr2 | 84900000 | 85000000 | -0.06 | -0.34 | B>B |
| chr2 | 85000000 | 85100000 | 0.09 | -0.19 | B>B |

| | | | | | |
|---|---|---|---|---|---|
| chr2 | 85100000 | 85200000 | 0.46 | 0.34 | A>A |
| chr2 | 85200000 | 85300000 | 0.70 | 0.56 | A>A |
| chr2 | 85300000 | 85400000 | 0.72 | 0.74 | A>A |
| chr2 | 85400000 | 85500000 | 0.82 | 0.84 | A>A |
| chr2 | 85500000 | 85600000 | 0.80 | 0.78 | A>A |
| chr2 | 85600000 | 85700000 | 1.16 | 1.20 | A>A |
| chr2 | 85700000 | 85800000 | 1.19 | 1.21 | A>A |
| chr2 | 85800000 | 85900000 | 1.06 | 1.01 | A>A |
| chr2 | 85900000 | 86000000 | 1.19 | 1.10 | A>A |
| chr2 | 86000000 | 86100000 | 0.93 | 1.02 | A>A |
| chr2 | 86100000 | 86200000 | 1.15 | 1.19 | A>A |
| chr2 | 86200000 | 86300000 | 1.20 | 1.10 | A>A |
| chr2 | 86300000 | 86400000 | 0.94 | 0.97 | A>A |
| chr2 | 86400000 | 86500000 | 1.03 | 1.11 | A>A |
| chr2 | 86500000 | 86600000 | 1.09 | 1.10 | A>A |
| chr2 | 86600000 | 86700000 | 0.86 | 0.93 | A>A |
| chr2 | 86700000 | 86800000 | 0.92 | 1.02 | A>A |
| chr2 | 86800000 | 86900000 | 0.93 | 0.98 | A>A |
| chr2 | 86900000 | 87000000 | 0.00 | 0.95 | NO |
| chr2 | 87000000 | 87100000 | 0.83 | 0.82 | A>A |
| chr2 | 87100000 | 87200000 | 0.00 | 0.00 | NO |
| chr2 | 87200000 | 87300000 | 0.00 | 0.00 | NO |
| chr2 | 87300000 | 87400000 | 0.00 | 0.00 | NO |
| chr2 | 87400000 | 87500000 | 0.00 | 0.00 | NO |
| chr2 | 87500000 | 87600000 | 0.00 | 0.00 | NO |
| chr2 | 87600000 | 87700000 | 0.00 | 0.00 | NO |
| chr2 | 87700000 | 87800000 | 0.00 | 0.00 | NO |
| chr2 | 87800000 | 87900000 | 0.00 | 0.00 | NO |
| chr2 | 87900000 | 88000000 | 0.00 | 0.00 | NO |
| chr2 | 88000000 | 88100000 | 0.00 | 0.00 | NO |
| chr2 | 88100000 | 88200000 | 0.00 | 0.00 | NO |
| chr2 | 88200000 | 88300000 | 0.00 | 0.00 | NO |
| chr2 | 88300000 | 88400000 | 0.44 | 0.20 | A>A |
| chr2 | 88400000 | 88500000 | 0.16 | -0.08 | A>A |
| chr2 | 88500000 | 88600000 | 0.04 | -0.23 | B>B |
| chr2 | 88600000 | 88700000 | -0.02 | -0.29 | B>B |
| chr2 | 88700000 | 88800000 | 0.16 | -0.11 | A>A |
| chr2 | 88800000 | 88900000 | 0.30 | 0.06 | A>A |
| chr2 | 88900000 | 89000000 | 0.41 | 0.17 | A>A |
| chr2 | 89000000 | 89100000 | 0.10 | -0.26 | B>B |
| chr2 | 89100000 | 89200000 | -0.29 | -0.82 | B>B |
| chr2 | 89200000 | 89300000 | 0.00 | -0.99 | NO |
| chr2 | 89300000 | 89400000 | 0.00 | 0.00 | NO |
| chr2 | 89400000 | 89500000 | 0.00 | 0.00 | NO |
| chr2 | 89500000 | 89600000 | 0.00 | 0.00 | NO |
| chr2 | 89600000 | 89700000 | 0.00 | 0.00 | NO |
| chr2 | 89700000 | 89800000 | 0.00 | 0.00 | NO |
| chr2 | 89800000 | 89900000 | 0.00 | 0.00 | NO |
| chr2 | 89900000 | 90000000 | 0.00 | 0.00 | NO |
| chr2 | 90000000 | 90100000 | 0.00 | 0.00 | NO |
| chr2 | 90100000 | 90200000 | 0.00 | 0.00 | NO |
| chr2 | 90200000 | 90300000 | 0.00 | 0.00 | NO |
| chr2 | 90300000 | 90400000 | 0.00 | 0.00 | NO |
| chr2 | 90400000 | 90500000 | 0.00 | 0.00 | NO |
| chr2 | 90500000 | 90600000 | 0.00 | 0.00 | NO |
| chr2 | 90600000 | 90700000 | 0.00 | 0.00 | NO |

| | | | | | |
|---|---|---|---|---|---|
| chr2 | 90700000 | 90800000 | 0.00 | 0.00 | NO |
| chr2 | 90800000 | 90900000 | 0.00 | 0.00 | NO |
| chr2 | 90900000 | 91000000 | 0.00 | 0.00 | NO |
| chr2 | 91000000 | 91100000 | 0.00 | 0.00 | NO |
| chr2 | 91100000 | 91200000 | 0.00 | 0.00 | NO |
| chr2 | 91200000 | 91300000 | 0.00 | 0.00 | NO |
| chr2 | 91300000 | 91400000 | 0.00 | 0.00 | NO |
| chr2 | 91400000 | 91500000 | 0.00 | 0.00 | NO |
| chr2 | 91500000 | 91600000 | 0.00 | 0.00 | NO |
| chr2 | 91600000 | 91700000 | 0.00 | 0.00 | NO |
| chr2 | 91700000 | 91800000 | -0.11 | -0.43 | B>B |
| chr2 | 91800000 | 91900000 | 0.00 | 0.00 | NO |
| chr2 | 91900000 | 92000000 | 0.00 | 0.00 | NO |
| chr2 | 92000000 | 92100000 | 0.00 | 0.00 | NO |
| chr2 | 92100000 | 92200000 | 0.00 | 0.00 | NO |
| chr2 | 92200000 | 92300000 | 0.00 | 0.00 | NO |
| chr2 | 92300000 | 92400000 | 0.00 | 0.00 | NO |
| chr2 | 92400000 | 92500000 | 0.00 | 0.00 | NO |
| chr2 | 92500000 | 92600000 | 0.00 | 0.00 | NO |
| chr2 | 92600000 | 92700000 | 0.00 | 0.00 | NO |
| chr2 | 92700000 | 92800000 | 0.00 | 0.00 | NO |
| chr2 | 92800000 | 92900000 | 0.00 | 0.00 | NO |
| chr2 | 92900000 | 93000000 | 0.00 | 0.00 | NO |
| chr2 | 93000000 | 93100000 | 0.00 | 0.00 | NO |
| chr2 | 93100000 | 93200000 | 0.00 | 0.00 | NO |
| chr2 | 93200000 | 93300000 | 0.00 | 0.00 | NO |
| chr2 | 93300000 | 93400000 | 0.00 | 0.00 | NO |
| chr2 | 93400000 | 93500000 | 0.00 | 0.00 | NO |
| chr2 | 93500000 | 93600000 | 0.00 | 0.00 | NO |
| chr2 | 93600000 | 93700000 | 0.00 | 0.00 | NO |
| chr2 | 93700000 | 93800000 | 0.00 | 0.00 | NO |
| chr2 | 93800000 | 93900000 | 0.00 | 0.00 | NO |
| chr2 | 93900000 | 94000000 | 0.00 | 0.00 | NO |
| chr2 | 94000000 | 94100000 | 0.00 | 0.00 | NO |
| chr2 | 94100000 | 94200000 | 0.00 | 0.00 | NO |
| chr2 | 94200000 | 94300000 | 0.00 | 0.00 | NO |
| chr2 | 94300000 | 94400000 | 0.00 | 0.00 | NO |
| chr2 | 94400000 | 94500000 | 0.00 | 0.00 | NO |
| chr2 | 94500000 | 94600000 | 0.00 | 0.00 | NO |
| chr2 | 94600000 | 94700000 | 0.00 | 0.00 | NO |
| chr2 | 94700000 | 94800000 | 0.00 | 0.00 | NO |
| chr2 | 94800000 | 94900000 | 0.00 | 0.00 | NO |
| chr2 | 94900000 | 95000000 | 0.00 | 0.00 | NO |
| chr2 | 95000000 | 95100000 | 0.00 | 0.00 | NO |
| chr2 | 95100000 | 95200000 | 0.00 | 0.00 | NO |
| chr2 | 95200000 | 95300000 | 0.00 | 0.00 | NO |
| chr2 | 95300000 | 95400000 | 0.00 | 0.00 | NO |
| chr2 | 95400000 | 95500000 | 0.00 | 0.00 | NO |
| chr2 | 95500000 | 95600000 | 0.00 | 0.00 | NO |
| chr2 | 95600000 | 95700000 | 0.11 | 0.19 | A>A |
| chr2 | 95700000 | 95800000 | 0.33 | 0.63 | A>A |
| chr2 | 95800000 | 95900000 | 0.15 | 0.94 | B>A |
| chr2 | 95900000 | 96000000 | 0.03 | 1.10 | B>A |
| chr2 | 96000000 | 96100000 | 0.23 | 0.86 | B>A |
| chr2 | 96100000 | 96200000 | 0.00 | 0.00 | NO |
| chr2 | 96200000 | 96300000 | 0.00 | 0.00 | NO |

| | | | | | |
|---|---|---|---|---|---|
| chr2 | 96300000 | 96400000 | 0.00 | 0.00 | NO |
| chr2 | 96400000 | 96500000 | 0.00 | 0.00 | NO |
| chr2 | 96500000 | 96600000 | 0.00 | 0.00 | NO |
| chr2 | 96600000 | 96700000 | 0.00 | 0.00 | NO |
| chr2 | 96700000 | 96800000 | 0.00 | 0.00 | NO |
| chr2 | 96800000 | 96900000 | 1.34 | 1.53 | A>A |
| chr2 | 96900000 | 97000000 | 1.48 | 1.62 | A>A |
| chr2 | 97000000 | 97100000 | 1.41 | 1.39 | A>A |
| chr2 | 97100000 | 97200000 | 1.19 | 1.40 | A>A |
| chr2 | 97200000 | 97300000 | 1.27 | 1.27 | A>A |
| chr2 | 97300000 | 97400000 | 1.70 | 1.76 | A>A |
| chr2 | 97400000 | 97500000 | 1.31 | 1.48 | A>A |
| chr2 | 97500000 | 97600000 | 1.45 | 1.44 | A>A |
| chr2 | 97600000 | 97700000 | 0.00 | 0.00 | NO |
| chr2 | 97700000 | 97800000 | 0.00 | 0.00 | NO |
| chr2 | 97800000 | 97900000 | 0.00 | 0.00 | NO |
| chr2 | 97900000 | 98000000 | 0.00 | 0.00 | NO |
| chr2 | 98000000 | 98100000 | 0.00 | 0.00 | NO |
| chr2 | 98100000 | 98200000 | 0.00 | 0.00 | NO |
| chr2 | 98200000 | 98300000 | 0.00 | 0.00 | NO |
| chr2 | 98300000 | 98400000 | 0.79 | 0.94 | A>A |
| chr2 | 98400000 | 98500000 | 0.82 | 0.94 | A>A |
| chr2 | 98500000 | 98600000 | 1.02 | 1.16 | A>A |
| chr2 | 98600000 | 98700000 | 0.79 | 0.85 | A>A |
| chr2 | 98700000 | 98800000 | 0.52 | 0.86 | A>A |
| chr2 | 98800000 | 98900000 | 0.47 | 0.90 | A>A |
| chr2 | 98900000 | 99000000 | 0.55 | 0.90 | A>A |
| chr2 | 99000000 | 99100000 | 0.79 | 0.86 | A>A |
| chr2 | 99100000 | 99200000 | 0.77 | 1.00 | A>A |
| chr2 | 99200000 | 99300000 | 0.61 | 0.70 | A>A |
| chr2 | 99300000 | 99400000 | 0.80 | 1.01 | A>A |
| chr2 | 99400000 | 99500000 | 0.52 | 0.86 | A>A |
| chr2 | 99500000 | 99600000 | 0.58 | 0.85 | A>A |
| chr2 | 99600000 | 99700000 | 0.33 | 0.61 | A>A |
| chr2 | 99700000 | 99800000 | 0.68 | 0.77 | A>A |
| chr2 | 99800000 | 99900000 | 0.68 | 0.85 | A>A |
| chr2 | 99900000 | 100000000 | 0.89 | 0.87 | A>A |
| chr2 | 1E+08 | 100100000 | 0.84 | 0.71 | A>A |
| chr2 | 1E+08 | 100200000 | 0.29 | 0.03 | A>A |
| chr2 | 1E+08 | 100300000 | 0.07 | -0.23 | B>B |
| chr2 | 1E+08 | 100400000 | 0.03 | -0.32 | B>B |
| chr2 | 1E+08 | 100500000 | -0.12 | -0.38 | B>B |
| chr2 | 1.01E+08 | 100600000 | -0.03 | -0.35 | B>B |
| chr2 | 1.01E+08 | 100700000 | -0.03 | -0.36 | B>B |
| chr2 | 1.01E+08 | 100800000 | 0.10 | -0.22 | B>B |
| chr2 | 1.01E+08 | 100900000 | 0.23 | -0.02 | A>A |
| chr2 | 1.01E+08 | 101000000 | 0.68 | 0.41 | A>A |
| chr2 | 1.01E+08 | 101100000 | 0.54 | 0.22 | A>A |
| chr2 | 1.01E+08 | 101200000 | 0.58 | 0.36 | A>A |
| chr2 | 1.01E+08 | 101300000 | 0.82 | 0.63 | A>A |
| chr2 | 1.01E+08 | 101400000 | 1.04 | 1.03 | A>A |
| chr2 | 1.01E+08 | 101500000 | 1.14 | 1.18 | A>A |
| chr2 | 1.02E+08 | 101600000 | 0.93 | 0.93 | A>A |
| chr2 | 1.02E+08 | 101700000 | 0.96 | 1.02 | A>A |
| chr2 | 1.02E+08 | 101800000 | 1.24 | 1.33 | A>A |
| chr2 | 1.02E+08 | 101900000 | 1.04 | 1.02 | A>A |

| | | | | | |
|---|---|---|---|---|---|
| chr2 | 1.02E+08 | 102000000 | 0.84 | 0.91 | A>A |
| chr2 | 1.02E+08 | 102100000 | 0.92 | 0.87 | A>A |
| chr2 | 1.02E+08 | 102200000 | 0.97 | 1.02 | A>A |
| chr2 | 1.02E+08 | 102300000 | 1.11 | 1.15 | A>A |
| chr2 | 1.02E+08 | 102400000 | 0.95 | 1.10 | A>A |
| chr2 | 1.02E+08 | 102500000 | 0.67 | 0.81 | A>A |
| chr2 | 1.03E+08 | 102600000 | 0.51 | 0.67 | A>A |
| chr2 | 1.03E+08 | 102700000 | 0.71 | 0.74 | A>A |
| chr2 | 1.03E+08 | 102800000 | 0.89 | 0.92 | A>A |
| chr2 | 1.03E+08 | 102900000 | 0.54 | 0.56 | A>A |
| chr2 | 1.03E+08 | 103000000 | 0.42 | 0.43 | A>A |
| chr2 | 1.03E+08 | 103100000 | 0.23 | 0.35 | A>A |
| chr2 | 1.03E+08 | 103200000 | 0.27 | 0.58 | A>A |
| chr2 | 1.03E+08 | 103300000 | 0.17 | 0.34 | A>A |
| chr2 | 1.03E+08 | 103400000 | 0.03 | 0.14 | A>A |
| chr2 | 1.03E+08 | 103500000 | -0.29 | -0.33 | B>B |
| chr2 | 1.04E+08 | 103600000 | -0.59 | -0.73 | B>B |
| chr2 | 1.04E+08 | 103700000 | -0.66 | -0.87 | B>B |
| chr2 | 1.04E+08 | 103800000 | -0.68 | -0.74 | B>B |
| chr2 | 1.04E+08 | 103900000 | -0.79 | -0.80 | B>B |
| chr2 | 1.04E+08 | 104000000 | -0.82 | -0.84 | B>B |
| chr2 | 1.04E+08 | 104100000 | -0.79 | -0.77 | B>B |
| chr2 | 1.04E+08 | 104200000 | -0.79 | -0.83 | B>B |
| chr2 | 1.04E+08 | 104300000 | -0.80 | -0.87 | B>B |
| chr2 | 1.04E+08 | 104400000 | -0.88 | -0.78 | B>B |
| chr2 | 1.04E+08 | 104500000 | -0.81 | -0.66 | B>B |
| chr2 | 1.05E+08 | 104600000 | -0.84 | -0.47 | B>B |
| chr2 | 1.05E+08 | 104700000 | -0.84 | -0.11 | B>A |
| chr2 | 1.05E+08 | 104800000 | -0.82 | 0.27 | B>A |
| chr2 | 1.05E+08 | 104900000 | -0.80 | 0.19 | B>A |
| chr2 | 1.05E+08 | 105000000 | -0.79 | 0.17 | B>A |
| chr2 | 1.05E+08 | 105100000 | -0.77 | -0.05 | B>A |
| chr2 | 1.05E+08 | 105200000 | -0.75 | -0.44 | B>B |
| chr2 | 1.05E+08 | 105300000 | -0.70 | -0.55 | B>B |
| chr2 | 1.05E+08 | 105400000 | -0.55 | -0.52 | B>B |
| chr2 | 1.05E+08 | 105500000 | -0.43 | -0.40 | B>B |
| chr2 | 1.06E+08 | 105600000 | -0.30 | -0.29 | B>B |
| chr2 | 1.06E+08 | 105700000 | -0.02 | -0.10 | B>B |
| chr2 | 1.06E+08 | 105800000 | 0.22 | 0.03 | A>A |
| chr2 | 1.06E+08 | 105900000 | 0.38 | 0.16 | A>A |
| chr2 | 1.06E+08 | 106000000 | 0.69 | 0.43 | A>A |
| chr2 | 1.06E+08 | 106100000 | 0.99 | 0.74 | A>A |
| chr2 | 1.06E+08 | 106200000 | 0.65 | 0.43 | A>A |
| chr2 | 1.06E+08 | 106300000 | 0.68 | 0.38 | A>A |
| chr2 | 1.06E+08 | 106400000 | 0.75 | 0.40 | A>A |
| chr2 | 1.06E+08 | 106500000 | 0.66 | 0.36 | A>A |
| chr2 | 1.07E+08 | 106600000 | 0.44 | 0.09 | A>A |
| chr2 | 1.07E+08 | 106700000 | 0.57 | -0.10 | A>B |
| chr2 | 1.07E+08 | 106800000 | 0.59 | 0.16 | A>A |
| chr2 | 1.07E+08 | 106900000 | 0.00 | -0.02 | NO |
| chr2 | 1.07E+08 | 107000000 | 0.44 | -0.27 | A>B |
| chr2 | 1.07E+08 | 107100000 | 0.00 | 0.00 | NO |
| chr2 | 1.07E+08 | 107200000 | 0.43 | -0.10 | A>A |
| chr2 | 1.07E+08 | 107300000 | 0.15 | -0.37 | B>B |
| chr2 | 1.07E+08 | 107400000 | 0.14 | -0.51 | A>B |
| chr2 | 1.07E+08 | 107500000 | 0.29 | -0.38 | A>B |

| | | | | | |
|---|---|---|---|---|---|
| chr2 | 1.08E+08 | 107600000 | 0.51 | -0.32 | A>B |
| chr2 | 1.08E+08 | 107700000 | 0.66 | -0.25 | A>B |
| chr2 | 1.08E+08 | 107800000 | 0.40 | -0.43 | A>B |
| chr2 | 1.08E+08 | 107900000 | 0.34 | -0.55 | A>B |
| chr2 | 1.08E+08 | 108000000 | 0.17 | -0.49 | A>B |
| chr2 | 1.08E+08 | 108100000 | 0.18 | -0.54 | A>B |
| chr2 | 1.08E+08 | 108200000 | 0.13 | -0.39 | B>B |
| chr2 | 1.08E+08 | 108300000 | 0.13 | -0.24 | B>B |
| chr2 | 1.08E+08 | 108400000 | 0.13 | -0.28 | B>B |
| chr2 | 1.08E+08 | 108500000 | 0.00 | 0.00 | NO |
| chr2 | 1.09E+08 | 108600000 | 0.00 | 0.00 | NO |
| chr2 | 1.09E+08 | 108700000 | 0.04 | -0.50 | B>B |
| chr2 | 1.09E+08 | 108800000 | 0.19 | -0.37 | B>B |
| chr2 | 1.09E+08 | 108900000 | 0.70 | 0.42 | A>A |
| chr2 | 1.09E+08 | 109000000 | 0.98 | 0.72 | A>A |
| chr2 | 1.09E+08 | 109100000 | 1.09 | 0.85 | A>A |
| chr2 | 1.09E+08 | 109200000 | 0.00 | 0.00 | NO |
| chr2 | 1.09E+08 | 109300000 | 0.00 | 0.00 | NO |
| chr2 | 1.09E+08 | 109400000 | 0.00 | 0.00 | NO |
| chr2 | 1.09E+08 | 109500000 | 1.06 | 1.02 | A>A |
| chr2 | 1.1E+08 | 109600000 | 1.06 | 1.27 | A>A |
| chr2 | 1.1E+08 | 109700000 | 1.09 | 1.01 | A>A |
| chr2 | 1.1E+08 | 109800000 | 0.96 | 1.01 | A>A |
| chr2 | 1.1E+08 | 109900000 | 1.15 | 1.02 | A>A |
| chr2 | 1.1E+08 | 110000000 | 1.09 | 1.13 | A>A |
| chr2 | 1.1E+08 | 110100000 | 0.99 | 0.90 | A>A |
| chr2 | 1.1E+08 | 110200000 | 0.00 | 0.00 | NO |
| chr2 | 1.1E+08 | 110300000 | 0.00 | 0.00 | NO |
| chr2 | 1.1E+08 | 110400000 | 1.58 | 1.66 | A>A |
| chr2 | 1.1E+08 | 110500000 | 1.47 | 1.46 | A>A |
| chr2 | 1.11E+08 | 110600000 | 0.00 | 0.00 | NO |
| chr2 | 1.11E+08 | 110700000 | 0.00 | 0.00 | NO |
| chr2 | 1.11E+08 | 110800000 | 0.00 | 0.00 | NO |
| chr2 | 1.11E+08 | 110900000 | 0.00 | 0.00 | NO |
| chr2 | 1.11E+08 | 111000000 | 0.00 | 0.00 | NO |
| chr2 | 1.11E+08 | 111100000 | 0.00 | 0.00 | NO |
| chr2 | 1.11E+08 | 111200000 | 0.00 | 0.00 | NO |
| chr2 | 1.11E+08 | 111300000 | 0.00 | 0.00 | NO |
| chr2 | 1.11E+08 | 111400000 | 0.00 | 0.00 | NO |
| chr2 | 1.11E+08 | 111500000 | 1.61 | 1.68 | A>A |
| chr2 | 1.12E+08 | 111600000 | 1.25 | 1.34 | A>A |
| chr2 | 1.12E+08 | 111700000 | 1.21 | 1.41 | A>A |
| chr2 | 1.12E+08 | 111800000 | 1.34 | 1.46 | A>A |
| chr2 | 1.12E+08 | 111900000 | 1.23 | 1.44 | A>A |
| chr2 | 1.12E+08 | 112000000 | 1.45 | 1.72 | A>A |
| chr2 | 1.12E+08 | 112100000 | 0.00 | 0.00 | NO |
| chr2 | 1.12E+08 | 112200000 | 0.00 | 0.00 | NO |
| chr2 | 1.12E+08 | 112300000 | 0.00 | 0.00 | NO |
| chr2 | 1.12E+08 | 112400000 | 0.00 | 0.00 | NO |
| chr2 | 1.12E+08 | 112500000 | 0.00 | 0.00 | NO |
| chr2 | 1.13E+08 | 112600000 | 0.00 | 0.00 | NO |
| chr2 | 1.13E+08 | 112700000 | 0.00 | 0.00 | NO |
| chr2 | 1.13E+08 | 112800000 | 1.88 | 1.89 | A>A |
| chr2 | 1.13E+08 | 112900000 | 1.47 | 1.73 | A>A |
| chr2 | 1.13E+08 | 113000000 | 1.69 | 1.79 | A>A |
| chr2 | 1.13E+08 | 113100000 | 1.59 | 1.75 | A>A |

| | | | | | |
|---|---|---|---|---|---|
| chr2 | 1.13E+08 | 113200000 | 0.00 | 0.00 | NO |
| chr2 | 1.13E+08 | 113300000 | 1.41 | 1.45 | A>A |
| chr2 | 1.13E+08 | 113400000 | 1.48 | 1.49 | A>A |
| chr2 | 1.13E+08 | 113500000 | 1.47 | 1.59 | A>A |
| chr2 | 1.14E+08 | 113600000 | 1.42 | 1.47 | A>A |
| chr2 | 1.14E+08 | 113700000 | 1.24 | 1.48 | A>A |
| chr2 | 1.14E+08 | 113800000 | 1.12 | 1.18 | A>A |
| chr2 | 1.14E+08 | 113900000 | 1.00 | 1.26 | A>A |
| chr2 | 1.14E+08 | 114000000 | 1.25 | 1.45 | A>A |
| chr2 | 1.14E+08 | 114100000 | 0.00 | 0.00 | NO |
| chr2 | 1.14E+08 | 114200000 | 0.00 | 0.00 | NO |
| chr2 | 1.14E+08 | 114300000 | 0.00 | 0.00 | NO |
| chr2 | 1.14E+08 | 114400000 | 0.00 | 0.00 | NO |
| chr2 | 1.14E+08 | 114500000 | 0.00 | 0.00 | NO |
| chr2 | 1.15E+08 | 114600000 | 0.48 | 0.74 | A>A |
| chr2 | 1.15E+08 | 114700000 | 0.52 | 0.69 | A>A |
| chr2 | 1.15E+08 | 114800000 | -0.06 | -0.15 | B>B |
| chr2 | 1.15E+08 | 114900000 | -0.45 | -0.78 | B>B |
| chr2 | 1.15E+08 | 115000000 | -0.56 | -0.92 | B>B |
| chr2 | 1.15E+08 | 115100000 | -0.65 | -0.93 | B>B |
| chr2 | 1.15E+08 | 115200000 | -0.70 | -0.90 | B>B |
| chr2 | 1.15E+08 | 115300000 | -0.62 | -0.93 | B>B |
| chr2 | 1.15E+08 | 115400000 | -0.61 | -0.91 | B>B |
| chr2 | 1.15E+08 | 115500000 | -0.62 | -0.86 | B>B |
| chr2 | 1.16E+08 | 115600000 | -0.53 | -0.92 | B>B |
| chr2 | 1.16E+08 | 115700000 | -0.64 | -0.85 | B>B |
| chr2 | 1.16E+08 | 115800000 | -0.68 | -0.85 | B>B |
| chr2 | 1.16E+08 | 115900000 | -0.69 | -0.83 | B>B |
| chr2 | 1.16E+08 | 116000000 | -0.71 | -0.84 | B>B |
| chr2 | 1.16E+08 | 116100000 | -0.84 | -0.82 | B>B |
| chr2 | 1.16E+08 | 116200000 | -0.85 | -0.89 | B>B |
| chr2 | 1.16E+08 | 116300000 | -0.89 | -0.86 | B>B |
| chr2 | 1.16E+08 | 116400000 | -0.82 | -0.85 | B>B |
| chr2 | 1.16E+08 | 116500000 | -0.83 | -0.88 | B>B |
| chr2 | 1.17E+08 | 116600000 | -0.84 | -0.82 | B>B |
| chr2 | 1.17E+08 | 116700000 | -0.78 | -0.81 | B>B |
| chr2 | 1.17E+08 | 116800000 | -0.82 | -0.85 | B>B |
| chr2 | 1.17E+08 | 116900000 | -0.77 | -0.84 | B>B |
| chr2 | 1.17E+08 | 117000000 | -0.78 | -0.81 | B>B |
| chr2 | 1.17E+08 | 117100000 | -0.82 | -0.89 | B>B |
| chr2 | 1.17E+08 | 117200000 | -0.81 | -0.88 | B>B |
| chr2 | 1.17E+08 | 117300000 | -0.84 | -0.81 | B>B |
| chr2 | 1.17E+08 | 117400000 | -0.82 | -0.85 | B>B |
| chr2 | 1.17E+08 | 117500000 | -0.84 | -0.85 | B>B |
| chr2 | 1.18E+08 | 117600000 | -0.83 | -0.87 | B>B |
| chr2 | 1.18E+08 | 117700000 | -0.83 | -0.80 | B>B |
| chr2 | 1.18E+08 | 117800000 | -0.81 | -0.83 | B>B |
| chr2 | 1.18E+08 | 117900000 | -0.79 | -0.79 | B>B |
| chr2 | 1.18E+08 | 118000000 | -0.78 | -0.88 | B>B |
| chr2 | 1.18E+08 | 118100000 | -0.79 | -0.87 | B>B |
| chr2 | 1.18E+08 | 118200000 | -0.76 | -0.85 | B>B |
| chr2 | 1.18E+08 | 118300000 | -0.75 | -0.85 | B>B |
| chr2 | 1.18E+08 | 118400000 | -0.75 | -0.83 | B>B |
| chr2 | 1.18E+08 | 118500000 | -0.57 | -0.82 | B>B |
| chr2 | 1.19E+08 | 118600000 | -0.23 | -0.44 | B>B |
| chr2 | 1.19E+08 | 118700000 | 0.13 | -0.04 | A>A |

| | | | | | |
|---|---|---|---|---|---|
| chr2 | 1.19E+08 | 118800000 | 0.16 | 0.11 | A>A |
| chr2 | 1.19E+08 | 118900000 | 0.26 | 0.23 | A>A |
| chr2 | 1.19E+08 | 119000000 | 0.12 | 0.05 | A>A |
| chr2 | 1.19E+08 | 119100000 | -0.16 | -0.22 | B>B |
| chr2 | 1.19E+08 | 119200000 | -0.26 | -0.38 | B>B |
| chr2 | 1.19E+08 | 119300000 | -0.22 | -0.38 | B>B |
| chr2 | 1.19E+08 | 119400000 | -0.05 | -0.46 | B>B |
| chr2 | 1.19E+08 | 119500000 | 0.12 | -0.30 | B>B |
| chr2 | 1.2E+08 | 119600000 | 0.16 | -0.21 | B>B |
| chr2 | 1.2E+08 | 119700000 | -0.06 | -0.04 | B>B |
| chr2 | 1.2E+08 | 119800000 | 0.03 | 0.15 | A>A |
| chr2 | 1.2E+08 | 119900000 | 0.23 | 0.70 | A>A |
| chr2 | 1.2E+08 | 120000000 | 0.47 | 0.61 | A>A |
| chr2 | 1.2E+08 | 120100000 | 0.47 | 0.53 | A>A |
| chr2 | 1.2E+08 | 120200000 | 0.85 | 0.87 | A>A |
| chr2 | 1.2E+08 | 120300000 | 0.96 | 0.97 | A>A |
| chr2 | 1.2E+08 | 120400000 | 0.95 | 1.11 | A>A |
| chr2 | 1.2E+08 | 120500000 | 0.92 | 0.99 | A>A |
| chr2 | 1.21E+08 | 120600000 | 1.14 | 1.37 | A>A |
| chr2 | 1.21E+08 | 120700000 | 1.12 | 1.43 | A>A |
| chr2 | 1.21E+08 | 120800000 | 1.15 | 1.31 | A>A |
| chr2 | 1.21E+08 | 120900000 | 1.22 | 1.37 | A>A |
| chr2 | 1.21E+08 | 121000000 | 1.09 | 1.23 | A>A |
| chr2 | 1.21E+08 | 121100000 | 0.98 | 1.15 | A>A |
| chr2 | 1.21E+08 | 121200000 | 1.50 | 1.54 | A>A |
| chr2 | 1.21E+08 | 121300000 | 1.47 | 1.31 | A>A |
| chr2 | 1.21E+08 | 121400000 | 1.26 | 1.24 | A>A |
| chr2 | 1.21E+08 | 121500000 | 0.81 | 0.87 | A>A |
| chr2 | 1.22E+08 | 121600000 | 1.26 | 1.34 | A>A |
| chr2 | 1.22E+08 | 121700000 | 1.22 | 1.42 | A>A |
| chr2 | 1.22E+08 | 121800000 | 1.19 | 1.12 | A>A |
| chr2 | 1.22E+08 | 121900000 | 0.68 | 0.67 | A>A |
| chr2 | 1.22E+08 | 122000000 | 1.10 | 1.05 | A>A |
| chr2 | 1.22E+08 | 122100000 | 1.00 | 0.92 | A>A |
| chr2 | 1.22E+08 | 122200000 | 0.77 | 0.81 | A>A |
| chr2 | 1.22E+08 | 122300000 | 1.02 | 0.89 | A>A |
| chr2 | 1.22E+08 | 122400000 | 1.03 | 0.98 | A>A |
| chr2 | 1.22E+08 | 122500000 | 0.80 | 0.66 | A>A |
| chr2 | 1.23E+08 | 122600000 | 0.20 | -0.05 | A>A |
| chr2 | 1.23E+08 | 122700000 | -0.06 | -0.41 | B>B |
| chr2 | 1.23E+08 | 122800000 | -0.33 | -0.59 | B>B |
| chr2 | 1.23E+08 | 122900000 | -0.42 | -0.75 | B>B |
| chr2 | 1.23E+08 | 123000000 | -0.47 | -0.74 | B>B |
| chr2 | 1.23E+08 | 123100000 | -0.59 | -0.79 | B>B |
| chr2 | 1.23E+08 | 123200000 | -0.69 | -0.80 | B>B |
| chr2 | 1.23E+08 | 123300000 | -0.76 | -0.80 | B>B |
| chr2 | 1.23E+08 | 123400000 | -0.78 | -0.77 | B>B |
| chr2 | 1.23E+08 | 123500000 | -0.78 | -0.80 | B>B |
| chr2 | 1.24E+08 | 123600000 | -0.85 | -0.87 | B>B |
| chr2 | 1.24E+08 | 123700000 | -0.80 | -0.80 | B>B |
| chr2 | 1.24E+08 | 123800000 | -0.81 | -0.83 | B>B |
| chr2 | 1.24E+08 | 123900000 | -0.83 | -0.80 | B>B |
| chr2 | 1.24E+08 | 124000000 | -0.82 | -0.75 | B>B |
| chr2 | 1.24E+08 | 124100000 | -0.83 | -0.79 | B>B |
| chr2 | 1.24E+08 | 124200000 | -0.74 | -0.75 | B>B |
| chr2 | 1.24E+08 | 124300000 | -0.82 | -0.81 | B>B |

| | | | | | |
|---|---|---|---|---|---|
| chr2 | 1.24E+08 | 124400000 | -0.83 | -0.81 | B>B |
| chr2 | 1.24E+08 | 124500000 | -0.81 | -0.76 | B>B |
| chr2 | 1.25E+08 | 124600000 | -0.78 | -0.76 | B>B |
| chr2 | 1.25E+08 | 124700000 | -0.78 | -0.76 | B>B |
| chr2 | 1.25E+08 | 124800000 | -0.71 | -0.73 | B>B |
| chr2 | 1.25E+08 | 124900000 | -0.74 | -0.72 | B>B |
| chr2 | 1.25E+08 | 125000000 | -0.75 | -0.77 | B>B |
| chr2 | 1.25E+08 | 125100000 | -0.76 | -0.76 | B>B |
| chr2 | 1.25E+08 | 125200000 | -0.72 | -0.70 | B>B |
| chr2 | 1.25E+08 | 125300000 | -0.72 | -0.72 | B>B |
| chr2 | 1.25E+08 | 125400000 | -0.69 | -0.69 | B>B |
| chr2 | 1.25E+08 | 125500000 | -0.72 | -0.72 | B>B |
| chr2 | 1.26E+08 | 125600000 | -0.69 | -0.72 | B>B |
| chr2 | 1.26E+08 | 125700000 | -0.73 | -0.72 | B>B |
| chr2 | 1.26E+08 | 125800000 | -0.73 | -0.67 | B>B |
| chr2 | 1.26E+08 | 125900000 | -0.76 | -0.76 | B>B |
| chr2 | 1.26E+08 | 126000000 | -0.74 | -0.72 | B>B |
| chr2 | 1.26E+08 | 126100000 | -0.67 | -0.70 | B>B |
| chr2 | 1.26E+08 | 126200000 | -0.58 | -0.57 | B>B |
| chr2 | 1.26E+08 | 126300000 | -0.64 | -0.68 | B>B |
| chr2 | 1.26E+08 | 126400000 | -0.74 | -0.74 | B>B |
| chr2 | 1.26E+08 | 126500000 | -0.74 | -0.73 | B>B |
| chr2 | 1.27E+08 | 126600000 | -0.72 | -0.67 | B>B |
| chr2 | 1.27E+08 | 126700000 | -0.75 | -0.72 | B>B |
| chr2 | 1.27E+08 | 126800000 | -0.82 | -0.77 | B>B |
| chr2 | 1.27E+08 | 126900000 | -0.79 | -0.79 | B>B |
| chr2 | 1.27E+08 | 127000000 | -0.79 | -0.79 | B>B |
| chr2 | 1.27E+08 | 127100000 | -0.77 | -0.78 | B>B |
| chr2 | 1.27E+08 | 127200000 | -0.72 | -0.76 | B>B |
| chr2 | 1.27E+08 | 127300000 | -0.66 | -0.78 | B>B |
| chr2 | 1.27E+08 | 127400000 | -0.56 | -0.76 | B>B |
| chr2 | 1.27E+08 | 127500000 | -0.36 | -0.68 | B>B |
| chr2 | 1.28E+08 | 127600000 | -0.23 | -0.52 | B>B |
| chr2 | 1.28E+08 | 127700000 | -0.01 | -0.38 | B>B |
| chr2 | 1.28E+08 | 127800000 | 0.25 | -0.07 | A>A |
| chr2 | 1.28E+08 | 127900000 | 0.82 | 0.61 | A>A |
| chr2 | 1.28E+08 | 128000000 | 0.96 | 0.82 | A>A |
| chr2 | 1.28E+08 | 128100000 | 0.97 | 1.07 | A>A |
| chr2 | 1.28E+08 | 128200000 | 1.36 | 1.27 | A>A |
| chr2 | 1.28E+08 | 128300000 | 1.10 | 1.11 | A>A |
| chr2 | 1.28E+08 | 128400000 | 1.26 | 1.33 | A>A |
| chr2 | 1.28E+08 | 128500000 | 1.36 | 1.35 | A>A |
| chr2 | 1.29E+08 | 128600000 | 1.37 | 1.36 | A>A |
| chr2 | 1.29E+08 | 128700000 | 1.35 | 1.23 | A>A |
| chr2 | 1.29E+08 | 128800000 | 1.27 | 1.10 | A>A |
| chr2 | 1.29E+08 | 128900000 | 0.99 | 0.86 | A>A |
| chr2 | 1.29E+08 | 129000000 | 0.78 | 0.58 | A>A |
| chr2 | 1.29E+08 | 129100000 | 0.55 | 0.43 | A>A |
| chr2 | 1.29E+08 | 129200000 | 0.50 | 0.44 | A>A |
| chr2 | 1.29E+08 | 129300000 | 0.36 | 0.24 | A>A |
| chr2 | 1.29E+08 | 129400000 | 0.25 | 0.06 | A>A |
| chr2 | 1.29E+08 | 129500000 | -0.03 | -0.20 | B>B |
| chr2 | 1.3E+08 | 129600000 | -0.28 | -0.55 | B>B |
| chr2 | 1.3E+08 | 129700000 | -0.51 | -0.83 | B>B |
| chr2 | 1.3E+08 | 129800000 | -0.60 | -0.83 | B>B |
| chr2 | 1.3E+08 | 129900000 | -0.56 | -0.88 | B>B |

| | | | | | |
|---|---|---|---|---|---|
| chr2 | 1.3E+08 | 130000000 | -0.53 | -0.90 | B>B |
| chr2 | 1.3E+08 | 130100000 | -0.57 | -0.88 | B>B |
| chr2 | 1.3E+08 | 130200000 | -0.54 | -0.88 | B>B |
| chr2 | 1.3E+08 | 130300000 | -0.56 | -0.87 | B>B |
| chr2 | 1.3E+08 | 130400000 | -0.60 | -0.93 | B>B |
| chr2 | 1.3E+08 | 130500000 | -0.39 | -0.82 | B>B |
| chr2 | 1.31E+08 | 130600000 | -0.25 | -0.64 | B>B |
| chr2 | 1.31E+08 | 130700000 | -0.09 | -0.37 | B>B |
| chr2 | 1.31E+08 | 130800000 | 0.00 | 0.00 | NO |
| chr2 | 1.31E+08 | 130900000 | 0.00 | 0.00 | NO |
| chr2 | 1.31E+08 | 131000000 | 0.00 | 0.00 | NO |
| chr2 | 1.31E+08 | 131100000 | 0.86 | 0.18 | A>B |
| chr2 | 1.31E+08 | 131200000 | 0.00 | 0.00 | NO |
| chr2 | 1.31E+08 | 131300000 | 0.00 | 0.00 | NO |
| chr2 | 1.31E+08 | 131400000 | 0.00 | 0.00 | NO |
| chr2 | 1.31E+08 | 131500000 | 0.00 | 0.00 | NO |
| chr2 | 1.32E+08 | 131600000 | 0.83 | 0.54 | A>A |
| chr2 | 1.32E+08 | 131700000 | 0.98 | 0.71 | A>A |
| chr2 | 1.32E+08 | 131800000 | 0.95 | 0.88 | A>A |
| chr2 | 1.32E+08 | 131900000 | 0.80 | 0.55 | A>A |
| chr2 | 1.32E+08 | 132000000 | 0.00 | 0.00 | NO |
| chr2 | 1.32E+08 | 132100000 | 0.00 | 0.00 | NO |
| chr2 | 1.32E+08 | 132200000 | 0.00 | 0.00 | NO |
| chr2 | 1.32E+08 | 132300000 | 0.00 | 0.00 | NO |
| chr2 | 1.32E+08 | 132400000 | 0.00 | 0.00 | NO |
| chr2 | 1.32E+08 | 132500000 | 0.00 | 0.00 | NO |
| chr2 | 1.33E+08 | 132600000 | 0.00 | 0.00 | NO |
| chr2 | 1.33E+08 | 132700000 | 0.00 | 0.00 | NO |
| chr2 | 1.33E+08 | 132800000 | 0.00 | 0.00 | NO |
| chr2 | 1.33E+08 | 132900000 | 0.00 | 0.00 | NO |
| chr2 | 1.33E+08 | 133000000 | 0.00 | 0.00 | NO |
| chr2 | 1.33E+08 | 133100000 | 0.04 | -0.16 | B>B |
| chr2 | 1.33E+08 | 133200000 | 0.40 | 0.30 | A>A |
| chr2 | 1.33E+08 | 133300000 | 0.31 | 0.26 | A>A |
| chr2 | 1.33E+08 | 133400000 | 0.24 | 0.04 | A>A |
| chr2 | 1.33E+08 | 133500000 | 0.18 | -0.27 | B>B |
| chr2 | 1.34E+08 | 133600000 | 0.23 | -0.53 | A>B |
| chr2 | 1.34E+08 | 133700000 | 0.16 | -0.47 | A>B |
| chr2 | 1.34E+08 | 133800000 | 0.13 | -0.58 | A>B |
| chr2 | 1.34E+08 | 133900000 | 0.17 | -0.49 | A>B |
| chr2 | 1.34E+08 | 134000000 | 0.34 | -0.36 | A>B |
| chr2 | 1.34E+08 | 134100000 | 0.69 | -0.34 | A>B |
| chr2 | 1.34E+08 | 134200000 | 0.74 | -0.32 | A>B |
| chr2 | 1.34E+08 | 134300000 | 0.56 | -0.26 | A>B |
| chr2 | 1.34E+08 | 134400000 | 0.46 | -0.32 | A>B |
| chr2 | 1.34E+08 | 134500000 | 0.57 | -0.24 | A>B |
| chr2 | 1.35E+08 | 134600000 | 0.34 | -0.27 | A>B |
| chr2 | 1.35E+08 | 134700000 | 0.18 | -0.28 | B>B |
| chr2 | 1.35E+08 | 134800000 | 0.38 | -0.17 | A>A |
| chr2 | 1.35E+08 | 134900000 | 0.47 | 0.10 | A>A |
| chr2 | 1.35E+08 | 135000000 | 0.63 | 0.27 | A>A |
| chr2 | 1.35E+08 | 135100000 | 0.54 | 0.28 | A>A |
| chr2 | 1.35E+08 | 135200000 | 0.46 | 0.28 | A>A |
| chr2 | 1.35E+08 | 135300000 | 0.28 | 0.12 | A>A |
| chr2 | 1.35E+08 | 135400000 | 0.29 | 0.06 | A>A |
| chr2 | 1.35E+08 | 135500000 | 0.33 | 0.20 | A>A |

| | | | | | |
|---|---|---|---|---|---|
| chr2 | 1.36E+08 | 135600000 | 0.35 | 0.23 | A>A |
| chr2 | 1.36E+08 | 135700000 | 0.43 | 0.30 | A>A |
| chr2 | 1.36E+08 | 135800000 | 0.51 | 0.53 | A>A |
| chr2 | 1.36E+08 | 135900000 | 0.91 | 1.01 | A>A |
| chr2 | 1.36E+08 | 136000000 | 0.72 | 1.02 | A>A |
| chr2 | 1.36E+08 | 136100000 | 0.46 | 0.91 | A>A |
| chr2 | 1.36E+08 | 136200000 | 0.44 | 0.79 | A>A |
| chr2 | 1.36E+08 | 136300000 | 0.69 | 0.96 | A>A |
| chr2 | 1.36E+08 | 136400000 | 0.93 | 1.02 | A>A |
| chr2 | 1.36E+08 | 136500000 | 0.83 | 0.95 | A>A |
| chr2 | 1.37E+08 | 136600000 | 0.79 | 0.54 | A>A |
| chr2 | 1.37E+08 | 136700000 | 0.48 | 0.12 | A>A |
| chr2 | 1.37E+08 | 136800000 | 0.51 | 0.17 | A>A |
| chr2 | 1.37E+08 | 136900000 | 0.46 | -0.10 | A>A |
| chr2 | 1.37E+08 | 137000000 | 0.35 | -0.19 | A>A |
| chr2 | 1.37E+08 | 137100000 | 0.28 | -0.17 | A>A |
| chr2 | 1.37E+08 | 137200000 | -0.10 | -0.36 | B>B |
| chr2 | 1.37E+08 | 137300000 | -0.52 | -0.77 | B>B |
| chr2 | 1.37E+08 | 137400000 | -0.71 | -0.87 | B>B |
| chr2 | 1.37E+08 | 137500000 | -0.78 | -0.87 | B>B |
| chr2 | 1.38E+08 | 137600000 | -0.75 | -0.85 | B>B |
| chr2 | 1.38E+08 | 137700000 | -0.86 | -0.85 | B>B |
| chr2 | 1.38E+08 | 137800000 | -0.89 | -0.88 | B>B |
| chr2 | 1.38E+08 | 137900000 | -0.92 | -0.88 | B>B |
| chr2 | 1.38E+08 | 138000000 | -0.94 | -0.87 | B>B |
| chr2 | 1.38E+08 | 138100000 | -0.91 | -0.88 | B>B |
| chr2 | 1.38E+08 | 138200000 | -0.90 | -0.86 | B>B |
| chr2 | 1.38E+08 | 138300000 | -0.89 | -0.89 | B>B |
| chr2 | 1.38E+08 | 138400000 | -0.84 | -0.87 | B>B |
| chr2 | 1.38E+08 | 138500000 | -0.78 | -0.78 | B>B |
| chr2 | 1.39E+08 | 138600000 | -0.69 | -0.66 | B>B |
| chr2 | 1.39E+08 | 138700000 | -0.23 | -0.14 | B>B |
| chr2 | 1.39E+08 | 138800000 | 0.21 | 0.43 | A>A |
| chr2 | 1.39E+08 | 138900000 | 0.22 | 0.49 | A>A |
| chr2 | 1.39E+08 | 139000000 | 0.31 | 0.52 | A>A |
| chr2 | 1.39E+08 | 139100000 | 0.24 | 0.39 | A>A |
| chr2 | 1.39E+08 | 139200000 | 0.12 | 0.34 | A>A |
| chr2 | 1.39E+08 | 139300000 | 0.32 | 0.48 | A>A |
| chr2 | 1.39E+08 | 139400000 | 0.22 | 0.38 | A>A |
| chr2 | 1.39E+08 | 139500000 | 0.01 | 0.22 | A>A |
| chr2 | 1.4E+08 | 139600000 | -0.17 | -0.08 | B>B |
| chr2 | 1.4E+08 | 139700000 | -0.58 | -0.54 | B>B |
| chr2 | 1.4E+08 | 139800000 | -0.75 | -0.69 | B>B |
| chr2 | 1.4E+08 | 139900000 | -0.82 | -0.79 | B>B |
| chr2 | 1.4E+08 | 140000000 | -0.87 | -0.80 | B>B |
| chr2 | 1.4E+08 | 140100000 | -0.80 | -0.74 | B>B |
| chr2 | 1.4E+08 | 140200000 | -0.79 | -0.78 | B>B |
| chr2 | 1.4E+08 | 140300000 | -0.78 | -0.77 | B>B |
| chr2 | 1.4E+08 | 140400000 | -0.74 | -0.74 | B>B |
| chr2 | 1.4E+08 | 140500000 | -0.77 | -0.71 | B>B |
| chr2 | 1.41E+08 | 140600000 | -0.72 | -0.71 | B>B |
| chr2 | 1.41E+08 | 140700000 | -0.74 | -0.71 | B>B |
| chr2 | 1.41E+08 | 140800000 | -0.75 | -0.70 | B>B |
| chr2 | 1.41E+08 | 140900000 | -0.75 | -0.71 | B>B |
| chr2 | 1.41E+08 | 141000000 | -0.73 | -0.65 | B>B |
| chr2 | 1.41E+08 | 141100000 | -0.67 | -0.51 | B>B |

| | | | | | |
|---|---|---|---|---|---|
| chr2 | 1.41E+08 | 141200000 | -0.50 | -0.29 | B>B |
| chr2 | 1.41E+08 | 141300000 | -0.29 | 0.01 | B>B |
| chr2 | 1.41E+08 | 141400000 | -0.12 | 0.00 | B>B |
| chr2 | 1.41E+08 | 141500000 | -0.12 | -0.19 | B>B |
| chr2 | 1.42E+08 | 141600000 | -0.27 | -0.36 | B>B |
| chr2 | 1.42E+08 | 141700000 | -0.55 | -0.59 | B>B |
| chr2 | 1.42E+08 | 141800000 | -0.68 | -0.69 | B>B |
| chr2 | 1.42E+08 | 141900000 | -0.73 | -0.79 | B>B |
| chr2 | 1.42E+08 | 142000000 | -0.74 | -0.77 | B>B |
| chr2 | 1.42E+08 | 142100000 | -0.68 | -0.78 | B>B |
| chr2 | 1.42E+08 | 142200000 | -0.64 | -0.71 | B>B |
| chr2 | 1.42E+08 | 142300000 | -0.61 | -0.81 | B>B |
| chr2 | 1.42E+08 | 142400000 | -0.67 | -0.79 | B>B |
| chr2 | 1.42E+08 | 142500000 | -0.79 | -0.80 | B>B |
| chr2 | 1.43E+08 | 142600000 | -0.75 | -0.80 | B>B |
| chr2 | 1.43E+08 | 142700000 | -0.68 | -0.76 | B>B |
| chr2 | 1.43E+08 | 142800000 | -0.65 | -0.82 | B>B |
| chr2 | 1.43E+08 | 142900000 | -0.52 | -0.66 | B>B |
| chr2 | 1.43E+08 | 143000000 | -0.32 | -0.25 | B>B |
| chr2 | 1.43E+08 | 143100000 | -0.36 | -0.35 | B>B |
| chr2 | 1.43E+08 | 143200000 | -0.31 | -0.39 | B>B |
| chr2 | 1.43E+08 | 143300000 | -0.14 | -0.29 | B>B |
| chr2 | 1.43E+08 | 143400000 | -0.13 | -0.26 | B>B |
| chr2 | 1.43E+08 | 143500000 | 0.37 | 0.21 | A>A |
| chr2 | 1.44E+08 | 143600000 | 0.83 | 0.60 | A>A |
| chr2 | 1.44E+08 | 143700000 | 0.66 | 0.56 | A>A |
| chr2 | 1.44E+08 | 143800000 | 0.15 | 0.16 | A>A |
| chr2 | 1.44E+08 | 143900000 | -0.39 | -0.50 | B>B |
| chr2 | 1.44E+08 | 144000000 | -0.46 | -0.67 | B>B |
| chr2 | 1.44E+08 | 144100000 | -0.54 | -0.78 | B>B |
| chr2 | 1.44E+08 | 144200000 | -0.52 | -0.86 | B>B |
| chr2 | 1.44E+08 | 144300000 | -0.50 | -0.85 | B>B |
| chr2 | 1.44E+08 | 144400000 | -0.53 | -0.84 | B>B |
| chr2 | 1.44E+08 | 144500000 | -0.46 | -0.73 | B>B |
| chr2 | 1.45E+08 | 144600000 | -0.33 | -0.71 | B>B |
| chr2 | 1.45E+08 | 144700000 | -0.22 | -0.56 | B>B |
| chr2 | 1.45E+08 | 144800000 | -0.08 | -0.48 | B>B |
| chr2 | 1.45E+08 | 144900000 | 0.10 | -0.39 | B>B |
| chr2 | 1.45E+08 | 145000000 | 0.23 | -0.22 | A>A |
| chr2 | 1.45E+08 | 145100000 | 0.31 | 0.03 | A>A |
| chr2 | 1.45E+08 | 145200000 | 0.38 | 0.00 | A>A |
| chr2 | 1.45E+08 | 145300000 | 0.20 | -0.04 | A>A |
| chr2 | 1.45E+08 | 145400000 | 0.28 | -0.10 | A>A |
| chr2 | 1.45E+08 | 145500000 | 0.25 | -0.13 | A>A |
| chr2 | 1.46E+08 | 145600000 | 0.24 | -0.22 | A>A |
| chr2 | 1.46E+08 | 145700000 | 0.35 | -0.09 | A>A |
| chr2 | 1.46E+08 | 145800000 | 0.39 | -0.03 | A>A |
| chr2 | 1.46E+08 | 145900000 | 0.29 | -0.10 | A>A |
| chr2 | 1.46E+08 | 146000000 | 0.19 | -0.06 | A>A |
| chr2 | 1.46E+08 | 146100000 | 0.04 | -0.05 | B>B |
| chr2 | 1.46E+08 | 146200000 | -0.04 | -0.11 | B>B |
| chr2 | 1.46E+08 | 146300000 | -0.08 | 0.04 | B>B |
| chr2 | 1.46E+08 | 146400000 | -0.12 | 0.04 | B>B |
| chr2 | 1.46E+08 | 146500000 | -0.16 | 0.16 | A>A |
| chr2 | 1.47E+08 | 146600000 | -0.17 | 0.24 | A>A |
| chr2 | 1.47E+08 | 146700000 | -0.27 | 0.30 | B>A |

| | | | | | |
|---|---|---|---|---|---|
| chr2 | 1.47E+08 | 146800000 | -0.42 | 0.32 | B>A |
| chr2 | 1.47E+08 | 146900000 | -0.53 | 0.39 | B>A |
| chr2 | 1.47E+08 | 147000000 | -0.53 | 0.35 | B>A |
| chr2 | 1.47E+08 | 147100000 | -0.51 | 0.21 | B>A |
| chr2 | 1.47E+08 | 147200000 | -0.49 | 0.16 | B>A |
| chr2 | 1.47E+08 | 147300000 | -0.43 | 0.02 | B>B |
| chr2 | 1.47E+08 | 147400000 | -0.30 | -0.06 | B>B |
| chr2 | 1.47E+08 | 147500000 | -0.51 | -0.40 | B>B |
| chr2 | 1.48E+08 | 147600000 | -0.73 | -0.63 | B>B |
| chr2 | 1.48E+08 | 147700000 | -0.82 | -0.74 | B>B |
| chr2 | 1.48E+08 | 147800000 | -0.85 | -0.87 | B>B |
| chr2 | 1.48E+08 | 147900000 | -0.80 | -0.87 | B>B |
| chr2 | 1.48E+08 | 148000000 | -0.89 | -0.89 | B>B |
| chr2 | 1.48E+08 | 148100000 | -0.88 | -0.85 | B>B |
| chr2 | 1.48E+08 | 148200000 | -0.79 | -0.82 | B>B |
| chr2 | 1.48E+08 | 148300000 | -0.66 | -0.81 | B>B |
| chr2 | 1.48E+08 | 148400000 | -0.63 | -0.78 | B>B |
| chr2 | 1.48E+08 | 148500000 | -0.60 | -0.80 | B>B |
| chr2 | 1.49E+08 | 148600000 | -0.39 | -0.57 | B>B |
| chr2 | 1.49E+08 | 148700000 | -0.21 | -0.35 | B>B |
| chr2 | 1.49E+08 | 148800000 | 0.11 | -0.01 | A>A |
| chr2 | 1.49E+08 | 148900000 | 0.06 | 0.03 | A>A |
| chr2 | 1.49E+08 | 149000000 | -0.11 | -0.12 | B>B |
| chr2 | 1.49E+08 | 149100000 | -0.23 | -0.18 | B>B |
| chr2 | 1.49E+08 | 149200000 | -0.31 | -0.30 | B>B |
| chr2 | 1.49E+08 | 149300000 | -0.21 | -0.25 | B>B |
| chr2 | 1.49E+08 | 149400000 | 0.13 | -0.15 | B>B |
| chr2 | 1.49E+08 | 149500000 | 0.39 | 0.15 | A>A |
| chr2 | 1.5E+08 | 149600000 | 0.22 | -0.06 | A>A |
| chr2 | 1.5E+08 | 149700000 | -0.03 | -0.36 | B>B |
| chr2 | 1.5E+08 | 149800000 | 0.00 | 0.00 | NO |
| chr2 | 1.5E+08 | 149900000 | -0.16 | -0.42 | B>B |
| chr2 | 1.5E+08 | 150000000 | -0.34 | -0.82 | B>B |
| chr2 | 1.5E+08 | 150100000 | -0.40 | -0.84 | B>B |
| chr2 | 1.5E+08 | 150200000 | -0.45 | -0.76 | B>B |
| chr2 | 1.5E+08 | 150300000 | -0.41 | -0.72 | B>B |
| chr2 | 1.5E+08 | 150400000 | -0.35 | -0.52 | B>B |
| chr2 | 1.5E+08 | 150500000 | 0.22 | 0.20 | A>A |
| chr2 | 1.51E+08 | 150600000 | 0.58 | 0.72 | A>A |
| chr2 | 1.51E+08 | 150700000 | 0.69 | 0.79 | A>A |
| chr2 | 1.51E+08 | 150800000 | 0.86 | 0.98 | A>A |
| chr2 | 1.51E+08 | 150900000 | 0.80 | 0.92 | A>A |
| chr2 | 1.51E+08 | 151000000 | 0.83 | 0.99 | A>A |
| chr2 | 1.51E+08 | 151100000 | 0.67 | 0.77 | A>A |
| chr2 | 1.51E+08 | 151200000 | 0.83 | 1.05 | A>A |
| chr2 | 1.51E+08 | 151300000 | 0.75 | 0.98 | A>A |
| chr2 | 1.51E+08 | 151400000 | 0.52 | 0.66 | A>A |
| chr2 | 1.51E+08 | 151500000 | 0.44 | 0.63 | A>A |
| chr2 | 1.52E+08 | 151600000 | 0.23 | 0.53 | A>A |
| chr2 | 1.52E+08 | 151700000 | 0.07 | 0.46 | A>A |
| chr2 | 1.52E+08 | 151800000 | -0.05 | 0.45 | A>A |
| chr2 | 1.52E+08 | 151900000 | -0.04 | 0.32 | A>A |
| chr2 | 1.52E+08 | 152000000 | -0.03 | 0.29 | A>A |
| chr2 | 1.52E+08 | 152100000 | 0.11 | 0.32 | A>A |
| chr2 | 1.52E+08 | 152200000 | 0.64 | 0.65 | A>A |
| chr2 | 1.52E+08 | 152300000 | 0.87 | 0.75 | A>A |

| | | | | | |
|---|---|---|---|---|---|
| chr2 | 1.52E+08 | 152400000 | 0.71 | 0.55 | A>A |
| chr2 | 1.52E+08 | 152500000 | 0.74 | 0.32 | A>A |
| chr2 | 1.53E+08 | 152600000 | 0.81 | 0.48 | A>A |
| chr2 | 1.53E+08 | 152700000 | 0.94 | 0.43 | A>A |
| chr2 | 1.53E+08 | 152800000 | 0.85 | 0.30 | A>A |
| chr2 | 1.53E+08 | 152900000 | 0.65 | 0.25 | A>A |
| chr2 | 1.53E+08 | 153000000 | 0.91 | 0.61 | A>A |
| chr2 | 1.53E+08 | 153100000 | 0.93 | 0.75 | A>A |
| chr2 | 1.53E+08 | 153200000 | 1.11 | 1.11 | A>A |
| chr2 | 1.53E+08 | 153300000 | 1.19 | 1.13 | A>A |
| chr2 | 1.53E+08 | 153400000 | 1.18 | 1.17 | A>A |
| chr2 | 1.53E+08 | 153500000 | 0.98 | 0.85 | A>A |
| chr2 | 1.54E+08 | 153600000 | 0.55 | 0.38 | A>A |
| chr2 | 1.54E+08 | 153700000 | -0.25 | -0.44 | B>B |
| chr2 | 1.54E+08 | 153800000 | -0.48 | -0.62 | B>B |
| chr2 | 1.54E+08 | 153900000 | -0.64 | -0.75 | B>B |
| chr2 | 1.54E+08 | 154000000 | -0.47 | -0.63 | B>B |
| chr2 | 1.54E+08 | 154100000 | -0.71 | -0.78 | B>B |
| chr2 | 1.54E+08 | 154200000 | -0.74 | -0.82 | B>B |
| chr2 | 1.54E+08 | 154300000 | -0.78 | -0.85 | B>B |
| chr2 | 1.54E+08 | 154400000 | -0.46 | -0.63 | B>B |
| chr2 | 1.54E+08 | 154500000 | -0.63 | -0.82 | B>B |
| chr2 | 1.55E+08 | 154600000 | -0.74 | -0.86 | B>B |
| chr2 | 1.55E+08 | 154700000 | -0.66 | -0.80 | B>B |
| chr2 | 1.55E+08 | 154800000 | -0.08 | -0.51 | B>B |
| chr2 | 1.55E+08 | 154900000 | 0.02 | -0.43 | B>B |
| chr2 | 1.55E+08 | 155000000 | -0.05 | -0.58 | B>B |
| chr2 | 1.55E+08 | 155100000 | -0.06 | -0.50 | B>B |
| chr2 | 1.55E+08 | 155200000 | -0.14 | -0.57 | B>B |
| chr2 | 1.55E+08 | 155300000 | -0.35 | -0.52 | B>B |
| chr2 | 1.55E+08 | 155400000 | -0.37 | -0.50 | B>B |
| chr2 | 1.55E+08 | 155500000 | -0.54 | -0.67 | B>B |
| chr2 | 1.56E+08 | 155600000 | -0.62 | -0.72 | B>B |
| chr2 | 1.56E+08 | 155700000 | -0.78 | -0.78 | B>B |
| chr2 | 1.56E+08 | 155800000 | -0.87 | -0.78 | B>B |
| chr2 | 1.56E+08 | 155900000 | -0.84 | -0.80 | B>B |
| chr2 | 1.56E+08 | 156000000 | -0.91 | -0.81 | B>B |
| chr2 | 1.56E+08 | 156100000 | -0.81 | -0.76 | B>B |
| chr2 | 1.56E+08 | 156200000 | -0.72 | -0.65 | B>B |
| chr2 | 1.56E+08 | 156300000 | -0.67 | -0.58 | B>B |
| chr2 | 1.56E+08 | 156400000 | -0.88 | -0.80 | B>B |
| chr2 | 1.56E+08 | 156500000 | -0.82 | -0.83 | B>B |
| chr2 | 1.57E+08 | 156600000 | -0.89 | -0.77 | B>B |
| chr2 | 1.57E+08 | 156700000 | -0.82 | -0.80 | B>B |
| chr2 | 1.57E+08 | 156800000 | -0.84 | -0.74 | B>B |
| chr2 | 1.57E+08 | 156900000 | -0.66 | -0.59 | B>B |
| chr2 | 1.57E+08 | 157000000 | -0.53 | -0.48 | B>B |
| chr2 | 1.57E+08 | 157100000 | -0.44 | -0.41 | B>B |
| chr2 | 1.57E+08 | 157200000 | -0.22 | -0.28 | B>B |
| chr2 | 1.57E+08 | 157300000 | 0.19 | 0.14 | A>A |
| chr2 | 1.57E+08 | 157400000 | 0.20 | 0.25 | A>A |
| chr2 | 1.57E+08 | 157500000 | -0.13 | -0.25 | B>B |
| chr2 | 1.58E+08 | 157600000 | -0.38 | -0.56 | B>B |
| chr2 | 1.58E+08 | 157700000 | -0.43 | -0.58 | B>B |
| chr2 | 1.58E+08 | 157800000 | -0.36 | -0.32 | B>B |
| chr2 | 1.58E+08 | 157900000 | -0.21 | -0.14 | B>B |

| | | | | | |
|---|---|---|---|---|---|
| chr2 | 1.58E+08 | 158000000 | -0.04 | -0.06 | B>B |
| chr2 | 1.58E+08 | 158100000 | -0.01 | -0.04 | B>B |
| chr2 | 1.58E+08 | 158200000 | 0.05 | 0.10 | A>A |
| chr2 | 1.58E+08 | 158300000 | 0.06 | 0.14 | A>A |
| chr2 | 1.58E+08 | 158400000 | 0.03 | 0.11 | A>A |
| chr2 | 1.58E+08 | 158500000 | -0.13 | 0.08 | B>B |
| chr2 | 1.59E+08 | 158600000 | -0.01 | 0.22 | A>A |
| chr2 | 1.59E+08 | 158700000 | 0.32 | 0.43 | A>A |
| chr2 | 1.59E+08 | 158800000 | 0.46 | 0.51 | A>A |
| chr2 | 1.59E+08 | 158900000 | 0.12 | 0.22 | A>A |
| chr2 | 1.59E+08 | 159000000 | 0.19 | 0.43 | A>A |
| chr2 | 1.59E+08 | 159100000 | 0.20 | 0.41 | A>A |
| chr2 | 1.59E+08 | 159200000 | 0.23 | 0.43 | A>A |
| chr2 | 1.59E+08 | 159300000 | 0.39 | 0.49 | A>A |
| chr2 | 1.59E+08 | 159400000 | 0.52 | 0.72 | A>A |
| chr2 | 1.59E+08 | 159500000 | 0.45 | 0.75 | A>A |
| chr2 | 1.6E+08 | 159600000 | 0.57 | 0.66 | A>A |
| chr2 | 1.6E+08 | 159700000 | 0.51 | 0.69 | A>A |
| chr2 | 1.6E+08 | 159800000 | 0.91 | 0.99 | A>A |
| chr2 | 1.6E+08 | 159900000 | 1.33 | 1.43 | A>A |
| chr2 | 1.6E+08 | 160000000 | 1.12 | 1.36 | A>A |
| chr2 | 1.6E+08 | 160100000 | 0.94 | 1.28 | A>A |
| chr2 | 1.6E+08 | 160200000 | 1.14 | 1.37 | A>A |
| chr2 | 1.6E+08 | 160300000 | 1.12 | 1.40 | A>A |
| chr2 | 1.6E+08 | 160400000 | 1.23 | 1.52 | A>A |
| chr2 | 1.6E+08 | 160500000 | 1.35 | 1.65 | A>A |
| chr2 | 1.61E+08 | 160600000 | 1.21 | 1.45 | A>A |
| chr2 | 1.61E+08 | 160700000 | 0.97 | 1.25 | A>A |
| chr2 | 1.61E+08 | 160800000 | 1.21 | 1.27 | A>A |
| chr2 | 1.61E+08 | 160900000 | 0.93 | 1.03 | A>A |
| chr2 | 1.61E+08 | 161000000 | 0.79 | 0.90 | A>A |
| chr2 | 1.61E+08 | 161100000 | 1.36 | 1.37 | A>A |
| chr2 | 1.61E+08 | 161200000 | 0.97 | 1.11 | A>A |
| chr2 | 1.61E+08 | 161300000 | 1.12 | 1.27 | A>A |
| chr2 | 1.61E+08 | 161400000 | 1.22 | 1.28 | A>A |
| chr2 | 1.61E+08 | 161500000 | 1.21 | 1.30 | A>A |
| chr2 | 1.62E+08 | 161600000 | 1.29 | 1.40 | A>A |
| chr2 | 1.62E+08 | 161700000 | 1.30 | 1.48 | A>A |
| chr2 | 1.62E+08 | 161800000 | 1.23 | 1.17 | A>A |
| chr2 | 1.62E+08 | 161900000 | 0.66 | 0.75 | A>A |
| chr2 | 1.62E+08 | 162000000 | 0.92 | 1.00 | A>A |
| chr2 | 1.62E+08 | 162100000 | 0.70 | 0.76 | A>A |
| chr2 | 1.62E+08 | 162200000 | 0.32 | 0.29 | A>A |
| chr2 | 1.62E+08 | 162300000 | 0.14 | -0.08 | A>A |
| chr2 | 1.62E+08 | 162400000 | -0.48 | -0.67 | B>B |
| chr2 | 1.62E+08 | 162500000 | -0.59 | -0.76 | B>B |
| chr2 | 1.63E+08 | 162600000 | -0.80 | -0.91 | B>B |
| chr2 | 1.63E+08 | 162700000 | -0.77 | -0.92 | B>B |
| chr2 | 1.63E+08 | 162800000 | -0.74 | -0.62 | B>B |
| chr2 | 1.63E+08 | 162900000 | -0.42 | -0.27 | B>B |
| chr2 | 1.63E+08 | 163000000 | -0.38 | -0.27 | B>B |
| chr2 | 1.63E+08 | 163100000 | -0.58 | -0.71 | B>B |
| chr2 | 1.63E+08 | 163200000 | -0.41 | -0.47 | B>B |
| chr2 | 1.63E+08 | 163300000 | -0.58 | -0.65 | B>B |
| chr2 | 1.63E+08 | 163400000 | -0.69 | -0.74 | B>B |
| chr2 | 1.63E+08 | 163500000 | -0.67 | -0.73 | B>B |

| | | | | | |
|---|---|---|---|---|---|
| chr2 | 1.64E+08 | 163600000 | -0.79 | -0.60 | B>B |
| chr2 | 1.64E+08 | 163700000 | -0.84 | -0.64 | B>B |
| chr2 | 1.64E+08 | 163800000 | -0.71 | -0.44 | B>B |
| chr2 | 1.64E+08 | 163900000 | -0.88 | -0.56 | B>B |
| chr2 | 1.64E+08 | 164000000 | -0.59 | -0.46 | B>B |
| chr2 | 1.64E+08 | 164100000 | -0.07 | 0.05 | B>B |
| chr2 | 1.64E+08 | 164200000 | 0.19 | 0.34 | A>A |
| chr2 | 1.64E+08 | 164300000 | 0.43 | 0.46 | A>A |
| chr2 | 1.64E+08 | 164400000 | 0.49 | 0.55 | A>A |
| chr2 | 1.64E+08 | 164500000 | 0.59 | 0.63 | A>A |
| chr2 | 1.65E+08 | 164600000 | 0.60 | 0.56 | A>A |
| chr2 | 1.65E+08 | 164700000 | 0.56 | 0.65 | A>A |
| chr2 | 1.65E+08 | 164800000 | 0.51 | 0.56 | A>A |
| chr2 | 1.65E+08 | 164900000 | 0.46 | 0.50 | A>A |
| chr2 | 1.65E+08 | 165000000 | 0.27 | 0.33 | A>A |
| chr2 | 1.65E+08 | 165100000 | 0.22 | 0.31 | A>A |
| chr2 | 1.65E+08 | 165200000 | 0.15 | 0.24 | A>A |
| chr2 | 1.65E+08 | 165300000 | 0.10 | 0.26 | A>A |
| chr2 | 1.65E+08 | 165400000 | 0.12 | 0.18 | A>A |
| chr2 | 1.65E+08 | 165500000 | 0.49 | 0.55 | A>A |
| chr2 | 1.66E+08 | 165600000 | 0.38 | 0.49 | A>A |
| chr2 | 1.66E+08 | 165700000 | 0.32 | 0.53 | A>A |
| chr2 | 1.66E+08 | 165800000 | 0.31 | 0.47 | A>A |
| chr2 | 1.66E+08 | 165900000 | -0.08 | 0.07 | B>B |
| chr2 | 1.66E+08 | 166000000 | -0.36 | 0.00 | B>B |
| chr2 | 1.66E+08 | 166100000 | -0.50 | -0.13 | B>B |
| chr2 | 1.66E+08 | 166200000 | -0.51 | -0.15 | B>B |
| chr2 | 1.66E+08 | 166300000 | -0.52 | -0.06 | B>B |
| chr2 | 1.66E+08 | 166400000 | -0.67 | 0.03 | B>A |
| chr2 | 1.66E+08 | 166500000 | -0.62 | 0.19 | B>A |
| chr2 | 1.67E+08 | 166600000 | -0.56 | 0.04 | B>A |
| chr2 | 1.67E+08 | 166700000 | -0.45 | -0.10 | B>B |
| chr2 | 1.67E+08 | 166800000 | -0.17 | -0.21 | B>B |
| chr2 | 1.67E+08 | 166900000 | -0.47 | -0.66 | B>B |
| chr2 | 1.67E+08 | 167000000 | -0.51 | -0.69 | B>B |
| chr2 | 1.67E+08 | 167100000 | -0.56 | -0.71 | B>B |
| chr2 | 1.67E+08 | 167200000 | -0.53 | -0.82 | B>B |
| chr2 | 1.67E+08 | 167300000 | -0.45 | -0.69 | B>B |
| chr2 | 1.67E+08 | 167400000 | -0.68 | -0.82 | B>B |
| chr2 | 1.67E+08 | 167500000 | -0.85 | -0.90 | B>B |
| chr2 | 1.68E+08 | 167600000 | -0.90 | -0.95 | B>B |
| chr2 | 1.68E+08 | 167700000 | -0.96 | -0.93 | B>B |
| chr2 | 1.68E+08 | 167800000 | -0.94 | -0.95 | B>B |
| chr2 | 1.68E+08 | 167900000 | -0.95 | -0.92 | B>B |
| chr2 | 1.68E+08 | 168000000 | -0.89 | -0.91 | B>B |
| chr2 | 1.68E+08 | 168100000 | -0.84 | -0.90 | B>B |
| chr2 | 1.68E+08 | 168200000 | -0.80 | -0.88 | B>B |
| chr2 | 1.68E+08 | 168300000 | -0.81 | -0.89 | B>B |
| chr2 | 1.68E+08 | 168400000 | -0.83 | -0.89 | B>B |
| chr2 | 1.68E+08 | 168500000 | -0.84 | -0.92 | B>B |
| chr2 | 1.69E+08 | 168600000 | -0.81 | -0.94 | B>B |
| chr2 | 1.69E+08 | 168700000 | -0.66 | -0.93 | B>B |
| chr2 | 1.69E+08 | 168800000 | -0.51 | -0.82 | B>B |
| chr2 | 1.69E+08 | 168900000 | -0.22 | -0.38 | B>B |
| chr2 | 1.69E+08 | 169000000 | -0.14 | -0.21 | B>B |
| chr2 | 1.69E+08 | 169100000 | 0.21 | 0.08 | A>A |

| | | | | | |
|---|---|---|---|---|---|
| chr2 | 1.69E+08 | 169200000 | 0.13 | -0.01 | A>A |
| chr2 | 1.69E+08 | 169300000 | 0.19 | -0.01 | A>A |
| chr2 | 1.69E+08 | 169400000 | 0.56 | 0.28 | A>A |
| chr2 | 1.69E+08 | 169500000 | 0.51 | 0.29 | A>A |
| chr2 | 1.7E+08 | 169600000 | 0.30 | 0.08 | A>A |
| chr2 | 1.7E+08 | 169700000 | 0.36 | 0.06 | A>A |
| chr2 | 1.7E+08 | 169800000 | 0.21 | -0.04 | A>A |
| chr2 | 1.7E+08 | 169900000 | 0.20 | 0.00 | A>A |
| chr2 | 1.7E+08 | 170000000 | 0.24 | 0.02 | A>A |
| chr2 | 1.7E+08 | 170100000 | 0.01 | -0.38 | B>B |
| chr2 | 1.7E+08 | 170200000 | 0.06 | -0.40 | B>B |
| chr2 | 1.7E+08 | 170300000 | 0.24 | -0.25 | B>B |
| chr2 | 1.7E+08 | 170400000 | 0.76 | 0.43 | A>A |
| chr2 | 1.7E+08 | 170500000 | 1.22 | 0.78 | A>A |
| chr2 | 1.71E+08 | 170600000 | 1.22 | 0.80 | A>A |
| chr2 | 1.71E+08 | 170700000 | 1.06 | 0.75 | A>A |
| chr2 | 1.71E+08 | 170800000 | 0.97 | 0.67 | A>A |
| chr2 | 1.71E+08 | 170900000 | 0.89 | 0.45 | A>A |
| chr2 | 1.71E+08 | 171000000 | 0.92 | 0.48 | A>A |
| chr2 | 1.71E+08 | 171100000 | 0.73 | 0.22 | A>A |
| chr2 | 1.71E+08 | 171200000 | 0.45 | -0.12 | A>A |
| chr2 | 1.71E+08 | 171300000 | 0.25 | -0.25 | A>A |
| chr2 | 1.71E+08 | 171400000 | 0.41 | -0.36 | A>B |
| chr2 | 1.71E+08 | 171500000 | 0.31 | -0.47 | A>B |
| chr2 | 1.72E+08 | 171600000 | 0.44 | -0.31 | A>B |
| chr2 | 1.72E+08 | 171700000 | 0.42 | -0.18 | A>B |
| chr2 | 1.72E+08 | 171800000 | 0.77 | 0.23 | A>A |
| chr2 | 1.72E+08 | 171900000 | 0.99 | 0.97 | A>A |
| chr2 | 1.72E+08 | 172000000 | 1.45 | 1.27 | A>A |
| chr2 | 1.72E+08 | 172100000 | 1.33 | 1.22 | A>A |
| chr2 | 1.72E+08 | 172200000 | 1.33 | 1.18 | A>A |
| chr2 | 1.72E+08 | 172300000 | 1.13 | 1.15 | A>A |
| chr2 | 1.72E+08 | 172400000 | 1.34 | 1.38 | A>A |
| chr2 | 1.72E+08 | 172500000 | 1.32 | 1.28 | A>A |
| chr2 | 1.73E+08 | 172600000 | 1.33 | 1.39 | A>A |
| chr2 | 1.73E+08 | 172700000 | 1.37 | 1.44 | A>A |
| chr2 | 1.73E+08 | 172800000 | 1.64 | 1.78 | A>A |
| chr2 | 1.73E+08 | 172900000 | 1.67 | 1.80 | A>A |
| chr2 | 1.73E+08 | 173000000 | 1.44 | 1.54 | A>A |
| chr2 | 1.73E+08 | 173100000 | 1.11 | 1.25 | A>A |
| chr2 | 1.73E+08 | 173200000 | 1.62 | 1.72 | A>A |
| chr2 | 1.73E+08 | 173300000 | 1.67 | 1.74 | A>A |
| chr2 | 1.73E+08 | 173400000 | 1.18 | 1.36 | A>A |
| chr2 | 1.73E+08 | 173500000 | 1.47 | 1.58 | A>A |
| chr2 | 1.74E+08 | 173600000 | 1.54 | 1.49 | A>A |
| chr2 | 1.74E+08 | 173700000 | 1.21 | 1.26 | A>A |
| chr2 | 1.74E+08 | 173800000 | 1.40 | 1.45 | A>A |
| chr2 | 1.74E+08 | 173900000 | 1.29 | 1.28 | A>A |
| chr2 | 1.74E+08 | 174000000 | 1.61 | 1.70 | A>A |
| chr2 | 1.74E+08 | 174100000 | 1.45 | 1.64 | A>A |
| chr2 | 1.74E+08 | 174200000 | 1.30 | 1.36 | A>A |
| chr2 | 1.74E+08 | 174300000 | 1.05 | 1.09 | A>A |
| chr2 | 1.74E+08 | 174400000 | 1.28 | 1.36 | A>A |
| chr2 | 1.74E+08 | 174500000 | 1.22 | 1.42 | A>A |
| chr2 | 1.75E+08 | 174600000 | 1.30 | 1.26 | A>A |
| chr2 | 1.75E+08 | 174700000 | 1.21 | 1.28 | A>A |

| | | | | | |
|---|---|---|---|---|---|
| chr2 | 1.75E+08 | 174800000 | 1.18 | 1.24 | A>A |
| chr2 | 1.75E+08 | 174900000 | 1.50 | 1.73 | A>A |
| chr2 | 1.75E+08 | 175000000 | 1.19 | 1.42 | A>A |
| chr2 | 1.75E+08 | 175100000 | 1.46 | 1.55 | A>A |
| chr2 | 1.75E+08 | 175200000 | 1.33 | 1.54 | A>A |
| chr2 | 1.75E+08 | 175300000 | 1.04 | 1.11 | A>A |
| chr2 | 1.75E+08 | 175400000 | 1.16 | 1.14 | A>A |
| chr2 | 1.75E+08 | 175500000 | 1.05 | 1.03 | A>A |
| chr2 | 1.76E+08 | 175600000 | 0.97 | 0.84 | A>A |
| chr2 | 1.76E+08 | 175700000 | 0.85 | 0.50 | A>A |
| chr2 | 1.76E+08 | 175800000 | 0.88 | 0.65 | A>A |
| chr2 | 1.76E+08 | 175900000 | 0.74 | 0.51 | A>A |
| chr2 | 1.76E+08 | 176000000 | 0.65 | 0.38 | A>A |
| chr2 | 1.76E+08 | 176100000 | 0.42 | -0.08 | A>A |
| chr2 | 1.76E+08 | 176200000 | -0.11 | -0.56 | B>B |
| chr2 | 1.76E+08 | 176300000 | -0.37 | -0.78 | B>B |
| chr2 | 1.76E+08 | 176400000 | -0.62 | -0.98 | B>B |
| chr2 | 1.76E+08 | 176500000 | -0.65 | -0.99 | B>B |
| chr2 | 1.77E+08 | 176600000 | -0.53 | -0.94 | B>B |
| chr2 | 1.77E+08 | 176700000 | -0.56 | -0.98 | B>B |
| chr2 | 1.77E+08 | 176800000 | -0.39 | -0.80 | B>B |
| chr2 | 1.77E+08 | 176900000 | -0.08 | -0.36 | B>B |
| chr2 | 1.77E+08 | 177000000 | -0.13 | -0.30 | B>B |
| chr2 | 1.77E+08 | 177100000 | 0.15 | -0.30 | B>B |
| chr2 | 1.77E+08 | 177200000 | 0.45 | -0.07 | A>A |
| chr2 | 1.77E+08 | 177300000 | 0.26 | -0.18 | A>A |
| chr2 | 1.77E+08 | 177400000 | 0.20 | -0.39 | A>B |
| chr2 | 1.77E+08 | 177500000 | 0.23 | -0.29 | B>B |
| chr2 | 1.78E+08 | 177600000 | -0.22 | -0.63 | B>B |
| chr2 | 1.78E+08 | 177700000 | -0.36 | -0.74 | B>B |
| chr2 | 1.78E+08 | 177800000 | -0.47 | -0.71 | B>B |
| chr2 | 1.78E+08 | 177900000 | -0.44 | -0.80 | B>B |
| chr2 | 1.78E+08 | 178000000 | -0.35 | -0.69 | B>B |
| chr2 | 1.78E+08 | 178100000 | 0.36 | 0.08 | A>A |
| chr2 | 1.78E+08 | 178200000 | 0.57 | 0.31 | A>A |
| chr2 | 1.78E+08 | 178300000 | 0.41 | 0.14 | A>A |
| chr2 | 1.78E+08 | 178400000 | 0.41 | 0.11 | A>A |
| chr2 | 1.78E+08 | 178500000 | 0.32 | -0.24 | A>A |
| chr2 | 1.79E+08 | 178600000 | 0.12 | -0.40 | B>B |
| chr2 | 1.79E+08 | 178700000 | -0.26 | -0.70 | B>B |
| chr2 | 1.79E+08 | 178800000 | -0.44 | -0.96 | B>B |
| chr2 | 1.79E+08 | 178900000 | -0.30 | -0.93 | A>B |
| chr2 | 1.79E+08 | 179000000 | 0.01 | -0.68 | A>B |
| chr2 | 1.79E+08 | 179100000 | 0.44 | -0.58 | A>B |
| chr2 | 1.79E+08 | 179200000 | 0.42 | -0.66 | A>B |
| chr2 | 1.79E+08 | 179300000 | 0.26 | -0.51 | A>B |
| chr2 | 1.79E+08 | 179400000 | 0.29 | -0.15 | A>A |
| chr2 | 1.79E+08 | 179500000 | -0.32 | -0.63 | B>B |
| chr2 | 1.8E+08 | 179600000 | -0.63 | -0.53 | B>B |
| chr2 | 1.8E+08 | 179700000 | -0.52 | -0.04 | B>B |
| chr2 | 1.8E+08 | 179800000 | -0.50 | 0.13 | B>A |
| chr2 | 1.8E+08 | 179900000 | -0.57 | 0.24 | B>A |
| chr2 | 1.8E+08 | 180000000 | -0.32 | 0.29 | B>A |
| chr2 | 1.8E+08 | 180100000 | -0.01 | 0.39 | A>A |
| chr2 | 1.8E+08 | 180200000 | 0.12 | 0.76 | B>A |
| chr2 | 1.8E+08 | 180300000 | -0.36 | 0.37 | B>A |

| | | | | | |
|---|---|---|---|---|---|
| chr2 | 1.8E+08 | 180400000 | -0.64 | 0.62 | B>A |
| chr2 | 1.8E+08 | 180500000 | -0.75 | 0.79 | B>A |
| chr2 | 1.81E+08 | 180600000 | -0.72 | 0.68 | B>A |
| chr2 | 1.81E+08 | 180700000 | -0.63 | 0.30 | B>A |
| chr2 | 1.81E+08 | 180800000 | -0.49 | 0.37 | B>A |
| chr2 | 1.81E+08 | 180900000 | -0.19 | 0.34 | B>A |
| chr2 | 1.81E+08 | 181000000 | -0.33 | 0.13 | B>B |
| chr2 | 1.81E+08 | 181100000 | -0.02 | 0.32 | A>A |
| chr2 | 1.81E+08 | 181200000 | 0.08 | 0.41 | A>A |
| chr2 | 1.81E+08 | 181300000 | 0.19 | 0.59 | A>A |
| chr2 | 1.81E+08 | 181400000 | 0.16 | 0.49 | A>A |
| chr2 | 1.81E+08 | 181500000 | 0.30 | 0.59 | A>A |
| chr2 | 1.82E+08 | 181600000 | 0.41 | 0.77 | A>A |
| chr2 | 1.82E+08 | 181700000 | 0.37 | 0.75 | A>A |
| chr2 | 1.82E+08 | 181800000 | 0.31 | 0.58 | A>A |
| chr2 | 1.82E+08 | 181900000 | 0.28 | 0.52 | A>A |
| chr2 | 1.82E+08 | 182000000 | 0.01 | 0.18 | A>A |
| chr2 | 1.82E+08 | 182100000 | -0.15 | 0.28 | A>A |
| chr2 | 1.82E+08 | 182200000 | -0.21 | 0.19 | B>B |
| chr2 | 1.82E+08 | 182300000 | -0.14 | 0.20 | A>A |
| chr2 | 1.82E+08 | 182400000 | -0.11 | 0.19 | A>A |
| chr2 | 1.82E+08 | 182500000 | 0.11 | 0.21 | A>A |
| chr2 | 1.83E+08 | 182600000 | 0.07 | 0.19 | A>A |
| chr2 | 1.83E+08 | 182700000 | 0.48 | 0.57 | A>A |
| chr2 | 1.83E+08 | 182800000 | 0.54 | 0.62 | A>A |
| chr2 | 1.83E+08 | 182900000 | 0.16 | 0.36 | A>A |
| chr2 | 1.83E+08 | 183000000 | 0.05 | 0.15 | A>A |
| chr2 | 1.83E+08 | 183100000 | 0.08 | -0.04 | A>A |
| chr2 | 1.83E+08 | 183200000 | 0.33 | 0.03 | A>A |
| chr2 | 1.83E+08 | 183300000 | 0.51 | 0.09 | A>A |
| chr2 | 1.83E+08 | 183400000 | 0.46 | 0.05 | A>A |
| chr2 | 1.83E+08 | 183500000 | 0.39 | 0.02 | A>A |
| chr2 | 1.84E+08 | 183600000 | 0.27 | 0.02 | A>A |
| chr2 | 1.84E+08 | 183700000 | 0.55 | 0.13 | A>A |
| chr2 | 1.84E+08 | 183800000 | 0.73 | 0.26 | A>A |
| chr2 | 1.84E+08 | 183900000 | 0.69 | 0.47 | A>A |
| chr2 | 1.84E+08 | 184000000 | 0.60 | 0.31 | A>A |
| chr2 | 1.84E+08 | 184100000 | 0.16 | -0.11 | A>A |
| chr2 | 1.84E+08 | 184200000 | -0.45 | -0.64 | B>B |
| chr2 | 1.84E+08 | 184300000 | -0.75 | -0.81 | B>B |
| chr2 | 1.84E+08 | 184400000 | -0.85 | -0.88 | B>B |
| chr2 | 1.84E+08 | 184500000 | -0.84 | -0.92 | B>B |
| chr2 | 1.85E+08 | 184600000 | -0.85 | -0.88 | B>B |
| chr2 | 1.85E+08 | 184700000 | -0.81 | -0.82 | B>B |
| chr2 | 1.85E+08 | 184800000 | -0.78 | -0.83 | B>B |
| chr2 | 1.85E+08 | 184900000 | -0.81 | -0.88 | B>B |
| chr2 | 1.85E+08 | 185000000 | -0.75 | -0.81 | B>B |
| chr2 | 1.85E+08 | 185100000 | -0.82 | -0.87 | B>B |
| chr2 | 1.85E+08 | 185200000 | -0.83 | -0.85 | B>B |
| chr2 | 1.85E+08 | 185300000 | -0.78 | -0.84 | B>B |
| chr2 | 1.85E+08 | 185400000 | -0.74 | -0.83 | B>B |
| chr2 | 1.85E+08 | 185500000 | -0.67 | -0.81 | B>B |
| chr2 | 1.86E+08 | 185600000 | -0.75 | -0.79 | B>B |
| chr2 | 1.86E+08 | 185700000 | -0.74 | -0.79 | B>B |
| chr2 | 1.86E+08 | 185800000 | -0.74 | -0.79 | B>B |
| chr2 | 1.86E+08 | 185900000 | -0.75 | -0.79 | B>B |

| | | | | | |
|---|---|---|---|---|---|
| chr2 | 1.86E+08 | 186000000 | -0.72 | -0.80 | B>B |
| chr2 | 1.86E+08 | 186100000 | -0.72 | -0.75 | B>B |
| chr2 | 1.86E+08 | 186200000 | -0.73 | -0.71 | B>B |
| chr2 | 1.86E+08 | 186300000 | -0.71 | -0.68 | B>B |
| chr2 | 1.86E+08 | 186400000 | -0.69 | -0.52 | B>B |
| chr2 | 1.86E+08 | 186500000 | -0.67 | -0.34 | B>B |
| chr2 | 1.87E+08 | 186600000 | -0.71 | -0.41 | B>B |
| chr2 | 1.87E+08 | 186700000 | -0.64 | -0.37 | B>B |
| chr2 | 1.87E+08 | 186800000 | -0.63 | -0.39 | B>B |
| chr2 | 1.87E+08 | 186900000 | -0.68 | -0.40 | B>B |
| chr2 | 1.87E+08 | 187000000 | -0.64 | -0.41 | B>B |
| chr2 | 1.87E+08 | 187100000 | -0.53 | -0.40 | B>B |
| chr2 | 1.87E+08 | 187200000 | -0.39 | -0.36 | B>B |
| chr2 | 1.87E+08 | 187300000 | -0.28 | -0.21 | B>B |
| chr2 | 1.87E+08 | 187400000 | -0.03 | -0.04 | B>B |
| chr2 | 1.87E+08 | 187500000 | 0.44 | 0.25 | A>A |
| chr2 | 1.88E+08 | 187600000 | 0.08 | 0.02 | A>A |
| chr2 | 1.88E+08 | 187700000 | -0.26 | -0.18 | B>B |
| chr2 | 1.88E+08 | 187800000 | -0.37 | -0.45 | B>B |
| chr2 | 1.88E+08 | 187900000 | -0.40 | -0.69 | B>B |
| chr2 | 1.88E+08 | 188000000 | -0.57 | -0.74 | B>B |
| chr2 | 1.88E+08 | 188100000 | -0.55 | -0.74 | B>B |
| chr2 | 1.88E+08 | 188200000 | -0.49 | -0.61 | B>B |
| chr2 | 1.88E+08 | 188300000 | -0.31 | -0.41 | B>B |
| chr2 | 1.88E+08 | 188400000 | -0.10 | 0.01 | B>B |
| chr2 | 1.88E+08 | 188500000 | 0.14 | 0.27 | A>A |
| chr2 | 1.89E+08 | 188600000 | -0.04 | 0.25 | A>A |
| chr2 | 1.89E+08 | 188700000 | -0.05 | 0.30 | A>A |
| chr2 | 1.89E+08 | 188800000 | -0.05 | 0.26 | A>A |
| chr2 | 1.89E+08 | 188900000 | 0.03 | 0.53 | A>A |
| chr2 | 1.89E+08 | 189000000 | 0.10 | 0.52 | A>A |
| chr2 | 1.89E+08 | 189100000 | 0.18 | 0.70 | B>A |
| chr2 | 1.89E+08 | 189200000 | 0.32 | 0.71 | A>A |
| chr2 | 1.89E+08 | 189300000 | 0.32 | 0.72 | A>A |
| chr2 | 1.89E+08 | 189400000 | 0.19 | 0.51 | A>A |
| chr2 | 1.89E+08 | 189500000 | 0.21 | 0.57 | A>A |
| chr2 | 1.9E+08 | 189600000 | 0.24 | 0.58 | A>A |
| chr2 | 1.9E+08 | 189700000 | 0.14 | 0.66 | B>A |
| chr2 | 1.9E+08 | 189800000 | 0.24 | 0.65 | A>A |
| chr2 | 1.9E+08 | 189900000 | 0.29 | 0.53 | A>A |
| chr2 | 1.9E+08 | 190000000 | 0.55 | 0.80 | A>A |
| chr2 | 1.9E+08 | 190100000 | 1.04 | 1.28 | A>A |
| chr2 | 1.9E+08 | 190200000 | 0.75 | 0.98 | A>A |
| chr2 | 1.9E+08 | 190300000 | 0.51 | 0.76 | A>A |
| chr2 | 1.9E+08 | 190400000 | 0.99 | 1.19 | A>A |
| chr2 | 1.9E+08 | 190500000 | 1.30 | 1.45 | A>A |
| chr2 | 1.91E+08 | 190600000 | 1.04 | 1.25 | A>A |
| chr2 | 1.91E+08 | 190700000 | 1.06 | 1.34 | A>A |
| chr2 | 1.91E+08 | 190800000 | 1.26 | 1.40 | A>A |
| chr2 | 1.91E+08 | 190900000 | 1.14 | 1.26 | A>A |
| chr2 | 1.91E+08 | 191000000 | 0.89 | 1.10 | A>A |
| chr2 | 1.91E+08 | 191100000 | 1.21 | 1.37 | A>A |
| chr2 | 1.91E+08 | 191200000 | 1.12 | 1.14 | A>A |
| chr2 | 1.91E+08 | 191300000 | 1.05 | 1.25 | A>A |
| chr2 | 1.91E+08 | 191400000 | 1.24 | 1.44 | A>A |
| chr2 | 1.91E+08 | 191500000 | 1.34 | 1.41 | A>A |

| | | | | | |
|---|---|---|---|---|---|
| chr2 | 1.92E+08 | 191600000 | 1.13 | 1.35 | A>A |
| chr2 | 1.92E+08 | 191700000 | 0.91 | 1.12 | A>A |
| chr2 | 1.92E+08 | 191800000 | 1.36 | 1.50 | A>A |
| chr2 | 1.92E+08 | 191900000 | 0.93 | 1.21 | A>A |
| chr2 | 1.92E+08 | 192000000 | 0.77 | 0.97 | A>A |
| chr2 | 1.92E+08 | 192100000 | 0.74 | 0.89 | A>A |
| chr2 | 1.92E+08 | 192200000 | 0.89 | 1.04 | A>A |
| chr2 | 1.92E+08 | 192300000 | 0.82 | 1.12 | A>A |
| chr2 | 1.92E+08 | 192400000 | 0.71 | 0.79 | A>A |
| chr2 | 1.92E+08 | 192500000 | 0.40 | 0.58 | A>A |
| chr2 | 1.93E+08 | 192600000 | 0.44 | 0.66 | A>A |
| chr2 | 1.93E+08 | 192700000 | 0.06 | 0.31 | A>A |
| chr2 | 1.93E+08 | 192800000 | 0.27 | 0.45 | A>A |
| chr2 | 1.93E+08 | 192900000 | 0.27 | 0.55 | A>A |
| chr2 | 1.93E+08 | 193000000 | 0.11 | 0.35 | A>A |
| chr2 | 1.93E+08 | 193100000 | -0.15 | 0.09 | B>B |
| chr2 | 1.93E+08 | 193200000 | -0.34 | 0.11 | B>B |
| chr2 | 1.93E+08 | 193300000 | -0.40 | 0.22 | B>A |
| chr2 | 1.93E+08 | 193400000 | -0.33 | 0.19 | B>A |
| chr2 | 1.93E+08 | 193500000 | -0.37 | 0.19 | B>A |
| chr2 | 1.94E+08 | 193600000 | -0.53 | -0.02 | B>A |
| chr2 | 1.94E+08 | 193700000 | -0.59 | -0.12 | B>B |
| chr2 | 1.94E+08 | 193800000 | -0.62 | -0.08 | B>A |
| chr2 | 1.94E+08 | 193900000 | -0.64 | -0.13 | B>A |
| chr2 | 1.94E+08 | 194000000 | -0.55 | -0.11 | B>B |
| chr2 | 1.94E+08 | 194100000 | -0.52 | -0.07 | B>B |
| chr2 | 1.94E+08 | 194200000 | -0.53 | -0.17 | B>B |
| chr2 | 1.94E+08 | 194300000 | -0.54 | -0.10 | B>B |
| chr2 | 1.94E+08 | 194400000 | -0.51 | -0.07 | B>B |
| chr2 | 1.94E+08 | 194500000 | -0.58 | -0.24 | B>B |
| chr2 | 1.95E+08 | 194600000 | -0.57 | -0.38 | B>B |
| chr2 | 1.95E+08 | 194700000 | -0.53 | -0.39 | B>B |
| chr2 | 1.95E+08 | 194800000 | -0.59 | -0.38 | B>B |
| chr2 | 1.95E+08 | 194900000 | -0.60 | -0.51 | B>B |
| chr2 | 1.95E+08 | 195000000 | -0.59 | -0.63 | B>B |
| chr2 | 1.95E+08 | 195100000 | -0.61 | -0.68 | B>B |
| chr2 | 1.95E+08 | 195200000 | -0.61 | -0.70 | B>B |
| chr2 | 1.95E+08 | 195300000 | -0.60 | -0.68 | B>B |
| chr2 | 1.95E+08 | 195400000 | -0.55 | -0.60 | B>B |
| chr2 | 1.95E+08 | 195500000 | -0.54 | -0.52 | B>B |
| chr2 | 1.96E+08 | 195600000 | -0.49 | -0.54 | B>B |
| chr2 | 1.96E+08 | 195700000 | -0.42 | -0.53 | B>B |
| chr2 | 1.96E+08 | 195800000 | -0.37 | -0.46 | B>B |
| chr2 | 1.96E+08 | 195900000 | -0.30 | -0.43 | B>B |
| chr2 | 1.96E+08 | 196000000 | -0.28 | -0.40 | B>B |
| chr2 | 1.96E+08 | 196100000 | -0.33 | -0.38 | B>B |
| chr2 | 1.96E+08 | 196200000 | -0.19 | -0.21 | B>B |
| chr2 | 1.96E+08 | 196300000 | 0.13 | 0.15 | A>A |
| chr2 | 1.96E+08 | 196400000 | 0.21 | 0.20 | A>A |
| chr2 | 1.96E+08 | 196500000 | 0.28 | 0.25 | A>A |
| chr2 | 1.97E+08 | 196600000 | 0.14 | 0.11 | A>A |
| chr2 | 1.97E+08 | 196700000 | 0.01 | 0.04 | A>A |
| chr2 | 1.97E+08 | 196800000 | -0.14 | -0.01 | B>B |
| chr2 | 1.97E+08 | 196900000 | -0.25 | -0.05 | B>B |
| chr2 | 1.97E+08 | 197000000 | -0.01 | 0.14 | A>A |
| chr2 | 1.97E+08 | 197100000 | 0.24 | 0.58 | A>A |

| | | | | | |
|---|---|---|---|---|---|
| chr2 | 1.97E+08 | 197200000 | 0.26 | 0.52 | A>A |
| chr2 | 1.97E+08 | 197300000 | 0.18 | 0.55 | A>A |
| chr2 | 1.97E+08 | 197400000 | 0.24 | 0.69 | A>A |
| chr2 | 1.97E+08 | 197500000 | 0.39 | 0.74 | A>A |
| chr2 | 1.98E+08 | 197600000 | 0.45 | 0.73 | A>A |
| chr2 | 1.98E+08 | 197700000 | 0.40 | 0.59 | A>A |
| chr2 | 1.98E+08 | 197800000 | 0.58 | 0.68 | A>A |
| chr2 | 1.98E+08 | 197900000 | 0.51 | 0.53 | A>A |
| chr2 | 1.98E+08 | 198000000 | 0.43 | 0.47 | A>A |
| chr2 | 1.98E+08 | 198100000 | 0.46 | 0.49 | A>A |
| chr2 | 1.98E+08 | 198200000 | 0.60 | 0.56 | A>A |
| chr2 | 1.98E+08 | 198300000 | 0.50 | 0.45 | A>A |
| chr2 | 1.98E+08 | 198400000 | 0.70 | 0.59 | A>A |
| chr2 | 1.98E+08 | 198500000 | 0.48 | 0.30 | A>A |
| chr2 | 1.99E+08 | 198600000 | 0.19 | 0.05 | A>A |
| chr2 | 1.99E+08 | 198700000 | -0.21 | -0.40 | B>B |
| chr2 | 1.99E+08 | 198800000 | -0.53 | -0.85 | B>B |
| chr2 | 1.99E+08 | 198900000 | -0.69 | -0.94 | B>B |
| chr2 | 1.99E+08 | 199000000 | -0.59 | -0.94 | B>B |
| chr2 | 1.99E+08 | 199100000 | -0.68 | -0.89 | B>B |
| chr2 | 1.99E+08 | 199200000 | -0.46 | -0.65 | B>B |
| chr2 | 1.99E+08 | 199300000 | -0.58 | -0.78 | B>B |
| chr2 | 1.99E+08 | 199400000 | -0.79 | -0.87 | B>B |
| chr2 | 1.99E+08 | 199500000 | -0.69 | -0.81 | B>B |
| chr2 | 2E+08 | 199600000 | -0.71 | -0.82 | B>B |
| chr2 | 2E+08 | 199700000 | -0.81 | -0.86 | B>B |
| chr2 | 2E+08 | 199800000 | -0.76 | -0.82 | B>B |
| chr2 | 2E+08 | 199900000 | -0.70 | -0.80 | B>B |
| chr2 | 2E+08 | 200000000 | -0.63 | -0.83 | B>B |
| chr2 | 2E+08 | 200100000 | -0.53 | -0.71 | B>B |
| chr2 | 2E+08 | 200200000 | -0.27 | -0.55 | B>B |
| chr2 | 2E+08 | 200300000 | 0.01 | -0.29 | B>B |
| chr2 | 2E+08 | 200400000 | -0.15 | -0.48 | B>B |
| chr2 | 2E+08 | 200500000 | -0.25 | -0.73 | B>B |
| chr2 | 2.01E+08 | 200600000 | -0.09 | -0.44 | B>B |
| chr2 | 2.01E+08 | 200700000 | 0.04 | -0.30 | B>B |
| chr2 | 2.01E+08 | 200800000 | 0.35 | 0.01 | A>A |
| chr2 | 2.01E+08 | 200900000 | 0.46 | 0.46 | A>A |
| chr2 | 2.01E+08 | 201000000 | 0.71 | 0.82 | A>A |
| chr2 | 2.01E+08 | 201100000 | 0.88 | 0.78 | A>A |
| chr2 | 2.01E+08 | 201200000 | 0.80 | 0.68 | A>A |
| chr2 | 2.01E+08 | 201300000 | 0.81 | 0.78 | A>A |
| chr2 | 2.01E+08 | 201400000 | 0.63 | 0.67 | A>A |
| chr2 | 2.01E+08 | 201500000 | 0.95 | 0.88 | A>A |
| chr2 | 2.02E+08 | 201600000 | 0.84 | 0.95 | A>A |
| chr2 | 2.02E+08 | 201700000 | 1.17 | 1.11 | A>A |
| chr2 | 2.02E+08 | 201800000 | 0.99 | 1.10 | A>A |
| chr2 | 2.02E+08 | 201900000 | 1.27 | 1.33 | A>A |
| chr2 | 2.02E+08 | 202000000 | 1.37 | 1.31 | A>A |
| chr2 | 2.02E+08 | 202100000 | 0.98 | 1.06 | A>A |
| chr2 | 2.02E+08 | 202200000 | 1.05 | 1.11 | A>A |
| chr2 | 2.02E+08 | 202300000 | 1.04 | 1.10 | A>A |
| chr2 | 2.02E+08 | 202400000 | 1.05 | 1.09 | A>A |
| chr2 | 2.02E+08 | 202500000 | 1.05 | 1.04 | A>A |
| chr2 | 2.03E+08 | 202600000 | 1.10 | 0.98 | A>A |
| chr2 | 2.03E+08 | 202700000 | 1.13 | 1.04 | A>A |

| | | | | | |
|---|---|---|---|---|---|
| chr2 | 2.03E+08 | 202800000 | 1.18 | 0.97 | A>A |
| chr2 | 2.03E+08 | 202900000 | 1.22 | 1.17 | A>A |
| chr2 | 2.03E+08 | 203000000 | 1.16 | 1.10 | A>A |
| chr2 | 2.03E+08 | 203100000 | 1.02 | 0.98 | A>A |
| chr2 | 2.03E+08 | 203200000 | 1.38 | 1.18 | A>A |
| chr2 | 2.03E+08 | 203300000 | 1.25 | 1.30 | A>A |
| chr2 | 2.03E+08 | 203400000 | 1.14 | 1.16 | A>A |
| chr2 | 2.03E+08 | 203500000 | 0.88 | 0.86 | A>A |
| chr2 | 2.04E+08 | 203600000 | 1.10 | 1.03 | A>A |
| chr2 | 2.04E+08 | 203700000 | 0.95 | 0.94 | A>A |
| chr2 | 2.04E+08 | 203800000 | 0.88 | 0.98 | A>A |
| chr2 | 2.04E+08 | 203900000 | 1.21 | 1.03 | A>A |
| chr2 | 2.04E+08 | 204000000 | 1.06 | 1.03 | A>A |
| chr2 | 2.04E+08 | 204100000 | 0.90 | 0.76 | A>A |
| chr2 | 2.04E+08 | 204200000 | 1.01 | 0.87 | A>A |
| chr2 | 2.04E+08 | 204300000 | 0.96 | 0.96 | A>A |
| chr2 | 2.04E+08 | 204400000 | 0.72 | 0.65 | A>A |
| chr2 | 2.04E+08 | 204500000 | 1.01 | 0.89 | A>A |
| chr2 | 2.05E+08 | 204600000 | 0.76 | 0.54 | A>A |
| chr2 | 2.05E+08 | 204700000 | 0.54 | 0.44 | A>A |
| chr2 | 2.05E+08 | 204800000 | 0.41 | 0.12 | A>A |
| chr2 | 2.05E+08 | 204900000 | 0.31 | -0.12 | A>A |
| chr2 | 2.05E+08 | 205000000 | 0.25 | -0.10 | A>A |
| chr2 | 2.05E+08 | 205100000 | 0.30 | -0.10 | A>A |
| chr2 | 2.05E+08 | 205200000 | 0.25 | -0.01 | A>A |
| chr2 | 2.05E+08 | 205300000 | 0.47 | 0.12 | A>A |
| chr2 | 2.05E+08 | 205400000 | 0.57 | 0.26 | A>A |
| chr2 | 2.05E+08 | 205500000 | 0.57 | 0.32 | A>A |
| chr2 | 2.06E+08 | 205600000 | 0.48 | 0.13 | A>A |
| chr2 | 2.06E+08 | 205700000 | 0.44 | 0.19 | A>A |
| chr2 | 2.06E+08 | 205800000 | 0.36 | 0.20 | A>A |
| chr2 | 2.06E+08 | 205900000 | 0.51 | 0.29 | A>A |
| chr2 | 2.06E+08 | 206000000 | 0.37 | 0.26 | A>A |
| chr2 | 2.06E+08 | 206100000 | 0.40 | 0.17 | A>A |
| chr2 | 2.06E+08 | 206200000 | 0.39 | 0.13 | A>A |
| chr2 | 2.06E+08 | 206300000 | 0.41 | 0.05 | A>A |
| chr2 | 2.06E+08 | 206400000 | 0.40 | 0.05 | A>A |
| chr2 | 2.06E+08 | 206500000 | 0.30 | 0.01 | A>A |
| chr2 | 2.07E+08 | 206600000 | 0.46 | 0.21 | A>A |
| chr2 | 2.07E+08 | 206700000 | 0.66 | 0.46 | A>A |
| chr2 | 2.07E+08 | 206800000 | 0.47 | 0.23 | A>A |
| chr2 | 2.07E+08 | 206900000 | 0.78 | 0.60 | A>A |
| chr2 | 2.07E+08 | 207000000 | 1.01 | 0.67 | A>A |
| chr2 | 2.07E+08 | 207100000 | 0.74 | 0.32 | A>A |
| chr2 | 2.07E+08 | 207200000 | 0.56 | 0.04 | A>A |
| chr2 | 2.07E+08 | 207300000 | 0.32 | -0.26 | A>A |
| chr2 | 2.07E+08 | 207400000 | 0.36 | -0.39 | A>B |
| chr2 | 2.07E+08 | 207500000 | 0.29 | -0.42 | A>B |
| chr2 | 2.08E+08 | 207600000 | 0.24 | -0.40 | A>B |
| chr2 | 2.08E+08 | 207700000 | 0.41 | -0.27 | A>B |
| chr2 | 2.08E+08 | 207800000 | 0.28 | -0.35 | A>B |
| chr2 | 2.08E+08 | 207900000 | 0.30 | -0.12 | A>A |
| chr2 | 2.08E+08 | 208000000 | 0.53 | 0.09 | A>A |
| chr2 | 2.08E+08 | 208100000 | 0.65 | 0.30 | A>A |
| chr2 | 2.08E+08 | 208200000 | 0.74 | 0.37 | A>A |
| chr2 | 2.08E+08 | 208300000 | 0.71 | 0.29 | A>A |

| | | | | | |
|---|---|---|---|---|---|
| chr2 | 2.08E+08 | 208400000 | 0.48 | 0.22 | A>A |
| chr2 | 2.08E+08 | 208500000 | 0.69 | 0.46 | A>A |
| chr2 | 2.09E+08 | 208600000 | 0.93 | 0.65 | A>A |
| chr2 | 2.09E+08 | 208700000 | 0.95 | 0.70 | A>A |
| chr2 | 2.09E+08 | 208800000 | 1.08 | 0.95 | A>A |
| chr2 | 2.09E+08 | 208900000 | 0.87 | 0.68 | A>A |
| chr2 | 2.09E+08 | 209000000 | 0.57 | 0.46 | A>A |
| chr2 | 2.09E+08 | 209100000 | 0.61 | 0.40 | A>A |
| chr2 | 2.09E+08 | 209200000 | 0.48 | 0.33 | A>A |
| chr2 | 2.09E+08 | 209300000 | -0.11 | -0.23 | B>B |
| chr2 | 2.09E+08 | 209400000 | -0.46 | -0.45 | B>B |
| chr2 | 2.09E+08 | 209500000 | -0.47 | -0.29 | B>B |
| chr2 | 2.1E+08 | 209600000 | -0.58 | -0.25 | B>B |
| chr2 | 2.1E+08 | 209700000 | -0.65 | -0.41 | B>B |
| chr2 | 2.1E+08 | 209800000 | -0.69 | -0.74 | B>B |
| chr2 | 2.1E+08 | 209900000 | -0.68 | -0.79 | B>B |
| chr2 | 2.1E+08 | 210000000 | -0.69 | -0.80 | B>B |
| chr2 | 2.1E+08 | 210100000 | -0.61 | -0.68 | B>B |
| chr2 | 2.1E+08 | 210200000 | -0.48 | -0.55 | B>B |
| chr2 | 2.1E+08 | 210300000 | -0.25 | -0.15 | B>B |
| chr2 | 2.1E+08 | 210400000 | -0.07 | -0.07 | B>B |
| chr2 | 2.1E+08 | 210500000 | -0.28 | -0.33 | B>B |
| chr2 | 2.11E+08 | 210600000 | -0.53 | -0.64 | B>B |
| chr2 | 2.11E+08 | 210700000 | -0.59 | -0.66 | B>B |
| chr2 | 2.11E+08 | 210800000 | -0.55 | -0.67 | B>B |
| chr2 | 2.11E+08 | 210900000 | -0.13 | -0.35 | B>B |
| chr2 | 2.11E+08 | 211000000 | -0.18 | -0.06 | B>B |
| chr2 | 2.11E+08 | 211100000 | -0.22 | -0.28 | B>B |
| chr2 | 2.11E+08 | 211200000 | -0.52 | -0.63 | B>B |
| chr2 | 2.11E+08 | 211300000 | -0.45 | -0.64 | B>B |
| chr2 | 2.11E+08 | 211400000 | -0.26 | -0.50 | B>B |
| chr2 | 2.11E+08 | 211500000 | -0.44 | -0.69 | B>B |
| chr2 | 2.12E+08 | 211600000 | -0.62 | -0.76 | B>B |
| chr2 | 2.12E+08 | 211700000 | -0.74 | -0.81 | B>B |
| chr2 | 2.12E+08 | 211800000 | -0.70 | -0.79 | B>B |
| chr2 | 2.12E+08 | 211900000 | -0.65 | -0.73 | B>B |
| chr2 | 2.12E+08 | 212000000 | -0.59 | -0.72 | B>B |
| chr2 | 2.12E+08 | 212100000 | -0.62 | -0.72 | B>B |
| chr2 | 2.12E+08 | 212200000 | -0.63 | -0.78 | B>B |
| chr2 | 2.12E+08 | 212300000 | -0.58 | -0.76 | B>B |
| chr2 | 2.12E+08 | 212400000 | -0.62 | -0.71 | B>B |
| chr2 | 2.12E+08 | 212500000 | -0.68 | -0.73 | B>B |
| chr2 | 2.13E+08 | 212600000 | -0.66 | -0.69 | B>B |
| chr2 | 2.13E+08 | 212700000 | -0.70 | -0.75 | B>B |
| chr2 | 2.13E+08 | 212800000 | -0.70 | -0.79 | B>B |
| chr2 | 2.13E+08 | 212900000 | -0.68 | -0.68 | B>B |
| chr2 | 2.13E+08 | 213000000 | -0.65 | -0.56 | B>B |
| chr2 | 2.13E+08 | 213100000 | -0.66 | -0.54 | B>B |
| chr2 | 2.13E+08 | 213200000 | -0.68 | -0.52 | B>B |
| chr2 | 2.13E+08 | 213300000 | -0.65 | -0.49 | B>B |
| chr2 | 2.13E+08 | 213400000 | -0.60 | -0.50 | B>B |
| chr2 | 2.13E+08 | 213500000 | -0.51 | -0.56 | B>B |
| chr2 | 2.14E+08 | 213600000 | -0.58 | -0.61 | B>B |
| chr2 | 2.14E+08 | 213700000 | -0.62 | -0.66 | B>B |
| chr2 | 2.14E+08 | 213800000 | -0.54 | -0.38 | B>B |
| chr2 | 2.14E+08 | 213900000 | -0.52 | -0.35 | B>B |

| | | | | | |
|---|---|---|---|---|---|
| chr2 | 2.14E+08 | 214000000 | -0.54 | -0.35 | B>B |
| chr2 | 2.14E+08 | 214100000 | -0.33 | -0.29 | B>B |
| chr2 | 2.14E+08 | 214200000 | -0.19 | -0.10 | B>B |
| chr2 | 2.14E+08 | 214300000 | -0.47 | -0.24 | B>B |
| chr2 | 2.14E+08 | 214400000 | -0.55 | -0.29 | B>B |
| chr2 | 2.14E+08 | 214500000 | -0.58 | -0.30 | B>B |
| chr2 | 2.15E+08 | 214600000 | -0.52 | -0.23 | B>B |
| chr2 | 2.15E+08 | 214700000 | -0.46 | -0.28 | B>B |
| chr2 | 2.15E+08 | 214800000 | -0.48 | -0.31 | B>B |
| chr2 | 2.15E+08 | 214900000 | -0.30 | -0.17 | B>B |
| chr2 | 2.15E+08 | 215000000 | -0.29 | -0.16 | B>B |
| chr2 | 2.15E+08 | 215100000 | -0.48 | -0.31 | B>B |
| chr2 | 2.15E+08 | 215200000 | -0.50 | -0.33 | B>B |
| chr2 | 2.15E+08 | 215300000 | -0.52 | -0.36 | B>B |
| chr2 | 2.15E+08 | 215400000 | -0.50 | -0.35 | B>B |
| chr2 | 2.15E+08 | 215500000 | -0.44 | -0.34 | B>B |
| chr2 | 2.16E+08 | 215600000 | -0.25 | -0.22 | B>B |
| chr2 | 2.16E+08 | 215700000 | -0.04 | -0.03 | B>B |
| chr2 | 2.16E+08 | 215800000 | -0.24 | -0.18 | B>B |
| chr2 | 2.16E+08 | 215900000 | -0.22 | -0.04 | B>B |
| chr2 | 2.16E+08 | 216000000 | -0.07 | 0.08 | A>A |
| chr2 | 2.16E+08 | 216100000 | 0.02 | 0.17 | A>A |
| chr2 | 2.16E+08 | 216200000 | 0.14 | 0.09 | A>A |
| chr2 | 2.16E+08 | 216300000 | 0.37 | 0.37 | A>A |
| chr2 | 2.16E+08 | 216400000 | 0.44 | 0.31 | A>A |
| chr2 | 2.16E+08 | 216500000 | 0.51 | 0.43 | A>A |
| chr2 | 2.17E+08 | 216600000 | 0.49 | 0.53 | A>A |
| chr2 | 2.17E+08 | 216700000 | 0.46 | 0.44 | A>A |
| chr2 | 2.17E+08 | 216800000 | 0.25 | 0.21 | A>A |
| chr2 | 2.17E+08 | 216900000 | 0.38 | 0.37 | A>A |
| chr2 | 2.17E+08 | 217000000 | 0.82 | 0.85 | A>A |
| chr2 | 2.17E+08 | 217100000 | 0.80 | 0.90 | A>A |
| chr2 | 2.17E+08 | 217200000 | 0.95 | 0.78 | A>A |
| chr2 | 2.17E+08 | 217300000 | 0.93 | 0.85 | A>A |
| chr2 | 2.17E+08 | 217400000 | 0.68 | 0.82 | A>A |
| chr2 | 2.17E+08 | 217500000 | 0.70 | 0.97 | A>A |
| chr2 | 2.18E+08 | 217600000 | 0.46 | 0.53 | A>A |
| chr2 | 2.18E+08 | 217700000 | 0.44 | 0.77 | A>A |
| chr2 | 2.18E+08 | 217800000 | 0.47 | 0.92 | A>A |
| chr2 | 2.18E+08 | 217900000 | 0.48 | 0.78 | A>A |
| chr2 | 2.18E+08 | 218000000 | 0.46 | 0.90 | A>A |
| chr2 | 2.18E+08 | 218100000 | 0.43 | 0.88 | A>A |
| chr2 | 2.18E+08 | 218200000 | 0.63 | 1.04 | A>A |
| chr2 | 2.18E+08 | 218300000 | 0.65 | 1.00 | A>A |
| chr2 | 2.18E+08 | 218400000 | 0.42 | 0.75 | A>A |
| chr2 | 2.18E+08 | 218500000 | 0.48 | 0.79 | A>A |
| chr2 | 2.19E+08 | 218600000 | 0.45 | 0.64 | A>A |
| chr2 | 2.19E+08 | 218700000 | 0.55 | 0.63 | A>A |
| chr2 | 2.19E+08 | 218800000 | 0.84 | 1.00 | A>A |
| chr2 | 2.19E+08 | 218900000 | 0.83 | 0.98 | A>A |
| chr2 | 2.19E+08 | 219000000 | 0.87 | 0.96 | A>A |
| chr2 | 2.19E+08 | 219100000 | 1.04 | 1.01 | A>A |
| chr2 | 2.19E+08 | 219200000 | 1.14 | 1.14 | A>A |
| chr2 | 2.19E+08 | 219300000 | 1.18 | 1.28 | A>A |
| chr2 | 2.19E+08 | 219400000 | 1.26 | 1.21 | A>A |
| chr2 | 2.19E+08 | 219500000 | 1.23 | 1.25 | A>A |

| | | | | | |
|---|---|---|---|---|---|
| chr2 | 2.2E+08 | 219600000 | 1.10 | 1.15 | A>A |
| chr2 | 2.2E+08 | 219700000 | 1.26 | 1.15 | A>A |
| chr2 | 2.2E+08 | 219800000 | 1.04 | 1.07 | A>A |
| chr2 | 2.2E+08 | 219900000 | 1.08 | 0.97 | A>A |
| chr2 | 2.2E+08 | 220000000 | 1.10 | 1.11 | A>A |
| chr2 | 2.2E+08 | 220100000 | 1.05 | 1.10 | A>A |
| chr2 | 2.2E+08 | 220200000 | 1.09 | 1.16 | A>A |
| chr2 | 2.2E+08 | 220300000 | 0.94 | 0.98 | A>A |
| chr2 | 2.2E+08 | 220400000 | 0.97 | 1.07 | A>A |
| chr2 | 2.2E+08 | 220500000 | 0.73 | 0.81 | A>A |
| chr2 | 2.21E+08 | 220600000 | 0.33 | 0.50 | A>A |
| chr2 | 2.21E+08 | 220700000 | 0.07 | 0.35 | A>A |
| chr2 | 2.21E+08 | 220800000 | 0.17 | 0.34 | A>A |
| chr2 | 2.21E+08 | 220900000 | 0.19 | 0.41 | A>A |
| chr2 | 2.21E+08 | 221000000 | 0.23 | 0.43 | A>A |
| chr2 | 2.21E+08 | 221100000 | 0.24 | 0.60 | A>A |
| chr2 | 2.21E+08 | 221200000 | 0.21 | 0.47 | A>A |
| chr2 | 2.21E+08 | 221300000 | 0.24 | 0.55 | A>A |
| chr2 | 2.21E+08 | 221400000 | 0.18 | 0.56 | A>A |
| chr2 | 2.21E+08 | 221500000 | 0.18 | 0.61 | A>A |
| chr2 | 2.22E+08 | 221600000 | 0.05 | 0.50 | A>A |
| chr2 | 2.22E+08 | 221700000 | 0.07 | 0.51 | A>A |
| chr2 | 2.22E+08 | 221800000 | -0.03 | 0.55 | B>A |
| chr2 | 2.22E+08 | 221900000 | -0.12 | 0.57 | B>A |
| chr2 | 2.22E+08 | 222000000 | -0.20 | 0.42 | B>A |
| chr2 | 2.22E+08 | 222100000 | -0.29 | 0.49 | B>A |
| chr2 | 2.22E+08 | 222200000 | -0.41 | 0.52 | B>A |
| chr2 | 2.22E+08 | 222300000 | -0.44 | 0.48 | B>A |
| chr2 | 2.22E+08 | 222400000 | -0.37 | 0.48 | B>A |
| chr2 | 2.22E+08 | 222500000 | -0.34 | 0.16 | B>B |
| chr2 | 2.23E+08 | 222600000 | -0.49 | 0.02 | B>A |
| chr2 | 2.23E+08 | 222700000 | -0.53 | 0.04 | B>A |
| chr2 | 2.23E+08 | 222800000 | -0.59 | -0.01 | B>A |
| chr2 | 2.23E+08 | 222900000 | -0.44 | -0.02 | B>B |
| chr2 | 2.23E+08 | 223000000 | -0.45 | -0.03 | B>B |
| chr2 | 2.23E+08 | 223100000 | -0.59 | -0.14 | B>B |
| chr2 | 2.23E+08 | 223200000 | -0.31 | 0.06 | B>B |
| chr2 | 2.23E+08 | 223300000 | -0.35 | 0.41 | B>A |
| chr2 | 2.23E+08 | 223400000 | -0.27 | 0.36 | B>A |
| chr2 | 2.23E+08 | 223500000 | 0.02 | 0.31 | A>A |
| chr2 | 2.24E+08 | 223600000 | 0.51 | 0.56 | A>A |
| chr2 | 2.24E+08 | 223700000 | 0.86 | 1.08 | A>A |
| chr2 | 2.24E+08 | 223800000 | 0.73 | 0.91 | A>A |
| chr2 | 2.24E+08 | 223900000 | 0.44 | 0.70 | A>A |
| chr2 | 2.24E+08 | 224000000 | 0.55 | 0.85 | A>A |
| chr2 | 2.24E+08 | 224100000 | 0.56 | 0.77 | A>A |
| chr2 | 2.24E+08 | 224200000 | 0.57 | 0.70 | A>A |
| chr2 | 2.24E+08 | 224300000 | 0.54 | 0.84 | A>A |
| chr2 | 2.24E+08 | 224400000 | 0.70 | 0.89 | A>A |
| chr2 | 2.24E+08 | 224500000 | 0.68 | 0.89 | A>A |
| chr2 | 2.25E+08 | 224600000 | 0.67 | 0.88 | A>A |
| chr2 | 2.25E+08 | 224700000 | 0.69 | 0.87 | A>A |
| chr2 | 2.25E+08 | 224800000 | 0.83 | 0.97 | A>A |
| chr2 | 2.25E+08 | 224900000 | 0.66 | 0.71 | A>A |
| chr2 | 2.25E+08 | 225000000 | 0.86 | 0.90 | A>A |
| chr2 | 2.25E+08 | 225100000 | 0.93 | 0.99 | A>A |

| | | | | | |
|---|---|---|---|---|---|
| chr2 | 2.25E+08 | 225200000 | 0.55 | 0.63 | A>A |
| chr2 | 2.25E+08 | 225300000 | 0.53 | 0.49 | A>A |
| chr2 | 2.25E+08 | 225400000 | 0.86 | 0.87 | A>A |
| chr2 | 2.25E+08 | 225500000 | 0.86 | 0.83 | A>A |
| chr2 | 2.26E+08 | 225600000 | 0.57 | 0.72 | A>A |
| chr2 | 2.26E+08 | 225700000 | 0.53 | 0.54 | A>A |
| chr2 | 2.26E+08 | 225800000 | 0.59 | 0.57 | A>A |
| chr2 | 2.26E+08 | 225900000 | 0.52 | 0.53 | A>A |
| chr2 | 2.26E+08 | 226000000 | 0.50 | 0.39 | A>A |
| chr2 | 2.26E+08 | 226100000 | 0.42 | 0.33 | A>A |
| chr2 | 2.26E+08 | 226200000 | 0.29 | 0.25 | A>A |
| chr2 | 2.26E+08 | 226300000 | 0.35 | 0.25 | A>A |
| chr2 | 2.26E+08 | 226400000 | 0.39 | 0.22 | A>A |
| chr2 | 2.26E+08 | 226500000 | 0.28 | 0.15 | A>A |
| chr2 | 2.27E+08 | 226600000 | 0.19 | 0.19 | A>A |
| chr2 | 2.27E+08 | 226700000 | 0.51 | 0.40 | A>A |
| chr2 | 2.27E+08 | 226800000 | 0.52 | 0.47 | A>A |
| chr2 | 2.27E+08 | 226900000 | 0.58 | 0.63 | A>A |
| chr2 | 2.27E+08 | 227000000 | 0.62 | 0.62 | A>A |
| chr2 | 2.27E+08 | 227100000 | 0.53 | 0.59 | A>A |
| chr2 | 2.27E+08 | 227200000 | 0.59 | 0.74 | A>A |
| chr2 | 2.27E+08 | 227300000 | 0.57 | 0.68 | A>A |
| chr2 | 2.27E+08 | 227400000 | 0.58 | 0.82 | A>A |
| chr2 | 2.27E+08 | 227500000 | 0.69 | 0.72 | A>A |
| chr2 | 2.28E+08 | 227600000 | 0.63 | 0.69 | A>A |
| chr2 | 2.28E+08 | 227700000 | 0.42 | 0.44 | A>A |
| chr2 | 2.28E+08 | 227800000 | 0.52 | 0.48 | A>A |
| chr2 | 2.28E+08 | 227900000 | 0.35 | 0.31 | A>A |
| chr2 | 2.28E+08 | 228000000 | 0.32 | 0.38 | A>A |
| chr2 | 2.28E+08 | 228100000 | 0.69 | 0.55 | A>A |
| chr2 | 2.28E+08 | 228200000 | 0.64 | 0.54 | A>A |
| chr2 | 2.28E+08 | 228300000 | 0.58 | 0.47 | A>A |
| chr2 | 2.28E+08 | 228400000 | 0.70 | 0.49 | A>A |
| chr2 | 2.28E+08 | 228500000 | 0.61 | 0.44 | A>A |
| chr2 | 2.29E+08 | 228600000 | 0.40 | 0.27 | A>A |
| chr2 | 2.29E+08 | 228700000 | 0.29 | 0.13 | A>A |
| chr2 | 2.29E+08 | 228800000 | 0.04 | -0.31 | B>B |
| chr2 | 2.29E+08 | 228900000 | -0.26 | -0.74 | B>B |
| chr2 | 2.29E+08 | 229000000 | -0.38 | -0.84 | B>B |
| chr2 | 2.29E+08 | 229100000 | -0.52 | -0.90 | B>B |
| chr2 | 2.29E+08 | 229200000 | -0.58 | -0.85 | B>B |
| chr2 | 2.29E+08 | 229300000 | -0.66 | -0.81 | B>B |
| chr2 | 2.29E+08 | 229400000 | -0.64 | -0.84 | B>B |
| chr2 | 2.29E+08 | 229500000 | -0.59 | -0.83 | B>B |
| chr2 | 2.3E+08 | 229600000 | -0.47 | -0.75 | B>B |
| chr2 | 2.3E+08 | 229700000 | -0.69 | -0.84 | B>B |
| chr2 | 2.3E+08 | 229800000 | -0.73 | -0.93 | B>B |
| chr2 | 2.3E+08 | 229900000 | -0.59 | -0.80 | B>B |
| chr2 | 2.3E+08 | 230000000 | -0.45 | -0.76 | B>B |
| chr2 | 2.3E+08 | 230100000 | -0.28 | -0.75 | B>B |
| chr2 | 2.3E+08 | 230200000 | -0.12 | -0.52 | B>B |
| chr2 | 2.3E+08 | 230300000 | 0.05 | -0.25 | B>B |
| chr2 | 2.3E+08 | 230400000 | 0.08 | -0.46 | B>B |
| chr2 | 2.3E+08 | 230500000 | -0.05 | -0.57 | B>B |
| chr2 | 2.31E+08 | 230600000 | -0.01 | -0.55 | B>B |
| chr2 | 2.31E+08 | 230700000 | 0.38 | 0.44 | A>A |

| | | | | | |
|---|---|---|---|---|---|
| chr2 | 2.31E+08 | 230800000 | 0.70 | 0.85 | A>A |
| chr2 | 2.31E+08 | 230900000 | 0.38 | 0.73 | A>A |
| chr2 | 2.31E+08 | 231000000 | 0.22 | 0.80 | B>A |
| chr2 | 2.31E+08 | 231100000 | 0.10 | 0.68 | B>A |
| chr2 | 2.31E+08 | 231200000 | 0.01 | 0.70 | B>A |
| chr2 | 2.31E+08 | 231300000 | 0.30 | 0.84 | B>A |
| chr2 | 2.31E+08 | 231400000 | 0.21 | 0.82 | B>A |
| chr2 | 2.31E+08 | 231500000 | 0.24 | 0.76 | B>A |
| chr2 | 2.32E+08 | 231600000 | 0.71 | 1.03 | A>A |
| chr2 | 2.32E+08 | 231700000 | 0.77 | 1.04 | A>A |
| chr2 | 2.32E+08 | 231800000 | 0.95 | 1.19 | A>A |
| chr2 | 2.32E+08 | 231900000 | 1.12 | 1.31 | A>A |
| chr2 | 2.32E+08 | 232000000 | 1.13 | 1.34 | A>A |
| chr2 | 2.32E+08 | 232100000 | 1.23 | 1.34 | A>A |
| chr2 | 2.32E+08 | 232200000 | 1.34 | 1.46 | A>A |
| chr2 | 2.32E+08 | 232300000 | 1.30 | 1.46 | A>A |
| chr2 | 2.32E+08 | 232400000 | 1.32 | 1.27 | A>A |
| chr2 | 2.32E+08 | 232500000 | 1.41 | 1.39 | A>A |
| chr2 | 2.33E+08 | 232600000 | 1.45 | 1.52 | A>A |
| chr2 | 2.33E+08 | 232700000 | 1.30 | 1.32 | A>A |
| chr2 | 2.33E+08 | 232800000 | 1.05 | 1.04 | A>A |
| chr2 | 2.33E+08 | 232900000 | 1.30 | 1.41 | A>A |
| chr2 | 2.33E+08 | 233000000 | 1.30 | 1.40 | A>A |
| chr2 | 2.33E+08 | 233100000 | 1.26 | 1.37 | A>A |
| chr2 | 2.33E+08 | 233200000 | 1.21 | 1.25 | A>A |
| chr2 | 2.33E+08 | 233300000 | 1.30 | 1.24 | A>A |
| chr2 | 2.33E+08 | 233400000 | 1.17 | 1.15 | A>A |
| chr2 | 2.33E+08 | 233500000 | 1.28 | 1.21 | A>A |
| chr2 | 2.34E+08 | 233600000 | 1.36 | 1.43 | A>A |
| chr2 | 2.34E+08 | 233700000 | 1.19 | 1.18 | A>A |
| chr2 | 2.34E+08 | 233800000 | 1.05 | 1.15 | A>A |
| chr2 | 2.34E+08 | 233900000 | 1.32 | 1.21 | A>A |
| chr2 | 2.34E+08 | 234000000 | 1.05 | 0.90 | A>A |
| chr2 | 2.34E+08 | 234100000 | 0.00 | 0.00 | NO |
| chr2 | 2.34E+08 | 234200000 | 0.86 | 0.85 | A>A |
| chr2 | 2.34E+08 | 234300000 | 1.04 | 1.01 | A>A |
| chr2 | 2.34E+08 | 234400000 | 0.74 | 0.84 | A>A |
| chr2 | 2.34E+08 | 234500000 | 1.07 | 1.14 | A>A |
| chr2 | 2.35E+08 | 234600000 | 1.07 | 1.18 | A>A |
| chr2 | 2.35E+08 | 234700000 | 0.91 | 1.00 | A>A |
| chr2 | 2.35E+08 | 234800000 | 0.64 | 0.66 | A>A |
| chr2 | 2.35E+08 | 234900000 | 0.73 | 0.74 | A>A |
| chr2 | 2.35E+08 | 235000000 | 0.95 | 1.01 | A>A |
| chr2 | 2.35E+08 | 235100000 | 0.87 | 0.85 | A>A |
| chr2 | 2.35E+08 | 235200000 | 0.99 | 0.94 | A>A |
| chr2 | 2.35E+08 | 235300000 | 1.02 | 0.97 | A>A |
| chr2 | 2.35E+08 | 235400000 | 1.05 | 0.99 | A>A |
| chr2 | 2.35E+08 | 235500000 | 0.41 | 0.52 | A>A |
| chr2 | 2.36E+08 | 235600000 | 0.91 | 0.92 | A>A |
| chr2 | 2.36E+08 | 235700000 | 1.10 | 1.03 | A>A |
| chr2 | 2.36E+08 | 235800000 | 0.94 | 0.97 | A>A |
| chr2 | 2.36E+08 | 235900000 | 1.00 | 0.94 | A>A |
| chr2 | 2.36E+08 | 236000000 | 0.90 | 0.90 | A>A |
| chr2 | 2.36E+08 | 236100000 | 0.60 | 0.56 | A>A |
| chr2 | 2.36E+08 | 236200000 | 1.02 | 0.86 | A>A |
| chr2 | 2.36E+08 | 236300000 | 1.01 | 0.91 | A>A |

| | | | | | |
|---|---|---|---|---|---|
| chr2 | 2.36E+08 | 236400000 | 1.13 | 1.00 | A>A |
| chr2 | 2.36E+08 | 236500000 | 0.96 | 0.97 | A>A |
| chr2 | 2.37E+08 | 236600000 | 1.03 | 0.95 | A>A |
| chr2 | 2.37E+08 | 236700000 | 0.91 | 0.78 | A>A |
| chr2 | 2.37E+08 | 236800000 | 0.92 | 0.76 | A>A |
| chr2 | 2.37E+08 | 236900000 | 0.88 | 0.68 | A>A |
| chr2 | 2.37E+08 | 237000000 | 1.09 | 0.78 | A>A |
| chr2 | 2.37E+08 | 237100000 | 0.93 | 0.61 | A>A |
| chr2 | 2.37E+08 | 237200000 | 0.67 | 0.28 | A>A |
| chr2 | 2.37E+08 | 237300000 | 0.39 | -0.02 | A>A |
| chr2 | 2.37E+08 | 237400000 | 0.46 | -0.20 | A>B |
| chr2 | 2.37E+08 | 237500000 | 0.46 | -0.21 | A>B |
| chr2 | 2.38E+08 | 237600000 | 0.47 | -0.18 | A>B |
| chr2 | 2.38E+08 | 237700000 | 0.63 | -0.16 | A>B |
| chr2 | 2.38E+08 | 237800000 | 0.51 | -0.13 | A>B |
| chr2 | 2.38E+08 | 237900000 | 0.52 | -0.13 | A>B |
| chr2 | 2.38E+08 | 238000000 | 0.36 | 0.02 | A>A |
| chr2 | 2.38E+08 | 238100000 | 0.76 | 0.56 | A>A |
| chr2 | 2.38E+08 | 238200000 | 1.14 | 0.82 | A>A |
| chr2 | 2.38E+08 | 238300000 | 1.13 | 0.79 | A>A |
| chr2 | 2.38E+08 | 238400000 | 1.04 | 0.86 | A>A |
| chr2 | 2.38E+08 | 238500000 | 1.16 | 1.00 | A>A |
| chr2 | 2.39E+08 | 238600000 | 1.10 | 0.96 | A>A |
| chr2 | 2.39E+08 | 238700000 | 1.23 | 1.15 | A>A |
| chr2 | 2.39E+08 | 238800000 | 1.04 | 1.14 | A>A |
| chr2 | 2.39E+08 | 238900000 | 1.20 | 1.22 | A>A |
| chr2 | 2.39E+08 | 239000000 | 1.23 | 1.22 | A>A |
| chr2 | 2.39E+08 | 239100000 | 1.22 | 1.18 | A>A |
| chr2 | 2.39E+08 | 239200000 | 1.11 | 1.19 | A>A |
| chr2 | 2.39E+08 | 239300000 | 0.99 | 1.09 | A>A |
| chr2 | 2.39E+08 | 239400000 | 0.95 | 1.00 | A>A |
| chr2 | 2.39E+08 | 239500000 | 0.92 | 1.12 | A>A |
| chr2 | 2.4E+08 | 239600000 | 1.02 | 1.27 | A>A |
| chr2 | 2.4E+08 | 239700000 | 0.76 | 0.88 | A>A |
| chr2 | 2.4E+08 | 239800000 | 0.96 | 1.12 | A>A |
| chr2 | 2.4E+08 | 239900000 | 0.86 | 0.93 | A>A |
| chr2 | 2.4E+08 | 240000000 | 0.82 | 0.96 | A>A |
| chr2 | 2.4E+08 | 240100000 | 0.78 | 0.89 | A>A |
| chr2 | 2.4E+08 | 240200000 | 1.16 | 1.20 | A>A |
| chr2 | 2.4E+08 | 240300000 | 1.13 | 1.06 | A>A |
| chr2 | 2.4E+08 | 240400000 | 0.83 | 0.91 | A>A |
| chr2 | 2.4E+08 | 240500000 | 0.84 | 1.02 | A>A |
| chr2 | 2.41E+08 | 240600000 | 0.82 | 1.01 | A>A |
| chr2 | 2.41E+08 | 240700000 | 0.79 | 1.01 | A>A |
| chr2 | 2.41E+08 | 240800000 | 0.87 | 0.91 | A>A |
| chr2 | 2.41E+08 | 240900000 | 0.90 | 1.00 | A>A |
| chr2 | 2.41E+08 | 241000000 | 0.81 | 0.96 | A>A |
| chr2 | 2.41E+08 | 241100000 | 0.79 | 0.91 | A>A |
| chr2 | 2.41E+08 | 241200000 | 0.84 | 1.05 | A>A |
| chr2 | 2.41E+08 | 241300000 | 1.10 | 1.04 | A>A |
| chr2 | 2.41E+08 | 241400000 | 1.13 | 1.08 | A>A |
| chr2 | 2.41E+08 | 241500000 | 1.05 | 1.02 | A>A |
| chr2 | 2.42E+08 | 241600000 | 1.01 | 1.00 | A>A |
| chr2 | 2.42E+08 | 241700000 | 0.99 | 0.95 | A>A |
| chr2 | 2.42E+08 | 241800000 | 0.83 | 0.93 | A>A |
| chr2 | 2.42E+08 | 241900000 | 0.85 | 0.99 | A>A |

| | | | | | |
|---|---|---|---|---|---|
| chr2 | 2.42E+08 | 242000000 | 0.98 | 1.08 | A>A |
| chr2 | 2.42E+08 | 242100000 | 1.10 | 0.97 | A>A |
| chr2 | 2.42E+08 | 242200000 | 1.02 | 0.89 | A>A |
| chr2 | 2.42E+08 | 242300000 | 1.21 | 1.22 | A>A |
| chr2 | 2.42E+08 | 242400000 | 1.10 | 1.04 | A>A |
| chr2 | 2.42E+08 | 242500000 | 1.07 | 1.03 | A>A |
| chr2 | 2.43E+08 | 242600000 | 1.13 | 1.04 | A>A |
| chr2 | 2.43E+08 | 242700000 | 1.04 | 1.03 | A>A |
| chr2 | 2.43E+08 | 242800000 | 1.16 | 1.05 | A>A |
| chr2 | 2.43E+08 | 242900000 | 1.02 | 1.04 | A>A |
| chr2 | 2.43E+08 | 243000000 | 1.06 | 1.14 | A>A |
| chr2 | 2.43E+08 | 243100000 | 0.00 | 0.00 | NO |
| chr2 | 2.43E+08 | 243199373 | 0.00 | 0.00 | NO |
| chr3 | 0 | 100000 | 0.00 | 0.00 | NO |
| chr3 | 100000 | 200000 | -0.97 | -1.45 | B>B |
| chr3 | 200000 | 300000 | -0.86 | -1.30 | B>B |
| chr3 | 300000 | 400000 | -0.80 | -1.18 | B>B |
| chr3 | 400000 | 500000 | -0.85 | -1.19 | B>B |
| chr3 | 500000 | 600000 | -1.04 | -1.21 | B>B |
| chr3 | 600000 | 700000 | -0.92 | -1.13 | B>B |
| chr3 | 700000 | 800000 | -0.96 | -1.27 | B>B |
| chr3 | 800000 | 900000 | -0.89 | -1.26 | B>B |
| chr3 | 900000 | 1000000 | -0.82 | -1.18 | B>B |
| chr3 | 1000000 | 1100000 | -0.85 | -1.10 | B>B |
| chr3 | 1100000 | 1200000 | -0.80 | -1.19 | B>B |
| chr3 | 1200000 | 1300000 | -0.83 | -1.15 | B>B |
| chr3 | 1300000 | 1400000 | -0.79 | -1.20 | B>B |
| chr3 | 1400000 | 1500000 | -0.83 | -1.14 | B>B |
| chr3 | 1500000 | 1600000 | -0.81 | -1.16 | B>B |
| chr3 | 1600000 | 1700000 | -0.78 | -1.15 | B>B |
| chr3 | 1700000 | 1800000 | -0.78 | -1.13 | B>B |
| chr3 | 1800000 | 1900000 | -0.72 | -1.13 | B>B |
| chr3 | 1900000 | 2000000 | -0.70 | -1.12 | B>B |
| chr3 | 2000000 | 2100000 | -0.71 | -1.11 | B>B |
| chr3 | 2100000 | 2200000 | -0.67 | -1.10 | B>B |
| chr3 | 2200000 | 2300000 | -0.78 | -1.09 | B>B |
| chr3 | 2300000 | 2400000 | -0.74 | -1.08 | B>B |
| chr3 | 2400000 | 2500000 | -0.75 | -1.10 | B>B |
| chr3 | 2500000 | 2600000 | -0.78 | -1.08 | B>B |
| chr3 | 2600000 | 2700000 | -0.68 | -1.06 | B>B |
| chr3 | 2700000 | 2800000 | -0.68 | -1.04 | B>B |
| chr3 | 2800000 | 2900000 | -0.53 | -1.05 | B>B |
| chr3 | 2900000 | 3000000 | -0.60 | -1.06 | B>B |
| chr3 | 3000000 | 3100000 | -0.34 | -1.04 | A>B |
| chr3 | 3100000 | 3200000 | 0.13 | -0.57 | A>B |
| chr3 | 3200000 | 3300000 | -0.33 | -0.64 | B>B |
| chr3 | 3300000 | 3400000 | -0.67 | -1.01 | B>B |
| chr3 | 3400000 | 3500000 | -0.72 | -1.04 | B>B |
| chr3 | 3500000 | 3600000 | -0.80 | -1.08 | B>B |
| chr3 | 3600000 | 3700000 | -0.66 | -0.99 | B>B |
| chr3 | 3700000 | 3800000 | -0.66 | -0.96 | B>B |
| chr3 | 3800000 | 3900000 | -0.68 | -1.03 | B>B |
| chr3 | 3900000 | 4000000 | -0.71 | -1.03 | B>B |
| chr3 | 4000000 | 4100000 | -0.66 | -1.01 | B>B |
| chr3 | 4100000 | 4200000 | -0.65 | -1.04 | B>B |
| chr3 | 4200000 | 4300000 | -0.61 | -0.93 | B>B |

| | | | | | |
|---|---|---|---|---|---|
| chr3 | 4300000 | 4400000 | -0.06 | -0.39 | B>B |
| chr3 | 4400000 | 4500000 | 0.42 | 0.04 | A>A |
| chr3 | 4500000 | 4600000 | 0.48 | -0.04 | A>A |
| chr3 | 4600000 | 4700000 | 0.30 | -0.10 | A>A |
| chr3 | 4700000 | 4800000 | 0.66 | 0.50 | A>A |
| chr3 | 4800000 | 4900000 | 1.23 | 1.04 | A>A |
| chr3 | 4900000 | 5000000 | 1.17 | 1.08 | A>A |
| chr3 | 5000000 | 5100000 | 1.11 | 0.81 | A>A |
| chr3 | 5100000 | 5200000 | 0.68 | 0.46 | A>A |
| chr3 | 5200000 | 5300000 | 0.37 | 0.24 | A>A |
| chr3 | 5300000 | 5400000 | 0.14 | 0.13 | A>A |
| chr3 | 5400000 | 5500000 | 0.15 | 0.18 | A>A |
| chr3 | 5500000 | 5600000 | 0.14 | 0.11 | A>A |
| chr3 | 5600000 | 5700000 | -0.13 | 0.02 | B>B |
| chr3 | 5700000 | 5800000 | -0.15 | -0.02 | B>B |
| chr3 | 5800000 | 5900000 | -0.25 | -0.14 | B>B |
| chr3 | 5900000 | 6000000 | -0.36 | -0.09 | B>B |
| chr3 | 6000000 | 6100000 | -0.53 | -0.09 | B>B |
| chr3 | 6100000 | 6200000 | -0.54 | -0.24 | B>B |
| chr3 | 6200000 | 6300000 | -0.55 | -0.43 | B>B |
| chr3 | 6300000 | 6400000 | -0.64 | -0.55 | B>B |
| chr3 | 6400000 | 6500000 | -0.68 | -0.73 | B>B |
| chr3 | 6500000 | 6600000 | -0.75 | -0.75 | B>B |
| chr3 | 6600000 | 6700000 | -0.73 | -0.76 | B>B |
| chr3 | 6700000 | 6800000 | -0.66 | -0.76 | B>B |
| chr3 | 6800000 | 6900000 | -0.60 | -0.62 | B>B |
| chr3 | 6900000 | 7000000 | -0.70 | -0.76 | B>B |
| chr3 | 7000000 | 7100000 | -0.75 | -0.76 | B>B |
| chr3 | 7100000 | 7200000 | -0.74 | -0.71 | B>B |
| chr3 | 7200000 | 7300000 | -0.72 | -0.49 | B>B |
| chr3 | 7300000 | 7400000 | -0.63 | -0.25 | B>B |
| chr3 | 7400000 | 7500000 | -0.51 | -0.13 | B>B |
| chr3 | 7500000 | 7600000 | -0.36 | -0.08 | B>B |
| chr3 | 7600000 | 7700000 | -0.52 | -0.33 | B>B |
| chr3 | 7700000 | 7800000 | -0.56 | -0.48 | B>B |
| chr3 | 7800000 | 7900000 | -0.56 | -0.61 | B>B |
| chr3 | 7900000 | 8000000 | -0.64 | -0.78 | B>B |
| chr3 | 8000000 | 8100000 | -0.48 | -0.79 | B>B |
| chr3 | 8100000 | 8200000 | -0.39 | -0.67 | B>B |
| chr3 | 8200000 | 8300000 | -0.28 | -0.69 | B>B |
| chr3 | 8300000 | 8400000 | -0.12 | -0.45 | B>B |
| chr3 | 8400000 | 8500000 | 0.21 | -0.04 | A>A |
| chr3 | 8500000 | 8600000 | 0.63 | 0.47 | A>A |
| chr3 | 8600000 | 8700000 | 0.48 | 0.36 | A>A |
| chr3 | 8700000 | 8800000 | 0.81 | 0.78 | A>A |
| chr3 | 8800000 | 8900000 | 0.81 | 0.87 | A>A |
| chr3 | 8900000 | 9000000 | 0.94 | 0.89 | A>A |
| chr3 | 9000000 | 9100000 | 0.93 | 0.85 | A>A |
| chr3 | 9100000 | 9200000 | 1.12 | 1.07 | A>A |
| chr3 | 9200000 | 9300000 | 1.11 | 1.06 | A>A |
| chr3 | 9300000 | 9400000 | 0.88 | 0.85 | A>A |
| chr3 | 9400000 | 9500000 | 1.25 | 1.12 | A>A |
| chr3 | 9500000 | 9600000 | 1.04 | 1.02 | A>A |
| chr3 | 9600000 | 9700000 | 1.23 | 1.09 | A>A |
| chr3 | 9700000 | 9800000 | 1.13 | 1.06 | A>A |
| chr3 | 9800000 | 9900000 | 1.20 | 1.07 | A>A |

| | | | | | |
|---|---|---|---|---|---|
| chr3 | 9900000 | 10000000 | 1.27 | 1.24 | A>A |
| chr3 | 10000000 | 10100000 | 1.22 | 1.08 | A>A |
| chr3 | 10100000 | 10200000 | 1.15 | 1.08 | A>A |
| chr3 | 10200000 | 10300000 | 1.19 | 1.15 | A>A |
| chr3 | 10300000 | 10400000 | 0.91 | 0.80 | A>A |
| chr3 | 10400000 | 10500000 | 0.79 | 0.63 | A>A |
| chr3 | 10500000 | 10600000 | 0.83 | 0.58 | A>A |
| chr3 | 10600000 | 10700000 | 0.70 | 0.49 | A>A |
| chr3 | 10700000 | 10800000 | 0.60 | 0.47 | A>A |
| chr3 | 10800000 | 10900000 | 0.66 | 0.45 | A>A |
| chr3 | 10900000 | 11000000 | 0.63 | 0.47 | A>A |
| chr3 | 11000000 | 11100000 | 0.77 | 0.64 | A>A |
| chr3 | 11100000 | 11200000 | 0.88 | 0.78 | A>A |
| chr3 | 11200000 | 11300000 | 1.00 | 0.79 | A>A |
| chr3 | 11300000 | 11400000 | 0.96 | 0.87 | A>A |
| chr3 | 11400000 | 11500000 | 0.79 | 0.78 | A>A |
| chr3 | 11500000 | 11600000 | 0.83 | 0.85 | A>A |
| chr3 | 11600000 | 11700000 | 0.80 | 0.86 | A>A |
| chr3 | 11700000 | 11800000 | 0.93 | 0.90 | A>A |
| chr3 | 11800000 | 11900000 | 0.73 | 0.68 | A>A |
| chr3 | 11900000 | 12000000 | 0.59 | 0.50 | A>A |
| chr3 | 12000000 | 12100000 | 0.48 | 0.47 | A>A |
| chr3 | 12100000 | 12200000 | 0.48 | 0.48 | A>A |
| chr3 | 12200000 | 12300000 | 0.81 | 0.84 | A>A |
| chr3 | 12300000 | 12400000 | 1.05 | 0.95 | A>A |
| chr3 | 12400000 | 12500000 | 0.97 | 0.88 | A>A |
| chr3 | 12500000 | 12600000 | 0.78 | 0.82 | A>A |
| chr3 | 12600000 | 12700000 | 1.16 | 1.10 | A>A |
| chr3 | 12700000 | 12800000 | 1.25 | 1.13 | A>A |
| chr3 | 12800000 | 12900000 | 1.10 | 1.15 | A>A |
| chr3 | 12900000 | 13000000 | 1.01 | 1.07 | A>A |
| chr3 | 13000000 | 13100000 | 1.06 | 1.05 | A>A |
| chr3 | 13100000 | 13200000 | 0.89 | 0.97 | A>A |
| chr3 | 13200000 | 13300000 | 0.97 | 1.08 | A>A |
| chr3 | 13300000 | 13400000 | 1.00 | 1.06 | A>A |
| chr3 | 13400000 | 13500000 | 1.08 | 1.02 | A>A |
| chr3 | 13500000 | 13600000 | 0.81 | 1.00 | A>A |
| chr3 | 13600000 | 13700000 | 0.71 | 1.02 | A>A |
| chr3 | 13700000 | 13800000 | 0.44 | 0.93 | A>A |
| chr3 | 13800000 | 13900000 | 0.44 | 0.94 | A>A |
| chr3 | 13900000 | 14000000 | 0.25 | 0.82 | B>A |
| chr3 | 14000000 | 14100000 | 0.28 | 1.06 | B>A |
| chr3 | 14100000 | 14200000 | 0.67 | 1.06 | A>A |
| chr3 | 14200000 | 14300000 | 0.75 | 0.92 | A>A |
| chr3 | 14300000 | 14400000 | 1.05 | 1.17 | A>A |
| chr3 | 14400000 | 14500000 | 1.13 | 1.26 | A>A |
| chr3 | 14500000 | 14600000 | 0.79 | 1.04 | A>A |
| chr3 | 14600000 | 14700000 | 0.96 | 1.04 | A>A |
| chr3 | 14700000 | 14800000 | 0.97 | 0.97 | A>A |
| chr3 | 14800000 | 14900000 | 1.02 | 1.09 | A>A |
| chr3 | 14900000 | 15000000 | 0.94 | 1.06 | A>A |
| chr3 | 15000000 | 15100000 | 1.20 | 1.17 | A>A |
| chr3 | 15100000 | 15200000 | 1.14 | 1.03 | A>A |
| chr3 | 15200000 | 15300000 | 1.19 | 1.14 | A>A |
| chr3 | 15300000 | 15400000 | 1.31 | 1.23 | A>A |
| chr3 | 15400000 | 15500000 | 1.14 | 1.14 | A>A |

| | | | | | |
|---|---|---|---|---|---|
| chr3 | 15500000 | 15600000 | 1.13 | 1.12 | A>A |
| chr3 | 15600000 | 15700000 | 1.05 | 1.04 | A>A |
| chr3 | 15700000 | 15800000 | 1.07 | 1.05 | A>A |
| chr3 | 15800000 | 15900000 | 0.96 | 0.91 | A>A |
| chr3 | 15900000 | 16000000 | 0.99 | 0.86 | A>A |
| chr3 | 16000000 | 16100000 | 0.93 | 0.73 | A>A |
| chr3 | 16100000 | 16200000 | 1.29 | 1.01 | A>A |
| chr3 | 16200000 | 16300000 | 0.83 | 0.71 | A>A |
| chr3 | 16300000 | 16400000 | 0.66 | 0.69 | A>A |
| chr3 | 16400000 | 16500000 | 0.87 | 0.82 | A>A |
| chr3 | 16500000 | 16600000 | 0.74 | 0.74 | A>A |
| chr3 | 16600000 | 16700000 | 0.34 | 0.46 | A>A |
| chr3 | 16700000 | 16800000 | 0.39 | 0.58 | A>A |
| chr3 | 16800000 | 16900000 | 0.47 | 0.65 | A>A |
| chr3 | 16900000 | 17000000 | 0.69 | 0.78 | A>A |
| chr3 | 17000000 | 17100000 | 0.56 | 0.76 | A>A |
| chr3 | 17100000 | 17200000 | 0.47 | 0.62 | A>A |
| chr3 | 17200000 | 17300000 | 0.31 | 0.49 | A>A |
| chr3 | 17300000 | 17400000 | 0.36 | 0.39 | A>A |
| chr3 | 17400000 | 17500000 | 0.29 | 0.41 | A>A |
| chr3 | 17500000 | 17600000 | 0.36 | 0.43 | A>A |
| chr3 | 17600000 | 17700000 | 0.61 | 0.53 | A>A |
| chr3 | 17700000 | 17800000 | 0.54 | 0.53 | A>A |
| chr3 | 17800000 | 17900000 | 0.20 | 0.22 | A>A |
| chr3 | 17900000 | 18000000 | -0.20 | 0.03 | B>B |
| chr3 | 18000000 | 18100000 | -0.24 | 0.06 | B>B |
| chr3 | 18100000 | 18200000 | -0.19 | 0.17 | B>B |
| chr3 | 18200000 | 18300000 | -0.41 | 0.03 | B>B |
| chr3 | 18300000 | 18400000 | -0.47 | -0.07 | B>B |
| chr3 | 18400000 | 18500000 | -0.44 | -0.27 | B>B |
| chr3 | 18500000 | 18600000 | -0.75 | -1.04 | B>B |
| chr3 | 18600000 | 18700000 | -0.90 | -1.19 | B>B |
| chr3 | 18700000 | 18800000 | -0.89 | -1.17 | B>B |
| chr3 | 18800000 | 18900000 | -0.76 | -1.09 | B>B |
| chr3 | 18900000 | 19000000 | -0.95 | -1.09 | B>B |
| chr3 | 19000000 | 19100000 | -0.78 | -0.98 | B>B |
| chr3 | 19100000 | 19200000 | -0.66 | -0.92 | B>B |
| chr3 | 19200000 | 19300000 | -0.75 | -1.11 | B>B |
| chr3 | 19300000 | 19400000 | -0.70 | -1.03 | B>B |
| chr3 | 19400000 | 19500000 | -0.65 | -1.07 | B>B |
| chr3 | 19500000 | 19600000 | -0.53 | -1.05 | B>B |
| chr3 | 19600000 | 19700000 | -0.59 | -1.01 | B>B |
| chr3 | 19700000 | 19800000 | -0.45 | -1.00 | B>B |
| chr3 | 19800000 | 19900000 | -0.40 | -0.84 | B>B |
| chr3 | 19900000 | 20000000 | 0.30 | -0.11 | A>A |
| chr3 | 20000000 | 20100000 | 0.58 | 0.18 | A>A |
| chr3 | 20100000 | 20200000 | 0.44 | 0.10 | A>A |
| chr3 | 20200000 | 20300000 | 0.02 | -0.45 | B>B |
| chr3 | 20300000 | 20400000 | -0.42 | -0.87 | B>B |
| chr3 | 20400000 | 20500000 | -0.61 | -0.94 | B>B |
| chr3 | 20500000 | 20600000 | -0.68 | -0.95 | B>B |
| chr3 | 20600000 | 20700000 | -0.78 | -1.01 | B>B |
| chr3 | 20700000 | 20800000 | -0.86 | -1.01 | B>B |
| chr3 | 20800000 | 20900000 | -0.88 | -1.02 | B>B |
| chr3 | 20900000 | 21000000 | -0.86 | -0.97 | B>B |
| chr3 | 21000000 | 21100000 | -0.91 | -1.00 | B>B |

| | | | | | |
|---|---|---|---|---|---|
| chr3 | 21100000 | 21200000 | -0.91 | -0.93 | B>B |
| chr3 | 21200000 | 21300000 | -0.79 | -0.92 | B>B |
| chr3 | 21300000 | 21400000 | -0.83 | -0.69 | B>B |
| chr3 | 21400000 | 21500000 | -0.74 | -0.53 | B>B |
| chr3 | 21500000 | 21600000 | -0.80 | -0.84 | B>B |
| chr3 | 21600000 | 21700000 | -0.79 | -1.04 | B>B |
| chr3 | 21700000 | 21800000 | -0.78 | -1.07 | B>B |
| chr3 | 21800000 | 21900000 | -0.86 | -1.01 | B>B |
| chr3 | 21900000 | 22000000 | -0.84 | -0.87 | B>B |
| chr3 | 22000000 | 22100000 | -0.83 | -0.86 | B>B |
| chr3 | 22100000 | 22200000 | -0.68 | -0.92 | B>B |
| chr3 | 22200000 | 22300000 | -0.73 | -1.13 | B>B |
| chr3 | 22300000 | 22400000 | -0.78 | -1.21 | B>B |
| chr3 | 22400000 | 22500000 | -0.95 | -1.26 | B>B |
| chr3 | 22500000 | 22600000 | -0.80 | -1.16 | B>B |
| chr3 | 22600000 | 22700000 | -0.76 | -1.13 | B>B |
| chr3 | 22700000 | 22800000 | -0.72 | -1.06 | B>B |
| chr3 | 22800000 | 22900000 | -0.64 | -1.03 | B>B |
| chr3 | 22900000 | 23000000 | -0.45 | -0.84 | B>B |
| chr3 | 23000000 | 23100000 | -0.20 | -0.56 | B>B |
| chr3 | 23100000 | 23200000 | 0.14 | -0.22 | B>B |
| chr3 | 23200000 | 23300000 | 0.36 | 0.10 | A>A |
| chr3 | 23300000 | 23400000 | 0.48 | 0.13 | A>A |
| chr3 | 23400000 | 23500000 | 0.47 | 0.15 | A>A |
| chr3 | 23500000 | 23600000 | 0.57 | 0.28 | A>A |
| chr3 | 23600000 | 23700000 | 0.60 | 0.43 | A>A |
| chr3 | 23700000 | 23800000 | 0.82 | 0.47 | A>A |
| chr3 | 23800000 | 23900000 | 0.64 | 0.42 | A>A |
| chr3 | 23900000 | 24000000 | 0.69 | 0.45 | A>A |
| chr3 | 24000000 | 24100000 | 0.48 | 0.41 | A>A |
| chr3 | 24100000 | 24200000 | 0.68 | 0.59 | A>A |
| chr3 | 24200000 | 24300000 | 0.48 | 0.51 | A>A |
| chr3 | 24300000 | 24400000 | 0.52 | 0.69 | A>A |
| chr3 | 24400000 | 24500000 | 0.59 | 0.70 | A>A |
| chr3 | 24500000 | 24600000 | 0.50 | 0.63 | A>A |
| chr3 | 24600000 | 24700000 | 0.34 | 0.53 | A>A |
| chr3 | 24700000 | 24800000 | -0.10 | -0.09 | B>B |
| chr3 | 24800000 | 24900000 | -0.45 | -0.66 | B>B |
| chr3 | 24900000 | 25000000 | -0.56 | -0.95 | B>B |
| chr3 | 25000000 | 25100000 | -0.33 | -0.91 | A>B |
| chr3 | 25100000 | 25200000 | -0.29 | -0.96 | A>B |
| chr3 | 25200000 | 25300000 | -0.21 | -0.90 | A>B |
| chr3 | 25300000 | 25400000 | 0.07 | -0.70 | A>B |
| chr3 | 25400000 | 25500000 | 0.37 | -0.32 | A>B |
| chr3 | 25500000 | 25600000 | 0.32 | -0.18 | A>A |
| chr3 | 25600000 | 25700000 | 0.08 | -0.15 | B>B |
| chr3 | 25700000 | 25800000 | 0.29 | -0.13 | A>A |
| chr3 | 25800000 | 25900000 | 0.03 | -0.44 | B>B |
| chr3 | 25900000 | 26000000 | -0.51 | -1.01 | B>B |
| chr3 | 26000000 | 26100000 | -0.83 | -1.19 | B>B |
| chr3 | 26100000 | 26200000 | -0.92 | -1.16 | B>B |
| chr3 | 26200000 | 26300000 | -1.01 | -1.20 | B>B |
| chr3 | 26300000 | 26400000 | -0.96 | -1.18 | B>B |
| chr3 | 26400000 | 26500000 | -0.87 | -1.05 | B>B |
| chr3 | 26500000 | 26600000 | -0.93 | -1.16 | B>B |
| chr3 | 26600000 | 26700000 | -0.82 | -1.16 | B>B |

| | | | | | |
|---|---|---|---|---|---|
| chr3 | 26700000 | 26800000 | -0.79 | -1.20 | B>B |
| chr3 | 26800000 | 26900000 | -0.91 | -1.24 | B>B |
| chr3 | 26900000 | 27000000 | -0.83 | -1.25 | B>B |
| chr3 | 27000000 | 27100000 | -0.57 | -1.17 | A>B |
| chr3 | 27100000 | 27200000 | -0.18 | -0.67 | B>B |
| chr3 | 27200000 | 27300000 | 0.02 | 0.02 | A>A |
| chr3 | 27300000 | 27400000 | 0.29 | 0.10 | A>A |
| chr3 | 27400000 | 27500000 | 0.81 | 0.58 | A>A |
| chr3 | 27500000 | 27600000 | 0.89 | 0.58 | A>A |
| chr3 | 27600000 | 27700000 | 0.94 | 0.63 | A>A |
| chr3 | 27700000 | 27800000 | 0.28 | 0.04 | A>A |
| chr3 | 27800000 | 27900000 | -0.07 | -0.49 | B>B |
| chr3 | 27900000 | 28000000 | -0.20 | -0.44 | B>B |
| chr3 | 28000000 | 28100000 | -0.05 | -0.40 | B>B |
| chr3 | 28100000 | 28200000 | -0.14 | -0.33 | B>B |
| chr3 | 28200000 | 28300000 | -0.03 | -0.16 | B>B |
| chr3 | 28300000 | 28400000 | 0.02 | -0.10 | B>B |
| chr3 | 28400000 | 28500000 | -0.71 | -1.14 | B>B |
| chr3 | 28500000 | 28600000 | -0.64 | -1.23 | B>B |
| chr3 | 28600000 | 28700000 | -0.53 | -1.30 | A>B |
| chr3 | 28700000 | 28800000 | -0.50 | -1.21 | A>B |
| chr3 | 28800000 | 28900000 | -0.45 | -1.04 | A>B |
| chr3 | 28900000 | 29000000 | -0.33 | -0.61 | B>B |
| chr3 | 29000000 | 29100000 | -0.31 | -0.43 | B>B |
| chr3 | 29100000 | 29200000 | -0.12 | -0.20 | B>B |
| chr3 | 29200000 | 29300000 | 0.06 | -0.09 | B>B |
| chr3 | 29300000 | 29400000 | 0.20 | 0.06 | A>A |
| chr3 | 29400000 | 29500000 | 0.23 | 0.01 | A>A |
| chr3 | 29500000 | 29600000 | 0.28 | 0.08 | A>A |
| chr3 | 29600000 | 29700000 | 0.33 | 0.06 | A>A |
| chr3 | 29700000 | 29800000 | 0.25 | 0.00 | A>A |
| chr3 | 29800000 | 29900000 | 0.10 | -0.08 | A>A |
| chr3 | 29900000 | 30000000 | 0.09 | -0.12 | B>B |
| chr3 | 30000000 | 30100000 | 0.21 | 0.09 | A>A |
| chr3 | 30100000 | 30200000 | 0.62 | 0.53 | A>A |
| chr3 | 30200000 | 30300000 | 0.85 | 0.67 | A>A |
| chr3 | 30300000 | 30400000 | 0.91 | 0.84 | A>A |
| chr3 | 30400000 | 30500000 | 0.86 | 0.84 | A>A |
| chr3 | 30500000 | 30600000 | 0.87 | 0.82 | A>A |
| chr3 | 30600000 | 30700000 | 0.88 | 0.74 | A>A |
| chr3 | 30700000 | 30800000 | 0.50 | 0.53 | A>A |
| chr3 | 30800000 | 30900000 | 0.37 | 0.48 | A>A |
| chr3 | 30900000 | 31000000 | 0.35 | 0.51 | A>A |
| chr3 | 31000000 | 31100000 | 0.24 | 0.62 | A>A |
| chr3 | 31100000 | 31200000 | 0.25 | 0.57 | A>A |
| chr3 | 31200000 | 31300000 | 0.54 | 0.77 | A>A |
| chr3 | 31300000 | 31400000 | 0.50 | 0.80 | A>A |
| chr3 | 31400000 | 31500000 | 0.47 | 0.83 | A>A |
| chr3 | 31500000 | 31600000 | 0.55 | 0.78 | A>A |
| chr3 | 31600000 | 31700000 | 0.80 | 0.82 | A>A |
| chr3 | 31700000 | 31800000 | 0.75 | 0.84 | A>A |
| chr3 | 31800000 | 31900000 | 0.86 | 0.97 | A>A |
| chr3 | 31900000 | 32000000 | 1.06 | 1.15 | A>A |
| chr3 | 32000000 | 32100000 | 0.96 | 1.06 | A>A |
| chr3 | 32100000 | 32200000 | 0.98 | 1.04 | A>A |
| chr3 | 32200000 | 32300000 | 1.07 | 1.12 | A>A |

| | | | | | |
|---|---|---|---|---|---|
| chr3 | 32300000 | 32400000 | 1.26 | 1.14 | A>A |
| chr3 | 32400000 | 32500000 | 1.25 | 1.22 | A>A |
| chr3 | 32500000 | 32600000 | 1.21 | 1.19 | A>A |
| chr3 | 32600000 | 32700000 | 1.55 | 1.35 | A>A |
| chr3 | 32700000 | 32800000 | 1.22 | 1.24 | A>A |
| chr3 | 32800000 | 32900000 | 1.57 | 1.47 | A>A |
| chr3 | 32900000 | 33000000 | 1.44 | 1.27 | A>A |
| chr3 | 33000000 | 33100000 | 1.45 | 1.35 | A>A |
| chr3 | 33100000 | 33200000 | 1.34 | 1.27 | A>A |
| chr3 | 33200000 | 33300000 | 1.43 | 1.33 | A>A |
| chr3 | 33300000 | 33400000 | 1.38 | 1.24 | A>A |
| chr3 | 33400000 | 33500000 | 1.17 | 1.24 | A>A |
| chr3 | 33500000 | 33600000 | 1.40 | 1.26 | A>A |
| chr3 | 33600000 | 33700000 | 1.28 | 1.10 | A>A |
| chr3 | 33700000 | 33800000 | 1.29 | 1.08 | A>A |
| chr3 | 33800000 | 33900000 | 0.97 | 0.80 | A>A |
| chr3 | 33900000 | 34000000 | 0.15 | -0.35 | B>B |
| chr3 | 34000000 | 34100000 | 0.30 | -0.20 | A>A |
| chr3 | 34100000 | 34200000 | 0.40 | -0.10 | A>A |
| chr3 | 34200000 | 34300000 | 0.33 | -0.26 | A>B |
| chr3 | 34300000 | 34400000 | -0.12 | -0.76 | A>B |
| chr3 | 34400000 | 34500000 | -0.43 | -1.09 | A>B |
| chr3 | 34500000 | 34600000 | -0.60 | -1.20 | A>B |
| chr3 | 34600000 | 34700000 | -0.67 | -1.24 | B>B |
| chr3 | 34700000 | 34800000 | -0.63 | -1.13 | B>B |
| chr3 | 34800000 | 34900000 | -0.79 | -1.28 | B>B |
| chr3 | 34900000 | 35000000 | -0.77 | -1.29 | B>B |
| chr3 | 35000000 | 35100000 | -0.79 | -1.25 | B>B |
| chr3 | 35100000 | 35200000 | -0.76 | -1.31 | B>B |
| chr3 | 35200000 | 35300000 | -0.63 | -1.34 | A>B |
| chr3 | 35300000 | 35400000 | -0.52 | -1.31 | A>B |
| chr3 | 35400000 | 35500000 | -0.22 | -1.32 | A>B |
| chr3 | 35500000 | 35600000 | -0.16 | -1.28 | A>B |
| chr3 | 35600000 | 35700000 | 0.03 | -1.35 | A>B |
| chr3 | 35700000 | 35800000 | 0.19 | -1.37 | A>B |
| chr3 | 35800000 | 35900000 | 0.30 | -1.31 | A>B |
| chr3 | 35900000 | 36000000 | 0.14 | -0.69 | A>B |
| chr3 | 36000000 | 36100000 | -0.14 | -0.80 | A>B |
| chr3 | 36100000 | 36200000 | 0.00 | -0.52 | B>B |
| chr3 | 36200000 | 36300000 | 0.20 | -0.03 | A>A |
| chr3 | 36300000 | 36400000 | 0.33 | 0.05 | A>A |
| chr3 | 36400000 | 36500000 | 0.38 | 0.01 | A>A |
| chr3 | 36500000 | 36600000 | 0.36 | -0.08 | A>A |
| chr3 | 36600000 | 36700000 | 0.24 | -0.12 | A>A |
| chr3 | 36700000 | 36800000 | 0.31 | -0.10 | A>A |
| chr3 | 36800000 | 36900000 | 0.62 | -0.04 | A>B |
| chr3 | 36900000 | 37000000 | 0.86 | 0.48 | A>A |
| chr3 | 37000000 | 37100000 | 1.20 | 0.83 | A>A |
| chr3 | 37100000 | 37200000 | 1.52 | 1.14 | A>A |
| chr3 | 37200000 | 37300000 | 1.49 | 1.17 | A>A |
| chr3 | 37300000 | 37400000 | 1.33 | 0.94 | A>A |
| chr3 | 37400000 | 37500000 | 1.02 | 0.56 | A>A |
| chr3 | 37500000 | 37600000 | 0.92 | 0.24 | A>B |
| chr3 | 37600000 | 37700000 | 0.89 | 0.23 | A>B |
| chr3 | 37700000 | 37800000 | 0.91 | 0.28 | A>B |
| chr3 | 37800000 | 37900000 | 0.91 | 0.53 | A>A |

| | | | | | |
|---|---|---|---|---|---|
| chr3 | 37900000 | 38000000 | 1.34 | 0.96 | A>A |
| chr3 | 38000000 | 38100000 | 1.37 | 0.92 | A>A |
| chr3 | 38100000 | 38200000 | 1.32 | 0.93 | A>A |
| chr3 | 38200000 | 38300000 | 1.43 | 0.93 | A>A |
| chr3 | 38300000 | 38400000 | 1.12 | 0.70 | A>A |
| chr3 | 38400000 | 38500000 | 0.92 | 0.55 | A>A |
| chr3 | 38500000 | 38600000 | 0.72 | 0.10 | A>B |
| chr3 | 38600000 | 38700000 | 0.54 | -0.34 | A>B |
| chr3 | 38700000 | 38800000 | 0.08 | -0.98 | A>B |
| chr3 | 38800000 | 38900000 | -0.11 | -1.33 | A>B |
| chr3 | 38900000 | 39000000 | -0.15 | -1.21 | A>B |
| chr3 | 39000000 | 39100000 | 0.27 | -0.69 | A>B |
| chr3 | 39100000 | 39200000 | 0.88 | 0.30 | A>B |
| chr3 | 39200000 | 39300000 | 0.92 | 0.30 | A>B |
| chr3 | 39300000 | 39400000 | 0.55 | 0.14 | A>A |
| chr3 | 39400000 | 39500000 | 0.65 | 0.02 | A>B |
| chr3 | 39500000 | 39600000 | 0.24 | -0.48 | A>B |
| chr3 | 39600000 | 39700000 | 0.13 | -0.65 | A>B |
| chr3 | 39700000 | 39800000 | 0.00 | -0.68 | A>B |
| chr3 | 39800000 | 39900000 | -0.11 | -0.66 | B>B |
| chr3 | 39900000 | 40000000 | -0.19 | -0.62 | B>B |
| chr3 | 40000000 | 40100000 | -0.50 | -0.56 | B>B |
| chr3 | 40100000 | 40200000 | -0.16 | -0.25 | B>B |
| chr3 | 40200000 | 40300000 | -0.20 | -0.41 | B>B |
| chr3 | 40300000 | 40400000 | -0.02 | -0.26 | B>B |
| chr3 | 40400000 | 40500000 | 0.22 | -0.18 | A>A |
| chr3 | 40500000 | 40600000 | 0.40 | 0.25 | A>A |
| chr3 | 40600000 | 40700000 | -0.09 | -0.22 | B>B |
| chr3 | 40700000 | 40800000 | -0.16 | -0.37 | B>B |
| chr3 | 40800000 | 40900000 | -0.08 | -0.21 | B>B |
| chr3 | 40900000 | 41000000 | -0.01 | -0.05 | B>B |
| chr3 | 41000000 | 41100000 | 0.08 | -0.02 | A>A |
| chr3 | 41100000 | 41200000 | 0.09 | -0.12 | B>B |
| chr3 | 41200000 | 41300000 | 0.21 | 0.09 | A>A |
| chr3 | 41300000 | 41400000 | 0.30 | 0.13 | A>A |
| chr3 | 41400000 | 41500000 | 0.08 | -0.11 | B>B |
| chr3 | 41500000 | 41600000 | -0.04 | -0.24 | B>B |
| chr3 | 41600000 | 41700000 | 0.01 | -0.18 | B>B |
| chr3 | 41700000 | 41800000 | 0.11 | -0.04 | A>A |
| chr3 | 41800000 | 41900000 | 0.00 | -0.05 | NO |
| chr3 | 41900000 | 42000000 | 0.37 | 0.36 | A>A |
| chr3 | 42000000 | 42100000 | 0.99 | 0.76 | A>A |
| chr3 | 42100000 | 42200000 | 0.76 | 0.68 | A>A |
| chr3 | 42200000 | 42300000 | 0.57 | 0.52 | A>A |
| chr3 | 42300000 | 42400000 | 0.39 | 0.47 | A>A |
| chr3 | 42400000 | 42500000 | 0.50 | 0.41 | A>A |
| chr3 | 42500000 | 42600000 | 0.67 | 0.69 | A>A |
| chr3 | 42600000 | 42700000 | 1.12 | 1.05 | A>A |
| chr3 | 42700000 | 42800000 | 1.11 | 0.91 | A>A |
| chr3 | 42800000 | 42900000 | 0.83 | 0.84 | A>A |
| chr3 | 42900000 | 43000000 | 0.45 | 0.60 | A>A |
| chr3 | 43000000 | 43100000 | 0.45 | 0.56 | A>A |
| chr3 | 43100000 | 43200000 | 0.56 | 0.61 | A>A |
| chr3 | 43200000 | 43300000 | 0.37 | 0.61 | A>A |
| chr3 | 43300000 | 43400000 | 0.64 | 0.80 | A>A |
| chr3 | 43400000 | 43500000 | 0.54 | 0.69 | A>A |

| | | | | | |
|---|---|---|---|---|---|
| chr3 | 43500000 | 43600000 | 0.47 | 0.58 | A>A |
| chr3 | 43600000 | 43700000 | 0.85 | 0.91 | A>A |
| chr3 | 43700000 | 43800000 | 0.75 | 0.77 | A>A |
| chr3 | 43800000 | 43900000 | 0.83 | 0.82 | A>A |
| chr3 | 43900000 | 44000000 | 1.09 | 1.10 | A>A |
| chr3 | 44000000 | 44100000 | 0.90 | 1.06 | A>A |
| chr3 | 44100000 | 44200000 | 0.64 | 0.86 | A>A |
| chr3 | 44200000 | 44300000 | 0.38 | 0.60 | A>A |
| chr3 | 44300000 | 44400000 | 0.43 | 0.88 | A>A |
| chr3 | 44400000 | 44500000 | 0.53 | 1.13 | B>A |
| chr3 | 44500000 | 44600000 | 0.40 | 0.80 | A>A |
| chr3 | 44600000 | 44700000 | 0.27 | 0.93 | B>A |
| chr3 | 44700000 | 44800000 | 0.36 | 1.00 | B>A |
| chr3 | 44800000 | 44900000 | 0.82 | 1.08 | A>A |
| chr3 | 44900000 | 45000000 | 0.88 | 0.99 | A>A |
| chr3 | 45000000 | 45100000 | 0.71 | 0.77 | A>A |
| chr3 | 45100000 | 45200000 | 1.02 | 1.08 | A>A |
| chr3 | 45200000 | 45300000 | 0.83 | 0.90 | A>A |
| chr3 | 45300000 | 45400000 | 0.84 | 0.86 | A>A |
| chr3 | 45400000 | 45500000 | 0.95 | 0.87 | A>A |
| chr3 | 45500000 | 45600000 | 0.71 | 0.75 | A>A |
| chr3 | 45600000 | 45700000 | 1.19 | 1.05 | A>A |
| chr3 | 45700000 | 45800000 | 0.86 | 0.80 | A>A |
| chr3 | 45800000 | 45900000 | 0.77 | 0.82 | A>A |
| chr3 | 45900000 | 46000000 | 0.66 | 0.60 | A>A |
| chr3 | 46000000 | 46100000 | 0.79 | 0.60 | A>A |
| chr3 | 46100000 | 46200000 | 0.41 | 0.45 | A>A |
| chr3 | 46200000 | 46300000 | 0.00 | 0.35 | NO |
| chr3 | 46300000 | 46400000 | 0.44 | 0.37 | A>A |
| chr3 | 46400000 | 46500000 | 0.60 | 0.42 | A>A |
| chr3 | 46500000 | 46600000 | 0.81 | 0.56 | A>A |
| chr3 | 46600000 | 46700000 | 0.00 | 0.72 | NO |
| chr3 | 46700000 | 46800000 | 0.79 | 0.80 | A>A |
| chr3 | 46800000 | 46900000 | 0.00 | 0.84 | NO |
| chr3 | 46900000 | 47000000 | 0.89 | 0.86 | A>A |
| chr3 | 47000000 | 47100000 | 1.08 | 0.97 | A>A |
| chr3 | 47100000 | 47200000 | 1.15 | 1.08 | A>A |
| chr3 | 47200000 | 47300000 | 1.07 | 1.05 | A>A |
| chr3 | 47300000 | 47400000 | 1.02 | 0.93 | A>A |
| chr3 | 47400000 | 47500000 | 0.91 | 0.98 | A>A |
| chr3 | 47500000 | 47600000 | 0.93 | 0.96 | A>A |
| chr3 | 47600000 | 47700000 | 1.31 | 1.09 | A>A |
| chr3 | 47700000 | 47800000 | 1.22 | 0.00 | NO |
| chr3 | 47800000 | 47900000 | 0.88 | 0.81 | A>A |
| chr3 | 47900000 | 48000000 | 1.14 | 1.16 | A>A |
| chr3 | 48000000 | 48100000 | 0.00 | 0.00 | NO |
| chr3 | 48100000 | 48200000 | 0.00 | 0.00 | NO |
| chr3 | 48200000 | 48300000 | 0.90 | 0.78 | A>A |
| chr3 | 48300000 | 48400000 | 0.00 | 0.00 | NO |
| chr3 | 48400000 | 48500000 | 0.00 | 0.77 | NO |
| chr3 | 48500000 | 48600000 | 0.00 | 0.00 | NO |
| chr3 | 48600000 | 48700000 | 0.97 | 0.85 | A>A |
| chr3 | 48700000 | 48800000 | 1.16 | 0.00 | NO |
| chr3 | 48800000 | 48900000 | 0.00 | 0.00 | NO |
| chr3 | 48900000 | 49000000 | 0.00 | 0.00 | NO |
| chr3 | 49000000 | 49100000 | 0.96 | 0.98 | A>A |

| | | | | | |
|---|---|---|---|---|---|
| chr3 | 49100000 | 49200000 | 0.00 | 0.00 | NO |
| chr3 | 49200000 | 49300000 | 0.00 | 0.00 | NO |
| chr3 | 49300000 | 49400000 | 0.00 | 0.00 | NO |
| chr3 | 49400000 | 49500000 | 0.00 | 0.00 | NO |
| chr3 | 49500000 | 49600000 | 1.06 | 0.94 | A>A |
| chr3 | 49600000 | 49700000 | 0.00 | 1.09 | NO |
| chr3 | 49700000 | 49800000 | 0.00 | 0.80 | NO |
| chr3 | 49800000 | 49900000 | 0.95 | 0.96 | A>A |
| chr3 | 49900000 | 50000000 | 1.15 | 0.96 | A>A |
| chr3 | 50000000 | 50100000 | 1.25 | 1.10 | A>A |
| chr3 | 50100000 | 50200000 | 1.07 | 0.99 | A>A |
| chr3 | 50200000 | 50300000 | 1.10 | 1.10 | A>A |
| chr3 | 50300000 | 50400000 | 1.04 | 1.02 | A>A |
| chr3 | 50400000 | 50500000 | 1.04 | 0.84 | A>A |
| chr3 | 50500000 | 50600000 | 1.02 | 0.78 | A>A |
| chr3 | 50600000 | 50700000 | 0.68 | 0.43 | A>A |
| chr3 | 50700000 | 50800000 | 0.42 | -0.03 | A>A |
| chr3 | 50800000 | 50900000 | 0.00 | -0.37 | B>B |
| chr3 | 50900000 | 51000000 | 0.00 | -0.28 | NO |
| chr3 | 51000000 | 51100000 | 0.00 | -0.34 | NO |
| chr3 | 51100000 | 51200000 | 0.00 | -0.25 | NO |
| chr3 | 51200000 | 51300000 | 0.00 | -0.32 | NO |
| chr3 | 51300000 | 51400000 | 0.37 | -0.12 | A>A |
| chr3 | 51400000 | 51500000 | 0.75 | 0.45 | A>A |
| chr3 | 51500000 | 51600000 | 1.01 | 0.83 | A>A |
| chr3 | 51600000 | 51700000 | 0.00 | 0.96 | NO |
| chr3 | 51700000 | 51800000 | 0.86 | 0.64 | A>A |
| chr3 | 51800000 | 51900000 | 0.84 | 0.80 | A>A |
| chr3 | 51900000 | 52000000 | 0.87 | 0.80 | A>A |
| chr3 | 52000000 | 52100000 | 1.07 | 1.00 | A>A |
| chr3 | 52100000 | 52200000 | 1.13 | 1.05 | A>A |
| chr3 | 52200000 | 52300000 | 1.10 | 1.02 | A>A |
| chr3 | 52300000 | 52400000 | 1.12 | 1.15 | A>A |
| chr3 | 52400000 | 52500000 | 0.97 | 1.05 | A>A |
| chr3 | 52500000 | 52600000 | 1.20 | 1.13 | A>A |
| chr3 | 52600000 | 52700000 | 1.27 | 1.20 | A>A |
| chr3 | 52700000 | 52800000 | 0.95 | 1.08 | A>A |
| chr3 | 52800000 | 52900000 | 1.05 | 1.19 | A>A |
| chr3 | 52900000 | 53000000 | 1.13 | 1.24 | A>A |
| chr3 | 53000000 | 53100000 | 1.08 | 1.17 | A>A |
| chr3 | 53100000 | 53200000 | 0.98 | 0.98 | A>A |
| chr3 | 53200000 | 53300000 | 1.08 | 1.16 | A>A |
| chr3 | 53300000 | 53400000 | 0.79 | 0.76 | A>A |
| chr3 | 53400000 | 53500000 | 0.65 | 0.71 | A>A |
| chr3 | 53500000 | 53600000 | 0.90 | 0.81 | A>A |
| chr3 | 53600000 | 53700000 | 0.87 | 0.80 | A>A |
| chr3 | 53700000 | 53800000 | 0.62 | 0.60 | A>A |
| chr3 | 53800000 | 53900000 | 0.69 | 0.62 | A>A |
| chr3 | 53900000 | 54000000 | 0.32 | 0.11 | A>A |
| chr3 | 54000000 | 54100000 | 0.19 | -0.33 | B>B |
| chr3 | 54100000 | 54200000 | 0.06 | -0.39 | B>B |
| chr3 | 54200000 | 54300000 | 0.00 | -0.47 | B>B |
| chr3 | 54300000 | 54400000 | -0.15 | -0.60 | B>B |
| chr3 | 54400000 | 54500000 | -0.22 | -0.65 | B>B |
| chr3 | 54500000 | 54600000 | -0.29 | -0.86 | B>B |
| chr3 | 54600000 | 54700000 | -0.31 | -0.89 | B>B |

| | | | | | |
|---|---|---|---|---|---|
| chr3 | 54700000 | 54800000 | -0.32 | -0.83 | B>B |
| chr3 | 54800000 | 54900000 | -0.18 | -0.48 | B>B |
| chr3 | 54900000 | 55000000 | 0.30 | -0.04 | A>A |
| chr3 | 55000000 | 55100000 | 0.06 | -0.13 | B>B |
| chr3 | 55100000 | 55200000 | -0.02 | -0.24 | B>B |
| chr3 | 55200000 | 55300000 | 0.04 | -0.20 | B>B |
| chr3 | 55300000 | 55400000 | -0.06 | -0.30 | B>B |
| chr3 | 55400000 | 55500000 | 0.05 | -0.28 | B>B |
| chr3 | 55500000 | 55600000 | 0.05 | -0.51 | B>B |
| chr3 | 55600000 | 55700000 | 0.06 | -0.80 | A>B |
| chr3 | 55700000 | 55800000 | -0.17 | -0.76 | A>B |
| chr3 | 55800000 | 55900000 | -0.14 | -0.74 | A>B |
| chr3 | 55900000 | 56000000 | -0.11 | -0.63 | B>B |
| chr3 | 56000000 | 56100000 | -0.05 | -0.49 | B>B |
| chr3 | 56100000 | 56200000 | 0.18 | -0.12 | A>A |
| chr3 | 56200000 | 56300000 | 0.02 | -0.32 | B>B |
| chr3 | 56300000 | 56400000 | -0.06 | -0.56 | B>B |
| chr3 | 56400000 | 56500000 | -0.05 | -0.67 | A>B |
| chr3 | 56500000 | 56600000 | 0.21 | -0.34 | B>B |
| chr3 | 56600000 | 56700000 | 0.00 | 0.50 | NO |
| chr3 | 56700000 | 56800000 | 0.86 | 0.79 | A>A |
| chr3 | 56800000 | 56900000 | 1.03 | 0.86 | A>A |
| chr3 | 56900000 | 57000000 | 0.98 | 0.76 | A>A |
| chr3 | 57000000 | 57100000 | 0.86 | 0.77 | A>A |
| chr3 | 57100000 | 57200000 | 0.97 | 0.91 | A>A |
| chr3 | 57200000 | 57300000 | 0.96 | 0.84 | A>A |
| chr3 | 57300000 | 57400000 | 0.00 | 0.00 | NO |
| chr3 | 57400000 | 57500000 | 0.00 | 0.00 | NO |
| chr3 | 57500000 | 57600000 | 1.10 | 0.97 | A>A |
| chr3 | 57600000 | 57700000 | 1.11 | 1.07 | A>A |
| chr3 | 57700000 | 57800000 | 1.40 | 1.17 | A>A |
| chr3 | 57800000 | 57900000 | 0.91 | 0.93 | A>A |
| chr3 | 57900000 | 58000000 | 1.01 | 0.96 | A>A |
| chr3 | 58000000 | 58100000 | 1.26 | 1.14 | A>A |
| chr3 | 58100000 | 58200000 | 0.93 | 0.95 | A>A |
| chr3 | 58200000 | 58300000 | 0.89 | 0.83 | A>A |
| chr3 | 58300000 | 58400000 | 0.83 | 0.58 | A>A |
| chr3 | 58400000 | 58500000 | 0.84 | 0.67 | A>A |
| chr3 | 58500000 | 58600000 | 0.63 | 0.28 | A>A |
| chr3 | 58600000 | 58700000 | 0.20 | -0.22 | B>B |
| chr3 | 58700000 | 58800000 | -0.06 | -0.56 | B>B |
| chr3 | 58800000 | 58900000 | 0.00 | -0.64 | NO |
| chr3 | 58900000 | 59000000 | -0.12 | -0.55 | B>B |
| chr3 | 59000000 | 59100000 | -0.05 | -0.49 | B>B |
| chr3 | 59100000 | 59200000 | -0.24 | -0.82 | B>B |
| chr3 | 59200000 | 59300000 | -0.17 | -0.82 | A>B |
| chr3 | 59300000 | 59400000 | -0.20 | -0.85 | A>B |
| chr3 | 59400000 | 59500000 | -0.17 | -0.80 | A>B |
| chr3 | 59500000 | 59600000 | -0.08 | -0.72 | A>B |
| chr3 | 59600000 | 59700000 | 0.19 | -0.70 | A>B |
| chr3 | 59700000 | 59800000 | 0.16 | -0.68 | A>B |
| chr3 | 59800000 | 59900000 | -0.19 | 0.00 | NO |
| chr3 | 59900000 | 60000000 | -0.46 | 0.00 | NO |
| chr3 | 60000000 | 60100000 | -0.60 | 0.00 | NO |
| chr3 | 60100000 | 60200000 | -0.66 | 0.00 | NO |
| chr3 | 60200000 | 60300000 | -0.69 | -0.79 | B>B |

| | | | | | |
|---|---|---|---|---|---|
| chr3 | 60300000 | 60400000 | -0.68 | -0.80 | B>B |
| chr3 | 60400000 | 60500000 | 0.00 | -0.77 | NO |
| chr3 | 60500000 | 60600000 | -0.67 | -0.76 | B>B |
| chr3 | 60600000 | 60700000 | -0.77 | -0.72 | B>B |
| chr3 | 60700000 | 60800000 | -0.77 | -0.72 | B>B |
| chr3 | 60800000 | 60900000 | -0.72 | -0.70 | B>B |
| chr3 | 60900000 | 61000000 | -0.76 | -0.77 | B>B |
| chr3 | 61000000 | 61100000 | -0.63 | -0.66 | B>B |
| chr3 | 61100000 | 61200000 | -0.48 | -0.64 | B>B |
| chr3 | 61200000 | 61300000 | -0.27 | -0.52 | B>B |
| chr3 | 61300000 | 61400000 | -0.31 | -0.52 | B>B |
| chr3 | 61400000 | 61500000 | -0.08 | -0.46 | B>B |
| chr3 | 61500000 | 61600000 | 0.24 | -0.02 | A>A |
| chr3 | 61600000 | 61700000 | 0.54 | 0.13 | A>A |
| chr3 | 61700000 | 61800000 | 0.32 | -0.05 | A>A |
| chr3 | 61800000 | 61900000 | 0.22 | -0.19 | A>A |
| chr3 | 61900000 | 62000000 | 0.29 | -0.34 | A>B |
| chr3 | 62000000 | 62100000 | 0.17 | -0.51 | A>B |
| chr3 | 62100000 | 62200000 | -0.02 | -0.38 | B>B |
| chr3 | 62200000 | 62300000 | -0.06 | -0.44 | B>B |
| chr3 | 62300000 | 62400000 | -0.11 | -0.58 | B>B |
| chr3 | 62400000 | 62500000 | -0.23 | -0.77 | B>B |
| chr3 | 62500000 | 62600000 | -0.30 | -0.70 | B>B |
| chr3 | 62600000 | 62700000 | -0.45 | -0.74 | B>B |
| chr3 | 62700000 | 62800000 | -0.53 | -0.82 | B>B |
| chr3 | 62800000 | 62900000 | -0.52 | -0.88 | B>B |
| chr3 | 62900000 | 63000000 | -0.57 | -0.90 | B>B |
| chr3 | 63000000 | 63100000 | -0.67 | -0.86 | B>B |
| chr3 | 63100000 | 63200000 | -0.61 | -0.90 | B>B |
| chr3 | 63200000 | 63300000 | -0.46 | -0.79 | B>B |
| chr3 | 63300000 | 63400000 | -0.40 | -0.69 | B>B |
| chr3 | 63400000 | 63500000 | -0.50 | -0.86 | B>B |
| chr3 | 63500000 | 63600000 | -0.49 | -0.85 | B>B |
| chr3 | 63600000 | 63700000 | -0.41 | -0.78 | B>B |
| chr3 | 63700000 | 63800000 | -0.28 | -0.61 | B>B |
| chr3 | 63800000 | 63900000 | 0.29 | 0.10 | A>A |
| chr3 | 63900000 | 64000000 | 0.55 | 0.40 | A>A |
| chr3 | 64000000 | 64100000 | 0.63 | 0.52 | A>A |
| chr3 | 64100000 | 64200000 | 0.74 | 0.71 | A>A |
| chr3 | 64200000 | 64300000 | 0.72 | 0.58 | A>A |
| chr3 | 64300000 | 64400000 | 0.68 | 0.55 | A>A |
| chr3 | 64400000 | 64500000 | 0.33 | 0.22 | A>A |
| chr3 | 64500000 | 64600000 | -0.19 | -0.23 | B>B |
| chr3 | 64600000 | 64700000 | -0.23 | -0.36 | B>B |
| chr3 | 64700000 | 64800000 | -0.43 | -0.55 | B>B |
| chr3 | 64800000 | 64900000 | -0.52 | -0.62 | B>B |
| chr3 | 64900000 | 65000000 | -0.54 | -0.66 | B>B |
| chr3 | 65000000 | 65100000 | -0.58 | -0.73 | B>B |
| chr3 | 65100000 | 65200000 | -0.52 | -0.75 | B>B |
| chr3 | 65200000 | 65300000 | -0.41 | -0.65 | B>B |
| chr3 | 65300000 | 65400000 | -0.22 | -0.48 | B>B |
| chr3 | 65400000 | 65500000 | -0.18 | -0.33 | B>B |
| chr3 | 65500000 | 65600000 | -0.07 | -0.35 | B>B |
| chr3 | 65600000 | 65700000 | -0.12 | -0.25 | B>B |
| chr3 | 65700000 | 65800000 | 0.10 | -0.02 | A>A |
| chr3 | 65800000 | 65900000 | 0.52 | 0.28 | A>A |

| | | | | | |
|---|---|---|---|---|---|
| chr3 | 65900000 | 66000000 | 0.63 | 0.36 | A>A |
| chr3 | 66000000 | 66100000 | 0.51 | 0.36 | A>A |
| chr3 | 66100000 | 66200000 | 0.00 | 0.04 | NO |
| chr3 | 66200000 | 66300000 | 0.00 | 0.00 | NO |
| chr3 | 66300000 | 66400000 | 0.19 | -0.02 | A>A |
| chr3 | 66400000 | 66500000 | 0.40 | 0.15 | A>A |
| chr3 | 66500000 | 66600000 | 0.68 | 0.32 | A>A |
| chr3 | 66600000 | 66700000 | 0.67 | 0.29 | A>A |
| chr3 | 66700000 | 66800000 | 0.48 | 0.06 | A>A |
| chr3 | 66800000 | 66900000 | 0.18 | -0.10 | A>A |
| chr3 | 66900000 | 67000000 | -0.26 | -0.51 | B>B |
| chr3 | 67000000 | 67100000 | -0.17 | -0.41 | B>B |
| chr3 | 67100000 | 67200000 | -0.37 | -0.60 | B>B |
| chr3 | 67200000 | 67300000 | -0.55 | -0.77 | B>B |
| chr3 | 67300000 | 67400000 | -0.40 | -0.64 | B>B |
| chr3 | 67400000 | 67500000 | -0.03 | -0.16 | B>B |
| chr3 | 67500000 | 67600000 | 0.27 | 0.15 | A>A |
| chr3 | 67600000 | 67700000 | 0.28 | 0.18 | A>A |
| chr3 | 67700000 | 67800000 | 0.17 | 0.05 | A>A |
| chr3 | 67800000 | 67900000 | -0.04 | -0.26 | B>B |
| chr3 | 67900000 | 68000000 | -0.21 | -0.55 | B>B |
| chr3 | 68000000 | 68100000 | -0.33 | -0.72 | B>B |
| chr3 | 68100000 | 68200000 | -0.49 | -0.78 | B>B |
| chr3 | 68200000 | 68300000 | -0.56 | -0.75 | B>B |
| chr3 | 68300000 | 68400000 | -0.58 | -0.78 | B>B |
| chr3 | 68400000 | 68500000 | -0.61 | -0.89 | B>B |
| chr3 | 68500000 | 68600000 | -0.60 | -0.85 | B>B |
| chr3 | 68600000 | 68700000 | -0.58 | -0.89 | B>B |
| chr3 | 68700000 | 68800000 | -0.62 | -0.83 | B>B |
| chr3 | 68800000 | 68900000 | -0.52 | -0.74 | B>B |
| chr3 | 68900000 | 69000000 | -0.24 | -0.36 | B>B |
| chr3 | 69000000 | 69100000 | 0.35 | 0.17 | A>A |
| chr3 | 69100000 | 69200000 | 0.78 | 0.64 | A>A |
| chr3 | 69200000 | 69300000 | 0.92 | 0.85 | A>A |
| chr3 | 69300000 | 69400000 | 0.71 | 0.70 | A>A |
| chr3 | 69400000 | 69500000 | 0.55 | 0.43 | A>A |
| chr3 | 69500000 | 69600000 | 0.35 | 0.27 | A>A |
| chr3 | 69600000 | 69700000 | 0.45 | 0.22 | A>A |
| chr3 | 69700000 | 69800000 | 0.46 | 0.49 | A>A |
| chr3 | 69800000 | 69900000 | 0.63 | 0.64 | A>A |
| chr3 | 69900000 | 70000000 | 0.56 | 0.64 | A>A |
| chr3 | 70000000 | 70100000 | 0.14 | 0.42 | A>A |
| chr3 | 70100000 | 70200000 | 0.08 | 0.27 | A>A |
| chr3 | 70200000 | 70300000 | 0.39 | 0.67 | A>A |
| chr3 | 70300000 | 70400000 | 0.39 | 0.61 | A>A |
| chr3 | 70400000 | 70500000 | 0.10 | 0.39 | A>A |
| chr3 | 70500000 | 70600000 | 0.03 | 0.28 | A>A |
| chr3 | 70600000 | 70700000 | -0.12 | 0.22 | A>A |
| chr3 | 70700000 | 70800000 | 0.05 | 0.26 | A>A |
| chr3 | 70800000 | 70900000 | 0.16 | 0.36 | A>A |
| chr3 | 70900000 | 71000000 | 0.14 | 0.30 | A>A |
| chr3 | 71000000 | 71100000 | 0.18 | 0.38 | A>A |
| chr3 | 71100000 | 71200000 | 0.41 | 0.56 | A>A |
| chr3 | 71200000 | 71300000 | 0.50 | 0.67 | A>A |
| chr3 | 71300000 | 71400000 | 0.45 | 0.68 | A>A |
| chr3 | 71400000 | 71500000 | 0.41 | 0.61 | A>A |

| | | | | | |
|---|---|---|---|---|---|
| chr3 | 71500000 | 71600000 | 0.50 | 0.59 | A>A |
| chr3 | 71600000 | 71700000 | 0.23 | 0.33 | A>A |
| chr3 | 71700000 | 71800000 | 0.30 | 0.24 | A>A |
| chr3 | 71800000 | 71900000 | 0.30 | 0.16 | A>A |
| chr3 | 71900000 | 72000000 | 0.17 | 0.14 | A>A |
| chr3 | 72000000 | 72100000 | 0.29 | 0.22 | A>A |
| chr3 | 72100000 | 72200000 | 0.44 | 0.35 | A>A |
| chr3 | 72200000 | 72300000 | 0.36 | 0.33 | A>A |
| chr3 | 72300000 | 72400000 | 0.36 | 0.37 | A>A |
| chr3 | 72400000 | 72500000 | 0.55 | 0.49 | A>A |
| chr3 | 72500000 | 72600000 | 0.55 | 0.53 | A>A |
| chr3 | 72600000 | 72700000 | 0.56 | 0.53 | A>A |
| chr3 | 72700000 | 72800000 | 0.53 | 0.54 | A>A |
| chr3 | 72800000 | 72900000 | 0.40 | 0.37 | A>A |
| chr3 | 72900000 | 73000000 | 0.47 | 0.26 | A>A |
| chr3 | 73000000 | 73100000 | 0.67 | 0.34 | A>A |
| chr3 | 73100000 | 73200000 | 0.09 | -0.18 | B>B |
| chr3 | 73200000 | 73300000 | -0.21 | -0.69 | B>B |
| chr3 | 73300000 | 73400000 | -0.35 | -0.79 | B>B |
| chr3 | 73400000 | 73500000 | -0.31 | -0.79 | B>B |
| chr3 | 73500000 | 73600000 | -0.34 | -0.70 | B>B |
| chr3 | 73600000 | 73700000 | -0.53 | -0.78 | B>B |
| chr3 | 73700000 | 73800000 | -0.61 | -0.85 | B>B |
| chr3 | 73800000 | 73900000 | -0.66 | -0.82 | B>B |
| chr3 | 73900000 | 74000000 | -0.69 | -0.78 | B>B |
| chr3 | 74000000 | 74100000 | -0.71 | -0.81 | B>B |
| chr3 | 74100000 | 74200000 | -0.69 | -0.77 | B>B |
| chr3 | 74200000 | 74300000 | -0.68 | -0.82 | B>B |
| chr3 | 74300000 | 74400000 | -0.67 | -0.77 | B>B |
| chr3 | 74400000 | 74500000 | -0.75 | -0.76 | B>B |
| chr3 | 74500000 | 74600000 | -0.72 | -0.76 | B>B |
| chr3 | 74600000 | 74700000 | -0.68 | -0.72 | B>B |
| chr3 | 74700000 | 74800000 | -0.69 | -0.79 | B>B |
| chr3 | 74800000 | 74900000 | -0.74 | -0.73 | B>B |
| chr3 | 74900000 | 75000000 | -0.67 | -0.80 | B>B |
| chr3 | 75000000 | 75100000 | -0.69 | -0.77 | B>B |
| chr3 | 75100000 | 75200000 | -0.66 | -0.66 | B>B |
| chr3 | 75200000 | 75300000 | -0.82 | -1.05 | B>B |
| chr3 | 75300000 | 75400000 | 0.00 | 0.00 | NO |
| chr3 | 75400000 | 75500000 | 0.00 | 0.00 | NO |
| chr3 | 75500000 | 75600000 | 0.00 | 0.00 | NO |
| chr3 | 75600000 | 75700000 | 0.00 | 0.00 | NO |
| chr3 | 75700000 | 75800000 | 0.00 | 0.00 | NO |
| chr3 | 75800000 | 75900000 | 0.00 | 0.00 | NO |
| chr3 | 75900000 | 76000000 | 0.00 | -0.76 | NO |
| chr3 | 76000000 | 76100000 | -0.75 | -0.78 | B>B |
| chr3 | 76100000 | 76200000 | -0.72 | -0.74 | B>B |
| chr3 | 76200000 | 76300000 | -0.74 | -0.82 | B>B |
| chr3 | 76300000 | 76400000 | -0.68 | -0.73 | B>B |
| chr3 | 76400000 | 76500000 | -0.73 | -0.81 | B>B |
| chr3 | 76500000 | 76600000 | -0.68 | -0.75 | B>B |
| chr3 | 76600000 | 76700000 | -0.65 | -0.71 | B>B |
| chr3 | 76700000 | 76800000 | -0.66 | -0.74 | B>B |
| chr3 | 76800000 | 76900000 | -0.64 | -0.78 | B>B |
| chr3 | 76900000 | 77000000 | -0.69 | -0.74 | B>B |
| chr3 | 77000000 | 77100000 | -0.61 | -0.64 | B>B |

| | | | | | |
|---|---|---|---|---|---|
| chr3 | 77100000 | 77200000 | -0.63 | -0.70 | B>B |
| chr3 | 77200000 | 77300000 | -0.65 | -0.73 | B>B |
| chr3 | 77300000 | 77400000 | -0.57 | -0.66 | B>B |
| chr3 | 77400000 | 77500000 | -0.58 | -0.65 | B>B |
| chr3 | 77500000 | 77600000 | -0.66 | -0.65 | B>B |
| chr3 | 77600000 | 77700000 | -0.62 | -0.60 | B>B |
| chr3 | 77700000 | 77800000 | -0.61 | -0.55 | B>B |
| chr3 | 77800000 | 77900000 | -0.58 | -0.50 | B>B |
| chr3 | 77900000 | 78000000 | -0.62 | -0.55 | B>B |
| chr3 | 78000000 | 78100000 | -0.63 | -0.55 | B>B |
| chr3 | 78100000 | 78200000 | -0.60 | -0.53 | B>B |
| chr3 | 78200000 | 78300000 | -0.57 | -0.57 | B>B |
| chr3 | 78300000 | 78400000 | -0.55 | -0.61 | B>B |
| chr3 | 78400000 | 78500000 | -0.62 | -0.63 | B>B |
| chr3 | 78500000 | 78600000 | -0.55 | -0.60 | B>B |
| chr3 | 78600000 | 78700000 | -0.38 | -0.41 | B>B |
| chr3 | 78700000 | 78800000 | -0.12 | 0.03 | B>B |
| chr3 | 78800000 | 78900000 | -0.02 | 0.15 | A>A |
| chr3 | 78900000 | 79000000 | -0.02 | 0.17 | A>A |
| chr3 | 79000000 | 79100000 | -0.18 | -0.06 | B>B |
| chr3 | 79100000 | 79200000 | -0.53 | -0.44 | B>B |
| chr3 | 79200000 | 79300000 | -0.58 | -0.54 | B>B |
| chr3 | 79300000 | 79400000 | -0.63 | -0.52 | B>B |
| chr3 | 79400000 | 79500000 | -0.58 | -0.50 | B>B |
| chr3 | 79500000 | 79600000 | -0.56 | -0.43 | B>B |
| chr3 | 79600000 | 79700000 | -0.60 | -0.39 | B>B |
| chr3 | 79700000 | 79800000 | -0.61 | -0.44 | B>B |
| chr3 | 79800000 | 79900000 | -0.63 | -0.46 | B>B |
| chr3 | 79900000 | 80000000 | -0.60 | -0.43 | B>B |
| chr3 | 80000000 | 80100000 | -0.63 | -0.37 | B>B |
| chr3 | 80100000 | 80200000 | -0.61 | -0.41 | B>B |
| chr3 | 80200000 | 80300000 | -0.61 | -0.46 | B>B |
| chr3 | 80300000 | 80400000 | -0.66 | -0.49 | B>B |
| chr3 | 80400000 | 80500000 | -0.64 | -0.47 | B>B |
| chr3 | 80500000 | 80600000 | -0.64 | -0.52 | B>B |
| chr3 | 80600000 | 80700000 | -0.56 | -0.58 | B>B |
| chr3 | 80700000 | 80800000 | -0.62 | -0.60 | B>B |
| chr3 | 80800000 | 80900000 | -0.56 | -0.48 | B>B |
| chr3 | 80900000 | 81000000 | -0.66 | -0.17 | B>B |
| chr3 | 81000000 | 81100000 | -0.66 | -0.31 | B>B |
| chr3 | 81100000 | 81200000 | -0.65 | -0.27 | B>B |
| chr3 | 81200000 | 81300000 | -0.58 | -0.25 | B>B |
| chr3 | 81300000 | 81400000 | -0.52 | -0.22 | B>B |
| chr3 | 81400000 | 81500000 | -0.35 | -0.17 | B>B |
| chr3 | 81500000 | 81600000 | -0.12 | -0.05 | B>B |
| chr3 | 81600000 | 81700000 | 0.03 | 0.03 | A>A |
| chr3 | 81700000 | 81800000 | 0.23 | 0.27 | A>A |
| chr3 | 81800000 | 81900000 | 0.17 | 0.06 | A>A |
| chr3 | 81900000 | 82000000 | -0.16 | -0.12 | B>B |
| chr3 | 82000000 | 82100000 | -0.34 | -0.22 | B>B |
| chr3 | 82100000 | 82200000 | -0.40 | -0.21 | B>B |
| chr3 | 82200000 | 82300000 | -0.53 | -0.19 | B>B |
| chr3 | 82300000 | 82400000 | -0.61 | -0.20 | B>B |
| chr3 | 82400000 | 82500000 | -0.58 | -0.19 | B>B |
| chr3 | 82500000 | 82600000 | -0.62 | -0.21 | B>B |
| chr3 | 82600000 | 82700000 | -0.59 | -0.12 | B>B |

| | | | | | |
|---|---|---|---|---|---|
| chr3 | 82700000 | 82800000 | -0.55 | -0.06 | B>B |
| chr3 | 82800000 | 82900000 | -0.56 | -0.07 | B>B |
| chr3 | 82900000 | 83000000 | -0.51 | 0.00 | B>A |
| chr3 | 83000000 | 83100000 | -0.54 | 0.05 | B>A |
| chr3 | 83100000 | 83200000 | -0.55 | 0.04 | B>A |
| chr3 | 83200000 | 83300000 | -0.52 | -0.03 | B>B |
| chr3 | 83300000 | 83400000 | -0.54 | 0.01 | B>A |
| chr3 | 83400000 | 83500000 | -0.55 | 0.01 | B>A |
| chr3 | 83500000 | 83600000 | -0.53 | -0.02 | B>A |
| chr3 | 83600000 | 83700000 | -0.53 | 0.09 | B>A |
| chr3 | 83700000 | 83800000 | -0.57 | 0.11 | B>A |
| chr3 | 83800000 | 83900000 | -0.52 | 0.09 | B>A |
| chr3 | 83900000 | 84000000 | -0.50 | 0.09 | B>A |
| chr3 | 84000000 | 84100000 | -0.57 | 0.20 | B>A |
| chr3 | 84100000 | 84200000 | -0.55 | 0.20 | B>A |
| chr3 | 84200000 | 84300000 | -0.55 | 0.16 | B>A |
| chr3 | 84300000 | 84400000 | -0.53 | 0.18 | B>A |
| chr3 | 84400000 | 84500000 | -0.63 | 0.19 | B>A |
| chr3 | 84500000 | 84600000 | -0.54 | 0.05 | B>A |
| chr3 | 84600000 | 84700000 | -0.58 | 0.05 | B>A |
| chr3 | 84700000 | 84800000 | -0.57 | 0.10 | B>A |
| chr3 | 84800000 | 84900000 | -0.55 | 0.02 | B>A |
| chr3 | 84900000 | 85000000 | -0.52 | 0.04 | B>A |
| chr3 | 85000000 | 85100000 | -0.49 | 0.10 | B>A |
| chr3 | 85100000 | 85200000 | -0.57 | 0.15 | B>A |
| chr3 | 85200000 | 85300000 | -0.57 | 0.10 | B>A |
| chr3 | 85300000 | 85400000 | -0.58 | 0.07 | B>A |
| chr3 | 85400000 | 85500000 | -0.58 | 0.07 | B>A |
| chr3 | 85500000 | 85600000 | -0.62 | 0.06 | B>A |
| chr3 | 85600000 | 85700000 | -0.59 | 0.00 | B>A |
| chr3 | 85700000 | 85800000 | -0.61 | -0.04 | B>A |
| chr3 | 85800000 | 85900000 | -0.56 | -0.03 | B>A |
| chr3 | 85900000 | 86000000 | -0.53 | -0.08 | B>B |
| chr3 | 86000000 | 86100000 | -0.53 | -0.08 | B>B |
| chr3 | 86100000 | 86200000 | -0.49 | -0.09 | B>B |
| chr3 | 86200000 | 86300000 | -0.47 | -0.10 | B>B |
| chr3 | 86300000 | 86400000 | -0.59 | -0.31 | B>B |
| chr3 | 86400000 | 86500000 | -0.62 | -0.34 | B>B |
| chr3 | 86500000 | 86600000 | -0.65 | -0.39 | B>B |
| chr3 | 86600000 | 86700000 | -0.60 | -0.33 | B>B |
| chr3 | 86700000 | 86800000 | -0.47 | -0.22 | B>B |
| chr3 | 86800000 | 86900000 | -0.14 | 0.04 | B>B |
| chr3 | 86900000 | 87000000 | 0.06 | 0.17 | A>A |
| chr3 | 87000000 | 87100000 | 0.22 | 0.30 | A>A |
| chr3 | 87100000 | 87200000 | 0.24 | 0.31 | A>A |
| chr3 | 87200000 | 87300000 | 0.37 | 0.40 | A>A |
| chr3 | 87300000 | 87400000 | 0.34 | 0.42 | A>A |
| chr3 | 87400000 | 87500000 | 0.43 | 0.55 | A>A |
| chr3 | 87500000 | 87600000 | 0.27 | 0.49 | A>A |
| chr3 | 87600000 | 87700000 | 0.25 | 0.40 | A>A |
| chr3 | 87700000 | 87800000 | 0.26 | 0.43 | A>A |
| chr3 | 87800000 | 87900000 | 0.30 | 0.43 | A>A |
| chr3 | 87900000 | 88000000 | 0.09 | 0.29 | A>A |
| chr3 | 88000000 | 88100000 | 0.11 | 0.21 | A>A |
| chr3 | 88100000 | 88200000 | 0.20 | 0.23 | A>A |
| chr3 | 88200000 | 88300000 | -0.33 | -0.41 | B>B |

| | | | | | |
|---|---|---|---|---|---|
| chr3 | 88300000 | 88400000 | -0.59 | -0.71 | B>B |
| chr3 | 88400000 | 88500000 | -0.66 | -0.76 | B>B |
| chr3 | 88500000 | 88600000 | -0.65 | -0.69 | B>B |
| chr3 | 88600000 | 88700000 | -0.63 | -0.75 | B>B |
| chr3 | 88700000 | 88800000 | -0.69 | -0.75 | B>B |
| chr3 | 88800000 | 88900000 | -0.65 | -0.76 | B>B |
| chr3 | 88900000 | 89000000 | -0.67 | -0.83 | B>B |
| chr3 | 89000000 | 89100000 | -0.64 | -0.73 | B>B |
| chr3 | 89100000 | 89200000 | -0.59 | -0.61 | B>B |
| chr3 | 89200000 | 89300000 | -0.60 | -0.71 | B>B |
| chr3 | 89300000 | 89400000 | -0.65 | -0.76 | B>B |
| chr3 | 89400000 | 89500000 | -0.66 | -0.78 | B>B |
| chr3 | 89500000 | 89600000 | -0.69 | -0.76 | B>B |
| chr3 | 89600000 | 89700000 | -0.64 | -0.76 | B>B |
| chr3 | 89700000 | 89800000 | -0.66 | -0.76 | B>B |
| chr3 | 89800000 | 89900000 | -0.69 | -0.76 | B>B |
| chr3 | 89900000 | 90000000 | -0.68 | -0.75 | B>B |
| chr3 | 90000000 | 90100000 | -0.70 | -0.74 | B>B |
| chr3 | 90100000 | 90200000 | -0.68 | -0.65 | B>B |
| chr3 | 90200000 | 90300000 | 0.00 | 0.00 | NO |
| chr3 | 90300000 | 90400000 | 0.00 | 0.00 | NO |
| chr3 | 90400000 | 90500000 | 0.00 | 0.00 | NO |
| chr3 | 90500000 | 90600000 | 0.00 | 0.00 | NO |
| chr3 | 90600000 | 90700000 | 0.00 | 0.00 | NO |
| chr3 | 90700000 | 90800000 | 0.00 | 0.00 | NO |
| chr3 | 90800000 | 90900000 | 0.00 | 0.00 | NO |
| chr3 | 90900000 | 91000000 | 0.00 | 0.00 | NO |
| chr3 | 91000000 | 91100000 | 0.00 | 0.00 | NO |
| chr3 | 91100000 | 91200000 | 0.00 | 0.00 | NO |
| chr3 | 91200000 | 91300000 | 0.00 | 0.00 | NO |
| chr3 | 91300000 | 91400000 | 0.00 | 0.00 | NO |
| chr3 | 91400000 | 91500000 | 0.00 | 0.00 | NO |
| chr3 | 91500000 | 91600000 | 0.00 | 0.00 | NO |
| chr3 | 91600000 | 91700000 | 0.00 | 0.00 | NO |
| chr3 | 91700000 | 91800000 | 0.00 | 0.00 | NO |
| chr3 | 91800000 | 91900000 | 0.00 | 0.00 | NO |
| chr3 | 91900000 | 92000000 | 0.00 | 0.00 | NO |
| chr3 | 92000000 | 92100000 | 0.00 | 0.00 | NO |
| chr3 | 92100000 | 92200000 | 0.00 | 0.00 | NO |
| chr3 | 92200000 | 92300000 | 0.00 | 0.00 | NO |
| chr3 | 92300000 | 92400000 | 0.00 | 0.00 | NO |
| chr3 | 92400000 | 92500000 | 0.00 | 0.00 | NO |
| chr3 | 92500000 | 92600000 | 0.00 | 0.00 | NO |
| chr3 | 92600000 | 92700000 | 0.00 | 0.00 | NO |
| chr3 | 92700000 | 92800000 | 0.00 | 0.00 | NO |
| chr3 | 92800000 | 92900000 | 0.00 | 0.00 | NO |
| chr3 | 92900000 | 93000000 | 0.00 | 0.00 | NO |
| chr3 | 93000000 | 93100000 | 0.00 | 0.00 | NO |
| chr3 | 93100000 | 93200000 | 0.00 | 0.00 | NO |
| chr3 | 93200000 | 93300000 | 0.00 | 0.00 | NO |
| chr3 | 93300000 | 93400000 | 0.00 | 0.00 | NO |
| chr3 | 93400000 | 93500000 | 0.00 | 0.00 | NO |
| chr3 | 93500000 | 93600000 | 0.00 | 0.00 | NO |
| chr3 | 93600000 | 93700000 | 0.00 | 0.16 | NO |
| chr3 | 93700000 | 93800000 | 0.42 | 0.46 | A>A |
| chr3 | 93800000 | 93900000 | 0.05 | 0.48 | A>A |

| | | | | | |
|---|---|---|---|---|---|
| chr3 | 93900000 | 94000000 | -0.36 | 0.52 | B>A |
| chr3 | 94000000 | 94100000 | -0.52 | 0.55 | B>A |
| chr3 | 94100000 | 94200000 | -0.53 | 0.77 | B>A |
| chr3 | 94200000 | 94300000 | -0.41 | 0.67 | B>A |
| chr3 | 94300000 | 94400000 | -0.15 | 0.79 | B>A |
| chr3 | 94400000 | 94500000 | -0.39 | 0.66 | B>A |
| chr3 | 94500000 | 94600000 | -0.58 | 0.55 | B>A |
| chr3 | 94600000 | 94700000 | -0.66 | 0.60 | B>A |
| chr3 | 94700000 | 94800000 | -0.63 | 0.67 | B>A |
| chr3 | 94800000 | 94900000 | -0.66 | 0.72 | B>A |
| chr3 | 94900000 | 95000000 | -0.65 | 0.66 | B>A |
| chr3 | 95000000 | 95100000 | -0.70 | 0.68 | B>A |
| chr3 | 95100000 | 95200000 | -0.70 | 0.73 | B>A |
| chr3 | 95200000 | 95300000 | -0.69 | 0.54 | B>A |
| chr3 | 95300000 | 95400000 | -0.69 | 0.67 | B>A |
| chr3 | 95400000 | 95500000 | -0.67 | 0.58 | B>A |
| chr3 | 95500000 | 95600000 | -0.64 | 0.58 | B>A |
| chr3 | 95600000 | 95700000 | -0.65 | 0.61 | B>A |
| chr3 | 95700000 | 95800000 | -0.71 | 0.61 | B>A |
| chr3 | 95800000 | 95900000 | -0.71 | 0.51 | B>A |
| chr3 | 95900000 | 96000000 | -0.72 | 0.48 | B>A |
| chr3 | 96000000 | 96100000 | -0.74 | 0.42 | B>A |
| chr3 | 96100000 | 96200000 | -0.72 | 0.47 | B>A |
| chr3 | 96200000 | 96300000 | -0.63 | 0.47 | B>A |
| chr3 | 96300000 | 96400000 | -0.72 | 0.39 | B>A |
| chr3 | 96400000 | 96500000 | -0.66 | 0.37 | B>A |
| chr3 | 96500000 | 96600000 | -0.67 | 0.09 | B>A |
| chr3 | 96600000 | 96700000 | -0.74 | 0.06 | B>A |
| chr3 | 96700000 | 96800000 | -0.75 | -0.02 | B>A |
| chr3 | 96800000 | 96900000 | -0.80 | -0.19 | B>A |
| chr3 | 96900000 | 97000000 | -0.76 | -0.35 | B>B |
| chr3 | 97000000 | 97100000 | -0.79 | -0.47 | B>B |
| chr3 | 97100000 | 97200000 | -0.79 | -0.54 | B>B |
| chr3 | 97200000 | 97300000 | -0.74 | -0.49 | B>B |
| chr3 | 97300000 | 97400000 | -0.61 | -0.36 | B>B |
| chr3 | 97400000 | 97500000 | -0.34 | -0.10 | B>B |
| chr3 | 97500000 | 97600000 | 0.18 | 0.10 | A>A |
| chr3 | 97600000 | 97700000 | 0.28 | 0.14 | A>A |
| chr3 | 97700000 | 97800000 | -0.16 | -0.24 | B>B |
| chr3 | 97800000 | 97900000 | -0.54 | -0.92 | B>B |
| chr3 | 97900000 | 98000000 | -0.59 | -0.93 | B>B |
| chr3 | 98000000 | 98100000 | -0.50 | -0.71 | B>B |
| chr3 | 98100000 | 98200000 | -0.57 | -0.73 | B>B |
| chr3 | 98200000 | 98300000 | -0.17 | -0.40 | B>B |
| chr3 | 98300000 | 98400000 | -0.02 | -0.15 | B>B |
| chr3 | 98400000 | 98500000 | 0.08 | -0.12 | B>B |
| chr3 | 98500000 | 98600000 | 0.33 | 0.08 | A>A |
| chr3 | 98600000 | 98700000 | 0.64 | 0.36 | A>A |
| chr3 | 98700000 | 98800000 | -0.20 | -0.43 | B>B |
| chr3 | 98800000 | 98900000 | -0.53 | -0.70 | B>B |
| chr3 | 98900000 | 99000000 | -0.57 | -0.78 | B>B |
| chr3 | 99000000 | 99100000 | -0.67 | -0.92 | B>B |
| chr3 | 99100000 | 99200000 | -0.61 | -0.96 | B>B |
| chr3 | 99200000 | 99300000 | -0.62 | -0.97 | B>B |
| chr3 | 99300000 | 99400000 | -0.55 | -0.77 | B>B |
| chr3 | 99400000 | 99500000 | -0.38 | -0.60 | B>B |

| | | | | | |
|---|---|---|---|---|---|
| chr3 | 99500000 | 99600000 | 0.11 | 0.14 | A>A |
| chr3 | 99600000 | 99700000 | 0.52 | 0.67 | A>A |
| chr3 | 99700000 | 99800000 | 0.56 | 0.55 | A>A |
| chr3 | 99800000 | 99900000 | 0.51 | 0.55 | A>A |
| chr3 | 99900000 | 100000000 | 0.41 | 0.38 | A>A |
| chr3 | 1E+08 | 100100000 | 0.65 | 0.63 | A>A |
| chr3 | 1E+08 | 100200000 | 0.72 | 0.61 | A>A |
| chr3 | 1E+08 | 100300000 | 0.52 | 0.71 | A>A |
| chr3 | 1E+08 | 100400000 | 0.46 | 0.64 | A>A |
| chr3 | 1E+08 | 100500000 | 0.39 | 0.65 | A>A |
| chr3 | 1.01E+08 | 100600000 | 0.21 | 0.57 | A>A |
| chr3 | 1.01E+08 | 100700000 | 0.26 | 0.78 | B>A |
| chr3 | 1.01E+08 | 100800000 | 0.42 | 0.77 | A>A |
| chr3 | 1.01E+08 | 100900000 | 0.41 | 0.82 | A>A |
| chr3 | 1.01E+08 | 101000000 | 0.15 | 0.61 | A>A |
| chr3 | 1.01E+08 | 101100000 | 0.24 | 0.61 | A>A |
| chr3 | 1.01E+08 | 101200000 | 0.41 | 0.67 | A>A |
| chr3 | 1.01E+08 | 101300000 | 0.65 | 0.89 | A>A |
| chr3 | 1.01E+08 | 101400000 | 0.81 | 0.92 | A>A |
| chr3 | 1.01E+08 | 101500000 | 0.62 | 0.78 | A>A |
| chr3 | 1.02E+08 | 101600000 | 0.27 | 0.47 | A>A |
| chr3 | 1.02E+08 | 101700000 | 0.49 | 0.68 | A>A |
| chr3 | 1.02E+08 | 101800000 | 0.50 | 0.74 | A>A |
| chr3 | 1.02E+08 | 101900000 | 0.40 | 0.67 | A>A |
| chr3 | 1.02E+08 | 102000000 | -0.12 | 0.20 | A>A |
| chr3 | 1.02E+08 | 102100000 | -0.53 | -0.07 | B>B |
| chr3 | 1.02E+08 | 102200000 | -0.47 | -0.01 | B>B |
| chr3 | 1.02E+08 | 102300000 | -0.48 | -0.02 | B>B |
| chr3 | 1.02E+08 | 102400000 | -0.69 | -0.19 | B>B |
| chr3 | 1.02E+08 | 102500000 | -0.74 | -0.13 | B>A |
| chr3 | 1.03E+08 | 102600000 | -0.83 | -0.03 | B>A |
| chr3 | 1.03E+08 | 102700000 | -0.88 | 0.11 | B>A |
| chr3 | 1.03E+08 | 102800000 | -0.89 | 0.13 | B>A |
| chr3 | 1.03E+08 | 102900000 | -0.87 | 0.03 | B>A |
| chr3 | 1.03E+08 | 103000000 | -0.84 | 0.03 | B>A |
| chr3 | 1.03E+08 | 103100000 | -0.84 | -0.20 | B>A |
| chr3 | 1.03E+08 | 103200000 | -0.82 | -0.16 | B>A |
| chr3 | 1.03E+08 | 103300000 | -0.79 | -0.31 | B>B |
| chr3 | 1.03E+08 | 103400000 | -0.81 | -0.55 | B>B |
| chr3 | 1.03E+08 | 103500000 | -0.85 | -0.68 | B>B |
| chr3 | 1.04E+08 | 103600000 | -0.83 | -0.74 | B>B |
| chr3 | 1.04E+08 | 103700000 | -0.79 | -0.79 | B>B |
| chr3 | 1.04E+08 | 103800000 | -0.76 | -0.76 | B>B |
| chr3 | 1.04E+08 | 103900000 | -0.66 | -0.70 | B>B |
| chr3 | 1.04E+08 | 104000000 | -0.60 | -0.58 | B>B |
| chr3 | 1.04E+08 | 104100000 | -0.50 | -0.34 | B>B |
| chr3 | 1.04E+08 | 104200000 | -0.48 | -0.38 | B>B |
| chr3 | 1.04E+08 | 104300000 | -0.51 | -0.40 | B>B |
| chr3 | 1.04E+08 | 104400000 | -0.51 | -0.27 | B>B |
| chr3 | 1.04E+08 | 104500000 | -0.61 | -0.34 | B>B |
| chr3 | 1.05E+08 | 104600000 | -0.55 | -0.34 | B>B |
| chr3 | 1.05E+08 | 104700000 | -0.45 | -0.22 | B>B |
| chr3 | 1.05E+08 | 104800000 | -0.44 | -0.14 | B>B |
| chr3 | 1.05E+08 | 104900000 | -0.29 | -0.05 | B>B |
| chr3 | 1.05E+08 | 105000000 | -0.22 | 0.03 | B>B |
| chr3 | 1.05E+08 | 105100000 | -0.18 | -0.05 | B>B |

| | | | | | |
|---|---|---|---|---|---|
| chr3 | 1.05E+08 | 105200000 | 0.20 | 0.17 | A>A |
| chr3 | 1.05E+08 | 105300000 | -0.03 | 0.02 | B>B |
| chr3 | 1.05E+08 | 105400000 | -0.05 | 0.01 | B>B |
| chr3 | 1.05E+08 | 105500000 | 0.22 | 0.21 | A>A |
| chr3 | 1.06E+08 | 105600000 | 0.41 | 0.32 | A>A |
| chr3 | 1.06E+08 | 105700000 | 0.27 | 0.35 | A>A |
| chr3 | 1.06E+08 | 105800000 | 0.22 | 0.40 | A>A |
| chr3 | 1.06E+08 | 105900000 | 0.23 | 0.31 | A>A |
| chr3 | 1.06E+08 | 106000000 | 0.00 | 0.10 | A>A |
| chr3 | 1.06E+08 | 106100000 | -0.02 | 0.04 | A>A |
| chr3 | 1.06E+08 | 106200000 | 0.00 | -0.04 | B>B |
| chr3 | 1.06E+08 | 106300000 | 0.00 | -0.09 | B>B |
| chr3 | 1.06E+08 | 106400000 | -0.28 | -0.28 | B>B |
| chr3 | 1.06E+08 | 106500000 | -0.45 | -0.56 | B>B |
| chr3 | 1.07E+08 | 106600000 | -0.63 | -0.68 | B>B |
| chr3 | 1.07E+08 | 106700000 | -0.68 | -0.69 | B>B |
| chr3 | 1.07E+08 | 106800000 | -0.59 | -0.55 | B>B |
| chr3 | 1.07E+08 | 106900000 | -0.47 | -0.44 | B>B |
| chr3 | 1.07E+08 | 107000000 | -0.05 | -0.14 | B>B |
| chr3 | 1.07E+08 | 107100000 | 0.02 | -0.06 | B>B |
| chr3 | 1.07E+08 | 107200000 | 0.31 | 0.07 | A>A |
| chr3 | 1.07E+08 | 107300000 | 0.50 | 0.25 | A>A |
| chr3 | 1.07E+08 | 107400000 | 0.48 | 0.36 | A>A |
| chr3 | 1.07E+08 | 107500000 | 0.43 | 0.26 | A>A |
| chr3 | 1.08E+08 | 107600000 | 0.38 | 0.18 | A>A |
| chr3 | 1.08E+08 | 107700000 | 0.43 | 0.19 | A>A |
| chr3 | 1.08E+08 | 107800000 | 0.59 | 0.36 | A>A |
| chr3 | 1.08E+08 | 107900000 | 0.58 | 0.23 | A>A |
| chr3 | 1.08E+08 | 108000000 | 0.42 | 0.18 | A>A |
| chr3 | 1.08E+08 | 108100000 | 0.19 | 0.04 | A>A |
| chr3 | 1.08E+08 | 108200000 | 0.24 | -0.01 | A>A |
| chr3 | 1.08E+08 | 108300000 | 0.25 | 0.06 | A>A |
| chr3 | 1.08E+08 | 108400000 | 0.04 | -0.10 | B>B |
| chr3 | 1.08E+08 | 108500000 | -0.39 | -0.47 | B>B |
| chr3 | 1.09E+08 | 108600000 | -0.47 | -0.55 | B>B |
| chr3 | 1.09E+08 | 108700000 | -0.39 | -0.45 | B>B |
| chr3 | 1.09E+08 | 108800000 | -0.43 | -0.44 | B>B |
| chr3 | 1.09E+08 | 108900000 | -0.50 | -0.41 | B>B |
| chr3 | 1.09E+08 | 109000000 | -0.41 | -0.34 | B>B |
| chr3 | 1.09E+08 | 109100000 | -0.26 | -0.06 | B>B |
| chr3 | 1.09E+08 | 109200000 | -0.35 | -0.07 | B>B |
| chr3 | 1.09E+08 | 109300000 | -0.32 | 0.02 | B>B |
| chr3 | 1.09E+08 | 109400000 | -0.16 | 0.21 | A>A |
| chr3 | 1.09E+08 | 109500000 | -0.12 | 0.12 | B>B |
| chr3 | 1.1E+08 | 109600000 | -0.09 | 0.09 | A>A |
| chr3 | 1.1E+08 | 109700000 | 0.07 | 0.13 | A>A |
| chr3 | 1.1E+08 | 109800000 | 0.08 | 0.13 | A>A |
| chr3 | 1.1E+08 | 109900000 | 0.12 | 0.11 | A>A |
| chr3 | 1.1E+08 | 110000000 | 0.28 | 0.19 | A>A |
| chr3 | 1.1E+08 | 110100000 | 0.18 | 0.26 | A>A |
| chr3 | 1.1E+08 | 110200000 | 0.25 | 0.36 | A>A |
| chr3 | 1.1E+08 | 110300000 | 0.30 | 0.41 | A>A |
| chr3 | 1.1E+08 | 110400000 | 0.30 | 0.38 | A>A |
| chr3 | 1.1E+08 | 110500000 | 0.24 | 0.32 | A>A |
| chr3 | 1.11E+08 | 110600000 | 0.26 | 0.34 | A>A |
| chr3 | 1.11E+08 | 110700000 | 0.07 | 0.45 | A>A |

| | | | | | |
|---|---|---|---|---|---|
| chr3 | 1.11E+08 | 110800000 | 0.06 | 0.26 | A>A |
| chr3 | 1.11E+08 | 110900000 | 0.22 | 0.09 | A>A |
| chr3 | 1.11E+08 | 111000000 | -0.04 | -0.11 | B>B |
| chr3 | 1.11E+08 | 111100000 | -0.24 | -0.16 | B>B |
| chr3 | 1.11E+08 | 111200000 | -0.34 | -0.28 | B>B |
| chr3 | 1.11E+08 | 111300000 | -0.17 | -0.10 | B>B |
| chr3 | 1.11E+08 | 111400000 | 0.27 | 0.18 | A>A |
| chr3 | 1.11E+08 | 111500000 | 0.60 | 0.47 | A>A |
| chr3 | 1.12E+08 | 111600000 | 0.73 | 0.60 | A>A |
| chr3 | 1.12E+08 | 111700000 | 0.71 | 0.53 | A>A |
| chr3 | 1.12E+08 | 111800000 | 0.57 | 0.40 | A>A |
| chr3 | 1.12E+08 | 111900000 | 0.21 | 0.01 | A>A |
| chr3 | 1.12E+08 | 112000000 | -0.17 | -0.25 | B>B |
| chr3 | 1.12E+08 | 112100000 | 0.08 | -0.17 | B>B |
| chr3 | 1.12E+08 | 112200000 | 0.00 | -0.29 | B>B |
| chr3 | 1.12E+08 | 112300000 | 0.32 | 0.15 | A>A |
| chr3 | 1.12E+08 | 112400000 | 0.85 | 0.71 | A>A |
| chr3 | 1.12E+08 | 112500000 | 1.01 | 0.92 | A>A |
| chr3 | 1.13E+08 | 112600000 | 1.13 | 0.95 | A>A |
| chr3 | 1.13E+08 | 112700000 | 1.03 | 0.80 | A>A |
| chr3 | 1.13E+08 | 112800000 | 0.74 | 0.68 | A>A |
| chr3 | 1.13E+08 | 112900000 | 1.06 | 0.77 | A>A |
| chr3 | 1.13E+08 | 113000000 | 1.09 | 0.79 | A>A |
| chr3 | 1.13E+08 | 113100000 | 0.78 | 0.76 | A>A |
| chr3 | 1.13E+08 | 113200000 | 0.91 | 0.86 | A>A |
| chr3 | 1.13E+08 | 113300000 | 1.01 | 1.01 | A>A |
| chr3 | 1.13E+08 | 113400000 | 0.83 | 0.80 | A>A |
| chr3 | 1.13E+08 | 113500000 | 1.00 | 0.96 | A>A |
| chr3 | 1.14E+08 | 113600000 | 0.90 | 0.90 | A>A |
| chr3 | 1.14E+08 | 113700000 | 0.79 | 0.83 | A>A |
| chr3 | 1.14E+08 | 113800000 | 0.79 | 0.76 | A>A |
| chr3 | 1.14E+08 | 113900000 | 0.62 | 0.50 | A>A |
| chr3 | 1.14E+08 | 114000000 | 0.29 | 0.42 | A>A |
| chr3 | 1.14E+08 | 114100000 | 0.33 | 0.53 | A>A |
| chr3 | 1.14E+08 | 114200000 | 0.35 | 0.57 | A>A |
| chr3 | 1.14E+08 | 114300000 | 0.47 | 0.60 | A>A |
| chr3 | 1.14E+08 | 114400000 | 0.51 | 0.63 | A>A |
| chr3 | 1.14E+08 | 114500000 | 0.51 | 0.64 | A>A |
| chr3 | 1.15E+08 | 114600000 | 0.31 | 0.54 | A>A |
| chr3 | 1.15E+08 | 114700000 | 0.31 | 0.65 | A>A |
| chr3 | 1.15E+08 | 114800000 | 0.25 | 0.57 | A>A |
| chr3 | 1.15E+08 | 114900000 | 0.19 | 0.41 | A>A |
| chr3 | 1.15E+08 | 115000000 | -0.05 | -0.03 | B>B |
| chr3 | 1.15E+08 | 115100000 | -0.32 | -0.19 | B>B |
| chr3 | 1.15E+08 | 115200000 | -0.40 | -0.19 | B>B |
| chr3 | 1.15E+08 | 115300000 | -0.45 | -0.25 | B>B |
| chr3 | 1.15E+08 | 115400000 | -0.31 | -0.12 | B>B |
| chr3 | 1.15E+08 | 115500000 | -0.13 | 0.00 | B>B |
| chr3 | 1.16E+08 | 115600000 | -0.17 | -0.05 | B>B |
| chr3 | 1.16E+08 | 115700000 | -0.44 | -0.19 | B>B |
| chr3 | 1.16E+08 | 115800000 | -0.27 | -0.23 | B>B |
| chr3 | 1.16E+08 | 115900000 | -0.43 | -0.38 | B>B |
| chr3 | 1.16E+08 | 116000000 | -0.68 | -0.74 | B>B |
| chr3 | 1.16E+08 | 116100000 | -0.72 | -0.85 | B>B |
| chr3 | 1.16E+08 | 116200000 | -0.80 | -0.88 | B>B |
| chr3 | 1.16E+08 | 116300000 | -0.90 | -0.90 | B>B |

| | | | | | |
|---|---|---|---|---|---|
| chr3 | 1.16E+08 | 116400000 | -0.81 | -0.90 | B>B |
| chr3 | 1.16E+08 | 116500000 | -0.64 | -0.62 | B>B |
| chr3 | 1.17E+08 | 116600000 | -0.55 | -0.24 | B>B |
| chr3 | 1.17E+08 | 116700000 | -0.48 | -0.16 | B>B |
| chr3 | 1.17E+08 | 116800000 | -0.11 | -0.03 | B>B |
| chr3 | 1.17E+08 | 116900000 | 0.11 | 0.10 | A>A |
| chr3 | 1.17E+08 | 117000000 | 0.18 | 0.15 | A>A |
| chr3 | 1.17E+08 | 117100000 | 0.04 | 0.03 | A>A |
| chr3 | 1.17E+08 | 117200000 | 0.00 | 0.01 | A>A |
| chr3 | 1.17E+08 | 117300000 | 0.07 | 0.05 | A>A |
| chr3 | 1.17E+08 | 117400000 | 0.07 | 0.08 | A>A |
| chr3 | 1.17E+08 | 117500000 | 0.01 | -0.02 | B>B |
| chr3 | 1.18E+08 | 117600000 | 0.10 | 0.06 | A>A |
| chr3 | 1.18E+08 | 117700000 | 0.00 | -0.05 | B>B |
| chr3 | 1.18E+08 | 117800000 | -0.43 | -0.29 | B>B |
| chr3 | 1.18E+08 | 117900000 | -0.63 | -0.50 | B>B |
| chr3 | 1.18E+08 | 118000000 | -0.77 | -0.70 | B>B |
| chr3 | 1.18E+08 | 118100000 | -0.87 | -0.81 | B>B |
| chr3 | 1.18E+08 | 118200000 | -0.77 | -0.93 | B>B |
| chr3 | 1.18E+08 | 118300000 | -0.73 | -0.85 | B>B |
| chr3 | 1.18E+08 | 118400000 | -0.61 | -0.67 | B>B |
| chr3 | 1.18E+08 | 118500000 | -0.52 | -0.77 | B>B |
| chr3 | 1.19E+08 | 118600000 | -0.21 | -0.40 | B>B |
| chr3 | 1.19E+08 | 118700000 | 0.35 | -0.05 | A>A |
| chr3 | 1.19E+08 | 118800000 | 0.38 | 0.06 | A>A |
| chr3 | 1.19E+08 | 118900000 | 0.51 | 0.24 | A>A |
| chr3 | 1.19E+08 | 119000000 | 0.71 | 0.49 | A>A |
| chr3 | 1.19E+08 | 119100000 | 0.77 | 0.49 | A>A |
| chr3 | 1.19E+08 | 119200000 | 0.82 | 0.48 | A>A |
| chr3 | 1.19E+08 | 119300000 | 0.89 | 0.59 | A>A |
| chr3 | 1.19E+08 | 119400000 | 0.76 | 0.49 | A>A |
| chr3 | 1.19E+08 | 119500000 | 0.63 | 0.31 | A>A |
| chr3 | 1.2E+08 | 119600000 | 0.70 | 0.41 | A>A |
| chr3 | 1.2E+08 | 119700000 | 0.92 | 0.64 | A>A |
| chr3 | 1.2E+08 | 119800000 | 0.97 | 0.68 | A>A |
| chr3 | 1.2E+08 | 119900000 | 0.79 | 0.58 | A>A |
| chr3 | 1.2E+08 | 120000000 | 0.43 | 0.19 | A>A |
| chr3 | 1.2E+08 | 120100000 | 0.58 | 0.17 | A>A |
| chr3 | 1.2E+08 | 120200000 | 0.41 | -0.08 | A>A |
| chr3 | 1.2E+08 | 120300000 | 0.19 | -0.26 | B>B |
| chr3 | 1.2E+08 | 120400000 | 0.42 | -0.02 | A>A |
| chr3 | 1.2E+08 | 120500000 | 0.59 | -0.01 | A>B |
| chr3 | 1.21E+08 | 120600000 | -0.07 | -0.65 | B>B |
| chr3 | 1.21E+08 | 120700000 | -0.44 | -0.94 | B>B |
| chr3 | 1.21E+08 | 120800000 | -0.72 | -1.14 | B>B |
| chr3 | 1.21E+08 | 120900000 | -0.60 | -1.20 | A>B |
| chr3 | 1.21E+08 | 121000000 | -0.71 | -1.15 | B>B |
| chr3 | 1.21E+08 | 121100000 | -0.56 | -1.03 | B>B |
| chr3 | 1.21E+08 | 121200000 | 0.02 | -0.46 | B>B |
| chr3 | 1.21E+08 | 121300000 | 0.44 | 0.07 | A>A |
| chr3 | 1.21E+08 | 121400000 | 0.57 | 0.18 | A>A |
| chr3 | 1.21E+08 | 121500000 | 0.59 | 0.27 | A>A |
| chr3 | 1.22E+08 | 121600000 | 0.43 | 0.00 | A>A |
| chr3 | 1.22E+08 | 121700000 | 0.12 | -0.42 | B>B |
| chr3 | 1.22E+08 | 121800000 | -0.03 | -0.40 | B>B |
| chr3 | 1.22E+08 | 121900000 | -0.15 | -0.54 | B>B |

| | | | | | |
|---|---|---|---|---|---|
| chr3 | 1.22E+08 | 122000000 | 0.07 | -0.43 | B>B |
| chr3 | 1.22E+08 | 122100000 | 0.51 | 0.13 | A>A |
| chr3 | 1.22E+08 | 122200000 | 0.90 | 0.66 | A>A |
| chr3 | 1.22E+08 | 122300000 | 1.09 | 0.97 | A>A |
| chr3 | 1.22E+08 | 122400000 | 1.14 | 0.92 | A>A |
| chr3 | 1.22E+08 | 122500000 | 1.01 | 0.97 | A>A |
| chr3 | 1.23E+08 | 122600000 | 1.04 | 0.98 | A>A |
| chr3 | 1.23E+08 | 122700000 | 1.05 | 0.94 | A>A |
| chr3 | 1.23E+08 | 122800000 | 1.17 | 0.94 | A>A |
| chr3 | 1.23E+08 | 122900000 | 1.16 | 1.21 | A>A |
| chr3 | 1.23E+08 | 123000000 | 1.15 | 1.03 | A>A |
| chr3 | 1.23E+08 | 123100000 | 1.12 | 1.12 | A>A |
| chr3 | 1.23E+08 | 123200000 | 1.29 | 1.40 | A>A |
| chr3 | 1.23E+08 | 123300000 | 1.16 | 1.04 | A>A |
| chr3 | 1.23E+08 | 123400000 | 1.17 | 1.01 | A>A |
| chr3 | 1.23E+08 | 123500000 | 1.16 | 0.96 | A>A |
| chr3 | 1.24E+08 | 123600000 | 1.32 | 1.01 | A>A |
| chr3 | 1.24E+08 | 123700000 | 1.13 | 0.79 | A>A |
| chr3 | 1.24E+08 | 123800000 | 0.86 | 0.43 | A>A |
| chr3 | 1.24E+08 | 123900000 | 0.89 | 0.38 | A>A |
| chr3 | 1.24E+08 | 124000000 | 0.95 | 0.38 | A>A |
| chr3 | 1.24E+08 | 124100000 | 0.81 | 0.37 | A>A |
| chr3 | 1.24E+08 | 124200000 | 0.90 | 0.33 | A>A |
| chr3 | 1.24E+08 | 124300000 | 0.78 | 0.47 | A>A |
| chr3 | 1.24E+08 | 124400000 | 1.08 | 0.68 | A>A |
| chr3 | 1.24E+08 | 124500000 | 1.01 | 0.71 | A>A |
| chr3 | 1.25E+08 | 124600000 | 1.42 | 1.22 | A>A |
| chr3 | 1.25E+08 | 124700000 | 0.91 | 0.81 | A>A |
| chr3 | 1.25E+08 | 124800000 | 1.10 | 1.11 | A>A |
| chr3 | 1.25E+08 | 124900000 | 1.29 | 1.24 | A>A |
| chr3 | 1.25E+08 | 125000000 | 1.17 | 1.13 | A>A |
| chr3 | 1.25E+08 | 125100000 | 1.31 | 1.33 | A>A |
| chr3 | 1.25E+08 | 125200000 | 1.09 | 1.07 | A>A |
| chr3 | 1.25E+08 | 125300000 | 1.25 | 1.18 | A>A |
| chr3 | 1.25E+08 | 125400000 | 0.00 | 1.11 | NO |
| chr3 | 1.25E+08 | 125500000 | 0.00 | 0.00 | NO |
| chr3 | 1.26E+08 | 125600000 | 0.00 | 0.00 | NO |
| chr3 | 1.26E+08 | 125700000 | 0.00 | 0.00 | NO |
| chr3 | 1.26E+08 | 125800000 | 0.00 | 1.12 | NO |
| chr3 | 1.26E+08 | 125900000 | 1.04 | 1.01 | A>A |
| chr3 | 1.26E+08 | 126000000 | 1.14 | 1.06 | A>A |
| chr3 | 1.26E+08 | 126100000 | 0.00 | 1.12 | NO |
| chr3 | 1.26E+08 | 126200000 | 1.07 | 1.07 | A>A |
| chr3 | 1.26E+08 | 126300000 | 1.13 | 1.07 | A>A |
| chr3 | 1.26E+08 | 126400000 | 0.90 | 0.92 | A>A |
| chr3 | 1.26E+08 | 126500000 | 0.81 | 0.92 | A>A |
| chr3 | 1.27E+08 | 126600000 | 1.19 | 1.15 | A>A |
| chr3 | 1.27E+08 | 126700000 | 1.00 | 1.06 | A>A |
| chr3 | 1.27E+08 | 126800000 | 0.92 | 1.02 | A>A |
| chr3 | 1.27E+08 | 126900000 | 0.93 | 1.03 | A>A |
| chr3 | 1.27E+08 | 127000000 | 0.85 | 0.89 | A>A |
| chr3 | 1.27E+08 | 127100000 | 0.88 | 0.98 | A>A |
| chr3 | 1.27E+08 | 127200000 | 0.70 | 0.84 | A>A |
| chr3 | 1.27E+08 | 127300000 | 0.85 | 0.94 | A>A |
| chr3 | 1.27E+08 | 127400000 | 1.05 | 1.11 | A>A |
| chr3 | 1.27E+08 | 127500000 | 1.35 | 1.28 | A>A |

| | | | | | |
|---|---|---|---|---|---|
| chr3 | 1.28E+08 | 127600000 | 1.30 | 1.29 | A>A |
| chr3 | 1.28E+08 | 127700000 | 1.09 | 1.14 | A>A |
| chr3 | 1.28E+08 | 127800000 | 1.03 | 1.19 | A>A |
| chr3 | 1.28E+08 | 127900000 | 1.09 | 1.14 | A>A |
| chr3 | 1.28E+08 | 128000000 | 1.18 | 1.17 | A>A |
| chr3 | 1.28E+08 | 128100000 | 1.24 | 1.33 | A>A |
| chr3 | 1.28E+08 | 128200000 | 1.23 | 1.33 | A>A |
| chr3 | 1.28E+08 | 128300000 | 1.16 | 1.13 | A>A |
| chr3 | 1.28E+08 | 128400000 | 1.21 | 1.20 | A>A |
| chr3 | 1.28E+08 | 128500000 | 1.55 | 1.51 | A>A |
| chr3 | 1.29E+08 | 128600000 | 1.50 | 1.32 | A>A |
| chr3 | 1.29E+08 | 128700000 | 0.00 | 1.17 | NO |
| chr3 | 1.29E+08 | 128800000 | 1.37 | 1.37 | A>A |
| chr3 | 1.29E+08 | 128900000 | 0.00 | 1.39 | NO |
| chr3 | 1.29E+08 | 129000000 | 1.40 | 1.33 | A>A |
| chr3 | 1.29E+08 | 129100000 | 1.36 | 1.18 | A>A |
| chr3 | 1.29E+08 | 129200000 | 1.42 | 1.20 | A>A |
| chr3 | 1.29E+08 | 129300000 | 1.41 | 1.33 | A>A |
| chr3 | 1.29E+08 | 129400000 | 1.23 | 1.13 | A>A |
| chr3 | 1.29E+08 | 129500000 | 1.46 | 1.38 | A>A |
| chr3 | 1.3E+08 | 129600000 | 1.12 | 1.10 | A>A |
| chr3 | 1.3E+08 | 129700000 | 1.02 | 0.95 | A>A |
| chr3 | 1.3E+08 | 129800000 | 0.00 | 0.00 | NO |
| chr3 | 1.3E+08 | 129900000 | 0.00 | 0.00 | NO |
| chr3 | 1.3E+08 | 130000000 | 0.00 | -0.33 | NO |
| chr3 | 1.3E+08 | 130100000 | -0.11 | -0.43 | B>B |
| chr3 | 1.3E+08 | 130200000 | -0.05 | -0.47 | B>B |
| chr3 | 1.3E+08 | 130300000 | -0.01 | -0.43 | B>B |
| chr3 | 1.3E+08 | 130400000 | 0.11 | -0.34 | B>B |
| chr3 | 1.3E+08 | 130500000 | 0.48 | 0.13 | A>A |
| chr3 | 1.31E+08 | 130600000 | 0.00 | -0.05 | NO |
| chr3 | 1.31E+08 | 130700000 | 0.70 | 0.38 | A>A |
| chr3 | 1.31E+08 | 130800000 | 0.61 | 0.41 | A>A |
| chr3 | 1.31E+08 | 130900000 | 0.56 | 0.47 | A>A |
| chr3 | 1.31E+08 | 131000000 | 0.00 | 0.50 | NO |
| chr3 | 1.31E+08 | 131100000 | 0.60 | 0.55 | A>A |
| chr3 | 1.31E+08 | 131200000 | 0.62 | 0.48 | A>A |
| chr3 | 1.31E+08 | 131300000 | 0.64 | 0.43 | A>A |
| chr3 | 1.31E+08 | 131400000 | 0.51 | 0.27 | A>A |
| chr3 | 1.31E+08 | 131500000 | 0.45 | 0.22 | A>A |
| chr3 | 1.32E+08 | 131600000 | 0.42 | 0.25 | A>A |
| chr3 | 1.32E+08 | 131700000 | 0.61 | 0.33 | A>A |
| chr3 | 1.32E+08 | 131800000 | 0.62 | 0.45 | A>A |
| chr3 | 1.32E+08 | 131900000 | 0.00 | 0.69 | NO |
| chr3 | 1.32E+08 | 132000000 | 0.56 | 0.39 | A>A |
| chr3 | 1.32E+08 | 132100000 | 0.00 | 0.44 | NO |
| chr3 | 1.32E+08 | 132200000 | 1.00 | 0.55 | A>A |
| chr3 | 1.32E+08 | 132300000 | 0.91 | 0.54 | A>A |
| chr3 | 1.32E+08 | 132400000 | 0.00 | 0.47 | NO |
| chr3 | 1.32E+08 | 132500000 | 0.07 | -0.18 | B>B |
| chr3 | 1.33E+08 | 132600000 | -0.45 | -0.91 | B>B |
| chr3 | 1.33E+08 | 132700000 | -0.54 | -0.86 | B>B |
| chr3 | 1.33E+08 | 132800000 | -0.29 | -0.80 | B>B |
| chr3 | 1.33E+08 | 132900000 | -0.38 | -0.90 | B>B |
| chr3 | 1.33E+08 | 133000000 | -0.41 | -0.85 | B>B |
| chr3 | 1.33E+08 | 133100000 | -0.31 | -0.70 | B>B |

| | | | | | |
|---|---|---|---|---|---|
| chr3 | 1.33E+08 | 133200000 | 0.21 | -0.09 | A>A |
| chr3 | 1.33E+08 | 133300000 | 0.84 | 0.47 | A>A |
| chr3 | 1.33E+08 | 133400000 | 0.75 | 0.34 | A>A |
| chr3 | 1.33E+08 | 133500000 | 0.00 | 0.29 | NO |
| chr3 | 1.34E+08 | 133600000 | 0.67 | 0.26 | A>A |
| chr3 | 1.34E+08 | 133700000 | 0.64 | 0.21 | A>A |
| chr3 | 1.34E+08 | 133800000 | 0.61 | 0.20 | A>A |
| chr3 | 1.34E+08 | 133900000 | 0.56 | 0.28 | A>A |
| chr3 | 1.34E+08 | 134000000 | 0.79 | 0.49 | A>A |
| chr3 | 1.34E+08 | 134100000 | 1.10 | 0.77 | A>A |
| chr3 | 1.34E+08 | 134200000 | 1.05 | 0.67 | A>A |
| chr3 | 1.34E+08 | 134300000 | 0.45 | 0.08 | A>A |
| chr3 | 1.34E+08 | 134400000 | 0.00 | -0.33 | B>B |
| chr3 | 1.34E+08 | 134500000 | -0.09 | -0.61 | B>B |
| chr3 | 1.35E+08 | 134600000 | -0.18 | -0.60 | B>B |
| chr3 | 1.35E+08 | 134700000 | -0.23 | -0.73 | B>B |
| chr3 | 1.35E+08 | 134800000 | -0.26 | -0.76 | B>B |
| chr3 | 1.35E+08 | 134900000 | -0.39 | -0.79 | B>B |
| chr3 | 1.35E+08 | 135000000 | -0.46 | -0.79 | B>B |
| chr3 | 1.35E+08 | 135100000 | -0.55 | -0.77 | B>B |
| chr3 | 1.35E+08 | 135200000 | -0.50 | -0.73 | B>B |
| chr3 | 1.35E+08 | 135300000 | -0.47 | -0.74 | B>B |
| chr3 | 1.35E+08 | 135400000 | 0.00 | -0.81 | NO |
| chr3 | 1.35E+08 | 135500000 | -0.50 | -0.77 | B>B |
| chr3 | 1.36E+08 | 135600000 | -0.46 | -0.72 | B>B |
| chr3 | 1.36E+08 | 135700000 | -0.16 | -0.43 | B>B |
| chr3 | 1.36E+08 | 135800000 | -0.01 | -0.14 | B>B |
| chr3 | 1.36E+08 | 135900000 | 0.57 | 0.39 | A>A |
| chr3 | 1.36E+08 | 136000000 | 0.74 | 0.56 | A>A |
| chr3 | 1.36E+08 | 136100000 | 0.61 | 0.38 | A>A |
| chr3 | 1.36E+08 | 136200000 | 0.61 | 0.38 | A>A |
| chr3 | 1.36E+08 | 136300000 | 0.65 | 0.53 | A>A |
| chr3 | 1.36E+08 | 136400000 | 0.00 | 0.70 | NO |
| chr3 | 1.36E+08 | 136500000 | 0.55 | 0.45 | A>A |
| chr3 | 1.37E+08 | 136600000 | 0.66 | 0.50 | A>A |
| chr3 | 1.37E+08 | 136700000 | 0.00 | 0.45 | NO |
| chr3 | 1.37E+08 | 136800000 | 0.06 | -0.11 | B>B |
| chr3 | 1.37E+08 | 136900000 | -0.33 | -0.59 | B>B |
| chr3 | 1.37E+08 | 137000000 | -0.57 | -0.86 | B>B |
| chr3 | 1.37E+08 | 137100000 | -0.74 | -0.86 | B>B |
| chr3 | 1.37E+08 | 137200000 | 0.00 | -0.86 | NO |
| chr3 | 1.37E+08 | 137300000 | -0.65 | -0.89 | B>B |
| chr3 | 1.37E+08 | 137400000 | 0.00 | -0.84 | NO |
| chr3 | 1.37E+08 | 137500000 | -0.46 | -0.79 | B>B |
| chr3 | 1.38E+08 | 137600000 | -0.41 | -0.77 | B>B |
| chr3 | 1.38E+08 | 137700000 | -0.37 | -0.53 | B>B |
| chr3 | 1.38E+08 | 137800000 | 0.14 | -0.09 | A>A |
| chr3 | 1.38E+08 | 137900000 | 0.63 | 0.37 | A>A |
| chr3 | 1.38E+08 | 138000000 | 0.93 | 0.71 | A>A |
| chr3 | 1.38E+08 | 138100000 | 0.90 | 0.76 | A>A |
| chr3 | 1.38E+08 | 138200000 | 0.85 | 0.79 | A>A |
| chr3 | 1.38E+08 | 138300000 | 0.00 | 0.83 | NO |
| chr3 | 1.38E+08 | 138400000 | 0.97 | 0.84 | A>A |
| chr3 | 1.38E+08 | 138500000 | 1.19 | 1.02 | A>A |
| chr3 | 1.39E+08 | 138600000 | 1.17 | 1.13 | A>A |
| chr3 | 1.39E+08 | 138700000 | 0.74 | 0.53 | A>A |

| | | | | | |
|---|---|---|---|---|---|
| chr3 | 1.39E+08 | 138800000 | 0.52 | 0.37 | A>A |
| chr3 | 1.39E+08 | 138900000 | 0.58 | 0.33 | A>A |
| chr3 | 1.39E+08 | 139000000 | 0.73 | 0.40 | A>A |
| chr3 | 1.39E+08 | 139100000 | 0.51 | 0.37 | A>A |
| chr3 | 1.39E+08 | 139200000 | 0.06 | -0.16 | B>B |
| chr3 | 1.39E+08 | 139300000 | -0.06 | -0.29 | B>B |
| chr3 | 1.39E+08 | 139400000 | -0.18 | -0.40 | B>B |
| chr3 | 1.39E+08 | 139500000 | -0.27 | -0.70 | B>B |
| chr3 | 1.4E+08 | 139600000 | -0.47 | -0.77 | B>B |
| chr3 | 1.4E+08 | 139700000 | -0.40 | -0.72 | B>B |
| chr3 | 1.4E+08 | 139800000 | -0.40 | -0.72 | B>B |
| chr3 | 1.4E+08 | 139900000 | -0.42 | -0.82 | B>B |
| chr3 | 1.4E+08 | 140000000 | -0.46 | -0.77 | B>B |
| chr3 | 1.4E+08 | 140100000 | -0.36 | -0.80 | B>B |
| chr3 | 1.4E+08 | 140200000 | -0.30 | -0.87 | B>B |
| chr3 | 1.4E+08 | 140300000 | -0.40 | -0.79 | B>B |
| chr3 | 1.4E+08 | 140400000 | -0.48 | -0.81 | B>B |
| chr3 | 1.4E+08 | 140500000 | -0.42 | -0.79 | B>B |
| chr3 | 1.41E+08 | 140600000 | -0.30 | -0.64 | B>B |
| chr3 | 1.41E+08 | 140700000 | 0.17 | -0.24 | B>B |
| chr3 | 1.41E+08 | 140800000 | 0.41 | -0.23 | A>B |
| chr3 | 1.41E+08 | 140900000 | 0.27 | -0.27 | A>A |
| chr3 | 1.41E+08 | 141000000 | 0.80 | 0.06 | A>B |
| chr3 | 1.41E+08 | 141100000 | 1.27 | 0.71 | A>A |
| chr3 | 1.41E+08 | 141200000 | 1.43 | 0.88 | A>A |
| chr3 | 1.41E+08 | 141300000 | 1.39 | 0.90 | A>A |
| chr3 | 1.41E+08 | 141400000 | 1.37 | 0.99 | A>A |
| chr3 | 1.41E+08 | 141500000 | 1.50 | 1.16 | A>A |
| chr3 | 1.42E+08 | 141600000 | 1.63 | 1.26 | A>A |
| chr3 | 1.42E+08 | 141700000 | 1.34 | 1.18 | A>A |
| chr3 | 1.42E+08 | 141800000 | 1.47 | 1.39 | A>A |
| chr3 | 1.42E+08 | 141900000 | 1.41 | 1.37 | A>A |
| chr3 | 1.42E+08 | 142000000 | 1.58 | 1.36 | A>A |
| chr3 | 1.42E+08 | 142100000 | 1.45 | 1.37 | A>A |
| chr3 | 1.42E+08 | 142200000 | 1.48 | 1.41 | A>A |
| chr3 | 1.42E+08 | 142300000 | 1.50 | 1.23 | A>A |
| chr3 | 1.42E+08 | 142400000 | 1.57 | 1.38 | A>A |
| chr3 | 1.42E+08 | 142500000 | 1.47 | 1.21 | A>A |
| chr3 | 1.43E+08 | 142600000 | 1.35 | 0.97 | A>A |
| chr3 | 1.43E+08 | 142700000 | 1.44 | 1.08 | A>A |
| chr3 | 1.43E+08 | 142800000 | 1.02 | 0.35 | A>B |
| chr3 | 1.43E+08 | 142900000 | 0.39 | -0.86 | A>B |
| chr3 | 1.43E+08 | 143000000 | -0.01 | -1.32 | A>B |
| chr3 | 1.43E+08 | 143100000 | -0.24 | -1.34 | A>B |
| chr3 | 1.43E+08 | 143200000 | -0.23 | -1.14 | A>B |
| chr3 | 1.43E+08 | 143300000 | -0.39 | -1.23 | A>B |
| chr3 | 1.43E+08 | 143400000 | -0.45 | -1.22 | A>B |
| chr3 | 1.43E+08 | 143500000 | -0.61 | -1.20 | A>B |
| chr3 | 1.44E+08 | 143600000 | -0.48 | -0.98 | B>B |
| chr3 | 1.44E+08 | 143700000 | -0.37 | -0.95 | A>B |
| chr3 | 1.44E+08 | 143800000 | -0.46 | -1.15 | A>B |
| chr3 | 1.44E+08 | 143900000 | -0.79 | -1.24 | B>B |
| chr3 | 1.44E+08 | 144000000 | -1.02 | -1.29 | B>B |
| chr3 | 1.44E+08 | 144100000 | -1.01 | -1.33 | B>B |
| chr3 | 1.44E+08 | 144200000 | -0.97 | -1.30 | B>B |
| chr3 | 1.44E+08 | 144300000 | -1.02 | -1.25 | B>B |

| | | | | | |
|---|---|---|---|---|---|
| chr3 | 1.44E+08 | 144400000 | -0.96 | -1.29 | B>B |
| chr3 | 1.44E+08 | 144500000 | -0.94 | -1.22 | B>B |
| chr3 | 1.45E+08 | 144600000 | -0.94 | -1.19 | B>B |
| chr3 | 1.45E+08 | 144700000 | -0.98 | -1.17 | B>B |
| chr3 | 1.45E+08 | 144800000 | -0.97 | -1.17 | B>B |
| chr3 | 1.45E+08 | 144900000 | -0.91 | -1.00 | B>B |
| chr3 | 1.45E+08 | 145000000 | -0.94 | -1.01 | B>B |
| chr3 | 1.45E+08 | 145100000 | -0.94 | -1.12 | B>B |
| chr3 | 1.45E+08 | 145200000 | -0.95 | -1.13 | B>B |
| chr3 | 1.45E+08 | 145300000 | -0.86 | -1.22 | B>B |
| chr3 | 1.45E+08 | 145400000 | -0.84 | -1.10 | B>B |
| chr3 | 1.45E+08 | 145500000 | -0.66 | -1.15 | B>B |
| chr3 | 1.46E+08 | 145600000 | -0.66 | -1.18 | B>B |
| chr3 | 1.46E+08 | 145700000 | -0.55 | -1.12 | B>B |
| chr3 | 1.46E+08 | 145800000 | -0.02 | -0.60 | B>B |
| chr3 | 1.46E+08 | 145900000 | 0.57 | 0.12 | A>A |
| chr3 | 1.46E+08 | 146000000 | 0.40 | 0.19 | A>A |
| chr3 | 1.46E+08 | 146100000 | 0.47 | 0.24 | A>A |
| chr3 | 1.46E+08 | 146200000 | 0.68 | 0.43 | A>A |
| chr3 | 1.46E+08 | 146300000 | 0.57 | 0.32 | A>A |
| chr3 | 1.46E+08 | 146400000 | -0.24 | -0.42 | B>B |
| chr3 | 1.46E+08 | 146500000 | -0.78 | -1.05 | B>B |
| chr3 | 1.47E+08 | 146600000 | -0.75 | -1.10 | B>B |
| chr3 | 1.47E+08 | 146700000 | -0.93 | -1.20 | B>B |
| chr3 | 1.47E+08 | 146800000 | -1.01 | -1.10 | B>B |
| chr3 | 1.47E+08 | 146900000 | -0.97 | -1.07 | B>B |
| chr3 | 1.47E+08 | 147000000 | -0.88 | -1.06 | B>B |
| chr3 | 1.47E+08 | 147100000 | -0.67 | -1.07 | B>B |
| chr3 | 1.47E+08 | 147200000 | -0.75 | -1.07 | B>B |
| chr3 | 1.47E+08 | 147300000 | -0.97 | -1.07 | B>B |
| chr3 | 1.47E+08 | 147400000 | -0.97 | -1.11 | B>B |
| chr3 | 1.47E+08 | 147500000 | -1.00 | -1.14 | B>B |
| chr3 | 1.48E+08 | 147600000 | -0.97 | -1.10 | B>B |
| chr3 | 1.48E+08 | 147700000 | -0.90 | -1.06 | B>B |
| chr3 | 1.48E+08 | 147800000 | -0.84 | -1.09 | B>B |
| chr3 | 1.48E+08 | 147900000 | -0.89 | -1.14 | B>B |
| chr3 | 1.48E+08 | 148000000 | -0.83 | -1.15 | B>B |
| chr3 | 1.48E+08 | 148100000 | -0.52 | -0.77 | B>B |
| chr3 | 1.48E+08 | 148200000 | -0.03 | -0.46 | B>B |
| chr3 | 1.48E+08 | 148300000 | 0.20 | -0.17 | A>A |
| chr3 | 1.48E+08 | 148400000 | 0.15 | -0.09 | A>A |
| chr3 | 1.48E+08 | 148500000 | -0.01 | -0.23 | B>B |
| chr3 | 1.49E+08 | 148600000 | 0.02 | -0.23 | B>B |
| chr3 | 1.49E+08 | 148700000 | 0.25 | -0.05 | A>A |
| chr3 | 1.49E+08 | 148800000 | 0.82 | 0.61 | A>A |
| chr3 | 1.49E+08 | 148900000 | 0.98 | 0.88 | A>A |
| chr3 | 1.49E+08 | 149000000 | 1.13 | 0.97 | A>A |
| chr3 | 1.49E+08 | 149100000 | 1.08 | 0.96 | A>A |
| chr3 | 1.49E+08 | 149200000 | 1.29 | 1.14 | A>A |
| chr3 | 1.49E+08 | 149300000 | 1.07 | 1.01 | A>A |
| chr3 | 1.49E+08 | 149400000 | 1.37 | 1.38 | A>A |
| chr3 | 1.49E+08 | 149500000 | 1.12 | 1.07 | A>A |
| chr3 | 1.5E+08 | 149600000 | 1.29 | 1.20 | A>A |
| chr3 | 1.5E+08 | 149700000 | 1.02 | 1.07 | A>A |
| chr3 | 1.5E+08 | 149800000 | 1.35 | 1.19 | A>A |
| chr3 | 1.5E+08 | 149900000 | 1.31 | 1.23 | A>A |

| | | | | | |
|---|---|---|---|---|---|
| chr3 | 1.5E+08 | 150000000 | 1.01 | 0.95 | A>A |
| chr3 | 1.5E+08 | 150100000 | 1.53 | 1.23 | A>A |
| chr3 | 1.5E+08 | 150200000 | 1.33 | 1.17 | A>A |
| chr3 | 1.5E+08 | 150300000 | 1.06 | 0.99 | A>A |
| chr3 | 1.5E+08 | 150400000 | 1.21 | 0.84 | A>A |
| chr3 | 1.5E+08 | 150500000 | 1.13 | 0.65 | A>A |
| chr3 | 1.51E+08 | 150600000 | 0.97 | 0.41 | A>A |
| chr3 | 1.51E+08 | 150700000 | 0.80 | 0.15 | A>B |
| chr3 | 1.51E+08 | 150800000 | 0.76 | 0.18 | A>A |
| chr3 | 1.51E+08 | 150900000 | 0.71 | 0.30 | A>A |
| chr3 | 1.51E+08 | 151000000 | 0.64 | 0.43 | A>A |
| chr3 | 1.51E+08 | 151100000 | 0.70 | 0.53 | A>A |
| chr3 | 1.51E+08 | 151200000 | 0.72 | 0.52 | A>A |
| chr3 | 1.51E+08 | 151300000 | 0.98 | 0.61 | A>A |
| chr3 | 1.51E+08 | 151400000 | 1.03 | 0.65 | A>A |
| chr3 | 1.51E+08 | 151500000 | 1.01 | 0.74 | A>A |
| chr3 | 1.52E+08 | 151600000 | 0.81 | 0.73 | A>A |
| chr3 | 1.52E+08 | 151700000 | 0.79 | 0.68 | A>A |
| chr3 | 1.52E+08 | 151800000 | 0.78 | 0.72 | A>A |
| chr3 | 1.52E+08 | 151900000 | 0.85 | 0.77 | A>A |
| chr3 | 1.52E+08 | 152000000 | 0.92 | 0.83 | A>A |
| chr3 | 1.52E+08 | 152100000 | 0.89 | 0.85 | A>A |
| chr3 | 1.52E+08 | 152200000 | 0.60 | 0.50 | A>A |
| chr3 | 1.52E+08 | 152300000 | 0.29 | 0.01 | A>A |
| chr3 | 1.52E+08 | 152400000 | -0.07 | -0.43 | B>B |
| chr3 | 1.52E+08 | 152500000 | -0.29 | -0.60 | B>B |
| chr3 | 1.53E+08 | 152600000 | -0.08 | -0.42 | B>B |
| chr3 | 1.53E+08 | 152700000 | 0.18 | -0.04 | A>A |
| chr3 | 1.53E+08 | 152800000 | 0.10 | 0.27 | A>A |
| chr3 | 1.53E+08 | 152900000 | 0.46 | 0.56 | A>A |
| chr3 | 1.53E+08 | 153000000 | 0.26 | 0.32 | A>A |
| chr3 | 1.53E+08 | 153100000 | -0.04 | 0.13 | A>A |
| chr3 | 1.53E+08 | 153200000 | -0.38 | -0.09 | B>B |
| chr3 | 1.53E+08 | 153300000 | -0.62 | -0.05 | B>A |
| chr3 | 1.53E+08 | 153400000 | -0.57 | 0.14 | B>A |
| chr3 | 1.53E+08 | 153500000 | -0.64 | 0.29 | B>A |
| chr3 | 1.54E+08 | 153600000 | -0.68 | 0.00 | B>A |
| chr3 | 1.54E+08 | 153700000 | -0.62 | -0.08 | B>A |
| chr3 | 1.54E+08 | 153800000 | -0.27 | -0.16 | B>B |
| chr3 | 1.54E+08 | 153900000 | 0.15 | -0.06 | A>A |
| chr3 | 1.54E+08 | 154000000 | 0.12 | -0.04 | A>A |
| chr3 | 1.54E+08 | 154100000 | -0.27 | -0.59 | B>B |
| chr3 | 1.54E+08 | 154200000 | -0.79 | -1.22 | B>B |
| chr3 | 1.54E+08 | 154300000 | -0.98 | -1.20 | B>B |
| chr3 | 1.54E+08 | 154400000 | -0.92 | -1.21 | B>B |
| chr3 | 1.54E+08 | 154500000 | -0.78 | -1.10 | B>B |
| chr3 | 1.55E+08 | 154600000 | -0.89 | -1.16 | B>B |
| chr3 | 1.55E+08 | 154700000 | -0.83 | -1.17 | B>B |
| chr3 | 1.55E+08 | 154800000 | -0.68 | -1.11 | B>B |
| chr3 | 1.55E+08 | 154900000 | -0.54 | -1.10 | B>B |
| chr3 | 1.55E+08 | 155000000 | -0.61 | -1.09 | B>B |
| chr3 | 1.55E+08 | 155100000 | -0.17 | -0.43 | B>B |
| chr3 | 1.55E+08 | 155200000 | 0.23 | 0.20 | A>A |
| chr3 | 1.55E+08 | 155300000 | 0.48 | 0.47 | A>A |
| chr3 | 1.55E+08 | 155400000 | 0.50 | 0.40 | A>A |
| chr3 | 1.55E+08 | 155500000 | 0.50 | 0.32 | A>A |

| | | | | | |
|---|---|---|---|---|---|
| chr3 | 1.56E+08 | 155600000 | 0.84 | 0.53 | A>A |
| chr3 | 1.56E+08 | 155700000 | 0.63 | 0.22 | A>A |
| chr3 | 1.56E+08 | 155800000 | 0.24 | -0.38 | A>B |
| chr3 | 1.56E+08 | 155900000 | 0.03 | -0.49 | B>B |
| chr3 | 1.56E+08 | 156000000 | -0.17 | -0.75 | B>B |
| chr3 | 1.56E+08 | 156100000 | -0.06 | -0.69 | A>B |
| chr3 | 1.56E+08 | 156200000 | -0.10 | -0.63 | B>B |
| chr3 | 1.56E+08 | 156300000 | 0.12 | -0.24 | B>B |
| chr3 | 1.56E+08 | 156400000 | 0.42 | -0.04 | A>A |
| chr3 | 1.56E+08 | 156500000 | 0.64 | 0.32 | A>A |
| chr3 | 1.57E+08 | 156600000 | 0.53 | 0.18 | A>A |
| chr3 | 1.57E+08 | 156700000 | 0.19 | -0.30 | B>B |
| chr3 | 1.57E+08 | 156800000 | 0.15 | -0.28 | B>B |
| chr3 | 1.57E+08 | 156900000 | 0.44 | 0.13 | A>A |
| chr3 | 1.57E+08 | 157000000 | 0.48 | 0.08 | A>A |
| chr3 | 1.57E+08 | 157100000 | 0.60 | 0.11 | A>A |
| chr3 | 1.57E+08 | 157200000 | 0.43 | -0.04 | A>A |
| chr3 | 1.57E+08 | 157300000 | 0.16 | -0.16 | B>B |
| chr3 | 1.57E+08 | 157400000 | -0.21 | -0.53 | B>B |
| chr3 | 1.57E+08 | 157500000 | -0.39 | -0.74 | B>B |
| chr3 | 1.58E+08 | 157600000 | -0.15 | -0.66 | B>B |
| chr3 | 1.58E+08 | 157700000 | -0.36 | -0.82 | B>B |
| chr3 | 1.58E+08 | 157800000 | -0.36 | -0.66 | B>B |
| chr3 | 1.58E+08 | 157900000 | 0.32 | 0.05 | A>A |
| chr3 | 1.58E+08 | 158000000 | 0.34 | 0.03 | A>A |
| chr3 | 1.58E+08 | 158100000 | 0.18 | -0.09 | A>A |
| chr3 | 1.58E+08 | 158200000 | 0.12 | -0.23 | B>B |
| chr3 | 1.58E+08 | 158300000 | 0.12 | -0.12 | B>B |
| chr3 | 1.58E+08 | 158400000 | 0.40 | 0.10 | A>A |
| chr3 | 1.58E+08 | 158500000 | 0.65 | 0.37 | A>A |
| chr3 | 1.59E+08 | 158600000 | -0.02 | -0.36 | B>B |
| chr3 | 1.59E+08 | 158700000 | -0.46 | -0.93 | B>B |
| chr3 | 1.59E+08 | 158800000 | -0.46 | -1.03 | B>B |
| chr3 | 1.59E+08 | 158900000 | -0.18 | -0.83 | A>B |
| chr3 | 1.59E+08 | 159000000 | -0.21 | -0.59 | B>B |
| chr3 | 1.59E+08 | 159100000 | -0.34 | -0.53 | B>B |
| chr3 | 1.59E+08 | 159200000 | -0.31 | -0.41 | B>B |
| chr3 | 1.59E+08 | 159300000 | -0.17 | -0.45 | B>B |
| chr3 | 1.59E+08 | 159400000 | 0.50 | 0.21 | A>A |
| chr3 | 1.59E+08 | 159500000 | 0.71 | 0.34 | A>A |
| chr3 | 1.6E+08 | 159600000 | 0.50 | 0.36 | A>A |
| chr3 | 1.6E+08 | 159700000 | 0.67 | 0.48 | A>A |
| chr3 | 1.6E+08 | 159800000 | 0.56 | 0.49 | A>A |
| chr3 | 1.6E+08 | 159900000 | 0.78 | 0.69 | A>A |
| chr3 | 1.6E+08 | 160000000 | 0.89 | 0.75 | A>A |
| chr3 | 1.6E+08 | 160100000 | 0.91 | 0.82 | A>A |
| chr3 | 1.6E+08 | 160200000 | 1.08 | 0.86 | A>A |
| chr3 | 1.6E+08 | 160300000 | 0.78 | 0.72 | A>A |
| chr3 | 1.6E+08 | 160400000 | 0.52 | 0.51 | A>A |
| chr3 | 1.6E+08 | 160500000 | 0.29 | 0.31 | A>A |
| chr3 | 1.61E+08 | 160600000 | 0.03 | 0.16 | A>A |
| chr3 | 1.61E+08 | 160700000 | -0.18 | 0.29 | A>A |
| chr3 | 1.61E+08 | 160800000 | -0.30 | 0.50 | B>A |
| chr3 | 1.61E+08 | 160900000 | -0.23 | 0.54 | B>A |
| chr3 | 1.61E+08 | 161000000 | 0.11 | 0.55 | A>A |
| chr3 | 1.61E+08 | 161100000 | 0.17 | 0.67 | A>A |

| | | | | | |
|---|---|---|---|---|---|
| chr3 | 1.61E+08 | 161200000 | -0.24 | 0.15 | B>B |
| chr3 | 1.61E+08 | 161300000 | -0.66 | -0.47 | B>B |
| chr3 | 1.61E+08 | 161400000 | -0.71 | -0.72 | B>B |
| chr3 | 1.61E+08 | 161500000 | -0.69 | -0.76 | B>B |
| chr3 | 1.62E+08 | 161600000 | -0.57 | -0.61 | B>B |
| chr3 | 1.62E+08 | 161700000 | -0.53 | -0.57 | B>B |
| chr3 | 1.62E+08 | 161800000 | -0.49 | -0.54 | B>B |
| chr3 | 1.62E+08 | 161900000 | -0.33 | -0.53 | B>B |
| chr3 | 1.62E+08 | 162000000 | -0.25 | -0.51 | B>B |
| chr3 | 1.62E+08 | 162100000 | -0.04 | -0.32 | B>B |
| chr3 | 1.62E+08 | 162200000 | -0.07 | -0.34 | B>B |
| chr3 | 1.62E+08 | 162300000 | -0.04 | -0.27 | B>B |
| chr3 | 1.62E+08 | 162400000 | -0.01 | -0.25 | B>B |
| chr3 | 1.62E+08 | 162500000 | 0.03 | -0.23 | B>B |
| chr3 | 1.63E+08 | 162600000 | 0.15 | 0.22 | A>A |
| chr3 | 1.63E+08 | 162700000 | -0.08 | -0.11 | B>B |
| chr3 | 1.63E+08 | 162800000 | -0.33 | -0.27 | B>B |
| chr3 | 1.63E+08 | 162900000 | -0.41 | -0.33 | B>B |
| chr3 | 1.63E+08 | 163000000 | -0.56 | -0.32 | B>B |
| chr3 | 1.63E+08 | 163100000 | -0.56 | -0.41 | B>B |
| chr3 | 1.63E+08 | 163200000 | -0.63 | -0.39 | B>B |
| chr3 | 1.63E+08 | 163300000 | -0.68 | -0.64 | B>B |
| chr3 | 1.63E+08 | 163400000 | -0.84 | -0.76 | B>B |
| chr3 | 1.63E+08 | 163500000 | -0.92 | -0.80 | B>B |
| chr3 | 1.64E+08 | 163600000 | -0.89 | -0.74 | B>B |
| chr3 | 1.64E+08 | 163700000 | -0.82 | -0.67 | B>B |
| chr3 | 1.64E+08 | 163800000 | -0.87 | -0.60 | B>B |
| chr3 | 1.64E+08 | 163900000 | -0.93 | -0.45 | B>B |
| chr3 | 1.64E+08 | 164000000 | -0.90 | -0.20 | B>A |
| chr3 | 1.64E+08 | 164100000 | -0.88 | -0.05 | B>A |
| chr3 | 1.64E+08 | 164200000 | -0.84 | 0.06 | B>A |
| chr3 | 1.64E+08 | 164300000 | -0.86 | 0.09 | B>A |
| chr3 | 1.64E+08 | 164400000 | -0.92 | 0.08 | B>A |
| chr3 | 1.64E+08 | 164500000 | -0.94 | -0.28 | B>A |
| chr3 | 1.65E+08 | 164600000 | -0.96 | -0.46 | B>B |
| chr3 | 1.65E+08 | 164700000 | -0.97 | -0.61 | B>B |
| chr3 | 1.65E+08 | 164800000 | -1.00 | -0.69 | B>B |
| chr3 | 1.65E+08 | 164900000 | -0.85 | -0.60 | B>B |
| chr3 | 1.65E+08 | 165000000 | -0.72 | -0.57 | B>B |
| chr3 | 1.65E+08 | 165100000 | -0.90 | -0.86 | B>B |
| chr3 | 1.65E+08 | 165200000 | -0.86 | -0.94 | B>B |
| chr3 | 1.65E+08 | 165300000 | -0.93 | -0.94 | B>B |
| chr3 | 1.65E+08 | 165400000 | -0.96 | -0.88 | B>B |
| chr3 | 1.65E+08 | 165500000 | -0.90 | -0.90 | B>B |
| chr3 | 1.66E+08 | 165600000 | -0.73 | -0.64 | B>B |
| chr3 | 1.66E+08 | 165700000 | -0.82 | -0.86 | B>B |
| chr3 | 1.66E+08 | 165800000 | -0.64 | -0.85 | B>B |
| chr3 | 1.66E+08 | 165900000 | -0.69 | -0.79 | B>B |
| chr3 | 1.66E+08 | 166000000 | -0.67 | -0.74 | B>B |
| chr3 | 1.66E+08 | 166100000 | -0.67 | -0.73 | B>B |
| chr3 | 1.66E+08 | 166200000 | -0.65 | -0.67 | B>B |
| chr3 | 1.66E+08 | 166300000 | -0.63 | -0.68 | B>B |
| chr3 | 1.66E+08 | 166400000 | -0.64 | -0.73 | B>B |
| chr3 | 1.66E+08 | 166500000 | -0.68 | -0.74 | B>B |
| chr3 | 1.67E+08 | 166600000 | -0.69 | -0.69 | B>B |
| chr3 | 1.67E+08 | 166700000 | -0.67 | -0.69 | B>B |

| | | | | | |
|---|---|---|---|---|---|
| chr3 | 1.67E+08 | 166800000 | -0.67 | -0.73 | B>B |
| chr3 | 1.67E+08 | 166900000 | -0.70 | -0.72 | B>B |
| chr3 | 1.67E+08 | 167000000 | -0.75 | -0.74 | B>B |
| chr3 | 1.67E+08 | 167100000 | -0.61 | -0.72 | B>B |
| chr3 | 1.67E+08 | 167200000 | -0.65 | -0.73 | B>B |
| chr3 | 1.67E+08 | 167300000 | -0.70 | -0.81 | B>B |
| chr3 | 1.67E+08 | 167400000 | -0.45 | -0.66 | B>B |
| chr3 | 1.67E+08 | 167500000 | 0.08 | -0.39 | B>B |
| chr3 | 1.68E+08 | 167600000 | 0.04 | -0.75 | A>B |
| chr3 | 1.68E+08 | 167700000 | -0.23 | -0.73 | B>B |
| chr3 | 1.68E+08 | 167800000 | -0.10 | -0.48 | B>B |
| chr3 | 1.68E+08 | 167900000 | -0.11 | -0.54 | B>B |
| chr3 | 1.68E+08 | 168000000 | -0.41 | -0.63 | B>B |
| chr3 | 1.68E+08 | 168100000 | -0.64 | -0.80 | B>B |
| chr3 | 1.68E+08 | 168200000 | -0.78 | -0.83 | B>B |
| chr3 | 1.68E+08 | 168300000 | -0.81 | -0.83 | B>B |
| chr3 | 1.68E+08 | 168400000 | -0.77 | -0.88 | B>B |
| chr3 | 1.68E+08 | 168500000 | -0.73 | -0.85 | B>B |
| chr3 | 1.69E+08 | 168600000 | -0.69 | -0.74 | B>B |
| chr3 | 1.69E+08 | 168700000 | -0.78 | -0.83 | B>B |
| chr3 | 1.69E+08 | 168800000 | -0.77 | -0.97 | B>B |
| chr3 | 1.69E+08 | 168900000 | -0.56 | -0.67 | B>B |
| chr3 | 1.69E+08 | 169000000 | -0.70 | -0.71 | B>B |
| chr3 | 1.69E+08 | 169100000 | -0.75 | -0.77 | B>B |
| chr3 | 1.69E+08 | 169200000 | -0.77 | -0.95 | B>B |
| chr3 | 1.69E+08 | 169300000 | -0.59 | -0.94 | B>B |
| chr3 | 1.69E+08 | 169400000 | -0.37 | -0.74 | B>B |
| chr3 | 1.69E+08 | 169500000 | -0.08 | -0.47 | B>B |
| chr3 | 1.7E+08 | 169600000 | 0.56 | 0.10 | A>A |
| chr3 | 1.7E+08 | 169700000 | 0.57 | 0.01 | A>A |
| chr3 | 1.7E+08 | 169800000 | 0.86 | 0.39 | A>A |
| chr3 | 1.7E+08 | 169900000 | 1.12 | 0.80 | A>A |
| chr3 | 1.7E+08 | 170000000 | 1.16 | 0.92 | A>A |
| chr3 | 1.7E+08 | 170100000 | 1.01 | 0.76 | A>A |
| chr3 | 1.7E+08 | 170200000 | 0.57 | 0.22 | A>A |
| chr3 | 1.7E+08 | 170300000 | 0.49 | -0.17 | A>B |
| chr3 | 1.7E+08 | 170400000 | 0.64 | 0.16 | A>A |
| chr3 | 1.7E+08 | 170500000 | 1.01 | 0.54 | A>A |
| chr3 | 1.71E+08 | 170600000 | 0.70 | 0.51 | A>A |
| chr3 | 1.71E+08 | 170700000 | 0.00 | 0.67 | NO |
| chr3 | 1.71E+08 | 170800000 | 0.98 | 0.79 | A>A |
| chr3 | 1.71E+08 | 170900000 | 0.97 | 0.69 | A>A |
| chr3 | 1.71E+08 | 171000000 | 0.80 | 0.59 | A>A |
| chr3 | 1.71E+08 | 171100000 | 0.97 | 0.68 | A>A |
| chr3 | 1.71E+08 | 171200000 | 0.87 | 0.64 | A>A |
| chr3 | 1.71E+08 | 171300000 | 0.75 | 0.62 | A>A |
| chr3 | 1.71E+08 | 171400000 | 0.76 | 0.72 | A>A |
| chr3 | 1.71E+08 | 171500000 | 0.97 | 0.92 | A>A |
| chr3 | 1.72E+08 | 171600000 | 0.84 | 0.98 | A>A |
| chr3 | 1.72E+08 | 171700000 | 1.13 | 1.09 | A>A |
| chr3 | 1.72E+08 | 171800000 | 0.92 | 0.97 | A>A |
| chr3 | 1.72E+08 | 171900000 | 0.97 | 1.01 | A>A |
| chr3 | 1.72E+08 | 172000000 | 0.98 | 1.00 | A>A |
| chr3 | 1.72E+08 | 172100000 | 0.71 | 0.78 | A>A |
| chr3 | 1.72E+08 | 172200000 | 0.62 | 0.74 | A>A |
| chr3 | 1.72E+08 | 172300000 | 0.86 | 0.85 | A>A |

| | | | | | |
|---|---|---|---|---|---|
| chr3 | 1.72E+08 | 172400000 | 0.66 | 0.71 | A>A |
| chr3 | 1.72E+08 | 172500000 | 1.16 | 0.77 | A>A |
| chr3 | 1.73E+08 | 172600000 | 0.49 | -0.26 | A>B |
| chr3 | 1.73E+08 | 172700000 | -0.24 | -0.93 | A>B |
| chr3 | 1.73E+08 | 172800000 | -0.41 | -1.02 | A>B |
| chr3 | 1.73E+08 | 172900000 | -0.67 | -1.09 | B>B |
| chr3 | 1.73E+08 | 173000000 | -0.72 | -1.08 | B>B |
| chr3 | 1.73E+08 | 173100000 | -0.58 | -0.73 | B>B |
| chr3 | 1.73E+08 | 173200000 | -0.59 | -0.64 | B>B |
| chr3 | 1.73E+08 | 173300000 | -0.62 | -0.68 | B>B |
| chr3 | 1.73E+08 | 173400000 | -0.67 | -0.68 | B>B |
| chr3 | 1.73E+08 | 173500000 | -0.69 | -0.73 | B>B |
| chr3 | 1.74E+08 | 173600000 | -0.66 | -0.73 | B>B |
| chr3 | 1.74E+08 | 173700000 | -0.63 | -0.74 | B>B |
| chr3 | 1.74E+08 | 173800000 | -0.65 | -0.72 | B>B |
| chr3 | 1.74E+08 | 173900000 | -0.62 | -0.62 | B>B |
| chr3 | 1.74E+08 | 174000000 | -0.58 | -0.55 | B>B |
| chr3 | 1.74E+08 | 174100000 | -0.43 | -0.30 | B>B |
| chr3 | 1.74E+08 | 174200000 | -0.32 | -0.01 | B>B |
| chr3 | 1.74E+08 | 174300000 | -0.48 | 0.01 | B>B |
| chr3 | 1.74E+08 | 174400000 | -0.54 | 0.06 | B>A |
| chr3 | 1.74E+08 | 174500000 | -0.45 | 0.08 | B>A |
| chr3 | 1.75E+08 | 174600000 | -0.41 | 0.15 | B>A |
| chr3 | 1.75E+08 | 174700000 | -0.22 | 0.18 | B>B |
| chr3 | 1.75E+08 | 174800000 | -0.36 | 0.30 | B>A |
| chr3 | 1.75E+08 | 174900000 | -0.43 | 0.21 | B>A |
| chr3 | 1.75E+08 | 175000000 | -0.44 | 0.22 | B>A |
| chr3 | 1.75E+08 | 175100000 | -0.44 | 0.25 | B>A |
| chr3 | 1.75E+08 | 175200000 | -0.56 | 0.20 | B>A |
| chr3 | 1.75E+08 | 175300000 | -0.55 | 0.26 | B>A |
| chr3 | 1.75E+08 | 175400000 | -0.60 | 0.25 | B>A |
| chr3 | 1.75E+08 | 175500000 | -0.59 | 0.23 | B>A |
| chr3 | 1.76E+08 | 175600000 | -0.65 | 0.18 | B>A |
| chr3 | 1.76E+08 | 175700000 | -0.59 | 0.06 | B>A |
| chr3 | 1.76E+08 | 175800000 | -0.63 | -0.05 | B>A |
| chr3 | 1.76E+08 | 175900000 | -0.66 | -0.18 | B>B |
| chr3 | 1.76E+08 | 176000000 | -0.67 | -0.21 | B>B |
| chr3 | 1.76E+08 | 176100000 | -0.64 | -0.22 | B>B |
| chr3 | 1.76E+08 | 176200000 | -0.63 | -0.19 | B>B |
| chr3 | 1.76E+08 | 176300000 | -0.63 | -0.35 | B>B |
| chr3 | 1.76E+08 | 176400000 | -0.64 | -0.49 | B>B |
| chr3 | 1.76E+08 | 176500000 | -0.62 | -0.62 | B>B |
| chr3 | 1.77E+08 | 176600000 | -0.41 | -0.52 | B>B |
| chr3 | 1.77E+08 | 176700000 | -0.21 | -0.20 | B>B |
| chr3 | 1.77E+08 | 176800000 | 0.20 | 0.20 | A>A |
| chr3 | 1.77E+08 | 176900000 | 0.59 | 0.55 | A>A |
| chr3 | 1.77E+08 | 177000000 | 0.47 | 0.53 | A>A |
| chr3 | 1.77E+08 | 177100000 | 0.46 | 0.46 | A>A |
| chr3 | 1.77E+08 | 177200000 | 0.49 | 0.52 | A>A |
| chr3 | 1.77E+08 | 177300000 | 0.45 | 0.54 | A>A |
| chr3 | 1.77E+08 | 177400000 | 0.46 | 0.40 | A>A |
| chr3 | 1.77E+08 | 177500000 | 0.44 | 0.48 | A>A |
| chr3 | 1.78E+08 | 177600000 | 0.37 | 0.39 | A>A |
| chr3 | 1.78E+08 | 177700000 | 0.40 | 0.44 | A>A |
| chr3 | 1.78E+08 | 177800000 | 0.36 | 0.38 | A>A |
| chr3 | 1.78E+08 | 177900000 | 0.33 | 0.32 | A>A |

| | | | | | |
|---|---|---|---|---|---|
| chr3 | 1.78E+08 | 178000000 | 0.25 | 0.21 | A>A |
| chr3 | 1.78E+08 | 178100000 | 0.06 | 0.09 | A>A |
| chr3 | 1.78E+08 | 178200000 | -0.28 | -0.32 | B>B |
| chr3 | 1.78E+08 | 178300000 | -0.52 | -0.79 | B>B |
| chr3 | 1.78E+08 | 178400000 | -0.69 | -0.86 | B>B |
| chr3 | 1.78E+08 | 178500000 | -0.78 | -0.91 | B>B |
| chr3 | 1.79E+08 | 178600000 | -0.75 | -0.92 | B>B |
| chr3 | 1.79E+08 | 178700000 | -0.60 | -0.65 | B>B |
| chr3 | 1.79E+08 | 178800000 | -0.07 | -0.23 | B>B |
| chr3 | 1.79E+08 | 178900000 | 0.25 | -0.09 | A>A |
| chr3 | 1.79E+08 | 179000000 | 0.37 | 0.05 | A>A |
| chr3 | 1.79E+08 | 179100000 | 0.28 | 0.06 | A>A |
| chr3 | 1.79E+08 | 179200000 | 0.46 | 0.06 | A>A |
| chr3 | 1.79E+08 | 179300000 | 0.41 | -0.06 | A>A |
| chr3 | 1.79E+08 | 179400000 | 0.57 | 0.17 | A>A |
| chr3 | 1.79E+08 | 179500000 | 0.29 | -0.11 | A>A |
| chr3 | 1.8E+08 | 179600000 | -0.09 | -0.59 | B>B |
| chr3 | 1.8E+08 | 179700000 | -0.15 | -0.82 | A>B |
| chr3 | 1.8E+08 | 179800000 | -0.19 | -0.91 | A>B |
| chr3 | 1.8E+08 | 179900000 | -0.53 | -0.91 | B>B |
| chr3 | 1.8E+08 | 180000000 | -0.62 | -0.96 | B>B |
| chr3 | 1.8E+08 | 180100000 | -0.63 | -0.96 | B>B |
| chr3 | 1.8E+08 | 180200000 | -0.70 | -0.89 | B>B |
| chr3 | 1.8E+08 | 180300000 | -0.58 | -0.78 | B>B |
| chr3 | 1.8E+08 | 180400000 | -0.24 | -0.41 | B>B |
| chr3 | 1.8E+08 | 180500000 | -0.49 | -0.52 | B>B |
| chr3 | 1.81E+08 | 180600000 | -0.41 | -0.61 | B>B |
| chr3 | 1.81E+08 | 180700000 | 0.06 | -0.15 | B>B |
| chr3 | 1.81E+08 | 180800000 | -0.45 | -0.63 | B>B |
| chr3 | 1.81E+08 | 180900000 | -0.63 | -0.78 | B>B |
| chr3 | 1.81E+08 | 181000000 | -0.52 | -0.69 | B>B |
| chr3 | 1.81E+08 | 181100000 | -0.57 | -0.82 | B>B |
| chr3 | 1.81E+08 | 181200000 | -0.67 | -0.81 | B>B |
| chr3 | 1.81E+08 | 181300000 | -0.58 | -0.72 | B>B |
| chr3 | 1.81E+08 | 181400000 | -0.32 | -0.39 | B>B |
| chr3 | 1.81E+08 | 181500000 | 0.19 | 0.15 | A>A |
| chr3 | 1.82E+08 | 181600000 | 0.51 | 0.47 | A>A |
| chr3 | 1.82E+08 | 181700000 | 0.47 | 0.38 | A>A |
| chr3 | 1.82E+08 | 181800000 | 0.54 | 0.51 | A>A |
| chr3 | 1.82E+08 | 181900000 | 0.52 | 0.55 | A>A |
| chr3 | 1.82E+08 | 182000000 | 0.21 | 0.31 | A>A |
| chr3 | 1.82E+08 | 182100000 | 0.04 | 0.19 | A>A |
| chr3 | 1.82E+08 | 182200000 | -0.06 | 0.18 | A>A |
| chr3 | 1.82E+08 | 182300000 | -0.17 | 0.16 | B>B |
| chr3 | 1.82E+08 | 182400000 | -0.16 | 0.11 | B>B |
| chr3 | 1.82E+08 | 182500000 | 0.28 | 0.33 | A>A |
| chr3 | 1.83E+08 | 182600000 | 0.70 | 0.59 | A>A |
| chr3 | 1.83E+08 | 182700000 | 0.81 | 0.86 | A>A |
| chr3 | 1.83E+08 | 182800000 | 1.09 | 1.02 | A>A |
| chr3 | 1.83E+08 | 182900000 | 0.85 | 0.98 | A>A |
| chr3 | 1.83E+08 | 183000000 | 1.08 | 1.13 | A>A |
| chr3 | 1.83E+08 | 183100000 | 1.16 | 1.20 | A>A |
| chr3 | 1.83E+08 | 183200000 | 0.81 | 0.93 | A>A |
| chr3 | 1.83E+08 | 183300000 | 1.05 | 1.19 | A>A |
| chr3 | 1.83E+08 | 183400000 | 1.25 | 1.29 | A>A |
| chr3 | 1.83E+08 | 183500000 | 1.09 | 1.16 | A>A |

| | | | | | |
|---|---|---|---|---|---|
| chr3 | 1.84E+08 | 183600000 | 1.11 | 1.18 | A>A |
| chr3 | 1.84E+08 | 183700000 | 1.24 | 1.24 | A>A |
| chr3 | 1.84E+08 | 183800000 | 1.36 | 1.30 | A>A |
| chr3 | 1.84E+08 | 183900000 | 1.23 | 1.28 | A>A |
| chr3 | 1.84E+08 | 184000000 | 1.20 | 1.24 | A>A |
| chr3 | 1.84E+08 | 184100000 | 1.01 | 1.12 | A>A |
| chr3 | 1.84E+08 | 184200000 | 1.26 | 1.15 | A>A |
| chr3 | 1.84E+08 | 184300000 | 1.04 | 1.06 | A>A |
| chr3 | 1.84E+08 | 184400000 | 1.27 | 1.25 | A>A |
| chr3 | 1.84E+08 | 184500000 | 1.20 | 1.14 | A>A |
| chr3 | 1.85E+08 | 184600000 | 1.06 | 1.03 | A>A |
| chr3 | 1.85E+08 | 184700000 | 1.18 | 1.05 | A>A |
| chr3 | 1.85E+08 | 184800000 | 1.09 | 1.16 | A>A |
| chr3 | 1.85E+08 | 184900000 | 1.10 | 1.16 | A>A |
| chr3 | 1.85E+08 | 185000000 | 1.11 | 1.03 | A>A |
| chr3 | 1.85E+08 | 185100000 | 1.23 | 1.27 | A>A |
| chr3 | 1.85E+08 | 185200000 | 1.21 | 1.19 | A>A |
| chr3 | 1.85E+08 | 185300000 | 0.00 | 0.00 | NO |
| chr3 | 1.85E+08 | 185400000 | 1.02 | 1.07 | A>A |
| chr3 | 1.85E+08 | 185500000 | 1.22 | 1.16 | A>A |
| chr3 | 1.86E+08 | 185600000 | 1.30 | 1.27 | A>A |
| chr3 | 1.86E+08 | 185700000 | 1.08 | 1.01 | A>A |
| chr3 | 1.86E+08 | 185800000 | 0.92 | 0.86 | A>A |
| chr3 | 1.86E+08 | 185900000 | 1.07 | 1.11 | A>A |
| chr3 | 1.86E+08 | 186000000 | 0.85 | 0.82 | A>A |
| chr3 | 1.86E+08 | 186100000 | 0.88 | 0.76 | A>A |
| chr3 | 1.86E+08 | 186200000 | 0.97 | 0.85 | A>A |
| chr3 | 1.86E+08 | 186300000 | 0.84 | 0.77 | A>A |
| chr3 | 1.86E+08 | 186400000 | 1.00 | 0.96 | A>A |
| chr3 | 1.86E+08 | 186500000 | 0.00 | 0.00 | NO |
| chr3 | 1.87E+08 | 186600000 | 0.79 | 0.77 | A>A |
| chr3 | 1.87E+08 | 186700000 | 0.92 | 0.95 | A>A |
| chr3 | 1.87E+08 | 186800000 | 0.86 | 0.94 | A>A |
| chr3 | 1.87E+08 | 186900000 | 0.62 | 0.76 | A>A |
| chr3 | 1.87E+08 | 187000000 | 0.50 | 0.67 | A>A |
| chr3 | 1.87E+08 | 187100000 | 0.45 | 0.58 | A>A |
| chr3 | 1.87E+08 | 187200000 | 0.46 | 0.73 | A>A |
| chr3 | 1.87E+08 | 187300000 | 0.27 | 0.66 | A>A |
| chr3 | 1.87E+08 | 187400000 | 0.37 | 0.56 | A>A |
| chr3 | 1.87E+08 | 187500000 | 0.29 | 0.51 | A>A |
| chr3 | 1.88E+08 | 187600000 | 0.84 | 0.94 | A>A |
| chr3 | 1.88E+08 | 187700000 | 0.73 | 0.88 | A>A |
| chr3 | 1.88E+08 | 187800000 | 0.66 | 0.77 | A>A |
| chr3 | 1.88E+08 | 187900000 | 0.78 | 0.80 | A>A |
| chr3 | 1.88E+08 | 188000000 | 0.57 | 0.82 | A>A |
| chr3 | 1.88E+08 | 188100000 | 0.61 | 0.78 | A>A |
| chr3 | 1.88E+08 | 188200000 | 0.62 | 0.81 | A>A |
| chr3 | 1.88E+08 | 188300000 | 0.62 | 0.70 | A>A |
| chr3 | 1.88E+08 | 188400000 | 0.52 | 0.70 | A>A |
| chr3 | 1.88E+08 | 188500000 | 0.55 | 0.71 | A>A |
| chr3 | 1.89E+08 | 188600000 | 0.58 | 0.76 | A>A |
| chr3 | 1.89E+08 | 188700000 | 0.46 | 0.62 | A>A |
| chr3 | 1.89E+08 | 188800000 | 0.49 | 0.72 | A>A |
| chr3 | 1.89E+08 | 188900000 | 0.44 | 0.70 | A>A |
| chr3 | 1.89E+08 | 189000000 | 0.51 | 0.57 | A>A |
| chr3 | 1.89E+08 | 189100000 | 0.60 | 0.75 | A>A |

| | | | | | |
|---|---|---|---|---|---|
| chr3 | 1.89E+08 | 189200000 | 0.81 | 0.88 | A>A |
| chr3 | 1.89E+08 | 189300000 | 0.72 | 0.86 | A>A |
| chr3 | 1.89E+08 | 189400000 | 0.73 | 0.79 | A>A |
| chr3 | 1.89E+08 | 189500000 | 0.77 | 0.74 | A>A |
| chr3 | 1.9E+08 | 189600000 | 0.72 | 0.82 | A>A |
| chr3 | 1.9E+08 | 189700000 | 0.41 | 0.57 | A>A |
| chr3 | 1.9E+08 | 189800000 | 0.65 | 0.66 | A>A |
| chr3 | 1.9E+08 | 189900000 | 0.85 | 0.78 | A>A |
| chr3 | 1.9E+08 | 190000000 | 0.59 | 0.50 | A>A |
| chr3 | 1.9E+08 | 190100000 | 0.65 | 0.61 | A>A |
| chr3 | 1.9E+08 | 190200000 | 0.45 | 0.39 | A>A |
| chr3 | 1.9E+08 | 190300000 | 0.43 | 0.27 | A>A |
| chr3 | 1.9E+08 | 190400000 | 0.34 | 0.13 | A>A |
| chr3 | 1.9E+08 | 190500000 | -0.01 | -0.31 | B>B |
| chr3 | 1.91E+08 | 190600000 | -0.24 | -0.50 | B>B |
| chr3 | 1.91E+08 | 190700000 | -0.38 | -0.77 | B>B |
| chr3 | 1.91E+08 | 190800000 | -0.33 | -0.68 | B>B |
| chr3 | 1.91E+08 | 190900000 | -0.30 | -0.68 | B>B |
| chr3 | 1.91E+08 | 191000000 | -0.12 | -0.50 | B>B |
| chr3 | 1.91E+08 | 191100000 | 0.11 | -0.20 | B>B |
| chr3 | 1.91E+08 | 191200000 | -0.37 | -0.23 | B>B |
| chr3 | 1.91E+08 | 191300000 | -0.63 | -0.24 | B>B |
| chr3 | 1.91E+08 | 191400000 | -0.67 | -0.17 | B>B |
| chr3 | 1.91E+08 | 191500000 | -0.61 | -0.18 | B>B |
| chr3 | 1.92E+08 | 191600000 | -0.60 | -0.28 | B>B |
| chr3 | 1.92E+08 | 191700000 | -0.62 | -0.26 | B>B |
| chr3 | 1.92E+08 | 191800000 | -0.69 | -0.21 | B>B |
| chr3 | 1.92E+08 | 191900000 | -0.52 | -0.14 | B>B |
| chr3 | 1.92E+08 | 192000000 | -0.53 | 0.07 | B>A |
| chr3 | 1.92E+08 | 192100000 | -0.45 | 0.17 | B>A |
| chr3 | 1.92E+08 | 192200000 | -0.25 | -0.09 | B>B |
| chr3 | 1.92E+08 | 192300000 | -0.49 | -0.60 | B>B |
| chr3 | 1.92E+08 | 192400000 | -0.55 | -0.96 | B>B |
| chr3 | 1.92E+08 | 192500000 | -0.52 | -0.84 | B>B |
| chr3 | 1.93E+08 | 192600000 | 0.09 | -0.21 | B>B |
| chr3 | 1.93E+08 | 192700000 | 0.43 | -0.16 | A>B |
| chr3 | 1.93E+08 | 192800000 | -0.05 | -0.54 | B>B |
| chr3 | 1.93E+08 | 192900000 | -0.53 | -0.92 | B>B |
| chr3 | 1.93E+08 | 193000000 | -0.49 | -0.92 | B>B |
| chr3 | 1.93E+08 | 193100000 | -0.59 | -1.07 | B>B |
| chr3 | 1.93E+08 | 193200000 | -0.59 | -0.91 | B>B |
| chr3 | 1.93E+08 | 193300000 | -0.34 | -0.69 | B>B |
| chr3 | 1.93E+08 | 193400000 | 0.68 | 0.35 | A>A |
| chr3 | 1.93E+08 | 193500000 | 0.83 | 0.68 | A>A |
| chr3 | 1.94E+08 | 193600000 | 0.86 | 0.90 | A>A |
| chr3 | 1.94E+08 | 193700000 | 1.03 | 0.99 | A>A |
| chr3 | 1.94E+08 | 193800000 | 1.05 | 0.98 | A>A |
| chr3 | 1.94E+08 | 193900000 | 0.95 | 0.89 | A>A |
| chr3 | 1.94E+08 | 194000000 | 1.14 | 0.91 | A>A |
| chr3 | 1.94E+08 | 194100000 | 0.77 | 0.82 | A>A |
| chr3 | 1.94E+08 | 194200000 | 1.17 | 1.00 | A>A |
| chr3 | 1.94E+08 | 194300000 | 1.49 | 1.28 | A>A |
| chr3 | 1.94E+08 | 194400000 | 1.03 | 0.98 | A>A |
| chr3 | 1.94E+08 | 194500000 | 1.08 | 1.13 | A>A |
| chr3 | 1.95E+08 | 194600000 | 1.35 | 1.30 | A>A |
| chr3 | 1.95E+08 | 194700000 | 1.17 | 1.30 | A>A |

| | | | | | |
|---|---|---|---|---|---|
| chr3 | 1.95E+08 | 194800000 | 1.20 | 1.35 | A>A |
| chr3 | 1.95E+08 | 194900000 | 1.21 | 1.28 | A>A |
| chr3 | 1.95E+08 | 195000000 | 1.13 | 1.26 | A>A |
| chr3 | 1.95E+08 | 195100000 | 1.56 | 1.48 | A>A |
| chr3 | 1.95E+08 | 195200000 | 1.55 | 1.54 | A>A |
| chr3 | 1.95E+08 | 195300000 | 0.00 | 0.00 | NO |
| chr3 | 1.95E+08 | 195400000 | 0.00 | 0.00 | NO |
| chr3 | 1.95E+08 | 195500000 | 1.98 | 1.75 | A>A |
| chr3 | 1.96E+08 | 195600000 | 0.00 | 0.00 | NO |
| chr3 | 1.96E+08 | 195700000 | 0.00 | 0.00 | NO |
| chr3 | 1.96E+08 | 195800000 | 0.00 | 0.00 | NO |
| chr3 | 1.96E+08 | 195900000 | 0.00 | 0.00 | NO |
| chr3 | 1.96E+08 | 196000000 | 1.64 | 1.40 | A>A |
| chr3 | 1.96E+08 | 196100000 | 0.00 | 0.00 | NO |
| chr3 | 1.96E+08 | 196200000 | 0.00 | 0.00 | NO |
| chr3 | 1.96E+08 | 196300000 | 0.00 | 1.50 | NO |
| chr3 | 1.96E+08 | 196400000 | 1.55 | 1.53 | A>A |
| chr3 | 1.96E+08 | 196500000 | 0.00 | 0.00 | NO |
| chr3 | 1.97E+08 | 196600000 | 0.00 | 0.00 | NO |
| chr3 | 1.97E+08 | 196700000 | 1.63 | 1.41 | A>A |
| chr3 | 1.97E+08 | 196800000 | 1.55 | 1.41 | A>A |
| chr3 | 1.97E+08 | 196900000 | 1.65 | 1.43 | A>A |
| chr3 | 1.97E+08 | 197000000 | 1.44 | 1.39 | A>A |
| chr3 | 1.97E+08 | 197100000 | 1.54 | 1.47 | A>A |
| chr3 | 1.97E+08 | 197200000 | 0.00 | 0.00 | NO |
| chr3 | 1.97E+08 | 197300000 | 1.35 | 1.31 | A>A |
| chr3 | 1.97E+08 | 197400000 | 0.00 | 0.00 | NO |
| chr3 | 1.97E+08 | 197500000 | 1.89 | 1.51 | A>A |
| chr3 | 1.98E+08 | 197600000 | 1.86 | 1.59 | A>A |
| chr3 | 1.98E+08 | 197700000 | 0.00 | 1.19 | NO |
| chr3 | 1.98E+08 | 197800000 | 0.00 | 0.00 | NO |
| chr3 | 1.98E+08 | 197900000 | 0.00 | 0.00 | NO |
| chr3 | 1.98E+08 | 198000000 | 0.00 | 0.00 | NO |
| chr3 | 1.98E+08 | 198022430 | 0.00 | 0.00 | NO |
| chr4 | 0 | 100000 | -0.42 | 0.51 | B>A |
| chr4 | 100000 | 200000 | -0.53 | 1.56 | B>A |
| chr4 | 200000 | 300000 | -0.82 | 1.22 | B>A |
| chr4 | 300000 | 400000 | 0.06 | 1.20 | B>A |
| chr4 | 400000 | 500000 | 0.47 | 1.31 | B>A |
| chr4 | 500000 | 600000 | 0.60 | 1.15 | B>A |
| chr4 | 600000 | 700000 | 0.62 | 0.95 | A>A |
| chr4 | 700000 | 800000 | 1.36 | 0.93 | A>A |
| chr4 | 800000 | 900000 | 1.36 | 0.81 | A>A |
| chr4 | 900000 | 1000000 | 1.39 | 0.92 | A>A |
| chr4 | 1000000 | 1100000 | 1.19 | 1.02 | A>A |
| chr4 | 1100000 | 1200000 | 1.23 | 1.39 | A>A |
| chr4 | 1200000 | 1300000 | 1.21 | 1.17 | A>A |
| chr4 | 1300000 | 1400000 | 0.98 | 0.72 | A>A |
| chr4 | 1400000 | 1500000 | 0.00 | 0.00 | NO |
| chr4 | 1500000 | 1600000 | 1.13 | 0.83 | A>A |
| chr4 | 1600000 | 1700000 | 1.11 | 0.80 | A>A |
| chr4 | 1700000 | 1800000 | 1.50 | 0.94 | A>A |
| chr4 | 1800000 | 1900000 | 1.44 | 0.87 | A>A |
| chr4 | 1900000 | 2000000 | 1.49 | 0.86 | A>B |
| chr4 | 2000000 | 2100000 | 1.28 | 1.00 | A>A |
| chr4 | 2100000 | 2200000 | 1.41 | 0.95 | A>A |

| | | | | | |
|---|---|---|---|---|---|
| chr4 | 2200000 | 2300000 | 0.97 | 0.89 | A>A |
| chr4 | 2300000 | 2400000 | 1.45 | 1.06 | A>A |
| chr4 | 2400000 | 2500000 | 1.42 | 0.95 | A>A |
| chr4 | 2500000 | 2600000 | 1.49 | 0.97 | A>A |
| chr4 | 2600000 | 2700000 | 1.56 | 1.04 | A>A |
| chr4 | 2700000 | 2800000 | 1.22 | 0.87 | A>A |
| chr4 | 2800000 | 2900000 | 1.41 | 0.95 | A>A |
| chr4 | 2900000 | 3000000 | 1.13 | 0.77 | A>A |
| chr4 | 3000000 | 3100000 | 1.17 | 0.83 | A>A |
| chr4 | 3100000 | 3200000 | 1.12 | 0.77 | A>A |
| chr4 | 3200000 | 3300000 | 1.18 | 0.93 | A>A |
| chr4 | 3300000 | 3400000 | 1.40 | 0.95 | A>A |
| chr4 | 3400000 | 3500000 | 1.11 | 0.81 | A>A |
| chr4 | 3500000 | 3600000 | 0.73 | 0.39 | A>A |
| chr4 | 3600000 | 3700000 | 0.92 | 0.80 | A>A |
| chr4 | 3700000 | 3800000 | 1.14 | 0.91 | A>A |
| chr4 | 3800000 | 3900000 | 0.94 | 0.76 | A>A |
| chr4 | 3900000 | 4000000 | 0.00 | 0.00 | NO |
| chr4 | 4000000 | 4100000 | 0.00 | 0.00 | NO |
| chr4 | 4100000 | 4200000 | 0.00 | 0.00 | NO |
| chr4 | 4200000 | 4300000 | 1.34 | 0.99 | A>A |
| chr4 | 4300000 | 4400000 | 1.37 | 1.14 | A>A |
| chr4 | 4400000 | 4500000 | 1.29 | 1.00 | A>A |
| chr4 | 4500000 | 4600000 | 1.25 | 1.09 | A>A |
| chr4 | 4600000 | 4700000 | 1.15 | 1.08 | A>A |
| chr4 | 4700000 | 4800000 | 1.06 | 1.00 | A>A |
| chr4 | 4800000 | 4900000 | 1.02 | 0.82 | A>A |
| chr4 | 4900000 | 5000000 | 0.86 | 0.81 | A>A |
| chr4 | 5000000 | 5100000 | 1.08 | 1.04 | A>A |
| chr4 | 5100000 | 5200000 | 1.62 | 1.39 | A>A |
| chr4 | 5200000 | 5300000 | 1.47 | 1.21 | A>A |
| chr4 | 5300000 | 5400000 | 1.31 | 1.13 | A>A |
| chr4 | 5400000 | 5500000 | 1.11 | 0.98 | A>A |
| chr4 | 5500000 | 5600000 | 1.10 | 1.09 | A>A |
| chr4 | 5600000 | 5700000 | 1.20 | 1.07 | A>A |
| chr4 | 5700000 | 5800000 | 1.33 | 1.09 | A>A |
| chr4 | 5800000 | 5900000 | 0.99 | 0.83 | A>A |
| chr4 | 5900000 | 6000000 | 1.09 | 0.87 | A>A |
| chr4 | 6000000 | 6100000 | 1.24 | 0.96 | A>A |
| chr4 | 6100000 | 6200000 | 1.17 | 0.97 | A>A |
| chr4 | 6200000 | 6300000 | 1.30 | 1.11 | A>A |
| chr4 | 6300000 | 6400000 | 1.29 | 1.08 | A>A |
| chr4 | 6400000 | 6500000 | 1.53 | 1.37 | A>A |
| chr4 | 6500000 | 6600000 | 1.44 | 1.03 | A>A |
| chr4 | 6600000 | 6700000 | 1.54 | 1.16 | A>A |
| chr4 | 6700000 | 6800000 | 1.68 | 1.32 | A>A |
| chr4 | 6800000 | 6900000 | 1.71 | 1.43 | A>A |
| chr4 | 6900000 | 7000000 | 1.73 | 1.41 | A>A |
| chr4 | 7000000 | 7100000 | 1.51 | 1.24 | A>A |
| chr4 | 7100000 | 7200000 | 1.07 | 0.93 | A>A |
| chr4 | 7200000 | 7300000 | 1.29 | 1.10 | A>A |
| chr4 | 7300000 | 7400000 | 1.03 | 1.08 | A>A |
| chr4 | 7400000 | 7500000 | 1.24 | 1.02 | A>A |
| chr4 | 7500000 | 7600000 | 1.14 | 1.27 | A>A |
| chr4 | 7600000 | 7700000 | 1.05 | 1.06 | A>A |
| chr4 | 7700000 | 7800000 | 1.00 | 0.98 | A>A |

| | | | | | |
|---|---|---|---|---|---|
| chr4 | 7800000 | 7900000 | 1.71 | 1.49 | A>A |
| chr4 | 7900000 | 8000000 | 1.61 | 1.37 | A>A |
| chr4 | 8000000 | 8100000 | 1.15 | 1.06 | A>A |
| chr4 | 8100000 | 8200000 | 1.31 | 1.32 | A>A |
| chr4 | 8200000 | 8300000 | 1.18 | 1.20 | A>A |
| chr4 | 8300000 | 8400000 | 1.34 | 1.24 | A>A |
| chr4 | 8400000 | 8500000 | 1.11 | 1.09 | A>A |
| chr4 | 8500000 | 8600000 | 1.22 | 1.09 | A>A |
| chr4 | 8600000 | 8700000 | 0.94 | 0.83 | A>A |
| chr4 | 8700000 | 8800000 | 1.31 | 0.85 | A>A |
| chr4 | 8800000 | 8900000 | 1.32 | 0.68 | A>B |
| chr4 | 8900000 | 9000000 | 0.00 | 0.00 | NO |
| chr4 | 9000000 | 9100000 | 0.00 | 0.00 | NO |
| chr4 | 9100000 | 9200000 | 0.00 | 0.00 | NO |
| chr4 | 9200000 | 9300000 | 0.00 | 0.00 | NO |
| chr4 | 9300000 | 9400000 | 0.00 | 0.00 | NO |
| chr4 | 9400000 | 9500000 | 0.00 | 0.00 | NO |
| chr4 | 9500000 | 9600000 | 0.00 | 0.00 | NO |
| chr4 | 9600000 | 9700000 | 0.00 | 0.00 | NO |
| chr4 | 9700000 | 9800000 | 0.00 | 0.00 | NO |
| chr4 | 9800000 | 9900000 | 0.29 | -0.44 | A>B |
| chr4 | 9900000 | 10000000 | 0.56 | -0.05 | A>B |
| chr4 | 10000000 | 10100000 | 0.60 | 0.15 | A>A |
| chr4 | 10100000 | 10200000 | 0.18 | -0.51 | A>B |
| chr4 | 10200000 | 10300000 | -0.37 | -0.94 | B>B |
| chr4 | 10300000 | 10400000 | -0.36 | -0.96 | A>B |
| chr4 | 10400000 | 10500000 | -0.17 | -0.48 | B>B |
| chr4 | 10500000 | 10600000 | -0.19 | -0.55 | B>B |
| chr4 | 10600000 | 10700000 | -0.37 | -0.75 | B>B |
| chr4 | 10700000 | 10800000 | -0.48 | -0.76 | B>B |
| chr4 | 10800000 | 10900000 | -0.51 | -0.89 | B>B |
| chr4 | 10900000 | 11000000 | -0.60 | -0.83 | B>B |
| chr4 | 11000000 | 11100000 | -0.46 | -0.82 | B>B |
| chr4 | 11100000 | 11200000 | -0.40 | -0.77 | B>B |
| chr4 | 11200000 | 11300000 | -0.11 | -0.72 | A>B |
| chr4 | 11300000 | 11400000 | 0.02 | -0.58 | A>B |
| chr4 | 11400000 | 11500000 | 0.24 | 0.08 | A>A |
| chr4 | 11500000 | 11600000 | 0.52 | 0.39 | A>A |
| chr4 | 11600000 | 11700000 | 0.51 | 0.26 | A>A |
| chr4 | 11700000 | 11800000 | 0.16 | -0.18 | B>B |
| chr4 | 11800000 | 11900000 | -0.26 | -0.75 | B>B |
| chr4 | 11900000 | 12000000 | -0.48 | -0.85 | B>B |
| chr4 | 12000000 | 12100000 | -0.65 | -0.89 | B>B |
| chr4 | 12100000 | 12200000 | -0.85 | -1.00 | B>B |
| chr4 | 12200000 | 12300000 | -0.84 | -1.00 | B>B |
| chr4 | 12300000 | 12400000 | -0.89 | -1.06 | B>B |
| chr4 | 12400000 | 12500000 | -0.87 | -0.98 | B>B |
| chr4 | 12500000 | 12600000 | -0.76 | -0.97 | B>B |
| chr4 | 12600000 | 12700000 | -0.56 | -0.73 | B>B |
| chr4 | 12700000 | 12800000 | -0.26 | -0.76 | B>B |
| chr4 | 12800000 | 12900000 | -0.30 | -0.74 | B>B |
| chr4 | 12900000 | 13000000 | -0.45 | -1.11 | A>B |
| chr4 | 13000000 | 13100000 | -0.30 | -1.03 | A>B |
| chr4 | 13100000 | 13200000 | -0.07 | -0.98 | A>B |
| chr4 | 13200000 | 13300000 | -0.17 | -0.83 | A>B |
| chr4 | 13300000 | 13400000 | 0.20 | -0.39 | A>B |

| | | | | | |
|---|---|---|---|---|---|
| chr4 | 13400000 | 13500000 | 0.45 | 0.00 | A>A |
| chr4 | 13500000 | 13600000 | 0.23 | -0.14 | A>A |
| chr4 | 13600000 | 13700000 | 0.41 | 0.35 | A>A |
| chr4 | 13700000 | 13800000 | 0.87 | 0.89 | A>A |
| chr4 | 13800000 | 13900000 | 0.87 | 0.67 | A>A |
| chr4 | 13900000 | 14000000 | 0.76 | 0.67 | A>A |
| chr4 | 14000000 | 14100000 | 0.96 | 0.87 | A>A |
| chr4 | 14100000 | 14200000 | 0.72 | 0.90 | A>A |
| chr4 | 14200000 | 14300000 | 0.71 | 0.82 | A>A |
| chr4 | 14300000 | 14400000 | 0.78 | 0.67 | A>A |
| chr4 | 14400000 | 14500000 | 0.77 | 0.70 | A>A |
| chr4 | 14500000 | 14600000 | 0.62 | 0.75 | A>A |
| chr4 | 14600000 | 14700000 | 0.70 | 0.83 | A>A |
| chr4 | 14700000 | 14800000 | 0.49 | 0.64 | A>A |
| chr4 | 14800000 | 14900000 | 0.81 | 0.91 | A>A |
| chr4 | 14900000 | 15000000 | 0.82 | 0.93 | A>A |
| chr4 | 15000000 | 15100000 | 0.67 | 0.77 | A>A |
| chr4 | 15100000 | 15200000 | 0.70 | 0.71 | A>A |
| chr4 | 15200000 | 15300000 | 0.63 | 0.25 | A>A |
| chr4 | 15300000 | 15400000 | 0.70 | 0.15 | A>A |
| chr4 | 15400000 | 15500000 | 0.49 | 0.21 | A>A |
| chr4 | 15500000 | 15600000 | 0.84 | 0.72 | A>A |
| chr4 | 15600000 | 15700000 | 0.96 | 0.80 | A>A |
| chr4 | 15700000 | 15800000 | 0.88 | 0.47 | A>A |
| chr4 | 15800000 | 15900000 | 0.66 | 0.50 | A>A |
| chr4 | 15900000 | 16000000 | 0.91 | 0.47 | A>A |
| chr4 | 16000000 | 16100000 | 0.49 | 0.32 | A>A |
| chr4 | 16100000 | 16200000 | 0.65 | 0.48 | A>A |
| chr4 | 16200000 | 16300000 | 0.37 | 0.02 | A>A |
| chr4 | 16300000 | 16400000 | -0.16 | -0.92 | A>B |
| chr4 | 16400000 | 16500000 | -0.29 | -0.98 | A>B |
| chr4 | 16500000 | 16600000 | -0.41 | -1.09 | A>B |
| chr4 | 16600000 | 16700000 | -0.34 | -1.01 | A>B |
| chr4 | 16700000 | 16800000 | -0.33 | -1.03 | A>B |
| chr4 | 16800000 | 16900000 | -0.34 | -1.02 | A>B |
| chr4 | 16900000 | 17000000 | -0.27 | -0.96 | A>B |
| chr4 | 17000000 | 17100000 | -0.03 | -1.04 | A>B |
| chr4 | 17100000 | 17200000 | -0.24 | -1.08 | A>B |
| chr4 | 17200000 | 17300000 | -0.35 | -1.03 | A>B |
| chr4 | 17300000 | 17400000 | -0.24 | -0.92 | A>B |
| chr4 | 17400000 | 17500000 | -0.04 | -0.56 | B>B |
| chr4 | 17500000 | 17600000 | 0.50 | 0.14 | A>A |
| chr4 | 17600000 | 17700000 | 0.52 | -0.11 | A>B |
| chr4 | 17700000 | 17800000 | 0.53 | -0.28 | A>B |
| chr4 | 17800000 | 17900000 | 0.24 | 0.00 | A>A |
| chr4 | 17900000 | 18000000 | 0.32 | -0.16 | A>A |
| chr4 | 18000000 | 18100000 | -0.04 | -0.54 | B>B |
| chr4 | 18100000 | 18200000 | -0.51 | -0.92 | B>B |
| chr4 | 18200000 | 18300000 | -0.58 | -0.98 | B>B |
| chr4 | 18300000 | 18400000 | -0.72 | -0.99 | B>B |
| chr4 | 18400000 | 18500000 | -0.81 | -1.02 | B>B |
| chr4 | 18500000 | 18600000 | -0.62 | -0.82 | B>B |
| chr4 | 18600000 | 18700000 | -0.61 | -0.82 | B>B |
| chr4 | 18700000 | 18800000 | -0.74 | -0.92 | B>B |
| chr4 | 18800000 | 18900000 | -0.91 | -0.96 | B>B |
| chr4 | 18900000 | 19000000 | -0.91 | -0.96 | B>B |

| | | | | | |
|---|---|---|---|---|---|
| chr4 | 19000000 | 19100000 | -0.82 | -0.93 | B>B |
| chr4 | 19100000 | 19200000 | -0.80 | -0.94 | B>B |
| chr4 | 19200000 | 19300000 | -0.67 | -0.26 | B>B |
| chr4 | 19300000 | 19400000 | -0.75 | -0.03 | B>A |
| chr4 | 19400000 | 19500000 | -0.68 | 0.26 | B>A |
| chr4 | 19500000 | 19600000 | -0.62 | 0.35 | B>A |
| chr4 | 19600000 | 19700000 | -0.49 | 0.30 | B>A |
| chr4 | 19700000 | 19800000 | -0.36 | 0.41 | B>A |
| chr4 | 19800000 | 19900000 | -0.49 | 0.41 | B>A |
| chr4 | 19900000 | 20000000 | -0.66 | 0.39 | B>A |
| chr4 | 20000000 | 20100000 | -0.69 | 0.40 | B>A |
| chr4 | 20100000 | 20200000 | -0.57 | 0.18 | B>A |
| chr4 | 20200000 | 20300000 | -0.46 | 0.00 | B>B |
| chr4 | 20300000 | 20400000 | -0.49 | -0.07 | B>B |
| chr4 | 20400000 | 20500000 | -0.45 | -0.19 | B>B |
| chr4 | 20500000 | 20600000 | -0.44 | -0.28 | B>B |
| chr4 | 20600000 | 20700000 | -0.33 | -0.32 | B>B |
| chr4 | 20700000 | 20800000 | -0.35 | -0.26 | B>B |
| chr4 | 20800000 | 20900000 | -0.71 | -0.02 | B>A |
| chr4 | 20900000 | 21000000 | -0.73 | 0.09 | B>A |
| chr4 | 21000000 | 21100000 | -0.84 | 0.11 | B>A |
| chr4 | 21100000 | 21200000 | -0.79 | -0.31 | B>B |
| chr4 | 21200000 | 21300000 | -0.82 | -0.80 | B>B |
| chr4 | 21300000 | 21400000 | -0.87 | -0.77 | B>B |
| chr4 | 21400000 | 21500000 | -0.72 | -0.69 | B>B |
| chr4 | 21500000 | 21600000 | -0.77 | -0.38 | B>B |
| chr4 | 21600000 | 21700000 | -0.75 | -0.28 | B>B |
| chr4 | 21700000 | 21800000 | -0.53 | -0.31 | B>B |
| chr4 | 21800000 | 21900000 | -0.54 | -0.13 | B>B |
| chr4 | 21900000 | 22000000 | -0.65 | -0.14 | B>A |
| chr4 | 22000000 | 22100000 | -0.56 | -0.12 | B>B |
| chr4 | 22100000 | 22200000 | -0.57 | -0.28 | B>B |
| chr4 | 22200000 | 22300000 | -0.49 | -0.34 | B>B |
| chr4 | 22300000 | 22400000 | -0.12 | -0.11 | B>B |
| chr4 | 22400000 | 22500000 | 0.34 | 0.12 | A>A |
| chr4 | 22500000 | 22600000 | 0.15 | 0.18 | A>A |
| chr4 | 22600000 | 22700000 | -0.08 | 0.10 | A>A |
| chr4 | 22700000 | 22800000 | -0.28 | 0.06 | B>B |
| chr4 | 22800000 | 22900000 | -0.31 | 0.01 | B>B |
| chr4 | 22900000 | 23000000 | -0.30 | 0.16 | B>B |
| chr4 | 23000000 | 23100000 | -0.35 | 0.27 | B>A |
| chr4 | 23100000 | 23200000 | -0.34 | 0.23 | B>A |
| chr4 | 23200000 | 23300000 | -0.22 | 0.25 | A>A |
| chr4 | 23300000 | 23400000 | -0.27 | 0.00 | B>B |
| chr4 | 23400000 | 23500000 | -0.47 | -0.16 | B>B |
| chr4 | 23500000 | 23600000 | -0.48 | -0.20 | B>B |
| chr4 | 23600000 | 23700000 | -0.45 | -0.27 | B>B |
| chr4 | 23700000 | 23800000 | -0.47 | -0.57 | B>B |
| chr4 | 23800000 | 23900000 | -0.42 | -0.68 | B>B |
| chr4 | 23900000 | 24000000 | -0.48 | -0.84 | B>B |
| chr4 | 24000000 | 24100000 | -0.46 | -0.82 | B>B |
| chr4 | 24100000 | 24200000 | -0.41 | -0.81 | B>B |
| chr4 | 24200000 | 24300000 | -0.21 | -0.75 | B>B |
| chr4 | 24300000 | 24400000 | -0.08 | -0.56 | B>B |
| chr4 | 24400000 | 24500000 | 0.14 | -0.13 | A>A |
| chr4 | 24500000 | 24600000 | 0.90 | 0.70 | A>A |

| | | | | | |
|---|---|---|---|---|---|
| chr4 | 24600000 | 24700000 | 0.90 | 0.78 | A>A |
| chr4 | 24700000 | 24800000 | 0.75 | 0.46 | A>A |
| chr4 | 24800000 | 24900000 | 0.65 | 0.44 | A>A |
| chr4 | 24900000 | 25000000 | 0.61 | 0.38 | A>A |
| chr4 | 25000000 | 25100000 | 0.61 | 0.32 | A>A |
| chr4 | 25100000 | 25200000 | 0.74 | 0.63 | A>A |
| chr4 | 25200000 | 25300000 | 0.88 | 0.73 | A>A |
| chr4 | 25300000 | 25400000 | 0.95 | 0.86 | A>A |
| chr4 | 25400000 | 25500000 | 0.77 | 0.85 | A>A |
| chr4 | 25500000 | 25600000 | 1.03 | 1.22 | A>A |
| chr4 | 25600000 | 25700000 | 1.16 | 1.16 | A>A |
| chr4 | 25700000 | 25800000 | 0.92 | 0.83 | A>A |
| chr4 | 25800000 | 25900000 | 1.01 | 0.90 | A>A |
| chr4 | 25900000 | 26000000 | 0.74 | 0.63 | A>A |
| chr4 | 26000000 | 26100000 | 0.62 | 0.56 | A>A |
| chr4 | 26100000 | 26200000 | 0.40 | 0.44 | A>A |
| chr4 | 26200000 | 26300000 | 0.76 | 0.55 | A>A |
| chr4 | 26300000 | 26400000 | 0.78 | 0.55 | A>A |
| chr4 | 26400000 | 26500000 | 0.34 | 0.29 | A>A |
| chr4 | 26500000 | 26600000 | 0.45 | 0.46 | A>A |
| chr4 | 26600000 | 26700000 | 0.55 | 0.34 | A>A |
| chr4 | 26700000 | 26800000 | 0.30 | 0.17 | A>A |
| chr4 | 26800000 | 26900000 | 0.54 | 0.28 | A>A |
| chr4 | 26900000 | 27000000 | 0.35 | 0.24 | A>A |
| chr4 | 27000000 | 27100000 | -0.11 | -0.19 | B>B |
| chr4 | 27100000 | 27200000 | -0.32 | -0.67 | B>B |
| chr4 | 27200000 | 27300000 | -0.55 | -0.83 | B>B |
| chr4 | 27300000 | 27400000 | -0.64 | -0.90 | B>B |
| chr4 | 27400000 | 27500000 | -0.70 | -0.99 | B>B |
| chr4 | 27500000 | 27600000 | -0.69 | -1.00 | B>B |
| chr4 | 27600000 | 27700000 | -0.61 | -0.72 | B>B |
| chr4 | 27700000 | 27800000 | -0.57 | -0.35 | B>B |
| chr4 | 27800000 | 27900000 | -0.39 | -0.24 | B>B |
| chr4 | 27900000 | 28000000 | -0.17 | 0.07 | B>B |
| chr4 | 28000000 | 28100000 | 0.00 | 0.19 | A>A |
| chr4 | 28100000 | 28200000 | 0.11 | 0.14 | A>A |
| chr4 | 28200000 | 28300000 | 0.05 | 0.04 | A>A |
| chr4 | 28300000 | 28400000 | 0.12 | 0.15 | A>A |
| chr4 | 28400000 | 28500000 | 0.13 | 0.12 | A>A |
| chr4 | 28500000 | 28600000 | -0.16 | -0.25 | B>B |
| chr4 | 28600000 | 28700000 | -0.27 | -0.36 | B>B |
| chr4 | 28700000 | 28800000 | -0.49 | -0.34 | B>B |
| chr4 | 28800000 | 28900000 | -0.56 | -0.23 | B>B |
| chr4 | 28900000 | 29000000 | -0.61 | -0.26 | B>B |
| chr4 | 29000000 | 29100000 | -0.69 | -0.26 | B>B |
| chr4 | 29100000 | 29200000 | -0.68 | -0.23 | B>B |
| chr4 | 29200000 | 29300000 | -0.69 | -0.21 | B>B |
| chr4 | 29300000 | 29400000 | -0.65 | -0.21 | B>B |
| chr4 | 29400000 | 29500000 | -0.70 | -0.18 | B>A |
| chr4 | 29500000 | 29600000 | -0.67 | -0.25 | B>B |
| chr4 | 29600000 | 29700000 | -0.61 | -0.19 | B>B |
| chr4 | 29700000 | 29800000 | -0.70 | -0.27 | B>B |
| chr4 | 29800000 | 29900000 | -0.65 | -0.33 | B>B |
| chr4 | 29900000 | 30000000 | -0.66 | -0.34 | B>B |
| chr4 | 30000000 | 30100000 | -0.67 | -0.28 | B>B |
| chr4 | 30100000 | 30200000 | -0.68 | -0.25 | B>B |

| | | | | | |
|---|---|---|---|---|---|
| chr4 | 30200000 | 30300000 | -0.66 | -0.26 | B>B |
| chr4 | 30300000 | 30400000 | -0.78 | -0.10 | B>A |
| chr4 | 30400000 | 30500000 | -0.73 | 0.11 | B>A |
| chr4 | 30500000 | 30600000 | -0.75 | 0.17 | B>A |
| chr4 | 30600000 | 30700000 | -0.66 | 0.09 | B>A |
| chr4 | 30700000 | 30800000 | -0.53 | 0.40 | B>A |
| chr4 | 30800000 | 30900000 | -0.57 | 0.53 | B>A |
| chr4 | 30900000 | 31000000 | -0.42 | 0.51 | B>A |
| chr4 | 31000000 | 31100000 | -0.52 | 0.41 | B>A |
| chr4 | 31100000 | 31200000 | -0.52 | 0.56 | B>A |
| chr4 | 31200000 | 31300000 | -0.56 | 0.54 | B>A |
| chr4 | 31300000 | 31400000 | -0.63 | 0.40 | B>A |
| chr4 | 31400000 | 31500000 | -0.66 | 0.41 | B>A |
| chr4 | 31500000 | 31600000 | -0.64 | 0.47 | B>A |
| chr4 | 31600000 | 31700000 | -0.58 | 0.42 | B>A |
| chr4 | 31700000 | 31800000 | -0.53 | 0.22 | B>A |
| chr4 | 31800000 | 31900000 | -0.31 | 0.36 | B>A |
| chr4 | 31900000 | 32000000 | -0.08 | 0.39 | A>A |
| chr4 | 32000000 | 32100000 | 0.16 | 0.34 | A>A |
| chr4 | 32100000 | 32200000 | -0.15 | 0.23 | A>A |
| chr4 | 32200000 | 32300000 | -0.33 | 0.27 | B>A |
| chr4 | 32300000 | 32400000 | -0.53 | 0.28 | B>A |
| chr4 | 32400000 | 32500000 | -0.62 | 0.06 | B>A |
| chr4 | 32500000 | 32600000 | -0.61 | 0.13 | B>A |
| chr4 | 32600000 | 32700000 | -0.60 | 0.04 | B>A |
| chr4 | 32700000 | 32800000 | -0.59 | -0.03 | B>A |
| chr4 | 32800000 | 32900000 | -0.64 | -0.08 | B>A |
| chr4 | 32900000 | 33000000 | -0.63 | -0.15 | B>B |
| chr4 | 33000000 | 33100000 | -0.53 | -0.16 | B>B |
| chr4 | 33100000 | 33200000 | -0.42 | -0.19 | B>B |
| chr4 | 33200000 | 33300000 | -0.32 | -0.10 | B>B |
| chr4 | 33300000 | 33400000 | -0.17 | -0.03 | B>B |
| chr4 | 33400000 | 33500000 | -0.02 | -0.01 | B>B |
| chr4 | 33500000 | 33600000 | -0.03 | -0.03 | B>B |
| chr4 | 33600000 | 33700000 | 0.13 | -0.13 | A>A |
| chr4 | 33700000 | 33800000 | 0.30 | -0.05 | A>A |
| chr4 | 33800000 | 33900000 | 0.20 | -0.15 | A>A |
| chr4 | 33900000 | 34000000 | 0.40 | 0.07 | A>A |
| chr4 | 34000000 | 34100000 | 0.40 | -0.10 | A>A |
| chr4 | 34100000 | 34200000 | 0.31 | -0.02 | A>A |
| chr4 | 34200000 | 34300000 | 0.30 | -0.05 | A>A |
| chr4 | 34300000 | 34400000 | 0.26 | -0.07 | A>A |
| chr4 | 34400000 | 34500000 | 0.12 | -0.04 | A>A |
| chr4 | 34500000 | 34600000 | -0.05 | 0.04 | B>B |
| chr4 | 34600000 | 34700000 | -0.11 | -0.07 | B>B |
| chr4 | 34700000 | 34800000 | -0.26 | -0.17 | B>B |
| chr4 | 34800000 | 34900000 | -0.34 | -0.08 | B>B |
| chr4 | 34900000 | 35000000 | -0.25 | -0.16 | B>B |
| chr4 | 35000000 | 35100000 | -0.25 | -0.18 | B>B |
| chr4 | 35100000 | 35200000 | -0.43 | -0.20 | B>B |
| chr4 | 35200000 | 35300000 | -0.60 | -0.17 | B>B |
| chr4 | 35300000 | 35400000 | -0.61 | -0.19 | B>B |
| chr4 | 35400000 | 35500000 | -0.72 | -0.25 | B>B |
| chr4 | 35500000 | 35600000 | -0.71 | -0.17 | B>A |
| chr4 | 35600000 | 35700000 | -0.72 | -0.14 | B>A |
| chr4 | 35700000 | 35800000 | -0.73 | -0.18 | B>A |

| | | | | | |
|---|---|---|---|---|---|
| chr4 | 35800000 | 35900000 | -0.66 | -0.21 | B>B |
| chr4 | 35900000 | 36000000 | -0.62 | -0.07 | B>A |
| chr4 | 36000000 | 36100000 | -0.66 | 0.22 | B>A |
| chr4 | 36100000 | 36200000 | -0.52 | 0.07 | B>A |
| chr4 | 36200000 | 36300000 | -0.58 | 0.13 | B>A |
| chr4 | 36300000 | 36400000 | -0.58 | -0.14 | B>B |
| chr4 | 36400000 | 36500000 | -0.67 | -0.71 | B>B |
| chr4 | 36500000 | 36600000 | -0.81 | -0.97 | B>B |
| chr4 | 36600000 | 36700000 | -0.71 | -1.01 | B>B |
| chr4 | 36700000 | 36800000 | -0.66 | -0.96 | B>B |
| chr4 | 36800000 | 36900000 | -0.75 | -0.98 | B>B |
| chr4 | 36900000 | 37000000 | -0.58 | -0.97 | B>B |
| chr4 | 37000000 | 37100000 | -0.50 | -0.93 | B>B |
| chr4 | 37100000 | 37200000 | -0.57 | -1.01 | B>B |
| chr4 | 37200000 | 37300000 | -0.50 | -0.89 | B>B |
| chr4 | 37300000 | 37400000 | -0.46 | -0.68 | B>B |
| chr4 | 37400000 | 37500000 | 0.02 | -0.17 | B>B |
| chr4 | 37500000 | 37600000 | 0.26 | -0.14 | A>A |
| chr4 | 37600000 | 37700000 | 0.78 | 0.37 | A>A |
| chr4 | 37700000 | 37800000 | 0.52 | 0.36 | A>A |
| chr4 | 37800000 | 37900000 | 0.65 | 0.44 | A>A |
| chr4 | 37900000 | 38000000 | 0.83 | 0.53 | A>A |
| chr4 | 38000000 | 38100000 | 0.70 | 0.56 | A>A |
| chr4 | 38100000 | 38200000 | 0.84 | 0.83 | A>A |
| chr4 | 38200000 | 38300000 | 0.95 | 0.93 | A>A |
| chr4 | 38300000 | 38400000 | 0.88 | 0.87 | A>A |
| chr4 | 38400000 | 38500000 | 1.03 | 0.91 | A>A |
| chr4 | 38500000 | 38600000 | 1.04 | 1.07 | A>A |
| chr4 | 38600000 | 38700000 | 0.90 | 0.80 | A>A |
| chr4 | 38700000 | 38800000 | 0.69 | 0.63 | A>A |
| chr4 | 38800000 | 38900000 | 0.94 | 0.96 | A>A |
| chr4 | 38900000 | 39000000 | 1.15 | 0.95 | A>A |
| chr4 | 39000000 | 39100000 | 1.27 | 1.28 | A>A |
| chr4 | 39100000 | 39200000 | 0.98 | 0.97 | A>A |
| chr4 | 39200000 | 39300000 | 1.04 | 1.02 | A>A |
| chr4 | 39300000 | 39400000 | 1.23 | 1.22 | A>A |
| chr4 | 39400000 | 39500000 | 1.11 | 1.12 | A>A |
| chr4 | 39500000 | 39600000 | 1.38 | 1.30 | A>A |
| chr4 | 39600000 | 39700000 | 1.28 | 1.14 | A>A |
| chr4 | 39700000 | 39800000 | 1.28 | 0.99 | A>A |
| chr4 | 39800000 | 39900000 | 1.25 | 1.05 | A>A |
| chr4 | 39900000 | 40000000 | 1.12 | 0.99 | A>A |
| chr4 | 40000000 | 40100000 | 1.07 | 0.92 | A>A |
| chr4 | 40100000 | 40200000 | 1.12 | 1.03 | A>A |
| chr4 | 40200000 | 40300000 | 1.29 | 1.22 | A>A |
| chr4 | 40300000 | 40400000 | 0.97 | 0.95 | A>A |
| chr4 | 40400000 | 40500000 | 1.31 | 1.17 | A>A |
| chr4 | 40500000 | 40600000 | 1.29 | 1.07 | A>A |
| chr4 | 40600000 | 40700000 | 1.34 | 1.23 | A>A |
| chr4 | 40700000 | 40800000 | 0.94 | 1.05 | A>A |
| chr4 | 40800000 | 40900000 | 1.21 | 1.12 | A>A |
| chr4 | 40900000 | 41000000 | 1.03 | 1.03 | A>A |
| chr4 | 41000000 | 41100000 | 1.50 | 1.34 | A>A |
| chr4 | 41100000 | 41200000 | 1.27 | 1.27 | A>A |
| chr4 | 41200000 | 41300000 | 0.91 | 0.90 | A>A |
| chr4 | 41300000 | 41400000 | 1.05 | 1.11 | A>A |

| | | | | | |
|---|---|---|---|---|---|
| chr4 | 41400000 | 41500000 | 1.32 | 1.26 | A>A |
| chr4 | 41500000 | 41600000 | 1.26 | 1.21 | A>A |
| chr4 | 41600000 | 41700000 | 0.97 | 0.95 | A>A |
| chr4 | 41700000 | 41800000 | 0.96 | 0.75 | A>A |
| chr4 | 41800000 | 41900000 | 1.00 | 0.87 | A>A |
| chr4 | 41900000 | 42000000 | 0.94 | 0.65 | A>A |
| chr4 | 42000000 | 42100000 | 0.66 | 0.49 | A>A |
| chr4 | 42100000 | 42200000 | 0.47 | 0.34 | A>A |
| chr4 | 42200000 | 42300000 | 0.37 | 0.17 | A>A |
| chr4 | 42300000 | 42400000 | 0.26 | 0.01 | A>A |
| chr4 | 42400000 | 42500000 | 0.11 | 0.04 | A>A |
| chr4 | 42500000 | 42600000 | -0.11 | -0.44 | B>B |
| chr4 | 42600000 | 42700000 | -0.09 | -0.81 | A>B |
| chr4 | 42700000 | 42800000 | -0.52 | -1.03 | B>B |
| chr4 | 42800000 | 42900000 | -0.55 | -0.93 | B>B |
| chr4 | 42900000 | 43000000 | -0.76 | -0.98 | B>B |
| chr4 | 43000000 | 43100000 | -0.68 | -0.98 | B>B |
| chr4 | 43100000 | 43200000 | -0.76 | -0.93 | B>B |
| chr4 | 43200000 | 43300000 | -0.72 | -0.99 | B>B |
| chr4 | 43300000 | 43400000 | -0.80 | -0.89 | B>B |
| chr4 | 43400000 | 43500000 | -0.76 | -0.97 | B>B |
| chr4 | 43500000 | 43600000 | -0.79 | -0.87 | B>B |
| chr4 | 43600000 | 43700000 | -0.75 | -0.86 | B>B |
| chr4 | 43700000 | 43800000 | -0.72 | -0.90 | B>B |
| chr4 | 43800000 | 43900000 | -0.69 | -0.86 | B>B |
| chr4 | 43900000 | 44000000 | -0.73 | -0.85 | B>B |
| chr4 | 44000000 | 44100000 | -0.77 | -0.91 | B>B |
| chr4 | 44100000 | 44200000 | -0.74 | -0.89 | B>B |
| chr4 | 44200000 | 44300000 | -0.73 | -0.85 | B>B |
| chr4 | 44300000 | 44400000 | -0.69 | -0.83 | B>B |
| chr4 | 44400000 | 44500000 | -0.56 | -0.86 | B>B |
| chr4 | 44500000 | 44600000 | -0.67 | -0.86 | B>B |
| chr4 | 44600000 | 44700000 | -0.28 | -0.54 | B>B |
| chr4 | 44700000 | 44800000 | -0.30 | -0.50 | B>B |
| chr4 | 44800000 | 44900000 | -0.61 | -0.82 | B>B |
| chr4 | 44900000 | 45000000 | -0.70 | -0.79 | B>B |
| chr4 | 45000000 | 45100000 | -0.73 | -0.83 | B>B |
| chr4 | 45100000 | 45200000 | -0.72 | -0.80 | B>B |
| chr4 | 45200000 | 45300000 | -0.73 | -0.86 | B>B |
| chr4 | 45300000 | 45400000 | -0.80 | -0.80 | B>B |
| chr4 | 45400000 | 45500000 | -0.62 | -0.77 | B>B |
| chr4 | 45500000 | 45600000 | -0.71 | -0.67 | B>B |
| chr4 | 45600000 | 45700000 | -0.66 | -0.58 | B>B |
| chr4 | 45700000 | 45800000 | -0.78 | -0.78 | B>B |
| chr4 | 45800000 | 45900000 | -0.76 | -0.72 | B>B |
| chr4 | 45900000 | 46000000 | -0.74 | -0.79 | B>B |
| chr4 | 46000000 | 46100000 | -0.73 | -0.75 | B>B |
| chr4 | 46100000 | 46200000 | -0.74 | -0.77 | B>B |
| chr4 | 46200000 | 46300000 | -0.69 | -0.76 | B>B |
| chr4 | 46300000 | 46400000 | -0.65 | -0.80 | B>B |
| chr4 | 46400000 | 46500000 | -0.67 | -0.87 | B>B |
| chr4 | 46500000 | 46600000 | -0.72 | -0.91 | B>B |
| chr4 | 46600000 | 46700000 | -0.69 | -0.85 | B>B |
| chr4 | 46700000 | 46800000 | -0.73 | -0.85 | B>B |
| chr4 | 46800000 | 46900000 | -0.59 | -0.84 | B>B |
| chr4 | 46900000 | 47000000 | -0.55 | -0.93 | B>B |

| | | | | | |
|---|---|---|---|---|---|
| chr4 | 47000000 | 47100000 | -0.57 | -0.89 | B>B |
| chr4 | 47100000 | 47200000 | -0.59 | -0.89 | B>B |
| chr4 | 47200000 | 47300000 | -0.68 | -0.94 | B>B |
| chr4 | 47300000 | 47400000 | -0.43 | -0.85 | B>B |
| chr4 | 47400000 | 47500000 | 0.04 | -0.49 | B>B |
| chr4 | 47500000 | 47600000 | 0.13 | -0.30 | B>B |
| chr4 | 47600000 | 47700000 | -0.05 | -0.53 | B>B |
| chr4 | 47700000 | 47800000 | 0.17 | -0.39 | B>B |
| chr4 | 47800000 | 47900000 | 0.46 | -0.08 | A>A |
| chr4 | 47900000 | 48000000 | 0.54 | 0.03 | A>A |
| chr4 | 48000000 | 48100000 | 0.65 | 0.36 | A>A |
| chr4 | 48100000 | 48200000 | 0.64 | 0.49 | A>A |
| chr4 | 48200000 | 48300000 | 0.99 | 0.72 | A>A |
| chr4 | 48300000 | 48400000 | 1.00 | 0.78 | A>A |
| chr4 | 48400000 | 48500000 | 0.88 | 0.68 | A>A |
| chr4 | 48500000 | 48600000 | 0.76 | 0.57 | A>A |
| chr4 | 48600000 | 48700000 | 1.00 | 0.73 | A>A |
| chr4 | 48700000 | 48800000 | 1.04 | 0.80 | A>A |
| chr4 | 48800000 | 48900000 | 0.84 | 0.59 | A>A |
| chr4 | 48900000 | 49000000 | 0.44 | 0.18 | A>A |
| chr4 | 49000000 | 49100000 | 0.00 | -0.69 | NO |
| chr4 | 49100000 | 49200000 | 0.00 | 0.00 | NO |
| chr4 | 49200000 | 49300000 | 0.00 | 0.00 | NO |
| chr4 | 49300000 | 49400000 | 0.00 | 0.00 | NO |
| chr4 | 49400000 | 49500000 | 0.00 | 0.00 | NO |
| chr4 | 49500000 | 49600000 | 0.00 | 0.00 | NO |
| chr4 | 49600000 | 49700000 | 0.00 | 0.00 | NO |
| chr4 | 49700000 | 49800000 | 0.00 | 0.00 | NO |
| chr4 | 49800000 | 49900000 | 0.00 | 0.00 | NO |
| chr4 | 49900000 | 50000000 | 0.00 | 0.00 | NO |
| chr4 | 50000000 | 50100000 | 0.00 | 0.00 | NO |
| chr4 | 50100000 | 50200000 | 0.00 | 0.00 | NO |
| chr4 | 50200000 | 50300000 | 0.00 | 0.00 | NO |
| chr4 | 50300000 | 50400000 | 0.00 | 0.00 | NO |
| chr4 | 50400000 | 50500000 | 0.00 | 0.00 | NO |
| chr4 | 50500000 | 50600000 | 0.00 | 0.00 | NO |
| chr4 | 50600000 | 50700000 | 0.00 | 0.00 | NO |
| chr4 | 50700000 | 50800000 | 0.00 | 0.00 | NO |
| chr4 | 50800000 | 50900000 | 0.00 | 0.00 | NO |
| chr4 | 50900000 | 51000000 | 0.00 | 0.00 | NO |
| chr4 | 51000000 | 51100000 | 0.00 | 0.00 | NO |
| chr4 | 51100000 | 51200000 | 0.00 | 0.00 | NO |
| chr4 | 51200000 | 51300000 | 0.00 | 0.00 | NO |
| chr4 | 51300000 | 51400000 | 0.00 | 0.00 | NO |
| chr4 | 51400000 | 51500000 | 0.00 | 0.00 | NO |
| chr4 | 51500000 | 51600000 | 0.00 | 0.00 | NO |
| chr4 | 51600000 | 51700000 | 0.00 | 0.00 | NO |
| chr4 | 51700000 | 51800000 | 0.00 | 0.00 | NO |
| chr4 | 51800000 | 51900000 | 0.00 | 0.00 | NO |
| chr4 | 51900000 | 52000000 | 0.00 | 0.00 | NO |
| chr4 | 52000000 | 52100000 | 0.00 | 0.00 | NO |
| chr4 | 52100000 | 52200000 | 0.00 | 0.00 | NO |
| chr4 | 52200000 | 52300000 | 0.00 | 0.00 | NO |
| chr4 | 52300000 | 52400000 | 0.00 | 0.00 | NO |
| chr4 | 52400000 | 52500000 | 0.00 | 0.00 | NO |
| chr4 | 52500000 | 52600000 | 0.00 | 0.00 | NO |

| | | | | | |
|---|---|---|---|---|---|
| chr4 | 52600000 | 52700000 | 0.00 | 0.00 | NO |
| chr4 | 52700000 | 52800000 | 0.41 | 0.10 | A>A |
| chr4 | 52800000 | 52900000 | 0.34 | -0.03 | A>A |
| chr4 | 52900000 | 53000000 | 0.10 | -0.42 | B>B |
| chr4 | 53000000 | 53100000 | 0.07 | -0.63 | A>B |
| chr4 | 53100000 | 53200000 | -0.19 | -0.77 | B>B |
| chr4 | 53200000 | 53300000 | -0.10 | -0.78 | A>B |
| chr4 | 53300000 | 53400000 | 0.00 | -0.69 | NO |
| chr4 | 53400000 | 53500000 | 0.22 | -0.22 | B>B |
| chr4 | 53500000 | 53600000 | 0.53 | -0.07 | A>B |
| chr4 | 53600000 | 53700000 | 0.41 | -0.09 | A>A |
| chr4 | 53700000 | 53800000 | 0.25 | -0.17 | A>A |
| chr4 | 53800000 | 53900000 | 0.15 | -0.35 | B>B |
| chr4 | 53900000 | 54000000 | 0.28 | -0.16 | A>A |
| chr4 | 54000000 | 54100000 | 0.29 | -0.12 | A>A |
| chr4 | 54100000 | 54200000 | 0.55 | 0.20 | A>A |
| chr4 | 54200000 | 54300000 | 0.78 | 0.37 | A>A |
| chr4 | 54300000 | 54400000 | 0.64 | 0.26 | A>A |
| chr4 | 54400000 | 54500000 | 0.77 | 0.23 | A>A |
| chr4 | 54500000 | 54600000 | 0.73 | 0.16 | A>A |
| chr4 | 54600000 | 54700000 | 0.76 | 0.18 | A>A |
| chr4 | 54700000 | 54800000 | 0.62 | 0.16 | A>A |
| chr4 | 54800000 | 54900000 | 0.76 | 0.30 | A>A |
| chr4 | 54900000 | 55000000 | 0.91 | 0.44 | A>A |
| chr4 | 55000000 | 55100000 | 0.48 | 0.07 | A>A |
| chr4 | 55100000 | 55200000 | 0.05 | -0.54 | A>B |
| chr4 | 55200000 | 55300000 | -0.12 | -0.90 | A>B |
| chr4 | 55300000 | 55400000 | -0.04 | -0.97 | A>B |
| chr4 | 55400000 | 55500000 | -0.06 | -1.06 | A>B |
| chr4 | 55500000 | 55600000 | -0.18 | -0.91 | A>B |
| chr4 | 55600000 | 55700000 | -0.39 | -0.97 | B>B |
| chr4 | 55700000 | 55800000 | -0.40 | -0.93 | B>B |
| chr4 | 55800000 | 55900000 | -0.26 | -0.76 | B>B |
| chr4 | 55900000 | 56000000 | -0.17 | -0.73 | B>B |
| chr4 | 56000000 | 56100000 | -0.05 | -0.48 | B>B |
| chr4 | 56100000 | 56200000 | -0.12 | -0.42 | B>B |
| chr4 | 56200000 | 56300000 | 0.47 | 0.20 | A>A |
| chr4 | 56300000 | 56400000 | 0.77 | 0.24 | A>A |
| chr4 | 56400000 | 56500000 | 0.45 | -0.08 | A>A |
| chr4 | 56500000 | 56600000 | 0.20 | -0.31 | B>B |
| chr4 | 56600000 | 56700000 | 0.36 | -0.30 | A>B |
| chr4 | 56700000 | 56800000 | 0.64 | 0.06 | A>A |
| chr4 | 56800000 | 56900000 | 0.87 | 0.15 | A>B |
| chr4 | 56900000 | 57000000 | 0.92 | -0.11 | A>B |
| chr4 | 57000000 | 57100000 | 0.88 | -0.30 | A>B |
| chr4 | 57100000 | 57200000 | 0.89 | -0.14 | A>B |
| chr4 | 57200000 | 57300000 | 1.13 | 0.52 | A>B |
| chr4 | 57300000 | 57400000 | 1.27 | 0.79 | A>A |
| chr4 | 57400000 | 57500000 | 0.98 | 0.45 | A>A |
| chr4 | 57500000 | 57600000 | 1.07 | 0.68 | A>A |
| chr4 | 57600000 | 57700000 | 0.90 | 0.60 | A>A |
| chr4 | 57700000 | 57800000 | 1.02 | 0.68 | A>A |
| chr4 | 57800000 | 57900000 | 0.82 | 0.70 | A>A |
| chr4 | 57900000 | 58000000 | 0.68 | 0.52 | A>A |
| chr4 | 58000000 | 58100000 | 0.16 | 0.04 | A>A |
| chr4 | 58100000 | 58200000 | -0.06 | -0.26 | B>B |

| | | | | | |
|---|---|---|---|---|---|
| chr4 | 58200000 | 58300000 | -0.37 | -0.18 | B>B |
| chr4 | 58300000 | 58400000 | -0.49 | -0.24 | B>B |
| chr4 | 58400000 | 58500000 | -0.66 | -0.27 | B>B |
| chr4 | 58500000 | 58600000 | -0.69 | -0.26 | B>B |
| chr4 | 58600000 | 58700000 | -0.71 | -0.29 | B>B |
| chr4 | 58700000 | 58800000 | -0.68 | -0.28 | B>B |
| chr4 | 58800000 | 58900000 | -0.68 | -0.26 | B>B |
| chr4 | 58900000 | 59000000 | -0.70 | -0.21 | B>B |
| chr4 | 59000000 | 59100000 | -0.71 | -0.23 | B>B |
| chr4 | 59100000 | 59200000 | -0.63 | -0.20 | B>B |
| chr4 | 59200000 | 59300000 | -0.62 | -0.18 | B>B |
| chr4 | 59300000 | 59400000 | -0.63 | -0.07 | B>A |
| chr4 | 59400000 | 59500000 | -0.53 | 0.02 | B>A |
| chr4 | 59500000 | 59600000 | -0.46 | 0.19 | B>A |
| chr4 | 59600000 | 59700000 | -0.31 | 0.28 | B>A |
| chr4 | 59700000 | 59800000 | 0.00 | 0.00 | NO |
| chr4 | 59800000 | 59900000 | -0.44 | 0.39 | B>A |
| chr4 | 59900000 | 60000000 | -0.48 | 0.26 | B>A |
| chr4 | 60000000 | 60100000 | -0.52 | 0.21 | B>A |
| chr4 | 60100000 | 60200000 | -0.66 | -0.03 | B>A |
| chr4 | 60200000 | 60300000 | -0.67 | -0.01 | B>A |
| chr4 | 60300000 | 60400000 | -0.59 | 0.12 | B>A |
| chr4 | 60400000 | 60500000 | -0.51 | 0.11 | B>A |
| chr4 | 60500000 | 60600000 | -0.63 | 0.00 | B>A |
| chr4 | 60600000 | 60700000 | -0.67 | -0.05 | B>A |
| chr4 | 60700000 | 60800000 | -0.60 | -0.06 | B>A |
| chr4 | 60800000 | 60900000 | -0.60 | -0.13 | B>B |
| chr4 | 60900000 | 61000000 | -0.67 | -0.05 | B>A |
| chr4 | 61000000 | 61100000 | -0.62 | -0.11 | B>A |
| chr4 | 61100000 | 61200000 | -0.59 | -0.12 | B>B |
| chr4 | 61200000 | 61300000 | -0.72 | -0.12 | B>A |
| chr4 | 61300000 | 61400000 | -0.61 | -0.20 | B>B |
| chr4 | 61400000 | 61500000 | -0.64 | -0.11 | B>A |
| chr4 | 61500000 | 61600000 | -0.64 | -0.03 | B>A |
| chr4 | 61600000 | 61700000 | -0.69 | 0.01 | B>A |
| chr4 | 61700000 | 61800000 | -0.62 | -0.04 | B>A |
| chr4 | 61800000 | 61900000 | -0.68 | -0.03 | B>A |
| chr4 | 61900000 | 62000000 | -0.63 | 0.08 | B>A |
| chr4 | 62000000 | 62100000 | -0.62 | 0.07 | B>A |
| chr4 | 62100000 | 62200000 | -0.62 | 0.07 | B>A |
| chr4 | 62200000 | 62300000 | -0.66 | 0.20 | B>A |
| chr4 | 62300000 | 62400000 | -0.67 | 0.15 | B>A |
| chr4 | 62400000 | 62500000 | -0.72 | 0.13 | B>A |
| chr4 | 62500000 | 62600000 | -0.63 | 0.06 | B>A |
| chr4 | 62600000 | 62700000 | -0.65 | 0.01 | B>A |
| chr4 | 62700000 | 62800000 | -0.65 | 0.00 | B>A |
| chr4 | 62800000 | 62900000 | -0.67 | -0.03 | B>A |
| chr4 | 62900000 | 63000000 | -0.70 | -0.07 | B>A |
| chr4 | 63000000 | 63100000 | -0.67 | -0.16 | B>A |
| chr4 | 63100000 | 63200000 | -0.66 | -0.07 | B>A |
| chr4 | 63200000 | 63300000 | -0.68 | -0.10 | B>A |
| chr4 | 63300000 | 63400000 | -0.65 | -0.08 | B>A |
| chr4 | 63400000 | 63500000 | -0.65 | 0.09 | B>A |
| chr4 | 63500000 | 63600000 | -0.67 | 0.05 | B>A |
| chr4 | 63600000 | 63700000 | -0.66 | 0.31 | B>A |
| chr4 | 63700000 | 63800000 | -0.67 | 0.42 | B>A |

| | | | | | |
|---|---|---|---|---|---|
| chr4 | 63800000 | 63900000 | -0.73 | 0.42 | B>A |
| chr4 | 63900000 | 64000000 | -0.75 | 0.42 | B>A |
| chr4 | 64000000 | 64100000 | -0.69 | 0.44 | B>A |
| chr4 | 64100000 | 64200000 | -0.72 | 0.31 | B>A |
| chr4 | 64200000 | 64300000 | -0.71 | 0.32 | B>A |
| chr4 | 64300000 | 64400000 | -0.70 | 0.24 | B>A |
| chr4 | 64400000 | 64500000 | -0.66 | 0.26 | B>A |
| chr4 | 64500000 | 64600000 | -0.74 | 0.37 | B>A |
| chr4 | 64600000 | 64700000 | -0.69 | 0.41 | B>A |
| chr4 | 64700000 | 64800000 | -0.73 | 0.44 | B>A |
| chr4 | 64800000 | 64900000 | -0.78 | 0.59 | B>A |
| chr4 | 64900000 | 65000000 | -0.78 | 0.33 | B>A |
| chr4 | 65000000 | 65100000 | -0.74 | 0.23 | B>A |
| chr4 | 65100000 | 65200000 | -0.77 | 0.11 | B>A |
| chr4 | 65200000 | 65300000 | -0.70 | -0.02 | B>A |
| chr4 | 65300000 | 65400000 | -0.76 | -0.01 | B>A |
| chr4 | 65400000 | 65500000 | -0.72 | -0.16 | B>A |
| chr4 | 65500000 | 65600000 | -0.81 | -0.41 | B>B |
| chr4 | 65600000 | 65700000 | -0.71 | -0.38 | B>B |
| chr4 | 65700000 | 65800000 | -0.74 | -0.42 | B>B |
| chr4 | 65800000 | 65900000 | -0.75 | -0.38 | B>B |
| chr4 | 65900000 | 66000000 | -0.70 | -0.14 | B>A |
| chr4 | 66000000 | 66100000 | -0.68 | 0.01 | B>A |
| chr4 | 66100000 | 66200000 | -0.67 | 0.10 | B>A |
| chr4 | 66200000 | 66300000 | -0.54 | 0.16 | B>A |
| chr4 | 66300000 | 66400000 | -0.58 | 0.11 | B>A |
| chr4 | 66400000 | 66500000 | -0.50 | 0.03 | B>A |
| chr4 | 66500000 | 66600000 | -0.46 | 0.19 | B>A |
| chr4 | 66600000 | 66700000 | -0.44 | -0.12 | B>B |
| chr4 | 66700000 | 66800000 | -0.28 | -0.08 | B>B |
| chr4 | 66800000 | 66900000 | -0.13 | -0.19 | B>B |
| chr4 | 66900000 | 67000000 | -0.14 | -0.25 | B>B |
| chr4 | 67000000 | 67100000 | -0.19 | -0.25 | B>B |
| chr4 | 67100000 | 67200000 | -0.02 | -0.20 | B>B |
| chr4 | 67200000 | 67300000 | 0.11 | -0.15 | B>B |
| chr4 | 67300000 | 67400000 | 0.24 | -0.02 | A>A |
| chr4 | 67400000 | 67500000 | 0.30 | 0.06 | A>A |
| chr4 | 67500000 | 67600000 | 0.06 | -0.23 | B>B |
| chr4 | 67600000 | 67700000 | -0.17 | -0.54 | B>B |
| chr4 | 67700000 | 67800000 | -0.32 | -0.57 | B>B |
| chr4 | 67800000 | 67900000 | -0.46 | -0.61 | B>B |
| chr4 | 67900000 | 68000000 | -0.54 | -0.68 | B>B |
| chr4 | 68000000 | 68100000 | -0.60 | -0.66 | B>B |
| chr4 | 68100000 | 68200000 | -0.51 | -0.64 | B>B |
| chr4 | 68200000 | 68300000 | -0.35 | -0.40 | B>B |
| chr4 | 68300000 | 68400000 | 0.26 | -0.15 | A>A |
| chr4 | 68400000 | 68500000 | 0.48 | -0.15 | A>B |
| chr4 | 68500000 | 68600000 | 0.52 | 0.02 | A>A |
| chr4 | 68600000 | 68700000 | -0.14 | -0.62 | B>B |
| chr4 | 68700000 | 68800000 | -0.41 | -0.63 | B>B |
| chr4 | 68800000 | 68900000 | -0.42 | -0.73 | B>B |
| chr4 | 68900000 | 69000000 | -0.45 | -0.74 | B>B |
| chr4 | 69000000 | 69100000 | -0.23 | -0.48 | B>B |
| chr4 | 69100000 | 69200000 | 0.00 | 0.00 | NO |
| chr4 | 69200000 | 69300000 | 0.00 | 0.00 | NO |
| chr4 | 69300000 | 69400000 | 0.00 | 0.00 | NO |

| | | | | | |
|---|---|---|---|---|---|
| chr4 | 69400000 | 69500000 | 0.00 | 0.00 | NO |
| chr4 | 69500000 | 69600000 | 0.00 | 0.00 | NO |
| chr4 | 69600000 | 69700000 | 0.00 | 0.00 | NO |
| chr4 | 69700000 | 69800000 | 0.00 | 0.00 | NO |
| chr4 | 69800000 | 69900000 | 0.00 | 0.00 | NO |
| chr4 | 69900000 | 70000000 | 0.00 | 0.89 | NO |
| chr4 | 70000000 | 70100000 | 0.00 | 0.00 | NO |
| chr4 | 70100000 | 70200000 | 0.00 | 0.00 | NO |
| chr4 | 70200000 | 70300000 | 0.00 | 0.00 | NO |
| chr4 | 70300000 | 70400000 | -0.22 | 0.30 | B>A |
| chr4 | 70400000 | 70500000 | -0.13 | 0.09 | B>B |
| chr4 | 70500000 | 70600000 | -0.35 | -0.59 | B>B |
| chr4 | 70600000 | 70700000 | -0.78 | -0.98 | B>B |
| chr4 | 70700000 | 70800000 | -0.66 | -0.98 | B>B |
| chr4 | 70800000 | 70900000 | -0.71 | -1.03 | B>B |
| chr4 | 70900000 | 71000000 | -0.80 | -1.07 | B>B |
| chr4 | 71000000 | 71100000 | -0.73 | -1.10 | B>B |
| chr4 | 71100000 | 71200000 | -0.73 | -1.00 | B>B |
| chr4 | 71200000 | 71300000 | -0.74 | -0.96 | B>B |
| chr4 | 71300000 | 71400000 | -0.46 | -0.86 | B>B |
| chr4 | 71400000 | 71500000 | -0.01 | -0.31 | B>B |
| chr4 | 71500000 | 71600000 | 0.61 | 0.31 | A>A |
| chr4 | 71600000 | 71700000 | 0.89 | 0.57 | A>A |
| chr4 | 71700000 | 71800000 | 0.87 | 0.52 | A>A |
| chr4 | 71800000 | 71900000 | 0.74 | 0.46 | A>A |
| chr4 | 71900000 | 72000000 | 0.33 | 0.18 | A>A |
| chr4 | 72000000 | 72100000 | 0.23 | 0.19 | A>A |
| chr4 | 72100000 | 72200000 | 0.27 | 0.26 | A>A |
| chr4 | 72200000 | 72300000 | 0.37 | 0.38 | A>A |
| chr4 | 72300000 | 72400000 | 0.38 | 0.34 | A>A |
| chr4 | 72400000 | 72500000 | 0.51 | 0.37 | A>A |
| chr4 | 72500000 | 72600000 | 0.22 | 0.25 | A>A |
| chr4 | 72600000 | 72700000 | -0.39 | -0.32 | B>B |
| chr4 | 72700000 | 72800000 | -0.60 | -0.20 | B>B |
| chr4 | 72800000 | 72900000 | -0.57 | -0.17 | B>B |
| chr4 | 72900000 | 73000000 | -0.72 | -0.09 | B>A |
| chr4 | 73000000 | 73100000 | -0.71 | -0.19 | B>A |
| chr4 | 73100000 | 73200000 | -0.50 | -0.40 | B>B |
| chr4 | 73200000 | 73300000 | -0.65 | -0.93 | B>B |
| chr4 | 73300000 | 73400000 | -0.68 | -0.97 | B>B |
| chr4 | 73400000 | 73500000 | -0.44 | -0.92 | B>B |
| chr4 | 73500000 | 73600000 | -0.47 | -0.95 | B>B |
| chr4 | 73600000 | 73700000 | -0.40 | -0.99 | A>B |
| chr4 | 73700000 | 73800000 | -0.33 | -1.00 | A>B |
| chr4 | 73800000 | 73900000 | -0.09 | -0.60 | B>B |
| chr4 | 73900000 | 74000000 | 0.51 | 0.22 | A>A |
| chr4 | 74000000 | 74100000 | 0.99 | 0.69 | A>A |
| chr4 | 74100000 | 74200000 | 0.71 | 0.59 | A>A |
| chr4 | 74200000 | 74300000 | 0.74 | 0.81 | A>A |
| chr4 | 74300000 | 74400000 | 0.86 | 0.68 | A>A |
| chr4 | 74400000 | 74500000 | 0.83 | 0.70 | A>A |
| chr4 | 74500000 | 74600000 | 0.82 | 0.71 | A>A |
| chr4 | 74600000 | 74700000 | 1.11 | 1.16 | A>A |
| chr4 | 74700000 | 74800000 | 1.02 | 0.91 | A>A |
| chr4 | 74800000 | 74900000 | 1.02 | 0.91 | A>A |
| chr4 | 74900000 | 75000000 | 1.13 | 0.88 | A>A |

| | | | | | |
|---|---|---|---|---|---|
| chr4 | 75000000 | 75100000 | 0.68 | 0.51 | A>A |
| chr4 | 75100000 | 75200000 | 0.66 | 0.49 | A>A |
| chr4 | 75200000 | 75300000 | 1.01 | 0.61 | A>A |
| chr4 | 75300000 | 75400000 | 1.08 | 0.69 | A>A |
| chr4 | 75400000 | 75500000 | 0.00 | 0.00 | NO |
| chr4 | 75500000 | 75600000 | 0.99 | 0.60 | A>A |
| chr4 | 75600000 | 75700000 | 0.80 | 0.21 | A>B |
| chr4 | 75700000 | 75800000 | 0.57 | -0.20 | A>B |
| chr4 | 75800000 | 75900000 | 0.32 | -0.31 | A>B |
| chr4 | 75900000 | 76000000 | 0.45 | -0.43 | A>B |
| chr4 | 76000000 | 76100000 | 0.24 | -0.40 | A>B |
| chr4 | 76100000 | 76200000 | 0.15 | -0.57 | A>B |
| chr4 | 76200000 | 76300000 | 0.19 | -0.60 | A>B |
| chr4 | 76300000 | 76400000 | 0.23 | -0.52 | A>B |
| chr4 | 76400000 | 76500000 | 0.82 | 0.15 | A>B |
| chr4 | 76500000 | 76600000 | 1.15 | 0.64 | A>A |
| chr4 | 76600000 | 76700000 | 1.50 | 0.92 | A>A |
| chr4 | 76700000 | 76800000 | 1.24 | 0.86 | A>A |
| chr4 | 76800000 | 76900000 | 1.26 | 0.96 | A>A |
| chr4 | 76900000 | 77000000 | 1.28 | 0.80 | A>A |
| chr4 | 77000000 | 77100000 | 1.25 | 0.00 | NO |
| chr4 | 77100000 | 77200000 | 1.21 | 0.92 | A>A |
| chr4 | 77200000 | 77300000 | 1.13 | 1.03 | A>A |
| chr4 | 77300000 | 77400000 | 0.95 | 0.75 | A>A |
| chr4 | 77400000 | 77500000 | 1.04 | 0.86 | A>A |
| chr4 | 77500000 | 77600000 | 1.50 | 1.21 | A>A |
| chr4 | 77600000 | 77700000 | 1.09 | 0.80 | A>A |
| chr4 | 77700000 | 77800000 | 1.16 | 0.92 | A>A |
| chr4 | 77800000 | 77900000 | 1.22 | 0.91 | A>A |
| chr4 | 77900000 | 78000000 | 0.86 | 0.81 | A>A |
| chr4 | 78000000 | 78100000 | 1.06 | 0.91 | A>A |
| chr4 | 78100000 | 78200000 | 0.75 | 0.49 | A>A |
| chr4 | 78200000 | 78300000 | 0.52 | 0.43 | A>A |
| chr4 | 78300000 | 78400000 | 0.51 | 0.39 | A>A |
| chr4 | 78400000 | 78500000 | 0.71 | 0.61 | A>A |
| chr4 | 78500000 | 78600000 | 0.54 | 0.53 | A>A |
| chr4 | 78600000 | 78700000 | 0.81 | 0.61 | A>A |
| chr4 | 78700000 | 78800000 | 1.03 | 0.73 | A>A |
| chr4 | 78800000 | 78900000 | 0.79 | 0.59 | A>A |
| chr4 | 78900000 | 79000000 | 0.74 | 0.61 | A>A |
| chr4 | 79000000 | 79100000 | 0.82 | 0.81 | A>A |
| chr4 | 79100000 | 79200000 | 0.92 | 0.88 | A>A |
| chr4 | 79200000 | 79300000 | 0.86 | 0.72 | A>A |
| chr4 | 79300000 | 79400000 | 0.84 | 0.63 | A>A |
| chr4 | 79400000 | 79500000 | 0.75 | 0.59 | A>A |
| chr4 | 79500000 | 79600000 | 0.84 | 0.53 | A>A |
| chr4 | 79600000 | 79700000 | 1.08 | 0.57 | A>A |
| chr4 | 79700000 | 79800000 | 0.73 | 0.47 | A>A |
| chr4 | 79800000 | 79900000 | 0.22 | -0.15 | A>A |
| chr4 | 79900000 | 80000000 | -0.34 | -0.88 | B>B |
| chr4 | 80000000 | 80100000 | -0.51 | -1.01 | B>B |
| chr4 | 80100000 | 80200000 | -0.54 | -1.11 | B>B |
| chr4 | 80200000 | 80300000 | -0.41 | -1.13 | A>B |
| chr4 | 80300000 | 80400000 | -0.56 | -1.09 | B>B |
| chr4 | 80400000 | 80500000 | -0.47 | -1.02 | B>B |
| chr4 | 80500000 | 80600000 | -0.55 | -1.06 | B>B |

| | | | | | |
|---|---|---|---|---|---|
| chr4 | 80600000 | 80700000 | -0.56 | -1.05 | B>B |
| chr4 | 80700000 | 80800000 | -0.59 | -0.96 | B>B |
| chr4 | 80800000 | 80900000 | -0.30 | -0.75 | B>B |
| chr4 | 80900000 | 81000000 | 0.05 | -0.41 | B>B |
| chr4 | 81000000 | 81100000 | 0.13 | -0.24 | B>B |
| chr4 | 81100000 | 81200000 | 0.00 | -0.29 | B>B |
| chr4 | 81200000 | 81300000 | -0.23 | -0.44 | B>B |
| chr4 | 81300000 | 81400000 | -0.63 | -0.61 | B>B |
| chr4 | 81400000 | 81500000 | -0.68 | -0.77 | B>B |
| chr4 | 81500000 | 81600000 | -0.76 | -0.78 | B>B |
| chr4 | 81600000 | 81700000 | -0.91 | -0.78 | B>B |
| chr4 | 81700000 | 81800000 | -0.84 | -0.75 | B>B |
| chr4 | 81800000 | 81900000 | -0.70 | -0.63 | B>B |
| chr4 | 81900000 | 82000000 | -0.42 | -0.57 | B>B |
| chr4 | 82000000 | 82100000 | 0.05 | -0.46 | B>B |
| chr4 | 82100000 | 82200000 | -0.23 | -0.68 | B>B |
| chr4 | 82200000 | 82300000 | -0.32 | -0.72 | B>B |
| chr4 | 82300000 | 82400000 | -0.32 | -0.78 | B>B |
| chr4 | 82400000 | 82500000 | -0.46 | -1.00 | B>B |
| chr4 | 82500000 | 82600000 | -0.54 | -1.05 | B>B |
| chr4 | 82600000 | 82700000 | -0.42 | -1.15 | A>B |
| chr4 | 82700000 | 82800000 | -0.38 | -1.10 | A>B |
| chr4 | 82800000 | 82900000 | -0.39 | -1.17 | A>B |
| chr4 | 82900000 | 83000000 | -0.36 | -1.03 | A>B |
| chr4 | 83000000 | 83100000 | -0.24 | -1.03 | A>B |
| chr4 | 83100000 | 83200000 | -0.11 | -0.87 | A>B |
| chr4 | 83200000 | 83300000 | 0.61 | -0.13 | A>B |
| chr4 | 83300000 | 83400000 | 1.02 | 0.57 | A>A |
| chr4 | 83400000 | 83500000 | 1.17 | 0.61 | A>A |
| chr4 | 83500000 | 83600000 | 1.09 | 0.77 | A>A |
| chr4 | 83600000 | 83700000 | 1.06 | 0.61 | A>A |
| chr4 | 83700000 | 83800000 | 1.14 | 0.87 | A>A |
| chr4 | 83800000 | 83900000 | 1.51 | 1.09 | A>A |
| chr4 | 83900000 | 84000000 | 1.29 | 1.11 | A>A |
| chr4 | 84000000 | 84100000 | 1.68 | 1.22 | A>A |
| chr4 | 84100000 | 84200000 | 1.38 | 1.10 | A>A |
| chr4 | 84200000 | 84300000 | 1.18 | 0.88 | A>A |
| chr4 | 84300000 | 84400000 | 1.25 | 0.84 | A>A |
| chr4 | 84400000 | 84500000 | 1.01 | 0.69 | A>A |
| chr4 | 84500000 | 84600000 | 0.81 | 0.43 | A>A |
| chr4 | 84600000 | 84700000 | 0.59 | -0.06 | A>B |
| chr4 | 84700000 | 84800000 | 0.75 | 0.23 | A>A |
| chr4 | 84800000 | 84900000 | 0.65 | 0.09 | A>A |
| chr4 | 84900000 | 85000000 | 0.49 | -0.02 | A>A |
| chr4 | 85000000 | 85100000 | 0.45 | 0.05 | A>A |
| chr4 | 85100000 | 85200000 | 0.38 | 0.00 | A>A |
| chr4 | 85200000 | 85300000 | 0.30 | -0.03 | A>A |
| chr4 | 85300000 | 85400000 | 0.31 | -0.15 | A>A |
| chr4 | 85400000 | 85500000 | 0.49 | 0.11 | A>A |
| chr4 | 85500000 | 85600000 | 0.82 | 0.45 | A>A |
| chr4 | 85600000 | 85700000 | 0.86 | 0.62 | A>A |
| chr4 | 85700000 | 85800000 | 1.29 | 0.77 | A>A |
| chr4 | 85800000 | 85900000 | 1.19 | 0.80 | A>A |
| chr4 | 85900000 | 86000000 | 0.77 | 0.31 | A>A |
| chr4 | 86000000 | 86100000 | 0.64 | 0.21 | A>A |
| chr4 | 86100000 | 86200000 | 0.94 | 0.21 | A>B |

| | | | | | |
|---|---|---|---|---|---|
| chr4 | 86200000 | 86300000 | 0.99 | 0.16 | A>B |
| chr4 | 86300000 | 86400000 | 0.99 | 0.23 | A>B |
| chr4 | 86400000 | 86500000 | 1.15 | 0.40 | A>B |
| chr4 | 86500000 | 86600000 | 1.13 | 0.34 | A>B |
| chr4 | 86600000 | 86700000 | 0.88 | 0.23 | A>B |
| chr4 | 86700000 | 86800000 | 0.92 | 0.60 | A>A |
| chr4 | 86800000 | 86900000 | 1.14 | 0.84 | A>A |
| chr4 | 86900000 | 87000000 | 0.80 | 0.32 | A>A |
| chr4 | 87000000 | 87100000 | 1.06 | 0.50 | A>A |
| chr4 | 87100000 | 87200000 | 0.95 | 0.40 | A>A |
| chr4 | 87200000 | 87300000 | 1.12 | 0.50 | A>B |
| chr4 | 87300000 | 87400000 | 0.83 | 0.43 | A>A |
| chr4 | 87400000 | 87500000 | 0.91 | 0.61 | A>A |
| chr4 | 87500000 | 87600000 | 1.11 | 0.80 | A>A |
| chr4 | 87600000 | 87700000 | 1.26 | 0.79 | A>A |
| chr4 | 87700000 | 87800000 | 1.07 | 0.72 | A>A |
| chr4 | 87800000 | 87900000 | 0.98 | 0.79 | A>A |
| chr4 | 87900000 | 88000000 | 1.44 | 1.06 | A>A |
| chr4 | 88000000 | 88100000 | 1.00 | 0.84 | A>A |
| chr4 | 88100000 | 88200000 | 1.10 | 0.95 | A>A |
| chr4 | 88200000 | 88300000 | 1.22 | 1.02 | A>A |
| chr4 | 88300000 | 88400000 | 1.43 | 0.97 | A>A |
| chr4 | 88400000 | 88500000 | 0.97 | 0.85 | A>A |
| chr4 | 88500000 | 88600000 | 1.06 | 0.97 | A>A |
| chr4 | 88600000 | 88700000 | 1.05 | 0.86 | A>A |
| chr4 | 88700000 | 88800000 | 0.87 | 0.76 | A>A |
| chr4 | 88800000 | 88900000 | 1.22 | 1.11 | A>A |
| chr4 | 88900000 | 89000000 | 1.00 | 0.85 | A>A |
| chr4 | 89000000 | 89100000 | 1.44 | 1.26 | A>A |
| chr4 | 89100000 | 89200000 | 1.12 | 0.77 | A>A |
| chr4 | 89200000 | 89300000 | 1.35 | 0.98 | A>A |
| chr4 | 89300000 | 89400000 | 1.27 | 1.03 | A>A |
| chr4 | 89400000 | 89500000 | 0.78 | 0.80 | A>A |
| chr4 | 89500000 | 89600000 | 1.09 | 1.03 | A>A |
| chr4 | 89600000 | 89700000 | 1.05 | 0.90 | A>A |
| chr4 | 89700000 | 89800000 | 0.85 | 0.99 | A>A |
| chr4 | 89800000 | 89900000 | 1.11 | 1.12 | A>A |
| chr4 | 89900000 | 90000000 | 0.74 | 0.75 | A>A |
| chr4 | 90000000 | 90100000 | 0.70 | 0.90 | A>A |
| chr4 | 90100000 | 90200000 | 0.48 | 0.87 | A>A |
| chr4 | 90200000 | 90300000 | 0.75 | 1.00 | A>A |
| chr4 | 90300000 | 90400000 | 0.49 | 0.80 | A>A |
| chr4 | 90400000 | 90500000 | 0.17 | 0.84 | B>A |
| chr4 | 90500000 | 90600000 | 0.00 | 0.75 | B>A |
| chr4 | 90600000 | 90700000 | -0.10 | 0.94 | B>A |
| chr4 | 90700000 | 90800000 | -0.07 | 0.89 | B>A |
| chr4 | 90800000 | 90900000 | -0.11 | 0.60 | B>A |
| chr4 | 90900000 | 91000000 | -0.28 | 0.80 | B>A |
| chr4 | 91000000 | 91100000 | -0.39 | 0.78 | B>A |
| chr4 | 91100000 | 91200000 | -0.50 | 0.76 | B>A |
| chr4 | 91200000 | 91300000 | -0.31 | 0.44 | B>A |
| chr4 | 91300000 | 91400000 | -0.26 | 0.53 | B>A |
| chr4 | 91400000 | 91500000 | -0.27 | 0.76 | B>A |
| chr4 | 91500000 | 91600000 | -0.21 | 0.85 | B>A |
| chr4 | 91600000 | 91700000 | -0.22 | 0.75 | B>A |
| chr4 | 91700000 | 91800000 | -0.02 | 0.76 | B>A |

| | | | | | |
|---|---|---|---|---|---|
| chr4 | 91800000 | 91900000 | -0.10 | 0.82 | B>A |
| chr4 | 91900000 | 92000000 | -0.12 | 0.83 | B>A |
| chr4 | 92000000 | 92100000 | -0.40 | 0.41 | B>A |
| chr4 | 92100000 | 92200000 | -0.51 | 0.75 | B>A |
| chr4 | 92200000 | 92300000 | -0.68 | 0.82 | B>A |
| chr4 | 92300000 | 92400000 | -0.77 | 0.74 | B>A |
| chr4 | 92400000 | 92500000 | -0.77 | 0.60 | B>A |
| chr4 | 92500000 | 92600000 | -0.79 | 0.68 | B>A |
| chr4 | 92600000 | 92700000 | -0.82 | 0.65 | B>A |
| chr4 | 92700000 | 92800000 | -0.82 | 0.71 | B>A |
| chr4 | 92800000 | 92900000 | -0.70 | 0.73 | B>A |
| chr4 | 92900000 | 93000000 | -0.70 | 0.58 | B>A |
| chr4 | 93000000 | 93100000 | -0.62 | 0.47 | B>A |
| chr4 | 93100000 | 93200000 | -0.55 | 0.70 | B>A |
| chr4 | 93200000 | 93300000 | -0.31 | 0.56 | B>A |
| chr4 | 93300000 | 93400000 | -0.21 | 0.62 | B>A |
| chr4 | 93400000 | 93500000 | -0.22 | 0.65 | B>A |
| chr4 | 93500000 | 93600000 | -0.01 | 0.67 | B>A |
| chr4 | 93600000 | 93700000 | 0.13 | 0.66 | B>A |
| chr4 | 93700000 | 93800000 | 0.15 | 0.68 | B>A |
| chr4 | 93800000 | 93900000 | 0.36 | 0.70 | A>A |
| chr4 | 93900000 | 94000000 | 0.38 | 0.74 | A>A |
| chr4 | 94000000 | 94100000 | 0.34 | 0.65 | A>A |
| chr4 | 94100000 | 94200000 | 0.17 | 0.45 | A>A |
| chr4 | 94200000 | 94300000 | 0.19 | 0.62 | A>A |
| chr4 | 94300000 | 94400000 | 0.33 | 0.67 | A>A |
| chr4 | 94400000 | 94500000 | 0.16 | 0.57 | A>A |
| chr4 | 94500000 | 94600000 | 0.06 | 0.59 | B>A |
| chr4 | 94600000 | 94700000 | -0.05 | 0.39 | A>A |
| chr4 | 94700000 | 94800000 | -0.14 | 0.43 | B>A |
| chr4 | 94800000 | 94900000 | -0.32 | 0.41 | B>A |
| chr4 | 94900000 | 95000000 | -0.28 | 0.47 | B>A |
| chr4 | 95000000 | 95100000 | -0.12 | 0.44 | B>A |
| chr4 | 95100000 | 95200000 | 0.47 | 0.70 | A>A |
| chr4 | 95200000 | 95300000 | 0.57 | 0.70 | A>A |
| chr4 | 95300000 | 95400000 | 1.03 | 0.92 | A>A |
| chr4 | 95400000 | 95500000 | 1.10 | 1.11 | A>A |
| chr4 | 95500000 | 95600000 | 0.80 | 0.81 | A>A |
| chr4 | 95600000 | 95700000 | 0.90 | 0.90 | A>A |
| chr4 | 95700000 | 95800000 | 0.99 | 1.16 | A>A |
| chr4 | 95800000 | 95900000 | 1.07 | 1.05 | A>A |
| chr4 | 95900000 | 96000000 | 0.97 | 1.02 | A>A |
| chr4 | 96000000 | 96100000 | 0.35 | 0.68 | A>A |
| chr4 | 96100000 | 96200000 | 0.15 | 0.54 | A>A |
| chr4 | 96200000 | 96300000 | 0.29 | 0.74 | A>A |
| chr4 | 96300000 | 96400000 | 0.30 | 0.73 | A>A |
| chr4 | 96400000 | 96500000 | 0.14 | 0.54 | A>A |
| chr4 | 96500000 | 96600000 | -0.09 | 0.45 | B>A |
| chr4 | 96600000 | 96700000 | -0.30 | 0.57 | B>A |
| chr4 | 96700000 | 96800000 | -0.52 | 0.47 | B>A |
| chr4 | 96800000 | 96900000 | -0.65 | 0.57 | B>A |
| chr4 | 96900000 | 97000000 | -0.66 | 0.52 | B>A |
| chr4 | 97000000 | 97100000 | -0.72 | 0.53 | B>A |
| chr4 | 97100000 | 97200000 | -0.71 | 0.40 | B>A |
| chr4 | 97200000 | 97300000 | -0.66 | 0.31 | B>A |
| chr4 | 97300000 | 97400000 | -0.76 | 0.35 | B>A |

| | | | | | |
|---|---|---|---|---|---|
| chr4 | 97400000 | 97500000 | -0.70 | 0.37 | B>A |
| chr4 | 97500000 | 97600000 | -0.79 | 0.36 | B>A |
| chr4 | 97600000 | 97700000 | -0.73 | 0.42 | B>A |
| chr4 | 97700000 | 97800000 | -0.74 | 0.40 | B>A |
| chr4 | 97800000 | 97900000 | -0.69 | 0.36 | B>A |
| chr4 | 97900000 | 98000000 | -0.62 | 0.44 | B>A |
| chr4 | 98000000 | 98100000 | -0.71 | 0.45 | B>A |
| chr4 | 98100000 | 98200000 | -0.63 | 0.42 | B>A |
| chr4 | 98200000 | 98300000 | -0.62 | 0.59 | B>A |
| chr4 | 98300000 | 98400000 | -0.54 | 0.45 | B>A |
| chr4 | 98400000 | 98500000 | -0.56 | 0.57 | B>A |
| chr4 | 98500000 | 98600000 | -0.65 | 0.52 | B>A |
| chr4 | 98600000 | 98700000 | -0.73 | 0.43 | B>A |
| chr4 | 98700000 | 98800000 | -0.73 | 0.55 | B>A |
| chr4 | 98800000 | 98900000 | -0.70 | 0.47 | B>A |
| chr4 | 98900000 | 99000000 | -0.56 | 0.33 | B>A |
| chr4 | 99000000 | 99100000 | -0.13 | 0.34 | A>A |
| chr4 | 99100000 | 99200000 | 0.34 | 0.50 | A>A |
| chr4 | 99200000 | 99300000 | 0.53 | 0.64 | A>A |
| chr4 | 99300000 | 99400000 | 0.47 | 0.52 | A>A |
| chr4 | 99400000 | 99500000 | 0.63 | 0.55 | A>A |
| chr4 | 99500000 | 99600000 | 0.82 | 0.67 | A>A |
| chr4 | 99600000 | 99700000 | 0.61 | 0.57 | A>A |
| chr4 | 99700000 | 99800000 | 0.65 | 0.54 | A>A |
| chr4 | 99800000 | 99900000 | 0.82 | 0.64 | A>A |
| chr4 | 99900000 | 100000000 | 0.75 | 0.67 | A>A |
| chr4 | 1E+08 | 100100000 | 0.22 | 0.39 | A>A |
| chr4 | 1E+08 | 100200000 | -0.14 | 0.21 | A>A |
| chr4 | 1E+08 | 100300000 | -0.07 | 0.24 | A>A |
| chr4 | 1E+08 | 100400000 | 0.03 | 0.42 | A>A |
| chr4 | 1E+08 | 100500000 | 0.17 | 0.31 | A>A |
| chr4 | 1.01E+08 | 100600000 | 0.28 | 0.64 | A>A |
| chr4 | 1.01E+08 | 100700000 | 0.25 | 0.64 | A>A |
| chr4 | 1.01E+08 | 100800000 | 0.27 | 0.47 | A>A |
| chr4 | 1.01E+08 | 100900000 | 0.56 | 0.69 | A>A |
| chr4 | 1.01E+08 | 101000000 | 0.28 | 0.59 | A>A |
| chr4 | 1.01E+08 | 101100000 | 0.13 | 0.61 | A>A |
| chr4 | 1.01E+08 | 101200000 | 0.16 | 0.81 | B>A |
| chr4 | 1.01E+08 | 101300000 | 0.20 | 0.70 | A>A |
| chr4 | 1.01E+08 | 101400000 | 0.09 | 0.54 | A>A |
| chr4 | 1.01E+08 | 101500000 | 0.35 | 0.69 | A>A |
| chr4 | 1.02E+08 | 101600000 | 0.32 | 0.78 | A>A |
| chr4 | 1.02E+08 | 101700000 | 0.29 | 0.61 | A>A |
| chr4 | 1.02E+08 | 101800000 | 0.49 | 0.71 | A>A |
| chr4 | 1.02E+08 | 101900000 | 0.60 | 0.59 | A>A |
| chr4 | 1.02E+08 | 102000000 | 0.85 | 0.81 | A>A |
| chr4 | 1.02E+08 | 102100000 | 0.72 | 0.81 | A>A |
| chr4 | 1.02E+08 | 102200000 | 0.89 | 0.85 | A>A |
| chr4 | 1.02E+08 | 102300000 | 0.84 | 0.79 | A>A |
| chr4 | 1.02E+08 | 102400000 | 0.90 | 0.80 | A>A |
| chr4 | 1.02E+08 | 102500000 | 0.92 | 0.84 | A>A |
| chr4 | 1.03E+08 | 102600000 | 0.73 | 0.77 | A>A |
| chr4 | 1.03E+08 | 102700000 | 0.69 | 0.71 | A>A |
| chr4 | 1.03E+08 | 102800000 | 0.75 | 0.88 | A>A |
| chr4 | 1.03E+08 | 102900000 | 0.96 | 1.02 | A>A |
| chr4 | 1.03E+08 | 103000000 | 1.07 | 0.81 | A>A |

| | | | | | |
|---|---|---|---|---|---|
| chr4 | 1.03E+08 | 103100000 | 0.94 | 0.80 | A>A |
| chr4 | 1.03E+08 | 103200000 | 0.73 | 0.77 | A>A |
| chr4 | 1.03E+08 | 103300000 | 0.62 | 0.69 | A>A |
| chr4 | 1.03E+08 | 103400000 | 0.71 | 0.76 | A>A |
| chr4 | 1.03E+08 | 103500000 | 0.94 | 1.09 | A>A |
| chr4 | 1.04E+08 | 103600000 | 0.81 | 0.70 | A>A |
| chr4 | 1.04E+08 | 103700000 | 0.91 | 1.03 | A>A |
| chr4 | 1.04E+08 | 103800000 | 0.94 | 1.05 | A>A |
| chr4 | 1.04E+08 | 103900000 | 0.00 | 0.00 | NO |
| chr4 | 1.04E+08 | 104000000 | 0.47 | 0.61 | A>A |
| chr4 | 1.04E+08 | 104100000 | 0.32 | 0.61 | A>A |
| chr4 | 1.04E+08 | 104200000 | -0.28 | 0.41 | B>A |
| chr4 | 1.04E+08 | 104300000 | -0.41 | 0.61 | B>A |
| chr4 | 1.04E+08 | 104400000 | -0.24 | 0.69 | B>A |
| chr4 | 1.04E+08 | 104500000 | -0.58 | 0.53 | B>A |
| chr4 | 1.05E+08 | 104600000 | -0.63 | 0.46 | B>A |
| chr4 | 1.05E+08 | 104700000 | -0.55 | 0.41 | B>A |
| chr4 | 1.05E+08 | 104800000 | -0.67 | 0.32 | B>A |
| chr4 | 1.05E+08 | 104900000 | -0.45 | 0.28 | B>A |
| chr4 | 1.05E+08 | 105000000 | -0.42 | 0.26 | B>A |
| chr4 | 1.05E+08 | 105100000 | -0.52 | 0.58 | B>A |
| chr4 | 1.05E+08 | 105200000 | -0.52 | 0.45 | B>A |
| chr4 | 1.05E+08 | 105300000 | -0.56 | 0.49 | B>A |
| chr4 | 1.05E+08 | 105400000 | -0.20 | 0.66 | B>A |
| chr4 | 1.05E+08 | 105500000 | -0.16 | 0.59 | B>A |
| chr4 | 1.06E+08 | 105600000 | -0.20 | 0.55 | B>A |
| chr4 | 1.06E+08 | 105700000 | 0.05 | 0.68 | B>A |
| chr4 | 1.06E+08 | 105800000 | -0.13 | 0.48 | B>A |
| chr4 | 1.06E+08 | 105900000 | -0.29 | 0.36 | B>A |
| chr4 | 1.06E+08 | 106000000 | -0.37 | 0.40 | B>A |
| chr4 | 1.06E+08 | 106100000 | -0.01 | 0.51 | B>A |
| chr4 | 1.06E+08 | 106200000 | 0.29 | 0.56 | A>A |
| chr4 | 1.06E+08 | 106300000 | 0.17 | 0.55 | A>A |
| chr4 | 1.06E+08 | 106400000 | 0.43 | 0.61 | A>A |
| chr4 | 1.06E+08 | 106500000 | 0.64 | 0.85 | A>A |
| chr4 | 1.07E+08 | 106600000 | 0.72 | 0.92 | A>A |
| chr4 | 1.07E+08 | 106700000 | 0.43 | 0.56 | A>A |
| chr4 | 1.07E+08 | 106800000 | 0.28 | 0.65 | A>A |
| chr4 | 1.07E+08 | 106900000 | 0.15 | 0.58 | A>A |
| chr4 | 1.07E+08 | 107000000 | -0.05 | 0.56 | B>A |
| chr4 | 1.07E+08 | 107100000 | 0.12 | 0.40 | A>A |
| chr4 | 1.07E+08 | 107200000 | 0.07 | 0.29 | A>A |
| chr4 | 1.07E+08 | 107300000 | 0.08 | 0.09 | A>A |
| chr4 | 1.07E+08 | 107400000 | -0.51 | -0.26 | B>B |
| chr4 | 1.07E+08 | 107500000 | -0.65 | -0.18 | B>B |
| chr4 | 1.08E+08 | 107600000 | -0.89 | -0.16 | B>A |
| chr4 | 1.08E+08 | 107700000 | -0.92 | -0.32 | B>A |
| chr4 | 1.08E+08 | 107800000 | -0.91 | -0.48 | B>B |
| chr4 | 1.08E+08 | 107900000 | -0.89 | -0.71 | B>B |
| chr4 | 1.08E+08 | 108000000 | -0.77 | -0.74 | B>B |
| chr4 | 1.08E+08 | 108100000 | -0.75 | -0.68 | B>B |
| chr4 | 1.08E+08 | 108200000 | -0.55 | -0.46 | B>B |
| chr4 | 1.08E+08 | 108300000 | -0.48 | -0.45 | B>B |
| chr4 | 1.08E+08 | 108400000 | -0.48 | -0.50 | B>B |
| chr4 | 1.08E+08 | 108500000 | -0.27 | -0.29 | B>B |
| chr4 | 1.09E+08 | 108600000 | 0.07 | 0.06 | A>A |

| | | | | | |
|---|---|---|---|---|---|
| chr4 | 1.09E+08 | 108700000 | 0.31 | 0.13 | A>A |
| chr4 | 1.09E+08 | 108800000 | 0.70 | 0.43 | A>A |
| chr4 | 1.09E+08 | 108900000 | 0.41 | 0.12 | A>A |
| chr4 | 1.09E+08 | 109000000 | 0.17 | -0.45 | A>B |
| chr4 | 1.09E+08 | 109100000 | 0.05 | -0.52 | B>B |
| chr4 | 1.09E+08 | 109200000 | -0.18 | -1.00 | A>B |
| chr4 | 1.09E+08 | 109300000 | -0.23 | -1.12 | A>B |
| chr4 | 1.09E+08 | 109400000 | -0.38 | -1.13 | A>B |
| chr4 | 1.09E+08 | 109500000 | -0.31 | -0.95 | A>B |
| chr4 | 1.1E+08 | 109600000 | 0.30 | -0.32 | A>B |
| chr4 | 1.1E+08 | 109700000 | 0.14 | -0.76 | A>B |
| chr4 | 1.1E+08 | 109800000 | -0.15 | -1.02 | A>B |
| chr4 | 1.1E+08 | 109900000 | -0.36 | -1.20 | A>B |
| chr4 | 1.1E+08 | 110000000 | -0.52 | -1.28 | A>B |
| chr4 | 1.1E+08 | 110100000 | -0.53 | -1.28 | A>B |
| chr4 | 1.1E+08 | 110200000 | -0.46 | -1.17 | A>B |
| chr4 | 1.1E+08 | 110300000 | -0.18 | -1.01 | A>B |
| chr4 | 1.1E+08 | 110400000 | 0.42 | -0.16 | A>B |
| chr4 | 1.1E+08 | 110500000 | 0.96 | 0.45 | A>A |
| chr4 | 1.11E+08 | 110600000 | 0.94 | 0.38 | A>A |
| chr4 | 1.11E+08 | 110700000 | 0.74 | 0.19 | A>A |
| chr4 | 1.11E+08 | 110800000 | 0.45 | -0.23 | A>B |
| chr4 | 1.11E+08 | 110900000 | 0.26 | -0.49 | A>B |
| chr4 | 1.11E+08 | 111000000 | 0.31 | 0.02 | A>A |
| chr4 | 1.11E+08 | 111100000 | 1.04 | 0.53 | A>A |
| chr4 | 1.11E+08 | 111200000 | 0.76 | 0.04 | A>B |
| chr4 | 1.11E+08 | 111300000 | 0.16 | -0.39 | B>B |
| chr4 | 1.11E+08 | 111400000 | 0.02 | -0.41 | B>B |
| chr4 | 1.11E+08 | 111500000 | 0.01 | -0.16 | B>B |
| chr4 | 1.12E+08 | 111600000 | -0.17 | -0.31 | B>B |
| chr4 | 1.12E+08 | 111700000 | -0.39 | -0.73 | B>B |
| chr4 | 1.12E+08 | 111800000 | -0.48 | -0.77 | B>B |
| chr4 | 1.12E+08 | 111900000 | -0.49 | -0.86 | B>B |
| chr4 | 1.12E+08 | 112000000 | -0.64 | -0.90 | B>B |
| chr4 | 1.12E+08 | 112100000 | -0.76 | -0.94 | B>B |
| chr4 | 1.12E+08 | 112200000 | -0.75 | -0.80 | B>B |
| chr4 | 1.12E+08 | 112300000 | -0.52 | -0.67 | B>B |
| chr4 | 1.12E+08 | 112400000 | -0.53 | -0.81 | B>B |
| chr4 | 1.12E+08 | 112500000 | -0.28 | -0.69 | B>B |
| chr4 | 1.13E+08 | 112600000 | -0.27 | -0.47 | B>B |
| chr4 | 1.13E+08 | 112700000 | -0.32 | -0.57 | B>B |
| chr4 | 1.13E+08 | 112800000 | -0.30 | -0.53 | B>B |
| chr4 | 1.13E+08 | 112900000 | -0.08 | -0.20 | B>B |
| chr4 | 1.13E+08 | 113000000 | 0.27 | 0.05 | A>A |
| chr4 | 1.13E+08 | 113100000 | 1.02 | 0.65 | A>A |
| chr4 | 1.13E+08 | 113200000 | 0.84 | 0.56 | A>A |
| chr4 | 1.13E+08 | 113300000 | 0.57 | 0.33 | A>A |
| chr4 | 1.13E+08 | 113400000 | 0.39 | 0.12 | A>A |
| chr4 | 1.13E+08 | 113500000 | 0.68 | 0.24 | A>A |
| chr4 | 1.14E+08 | 113600000 | 0.83 | 0.30 | A>A |
| chr4 | 1.14E+08 | 113700000 | 0.93 | 0.20 | A>B |
| chr4 | 1.14E+08 | 113800000 | 0.70 | 0.00 | A>B |
| chr4 | 1.14E+08 | 113900000 | 0.40 | -0.08 | A>A |
| chr4 | 1.14E+08 | 114000000 | 0.38 | 0.03 | A>A |
| chr4 | 1.14E+08 | 114100000 | 0.31 | 0.21 | A>A |
| chr4 | 1.14E+08 | 114200000 | 0.21 | 0.23 | A>A |

| | | | | | |
|---|---|---|---|---|---|
| chr4 | 1.14E+08 | 114300000 | 0.25 | 0.23 | A>A |
| chr4 | 1.14E+08 | 114400000 | 0.34 | 0.50 | A>A |
| chr4 | 1.14E+08 | 114500000 | 0.70 | 0.68 | A>A |
| chr4 | 1.15E+08 | 114600000 | 0.91 | 0.87 | A>A |
| chr4 | 1.15E+08 | 114700000 | 0.80 | 0.77 | A>A |
| chr4 | 1.15E+08 | 114800000 | 0.59 | 0.41 | A>A |
| chr4 | 1.15E+08 | 114900000 | 0.67 | 0.48 | A>A |
| chr4 | 1.15E+08 | 115000000 | 0.83 | 0.68 | A>A |
| chr4 | 1.15E+08 | 115100000 | 0.60 | 0.51 | A>A |
| chr4 | 1.15E+08 | 115200000 | 0.38 | 0.54 | A>A |
| chr4 | 1.15E+08 | 115300000 | 0.24 | 0.41 | A>A |
| chr4 | 1.15E+08 | 115400000 | -0.36 | 0.17 | B>A |
| chr4 | 1.15E+08 | 115500000 | -0.48 | 0.37 | B>A |
| chr4 | 1.16E+08 | 115600000 | -0.34 | 0.33 | B>A |
| chr4 | 1.16E+08 | 115700000 | -0.53 | 0.17 | B>A |
| chr4 | 1.16E+08 | 115800000 | -0.79 | 0.28 | B>A |
| chr4 | 1.16E+08 | 115900000 | -0.95 | 0.20 | B>A |
| chr4 | 1.16E+08 | 116000000 | -0.92 | 0.21 | B>A |
| chr4 | 1.16E+08 | 116100000 | -0.77 | 0.24 | B>A |
| chr4 | 1.16E+08 | 116200000 | -0.95 | 0.39 | B>A |
| chr4 | 1.16E+08 | 116300000 | -0.89 | 0.51 | B>A |
| chr4 | 1.16E+08 | 116400000 | -0.89 | 0.47 | B>A |
| chr4 | 1.16E+08 | 116500000 | -0.89 | 0.40 | B>A |
| chr4 | 1.17E+08 | 116600000 | -0.86 | 0.58 | B>A |
| chr4 | 1.17E+08 | 116700000 | -0.79 | 0.51 | B>A |
| chr4 | 1.17E+08 | 116800000 | -0.87 | 0.40 | B>A |
| chr4 | 1.17E+08 | 116900000 | -0.84 | 0.38 | B>A |
| chr4 | 1.17E+08 | 117000000 | -0.75 | 0.36 | B>A |
| chr4 | 1.17E+08 | 117100000 | -0.74 | 0.28 | B>A |
| chr4 | 1.17E+08 | 117200000 | -0.66 | 0.25 | B>A |
| chr4 | 1.17E+08 | 117300000 | -0.56 | 0.22 | B>A |
| chr4 | 1.17E+08 | 117400000 | -0.67 | 0.23 | B>A |
| chr4 | 1.17E+08 | 117500000 | -0.74 | 0.29 | B>A |
| chr4 | 1.18E+08 | 117600000 | -0.69 | 0.31 | B>A |
| chr4 | 1.18E+08 | 117700000 | -0.79 | 0.25 | B>A |
| chr4 | 1.18E+08 | 117800000 | -0.80 | 0.28 | B>A |
| chr4 | 1.18E+08 | 117900000 | -0.80 | 0.38 | B>A |
| chr4 | 1.18E+08 | 118000000 | -0.68 | 0.47 | B>A |
| chr4 | 1.18E+08 | 118100000 | -0.70 | 0.24 | B>A |
| chr4 | 1.18E+08 | 118200000 | -0.79 | 0.34 | B>A |
| chr4 | 1.18E+08 | 118300000 | -0.75 | 0.49 | B>A |
| chr4 | 1.18E+08 | 118400000 | -0.82 | 0.46 | B>A |
| chr4 | 1.18E+08 | 118500000 | -0.81 | 0.38 | B>A |
| chr4 | 1.19E+08 | 118600000 | -0.83 | 0.44 | B>A |
| chr4 | 1.19E+08 | 118700000 | -0.81 | 0.45 | B>A |
| chr4 | 1.19E+08 | 118800000 | -0.76 | 0.59 | B>A |
| chr4 | 1.19E+08 | 118900000 | -0.67 | 0.51 | B>A |
| chr4 | 1.19E+08 | 119000000 | -0.52 | 0.25 | B>A |
| chr4 | 1.19E+08 | 119100000 | -0.51 | 0.34 | B>A |
| chr4 | 1.19E+08 | 119200000 | -0.38 | 0.22 | B>A |
| chr4 | 1.19E+08 | 119300000 | 0.19 | 0.41 | A>A |
| chr4 | 1.19E+08 | 119400000 | 0.00 | 0.00 | NO |
| chr4 | 1.19E+08 | 119500000 | 0.00 | 0.00 | NO |
| chr4 | 1.2E+08 | 119600000 | 0.00 | 0.00 | NO |
| chr4 | 1.2E+08 | 119700000 | 1.01 | 0.63 | A>A |
| chr4 | 1.2E+08 | 119800000 | 1.31 | 0.80 | A>A |

| | | | | | |
|---|---|---|---|---|---|
| chr4 | 1.2E+08 | 119900000 | 0.79 | 0.62 | A>A |
| chr4 | 1.2E+08 | 120000000 | 1.19 | 0.80 | A>A |
| chr4 | 1.2E+08 | 120100000 | 1.80 | 1.06 | A>B |
| chr4 | 1.2E+08 | 120200000 | 1.38 | 0.98 | A>A |
| chr4 | 1.2E+08 | 120300000 | 0.50 | 0.35 | A>A |
| chr4 | 1.2E+08 | 120400000 | 0.00 | 0.00 | NO |
| chr4 | 1.2E+08 | 120500000 | 0.22 | 0.15 | A>A |
| chr4 | 1.21E+08 | 120600000 | 0.19 | 0.31 | A>A |
| chr4 | 1.21E+08 | 120700000 | 0.29 | 0.47 | A>A |
| chr4 | 1.21E+08 | 120800000 | -0.09 | 0.24 | A>A |
| chr4 | 1.21E+08 | 120900000 | -0.25 | 0.17 | B>B |
| chr4 | 1.21E+08 | 121000000 | -0.09 | 0.02 | B>B |
| chr4 | 1.21E+08 | 121100000 | -0.34 | -0.72 | B>B |
| chr4 | 1.21E+08 | 121200000 | -0.55 | -0.97 | B>B |
| chr4 | 1.21E+08 | 121300000 | -0.68 | -1.10 | B>B |
| chr4 | 1.21E+08 | 121400000 | -0.71 | -0.97 | B>B |
| chr4 | 1.21E+08 | 121500000 | -0.53 | -0.73 | B>B |
| chr4 | 1.22E+08 | 121600000 | -0.49 | -0.76 | B>B |
| chr4 | 1.22E+08 | 121700000 | -0.05 | -0.43 | B>B |
| chr4 | 1.22E+08 | 121800000 | 0.14 | -0.37 | B>B |
| chr4 | 1.22E+08 | 121900000 | 0.10 | -0.58 | A>B |
| chr4 | 1.22E+08 | 122000000 | -0.22 | -0.73 | B>B |
| chr4 | 1.22E+08 | 122100000 | -0.13 | -0.56 | B>B |
| chr4 | 1.22E+08 | 122200000 | -0.09 | -0.64 | B>B |
| chr4 | 1.22E+08 | 122300000 | -0.24 | -0.72 | B>B |
| chr4 | 1.22E+08 | 122400000 | -0.30 | -0.57 | B>B |
| chr4 | 1.22E+08 | 122500000 | -0.33 | -0.81 | B>B |
| chr4 | 1.23E+08 | 122600000 | 0.17 | -0.56 | A>B |
| chr4 | 1.23E+08 | 122700000 | 0.43 | -0.07 | A>A |
| chr4 | 1.23E+08 | 122800000 | 0.63 | 0.16 | A>A |
| chr4 | 1.23E+08 | 122900000 | 0.33 | -0.05 | A>A |
| chr4 | 1.23E+08 | 123000000 | 0.20 | 0.05 | A>A |
| chr4 | 1.23E+08 | 123100000 | 0.37 | 0.11 | A>A |
| chr4 | 1.23E+08 | 123200000 | 0.57 | 0.24 | A>A |
| chr4 | 1.23E+08 | 123300000 | 0.32 | 0.14 | A>A |
| chr4 | 1.23E+08 | 123400000 | 0.16 | -0.11 | A>A |
| chr4 | 1.23E+08 | 123500000 | 0.05 | -0.19 | B>B |
| chr4 | 1.24E+08 | 123600000 | 0.36 | 0.12 | A>A |
| chr4 | 1.24E+08 | 123700000 | 0.88 | 0.38 | A>A |
| chr4 | 1.24E+08 | 123800000 | 1.39 | 0.85 | A>A |
| chr4 | 1.24E+08 | 123900000 | 0.84 | 0.50 | A>A |
| chr4 | 1.24E+08 | 124000000 | 0.78 | 0.66 | A>A |
| chr4 | 1.24E+08 | 124100000 | 0.45 | 0.52 | A>A |
| chr4 | 1.24E+08 | 124200000 | 0.52 | 0.75 | A>A |
| chr4 | 1.24E+08 | 124300000 | 0.56 | 0.68 | A>A |
| chr4 | 1.24E+08 | 124400000 | 0.76 | 0.85 | A>A |
| chr4 | 1.24E+08 | 124500000 | 1.02 | 0.89 | A>A |
| chr4 | 1.25E+08 | 124600000 | 0.85 | 0.91 | A>A |
| chr4 | 1.25E+08 | 124700000 | 0.20 | 0.57 | A>A |
| chr4 | 1.25E+08 | 124800000 | -0.06 | 0.65 | B>A |
| chr4 | 1.25E+08 | 124900000 | -0.19 | 0.81 | B>A |
| chr4 | 1.25E+08 | 125000000 | -0.16 | 0.76 | B>A |
| chr4 | 1.25E+08 | 125100000 | -0.08 | 0.57 | B>A |
| chr4 | 1.25E+08 | 125200000 | 0.10 | 0.54 | A>A |
| chr4 | 1.25E+08 | 125300000 | 0.14 | 0.43 | A>A |
| chr4 | 1.25E+08 | 125400000 | 0.48 | 0.64 | A>A |

| | | | | | |
|---|---|---|---|---|---|
| chr4 | 1.25E+08 | 125500000 | 0.44 | 0.66 | A>A |
| chr4 | 1.26E+08 | 125600000 | 0.49 | 0.63 | A>A |
| chr4 | 1.26E+08 | 125700000 | 0.37 | 0.50 | A>A |
| chr4 | 1.26E+08 | 125800000 | 0.34 | 0.51 | A>A |
| chr4 | 1.26E+08 | 125900000 | 0.25 | 0.51 | A>A |
| chr4 | 1.26E+08 | 126000000 | 0.32 | 0.52 | A>A |
| chr4 | 1.26E+08 | 126100000 | 0.17 | 0.40 | A>A |
| chr4 | 1.26E+08 | 126200000 | 0.06 | 0.26 | A>A |
| chr4 | 1.26E+08 | 126300000 | 0.04 | -0.12 | B>B |
| chr4 | 1.26E+08 | 126400000 | 0.22 | -0.18 | A>A |
| chr4 | 1.26E+08 | 126500000 | 0.20 | -0.32 | B>B |
| chr4 | 1.27E+08 | 126600000 | 0.38 | -0.18 | A>A |
| chr4 | 1.27E+08 | 126700000 | 0.14 | -0.14 | A>A |
| chr4 | 1.27E+08 | 126800000 | -0.30 | -0.50 | B>B |
| chr4 | 1.27E+08 | 126900000 | -0.65 | -0.89 | B>B |
| chr4 | 1.27E+08 | 127000000 | -0.71 | -1.02 | B>B |
| chr4 | 1.27E+08 | 127100000 | -0.72 | -0.95 | B>B |
| chr4 | 1.27E+08 | 127200000 | -0.79 | -0.98 | B>B |
| chr4 | 1.27E+08 | 127300000 | -0.88 | -1.02 | B>B |
| chr4 | 1.27E+08 | 127400000 | -0.76 | -0.96 | B>B |
| chr4 | 1.27E+08 | 127500000 | -0.73 | -0.87 | B>B |
| chr4 | 1.28E+08 | 127600000 | -0.61 | -0.47 | B>B |
| chr4 | 1.28E+08 | 127700000 | -0.47 | -0.22 | B>B |
| chr4 | 1.28E+08 | 127800000 | -0.28 | -0.10 | B>B |
| chr4 | 1.28E+08 | 127900000 | -0.32 | -0.21 | B>B |
| chr4 | 1.28E+08 | 128000000 | -0.23 | -0.26 | B>B |
| chr4 | 1.28E+08 | 128100000 | -0.37 | -0.43 | B>B |
| chr4 | 1.28E+08 | 128200000 | -0.38 | -0.59 | B>B |
| chr4 | 1.28E+08 | 128300000 | -0.32 | -0.67 | B>B |
| chr4 | 1.28E+08 | 128400000 | -0.16 | -0.36 | B>B |
| chr4 | 1.28E+08 | 128500000 | -0.03 | -0.15 | B>B |
| chr4 | 1.29E+08 | 128600000 | 0.59 | 0.27 | A>A |
| chr4 | 1.29E+08 | 128700000 | 0.67 | 0.35 | A>A |
| chr4 | 1.29E+08 | 128800000 | 1.11 | 0.71 | A>A |
| chr4 | 1.29E+08 | 128900000 | 1.39 | 1.08 | A>A |
| chr4 | 1.29E+08 | 129000000 | 1.35 | 1.13 | A>A |
| chr4 | 1.29E+08 | 129100000 | 1.17 | 1.16 | A>A |
| chr4 | 1.29E+08 | 129200000 | 1.05 | 1.01 | A>A |
| chr4 | 1.29E+08 | 129300000 | 1.39 | 1.28 | A>A |
| chr4 | 1.29E+08 | 129400000 | 1.12 | 0.88 | A>A |
| chr4 | 1.29E+08 | 129500000 | 1.31 | 1.23 | A>A |
| chr4 | 1.3E+08 | 129600000 | 1.47 | 1.36 | A>A |
| chr4 | 1.3E+08 | 129700000 | 1.69 | 1.37 | A>A |
| chr4 | 1.3E+08 | 129800000 | 1.06 | 0.95 | A>A |
| chr4 | 1.3E+08 | 129900000 | 0.79 | 0.81 | A>A |
| chr4 | 1.3E+08 | 130000000 | 0.71 | 0.66 | A>A |
| chr4 | 1.3E+08 | 130100000 | 0.64 | 0.43 | A>A |
| chr4 | 1.3E+08 | 130200000 | 0.56 | 0.30 | A>A |
| chr4 | 1.3E+08 | 130300000 | 0.18 | 0.22 | A>A |
| chr4 | 1.3E+08 | 130400000 | 0.19 | 0.28 | A>A |
| chr4 | 1.3E+08 | 130500000 | 0.17 | 0.23 | A>A |
| chr4 | 1.31E+08 | 130600000 | 0.07 | 0.26 | A>A |
| chr4 | 1.31E+08 | 130700000 | 0.21 | 0.40 | A>A |
| chr4 | 1.31E+08 | 130800000 | 0.22 | 0.40 | A>A |
| chr4 | 1.31E+08 | 130900000 | 0.26 | 0.29 | A>A |
| chr4 | 1.31E+08 | 131000000 | 0.41 | 0.68 | A>A |

| | | | | | |
|---|---|---|---|---|---|
| chr4 | 1.31E+08 | 131100000 | 0.47 | 0.54 | A>A |
| chr4 | 1.31E+08 | 131200000 | 0.45 | 0.58 | A>A |
| chr4 | 1.31E+08 | 131300000 | 0.34 | 0.38 | A>A |
| chr4 | 1.31E+08 | 131400000 | 0.27 | 0.35 | A>A |
| chr4 | 1.31E+08 | 131500000 | 0.26 | 0.29 | A>A |
| chr4 | 1.32E+08 | 131600000 | 0.40 | 0.43 | A>A |
| chr4 | 1.32E+08 | 131700000 | 0.28 | 0.36 | A>A |
| chr4 | 1.32E+08 | 131800000 | 0.25 | 0.26 | A>A |
| chr4 | 1.32E+08 | 131900000 | 0.20 | 0.35 | A>A |
| chr4 | 1.32E+08 | 132000000 | 0.10 | 0.26 | A>A |
| chr4 | 1.32E+08 | 132100000 | 0.20 | 0.27 | A>A |
| chr4 | 1.32E+08 | 132200000 | 0.22 | 0.47 | A>A |
| chr4 | 1.32E+08 | 132300000 | 0.45 | 0.55 | A>A |
| chr4 | 1.32E+08 | 132400000 | 0.23 | 0.34 | A>A |
| chr4 | 1.32E+08 | 132500000 | 0.37 | 0.47 | A>A |
| chr4 | 1.33E+08 | 132600000 | 0.16 | 0.39 | A>A |
| chr4 | 1.33E+08 | 132700000 | 0.00 | 0.00 | NO |
| chr4 | 1.33E+08 | 132800000 | 0.00 | 0.00 | NO |
| chr4 | 1.33E+08 | 132900000 | 0.00 | 0.00 | NO |
| chr4 | 1.33E+08 | 133000000 | 0.31 | 0.41 | A>A |
| chr4 | 1.33E+08 | 133100000 | 0.15 | 0.39 | A>A |
| chr4 | 1.33E+08 | 133200000 | -0.01 | 0.25 | A>A |
| chr4 | 1.33E+08 | 133300000 | -0.01 | 0.34 | A>A |
| chr4 | 1.33E+08 | 133400000 | -0.13 | 0.36 | A>A |
| chr4 | 1.33E+08 | 133500000 | -0.07 | 0.26 | A>A |
| chr4 | 1.34E+08 | 133600000 | -0.08 | 0.01 | B>B |
| chr4 | 1.34E+08 | 133700000 | -0.12 | -0.20 | B>B |
| chr4 | 1.34E+08 | 133800000 | -0.26 | -0.44 | B>B |
| chr4 | 1.34E+08 | 133900000 | -0.25 | -0.47 | B>B |
| chr4 | 1.34E+08 | 134000000 | -0.40 | -0.47 | B>B |
| chr4 | 1.34E+08 | 134100000 | -0.25 | -0.31 | B>B |
| chr4 | 1.34E+08 | 134200000 | -0.17 | -0.34 | B>B |
| chr4 | 1.34E+08 | 134300000 | -0.20 | -0.31 | B>B |
| chr4 | 1.34E+08 | 134400000 | -0.30 | -0.35 | B>B |
| chr4 | 1.34E+08 | 134500000 | -0.22 | -0.24 | B>B |
| chr4 | 1.35E+08 | 134600000 | -0.20 | -0.28 | B>B |
| chr4 | 1.35E+08 | 134700000 | -0.30 | -0.21 | B>B |
| chr4 | 1.35E+08 | 134800000 | -0.36 | -0.33 | B>B |
| chr4 | 1.35E+08 | 134900000 | -0.23 | -0.34 | B>B |
| chr4 | 1.35E+08 | 135000000 | -0.21 | -0.29 | B>B |
| chr4 | 1.35E+08 | 135100000 | -0.25 | -0.32 | B>B |
| chr4 | 1.35E+08 | 135200000 | -0.35 | -0.32 | B>B |
| chr4 | 1.35E+08 | 135300000 | -0.25 | -0.34 | B>B |
| chr4 | 1.35E+08 | 135400000 | -0.42 | -0.36 | B>B |
| chr4 | 1.35E+08 | 135500000 | -0.43 | -0.33 | B>B |
| chr4 | 1.36E+08 | 135600000 | -0.38 | -0.31 | B>B |
| chr4 | 1.36E+08 | 135700000 | -0.33 | -0.36 | B>B |
| chr4 | 1.36E+08 | 135800000 | -0.49 | -0.38 | B>B |
| chr4 | 1.36E+08 | 135900000 | -0.37 | -0.42 | B>B |
| chr4 | 1.36E+08 | 136000000 | -0.28 | -0.34 | B>B |
| chr4 | 1.36E+08 | 136100000 | -0.16 | -0.36 | B>B |
| chr4 | 1.36E+08 | 136200000 | -0.05 | -0.30 | B>B |
| chr4 | 1.36E+08 | 136300000 | -0.06 | -0.33 | B>B |
| chr4 | 1.36E+08 | 136400000 | -0.14 | -0.30 | B>B |
| chr4 | 1.36E+08 | 136500000 | -0.10 | -0.07 | B>B |
| chr4 | 1.37E+08 | 136600000 | -0.11 | -0.08 | B>B |

| | | | | | |
|---|---|---|---|---|---|
| chr4 | 1.37E+08 | 136700000 | 0.09 | 0.02 | A>A |
| chr4 | 1.37E+08 | 136800000 | 0.04 | 0.08 | A>A |
| chr4 | 1.37E+08 | 136900000 | 0.06 | 0.07 | A>A |
| chr4 | 1.37E+08 | 137000000 | 0.10 | 0.23 | A>A |
| chr4 | 1.37E+08 | 137100000 | 0.20 | 0.33 | A>A |
| chr4 | 1.37E+08 | 137200000 | 0.10 | 0.27 | A>A |
| chr4 | 1.37E+08 | 137300000 | 0.12 | 0.28 | A>A |
| chr4 | 1.37E+08 | 137400000 | -0.02 | 0.30 | A>A |
| chr4 | 1.37E+08 | 137500000 | -0.04 | 0.24 | A>A |
| chr4 | 1.38E+08 | 137600000 | -0.12 | 0.17 | A>A |
| chr4 | 1.38E+08 | 137700000 | 0.02 | 0.41 | A>A |
| chr4 | 1.38E+08 | 137800000 | -0.16 | 0.17 | A>A |
| chr4 | 1.38E+08 | 137900000 | -0.36 | 0.27 | B>A |
| chr4 | 1.38E+08 | 138000000 | -0.40 | 0.42 | B>A |
| chr4 | 1.38E+08 | 138100000 | -0.36 | 0.28 | B>A |
| chr4 | 1.38E+08 | 138200000 | -0.46 | 0.31 | B>A |
| chr4 | 1.38E+08 | 138300000 | -0.41 | 0.37 | B>A |
| chr4 | 1.38E+08 | 138400000 | -0.33 | 0.43 | B>A |
| chr4 | 1.38E+08 | 138500000 | -0.17 | 0.43 | B>A |
| chr4 | 1.39E+08 | 138600000 | -0.16 | 0.36 | B>A |
| chr4 | 1.39E+08 | 138700000 | -0.05 | 0.17 | A>A |
| chr4 | 1.39E+08 | 138800000 | 0.06 | 0.03 | A>A |
| chr4 | 1.39E+08 | 138900000 | 0.15 | 0.13 | A>A |
| chr4 | 1.39E+08 | 139000000 | 0.27 | 0.15 | A>A |
| chr4 | 1.39E+08 | 139100000 | 0.44 | 0.34 | A>A |
| chr4 | 1.39E+08 | 139200000 | 0.84 | 0.70 | A>A |
| chr4 | 1.39E+08 | 139300000 | 1.16 | 0.61 | A>A |
| chr4 | 1.39E+08 | 139400000 | 1.00 | 0.58 | A>A |
| chr4 | 1.39E+08 | 139500000 | 0.61 | 0.11 | A>A |
| chr4 | 1.4E+08 | 139600000 | 0.41 | 0.03 | A>A |
| chr4 | 1.4E+08 | 139700000 | 0.43 | 0.02 | A>A |
| chr4 | 1.4E+08 | 139800000 | 0.64 | 0.16 | A>A |
| chr4 | 1.4E+08 | 139900000 | 0.71 | 0.52 | A>A |
| chr4 | 1.4E+08 | 140000000 | 1.24 | 1.02 | A>A |
| chr4 | 1.4E+08 | 140100000 | 1.58 | 1.24 | A>A |
| chr4 | 1.4E+08 | 140200000 | 1.59 | 1.35 | A>A |
| chr4 | 1.4E+08 | 140300000 | 1.76 | 1.46 | A>A |
| chr4 | 1.4E+08 | 140400000 | 1.50 | 1.51 | A>A |
| chr4 | 1.4E+08 | 140500000 | 1.63 | 1.47 | A>A |
| chr4 | 1.41E+08 | 140600000 | 1.29 | 1.26 | A>A |
| chr4 | 1.41E+08 | 140700000 | 1.13 | 1.21 | A>A |
| chr4 | 1.41E+08 | 140800000 | 1.11 | 1.30 | A>A |
| chr4 | 1.41E+08 | 140900000 | 1.11 | 1.25 | A>A |
| chr4 | 1.41E+08 | 141000000 | 1.21 | 1.31 | A>A |
| chr4 | 1.41E+08 | 141100000 | 1.43 | 1.45 | A>A |
| chr4 | 1.41E+08 | 141200000 | 1.36 | 1.43 | A>A |
| chr4 | 1.41E+08 | 141300000 | 0.95 | 0.96 | A>A |
| chr4 | 1.41E+08 | 141400000 | 0.92 | 0.98 | A>A |
| chr4 | 1.41E+08 | 141500000 | 0.92 | 0.94 | A>A |
| chr4 | 1.42E+08 | 141600000 | 0.90 | 0.83 | A>A |
| chr4 | 1.42E+08 | 141700000 | 1.16 | 1.14 | A>A |
| chr4 | 1.42E+08 | 141800000 | 0.66 | 0.62 | A>A |
| chr4 | 1.42E+08 | 141900000 | 0.57 | 0.68 | A>A |
| chr4 | 1.42E+08 | 142000000 | 0.34 | 0.76 | A>A |
| chr4 | 1.42E+08 | 142100000 | 0.25 | 0.46 | A>A |
| chr4 | 1.42E+08 | 142200000 | 0.39 | 0.53 | A>A |

| | | | | | |
|---|---|---|---|---|---|
| chr4 | 1.42E+08 | 142300000 | 0.40 | 0.55 | A>A |
| chr4 | 1.42E+08 | 142400000 | 0.95 | 1.06 | A>A |
| chr4 | 1.42E+08 | 142500000 | 0.87 | 1.10 | A>A |
| chr4 | 1.43E+08 | 142600000 | 0.81 | 0.87 | A>A |
| chr4 | 1.43E+08 | 142700000 | 0.70 | 0.76 | A>A |
| chr4 | 1.43E+08 | 142800000 | 0.71 | 0.73 | A>A |
| chr4 | 1.43E+08 | 142900000 | 0.66 | 0.85 | A>A |
| chr4 | 1.43E+08 | 143000000 | 0.50 | 0.75 | A>A |
| chr4 | 1.43E+08 | 143100000 | 0.43 | 0.66 | A>A |
| chr4 | 1.43E+08 | 143200000 | 0.64 | 0.89 | A>A |
| chr4 | 1.43E+08 | 143300000 | 0.41 | 0.85 | A>A |
| chr4 | 1.43E+08 | 143400000 | 0.68 | 1.08 | A>A |
| chr4 | 1.43E+08 | 143500000 | 0.59 | 0.91 | A>A |
| chr4 | 1.44E+08 | 143600000 | 0.47 | 0.96 | A>A |
| chr4 | 1.44E+08 | 143700000 | 0.27 | 0.91 | B>A |
| chr4 | 1.44E+08 | 143800000 | 0.08 | 0.60 | B>A |
| chr4 | 1.44E+08 | 143900000 | -0.12 | 0.32 | A>A |
| chr4 | 1.44E+08 | 144000000 | -0.15 | 0.19 | A>A |
| chr4 | 1.44E+08 | 144100000 | 0.03 | 0.17 | A>A |
| chr4 | 1.44E+08 | 144200000 | 0.31 | 0.37 | A>A |
| chr4 | 1.44E+08 | 144300000 | 0.61 | 0.36 | A>A |
| chr4 | 1.44E+08 | 144400000 | 0.47 | 0.23 | A>A |
| chr4 | 1.44E+08 | 144500000 | 0.13 | -0.15 | B>B |
| chr4 | 1.45E+08 | 144600000 | -0.36 | -1.32 | A>B |
| chr4 | 1.45E+08 | 144700000 | -0.52 | -1.31 | A>B |
| chr4 | 1.45E+08 | 144800000 | 0.00 | 0.00 | NO |
| chr4 | 1.45E+08 | 144900000 | 0.00 | 0.00 | NO |
| chr4 | 1.45E+08 | 145000000 | 0.00 | 0.00 | NO |
| chr4 | 1.45E+08 | 145100000 | 0.00 | 0.00 | NO |
| chr4 | 1.45E+08 | 145200000 | -0.09 | -0.97 | A>B |
| chr4 | 1.45E+08 | 145300000 | 0.21 | -0.75 | A>B |
| chr4 | 1.45E+08 | 145400000 | 0.23 | -0.66 | A>B |
| chr4 | 1.45E+08 | 145500000 | 0.31 | -0.62 | A>B |
| chr4 | 1.46E+08 | 145600000 | 0.63 | -0.23 | A>B |
| chr4 | 1.46E+08 | 145700000 | 0.37 | -0.40 | A>B |
| chr4 | 1.46E+08 | 145800000 | 0.11 | -0.79 | A>B |
| chr4 | 1.46E+08 | 145900000 | 0.05 | -0.91 | A>B |
| chr4 | 1.46E+08 | 146000000 | 0.27 | -0.34 | A>B |
| chr4 | 1.46E+08 | 146100000 | 0.65 | 0.00 | A>B |
| chr4 | 1.46E+08 | 146200000 | 0.36 | -0.34 | A>B |
| chr4 | 1.46E+08 | 146300000 | 0.14 | -0.77 | A>B |
| chr4 | 1.46E+08 | 146400000 | 0.16 | -0.55 | A>B |
| chr4 | 1.46E+08 | 146500000 | 0.52 | -0.17 | A>B |
| chr4 | 1.47E+08 | 146600000 | 0.46 | -0.23 | A>B |
| chr4 | 1.47E+08 | 146700000 | 0.86 | 0.02 | A>B |
| chr4 | 1.47E+08 | 146800000 | 0.81 | 0.08 | A>B |
| chr4 | 1.47E+08 | 146900000 | 0.47 | -0.23 | A>B |
| chr4 | 1.47E+08 | 147000000 | 0.18 | -0.61 | A>B |
| chr4 | 1.47E+08 | 147100000 | 0.14 | -0.61 | A>B |
| chr4 | 1.47E+08 | 147200000 | 0.21 | -0.26 | B>B |
| chr4 | 1.47E+08 | 147300000 | 0.07 | -0.33 | B>B |
| chr4 | 1.47E+08 | 147400000 | 0.18 | -0.34 | B>B |
| chr4 | 1.47E+08 | 147500000 | 0.09 | -0.46 | B>B |
| chr4 | 1.48E+08 | 147600000 | -0.19 | -0.67 | B>B |
| chr4 | 1.48E+08 | 147700000 | -0.36 | -0.84 | B>B |
| chr4 | 1.48E+08 | 147800000 | -0.34 | -0.78 | B>B |

| | | | | | |
|---|---|---|---|---|---|
| chr4 | 1.48E+08 | 147900000 | -0.05 | -0.30 | B>B |
| chr4 | 1.48E+08 | 148000000 | 0.16 | -0.07 | A>A |
| chr4 | 1.48E+08 | 148100000 | 0.40 | 0.16 | A>A |
| chr4 | 1.48E+08 | 148200000 | 0.88 | 0.60 | A>A |
| chr4 | 1.48E+08 | 148300000 | 0.91 | 0.74 | A>A |
| chr4 | 1.48E+08 | 148400000 | 0.69 | 0.63 | A>A |
| chr4 | 1.48E+08 | 148500000 | 0.59 | 0.51 | A>A |
| chr4 | 1.49E+08 | 148600000 | 0.58 | 0.34 | A>A |
| chr4 | 1.49E+08 | 148700000 | 0.91 | 0.61 | A>A |
| chr4 | 1.49E+08 | 148800000 | 0.89 | 0.55 | A>A |
| chr4 | 1.49E+08 | 148900000 | 0.32 | 0.01 | A>A |
| chr4 | 1.49E+08 | 149000000 | 0.22 | -0.21 | A>A |
| chr4 | 1.49E+08 | 149100000 | -0.16 | -0.38 | B>B |
| chr4 | 1.49E+08 | 149200000 | -0.33 | -0.45 | B>B |
| chr4 | 1.49E+08 | 149300000 | -0.47 | -0.37 | B>B |
| chr4 | 1.49E+08 | 149400000 | -0.36 | -0.37 | B>B |
| chr4 | 1.49E+08 | 149500000 | -0.60 | -0.88 | B>B |
| chr4 | 1.5E+08 | 149600000 | -0.80 | -1.08 | B>B |
| chr4 | 1.5E+08 | 149700000 | -0.66 | -1.00 | B>B |
| chr4 | 1.5E+08 | 149800000 | -0.68 | -1.02 | B>B |
| chr4 | 1.5E+08 | 149900000 | -0.73 | -1.06 | B>B |
| chr4 | 1.5E+08 | 150000000 | -0.69 | -1.15 | B>B |
| chr4 | 1.5E+08 | 150100000 | -0.77 | -1.21 | B>B |
| chr4 | 1.5E+08 | 150200000 | -0.77 | -0.86 | B>B |
| chr4 | 1.5E+08 | 150300000 | -0.63 | -0.41 | B>B |
| chr4 | 1.5E+08 | 150400000 | -0.63 | 0.00 | B>A |
| chr4 | 1.5E+08 | 150500000 | -0.63 | -0.03 | B>A |
| chr4 | 1.51E+08 | 150600000 | -0.54 | -0.04 | B>B |
| chr4 | 1.51E+08 | 150700000 | -0.32 | -0.09 | B>B |
| chr4 | 1.51E+08 | 150800000 | -0.18 | -0.09 | B>B |
| chr4 | 1.51E+08 | 150900000 | 0.15 | 0.09 | A>A |
| chr4 | 1.51E+08 | 151000000 | 0.76 | 0.71 | A>A |
| chr4 | 1.51E+08 | 151100000 | 1.00 | 0.97 | A>A |
| chr4 | 1.51E+08 | 151200000 | 0.80 | 0.78 | A>A |
| chr4 | 1.51E+08 | 151300000 | 0.74 | 0.84 | A>A |
| chr4 | 1.51E+08 | 151400000 | 0.85 | 1.00 | A>A |
| chr4 | 1.51E+08 | 151500000 | 0.85 | 1.04 | A>A |
| chr4 | 1.52E+08 | 151600000 | 0.78 | 0.97 | A>A |
| chr4 | 1.52E+08 | 151700000 | 0.77 | 0.98 | A>A |
| chr4 | 1.52E+08 | 151800000 | 0.67 | 0.88 | A>A |
| chr4 | 1.52E+08 | 151900000 | 1.23 | 0.91 | A>A |
| chr4 | 1.52E+08 | 152000000 | 1.32 | 0.99 | A>A |
| chr4 | 1.52E+08 | 152100000 | 0.99 | 0.60 | A>A |
| chr4 | 1.52E+08 | 152200000 | 1.26 | 0.68 | A>B |
| chr4 | 1.52E+08 | 152300000 | 1.23 | 0.55 | A>B |
| chr4 | 1.52E+08 | 152400000 | 1.03 | 0.21 | A>B |
| chr4 | 1.52E+08 | 152500000 | 0.93 | 0.27 | A>B |
| chr4 | 1.53E+08 | 152600000 | 0.73 | 0.11 | A>B |
| chr4 | 1.53E+08 | 152700000 | 0.83 | 0.07 | A>B |
| chr4 | 1.53E+08 | 152800000 | 0.89 | -0.11 | A>B |
| chr4 | 1.53E+08 | 152900000 | 0.84 | -0.21 | A>B |
| chr4 | 1.53E+08 | 153000000 | 0.83 | 0.04 | A>B |
| chr4 | 1.53E+08 | 153100000 | 0.93 | 0.11 | A>B |
| chr4 | 1.53E+08 | 153200000 | 0.88 | 0.07 | A>B |
| chr4 | 1.53E+08 | 153300000 | 0.92 | 0.25 | A>B |
| chr4 | 1.53E+08 | 153400000 | 0.94 | 0.54 | A>A |

| | | | | | |
|---|---|---|---|---|---|
| chr4 | 1.53E+08 | 153500000 | 1.44 | 0.92 | A>A |
| chr4 | 1.54E+08 | 153600000 | 1.33 | 0.84 | A>A |
| chr4 | 1.54E+08 | 153700000 | 1.08 | 0.65 | A>A |
| chr4 | 1.54E+08 | 153800000 | 1.07 | 0.66 | A>A |
| chr4 | 1.54E+08 | 153900000 | 1.27 | 0.81 | A>A |
| chr4 | 1.54E+08 | 154000000 | 1.19 | 0.94 | A>A |
| chr4 | 1.54E+08 | 154100000 | 0.93 | 0.72 | A>A |
| chr4 | 1.54E+08 | 154200000 | 0.97 | 0.71 | A>A |
| chr4 | 1.54E+08 | 154300000 | 0.89 | 0.47 | A>A |
| chr4 | 1.54E+08 | 154400000 | 1.01 | 0.52 | A>A |
| chr4 | 1.54E+08 | 154500000 | 0.78 | 0.49 | A>A |
| chr4 | 1.55E+08 | 154600000 | 0.56 | -0.11 | A>B |
| chr4 | 1.55E+08 | 154700000 | 0.17 | -0.60 | A>B |
| chr4 | 1.55E+08 | 154800000 | -0.36 | -0.95 | A>B |
| chr4 | 1.55E+08 | 154900000 | -0.55 | -1.15 | A>B |
| chr4 | 1.55E+08 | 155000000 | -0.50 | -1.16 | A>B |
| chr4 | 1.55E+08 | 155100000 | -0.48 | -1.21 | A>B |
| chr4 | 1.55E+08 | 155200000 | -0.62 | -1.14 | B>B |
| chr4 | 1.55E+08 | 155300000 | -0.30 | -0.95 | A>B |
| chr4 | 1.55E+08 | 155400000 | -0.20 | -0.82 | A>B |
| chr4 | 1.55E+08 | 155500000 | 0.09 | -0.39 | B>B |
| chr4 | 1.56E+08 | 155600000 | 0.32 | -0.19 | A>A |
| chr4 | 1.56E+08 | 155700000 | -0.16 | -0.62 | B>B |
| chr4 | 1.56E+08 | 155800000 | -0.07 | -0.42 | B>B |
| chr4 | 1.56E+08 | 155900000 | -0.09 | -0.41 | B>B |
| chr4 | 1.56E+08 | 156000000 | -0.07 | -0.37 | B>B |
| chr4 | 1.56E+08 | 156100000 | 0.01 | -0.34 | B>B |
| chr4 | 1.56E+08 | 156200000 | 0.37 | -0.18 | A>A |
| chr4 | 1.56E+08 | 156300000 | 0.31 | -0.14 | A>A |
| chr4 | 1.56E+08 | 156400000 | -0.02 | -0.43 | B>B |
| chr4 | 1.56E+08 | 156500000 | -0.38 | -0.83 | B>B |
| chr4 | 1.57E+08 | 156600000 | -0.33 | -0.84 | B>B |
| chr4 | 1.57E+08 | 156700000 | -0.25 | -0.68 | B>B |
| chr4 | 1.57E+08 | 156800000 | 0.07 | -0.13 | B>B |
| chr4 | 1.57E+08 | 156900000 | 0.10 | -0.25 | B>B |
| chr4 | 1.57E+08 | 157000000 | -0.56 | -0.79 | B>B |
| chr4 | 1.57E+08 | 157100000 | -0.60 | -0.94 | B>B |
| chr4 | 1.57E+08 | 157200000 | -0.69 | -0.90 | B>B |
| chr4 | 1.57E+08 | 157300000 | -0.58 | -0.90 | B>B |
| chr4 | 1.57E+08 | 157400000 | -0.53 | -0.76 | B>B |
| chr4 | 1.57E+08 | 157500000 | -0.30 | -0.25 | B>B |
| chr4 | 1.58E+08 | 157600000 | -0.16 | -0.26 | B>B |
| chr4 | 1.58E+08 | 157700000 | 0.10 | -0.01 | A>A |
| chr4 | 1.58E+08 | 157800000 | 0.29 | 0.12 | A>A |
| chr4 | 1.58E+08 | 157900000 | 0.26 | 0.03 | A>A |
| chr4 | 1.58E+08 | 158000000 | 0.17 | -0.16 | A>A |
| chr4 | 1.58E+08 | 158100000 | -0.14 | -0.36 | B>B |
| chr4 | 1.58E+08 | 158200000 | -0.36 | -0.82 | B>B |
| chr4 | 1.58E+08 | 158300000 | -0.53 | -0.99 | B>B |
| chr4 | 1.58E+08 | 158400000 | -0.62 | -1.01 | B>B |
| chr4 | 1.58E+08 | 158500000 | -0.69 | -1.02 | B>B |
| chr4 | 1.59E+08 | 158600000 | -0.77 | -0.98 | B>B |
| chr4 | 1.59E+08 | 158700000 | -0.82 | -0.93 | B>B |
| chr4 | 1.59E+08 | 158800000 | -0.65 | -0.81 | B>B |
| chr4 | 1.59E+08 | 158900000 | -0.44 | -0.58 | B>B |
| chr4 | 1.59E+08 | 159000000 | 0.34 | 0.16 | A>A |

| | | | | | |
|---|---|---|---|---|---|
| chr4 | 1.59E+08 | 159100000 | 0.47 | 0.20 | A>A |
| chr4 | 1.59E+08 | 159200000 | 0.28 | 0.28 | A>A |
| chr4 | 1.59E+08 | 159300000 | 0.86 | 0.81 | A>A |
| chr4 | 1.59E+08 | 159400000 | 0.78 | 0.77 | A>A |
| chr4 | 1.59E+08 | 159500000 | 0.81 | 0.91 | A>A |
| chr4 | 1.6E+08 | 159600000 | 0.62 | 0.57 | A>A |
| chr4 | 1.6E+08 | 159700000 | 0.91 | 0.87 | A>A |
| chr4 | 1.6E+08 | 159800000 | 1.03 | 0.92 | A>A |
| chr4 | 1.6E+08 | 159900000 | 1.20 | 1.10 | A>A |
| chr4 | 1.6E+08 | 160000000 | 1.29 | 1.09 | A>A |
| chr4 | 1.6E+08 | 160100000 | 1.15 | 1.02 | A>A |
| chr4 | 1.6E+08 | 160200000 | 0.78 | 0.85 | A>A |
| chr4 | 1.6E+08 | 160300000 | 0.66 | 0.77 | A>A |
| chr4 | 1.6E+08 | 160400000 | 0.29 | 0.41 | A>A |
| chr4 | 1.6E+08 | 160500000 | -0.43 | -0.17 | B>B |
| chr4 | 1.61E+08 | 160600000 | -0.70 | -0.25 | B>B |
| chr4 | 1.61E+08 | 160700000 | -0.62 | -0.12 | B>B |
| chr4 | 1.61E+08 | 160800000 | -0.61 | 0.11 | B>A |
| chr4 | 1.61E+08 | 160900000 | -0.28 | 0.01 | B>B |
| chr4 | 1.61E+08 | 161000000 | 0.08 | 0.15 | A>A |
| chr4 | 1.61E+08 | 161100000 | 0.11 | 0.11 | A>A |
| chr4 | 1.61E+08 | 161200000 | 0.21 | 0.17 | A>A |
| chr4 | 1.61E+08 | 161300000 | 0.26 | 0.19 | A>A |
| chr4 | 1.61E+08 | 161400000 | 0.21 | 0.05 | A>A |
| chr4 | 1.61E+08 | 161500000 | 0.05 | -0.09 | B>B |
| chr4 | 1.62E+08 | 161600000 | -0.20 | -0.05 | B>B |
| chr4 | 1.62E+08 | 161700000 | -0.35 | -0.02 | B>B |
| chr4 | 1.62E+08 | 161800000 | -0.39 | 0.04 | B>B |
| chr4 | 1.62E+08 | 161900000 | -0.56 | -0.07 | B>B |
| chr4 | 1.62E+08 | 162000000 | -0.72 | 0.04 | B>A |
| chr4 | 1.62E+08 | 162100000 | -0.66 | -0.04 | B>A |
| chr4 | 1.62E+08 | 162200000 | -0.52 | -0.05 | B>B |
| chr4 | 1.62E+08 | 162300000 | -0.29 | 0.20 | B>B |
| chr4 | 1.62E+08 | 162400000 | -0.24 | 0.28 | B>A |
| chr4 | 1.62E+08 | 162500000 | -0.16 | 0.31 | A>A |
| chr4 | 1.63E+08 | 162600000 | 0.00 | 0.41 | A>A |
| chr4 | 1.63E+08 | 162700000 | 0.10 | 0.53 | A>A |
| chr4 | 1.63E+08 | 162800000 | 0.16 | 0.43 | A>A |
| chr4 | 1.63E+08 | 162900000 | 0.24 | 0.35 | A>A |
| chr4 | 1.63E+08 | 163000000 | -0.07 | 0.30 | A>A |
| chr4 | 1.63E+08 | 163100000 | -0.17 | 0.26 | A>A |
| chr4 | 1.63E+08 | 163200000 | -0.49 | 0.29 | B>A |
| chr4 | 1.63E+08 | 163300000 | -0.52 | 0.25 | B>A |
| chr4 | 1.63E+08 | 163400000 | -0.44 | 0.02 | B>B |
| chr4 | 1.63E+08 | 163500000 | -0.11 | -0.28 | B>B |
| chr4 | 1.64E+08 | 163600000 | 0.13 | -0.38 | B>B |
| chr4 | 1.64E+08 | 163700000 | 0.33 | -0.11 | A>A |
| chr4 | 1.64E+08 | 163800000 | 0.27 | -0.20 | A>A |
| chr4 | 1.64E+08 | 163900000 | 0.09 | -0.28 | B>B |
| chr4 | 1.64E+08 | 164000000 | 0.00 | -0.38 | B>B |
| chr4 | 1.64E+08 | 164100000 | 0.13 | -0.29 | B>B |
| chr4 | 1.64E+08 | 164200000 | -0.23 | -0.70 | B>B |
| chr4 | 1.64E+08 | 164300000 | -0.33 | -0.61 | B>B |
| chr4 | 1.64E+08 | 164400000 | -0.16 | -0.70 | B>B |
| chr4 | 1.64E+08 | 164500000 | -0.05 | -0.42 | B>B |
| chr4 | 1.65E+08 | 164600000 | -0.61 | -0.90 | B>B |

| | | | | | |
|---|---|---|---|---|---|
| chr4 | 1.65E+08 | 164700000 | -0.66 | -0.98 | B>B |
| chr4 | 1.65E+08 | 164800000 | -0.76 | -1.07 | B>B |
| chr4 | 1.65E+08 | 164900000 | -0.82 | -1.06 | B>B |
| chr4 | 1.65E+08 | 165000000 | -0.77 | -1.04 | B>B |
| chr4 | 1.65E+08 | 165100000 | -0.78 | -1.13 | B>B |
| chr4 | 1.65E+08 | 165200000 | -0.79 | -1.06 | B>B |
| chr4 | 1.65E+08 | 165300000 | -0.79 | -1.13 | B>B |
| chr4 | 1.65E+08 | 165400000 | -0.81 | -1.04 | B>B |
| chr4 | 1.65E+08 | 165500000 | -0.87 | -1.06 | B>B |
| chr4 | 1.66E+08 | 165600000 | -0.66 | -1.06 | B>B |
| chr4 | 1.66E+08 | 165700000 | -0.35 | -0.67 | B>B |
| chr4 | 1.66E+08 | 165800000 | -0.19 | -0.88 | A>B |
| chr4 | 1.66E+08 | 165900000 | 0.05 | -0.96 | A>B |
| chr4 | 1.66E+08 | 166000000 | 0.26 | -0.53 | A>B |
| chr4 | 1.66E+08 | 166100000 | 0.18 | -0.01 | A>A |
| chr4 | 1.66E+08 | 166200000 | 0.00 | 0.00 | NO |
| chr4 | 1.66E+08 | 166300000 | 0.54 | 0.25 | A>A |
| chr4 | 1.66E+08 | 166400000 | 0.21 | 0.25 | A>A |
| chr4 | 1.66E+08 | 166500000 | -0.11 | 0.21 | A>A |
| chr4 | 1.67E+08 | 166600000 | -0.11 | 0.62 | B>A |
| chr4 | 1.67E+08 | 166700000 | 0.02 | 0.67 | B>A |
| chr4 | 1.67E+08 | 166800000 | -0.17 | 0.52 | B>A |
| chr4 | 1.67E+08 | 166900000 | -0.17 | 0.50 | B>A |
| chr4 | 1.67E+08 | 167000000 | -0.20 | 0.43 | B>A |
| chr4 | 1.67E+08 | 167100000 | -0.41 | 0.09 | B>B |
| chr4 | 1.67E+08 | 167200000 | -0.56 | -0.09 | B>B |
| chr4 | 1.67E+08 | 167300000 | -0.70 | -0.11 | B>A |
| chr4 | 1.67E+08 | 167400000 | -0.66 | -0.02 | B>A |
| chr4 | 1.67E+08 | 167500000 | -0.70 | -0.17 | B>A |
| chr4 | 1.68E+08 | 167600000 | -0.50 | -0.04 | B>B |
| chr4 | 1.68E+08 | 167700000 | -0.39 | 0.25 | B>A |
| chr4 | 1.68E+08 | 167800000 | -0.16 | 0.24 | A>A |
| chr4 | 1.68E+08 | 167900000 | 0.11 | 0.30 | A>A |
| chr4 | 1.68E+08 | 168000000 | 0.08 | 0.34 | A>A |
| chr4 | 1.68E+08 | 168100000 | 0.37 | 0.63 | A>A |
| chr4 | 1.68E+08 | 168200000 | 0.25 | 0.54 | A>A |
| chr4 | 1.68E+08 | 168300000 | 0.02 | 0.24 | A>A |
| chr4 | 1.68E+08 | 168400000 | -0.21 | 0.26 | A>A |
| chr4 | 1.68E+08 | 168500000 | -0.22 | 0.34 | B>A |
| chr4 | 1.69E+08 | 168600000 | -0.16 | 0.38 | B>A |
| chr4 | 1.69E+08 | 168700000 | -0.25 | 0.33 | B>A |
| chr4 | 1.69E+08 | 168800000 | -0.15 | 0.44 | B>A |
| chr4 | 1.69E+08 | 168900000 | -0.13 | 0.37 | A>A |
| chr4 | 1.69E+08 | 169000000 | -0.15 | 0.37 | B>A |
| chr4 | 1.69E+08 | 169100000 | 0.16 | 0.51 | A>A |
| chr4 | 1.69E+08 | 169200000 | 0.58 | 0.69 | A>A |
| chr4 | 1.69E+08 | 169300000 | 0.97 | 0.90 | A>A |
| chr4 | 1.69E+08 | 169400000 | 1.04 | 1.03 | A>A |
| chr4 | 1.69E+08 | 169500000 | 1.00 | 0.82 | A>A |
| chr4 | 1.7E+08 | 169600000 | 1.05 | 1.08 | A>A |
| chr4 | 1.7E+08 | 169700000 | 1.06 | 1.06 | A>A |
| chr4 | 1.7E+08 | 169800000 | 1.00 | 1.03 | A>A |
| chr4 | 1.7E+08 | 169900000 | 0.60 | 0.56 | A>A |
| chr4 | 1.7E+08 | 170000000 | 0.91 | 0.84 | A>A |
| chr4 | 1.7E+08 | 170100000 | 1.14 | 1.07 | A>A |
| chr4 | 1.7E+08 | 170200000 | 1.38 | 1.10 | A>A |

| | | | | | |
|---|---|---|---|---|---|
| chr4 | 1.7E+08 | 170300000 | 1.11 | 1.09 | A>A |
| chr4 | 1.7E+08 | 170400000 | 0.89 | 0.98 | A>A |
| chr4 | 1.7E+08 | 170500000 | 0.56 | 0.50 | A>A |
| chr4 | 1.71E+08 | 170600000 | 0.86 | 0.76 | A>A |
| chr4 | 1.71E+08 | 170700000 | 0.75 | 0.49 | A>A |
| chr4 | 1.71E+08 | 170800000 | 0.52 | 0.14 | A>A |
| chr4 | 1.71E+08 | 170900000 | 0.44 | 0.18 | A>A |
| chr4 | 1.71E+08 | 171000000 | 0.39 | 0.05 | A>A |
| chr4 | 1.71E+08 | 171100000 | -0.18 | -0.51 | B>B |
| chr4 | 1.71E+08 | 171200000 | -0.60 | -0.88 | B>B |
| chr4 | 1.71E+08 | 171300000 | -0.68 | -1.01 | B>B |
| chr4 | 1.71E+08 | 171400000 | -0.67 | -1.00 | B>B |
| chr4 | 1.71E+08 | 171500000 | -0.71 | -1.13 | B>B |
| chr4 | 1.72E+08 | 171600000 | -0.73 | -1.06 | B>B |
| chr4 | 1.72E+08 | 171700000 | -0.75 | -1.02 | B>B |
| chr4 | 1.72E+08 | 171800000 | -0.70 | -0.98 | B>B |
| chr4 | 1.72E+08 | 171900000 | -0.76 | -0.96 | B>B |
| chr4 | 1.72E+08 | 172000000 | -0.60 | -0.95 | B>B |
| chr4 | 1.72E+08 | 172100000 | -0.63 | -0.95 | B>B |
| chr4 | 1.72E+08 | 172200000 | -0.62 | -0.91 | B>B |
| chr4 | 1.72E+08 | 172300000 | -0.68 | -0.94 | B>B |
| chr4 | 1.72E+08 | 172400000 | -0.65 | -0.93 | B>B |
| chr4 | 1.72E+08 | 172500000 | -0.64 | -0.91 | B>B |
| chr4 | 1.73E+08 | 172600000 | -0.70 | -0.92 | B>B |
| chr4 | 1.73E+08 | 172700000 | -0.52 | -0.93 | B>B |
| chr4 | 1.73E+08 | 172800000 | -0.32 | -0.84 | B>B |
| chr4 | 1.73E+08 | 172900000 | -0.26 | -0.89 | A>B |
| chr4 | 1.73E+08 | 173000000 | -0.12 | -0.85 | A>B |
| chr4 | 1.73E+08 | 173100000 | -0.02 | -0.65 | A>B |
| chr4 | 1.73E+08 | 173200000 | 0.01 | -0.55 | B>B |
| chr4 | 1.73E+08 | 173300000 | 0.18 | -0.49 | A>B |
| chr4 | 1.73E+08 | 173400000 | 0.34 | -0.43 | A>B |
| chr4 | 1.73E+08 | 173500000 | 0.43 | -0.22 | A>B |
| chr4 | 1.74E+08 | 173600000 | 0.52 | -0.04 | A>A |
| chr4 | 1.74E+08 | 173700000 | 0.35 | -0.04 | A>A |
| chr4 | 1.74E+08 | 173800000 | 0.26 | -0.09 | A>A |
| chr4 | 1.74E+08 | 173900000 | 0.54 | 0.27 | A>A |
| chr4 | 1.74E+08 | 174000000 | 0.46 | 0.38 | A>A |
| chr4 | 1.74E+08 | 174100000 | 0.87 | 0.60 | A>A |
| chr4 | 1.74E+08 | 174200000 | 0.65 | 0.39 | A>A |
| chr4 | 1.74E+08 | 174300000 | 0.57 | 0.31 | A>A |
| chr4 | 1.74E+08 | 174400000 | 0.39 | 0.23 | A>A |
| chr4 | 1.74E+08 | 174500000 | 0.14 | 0.21 | A>A |
| chr4 | 1.75E+08 | 174600000 | 0.17 | 0.42 | A>A |
| chr4 | 1.75E+08 | 174700000 | -0.10 | 0.22 | A>A |
| chr4 | 1.75E+08 | 174800000 | -0.10 | 0.24 | A>A |
| chr4 | 1.75E+08 | 174900000 | -0.01 | 0.14 | A>A |
| chr4 | 1.75E+08 | 175000000 | -0.03 | 0.16 | A>A |
| chr4 | 1.75E+08 | 175100000 | 0.27 | 0.31 | A>A |
| chr4 | 1.75E+08 | 175200000 | 0.44 | 0.57 | A>A |
| chr4 | 1.75E+08 | 175300000 | 0.49 | 0.55 | A>A |
| chr4 | 1.75E+08 | 175400000 | 1.00 | 1.04 | A>A |
| chr4 | 1.75E+08 | 175500000 | 0.95 | 0.95 | A>A |
| chr4 | 1.76E+08 | 175600000 | 0.68 | 0.79 | A>A |
| chr4 | 1.76E+08 | 175700000 | 0.29 | 0.54 | A>A |
| chr4 | 1.76E+08 | 175800000 | 0.13 | 0.52 | A>A |

| | | | | | |
|---|---|---|---|---|---|
| chr4 | 1.76E+08 | 175900000 | 0.00 | 0.66 | B>A |
| chr4 | 1.76E+08 | 176000000 | 0.02 | 0.84 | B>A |
| chr4 | 1.76E+08 | 176100000 | 0.01 | 0.72 | B>A |
| chr4 | 1.76E+08 | 176200000 | 0.03 | 0.57 | B>A |
| chr4 | 1.76E+08 | 176300000 | 0.28 | 0.75 | A>A |
| chr4 | 1.76E+08 | 176400000 | 0.26 | 0.76 | A>A |
| chr4 | 1.76E+08 | 176500000 | -0.03 | 0.41 | A>A |
| chr4 | 1.77E+08 | 176600000 | -0.08 | 0.49 | B>A |
| chr4 | 1.77E+08 | 176700000 | -0.11 | 0.58 | B>A |
| chr4 | 1.77E+08 | 176800000 | -0.24 | 0.56 | B>A |
| chr4 | 1.77E+08 | 176900000 | -0.16 | 0.53 | B>A |
| chr4 | 1.77E+08 | 177000000 | 0.08 | 0.58 | A>A |
| chr4 | 1.77E+08 | 177100000 | 0.46 | 0.67 | A>A |
| chr4 | 1.77E+08 | 177200000 | 0.42 | 0.76 | A>A |
| chr4 | 1.77E+08 | 177300000 | 0.35 | 0.67 | A>A |
| chr4 | 1.77E+08 | 177400000 | 0.55 | 0.91 | A>A |
| chr4 | 1.77E+08 | 177500000 | 0.63 | 0.96 | A>A |
| chr4 | 1.78E+08 | 177600000 | 0.73 | 0.94 | A>A |
| chr4 | 1.78E+08 | 177700000 | 0.79 | 1.00 | A>A |
| chr4 | 1.78E+08 | 177800000 | 0.75 | 0.99 | A>A |
| chr4 | 1.78E+08 | 177900000 | 0.85 | 1.15 | A>A |
| chr4 | 1.78E+08 | 178000000 | 0.47 | 0.85 | A>A |
| chr4 | 1.78E+08 | 178100000 | 0.42 | 0.68 | A>A |
| chr4 | 1.78E+08 | 178200000 | 0.36 | 0.73 | A>A |
| chr4 | 1.78E+08 | 178300000 | 0.61 | 0.70 | A>A |
| chr4 | 1.78E+08 | 178400000 | 0.41 | 0.74 | A>A |
| chr4 | 1.78E+08 | 178500000 | 0.08 | 0.57 | A>A |
| chr4 | 1.79E+08 | 178600000 | -0.10 | 0.40 | A>A |
| chr4 | 1.79E+08 | 178700000 | -0.11 | 0.59 | B>A |
| chr4 | 1.79E+08 | 178800000 | -0.24 | 0.52 | B>A |
| chr4 | 1.79E+08 | 178900000 | -0.21 | 0.68 | B>A |
| chr4 | 1.79E+08 | 179000000 | -0.30 | 0.66 | B>A |
| chr4 | 1.79E+08 | 179100000 | -0.29 | 0.61 | B>A |
| chr4 | 1.79E+08 | 179200000 | -0.29 | 0.58 | B>A |
| chr4 | 1.79E+08 | 179300000 | -0.31 | 0.52 | B>A |
| chr4 | 1.79E+08 | 179400000 | -0.42 | 0.55 | B>A |
| chr4 | 1.79E+08 | 179500000 | -0.28 | 0.53 | B>A |
| chr4 | 1.8E+08 | 179600000 | -0.30 | 0.46 | B>A |
| chr4 | 1.8E+08 | 179700000 | -0.27 | 0.64 | B>A |
| chr4 | 1.8E+08 | 179800000 | -0.29 | 0.65 | B>A |
| chr4 | 1.8E+08 | 179900000 | -0.36 | 0.55 | B>A |
| chr4 | 1.8E+08 | 180000000 | -0.26 | 0.50 | B>A |
| chr4 | 1.8E+08 | 180100000 | -0.36 | 0.36 | B>A |
| chr4 | 1.8E+08 | 180200000 | -0.29 | 0.46 | B>A |
| chr4 | 1.8E+08 | 180300000 | -0.32 | 0.40 | B>A |
| chr4 | 1.8E+08 | 180400000 | -0.25 | 0.47 | B>A |
| chr4 | 1.8E+08 | 180500000 | -0.28 | 0.50 | B>A |
| chr4 | 1.81E+08 | 180600000 | -0.26 | 0.37 | B>A |
| chr4 | 1.81E+08 | 180700000 | -0.34 | 0.44 | B>A |
| chr4 | 1.81E+08 | 180800000 | -0.41 | 0.37 | B>A |
| chr4 | 1.81E+08 | 180900000 | -0.40 | 0.24 | B>A |
| chr4 | 1.81E+08 | 181000000 | -0.47 | 0.20 | B>A |
| chr4 | 1.81E+08 | 181100000 | -0.44 | 0.29 | B>A |
| chr4 | 1.81E+08 | 181200000 | -0.58 | 0.24 | B>A |
| chr4 | 1.81E+08 | 181300000 | -0.70 | 0.29 | B>A |
| chr4 | 1.81E+08 | 181400000 | -0.55 | 0.11 | B>A |

| | | | | | |
|---|---|---|---|---|---|
| chr4 | 1.81E+08 | 181500000 | -0.65 | 0.03 | B>A |
| chr4 | 1.82E+08 | 181600000 | -0.56 | -0.15 | B>B |
| chr4 | 1.82E+08 | 181700000 | -0.74 | -0.68 | B>B |
| chr4 | 1.82E+08 | 181800000 | -0.64 | -0.78 | B>B |
| chr4 | 1.82E+08 | 181900000 | -0.78 | -1.07 | B>B |
| chr4 | 1.82E+08 | 182000000 | -0.76 | -1.09 | B>B |
| chr4 | 1.82E+08 | 182100000 | -0.54 | -0.91 | B>B |
| chr4 | 1.82E+08 | 182200000 | -0.56 | -1.04 | B>B |
| chr4 | 1.82E+08 | 182300000 | -0.66 | -1.05 | B>B |
| chr4 | 1.82E+08 | 182400000 | -0.62 | -1.08 | B>B |
| chr4 | 1.82E+08 | 182500000 | -0.49 | -1.04 | B>B |
| chr4 | 1.83E+08 | 182600000 | -0.51 | -1.08 | B>B |
| chr4 | 1.83E+08 | 182700000 | -0.49 | -1.03 | B>B |
| chr4 | 1.83E+08 | 182800000 | -0.39 | -1.07 | A>B |
| chr4 | 1.83E+08 | 182900000 | -0.35 | -1.00 | A>B |
| chr4 | 1.83E+08 | 183000000 | -0.27 | -0.99 | A>B |
| chr4 | 1.83E+08 | 183100000 | 0.02 | -0.81 | A>B |
| chr4 | 1.83E+08 | 183200000 | 0.17 | -0.87 | A>B |
| chr4 | 1.83E+08 | 183300000 | -0.04 | -0.97 | A>B |
| chr4 | 1.83E+08 | 183400000 | 0.02 | -0.85 | A>B |
| chr4 | 1.83E+08 | 183500000 | 0.05 | -0.86 | A>B |
| chr4 | 1.84E+08 | 183600000 | 0.08 | -0.96 | A>B |
| chr4 | 1.84E+08 | 183700000 | 0.16 | -0.90 | A>B |
| chr4 | 1.84E+08 | 183800000 | 0.32 | -0.33 | A>B |
| chr4 | 1.84E+08 | 183900000 | 0.62 | 0.25 | A>A |
| chr4 | 1.84E+08 | 184000000 | 0.83 | 0.32 | A>A |
| chr4 | 1.84E+08 | 184100000 | 1.19 | 0.72 | A>A |
| chr4 | 1.84E+08 | 184200000 | 0.79 | 0.47 | A>A |
| chr4 | 1.84E+08 | 184300000 | 1.16 | 0.78 | A>A |
| chr4 | 1.84E+08 | 184400000 | 1.24 | 0.84 | A>A |
| chr4 | 1.84E+08 | 184500000 | 1.43 | 1.05 | A>A |
| chr4 | 1.85E+08 | 184600000 | 1.43 | 1.07 | A>A |
| chr4 | 1.85E+08 | 184700000 | 1.30 | 1.24 | A>A |
| chr4 | 1.85E+08 | 184800000 | 1.09 | 0.95 | A>A |
| chr4 | 1.85E+08 | 184900000 | 1.24 | 1.24 | A>A |
| chr4 | 1.85E+08 | 185000000 | 1.29 | 1.26 | A>A |
| chr4 | 1.85E+08 | 185100000 | 1.05 | 0.98 | A>A |
| chr4 | 1.85E+08 | 185200000 | 0.96 | 0.68 | A>A |
| chr4 | 1.85E+08 | 185300000 | 1.42 | 1.19 | A>A |
| chr4 | 1.85E+08 | 185400000 | 1.54 | 1.18 | A>A |
| chr4 | 1.85E+08 | 185500000 | 1.13 | 1.04 | A>A |
| chr4 | 1.86E+08 | 185600000 | 1.17 | 0.99 | A>A |
| chr4 | 1.86E+08 | 185700000 | 1.14 | 0.90 | A>A |
| chr4 | 1.86E+08 | 185800000 | 1.35 | 0.95 | A>A |
| chr4 | 1.86E+08 | 185900000 | 1.25 | 0.87 | A>A |
| chr4 | 1.86E+08 | 186000000 | 1.15 | 0.83 | A>A |
| chr4 | 1.86E+08 | 186100000 | 1.10 | 0.83 | A>A |
| chr4 | 1.86E+08 | 186200000 | 1.48 | 0.91 | A>A |
| chr4 | 1.86E+08 | 186300000 | 1.35 | 0.96 | A>A |
| chr4 | 1.86E+08 | 186400000 | 1.14 | 0.83 | A>A |
| chr4 | 1.86E+08 | 186500000 | 1.09 | 0.86 | A>A |
| chr4 | 1.87E+08 | 186600000 | 0.86 | 0.65 | A>A |
| chr4 | 1.87E+08 | 186700000 | 0.99 | 0.82 | A>A |
| chr4 | 1.87E+08 | 186800000 | 1.26 | 0.95 | A>A |
| chr4 | 1.87E+08 | 186900000 | 1.15 | 0.83 | A>A |
| chr4 | 1.87E+08 | 187000000 | 0.76 | 0.58 | A>A |

| | | | | | |
|---|---|---|---|---|---|
| chr4 | 1.87E+08 | 187100000 | 0.97 | 0.68 | A>A |
| chr4 | 1.87E+08 | 187200000 | 0.83 | 0.51 | A>A |
| chr4 | 1.87E+08 | 187300000 | 0.45 | 0.15 | A>A |
| chr4 | 1.87E+08 | 187400000 | 0.71 | 0.37 | A>A |
| chr4 | 1.87E+08 | 187500000 | 0.57 | 0.20 | A>A |
| chr4 | 1.88E+08 | 187600000 | 0.74 | 0.29 | A>A |
| chr4 | 1.88E+08 | 187700000 | 0.73 | 0.23 | A>A |
| chr4 | 1.88E+08 | 187800000 | 0.89 | 0.35 | A>A |
| chr4 | 1.88E+08 | 187900000 | 0.73 | 0.10 | A>B |
| chr4 | 1.88E+08 | 188000000 | 0.54 | -0.34 | A>B |
| chr4 | 1.88E+08 | 188100000 | 0.55 | -0.35 | A>B |
| chr4 | 1.88E+08 | 188200000 | 0.59 | -0.30 | A>B |
| chr4 | 1.88E+08 | 188300000 | 0.48 | -0.27 | A>B |
| chr4 | 1.88E+08 | 188400000 | 0.46 | -0.39 | A>B |
| chr4 | 1.88E+08 | 188500000 | 0.46 | -0.40 | A>B |
| chr4 | 1.89E+08 | 188600000 | 0.32 | -0.37 | A>B |
| chr4 | 1.89E+08 | 188700000 | 0.33 | -0.40 | A>B |
| chr4 | 1.89E+08 | 188800000 | 0.49 | -0.40 | A>B |
| chr4 | 1.89E+08 | 188900000 | 0.60 | -0.29 | A>B |
| chr4 | 1.89E+08 | 189000000 | 0.69 | 0.02 | A>B |
| chr4 | 1.89E+08 | 189100000 | 0.69 | 0.17 | A>A |
| chr4 | 1.89E+08 | 189200000 | 0.79 | 0.07 | A>B |
| chr4 | 1.89E+08 | 189300000 | 0.73 | -0.02 | A>B |
| chr4 | 1.89E+08 | 189400000 | 0.69 | 0.22 | A>A |
| chr4 | 1.89E+08 | 189500000 | 0.61 | -0.13 | A>B |
| chr4 | 1.9E+08 | 189600000 | 0.67 | -0.30 | A>B |
| chr4 | 1.9E+08 | 189700000 | 0.52 | -0.23 | A>B |
| chr4 | 1.9E+08 | 189800000 | 0.56 | -0.18 | A>B |
| chr4 | 1.9E+08 | 189900000 | 0.67 | 0.01 | A>B |
| chr4 | 1.9E+08 | 190000000 | 0.65 | 0.27 | A>A |
| chr4 | 1.9E+08 | 190100000 | 0.45 | -0.02 | A>A |
| chr4 | 1.9E+08 | 190200000 | 0.03 | -0.61 | A>B |
| chr4 | 1.9E+08 | 190300000 | -0.15 | -0.86 | A>B |
| chr4 | 1.9E+08 | 190400000 | -0.09 | -1.04 | A>B |
| chr4 | 1.9E+08 | 190500000 | -0.20 | -1.03 | A>B |
| chr4 | 1.91E+08 | 190600000 | -0.05 | -0.75 | A>B |
| chr4 | 1.91E+08 | 190700000 | -0.17 | -0.78 | A>B |
| chr4 | 1.91E+08 | 190800000 | 0.02 | -0.64 | A>B |
| chr4 | 1.91E+08 | 190900000 | 0.00 | 0.00 | NO |
| chr4 | 1.91E+08 | 191000000 | 0.00 | 0.00 | NO |
| chr4 | 1.91E+08 | 191100000 | 0.00 | 0.00 | NO |
| chr4 | 1.91E+08 | 191154276 | 0.00 | 0.00 | NO |
| chr5 | 0 | 100000 | 0.00 | 0.00 | NO |
| chr5 | 100000 | 200000 | 1.52 | 1.53 | A>A |
| chr5 | 200000 | 300000 | 0.00 | 0.00 | NO |
| chr5 | 300000 | 400000 | 1.62 | 1.55 | A>A |
| chr5 | 400000 | 500000 | 1.65 | 1.69 | A>A |
| chr5 | 500000 | 600000 | 1.74 | 1.62 | A>A |
| chr5 | 600000 | 700000 | 1.53 | 1.52 | A>A |
| chr5 | 700000 | 800000 | 0.00 | 0.00 | NO |
| chr5 | 800000 | 900000 | 0.00 | 0.00 | NO |
| chr5 | 900000 | 1000000 | 1.30 | 1.46 | A>A |
| chr5 | 1000000 | 1100000 | 1.58 | 1.66 | A>A |
| chr5 | 1100000 | 1200000 | 1.27 | 1.45 | A>A |
| chr5 | 1200000 | 1300000 | 1.43 | 1.45 | A>A |
| chr5 | 1300000 | 1400000 | 1.40 | 1.60 | A>A |

| | | | | | |
|---|---|---|---|---|---|
| chr5 | 1400000 | 1500000 | 1.19 | 1.47 | A>A |
| chr5 | 1500000 | 1600000 | 1.34 | 1.61 | A>A |
| chr5 | 1600000 | 1700000 | 0.80 | 1.00 | A>A |
| chr5 | 1700000 | 1800000 | 0.83 | 0.75 | A>A |
| chr5 | 1800000 | 1900000 | 1.03 | 0.75 | A>A |
| chr5 | 1900000 | 2000000 | 0.64 | 0.37 | A>A |
| chr5 | 2000000 | 2100000 | 0.37 | 0.09 | A>A |
| chr5 | 2100000 | 2200000 | 0.27 | -0.02 | A>A |
| chr5 | 2200000 | 2300000 | 0.07 | -0.39 | B>B |
| chr5 | 2300000 | 2400000 | -0.17 | -0.68 | B>B |
| chr5 | 2400000 | 2500000 | -0.24 | -0.67 | B>B |
| chr5 | 2500000 | 2600000 | -0.32 | -0.62 | B>B |
| chr5 | 2600000 | 2700000 | -0.11 | -0.51 | B>B |
| chr5 | 2700000 | 2800000 | 0.01 | -0.36 | B>B |
| chr5 | 2800000 | 2900000 | -0.36 | -0.62 | B>B |
| chr5 | 2900000 | 3000000 | -0.65 | -0.68 | B>B |
| chr5 | 3000000 | 3100000 | -0.59 | -0.67 | B>B |
| chr5 | 3100000 | 3200000 | -0.57 | -0.56 | B>B |
| chr5 | 3200000 | 3300000 | -0.67 | -0.47 | B>B |
| chr5 | 3300000 | 3400000 | -0.62 | -0.27 | B>B |
| chr5 | 3400000 | 3500000 | -0.71 | -0.42 | B>B |
| chr5 | 3500000 | 3600000 | -0.77 | -0.22 | B>A |
| chr5 | 3600000 | 3700000 | -0.74 | -0.11 | B>A |
| chr5 | 3700000 | 3800000 | -0.79 | 0.03 | B>A |
| chr5 | 3800000 | 3900000 | -0.80 | -0.12 | B>A |
| chr5 | 3900000 | 4000000 | -0.92 | -0.29 | B>A |
| chr5 | 4000000 | 4100000 | -0.88 | -0.47 | B>B |
| chr5 | 4100000 | 4200000 | -0.84 | -0.53 | B>B |
| chr5 | 4200000 | 4300000 | -0.85 | -0.64 | B>B |
| chr5 | 4300000 | 4400000 | -1.02 | -0.79 | B>B |
| chr5 | 4400000 | 4500000 | -0.95 | -0.81 | B>B |
| chr5 | 4500000 | 4600000 | -0.97 | -0.78 | B>B |
| chr5 | 4600000 | 4700000 | -0.87 | -0.75 | B>B |
| chr5 | 4700000 | 4800000 | -0.88 | -0.82 | B>B |
| chr5 | 4800000 | 4900000 | -0.42 | -0.31 | B>B |
| chr5 | 4900000 | 5000000 | 0.00 | 0.02 | A>A |
| chr5 | 5000000 | 5100000 | 0.13 | 0.18 | A>A |
| chr5 | 5100000 | 5200000 | 0.27 | 0.62 | A>A |
| chr5 | 5200000 | 5300000 | 0.58 | 1.06 | A>A |
| chr5 | 5300000 | 5400000 | 0.59 | 0.81 | A>A |
| chr5 | 5400000 | 5500000 | 0.21 | 0.21 | A>A |
| chr5 | 5500000 | 5600000 | -0.01 | -0.43 | B>B |
| chr5 | 5600000 | 5700000 | -0.12 | -0.62 | B>B |
| chr5 | 5700000 | 5800000 | -0.05 | -0.69 | A>B |
| chr5 | 5800000 | 5900000 | 0.09 | -0.47 | B>B |
| chr5 | 5900000 | 6000000 | 0.24 | -0.45 | A>B |
| chr5 | 6000000 | 6100000 | 0.45 | -0.28 | A>B |
| chr5 | 6100000 | 6200000 | 0.48 | -0.20 | A>B |
| chr5 | 6200000 | 6300000 | 0.31 | -0.27 | A>A |
| chr5 | 6300000 | 6400000 | 0.54 | -0.04 | A>A |
| chr5 | 6400000 | 6500000 | 0.58 | 0.21 | A>A |
| chr5 | 6500000 | 6600000 | 0.50 | 0.12 | A>A |
| chr5 | 6600000 | 6700000 | 0.84 | 0.69 | A>A |
| chr5 | 6700000 | 6800000 | 0.42 | 0.43 | A>A |
| chr5 | 6800000 | 6900000 | 0.07 | 0.16 | A>A |
| chr5 | 6900000 | 7000000 | -0.11 | 0.13 | A>A |

| | | | | | |
|---|---|---|---|---|---|
| chr5 | 7000000 | 7100000 | -0.40 | -0.29 | B>B |
| chr5 | 7100000 | 7200000 | -0.56 | -0.55 | B>B |
| chr5 | 7200000 | 7300000 | -0.38 | -0.64 | B>B |
| chr5 | 7300000 | 7400000 | -0.37 | -0.58 | B>B |
| chr5 | 7400000 | 7500000 | -0.38 | -0.68 | B>B |
| chr5 | 7500000 | 7600000 | -0.43 | -0.79 | B>B |
| chr5 | 7600000 | 7700000 | -0.49 | -0.73 | B>B |
| chr5 | 7700000 | 7800000 | -0.18 | -0.41 | B>B |
| chr5 | 7800000 | 7900000 | 0.16 | -0.01 | A>A |
| chr5 | 7900000 | 8000000 | -0.29 | -0.43 | B>B |
| chr5 | 8000000 | 8100000 | -0.46 | -0.59 | B>B |
| chr5 | 8100000 | 8200000 | -0.57 | -0.69 | B>B |
| chr5 | 8200000 | 8300000 | -0.68 | -0.73 | B>B |
| chr5 | 8300000 | 8400000 | -0.66 | -0.65 | B>B |
| chr5 | 8400000 | 8500000 | -0.62 | -0.60 | B>B |
| chr5 | 8500000 | 8600000 | -0.68 | -0.66 | B>B |
| chr5 | 8600000 | 8700000 | -0.76 | -0.66 | B>B |
| chr5 | 8700000 | 8800000 | -0.81 | -0.63 | B>B |
| chr5 | 8800000 | 8900000 | -0.67 | -0.67 | B>B |
| chr5 | 8900000 | 9000000 | -0.71 | -0.70 | B>B |
| chr5 | 9000000 | 9100000 | -0.84 | -0.62 | B>B |
| chr5 | 9100000 | 9200000 | -0.60 | -0.63 | B>B |
| chr5 | 9200000 | 9300000 | -0.61 | -0.57 | B>B |
| chr5 | 9300000 | 9400000 | -0.61 | -0.64 | B>B |
| chr5 | 9400000 | 9500000 | -0.53 | -0.72 | B>B |
| chr5 | 9500000 | 9600000 | -0.38 | -0.64 | B>B |
| chr5 | 9600000 | 9700000 | -0.51 | -0.68 | B>B |
| chr5 | 9700000 | 9800000 | -0.39 | -0.75 | B>B |
| chr5 | 9800000 | 9900000 | -0.28 | -0.70 | B>B |
| chr5 | 9900000 | 10000000 | -0.16 | -0.71 | B>B |
| chr5 | 10000000 | 10100000 | -0.16 | -0.63 | B>B |
| chr5 | 10100000 | 10200000 | 0.09 | -0.31 | B>B |
| chr5 | 10200000 | 10300000 | 0.56 | 0.44 | A>A |
| chr5 | 10300000 | 10400000 | 1.07 | 0.91 | A>A |
| chr5 | 10400000 | 10500000 | 0.93 | 0.74 | A>A |
| chr5 | 10500000 | 10600000 | 1.43 | 1.06 | A>A |
| chr5 | 10600000 | 10700000 | 0.63 | 0.35 | A>A |
| chr5 | 10700000 | 10800000 | 0.45 | 0.17 | A>A |
| chr5 | 10800000 | 10900000 | 0.23 | -0.07 | A>A |
| chr5 | 10900000 | 11000000 | 0.06 | -0.25 | B>B |
| chr5 | 11000000 | 11100000 | -0.08 | -0.49 | B>B |
| chr5 | 11100000 | 11200000 | -0.09 | -0.51 | B>B |
| chr5 | 11200000 | 11300000 | -0.25 | -0.56 | B>B |
| chr5 | 11300000 | 11400000 | -0.36 | -0.59 | B>B |
| chr5 | 11400000 | 11500000 | -0.34 | -0.62 | B>B |
| chr5 | 11500000 | 11600000 | -0.47 | -0.64 | B>B |
| chr5 | 11600000 | 11700000 | -0.70 | -0.62 | B>B |
| chr5 | 11700000 | 11800000 | -0.73 | -0.66 | B>B |
| chr5 | 11800000 | 11900000 | -0.74 | -0.67 | B>B |
| chr5 | 11900000 | 12000000 | -0.71 | -0.64 | B>B |
| chr5 | 12000000 | 12100000 | -0.73 | -0.59 | B>B |
| chr5 | 12100000 | 12200000 | -0.87 | -0.65 | B>B |
| chr5 | 12200000 | 12300000 | -0.84 | -0.66 | B>B |
| chr5 | 12300000 | 12400000 | -0.85 | -0.70 | B>B |
| chr5 | 12400000 | 12500000 | -0.77 | -0.74 | B>B |
| chr5 | 12500000 | 12600000 | -0.81 | -0.67 | B>B |

| | | | | | |
|---|---|---|---|---|---|
| chr5 | 12600000 | 12700000 | -0.84 | -0.59 | B>B |
| chr5 | 12700000 | 12800000 | -0.90 | -0.47 | B>B |
| chr5 | 12800000 | 12900000 | -0.83 | -0.44 | B>B |
| chr5 | 12900000 | 13000000 | -0.82 | -0.49 | B>B |
| chr5 | 13000000 | 13100000 | -0.85 | -0.60 | B>B |
| chr5 | 13100000 | 13200000 | -0.75 | -0.56 | B>B |
| chr5 | 13200000 | 13300000 | -0.68 | -0.44 | B>B |
| chr5 | 13300000 | 13400000 | -0.60 | -0.36 | B>B |
| chr5 | 13400000 | 13500000 | -0.35 | -0.14 | B>B |
| chr5 | 13500000 | 13600000 | -0.22 | 0.12 | B>B |
| chr5 | 13600000 | 13700000 | -0.08 | 0.09 | A>A |
| chr5 | 13700000 | 13800000 | -0.14 | -0.03 | B>B |
| chr5 | 13800000 | 13900000 | -0.09 | -0.01 | B>B |
| chr5 | 13900000 | 14000000 | 0.05 | 0.11 | A>A |
| chr5 | 14000000 | 14100000 | 1.01 | 0.98 | A>A |
| chr5 | 14100000 | 14200000 | 1.00 | 1.11 | A>A |
| chr5 | 14200000 | 14300000 | 0.88 | 1.10 | A>A |
| chr5 | 14300000 | 14400000 | 0.59 | 0.75 | A>A |
| chr5 | 14400000 | 14500000 | 0.82 | 0.80 | A>A |
| chr5 | 14500000 | 14600000 | 0.69 | 0.87 | A>A |
| chr5 | 14600000 | 14700000 | 0.47 | 0.45 | A>A |
| chr5 | 14700000 | 14800000 | 0.49 | 0.42 | A>A |
| chr5 | 14800000 | 14900000 | 0.76 | 0.59 | A>A |
| chr5 | 14900000 | 15000000 | 0.65 | 0.43 | A>A |
| chr5 | 15000000 | 15100000 | -0.04 | -0.31 | B>B |
| chr5 | 15100000 | 15200000 | -0.30 | -0.71 | B>B |
| chr5 | 15200000 | 15300000 | -0.54 | -0.78 | B>B |
| chr5 | 15300000 | 15400000 | -0.50 | -0.70 | B>B |
| chr5 | 15400000 | 15500000 | -0.38 | -0.63 | B>B |
| chr5 | 15500000 | 15600000 | -0.28 | -0.53 | B>B |
| chr5 | 15600000 | 15700000 | -0.32 | -0.53 | B>B |
| chr5 | 15700000 | 15800000 | -0.33 | -0.49 | B>B |
| chr5 | 15800000 | 15900000 | -0.34 | -0.52 | B>B |
| chr5 | 15900000 | 16000000 | -0.41 | -0.57 | B>B |
| chr5 | 16000000 | 16100000 | -0.40 | -0.67 | B>B |
| chr5 | 16100000 | 16200000 | -0.35 | -0.74 | B>B |
| chr5 | 16200000 | 16300000 | -0.40 | -0.80 | B>B |
| chr5 | 16300000 | 16400000 | -0.41 | -0.66 | B>B |
| chr5 | 16400000 | 16500000 | 0.10 | -0.08 | A>A |
| chr5 | 16500000 | 16600000 | 0.58 | 0.42 | A>A |
| chr5 | 16600000 | 16700000 | 0.70 | 0.63 | A>A |
| chr5 | 16700000 | 16800000 | 0.79 | 0.80 | A>A |
| chr5 | 16800000 | 16900000 | 1.04 | 1.12 | A>A |
| chr5 | 16900000 | 17000000 | 1.06 | 1.15 | A>A |
| chr5 | 17000000 | 17100000 | 1.25 | 1.11 | A>A |
| chr5 | 17100000 | 17200000 | 0.78 | 0.76 | A>A |
| chr5 | 17200000 | 17300000 | 0.62 | 0.81 | A>A |
| chr5 | 17300000 | 17400000 | 0.87 | 1.06 | A>A |
| chr5 | 17400000 | 17500000 | 1.00 | 1.28 | A>A |
| chr5 | 17500000 | 17600000 | 0.00 | 0.00 | NO |
| chr5 | 17600000 | 17700000 | 0.88 | 0.95 | A>A |
| chr5 | 17700000 | 17800000 | 0.85 | 1.03 | A>A |
| chr5 | 17800000 | 17900000 | 0.86 | 0.95 | A>A |
| chr5 | 17900000 | 18000000 | 0.77 | 0.86 | A>A |
| chr5 | 18000000 | 18100000 | 0.73 | 0.82 | A>A |
| chr5 | 18100000 | 18200000 | 0.53 | 0.51 | A>A |

| | | | | | |
|---|---|---|---|---|---|
| chr5 | 18200000 | 18300000 | 0.21 | 0.28 | A>A |
| chr5 | 18300000 | 18400000 | -0.14 | -0.28 | B>B |
| chr5 | 18400000 | 18500000 | -0.56 | -0.64 | B>B |
| chr5 | 18500000 | 18600000 | -0.61 | -0.71 | B>B |
| chr5 | 18600000 | 18700000 | -0.59 | -0.66 | B>B |
| chr5 | 18700000 | 18800000 | -0.46 | -0.48 | B>B |
| chr5 | 18800000 | 18900000 | -0.56 | -0.67 | B>B |
| chr5 | 18900000 | 19000000 | -0.61 | -0.69 | B>B |
| chr5 | 19000000 | 19100000 | -0.61 | -0.71 | B>B |
| chr5 | 19100000 | 19200000 | -0.60 | -0.66 | B>B |
| chr5 | 19200000 | 19300000 | -0.57 | -0.60 | B>B |
| chr5 | 19300000 | 19400000 | -0.60 | -0.49 | B>B |
| chr5 | 19400000 | 19500000 | -0.59 | -0.52 | B>B |
| chr5 | 19500000 | 19600000 | -0.56 | -0.55 | B>B |
| chr5 | 19600000 | 19700000 | -0.57 | -0.59 | B>B |
| chr5 | 19700000 | 19800000 | -0.54 | -0.67 | B>B |
| chr5 | 19800000 | 19900000 | -0.55 | -0.65 | B>B |
| chr5 | 19900000 | 20000000 | -0.53 | -0.60 | B>B |
| chr5 | 20000000 | 20100000 | -0.55 | -0.71 | B>B |
| chr5 | 20100000 | 20200000 | -0.59 | -0.68 | B>B |
| chr5 | 20200000 | 20300000 | -0.50 | -0.57 | B>B |
| chr5 | 20300000 | 20400000 | -0.47 | -0.62 | B>B |
| chr5 | 20400000 | 20500000 | -0.54 | -0.70 | B>B |
| chr5 | 20500000 | 20600000 | -0.57 | -0.64 | B>B |
| chr5 | 20600000 | 20700000 | -0.53 | -0.67 | B>B |
| chr5 | 20700000 | 20800000 | 0.00 | 0.00 | NO |
| chr5 | 20800000 | 20900000 | 0.00 | 0.00 | NO |
| chr5 | 20900000 | 21000000 | -0.65 | -0.19 | B>B |
| chr5 | 21000000 | 21100000 | -0.67 | 0.17 | B>A |
| chr5 | 21100000 | 21200000 | -0.63 | 0.28 | B>A |
| chr5 | 21200000 | 21300000 | -0.59 | 0.15 | B>A |
| chr5 | 21300000 | 21400000 | 0.00 | 0.00 | NO |
| chr5 | 21400000 | 21500000 | 0.00 | 0.00 | NO |
| chr5 | 21500000 | 21600000 | -0.52 | -0.35 | B>B |
| chr5 | 21600000 | 21700000 | -0.82 | -0.55 | B>B |
| chr5 | 21700000 | 21800000 | -0.75 | -0.51 | B>B |
| chr5 | 21800000 | 21900000 | -0.69 | -0.61 | B>B |
| chr5 | 21900000 | 22000000 | 0.00 | 0.00 | NO |
| chr5 | 22000000 | 22100000 | -0.77 | -0.70 | B>B |
| chr5 | 22100000 | 22200000 | 0.00 | 0.00 | NO |
| chr5 | 22200000 | 22300000 | -0.73 | -0.72 | B>B |
| chr5 | 22300000 | 22400000 | -0.80 | -0.83 | B>B |
| chr5 | 22400000 | 22500000 | -0.79 | -0.75 | B>B |
| chr5 | 22500000 | 22600000 | -0.78 | -0.82 | B>B |
| chr5 | 22600000 | 22700000 | -0.73 | -0.78 | B>B |
| chr5 | 22700000 | 22800000 | -0.75 | -0.74 | B>B |
| chr5 | 22800000 | 22900000 | -0.73 | -0.62 | B>B |
| chr5 | 22900000 | 23000000 | -0.77 | -0.49 | B>B |
| chr5 | 23000000 | 23100000 | -0.74 | -0.41 | B>B |
| chr5 | 23100000 | 23200000 | -0.69 | -0.40 | B>B |
| chr5 | 23200000 | 23300000 | -0.64 | -0.09 | B>A |
| chr5 | 23300000 | 23400000 | -0.68 | 0.35 | B>A |
| chr5 | 23400000 | 23500000 | -0.64 | 0.68 | B>A |
| chr5 | 23500000 | 23600000 | -0.65 | 0.50 | B>A |
| chr5 | 23600000 | 23700000 | -0.71 | 0.59 | B>A |
| chr5 | 23700000 | 23800000 | -0.70 | 0.27 | B>A |

| | | | | | |
|---|---|---|---|---|---|
| chr5 | 23800000 | 23900000 | -0.74 | -0.08 | B>A |
| chr5 | 23900000 | 24000000 | -0.67 | -0.13 | B>A |
| chr5 | 24000000 | 24100000 | -0.75 | -0.31 | B>B |
| chr5 | 24100000 | 24200000 | -0.76 | -0.35 | B>B |
| chr5 | 24200000 | 24300000 | -0.77 | -0.34 | B>B |
| chr5 | 24300000 | 24400000 | -0.76 | -0.28 | B>B |
| chr5 | 24400000 | 24500000 | -0.74 | -0.26 | B>B |
| chr5 | 24500000 | 24600000 | -0.73 | -0.27 | B>B |
| chr5 | 24600000 | 24700000 | -0.65 | -0.29 | B>B |
| chr5 | 24700000 | 24800000 | -0.65 | -0.29 | B>B |
| chr5 | 24800000 | 24900000 | -0.72 | -0.36 | B>B |
| chr5 | 24900000 | 25000000 | -0.71 | -0.34 | B>B |
| chr5 | 25000000 | 25100000 | -0.65 | -0.36 | B>B |
| chr5 | 25100000 | 25200000 | -0.71 | -0.28 | B>B |
| chr5 | 25200000 | 25300000 | -0.67 | -0.41 | B>B |
| chr5 | 25300000 | 25400000 | -0.70 | -0.35 | B>B |
| chr5 | 25400000 | 25500000 | -0.67 | -0.33 | B>B |
| chr5 | 25500000 | 25600000 | -0.69 | -0.32 | B>B |
| chr5 | 25600000 | 25700000 | -0.65 | -0.37 | B>B |
| chr5 | 25700000 | 25800000 | -0.66 | -0.42 | B>B |
| chr5 | 25800000 | 25900000 | -0.65 | -0.37 | B>B |
| chr5 | 25900000 | 26000000 | -0.66 | -0.42 | B>B |
| chr5 | 26000000 | 26100000 | -0.69 | -0.55 | B>B |
| chr5 | 26100000 | 26200000 | -0.74 | -0.60 | B>B |
| chr5 | 26200000 | 26300000 | -0.68 | -0.52 | B>B |
| chr5 | 26300000 | 26400000 | -0.67 | -0.53 | B>B |
| chr5 | 26400000 | 26500000 | -0.67 | -0.46 | B>B |
| chr5 | 26500000 | 26600000 | -0.72 | -0.46 | B>B |
| chr5 | 26600000 | 26700000 | -0.65 | -0.33 | B>B |
| chr5 | 26700000 | 26800000 | -0.65 | -0.26 | B>B |
| chr5 | 26800000 | 26900000 | -0.60 | -0.15 | B>B |
| chr5 | 26900000 | 27000000 | -0.56 | -0.19 | B>B |
| chr5 | 27000000 | 27100000 | -0.59 | -0.14 | B>B |
| chr5 | 27100000 | 27200000 | -0.60 | -0.04 | B>A |
| chr5 | 27200000 | 27300000 | -0.58 | 0.04 | B>A |
| chr5 | 27300000 | 27400000 | -0.50 | 0.05 | B>A |
| chr5 | 27400000 | 27500000 | -0.24 | 0.22 | B>B |
| chr5 | 27500000 | 27600000 | -0.40 | 0.04 | B>B |
| chr5 | 27600000 | 27700000 | -0.52 | 0.03 | B>A |
| chr5 | 27700000 | 27800000 | -0.51 | 0.07 | B>A |
| chr5 | 27800000 | 27900000 | -0.45 | 0.11 | B>A |
| chr5 | 27900000 | 28000000 | -0.48 | 0.01 | B>B |
| chr5 | 28000000 | 28100000 | -0.45 | -0.19 | B>B |
| chr5 | 28100000 | 28200000 | -0.45 | -0.29 | B>B |
| chr5 | 28200000 | 28300000 | -0.45 | -0.25 | B>B |
| chr5 | 28300000 | 28400000 | -0.45 | -0.21 | B>B |
| chr5 | 28400000 | 28500000 | -0.52 | -0.26 | B>B |
| chr5 | 28500000 | 28600000 | -0.49 | -0.29 | B>B |
| chr5 | 28600000 | 28700000 | -0.54 | -0.22 | B>B |
| chr5 | 28700000 | 28800000 | -0.62 | -0.31 | B>B |
| chr5 | 28800000 | 28900000 | -0.68 | -0.16 | B>A |
| chr5 | 28900000 | 29000000 | -0.63 | -0.18 | B>B |
| chr5 | 29000000 | 29100000 | -0.62 | -0.17 | B>B |
| chr5 | 29100000 | 29200000 | -0.57 | -0.20 | B>B |
| chr5 | 29200000 | 29300000 | -0.57 | -0.23 | B>B |
| chr5 | 29300000 | 29400000 | -0.63 | -0.15 | B>B |

| | | | | | |
|---|---|---|---|---|---|
| chr5 | 29400000 | 29500000 | -0.64 | -0.15 | B>B |
| chr5 | 29500000 | 29600000 | -0.64 | -0.05 | B>A |
| chr5 | 29600000 | 29700000 | -0.56 | 0.03 | B>A |
| chr5 | 29700000 | 29800000 | -0.57 | 0.10 | B>A |
| chr5 | 29800000 | 29900000 | -0.53 | 0.01 | B>A |
| chr5 | 29900000 | 30000000 | -0.58 | 0.09 | B>A |
| chr5 | 30000000 | 30100000 | -0.37 | 0.28 | B>A |
| chr5 | 30100000 | 30200000 | -0.33 | 0.38 | B>A |
| chr5 | 30200000 | 30300000 | -0.45 | 0.33 | B>A |
| chr5 | 30300000 | 30400000 | -0.60 | 0.20 | B>A |
| chr5 | 30400000 | 30500000 | -0.65 | 0.12 | B>A |
| chr5 | 30500000 | 30600000 | -0.73 | 0.22 | B>A |
| chr5 | 30600000 | 30700000 | -0.77 | 0.02 | B>A |
| chr5 | 30700000 | 30800000 | -0.74 | -0.08 | B>A |
| chr5 | 30800000 | 30900000 | -0.74 | -0.10 | B>A |
| chr5 | 30900000 | 31000000 | -0.77 | -0.09 | B>A |
| chr5 | 31000000 | 31100000 | -0.74 | -0.04 | B>A |
| chr5 | 31100000 | 31200000 | -0.74 | -0.05 | B>A |
| chr5 | 31200000 | 31300000 | -0.62 | -0.09 | B>A |
| chr5 | 31300000 | 31400000 | -0.45 | -0.12 | B>B |
| chr5 | 31400000 | 31500000 | 0.29 | 0.54 | A>A |
| chr5 | 31500000 | 31600000 | 0.49 | 0.69 | A>A |
| chr5 | 31600000 | 31700000 | 0.66 | 0.62 | A>A |
| chr5 | 31700000 | 31800000 | 0.66 | 0.60 | A>A |
| chr5 | 31800000 | 31900000 | 0.65 | 0.70 | A>A |
| chr5 | 31900000 | 32000000 | 0.93 | 0.90 | A>A |
| chr5 | 32000000 | 32100000 | 0.85 | 0.89 | A>A |
| chr5 | 32100000 | 32200000 | 0.99 | 1.11 | A>A |
| chr5 | 32200000 | 32300000 | 1.12 | 1.19 | A>A |
| chr5 | 32300000 | 32400000 | 1.02 | 1.02 | A>A |
| chr5 | 32400000 | 32500000 | 1.02 | 0.98 | A>A |
| chr5 | 32500000 | 32600000 | 1.01 | 0.93 | A>A |
| chr5 | 32600000 | 32700000 | 0.57 | 0.57 | A>A |
| chr5 | 32700000 | 32800000 | 0.77 | 0.41 | A>A |
| chr5 | 32800000 | 32900000 | 0.62 | 0.23 | A>A |
| chr5 | 32900000 | 33000000 | 0.47 | 0.11 | A>A |
| chr5 | 33000000 | 33100000 | 0.53 | 0.16 | A>A |
| chr5 | 33100000 | 33200000 | 0.44 | 0.26 | A>A |
| chr5 | 33200000 | 33300000 | 0.50 | 0.28 | A>A |
| chr5 | 33300000 | 33400000 | 0.48 | 0.33 | A>A |
| chr5 | 33400000 | 33500000 | 0.48 | 0.02 | A>A |
| chr5 | 33500000 | 33600000 | 0.32 | -0.59 | A>B |
| chr5 | 33600000 | 33700000 | 0.26 | -0.79 | A>B |
| chr5 | 33700000 | 33800000 | 0.29 | -0.76 | A>B |
| chr5 | 33800000 | 33900000 | 0.36 | -0.69 | A>B |
| chr5 | 33900000 | 34000000 | 0.30 | -0.42 | A>B |
| chr5 | 34000000 | 34100000 | 0.33 | -0.21 | A>A |
| chr5 | 34100000 | 34200000 | 0.00 | 0.00 | NO |
| chr5 | 34200000 | 34300000 | 0.00 | 0.00 | NO |
| chr5 | 34300000 | 34400000 | 0.00 | 0.00 | NO |
| chr5 | 34400000 | 34500000 | 0.28 | 0.17 | A>A |
| chr5 | 34500000 | 34600000 | 0.78 | 0.66 | A>A |
| chr5 | 34600000 | 34700000 | 0.85 | 0.89 | A>A |
| chr5 | 34700000 | 34800000 | 0.74 | 0.63 | A>A |
| chr5 | 34800000 | 34900000 | 0.60 | 0.45 | A>A |
| chr5 | 34900000 | 35000000 | 0.36 | -0.01 | A>A |

| | | | | | |
|---|---|---|---|---|---|
| chr5 | 35000000 | 35100000 | -0.22 | -0.64 | B>B |
| chr5 | 35100000 | 35200000 | -0.37 | -0.85 | B>B |
| chr5 | 35200000 | 35300000 | -0.40 | -0.74 | B>B |
| chr5 | 35300000 | 35400000 | -0.60 | -0.80 | B>B |
| chr5 | 35400000 | 35500000 | -0.63 | -0.81 | B>B |
| chr5 | 35500000 | 35600000 | -0.61 | -0.84 | B>B |
| chr5 | 35600000 | 35700000 | -0.28 | -0.54 | B>B |
| chr5 | 35700000 | 35800000 | -0.33 | -0.73 | B>B |
| chr5 | 35800000 | 35900000 | -0.50 | -0.79 | B>B |
| chr5 | 35900000 | 36000000 | -0.54 | -0.75 | B>B |
| chr5 | 36000000 | 36100000 | -0.41 | -0.63 | B>B |
| chr5 | 36100000 | 36200000 | 0.10 | -0.07 | A>A |
| chr5 | 36200000 | 36300000 | 0.08 | -0.38 | B>B |
| chr5 | 36300000 | 36400000 | -0.29 | -0.86 | B>B |
| chr5 | 36400000 | 36500000 | -0.24 | -0.84 | A>B |
| chr5 | 36500000 | 36600000 | -0.28 | -0.86 | B>B |
| chr5 | 36600000 | 36700000 | -0.10 | -0.69 | A>B |
| chr5 | 36700000 | 36800000 | 0.08 | -0.25 | B>B |
| chr5 | 36800000 | 36900000 | 0.31 | -0.07 | A>A |
| chr5 | 36900000 | 37000000 | 0.67 | 0.35 | A>A |
| chr5 | 37000000 | 37100000 | 0.53 | 0.58 | A>A |
| chr5 | 37100000 | 37200000 | 0.77 | 0.84 | A>A |
| chr5 | 37200000 | 37300000 | 1.09 | 1.10 | A>A |
| chr5 | 37300000 | 37400000 | 0.90 | 0.74 | A>A |
| chr5 | 37400000 | 37500000 | 0.54 | 0.41 | A>A |
| chr5 | 37500000 | 37600000 | 0.47 | 0.41 | A>A |
| chr5 | 37600000 | 37700000 | 0.45 | 0.30 | A>A |
| chr5 | 37700000 | 37800000 | 0.23 | 0.03 | A>A |
| chr5 | 37800000 | 37900000 | 0.05 | -0.39 | B>B |
| chr5 | 37900000 | 38000000 | -0.10 | -0.78 | A>B |
| chr5 | 38000000 | 38100000 | -0.17 | -0.82 | A>B |
| chr5 | 38100000 | 38200000 | -0.12 | -0.61 | B>B |
| chr5 | 38200000 | 38300000 | 0.07 | -0.48 | B>B |
| chr5 | 38300000 | 38400000 | 0.22 | -0.37 | A>B |
| chr5 | 38400000 | 38500000 | 0.47 | 0.19 | A>A |
| chr5 | 38500000 | 38600000 | 0.69 | 0.37 | A>A |
| chr5 | 38600000 | 38700000 | 0.75 | 0.41 | A>A |
| chr5 | 38700000 | 38800000 | 0.60 | 0.31 | A>A |
| chr5 | 38800000 | 38900000 | 0.60 | 0.50 | A>A |
| chr5 | 38900000 | 39000000 | 0.58 | 0.53 | A>A |
| chr5 | 39000000 | 39100000 | 1.04 | 0.97 | A>A |
| chr5 | 39100000 | 39200000 | 1.00 | 0.89 | A>A |
| chr5 | 39200000 | 39300000 | 0.76 | 0.92 | A>A |
| chr5 | 39300000 | 39400000 | 0.66 | 0.75 | A>A |
| chr5 | 39400000 | 39500000 | 0.92 | 0.92 | A>A |
| chr5 | 39500000 | 39600000 | 0.49 | 0.55 | A>A |
| chr5 | 39600000 | 39700000 | 0.52 | 0.59 | A>A |
| chr5 | 39700000 | 39800000 | 0.47 | 0.47 | A>A |
| chr5 | 39800000 | 39900000 | 0.36 | 0.61 | A>A |
| chr5 | 39900000 | 40000000 | 0.28 | 0.42 | A>A |
| chr5 | 40000000 | 40100000 | 0.26 | 0.48 | A>A |
| chr5 | 40100000 | 40200000 | 0.29 | 0.41 | A>A |
| chr5 | 40200000 | 40300000 | 0.31 | 0.44 | A>A |
| chr5 | 40300000 | 40400000 | 0.44 | 0.47 | A>A |
| chr5 | 40400000 | 40500000 | 0.51 | 0.51 | A>A |
| chr5 | 40500000 | 40600000 | 0.47 | 0.54 | A>A |

| | | | | | |
|---|---|---|---|---|---|
| chr5 | 40600000 | 40700000 | 0.39 | 0.48 | A>A |
| chr5 | 40700000 | 40800000 | 0.46 | 0.50 | A>A |
| chr5 | 40800000 | 40900000 | 0.43 | 0.40 | A>A |
| chr5 | 40900000 | 41000000 | -0.11 | -0.28 | B>B |
| chr5 | 41000000 | 41100000 | -0.19 | -0.44 | B>B |
| chr5 | 41100000 | 41200000 | -0.20 | -0.50 | B>B |
| chr5 | 41200000 | 41300000 | -0.08 | -0.31 | B>B |
| chr5 | 41300000 | 41400000 | 0.11 | 0.00 | A>A |
| chr5 | 41400000 | 41500000 | 0.23 | 0.10 | A>A |
| chr5 | 41500000 | 41600000 | -0.13 | -0.33 | B>B |
| chr5 | 41600000 | 41700000 | -0.38 | -0.49 | B>B |
| chr5 | 41700000 | 41800000 | -0.26 | -0.30 | B>B |
| chr5 | 41800000 | 41900000 | -0.12 | -0.26 | B>B |
| chr5 | 41900000 | 42000000 | -0.17 | -0.37 | B>B |
| chr5 | 42000000 | 42100000 | -0.50 | -0.73 | B>B |
| chr5 | 42100000 | 42200000 | -0.65 | -0.77 | B>B |
| chr5 | 42200000 | 42300000 | -0.71 | -0.84 | B>B |
| chr5 | 42300000 | 42400000 | -0.58 | -0.80 | B>B |
| chr5 | 42400000 | 42500000 | -0.49 | -0.79 | B>B |
| chr5 | 42500000 | 42600000 | -0.45 | -0.69 | B>B |
| chr5 | 42600000 | 42700000 | -0.39 | -0.69 | B>B |
| chr5 | 42700000 | 42800000 | -0.33 | -0.42 | B>B |
| chr5 | 42800000 | 42900000 | -0.05 | -0.17 | B>B |
| chr5 | 42900000 | 43000000 | 0.59 | 0.41 | A>A |
| chr5 | 43000000 | 43100000 | 0.71 | 0.61 | A>A |
| chr5 | 43100000 | 43200000 | 0.71 | 0.78 | A>A |
| chr5 | 43200000 | 43300000 | 0.68 | 0.85 | A>A |
| chr5 | 43300000 | 43400000 | 0.79 | 0.91 | A>A |
| chr5 | 43400000 | 43500000 | 0.67 | 0.84 | A>A |
| chr5 | 43500000 | 43600000 | 0.42 | 0.57 | A>A |
| chr5 | 43600000 | 43700000 | -0.06 | 0.11 | A>A |
| chr5 | 43700000 | 43800000 | -0.41 | -0.38 | B>B |
| chr5 | 43800000 | 43900000 | -0.63 | -0.57 | B>B |
| chr5 | 43900000 | 44000000 | -0.74 | -0.46 | B>B |
| chr5 | 44000000 | 44100000 | -0.71 | -0.36 | B>B |
| chr5 | 44100000 | 44200000 | -0.78 | -0.59 | B>B |
| chr5 | 44200000 | 44300000 | -0.83 | -0.66 | B>B |
| chr5 | 44300000 | 44400000 | -0.73 | -0.67 | B>B |
| chr5 | 44400000 | 44500000 | -0.86 | -0.76 | B>B |
| chr5 | 44500000 | 44600000 | -0.89 | -0.82 | B>B |
| chr5 | 44600000 | 44700000 | -0.83 | -0.74 | B>B |
| chr5 | 44700000 | 44800000 | -0.58 | -0.44 | B>B |
| chr5 | 44800000 | 44900000 | -0.46 | 0.23 | B>A |
| chr5 | 44900000 | 45000000 | -0.80 | 0.23 | B>A |
| chr5 | 45000000 | 45100000 | -0.91 | 0.12 | B>A |
| chr5 | 45100000 | 45200000 | -0.92 | 0.24 | B>A |
| chr5 | 45200000 | 45300000 | -0.95 | 0.28 | B>A |
| chr5 | 45300000 | 45400000 | -1.00 | 0.26 | B>A |
| chr5 | 45400000 | 45500000 | -0.97 | 0.22 | B>A |
| chr5 | 45500000 | 45600000 | -1.03 | 0.12 | B>A |
| chr5 | 45600000 | 45700000 | -0.99 | 0.14 | B>A |
| chr5 | 45700000 | 45800000 | -0.97 | 0.23 | B>A |
| chr5 | 45800000 | 45900000 | -1.06 | 0.35 | B>A |
| chr5 | 45900000 | 46000000 | 0.00 | 0.00 | NO |
| chr5 | 46000000 | 46100000 | 0.00 | 0.00 | NO |
| chr5 | 46100000 | 46200000 | 0.00 | 0.00 | NO |

| | | | | | |
|---|---|---|---|---|---|
| chr5 | 46200000 | 46300000 | 0.00 | 0.00 | NO |
| chr5 | 46300000 | 46400000 | 0.00 | 0.00 | NO |
| chr5 | 46400000 | 46500000 | 0.00 | 0.00 | NO |
| chr5 | 46500000 | 46600000 | 0.00 | 0.00 | NO |
| chr5 | 46600000 | 46700000 | 0.00 | 0.00 | NO |
| chr5 | 46700000 | 46800000 | 0.00 | 0.00 | NO |
| chr5 | 46800000 | 46900000 | 0.00 | 0.00 | NO |
| chr5 | 46900000 | 47000000 | 0.00 | 0.00 | NO |
| chr5 | 47000000 | 47100000 | 0.00 | 0.00 | NO |
| chr5 | 47100000 | 47200000 | 0.00 | 0.00 | NO |
| chr5 | 47200000 | 47300000 | 0.00 | 0.00 | NO |
| chr5 | 47300000 | 47400000 | 0.00 | 0.00 | NO |
| chr5 | 47400000 | 47500000 | 0.00 | 0.00 | NO |
| chr5 | 47500000 | 47600000 | 0.00 | 0.00 | NO |
| chr5 | 47600000 | 47700000 | 0.00 | 0.00 | NO |
| chr5 | 47700000 | 47800000 | 0.00 | 0.00 | NO |
| chr5 | 47800000 | 47900000 | 0.00 | 0.00 | NO |
| chr5 | 47900000 | 48000000 | 0.00 | 0.00 | NO |
| chr5 | 48000000 | 48100000 | 0.00 | 0.00 | NO |
| chr5 | 48100000 | 48200000 | 0.00 | 0.00 | NO |
| chr5 | 48200000 | 48300000 | 0.00 | 0.00 | NO |
| chr5 | 48300000 | 48400000 | 0.00 | 0.00 | NO |
| chr5 | 48400000 | 48500000 | 0.00 | 0.00 | NO |
| chr5 | 48500000 | 48600000 | 0.00 | 0.00 | NO |
| chr5 | 48600000 | 48700000 | 0.00 | 0.00 | NO |
| chr5 | 48700000 | 48800000 | 0.00 | 0.00 | NO |
| chr5 | 48800000 | 48900000 | 0.00 | 0.00 | NO |
| chr5 | 48900000 | 49000000 | 0.00 | 0.00 | NO |
| chr5 | 49000000 | 49100000 | 0.00 | 0.00 | NO |
| chr5 | 49100000 | 49200000 | 0.00 | 0.00 | NO |
| chr5 | 49200000 | 49300000 | 0.00 | 0.00 | NO |
| chr5 | 49300000 | 49400000 | 0.00 | 0.00 | NO |
| chr5 | 49400000 | 49500000 | 0.00 | 0.00 | NO |
| chr5 | 49500000 | 49600000 | 0.00 | 0.00 | NO |
| chr5 | 49600000 | 49700000 | 0.00 | 0.00 | NO |
| chr5 | 49700000 | 49800000 | 0.00 | 0.00 | NO |
| chr5 | 49800000 | 49900000 | 0.00 | 0.00 | NO |
| chr5 | 49900000 | 50000000 | 0.00 | 0.00 | NO |
| chr5 | 50000000 | 50100000 | -0.89 | 0.11 | B>A |
| chr5 | 50100000 | 50200000 | -1.04 | -0.36 | B>A |
| chr5 | 50200000 | 50300000 | -0.71 | -0.64 | B>B |
| chr5 | 50300000 | 50400000 | -0.95 | -0.93 | B>B |
| chr5 | 50400000 | 50500000 | -0.97 | -1.00 | B>B |
| chr5 | 50500000 | 50600000 | -0.94 | -0.96 | B>B |
| chr5 | 50600000 | 50700000 | -0.80 | -0.82 | B>B |
| chr5 | 50700000 | 50800000 | -0.88 | -0.89 | B>B |
| chr5 | 50800000 | 50900000 | -0.97 | -0.91 | B>B |
| chr5 | 50900000 | 51000000 | -0.95 | -0.90 | B>B |
| chr5 | 51000000 | 51100000 | -0.95 | -0.77 | B>B |
| chr5 | 51100000 | 51200000 | -0.97 | -0.75 | B>B |
| chr5 | 51200000 | 51300000 | -0.96 | -0.84 | B>B |
| chr5 | 51300000 | 51400000 | -0.94 | -0.95 | B>B |
| chr5 | 51400000 | 51500000 | -0.98 | -1.01 | B>B |
| chr5 | 51500000 | 51600000 | -0.95 | -0.99 | B>B |
| chr5 | 51600000 | 51700000 | -0.89 | -1.04 | B>B |
| chr5 | 51700000 | 51800000 | -0.85 | -1.05 | B>B |

| | | | | | |
|---|---|---|---|---|---|
| chr5 | 51800000 | 51900000 | -0.69 | -0.97 | B>B |
| chr5 | 51900000 | 52000000 | -0.47 | -0.80 | B>B |
| chr5 | 52000000 | 52100000 | -0.05 | -0.56 | B>B |
| chr5 | 52100000 | 52200000 | 0.22 | -0.20 | A>A |
| chr5 | 52200000 | 52300000 | 0.49 | 0.22 | A>A |
| chr5 | 52300000 | 52400000 | 0.69 | 0.43 | A>A |
| chr5 | 52400000 | 52500000 | 0.40 | 0.33 | A>A |
| chr5 | 52500000 | 52600000 | 0.56 | 0.34 | A>A |
| chr5 | 52600000 | 52700000 | 0.64 | 0.54 | A>A |
| chr5 | 52700000 | 52800000 | 0.53 | 0.40 | A>A |
| chr5 | 52800000 | 52900000 | 0.70 | 0.51 | A>A |
| chr5 | 52900000 | 53000000 | 0.71 | 0.52 | A>A |
| chr5 | 53000000 | 53100000 | 0.67 | 0.46 | A>A |
| chr5 | 53100000 | 53200000 | 0.65 | 0.47 | A>A |
| chr5 | 53200000 | 53300000 | 0.73 | 0.62 | A>A |
| chr5 | 53300000 | 53400000 | 0.70 | 0.66 | A>A |
| chr5 | 53400000 | 53500000 | 0.58 | 0.45 | A>A |
| chr5 | 53500000 | 53600000 | 0.29 | 0.15 | A>A |
| chr5 | 53600000 | 53700000 | 0.31 | 0.02 | A>A |
| chr5 | 53700000 | 53800000 | 0.34 | 0.08 | A>A |
| chr5 | 53800000 | 53900000 | 0.20 | -0.19 | A>A |
| chr5 | 53900000 | 54000000 | 0.15 | -0.46 | A>B |
| chr5 | 54000000 | 54100000 | 0.09 | -0.42 | B>B |
| chr5 | 54100000 | 54200000 | 0.16 | -0.20 | B>B |
| chr5 | 54200000 | 54300000 | 0.16 | -0.32 | B>B |
| chr5 | 54300000 | 54400000 | 0.12 | -0.39 | B>B |
| chr5 | 54400000 | 54500000 | 0.35 | 0.14 | A>A |
| chr5 | 54500000 | 54600000 | 1.02 | 0.85 | A>A |
| chr5 | 54600000 | 54700000 | 0.99 | 0.85 | A>A |
| chr5 | 54700000 | 54800000 | 1.06 | 1.17 | A>A |
| chr5 | 54800000 | 54900000 | 1.14 | 1.15 | A>A |
| chr5 | 54900000 | 55000000 | 1.01 | 0.93 | A>A |
| chr5 | 55000000 | 55100000 | 0.97 | 1.02 | A>A |
| chr5 | 55100000 | 55200000 | 1.10 | 1.08 | A>A |
| chr5 | 55200000 | 55300000 | 0.88 | 0.95 | A>A |
| chr5 | 55300000 | 55400000 | 1.27 | 1.28 | A>A |
| chr5 | 55400000 | 55500000 | 1.14 | 1.17 | A>A |
| chr5 | 55500000 | 55600000 | 0.70 | 0.82 | A>A |
| chr5 | 55600000 | 55700000 | 0.85 | 1.12 | A>A |
| chr5 | 55700000 | 55800000 | 0.97 | 1.13 | A>A |
| chr5 | 55800000 | 55900000 | 1.00 | 1.19 | A>A |
| chr5 | 55900000 | 56000000 | 0.89 | 1.26 | A>A |
| chr5 | 56000000 | 56100000 | 0.94 | 1.13 | A>A |
| chr5 | 56100000 | 56200000 | 0.63 | 0.89 | A>A |
| chr5 | 56200000 | 56300000 | 1.05 | 1.13 | A>A |
| chr5 | 56300000 | 56400000 | 1.15 | 1.13 | A>A |
| chr5 | 56400000 | 56500000 | 0.96 | 1.11 | A>A |
| chr5 | 56500000 | 56600000 | 0.72 | 0.68 | A>A |
| chr5 | 56600000 | 56700000 | 0.91 | 0.92 | A>A |
| chr5 | 56700000 | 56800000 | 0.83 | 0.84 | A>A |
| chr5 | 56800000 | 56900000 | 0.83 | 0.81 | A>A |
| chr5 | 56900000 | 57000000 | 0.96 | 0.86 | A>A |
| chr5 | 57000000 | 57100000 | 0.76 | 0.74 | A>A |
| chr5 | 57100000 | 57200000 | 0.79 | 0.71 | A>A |
| chr5 | 57200000 | 57300000 | 1.09 | 0.92 | A>A |
| chr5 | 57300000 | 57400000 | 1.11 | 1.07 | A>A |

| | | | | | |
|---|---|---|---|---|---|
| chr5 | 57400000 | 57500000 | 0.90 | 1.06 | A>A |
| chr5 | 57500000 | 57600000 | 1.12 | 1.00 | A>A |
| chr5 | 57600000 | 57700000 | 1.15 | 1.00 | A>A |
| chr5 | 57700000 | 57800000 | 0.61 | 0.60 | A>A |
| chr5 | 57800000 | 57900000 | 0.70 | 0.73 | A>A |
| chr5 | 57900000 | 58000000 | 0.90 | 0.73 | A>A |
| chr5 | 58000000 | 58100000 | 0.81 | 0.69 | A>A |
| chr5 | 58100000 | 58200000 | 0.66 | 0.64 | A>A |
| chr5 | 58200000 | 58300000 | 1.22 | 1.06 | A>A |
| chr5 | 58300000 | 58400000 | 1.02 | 1.05 | A>A |
| chr5 | 58400000 | 58500000 | 0.93 | 0.86 | A>A |
| chr5 | 58500000 | 58600000 | 0.83 | 0.83 | A>A |
| chr5 | 58600000 | 58700000 | 0.88 | 0.79 | A>A |
| chr5 | 58700000 | 58800000 | 0.96 | 0.85 | A>A |
| chr5 | 58800000 | 58900000 | 1.07 | 0.92 | A>A |
| chr5 | 58900000 | 59000000 | 0.87 | 0.81 | A>A |
| chr5 | 59000000 | 59100000 | 0.79 | 0.62 | A>A |
| chr5 | 59100000 | 59200000 | 0.38 | -0.02 | A>A |
| chr5 | 59200000 | 59300000 | 0.36 | -0.20 | A>A |
| chr5 | 59300000 | 59400000 | 0.25 | -0.18 | A>A |
| chr5 | 59400000 | 59500000 | 0.32 | -0.19 | A>A |
| chr5 | 59500000 | 59600000 | 0.36 | -0.19 | A>A |
| chr5 | 59600000 | 59700000 | 0.36 | -0.22 | A>A |
| chr5 | 59700000 | 59800000 | 0.57 | -0.18 | A>B |
| chr5 | 59800000 | 59900000 | 0.38 | -0.12 | A>A |
| chr5 | 59900000 | 60000000 | 0.51 | 0.22 | A>A |
| chr5 | 60000000 | 60100000 | 0.92 | 0.49 | A>A |
| chr5 | 60100000 | 60200000 | 0.77 | 0.42 | A>A |
| chr5 | 60200000 | 60300000 | 0.55 | 0.28 | A>A |
| chr5 | 60300000 | 60400000 | 0.56 | 0.39 | A>A |
| chr5 | 60400000 | 60500000 | 0.62 | 0.41 | A>A |
| chr5 | 60500000 | 60600000 | 0.92 | 0.58 | A>A |
| chr5 | 60600000 | 60700000 | 0.92 | 0.57 | A>A |
| chr5 | 60700000 | 60800000 | 0.92 | 0.70 | A>A |
| chr5 | 60800000 | 60900000 | 0.94 | 0.72 | A>A |
| chr5 | 60900000 | 61000000 | 0.75 | 0.52 | A>A |
| chr5 | 61000000 | 61100000 | 0.66 | 0.18 | A>A |
| chr5 | 61100000 | 61200000 | 0.36 | -0.01 | A>A |
| chr5 | 61200000 | 61300000 | 0.22 | -0.19 | A>A |
| chr5 | 61300000 | 61400000 | 0.32 | -0.24 | A>A |
| chr5 | 61400000 | 61500000 | 0.35 | -0.13 | A>A |
| chr5 | 61500000 | 61600000 | 0.72 | 0.34 | A>A |
| chr5 | 61600000 | 61700000 | 0.73 | 0.36 | A>A |
| chr5 | 61700000 | 61800000 | 0.38 | 0.17 | A>A |
| chr5 | 61800000 | 61900000 | 0.46 | 0.19 | A>A |
| chr5 | 61900000 | 62000000 | 0.09 | -0.02 | A>A |
| chr5 | 62000000 | 62100000 | -0.09 | -0.25 | B>B |
| chr5 | 62100000 | 62200000 | -0.59 | -0.73 | B>B |
| chr5 | 62200000 | 62300000 | -0.78 | -0.72 | B>B |
| chr5 | 62300000 | 62400000 | -0.73 | -0.67 | B>B |
| chr5 | 62400000 | 62500000 | -0.78 | -0.72 | B>B |
| chr5 | 62500000 | 62600000 | -0.85 | -0.76 | B>B |
| chr5 | 62600000 | 62700000 | -0.95 | -0.78 | B>B |
| chr5 | 62700000 | 62800000 | -0.97 | -0.90 | B>B |
| chr5 | 62800000 | 62900000 | -1.08 | -0.95 | B>B |
| chr5 | 62900000 | 63000000 | -1.08 | -1.05 | B>B |

| | | | | | |
|---|---|---|---|---|---|
| chr5 | 63000000 | 63100000 | -1.07 | -0.96 | B>B |
| chr5 | 63100000 | 63200000 | -1.06 | -1.03 | B>B |
| chr5 | 63200000 | 63300000 | -1.00 | -0.93 | B>B |
| chr5 | 63300000 | 63400000 | -0.99 | -0.97 | B>B |
| chr5 | 63400000 | 63500000 | -0.65 | -0.74 | B>B |
| chr5 | 63500000 | 63600000 | -0.77 | -0.77 | B>B |
| chr5 | 63600000 | 63700000 | -0.91 | -1.01 | B>B |
| chr5 | 63700000 | 63800000 | -0.83 | -1.07 | B>B |
| chr5 | 63800000 | 63900000 | -0.56 | -0.86 | B>B |
| chr5 | 63900000 | 64000000 | -0.15 | -0.40 | B>B |
| chr5 | 64000000 | 64100000 | 0.06 | -0.09 | B>B |
| chr5 | 64100000 | 64200000 | 0.10 | 0.04 | A>A |
| chr5 | 64200000 | 64300000 | 0.09 | 0.12 | A>A |
| chr5 | 64300000 | 64400000 | 0.17 | -0.01 | A>A |
| chr5 | 64400000 | 64500000 | 0.01 | -0.15 | B>B |
| chr5 | 64500000 | 64600000 | -0.15 | -0.19 | B>B |
| chr5 | 64600000 | 64700000 | -0.21 | -0.29 | B>B |
| chr5 | 64700000 | 64800000 | -0.06 | -0.19 | B>B |
| chr5 | 64800000 | 64900000 | 0.55 | 0.38 | A>A |
| chr5 | 64900000 | 65000000 | 1.01 | 0.71 | A>A |
| chr5 | 65000000 | 65100000 | 0.92 | 0.82 | A>A |
| chr5 | 65100000 | 65200000 | 0.92 | 0.91 | A>A |
| chr5 | 65200000 | 65300000 | 1.04 | 0.96 | A>A |
| chr5 | 65300000 | 65400000 | 0.87 | 0.80 | A>A |
| chr5 | 65400000 | 65500000 | 0.58 | 0.34 | A>A |
| chr5 | 65500000 | 65600000 | 0.22 | -0.17 | A>A |
| chr5 | 65600000 | 65700000 | -0.16 | -0.68 | B>B |
| chr5 | 65700000 | 65800000 | -0.19 | -0.87 | A>B |
| chr5 | 65800000 | 65900000 | -0.24 | -0.77 | B>B |
| chr5 | 65900000 | 66000000 | -0.13 | -0.74 | A>B |
| chr5 | 66000000 | 66100000 | -0.02 | -0.37 | B>B |
| chr5 | 66100000 | 66200000 | 0.44 | 0.17 | A>A |
| chr5 | 66200000 | 66300000 | 0.62 | 0.39 | A>A |
| chr5 | 66300000 | 66400000 | 0.56 | 0.41 | A>A |
| chr5 | 66400000 | 66500000 | 0.29 | 0.17 | A>A |
| chr5 | 66500000 | 66600000 | -0.06 | -0.40 | B>B |
| chr5 | 66600000 | 66700000 | -0.06 | -0.79 | A>B |
| chr5 | 66700000 | 66800000 | -0.22 | -0.90 | A>B |
| chr5 | 66800000 | 66900000 | -0.38 | -0.92 | B>B |
| chr5 | 66900000 | 67000000 | -0.52 | -0.94 | B>B |
| chr5 | 67000000 | 67100000 | -0.51 | -0.93 | B>B |
| chr5 | 67100000 | 67200000 | -0.56 | -0.84 | B>B |
| chr5 | 67200000 | 67300000 | -0.39 | -0.88 | B>B |
| chr5 | 67300000 | 67400000 | -0.24 | -0.79 | B>B |
| chr5 | 67400000 | 67500000 | -0.03 | -0.51 | B>B |
| chr5 | 67500000 | 67600000 | 0.12 | -0.06 | A>A |
| chr5 | 67600000 | 67700000 | 0.43 | 0.48 | A>A |
| chr5 | 67700000 | 67800000 | 0.40 | 0.47 | A>A |
| chr5 | 67800000 | 67900000 | 0.34 | 0.45 | A>A |
| chr5 | 67900000 | 68000000 | 0.31 | 0.46 | A>A |
| chr5 | 68000000 | 68100000 | 0.31 | 0.40 | A>A |
| chr5 | 68100000 | 68200000 | 0.21 | 0.32 | A>A |
| chr5 | 68200000 | 68300000 | 0.34 | 0.28 | A>A |
| chr5 | 68300000 | 68400000 | 0.74 | 0.66 | A>A |
| chr5 | 68400000 | 68500000 | 1.32 | 1.17 | A>A |
| chr5 | 68500000 | 68600000 | 1.26 | 1.39 | A>A |

| | | | | | |
|---|---|---|---|---|---|
| chr5 | 68600000 | 68700000 | 1.20 | 1.37 | A>A |
| chr5 | 68700000 | 68800000 | 1.36 | 1.46 | A>A |
| chr5 | 68800000 | 68900000 | 0.00 | 0.00 | NO |
| chr5 | 68900000 | 69000000 | 0.00 | 0.00 | NO |
| chr5 | 69000000 | 69100000 | 0.00 | 0.00 | NO |
| chr5 | 69100000 | 69200000 | 0.00 | 0.00 | NO |
| chr5 | 69200000 | 69300000 | 0.00 | 0.00 | NO |
| chr5 | 69300000 | 69400000 | 0.00 | 0.00 | NO |
| chr5 | 69400000 | 69500000 | 0.00 | 0.00 | NO |
| chr5 | 69500000 | 69600000 | 0.00 | 0.00 | NO |
| chr5 | 69600000 | 69700000 | 0.00 | 0.00 | NO |
| chr5 | 69700000 | 69800000 | 0.00 | 0.00 | NO |
| chr5 | 69800000 | 69900000 | 0.00 | 0.00 | NO |
| chr5 | 69900000 | 70000000 | 0.00 | 0.00 | NO |
| chr5 | 70000000 | 70100000 | 0.00 | 0.00 | NO |
| chr5 | 70100000 | 70200000 | 0.00 | 0.00 | NO |
| chr5 | 70200000 | 70300000 | 0.00 | 0.00 | NO |
| chr5 | 70300000 | 70400000 | 0.00 | 0.00 | NO |
| chr5 | 70400000 | 70500000 | 0.00 | 0.00 | NO |
| chr5 | 70500000 | 70600000 | 0.00 | 0.00 | NO |
| chr5 | 70600000 | 70700000 | 0.00 | 0.00 | NO |
| chr5 | 70700000 | 70800000 | 0.00 | 0.00 | NO |
| chr5 | 70800000 | 70900000 | 1.44 | 1.36 | A>A |
| chr5 | 70900000 | 71000000 | 1.20 | 1.40 | A>A |
| chr5 | 71000000 | 71100000 | 0.87 | 0.98 | A>A |
| chr5 | 71100000 | 71200000 | 1.07 | 1.27 | A>A |
| chr5 | 71200000 | 71300000 | 1.16 | 1.27 | A>A |
| chr5 | 71300000 | 71400000 | 1.12 | 1.15 | A>A |
| chr5 | 71400000 | 71500000 | 1.17 | 1.23 | A>A |
| chr5 | 71500000 | 71600000 | 1.03 | 1.15 | A>A |
| chr5 | 71600000 | 71700000 | 0.96 | 1.14 | A>A |
| chr5 | 71700000 | 71800000 | 1.32 | 1.24 | A>A |
| chr5 | 71800000 | 71900000 | 1.02 | 1.06 | A>A |
| chr5 | 71900000 | 72000000 | 1.30 | 1.28 | A>A |
| chr5 | 72000000 | 72100000 | 1.28 | 1.22 | A>A |
| chr5 | 72100000 | 72200000 | 1.21 | 1.23 | A>A |
| chr5 | 72200000 | 72300000 | 1.11 | 1.18 | A>A |
| chr5 | 72300000 | 72400000 | 1.20 | 1.25 | A>A |
| chr5 | 72400000 | 72500000 | 1.17 | 1.23 | A>A |
| chr5 | 72500000 | 72600000 | 1.17 | 1.20 | A>A |
| chr5 | 72600000 | 72700000 | 1.20 | 1.25 | A>A |
| chr5 | 72700000 | 72800000 | 1.12 | 1.14 | A>A |
| chr5 | 72800000 | 72900000 | 0.94 | 1.11 | A>A |
| chr5 | 72900000 | 73000000 | 1.30 | 1.46 | A>A |
| chr5 | 73000000 | 73100000 | 1.29 | 1.34 | A>A |
| chr5 | 73100000 | 73200000 | 1.06 | 1.10 | A>A |
| chr5 | 73200000 | 73300000 | 0.83 | 0.88 | A>A |
| chr5 | 73300000 | 73400000 | 0.75 | 1.05 | A>A |
| chr5 | 73400000 | 73500000 | 0.88 | 1.00 | A>A |
| chr5 | 73500000 | 73600000 | 0.91 | 0.99 | A>A |
| chr5 | 73600000 | 73700000 | 1.00 | 1.02 | A>A |
| chr5 | 73700000 | 73800000 | 0.98 | 1.07 | A>A |
| chr5 | 73800000 | 73900000 | 1.00 | 1.06 | A>A |
| chr5 | 73900000 | 74000000 | 0.55 | 0.80 | A>A |
| chr5 | 74000000 | 74100000 | 0.86 | 0.89 | A>A |
| chr5 | 74100000 | 74200000 | 0.94 | 1.02 | A>A |

| | | | | | |
|---|---|---|---|---|---|
| chr5 | 74200000 | 74300000 | 0.61 | 0.61 | A>A |
| chr5 | 74300000 | 74400000 | 0.73 | 0.57 | A>A |
| chr5 | 74400000 | 74500000 | 0.48 | 0.44 | A>A |
| chr5 | 74500000 | 74600000 | 0.51 | 0.54 | A>A |
| chr5 | 74600000 | 74700000 | 0.76 | 0.70 | A>A |
| chr5 | 74700000 | 74800000 | 0.77 | 0.64 | A>A |
| chr5 | 74800000 | 74900000 | 0.88 | 0.63 | A>A |
| chr5 | 74900000 | 75000000 | 0.57 | 0.49 | A>A |
| chr5 | 75000000 | 75100000 | 0.29 | -0.23 | A>A |
| chr5 | 75100000 | 75200000 | -0.06 | -0.57 | B>B |
| chr5 | 75200000 | 75300000 | -0.22 | -0.64 | B>B |
| chr5 | 75300000 | 75400000 | -0.06 | -0.56 | B>B |
| chr5 | 75400000 | 75500000 | 0.08 | -0.47 | B>B |
| chr5 | 75500000 | 75600000 | 0.23 | -0.30 | B>B |
| chr5 | 75600000 | 75700000 | 0.55 | -0.12 | A>B |
| chr5 | 75700000 | 75800000 | 0.66 | -0.16 | A>B |
| chr5 | 75800000 | 75900000 | 0.55 | -0.11 | A>B |
| chr5 | 75900000 | 76000000 | 0.77 | 0.24 | A>A |
| chr5 | 76000000 | 76100000 | 0.96 | 0.46 | A>A |
| chr5 | 76100000 | 76200000 | 0.87 | 0.55 | A>A |
| chr5 | 76200000 | 76300000 | 0.98 | 0.42 | A>A |
| chr5 | 76300000 | 76400000 | 0.85 | 0.59 | A>A |
| chr5 | 76400000 | 76500000 | 1.00 | 0.87 | A>A |
| chr5 | 76500000 | 76600000 | 1.02 | 0.90 | A>A |
| chr5 | 76600000 | 76700000 | 0.99 | 0.84 | A>A |
| chr5 | 76700000 | 76800000 | 0.85 | 0.76 | A>A |
| chr5 | 76800000 | 76900000 | 0.82 | 0.62 | A>A |
| chr5 | 76900000 | 77000000 | 0.91 | 0.76 | A>A |
| chr5 | 77000000 | 77100000 | 1.08 | 0.89 | A>A |
| chr5 | 77100000 | 77200000 | 0.94 | 0.87 | A>A |
| chr5 | 77200000 | 77300000 | 0.84 | 0.80 | A>A |
| chr5 | 77300000 | 77400000 | 0.83 | 0.83 | A>A |
| chr5 | 77400000 | 77500000 | 1.03 | 0.88 | A>A |
| chr5 | 77500000 | 77600000 | 0.97 | 0.86 | A>A |
| chr5 | 77600000 | 77700000 | 1.00 | 0.99 | A>A |
| chr5 | 77700000 | 77800000 | 0.94 | 0.95 | A>A |
| chr5 | 77800000 | 77900000 | 1.07 | 1.01 | A>A |
| chr5 | 77900000 | 78000000 | 1.15 | 1.09 | A>A |
| chr5 | 78000000 | 78100000 | 1.13 | 1.15 | A>A |
| chr5 | 78100000 | 78200000 | 0.95 | 0.84 | A>A |
| chr5 | 78200000 | 78300000 | 0.79 | 0.83 | A>A |
| chr5 | 78300000 | 78400000 | 0.90 | 0.69 | A>A |
| chr5 | 78400000 | 78500000 | 0.80 | 0.72 | A>A |
| chr5 | 78500000 | 78600000 | 0.95 | 0.79 | A>A |
| chr5 | 78600000 | 78700000 | 1.00 | 0.86 | A>A |
| chr5 | 78700000 | 78800000 | 0.89 | 0.84 | A>A |
| chr5 | 78800000 | 78900000 | 0.83 | 0.72 | A>A |
| chr5 | 78900000 | 79000000 | 1.03 | 0.88 | A>A |
| chr5 | 79000000 | 79100000 | 0.78 | 0.46 | A>A |
| chr5 | 79100000 | 79200000 | 0.73 | 0.50 | A>A |
| chr5 | 79200000 | 79300000 | 0.79 | 0.53 | A>A |
| chr5 | 79300000 | 79400000 | 0.67 | 0.56 | A>A |
| chr5 | 79400000 | 79500000 | 1.14 | 1.03 | A>A |
| chr5 | 79500000 | 79600000 | 1.01 | 0.97 | A>A |
| chr5 | 79600000 | 79700000 | 0.78 | 0.69 | A>A |
| chr5 | 79700000 | 79800000 | 1.02 | 0.95 | A>A |

| | | | | | |
|---|---|---|---|---|---|
| chr5 | 79800000 | 79900000 | 1.04 | 0.92 | A>A |
| chr5 | 79900000 | 80000000 | 0.87 | 0.88 | A>A |
| chr5 | 80000000 | 80100000 | 0.82 | 0.88 | A>A |
| chr5 | 80100000 | 80200000 | 0.87 | 0.87 | A>A |
| chr5 | 80200000 | 80300000 | 0.92 | 0.87 | A>A |
| chr5 | 80300000 | 80400000 | 0.86 | 0.80 | A>A |
| chr5 | 80400000 | 80500000 | 0.68 | 0.68 | A>A |
| chr5 | 80500000 | 80600000 | 0.76 | 0.89 | A>A |
| chr5 | 80600000 | 80700000 | 0.67 | 0.85 | A>A |
| chr5 | 80700000 | 80800000 | 0.56 | 0.84 | A>A |
| chr5 | 80800000 | 80900000 | 0.54 | 0.84 | A>A |
| chr5 | 80900000 | 81000000 | 0.62 | 0.95 | A>A |
| chr5 | 81000000 | 81100000 | 0.63 | 0.90 | A>A |
| chr5 | 81100000 | 81200000 | 0.60 | 0.89 | A>A |
| chr5 | 81200000 | 81300000 | 0.71 | 0.96 | A>A |
| chr5 | 81300000 | 81400000 | 0.67 | 0.91 | A>A |
| chr5 | 81400000 | 81500000 | 0.56 | 0.80 | A>A |
| chr5 | 81500000 | 81600000 | 0.40 | 0.63 | A>A |
| chr5 | 81600000 | 81700000 | 0.07 | 0.39 | A>A |
| chr5 | 81700000 | 81800000 | 0.04 | 0.60 | B>A |
| chr5 | 81800000 | 81900000 | -0.07 | 0.57 | B>A |
| chr5 | 81900000 | 82000000 | 0.01 | 0.50 | A>A |
| chr5 | 82000000 | 82100000 | -0.04 | 0.61 | B>A |
| chr5 | 82100000 | 82200000 | -0.04 | 0.58 | B>A |
| chr5 | 82200000 | 82300000 | 0.19 | 0.70 | A>A |
| chr5 | 82300000 | 82400000 | 0.36 | 0.62 | A>A |
| chr5 | 82400000 | 82500000 | 0.43 | 0.73 | A>A |
| chr5 | 82500000 | 82600000 | 0.27 | 0.61 | A>A |
| chr5 | 82600000 | 82700000 | 0.02 | 0.58 | B>A |
| chr5 | 82700000 | 82800000 | -0.06 | 0.52 | B>A |
| chr5 | 82800000 | 82900000 | -0.06 | 0.49 | B>A |
| chr5 | 82900000 | 83000000 | -0.23 | 0.17 | B>B |
| chr5 | 83000000 | 83100000 | -0.42 | 0.00 | B>B |
| chr5 | 83100000 | 83200000 | -0.55 | 0.10 | B>A |
| chr5 | 83200000 | 83300000 | -0.56 | 0.02 | B>A |
| chr5 | 83300000 | 83400000 | -0.51 | 0.13 | B>A |
| chr5 | 83400000 | 83500000 | -0.38 | 0.21 | B>A |
| chr5 | 83500000 | 83600000 | -0.20 | 0.19 | B>B |
| chr5 | 83600000 | 83700000 | -0.25 | 0.21 | B>B |
| chr5 | 83700000 | 83800000 | -0.47 | 0.00 | B>B |
| chr5 | 83800000 | 83900000 | -0.75 | -0.21 | B>A |
| chr5 | 83900000 | 84000000 | -0.82 | -0.33 | B>B |
| chr5 | 84000000 | 84100000 | -0.83 | -0.48 | B>B |
| chr5 | 84100000 | 84200000 | -0.84 | -0.55 | B>B |
| chr5 | 84200000 | 84300000 | -0.85 | -0.58 | B>B |
| chr5 | 84300000 | 84400000 | -0.85 | -0.57 | B>B |
| chr5 | 84400000 | 84500000 | -0.87 | -0.56 | B>B |
| chr5 | 84500000 | 84600000 | -0.90 | -0.49 | B>B |
| chr5 | 84600000 | 84700000 | -0.86 | -0.48 | B>B |
| chr5 | 84700000 | 84800000 | -0.85 | -0.42 | B>B |
| chr5 | 84800000 | 84900000 | -0.86 | -0.45 | B>B |
| chr5 | 84900000 | 85000000 | -0.87 | -0.46 | B>B |
| chr5 | 85000000 | 85100000 | -0.79 | -0.48 | B>B |
| chr5 | 85100000 | 85200000 | -0.86 | -0.51 | B>B |
| chr5 | 85200000 | 85300000 | -0.82 | -0.47 | B>B |
| chr5 | 85300000 | 85400000 | -0.80 | -0.45 | B>B |

| | | | | | |
|---|---|---|---|---|---|
| chr5 | 85400000 | 85500000 | -0.61 | -0.22 | B>B |
| chr5 | 85500000 | 85600000 | -0.59 | -0.18 | B>B |
| chr5 | 85600000 | 85700000 | -0.51 | -0.27 | B>B |
| chr5 | 85700000 | 85800000 | -0.55 | -0.35 | B>B |
| chr5 | 85800000 | 85900000 | -0.41 | -0.40 | B>B |
| chr5 | 85900000 | 86000000 | -0.36 | -0.46 | B>B |
| chr5 | 86000000 | 86100000 | -0.68 | -0.73 | B>B |
| chr5 | 86100000 | 86200000 | -0.69 | -0.76 | B>B |
| chr5 | 86200000 | 86300000 | -0.49 | -0.46 | B>B |
| chr5 | 86300000 | 86400000 | -0.34 | -0.33 | B>B |
| chr5 | 86400000 | 86500000 | -0.12 | -0.36 | B>B |
| chr5 | 86500000 | 86600000 | 0.23 | -0.07 | A>A |
| chr5 | 86600000 | 86700000 | 0.26 | 0.11 | A>A |
| chr5 | 86700000 | 86800000 | -0.39 | -0.63 | B>B |
| chr5 | 86800000 | 86900000 | -0.62 | -0.84 | B>B |
| chr5 | 86900000 | 87000000 | -0.63 | -0.86 | B>B |
| chr5 | 87000000 | 87100000 | -0.43 | -0.59 | B>B |
| chr5 | 87100000 | 87200000 | -0.63 | -0.72 | B>B |
| chr5 | 87200000 | 87300000 | -0.54 | -0.84 | B>B |
| chr5 | 87300000 | 87400000 | -0.43 | -0.77 | B>B |
| chr5 | 87400000 | 87500000 | 0.02 | -0.37 | B>B |
| chr5 | 87500000 | 87600000 | 0.17 | -0.10 | A>A |
| chr5 | 87600000 | 87700000 | -0.24 | -0.73 | B>B |
| chr5 | 87700000 | 87800000 | -0.28 | -0.81 | B>B |
| chr5 | 87800000 | 87900000 | -0.31 | -0.76 | B>B |
| chr5 | 87900000 | 88000000 | -0.19 | -0.43 | B>B |
| chr5 | 88000000 | 88100000 | -0.21 | -0.21 | B>B |
| chr5 | 88100000 | 88200000 | -0.21 | -0.26 | B>B |
| chr5 | 88200000 | 88300000 | -0.46 | -0.60 | B>B |
| chr5 | 88300000 | 88400000 | -0.56 | -0.69 | B>B |
| chr5 | 88400000 | 88500000 | -0.70 | -0.64 | B>B |
| chr5 | 88500000 | 88600000 | -0.79 | -0.62 | B>B |
| chr5 | 88600000 | 88700000 | -0.88 | -0.55 | B>B |
| chr5 | 88700000 | 88800000 | -0.85 | -0.50 | B>B |
| chr5 | 88800000 | 88900000 | -0.86 | -0.52 | B>B |
| chr5 | 88900000 | 89000000 | -0.88 | -0.58 | B>B |
| chr5 | 89000000 | 89100000 | -0.86 | -0.64 | B>B |
| chr5 | 89100000 | 89200000 | -0.88 | -0.63 | B>B |
| chr5 | 89200000 | 89300000 | -0.88 | -0.48 | B>B |
| chr5 | 89300000 | 89400000 | -0.82 | -0.31 | B>B |
| chr5 | 89400000 | 89500000 | -0.74 | -0.48 | B>B |
| chr5 | 89500000 | 89600000 | -0.53 | -0.45 | B>B |
| chr5 | 89600000 | 89700000 | -0.30 | -0.45 | B>B |
| chr5 | 89700000 | 89800000 | 0.21 | 0.03 | A>A |
| chr5 | 89800000 | 89900000 | 0.01 | 0.01 | A>A |
| chr5 | 89900000 | 90000000 | -0.27 | -0.31 | B>B |
| chr5 | 90000000 | 90100000 | -0.23 | -0.23 | B>B |
| chr5 | 90100000 | 90200000 | 0.24 | 0.41 | A>A |
| chr5 | 90200000 | 90300000 | 0.73 | 0.84 | A>A |
| chr5 | 90300000 | 90400000 | 0.63 | 0.89 | A>A |
| chr5 | 90400000 | 90500000 | 0.69 | 0.88 | A>A |
| chr5 | 90500000 | 90600000 | 0.82 | 0.84 | A>A |
| chr5 | 90600000 | 90700000 | 0.86 | 0.74 | A>A |
| chr5 | 90700000 | 90800000 | 0.04 | 0.24 | A>A |
| chr5 | 90800000 | 90900000 | -0.18 | 0.04 | B>B |
| chr5 | 90900000 | 91000000 | -0.14 | -0.02 | B>B |

| | | | | | |
|---|---|---|---|---|---|
| chr5 | 91000000 | 91100000 | -0.06 | -0.01 | B>B |
| chr5 | 91100000 | 91200000 | 0.05 | 0.12 | A>A |
| chr5 | 91200000 | 91300000 | 0.02 | 0.17 | A>A |
| chr5 | 91300000 | 91400000 | -0.02 | 0.20 | A>A |
| chr5 | 91400000 | 91500000 | -0.01 | 0.08 | A>A |
| chr5 | 91500000 | 91600000 | 0.01 | 0.01 | A>A |
| chr5 | 91600000 | 91700000 | 0.00 | 0.00 | NO |
| chr5 | 91700000 | 91800000 | 0.39 | 0.42 | A>A |
| chr5 | 91800000 | 91900000 | 0.46 | 0.42 | A>A |
| chr5 | 91900000 | 92000000 | 0.58 | 0.51 | A>A |
| chr5 | 92000000 | 92100000 | 0.47 | 0.42 | A>A |
| chr5 | 92100000 | 92200000 | 0.50 | 0.48 | A>A |
| chr5 | 92200000 | 92300000 | 0.44 | 0.47 | A>A |
| chr5 | 92300000 | 92400000 | 0.58 | 0.68 | A>A |
| chr5 | 92400000 | 92500000 | 0.49 | 0.57 | A>A |
| chr5 | 92500000 | 92600000 | 0.52 | 0.67 | A>A |
| chr5 | 92600000 | 92700000 | 0.45 | 0.63 | A>A |
| chr5 | 92700000 | 92800000 | 0.38 | 0.45 | A>A |
| chr5 | 92800000 | 92900000 | 0.28 | 0.45 | A>A |
| chr5 | 92900000 | 93000000 | 0.21 | 0.26 | A>A |
| chr5 | 93000000 | 93100000 | 0.22 | 0.32 | A>A |
| chr5 | 93100000 | 93200000 | 0.06 | 0.18 | A>A |
| chr5 | 93200000 | 93300000 | 0.23 | 0.21 | A>A |
| chr5 | 93300000 | 93400000 | 0.30 | 0.21 | A>A |
| chr5 | 93400000 | 93500000 | 0.05 | -0.07 | B>B |
| chr5 | 93500000 | 93600000 | -0.19 | -0.44 | B>B |
| chr5 | 93600000 | 93700000 | -0.28 | -0.35 | B>B |
| chr5 | 93700000 | 93800000 | -0.26 | -0.37 | B>B |
| chr5 | 93800000 | 93900000 | -0.12 | -0.26 | B>B |
| chr5 | 93900000 | 94000000 | 0.15 | -0.05 | A>A |
| chr5 | 94000000 | 94100000 | 0.03 | -0.09 | B>B |
| chr5 | 94100000 | 94200000 | -0.15 | -0.35 | B>B |
| chr5 | 94200000 | 94300000 | -0.03 | -0.33 | B>B |
| chr5 | 94300000 | 94400000 | 0.01 | -0.31 | B>B |
| chr5 | 94400000 | 94500000 | 0.00 | -0.55 | B>B |
| chr5 | 94500000 | 94600000 | -0.24 | -0.66 | B>B |
| chr5 | 94600000 | 94700000 | -0.09 | -0.62 | B>B |
| chr5 | 94700000 | 94800000 | 0.05 | -0.50 | B>B |
| chr5 | 94800000 | 94900000 | 0.54 | 0.16 | A>A |
| chr5 | 94900000 | 95000000 | 0.74 | 0.62 | A>A |
| chr5 | 95000000 | 95100000 | 0.97 | 0.91 | A>A |
| chr5 | 95100000 | 95200000 | 0.87 | 0.80 | A>A |
| chr5 | 95200000 | 95300000 | 0.82 | 0.89 | A>A |
| chr5 | 95300000 | 95400000 | 1.09 | 1.13 | A>A |
| chr5 | 95400000 | 95500000 | 1.10 | 1.13 | A>A |
| chr5 | 95500000 | 95600000 | 1.04 | 1.09 | A>A |
| chr5 | 95600000 | 95700000 | 0.74 | 0.84 | A>A |
| chr5 | 95700000 | 95800000 | 0.68 | 0.90 | A>A |
| chr5 | 95800000 | 95900000 | 0.83 | 0.97 | A>A |
| chr5 | 95900000 | 96000000 | 0.75 | 0.84 | A>A |
| chr5 | 96000000 | 96100000 | 0.65 | 0.85 | A>A |
| chr5 | 96100000 | 96200000 | 0.59 | 0.57 | A>A |
| chr5 | 96200000 | 96300000 | 0.72 | 0.51 | A>A |
| chr5 | 96300000 | 96400000 | 0.57 | 0.39 | A>A |
| chr5 | 96400000 | 96500000 | 0.36 | 0.23 | A>A |
| chr5 | 96500000 | 96600000 | -0.25 | -0.31 | B>B |

| | | | | | |
|---|---|---|---|---|---|
| chr5 | 96600000 | 96700000 | -0.70 | -0.83 | B>B |
| chr5 | 96700000 | 96800000 | -0.82 | -0.88 | B>B |
| chr5 | 96800000 | 96900000 | -0.84 | -0.89 | B>B |
| chr5 | 96900000 | 97000000 | -0.85 | -0.90 | B>B |
| chr5 | 97000000 | 97100000 | -0.86 | -0.92 | B>B |
| chr5 | 97100000 | 97200000 | -0.86 | -0.89 | B>B |
| chr5 | 97200000 | 97300000 | -0.85 | -0.95 | B>B |
| chr5 | 97300000 | 97400000 | -0.87 | -0.84 | B>B |
| chr5 | 97400000 | 97500000 | -0.76 | -0.88 | B>B |
| chr5 | 97500000 | 97600000 | -0.84 | -0.83 | B>B |
| chr5 | 97600000 | 97700000 | -0.75 | -0.77 | B>B |
| chr5 | 97700000 | 97800000 | -0.72 | -0.79 | B>B |
| chr5 | 97800000 | 97900000 | -0.71 | -0.91 | B>B |
| chr5 | 97900000 | 98000000 | -0.74 | -0.81 | B>B |
| chr5 | 98000000 | 98100000 | -0.61 | -0.65 | B>B |
| chr5 | 98100000 | 98200000 | -0.06 | -0.12 | B>B |
| chr5 | 98200000 | 98300000 | 0.46 | 0.29 | A>A |
| chr5 | 98300000 | 98400000 | 0.30 | 0.31 | A>A |
| chr5 | 98400000 | 98500000 | -0.40 | -0.55 | B>B |
| chr5 | 98500000 | 98600000 | -0.75 | -0.81 | B>B |
| chr5 | 98600000 | 98700000 | -0.85 | -0.91 | B>B |
| chr5 | 98700000 | 98800000 | -0.87 | -0.85 | B>B |
| chr5 | 98800000 | 98900000 | 0.00 | 0.00 | NO |
| chr5 | 98900000 | 99000000 | -0.87 | -0.87 | B>B |
| chr5 | 99000000 | 99100000 | -0.87 | -0.83 | B>B |
| chr5 | 99100000 | 99200000 | -0.92 | -0.85 | B>B |
| chr5 | 99200000 | 99300000 | -0.90 | -0.86 | B>B |
| chr5 | 99300000 | 99400000 | -0.81 | -0.81 | B>B |
| chr5 | 99400000 | 99500000 | 0.00 | -0.82 | NO |
| chr5 | 99500000 | 99600000 | -0.78 | -0.80 | B>B |
| chr5 | 99600000 | 99700000 | 0.00 | -0.79 | NO |
| chr5 | 99700000 | 99800000 | -0.40 | -0.56 | B>B |
| chr5 | 99800000 | 99900000 | -0.28 | -0.32 | B>B |
| chr5 | 99900000 | 100000000 | -0.08 | -0.14 | B>B |
| chr5 | 1E+08 | 100100000 | -0.08 | -0.02 | B>B |
| chr5 | 1E+08 | 100200000 | 0.19 | 0.21 | A>A |
| chr5 | 1E+08 | 100300000 | 0.16 | 0.22 | A>A |
| chr5 | 1E+08 | 100400000 | 0.21 | 0.26 | A>A |
| chr5 | 1E+08 | 100500000 | 0.23 | 0.45 | A>A |
| chr5 | 1.01E+08 | 100600000 | 0.20 | 0.46 | A>A |
| chr5 | 1.01E+08 | 100700000 | 0.11 | 0.19 | A>A |
| chr5 | 1.01E+08 | 100800000 | -0.36 | -0.48 | B>B |
| chr5 | 1.01E+08 | 100900000 | -0.64 | -0.73 | B>B |
| chr5 | 1.01E+08 | 101000000 | -0.67 | -0.71 | B>B |
| chr5 | 1.01E+08 | 101100000 | -0.74 | -0.78 | B>B |
| chr5 | 1.01E+08 | 101200000 | -0.85 | -0.87 | B>B |
| chr5 | 1.01E+08 | 101300000 | -0.85 | -0.88 | B>B |
| chr5 | 1.01E+08 | 101400000 | -0.86 | -0.84 | B>B |
| chr5 | 1.01E+08 | 101500000 | -0.88 | -0.86 | B>B |
| chr5 | 1.02E+08 | 101600000 | -0.83 | -0.81 | B>B |
| chr5 | 1.02E+08 | 101700000 | -0.87 | -0.89 | B>B |
| chr5 | 1.02E+08 | 101800000 | -0.95 | -0.96 | B>B |
| chr5 | 1.02E+08 | 101900000 | -0.80 | -0.83 | B>B |
| chr5 | 1.02E+08 | 102000000 | -0.60 | -0.79 | B>B |
| chr5 | 1.02E+08 | 102100000 | -0.21 | -0.33 | B>B |
| chr5 | 1.02E+08 | 102200000 | 0.09 | 0.08 | A>A |

| | | | | | |
|---|---|---|---|---|---|
| chr5 | 1.02E+08 | 102300000 | 0.09 | 0.14 | A>A |
| chr5 | 1.02E+08 | 102400000 | -0.07 | -0.11 | B>B |
| chr5 | 1.02E+08 | 102500000 | 0.17 | 0.00 | A>A |
| chr5 | 1.03E+08 | 102600000 | 0.14 | -0.25 | B>B |
| chr5 | 1.03E+08 | 102700000 | 0.11 | -0.68 | A>B |
| chr5 | 1.03E+08 | 102800000 | -0.25 | -0.87 | A>B |
| chr5 | 1.03E+08 | 102900000 | -0.42 | -0.74 | B>B |
| chr5 | 1.03E+08 | 103000000 | -0.65 | -0.81 | B>B |
| chr5 | 1.03E+08 | 103100000 | -0.80 | -0.92 | B>B |
| chr5 | 1.03E+08 | 103200000 | -0.79 | -0.85 | B>B |
| chr5 | 1.03E+08 | 103300000 | -0.73 | -0.89 | B>B |
| chr5 | 1.03E+08 | 103400000 | -0.76 | -0.89 | B>B |
| chr5 | 1.03E+08 | 103500000 | -0.59 | -0.68 | B>B |
| chr5 | 1.04E+08 | 103600000 | -0.65 | -0.73 | B>B |
| chr5 | 1.04E+08 | 103700000 | -0.67 | -0.72 | B>B |
| chr5 | 1.04E+08 | 103800000 | -0.85 | -0.88 | B>B |
| chr5 | 1.04E+08 | 103900000 | -0.87 | -0.85 | B>B |
| chr5 | 1.04E+08 | 104000000 | -0.82 | -0.83 | B>B |
| chr5 | 1.04E+08 | 104100000 | -0.88 | -0.83 | B>B |
| chr5 | 1.04E+08 | 104200000 | -0.84 | -0.86 | B>B |
| chr5 | 1.04E+08 | 104300000 | -0.85 | -0.82 | B>B |
| chr5 | 1.04E+08 | 104400000 | -0.83 | -0.86 | B>B |
| chr5 | 1.04E+08 | 104500000 | -0.80 | -0.82 | B>B |
| chr5 | 1.05E+08 | 104600000 | -0.83 | -0.83 | B>B |
| chr5 | 1.05E+08 | 104700000 | -0.81 | -0.85 | B>B |
| chr5 | 1.05E+08 | 104800000 | -0.85 | -0.88 | B>B |
| chr5 | 1.05E+08 | 104900000 | -0.84 | -0.83 | B>B |
| chr5 | 1.05E+08 | 105000000 | -0.75 | -0.80 | B>B |
| chr5 | 1.05E+08 | 105100000 | -0.78 | -0.82 | B>B |
| chr5 | 1.05E+08 | 105200000 | -0.82 | -0.82 | B>B |
| chr5 | 1.05E+08 | 105300000 | -0.80 | -0.81 | B>B |
| chr5 | 1.05E+08 | 105400000 | -0.79 | -0.76 | B>B |
| chr5 | 1.05E+08 | 105500000 | -0.50 | -0.52 | B>B |
| chr5 | 1.06E+08 | 105600000 | -0.35 | -0.25 | B>B |
| chr5 | 1.06E+08 | 105700000 | -0.39 | -0.14 | B>B |
| chr5 | 1.06E+08 | 105800000 | -0.33 | -0.08 | B>B |
| chr5 | 1.06E+08 | 105900000 | -0.34 | -0.13 | B>B |
| chr5 | 1.06E+08 | 106000000 | -0.34 | -0.06 | B>B |
| chr5 | 1.06E+08 | 106100000 | -0.24 | -0.14 | B>B |
| chr5 | 1.06E+08 | 106200000 | 0.07 | -0.03 | A>A |
| chr5 | 1.06E+08 | 106300000 | 0.18 | 0.14 | A>A |
| chr5 | 1.06E+08 | 106400000 | 0.31 | 0.34 | A>A |
| chr5 | 1.06E+08 | 106500000 | 0.33 | 0.36 | A>A |
| chr5 | 1.07E+08 | 106600000 | 0.50 | 0.51 | A>A |
| chr5 | 1.07E+08 | 106700000 | 0.74 | 0.59 | A>A |
| chr5 | 1.07E+08 | 106800000 | 0.72 | 0.74 | A>A |
| chr5 | 1.07E+08 | 106900000 | 0.74 | 0.85 | A>A |
| chr5 | 1.07E+08 | 107000000 | 0.63 | 0.66 | A>A |
| chr5 | 1.07E+08 | 107100000 | 0.17 | 0.28 | A>A |
| chr5 | 1.07E+08 | 107200000 | 0.16 | 0.25 | A>A |
| chr5 | 1.07E+08 | 107300000 | 0.21 | 0.40 | A>A |
| chr5 | 1.07E+08 | 107400000 | 0.16 | 0.37 | A>A |
| chr5 | 1.07E+08 | 107500000 | 0.12 | 0.27 | A>A |
| chr5 | 1.08E+08 | 107600000 | 0.15 | 0.23 | A>A |
| chr5 | 1.08E+08 | 107700000 | 0.20 | 0.18 | A>A |
| chr5 | 1.08E+08 | 107800000 | -0.10 | -0.35 | B>B |

| | | | | | |
|---|---|---|---|---|---|
| chr5 | 1.08E+08 | 107900000 | -0.30 | -0.65 | B>B |
| chr5 | 1.08E+08 | 108000000 | -0.10 | -0.47 | B>B |
| chr5 | 1.08E+08 | 108100000 | 0.33 | 0.13 | A>A |
| chr5 | 1.08E+08 | 108200000 | 0.56 | 0.34 | A>A |
| chr5 | 1.08E+08 | 108300000 | 0.34 | 0.13 | A>A |
| chr5 | 1.08E+08 | 108400000 | 0.22 | 0.03 | A>A |
| chr5 | 1.08E+08 | 108500000 | 0.07 | -0.17 | B>B |
| chr5 | 1.09E+08 | 108600000 | -0.02 | -0.31 | B>B |
| chr5 | 1.09E+08 | 108700000 | 0.19 | -0.17 | A>A |
| chr5 | 1.09E+08 | 108800000 | 0.38 | 0.10 | A>A |
| chr5 | 1.09E+08 | 108900000 | -0.22 | -0.58 | B>B |
| chr5 | 1.09E+08 | 109000000 | -0.33 | -0.72 | B>B |
| chr5 | 1.09E+08 | 109100000 | 0.07 | -0.19 | B>B |
| chr5 | 1.09E+08 | 109200000 | 0.24 | -0.13 | A>A |
| chr5 | 1.09E+08 | 109300000 | -0.26 | -0.78 | B>B |
| chr5 | 1.09E+08 | 109400000 | -0.60 | -0.91 | B>B |
| chr5 | 1.09E+08 | 109500000 | -0.83 | -0.92 | B>B |
| chr5 | 1.1E+08 | 109600000 | -0.95 | -0.95 | B>B |
| chr5 | 1.1E+08 | 109700000 | -0.91 | -0.95 | B>B |
| chr5 | 1.1E+08 | 109800000 | -0.90 | -0.99 | B>B |
| chr5 | 1.1E+08 | 109900000 | -0.96 | -0.94 | B>B |
| chr5 | 1.1E+08 | 110000000 | -0.76 | -0.89 | B>B |
| chr5 | 1.1E+08 | 110100000 | -0.52 | -0.71 | B>B |
| chr5 | 1.1E+08 | 110200000 | -0.74 | -0.92 | B>B |
| chr5 | 1.1E+08 | 110300000 | -0.87 | -0.99 | B>B |
| chr5 | 1.1E+08 | 110400000 | -0.61 | -0.98 | B>B |
| chr5 | 1.1E+08 | 110500000 | -0.20 | -0.46 | B>B |
| chr5 | 1.11E+08 | 110600000 | -0.34 | -0.77 | B>B |
| chr5 | 1.11E+08 | 110700000 | -0.63 | -0.95 | B>B |
| chr5 | 1.11E+08 | 110800000 | -0.52 | -0.84 | B>B |
| chr5 | 1.11E+08 | 110900000 | -0.13 | -0.34 | B>B |
| chr5 | 1.11E+08 | 111000000 | -0.17 | -0.38 | B>B |
| chr5 | 1.11E+08 | 111100000 | -0.04 | -0.10 | B>B |
| chr5 | 1.11E+08 | 111200000 | -0.36 | -0.43 | B>B |
| chr5 | 1.11E+08 | 111300000 | -0.68 | -0.81 | B>B |
| chr5 | 1.11E+08 | 111400000 | -0.71 | -0.87 | B>B |
| chr5 | 1.11E+08 | 111500000 | -0.52 | -0.53 | B>B |
| chr5 | 1.12E+08 | 111600000 | 0.30 | 0.48 | A>A |
| chr5 | 1.12E+08 | 111700000 | 0.34 | 0.73 | A>A |
| chr5 | 1.12E+08 | 111800000 | 0.10 | 0.54 | A>A |
| chr5 | 1.12E+08 | 111900000 | 0.03 | 0.08 | A>A |
| chr5 | 1.12E+08 | 112000000 | 0.30 | -0.13 | A>A |
| chr5 | 1.12E+08 | 112100000 | 0.62 | 0.27 | A>A |
| chr5 | 1.12E+08 | 112200000 | 0.69 | 0.42 | A>A |
| chr5 | 1.12E+08 | 112300000 | 0.86 | 0.58 | A>A |
| chr5 | 1.12E+08 | 112400000 | 0.57 | -0.06 | A>B |
| chr5 | 1.12E+08 | 112500000 | 0.15 | -0.71 | A>B |
| chr5 | 1.13E+08 | 112600000 | 0.06 | -0.91 | A>B |
| chr5 | 1.13E+08 | 112700000 | 0.06 | -0.73 | A>B |
| chr5 | 1.13E+08 | 112800000 | 0.11 | -0.73 | A>B |
| chr5 | 1.13E+08 | 112900000 | 0.33 | -0.18 | A>A |
| chr5 | 1.13E+08 | 113000000 | -0.08 | -0.36 | B>B |
| chr5 | 1.13E+08 | 113100000 | -0.54 | -0.79 | B>B |
| chr5 | 1.13E+08 | 113200000 | -0.77 | -0.93 | B>B |
| chr5 | 1.13E+08 | 113300000 | -0.82 | -0.94 | B>B |
| chr5 | 1.13E+08 | 113400000 | -0.81 | -0.78 | B>B |

| | | | | | |
|---|---|---|---|---|---|
| chr5 | 1.13E+08 | 113500000 | -0.77 | -0.48 | B>B |
| chr5 | 1.14E+08 | 113600000 | -0.75 | -0.36 | B>B |
| chr5 | 1.14E+08 | 113700000 | -0.63 | -0.61 | B>B |
| chr5 | 1.14E+08 | 113800000 | -0.59 | -0.77 | B>B |
| chr5 | 1.14E+08 | 113900000 | -0.84 | -0.90 | B>B |
| chr5 | 1.14E+08 | 114000000 | -1.00 | -0.98 | B>B |
| chr5 | 1.14E+08 | 114100000 | -0.95 | -0.99 | B>B |
| chr5 | 1.14E+08 | 114200000 | -0.83 | -0.95 | B>B |
| chr5 | 1.14E+08 | 114300000 | -0.86 | -0.96 | B>B |
| chr5 | 1.14E+08 | 114400000 | -0.60 | -0.78 | B>B |
| chr5 | 1.14E+08 | 114500000 | -0.05 | -0.20 | B>B |
| chr5 | 1.15E+08 | 114600000 | 0.29 | 0.11 | A>A |
| chr5 | 1.15E+08 | 114700000 | 0.48 | 0.49 | A>A |
| chr5 | 1.15E+08 | 114800000 | 0.76 | 0.98 | A>A |
| chr5 | 1.15E+08 | 114900000 | 0.81 | 0.81 | A>A |
| chr5 | 1.15E+08 | 115000000 | 0.31 | 0.46 | A>A |
| chr5 | 1.15E+08 | 115100000 | -0.04 | 0.52 | B>A |
| chr5 | 1.15E+08 | 115200000 | 0.23 | 0.60 | A>A |
| chr5 | 1.15E+08 | 115300000 | 0.30 | 0.82 | B>A |
| chr5 | 1.15E+08 | 115400000 | 0.14 | 0.76 | B>A |
| chr5 | 1.15E+08 | 115500000 | 0.04 | 0.18 | A>A |
| chr5 | 1.16E+08 | 115600000 | -0.01 | 0.04 | A>A |
| chr5 | 1.16E+08 | 115700000 | -0.24 | 0.06 | B>B |
| chr5 | 1.16E+08 | 115800000 | -0.32 | -0.04 | B>B |
| chr5 | 1.16E+08 | 115900000 | -0.34 | -0.12 | B>B |
| chr5 | 1.16E+08 | 116000000 | -0.37 | -0.56 | B>B |
| chr5 | 1.16E+08 | 116100000 | -0.48 | -0.84 | B>B |
| chr5 | 1.16E+08 | 116200000 | -0.56 | -0.85 | B>B |
| chr5 | 1.16E+08 | 116300000 | -0.71 | -0.85 | B>B |
| chr5 | 1.16E+08 | 116400000 | -0.86 | -0.89 | B>B |
| chr5 | 1.16E+08 | 116500000 | -0.92 | -0.90 | B>B |
| chr5 | 1.17E+08 | 116600000 | -0.91 | -0.94 | B>B |
| chr5 | 1.17E+08 | 116700000 | -0.96 | -0.94 | B>B |
| chr5 | 1.17E+08 | 116800000 | -0.94 | -0.93 | B>B |
| chr5 | 1.17E+08 | 116900000 | -0.92 | -0.91 | B>B |
| chr5 | 1.17E+08 | 117000000 | -0.91 | -0.94 | B>B |
| chr5 | 1.17E+08 | 117100000 | -0.96 | -0.92 | B>B |
| chr5 | 1.17E+08 | 117200000 | -0.95 | -0.98 | B>B |
| chr5 | 1.17E+08 | 117300000 | -0.95 | -0.93 | B>B |
| chr5 | 1.17E+08 | 117400000 | -0.88 | -0.93 | B>B |
| chr5 | 1.17E+08 | 117500000 | -0.98 | -0.96 | B>B |
| chr5 | 1.18E+08 | 117600000 | -0.94 | -0.95 | B>B |
| chr5 | 1.18E+08 | 117700000 | -0.92 | -0.97 | B>B |
| chr5 | 1.18E+08 | 117800000 | -0.84 | -0.98 | B>B |
| chr5 | 1.18E+08 | 117900000 | -0.81 | -0.97 | B>B |
| chr5 | 1.18E+08 | 118000000 | -0.66 | -0.94 | B>B |
| chr5 | 1.18E+08 | 118100000 | -0.63 | -0.87 | B>B |
| chr5 | 1.18E+08 | 118200000 | -0.35 | -0.56 | B>B |
| chr5 | 1.18E+08 | 118300000 | -0.05 | -0.29 | B>B |
| chr5 | 1.18E+08 | 118400000 | 0.36 | 0.25 | A>A |
| chr5 | 1.18E+08 | 118500000 | 0.76 | 0.58 | A>A |
| chr5 | 1.19E+08 | 118600000 | 0.79 | 0.59 | A>A |
| chr5 | 1.19E+08 | 118700000 | 0.74 | 0.57 | A>A |
| chr5 | 1.19E+08 | 118800000 | 0.31 | 0.16 | A>A |
| chr5 | 1.19E+08 | 118900000 | -0.03 | -0.30 | B>B |
| chr5 | 1.19E+08 | 119000000 | -0.45 | -0.90 | B>B |

| | | | | | |
|---|---|---|---|---|---|
| chr5 | 1.19E+08 | 119100000 | -0.63 | -0.97 | B>B |
| chr5 | 1.19E+08 | 119200000 | -0.77 | -0.93 | B>B |
| chr5 | 1.19E+08 | 119300000 | -0.83 | -1.00 | B>B |
| chr5 | 1.19E+08 | 119400000 | -0.87 | -1.00 | B>B |
| chr5 | 1.19E+08 | 119500000 | -0.77 | -0.99 | B>B |
| chr5 | 1.2E+08 | 119600000 | -0.90 | -1.00 | B>B |
| chr5 | 1.2E+08 | 119700000 | -0.89 | -1.00 | B>B |
| chr5 | 1.2E+08 | 119800000 | -0.74 | -0.88 | B>B |
| chr5 | 1.2E+08 | 119900000 | -0.79 | -0.81 | B>B |
| chr5 | 1.2E+08 | 120000000 | -0.80 | -0.79 | B>B |
| chr5 | 1.2E+08 | 120100000 | -0.78 | -0.76 | B>B |
| chr5 | 1.2E+08 | 120200000 | -0.63 | -0.52 | B>B |
| chr5 | 1.2E+08 | 120300000 | -0.50 | -0.35 | B>B |
| chr5 | 1.2E+08 | 120400000 | -0.27 | -0.15 | B>B |
| chr5 | 1.2E+08 | 120500000 | 0.00 | 0.02 | A>A |
| chr5 | 1.21E+08 | 120600000 | 0.11 | 0.26 | A>A |
| chr5 | 1.21E+08 | 120700000 | 0.15 | 0.31 | A>A |
| chr5 | 1.21E+08 | 120800000 | 0.04 | 0.18 | A>A |
| chr5 | 1.21E+08 | 120900000 | -0.09 | 0.07 | B>B |
| chr5 | 1.21E+08 | 121000000 | -0.42 | -0.14 | B>B |
| chr5 | 1.21E+08 | 121100000 | -0.54 | -0.17 | B>B |
| chr5 | 1.21E+08 | 121200000 | -0.58 | -0.36 | B>B |
| chr5 | 1.21E+08 | 121300000 | -0.41 | -0.39 | B>B |
| chr5 | 1.21E+08 | 121400000 | 0.18 | -0.12 | A>A |
| chr5 | 1.21E+08 | 121500000 | -0.11 | -0.42 | B>B |
| chr5 | 1.22E+08 | 121600000 | -0.45 | -0.76 | B>B |
| chr5 | 1.22E+08 | 121700000 | -0.41 | -0.97 | B>B |
| chr5 | 1.22E+08 | 121800000 | -0.24 | -0.94 | A>B |
| chr5 | 1.22E+08 | 121900000 | -0.10 | -0.69 | A>B |
| chr5 | 1.22E+08 | 122000000 | -0.21 | -0.87 | A>B |
| chr5 | 1.22E+08 | 122100000 | 0.21 | -0.51 | A>B |
| chr5 | 1.22E+08 | 122200000 | 0.59 | 0.14 | A>A |
| chr5 | 1.22E+08 | 122300000 | 0.53 | 0.24 | A>A |
| chr5 | 1.22E+08 | 122400000 | 0.46 | 0.15 | A>A |
| chr5 | 1.22E+08 | 122500000 | 0.44 | 0.19 | A>A |
| chr5 | 1.23E+08 | 122600000 | 0.25 | 0.06 | A>A |
| chr5 | 1.23E+08 | 122700000 | 0.14 | 0.16 | A>A |
| chr5 | 1.23E+08 | 122800000 | 0.61 | 0.50 | A>A |
| chr5 | 1.23E+08 | 122900000 | 0.74 | 0.65 | A>A |
| chr5 | 1.23E+08 | 123000000 | 0.37 | 0.54 | A>A |
| chr5 | 1.23E+08 | 123100000 | 0.40 | 0.69 | A>A |
| chr5 | 1.23E+08 | 123200000 | 0.45 | 0.82 | A>A |
| chr5 | 1.23E+08 | 123300000 | 0.42 | 0.73 | A>A |
| chr5 | 1.23E+08 | 123400000 | 0.51 | 0.89 | A>A |
| chr5 | 1.23E+08 | 123500000 | 0.54 | 0.95 | A>A |
| chr5 | 1.24E+08 | 123600000 | 0.63 | 0.90 | A>A |
| chr5 | 1.24E+08 | 123700000 | 0.74 | 0.93 | A>A |
| chr5 | 1.24E+08 | 123800000 | 0.69 | 1.01 | A>A |
| chr5 | 1.24E+08 | 123900000 | 0.89 | 1.05 | A>A |
| chr5 | 1.24E+08 | 124000000 | 0.72 | 0.88 | A>A |
| chr5 | 1.24E+08 | 124100000 | 0.70 | 0.89 | A>A |
| chr5 | 1.24E+08 | 124200000 | 0.76 | 0.93 | A>A |
| chr5 | 1.24E+08 | 124300000 | 0.74 | 0.95 | A>A |
| chr5 | 1.24E+08 | 124400000 | 0.66 | 0.85 | A>A |
| chr5 | 1.24E+08 | 124500000 | 0.71 | 1.02 | A>A |
| chr5 | 1.25E+08 | 124600000 | 0.56 | 0.91 | A>A |

| | | | | | |
|---|---|---|---|---|---|
| chr5 | 1.25E+08 | 124700000 | 0.59 | 1.01 | A>A |
| chr5 | 1.25E+08 | 124800000 | 0.52 | 0.77 | A>A |
| chr5 | 1.25E+08 | 124900000 | 0.52 | 0.94 | A>A |
| chr5 | 1.25E+08 | 125000000 | 0.52 | 1.02 | A>A |
| chr5 | 1.25E+08 | 125100000 | 0.35 | 0.99 | B>A |
| chr5 | 1.25E+08 | 125200000 | 0.26 | 0.82 | B>A |
| chr5 | 1.25E+08 | 125300000 | 0.19 | 0.93 | B>A |
| chr5 | 1.25E+08 | 125400000 | 0.22 | 0.93 | B>A |
| chr5 | 1.25E+08 | 125500000 | 0.22 | 0.89 | B>A |
| chr5 | 1.26E+08 | 125600000 | 0.13 | 0.81 | B>A |
| chr5 | 1.26E+08 | 125700000 | 0.25 | 0.74 | A>A |
| chr5 | 1.26E+08 | 125800000 | 0.50 | 1.16 | B>A |
| chr5 | 1.26E+08 | 125900000 | 0.44 | 0.95 | A>A |
| chr5 | 1.26E+08 | 126000000 | 0.69 | 0.97 | A>A |
| chr5 | 1.26E+08 | 126100000 | 0.92 | 1.08 | A>A |
| chr5 | 1.26E+08 | 126200000 | 0.93 | 0.96 | A>A |
| chr5 | 1.26E+08 | 126300000 | 0.51 | 0.86 | A>A |
| chr5 | 1.26E+08 | 126400000 | 0.41 | 0.75 | A>A |
| chr5 | 1.26E+08 | 126500000 | 0.31 | 0.82 | B>A |
| chr5 | 1.27E+08 | 126600000 | 0.23 | 0.86 | B>A |
| chr5 | 1.27E+08 | 126700000 | 0.19 | 0.66 | A>A |
| chr5 | 1.27E+08 | 126800000 | 0.25 | 0.62 | A>A |
| chr5 | 1.27E+08 | 126900000 | 0.22 | 0.43 | A>A |
| chr5 | 1.27E+08 | 127000000 | 0.31 | 0.20 | A>A |
| chr5 | 1.27E+08 | 127100000 | 0.39 | 0.23 | A>A |
| chr5 | 1.27E+08 | 127200000 | 0.39 | 0.35 | A>A |
| chr5 | 1.27E+08 | 127300000 | 0.62 | 0.56 | A>A |
| chr5 | 1.27E+08 | 127400000 | 0.79 | 0.61 | A>A |
| chr5 | 1.27E+08 | 127500000 | 0.65 | 0.53 | A>A |
| chr5 | 1.28E+08 | 127600000 | 0.18 | -0.01 | A>A |
| chr5 | 1.28E+08 | 127700000 | 0.03 | -0.08 | B>B |
| chr5 | 1.28E+08 | 127800000 | 0.11 | -0.12 | B>B |
| chr5 | 1.28E+08 | 127900000 | 0.13 | -0.14 | B>B |
| chr5 | 1.28E+08 | 128000000 | 0.09 | -0.22 | B>B |
| chr5 | 1.28E+08 | 128100000 | -0.18 | -0.61 | B>B |
| chr5 | 1.28E+08 | 128200000 | -0.31 | -0.81 | B>B |
| chr5 | 1.28E+08 | 128300000 | -0.44 | -0.90 | B>B |
| chr5 | 1.28E+08 | 128400000 | -0.51 | -1.02 | B>B |
| chr5 | 1.28E+08 | 128500000 | -0.40 | -0.79 | B>B |
| chr5 | 1.29E+08 | 128600000 | -0.81 | -1.12 | B>B |
| chr5 | 1.29E+08 | 128700000 | -0.83 | -1.12 | B>B |
| chr5 | 1.29E+08 | 128800000 | -0.64 | -1.12 | B>B |
| chr5 | 1.29E+08 | 128900000 | -0.79 | -1.18 | B>B |
| chr5 | 1.29E+08 | 129000000 | -0.81 | -1.13 | B>B |
| chr5 | 1.29E+08 | 129100000 | -0.67 | -0.97 | B>B |
| chr5 | 1.29E+08 | 129200000 | -0.49 | -0.73 | B>B |
| chr5 | 1.29E+08 | 129300000 | -0.33 | -0.37 | B>B |
| chr5 | 1.29E+08 | 129400000 | -0.42 | -0.38 | B>B |
| chr5 | 1.29E+08 | 129500000 | -0.06 | -0.08 | B>B |
| chr5 | 1.3E+08 | 129600000 | 0.17 | 0.26 | A>A |
| chr5 | 1.3E+08 | 129700000 | 0.21 | 0.34 | A>A |
| chr5 | 1.3E+08 | 129800000 | 0.22 | 0.35 | A>A |
| chr5 | 1.3E+08 | 129900000 | 0.27 | 0.40 | A>A |
| chr5 | 1.3E+08 | 130000000 | 0.26 | 0.34 | A>A |
| chr5 | 1.3E+08 | 130100000 | 0.20 | 0.19 | A>A |
| chr5 | 1.3E+08 | 130200000 | 0.28 | 0.07 | A>A |

| | | | | | |
|---|---|---|---|---|---|
| chr5 | 1.3E+08 | 130300000 | 0.07 | -0.20 | B>B |
| chr5 | 1.3E+08 | 130400000 | 0.04 | -0.34 | B>B |
| chr5 | 1.3E+08 | 130500000 | 0.14 | -0.08 | A>A |
| chr5 | 1.31E+08 | 130600000 | 0.71 | 0.48 | A>A |
| chr5 | 1.31E+08 | 130700000 | 0.74 | 0.58 | A>A |
| chr5 | 1.31E+08 | 130800000 | 0.49 | 0.45 | A>A |
| chr5 | 1.31E+08 | 130900000 | 0.80 | 0.59 | A>A |
| chr5 | 1.31E+08 | 131000000 | 0.83 | 0.67 | A>A |
| chr5 | 1.31E+08 | 131100000 | 0.89 | 0.71 | A>A |
| chr5 | 1.31E+08 | 131200000 | 0.61 | 0.42 | A>A |
| chr5 | 1.31E+08 | 131300000 | 0.55 | 0.26 | A>A |
| chr5 | 1.31E+08 | 131400000 | 0.62 | 0.38 | A>A |
| chr5 | 1.31E+08 | 131500000 | 0.83 | 0.71 | A>A |
| chr5 | 1.32E+08 | 131600000 | 1.05 | 0.98 | A>A |
| chr5 | 1.32E+08 | 131700000 | 1.10 | 1.11 | A>A |
| chr5 | 1.32E+08 | 131800000 | 1.21 | 1.20 | A>A |
| chr5 | 1.32E+08 | 131900000 | 1.34 | 1.13 | A>A |
| chr5 | 1.32E+08 | 132000000 | 1.31 | 1.24 | A>A |
| chr5 | 1.32E+08 | 132100000 | 1.35 | 1.32 | A>A |
| chr5 | 1.32E+08 | 132200000 | 1.29 | 1.17 | A>A |
| chr5 | 1.32E+08 | 132300000 | 1.39 | 1.22 | A>A |
| chr5 | 1.32E+08 | 132400000 | 1.42 | 1.22 | A>A |
| chr5 | 1.32E+08 | 132500000 | 1.16 | 0.84 | A>A |
| chr5 | 1.33E+08 | 132600000 | 1.36 | 0.94 | A>A |
| chr5 | 1.33E+08 | 132700000 | 1.24 | 0.80 | A>A |
| chr5 | 1.33E+08 | 132800000 | 1.20 | 0.65 | A>A |
| chr5 | 1.33E+08 | 132900000 | 1.19 | 0.62 | A>A |
| chr5 | 1.33E+08 | 133000000 | 1.16 | 0.67 | A>A |
| chr5 | 1.33E+08 | 133100000 | 1.30 | 0.76 | A>A |
| chr5 | 1.33E+08 | 133200000 | 1.25 | 0.76 | A>A |
| chr5 | 1.33E+08 | 133300000 | 1.11 | 0.75 | A>A |
| chr5 | 1.33E+08 | 133400000 | 1.21 | 1.08 | A>A |
| chr5 | 1.33E+08 | 133500000 | 1.32 | 1.24 | A>A |
| chr5 | 1.34E+08 | 133600000 | 1.28 | 1.30 | A>A |
| chr5 | 1.34E+08 | 133700000 | 1.02 | 1.08 | A>A |
| chr5 | 1.34E+08 | 133800000 | 1.30 | 1.26 | A>A |
| chr5 | 1.34E+08 | 133900000 | 1.33 | 1.32 | A>A |
| chr5 | 1.34E+08 | 134000000 | 1.11 | 1.16 | A>A |
| chr5 | 1.34E+08 | 134100000 | 1.26 | 1.24 | A>A |
| chr5 | 1.34E+08 | 134200000 | 1.29 | 1.22 | A>A |
| chr5 | 1.34E+08 | 134300000 | 1.14 | 1.03 | A>A |
| chr5 | 1.34E+08 | 134400000 | 1.09 | 1.14 | A>A |
| chr5 | 1.34E+08 | 134500000 | 1.23 | 1.21 | A>A |
| chr5 | 1.35E+08 | 134600000 | 0.93 | 0.92 | A>A |
| chr5 | 1.35E+08 | 134700000 | 0.96 | 0.87 | A>A |
| chr5 | 1.35E+08 | 134800000 | 0.87 | 0.48 | A>A |
| chr5 | 1.35E+08 | 134900000 | 0.66 | -0.03 | A>B |
| chr5 | 1.35E+08 | 135000000 | 0.35 | -0.44 | A>B |
| chr5 | 1.35E+08 | 135100000 | 0.29 | -0.52 | A>B |
| chr5 | 1.35E+08 | 135200000 | 0.35 | -0.20 | A>A |
| chr5 | 1.35E+08 | 135300000 | 0.51 | 0.07 | A>A |
| chr5 | 1.35E+08 | 135400000 | 0.98 | 0.59 | A>A |
| chr5 | 1.35E+08 | 135500000 | 0.70 | 0.37 | A>A |
| chr5 | 1.36E+08 | 135600000 | 0.26 | -0.20 | A>A |
| chr5 | 1.36E+08 | 135700000 | -0.31 | -0.91 | A>B |
| chr5 | 1.36E+08 | 135800000 | -0.34 | -1.07 | A>B |

| | | | | | |
|---|---|---|---|---|---|
| chr5 | 1.36E+08 | 135900000 | -0.36 | -1.11 | A>B |
| chr5 | 1.36E+08 | 136000000 | -0.40 | -1.12 | A>B |
| chr5 | 1.36E+08 | 136100000 | -0.35 | -0.96 | A>B |
| chr5 | 1.36E+08 | 136200000 | -0.29 | -1.02 | A>B |
| chr5 | 1.36E+08 | 136300000 | -0.42 | -1.11 | A>B |
| chr5 | 1.36E+08 | 136400000 | -0.12 | -1.08 | A>B |
| chr5 | 1.36E+08 | 136500000 | 0.18 | -0.85 | A>B |
| chr5 | 1.37E+08 | 136600000 | 0.31 | -0.75 | A>B |
| chr5 | 1.37E+08 | 136700000 | 0.24 | -0.63 | A>B |
| chr5 | 1.37E+08 | 136800000 | 0.11 | -0.89 | A>B |
| chr5 | 1.37E+08 | 136900000 | 0.18 | -0.65 | A>B |
| chr5 | 1.37E+08 | 137000000 | 0.47 | -0.07 | A>A |
| chr5 | 1.37E+08 | 137100000 | 0.77 | 0.31 | A>A |
| chr5 | 1.37E+08 | 137200000 | 1.04 | 0.80 | A>A |
| chr5 | 1.37E+08 | 137300000 | 1.15 | 1.03 | A>A |
| chr5 | 1.37E+08 | 137400000 | 1.03 | 0.84 | A>A |
| chr5 | 1.37E+08 | 137500000 | 1.03 | 0.78 | A>A |
| chr5 | 1.38E+08 | 137600000 | 1.12 | 1.06 | A>A |
| chr5 | 1.38E+08 | 137700000 | 1.23 | 1.20 | A>A |
| chr5 | 1.38E+08 | 137800000 | 1.18 | 1.14 | A>A |
| chr5 | 1.38E+08 | 137900000 | 1.27 | 1.25 | A>A |
| chr5 | 1.38E+08 | 138000000 | 1.07 | 1.15 | A>A |
| chr5 | 1.38E+08 | 138100000 | 1.20 | 1.12 | A>A |
| chr5 | 1.38E+08 | 138200000 | 1.22 | 1.23 | A>A |
| chr5 | 1.38E+08 | 138300000 | 1.19 | 1.23 | A>A |
| chr5 | 1.38E+08 | 138400000 | 1.17 | 1.22 | A>A |
| chr5 | 1.38E+08 | 138500000 | 1.23 | 1.30 | A>A |
| chr5 | 1.39E+08 | 138600000 | 1.09 | 1.17 | A>A |
| chr5 | 1.39E+08 | 138700000 | 1.14 | 1.23 | A>A |
| chr5 | 1.39E+08 | 138800000 | 1.12 | 1.20 | A>A |
| chr5 | 1.39E+08 | 138900000 | 0.00 | 0.00 | NO |
| chr5 | 1.39E+08 | 139000000 | 1.26 | 1.20 | A>A |
| chr5 | 1.39E+08 | 139100000 | 1.44 | 1.38 | A>A |
| chr5 | 1.39E+08 | 139200000 | 1.14 | 1.20 | A>A |
| chr5 | 1.39E+08 | 139300000 | 1.36 | 1.20 | A>A |
| chr5 | 1.39E+08 | 139400000 | 1.23 | 1.21 | A>A |
| chr5 | 1.39E+08 | 139500000 | 1.19 | 1.31 | A>A |
| chr5 | 1.4E+08 | 139600000 | 1.11 | 1.22 | A>A |
| chr5 | 1.4E+08 | 139700000 | 1.15 | 1.30 | A>A |
| chr5 | 1.4E+08 | 139800000 | 1.04 | 1.15 | A>A |
| chr5 | 1.4E+08 | 139900000 | 1.15 | 1.33 | A>A |
| chr5 | 1.4E+08 | 140000000 | 1.04 | 1.23 | A>A |
| chr5 | 1.4E+08 | 140100000 | 0.97 | 1.16 | A>A |
| chr5 | 1.4E+08 | 140200000 | 0.18 | 1.06 | B>A |
| chr5 | 1.4E+08 | 140300000 | 0.10 | 1.17 | B>A |
| chr5 | 1.4E+08 | 140400000 | 0.56 | 1.35 | B>A |
| chr5 | 1.4E+08 | 140500000 | -0.03 | 1.18 | B>A |
| chr5 | 1.41E+08 | 140600000 | -0.32 | 1.17 | B>A |
| chr5 | 1.41E+08 | 140700000 | -0.39 | 1.19 | B>A |
| chr5 | 1.41E+08 | 140800000 | 0.05 | 1.22 | B>A |
| chr5 | 1.41E+08 | 140900000 | 0.24 | 1.20 | B>A |
| chr5 | 1.41E+08 | 141000000 | 0.52 | 1.12 | B>A |
| chr5 | 1.41E+08 | 141100000 | 0.88 | 1.06 | A>A |
| chr5 | 1.41E+08 | 141200000 | 1.16 | 1.29 | A>A |
| chr5 | 1.41E+08 | 141300000 | 1.03 | 1.27 | A>A |
| chr5 | 1.41E+08 | 141400000 | 0.98 | 1.17 | A>A |

| | | | | | |
|---|---|---|---|---|---|
| chr5 | 1.41E+08 | 141500000 | 1.17 | 1.22 | A>A |
| chr5 | 1.42E+08 | 141600000 | 0.84 | 1.02 | A>A |
| chr5 | 1.42E+08 | 141700000 | 1.13 | 1.12 | A>A |
| chr5 | 1.42E+08 | 141800000 | 1.12 | 1.14 | A>A |
| chr5 | 1.42E+08 | 141900000 | 1.18 | 1.26 | A>A |
| chr5 | 1.42E+08 | 142000000 | 0.98 | 0.98 | A>A |
| chr5 | 1.42E+08 | 142100000 | 0.79 | 0.93 | A>A |
| chr5 | 1.42E+08 | 142200000 | 1.03 | 1.12 | A>A |
| chr5 | 1.42E+08 | 142300000 | 0.81 | 1.04 | A>A |
| chr5 | 1.42E+08 | 142400000 | 0.60 | 0.67 | A>A |
| chr5 | 1.42E+08 | 142500000 | 0.68 | 0.66 | A>A |
| chr5 | 1.43E+08 | 142600000 | 0.69 | 0.68 | A>A |
| chr5 | 1.43E+08 | 142700000 | 0.62 | 0.62 | A>A |
| chr5 | 1.43E+08 | 142800000 | 0.59 | 0.72 | A>A |
| chr5 | 1.43E+08 | 142900000 | 0.82 | 0.82 | A>A |
| chr5 | 1.43E+08 | 143000000 | 0.67 | 0.75 | A>A |
| chr5 | 1.43E+08 | 143100000 | 0.73 | 0.74 | A>A |
| chr5 | 1.43E+08 | 143200000 | 0.81 | 0.73 | A>A |
| chr5 | 1.43E+08 | 143300000 | 0.65 | 0.59 | A>A |
| chr5 | 1.43E+08 | 143400000 | 0.48 | 0.53 | A>A |
| chr5 | 1.43E+08 | 143500000 | 0.46 | 0.26 | A>A |
| chr5 | 1.44E+08 | 143600000 | 0.20 | 0.08 | A>A |
| chr5 | 1.44E+08 | 143700000 | -0.32 | -0.74 | B>B |
| chr5 | 1.44E+08 | 143800000 | -0.40 | -0.97 | B>B |
| chr5 | 1.44E+08 | 143900000 | -0.52 | -0.99 | B>B |
| chr5 | 1.44E+08 | 144000000 | -0.55 | -0.93 | B>B |
| chr5 | 1.44E+08 | 144100000 | -0.65 | -0.99 | B>B |
| chr5 | 1.44E+08 | 144200000 | -0.74 | -0.95 | B>B |
| chr5 | 1.44E+08 | 144300000 | -0.74 | -0.97 | B>B |
| chr5 | 1.44E+08 | 144400000 | -0.71 | -0.92 | B>B |
| chr5 | 1.44E+08 | 144500000 | -0.65 | -0.99 | B>B |
| chr5 | 1.45E+08 | 144600000 | -0.58 | -0.92 | B>B |
| chr5 | 1.45E+08 | 144700000 | -0.48 | -0.84 | B>B |
| chr5 | 1.45E+08 | 144800000 | -0.56 | -0.71 | B>B |
| chr5 | 1.45E+08 | 144900000 | -0.34 | -0.57 | B>B |
| chr5 | 1.45E+08 | 145000000 | -0.24 | -0.43 | B>B |
| chr5 | 1.45E+08 | 145100000 | -0.10 | -0.28 | B>B |
| chr5 | 1.45E+08 | 145200000 | 0.11 | -0.12 | B>B |
| chr5 | 1.45E+08 | 145300000 | 0.25 | 0.12 | A>A |
| chr5 | 1.45E+08 | 145400000 | 0.39 | 0.32 | A>A |
| chr5 | 1.45E+08 | 145500000 | 0.27 | 0.19 | A>A |
| chr5 | 1.46E+08 | 145600000 | 0.70 | 0.38 | A>A |
| chr5 | 1.46E+08 | 145700000 | 0.74 | 0.30 | A>A |
| chr5 | 1.46E+08 | 145800000 | 0.44 | -0.15 | A>B |
| chr5 | 1.46E+08 | 145900000 | 0.32 | 0.02 | A>A |
| chr5 | 1.46E+08 | 146000000 | -0.25 | -0.60 | B>B |
| chr5 | 1.46E+08 | 146100000 | -0.33 | -0.66 | B>B |
| chr5 | 1.46E+08 | 146200000 | -0.39 | -0.59 | B>B |
| chr5 | 1.46E+08 | 146300000 | -0.33 | -0.83 | B>B |
| chr5 | 1.46E+08 | 146400000 | -0.50 | -0.72 | B>B |
| chr5 | 1.46E+08 | 146500000 | -0.66 | -0.98 | B>B |
| chr5 | 1.47E+08 | 146600000 | -0.60 | -0.75 | B>B |
| chr5 | 1.47E+08 | 146700000 | -0.40 | -0.49 | B>B |
| chr5 | 1.47E+08 | 146800000 | -0.30 | -0.57 | B>B |
| chr5 | 1.47E+08 | 146900000 | -0.19 | -0.61 | B>B |
| chr5 | 1.47E+08 | 147000000 | -0.26 | -0.69 | B>B |

| | | | | | |
|---|---|---|---|---|---|
| chr5 | 1.47E+08 | 147100000 | -0.54 | -0.90 | B>B |
| chr5 | 1.47E+08 | 147200000 | -0.52 | -0.95 | B>B |
| chr5 | 1.47E+08 | 147300000 | -0.33 | -0.77 | B>B |
| chr5 | 1.47E+08 | 147400000 | -0.62 | -0.87 | B>B |
| chr5 | 1.47E+08 | 147500000 | -0.69 | -0.97 | B>B |
| chr5 | 1.48E+08 | 147600000 | -0.52 | -0.87 | B>B |
| chr5 | 1.48E+08 | 147700000 | -0.55 | -0.85 | B>B |
| chr5 | 1.48E+08 | 147800000 | -0.27 | -0.54 | B>B |
| chr5 | 1.48E+08 | 147900000 | -0.20 | -0.52 | B>B |
| chr5 | 1.48E+08 | 148000000 | -0.65 | -1.00 | B>B |
| chr5 | 1.48E+08 | 148100000 | -0.46 | -0.89 | B>B |
| chr5 | 1.48E+08 | 148200000 | -0.09 | -0.29 | B>B |
| chr5 | 1.48E+08 | 148300000 | 0.11 | -0.09 | A>A |
| chr5 | 1.48E+08 | 148400000 | 0.15 | 0.04 | A>A |
| chr5 | 1.48E+08 | 148500000 | 0.05 | -0.16 | B>B |
| chr5 | 1.49E+08 | 148600000 | 0.45 | 0.30 | A>A |
| chr5 | 1.49E+08 | 148700000 | 0.62 | 0.43 | A>A |
| chr5 | 1.49E+08 | 148800000 | 0.93 | 0.85 | A>A |
| chr5 | 1.49E+08 | 148900000 | 0.96 | 0.93 | A>A |
| chr5 | 1.49E+08 | 149000000 | 0.90 | 0.92 | A>A |
| chr5 | 1.49E+08 | 149100000 | 0.87 | 1.01 | A>A |
| chr5 | 1.49E+08 | 149200000 | 1.24 | 1.13 | A>A |
| chr5 | 1.49E+08 | 149300000 | 1.15 | 1.15 | A>A |
| chr5 | 1.49E+08 | 149400000 | 1.09 | 1.10 | A>A |
| chr5 | 1.49E+08 | 149500000 | 1.13 | 1.15 | A>A |
| chr5 | 1.5E+08 | 149600000 | 1.13 | 1.17 | A>A |
| chr5 | 1.5E+08 | 149700000 | 1.17 | 1.33 | A>A |
| chr5 | 1.5E+08 | 149800000 | 1.11 | 1.20 | A>A |
| chr5 | 1.5E+08 | 149900000 | 0.91 | 1.10 | A>A |
| chr5 | 1.5E+08 | 150000000 | 0.85 | 1.17 | A>A |
| chr5 | 1.5E+08 | 150100000 | 0.78 | 1.16 | A>A |
| chr5 | 1.5E+08 | 150200000 | 0.33 | 0.93 | B>A |
| chr5 | 1.5E+08 | 150300000 | -0.27 | 0.77 | B>A |
| chr5 | 1.5E+08 | 150400000 | -0.24 | 0.75 | B>A |
| chr5 | 1.5E+08 | 150500000 | 0.38 | 0.88 | A>A |
| chr5 | 1.51E+08 | 150600000 | 0.41 | 0.79 | A>A |
| chr5 | 1.51E+08 | 150700000 | 0.40 | 0.61 | A>A |
| chr5 | 1.51E+08 | 150800000 | 0.07 | 0.28 | A>A |
| chr5 | 1.51E+08 | 150900000 | 0.24 | 0.37 | A>A |
| chr5 | 1.51E+08 | 151000000 | 0.34 | 0.39 | A>A |
| chr5 | 1.51E+08 | 151100000 | 0.40 | 0.50 | A>A |
| chr5 | 1.51E+08 | 151200000 | 0.34 | 0.27 | A>A |
| chr5 | 1.51E+08 | 151300000 | -0.26 | -0.64 | B>B |
| chr5 | 1.51E+08 | 151400000 | -0.54 | -0.87 | B>B |
| chr5 | 1.51E+08 | 151500000 | -0.51 | -0.93 | B>B |
| chr5 | 1.52E+08 | 151600000 | -0.51 | -0.94 | B>B |
| chr5 | 1.52E+08 | 151700000 | -0.64 | -0.95 | B>B |
| chr5 | 1.52E+08 | 151800000 | -0.71 | -0.86 | B>B |
| chr5 | 1.52E+08 | 151900000 | -0.56 | -0.74 | B>B |
| chr5 | 1.52E+08 | 152000000 | -0.56 | -0.79 | B>B |
| chr5 | 1.52E+08 | 152100000 | -0.78 | -0.95 | B>B |
| chr5 | 1.52E+08 | 152200000 | -0.89 | -1.01 | B>B |
| chr5 | 1.52E+08 | 152300000 | -0.87 | -1.03 | B>B |
| chr5 | 1.52E+08 | 152400000 | -0.93 | -0.99 | B>B |
| chr5 | 1.52E+08 | 152500000 | -0.83 | -1.01 | B>B |
| chr5 | 1.53E+08 | 152600000 | -0.81 | -0.98 | B>B |

| | | | | | |
|---|---|---|---|---|---|
| chr5 | 1.53E+08 | 152700000 | -0.78 | -0.96 | B>B |
| chr5 | 1.53E+08 | 152800000 | -0.69 | -0.97 | B>B |
| chr5 | 1.53E+08 | 152900000 | -0.58 | -0.94 | B>B |
| chr5 | 1.53E+08 | 153000000 | -0.48 | -0.93 | B>B |
| chr5 | 1.53E+08 | 153100000 | -0.53 | -0.99 | B>B |
| chr5 | 1.53E+08 | 153200000 | -0.54 | -0.95 | B>B |
| chr5 | 1.53E+08 | 153300000 | -0.41 | -0.90 | B>B |
| chr5 | 1.53E+08 | 153400000 | -0.18 | -0.36 | B>B |
| chr5 | 1.53E+08 | 153500000 | 0.18 | 0.11 | A>A |
| chr5 | 1.54E+08 | 153600000 | 0.38 | 0.26 | A>A |
| chr5 | 1.54E+08 | 153700000 | 0.45 | 0.59 | A>A |
| chr5 | 1.54E+08 | 153800000 | 0.36 | 0.42 | A>A |
| chr5 | 1.54E+08 | 153900000 | 0.46 | 0.43 | A>A |
| chr5 | 1.54E+08 | 154000000 | 0.64 | 0.50 | A>A |
| chr5 | 1.54E+08 | 154100000 | 0.62 | 0.61 | A>A |
| chr5 | 1.54E+08 | 154200000 | 0.82 | 0.82 | A>A |
| chr5 | 1.54E+08 | 154300000 | 0.74 | 0.61 | A>A |
| chr5 | 1.54E+08 | 154400000 | 0.47 | 0.06 | A>A |
| chr5 | 1.54E+08 | 154500000 | 0.26 | -0.41 | A>B |
| chr5 | 1.55E+08 | 154600000 | 0.02 | -0.58 | A>B |
| chr5 | 1.55E+08 | 154700000 | -0.05 | -0.84 | A>B |
| chr5 | 1.55E+08 | 154800000 | -0.03 | -0.90 | A>B |
| chr5 | 1.55E+08 | 154900000 | -0.02 | -0.88 | A>B |
| chr5 | 1.55E+08 | 155000000 | 0.06 | -0.77 | A>B |
| chr5 | 1.55E+08 | 155100000 | 0.14 | -0.52 | A>B |
| chr5 | 1.55E+08 | 155200000 | 0.00 | -0.42 | NO |
| chr5 | 1.55E+08 | 155300000 | 0.30 | -0.34 | A>B |
| chr5 | 1.55E+08 | 155400000 | 0.38 | 0.03 | A>A |
| chr5 | 1.55E+08 | 155500000 | 0.55 | 0.29 | A>A |
| chr5 | 1.56E+08 | 155600000 | 0.49 | 0.37 | A>A |
| chr5 | 1.56E+08 | 155700000 | 0.55 | 0.30 | A>A |
| chr5 | 1.56E+08 | 155800000 | 0.55 | 0.36 | A>A |
| chr5 | 1.56E+08 | 155900000 | 0.54 | 0.31 | A>A |
| chr5 | 1.56E+08 | 156000000 | 0.49 | 0.16 | A>A |
| chr5 | 1.56E+08 | 156100000 | 0.47 | 0.11 | A>A |
| chr5 | 1.56E+08 | 156200000 | 0.35 | 0.12 | A>A |
| chr5 | 1.56E+08 | 156300000 | 0.21 | -0.12 | A>A |
| chr5 | 1.56E+08 | 156400000 | 0.47 | 0.01 | A>A |
| chr5 | 1.56E+08 | 156500000 | 0.73 | 0.34 | A>A |
| chr5 | 1.57E+08 | 156600000 | 0.51 | 0.19 | A>A |
| chr5 | 1.57E+08 | 156700000 | 0.63 | 0.01 | A>B |
| chr5 | 1.57E+08 | 156800000 | 0.68 | 0.23 | A>A |
| chr5 | 1.57E+08 | 156900000 | 0.71 | 0.35 | A>A |
| chr5 | 1.57E+08 | 157000000 | 0.67 | 0.18 | A>A |
| chr5 | 1.57E+08 | 157100000 | 0.71 | 0.15 | A>A |
| chr5 | 1.57E+08 | 157200000 | 0.55 | 0.16 | A>A |
| chr5 | 1.57E+08 | 157300000 | 0.56 | 0.24 | A>A |
| chr5 | 1.57E+08 | 157400000 | 0.57 | 0.04 | A>A |
| chr5 | 1.57E+08 | 157500000 | 0.15 | -0.39 | B>B |
| chr5 | 1.58E+08 | 157600000 | -0.01 | -0.60 | A>B |
| chr5 | 1.58E+08 | 157700000 | -0.22 | -0.63 | B>B |
| chr5 | 1.58E+08 | 157800000 | -0.30 | -0.52 | B>B |
| chr5 | 1.58E+08 | 157900000 | -0.08 | -0.40 | B>B |
| chr5 | 1.58E+08 | 158000000 | -0.19 | -0.61 | B>B |
| chr5 | 1.58E+08 | 158100000 | -0.10 | -0.57 | B>B |
| chr5 | 1.58E+08 | 158200000 | 0.20 | -0.36 | B>B |

| | | | | | |
|---|---|---|---|---|---|
| chr5 | 1.58E+08 | 158300000 | 0.41 | -0.35 | A>B |
| chr5 | 1.58E+08 | 158400000 | 0.46 | -0.10 | A>A |
| chr5 | 1.58E+08 | 158500000 | 0.44 | -0.07 | A>A |
| chr5 | 1.59E+08 | 158600000 | 0.31 | 0.06 | A>A |
| chr5 | 1.59E+08 | 158700000 | 0.87 | 0.71 | A>A |
| chr5 | 1.59E+08 | 158800000 | 0.67 | 0.61 | A>A |
| chr5 | 1.59E+08 | 158900000 | 0.72 | 0.68 | A>A |
| chr5 | 1.59E+08 | 159000000 | 0.87 | 0.86 | A>A |
| chr5 | 1.59E+08 | 159100000 | 0.88 | 0.91 | A>A |
| chr5 | 1.59E+08 | 159200000 | 0.94 | 0.93 | A>A |
| chr5 | 1.59E+08 | 159300000 | 1.12 | 1.08 | A>A |
| chr5 | 1.59E+08 | 159400000 | 0.82 | 0.81 | A>A |
| chr5 | 1.59E+08 | 159500000 | 0.69 | 0.68 | A>A |
| chr5 | 1.6E+08 | 159600000 | 0.86 | 0.77 | A>A |
| chr5 | 1.6E+08 | 159700000 | 0.90 | 0.81 | A>A |
| chr5 | 1.6E+08 | 159800000 | 0.90 | 0.81 | A>A |
| chr5 | 1.6E+08 | 159900000 | 0.40 | 0.19 | A>A |
| chr5 | 1.6E+08 | 160000000 | -0.13 | -0.55 | B>B |
| chr5 | 1.6E+08 | 160100000 | -0.28 | -0.80 | B>B |
| chr5 | 1.6E+08 | 160200000 | -0.37 | -0.83 | B>B |
| chr5 | 1.6E+08 | 160300000 | -0.54 | -0.88 | B>B |
| chr5 | 1.6E+08 | 160400000 | -0.46 | -0.82 | B>B |
| chr5 | 1.6E+08 | 160500000 | -0.63 | -0.84 | B>B |
| chr5 | 1.61E+08 | 160600000 | -0.68 | -0.88 | B>B |
| chr5 | 1.61E+08 | 160700000 | -0.69 | -0.86 | B>B |
| chr5 | 1.61E+08 | 160800000 | -0.71 | -0.85 | B>B |
| chr5 | 1.61E+08 | 160900000 | -0.72 | -0.80 | B>B |
| chr5 | 1.61E+08 | 161000000 | -0.61 | -0.72 | B>B |
| chr5 | 1.61E+08 | 161100000 | -0.65 | -0.81 | B>B |
| chr5 | 1.61E+08 | 161200000 | -0.67 | -0.81 | B>B |
| chr5 | 1.61E+08 | 161300000 | -0.64 | -0.83 | B>B |
| chr5 | 1.61E+08 | 161400000 | -0.72 | -0.84 | B>B |
| chr5 | 1.61E+08 | 161500000 | -0.73 | -0.80 | B>B |
| chr5 | 1.62E+08 | 161600000 | -0.66 | -0.80 | B>B |
| chr5 | 1.62E+08 | 161700000 | -0.65 | -0.83 | B>B |
| chr5 | 1.62E+08 | 161800000 | -0.61 | -0.83 | B>B |
| chr5 | 1.62E+08 | 161900000 | -0.69 | -0.80 | B>B |
| chr5 | 1.62E+08 | 162000000 | -0.66 | -0.83 | B>B |
| chr5 | 1.62E+08 | 162100000 | -0.69 | -0.77 | B>B |
| chr5 | 1.62E+08 | 162200000 | -0.62 | -0.76 | B>B |
| chr5 | 1.62E+08 | 162300000 | -0.65 | -0.76 | B>B |
| chr5 | 1.62E+08 | 162400000 | -0.73 | -0.81 | B>B |
| chr5 | 1.62E+08 | 162500000 | -0.66 | -0.81 | B>B |
| chr5 | 1.63E+08 | 162600000 | -0.43 | -0.76 | B>B |
| chr5 | 1.63E+08 | 162700000 | -0.42 | -0.76 | B>B |
| chr5 | 1.63E+08 | 162800000 | -0.09 | -0.39 | B>B |
| chr5 | 1.63E+08 | 162900000 | 0.22 | 0.02 | A>A |
| chr5 | 1.63E+08 | 163000000 | 0.21 | -0.11 | A>A |
| chr5 | 1.63E+08 | 163100000 | -0.26 | -0.66 | B>B |
| chr5 | 1.63E+08 | 163200000 | -0.55 | -0.81 | B>B |
| chr5 | 1.63E+08 | 163300000 | -0.64 | -0.83 | B>B |
| chr5 | 1.63E+08 | 163400000 | -0.68 | -0.83 | B>B |
| chr5 | 1.63E+08 | 163500000 | -0.61 | -0.85 | B>B |
| chr5 | 1.64E+08 | 163600000 | -0.68 | -0.88 | B>B |
| chr5 | 1.64E+08 | 163700000 | -0.71 | -0.87 | B>B |
| chr5 | 1.64E+08 | 163800000 | -0.71 | -0.91 | B>B |

| | | | | | |
|---|---|---|---|---|---|
| chr5 | 1.64E+08 | 163900000 | -0.69 | -0.84 | B>B |
| chr5 | 1.64E+08 | 164000000 | -0.57 | -0.82 | B>B |
| chr5 | 1.64E+08 | 164100000 | -0.77 | -0.87 | B>B |
| chr5 | 1.64E+08 | 164200000 | -0.71 | -0.86 | B>B |
| chr5 | 1.64E+08 | 164300000 | -0.72 | -0.88 | B>B |
| chr5 | 1.64E+08 | 164400000 | -0.70 | -0.91 | B>B |
| chr5 | 1.64E+08 | 164500000 | -0.72 | -0.82 | B>B |
| chr5 | 1.65E+08 | 164600000 | -0.72 | -0.87 | B>B |
| chr5 | 1.65E+08 | 164700000 | -0.62 | -0.86 | B>B |
| chr5 | 1.65E+08 | 164800000 | -0.72 | -0.89 | B>B |
| chr5 | 1.65E+08 | 164900000 | -0.71 | -0.78 | B>B |
| chr5 | 1.65E+08 | 165000000 | -0.65 | -0.80 | B>B |
| chr5 | 1.65E+08 | 165100000 | -0.70 | -0.73 | B>B |
| chr5 | 1.65E+08 | 165200000 | -0.70 | -0.79 | B>B |
| chr5 | 1.65E+08 | 165300000 | -0.68 | -0.75 | B>B |
| chr5 | 1.65E+08 | 165400000 | -0.65 | -0.73 | B>B |
| chr5 | 1.65E+08 | 165500000 | -0.70 | -0.78 | B>B |
| chr5 | 1.66E+08 | 165600000 | -0.69 | -0.75 | B>B |
| chr5 | 1.66E+08 | 165700000 | -0.72 | -0.75 | B>B |
| chr5 | 1.66E+08 | 165800000 | -0.57 | -0.70 | B>B |
| chr5 | 1.66E+08 | 165900000 | -0.55 | -0.59 | B>B |
| chr5 | 1.66E+08 | 166000000 | -0.63 | -0.67 | B>B |
| chr5 | 1.66E+08 | 166100000 | -0.59 | -0.50 | B>B |
| chr5 | 1.66E+08 | 166200000 | -0.32 | -0.30 | B>B |
| chr5 | 1.66E+08 | 166300000 | -0.36 | -0.26 | B>B |
| chr5 | 1.66E+08 | 166400000 | -0.37 | -0.12 | B>B |
| chr5 | 1.66E+08 | 166500000 | -0.35 | -0.45 | B>B |
| chr5 | 1.67E+08 | 166600000 | -0.33 | -0.60 | B>B |
| chr5 | 1.67E+08 | 166700000 | -0.20 | -0.48 | B>B |
| chr5 | 1.67E+08 | 166800000 | -0.03 | -0.22 | B>B |
| chr5 | 1.67E+08 | 166900000 | -0.37 | -0.52 | B>B |
| chr5 | 1.67E+08 | 167000000 | -0.27 | -0.48 | B>B |
| chr5 | 1.67E+08 | 167100000 | 0.11 | 0.10 | A>A |
| chr5 | 1.67E+08 | 167200000 | 0.13 | 0.03 | A>A |
| chr5 | 1.67E+08 | 167300000 | 0.00 | -0.16 | B>B |
| chr5 | 1.67E+08 | 167400000 | -0.06 | -0.23 | B>B |
| chr5 | 1.67E+08 | 167500000 | -0.05 | -0.15 | B>B |
| chr5 | 1.68E+08 | 167600000 | 0.27 | 0.06 | A>A |
| chr5 | 1.68E+08 | 167700000 | 0.39 | 0.18 | A>A |
| chr5 | 1.68E+08 | 167800000 | 1.00 | 0.88 | A>A |
| chr5 | 1.68E+08 | 167900000 | 0.77 | 0.76 | A>A |
| chr5 | 1.68E+08 | 168000000 | 0.95 | 0.85 | A>A |
| chr5 | 1.68E+08 | 168100000 | 0.89 | 0.82 | A>A |
| chr5 | 1.68E+08 | 168200000 | 0.71 | 0.56 | A>A |
| chr5 | 1.68E+08 | 168300000 | 0.84 | 0.62 | A>A |
| chr5 | 1.68E+08 | 168400000 | 0.77 | 0.50 | A>A |
| chr5 | 1.68E+08 | 168500000 | 0.86 | 0.73 | A>A |
| chr5 | 1.69E+08 | 168600000 | 0.97 | 0.92 | A>A |
| chr5 | 1.69E+08 | 168700000 | 0.91 | 0.77 | A>A |
| chr5 | 1.69E+08 | 168800000 | 0.81 | 0.67 | A>A |
| chr5 | 1.69E+08 | 168900000 | 1.05 | 0.85 | A>A |
| chr5 | 1.69E+08 | 169000000 | 0.74 | 0.64 | A>A |
| chr5 | 1.69E+08 | 169100000 | 0.60 | 0.73 | A>A |
| chr5 | 1.69E+08 | 169200000 | 0.76 | 0.66 | A>A |
| chr5 | 1.69E+08 | 169300000 | 0.75 | 0.73 | A>A |
| chr5 | 1.69E+08 | 169400000 | 0.80 | 0.72 | A>A |

| | | | | | |
|---|---|---|---|---|---|
| chr5 | 1.69E+08 | 169500000 | 0.70 | 0.81 | A>A |
| chr5 | 1.7E+08 | 169600000 | 0.87 | 0.81 | A>A |
| chr5 | 1.7E+08 | 169700000 | 0.70 | 0.66 | A>A |
| chr5 | 1.7E+08 | 169800000 | 0.81 | 0.67 | A>A |
| chr5 | 1.7E+08 | 169900000 | 0.66 | 0.69 | A>A |
| chr5 | 1.7E+08 | 170000000 | 0.92 | 0.86 | A>A |
| chr5 | 1.7E+08 | 170100000 | 0.60 | 0.71 | A>A |
| chr5 | 1.7E+08 | 170200000 | 0.65 | 0.78 | A>A |
| chr5 | 1.7E+08 | 170300000 | 0.63 | 0.94 | A>A |
| chr5 | 1.7E+08 | 170400000 | 0.79 | 0.94 | A>A |
| chr5 | 1.7E+08 | 170500000 | 0.66 | 0.90 | A>A |
| chr5 | 1.71E+08 | 170600000 | 0.61 | 0.88 | A>A |
| chr5 | 1.71E+08 | 170700000 | 0.70 | 0.79 | A>A |
| chr5 | 1.71E+08 | 170800000 | 0.68 | 0.82 | A>A |
| chr5 | 1.71E+08 | 170900000 | 0.97 | 0.79 | A>A |
| chr5 | 1.71E+08 | 171000000 | 0.97 | 0.77 | A>A |
| chr5 | 1.71E+08 | 171100000 | 0.86 | 0.78 | A>A |
| chr5 | 1.71E+08 | 171200000 | 1.04 | 0.77 | A>A |
| chr5 | 1.71E+08 | 171300000 | 0.92 | 0.77 | A>A |
| chr5 | 1.71E+08 | 171400000 | 1.07 | 1.02 | A>A |
| chr5 | 1.71E+08 | 171500000 | 1.05 | 0.99 | A>A |
| chr5 | 1.72E+08 | 171600000 | 1.12 | 1.15 | A>A |
| chr5 | 1.72E+08 | 171700000 | 1.21 | 1.13 | A>A |
| chr5 | 1.72E+08 | 171800000 | 1.12 | 1.01 | A>A |
| chr5 | 1.72E+08 | 171900000 | 1.05 | 0.94 | A>A |
| chr5 | 1.72E+08 | 172000000 | 1.04 | 0.90 | A>A |
| chr5 | 1.72E+08 | 172100000 | 1.06 | 1.00 | A>A |
| chr5 | 1.72E+08 | 172200000 | 0.88 | 0.90 | A>A |
| chr5 | 1.72E+08 | 172300000 | 1.46 | 1.27 | A>A |
| chr5 | 1.72E+08 | 172400000 | 0.91 | 0.97 | A>A |
| chr5 | 1.72E+08 | 172500000 | 1.10 | 0.95 | A>A |
| chr5 | 1.73E+08 | 172600000 | 1.06 | 0.96 | A>A |
| chr5 | 1.73E+08 | 172700000 | 0.96 | 0.75 | A>A |
| chr5 | 1.73E+08 | 172800000 | 0.78 | 0.67 | A>A |
| chr5 | 1.73E+08 | 172900000 | 1.05 | 0.60 | A>A |
| chr5 | 1.73E+08 | 173000000 | 0.89 | 0.56 | A>A |
| chr5 | 1.73E+08 | 173100000 | 0.78 | 0.49 | A>A |
| chr5 | 1.73E+08 | 173200000 | 1.18 | 0.83 | A>A |
| chr5 | 1.73E+08 | 173300000 | 1.22 | 0.87 | A>A |
| chr5 | 1.73E+08 | 173400000 | 1.08 | 0.78 | A>A |
| chr5 | 1.73E+08 | 173500000 | 0.53 | 0.08 | A>A |
| chr5 | 1.74E+08 | 173600000 | 0.90 | -0.16 | A>B |
| chr5 | 1.74E+08 | 173700000 | 0.90 | -0.23 | A>B |
| chr5 | 1.74E+08 | 173800000 | 0.54 | -0.09 | A>B |
| chr5 | 1.74E+08 | 173900000 | 0.68 | 0.11 | A>A |
| chr5 | 1.74E+08 | 174000000 | 0.99 | 0.36 | A>B |
| chr5 | 1.74E+08 | 174100000 | 1.14 | 0.61 | A>A |
| chr5 | 1.74E+08 | 174200000 | 0.89 | 0.36 | A>A |
| chr5 | 1.74E+08 | 174300000 | 0.79 | -0.29 | A>B |
| chr5 | 1.74E+08 | 174400000 | 0.69 | -0.28 | A>B |
| chr5 | 1.74E+08 | 174500000 | 0.98 | -0.67 | A>B |
| chr5 | 1.75E+08 | 174600000 | 1.07 | -0.71 | A>B |
| chr5 | 1.75E+08 | 174700000 | 1.20 | -0.60 | A>B |
| chr5 | 1.75E+08 | 174800000 | 1.08 | -0.62 | A>B |
| chr5 | 1.75E+08 | 174900000 | 0.93 | -0.36 | A>B |
| chr5 | 1.75E+08 | 175000000 | 0.88 | 0.23 | A>B |

| | | | | | |
|---|---|---|---|---|---|
| chr5 | 1.75E+08 | 175100000 | 1.06 | 0.27 | A>B |
| chr5 | 1.75E+08 | 175200000 | 1.11 | 0.44 | A>B |
| chr5 | 1.75E+08 | 175300000 | 1.29 | 0.87 | A>A |
| chr5 | 1.75E+08 | 175400000 | 0.00 | 0.00 | NO |
| chr5 | 1.75E+08 | 175500000 | 0.00 | 0.00 | NO |
| chr5 | 1.76E+08 | 175600000 | 0.00 | 0.00 | NO |
| chr5 | 1.76E+08 | 175700000 | 0.00 | 0.00 | NO |
| chr5 | 1.76E+08 | 175800000 | 1.11 | 0.88 | A>A |
| chr5 | 1.76E+08 | 175900000 | 1.26 | 1.07 | A>A |
| chr5 | 1.76E+08 | 176000000 | 1.28 | 1.17 | A>A |
| chr5 | 1.76E+08 | 176100000 | 0.94 | 1.08 | A>A |
| chr5 | 1.76E+08 | 176200000 | 1.22 | 1.15 | A>A |
| chr5 | 1.76E+08 | 176300000 | 1.20 | 1.19 | A>A |
| chr5 | 1.76E+08 | 176400000 | 1.15 | 1.22 | A>A |
| chr5 | 1.76E+08 | 176500000 | 1.16 | 1.17 | A>A |
| chr5 | 1.77E+08 | 176600000 | 1.04 | 1.08 | A>A |
| chr5 | 1.77E+08 | 176700000 | 1.33 | 1.19 | A>A |
| chr5 | 1.77E+08 | 176800000 | 1.10 | 1.13 | A>A |
| chr5 | 1.77E+08 | 176900000 | 1.25 | 1.15 | A>A |
| chr5 | 1.77E+08 | 177000000 | 1.19 | 1.07 | A>A |
| chr5 | 1.77E+08 | 177100000 | 0.00 | 0.00 | NO |
| chr5 | 1.77E+08 | 177200000 | 0.00 | 0.00 | NO |
| chr5 | 1.77E+08 | 177300000 | 0.00 | 0.00 | NO |
| chr5 | 1.77E+08 | 177400000 | 0.00 | 0.00 | NO |
| chr5 | 1.77E+08 | 177500000 | 0.00 | 0.00 | NO |
| chr5 | 1.78E+08 | 177600000 | 1.15 | 1.30 | A>A |
| chr5 | 1.78E+08 | 177700000 | 0.66 | 1.09 | A>A |
| chr5 | 1.78E+08 | 177800000 | 0.02 | 1.16 | B>A |
| chr5 | 1.78E+08 | 177900000 | -0.29 | 1.37 | B>A |
| chr5 | 1.78E+08 | 178000000 | -0.25 | 1.18 | B>A |
| chr5 | 1.78E+08 | 178100000 | -0.03 | 1.29 | B>A |
| chr5 | 1.78E+08 | 178200000 | -0.19 | 1.29 | B>A |
| chr5 | 1.78E+08 | 178300000 | -0.11 | 1.17 | B>A |
| chr5 | 1.78E+08 | 178400000 | -0.38 | 1.13 | B>A |
| chr5 | 1.78E+08 | 178500000 | -0.42 | 1.19 | B>A |
| chr5 | 1.79E+08 | 178600000 | -0.32 | 1.10 | B>A |
| chr5 | 1.79E+08 | 178700000 | -0.25 | 1.04 | B>A |
| chr5 | 1.79E+08 | 178800000 | 0.05 | 1.13 | B>A |
| chr5 | 1.79E+08 | 178900000 | 0.14 | 1.12 | B>A |
| chr5 | 1.79E+08 | 179000000 | 0.51 | 0.95 | A>A |
| chr5 | 1.79E+08 | 179100000 | 0.97 | 0.88 | A>A |
| chr5 | 1.79E+08 | 179200000 | 1.19 | 1.06 | A>A |
| chr5 | 1.79E+08 | 179300000 | 1.14 | 1.04 | A>A |
| chr5 | 1.79E+08 | 179400000 | 1.07 | 1.11 | A>A |
| chr5 | 1.79E+08 | 179500000 | 1.10 | 1.29 | A>A |
| chr5 | 1.8E+08 | 179600000 | 0.85 | 1.06 | A>A |
| chr5 | 1.8E+08 | 179700000 | 1.07 | 1.18 | A>A |
| chr5 | 1.8E+08 | 179800000 | 1.17 | 1.24 | A>A |
| chr5 | 1.8E+08 | 179900000 | 1.24 | 1.30 | A>A |
| chr5 | 1.8E+08 | 180000000 | 1.18 | 1.17 | A>A |
| chr5 | 1.8E+08 | 180100000 | 1.13 | 1.20 | A>A |
| chr5 | 1.8E+08 | 180200000 | 0.90 | 0.95 | A>A |
| chr5 | 1.8E+08 | 180300000 | 0.93 | 1.08 | A>A |
| chr5 | 1.8E+08 | 180400000 | 0.76 | 0.95 | A>A |
| chr5 | 1.8E+08 | 180500000 | 1.12 | 1.03 | A>A |
| chr5 | 1.81E+08 | 180600000 | 1.12 | 1.03 | A>A |

| | | | | | |
|---|---|---|---|---|---|
| chr5 | 1.81E+08 | 180700000 | 1.34 | 1.14 | A>A |
| chr5 | 1.81E+08 | 180800000 | 0.00 | 0.00 | NO |
| chr5 | 1.81E+08 | 180900000 | 0.00 | 0.00 | NO |
| chr5 | 1.81E+08 | 180915260 | 0.00 | 0.00 | NO |
| chr6 | 0 | 100000 | 0.00 | 0.00 | NO |
| chr6 | 100000 | 200000 | 0.00 | 0.00 | NO |
| chr6 | 200000 | 300000 | 0.65 | 0.59 | A>A |
| chr6 | 300000 | 400000 | 0.84 | 0.78 | A>A |
| chr6 | 400000 | 500000 | 0.83 | 0.82 | A>A |
| chr6 | 500000 | 600000 | 0.57 | 0.61 | A>A |
| chr6 | 600000 | 700000 | 0.77 | 0.75 | A>A |
| chr6 | 700000 | 800000 | 0.68 | 0.55 | A>A |
| chr6 | 800000 | 900000 | 0.81 | 0.56 | A>A |
| chr6 | 900000 | 1000000 | 0.52 | 0.52 | A>A |
| chr6 | 1000000 | 1100000 | 0.65 | 0.73 | A>A |
| chr6 | 1100000 | 1200000 | 0.85 | 0.69 | A>A |
| chr6 | 1200000 | 1300000 | 0.82 | 0.70 | A>A |
| chr6 | 1300000 | 1400000 | 0.57 | 0.50 | A>A |
| chr6 | 1400000 | 1500000 | 0.50 | 0.58 | A>A |
| chr6 | 1500000 | 1600000 | 0.48 | 0.61 | A>A |
| chr6 | 1600000 | 1700000 | 0.44 | 0.80 | A>A |
| chr6 | 1700000 | 1800000 | 0.48 | 0.93 | A>A |
| chr6 | 1800000 | 1900000 | 0.39 | 0.92 | B>A |
| chr6 | 1900000 | 2000000 | 0.36 | 0.85 | A>A |
| chr6 | 2000000 | 2100000 | 0.38 | 0.83 | A>A |
| chr6 | 2100000 | 2200000 | 0.40 | 0.67 | A>A |
| chr6 | 2200000 | 2300000 | 0.46 | 0.81 | A>A |
| chr6 | 2300000 | 2400000 | 0.37 | 0.76 | A>A |
| chr6 | 2400000 | 2500000 | 0.52 | 0.79 | A>A |
| chr6 | 2500000 | 2600000 | 0.77 | 0.85 | A>A |
| chr6 | 2600000 | 2700000 | 0.51 | 0.68 | A>A |
| chr6 | 2700000 | 2800000 | 0.61 | 0.78 | A>A |
| chr6 | 2800000 | 2900000 | 1.12 | 1.04 | A>A |
| chr6 | 2900000 | 3000000 | 1.08 | 1.02 | A>A |
| chr6 | 3000000 | 3100000 | 0.99 | 1.00 | A>A |
| chr6 | 3100000 | 3200000 | 0.99 | 1.08 | A>A |
| chr6 | 3200000 | 3300000 | 0.74 | 0.82 | A>A |
| chr6 | 3300000 | 3400000 | 0.97 | 1.04 | A>A |
| chr6 | 3400000 | 3500000 | 0.89 | 0.90 | A>A |
| chr6 | 3500000 | 3600000 | 1.01 | 0.97 | A>A |
| chr6 | 3600000 | 3700000 | 0.96 | 1.12 | A>A |
| chr6 | 3700000 | 3800000 | 0.77 | 0.75 | A>A |
| chr6 | 3800000 | 3900000 | 1.08 | 1.11 | A>A |
| chr6 | 3900000 | 4000000 | 0.78 | 0.87 | A>A |
| chr6 | 4000000 | 4100000 | 0.83 | 0.97 | A>A |
| chr6 | 4100000 | 4200000 | 0.82 | 1.03 | A>A |
| chr6 | 4200000 | 4300000 | 0.73 | 0.75 | A>A |
| chr6 | 4300000 | 4400000 | 0.64 | 0.89 | A>A |
| chr6 | 4400000 | 4500000 | 0.71 | 0.84 | A>A |
| chr6 | 4500000 | 4600000 | 0.69 | 0.94 | A>A |
| chr6 | 4600000 | 4700000 | 0.75 | 0.91 | A>A |
| chr6 | 4700000 | 4800000 | 0.83 | 0.82 | A>A |
| chr6 | 4800000 | 4900000 | 0.81 | 0.85 | A>A |
| chr6 | 4900000 | 5000000 | 0.74 | 0.77 | A>A |
| chr6 | 5000000 | 5100000 | 0.65 | 0.63 | A>A |
| chr6 | 5100000 | 5200000 | 0.51 | 0.55 | A>A |

| | | | | | |
|---|---|---|---|---|---|
| chr6 | 5200000 | 5300000 | 0.72 | 0.60 | A>A |
| chr6 | 5300000 | 5400000 | 0.65 | 0.38 | A>A |
| chr6 | 5400000 | 5500000 | 0.29 | 0.10 | A>A |
| chr6 | 5500000 | 5600000 | 0.32 | 0.01 | A>A |
| chr6 | 5600000 | 5700000 | 0.13 | -0.05 | A>A |
| chr6 | 5700000 | 5800000 | 0.15 | -0.25 | B>B |
| chr6 | 5800000 | 5900000 | -0.05 | -0.30 | B>B |
| chr6 | 5900000 | 6000000 | -0.10 | -0.42 | B>B |
| chr6 | 6000000 | 6100000 | -0.32 | -0.64 | B>B |
| chr6 | 6100000 | 6200000 | -0.40 | -0.97 | B>B |
| chr6 | 6200000 | 6300000 | -0.39 | -1.01 | A>B |
| chr6 | 6300000 | 6400000 | -0.41 | -1.02 | A>B |
| chr6 | 6400000 | 6500000 | -0.50 | -1.01 | B>B |
| chr6 | 6500000 | 6600000 | -0.20 | -0.60 | B>B |
| chr6 | 6600000 | 6700000 | 0.27 | -0.04 | A>A |
| chr6 | 6700000 | 6800000 | 0.40 | 0.46 | A>A |
| chr6 | 6800000 | 6900000 | 0.70 | 0.64 | A>A |
| chr6 | 6900000 | 7000000 | 0.61 | 0.56 | A>A |
| chr6 | 7000000 | 7100000 | 0.68 | 0.68 | A>A |
| chr6 | 7100000 | 7200000 | 0.73 | 0.65 | A>A |
| chr6 | 7200000 | 7300000 | 0.46 | 0.50 | A>A |
| chr6 | 7300000 | 7400000 | 0.87 | 0.74 | A>A |
| chr6 | 7400000 | 7500000 | 0.87 | 0.92 | A>A |
| chr6 | 7500000 | 7600000 | 0.88 | 0.81 | A>A |
| chr6 | 7600000 | 7700000 | 0.57 | 0.52 | A>A |
| chr6 | 7700000 | 7800000 | 0.57 | 0.54 | A>A |
| chr6 | 7800000 | 7900000 | 0.47 | 0.52 | A>A |
| chr6 | 7900000 | 8000000 | 0.47 | 0.40 | A>A |
| chr6 | 8000000 | 8100000 | 0.55 | 0.47 | A>A |
| chr6 | 8100000 | 8200000 | 0.25 | 0.16 | A>A |
| chr6 | 8200000 | 8300000 | -0.06 | -0.02 | B>B |
| chr6 | 8300000 | 8400000 | -0.18 | -0.06 | B>B |
| chr6 | 8400000 | 8500000 | -0.03 | 0.08 | A>A |
| chr6 | 8500000 | 8600000 | -0.23 | 0.04 | B>B |
| chr6 | 8600000 | 8700000 | -0.29 | 0.03 | B>B |
| chr6 | 8700000 | 8800000 | -0.13 | 0.06 | B>B |
| chr6 | 8800000 | 8900000 | -0.10 | 0.07 | B>B |
| chr6 | 8900000 | 9000000 | 0.07 | 0.29 | A>A |
| chr6 | 9000000 | 9100000 | 0.21 | 0.33 | A>A |
| chr6 | 9100000 | 9200000 | 0.17 | 0.45 | A>A |
| chr6 | 9200000 | 9300000 | 0.15 | 0.31 | A>A |
| chr6 | 9300000 | 9400000 | 0.15 | 0.43 | A>A |
| chr6 | 9400000 | 9500000 | 0.30 | 0.45 | A>A |
| chr6 | 9500000 | 9600000 | 0.17 | 0.45 | A>A |
| chr6 | 9600000 | 9700000 | 0.22 | 0.38 | A>A |
| chr6 | 9700000 | 9800000 | 0.22 | 0.46 | A>A |
| chr6 | 9800000 | 9900000 | 0.21 | 0.31 | A>A |
| chr6 | 9900000 | 10000000 | 0.38 | 0.31 | A>A |
| chr6 | 10000000 | 10100000 | 0.31 | 0.47 | A>A |
| chr6 | 10100000 | 10200000 | 0.47 | 0.64 | A>A |
| chr6 | 10200000 | 10300000 | 0.48 | 0.43 | A>A |
| chr6 | 10300000 | 10400000 | 0.39 | 0.55 | A>A |
| chr6 | 10400000 | 10500000 | 0.30 | 0.49 | A>A |
| chr6 | 10500000 | 10600000 | 0.81 | 0.81 | A>A |
| chr6 | 10600000 | 10700000 | 0.83 | 0.93 | A>A |
| chr6 | 10700000 | 10800000 | 0.74 | 0.85 | A>A |

| | | | | | |
|---|---|---|---|---|---|
| chr6 | 10800000 | 10900000 | 0.73 | 0.71 | A>A |
| chr6 | 10900000 | 11000000 | 0.68 | 0.78 | A>A |
| chr6 | 11000000 | 11100000 | 0.71 | 0.75 | A>A |
| chr6 | 11100000 | 11200000 | 0.85 | 0.76 | A>A |
| chr6 | 11200000 | 11300000 | 1.01 | 0.96 | A>A |
| chr6 | 11300000 | 11400000 | 1.07 | 1.07 | A>A |
| chr6 | 11400000 | 11500000 | 0.84 | 0.82 | A>A |
| chr6 | 11500000 | 11600000 | 0.49 | 0.53 | A>A |
| chr6 | 11600000 | 11700000 | 0.46 | 0.44 | A>A |
| chr6 | 11700000 | 11800000 | 0.61 | 0.72 | A>A |
| chr6 | 11800000 | 11900000 | 0.74 | 0.71 | A>A |
| chr6 | 11900000 | 12000000 | 0.60 | 0.47 | A>A |
| chr6 | 12000000 | 12100000 | 0.58 | 0.51 | A>A |
| chr6 | 12100000 | 12200000 | 0.55 | 0.53 | A>A |
| chr6 | 12200000 | 12300000 | 0.63 | 0.46 | A>A |
| chr6 | 12300000 | 12400000 | 0.76 | 0.69 | A>A |
| chr6 | 12400000 | 12500000 | 0.88 | 0.77 | A>A |
| chr6 | 12500000 | 12600000 | 0.90 | 0.65 | A>A |
| chr6 | 12600000 | 12700000 | 0.35 | -0.01 | A>A |
| chr6 | 12700000 | 12800000 | 0.42 | -0.18 | A>B |
| chr6 | 12800000 | 12900000 | 0.33 | -0.12 | A>A |
| chr6 | 12900000 | 13000000 | 0.29 | -0.15 | A>A |
| chr6 | 13000000 | 13100000 | 0.42 | -0.13 | A>A |
| chr6 | 13100000 | 13200000 | 0.27 | -0.15 | A>A |
| chr6 | 13200000 | 13300000 | 0.50 | 0.01 | A>A |
| chr6 | 13300000 | 13400000 | 0.62 | 0.43 | A>A |
| chr6 | 13400000 | 13500000 | 0.87 | 0.67 | A>A |
| chr6 | 13500000 | 13600000 | 0.62 | 0.54 | A>A |
| chr6 | 13600000 | 13700000 | 0.75 | 0.82 | A>A |
| chr6 | 13700000 | 13800000 | 0.83 | 0.84 | A>A |
| chr6 | 13800000 | 13900000 | 0.67 | 0.85 | A>A |
| chr6 | 13900000 | 14000000 | 0.68 | 0.97 | A>A |
| chr6 | 14000000 | 14100000 | 0.65 | 0.84 | A>A |
| chr6 | 14100000 | 14200000 | 0.44 | 0.60 | A>A |
| chr6 | 14200000 | 14300000 | 0.52 | 0.83 | A>A |
| chr6 | 14300000 | 14400000 | 0.78 | 0.93 | A>A |
| chr6 | 14400000 | 14500000 | 0.65 | 1.00 | A>A |
| chr6 | 14500000 | 14600000 | 0.68 | 0.86 | A>A |
| chr6 | 14600000 | 14700000 | 0.68 | 0.87 | A>A |
| chr6 | 14700000 | 14800000 | 0.62 | 0.79 | A>A |
| chr6 | 14800000 | 14900000 | 0.66 | 0.89 | A>A |
| chr6 | 14900000 | 15000000 | 0.67 | 0.76 | A>A |
| chr6 | 15000000 | 15100000 | 0.65 | 0.82 | A>A |
| chr6 | 15100000 | 15200000 | 0.69 | 0.81 | A>A |
| chr6 | 15200000 | 15300000 | 0.70 | 0.81 | A>A |
| chr6 | 15300000 | 15400000 | 0.89 | 0.99 | A>A |
| chr6 | 15400000 | 15500000 | 0.61 | 0.81 | A>A |
| chr6 | 15500000 | 15600000 | 0.75 | 0.94 | A>A |
| chr6 | 15600000 | 15700000 | 0.61 | 0.85 | A>A |
| chr6 | 15700000 | 15800000 | 0.61 | 0.92 | A>A |
| chr6 | 15800000 | 15900000 | 0.61 | 0.90 | A>A |
| chr6 | 15900000 | 16000000 | 0.77 | 0.97 | A>A |
| chr6 | 16000000 | 16100000 | 0.68 | 0.95 | A>A |
| chr6 | 16100000 | 16200000 | 0.63 | 0.81 | A>A |
| chr6 | 16200000 | 16300000 | 0.66 | 0.78 | A>A |
| chr6 | 16300000 | 16400000 | 0.72 | 0.81 | A>A |

| | | | | | |
|---|---|---|---|---|---|
| chr6 | 16400000 | 16500000 | 0.92 | 0.94 | A>A |
| chr6 | 16500000 | 16600000 | 0.81 | 0.97 | A>A |
| chr6 | 16600000 | 16700000 | 1.00 | 1.04 | A>A |
| chr6 | 16700000 | 16800000 | 0.78 | 0.82 | A>A |
| chr6 | 16800000 | 16900000 | 0.86 | 0.79 | A>A |
| chr6 | 16900000 | 17000000 | 0.64 | 0.72 | A>A |
| chr6 | 17000000 | 17100000 | 0.55 | 0.74 | A>A |
| chr6 | 17100000 | 17200000 | 0.56 | 0.61 | A>A |
| chr6 | 17200000 | 17300000 | 0.85 | 0.88 | A>A |
| chr6 | 17300000 | 17400000 | 1.00 | 0.89 | A>A |
| chr6 | 17400000 | 17500000 | 0.94 | 1.03 | A>A |
| chr6 | 17500000 | 17600000 | 0.67 | 0.80 | A>A |
| chr6 | 17600000 | 17700000 | 0.99 | 0.94 | A>A |
| chr6 | 17700000 | 17800000 | 0.94 | 0.98 | A>A |
| chr6 | 17800000 | 17900000 | 0.79 | 0.88 | A>A |
| chr6 | 17900000 | 18000000 | 0.91 | 0.98 | A>A |
| chr6 | 18000000 | 18100000 | 0.93 | 0.94 | A>A |
| chr6 | 18100000 | 18200000 | 0.65 | 0.71 | A>A |
| chr6 | 18200000 | 18300000 | 0.60 | 0.59 | A>A |
| chr6 | 18300000 | 18400000 | 0.32 | 0.23 | A>A |
| chr6 | 18400000 | 18500000 | 0.30 | 0.35 | A>A |
| chr6 | 18500000 | 18600000 | -0.25 | -0.48 | B>B |
| chr6 | 18600000 | 18700000 | -0.69 | -0.95 | B>B |
| chr6 | 18700000 | 18800000 | -0.93 | -1.02 | B>B |
| chr6 | 18800000 | 18900000 | -0.95 | -1.03 | B>B |
| chr6 | 18900000 | 19000000 | -0.91 | -0.99 | B>B |
| chr6 | 19000000 | 19100000 | -0.87 | -1.14 | B>B |
| chr6 | 19100000 | 19200000 | -0.92 | -0.97 | B>B |
| chr6 | 19200000 | 19300000 | -0.88 | -1.02 | B>B |
| chr6 | 19300000 | 19400000 | -0.91 | -0.87 | B>B |
| chr6 | 19400000 | 19500000 | -0.81 | -0.86 | B>B |
| chr6 | 19500000 | 19600000 | -0.86 | -0.95 | B>B |
| chr6 | 19600000 | 19700000 | -0.80 | -0.96 | B>B |
| chr6 | 19700000 | 19800000 | -0.54 | -0.86 | B>B |
| chr6 | 19800000 | 19900000 | -0.43 | -0.75 | B>B |
| chr6 | 19900000 | 20000000 | -0.47 | -0.86 | B>B |
| chr6 | 20000000 | 20100000 | -0.27 | -0.54 | B>B |
| chr6 | 20100000 | 20200000 | 0.40 | 0.11 | A>A |
| chr6 | 20200000 | 20300000 | 0.37 | 0.21 | A>A |
| chr6 | 20300000 | 20400000 | 0.32 | 0.03 | A>A |
| chr6 | 20400000 | 20500000 | 0.44 | 0.30 | A>A |
| chr6 | 20500000 | 20600000 | 0.15 | 0.17 | A>A |
| chr6 | 20600000 | 20700000 | 0.49 | 0.46 | A>A |
| chr6 | 20700000 | 20800000 | 0.57 | 0.52 | A>A |
| chr6 | 20800000 | 20900000 | 0.38 | 0.41 | A>A |
| chr6 | 20900000 | 21000000 | 0.41 | 0.44 | A>A |
| chr6 | 21000000 | 21100000 | 0.46 | 0.45 | A>A |
| chr6 | 21100000 | 21200000 | 0.42 | 0.55 | A>A |
| chr6 | 21200000 | 21300000 | 0.41 | 0.45 | A>A |
| chr6 | 21300000 | 21400000 | 0.46 | 0.45 | A>A |
| chr6 | 21400000 | 21500000 | 0.36 | 0.50 | A>A |
| chr6 | 21500000 | 21600000 | 0.21 | 0.29 | A>A |
| chr6 | 21600000 | 21700000 | 0.38 | 0.42 | A>A |
| chr6 | 21700000 | 21800000 | 0.41 | 0.46 | A>A |
| chr6 | 21800000 | 21900000 | 0.51 | 0.53 | A>A |
| chr6 | 21900000 | 22000000 | 0.37 | 0.46 | A>A |

| | | | | | |
|---|---|---|---|---|---|
| chr6 | 22000000 | 22100000 | 0.35 | 0.48 | A>A |
| chr6 | 22100000 | 22200000 | 0.25 | 0.40 | A>A |
| chr6 | 22200000 | 22300000 | 0.24 | 0.33 | A>A |
| chr6 | 22300000 | 22400000 | 0.27 | 0.37 | A>A |
| chr6 | 22400000 | 22500000 | 0.24 | 0.40 | A>A |
| chr6 | 22500000 | 22600000 | 0.05 | 0.39 | A>A |
| chr6 | 22600000 | 22700000 | 0.07 | 0.40 | A>A |
| chr6 | 22700000 | 22800000 | -0.01 | 0.33 | A>A |
| chr6 | 22800000 | 22900000 | 0.02 | 0.14 | A>A |
| chr6 | 22900000 | 23000000 | -0.05 | 0.08 | A>A |
| chr6 | 23000000 | 23100000 | -0.28 | -0.10 | B>B |
| chr6 | 23100000 | 23200000 | -0.40 | -0.26 | B>B |
| chr6 | 23200000 | 23300000 | -0.48 | -0.47 | B>B |
| chr6 | 23300000 | 23400000 | -0.44 | -0.54 | B>B |
| chr6 | 23400000 | 23500000 | -0.49 | -0.63 | B>B |
| chr6 | 23500000 | 23600000 | -0.62 | -0.91 | B>B |
| chr6 | 23600000 | 23700000 | -0.61 | -1.00 | B>B |
| chr6 | 23700000 | 23800000 | -0.68 | -1.06 | B>B |
| chr6 | 23800000 | 23900000 | -0.64 | -0.97 | B>B |
| chr6 | 23900000 | 24000000 | -0.59 | -0.99 | B>B |
| chr6 | 24000000 | 24100000 | -0.42 | -0.85 | B>B |
| chr6 | 24100000 | 24200000 | -0.19 | -0.67 | B>B |
| chr6 | 24200000 | 24300000 | 0.17 | -0.09 | A>A |
| chr6 | 24300000 | 24400000 | 0.71 | 0.50 | A>A |
| chr6 | 24400000 | 24500000 | 0.75 | 0.71 | A>A |
| chr6 | 24500000 | 24600000 | 0.85 | 0.77 | A>A |
| chr6 | 24600000 | 24700000 | 0.98 | 0.98 | A>A |
| chr6 | 24700000 | 24800000 | 0.84 | 0.92 | A>A |
| chr6 | 24800000 | 24900000 | 0.72 | 0.83 | A>A |
| chr6 | 24900000 | 25000000 | 0.72 | 0.95 | A>A |
| chr6 | 25000000 | 25100000 | 0.61 | 0.82 | A>A |
| chr6 | 25100000 | 25200000 | 0.69 | 0.91 | A>A |
| chr6 | 25200000 | 25300000 | 0.63 | 0.75 | A>A |
| chr6 | 25300000 | 25400000 | 1.09 | 1.09 | A>A |
| chr6 | 25400000 | 25500000 | 0.95 | 1.03 | A>A |
| chr6 | 25500000 | 25600000 | 0.72 | 0.82 | A>A |
| chr6 | 25600000 | 25700000 | 0.75 | 0.82 | A>A |
| chr6 | 25700000 | 25800000 | 0.72 | 0.84 | A>A |
| chr6 | 25800000 | 25900000 | 0.63 | 0.89 | A>A |
| chr6 | 25900000 | 26000000 | 0.79 | 0.84 | A>A |
| chr6 | 26000000 | 26100000 | 1.00 | 0.89 | A>A |
| chr6 | 26100000 | 26200000 | 1.09 | 0.87 | A>A |
| chr6 | 26200000 | 26300000 | 1.12 | 0.80 | A>A |
| chr6 | 26300000 | 26400000 | 1.05 | 0.84 | A>A |
| chr6 | 26400000 | 26500000 | 1.30 | 1.13 | A>A |
| chr6 | 26500000 | 26600000 | 0.91 | 0.77 | A>A |
| chr6 | 26600000 | 26700000 | 0.00 | 0.00 | NO |
| chr6 | 26700000 | 26800000 | 0.00 | 0.00 | NO |
| chr6 | 26800000 | 26900000 | 0.00 | 0.00 | NO |
| chr6 | 26900000 | 27000000 | 0.00 | 0.00 | NO |
| chr6 | 27000000 | 27100000 | 0.76 | 0.93 | A>A |
| chr6 | 27100000 | 27200000 | 0.48 | 0.55 | A>A |
| chr6 | 27200000 | 27300000 | 0.60 | 0.80 | A>A |
| chr6 | 27300000 | 27400000 | 0.53 | 1.11 | B>A |
| chr6 | 27400000 | 27500000 | 0.65 | 0.90 | A>A |
| chr6 | 27500000 | 27600000 | 0.84 | 0.80 | A>A |

| | | | | | |
|---|---|---|---|---|---|
| chr6 | 27600000 | 27700000 | 0.95 | 0.86 | A>A |
| chr6 | 27700000 | 27800000 | 0.66 | 0.69 | A>A |
| chr6 | 27800000 | 27900000 | 0.80 | 0.69 | A>A |
| chr6 | 27900000 | 28000000 | 0.53 | 0.79 | A>A |
| chr6 | 28000000 | 28100000 | 0.63 | 0.88 | A>A |
| chr6 | 28100000 | 28200000 | 0.94 | 0.92 | A>A |
| chr6 | 28200000 | 28300000 | 0.87 | 1.04 | A>A |
| chr6 | 28300000 | 28400000 | 0.77 | 0.88 | A>A |
| chr6 | 28400000 | 28500000 | 0.00 | 0.00 | NO |
| chr6 | 28500000 | 28600000 | 0.00 | 0.00 | NO |
| chr6 | 28600000 | 28700000 | 0.00 | 0.00 | NO |
| chr6 | 28700000 | 28800000 | 0.00 | 0.00 | NO |
| chr6 | 28800000 | 28900000 | 0.00 | 0.00 | NO |
| chr6 | 28900000 | 29000000 | 0.00 | 0.00 | NO |
| chr6 | 29000000 | 29100000 | 0.00 | 0.00 | NO |
| chr6 | 29100000 | 29200000 | 0.00 | 0.00 | NO |
| chr6 | 29200000 | 29300000 | 0.00 | 0.00 | NO |
| chr6 | 29300000 | 29400000 | 0.00 | 0.00 | NO |
| chr6 | 29400000 | 29500000 | 0.00 | 0.00 | NO |
| chr6 | 29500000 | 29600000 | 0.00 | 0.00 | NO |
| chr6 | 29600000 | 29700000 | 0.00 | 0.00 | NO |
| chr6 | 29700000 | 29800000 | 0.00 | 0.00 | NO |
| chr6 | 29800000 | 29900000 | 0.00 | 0.00 | NO |
| chr6 | 29900000 | 30000000 | 0.00 | 0.00 | NO |
| chr6 | 30000000 | 30100000 | 0.00 | 0.00 | NO |
| chr6 | 30100000 | 30200000 | 0.00 | 0.00 | NO |
| chr6 | 30200000 | 30300000 | 0.00 | 0.00 | NO |
| chr6 | 30300000 | 30400000 | 0.00 | 0.00 | NO |
| chr6 | 30400000 | 30500000 | 0.00 | 0.00 | NO |
| chr6 | 30500000 | 30600000 | 0.00 | 0.00 | NO |
| chr6 | 30600000 | 30700000 | 0.00 | 0.00 | NO |
| chr6 | 30700000 | 30800000 | 0.00 | 0.00 | NO |
| chr6 | 30800000 | 30900000 | 0.00 | 0.00 | NO |
| chr6 | 30900000 | 31000000 | 0.00 | 0.00 | NO |
| chr6 | 31000000 | 31100000 | 0.00 | 0.00 | NO |
| chr6 | 31100000 | 31200000 | 0.00 | 0.00 | NO |
| chr6 | 31200000 | 31300000 | 0.00 | 0.00 | NO |
| chr6 | 31300000 | 31400000 | 0.00 | 0.00 | NO |
| chr6 | 31400000 | 31500000 | 0.00 | 0.00 | NO |
| chr6 | 31500000 | 31600000 | 0.00 | 0.00 | NO |
| chr6 | 31600000 | 31700000 | 0.00 | 0.00 | NO |
| chr6 | 31700000 | 31800000 | 0.00 | 0.00 | NO |
| chr6 | 31800000 | 31900000 | 0.00 | 0.00 | NO |
| chr6 | 31900000 | 32000000 | 0.00 | 0.00 | NO |
| chr6 | 32000000 | 32100000 | 0.00 | 0.00 | NO |
| chr6 | 32100000 | 32200000 | 0.00 | 0.00 | NO |
| chr6 | 32200000 | 32300000 | 0.00 | 0.00 | NO |
| chr6 | 32300000 | 32400000 | 0.00 | 0.00 | NO |
| chr6 | 32400000 | 32500000 | 0.00 | 0.00 | NO |
| chr6 | 32500000 | 32600000 | 0.00 | 0.00 | NO |
| chr6 | 32600000 | 32700000 | 0.00 | 0.00 | NO |
| chr6 | 32700000 | 32800000 | 0.00 | 0.00 | NO |
| chr6 | 32800000 | 32900000 | 0.00 | 0.00 | NO |
| chr6 | 32900000 | 33000000 | 0.00 | 0.00 | NO |
| chr6 | 33000000 | 33100000 | 0.00 | 0.00 | NO |
| chr6 | 33100000 | 33200000 | 0.00 | 0.00 | NO |

| | | | | | |
|---|---|---|---|---|---|
| chr6 | 33200000 | 33300000 | 0.00 | 0.00 | NO |
| chr6 | 33300000 | 33400000 | 0.00 | 0.00 | NO |
| chr6 | 33400000 | 33500000 | 0.00 | 0.00 | NO |
| chr6 | 33500000 | 33600000 | 0.64 | 0.57 | A>A |
| chr6 | 33600000 | 33700000 | 0.65 | 0.60 | A>A |
| chr6 | 33700000 | 33800000 | 0.63 | 0.53 | A>A |
| chr6 | 33800000 | 33900000 | 0.72 | 0.70 | A>A |
| chr6 | 33900000 | 34000000 | 0.68 | 0.59 | A>A |
| chr6 | 34000000 | 34100000 | 0.58 | 0.44 | A>A |
| chr6 | 34100000 | 34200000 | 0.44 | 0.44 | A>A |
| chr6 | 34200000 | 34300000 | 0.74 | 0.66 | A>A |
| chr6 | 34300000 | 34400000 | 0.74 | 0.66 | A>A |
| chr6 | 34400000 | 34500000 | 0.74 | 0.60 | A>A |
| chr6 | 34500000 | 34600000 | 0.86 | 0.75 | A>A |
| chr6 | 34600000 | 34700000 | 0.87 | 0.75 | A>A |
| chr6 | 34700000 | 34800000 | 0.76 | 0.72 | A>A |
| chr6 | 34800000 | 34900000 | 0.93 | 0.74 | A>A |
| chr6 | 34900000 | 35000000 | 0.90 | 0.81 | A>A |
| chr6 | 35000000 | 35100000 | 0.90 | 0.83 | A>A |
| chr6 | 35100000 | 35200000 | 0.90 | 0.75 | A>A |
| chr6 | 35200000 | 35300000 | 0.73 | 0.66 | A>A |
| chr6 | 35300000 | 35400000 | 0.94 | 0.86 | A>A |
| chr6 | 35400000 | 35500000 | 0.64 | 0.68 | A>A |
| chr6 | 35500000 | 35600000 | 0.79 | 0.79 | A>A |
| chr6 | 35600000 | 35700000 | 0.97 | 0.94 | A>A |
| chr6 | 35700000 | 35800000 | 0.72 | 0.72 | A>A |
| chr6 | 35800000 | 35900000 | 0.96 | 0.86 | A>A |
| chr6 | 35900000 | 36000000 | 0.99 | 0.98 | A>A |
| chr6 | 36000000 | 36100000 | 0.94 | 0.96 | A>A |
| chr6 | 36100000 | 36200000 | 0.98 | 0.96 | A>A |
| chr6 | 36200000 | 36300000 | 1.00 | 0.93 | A>A |
| chr6 | 36300000 | 36400000 | 0.78 | 0.86 | A>A |
| chr6 | 36400000 | 36500000 | 1.02 | 0.85 | A>A |
| chr6 | 36500000 | 36600000 | 0.94 | 0.84 | A>A |
| chr6 | 36600000 | 36700000 | 0.94 | 0.76 | A>A |
| chr6 | 36700000 | 36800000 | 0.79 | 0.83 | A>A |
| chr6 | 36800000 | 36900000 | 0.82 | 0.85 | A>A |
| chr6 | 36900000 | 37000000 | 0.69 | 0.75 | A>A |
| chr6 | 37000000 | 37100000 | 1.04 | 0.99 | A>A |
| chr6 | 37100000 | 37200000 | 0.82 | 0.80 | A>A |
| chr6 | 37200000 | 37300000 | 0.95 | 0.85 | A>A |
| chr6 | 37300000 | 37400000 | 0.83 | 0.89 | A>A |
| chr6 | 37400000 | 37500000 | 0.87 | 0.79 | A>A |
| chr6 | 37500000 | 37600000 | 0.96 | 0.99 | A>A |
| chr6 | 37600000 | 37700000 | 0.78 | 0.76 | A>A |
| chr6 | 37700000 | 37800000 | 0.84 | 0.85 | A>A |
| chr6 | 37800000 | 37900000 | 0.96 | 0.93 | A>A |
| chr6 | 37900000 | 38000000 | 0.95 | 0.83 | A>A |
| chr6 | 38000000 | 38100000 | 0.88 | 0.69 | A>A |
| chr6 | 38100000 | 38200000 | 0.67 | 0.50 | A>A |
| chr6 | 38200000 | 38300000 | 0.74 | 0.54 | A>A |
| chr6 | 38300000 | 38400000 | 0.73 | 0.55 | A>A |
| chr6 | 38400000 | 38500000 | 0.54 | 0.51 | A>A |
| chr6 | 38500000 | 38600000 | 0.70 | 0.49 | A>A |
| chr6 | 38600000 | 38700000 | 0.65 | 0.13 | A>A |
| chr6 | 38700000 | 38800000 | 0.07 | -0.42 | B>B |

| | | | | | |
|---|---|---|---|---|---|
| chr6 | 38800000 | 38900000 | -0.10 | -0.53 | B>B |
| chr6 | 38900000 | 39000000 | -0.01 | -0.54 | B>B |
| chr6 | 39000000 | 39100000 | 0.26 | -0.28 | B>B |
| chr6 | 39100000 | 39200000 | 0.54 | 0.27 | A>A |
| chr6 | 39200000 | 39300000 | 0.32 | 0.09 | A>A |
| chr6 | 39300000 | 39400000 | -0.02 | -0.24 | B>B |
| chr6 | 39400000 | 39500000 | -0.12 | -0.37 | B>B |
| chr6 | 39500000 | 39600000 | -0.17 | -0.56 | B>B |
| chr6 | 39600000 | 39700000 | -0.26 | -0.58 | B>B |
| chr6 | 39700000 | 39800000 | -0.02 | -0.59 | B>B |
| chr6 | 39800000 | 39900000 | -0.04 | -0.45 | B>B |
| chr6 | 39900000 | 40000000 | -0.24 | -0.52 | B>B |
| chr6 | 40000000 | 40100000 | -0.35 | -0.75 | B>B |
| chr6 | 40100000 | 40200000 | -0.26 | -0.78 | B>B |
| chr6 | 40200000 | 40300000 | -0.10 | -0.64 | B>B |
| chr6 | 40300000 | 40400000 | 0.16 | -0.69 | A>B |
| chr6 | 40400000 | 40500000 | 0.19 | -0.59 | A>B |
| chr6 | 40500000 | 40600000 | 0.27 | -0.34 | A>B |
| chr6 | 40600000 | 40700000 | 0.32 | -0.24 | A>A |
| chr6 | 40700000 | 40800000 | 0.39 | -0.25 | A>B |
| chr6 | 40800000 | 40900000 | 0.56 | 0.11 | A>A |
| chr6 | 40900000 | 41000000 | 0.82 | 0.39 | A>A |
| chr6 | 41000000 | 41100000 | 0.49 | 0.28 | A>A |
| chr6 | 41100000 | 41200000 | 0.57 | 0.17 | A>A |
| chr6 | 41200000 | 41300000 | 0.59 | 0.32 | A>A |
| chr6 | 41300000 | 41400000 | 0.68 | 0.47 | A>A |
| chr6 | 41400000 | 41500000 | 0.88 | 0.79 | A>A |
| chr6 | 41500000 | 41600000 | 0.82 | 0.70 | A>A |
| chr6 | 41600000 | 41700000 | 0.69 | 0.68 | A>A |
| chr6 | 41700000 | 41800000 | 0.75 | 0.68 | A>A |
| chr6 | 41800000 | 41900000 | 0.93 | 0.86 | A>A |
| chr6 | 41900000 | 42000000 | 1.02 | 0.83 | A>A |
| chr6 | 42000000 | 42100000 | 0.83 | 0.71 | A>A |
| chr6 | 42100000 | 42200000 | 0.82 | 0.76 | A>A |
| chr6 | 42200000 | 42300000 | 1.00 | 0.83 | A>A |
| chr6 | 42300000 | 42400000 | 1.18 | 1.03 | A>A |
| chr6 | 42400000 | 42500000 | 0.96 | 0.84 | A>A |
| chr6 | 42500000 | 42600000 | 0.82 | 0.76 | A>A |
| chr6 | 42600000 | 42700000 | 0.75 | 0.72 | A>A |
| chr6 | 42700000 | 42800000 | 0.96 | 0.91 | A>A |
| chr6 | 42800000 | 42900000 | 0.70 | 0.54 | A>A |
| chr6 | 42900000 | 43000000 | 0.63 | 0.64 | A>A |
| chr6 | 43000000 | 43100000 | 0.70 | 0.66 | A>A |
| chr6 | 43100000 | 43200000 | 0.63 | 0.61 | A>A |
| chr6 | 43200000 | 43300000 | 0.68 | 0.65 | A>A |
| chr6 | 43300000 | 43400000 | 0.77 | 0.68 | A>A |
| chr6 | 43400000 | 43500000 | 0.78 | 0.61 | A>A |
| chr6 | 43500000 | 43600000 | 0.71 | 0.62 | A>A |
| chr6 | 43600000 | 43700000 | 0.80 | 0.70 | A>A |
| chr6 | 43700000 | 43800000 | 0.70 | 0.68 | A>A |
| chr6 | 43800000 | 43900000 | 0.92 | 0.79 | A>A |
| chr6 | 43900000 | 44000000 | 1.02 | 0.86 | A>A |
| chr6 | 44000000 | 44100000 | 0.68 | 0.67 | A>A |
| chr6 | 44100000 | 44200000 | 0.91 | 0.81 | A>A |
| chr6 | 44200000 | 44300000 | 0.94 | 0.84 | A>A |
| chr6 | 44300000 | 44400000 | 0.97 | 0.81 | A>A |

| | | | | | |
|---|---|---|---|---|---|
| chr6 | 44400000 | 44500000 | 0.57 | 0.37 | A>A |
| chr6 | 44500000 | 44600000 | 0.33 | 0.06 | A>A |
| chr6 | 44600000 | 44700000 | 0.12 | -0.01 | A>A |
| chr6 | 44700000 | 44800000 | 0.13 | 0.12 | A>A |
| chr6 | 44800000 | 44900000 | 0.04 | 0.08 | A>A |
| chr6 | 44900000 | 45000000 | 0.07 | 0.24 | A>A |
| chr6 | 45000000 | 45100000 | 0.11 | 0.19 | A>A |
| chr6 | 45100000 | 45200000 | 0.29 | 0.31 | A>A |
| chr6 | 45200000 | 45300000 | 0.39 | 0.35 | A>A |
| chr6 | 45300000 | 45400000 | 0.38 | 0.29 | A>A |
| chr6 | 45400000 | 45500000 | 0.46 | 0.45 | A>A |
| chr6 | 45500000 | 45600000 | 0.45 | 0.45 | A>A |
| chr6 | 45600000 | 45700000 | 0.16 | 0.22 | A>A |
| chr6 | 45700000 | 45800000 | 0.02 | -0.13 | B>B |
| chr6 | 45800000 | 45900000 | -0.09 | -0.19 | B>B |
| chr6 | 45900000 | 46000000 | -0.20 | -0.23 | B>B |
| chr6 | 46000000 | 46100000 | -0.29 | -0.22 | B>B |
| chr6 | 46100000 | 46200000 | -0.32 | -0.37 | B>B |
| chr6 | 46200000 | 46300000 | -0.35 | -0.52 | B>B |
| chr6 | 46300000 | 46400000 | -0.26 | -0.44 | B>B |
| chr6 | 46400000 | 46500000 | -0.25 | -0.33 | B>B |
| chr6 | 46500000 | 46600000 | -0.25 | -0.33 | B>B |
| chr6 | 46600000 | 46700000 | -0.05 | 0.02 | B>B |
| chr6 | 46700000 | 46800000 | 0.10 | 0.11 | A>A |
| chr6 | 46800000 | 46900000 | 0.47 | 0.52 | A>A |
| chr6 | 46900000 | 47000000 | 0.48 | 0.47 | A>A |
| chr6 | 47000000 | 47100000 | 0.53 | 0.42 | A>A |
| chr6 | 47100000 | 47200000 | 0.92 | 0.76 | A>A |
| chr6 | 47200000 | 47300000 | 0.68 | 0.57 | A>A |
| chr6 | 47300000 | 47400000 | 0.53 | 0.53 | A>A |
| chr6 | 47400000 | 47500000 | 0.85 | 0.70 | A>A |
| chr6 | 47500000 | 47600000 | 0.63 | 0.46 | A>A |
| chr6 | 47600000 | 47700000 | 0.45 | 0.34 | A>A |
| chr6 | 47700000 | 47800000 | -0.02 | 0.11 | A>A |
| chr6 | 47800000 | 47900000 | -0.31 | -0.11 | B>B |
| chr6 | 47900000 | 48000000 | -0.29 | -0.04 | B>B |
| chr6 | 48000000 | 48100000 | -0.16 | 0.11 | B>B |
| chr6 | 48100000 | 48200000 | -0.13 | 0.32 | A>A |
| chr6 | 48200000 | 48300000 | -0.13 | 0.34 | A>A |
| chr6 | 48300000 | 48400000 | 0.09 | 0.36 | A>A |
| chr6 | 48400000 | 48500000 | 0.06 | 0.23 | A>A |
| chr6 | 48500000 | 48600000 | 0.01 | 0.25 | A>A |
| chr6 | 48600000 | 48700000 | 0.00 | 0.23 | A>A |
| chr6 | 48700000 | 48800000 | 0.17 | 0.11 | A>A |
| chr6 | 48800000 | 48900000 | 0.11 | 0.15 | A>A |
| chr6 | 48900000 | 49000000 | 0.10 | 0.14 | A>A |
| chr6 | 49000000 | 49100000 | -0.07 | 0.03 | B>B |
| chr6 | 49100000 | 49200000 | -0.09 | 0.00 | B>B |
| chr6 | 49200000 | 49300000 | -0.12 | 0.07 | B>B |
| chr6 | 49300000 | 49400000 | -0.08 | -0.05 | B>B |
| chr6 | 49400000 | 49500000 | 0.01 | -0.01 | A>A |
| chr6 | 49500000 | 49600000 | 0.14 | -0.03 | A>A |
| chr6 | 49600000 | 49700000 | -0.31 | -0.74 | B>B |
| chr6 | 49700000 | 49800000 | -0.31 | -0.86 | B>B |
| chr6 | 49800000 | 49900000 | -0.54 | -0.88 | B>B |
| chr6 | 49900000 | 50000000 | -0.72 | -1.23 | B>B |

| | | | | | |
|---|---|---|---|---|---|
| chr6 | 50000000 | 50100000 | -0.75 | -1.10 | B>B |
| chr6 | 50100000 | 50200000 | -0.95 | -1.14 | B>B |
| chr6 | 50200000 | 50300000 | -1.05 | -1.23 | B>B |
| chr6 | 50300000 | 50400000 | -1.09 | -1.26 | B>B |
| chr6 | 50400000 | 50500000 | -1.19 | -1.25 | B>B |
| chr6 | 50500000 | 50600000 | -1.00 | -1.11 | B>B |
| chr6 | 50600000 | 50700000 | -0.93 | -1.03 | B>B |
| chr6 | 50700000 | 50800000 | -0.71 | -0.76 | B>B |
| chr6 | 50800000 | 50900000 | -0.90 | -0.49 | B>B |
| chr6 | 50900000 | 51000000 | -1.01 | -0.20 | B>A |
| chr6 | 51000000 | 51100000 | -1.01 | -0.38 | B>A |
| chr6 | 51100000 | 51200000 | -1.00 | -0.72 | B>B |
| chr6 | 51200000 | 51300000 | -1.07 | -0.85 | B>B |
| chr6 | 51300000 | 51400000 | -0.91 | -0.95 | B>B |
| chr6 | 51400000 | 51500000 | -1.00 | -1.03 | B>B |
| chr6 | 51500000 | 51600000 | -1.00 | -1.12 | B>B |
| chr6 | 51600000 | 51700000 | -0.91 | -1.07 | B>B |
| chr6 | 51700000 | 51800000 | -0.67 | -0.94 | B>B |
| chr6 | 51800000 | 51900000 | -0.51 | -0.70 | B>B |
| chr6 | 51900000 | 52000000 | -0.33 | -0.30 | B>B |
| chr6 | 52000000 | 52100000 | 0.01 | -0.10 | B>B |
| chr6 | 52100000 | 52200000 | 0.57 | 0.33 | A>A |
| chr6 | 52200000 | 52300000 | 0.58 | 0.53 | A>A |
| chr6 | 52300000 | 52400000 | 0.82 | 0.89 | A>A |
| chr6 | 52400000 | 52500000 | 0.85 | 0.82 | A>A |
| chr6 | 52500000 | 52600000 | 0.67 | 0.63 | A>A |
| chr6 | 52600000 | 52700000 | 0.00 | 0.00 | NO |
| chr6 | 52700000 | 52800000 | 0.86 | 0.90 | A>A |
| chr6 | 52800000 | 52900000 | 1.07 | 0.97 | A>A |
| chr6 | 52900000 | 53000000 | 0.85 | 0.84 | A>A |
| chr6 | 53000000 | 53100000 | 0.78 | 0.85 | A>A |
| chr6 | 53100000 | 53200000 | 1.07 | 1.24 | A>A |
| chr6 | 53200000 | 53300000 | 0.99 | 0.99 | A>A |
| chr6 | 53300000 | 53400000 | 0.66 | 0.73 | A>A |
| chr6 | 53400000 | 53500000 | 1.07 | 1.12 | A>A |
| chr6 | 53500000 | 53600000 | 0.76 | 0.86 | A>A |
| chr6 | 53600000 | 53700000 | 0.66 | 0.89 | A>A |
| chr6 | 53700000 | 53800000 | 0.59 | 0.94 | A>A |
| chr6 | 53800000 | 53900000 | 0.10 | 0.63 | B>A |
| chr6 | 53900000 | 54000000 | 0.17 | 0.89 | B>A |
| chr6 | 54000000 | 54100000 | 0.09 | 0.92 | B>A |
| chr6 | 54100000 | 54200000 | 0.17 | 0.77 | B>A |
| chr6 | 54200000 | 54300000 | 0.07 | 0.90 | B>A |
| chr6 | 54300000 | 54400000 | -0.11 | 0.84 | B>A |
| chr6 | 54400000 | 54500000 | -0.16 | 0.76 | B>A |
| chr6 | 54500000 | 54600000 | -0.14 | 0.73 | B>A |
| chr6 | 54600000 | 54700000 | -0.08 | 0.80 | B>A |
| chr6 | 54700000 | 54800000 | 0.01 | 0.93 | B>A |
| chr6 | 54800000 | 54900000 | -0.17 | 0.85 | B>A |
| chr6 | 54900000 | 55000000 | -0.33 | 0.76 | B>A |
| chr6 | 55000000 | 55100000 | -0.29 | 0.70 | B>A |
| chr6 | 55100000 | 55200000 | -0.30 | 0.60 | B>A |
| chr6 | 55200000 | 55300000 | -0.19 | 0.70 | B>A |
| chr6 | 55300000 | 55400000 | 0.17 | 0.84 | B>A |
| chr6 | 55400000 | 55500000 | 0.26 | 0.98 | B>A |
| chr6 | 55500000 | 55600000 | 0.39 | 1.05 | B>A |

| | | | | | |
|---|---|---|---|---|---|
| chr6 | 55600000 | 55700000 | 0.37 | 1.05 | B>A |
| chr6 | 55700000 | 55800000 | 0.35 | 0.84 | A>A |
| chr6 | 55800000 | 55900000 | 0.25 | 0.63 | A>A |
| chr6 | 55900000 | 56000000 | 0.50 | 0.99 | A>A |
| chr6 | 56000000 | 56100000 | 0.62 | 0.92 | A>A |
| chr6 | 56100000 | 56200000 | 0.40 | 0.76 | A>A |
| chr6 | 56200000 | 56300000 | 0.69 | 0.86 | A>A |
| chr6 | 56300000 | 56400000 | 0.86 | 0.97 | A>A |
| chr6 | 56400000 | 56500000 | 0.74 | 0.81 | A>A |
| chr6 | 56500000 | 56600000 | 1.05 | 1.02 | A>A |
| chr6 | 56600000 | 56700000 | 1.07 | 1.05 | A>A |
| chr6 | 56700000 | 56800000 | 1.00 | 0.88 | A>A |
| chr6 | 56800000 | 56900000 | 1.01 | 0.94 | A>A |
| chr6 | 56900000 | 57000000 | 1.20 | 1.01 | A>A |
| chr6 | 57000000 | 57100000 | 1.17 | 1.00 | A>A |
| chr6 | 57100000 | 57200000 | 0.74 | 0.79 | A>A |
| chr6 | 57200000 | 57300000 | -0.20 | -0.37 | B>B |
| chr6 | 57300000 | 57400000 | -0.34 | -0.54 | B>B |
| chr6 | 57400000 | 57500000 | -0.60 | -0.67 | B>B |
| chr6 | 57500000 | 57600000 | -0.88 | -0.87 | B>B |
| chr6 | 57600000 | 57700000 | -1.72 | -1.69 | B>B |
| chr6 | 57700000 | 57800000 | 0.00 | 0.00 | NO |
| chr6 | 57800000 | 57900000 | 0.00 | 0.00 | NO |
| chr6 | 57900000 | 58000000 | 0.00 | 0.00 | NO |
| chr6 | 58000000 | 58100000 | 0.00 | 0.00 | NO |
| chr6 | 58100000 | 58200000 | 0.00 | 0.00 | NO |
| chr6 | 58200000 | 58300000 | 0.00 | 0.00 | NO |
| chr6 | 58300000 | 58400000 | 0.00 | 0.00 | NO |
| chr6 | 58400000 | 58500000 | 0.00 | 0.00 | NO |
| chr6 | 58500000 | 58600000 | 0.00 | 0.00 | NO |
| chr6 | 58600000 | 58700000 | 0.00 | 0.00 | NO |
| chr6 | 58700000 | 58800000 | 0.00 | 0.00 | NO |
| chr6 | 58800000 | 58900000 | 0.00 | 0.00 | NO |
| chr6 | 58900000 | 59000000 | 0.00 | 0.00 | NO |
| chr6 | 59000000 | 59100000 | 0.00 | 0.00 | NO |
| chr6 | 59100000 | 59200000 | 0.00 | 0.00 | NO |
| chr6 | 59200000 | 59300000 | 0.00 | 0.00 | NO |
| chr6 | 59300000 | 59400000 | 0.00 | 0.00 | NO |
| chr6 | 59400000 | 59500000 | 0.00 | 0.00 | NO |
| chr6 | 59500000 | 59600000 | 0.00 | 0.00 | NO |
| chr6 | 59600000 | 59700000 | 0.00 | 0.00 | NO |
| chr6 | 59700000 | 59800000 | 0.00 | 0.00 | NO |
| chr6 | 59800000 | 59900000 | 0.00 | 0.00 | NO |
| chr6 | 59900000 | 60000000 | 0.00 | 0.00 | NO |
| chr6 | 60000000 | 60100000 | 0.00 | 0.00 | NO |
| chr6 | 60100000 | 60200000 | 0.00 | 0.00 | NO |
| chr6 | 60200000 | 60300000 | 0.00 | 0.00 | NO |
| chr6 | 60300000 | 60400000 | 0.00 | 0.00 | NO |
| chr6 | 60400000 | 60500000 | 0.00 | 0.00 | NO |
| chr6 | 60500000 | 60600000 | 0.00 | 0.00 | NO |
| chr6 | 60600000 | 60700000 | 0.00 | 0.00 | NO |
| chr6 | 60700000 | 60800000 | 0.00 | 0.00 | NO |
| chr6 | 60800000 | 60900000 | 0.00 | 0.00 | NO |
| chr6 | 60900000 | 61000000 | 0.00 | 0.00 | NO |
| chr6 | 61000000 | 61100000 | 0.00 | 0.00 | NO |
| chr6 | 61100000 | 61200000 | 0.00 | 0.00 | NO |

| | | | | | |
|---|---|---|---|---|---|
| chr6 | 61200000 | 61300000 | 0.00 | 0.00 | NO |
| chr6 | 61300000 | 61400000 | 0.00 | 0.00 | NO |
| chr6 | 61400000 | 61500000 | 0.00 | 0.00 | NO |
| chr6 | 61500000 | 61600000 | 0.00 | 0.00 | NO |
| chr6 | 61600000 | 61700000 | 0.00 | 0.00 | NO |
| chr6 | 61700000 | 61800000 | 0.00 | 0.00 | NO |
| chr6 | 61800000 | 61900000 | 0.00 | 0.00 | NO |
| chr6 | 61900000 | 62000000 | 0.00 | 0.00 | NO |
| chr6 | 62000000 | 62100000 | 0.00 | -1.22 | NO |
| chr6 | 62100000 | 62200000 | 0.00 | 0.00 | NO |
| chr6 | 62200000 | 62300000 | -1.27 | -1.19 | B>B |
| chr6 | 62300000 | 62400000 | -1.28 | -1.26 | B>B |
| chr6 | 62400000 | 62500000 | -1.21 | -1.13 | B>B |
| chr6 | 62500000 | 62600000 | -1.16 | -1.09 | B>B |
| chr6 | 62600000 | 62700000 | -1.23 | -1.09 | B>B |
| chr6 | 62700000 | 62800000 | -1.17 | -1.07 | B>B |
| chr6 | 62800000 | 62900000 | -1.06 | -0.98 | B>B |
| chr6 | 62900000 | 63000000 | -1.02 | -0.79 | B>B |
| chr6 | 63000000 | 63100000 | -1.09 | -0.46 | B>A |
| chr6 | 63100000 | 63200000 | -1.03 | -0.26 | B>A |
| chr6 | 63200000 | 63300000 | -0.93 | 0.11 | B>A |
| chr6 | 63300000 | 63400000 | -0.82 | 0.06 | B>A |
| chr6 | 63400000 | 63500000 | -0.74 | 0.01 | B>A |
| chr6 | 63500000 | 63600000 | -0.64 | 0.12 | B>A |
| chr6 | 63600000 | 63700000 | -0.65 | -0.44 | B>B |
| chr6 | 63700000 | 63800000 | -0.57 | -0.48 | B>B |
| chr6 | 63800000 | 63900000 | -0.44 | -0.46 | B>B |
| chr6 | 63900000 | 64000000 | -0.32 | -0.16 | B>B |
| chr6 | 64000000 | 64100000 | 0.23 | 0.37 | A>A |
| chr6 | 64100000 | 64200000 | 0.93 | 0.82 | A>A |
| chr6 | 64200000 | 64300000 | 0.98 | 0.90 | A>A |
| chr6 | 64300000 | 64400000 | 0.67 | 0.46 | A>A |
| chr6 | 64400000 | 64500000 | 0.23 | 0.26 | A>A |
| chr6 | 64500000 | 64600000 | -0.35 | -0.34 | B>B |
| chr6 | 64600000 | 64700000 | -0.59 | -0.82 | B>B |
| chr6 | 64700000 | 64800000 | -0.82 | -0.95 | B>B |
| chr6 | 64800000 | 64900000 | -0.98 | -1.01 | B>B |
| chr6 | 64900000 | 65000000 | -1.01 | -0.95 | B>B |
| chr6 | 65000000 | 65100000 | -1.00 | -0.95 | B>B |
| chr6 | 65100000 | 65200000 | -1.05 | -0.90 | B>B |
| chr6 | 65200000 | 65300000 | -0.91 | -0.86 | B>B |
| chr6 | 65300000 | 65400000 | -0.97 | -0.91 | B>B |
| chr6 | 65400000 | 65500000 | -0.99 | -0.92 | B>B |
| chr6 | 65500000 | 65600000 | -1.00 | -0.87 | B>B |
| chr6 | 65600000 | 65700000 | -0.94 | -0.88 | B>B |
| chr6 | 65700000 | 65800000 | -0.93 | -0.84 | B>B |
| chr6 | 65800000 | 65900000 | -0.90 | -0.87 | B>B |
| chr6 | 65900000 | 66000000 | -0.85 | -0.84 | B>B |
| chr6 | 66000000 | 66100000 | -0.75 | -0.85 | B>B |
| chr6 | 66100000 | 66200000 | -0.70 | -0.78 | B>B |
| chr6 | 66200000 | 66300000 | -0.49 | -0.63 | B>B |
| chr6 | 66300000 | 66400000 | -0.26 | -0.47 | B>B |
| chr6 | 66400000 | 66500000 | -0.09 | -0.36 | B>B |
| chr6 | 66500000 | 66600000 | -0.13 | -0.28 | B>B |
| chr6 | 66600000 | 66700000 | -0.10 | -0.32 | B>B |
| chr6 | 66700000 | 66800000 | -0.29 | -0.49 | B>B |

| | | | | | |
|---|---|---|---|---|---|
| chr6 | 66800000 | 66900000 | -0.59 | -0.70 | B>B |
| chr6 | 66900000 | 67000000 | -0.65 | -0.77 | B>B |
| chr6 | 67000000 | 67100000 | -0.75 | -0.77 | B>B |
| chr6 | 67100000 | 67200000 | -0.85 | -0.82 | B>B |
| chr6 | 67200000 | 67300000 | -0.82 | -0.77 | B>B |
| chr6 | 67300000 | 67400000 | -0.89 | -0.80 | B>B |
| chr6 | 67400000 | 67500000 | -0.88 | -0.79 | B>B |
| chr6 | 67500000 | 67600000 | -0.93 | -0.82 | B>B |
| chr6 | 67600000 | 67700000 | -0.95 | -0.76 | B>B |
| chr6 | 67700000 | 67800000 | -0.90 | -0.58 | B>B |
| chr6 | 67800000 | 67900000 | -0.93 | -0.29 | B>A |
| chr6 | 67900000 | 68000000 | -0.96 | -0.50 | B>B |
| chr6 | 68000000 | 68100000 | -0.99 | -0.55 | B>B |
| chr6 | 68100000 | 68200000 | -0.92 | -0.66 | B>B |
| chr6 | 68200000 | 68300000 | -0.96 | -0.75 | B>B |
| chr6 | 68300000 | 68400000 | -1.01 | -0.73 | B>B |
| chr6 | 68400000 | 68500000 | -0.98 | -0.81 | B>B |
| chr6 | 68500000 | 68600000 | -0.98 | -0.79 | B>B |
| chr6 | 68600000 | 68700000 | -0.99 | -0.85 | B>B |
| chr6 | 68700000 | 68800000 | -0.99 | -0.80 | B>B |
| chr6 | 68800000 | 68900000 | -0.98 | -0.86 | B>B |
| chr6 | 68900000 | 69000000 | -1.02 | -0.86 | B>B |
| chr6 | 69000000 | 69100000 | -1.06 | -0.87 | B>B |
| chr6 | 69100000 | 69200000 | -1.06 | -0.88 | B>B |
| chr6 | 69200000 | 69300000 | -0.99 | -0.85 | B>B |
| chr6 | 69300000 | 69400000 | -0.94 | -0.77 | B>B |
| chr6 | 69400000 | 69500000 | -1.04 | -0.94 | B>B |
| chr6 | 69500000 | 69600000 | -1.03 | -0.91 | B>B |
| chr6 | 69600000 | 69700000 | -0.91 | -0.95 | B>B |
| chr6 | 69700000 | 69800000 | -0.82 | -0.95 | B>B |
| chr6 | 69800000 | 69900000 | -0.85 | -0.93 | B>B |
| chr6 | 69900000 | 70000000 | -0.97 | -0.97 | B>B |
| chr6 | 70000000 | 70100000 | -1.02 | -1.01 | B>B |
| chr6 | 70100000 | 70200000 | -1.04 | -0.98 | B>B |
| chr6 | 70200000 | 70300000 | -1.07 | -0.95 | B>B |
| chr6 | 70300000 | 70400000 | -0.69 | -0.79 | B>B |
| chr6 | 70400000 | 70500000 | -0.25 | -0.38 | B>B |
| chr6 | 70500000 | 70600000 | -0.38 | -0.75 | B>B |
| chr6 | 70600000 | 70700000 | -0.67 | -1.01 | B>B |
| chr6 | 70700000 | 70800000 | -0.66 | -1.01 | B>B |
| chr6 | 70800000 | 70900000 | -0.55 | -0.88 | B>B |
| chr6 | 70900000 | 71000000 | -0.15 | -0.63 | B>B |
| chr6 | 71000000 | 71100000 | 0.26 | -0.14 | A>A |
| chr6 | 71100000 | 71200000 | 0.67 | 0.34 | A>A |
| chr6 | 71200000 | 71300000 | 0.52 | 0.21 | A>A |
| chr6 | 71300000 | 71400000 | 0.57 | 0.13 | A>A |
| chr6 | 71400000 | 71500000 | 0.28 | 0.12 | A>A |
| chr6 | 71500000 | 71600000 | 0.12 | -0.06 | A>A |
| chr6 | 71600000 | 71700000 | -0.22 | -0.50 | B>B |
| chr6 | 71700000 | 71800000 | -0.23 | -0.61 | B>B |
| chr6 | 71800000 | 71900000 | -0.27 | -0.58 | B>B |
| chr6 | 71900000 | 72000000 | 0.22 | -0.18 | A>A |
| chr6 | 72000000 | 72100000 | 0.31 | 0.06 | A>A |
| chr6 | 72100000 | 72200000 | 0.54 | 0.29 | A>A |
| chr6 | 72200000 | 72300000 | -0.13 | -0.47 | B>B |
| chr6 | 72300000 | 72400000 | -0.51 | -0.90 | B>B |

| | | | | | |
|---|---|---|---|---|---|
| chr6 | 72400000 | 72500000 | -0.61 | -0.98 | B>B |
| chr6 | 72500000 | 72600000 | -0.70 | -1.01 | B>B |
| chr6 | 72600000 | 72700000 | -0.66 | -1.07 | B>B |
| chr6 | 72700000 | 72800000 | -0.69 | -0.98 | B>B |
| chr6 | 72800000 | 72900000 | -0.62 | -1.02 | B>B |
| chr6 | 72900000 | 73000000 | -0.52 | -1.15 | A>B |
| chr6 | 73000000 | 73100000 | -0.53 | -1.14 | A>B |
| chr6 | 73100000 | 73200000 | -0.33 | -1.02 | A>B |
| chr6 | 73200000 | 73300000 | -0.20 | -1.07 | A>B |
| chr6 | 73300000 | 73400000 | 0.01 | -1.03 | A>B |
| chr6 | 73400000 | 73500000 | 0.03 | -1.11 | A>B |
| chr6 | 73500000 | 73600000 | 0.04 | -1.15 | A>B |
| chr6 | 73600000 | 73700000 | 0.14 | -1.11 | A>B |
| chr6 | 73700000 | 73800000 | 0.18 | -1.08 | A>B |
| chr6 | 73800000 | 73900000 | 0.03 | -0.82 | A>B |
| chr6 | 73900000 | 74000000 | 0.54 | -0.16 | A>B |
| chr6 | 74000000 | 74100000 | 0.81 | 0.22 | A>B |
| chr6 | 74100000 | 74200000 | 1.23 | 0.73 | A>A |
| chr6 | 74200000 | 74300000 | 1.61 | 1.29 | A>A |
| chr6 | 74300000 | 74400000 | 0.94 | 0.93 | A>A |
| chr6 | 74400000 | 74500000 | 0.76 | 0.72 | A>A |
| chr6 | 74500000 | 74600000 | 0.37 | 0.45 | A>A |
| chr6 | 74600000 | 74700000 | -0.07 | 0.16 | A>A |
| chr6 | 74700000 | 74800000 | -0.11 | 0.13 | A>A |
| chr6 | 74800000 | 74900000 | -0.28 | 0.21 | B>B |
| chr6 | 74900000 | 75000000 | -0.29 | 0.40 | B>A |
| chr6 | 75000000 | 75100000 | -0.15 | 0.33 | A>A |
| chr6 | 75100000 | 75200000 | 0.30 | 0.35 | A>A |
| chr6 | 75200000 | 75300000 | 0.40 | 0.52 | A>A |
| chr6 | 75300000 | 75400000 | 0.26 | 0.31 | A>A |
| chr6 | 75400000 | 75500000 | 0.09 | 0.16 | A>A |
| chr6 | 75500000 | 75600000 | -0.11 | 0.00 | B>B |
| chr6 | 75600000 | 75700000 | -0.24 | -0.06 | B>B |
| chr6 | 75700000 | 75800000 | -0.14 | 0.06 | B>B |
| chr6 | 75800000 | 75900000 | 0.03 | 0.16 | A>A |
| chr6 | 75900000 | 76000000 | 0.31 | 0.40 | A>A |
| chr6 | 76000000 | 76100000 | 0.93 | 0.75 | A>A |
| chr6 | 76100000 | 76200000 | 1.31 | 1.10 | A>A |
| chr6 | 76200000 | 76300000 | 0.84 | 0.78 | A>A |
| chr6 | 76300000 | 76400000 | 1.06 | 0.92 | A>A |
| chr6 | 76400000 | 76500000 | 0.84 | 0.71 | A>A |
| chr6 | 76500000 | 76600000 | 0.27 | 0.25 | A>A |
| chr6 | 76600000 | 76700000 | -0.41 | -0.47 | B>B |
| chr6 | 76700000 | 76800000 | -0.71 | -0.98 | B>B |
| chr6 | 76800000 | 76900000 | -0.80 | -1.06 | B>B |
| chr6 | 76900000 | 77000000 | -0.99 | -1.03 | B>B |
| chr6 | 77000000 | 77100000 | -0.94 | -1.02 | B>B |
| chr6 | 77100000 | 77200000 | -0.87 | -1.05 | B>B |
| chr6 | 77200000 | 77300000 | -0.87 | -1.04 | B>B |
| chr6 | 77300000 | 77400000 | -0.57 | -0.79 | B>B |
| chr6 | 77400000 | 77500000 | -0.31 | -0.49 | B>B |
| chr6 | 77500000 | 77600000 | -0.18 | -0.32 | B>B |
| chr6 | 77600000 | 77700000 | -0.06 | -0.23 | B>B |
| chr6 | 77700000 | 77800000 | 0.26 | -0.09 | A>A |
| chr6 | 77800000 | 77900000 | 0.23 | -0.02 | A>A |
| chr6 | 77900000 | 78000000 | 0.32 | -0.02 | A>A |

| | | | | | |
|---|---|---|---|---|---|
| chr6 | 78000000 | 78100000 | 0.28 | 0.14 | A>A |
| chr6 | 78100000 | 78200000 | -0.01 | -0.20 | B>B |
| chr6 | 78200000 | 78300000 | -0.42 | -0.65 | B>B |
| chr6 | 78300000 | 78400000 | -0.52 | -0.74 | B>B |
| chr6 | 78400000 | 78500000 | -0.68 | -0.65 | B>B |
| chr6 | 78500000 | 78600000 | -0.81 | -0.75 | B>B |
| chr6 | 78600000 | 78700000 | -0.88 | -0.80 | B>B |
| chr6 | 78700000 | 78800000 | -0.87 | -0.76 | B>B |
| chr6 | 78800000 | 78900000 | -0.87 | -0.66 | B>B |
| chr6 | 78900000 | 79000000 | -0.75 | -0.46 | B>B |
| chr6 | 79000000 | 79100000 | -0.72 | -0.44 | B>B |
| chr6 | 79100000 | 79200000 | -0.59 | -0.19 | B>B |
| chr6 | 79200000 | 79300000 | -0.42 | -0.04 | B>B |
| chr6 | 79300000 | 79400000 | -0.22 | 0.06 | B>B |
| chr6 | 79400000 | 79500000 | -0.52 | -0.12 | B>B |
| chr6 | 79500000 | 79600000 | -0.25 | -0.15 | B>B |
| chr6 | 79600000 | 79700000 | 0.10 | 0.30 | A>A |
| chr6 | 79700000 | 79800000 | 0.47 | 0.54 | A>A |
| chr6 | 79800000 | 79900000 | 0.17 | 0.57 | A>A |
| chr6 | 79900000 | 80000000 | 0.06 | 0.48 | A>A |
| chr6 | 80000000 | 80100000 | -0.12 | 0.34 | A>A |
| chr6 | 80100000 | 80200000 | -0.03 | 0.39 | A>A |
| chr6 | 80200000 | 80300000 | 0.23 | 0.41 | A>A |
| chr6 | 80300000 | 80400000 | 0.29 | 0.46 | A>A |
| chr6 | 80400000 | 80500000 | -0.18 | 0.01 | B>B |
| chr6 | 80500000 | 80600000 | -0.29 | -0.11 | B>B |
| chr6 | 80600000 | 80700000 | -0.03 | -0.02 | B>B |
| chr6 | 80700000 | 80800000 | 0.38 | 0.25 | A>A |
| chr6 | 80800000 | 80900000 | 0.07 | 0.23 | A>A |
| chr6 | 80900000 | 81000000 | -0.09 | 0.15 | A>A |
| chr6 | 81000000 | 81100000 | -0.35 | 0.10 | B>B |
| chr6 | 81100000 | 81200000 | -0.52 | 0.18 | B>A |
| chr6 | 81200000 | 81300000 | -0.43 | 0.11 | B>A |
| chr6 | 81300000 | 81400000 | -0.55 | 0.19 | B>A |
| chr6 | 81400000 | 81500000 | -0.78 | 0.12 | B>A |
| chr6 | 81500000 | 81600000 | -0.87 | -0.24 | B>A |
| chr6 | 81600000 | 81700000 | -0.96 | -0.32 | B>A |
| chr6 | 81700000 | 81800000 | -0.91 | -0.30 | B>A |
| chr6 | 81800000 | 81900000 | -0.85 | -0.32 | B>A |
| chr6 | 81900000 | 82000000 | -0.83 | -0.23 | B>A |
| chr6 | 82000000 | 82100000 | -0.84 | -0.21 | B>A |
| chr6 | 82100000 | 82200000 | -0.68 | -0.06 | B>A |
| chr6 | 82200000 | 82300000 | -0.60 | 0.17 | B>A |
| chr6 | 82300000 | 82400000 | -0.34 | 0.36 | B>A |
| chr6 | 82400000 | 82500000 | -0.05 | 0.40 | A>A |
| chr6 | 82500000 | 82600000 | 0.18 | 0.71 | B>A |
| chr6 | 82600000 | 82700000 | 0.23 | 0.56 | A>A |
| chr6 | 82700000 | 82800000 | 0.13 | 0.42 | A>A |
| chr6 | 82800000 | 82900000 | 0.26 | 0.62 | A>A |
| chr6 | 82900000 | 83000000 | 0.47 | 0.66 | A>A |
| chr6 | 83000000 | 83100000 | 0.72 | 0.90 | A>A |
| chr6 | 83100000 | 83200000 | 0.94 | 1.06 | A>A |
| chr6 | 83200000 | 83300000 | 0.66 | 0.91 | A>A |
| chr6 | 83300000 | 83400000 | 0.35 | 0.83 | A>A |
| chr6 | 83400000 | 83500000 | 0.44 | 0.92 | A>A |
| chr6 | 83500000 | 83600000 | 0.48 | 0.84 | A>A |

| | | | | | |
|---|---|---|---|---|---|
| chr6 | 83600000 | 83700000 | 0.51 | 1.02 | A>A |
| chr6 | 83700000 | 83800000 | 0.39 | 0.66 | A>A |
| chr6 | 83800000 | 83900000 | 0.66 | 0.72 | A>A |
| chr6 | 83900000 | 84000000 | 0.54 | 0.69 | A>A |
| chr6 | 84000000 | 84100000 | 0.63 | 0.76 | A>A |
| chr6 | 84100000 | 84200000 | 0.67 | 0.76 | A>A |
| chr6 | 84200000 | 84300000 | 0.18 | 0.35 | A>A |
| chr6 | 84300000 | 84400000 | -0.25 | 0.14 | B>B |
| chr6 | 84400000 | 84500000 | -0.20 | 0.27 | A>A |
| chr6 | 84500000 | 84600000 | 0.09 | 0.34 | A>A |
| chr6 | 84600000 | 84700000 | 0.29 | 0.61 | A>A |
| chr6 | 84700000 | 84800000 | 0.46 | 0.77 | A>A |
| chr6 | 84800000 | 84900000 | 0.26 | 0.67 | A>A |
| chr6 | 84900000 | 85000000 | -0.16 | 0.24 | A>A |
| chr6 | 85000000 | 85100000 | -0.33 | 0.18 | B>A |
| chr6 | 85100000 | 85200000 | -0.04 | 0.45 | A>A |
| chr6 | 85200000 | 85300000 | 0.14 | 0.61 | A>A |
| chr6 | 85300000 | 85400000 | 0.21 | 0.63 | A>A |
| chr6 | 85400000 | 85500000 | 0.25 | 0.78 | B>A |
| chr6 | 85500000 | 85600000 | 0.19 | 0.89 | B>A |
| chr6 | 85600000 | 85700000 | 0.14 | 0.88 | B>A |
| chr6 | 85700000 | 85800000 | -0.02 | 0.90 | B>A |
| chr6 | 85800000 | 85900000 | -0.07 | 0.75 | B>A |
| chr6 | 85900000 | 86000000 | -0.12 | 0.72 | B>A |
| chr6 | 86000000 | 86100000 | 0.29 | 0.64 | A>A |
| chr6 | 86100000 | 86200000 | 1.02 | 1.11 | A>A |
| chr6 | 86200000 | 86300000 | 0.61 | 0.74 | A>A |
| chr6 | 86300000 | 86400000 | 0.67 | 0.57 | A>A |
| chr6 | 86400000 | 86500000 | -0.14 | -0.26 | B>B |
| chr6 | 86500000 | 86600000 | -0.51 | -0.70 | B>B |
| chr6 | 86600000 | 86700000 | -0.66 | -0.88 | B>B |
| chr6 | 86700000 | 86800000 | -0.81 | -1.00 | B>B |
| chr6 | 86800000 | 86900000 | -0.88 | -1.05 | B>B |
| chr6 | 86900000 | 87000000 | -0.94 | -1.04 | B>B |
| chr6 | 87000000 | 87100000 | -0.95 | -1.06 | B>B |
| chr6 | 87100000 | 87200000 | -0.95 | -0.98 | B>B |
| chr6 | 87200000 | 87300000 | -0.92 | -1.02 | B>B |
| chr6 | 87300000 | 87400000 | -0.80 | -0.94 | B>B |
| chr6 | 87400000 | 87500000 | -0.74 | -0.82 | B>B |
| chr6 | 87500000 | 87600000 | -0.79 | -0.86 | B>B |
| chr6 | 87600000 | 87700000 | -0.64 | -0.92 | B>B |
| chr6 | 87700000 | 87800000 | -0.33 | -0.59 | B>B |
| chr6 | 87800000 | 87900000 | 0.30 | -0.05 | A>A |
| chr6 | 87900000 | 88000000 | 0.40 | 0.13 | A>A |
| chr6 | 88000000 | 88100000 | 0.27 | 0.05 | A>A |
| chr6 | 88100000 | 88200000 | 0.43 | 0.25 | A>A |
| chr6 | 88200000 | 88300000 | 0.80 | 0.54 | A>A |
| chr6 | 88300000 | 88400000 | 0.85 | 0.75 | A>A |
| chr6 | 88400000 | 88500000 | 1.20 | 0.83 | A>A |
| chr6 | 88500000 | 88600000 | 0.54 | 0.05 | A>A |
| chr6 | 88600000 | 88700000 | 0.15 | -0.22 | B>B |
| chr6 | 88700000 | 88800000 | -0.06 | -0.29 | B>B |
| chr6 | 88800000 | 88900000 | -0.09 | -0.37 | B>B |
| chr6 | 88900000 | 89000000 | -0.18 | -0.44 | B>B |
| chr6 | 89000000 | 89100000 | -0.17 | -0.44 | B>B |
| chr6 | 89100000 | 89200000 | -0.13 | -0.38 | B>B |

| | | | | | |
|---|---|---|---|---|---|
| chr6 | 89200000 | 89300000 | 0.04 | -0.16 | B>B |
| chr6 | 89300000 | 89400000 | 0.43 | 0.14 | A>A |
| chr6 | 89400000 | 89500000 | 0.57 | 0.32 | A>A |
| chr6 | 89500000 | 89600000 | 0.52 | 0.32 | A>A |
| chr6 | 89600000 | 89700000 | 0.93 | 0.67 | A>A |
| chr6 | 89700000 | 89800000 | 1.42 | 1.18 | A>A |
| chr6 | 89800000 | 89900000 | 1.34 | 1.12 | A>A |
| chr6 | 89900000 | 90000000 | 1.39 | 1.00 | A>A |
| chr6 | 90000000 | 90100000 | 1.21 | 1.03 | A>A |
| chr6 | 90100000 | 90200000 | 1.24 | 0.84 | A>A |
| chr6 | 90200000 | 90300000 | 1.32 | 0.93 | A>A |
| chr6 | 90300000 | 90400000 | 1.24 | 0.87 | A>A |
| chr6 | 90400000 | 90500000 | 1.45 | 0.99 | A>A |
| chr6 | 90500000 | 90600000 | 1.11 | 0.59 | A>A |
| chr6 | 90600000 | 90700000 | 0.32 | -0.13 | A>A |
| chr6 | 90700000 | 90800000 | 0.15 | -0.32 | B>B |
| chr6 | 90800000 | 90900000 | 0.38 | -0.09 | A>A |
| chr6 | 90900000 | 91000000 | 0.53 | 0.11 | A>A |
| chr6 | 91000000 | 91100000 | 0.60 | 0.00 | A>B |
| chr6 | 91100000 | 91200000 | 0.26 | -0.10 | A>A |
| chr6 | 91200000 | 91300000 | 0.21 | -0.08 | A>A |
| chr6 | 91300000 | 91400000 | -0.29 | -0.83 | B>B |
| chr6 | 91400000 | 91500000 | -0.63 | -1.05 | B>B |
| chr6 | 91500000 | 91600000 | -0.70 | -1.08 | B>B |
| chr6 | 91600000 | 91700000 | -0.76 | -1.08 | B>B |
| chr6 | 91700000 | 91800000 | -0.77 | -1.01 | B>B |
| chr6 | 91800000 | 91900000 | -0.85 | -1.02 | B>B |
| chr6 | 91900000 | 92000000 | -1.00 | -1.03 | B>B |
| chr6 | 92000000 | 92100000 | -0.95 | -0.99 | B>B |
| chr6 | 92100000 | 92200000 | -0.94 | -1.01 | B>B |
| chr6 | 92200000 | 92300000 | -0.92 | -0.99 | B>B |
| chr6 | 92300000 | 92400000 | -0.97 | -1.03 | B>B |
| chr6 | 92400000 | 92500000 | -1.00 | -0.98 | B>B |
| chr6 | 92500000 | 92600000 | -0.98 | -0.98 | B>B |
| chr6 | 92600000 | 92700000 | -0.93 | -1.02 | B>B |
| chr6 | 92700000 | 92800000 | -0.97 | -1.03 | B>B |
| chr6 | 92800000 | 92900000 | -0.99 | -0.97 | B>B |
| chr6 | 92900000 | 93000000 | -0.99 | -0.97 | B>B |
| chr6 | 93000000 | 93100000 | -0.98 | -0.96 | B>B |
| chr6 | 93100000 | 93200000 | -0.97 | -0.93 | B>B |
| chr6 | 93200000 | 93300000 | -0.97 | -0.98 | B>B |
| chr6 | 93300000 | 93400000 | -0.94 | -0.93 | B>B |
| chr6 | 93400000 | 93500000 | -0.93 | -0.94 | B>B |
| chr6 | 93500000 | 93600000 | -0.92 | -0.94 | B>B |
| chr6 | 93600000 | 93700000 | -1.00 | -0.99 | B>B |
| chr6 | 93700000 | 93800000 | -0.95 | -0.91 | B>B |
| chr6 | 93800000 | 93900000 | -0.97 | -0.95 | B>B |
| chr6 | 93900000 | 94000000 | -0.95 | -0.95 | B>B |
| chr6 | 94000000 | 94100000 | -0.98 | -0.94 | B>B |
| chr6 | 94100000 | 94200000 | -0.88 | -0.78 | B>B |
| chr6 | 94200000 | 94300000 | -1.01 | -0.82 | B>B |
| chr6 | 94300000 | 94400000 | -0.99 | -0.70 | B>B |
| chr6 | 94400000 | 94500000 | -0.94 | -0.60 | B>B |
| chr6 | 94500000 | 94600000 | -0.98 | -0.55 | B>B |
| chr6 | 94600000 | 94700000 | -0.99 | -0.61 | B>B |
| chr6 | 94700000 | 94800000 | -1.04 | -0.68 | B>B |

| | | | | | |
|---|---|---|---|---|---|
| chr6 | 94800000 | 94900000 | -1.03 | -0.74 | B>B |
| chr6 | 94900000 | 95000000 | -1.05 | -0.84 | B>B |
| chr6 | 95000000 | 95100000 | -1.10 | -0.96 | B>B |
| chr6 | 95100000 | 95200000 | -1.03 | -0.98 | B>B |
| chr6 | 95200000 | 95300000 | -1.08 | -1.01 | B>B |
| chr6 | 95300000 | 95400000 | -0.99 | -0.98 | B>B |
| chr6 | 95400000 | 95500000 | -1.08 | -1.03 | B>B |
| chr6 | 95500000 | 95600000 | -1.15 | -1.00 | B>B |
| chr6 | 95600000 | 95700000 | -1.07 | -1.01 | B>B |
| chr6 | 95700000 | 95800000 | 0.00 | 0.00 | NO |
| chr6 | 95800000 | 95900000 | -0.82 | -0.84 | B>B |
| chr6 | 95900000 | 96000000 | -0.72 | -0.80 | B>B |
| chr6 | 96000000 | 96100000 | -0.44 | -0.47 | B>B |
| chr6 | 96100000 | 96200000 | -0.85 | -0.87 | B>B |
| chr6 | 96200000 | 96300000 | -0.96 | -0.90 | B>B |
| chr6 | 96300000 | 96400000 | -0.99 | -0.93 | B>B |
| chr6 | 96400000 | 96500000 | -0.97 | -0.99 | B>B |
| chr6 | 96500000 | 96600000 | -1.00 | -0.95 | B>B |
| chr6 | 96600000 | 96700000 | -1.00 | -0.97 | B>B |
| chr6 | 96700000 | 96800000 | -0.97 | -0.98 | B>B |
| chr6 | 96800000 | 96900000 | -0.88 | -0.93 | B>B |
| chr6 | 96900000 | 97000000 | -0.42 | -0.59 | B>B |
| chr6 | 97000000 | 97100000 | -0.41 | -0.75 | B>B |
| chr6 | 97100000 | 97200000 | -0.42 | -0.91 | B>B |
| chr6 | 97200000 | 97300000 | -0.03 | -0.65 | A>B |
| chr6 | 97300000 | 97400000 | -0.11 | -0.41 | B>B |
| chr6 | 97400000 | 97500000 | -0.35 | -0.73 | B>B |
| chr6 | 97500000 | 97600000 | -0.26 | -0.46 | B>B |
| chr6 | 97600000 | 97700000 | 0.22 | -0.07 | A>A |
| chr6 | 97700000 | 97800000 | 0.12 | -0.35 | B>B |
| chr6 | 97800000 | 97900000 | -0.41 | -0.71 | B>B |
| chr6 | 97900000 | 98000000 | -0.67 | -0.86 | B>B |
| chr6 | 98000000 | 98100000 | -0.85 | -0.94 | B>B |
| chr6 | 98100000 | 98200000 | -0.94 | -0.95 | B>B |
| chr6 | 98200000 | 98300000 | -0.86 | -0.92 | B>B |
| chr6 | 98300000 | 98400000 | -0.94 | -0.97 | B>B |
| chr6 | 98400000 | 98500000 | -0.90 | -0.91 | B>B |
| chr6 | 98500000 | 98600000 | -0.80 | -0.98 | B>B |
| chr6 | 98600000 | 98700000 | -0.88 | -0.90 | B>B |
| chr6 | 98700000 | 98800000 | -0.86 | -0.94 | B>B |
| chr6 | 98800000 | 98900000 | -0.84 | -0.94 | B>B |
| chr6 | 98900000 | 99000000 | -0.75 | -0.93 | B>B |
| chr6 | 99000000 | 99100000 | -0.65 | -0.95 | B>B |
| chr6 | 99100000 | 99200000 | -0.59 | -0.91 | B>B |
| chr6 | 99200000 | 99300000 | -0.35 | -0.57 | B>B |
| chr6 | 99300000 | 99400000 | 0.24 | 0.04 | A>A |
| chr6 | 99400000 | 99500000 | -0.04 | -0.78 | A>B |
| chr6 | 99500000 | 99600000 | -0.36 | -0.88 | B>B |
| chr6 | 99600000 | 99700000 | -0.33 | -0.78 | B>B |
| chr6 | 99700000 | 99800000 | 0.39 | -0.08 | A>A |
| chr6 | 99800000 | 99900000 | 0.92 | 0.48 | A>A |
| chr6 | 99900000 | 100000000 | 0.69 | 0.47 | A>A |
| chr6 | 1E+08 | 100100000 | 0.23 | -0.23 | A>A |
| chr6 | 1E+08 | 100200000 | -0.20 | -0.72 | B>B |
| chr6 | 1E+08 | 100300000 | -0.21 | -0.61 | B>B |
| chr6 | 1E+08 | 100400000 | -0.38 | -0.68 | B>B |

| | | | | | |
|---|---|---|---|---|---|
| chr6 | 1E+08 | 100500000 | -0.37 | -0.62 | B>B |
| chr6 | 1.01E+08 | 100600000 | 0.06 | -0.10 | B>B |
| chr6 | 1.01E+08 | 100700000 | 0.90 | 0.69 | A>A |
| chr6 | 1.01E+08 | 100800000 | 1.02 | 0.91 | A>A |
| chr6 | 1.01E+08 | 100900000 | 1.07 | 0.78 | A>A |
| chr6 | 1.01E+08 | 101000000 | 0.47 | 0.42 | A>A |
| chr6 | 1.01E+08 | 101100000 | 0.34 | 0.29 | A>A |
| chr6 | 1.01E+08 | 101200000 | 0.10 | 0.05 | A>A |
| chr6 | 1.01E+08 | 101300000 | 0.37 | 0.14 | A>A |
| chr6 | 1.01E+08 | 101400000 | -0.04 | 0.06 | A>A |
| chr6 | 1.01E+08 | 101500000 | -0.30 | -0.33 | B>B |
| chr6 | 1.02E+08 | 101600000 | -0.23 | -0.45 | B>B |
| chr6 | 1.02E+08 | 101700000 | -0.44 | -0.60 | B>B |
| chr6 | 1.02E+08 | 101800000 | -0.62 | -0.89 | B>B |
| chr6 | 1.02E+08 | 101900000 | -0.66 | -0.95 | B>B |
| chr6 | 1.02E+08 | 102000000 | -0.83 | -0.96 | B>B |
| chr6 | 1.02E+08 | 102100000 | -0.90 | -1.02 | B>B |
| chr6 | 1.02E+08 | 102200000 | -0.92 | -1.01 | B>B |
| chr6 | 1.02E+08 | 102300000 | -0.91 | -0.98 | B>B |
| chr6 | 1.02E+08 | 102400000 | -0.98 | -1.03 | B>B |
| chr6 | 1.02E+08 | 102500000 | -0.98 | -1.01 | B>B |
| chr6 | 1.03E+08 | 102600000 | -1.02 | -0.98 | B>B |
| chr6 | 1.03E+08 | 102700000 | -0.99 | -0.97 | B>B |
| chr6 | 1.03E+08 | 102800000 | -1.03 | -0.96 | B>B |
| chr6 | 1.03E+08 | 102900000 | -1.00 | -0.98 | B>B |
| chr6 | 1.03E+08 | 103000000 | -1.03 | -1.03 | B>B |
| chr6 | 1.03E+08 | 103100000 | -0.94 | -1.03 | B>B |
| chr6 | 1.03E+08 | 103200000 | -0.99 | -0.98 | B>B |
| chr6 | 1.03E+08 | 103300000 | -0.92 | -0.94 | B>B |
| chr6 | 1.03E+08 | 103400000 | -0.91 | -0.98 | B>B |
| chr6 | 1.03E+08 | 103500000 | -1.02 | -0.97 | B>B |
| chr6 | 1.04E+08 | 103600000 | -0.95 | -0.99 | B>B |
| chr6 | 1.04E+08 | 103700000 | -1.07 | -1.02 | B>B |
| chr6 | 1.04E+08 | 103800000 | -1.06 | -1.01 | B>B |
| chr6 | 1.04E+08 | 103900000 | -0.98 | -1.05 | B>B |
| chr6 | 1.04E+08 | 104000000 | -1.04 | -1.03 | B>B |
| chr6 | 1.04E+08 | 104100000 | -1.06 | -1.05 | B>B |
| chr6 | 1.04E+08 | 104200000 | -1.05 | -1.10 | B>B |
| chr6 | 1.04E+08 | 104300000 | -0.98 | -1.05 | B>B |
| chr6 | 1.04E+08 | 104400000 | -1.05 | -1.14 | B>B |
| chr6 | 1.04E+08 | 104500000 | -1.11 | -1.11 | B>B |
| chr6 | 1.05E+08 | 104600000 | -1.05 | -1.18 | B>B |
| chr6 | 1.05E+08 | 104700000 | -1.00 | -1.15 | B>B |
| chr6 | 1.05E+08 | 104800000 | -0.78 | -1.09 | B>B |
| chr6 | 1.05E+08 | 104900000 | -0.68 | -1.10 | B>B |
| chr6 | 1.05E+08 | 105000000 | -0.59 | -0.99 | B>B |
| chr6 | 1.05E+08 | 105100000 | -0.26 | -0.87 | A>B |
| chr6 | 1.05E+08 | 105200000 | 0.11 | -0.43 | B>B |
| chr6 | 1.05E+08 | 105300000 | 0.26 | 0.06 | A>A |
| chr6 | 1.05E+08 | 105400000 | 0.49 | 0.22 | A>A |
| chr6 | 1.05E+08 | 105500000 | 0.48 | 0.34 | A>A |
| chr6 | 1.06E+08 | 105600000 | 0.54 | 0.46 | A>A |
| chr6 | 1.06E+08 | 105700000 | 0.90 | 0.71 | A>A |
| chr6 | 1.06E+08 | 105800000 | 0.89 | 0.66 | A>A |
| chr6 | 1.06E+08 | 105900000 | 0.77 | 0.67 | A>A |
| chr6 | 1.06E+08 | 106000000 | 1.01 | 0.82 | A>A |

| | | | | | |
|---|---|---|---|---|---|
| chr6 | 1.06E+08 | 106100000 | 1.29 | 1.12 | A>A |
| chr6 | 1.06E+08 | 106200000 | 1.39 | 0.94 | A>A |
| chr6 | 1.06E+08 | 106300000 | 1.28 | 0.98 | A>A |
| chr6 | 1.06E+08 | 106400000 | 1.20 | 1.03 | A>A |
| chr6 | 1.06E+08 | 106500000 | 1.23 | 0.98 | A>A |
| chr6 | 1.07E+08 | 106600000 | 1.06 | 0.99 | A>A |
| chr6 | 1.07E+08 | 106700000 | 1.03 | 0.93 | A>A |
| chr6 | 1.07E+08 | 106800000 | 0.87 | 0.95 | A>A |
| chr6 | 1.07E+08 | 106900000 | 1.02 | 1.10 | A>A |
| chr6 | 1.07E+08 | 107000000 | 1.27 | 1.30 | A>A |
| chr6 | 1.07E+08 | 107100000 | 1.22 | 1.18 | A>A |
| chr6 | 1.07E+08 | 107200000 | 1.54 | 1.56 | A>A |
| chr6 | 1.07E+08 | 107300000 | 1.52 | 1.50 | A>A |
| chr6 | 1.07E+08 | 107400000 | 1.13 | 1.10 | A>A |
| chr6 | 1.07E+08 | 107500000 | 1.18 | 1.12 | A>A |
| chr6 | 1.08E+08 | 107600000 | 1.25 | 1.24 | A>A |
| chr6 | 1.08E+08 | 107700000 | 1.18 | 1.24 | A>A |
| chr6 | 1.08E+08 | 107800000 | 1.19 | 1.16 | A>A |
| chr6 | 1.08E+08 | 107900000 | 1.22 | 1.17 | A>A |
| chr6 | 1.08E+08 | 108000000 | 1.08 | 1.16 | A>A |
| chr6 | 1.08E+08 | 108100000 | 1.13 | 1.09 | A>A |
| chr6 | 1.08E+08 | 108200000 | 1.16 | 1.06 | A>A |
| chr6 | 1.08E+08 | 108300000 | 1.33 | 1.25 | A>A |
| chr6 | 1.08E+08 | 108400000 | 1.22 | 1.18 | A>A |
| chr6 | 1.08E+08 | 108500000 | 1.12 | 1.17 | A>A |
| chr6 | 1.09E+08 | 108600000 | 1.24 | 1.32 | A>A |
| chr6 | 1.09E+08 | 108700000 | 1.32 | 1.24 | A>A |
| chr6 | 1.09E+08 | 108800000 | 1.01 | 1.19 | A>A |
| chr6 | 1.09E+08 | 108900000 | 0.81 | 0.85 | A>A |
| chr6 | 1.09E+08 | 109000000 | 1.30 | 1.35 | A>A |
| chr6 | 1.09E+08 | 109100000 | 1.14 | 1.24 | A>A |
| chr6 | 1.09E+08 | 109200000 | 0.91 | 1.07 | A>A |
| chr6 | 1.09E+08 | 109300000 | 1.03 | 1.14 | A>A |
| chr6 | 1.09E+08 | 109400000 | 1.05 | 1.13 | A>A |
| chr6 | 1.09E+08 | 109500000 | 0.97 | 1.02 | A>A |
| chr6 | 1.1E+08 | 109600000 | 0.95 | 1.16 | A>A |
| chr6 | 1.1E+08 | 109700000 | 1.22 | 1.27 | A>A |
| chr6 | 1.1E+08 | 109800000 | 1.02 | 0.94 | A>A |
| chr6 | 1.1E+08 | 109900000 | 0.38 | 0.43 | A>A |
| chr6 | 1.1E+08 | 110000000 | 0.23 | 0.24 | A>A |
| chr6 | 1.1E+08 | 110100000 | 0.36 | 0.48 | A>A |
| chr6 | 1.1E+08 | 110200000 | 0.54 | 0.49 | A>A |
| chr6 | 1.1E+08 | 110300000 | 0.32 | 0.22 | A>A |
| chr6 | 1.1E+08 | 110400000 | 0.27 | 0.13 | A>A |
| chr6 | 1.1E+08 | 110500000 | 0.60 | 0.50 | A>A |
| chr6 | 1.11E+08 | 110600000 | 0.83 | 0.63 | A>A |
| chr6 | 1.11E+08 | 110700000 | 0.56 | 0.53 | A>A |
| chr6 | 1.11E+08 | 110800000 | 0.61 | 0.46 | A>A |
| chr6 | 1.11E+08 | 110900000 | 0.95 | 0.78 | A>A |
| chr6 | 1.11E+08 | 111000000 | 1.18 | 1.02 | A>A |
| chr6 | 1.11E+08 | 111100000 | 1.35 | 1.19 | A>A |
| chr6 | 1.11E+08 | 111200000 | 1.39 | 1.23 | A>A |
| chr6 | 1.11E+08 | 111300000 | 1.21 | 1.08 | A>A |
| chr6 | 1.11E+08 | 111400000 | 0.82 | 0.65 | A>A |
| chr6 | 1.11E+08 | 111500000 | 1.19 | 0.67 | A>A |
| chr6 | 1.12E+08 | 111600000 | 0.96 | 0.62 | A>A |

| | | | | | |
|---|---|---|---|---|---|
| chr6 | 1.12E+08 | 111700000 | 0.96 | 0.72 | A>A |
| chr6 | 1.12E+08 | 111800000 | 1.08 | 0.79 | A>A |
| chr6 | 1.12E+08 | 111900000 | 1.03 | 0.68 | A>A |
| chr6 | 1.12E+08 | 112000000 | 0.83 | 0.51 | A>A |
| chr6 | 1.12E+08 | 112100000 | 0.41 | -0.05 | A>A |
| chr6 | 1.12E+08 | 112200000 | 0.51 | -0.35 | A>B |
| chr6 | 1.12E+08 | 112300000 | 0.33 | -0.30 | A>B |
| chr6 | 1.12E+08 | 112400000 | 0.36 | -0.09 | A>A |
| chr6 | 1.12E+08 | 112500000 | 0.27 | -0.12 | A>A |
| chr6 | 1.13E+08 | 112600000 | 0.28 | 0.02 | A>A |
| chr6 | 1.13E+08 | 112700000 | 0.11 | -0.18 | B>B |
| chr6 | 1.13E+08 | 112800000 | -0.08 | -0.29 | B>B |
| chr6 | 1.13E+08 | 112900000 | -0.09 | -0.23 | B>B |
| chr6 | 1.13E+08 | 113000000 | 0.00 | -0.16 | B>B |
| chr6 | 1.13E+08 | 113100000 | 0.14 | -0.01 | A>A |
| chr6 | 1.13E+08 | 113200000 | 0.23 | 0.28 | A>A |
| chr6 | 1.13E+08 | 113300000 | 0.33 | 0.51 | A>A |
| chr6 | 1.13E+08 | 113400000 | 0.44 | 0.57 | A>A |
| chr6 | 1.13E+08 | 113500000 | 0.59 | 0.53 | A>A |
| chr6 | 1.14E+08 | 113600000 | 0.68 | 0.67 | A>A |
| chr6 | 1.14E+08 | 113700000 | 0.86 | 0.76 | A>A |
| chr6 | 1.14E+08 | 113800000 | 0.68 | 0.71 | A>A |
| chr6 | 1.14E+08 | 113900000 | 0.93 | 0.86 | A>A |
| chr6 | 1.14E+08 | 114000000 | 0.89 | 0.77 | A>A |
| chr6 | 1.14E+08 | 114100000 | 0.90 | 0.87 | A>A |
| chr6 | 1.14E+08 | 114200000 | 0.72 | 0.79 | A>A |
| chr6 | 1.14E+08 | 114300000 | 0.19 | 0.10 | A>A |
| chr6 | 1.14E+08 | 114400000 | -0.18 | -0.37 | B>B |
| chr6 | 1.14E+08 | 114500000 | -0.41 | -0.62 | B>B |
| chr6 | 1.15E+08 | 114600000 | -0.65 | -0.89 | B>B |
| chr6 | 1.15E+08 | 114700000 | -0.49 | -0.65 | B>B |
| chr6 | 1.15E+08 | 114800000 | -0.69 | -0.82 | B>B |
| chr6 | 1.15E+08 | 114900000 | -0.80 | -0.96 | B>B |
| chr6 | 1.15E+08 | 115000000 | -0.92 | -1.04 | B>B |
| chr6 | 1.15E+08 | 115100000 | -0.94 | -1.01 | B>B |
| chr6 | 1.15E+08 | 115200000 | -1.03 | -1.01 | B>B |
| chr6 | 1.15E+08 | 115300000 | -1.05 | -1.04 | B>B |
| chr6 | 1.15E+08 | 115400000 | -0.98 | -1.02 | B>B |
| chr6 | 1.15E+08 | 115500000 | -1.06 | -1.02 | B>B |
| chr6 | 1.16E+08 | 115600000 | -1.06 | -1.08 | B>B |
| chr6 | 1.16E+08 | 115700000 | -0.92 | -1.15 | B>B |
| chr6 | 1.16E+08 | 115800000 | -0.90 | -1.15 | B>B |
| chr6 | 1.16E+08 | 115900000 | -0.79 | -1.06 | B>B |
| chr6 | 1.16E+08 | 116000000 | -0.70 | -1.15 | B>B |
| chr6 | 1.16E+08 | 116100000 | -0.61 | -1.09 | B>B |
| chr6 | 1.16E+08 | 116200000 | -0.50 | -0.93 | B>B |
| chr6 | 1.16E+08 | 116300000 | -0.06 | -0.43 | B>B |
| chr6 | 1.16E+08 | 116400000 | 0.65 | 0.45 | A>A |
| chr6 | 1.16E+08 | 116500000 | 0.49 | 0.40 | A>A |
| chr6 | 1.17E+08 | 116600000 | 0.54 | 0.44 | A>A |
| chr6 | 1.17E+08 | 116700000 | 0.59 | 0.51 | A>A |
| chr6 | 1.17E+08 | 116800000 | 0.66 | 0.64 | A>A |
| chr6 | 1.17E+08 | 116900000 | 0.62 | 0.62 | A>A |
| chr6 | 1.17E+08 | 117000000 | 0.73 | 0.62 | A>A |
| chr6 | 1.17E+08 | 117100000 | 0.27 | 0.28 | A>A |
| chr6 | 1.17E+08 | 117200000 | -0.20 | -0.12 | B>B |

| | | | | | |
|---|---|---|---|---|---|
| chr6 | 1.17E+08 | 117300000 | -0.55 | -0.44 | B>B |
| chr6 | 1.17E+08 | 117400000 | -0.75 | -1.05 | B>B |
| chr6 | 1.17E+08 | 117500000 | -0.71 | -1.15 | B>B |
| chr6 | 1.18E+08 | 117600000 | -0.59 | -0.92 | B>B |
| chr6 | 1.18E+08 | 117700000 | -0.26 | -0.40 | B>B |
| chr6 | 1.18E+08 | 117800000 | 0.37 | 0.40 | A>A |
| chr6 | 1.18E+08 | 117900000 | 0.74 | 0.54 | A>A |
| chr6 | 1.18E+08 | 118000000 | 0.17 | -0.18 | B>B |
| chr6 | 1.18E+08 | 118100000 | -0.35 | -0.59 | B>B |
| chr6 | 1.18E+08 | 118200000 | -0.56 | -0.75 | B>B |
| chr6 | 1.18E+08 | 118300000 | -0.56 | -0.92 | B>B |
| chr6 | 1.18E+08 | 118400000 | -0.72 | -1.12 | B>B |
| chr6 | 1.18E+08 | 118500000 | -0.75 | -1.12 | B>B |
| chr6 | 1.19E+08 | 118600000 | -0.65 | -1.05 | B>B |
| chr6 | 1.19E+08 | 118700000 | -0.44 | -0.88 | B>B |
| chr6 | 1.19E+08 | 118800000 | -0.39 | -0.64 | B>B |
| chr6 | 1.19E+08 | 118900000 | -0.10 | -0.22 | B>B |
| chr6 | 1.19E+08 | 119000000 | 0.48 | 0.20 | A>A |
| chr6 | 1.19E+08 | 119100000 | 0.68 | 0.50 | A>A |
| chr6 | 1.19E+08 | 119200000 | 0.41 | 0.21 | A>A |
| chr6 | 1.19E+08 | 119300000 | 0.26 | 0.10 | A>A |
| chr6 | 1.19E+08 | 119400000 | -0.21 | -0.47 | B>B |
| chr6 | 1.19E+08 | 119500000 | -0.28 | -0.40 | B>B |
| chr6 | 1.2E+08 | 119600000 | -0.34 | -0.49 | B>B |
| chr6 | 1.2E+08 | 119700000 | -0.28 | -0.56 | B>B |
| chr6 | 1.2E+08 | 119800000 | -0.69 | -0.80 | B>B |
| chr6 | 1.2E+08 | 119900000 | -0.72 | -0.94 | B>B |
| chr6 | 1.2E+08 | 120000000 | -0.76 | -0.95 | B>B |
| chr6 | 1.2E+08 | 120100000 | -0.73 | -0.89 | B>B |
| chr6 | 1.2E+08 | 120200000 | -0.81 | -0.91 | B>B |
| chr6 | 1.2E+08 | 120300000 | -0.80 | -0.84 | B>B |
| chr6 | 1.2E+08 | 120400000 | -0.83 | -0.60 | B>B |
| chr6 | 1.2E+08 | 120500000 | -0.97 | -0.52 | B>B |
| chr6 | 1.21E+08 | 120600000 | -0.95 | -0.72 | B>B |
| chr6 | 1.21E+08 | 120700000 | -0.99 | -0.77 | B>B |
| chr6 | 1.21E+08 | 120800000 | -0.97 | -0.90 | B>B |
| chr6 | 1.21E+08 | 120900000 | -1.00 | -1.05 | B>B |
| chr6 | 1.21E+08 | 121000000 | -1.01 | -1.00 | B>B |
| chr6 | 1.21E+08 | 121100000 | -0.99 | -0.99 | B>B |
| chr6 | 1.21E+08 | 121200000 | -0.93 | -0.90 | B>B |
| chr6 | 1.21E+08 | 121300000 | -0.69 | -0.95 | B>B |
| chr6 | 1.21E+08 | 121400000 | -0.45 | -0.62 | B>B |
| chr6 | 1.21E+08 | 121500000 | -0.24 | -0.24 | B>B |
| chr6 | 1.22E+08 | 121600000 | -0.05 | -0.22 | B>B |
| chr6 | 1.22E+08 | 121700000 | 0.00 | -0.19 | B>B |
| chr6 | 1.22E+08 | 121800000 | 0.27 | 0.02 | A>A |
| chr6 | 1.22E+08 | 121900000 | 0.83 | 0.60 | A>A |
| chr6 | 1.22E+08 | 122000000 | 0.88 | 0.57 | A>A |
| chr6 | 1.22E+08 | 122100000 | 0.80 | 0.44 | A>A |
| chr6 | 1.22E+08 | 122200000 | 0.60 | 0.13 | A>A |
| chr6 | 1.22E+08 | 122300000 | 0.31 | -0.42 | A>B |
| chr6 | 1.22E+08 | 122400000 | 0.03 | -0.60 | A>B |
| chr6 | 1.22E+08 | 122500000 | -0.32 | -0.90 | B>B |
| chr6 | 1.23E+08 | 122600000 | -0.57 | -0.96 | B>B |
| chr6 | 1.23E+08 | 122700000 | -0.48 | -0.82 | B>B |
| chr6 | 1.23E+08 | 122800000 | 0.08 | -0.12 | B>B |

| | | | | | |
|---|---|---|---|---|---|
| chr6 | 1.23E+08 | 122900000 | -0.31 | -0.30 | B>B |
| chr6 | 1.23E+08 | 123000000 | -0.25 | -0.18 | B>B |
| chr6 | 1.23E+08 | 123100000 | -0.30 | -0.21 | B>B |
| chr6 | 1.23E+08 | 123200000 | -0.21 | -0.24 | B>B |
| chr6 | 1.23E+08 | 123300000 | -0.81 | -0.85 | B>B |
| chr6 | 1.23E+08 | 123400000 | -0.77 | -0.90 | B>B |
| chr6 | 1.23E+08 | 123500000 | -0.95 | -0.93 | B>B |
| chr6 | 1.24E+08 | 123600000 | -0.92 | -0.95 | B>B |
| chr6 | 1.24E+08 | 123700000 | -0.76 | -0.93 | B>B |
| chr6 | 1.24E+08 | 123800000 | -0.48 | -0.94 | B>B |
| chr6 | 1.24E+08 | 123900000 | -0.31 | -0.99 | A>B |
| chr6 | 1.24E+08 | 124000000 | -0.64 | -1.02 | B>B |
| chr6 | 1.24E+08 | 124100000 | -0.62 | -0.98 | B>B |
| chr6 | 1.24E+08 | 124200000 | -0.53 | -0.94 | B>B |
| chr6 | 1.24E+08 | 124300000 | -0.70 | -1.03 | B>B |
| chr6 | 1.24E+08 | 124400000 | -0.73 | -1.13 | B>B |
| chr6 | 1.24E+08 | 124500000 | -0.73 | -1.14 | B>B |
| chr6 | 1.25E+08 | 124600000 | -0.71 | -1.09 | B>B |
| chr6 | 1.25E+08 | 124700000 | -0.67 | -1.08 | B>B |
| chr6 | 1.25E+08 | 124800000 | -0.68 | -1.10 | B>B |
| chr6 | 1.25E+08 | 124900000 | -0.53 | -1.04 | B>B |
| chr6 | 1.25E+08 | 125000000 | -0.63 | -1.04 | B>B |
| chr6 | 1.25E+08 | 125100000 | -0.67 | -1.03 | B>B |
| chr6 | 1.25E+08 | 125200000 | -0.42 | -0.65 | B>B |
| chr6 | 1.25E+08 | 125300000 | 0.44 | 0.36 | A>A |
| chr6 | 1.25E+08 | 125400000 | 0.48 | 0.46 | A>A |
| chr6 | 1.25E+08 | 125500000 | 0.33 | 0.12 | A>A |
| chr6 | 1.26E+08 | 125600000 | 0.47 | 0.34 | A>A |
| chr6 | 1.26E+08 | 125700000 | -0.06 | -0.32 | B>B |
| chr6 | 1.26E+08 | 125800000 | -0.38 | -0.71 | B>B |
| chr6 | 1.26E+08 | 125900000 | -0.39 | -0.68 | B>B |
| chr6 | 1.26E+08 | 126000000 | -0.36 | -0.63 | B>B |
| chr6 | 1.26E+08 | 126100000 | -0.18 | -0.35 | B>B |
| chr6 | 1.26E+08 | 126200000 | 0.69 | 0.55 | A>A |
| chr6 | 1.26E+08 | 126300000 | 0.89 | 0.71 | A>A |
| chr6 | 1.26E+08 | 126400000 | 0.47 | 0.24 | A>A |
| chr6 | 1.26E+08 | 126500000 | 0.40 | 0.20 | A>A |
| chr6 | 1.27E+08 | 126600000 | 0.18 | -0.07 | A>A |
| chr6 | 1.27E+08 | 126700000 | 0.11 | -0.11 | B>B |
| chr6 | 1.27E+08 | 126800000 | -0.01 | -0.13 | B>B |
| chr6 | 1.27E+08 | 126900000 | 0.06 | -0.14 | B>B |
| chr6 | 1.27E+08 | 127000000 | 0.04 | -0.14 | B>B |
| chr6 | 1.27E+08 | 127100000 | 0.22 | -0.26 | B>B |
| chr6 | 1.27E+08 | 127200000 | 0.37 | -0.07 | A>A |
| chr6 | 1.27E+08 | 127300000 | 0.55 | 0.17 | A>A |
| chr6 | 1.27E+08 | 127400000 | 0.57 | 0.24 | A>A |
| chr6 | 1.27E+08 | 127500000 | 0.52 | -0.05 | A>A |
| chr6 | 1.28E+08 | 127600000 | 0.32 | -0.02 | A>A |
| chr6 | 1.28E+08 | 127700000 | 0.57 | 0.56 | A>A |
| chr6 | 1.28E+08 | 127800000 | 0.56 | 0.57 | A>A |
| chr6 | 1.28E+08 | 127900000 | 0.30 | 0.43 | A>A |
| chr6 | 1.28E+08 | 128000000 | -0.01 | 0.17 | A>A |
| chr6 | 1.28E+08 | 128100000 | -0.32 | 0.01 | B>B |
| chr6 | 1.28E+08 | 128200000 | -0.22 | 0.07 | B>B |
| chr6 | 1.28E+08 | 128300000 | -0.05 | 0.32 | A>A |
| chr6 | 1.28E+08 | 128400000 | 0.04 | 0.52 | A>A |

| | | | | | |
|---|---|---|---|---|---|
| chr6 | 1.28E+08 | 128500000 | 0.23 | 0.58 | A>A |
| chr6 | 1.29E+08 | 128600000 | 0.32 | 0.71 | A>A |
| chr6 | 1.29E+08 | 128700000 | 0.34 | 0.64 | A>A |
| chr6 | 1.29E+08 | 128800000 | 0.43 | 0.79 | A>A |
| chr6 | 1.29E+08 | 128900000 | 0.19 | 0.50 | A>A |
| chr6 | 1.29E+08 | 129000000 | 0.14 | 0.47 | A>A |
| chr6 | 1.29E+08 | 129100000 | -0.21 | 0.09 | B>B |
| chr6 | 1.29E+08 | 129200000 | -0.60 | -0.17 | B>B |
| chr6 | 1.29E+08 | 129300000 | -0.71 | -0.32 | B>B |
| chr6 | 1.29E+08 | 129400000 | -0.67 | -0.25 | B>B |
| chr6 | 1.29E+08 | 129500000 | -0.54 | -0.19 | B>B |
| chr6 | 1.3E+08 | 129600000 | -0.53 | -0.11 | B>B |
| chr6 | 1.3E+08 | 129700000 | -0.32 | 0.08 | B>B |
| chr6 | 1.3E+08 | 129800000 | -0.05 | 0.05 | A>A |
| chr6 | 1.3E+08 | 129900000 | 0.22 | 0.26 | A>A |
| chr6 | 1.3E+08 | 130000000 | 0.79 | 0.79 | A>A |
| chr6 | 1.3E+08 | 130100000 | 0.64 | 0.65 | A>A |
| chr6 | 1.3E+08 | 130200000 | 0.30 | 0.14 | A>A |
| chr6 | 1.3E+08 | 130300000 | 0.31 | 0.14 | A>A |
| chr6 | 1.3E+08 | 130400000 | 0.18 | 0.06 | A>A |
| chr6 | 1.3E+08 | 130500000 | 0.08 | -0.10 | B>B |
| chr6 | 1.31E+08 | 130600000 | -0.01 | -0.21 | B>B |
| chr6 | 1.31E+08 | 130700000 | -0.10 | -0.13 | B>B |
| chr6 | 1.31E+08 | 130800000 | -0.47 | -0.43 | B>B |
| chr6 | 1.31E+08 | 130900000 | -0.39 | -0.48 | B>B |
| chr6 | 1.31E+08 | 131000000 | -0.40 | -0.36 | B>B |
| chr6 | 1.31E+08 | 131100000 | -0.24 | -0.17 | B>B |
| chr6 | 1.31E+08 | 131200000 | 0.01 | 0.12 | A>A |
| chr6 | 1.31E+08 | 131300000 | 0.67 | 0.76 | A>A |
| chr6 | 1.31E+08 | 131400000 | 0.88 | 0.81 | A>A |
| chr6 | 1.31E+08 | 131500000 | 0.39 | 0.63 | A>A |
| chr6 | 1.32E+08 | 131600000 | 0.17 | 0.68 | A>A |
| chr6 | 1.32E+08 | 131700000 | 0.15 | 0.75 | B>A |
| chr6 | 1.32E+08 | 131800000 | 0.14 | 0.72 | B>A |
| chr6 | 1.32E+08 | 131900000 | 0.12 | 0.68 | B>A |
| chr6 | 1.32E+08 | 132000000 | 0.38 | 0.66 | A>A |
| chr6 | 1.32E+08 | 132100000 | 0.30 | 0.63 | A>A |
| chr6 | 1.32E+08 | 132200000 | 0.56 | 0.75 | A>A |
| chr6 | 1.32E+08 | 132300000 | 0.33 | 0.66 | A>A |
| chr6 | 1.32E+08 | 132400000 | 0.49 | 0.79 | A>A |
| chr6 | 1.32E+08 | 132500000 | 0.25 | 0.62 | A>A |
| chr6 | 1.33E+08 | 132600000 | 0.20 | 0.44 | A>A |
| chr6 | 1.33E+08 | 132700000 | 0.52 | 0.77 | A>A |
| chr6 | 1.33E+08 | 132800000 | 0.43 | 0.77 | A>A |
| chr6 | 1.33E+08 | 132900000 | 0.48 | 0.64 | A>A |
| chr6 | 1.33E+08 | 133000000 | 0.49 | 0.71 | A>A |
| chr6 | 1.33E+08 | 133100000 | 0.68 | 0.87 | A>A |
| chr6 | 1.33E+08 | 133200000 | 0.63 | 0.63 | A>A |
| chr6 | 1.33E+08 | 133300000 | 0.62 | 0.64 | A>A |
| chr6 | 1.33E+08 | 133400000 | 0.73 | 0.72 | A>A |
| chr6 | 1.33E+08 | 133500000 | 0.75 | 0.70 | A>A |
| chr6 | 1.34E+08 | 133600000 | 0.79 | 0.83 | A>A |
| chr6 | 1.34E+08 | 133700000 | 0.88 | 0.84 | A>A |
| chr6 | 1.34E+08 | 133800000 | 0.83 | 0.83 | A>A |
| chr6 | 1.34E+08 | 133900000 | 0.55 | 0.75 | A>A |
| chr6 | 1.34E+08 | 134000000 | 1.02 | 1.01 | A>A |

| | | | | | |
|---|---|---|---|---|---|
| chr6 | 1.34E+08 | 134100000 | 1.12 | 1.18 | A>A |
| chr6 | 1.34E+08 | 134200000 | 1.02 | 0.96 | A>A |
| chr6 | 1.34E+08 | 134300000 | 0.87 | 0.76 | A>A |
| chr6 | 1.34E+08 | 134400000 | 1.15 | 1.09 | A>A |
| chr6 | 1.34E+08 | 134500000 | 0.53 | 0.59 | A>A |
| chr6 | 1.35E+08 | 134600000 | 0.86 | 0.91 | A>A |
| chr6 | 1.35E+08 | 134700000 | 1.04 | 1.05 | A>A |
| chr6 | 1.35E+08 | 134800000 | 0.98 | 1.07 | A>A |
| chr6 | 1.35E+08 | 134900000 | 1.12 | 1.04 | A>A |
| chr6 | 1.35E+08 | 135000000 | 0.80 | 0.91 | A>A |
| chr6 | 1.35E+08 | 135100000 | 0.98 | 1.07 | A>A |
| chr6 | 1.35E+08 | 135200000 | 0.76 | 0.78 | A>A |
| chr6 | 1.35E+08 | 135300000 | 0.85 | 0.78 | A>A |
| chr6 | 1.35E+08 | 135400000 | 1.06 | 0.98 | A>A |
| chr6 | 1.35E+08 | 135500000 | 1.02 | 0.82 | A>A |
| chr6 | 1.36E+08 | 135600000 | 0.92 | 0.90 | A>A |
| chr6 | 1.36E+08 | 135700000 | 0.97 | 0.90 | A>A |
| chr6 | 1.36E+08 | 135800000 | 0.84 | 0.78 | A>A |
| chr6 | 1.36E+08 | 135900000 | 0.73 | 0.76 | A>A |
| chr6 | 1.36E+08 | 136000000 | 0.96 | 0.93 | A>A |
| chr6 | 1.36E+08 | 136100000 | 0.93 | 1.00 | A>A |
| chr6 | 1.36E+08 | 136200000 | 1.07 | 0.95 | A>A |
| chr6 | 1.36E+08 | 136300000 | 1.00 | 0.90 | A>A |
| chr6 | 1.36E+08 | 136400000 | 0.93 | 0.95 | A>A |
| chr6 | 1.36E+08 | 136500000 | 1.03 | 0.94 | A>A |
| chr6 | 1.37E+08 | 136600000 | 0.66 | 0.76 | A>A |
| chr6 | 1.37E+08 | 136700000 | 0.89 | 0.86 | A>A |
| chr6 | 1.37E+08 | 136800000 | 1.25 | 1.25 | A>A |
| chr6 | 1.37E+08 | 136900000 | 0.99 | 1.06 | A>A |
| chr6 | 1.37E+08 | 137000000 | 0.88 | 0.82 | A>A |
| chr6 | 1.37E+08 | 137100000 | 1.02 | 0.82 | A>A |
| chr6 | 1.37E+08 | 137200000 | 0.87 | 0.70 | A>A |
| chr6 | 1.37E+08 | 137300000 | 0.90 | 0.69 | A>A |
| chr6 | 1.37E+08 | 137400000 | 0.69 | 0.62 | A>A |
| chr6 | 1.37E+08 | 137500000 | 0.61 | 0.54 | A>A |
| chr6 | 1.38E+08 | 137600000 | 0.74 | 0.74 | A>A |
| chr6 | 1.38E+08 | 137700000 | 0.63 | 0.52 | A>A |
| chr6 | 1.38E+08 | 137800000 | 0.62 | 0.53 | A>A |
| chr6 | 1.38E+08 | 137900000 | 0.59 | 0.46 | A>A |
| chr6 | 1.38E+08 | 138000000 | 0.57 | 0.42 | A>A |
| chr6 | 1.38E+08 | 138100000 | 0.79 | 0.59 | A>A |
| chr6 | 1.38E+08 | 138200000 | 0.87 | 0.75 | A>A |
| chr6 | 1.38E+08 | 138300000 | 0.60 | 0.54 | A>A |
| chr6 | 1.38E+08 | 138400000 | 0.73 | 0.71 | A>A |
| chr6 | 1.38E+08 | 138500000 | 0.65 | 0.73 | A>A |
| chr6 | 1.39E+08 | 138600000 | 0.79 | 0.81 | A>A |
| chr6 | 1.39E+08 | 138700000 | 0.77 | 0.79 | A>A |
| chr6 | 1.39E+08 | 138800000 | 0.75 | 0.81 | A>A |
| chr6 | 1.39E+08 | 138900000 | 0.82 | 0.79 | A>A |
| chr6 | 1.39E+08 | 139000000 | 0.83 | 0.81 | A>A |
| chr6 | 1.39E+08 | 139100000 | 0.73 | 0.75 | A>A |
| chr6 | 1.39E+08 | 139200000 | 0.48 | 0.67 | A>A |
| chr6 | 1.39E+08 | 139300000 | 0.92 | 0.99 | A>A |
| chr6 | 1.39E+08 | 139400000 | 0.69 | 0.76 | A>A |
| chr6 | 1.39E+08 | 139500000 | 0.46 | 0.73 | A>A |
| chr6 | 1.4E+08 | 139600000 | 0.35 | 0.65 | A>A |

| | | | | | |
|---|---|---|---|---|---|
| chr6 | 1.4E+08 | 139700000 | 0.26 | 0.51 | A>A |
| chr6 | 1.4E+08 | 139800000 | 0.44 | 0.72 | A>A |
| chr6 | 1.4E+08 | 139900000 | 0.43 | 0.76 | A>A |
| chr6 | 1.4E+08 | 140000000 | 0.36 | 0.75 | A>A |
| chr6 | 1.4E+08 | 140100000 | 0.35 | 0.73 | A>A |
| chr6 | 1.4E+08 | 140200000 | 0.27 | 0.66 | A>A |
| chr6 | 1.4E+08 | 140300000 | 0.22 | 0.57 | A>A |
| chr6 | 1.4E+08 | 140400000 | 0.16 | 0.54 | A>A |
| chr6 | 1.4E+08 | 140500000 | 0.14 | 0.68 | B>A |
| chr6 | 1.41E+08 | 140600000 | 0.11 | 0.74 | B>A |
| chr6 | 1.41E+08 | 140700000 | 0.02 | 0.66 | B>A |
| chr6 | 1.41E+08 | 140800000 | 0.02 | 0.57 | B>A |
| chr6 | 1.41E+08 | 140900000 | 0.12 | 0.71 | B>A |
| chr6 | 1.41E+08 | 141000000 | 0.02 | 0.63 | B>A |
| chr6 | 1.41E+08 | 141100000 | -0.03 | 0.73 | B>A |
| chr6 | 1.41E+08 | 141200000 | 0.07 | 0.67 | B>A |
| chr6 | 1.41E+08 | 141300000 | -0.07 | 0.52 | B>A |
| chr6 | 1.41E+08 | 141400000 | 0.00 | 0.64 | B>A |
| chr6 | 1.41E+08 | 141500000 | 0.01 | 0.70 | B>A |
| chr6 | 1.42E+08 | 141600000 | 0.05 | 0.78 | B>A |
| chr6 | 1.42E+08 | 141700000 | 0.26 | 0.65 | A>A |
| chr6 | 1.42E+08 | 141800000 | 0.27 | 0.77 | A>A |
| chr6 | 1.42E+08 | 141900000 | 0.09 | 0.62 | B>A |
| chr6 | 1.42E+08 | 142000000 | -0.12 | 0.45 | B>A |
| chr6 | 1.42E+08 | 142100000 | -0.16 | 0.46 | B>A |
| chr6 | 1.42E+08 | 142200000 | -0.30 | 0.42 | B>A |
| chr6 | 1.42E+08 | 142300000 | -0.22 | 0.39 | B>A |
| chr6 | 1.42E+08 | 142400000 | 0.02 | 0.49 | A>A |
| chr6 | 1.42E+08 | 142500000 | 0.17 | 0.51 | A>A |
| chr6 | 1.43E+08 | 142600000 | 0.63 | 0.87 | A>A |
| chr6 | 1.43E+08 | 142700000 | 0.83 | 1.05 | A>A |
| chr6 | 1.43E+08 | 142800000 | 0.58 | 0.69 | A>A |
| chr6 | 1.43E+08 | 142900000 | 0.29 | 0.38 | A>A |
| chr6 | 1.43E+08 | 143000000 | 0.38 | 0.36 | A>A |
| chr6 | 1.43E+08 | 143100000 | 0.42 | 0.51 | A>A |
| chr6 | 1.43E+08 | 143200000 | 0.65 | 0.78 | A>A |
| chr6 | 1.43E+08 | 143300000 | 0.66 | 0.70 | A>A |
| chr6 | 1.43E+08 | 143400000 | 0.46 | 0.49 | A>A |
| chr6 | 1.43E+08 | 143500000 | 0.46 | 0.47 | A>A |
| chr6 | 1.44E+08 | 143600000 | 0.39 | 0.50 | A>A |
| chr6 | 1.44E+08 | 143700000 | 0.68 | 0.46 | A>A |
| chr6 | 1.44E+08 | 143800000 | 0.56 | 0.46 | A>A |
| chr6 | 1.44E+08 | 143900000 | 0.43 | 0.52 | A>A |
| chr6 | 1.44E+08 | 144000000 | 0.62 | 0.61 | A>A |
| chr6 | 1.44E+08 | 144100000 | 0.79 | 0.89 | A>A |
| chr6 | 1.44E+08 | 144200000 | 0.71 | 0.65 | A>A |
| chr6 | 1.44E+08 | 144300000 | 0.56 | 0.50 | A>A |
| chr6 | 1.44E+08 | 144400000 | 0.45 | 0.46 | A>A |
| chr6 | 1.44E+08 | 144500000 | 0.53 | 0.53 | A>A |
| chr6 | 1.45E+08 | 144600000 | 0.70 | 0.50 | A>A |
| chr6 | 1.45E+08 | 144700000 | 0.80 | 0.81 | A>A |
| chr6 | 1.45E+08 | 144800000 | 0.69 | 0.70 | A>A |
| chr6 | 1.45E+08 | 144900000 | 0.44 | 0.38 | A>A |
| chr6 | 1.45E+08 | 145000000 | 0.22 | 0.40 | A>A |
| chr6 | 1.45E+08 | 145100000 | 0.44 | 0.60 | A>A |
| chr6 | 1.45E+08 | 145200000 | 0.45 | 0.52 | A>A |

| | | | | | |
|---|---|---|---|---|---|
| chr6 | 1.45E+08 | 145300000 | 0.25 | 0.43 | A>A |
| chr6 | 1.45E+08 | 145400000 | 0.13 | 0.20 | A>A |
| chr6 | 1.45E+08 | 145500000 | -0.07 | 0.27 | A>A |
| chr6 | 1.46E+08 | 145600000 | -0.05 | 0.21 | A>A |
| chr6 | 1.46E+08 | 145700000 | -0.08 | 0.18 | A>A |
| chr6 | 1.46E+08 | 145800000 | -0.16 | 0.10 | B>B |
| chr6 | 1.46E+08 | 145900000 | -0.09 | 0.10 | A>A |
| chr6 | 1.46E+08 | 146000000 | -0.08 | 0.14 | A>A |
| chr6 | 1.46E+08 | 146100000 | 0.00 | 0.23 | A>A |
| chr6 | 1.46E+08 | 146200000 | 0.10 | 0.18 | A>A |
| chr6 | 1.46E+08 | 146300000 | -0.05 | -0.14 | B>B |
| chr6 | 1.46E+08 | 146400000 | -0.59 | -0.62 | B>B |
| chr6 | 1.46E+08 | 146500000 | -0.62 | -0.72 | B>B |
| chr6 | 1.47E+08 | 146600000 | -0.42 | -0.49 | B>B |
| chr6 | 1.47E+08 | 146700000 | -0.31 | -0.13 | B>B |
| chr6 | 1.47E+08 | 146800000 | -0.43 | -0.34 | B>B |
| chr6 | 1.47E+08 | 146900000 | -0.39 | -0.39 | B>B |
| chr6 | 1.47E+08 | 147000000 | -0.55 | -0.68 | B>B |
| chr6 | 1.47E+08 | 147100000 | -0.47 | -0.56 | B>B |
| chr6 | 1.47E+08 | 147200000 | -0.06 | -0.24 | B>B |
| chr6 | 1.47E+08 | 147300000 | 0.25 | 0.25 | A>A |
| chr6 | 1.47E+08 | 147400000 | 0.30 | 0.36 | A>A |
| chr6 | 1.47E+08 | 147500000 | 0.33 | 0.40 | A>A |
| chr6 | 1.48E+08 | 147600000 | 0.39 | 0.34 | A>A |
| chr6 | 1.48E+08 | 147700000 | 0.29 | 0.23 | A>A |
| chr6 | 1.48E+08 | 147800000 | 0.12 | 0.07 | A>A |
| chr6 | 1.48E+08 | 147900000 | 0.31 | 0.12 | A>A |
| chr6 | 1.48E+08 | 148000000 | 0.30 | 0.12 | A>A |
| chr6 | 1.48E+08 | 148100000 | 0.27 | 0.20 | A>A |
| chr6 | 1.48E+08 | 148200000 | 0.38 | 0.21 | A>A |
| chr6 | 1.48E+08 | 148300000 | 0.36 | 0.28 | A>A |
| chr6 | 1.48E+08 | 148400000 | 0.43 | 0.29 | A>A |
| chr6 | 1.48E+08 | 148500000 | 0.37 | 0.35 | A>A |
| chr6 | 1.49E+08 | 148600000 | 0.44 | 0.19 | A>A |
| chr6 | 1.49E+08 | 148700000 | 0.67 | 0.37 | A>A |
| chr6 | 1.49E+08 | 148800000 | 0.64 | 0.49 | A>A |
| chr6 | 1.49E+08 | 148900000 | 0.41 | 0.36 | A>A |
| chr6 | 1.49E+08 | 149000000 | 0.64 | 0.55 | A>A |
| chr6 | 1.49E+08 | 149100000 | 0.71 | 0.57 | A>A |
| chr6 | 1.49E+08 | 149200000 | 0.93 | 0.68 | A>A |
| chr6 | 1.49E+08 | 149300000 | 0.71 | 0.61 | A>A |
| chr6 | 1.49E+08 | 149400000 | 0.60 | 0.55 | A>A |
| chr6 | 1.49E+08 | 149500000 | 0.96 | 0.81 | A>A |
| chr6 | 1.5E+08 | 149600000 | 1.01 | 0.95 | A>A |
| chr6 | 1.5E+08 | 149700000 | 1.04 | 1.01 | A>A |
| chr6 | 1.5E+08 | 149800000 | 0.72 | 0.69 | A>A |
| chr6 | 1.5E+08 | 149900000 | 0.91 | 0.82 | A>A |
| chr6 | 1.5E+08 | 150000000 | 1.10 | 0.98 | A>A |
| chr6 | 1.5E+08 | 150100000 | 1.21 | 1.03 | A>A |
| chr6 | 1.5E+08 | 150200000 | 1.13 | 0.95 | A>A |
| chr6 | 1.5E+08 | 150300000 | 0.97 | 0.79 | A>A |
| chr6 | 1.5E+08 | 150400000 | 0.76 | 0.57 | A>A |
| chr6 | 1.5E+08 | 150500000 | 0.69 | 0.36 | A>A |
| chr6 | 1.51E+08 | 150600000 | 0.81 | 0.34 | A>A |
| chr6 | 1.51E+08 | 150700000 | 0.67 | 0.18 | A>A |
| chr6 | 1.51E+08 | 150800000 | 0.44 | 0.21 | A>A |

| | | | | | |
|---|---|---|---|---|---|
| chr6 | 1.51E+08 | 150900000 | 0.56 | 0.17 | A>A |
| chr6 | 1.51E+08 | 151000000 | 0.47 | 0.22 | A>A |
| chr6 | 1.51E+08 | 151100000 | 0.57 | 0.16 | A>A |
| chr6 | 1.51E+08 | 151200000 | 0.51 | 0.31 | A>A |
| chr6 | 1.51E+08 | 151300000 | 0.82 | 0.74 | A>A |
| chr6 | 1.51E+08 | 151400000 | 0.93 | 0.79 | A>A |
| chr6 | 1.51E+08 | 151500000 | 0.97 | 0.93 | A>A |
| chr6 | 1.52E+08 | 151600000 | 1.01 | 0.95 | A>A |
| chr6 | 1.52E+08 | 151700000 | 0.73 | 0.65 | A>A |
| chr6 | 1.52E+08 | 151800000 | 0.59 | 0.53 | A>A |
| chr6 | 1.52E+08 | 151900000 | 0.17 | 0.06 | A>A |
| chr6 | 1.52E+08 | 152000000 | -0.01 | -0.13 | B>B |
| chr6 | 1.52E+08 | 152100000 | -0.13 | -0.30 | B>B |
| chr6 | 1.52E+08 | 152200000 | -0.15 | -0.46 | B>B |
| chr6 | 1.52E+08 | 152300000 | -0.21 | -0.35 | B>B |
| chr6 | 1.52E+08 | 152400000 | -0.15 | -0.30 | B>B |
| chr6 | 1.52E+08 | 152500000 | 0.14 | -0.02 | A>A |
| chr6 | 1.53E+08 | 152600000 | 0.37 | 0.20 | A>A |
| chr6 | 1.53E+08 | 152700000 | 0.50 | 0.27 | A>A |
| chr6 | 1.53E+08 | 152800000 | 0.49 | 0.23 | A>A |
| chr6 | 1.53E+08 | 152900000 | 0.15 | -0.21 | B>B |
| chr6 | 1.53E+08 | 153000000 | -0.09 | -0.26 | B>B |
| chr6 | 1.53E+08 | 153100000 | -0.48 | -0.52 | B>B |
| chr6 | 1.53E+08 | 153200000 | -0.62 | -0.53 | B>B |
| chr6 | 1.53E+08 | 153300000 | -0.34 | -0.33 | B>B |
| chr6 | 1.53E+08 | 153400000 | 0.18 | 0.27 | A>A |
| chr6 | 1.53E+08 | 153500000 | 0.36 | 0.47 | A>A |
| chr6 | 1.54E+08 | 153600000 | 0.07 | 0.21 | A>A |
| chr6 | 1.54E+08 | 153700000 | -0.19 | 0.05 | B>B |
| chr6 | 1.54E+08 | 153800000 | -0.31 | -0.02 | B>B |
| chr6 | 1.54E+08 | 153900000 | -0.47 | -0.16 | B>B |
| chr6 | 1.54E+08 | 154000000 | -0.61 | -0.25 | B>B |
| chr6 | 1.54E+08 | 154100000 | -0.74 | -0.22 | B>A |
| chr6 | 1.54E+08 | 154200000 | -0.78 | -0.26 | B>A |
| chr6 | 1.54E+08 | 154300000 | -0.69 | -0.35 | B>B |
| chr6 | 1.54E+08 | 154400000 | -0.65 | -0.33 | B>B |
| chr6 | 1.54E+08 | 154500000 | -0.28 | -0.10 | B>B |
| chr6 | 1.55E+08 | 154600000 | 0.00 | 0.15 | A>A |
| chr6 | 1.55E+08 | 154700000 | 0.24 | 0.21 | A>A |
| chr6 | 1.55E+08 | 154800000 | 0.33 | 0.49 | A>A |
| chr6 | 1.55E+08 | 154900000 | 0.54 | 0.65 | A>A |
| chr6 | 1.55E+08 | 155000000 | 0.51 | 0.55 | A>A |
| chr6 | 1.55E+08 | 155100000 | 0.81 | 0.81 | A>A |
| chr6 | 1.55E+08 | 155200000 | 0.76 | 0.75 | A>A |
| chr6 | 1.55E+08 | 155300000 | 0.60 | 0.60 | A>A |
| chr6 | 1.55E+08 | 155400000 | 0.93 | 0.77 | A>A |
| chr6 | 1.55E+08 | 155500000 | 0.73 | 0.70 | A>A |
| chr6 | 1.56E+08 | 155600000 | 0.55 | 0.56 | A>A |
| chr6 | 1.56E+08 | 155700000 | 0.27 | -0.03 | A>A |
| chr6 | 1.56E+08 | 155800000 | 0.29 | -0.07 | A>A |
| chr6 | 1.56E+08 | 155900000 | 0.17 | -0.26 | B>B |
| chr6 | 1.56E+08 | 156000000 | 0.10 | -0.65 | A>B |
| chr6 | 1.56E+08 | 156100000 | 0.14 | -0.64 | A>B |
| chr6 | 1.56E+08 | 156200000 | 0.09 | -0.65 | A>B |
| chr6 | 1.56E+08 | 156300000 | 0.03 | -0.81 | A>B |
| chr6 | 1.56E+08 | 156400000 | -0.07 | -0.71 | A>B |

| | | | | | |
|---|---|---|---|---|---|
| chr6 | 1.56E+08 | 156500000 | -0.09 | -0.78 | A>B |
| chr6 | 1.57E+08 | 156600000 | -0.17 | -0.87 | A>B |
| chr6 | 1.57E+08 | 156700000 | 0.02 | -0.85 | A>B |
| chr6 | 1.57E+08 | 156800000 | 0.07 | -0.78 | A>B |
| chr6 | 1.57E+08 | 156900000 | 0.13 | -0.50 | A>B |
| chr6 | 1.57E+08 | 157000000 | 0.28 | -0.23 | A>A |
| chr6 | 1.57E+08 | 157100000 | 0.61 | 0.20 | A>A |
| chr6 | 1.57E+08 | 157200000 | 0.77 | 0.43 | A>A |
| chr6 | 1.57E+08 | 157300000 | 0.72 | 0.53 | A>A |
| chr6 | 1.57E+08 | 157400000 | 0.78 | 0.49 | A>A |
| chr6 | 1.57E+08 | 157500000 | 0.84 | 0.65 | A>A |
| chr6 | 1.58E+08 | 157600000 | 0.61 | 0.54 | A>A |
| chr6 | 1.58E+08 | 157700000 | 0.00 | 0.58 | NO |
| chr6 | 1.58E+08 | 157800000 | 0.83 | 0.76 | A>A |
| chr6 | 1.58E+08 | 157900000 | 1.05 | 0.89 | A>A |
| chr6 | 1.58E+08 | 158000000 | 1.16 | 0.92 | A>A |
| chr6 | 1.58E+08 | 158100000 | 0.78 | 0.83 | A>A |
| chr6 | 1.58E+08 | 158200000 | 0.91 | 0.97 | A>A |
| chr6 | 1.58E+08 | 158300000 | 1.18 | 1.11 | A>A |
| chr6 | 1.58E+08 | 158400000 | 0.99 | 0.85 | A>A |
| chr6 | 1.58E+08 | 158500000 | 1.32 | 1.14 | A>A |
| chr6 | 1.59E+08 | 158600000 | 0.94 | 0.90 | A>A |
| chr6 | 1.59E+08 | 158700000 | 0.80 | 0.83 | A>A |
| chr6 | 1.59E+08 | 158800000 | 1.16 | 0.98 | A>A |
| chr6 | 1.59E+08 | 158900000 | 1.08 | 1.04 | A>A |
| chr6 | 1.59E+08 | 159000000 | 0.96 | 0.98 | A>A |
| chr6 | 1.59E+08 | 159100000 | 0.92 | 0.76 | A>A |
| chr6 | 1.59E+08 | 159200000 | 0.78 | 0.67 | A>A |
| chr6 | 1.59E+08 | 159300000 | 1.14 | 0.86 | A>A |
| chr6 | 1.59E+08 | 159400000 | 0.85 | 0.65 | A>A |
| chr6 | 1.59E+08 | 159500000 | 0.62 | 0.23 | A>A |
| chr6 | 1.6E+08 | 159600000 | 0.39 | -0.08 | A>A |
| chr6 | 1.6E+08 | 159700000 | 0.01 | -0.57 | A>B |
| chr6 | 1.6E+08 | 159800000 | -0.26 | -0.81 | B>B |
| chr6 | 1.6E+08 | 159900000 | -0.29 | -0.78 | B>B |
| chr6 | 1.6E+08 | 160000000 | -0.13 | -0.72 | A>B |
| chr6 | 1.6E+08 | 160100000 | 0.03 | -0.45 | B>B |
| chr6 | 1.6E+08 | 160200000 | 0.72 | 0.37 | A>A |
| chr6 | 1.6E+08 | 160300000 | 0.54 | 0.25 | A>A |
| chr6 | 1.6E+08 | 160400000 | 0.43 | 0.12 | A>A |
| chr6 | 1.6E+08 | 160500000 | 0.48 | 0.24 | A>A |
| chr6 | 1.61E+08 | 160600000 | 0.43 | 0.30 | A>A |
| chr6 | 1.61E+08 | 160700000 | 0.68 | 0.63 | A>A |
| chr6 | 1.61E+08 | 160800000 | 0.74 | 0.68 | A>A |
| chr6 | 1.61E+08 | 160900000 | 0.41 | 0.40 | A>A |
| chr6 | 1.61E+08 | 161000000 | -0.04 | -0.19 | B>B |
| chr6 | 1.61E+08 | 161100000 | 0.00 | -0.63 | NO |
| chr6 | 1.61E+08 | 161200000 | -0.21 | -0.49 | B>B |
| chr6 | 1.61E+08 | 161300000 | -0.07 | -0.38 | B>B |
| chr6 | 1.61E+08 | 161400000 | -0.01 | -0.27 | B>B |
| chr6 | 1.61E+08 | 161500000 | 0.33 | 0.08 | A>A |
| chr6 | 1.62E+08 | 161600000 | 0.13 | 0.10 | A>A |
| chr6 | 1.62E+08 | 161700000 | 0.30 | 0.28 | A>A |
| chr6 | 1.62E+08 | 161800000 | 0.11 | 0.12 | A>A |
| chr6 | 1.62E+08 | 161900000 | -0.12 | -0.14 | B>B |
| chr6 | 1.62E+08 | 162000000 | -0.15 | -0.02 | B>B |

| | | | | | |
|---|---|---|---|---|---|
| chr6 | 1.62E+08 | 162100000 | -0.18 | -0.21 | B>B |
| chr6 | 1.62E+08 | 162200000 | -0.25 | -0.17 | B>B |
| chr6 | 1.62E+08 | 162300000 | -0.13 | -0.19 | B>B |
| chr6 | 1.62E+08 | 162400000 | -0.01 | -0.09 | B>B |
| chr6 | 1.62E+08 | 162500000 | -0.09 | -0.08 | B>B |
| chr6 | 1.63E+08 | 162600000 | -0.20 | -0.21 | B>B |
| chr6 | 1.63E+08 | 162700000 | -0.27 | -0.41 | B>B |
| chr6 | 1.63E+08 | 162800000 | -0.39 | -0.40 | B>B |
| chr6 | 1.63E+08 | 162900000 | -0.60 | -0.47 | B>B |
| chr6 | 1.63E+08 | 163000000 | -0.64 | -0.67 | B>B |
| chr6 | 1.63E+08 | 163100000 | -0.74 | -0.72 | B>B |
| chr6 | 1.63E+08 | 163200000 | -0.55 | -0.64 | B>B |
| chr6 | 1.63E+08 | 163300000 | -0.51 | -0.72 | B>B |
| chr6 | 1.63E+08 | 163400000 | -0.47 | -0.75 | B>B |
| chr6 | 1.63E+08 | 163500000 | -0.41 | -0.73 | B>B |
| chr6 | 1.64E+08 | 163600000 | -0.16 | -0.54 | B>B |
| chr6 | 1.64E+08 | 163700000 | 0.26 | 0.03 | A>A |
| chr6 | 1.64E+08 | 163800000 | 0.32 | 0.22 | A>A |
| chr6 | 1.64E+08 | 163900000 | 0.25 | 0.15 | A>A |
| chr6 | 1.64E+08 | 164000000 | 0.38 | 0.26 | A>A |
| chr6 | 1.64E+08 | 164100000 | 0.00 | -0.39 | B>B |
| chr6 | 1.64E+08 | 164200000 | -0.09 | -0.57 | B>B |
| chr6 | 1.64E+08 | 164300000 | -0.34 | -0.68 | B>B |
| chr6 | 1.64E+08 | 164400000 | -0.32 | -0.60 | B>B |
| chr6 | 1.64E+08 | 164500000 | -0.46 | -0.77 | B>B |
| chr6 | 1.65E+08 | 164600000 | -0.52 | -0.85 | B>B |
| chr6 | 1.65E+08 | 164700000 | -0.80 | -0.83 | B>B |
| chr6 | 1.65E+08 | 164800000 | -0.90 | -0.96 | B>B |
| chr6 | 1.65E+08 | 164900000 | -0.94 | -0.93 | B>B |
| chr6 | 1.65E+08 | 165000000 | -0.88 | -0.97 | B>B |
| chr6 | 1.65E+08 | 165100000 | -0.97 | -0.95 | B>B |
| chr6 | 1.65E+08 | 165200000 | -0.87 | -0.89 | B>B |
| chr6 | 1.65E+08 | 165300000 | -0.87 | -0.77 | B>B |
| chr6 | 1.65E+08 | 165400000 | -0.70 | -0.57 | B>B |
| chr6 | 1.65E+08 | 165500000 | -0.67 | -0.54 | B>B |
| chr6 | 1.66E+08 | 165600000 | -0.63 | -0.24 | B>B |
| chr6 | 1.66E+08 | 165700000 | -0.40 | -0.15 | B>B |
| chr6 | 1.66E+08 | 165800000 | 0.11 | 0.40 | A>A |
| chr6 | 1.66E+08 | 165900000 | 0.43 | 0.74 | A>A |
| chr6 | 1.66E+08 | 166000000 | 0.63 | 1.02 | A>A |
| chr6 | 1.66E+08 | 166100000 | 0.63 | 0.91 | A>A |
| chr6 | 1.66E+08 | 166200000 | 0.54 | 0.77 | A>A |
| chr6 | 1.66E+08 | 166300000 | 0.50 | 0.97 | A>A |
| chr6 | 1.66E+08 | 166400000 | 0.96 | 1.25 | A>A |
| chr6 | 1.66E+08 | 166500000 | 1.04 | 1.31 | A>A |
| chr6 | 1.67E+08 | 166600000 | 0.58 | 0.74 | A>A |
| chr6 | 1.67E+08 | 166700000 | 0.69 | 0.79 | A>A |
| chr6 | 1.67E+08 | 166800000 | 0.90 | 0.90 | A>A |
| chr6 | 1.67E+08 | 166900000 | 0.63 | 0.65 | A>A |
| chr6 | 1.67E+08 | 167000000 | 0.74 | 0.85 | A>A |
| chr6 | 1.67E+08 | 167100000 | 0.70 | 0.82 | A>A |
| chr6 | 1.67E+08 | 167200000 | 0.62 | 0.66 | A>A |
| chr6 | 1.67E+08 | 167300000 | 0.56 | 0.55 | A>A |
| chr6 | 1.67E+08 | 167400000 | 0.69 | 0.61 | A>A |
| chr6 | 1.67E+08 | 167500000 | 0.80 | 0.76 | A>A |
| chr6 | 1.68E+08 | 167600000 | 0.64 | 0.40 | A>A |

| | | | | | |
|---|---|---|---|---|---|
| chr6 | 1.68E+08 | 167700000 | 0.72 | 0.48 | A>A |
| chr6 | 1.68E+08 | 167800000 | 0.66 | 0.52 | A>A |
| chr6 | 1.68E+08 | 167900000 | 0.52 | 0.40 | A>A |
| chr6 | 1.68E+08 | 168000000 | 0.00 | 0.00 | NO |
| chr6 | 1.68E+08 | 168100000 | 0.00 | 0.60 | NO |
| chr6 | 1.68E+08 | 168200000 | 0.67 | 0.66 | A>A |
| chr6 | 1.68E+08 | 168300000 | 0.80 | 0.70 | A>A |
| chr6 | 1.68E+08 | 168400000 | 0.87 | 0.86 | A>A |
| chr6 | 1.68E+08 | 168500000 | 0.85 | 0.73 | A>A |
| chr6 | 1.69E+08 | 168600000 | 0.48 | 0.53 | A>A |
| chr6 | 1.69E+08 | 168700000 | 0.91 | 0.81 | A>A |
| chr6 | 1.69E+08 | 168800000 | 0.51 | 0.46 | A>A |
| chr6 | 1.69E+08 | 168900000 | 0.46 | 0.28 | A>A |
| chr6 | 1.69E+08 | 169000000 | 0.34 | 0.32 | A>A |
| chr6 | 1.69E+08 | 169100000 | 0.40 | 0.31 | A>A |
| chr6 | 1.69E+08 | 169200000 | 0.40 | 0.31 | A>A |
| chr6 | 1.69E+08 | 169300000 | 0.30 | 0.23 | A>A |
| chr6 | 1.69E+08 | 169400000 | 0.30 | 0.31 | A>A |
| chr6 | 1.69E+08 | 169500000 | 0.52 | 0.40 | A>A |
| chr6 | 1.7E+08 | 169600000 | 0.56 | 0.56 | A>A |
| chr6 | 1.7E+08 | 169700000 | 0.65 | 0.67 | A>A |
| chr6 | 1.7E+08 | 169800000 | 0.47 | 0.57 | A>A |
| chr6 | 1.7E+08 | 169900000 | 0.45 | 0.60 | A>A |
| chr6 | 1.7E+08 | 170000000 | 0.66 | 0.66 | A>A |
| chr6 | 1.7E+08 | 170100000 | 0.70 | 0.64 | A>A |
| chr6 | 1.7E+08 | 170200000 | 0.78 | 0.72 | A>A |
| chr6 | 1.7E+08 | 170300000 | 0.53 | 0.57 | A>A |
| chr6 | 1.7E+08 | 170400000 | 0.49 | 0.60 | A>A |
| chr6 | 1.7E+08 | 170500000 | 0.64 | 0.69 | A>A |
| chr6 | 1.71E+08 | 170600000 | 0.71 | 0.70 | A>A |
| chr6 | 1.71E+08 | 170700000 | 0.67 | 0.74 | A>A |
| chr6 | 1.71E+08 | 170800000 | 0.66 | 0.58 | A>A |
| chr6 | 1.71E+08 | 170900000 | 0.57 | 0.44 | A>A |
| chr6 | 1.71E+08 | 171000000 | 0.00 | 0.00 | NO |
| chr6 | 1.71E+08 | 171100000 | 0.00 | 0.00 | NO |
| chr6 | 1.71E+08 | 171115067 | 0.00 | 0.00 | NO |
| chr7 | 0 | 100000 | 0.00 | 0.00 | NO |
| chr7 | 100000 | 200000 | 1.05 | 1.04 | A>A |
| chr7 | 200000 | 300000 | 0.00 | 0.00 | NO |
| chr7 | 300000 | 400000 | 0.72 | 0.83 | A>A |
| chr7 | 400000 | 500000 | 1.12 | 1.11 | A>A |
| chr7 | 500000 | 600000 | 1.18 | 1.10 | A>A |
| chr7 | 600000 | 700000 | 1.01 | 0.96 | A>A |
| chr7 | 700000 | 800000 | 0.99 | 1.04 | A>A |
| chr7 | 800000 | 900000 | 0.95 | 1.02 | A>A |
| chr7 | 900000 | 1000000 | 0.95 | 1.05 | A>A |
| chr7 | 1000000 | 1100000 | 1.12 | 1.10 | A>A |
| chr7 | 1100000 | 1200000 | 1.10 | 1.09 | A>A |
| chr7 | 1200000 | 1300000 | 0.87 | 0.89 | A>A |
| chr7 | 1300000 | 1400000 | 1.08 | 1.04 | A>A |
| chr7 | 1400000 | 1500000 | 1.01 | 0.92 | A>A |
| chr7 | 1500000 | 1600000 | 0.98 | 0.88 | A>A |
| chr7 | 1600000 | 1700000 | 0.91 | 0.83 | A>A |
| chr7 | 1700000 | 1800000 | 0.91 | 0.91 | A>A |
| chr7 | 1800000 | 1900000 | 0.83 | 0.94 | A>A |
| chr7 | 1900000 | 2000000 | 0.94 | 0.89 | A>A |

| | | | | | |
|---|---|---|---|---|---|
| chr7 | 2000000 | 2100000 | 0.91 | 0.90 | A>A |
| chr7 | 2100000 | 2200000 | 1.07 | 0.99 | A>A |
| chr7 | 2200000 | 2300000 | 1.08 | 0.94 | A>A |
| chr7 | 2300000 | 2400000 | 1.13 | 0.98 | A>A |
| chr7 | 2400000 | 2500000 | 1.11 | 0.90 | A>A |
| chr7 | 2500000 | 2600000 | 1.30 | 1.20 | A>A |
| chr7 | 2600000 | 2700000 | 1.24 | 1.17 | A>A |
| chr7 | 2700000 | 2800000 | 0.97 | 0.95 | A>A |
| chr7 | 2800000 | 2900000 | 0.98 | 1.06 | A>A |
| chr7 | 2900000 | 3000000 | 0.71 | 0.84 | A>A |
| chr7 | 3000000 | 3100000 | 0.86 | 0.88 | A>A |
| chr7 | 3100000 | 3200000 | 0.73 | 0.90 | A>A |
| chr7 | 3200000 | 3300000 | 0.78 | 0.89 | A>A |
| chr7 | 3300000 | 3400000 | 0.55 | 0.86 | A>A |
| chr7 | 3400000 | 3500000 | 0.56 | 0.84 | A>A |
| chr7 | 3500000 | 3600000 | 0.42 | 0.88 | A>A |
| chr7 | 3600000 | 3700000 | 0.40 | 0.85 | A>A |
| chr7 | 3700000 | 3800000 | 0.40 | 0.75 | A>A |
| chr7 | 3800000 | 3900000 | 0.44 | 0.84 | A>A |
| chr7 | 3900000 | 4000000 | 0.43 | 0.79 | A>A |
| chr7 | 4000000 | 4100000 | 0.61 | 0.74 | A>A |
| chr7 | 4100000 | 4200000 | 0.57 | 0.84 | A>A |
| chr7 | 4200000 | 4300000 | 0.66 | 0.81 | A>A |
| chr7 | 4300000 | 4400000 | 0.58 | 0.87 | A>A |
| chr7 | 4400000 | 4500000 | 0.51 | 0.87 | A>A |
| chr7 | 4500000 | 4600000 | 0.52 | 0.76 | A>A |
| chr7 | 4600000 | 4700000 | 0.80 | 0.94 | A>A |
| chr7 | 4700000 | 4800000 | 1.15 | 1.20 | A>A |
| chr7 | 4800000 | 4900000 | 0.98 | 1.06 | A>A |
| chr7 | 4900000 | 5000000 | 1.10 | 1.20 | A>A |
| chr7 | 5000000 | 5100000 | 0.00 | 0.00 | NO |
| chr7 | 5100000 | 5200000 | 1.06 | 1.06 | A>A |
| chr7 | 5200000 | 5300000 | 1.19 | 1.13 | A>A |
| chr7 | 5300000 | 5400000 | 0.00 | 0.00 | NO |
| chr7 | 5400000 | 5500000 | 1.44 | 1.39 | A>A |
| chr7 | 5500000 | 5600000 | 1.32 | 1.31 | A>A |
| chr7 | 5600000 | 5700000 | 1.60 | 1.47 | A>A |
| chr7 | 5700000 | 5800000 | 1.44 | 1.39 | A>A |
| chr7 | 5800000 | 5900000 | 1.44 | 1.41 | A>A |
| chr7 | 5900000 | 6000000 | 0.00 | 0.00 | NO |
| chr7 | 6000000 | 6100000 | 0.00 | 0.00 | NO |
| chr7 | 6100000 | 6200000 | 1.46 | 1.48 | A>A |
| chr7 | 6200000 | 6300000 | 1.72 | 1.73 | A>A |
| chr7 | 6300000 | 6400000 | 1.59 | 1.54 | A>A |
| chr7 | 6400000 | 6500000 | 1.49 | 1.54 | A>A |
| chr7 | 6500000 | 6600000 | 1.52 | 1.70 | A>A |
| chr7 | 6600000 | 6700000 | 1.50 | 1.61 | A>A |
| chr7 | 6700000 | 6800000 | 0.00 | 0.00 | NO |
| chr7 | 6800000 | 6900000 | 0.00 | 0.00 | NO |
| chr7 | 6900000 | 7000000 | 0.00 | 0.00 | NO |
| chr7 | 7000000 | 7100000 | 0.00 | 0.00 | NO |
| chr7 | 7100000 | 7200000 | 0.41 | 0.72 | A>A |
| chr7 | 7200000 | 7300000 | 0.67 | 0.87 | A>A |
| chr7 | 7300000 | 7400000 | 0.33 | 0.58 | A>A |
| chr7 | 7400000 | 7500000 | 0.27 | 0.79 | B>A |
| chr7 | 7500000 | 7600000 | 0.53 | 0.89 | A>A |

| | | | | | |
|---|---|---|---|---|---|
| chr7 | 7600000 | 7700000 | 0.52 | 0.56 | A>A |
| chr7 | 7700000 | 7800000 | 0.42 | 0.56 | A>A |
| chr7 | 7800000 | 7900000 | 0.38 | 0.55 | A>A |
| chr7 | 7900000 | 8000000 | 0.37 | 0.43 | A>A |
| chr7 | 8000000 | 8100000 | 0.13 | 0.37 | A>A |
| chr7 | 8100000 | 8200000 | 0.04 | 0.29 | A>A |
| chr7 | 8200000 | 8300000 | 0.03 | 0.25 | A>A |
| chr7 | 8300000 | 8400000 | -0.20 | 0.01 | B>B |
| chr7 | 8400000 | 8500000 | -0.64 | -0.50 | B>B |
| chr7 | 8500000 | 8600000 | -1.05 | -0.92 | B>B |
| chr7 | 8600000 | 8700000 | -1.10 | -1.03 | B>B |
| chr7 | 8700000 | 8800000 | -0.99 | -1.09 | B>B |
| chr7 | 8800000 | 8900000 | -1.04 | -1.03 | B>B |
| chr7 | 8900000 | 9000000 | -1.00 | -1.08 | B>B |
| chr7 | 9000000 | 9100000 | -1.00 | -1.00 | B>B |
| chr7 | 9100000 | 9200000 | -0.96 | -0.90 | B>B |
| chr7 | 9200000 | 9300000 | -0.97 | -0.87 | B>B |
| chr7 | 9300000 | 9400000 | -0.92 | -0.95 | B>B |
| chr7 | 9400000 | 9500000 | -0.83 | -0.92 | B>B |
| chr7 | 9500000 | 9600000 | -0.84 | -0.77 | B>B |
| chr7 | 9600000 | 9700000 | -0.86 | -0.85 | B>B |
| chr7 | 9700000 | 9800000 | -0.89 | -0.86 | B>B |
| chr7 | 9800000 | 9900000 | -0.81 | -0.86 | B>B |
| chr7 | 9900000 | 10000000 | -0.88 | -0.87 | B>B |
| chr7 | 10000000 | 10100000 | -0.86 | -0.89 | B>B |
| chr7 | 10100000 | 10200000 | -0.83 | -0.80 | B>B |
| chr7 | 10200000 | 10300000 | -0.89 | -0.89 | B>B |
| chr7 | 10300000 | 10400000 | -0.83 | -0.84 | B>B |
| chr7 | 10400000 | 10500000 | -0.79 | -0.80 | B>B |
| chr7 | 10500000 | 10600000 | -0.89 | -0.86 | B>B |
| chr7 | 10600000 | 10700000 | -0.90 | -0.96 | B>B |
| chr7 | 10700000 | 10800000 | -0.85 | -1.00 | B>B |
| chr7 | 10800000 | 10900000 | -0.78 | -0.96 | B>B |
| chr7 | 10900000 | 11000000 | -0.33 | -0.52 | B>B |
| chr7 | 11000000 | 11100000 | 0.24 | 0.10 | A>A |
| chr7 | 11100000 | 11200000 | 0.18 | 0.05 | A>A |
| chr7 | 11200000 | 11300000 | -0.31 | -0.14 | B>B |
| chr7 | 11300000 | 11400000 | -0.52 | -0.22 | B>B |
| chr7 | 11400000 | 11500000 | -0.32 | 0.38 | B>A |
| chr7 | 11500000 | 11600000 | -0.21 | 0.62 | B>A |
| chr7 | 11600000 | 11700000 | -0.19 | 0.58 | B>A |
| chr7 | 11700000 | 11800000 | -0.05 | 0.63 | B>A |
| chr7 | 11800000 | 11900000 | -0.11 | 0.59 | B>A |
| chr7 | 11900000 | 12000000 | -0.23 | 0.68 | B>A |
| chr7 | 12000000 | 12100000 | -0.13 | 0.62 | B>A |
| chr7 | 12100000 | 12200000 | 0.24 | 0.71 | A>A |
| chr7 | 12200000 | 12300000 | 0.31 | 0.85 | B>A |
| chr7 | 12300000 | 12400000 | -0.04 | 0.59 | B>A |
| chr7 | 12400000 | 12500000 | -0.08 | 0.62 | B>A |
| chr7 | 12500000 | 12600000 | 0.07 | 0.59 | B>A |
| chr7 | 12600000 | 12700000 | 0.16 | 0.70 | B>A |
| chr7 | 12700000 | 12800000 | 0.43 | 0.69 | A>A |
| chr7 | 12800000 | 12900000 | 0.57 | 0.87 | A>A |
| chr7 | 12900000 | 13000000 | 0.52 | 0.90 | A>A |
| chr7 | 13000000 | 13100000 | 0.54 | 0.83 | A>A |
| chr7 | 13100000 | 13200000 | 0.40 | 0.66 | A>A |

| | | | | | |
|---|---|---|---|---|---|
| chr7 | 13200000 | 13300000 | 0.12 | 0.55 | A>A |
| chr7 | 13300000 | 13400000 | -0.55 | 0.27 | B>A |
| chr7 | 13400000 | 13500000 | -0.80 | 0.27 | B>A |
| chr7 | 13500000 | 13600000 | -0.82 | 0.17 | B>A |
| chr7 | 13600000 | 13700000 | -0.76 | 0.26 | B>A |
| chr7 | 13700000 | 13800000 | -0.78 | 0.21 | B>A |
| chr7 | 13800000 | 13900000 | -0.58 | 0.30 | B>A |
| chr7 | 13900000 | 14000000 | -0.42 | 0.39 | B>A |
| chr7 | 14000000 | 14100000 | -0.44 | 0.30 | B>A |
| chr7 | 14100000 | 14200000 | -0.70 | 0.18 | B>A |
| chr7 | 14200000 | 14300000 | -0.76 | 0.19 | B>A |
| chr7 | 14300000 | 14400000 | -0.81 | 0.25 | B>A |
| chr7 | 14400000 | 14500000 | -0.82 | 0.19 | B>A |
| chr7 | 14500000 | 14600000 | -0.80 | 0.14 | B>A |
| chr7 | 14600000 | 14700000 | -0.85 | 0.02 | B>A |
| chr7 | 14700000 | 14800000 | -0.68 | 0.01 | B>A |
| chr7 | 14800000 | 14900000 | -0.92 | -0.07 | B>A |
| chr7 | 14900000 | 15000000 | -0.88 | -0.09 | B>A |
| chr7 | 15000000 | 15100000 | -0.79 | 0.01 | B>A |
| chr7 | 15100000 | 15200000 | -0.75 | -0.03 | B>A |
| chr7 | 15200000 | 15300000 | -0.70 | 0.00 | B>A |
| chr7 | 15300000 | 15400000 | -0.57 | 0.06 | B>A |
| chr7 | 15400000 | 15500000 | -0.57 | 0.05 | B>A |
| chr7 | 15500000 | 15600000 | -0.53 | 0.04 | B>A |
| chr7 | 15600000 | 15700000 | -0.50 | 0.19 | B>A |
| chr7 | 15700000 | 15800000 | -0.54 | 0.00 | B>A |
| chr7 | 15800000 | 15900000 | -0.60 | -0.10 | B>B |
| chr7 | 15900000 | 16000000 | -0.66 | -0.02 | B>A |
| chr7 | 16000000 | 16100000 | -0.47 | -0.06 | B>B |
| chr7 | 16100000 | 16200000 | -0.04 | 0.26 | A>A |
| chr7 | 16200000 | 16300000 | 0.09 | 0.45 | A>A |
| chr7 | 16300000 | 16400000 | 0.10 | 0.55 | A>A |
| chr7 | 16400000 | 16500000 | 0.31 | 0.49 | A>A |
| chr7 | 16500000 | 16600000 | 0.28 | 0.55 | A>A |
| chr7 | 16600000 | 16700000 | 0.42 | 0.57 | A>A |
| chr7 | 16700000 | 16800000 | 0.72 | 0.72 | A>A |
| chr7 | 16800000 | 16900000 | 0.89 | 0.90 | A>A |
| chr7 | 16900000 | 17000000 | 0.59 | 0.62 | A>A |
| chr7 | 17000000 | 17100000 | 0.72 | 0.64 | A>A |
| chr7 | 17100000 | 17200000 | 0.91 | 0.84 | A>A |
| chr7 | 17200000 | 17300000 | 0.89 | 0.85 | A>A |
| chr7 | 17300000 | 17400000 | 0.71 | 0.74 | A>A |
| chr7 | 17400000 | 17500000 | 0.76 | 0.75 | A>A |
| chr7 | 17500000 | 17600000 | 0.76 | 0.79 | A>A |
| chr7 | 17600000 | 17700000 | 0.63 | 0.62 | A>A |
| chr7 | 17700000 | 17800000 | 0.25 | 0.20 | A>A |
| chr7 | 17800000 | 17900000 | 0.34 | 0.27 | A>A |
| chr7 | 17900000 | 18000000 | 0.44 | 0.32 | A>A |
| chr7 | 18000000 | 18100000 | 0.22 | 0.16 | A>A |
| chr7 | 18100000 | 18200000 | 0.12 | 0.08 | A>A |
| chr7 | 18200000 | 18300000 | 0.09 | -0.05 | A>A |
| chr7 | 18300000 | 18400000 | -0.02 | -0.03 | B>B |
| chr7 | 18400000 | 18500000 | -0.01 | 0.02 | A>A |
| chr7 | 18500000 | 18600000 | -0.01 | 0.08 | A>A |
| chr7 | 18600000 | 18700000 | -0.16 | -0.30 | B>B |
| chr7 | 18700000 | 18800000 | -0.31 | -0.55 | B>B |

| | | | | | |
|---|---|---|---|---|---|
| chr7 | 18800000 | 18900000 | -0.34 | -0.59 | B>B |
| chr7 | 18900000 | 19000000 | -0.17 | -0.48 | B>B |
| chr7 | 19000000 | 19100000 | -0.15 | -0.40 | B>B |
| chr7 | 19100000 | 19200000 | -0.31 | -0.39 | B>B |
| chr7 | 19200000 | 19300000 | -0.50 | -0.60 | B>B |
| chr7 | 19300000 | 19400000 | -0.38 | -0.46 | B>B |
| chr7 | 19400000 | 19500000 | -0.31 | -0.36 | B>B |
| chr7 | 19500000 | 19600000 | -0.55 | -0.59 | B>B |
| chr7 | 19600000 | 19700000 | -0.61 | -0.67 | B>B |
| chr7 | 19700000 | 19800000 | -0.11 | -0.04 | B>B |
| chr7 | 19800000 | 19900000 | 0.34 | 0.42 | A>A |
| chr7 | 19900000 | 20000000 | 0.45 | 0.55 | A>A |
| chr7 | 20000000 | 20100000 | 0.46 | 0.50 | A>A |
| chr7 | 20100000 | 20200000 | 0.38 | 0.46 | A>A |
| chr7 | 20200000 | 20300000 | 0.22 | 0.42 | A>A |
| chr7 | 20300000 | 20400000 | 0.20 | 0.41 | A>A |
| chr7 | 20400000 | 20500000 | 0.16 | 0.53 | A>A |
| chr7 | 20500000 | 20600000 | -0.08 | 0.39 | A>A |
| chr7 | 20600000 | 20700000 | -0.12 | 0.17 | A>A |
| chr7 | 20700000 | 20800000 | -0.20 | -0.12 | B>B |
| chr7 | 20800000 | 20900000 | -0.25 | -0.43 | B>B |
| chr7 | 20900000 | 21000000 | -0.60 | -0.82 | B>B |
| chr7 | 21000000 | 21100000 | -0.70 | -0.97 | B>B |
| chr7 | 21100000 | 21200000 | -0.72 | -0.94 | B>B |
| chr7 | 21200000 | 21300000 | -0.51 | -0.65 | B>B |
| chr7 | 21300000 | 21400000 | -0.15 | -0.20 | B>B |
| chr7 | 21400000 | 21500000 | 0.06 | -0.14 | B>B |
| chr7 | 21500000 | 21600000 | -0.04 | -0.12 | B>B |
| chr7 | 21600000 | 21700000 | 0.05 | 0.11 | A>A |
| chr7 | 21700000 | 21800000 | -0.05 | -0.10 | B>B |
| chr7 | 21800000 | 21900000 | -0.28 | -0.34 | B>B |
| chr7 | 21900000 | 22000000 | -0.03 | -0.17 | B>B |
| chr7 | 22000000 | 22100000 | -0.19 | -0.36 | B>B |
| chr7 | 22100000 | 22200000 | -0.35 | -0.53 | B>B |
| chr7 | 22200000 | 22300000 | -0.67 | -0.78 | B>B |
| chr7 | 22300000 | 22400000 | -0.71 | -0.78 | B>B |
| chr7 | 22400000 | 22500000 | -0.55 | -0.73 | B>B |
| chr7 | 22500000 | 22600000 | -0.10 | -0.30 | B>B |
| chr7 | 22600000 | 22700000 | 0.59 | 0.53 | A>A |
| chr7 | 22700000 | 22800000 | 0.52 | 0.46 | A>A |
| chr7 | 22800000 | 22900000 | 0.51 | 0.40 | A>A |
| chr7 | 22900000 | 23000000 | 0.53 | 0.35 | A>A |
| chr7 | 23000000 | 23100000 | 0.53 | 0.39 | A>A |
| chr7 | 23100000 | 23200000 | 0.60 | 0.48 | A>A |
| chr7 | 23200000 | 23300000 | 0.71 | 0.58 | A>A |
| chr7 | 23300000 | 23400000 | 0.74 | 0.79 | A>A |
| chr7 | 23400000 | 23500000 | 0.87 | 0.82 | A>A |
| chr7 | 23500000 | 23600000 | 0.58 | 0.60 | A>A |
| chr7 | 23600000 | 23700000 | 0.19 | 0.15 | A>A |
| chr7 | 23700000 | 23800000 | 0.02 | 0.05 | A>A |
| chr7 | 23800000 | 23900000 | -0.39 | -0.15 | B>B |
| chr7 | 23900000 | 24000000 | -0.52 | -0.60 | B>B |
| chr7 | 24000000 | 24100000 | -0.62 | -0.80 | B>B |
| chr7 | 24100000 | 24200000 | -0.63 | -0.87 | B>B |
| chr7 | 24200000 | 24300000 | -0.62 | -0.92 | B>B |
| chr7 | 24300000 | 24400000 | -0.38 | -0.69 | B>B |

| | | | | | |
|---|---|---|---|---|---|
| chr7 | 24400000 | 24500000 | -0.14 | -0.41 | B>B |
| chr7 | 24500000 | 24600000 | 0.01 | -0.37 | B>B |
| chr7 | 24600000 | 24700000 | 0.10 | -0.19 | B>B |
| chr7 | 24700000 | 24800000 | 0.15 | 0.00 | A>A |
| chr7 | 24800000 | 24900000 | 0.44 | 0.26 | A>A |
| chr7 | 24900000 | 25000000 | 0.68 | 0.45 | A>A |
| chr7 | 25000000 | 25100000 | 0.56 | 0.42 | A>A |
| chr7 | 25100000 | 25200000 | 0.30 | 0.04 | A>A |
| chr7 | 25200000 | 25300000 | -0.19 | -0.51 | B>B |
| chr7 | 25300000 | 25400000 | -0.57 | -0.93 | B>B |
| chr7 | 25400000 | 25500000 | -0.62 | -0.98 | B>B |
| chr7 | 25500000 | 25600000 | -0.62 | -0.91 | B>B |
| chr7 | 25600000 | 25700000 | -0.58 | -0.85 | B>B |
| chr7 | 25700000 | 25800000 | -0.64 | -1.03 | B>B |
| chr7 | 25800000 | 25900000 | -0.38 | -0.74 | B>B |
| chr7 | 25900000 | 26000000 | -0.02 | -0.30 | B>B |
| chr7 | 26000000 | 26100000 | 0.10 | -0.44 | B>B |
| chr7 | 26100000 | 26200000 | 0.53 | 0.01 | A>A |
| chr7 | 26200000 | 26300000 | 0.85 | 0.33 | A>A |
| chr7 | 26300000 | 26400000 | 0.42 | 0.03 | A>A |
| chr7 | 26400000 | 26500000 | 0.17 | -0.22 | B>B |
| chr7 | 26500000 | 26600000 | -0.06 | -0.33 | B>B |
| chr7 | 26600000 | 26700000 | 0.07 | -0.34 | B>B |
| chr7 | 26700000 | 26800000 | 0.22 | 0.11 | A>A |
| chr7 | 26800000 | 26900000 | 0.37 | 0.23 | A>A |
| chr7 | 26900000 | 27000000 | 0.53 | 0.41 | A>A |
| chr7 | 27000000 | 27100000 | 0.58 | 0.43 | A>A |
| chr7 | 27100000 | 27200000 | 0.55 | 0.43 | A>A |
| chr7 | 27200000 | 27300000 | 0.53 | 0.43 | A>A |
| chr7 | 27300000 | 27400000 | 0.74 | 0.80 | A>A |
| chr7 | 27400000 | 27500000 | 0.76 | 0.82 | A>A |
| chr7 | 27500000 | 27600000 | 0.64 | 0.66 | A>A |
| chr7 | 27600000 | 27700000 | 0.64 | 0.57 | A>A |
| chr7 | 27700000 | 27800000 | 0.84 | 0.79 | A>A |
| chr7 | 27800000 | 27900000 | 0.67 | 0.67 | A>A |
| chr7 | 27900000 | 28000000 | 0.63 | 0.53 | A>A |
| chr7 | 28000000 | 28100000 | 0.72 | 0.49 | A>A |
| chr7 | 28100000 | 28200000 | 0.44 | 0.16 | A>A |
| chr7 | 28200000 | 28300000 | 0.08 | -0.39 | B>B |
| chr7 | 28300000 | 28400000 | -0.03 | -0.50 | B>B |
| chr7 | 28400000 | 28500000 | -0.21 | -0.60 | B>B |
| chr7 | 28500000 | 28600000 | -0.28 | -0.78 | B>B |
| chr7 | 28600000 | 28700000 | -0.38 | -0.69 | B>B |
| chr7 | 28700000 | 28800000 | -0.25 | -0.62 | B>B |
| chr7 | 28800000 | 28900000 | -0.28 | -0.68 | B>B |
| chr7 | 28900000 | 29000000 | -0.31 | -0.39 | B>B |
| chr7 | 29000000 | 29100000 | -0.53 | -0.55 | B>B |
| chr7 | 29100000 | 29200000 | -0.34 | -0.43 | B>B |
| chr7 | 29200000 | 29300000 | -0.16 | -0.38 | B>B |
| chr7 | 29300000 | 29400000 | -0.06 | -0.43 | B>B |
| chr7 | 29400000 | 29500000 | 0.03 | -0.17 | B>B |
| chr7 | 29500000 | 29600000 | 0.35 | 0.22 | A>A |
| chr7 | 29600000 | 29700000 | 0.36 | 0.21 | A>A |
| chr7 | 29700000 | 29800000 | 0.00 | 0.00 | NO |
| chr7 | 29800000 | 29900000 | 0.58 | 0.44 | A>A |
| chr7 | 29900000 | 30000000 | 0.69 | 0.50 | A>A |

| | | | | | |
|---|---|---|---|---|---|
| chr7 | 30000000 | 30100000 | 0.93 | 0.78 | A>A |
| chr7 | 30100000 | 30200000 | 0.78 | 0.71 | A>A |
| chr7 | 30200000 | 30300000 | 0.79 | 0.84 | A>A |
| chr7 | 30300000 | 30400000 | 0.97 | 0.92 | A>A |
| chr7 | 30400000 | 30500000 | 0.97 | 0.89 | A>A |
| chr7 | 30500000 | 30600000 | 0.91 | 0.85 | A>A |
| chr7 | 30600000 | 30700000 | 0.99 | 0.73 | A>A |
| chr7 | 30700000 | 30800000 | 0.80 | 0.54 | A>A |
| chr7 | 30800000 | 30900000 | 0.69 | 0.26 | A>A |
| chr7 | 30900000 | 31000000 | 0.61 | 0.24 | A>A |
| chr7 | 31000000 | 31100000 | 0.43 | -0.14 | A>A |
| chr7 | 31100000 | 31200000 | 0.25 | -0.25 | A>A |
| chr7 | 31200000 | 31300000 | -0.08 | -0.56 | B>B |
| chr7 | 31300000 | 31400000 | 0.00 | -0.56 | B>B |
| chr7 | 31400000 | 31500000 | 0.30 | -0.32 | A>B |
| chr7 | 31500000 | 31600000 | -0.02 | -0.51 | B>B |
| chr7 | 31600000 | 31700000 | -0.10 | -0.60 | B>B |
| chr7 | 31700000 | 31800000 | 0.06 | -0.42 | B>B |
| chr7 | 31800000 | 31900000 | 0.56 | 0.00 | A>A |
| chr7 | 31900000 | 32000000 | 0.80 | 0.39 | A>A |
| chr7 | 32000000 | 32100000 | 0.77 | 0.29 | A>A |
| chr7 | 32100000 | 32200000 | 0.66 | 0.10 | A>A |
| chr7 | 32200000 | 32300000 | 0.20 | -0.29 | B>B |
| chr7 | 32300000 | 32400000 | 0.23 | -0.54 | A>B |
| chr7 | 32400000 | 32500000 | 0.16 | -0.50 | A>B |
| chr7 | 32500000 | 32600000 | 0.60 | 0.20 | A>A |
| chr7 | 32600000 | 32700000 | 0.73 | 0.51 | A>A |
| chr7 | 32700000 | 32800000 | 0.00 | 0.00 | NO |
| chr7 | 32800000 | 32900000 | 0.00 | 0.00 | NO |
| chr7 | 32900000 | 33000000 | 0.94 | 0.94 | A>A |
| chr7 | 33000000 | 33100000 | 0.00 | 0.00 | NO |
| chr7 | 33100000 | 33200000 | 0.54 | 0.39 | A>A |
| chr7 | 33200000 | 33300000 | -0.06 | -0.22 | B>B |
| chr7 | 33300000 | 33400000 | -0.21 | -0.39 | B>B |
| chr7 | 33400000 | 33500000 | -0.23 | -0.50 | B>B |
| chr7 | 33500000 | 33600000 | -0.32 | -0.68 | B>B |
| chr7 | 33600000 | 33700000 | -0.19 | -0.70 | B>B |
| chr7 | 33700000 | 33800000 | -0.11 | -0.59 | B>B |
| chr7 | 33800000 | 33900000 | 0.08 | -0.58 | A>B |
| chr7 | 33900000 | 34000000 | 0.14 | -0.47 | A>B |
| chr7 | 34000000 | 34100000 | 0.28 | -0.55 | A>B |
| chr7 | 34100000 | 34200000 | 0.16 | -0.59 | A>B |
| chr7 | 34200000 | 34300000 | 0.00 | -0.45 | B>B |
| chr7 | 34300000 | 34400000 | -0.23 | -0.50 | B>B |
| chr7 | 34400000 | 34500000 | -0.43 | -0.52 | B>B |
| chr7 | 34500000 | 34600000 | -0.39 | -0.37 | B>B |
| chr7 | 34600000 | 34700000 | -0.24 | -0.18 | B>B |
| chr7 | 34700000 | 34800000 | 0.11 | 0.08 | A>A |
| chr7 | 34800000 | 34900000 | 0.37 | 0.19 | A>A |
| chr7 | 34900000 | 35000000 | 0.00 | 0.00 | NO |
| chr7 | 35000000 | 35100000 | 0.00 | 0.32 | NO |
| chr7 | 35100000 | 35200000 | 0.00 | 0.00 | NO |
| chr7 | 35200000 | 35300000 | 0.00 | 0.00 | NO |
| chr7 | 35300000 | 35400000 | -0.11 | -0.56 | B>B |
| chr7 | 35400000 | 35500000 | -0.07 | -0.53 | B>B |
| chr7 | 35500000 | 35600000 | 0.05 | -0.29 | B>B |

| | | | | | |
|---|---|---|---|---|---|
| chr7 | 35600000 | 35700000 | 0.27 | -0.08 | A>A |
| chr7 | 35700000 | 35800000 | 0.79 | 0.41 | A>A |
| chr7 | 35800000 | 35900000 | 0.89 | 0.64 | A>A |
| chr7 | 35900000 | 36000000 | 0.00 | 0.00 | NO |
| chr7 | 36000000 | 36100000 | 0.90 | 0.78 | A>A |
| chr7 | 36100000 | 36200000 | 1.19 | 0.96 | A>A |
| chr7 | 36200000 | 36300000 | 1.21 | 1.05 | A>A |
| chr7 | 36300000 | 36400000 | 0.84 | 0.73 | A>A |
| chr7 | 36400000 | 36500000 | 0.69 | 0.35 | A>A |
| chr7 | 36500000 | 36600000 | 0.16 | 0.00 | A>A |
| chr7 | 36600000 | 36700000 | 0.14 | -0.06 | A>A |
| chr7 | 36700000 | 36800000 | -0.15 | -0.24 | B>B |
| chr7 | 36800000 | 36900000 | -0.36 | -0.38 | B>B |
| chr7 | 36900000 | 37000000 | -0.31 | -0.28 | B>B |
| chr7 | 37000000 | 37100000 | -0.07 | -0.16 | B>B |
| chr7 | 37100000 | 37200000 | -0.18 | -0.29 | B>B |
| chr7 | 37200000 | 37300000 | -0.16 | -0.40 | B>B |
| chr7 | 37300000 | 37400000 | -0.17 | -0.70 | B>B |
| chr7 | 37400000 | 37500000 | -0.12 | -0.79 | A>B |
| chr7 | 37500000 | 37600000 | -0.22 | -0.83 | A>B |
| chr7 | 37600000 | 37700000 | -0.21 | -0.99 | A>B |
| chr7 | 37700000 | 37800000 | -0.15 | -1.01 | A>B |
| chr7 | 37800000 | 37900000 | -0.10 | -0.75 | A>B |
| chr7 | 37900000 | 38000000 | 0.15 | -0.20 | B>B |
| chr7 | 38000000 | 38100000 | 0.23 | -0.04 | A>A |
| chr7 | 38100000 | 38200000 | 0.05 | -0.19 | B>B |
| chr7 | 38200000 | 38300000 | 0.16 | -0.14 | A>A |
| chr7 | 38300000 | 38400000 | 0.15 | -0.23 | B>B |
| chr7 | 38400000 | 38500000 | -0.36 | -0.63 | B>B |
| chr7 | 38500000 | 38600000 | -0.71 | -0.92 | B>B |
| chr7 | 38600000 | 38700000 | -0.58 | -0.94 | B>B |
| chr7 | 38700000 | 38800000 | -0.30 | -0.55 | B>B |
| chr7 | 38800000 | 38900000 | -0.06 | -0.20 | B>B |
| chr7 | 38900000 | 39000000 | -0.14 | -0.28 | B>B |
| chr7 | 39000000 | 39100000 | -0.65 | -0.75 | B>B |
| chr7 | 39100000 | 39200000 | -0.61 | -0.92 | B>B |
| chr7 | 39200000 | 39300000 | -0.69 | -0.92 | B>B |
| chr7 | 39300000 | 39400000 | -0.60 | -0.89 | B>B |
| chr7 | 39400000 | 39500000 | -0.28 | -0.66 | B>B |
| chr7 | 39500000 | 39600000 | -0.13 | -0.39 | B>B |
| chr7 | 39600000 | 39700000 | 0.47 | 0.23 | A>A |
| chr7 | 39700000 | 39800000 | 0.50 | 0.36 | A>A |
| chr7 | 39800000 | 39900000 | 0.00 | 0.00 | NO |
| chr7 | 39900000 | 40000000 | 0.48 | 0.33 | A>A |
| chr7 | 40000000 | 40100000 | 0.71 | 0.60 | A>A |
| chr7 | 40100000 | 40200000 | 0.24 | 0.16 | A>A |
| chr7 | 40200000 | 40300000 | -0.27 | -0.46 | B>B |
| chr7 | 40300000 | 40400000 | -0.52 | -0.67 | B>B |
| chr7 | 40400000 | 40500000 | -0.78 | -0.92 | B>B |
| chr7 | 40500000 | 40600000 | -0.76 | -0.88 | B>B |
| chr7 | 40600000 | 40700000 | -0.62 | -0.74 | B>B |
| chr7 | 40700000 | 40800000 | -0.52 | -0.52 | B>B |
| chr7 | 40800000 | 40900000 | -0.57 | -0.39 | B>B |
| chr7 | 40900000 | 41000000 | -0.49 | -0.38 | B>B |
| chr7 | 41000000 | 41100000 | -0.29 | -0.26 | B>B |
| chr7 | 41100000 | 41200000 | -0.22 | -0.08 | B>B |

| | | | | | |
|---|---|---|---|---|---|
| chr7 | 41200000 | 41300000 | -0.23 | -0.10 | B>B |
| chr7 | 41300000 | 41400000 | -0.35 | -0.26 | B>B |
| chr7 | 41400000 | 41500000 | -0.44 | -0.45 | B>B |
| chr7 | 41500000 | 41600000 | -0.40 | -0.70 | B>B |
| chr7 | 41600000 | 41700000 | -0.45 | -0.69 | B>B |
| chr7 | 41700000 | 41800000 | -0.44 | -0.55 | B>B |
| chr7 | 41800000 | 41900000 | -0.77 | -0.87 | B>B |
| chr7 | 41900000 | 42000000 | -0.61 | -0.64 | B>B |
| chr7 | 42000000 | 42100000 | -0.44 | -0.38 | B>B |
| chr7 | 42100000 | 42200000 | -0.41 | -0.27 | B>B |
| chr7 | 42200000 | 42300000 | -0.28 | -0.27 | B>B |
| chr7 | 42300000 | 42400000 | -0.71 | -0.71 | B>B |
| chr7 | 42400000 | 42500000 | -0.71 | -0.86 | B>B |
| chr7 | 42500000 | 42600000 | -0.75 | -0.86 | B>B |
| chr7 | 42600000 | 42700000 | -0.76 | -0.84 | B>B |
| chr7 | 42700000 | 42800000 | -0.66 | -0.80 | B>B |
| chr7 | 42800000 | 42900000 | -0.50 | -0.61 | B>B |
| chr7 | 42900000 | 43000000 | 0.08 | 0.01 | A>A |
| chr7 | 43000000 | 43100000 | 0.13 | -0.14 | B>B |
| chr7 | 43100000 | 43200000 | 0.23 | -0.16 | A>A |
| chr7 | 43200000 | 43300000 | 0.40 | -0.08 | A>A |
| chr7 | 43300000 | 43400000 | 0.36 | 0.11 | A>A |
| chr7 | 43400000 | 43500000 | 0.35 | 0.26 | A>A |
| chr7 | 43500000 | 43600000 | 0.60 | 0.48 | A>A |
| chr7 | 43600000 | 43700000 | 0.69 | 0.67 | A>A |
| chr7 | 43700000 | 43800000 | 0.83 | 0.75 | A>A |
| chr7 | 43800000 | 43900000 | 0.94 | 0.77 | A>A |
| chr7 | 43900000 | 44000000 | 1.21 | 1.08 | A>A |
| chr7 | 44000000 | 44100000 | 0.00 | 0.00 | NO |
| chr7 | 44100000 | 44200000 | 1.57 | 1.49 | A>A |
| chr7 | 44200000 | 44300000 | 1.31 | 1.32 | A>A |
| chr7 | 44300000 | 44400000 | 1.27 | 1.22 | A>A |
| chr7 | 44400000 | 44500000 | 1.26 | 1.20 | A>A |
| chr7 | 44500000 | 44600000 | 1.17 | 1.17 | A>A |
| chr7 | 44600000 | 44700000 | 1.25 | 1.21 | A>A |
| chr7 | 44700000 | 44800000 | 1.38 | 1.29 | A>A |
| chr7 | 44800000 | 44900000 | 1.17 | 1.17 | A>A |
| chr7 | 44900000 | 45000000 | 1.10 | 1.01 | A>A |
| chr7 | 45000000 | 45100000 | 1.27 | 1.06 | A>A |
| chr7 | 45100000 | 45200000 | 0.92 | 0.84 | A>A |
| chr7 | 45200000 | 45300000 | 0.61 | 0.48 | A>A |
| chr7 | 45300000 | 45400000 | 0.42 | 0.50 | A>A |
| chr7 | 45400000 | 45500000 | 0.42 | 0.43 | A>A |
| chr7 | 45500000 | 45600000 | 0.44 | 0.39 | A>A |
| chr7 | 45600000 | 45700000 | 0.48 | 0.45 | A>A |
| chr7 | 45700000 | 45800000 | 0.61 | 0.69 | A>A |
| chr7 | 45800000 | 45900000 | 0.00 | 0.00 | NO |
| chr7 | 45900000 | 46000000 | 0.37 | 0.52 | A>A |
| chr7 | 46000000 | 46100000 | 0.80 | 0.84 | A>A |
| chr7 | 46100000 | 46200000 | 0.70 | 0.81 | A>A |
| chr7 | 46200000 | 46300000 | 0.69 | 0.86 | A>A |
| chr7 | 46300000 | 46400000 | 0.52 | 0.73 | A>A |
| chr7 | 46400000 | 46500000 | 0.39 | 0.78 | A>A |
| chr7 | 46500000 | 46600000 | 0.45 | 0.85 | A>A |
| chr7 | 46600000 | 46700000 | 0.54 | 0.85 | A>A |
| chr7 | 46700000 | 46800000 | 0.48 | 0.86 | A>A |

| | | | | | |
|---|---|---|---|---|---|
| chr7 | 46800000 | 46900000 | 0.39 | 0.88 | A>A |
| chr7 | 46900000 | 47000000 | 0.34 | 0.90 | B>A |
| chr7 | 47000000 | 47100000 | 0.37 | 0.82 | A>A |
| chr7 | 47100000 | 47200000 | 0.30 | 0.64 | A>A |
| chr7 | 47200000 | 47300000 | 0.35 | 0.53 | A>A |
| chr7 | 47300000 | 47400000 | 0.50 | 0.66 | A>A |
| chr7 | 47400000 | 47500000 | 0.73 | 0.85 | A>A |
| chr7 | 47500000 | 47600000 | 0.83 | 0.94 | A>A |
| chr7 | 47600000 | 47700000 | 1.00 | 0.93 | A>A |
| chr7 | 47700000 | 47800000 | 1.08 | 1.06 | A>A |
| chr7 | 47800000 | 47900000 | 0.82 | 0.66 | A>A |
| chr7 | 47900000 | 48000000 | 0.48 | 0.55 | A>A |
| chr7 | 48000000 | 48100000 | 0.48 | 0.34 | A>A |
| chr7 | 48100000 | 48200000 | 0.32 | 0.30 | A>A |
| chr7 | 48200000 | 48300000 | -0.15 | -0.22 | B>B |
| chr7 | 48300000 | 48400000 | -0.36 | -0.42 | B>B |
| chr7 | 48400000 | 48500000 | -0.42 | -0.48 | B>B |
| chr7 | 48500000 | 48600000 | -0.54 | -0.50 | B>B |
| chr7 | 48600000 | 48700000 | -0.69 | -0.57 | B>B |
| chr7 | 48700000 | 48800000 | -0.71 | -0.53 | B>B |
| chr7 | 48800000 | 48900000 | -0.67 | -0.53 | B>B |
| chr7 | 48900000 | 49000000 | -0.59 | -0.31 | B>B |
| chr7 | 49000000 | 49100000 | -0.69 | -0.53 | B>B |
| chr7 | 49100000 | 49200000 | -0.78 | -0.62 | B>B |
| chr7 | 49200000 | 49300000 | -0.72 | -0.72 | B>B |
| chr7 | 49300000 | 49400000 | -0.83 | -0.80 | B>B |
| chr7 | 49400000 | 49500000 | -0.84 | -0.84 | B>B |
| chr7 | 49500000 | 49600000 | -0.89 | -0.84 | B>B |
| chr7 | 49600000 | 49700000 | -0.82 | -0.83 | B>B |
| chr7 | 49700000 | 49800000 | -0.82 | -0.83 | B>B |
| chr7 | 49800000 | 49900000 | -0.70 | -0.73 | B>B |
| chr7 | 49900000 | 50000000 | -0.75 | -0.78 | B>B |
| chr7 | 50000000 | 50100000 | -0.73 | -0.80 | B>B |
| chr7 | 50100000 | 50200000 | -0.35 | -0.50 | B>B |
| chr7 | 50200000 | 50300000 | -0.37 | -0.62 | B>B |
| chr7 | 50300000 | 50400000 | -0.31 | -0.62 | B>B |
| chr7 | 50400000 | 50500000 | -0.13 | -0.32 | B>B |
| chr7 | 50500000 | 50600000 | 0.19 | 0.14 | A>A |
| chr7 | 50600000 | 50700000 | 0.35 | 0.23 | A>A |
| chr7 | 50700000 | 50800000 | 0.29 | 0.24 | A>A |
| chr7 | 50800000 | 50900000 | 0.45 | 0.45 | A>A |
| chr7 | 50900000 | 51000000 | 0.58 | 0.51 | A>A |
| chr7 | 51000000 | 51100000 | 0.39 | 0.40 | A>A |
| chr7 | 51100000 | 51200000 | 0.20 | 0.20 | A>A |
| chr7 | 51200000 | 51300000 | 0.07 | 0.04 | A>A |
| chr7 | 51300000 | 51400000 | 0.17 | 0.19 | A>A |
| chr7 | 51400000 | 51500000 | 0.00 | -0.08 | NO |
| chr7 | 51500000 | 51600000 | -0.50 | -0.60 | B>B |
| chr7 | 51600000 | 51700000 | -0.62 | -0.78 | B>B |
| chr7 | 51700000 | 51800000 | -0.70 | -0.82 | B>B |
| chr7 | 51800000 | 51900000 | -0.75 | -0.85 | B>B |
| chr7 | 51900000 | 52000000 | -0.73 | -0.80 | B>B |
| chr7 | 52000000 | 52100000 | -0.71 | -0.85 | B>B |
| chr7 | 52100000 | 52200000 | -0.71 | -0.83 | B>B |
| chr7 | 52200000 | 52300000 | -0.66 | -0.82 | B>B |
| chr7 | 52300000 | 52400000 | -0.72 | -0.81 | B>B |

| | | | | | |
|---|---|---|---|---|---|
| chr7 | 52400000 | 52500000 | -0.69 | -0.76 | B>B |
| chr7 | 52500000 | 52600000 | -0.75 | -0.79 | B>B |
| chr7 | 52600000 | 52700000 | -0.73 | -0.82 | B>B |
| chr7 | 52700000 | 52800000 | -0.73 | -0.74 | B>B |
| chr7 | 52800000 | 52900000 | -0.67 | -0.81 | B>B |
| chr7 | 52900000 | 53000000 | -0.75 | -0.80 | B>B |
| chr7 | 53000000 | 53100000 | -0.76 | -0.80 | B>B |
| chr7 | 53100000 | 53200000 | -0.68 | -0.79 | B>B |
| chr7 | 53200000 | 53300000 | -0.70 | -0.77 | B>B |
| chr7 | 53300000 | 53400000 | -0.72 | -0.83 | B>B |
| chr7 | 53400000 | 53500000 | -0.74 | -0.81 | B>B |
| chr7 | 53500000 | 53600000 | -0.77 | -0.84 | B>B |
| chr7 | 53600000 | 53700000 | -0.72 | -0.85 | B>B |
| chr7 | 53700000 | 53800000 | -0.73 | -0.83 | B>B |
| chr7 | 53800000 | 53900000 | -0.76 | -0.83 | B>B |
| chr7 | 53900000 | 54000000 | -0.75 | -0.85 | B>B |
| chr7 | 54000000 | 54100000 | -0.75 | -0.86 | B>B |
| chr7 | 54100000 | 54200000 | -0.80 | -0.85 | B>B |
| chr7 | 54200000 | 54300000 | -0.79 | -0.89 | B>B |
| chr7 | 54300000 | 54400000 | -0.85 | -0.91 | B>B |
| chr7 | 54400000 | 54500000 | -0.71 | -0.87 | B>B |
| chr7 | 54500000 | 54600000 | -0.70 | -0.88 | B>B |
| chr7 | 54600000 | 54700000 | -0.41 | -0.71 | B>B |
| chr7 | 54700000 | 54800000 | -0.16 | -0.34 | B>B |
| chr7 | 54800000 | 54900000 | 0.27 | 0.20 | A>A |
| chr7 | 54900000 | 55000000 | 0.69 | 0.55 | A>A |
| chr7 | 55000000 | 55100000 | 0.71 | 0.71 | A>A |
| chr7 | 55100000 | 55200000 | 0.46 | 0.59 | A>A |
| chr7 | 55200000 | 55300000 | 0.43 | 0.47 | A>A |
| chr7 | 55300000 | 55400000 | 0.21 | 0.31 | A>A |
| chr7 | 55400000 | 55500000 | 0.35 | 0.46 | A>A |
| chr7 | 55500000 | 55600000 | 0.29 | 0.53 | A>A |
| chr7 | 55600000 | 55700000 | 0.27 | 0.64 | A>A |
| chr7 | 55700000 | 55800000 | 0.00 | 0.00 | NO |
| chr7 | 55800000 | 55900000 | 0.00 | 0.00 | NO |
| chr7 | 55900000 | 56000000 | 0.17 | 0.90 | B>A |
| chr7 | 56000000 | 56100000 | 0.49 | 0.98 | A>A |
| chr7 | 56100000 | 56200000 | 0.37 | 0.86 | A>A |
| chr7 | 56200000 | 56300000 | -0.52 | 0.31 | B>A |
| chr7 | 56300000 | 56400000 | 0.00 | 0.21 | NO |
| chr7 | 56400000 | 56500000 | 0.00 | 0.00 | NO |
| chr7 | 56500000 | 56600000 | -0.86 | -0.25 | B>A |
| chr7 | 56600000 | 56700000 | 0.00 | 0.00 | NO |
| chr7 | 56700000 | 56800000 | 0.00 | 0.00 | NO |
| chr7 | 56800000 | 56900000 | 0.00 | 0.00 | NO |
| chr7 | 56900000 | 57000000 | 0.00 | 0.00 | NO |
| chr7 | 57000000 | 57100000 | 0.00 | 0.00 | NO |
| chr7 | 57100000 | 57200000 | 0.00 | 0.00 | NO |
| chr7 | 57200000 | 57300000 | 0.00 | 0.00 | NO |
| chr7 | 57300000 | 57400000 | 0.00 | 0.00 | NO |
| chr7 | 57400000 | 57500000 | 0.00 | -0.11 | NO |
| chr7 | 57500000 | 57600000 | 0.00 | 0.00 | NO |
| chr7 | 57600000 | 57700000 | 0.00 | 0.00 | NO |
| chr7 | 57700000 | 57800000 | 0.00 | 0.00 | NO |
| chr7 | 57800000 | 57900000 | 0.00 | 0.00 | NO |
| chr7 | 57900000 | 58000000 | 0.00 | 0.00 | NO |

| | | | | | |
|---|---|---|---|---|---|
| chr7 | 58000000 | 58100000 | 0.00 | 0.00 | NO |
| chr7 | 58100000 | 58200000 | 0.00 | 0.00 | NO |
| chr7 | 58200000 | 58300000 | 0.00 | 0.00 | NO |
| chr7 | 58300000 | 58400000 | 0.00 | 0.00 | NO |
| chr7 | 58400000 | 58500000 | 0.00 | 0.00 | NO |
| chr7 | 58500000 | 58600000 | 0.00 | 0.00 | NO |
| chr7 | 58600000 | 58700000 | 0.00 | 0.00 | NO |
| chr7 | 58700000 | 58800000 | 0.00 | 0.00 | NO |
| chr7 | 58800000 | 58900000 | 0.00 | 0.00 | NO |
| chr7 | 58900000 | 59000000 | 0.00 | 0.00 | NO |
| chr7 | 59000000 | 59100000 | 0.00 | 0.00 | NO |
| chr7 | 59100000 | 59200000 | 0.00 | 0.00 | NO |
| chr7 | 59200000 | 59300000 | 0.00 | 0.00 | NO |
| chr7 | 59300000 | 59400000 | 0.00 | 0.00 | NO |
| chr7 | 59400000 | 59500000 | 0.00 | 0.00 | NO |
| chr7 | 59500000 | 59600000 | 0.00 | 0.00 | NO |
| chr7 | 59600000 | 59700000 | 0.00 | 0.00 | NO |
| chr7 | 59700000 | 59800000 | 0.00 | 0.00 | NO |
| chr7 | 59800000 | 59900000 | 0.00 | 0.00 | NO |
| chr7 | 59900000 | 60000000 | 0.00 | 0.00 | NO |
| chr7 | 60000000 | 60100000 | 0.00 | 0.00 | NO |
| chr7 | 60100000 | 60200000 | 0.00 | 0.00 | NO |
| chr7 | 60200000 | 60300000 | 0.00 | 0.00 | NO |
| chr7 | 60300000 | 60400000 | 0.00 | 0.00 | NO |
| chr7 | 60400000 | 60500000 | 0.00 | 0.00 | NO |
| chr7 | 60500000 | 60600000 | 0.00 | 0.00 | NO |
| chr7 | 60600000 | 60700000 | 0.00 | 0.00 | NO |
| chr7 | 60700000 | 60800000 | 0.00 | 0.00 | NO |
| chr7 | 60800000 | 60900000 | 0.00 | 0.00 | NO |
| chr7 | 60900000 | 61000000 | 0.00 | 0.00 | NO |
| chr7 | 61000000 | 61100000 | 0.00 | 0.00 | NO |
| chr7 | 61100000 | 61200000 | 0.00 | 0.00 | NO |
| chr7 | 61200000 | 61300000 | 0.00 | 0.00 | NO |
| chr7 | 61300000 | 61400000 | 0.00 | 0.00 | NO |
| chr7 | 61400000 | 61500000 | 0.00 | 0.00 | NO |
| chr7 | 61500000 | 61600000 | 0.00 | 0.00 | NO |
| chr7 | 61600000 | 61700000 | 0.00 | 0.00 | NO |
| chr7 | 61700000 | 61800000 | 0.00 | -0.84 | NO |
| chr7 | 61800000 | 61900000 | 0.00 | 0.00 | NO |
| chr7 | 61900000 | 62000000 | 0.00 | 0.00 | NO |
| chr7 | 62000000 | 62100000 | 0.00 | 0.00 | NO |
| chr7 | 62100000 | 62200000 | 0.00 | 0.00 | NO |
| chr7 | 62200000 | 62300000 | 0.00 | 0.00 | NO |
| chr7 | 62300000 | 62400000 | 0.00 | 0.00 | NO |
| chr7 | 62400000 | 62500000 | 0.00 | 0.00 | NO |
| chr7 | 62500000 | 62600000 | 0.00 | 0.00 | NO |
| chr7 | 62600000 | 62700000 | 0.00 | 0.00 | NO |
| chr7 | 62700000 | 62800000 | 0.00 | 0.00 | NO |
| chr7 | 62800000 | 62900000 | 0.00 | 0.00 | NO |
| chr7 | 62900000 | 63000000 | 0.00 | 0.00 | NO |
| chr7 | 63000000 | 63100000 | 0.00 | 0.00 | NO |
| chr7 | 63100000 | 63200000 | 0.00 | 0.00 | NO |
| chr7 | 63200000 | 63300000 | 0.00 | 0.00 | NO |
| chr7 | 63300000 | 63400000 | 0.00 | 0.00 | NO |
| chr7 | 63400000 | 63500000 | 0.00 | 0.00 | NO |
| chr7 | 63500000 | 63600000 | -0.94 | 1.02 | B>A |

| | | | | | |
|---|---|---|---|---|---|
| chr7 | 63600000 | 63700000 | 0.00 | 0.00 | NO |
| chr7 | 63700000 | 63800000 | 0.00 | 0.00 | NO |
| chr7 | 63800000 | 63900000 | 0.00 | 0.00 | NO |
| chr7 | 63900000 | 64000000 | -0.67 | 1.17 | B>A |
| chr7 | 64000000 | 64100000 | -0.51 | 1.21 | B>A |
| chr7 | 64100000 | 64200000 | -0.37 | 1.15 | B>A |
| chr7 | 64200000 | 64300000 | -0.34 | 0.00 | NO |
| chr7 | 64300000 | 64400000 | -0.22 | 1.26 | B>A |
| chr7 | 64400000 | 64500000 | 0.00 | 0.00 | NO |
| chr7 | 64500000 | 64600000 | 0.00 | 0.00 | NO |
| chr7 | 64600000 | 64700000 | 0.00 | 0.00 | NO |
| chr7 | 64700000 | 64800000 | 0.30 | 1.36 | B>A |
| chr7 | 64800000 | 64900000 | 0.00 | 0.00 | NO |
| chr7 | 64900000 | 65000000 | 0.00 | 0.00 | NO |
| chr7 | 65000000 | 65100000 | 0.00 | 0.00 | NO |
| chr7 | 65100000 | 65200000 | 0.00 | 0.00 | NO |
| chr7 | 65200000 | 65300000 | 0.00 | 0.00 | NO |
| chr7 | 65300000 | 65400000 | 0.00 | 0.00 | NO |
| chr7 | 65400000 | 65500000 | 0.98 | 1.37 | A>A |
| chr7 | 65500000 | 65600000 | 1.15 | 0.00 | NO |
| chr7 | 65600000 | 65700000 | 1.09 | 1.49 | A>A |
| chr7 | 65700000 | 65800000 | 1.07 | 1.50 | A>A |
| chr7 | 65800000 | 65900000 | 0.00 | 0.00 | NO |
| chr7 | 65900000 | 66000000 | 0.00 | 0.00 | NO |
| chr7 | 66000000 | 66100000 | 1.32 | 0.00 | NO |
| chr7 | 66100000 | 66200000 | 1.26 | 1.45 | A>A |
| chr7 | 66200000 | 66300000 | 0.00 | 0.00 | NO |
| chr7 | 66300000 | 66400000 | 0.00 | 0.00 | NO |
| chr7 | 66400000 | 66500000 | 0.00 | 0.00 | NO |
| chr7 | 66500000 | 66600000 | 0.00 | 0.00 | NO |
| chr7 | 66600000 | 66700000 | 0.00 | 0.00 | NO |
| chr7 | 66700000 | 66800000 | 0.00 | 0.00 | NO |
| chr7 | 66800000 | 66900000 | 0.38 | 0.54 | A>A |
| chr7 | 66900000 | 67000000 | 0.38 | 0.46 | A>A |
| chr7 | 67000000 | 67100000 | 0.47 | 0.47 | A>A |
| chr7 | 67100000 | 67200000 | 0.60 | 0.72 | A>A |
| chr7 | 67200000 | 67300000 | 0.62 | 0.65 | A>A |
| chr7 | 67300000 | 67400000 | 0.57 | 0.62 | A>A |
| chr7 | 67400000 | 67500000 | 0.72 | 0.66 | A>A |
| chr7 | 67500000 | 67600000 | 0.69 | 0.66 | A>A |
| chr7 | 67600000 | 67700000 | 0.40 | 0.58 | A>A |
| chr7 | 67700000 | 67800000 | 0.53 | 0.68 | A>A |
| chr7 | 67800000 | 67900000 | 0.55 | 0.70 | A>A |
| chr7 | 67900000 | 68000000 | 0.52 | 0.65 | A>A |
| chr7 | 68000000 | 68100000 | 0.61 | 0.74 | A>A |
| chr7 | 68100000 | 68200000 | 0.56 | 0.66 | A>A |
| chr7 | 68200000 | 68300000 | 0.51 | 0.71 | A>A |
| chr7 | 68300000 | 68400000 | 0.59 | 0.75 | A>A |
| chr7 | 68400000 | 68500000 | 0.53 | 0.73 | A>A |
| chr7 | 68500000 | 68600000 | 0.52 | 0.50 | A>A |
| chr7 | 68600000 | 68700000 | 0.48 | 0.56 | A>A |
| chr7 | 68700000 | 68800000 | 0.55 | 0.65 | A>A |
| chr7 | 68800000 | 68900000 | 0.51 | 0.57 | A>A |
| chr7 | 68900000 | 69000000 | 0.48 | 0.59 | A>A |
| chr7 | 69000000 | 69100000 | 0.52 | 0.61 | A>A |
| chr7 | 69100000 | 69200000 | 0.49 | 0.67 | A>A |

| | | | | | |
|---|---|---|---|---|---|
| chr7 | 69200000 | 69300000 | 0.45 | 0.66 | A>A |
| chr7 | 69300000 | 69400000 | 0.29 | 0.51 | A>A |
| chr7 | 69400000 | 69500000 | 0.17 | 0.30 | A>A |
| chr7 | 69500000 | 69600000 | 0.02 | 0.21 | A>A |
| chr7 | 69600000 | 69700000 | -0.08 | 0.02 | B>B |
| chr7 | 69700000 | 69800000 | -0.11 | -0.14 | B>B |
| chr7 | 69800000 | 69900000 | -0.08 | -0.15 | B>B |
| chr7 | 69900000 | 70000000 | -0.07 | -0.12 | B>B |
| chr7 | 70000000 | 70100000 | 0.00 | 0.04 | A>A |
| chr7 | 70100000 | 70200000 | 0.04 | 0.12 | A>A |
| chr7 | 70200000 | 70300000 | -0.12 | 0.10 | B>B |
| chr7 | 70300000 | 70400000 | -0.42 | -0.44 | B>B |
| chr7 | 70400000 | 70500000 | -0.61 | -0.81 | B>B |
| chr7 | 70500000 | 70600000 | -0.70 | -0.83 | B>B |
| chr7 | 70600000 | 70700000 | -0.64 | -0.79 | B>B |
| chr7 | 70700000 | 70800000 | -0.65 | -0.84 | B>B |
| chr7 | 70800000 | 70900000 | -0.64 | -0.81 | B>B |
| chr7 | 70900000 | 71000000 | -0.57 | -0.73 | B>B |
| chr7 | 71000000 | 71100000 | -0.58 | -0.56 | B>B |
| chr7 | 71100000 | 71200000 | -0.47 | -0.40 | B>B |
| chr7 | 71200000 | 71300000 | -0.48 | -0.53 | B>B |
| chr7 | 71300000 | 71400000 | -0.60 | -0.82 | B>B |
| chr7 | 71400000 | 71500000 | -0.56 | -0.83 | B>B |
| chr7 | 71500000 | 71600000 | -0.59 | -0.87 | B>B |
| chr7 | 71600000 | 71700000 | -0.54 | -0.76 | B>B |
| chr7 | 71700000 | 71800000 | -0.38 | -0.60 | B>B |
| chr7 | 71800000 | 71900000 | -0.21 | -0.44 | B>B |
| chr7 | 71900000 | 72000000 | 0.01 | -0.14 | B>B |
| chr7 | 72000000 | 72100000 | 0.00 | 0.00 | NO |
| chr7 | 72100000 | 72200000 | 0.00 | 0.00 | NO |
| chr7 | 72200000 | 72300000 | 0.00 | 0.00 | NO |
| chr7 | 72300000 | 72400000 | 0.00 | 0.00 | NO |
| chr7 | 72400000 | 72500000 | 0.00 | 0.00 | NO |
| chr7 | 72500000 | 72600000 | 0.00 | 0.00 | NO |
| chr7 | 72600000 | 72700000 | 0.00 | 0.00 | NO |
| chr7 | 72700000 | 72800000 | 0.00 | 0.00 | NO |
| chr7 | 72800000 | 72900000 | 1.33 | 0.00 | NO |
| chr7 | 72900000 | 73000000 | 1.31 | 1.30 | A>A |
| chr7 | 73000000 | 73100000 | 1.29 | 1.38 | A>A |
| chr7 | 73100000 | 73200000 | 1.25 | 1.23 | A>A |
| chr7 | 73200000 | 73300000 | 1.02 | 1.14 | A>A |
| chr7 | 73300000 | 73400000 | 1.21 | 1.21 | A>A |
| chr7 | 73400000 | 73500000 | 1.02 | 1.07 | A>A |
| chr7 | 73500000 | 73600000 | 1.16 | 1.18 | A>A |
| chr7 | 73600000 | 73700000 | 1.01 | 1.10 | A>A |
| chr7 | 73700000 | 73800000 | 1.17 | 1.19 | A>A |
| chr7 | 73800000 | 73900000 | 0.97 | 1.10 | A>A |
| chr7 | 73900000 | 74000000 | 1.36 | 1.23 | A>A |
| chr7 | 74000000 | 74100000 | 1.27 | 1.21 | A>A |
| chr7 | 74100000 | 74200000 | 0.00 | 0.00 | NO |
| chr7 | 74200000 | 74300000 | 0.00 | 0.00 | NO |
| chr7 | 74300000 | 74400000 | 0.00 | 0.00 | NO |
| chr7 | 74400000 | 74500000 | 0.00 | 0.00 | NO |
| chr7 | 74500000 | 74600000 | 0.00 | 0.00 | NO |
| chr7 | 74600000 | 74700000 | 0.00 | 0.00 | NO |
| chr7 | 74700000 | 74800000 | 0.00 | 0.00 | NO |

| | | | | | |
|---|---|---|---|---|---|
| chr7 | 74800000 | 74900000 | 0.00 | 0.00 | NO |
| chr7 | 74900000 | 75000000 | 0.00 | 0.00 | NO |
| chr7 | 75000000 | 75100000 | 0.00 | 0.00 | NO |
| chr7 | 75100000 | 75200000 | 0.00 | 0.00 | NO |
| chr7 | 75200000 | 75300000 | 1.36 | 1.26 | A>A |
| chr7 | 75300000 | 75400000 | 1.41 | 1.36 | A>A |
| chr7 | 75400000 | 75500000 | 1.28 | 1.25 | A>A |
| chr7 | 75500000 | 75600000 | 1.49 | 1.41 | A>A |
| chr7 | 75600000 | 75700000 | 1.09 | 1.21 | A>A |
| chr7 | 75700000 | 75800000 | 0.00 | 0.00 | NO |
| chr7 | 75800000 | 75900000 | 1.44 | 1.38 | A>A |
| chr7 | 75900000 | 76000000 | 1.36 | 1.35 | A>A |
| chr7 | 76000000 | 76100000 | 0.00 | 0.00 | NO |
| chr7 | 76100000 | 76200000 | 0.00 | 0.00 | NO |
| chr7 | 76200000 | 76300000 | 0.00 | 0.00 | NO |
| chr7 | 76300000 | 76400000 | 0.00 | 0.00 | NO |
| chr7 | 76400000 | 76500000 | 0.00 | 0.00 | NO |
| chr7 | 76500000 | 76600000 | 0.00 | 0.00 | NO |
| chr7 | 76600000 | 76700000 | 0.00 | 0.00 | NO |
| chr7 | 76700000 | 76800000 | 0.00 | 0.00 | NO |
| chr7 | 76800000 | 76900000 | 0.79 | 0.97 | A>A |
| chr7 | 76900000 | 77000000 | 0.87 | 0.94 | A>A |
| chr7 | 77000000 | 77100000 | 0.90 | 0.97 | A>A |
| chr7 | 77100000 | 77200000 | 1.06 | 0.99 | A>A |
| chr7 | 77200000 | 77300000 | 0.81 | 0.72 | A>A |
| chr7 | 77300000 | 77400000 | 0.80 | 0.55 | A>A |
| chr7 | 77400000 | 77500000 | 0.66 | 0.44 | A>A |
| chr7 | 77500000 | 77600000 | 0.38 | 0.13 | A>A |
| chr7 | 77600000 | 77700000 | 0.00 | -0.32 | B>B |
| chr7 | 77700000 | 77800000 | -0.38 | -0.69 | B>B |
| chr7 | 77800000 | 77900000 | -0.56 | -0.95 | B>B |
| chr7 | 77900000 | 78000000 | -0.72 | -1.01 | B>B |
| chr7 | 78000000 | 78100000 | -0.65 | -1.05 | B>B |
| chr7 | 78100000 | 78200000 | -0.51 | -0.97 | B>B |
| chr7 | 78200000 | 78300000 | -0.52 | -1.01 | B>B |
| chr7 | 78300000 | 78400000 | -0.65 | -1.05 | B>B |
| chr7 | 78400000 | 78500000 | -0.65 | -1.05 | B>B |
| chr7 | 78500000 | 78600000 | -0.63 | -1.00 | B>B |
| chr7 | 78600000 | 78700000 | -0.56 | -0.99 | B>B |
| chr7 | 78700000 | 78800000 | -0.40 | -0.80 | B>B |
| chr7 | 78800000 | 78900000 | -0.22 | -0.34 | B>B |
| chr7 | 78900000 | 79000000 | 0.12 | 0.03 | A>A |
| chr7 | 79000000 | 79100000 | 0.35 | 0.15 | A>A |
| chr7 | 79100000 | 79200000 | 0.24 | 0.11 | A>A |
| chr7 | 79200000 | 79300000 | 0.13 | -0.03 | A>A |
| chr7 | 79300000 | 79400000 | 0.00 | -0.03 | B>B |
| chr7 | 79400000 | 79500000 | -0.10 | -0.10 | B>B |
| chr7 | 79500000 | 79600000 | -0.15 | -0.12 | B>B |
| chr7 | 79600000 | 79700000 | 0.05 | 0.06 | A>A |
| chr7 | 79700000 | 79800000 | 0.36 | 0.43 | A>A |
| chr7 | 79800000 | 79900000 | 0.58 | 0.65 | A>A |
| chr7 | 79900000 | 80000000 | 0.55 | 0.53 | A>A |
| chr7 | 80000000 | 80100000 | 0.31 | 0.43 | A>A |
| chr7 | 80100000 | 80200000 | 0.49 | 0.65 | A>A |
| chr7 | 80200000 | 80300000 | 0.50 | 0.59 | A>A |
| chr7 | 80300000 | 80400000 | 0.50 | 0.56 | A>A |

| | | | | | |
|---|---|---|---|---|---|
| chr7 | 80400000 | 80500000 | 0.68 | 0.80 | A>A |
| chr7 | 80500000 | 80600000 | 0.65 | 0.67 | A>A |
| chr7 | 80600000 | 80700000 | 0.85 | 0.83 | A>A |
| chr7 | 80700000 | 80800000 | 0.84 | 0.86 | A>A |
| chr7 | 80800000 | 80900000 | 0.88 | 0.81 | A>A |
| chr7 | 80900000 | 81000000 | 0.81 | 0.85 | A>A |
| chr7 | 81000000 | 81100000 | 0.60 | 0.68 | A>A |
| chr7 | 81100000 | 81200000 | 0.36 | 0.65 | A>A |
| chr7 | 81200000 | 81300000 | 0.69 | 0.89 | A>A |
| chr7 | 81300000 | 81400000 | 0.63 | 0.80 | A>A |
| chr7 | 81400000 | 81500000 | 0.64 | 0.87 | A>A |
| chr7 | 81500000 | 81600000 | 0.48 | 0.74 | A>A |
| chr7 | 81600000 | 81700000 | 0.37 | 0.51 | A>A |
| chr7 | 81700000 | 81800000 | 0.61 | 0.68 | A>A |
| chr7 | 81800000 | 81900000 | 0.52 | 0.58 | A>A |
| chr7 | 81900000 | 82000000 | 0.66 | 0.65 | A>A |
| chr7 | 82000000 | 82100000 | 0.58 | 0.64 | A>A |
| chr7 | 82100000 | 82200000 | 0.51 | 0.61 | A>A |
| chr7 | 82200000 | 82300000 | 0.30 | 0.48 | A>A |
| chr7 | 82300000 | 82400000 | 0.19 | 0.45 | A>A |
| chr7 | 82400000 | 82500000 | 0.02 | 0.36 | A>A |
| chr7 | 82500000 | 82600000 | 0.03 | 0.40 | A>A |
| chr7 | 82600000 | 82700000 | 0.11 | 0.46 | A>A |
| chr7 | 82700000 | 82800000 | 0.06 | 0.44 | A>A |
| chr7 | 82800000 | 82900000 | -0.04 | 0.33 | A>A |
| chr7 | 82900000 | 83000000 | -0.07 | 0.31 | A>A |
| chr7 | 83000000 | 83100000 | 0.00 | 0.29 | A>A |
| chr7 | 83100000 | 83200000 | 0.19 | 0.44 | A>A |
| chr7 | 83200000 | 83300000 | 0.10 | 0.33 | A>A |
| chr7 | 83300000 | 83400000 | -0.17 | 0.15 | B>B |
| chr7 | 83400000 | 83500000 | -0.27 | 0.01 | B>B |
| chr7 | 83500000 | 83600000 | -0.22 | -0.01 | B>B |
| chr7 | 83600000 | 83700000 | 0.21 | 0.25 | A>A |
| chr7 | 83700000 | 83800000 | 0.49 | 0.42 | A>A |
| chr7 | 83800000 | 83900000 | 0.43 | 0.34 | A>A |
| chr7 | 83900000 | 84000000 | 0.40 | 0.40 | A>A |
| chr7 | 84000000 | 84100000 | 0.34 | 0.35 | A>A |
| chr7 | 84100000 | 84200000 | 0.38 | 0.43 | A>A |
| chr7 | 84200000 | 84300000 | 0.44 | 0.38 | A>A |
| chr7 | 84300000 | 84400000 | 0.38 | 0.31 | A>A |
| chr7 | 84400000 | 84500000 | 0.26 | 0.24 | A>A |
| chr7 | 84500000 | 84600000 | 0.16 | 0.18 | A>A |
| chr7 | 84600000 | 84700000 | -0.21 | -0.15 | B>B |
| chr7 | 84700000 | 84800000 | -0.60 | -0.54 | B>B |
| chr7 | 84800000 | 84900000 | -0.69 | -0.64 | B>B |
| chr7 | 84900000 | 85000000 | -0.87 | -0.73 | B>B |
| chr7 | 85000000 | 85100000 | -0.76 | -0.59 | B>B |
| chr7 | 85100000 | 85200000 | -0.62 | -0.25 | B>B |
| chr7 | 85200000 | 85300000 | -0.45 | -0.03 | B>B |
| chr7 | 85300000 | 85400000 | -0.40 | -0.10 | B>B |
| chr7 | 85400000 | 85500000 | -0.53 | -0.34 | B>B |
| chr7 | 85500000 | 85600000 | -0.76 | -0.82 | B>B |
| chr7 | 85600000 | 85700000 | -0.93 | -0.99 | B>B |
| chr7 | 85700000 | 85800000 | -0.98 | -1.10 | B>B |
| chr7 | 85800000 | 85900000 | -1.00 | -1.05 | B>B |
| chr7 | 85900000 | 86000000 | -0.95 | -1.10 | B>B |

| | | | | | |
|---|---|---|---|---|---|
| chr7 | 86000000 | 86100000 | -0.87 | -1.09 | B>B |
| chr7 | 86100000 | 86200000 | -0.85 | -1.03 | B>B |
| chr7 | 86200000 | 86300000 | -0.88 | -1.09 | B>B |
| chr7 | 86300000 | 86400000 | -0.90 | -1.11 | B>B |
| chr7 | 86400000 | 86500000 | -0.81 | -1.02 | B>B |
| chr7 | 86500000 | 86600000 | -0.56 | -0.83 | B>B |
| chr7 | 86600000 | 86700000 | -0.68 | -0.93 | B>B |
| chr7 | 86700000 | 86800000 | -0.40 | -0.68 | B>B |
| chr7 | 86800000 | 86900000 | -0.02 | -0.30 | B>B |
| chr7 | 86900000 | 87000000 | -0.02 | -0.13 | B>B |
| chr7 | 87000000 | 87100000 | -0.06 | -0.19 | B>B |
| chr7 | 87100000 | 87200000 | -0.35 | -0.33 | B>B |
| chr7 | 87200000 | 87300000 | -0.46 | -0.49 | B>B |
| chr7 | 87300000 | 87400000 | -0.58 | -0.55 | B>B |
| chr7 | 87400000 | 87500000 | -0.14 | -0.23 | B>B |
| chr7 | 87500000 | 87600000 | -0.01 | -0.14 | B>B |
| chr7 | 87600000 | 87700000 | -0.43 | -0.38 | B>B |
| chr7 | 87700000 | 87800000 | -0.27 | -0.27 | B>B |
| chr7 | 87800000 | 87900000 | -0.12 | -0.23 | B>B |
| chr7 | 87900000 | 88000000 | -0.32 | -0.48 | B>B |
| chr7 | 88000000 | 88100000 | -0.56 | -0.80 | B>B |
| chr7 | 88100000 | 88200000 | -0.68 | -0.87 | B>B |
| chr7 | 88200000 | 88300000 | -0.81 | -0.91 | B>B |
| chr7 | 88300000 | 88400000 | -0.74 | -0.87 | B>B |
| chr7 | 88400000 | 88500000 | -0.70 | -0.89 | B>B |
| chr7 | 88500000 | 88600000 | -0.92 | -0.90 | B>B |
| chr7 | 88600000 | 88700000 | -0.90 | -0.93 | B>B |
| chr7 | 88700000 | 88800000 | -0.97 | -0.95 | B>B |
| chr7 | 88800000 | 88900000 | -0.92 | -0.93 | B>B |
| chr7 | 88900000 | 89000000 | -0.88 | -0.92 | B>B |
| chr7 | 89000000 | 89100000 | -0.91 | -0.93 | B>B |
| chr7 | 89100000 | 89200000 | -0.93 | -0.94 | B>B |
| chr7 | 89200000 | 89300000 | -0.87 | -0.89 | B>B |
| chr7 | 89300000 | 89400000 | -0.84 | -0.89 | B>B |
| chr7 | 89400000 | 89500000 | -0.76 | -0.92 | B>B |
| chr7 | 89500000 | 89600000 | -0.66 | -0.99 | B>B |
| chr7 | 89600000 | 89700000 | -0.62 | -0.87 | B>B |
| chr7 | 89700000 | 89800000 | 0.00 | 0.00 | NO |
| chr7 | 89800000 | 89900000 | 0.27 | 0.13 | A>A |
| chr7 | 89900000 | 90000000 | 0.19 | -0.03 | A>A |
| chr7 | 90000000 | 90100000 | -0.08 | -0.14 | B>B |
| chr7 | 90100000 | 90200000 | -0.51 | -0.40 | B>B |
| chr7 | 90200000 | 90300000 | -0.51 | -0.30 | B>B |
| chr7 | 90300000 | 90400000 | -0.68 | -0.54 | B>B |
| chr7 | 90400000 | 90500000 | -0.77 | -0.69 | B>B |
| chr7 | 90500000 | 90600000 | -0.69 | -0.77 | B>B |
| chr7 | 90600000 | 90700000 | -0.61 | -0.70 | B>B |
| chr7 | 90700000 | 90800000 | -0.62 | -0.75 | B>B |
| chr7 | 90800000 | 90900000 | -0.53 | -0.71 | B>B |
| chr7 | 90900000 | 91000000 | -0.58 | -0.81 | B>B |
| chr7 | 91000000 | 91100000 | -0.69 | -0.89 | B>B |
| chr7 | 91100000 | 91200000 | -0.61 | -0.70 | B>B |
| chr7 | 91200000 | 91300000 | -0.58 | -0.74 | B>B |
| chr7 | 91300000 | 91400000 | -0.62 | -0.66 | B>B |
| chr7 | 91400000 | 91500000 | -0.40 | -0.39 | B>B |
| chr7 | 91500000 | 91600000 | 0.26 | 0.09 | A>A |

| | | | | | |
|---|---|---|---|---|---|
| chr7 | 91600000 | 91700000 | 0.43 | 0.40 | A>A |
| chr7 | 91700000 | 91800000 | 0.53 | 0.57 | A>A |
| chr7 | 91800000 | 91900000 | 0.61 | 0.59 | A>A |
| chr7 | 91900000 | 92000000 | 0.63 | 0.71 | A>A |
| chr7 | 92000000 | 92100000 | 0.38 | 0.49 | A>A |
| chr7 | 92100000 | 92200000 | 0.44 | 0.67 | A>A |
| chr7 | 92200000 | 92300000 | 0.35 | 0.59 | A>A |
| chr7 | 92300000 | 92400000 | 0.51 | 0.69 | A>A |
| chr7 | 92400000 | 92500000 | 0.36 | 0.73 | A>A |
| chr7 | 92500000 | 92600000 | -0.16 | 0.42 | B>A |
| chr7 | 92600000 | 92700000 | -0.18 | 0.29 | A>A |
| chr7 | 92700000 | 92800000 | -0.37 | -0.04 | B>B |
| chr7 | 92800000 | 92900000 | -0.27 | -0.11 | B>B |
| chr7 | 92900000 | 93000000 | -0.19 | -0.22 | B>B |
| chr7 | 93000000 | 93100000 | -0.55 | -0.50 | B>B |
| chr7 | 93100000 | 93200000 | -0.47 | -0.40 | B>B |
| chr7 | 93200000 | 93300000 | -0.19 | -0.19 | B>B |
| chr7 | 93300000 | 93400000 | -0.08 | -0.09 | B>B |
| chr7 | 93400000 | 93500000 | 0.18 | 0.09 | A>A |
| chr7 | 93500000 | 93600000 | 0.47 | 0.37 | A>A |
| chr7 | 93600000 | 93700000 | 0.46 | 0.40 | A>A |
| chr7 | 93700000 | 93800000 | 0.44 | 0.43 | A>A |
| chr7 | 93800000 | 93900000 | 0.38 | 0.29 | A>A |
| chr7 | 93900000 | 94000000 | 0.36 | 0.23 | A>A |
| chr7 | 94000000 | 94100000 | 0.29 | 0.22 | A>A |
| chr7 | 94100000 | 94200000 | 0.47 | 0.48 | A>A |
| chr7 | 94200000 | 94300000 | 0.48 | 0.54 | A>A |
| chr7 | 94300000 | 94400000 | 0.47 | 0.59 | A>A |
| chr7 | 94400000 | 94500000 | 0.43 | 0.59 | A>A |
| chr7 | 94500000 | 94600000 | 0.60 | 0.59 | A>A |
| chr7 | 94600000 | 94700000 | 0.63 | 0.60 | A>A |
| chr7 | 94700000 | 94800000 | 0.57 | 0.51 | A>A |
| chr7 | 94800000 | 94900000 | 0.44 | 0.37 | A>A |
| chr7 | 94900000 | 95000000 | 0.64 | 0.45 | A>A |
| chr7 | 95000000 | 95100000 | 0.64 | 0.49 | A>A |
| chr7 | 95100000 | 95200000 | 0.26 | 0.20 | A>A |
| chr7 | 95200000 | 95300000 | 0.25 | 0.28 | A>A |
| chr7 | 95300000 | 95400000 | -0.19 | -0.27 | B>B |
| chr7 | 95400000 | 95500000 | -0.40 | -0.64 | B>B |
| chr7 | 95500000 | 95600000 | -0.45 | -0.66 | B>B |
| chr7 | 95600000 | 95700000 | -0.47 | -0.70 | B>B |
| chr7 | 95700000 | 95800000 | -0.12 | -0.20 | B>B |
| chr7 | 95800000 | 95900000 | 0.13 | -0.14 | B>B |
| chr7 | 95900000 | 96000000 | 0.02 | -0.42 | B>B |
| chr7 | 96000000 | 96100000 | -0.40 | -0.81 | B>B |
| chr7 | 96100000 | 96200000 | -0.56 | -0.90 | B>B |
| chr7 | 96200000 | 96300000 | -0.55 | -0.79 | B>B |
| chr7 | 96300000 | 96400000 | -0.34 | -0.63 | B>B |
| chr7 | 96400000 | 96500000 | -0.65 | -0.98 | B>B |
| chr7 | 96500000 | 96600000 | -0.64 | -0.91 | B>B |
| chr7 | 96600000 | 96700000 | -0.34 | -0.54 | B>B |
| chr7 | 96700000 | 96800000 | -0.13 | -0.38 | B>B |
| chr7 | 96800000 | 96900000 | -0.48 | -0.88 | B>B |
| chr7 | 96900000 | 97000000 | -0.54 | -1.11 | B>B |
| chr7 | 97000000 | 97100000 | -0.54 | -1.16 | A>B |
| chr7 | 97100000 | 97200000 | -0.63 | -1.12 | B>B |

| | | | | | |
|---|---|---|---|---|---|
| chr7 | 97200000 | 97300000 | -0.72 | -1.13 | B>B |
| chr7 | 97300000 | 97400000 | -0.66 | -1.00 | B>B |
| chr7 | 97400000 | 97500000 | -0.30 | -0.51 | B>B |
| chr7 | 97500000 | 97600000 | 0.00 | 0.00 | NO |
| chr7 | 97600000 | 97700000 | 0.69 | 0.46 | A>A |
| chr7 | 97700000 | 97800000 | 0.85 | 0.64 | A>A |
| chr7 | 97800000 | 97900000 | 0.93 | 0.82 | A>A |
| chr7 | 97900000 | 98000000 | 1.04 | 1.21 | A>A |
| chr7 | 98000000 | 98100000 | 0.92 | 1.05 | A>A |
| chr7 | 98100000 | 98200000 | 0.97 | 1.02 | A>A |
| chr7 | 98200000 | 98300000 | 0.87 | 1.05 | A>A |
| chr7 | 98300000 | 98400000 | 1.00 | 1.00 | A>A |
| chr7 | 98400000 | 98500000 | 0.95 | 1.02 | A>A |
| chr7 | 98500000 | 98600000 | 1.25 | 1.28 | A>A |
| chr7 | 98600000 | 98700000 | 1.16 | 1.14 | A>A |
| chr7 | 98700000 | 98800000 | 1.30 | 1.38 | A>A |
| chr7 | 98800000 | 98900000 | 1.21 | 1.25 | A>A |
| chr7 | 98900000 | 99000000 | 1.12 | 1.21 | A>A |
| chr7 | 99000000 | 99100000 | 1.28 | 1.24 | A>A |
| chr7 | 99100000 | 99200000 | 1.19 | 1.34 | A>A |
| chr7 | 99200000 | 99300000 | 1.03 | 1.05 | A>A |
| chr7 | 99300000 | 99400000 | 0.00 | 0.00 | NO |
| chr7 | 99400000 | 99500000 | 0.97 | 1.04 | A>A |
| chr7 | 99500000 | 99600000 | 1.12 | 1.18 | A>A |
| chr7 | 99600000 | 99700000 | 1.17 | 1.21 | A>A |
| chr7 | 99700000 | 99800000 | 1.14 | 1.15 | A>A |
| chr7 | 99800000 | 99900000 | 0.00 | 0.00 | NO |
| chr7 | 99900000 | 100000000 | 0.00 | 0.00 | NO |
| chr7 | 1E+08 | 100100000 | 1.02 | 0.97 | A>A |
| chr7 | 1E+08 | 100200000 | 1.19 | 1.16 | A>A |
| chr7 | 1E+08 | 100300000 | 1.07 | 1.05 | A>A |
| chr7 | 1E+08 | 100400000 | 1.03 | 1.10 | A>A |
| chr7 | 1E+08 | 100500000 | 0.93 | 0.95 | A>A |
| chr7 | 1.01E+08 | 100600000 | 0.00 | 0.00 | NO |
| chr7 | 1.01E+08 | 100700000 | 1.02 | 1.10 | A>A |
| chr7 | 1.01E+08 | 100800000 | 1.03 | 1.08 | A>A |
| chr7 | 1.01E+08 | 100900000 | 1.12 | 1.22 | A>A |
| chr7 | 1.01E+08 | 101000000 | 0.91 | 1.03 | A>A |
| chr7 | 1.01E+08 | 101100000 | 1.12 | 1.21 | A>A |
| chr7 | 1.01E+08 | 101200000 | 0.93 | 1.10 | A>A |
| chr7 | 1.01E+08 | 101300000 | 0.88 | 1.01 | A>A |
| chr7 | 1.01E+08 | 101400000 | 1.01 | 1.13 | A>A |
| chr7 | 1.01E+08 | 101500000 | 0.90 | 1.08 | A>A |
| chr7 | 1.02E+08 | 101600000 | 1.01 | 1.21 | A>A |
| chr7 | 1.02E+08 | 101700000 | 1.03 | 1.22 | A>A |
| chr7 | 1.02E+08 | 101800000 | 0.98 | 1.21 | A>A |
| chr7 | 1.02E+08 | 101900000 | 1.00 | 1.08 | A>A |
| chr7 | 1.02E+08 | 102000000 | 1.02 | 1.16 | A>A |
| chr7 | 1.02E+08 | 102100000 | 1.32 | 1.42 | A>A |
| chr7 | 1.02E+08 | 102200000 | 0.00 | 0.00 | NO |
| chr7 | 1.02E+08 | 102300000 | 0.00 | 0.00 | NO |
| chr7 | 1.02E+08 | 102400000 | 0.00 | 0.00 | NO |
| chr7 | 1.02E+08 | 102500000 | 0.00 | 0.00 | NO |
| chr7 | 1.03E+08 | 102600000 | 0.93 | 1.10 | A>A |
| chr7 | 1.03E+08 | 102700000 | 0.00 | 1.05 | NO |
| chr7 | 1.03E+08 | 102800000 | 0.90 | 0.89 | A>A |

| | | | | | |
|---|---|---|---|---|---|
| chr7 | 1.03E+08 | 102900000 | 0.00 | 0.00 | NO |
| chr7 | 1.03E+08 | 103000000 | 0.66 | 0.46 | A>A |
| chr7 | 1.03E+08 | 103100000 | 0.16 | -0.08 | A>A |
| chr7 | 1.03E+08 | 103200000 | -0.31 | -0.58 | B>B |
| chr7 | 1.03E+08 | 103300000 | -0.38 | -0.60 | B>B |
| chr7 | 1.03E+08 | 103400000 | -0.51 | -0.80 | B>B |
| chr7 | 1.03E+08 | 103500000 | -0.67 | -0.96 | B>B |
| chr7 | 1.04E+08 | 103600000 | -0.65 | -1.07 | B>B |
| chr7 | 1.04E+08 | 103700000 | -0.61 | -1.05 | B>B |
| chr7 | 1.04E+08 | 103800000 | -0.40 | -0.76 | B>B |
| chr7 | 1.04E+08 | 103900000 | -0.20 | -0.46 | B>B |
| chr7 | 1.04E+08 | 104000000 | -0.71 | -1.17 | B>B |
| chr7 | 1.04E+08 | 104100000 | -0.78 | -1.23 | B>B |
| chr7 | 1.04E+08 | 104200000 | -0.92 | -1.31 | B>B |
| chr7 | 1.04E+08 | 104300000 | -0.80 | -1.37 | B>B |
| chr7 | 1.04E+08 | 104400000 | -0.82 | -1.24 | B>B |
| chr7 | 1.04E+08 | 104500000 | -0.46 | -0.95 | B>B |
| chr7 | 1.05E+08 | 104600000 | 0.09 | -0.11 | B>B |
| chr7 | 1.05E+08 | 104700000 | 0.70 | 0.58 | A>A |
| chr7 | 1.05E+08 | 104800000 | 0.65 | 0.65 | A>A |
| chr7 | 1.05E+08 | 104900000 | 0.88 | 0.86 | A>A |
| chr7 | 1.05E+08 | 105000000 | 0.81 | 0.87 | A>A |
| chr7 | 1.05E+08 | 105100000 | 0.82 | 0.73 | A>A |
| chr7 | 1.05E+08 | 105200000 | 0.94 | 0.80 | A>A |
| chr7 | 1.05E+08 | 105300000 | 0.83 | 0.62 | A>A |
| chr7 | 1.05E+08 | 105400000 | 0.69 | 0.54 | A>A |
| chr7 | 1.05E+08 | 105500000 | 0.85 | 0.61 | A>A |
| chr7 | 1.06E+08 | 105600000 | 0.83 | 0.62 | A>A |
| chr7 | 1.06E+08 | 105700000 | 0.63 | 0.39 | A>A |
| chr7 | 1.06E+08 | 105800000 | 0.70 | 0.27 | A>A |
| chr7 | 1.06E+08 | 105900000 | 0.56 | -0.13 | A>B |
| chr7 | 1.06E+08 | 106000000 | 0.83 | -0.29 | A>B |
| chr7 | 1.06E+08 | 106100000 | 0.70 | -0.37 | A>B |
| chr7 | 1.06E+08 | 106200000 | 0.50 | -0.62 | A>B |
| chr7 | 1.06E+08 | 106300000 | 0.47 | -0.75 | A>B |
| chr7 | 1.06E+08 | 106400000 | 0.42 | -0.53 | A>B |
| chr7 | 1.06E+08 | 106500000 | 0.40 | -0.56 | A>B |
| chr7 | 1.07E+08 | 106600000 | 0.35 | -0.80 | A>B |
| chr7 | 1.07E+08 | 106700000 | 0.52 | -0.67 | A>B |
| chr7 | 1.07E+08 | 106800000 | 0.58 | -0.45 | A>B |
| chr7 | 1.07E+08 | 106900000 | 0.56 | 0.01 | A>A |
| chr7 | 1.07E+08 | 107000000 | 0.51 | 0.03 | A>A |
| chr7 | 1.07E+08 | 107100000 | 0.59 | 0.07 | A>A |
| chr7 | 1.07E+08 | 107200000 | 0.57 | 0.21 | A>A |
| chr7 | 1.07E+08 | 107300000 | 0.74 | 0.15 | A>B |
| chr7 | 1.07E+08 | 107400000 | 0.60 | 0.01 | A>B |
| chr7 | 1.07E+08 | 107500000 | 0.73 | 0.01 | A>B |
| chr7 | 1.08E+08 | 107600000 | 0.79 | 0.52 | A>A |
| chr7 | 1.08E+08 | 107700000 | 1.08 | 0.89 | A>A |
| chr7 | 1.08E+08 | 107800000 | 0.73 | 0.62 | A>A |
| chr7 | 1.08E+08 | 107900000 | 0.57 | 0.35 | A>A |
| chr7 | 1.08E+08 | 108000000 | 0.64 | 0.35 | A>A |
| chr7 | 1.08E+08 | 108100000 | 0.53 | 0.26 | A>A |
| chr7 | 1.08E+08 | 108200000 | 0.37 | 0.27 | A>A |
| chr7 | 1.08E+08 | 108300000 | -0.14 | -0.13 | B>B |
| chr7 | 1.08E+08 | 108400000 | -0.63 | -0.64 | B>B |

| | | | | | |
|---|---|---|---|---|---|
| chr7 | 1.08E+08 | 108500000 | -0.70 | -0.74 | B>B |
| chr7 | 1.09E+08 | 108600000 | -0.77 | -0.72 | B>B |
| chr7 | 1.09E+08 | 108700000 | -0.74 | -0.70 | B>B |
| chr7 | 1.09E+08 | 108800000 | -0.79 | -0.67 | B>B |
| chr7 | 1.09E+08 | 108900000 | -0.78 | -0.58 | B>B |
| chr7 | 1.09E+08 | 109000000 | -0.73 | -0.45 | B>B |
| chr7 | 1.09E+08 | 109100000 | -0.59 | -0.28 | B>B |
| chr7 | 1.09E+08 | 109200000 | -0.35 | 0.06 | B>B |
| chr7 | 1.09E+08 | 109300000 | -0.31 | 0.12 | B>B |
| chr7 | 1.09E+08 | 109400000 | -0.30 | 0.14 | B>B |
| chr7 | 1.09E+08 | 109500000 | -0.24 | 0.05 | B>B |
| chr7 | 1.1E+08 | 109600000 | -0.12 | 0.14 | A>A |
| chr7 | 1.1E+08 | 109700000 | -0.07 | 0.23 | A>A |
| chr7 | 1.1E+08 | 109800000 | -0.04 | 0.15 | A>A |
| chr7 | 1.1E+08 | 109900000 | 0.06 | 0.19 | A>A |
| chr7 | 1.1E+08 | 110000000 | 0.15 | 0.30 | A>A |
| chr7 | 1.1E+08 | 110100000 | 0.13 | 0.26 | A>A |
| chr7 | 1.1E+08 | 110200000 | 0.21 | 0.25 | A>A |
| chr7 | 1.1E+08 | 110300000 | 0.23 | 0.35 | A>A |
| chr7 | 1.1E+08 | 110400000 | 0.15 | 0.27 | A>A |
| chr7 | 1.1E+08 | 110500000 | 0.13 | 0.32 | A>A |
| chr7 | 1.11E+08 | 110600000 | 0.13 | 0.37 | A>A |
| chr7 | 1.11E+08 | 110700000 | 0.13 | 0.40 | A>A |
| chr7 | 1.11E+08 | 110800000 | 0.14 | 0.29 | A>A |
| chr7 | 1.11E+08 | 110900000 | 0.23 | 0.38 | A>A |
| chr7 | 1.11E+08 | 111000000 | 0.25 | 0.42 | A>A |
| chr7 | 1.11E+08 | 111100000 | 0.24 | 0.46 | A>A |
| chr7 | 1.11E+08 | 111200000 | 0.25 | 0.39 | A>A |
| chr7 | 1.11E+08 | 111300000 | 0.37 | 0.50 | A>A |
| chr7 | 1.11E+08 | 111400000 | 0.17 | 0.18 | A>A |
| chr7 | 1.11E+08 | 111500000 | 0.21 | -0.08 | A>A |
| chr7 | 1.12E+08 | 111600000 | 0.30 | -0.14 | A>A |
| chr7 | 1.12E+08 | 111700000 | 0.57 | 0.00 | A>A |
| chr7 | 1.12E+08 | 111800000 | 0.66 | 0.04 | A>B |
| chr7 | 1.12E+08 | 111900000 | 0.58 | 0.10 | A>A |
| chr7 | 1.12E+08 | 112000000 | 0.47 | 0.09 | A>A |
| chr7 | 1.12E+08 | 112100000 | 0.34 | 0.02 | A>A |
| chr7 | 1.12E+08 | 112200000 | -0.08 | -0.33 | B>B |
| chr7 | 1.12E+08 | 112300000 | -0.51 | -0.93 | B>B |
| chr7 | 1.12E+08 | 112400000 | -0.54 | -0.98 | B>B |
| chr7 | 1.12E+08 | 112500000 | -0.32 | -0.55 | B>B |
| chr7 | 1.13E+08 | 112600000 | -0.24 | -0.50 | B>B |
| chr7 | 1.13E+08 | 112700000 | -0.69 | -1.00 | B>B |
| chr7 | 1.13E+08 | 112800000 | -0.58 | -0.94 | B>B |
| chr7 | 1.13E+08 | 112900000 | -0.88 | -1.08 | B>B |
| chr7 | 1.13E+08 | 113000000 | -0.97 | -1.18 | B>B |
| chr7 | 1.13E+08 | 113100000 | -0.87 | -1.06 | B>B |
| chr7 | 1.13E+08 | 113200000 | -1.04 | -1.16 | B>B |
| chr7 | 1.13E+08 | 113300000 | -1.08 | -1.17 | B>B |
| chr7 | 1.13E+08 | 113400000 | -0.91 | -1.04 | B>B |
| chr7 | 1.13E+08 | 113500000 | -1.11 | -1.15 | B>B |
| chr7 | 1.14E+08 | 113600000 | -1.02 | -1.17 | B>B |
| chr7 | 1.14E+08 | 113700000 | -0.71 | -0.61 | B>B |
| chr7 | 1.14E+08 | 113800000 | -0.56 | -0.29 | B>B |
| chr7 | 1.14E+08 | 113900000 | -0.71 | -0.46 | B>B |
| chr7 | 1.14E+08 | 114000000 | -0.66 | -0.33 | B>B |

| | | | | | |
|---|---|---|---|---|---|
| chr7 | 1.14E+08 | 114100000 | -0.52 | -0.05 | B>B |
| chr7 | 1.14E+08 | 114200000 | -0.54 | 0.11 | B>A |
| chr7 | 1.14E+08 | 114300000 | -0.40 | 0.05 | B>B |
| chr7 | 1.14E+08 | 114400000 | -0.15 | 0.09 | B>B |
| chr7 | 1.14E+08 | 114500000 | 0.02 | 0.25 | A>A |
| chr7 | 1.15E+08 | 114600000 | 0.20 | 0.36 | A>A |
| chr7 | 1.15E+08 | 114700000 | 0.22 | 0.40 | A>A |
| chr7 | 1.15E+08 | 114800000 | 0.10 | 0.43 | A>A |
| chr7 | 1.15E+08 | 114900000 | -0.05 | 0.50 | B>A |
| chr7 | 1.15E+08 | 115000000 | -0.18 | 0.43 | B>A |
| chr7 | 1.15E+08 | 115100000 | -0.50 | 0.37 | B>A |
| chr7 | 1.15E+08 | 115200000 | -0.56 | 0.44 | B>A |
| chr7 | 1.15E+08 | 115300000 | -0.44 | 0.40 | B>A |
| chr7 | 1.15E+08 | 115400000 | -0.47 | 0.29 | B>A |
| chr7 | 1.15E+08 | 115500000 | -0.63 | 0.19 | B>A |
| chr7 | 1.16E+08 | 115600000 | -0.64 | 0.05 | B>A |
| chr7 | 1.16E+08 | 115700000 | -0.60 | 0.05 | B>A |
| chr7 | 1.16E+08 | 115800000 | -0.23 | 0.12 | B>B |
| chr7 | 1.16E+08 | 115900000 | 0.29 | 0.44 | A>A |
| chr7 | 1.16E+08 | 116000000 | 0.78 | 0.89 | A>A |
| chr7 | 1.16E+08 | 116100000 | 1.01 | 0.98 | A>A |
| chr7 | 1.16E+08 | 116200000 | 0.91 | 0.87 | A>A |
| chr7 | 1.16E+08 | 116300000 | 0.60 | 0.74 | A>A |
| chr7 | 1.16E+08 | 116400000 | 0.81 | 0.83 | A>A |
| chr7 | 1.16E+08 | 116500000 | 0.56 | 0.68 | A>A |
| chr7 | 1.17E+08 | 116600000 | 0.65 | 0.79 | A>A |
| chr7 | 1.17E+08 | 116700000 | 0.59 | 0.64 | A>A |
| chr7 | 1.17E+08 | 116800000 | 0.41 | 0.57 | A>A |
| chr7 | 1.17E+08 | 116900000 | 0.14 | 0.29 | A>A |
| chr7 | 1.17E+08 | 117000000 | 0.01 | 0.16 | A>A |
| chr7 | 1.17E+08 | 117100000 | 0.05 | 0.17 | A>A |
| chr7 | 1.17E+08 | 117200000 | 0.37 | 0.43 | A>A |
| chr7 | 1.17E+08 | 117300000 | 0.45 | 0.47 | A>A |
| chr7 | 1.17E+08 | 117400000 | 0.15 | 0.17 | A>A |
| chr7 | 1.17E+08 | 117500000 | -0.09 | 0.01 | B>B |
| chr7 | 1.18E+08 | 117600000 | -0.14 | -0.14 | B>B |
| chr7 | 1.18E+08 | 117700000 | -0.38 | -0.24 | B>B |
| chr7 | 1.18E+08 | 117800000 | -0.29 | -0.21 | B>B |
| chr7 | 1.18E+08 | 117900000 | -0.37 | -0.32 | B>B |
| chr7 | 1.18E+08 | 118000000 | -0.86 | -0.71 | B>B |
| chr7 | 1.18E+08 | 118100000 | -0.94 | -0.82 | B>B |
| chr7 | 1.18E+08 | 118200000 | -1.02 | -0.99 | B>B |
| chr7 | 1.18E+08 | 118300000 | -0.98 | -0.98 | B>B |
| chr7 | 1.18E+08 | 118400000 | -0.97 | -0.97 | B>B |
| chr7 | 1.18E+08 | 118500000 | -0.96 | -0.89 | B>B |
| chr7 | 1.19E+08 | 118600000 | -0.95 | -0.66 | B>B |
| chr7 | 1.19E+08 | 118700000 | -0.93 | -0.58 | B>B |
| chr7 | 1.19E+08 | 118800000 | -0.88 | -0.31 | B>A |
| chr7 | 1.19E+08 | 118900000 | -0.83 | -0.46 | B>B |
| chr7 | 1.19E+08 | 119000000 | -0.85 | -0.48 | B>B |
| chr7 | 1.19E+08 | 119100000 | -0.95 | -0.39 | B>A |
| chr7 | 1.19E+08 | 119200000 | -0.90 | -0.28 | B>A |
| chr7 | 1.19E+08 | 119300000 | -0.89 | -0.27 | B>A |
| chr7 | 1.19E+08 | 119400000 | -0.89 | -0.25 | B>A |
| chr7 | 1.19E+08 | 119500000 | -0.93 | -0.38 | B>A |
| chr7 | 1.2E+08 | 119600000 | -0.89 | -0.51 | B>B |

| | | | | | |
|---|---|---|---|---|---|
| chr7 | 1.2E+08 | 119700000 | -0.88 | -0.73 | B>B |
| chr7 | 1.2E+08 | 119800000 | -0.94 | -0.81 | B>B |
| chr7 | 1.2E+08 | 119900000 | -0.88 | -0.88 | B>B |
| chr7 | 1.2E+08 | 120000000 | -0.82 | -0.85 | B>B |
| chr7 | 1.2E+08 | 120100000 | -0.95 | -0.96 | B>B |
| chr7 | 1.2E+08 | 120200000 | -0.96 | -0.99 | B>B |
| chr7 | 1.2E+08 | 120300000 | -0.87 | -1.01 | B>B |
| chr7 | 1.2E+08 | 120400000 | -0.73 | -0.87 | B>B |
| chr7 | 1.2E+08 | 120500000 | -0.44 | -0.80 | B>B |
| chr7 | 1.21E+08 | 120600000 | -0.44 | -0.75 | B>B |
| chr7 | 1.21E+08 | 120700000 | -0.40 | -0.73 | B>B |
| chr7 | 1.21E+08 | 120800000 | -0.52 | -0.68 | B>B |
| chr7 | 1.21E+08 | 120900000 | -0.74 | -0.82 | B>B |
| chr7 | 1.21E+08 | 121000000 | -0.50 | -0.45 | B>B |
| chr7 | 1.21E+08 | 121100000 | 0.07 | 0.13 | A>A |
| chr7 | 1.21E+08 | 121200000 | -0.08 | 0.09 | A>A |
| chr7 | 1.21E+08 | 121300000 | -0.57 | -0.68 | B>B |
| chr7 | 1.21E+08 | 121400000 | -0.87 | -0.95 | B>B |
| chr7 | 1.21E+08 | 121500000 | -0.84 | -1.01 | B>B |
| chr7 | 1.22E+08 | 121600000 | -0.71 | -0.99 | B>B |
| chr7 | 1.22E+08 | 121700000 | -0.66 | -0.85 | B>B |
| chr7 | 1.22E+08 | 121800000 | -0.73 | -0.92 | B>B |
| chr7 | 1.22E+08 | 121900000 | -0.73 | -0.87 | B>B |
| chr7 | 1.22E+08 | 122000000 | -0.67 | -0.88 | B>B |
| chr7 | 1.22E+08 | 122100000 | -0.79 | -0.87 | B>B |
| chr7 | 1.22E+08 | 122200000 | -0.75 | -0.81 | B>B |
| chr7 | 1.22E+08 | 122300000 | -0.94 | -0.75 | B>B |
| chr7 | 1.22E+08 | 122400000 | -0.85 | -0.73 | B>B |
| chr7 | 1.22E+08 | 122500000 | -0.90 | -0.74 | B>B |
| chr7 | 1.23E+08 | 122600000 | -0.81 | -0.75 | B>B |
| chr7 | 1.23E+08 | 122700000 | -0.93 | -0.98 | B>B |
| chr7 | 1.23E+08 | 122800000 | -0.87 | -0.91 | B>B |
| chr7 | 1.23E+08 | 122900000 | -0.80 | -1.02 | B>B |
| chr7 | 1.23E+08 | 123000000 | -0.92 | -1.07 | B>B |
| chr7 | 1.23E+08 | 123100000 | -0.83 | -1.04 | B>B |
| chr7 | 1.23E+08 | 123200000 | -0.40 | -0.61 | B>B |
| chr7 | 1.23E+08 | 123300000 | -0.10 | -0.34 | B>B |
| chr7 | 1.23E+08 | 123400000 | 0.36 | 0.15 | A>A |
| chr7 | 1.23E+08 | 123500000 | 0.21 | 0.06 | A>A |
| chr7 | 1.24E+08 | 123600000 | -0.35 | -0.56 | B>B |
| chr7 | 1.24E+08 | 123700000 | -0.47 | -0.76 | B>B |
| chr7 | 1.24E+08 | 123800000 | -0.49 | -0.58 | B>B |
| chr7 | 1.24E+08 | 123900000 | -0.64 | -0.34 | B>B |
| chr7 | 1.24E+08 | 124000000 | -0.69 | -0.24 | B>B |
| chr7 | 1.24E+08 | 124100000 | -0.62 | -0.15 | B>B |
| chr7 | 1.24E+08 | 124200000 | -0.52 | -0.13 | B>B |
| chr7 | 1.24E+08 | 124300000 | -0.55 | -0.18 | B>B |
| chr7 | 1.24E+08 | 124400000 | -0.35 | -0.26 | B>B |
| chr7 | 1.24E+08 | 124500000 | -0.19 | -0.28 | B>B |
| chr7 | 1.25E+08 | 124600000 | -0.19 | -0.51 | B>B |
| chr7 | 1.25E+08 | 124700000 | -0.67 | -0.99 | B>B |
| chr7 | 1.25E+08 | 124800000 | -0.87 | -1.01 | B>B |
| chr7 | 1.25E+08 | 124900000 | -0.91 | -1.04 | B>B |
| chr7 | 1.25E+08 | 125000000 | -0.99 | -1.05 | B>B |
| chr7 | 1.25E+08 | 125100000 | -0.93 | -1.05 | B>B |
| chr7 | 1.25E+08 | 125200000 | -0.96 | -1.02 | B>B |

| | | | | | |
|---|---|---|---|---|---|
| chr7 | 1.25E+08 | 125300000 | -0.96 | -1.06 | B>B |
| chr7 | 1.25E+08 | 125400000 | -0.96 | -1.07 | B>B |
| chr7 | 1.25E+08 | 125500000 | -0.95 | -1.06 | B>B |
| chr7 | 1.26E+08 | 125600000 | -0.93 | -1.03 | B>B |
| chr7 | 1.26E+08 | 125700000 | -0.97 | -1.08 | B>B |
| chr7 | 1.26E+08 | 125800000 | -0.94 | -1.08 | B>B |
| chr7 | 1.26E+08 | 125900000 | -0.98 | -1.01 | B>B |
| chr7 | 1.26E+08 | 126000000 | -0.97 | -1.07 | B>B |
| chr7 | 1.26E+08 | 126100000 | -0.94 | -1.06 | B>B |
| chr7 | 1.26E+08 | 126200000 | -0.85 | -1.06 | B>B |
| chr7 | 1.26E+08 | 126300000 | -0.79 | -1.06 | B>B |
| chr7 | 1.26E+08 | 126400000 | -0.56 | -1.03 | B>B |
| chr7 | 1.26E+08 | 126500000 | -0.72 | -0.96 | B>B |
| chr7 | 1.27E+08 | 126600000 | -0.83 | -1.04 | B>B |
| chr7 | 1.27E+08 | 126700000 | -0.87 | -1.14 | B>B |
| chr7 | 1.27E+08 | 126800000 | -0.84 | -1.10 | B>B |
| chr7 | 1.27E+08 | 126900000 | -0.66 | -0.96 | B>B |
| chr7 | 1.27E+08 | 127000000 | -0.43 | -0.85 | B>B |
| chr7 | 1.27E+08 | 127100000 | -0.20 | -0.60 | B>B |
| chr7 | 1.27E+08 | 127200000 | -0.30 | -0.79 | B>B |
| chr7 | 1.27E+08 | 127300000 | 0.30 | 0.00 | A>A |
| chr7 | 1.27E+08 | 127400000 | 0.57 | 0.49 | A>A |
| chr7 | 1.27E+08 | 127500000 | 0.66 | 0.66 | A>A |
| chr7 | 1.28E+08 | 127600000 | 0.58 | 0.65 | A>A |
| chr7 | 1.28E+08 | 127700000 | 0.51 | 0.55 | A>A |
| chr7 | 1.28E+08 | 127800000 | 0.57 | 0.64 | A>A |
| chr7 | 1.28E+08 | 127900000 | 0.72 | 0.60 | A>A |
| chr7 | 1.28E+08 | 128000000 | 1.21 | 1.04 | A>A |
| chr7 | 1.28E+08 | 128100000 | 1.30 | 1.28 | A>A |
| chr7 | 1.28E+08 | 128200000 | 0.00 | 0.00 | NO |
| chr7 | 1.28E+08 | 128300000 | 0.00 | 0.00 | NO |
| chr7 | 1.28E+08 | 128400000 | 1.30 | 1.37 | A>A |
| chr7 | 1.28E+08 | 128500000 | 1.33 | 1.48 | A>A |
| chr7 | 1.29E+08 | 128600000 | 1.30 | 1.28 | A>A |
| chr7 | 1.29E+08 | 128700000 | 1.26 | 1.37 | A>A |
| chr7 | 1.29E+08 | 128800000 | 1.11 | 1.20 | A>A |
| chr7 | 1.29E+08 | 128900000 | 1.08 | 1.19 | A>A |
| chr7 | 1.29E+08 | 129000000 | 0.99 | 1.10 | A>A |
| chr7 | 1.29E+08 | 129100000 | 1.00 | 1.20 | A>A |
| chr7 | 1.29E+08 | 129200000 | 1.07 | 1.17 | A>A |
| chr7 | 1.29E+08 | 129300000 | 1.03 | 1.14 | A>A |
| chr7 | 1.29E+08 | 129400000 | 1.04 | 1.14 | A>A |
| chr7 | 1.29E+08 | 129500000 | 1.02 | 1.25 | A>A |
| chr7 | 1.3E+08 | 129600000 | 1.21 | 1.26 | A>A |
| chr7 | 1.3E+08 | 129700000 | 0.96 | 1.05 | A>A |
| chr7 | 1.3E+08 | 129800000 | 0.86 | 1.01 | A>A |
| chr7 | 1.3E+08 | 129900000 | 0.81 | 0.93 | A>A |
| chr7 | 1.3E+08 | 130000000 | 0.96 | 0.95 | A>A |
| chr7 | 1.3E+08 | 130100000 | 0.89 | 1.03 | A>A |
| chr7 | 1.3E+08 | 130200000 | 0.00 | 0.87 | NO |
| chr7 | 1.3E+08 | 130300000 | 0.00 | 0.00 | NO |
| chr7 | 1.3E+08 | 130400000 | 0.70 | 0.82 | A>A |
| chr7 | 1.3E+08 | 130500000 | 0.65 | 0.81 | A>A |
| chr7 | 1.31E+08 | 130600000 | 0.76 | 0.99 | A>A |
| chr7 | 1.31E+08 | 130700000 | 0.87 | 1.02 | A>A |
| chr7 | 1.31E+08 | 130800000 | 0.87 | 0.93 | A>A |

| | | | | | |
|---|---|---|---|---|---|
| chr7 | 1.31E+08 | 130900000 | 1.03 | 1.16 | A>A |
| chr7 | 1.31E+08 | 131000000 | 1.08 | 1.06 | A>A |
| chr7 | 1.31E+08 | 131100000 | 0.99 | 1.01 | A>A |
| chr7 | 1.31E+08 | 131200000 | 0.67 | 0.81 | A>A |
| chr7 | 1.31E+08 | 131300000 | 0.85 | 0.82 | A>A |
| chr7 | 1.31E+08 | 131400000 | 0.57 | 0.45 | A>A |
| chr7 | 1.31E+08 | 131500000 | 0.59 | 0.20 | A>A |
| chr7 | 1.32E+08 | 131600000 | 0.56 | 0.24 | A>A |
| chr7 | 1.32E+08 | 131700000 | 0.68 | 0.38 | A>A |
| chr7 | 1.32E+08 | 131800000 | 0.78 | 0.43 | A>A |
| chr7 | 1.32E+08 | 131900000 | 0.75 | 0.30 | A>A |
| chr7 | 1.32E+08 | 132000000 | 0.52 | 0.10 | A>A |
| chr7 | 1.32E+08 | 132100000 | 0.47 | 0.04 | A>A |
| chr7 | 1.32E+08 | 132200000 | 0.50 | -0.01 | A>A |
| chr7 | 1.32E+08 | 132300000 | 0.41 | 0.03 | A>A |
| chr7 | 1.32E+08 | 132400000 | 0.47 | 0.10 | A>A |
| chr7 | 1.32E+08 | 132500000 | 0.46 | 0.31 | A>A |
| chr7 | 1.33E+08 | 132600000 | 0.53 | 0.37 | A>A |
| chr7 | 1.33E+08 | 132700000 | 0.61 | 0.48 | A>A |
| chr7 | 1.33E+08 | 132800000 | 0.60 | 0.39 | A>A |
| chr7 | 1.33E+08 | 132900000 | 0.50 | 0.19 | A>A |
| chr7 | 1.33E+08 | 133000000 | 0.50 | 0.35 | A>A |
| chr7 | 1.33E+08 | 133100000 | 0.52 | 0.40 | A>A |
| chr7 | 1.33E+08 | 133200000 | 0.47 | 0.38 | A>A |
| chr7 | 1.33E+08 | 133300000 | 0.53 | 0.49 | A>A |
| chr7 | 1.33E+08 | 133400000 | 0.42 | 0.52 | A>A |
| chr7 | 1.33E+08 | 133500000 | 0.47 | 0.39 | A>A |
| chr7 | 1.34E+08 | 133600000 | 0.44 | 0.39 | A>A |
| chr7 | 1.34E+08 | 133700000 | 0.57 | 0.46 | A>A |
| chr7 | 1.34E+08 | 133800000 | 0.56 | 0.40 | A>A |
| chr7 | 1.34E+08 | 133900000 | 0.66 | 0.45 | A>A |
| chr7 | 1.34E+08 | 134000000 | 0.86 | 0.55 | A>A |
| chr7 | 1.34E+08 | 134100000 | 0.90 | 0.65 | A>A |
| chr7 | 1.34E+08 | 134200000 | 0.95 | 0.75 | A>A |
| chr7 | 1.34E+08 | 134300000 | 1.10 | 0.94 | A>A |
| chr7 | 1.34E+08 | 134400000 | 0.80 | 0.56 | A>A |
| chr7 | 1.34E+08 | 134500000 | 0.65 | 0.40 | A>A |
| chr7 | 1.35E+08 | 134600000 | 0.78 | 0.43 | A>A |
| chr7 | 1.35E+08 | 134700000 | 0.49 | 0.35 | A>A |
| chr7 | 1.35E+08 | 134800000 | 0.36 | 0.04 | A>A |
| chr7 | 1.35E+08 | 134900000 | 0.55 | 0.33 | A>A |
| chr7 | 1.35E+08 | 135000000 | 0.51 | 0.35 | A>A |
| chr7 | 1.35E+08 | 135100000 | 0.55 | 0.33 | A>A |
| chr7 | 1.35E+08 | 135200000 | 0.74 | 0.51 | A>A |
| chr7 | 1.35E+08 | 135300000 | 0.75 | 0.37 | A>A |
| chr7 | 1.35E+08 | 135400000 | 0.46 | 0.01 | A>A |
| chr7 | 1.35E+08 | 135500000 | 0.09 | -0.41 | B>B |
| chr7 | 1.36E+08 | 135600000 | -0.07 | -0.75 | A>B |
| chr7 | 1.36E+08 | 135700000 | 0.05 | -0.38 | B>B |
| chr7 | 1.36E+08 | 135800000 | -0.42 | -0.89 | B>B |
| chr7 | 1.36E+08 | 135900000 | -0.79 | -1.08 | B>B |
| chr7 | 1.36E+08 | 136000000 | -0.95 | -1.12 | B>B |
| chr7 | 1.36E+08 | 136100000 | -1.01 | -1.10 | B>B |
| chr7 | 1.36E+08 | 136200000 | -1.04 | -1.11 | B>B |
| chr7 | 1.36E+08 | 136300000 | -1.10 | -1.10 | B>B |
| chr7 | 1.36E+08 | 136400000 | -1.08 | -1.10 | B>B |

| | | | | | |
|---|---|---|---|---|---|
| chr7 | 1.36E+08 | 136500000 | -1.02 | -1.08 | B>B |
| chr7 | 1.37E+08 | 136600000 | -1.01 | -1.09 | B>B |
| chr7 | 1.37E+08 | 136700000 | -1.05 | -1.06 | B>B |
| chr7 | 1.37E+08 | 136800000 | -1.10 | -1.16 | B>B |
| chr7 | 1.37E+08 | 136900000 | -1.06 | -1.10 | B>B |
| chr7 | 1.37E+08 | 137000000 | -1.01 | -1.10 | B>B |
| chr7 | 1.37E+08 | 137100000 | -1.00 | -1.10 | B>B |
| chr7 | 1.37E+08 | 137200000 | -0.82 | -1.04 | B>B |
| chr7 | 1.37E+08 | 137300000 | -0.58 | -0.72 | B>B |
| chr7 | 1.37E+08 | 137400000 | -0.35 | -0.24 | B>B |
| chr7 | 1.37E+08 | 137500000 | -0.43 | -0.32 | B>B |
| chr7 | 1.38E+08 | 137600000 | -0.11 | -0.01 | B>B |
| chr7 | 1.38E+08 | 137700000 | 0.35 | 0.37 | A>A |
| chr7 | 1.38E+08 | 137800000 | 0.39 | 0.56 | A>A |
| chr7 | 1.38E+08 | 137900000 | 0.51 | 0.71 | A>A |
| chr7 | 1.38E+08 | 138000000 | 0.63 | 0.77 | A>A |
| chr7 | 1.38E+08 | 138100000 | 0.66 | 0.83 | A>A |
| chr7 | 1.38E+08 | 138200000 | 0.59 | 0.82 | A>A |
| chr7 | 1.38E+08 | 138300000 | 0.66 | 0.82 | A>A |
| chr7 | 1.38E+08 | 138400000 | 0.63 | 0.84 | A>A |
| chr7 | 1.38E+08 | 138500000 | 0.55 | 0.75 | A>A |
| chr7 | 1.39E+08 | 138600000 | 0.77 | 0.86 | A>A |
| chr7 | 1.39E+08 | 138700000 | 0.89 | 1.08 | A>A |
| chr7 | 1.39E+08 | 138800000 | 1.09 | 1.20 | A>A |
| chr7 | 1.39E+08 | 138900000 | 1.16 | 1.27 | A>A |
| chr7 | 1.39E+08 | 139000000 | 1.25 | 1.25 | A>A |
| chr7 | 1.39E+08 | 139100000 | 1.23 | 1.27 | A>A |
| chr7 | 1.39E+08 | 139200000 | 1.05 | 1.25 | A>A |
| chr7 | 1.39E+08 | 139300000 | 0.98 | 0.93 | A>A |
| chr7 | 1.39E+08 | 139400000 | 0.96 | 1.04 | A>A |
| chr7 | 1.39E+08 | 139500000 | 1.10 | 1.16 | A>A |
| chr7 | 1.4E+08 | 139600000 | 1.14 | 1.15 | A>A |
| chr7 | 1.4E+08 | 139700000 | 1.04 | 1.04 | A>A |
| chr7 | 1.4E+08 | 139800000 | 0.91 | 0.92 | A>A |
| chr7 | 1.4E+08 | 139900000 | 1.05 | 1.05 | A>A |
| chr7 | 1.4E+08 | 140000000 | 1.12 | 1.18 | A>A |
| chr7 | 1.4E+08 | 140100000 | 1.00 | 0.96 | A>A |
| chr7 | 1.4E+08 | 140200000 | 0.92 | 0.94 | A>A |
| chr7 | 1.4E+08 | 140300000 | 1.10 | 0.85 | A>A |
| chr7 | 1.4E+08 | 140400000 | 0.91 | 0.71 | A>A |
| chr7 | 1.4E+08 | 140500000 | 0.53 | 0.31 | A>A |
| chr7 | 1.41E+08 | 140600000 | 0.53 | 0.32 | A>A |
| chr7 | 1.41E+08 | 140700000 | 0.43 | 0.18 | A>A |
| chr7 | 1.41E+08 | 140800000 | 0.08 | -0.38 | B>B |
| chr7 | 1.41E+08 | 140900000 | -0.07 | -0.61 | B>B |
| chr7 | 1.41E+08 | 141000000 | -0.09 | -0.83 | A>B |
| chr7 | 1.41E+08 | 141100000 | -0.20 | -1.02 | A>B |
| chr7 | 1.41E+08 | 141200000 | -0.16 | -0.98 | A>B |
| chr7 | 1.41E+08 | 141300000 | 0.23 | -0.40 | A>B |
| chr7 | 1.41E+08 | 141400000 | 0.37 | 0.10 | A>A |
| chr7 | 1.41E+08 | 141500000 | 0.20 | -0.11 | A>A |
| chr7 | 1.42E+08 | 141600000 | -0.42 | -1.01 | A>B |
| chr7 | 1.42E+08 | 141700000 | -0.27 | -0.77 | B>B |
| chr7 | 1.42E+08 | 141800000 | 0.44 | -0.19 | A>B |
| chr7 | 1.42E+08 | 141900000 | 0.32 | -0.16 | A>A |
| chr7 | 1.42E+08 | 142000000 | -0.12 | -0.86 | A>B |

| | | | | | |
|---|---|---|---|---|---|
| chr7 | 1.42E+08 | 142100000 | 0.00 | -0.83 | NO |
| chr7 | 1.42E+08 | 142200000 | 0.01 | -0.80 | A>B |
| chr7 | 1.42E+08 | 142300000 | -0.05 | -0.81 | A>B |
| chr7 | 1.42E+08 | 142400000 | 0.00 | 0.00 | NO |
| chr7 | 1.42E+08 | 142500000 | -0.48 | -0.99 | B>B |
| chr7 | 1.43E+08 | 142600000 | -0.24 | -0.82 | B>B |
| chr7 | 1.43E+08 | 142700000 | -0.28 | -0.97 | A>B |
| chr7 | 1.43E+08 | 142800000 | -0.32 | -0.90 | B>B |
| chr7 | 1.43E+08 | 142900000 | -0.21 | -0.79 | B>B |
| chr7 | 1.43E+08 | 143000000 | 0.19 | -0.07 | A>A |
| chr7 | 1.43E+08 | 143100000 | 0.61 | 0.31 | A>A |
| chr7 | 1.43E+08 | 143200000 | 0.16 | -0.11 | A>A |
| chr7 | 1.43E+08 | 143300000 | 0.00 | 0.00 | NO |
| chr7 | 1.43E+08 | 143400000 | 0.00 | 0.00 | NO |
| chr7 | 1.43E+08 | 143500000 | 0.00 | 0.00 | NO |
| chr7 | 1.44E+08 | 143600000 | 0.00 | 0.00 | NO |
| chr7 | 1.44E+08 | 143700000 | 0.01 | -0.48 | B>B |
| chr7 | 1.44E+08 | 143800000 | -0.47 | -1.00 | B>B |
| chr7 | 1.44E+08 | 143900000 | -0.83 | -1.08 | B>B |
| chr7 | 1.44E+08 | 144000000 | 0.00 | 0.00 | NO |
| chr7 | 1.44E+08 | 144100000 | 0.00 | 0.00 | NO |
| chr7 | 1.44E+08 | 144200000 | -0.92 | -0.77 | B>B |
| chr7 | 1.44E+08 | 144300000 | -1.12 | -0.72 | B>B |
| chr7 | 1.44E+08 | 144400000 | -1.06 | -0.66 | B>B |
| chr7 | 1.44E+08 | 144500000 | -0.76 | -0.49 | B>B |
| chr7 | 1.45E+08 | 144600000 | -0.63 | -0.61 | B>B |
| chr7 | 1.45E+08 | 144700000 | -0.86 | -0.89 | B>B |
| chr7 | 1.45E+08 | 144800000 | -0.90 | -0.88 | B>B |
| chr7 | 1.45E+08 | 144900000 | -0.88 | -0.94 | B>B |
| chr7 | 1.45E+08 | 145000000 | -0.93 | -0.97 | B>B |
| chr7 | 1.45E+08 | 145100000 | -0.93 | -0.96 | B>B |
| chr7 | 1.45E+08 | 145200000 | -0.94 | -0.97 | B>B |
| chr7 | 1.45E+08 | 145300000 | -0.97 | -0.94 | B>B |
| chr7 | 1.45E+08 | 145400000 | -0.93 | -0.93 | B>B |
| chr7 | 1.45E+08 | 145500000 | -0.95 | -0.87 | B>B |
| chr7 | 1.46E+08 | 145600000 | -0.93 | -0.68 | B>B |
| chr7 | 1.46E+08 | 145700000 | -1.00 | -0.54 | B>B |
| chr7 | 1.46E+08 | 145800000 | -0.85 | -0.56 | B>B |
| chr7 | 1.46E+08 | 145900000 | -0.96 | -0.86 | B>B |
| chr7 | 1.46E+08 | 146000000 | -0.92 | -0.92 | B>B |
| chr7 | 1.46E+08 | 146100000 | -0.91 | -0.93 | B>B |
| chr7 | 1.46E+08 | 146200000 | -0.92 | -0.90 | B>B |
| chr7 | 1.46E+08 | 146300000 | -0.96 | -0.87 | B>B |
| chr7 | 1.46E+08 | 146400000 | -0.91 | -0.83 | B>B |
| chr7 | 1.46E+08 | 146500000 | -0.92 | -0.78 | B>B |
| chr7 | 1.47E+08 | 146600000 | -0.92 | -0.70 | B>B |
| chr7 | 1.47E+08 | 146700000 | -0.93 | -0.70 | B>B |
| chr7 | 1.47E+08 | 146800000 | -0.92 | -0.75 | B>B |
| chr7 | 1.47E+08 | 146900000 | -0.94 | -0.79 | B>B |
| chr7 | 1.47E+08 | 147000000 | -0.88 | -0.87 | B>B |
| chr7 | 1.47E+08 | 147100000 | -0.98 | -1.02 | B>B |
| chr7 | 1.47E+08 | 147200000 | -0.97 | -1.00 | B>B |
| chr7 | 1.47E+08 | 147300000 | -0.95 | -1.03 | B>B |
| chr7 | 1.47E+08 | 147400000 | -0.98 | -1.05 | B>B |
| chr7 | 1.47E+08 | 147500000 | -0.98 | -1.09 | B>B |
| chr7 | 1.48E+08 | 147600000 | -0.99 | -1.03 | B>B |

| | | | | | |
|---|---|---|---|---|---|
| chr7 | 1.48E+08 | 147700000 | -0.96 | -1.02 | B>B |
| chr7 | 1.48E+08 | 147800000 | -0.97 | -0.98 | B>B |
| chr7 | 1.48E+08 | 147900000 | -0.83 | -0.94 | B>B |
| chr7 | 1.48E+08 | 148000000 | -0.49 | -0.67 | B>B |
| chr7 | 1.48E+08 | 148100000 | -0.23 | -0.52 | B>B |
| chr7 | 1.48E+08 | 148200000 | -0.05 | -0.34 | B>B |
| chr7 | 1.48E+08 | 148300000 | 0.26 | 0.02 | A>A |
| chr7 | 1.48E+08 | 148400000 | 0.62 | 0.27 | A>A |
| chr7 | 1.48E+08 | 148500000 | 0.69 | 0.54 | A>A |
| chr7 | 1.49E+08 | 148600000 | 0.87 | 0.68 | A>A |
| chr7 | 1.49E+08 | 148700000 | 0.77 | 0.57 | A>A |
| chr7 | 1.49E+08 | 148800000 | 0.96 | 0.83 | A>A |
| chr7 | 1.49E+08 | 148900000 | 1.04 | 1.00 | A>A |
| chr7 | 1.49E+08 | 149000000 | 1.00 | 0.96 | A>A |
| chr7 | 1.49E+08 | 149100000 | 0.84 | 0.64 | A>A |
| chr7 | 1.49E+08 | 149200000 | 0.83 | 0.67 | A>A |
| chr7 | 1.49E+08 | 149300000 | 0.50 | 0.50 | A>A |
| chr7 | 1.49E+08 | 149400000 | 0.71 | 0.43 | A>A |
| chr7 | 1.49E+08 | 149500000 | 0.85 | 0.46 | A>A |
| chr7 | 1.5E+08 | 149600000 | 0.54 | -0.02 | A>A |
| chr7 | 1.5E+08 | 149700000 | 0.00 | 0.00 | NO |
| chr7 | 1.5E+08 | 149800000 | 0.00 | 0.00 | NO |
| chr7 | 1.5E+08 | 149900000 | 0.00 | 0.00 | NO |
| chr7 | 1.5E+08 | 150000000 | 0.00 | 0.00 | NO |
| chr7 | 1.5E+08 | 150100000 | 0.37 | -0.11 | A>A |
| chr7 | 1.5E+08 | 150200000 | -0.06 | -0.53 | B>B |
| chr7 | 1.5E+08 | 150300000 | -0.70 | -1.11 | B>B |
| chr7 | 1.5E+08 | 150400000 | -0.55 | -0.99 | B>B |
| chr7 | 1.5E+08 | 150500000 | -0.27 | -0.67 | B>B |
| chr7 | 1.51E+08 | 150600000 | 0.19 | -0.34 | B>B |
| chr7 | 1.51E+08 | 150700000 | 0.80 | 0.29 | A>A |
| chr7 | 1.51E+08 | 150800000 | 1.37 | 1.00 | A>A |
| chr7 | 1.51E+08 | 150900000 | 1.32 | 1.10 | A>A |
| chr7 | 1.51E+08 | 151000000 | 1.53 | 1.25 | A>A |
| chr7 | 1.51E+08 | 151100000 | 1.37 | 1.20 | A>A |
| chr7 | 1.51E+08 | 151200000 | 1.38 | 1.39 | A>A |
| chr7 | 1.51E+08 | 151300000 | 1.24 | 1.30 | A>A |
| chr7 | 1.51E+08 | 151400000 | 1.51 | 1.51 | A>A |
| chr7 | 1.51E+08 | 151500000 | 1.27 | 1.24 | A>A |
| chr7 | 1.52E+08 | 151600000 | 1.28 | 1.27 | A>A |
| chr7 | 1.52E+08 | 151700000 | 1.08 | 1.11 | A>A |
| chr7 | 1.52E+08 | 151800000 | 0.95 | 1.07 | A>A |
| chr7 | 1.52E+08 | 151900000 | 0.74 | 0.89 | A>A |
| chr7 | 1.52E+08 | 152000000 | 0.00 | 0.00 | NO |
| chr7 | 1.52E+08 | 152100000 | 0.67 | 0.75 | A>A |
| chr7 | 1.52E+08 | 152200000 | 0.66 | 0.73 | A>A |
| chr7 | 1.52E+08 | 152300000 | 0.75 | 1.01 | A>A |
| chr7 | 1.52E+08 | 152400000 | 0.66 | 0.84 | A>A |
| chr7 | 1.52E+08 | 152500000 | 0.58 | 0.55 | A>A |
| chr7 | 1.53E+08 | 152600000 | 0.01 | -0.30 | B>B |
| chr7 | 1.53E+08 | 152700000 | -0.45 | -0.85 | B>B |
| chr7 | 1.53E+08 | 152800000 | -0.53 | -1.06 | B>B |
| chr7 | 1.53E+08 | 152900000 | -0.86 | -1.17 | B>B |
| chr7 | 1.53E+08 | 153000000 | -0.97 | -1.16 | B>B |
| chr7 | 1.53E+08 | 153100000 | -1.03 | -1.20 | B>B |
| chr7 | 1.53E+08 | 153200000 | -1.02 | -1.14 | B>B |

| | | | | | |
|---|---|---|---|---|---|
| chr7 | 1.53E+08 | 153300000 | -1.09 | -1.23 | B>B |
| chr7 | 1.53E+08 | 153400000 | -1.14 | -1.18 | B>B |
| chr7 | 1.53E+08 | 153500000 | 0.00 | 0.00 | NO |
| chr7 | 1.54E+08 | 153600000 | -1.10 | -1.19 | B>B |
| chr7 | 1.54E+08 | 153700000 | -0.93 | -1.18 | B>B |
| chr7 | 1.54E+08 | 153800000 | 0.00 | 0.00 | NO |
| chr7 | 1.54E+08 | 153900000 | 0.00 | 0.00 | NO |
| chr7 | 1.54E+08 | 154000000 | -0.97 | -1.18 | B>B |
| chr7 | 1.54E+08 | 154100000 | -0.99 | -1.22 | B>B |
| chr7 | 1.54E+08 | 154200000 | -1.08 | -1.22 | B>B |
| chr7 | 1.54E+08 | 154300000 | -0.95 | -1.13 | B>B |
| chr7 | 1.54E+08 | 154400000 | 0.00 | 0.00 | NO |
| chr7 | 1.54E+08 | 154500000 | -0.71 | -1.07 | B>B |
| chr7 | 1.55E+08 | 154600000 | -0.56 | -0.94 | B>B |
| chr7 | 1.55E+08 | 154700000 | -0.09 | -0.41 | B>B |
| chr7 | 1.55E+08 | 154800000 | 0.39 | 0.16 | A>A |
| chr7 | 1.55E+08 | 154900000 | 0.31 | -0.06 | A>A |
| chr7 | 1.55E+08 | 155000000 | 0.36 | 0.16 | A>A |
| chr7 | 1.55E+08 | 155100000 | 0.73 | 0.64 | A>A |
| chr7 | 1.55E+08 | 155200000 | 0.57 | 0.48 | A>A |
| chr7 | 1.55E+08 | 155300000 | 0.85 | 0.72 | A>A |
| chr7 | 1.55E+08 | 155400000 | 0.77 | 0.70 | A>A |
| chr7 | 1.55E+08 | 155500000 | 1.11 | 0.90 | A>A |
| chr7 | 1.56E+08 | 155600000 | 0.60 | 0.61 | A>A |
| chr7 | 1.56E+08 | 155700000 | 0.76 | 0.54 | A>A |
| chr7 | 1.56E+08 | 155800000 | 1.03 | 0.70 | A>A |
| chr7 | 1.56E+08 | 155900000 | 0.84 | 0.51 | A>A |
| chr7 | 1.56E+08 | 156000000 | 0.62 | 0.25 | A>A |
| chr7 | 1.56E+08 | 156100000 | 0.52 | 0.04 | A>A |
| chr7 | 1.56E+08 | 156200000 | 0.37 | -0.02 | A>A |
| chr7 | 1.56E+08 | 156300000 | 0.36 | 0.04 | A>A |
| chr7 | 1.56E+08 | 156400000 | 0.48 | 0.25 | A>A |
| chr7 | 1.56E+08 | 156500000 | 0.59 | 0.37 | A>A |
| chr7 | 1.57E+08 | 156600000 | 0.66 | 0.58 | A>A |
| chr7 | 1.57E+08 | 156700000 | 0.90 | 0.78 | A>A |
| chr7 | 1.57E+08 | 156800000 | 1.10 | 1.04 | A>A |
| chr7 | 1.57E+08 | 156900000 | 1.19 | 1.21 | A>A |
| chr7 | 1.57E+08 | 157000000 | 1.08 | 1.16 | A>A |
| chr7 | 1.57E+08 | 157100000 | 0.98 | 1.13 | A>A |
| chr7 | 1.57E+08 | 157200000 | 1.05 | 1.15 | A>A |
| chr7 | 1.57E+08 | 157300000 | 0.75 | 0.78 | A>A |
| chr7 | 1.57E+08 | 157400000 | 0.71 | 0.72 | A>A |
| chr7 | 1.57E+08 | 157500000 | 0.60 | 0.70 | A>A |
| chr7 | 1.58E+08 | 157600000 | 0.51 | 0.58 | A>A |
| chr7 | 1.58E+08 | 157700000 | 0.42 | 0.66 | A>A |
| chr7 | 1.58E+08 | 157800000 | 0.22 | 0.66 | A>A |
| chr7 | 1.58E+08 | 157900000 | 0.19 | 0.60 | A>A |
| chr7 | 1.58E+08 | 158000000 | -0.11 | 0.55 | B>A |
| chr7 | 1.58E+08 | 158100000 | -0.05 | 0.55 | B>A |
| chr7 | 1.58E+08 | 158200000 | 0.10 | 0.62 | B>A |
| chr7 | 1.58E+08 | 158300000 | 0.60 | 0.94 | A>A |
| chr7 | 1.58E+08 | 158400000 | 0.70 | 0.99 | A>A |
| chr7 | 1.58E+08 | 158500000 | 0.88 | 0.95 | A>A |
| chr7 | 1.59E+08 | 158600000 | 1.19 | 1.23 | A>A |
| chr7 | 1.59E+08 | 158700000 | 1.01 | 1.21 | A>A |
| chr7 | 1.59E+08 | 158800000 | 0.83 | 0.96 | A>A |

| | | | | | |
|---|---|---|---|---|---|
| chr7 | 1.59E+08 | 158900000 | 0.76 | 0.98 | A>A |
| chr7 | 1.59E+08 | 159000000 | 0.57 | 0.72 | A>A |
| chr7 | 1.59E+08 | 159100000 | 0.22 | 0.47 | A>A |
| chr7 | 1.59E+08 | 159138663 | 0.00 | 0.00 | NO |
| chr8 | 0 | 100000 | 0.00 | 0.00 | NO |
| chr8 | 100000 | 200000 | 0.00 | 0.00 | NO |
| chr8 | 200000 | 300000 | 0.01 | 0.60 | B>A |
| chr8 | 300000 | 400000 | 0.00 | 0.00 | NO |
| chr8 | 400000 | 500000 | 0.37 | 0.79 | A>A |
| chr8 | 500000 | 600000 | 0.18 | 0.84 | B>A |
| chr8 | 600000 | 700000 | 0.34 | 0.67 | A>A |
| chr8 | 700000 | 800000 | 0.05 | 0.17 | A>A |
| chr8 | 800000 | 900000 | -0.14 | 0.14 | B>B |
| chr8 | 900000 | 1000000 | -0.15 | 0.03 | B>B |
| chr8 | 1000000 | 1100000 | -0.27 | -0.15 | B>B |
| chr8 | 1100000 | 1200000 | -0.37 | -0.26 | B>B |
| chr8 | 1200000 | 1300000 | -0.26 | -0.16 | B>B |
| chr8 | 1300000 | 1400000 | 0.02 | -0.01 | A>A |
| chr8 | 1400000 | 1500000 | 0.20 | -0.03 | A>A |
| chr8 | 1500000 | 1600000 | 0.28 | 0.06 | A>A |
| chr8 | 1600000 | 1700000 | 0.68 | 0.49 | A>A |
| chr8 | 1700000 | 1800000 | 1.17 | 1.11 | A>A |
| chr8 | 1800000 | 1900000 | 0.99 | 0.92 | A>A |
| chr8 | 1900000 | 2000000 | 0.90 | 0.73 | A>A |
| chr8 | 2000000 | 2100000 | 0.45 | -0.02 | A>A |
| chr8 | 2100000 | 2200000 | 0.13 | -0.57 | A>B |
| chr8 | 2200000 | 2300000 | -0.24 | -1.16 | A>B |
| chr8 | 2300000 | 2400000 | -0.57 | -1.26 | A>B |
| chr8 | 2400000 | 2500000 | -0.66 | -1.34 | A>B |
| chr8 | 2500000 | 2600000 | -0.92 | -1.42 | B>B |
| chr8 | 2600000 | 2700000 | -1.05 | -1.42 | B>B |
| chr8 | 2700000 | 2800000 | -1.22 | -1.38 | B>B |
| chr8 | 2800000 | 2900000 | -1.25 | -1.39 | B>B |
| chr8 | 2900000 | 3000000 | -1.17 | -1.37 | B>B |
| chr8 | 3000000 | 3100000 | -1.02 | -1.30 | B>B |
| chr8 | 3100000 | 3200000 | -1.07 | -1.27 | B>B |
| chr8 | 3200000 | 3300000 | -1.11 | -1.31 | B>B |
| chr8 | 3300000 | 3400000 | -1.22 | -1.18 | B>B |
| chr8 | 3400000 | 3500000 | -1.14 | -1.32 | B>B |
| chr8 | 3500000 | 3600000 | -1.11 | -1.27 | B>B |
| chr8 | 3600000 | 3700000 | -1.11 | -1.29 | B>B |
| chr8 | 3700000 | 3800000 | -1.17 | -1.32 | B>B |
| chr8 | 3800000 | 3900000 | -1.07 | -1.26 | B>B |
| chr8 | 3900000 | 4000000 | -1.16 | -1.22 | B>B |
| chr8 | 4000000 | 4100000 | -1.03 | -1.21 | B>B |
| chr8 | 4100000 | 4200000 | -1.13 | -1.26 | B>B |
| chr8 | 4200000 | 4300000 | -1.10 | -1.24 | B>B |
| chr8 | 4300000 | 4400000 | -1.18 | -1.26 | B>B |
| chr8 | 4400000 | 4500000 | -1.19 | -1.29 | B>B |
| chr8 | 4500000 | 4600000 | -1.15 | -1.30 | B>B |
| chr8 | 4600000 | 4700000 | -1.05 | -1.19 | B>B |
| chr8 | 4700000 | 4800000 | -1.01 | -1.16 | B>B |
| chr8 | 4800000 | 4900000 | -0.92 | -1.04 | B>B |
| chr8 | 4900000 | 5000000 | -1.00 | -1.12 | B>B |
| chr8 | 5000000 | 5100000 | -1.11 | -1.23 | B>B |
| chr8 | 5100000 | 5200000 | -1.15 | -1.26 | B>B |

| | | | | | |
|---|---|---|---|---|---|
| chr8 | 5200000 | 5300000 | -1.07 | -1.25 | B>B |
| chr8 | 5300000 | 5400000 | -1.01 | -1.24 | B>B |
| chr8 | 5400000 | 5500000 | -1.06 | -1.20 | B>B |
| chr8 | 5500000 | 5600000 | -1.12 | -1.18 | B>B |
| chr8 | 5600000 | 5700000 | -1.11 | -1.15 | B>B |
| chr8 | 5700000 | 5800000 | -1.05 | -1.18 | B>B |
| chr8 | 5800000 | 5900000 | -1.07 | -1.12 | B>B |
| chr8 | 5900000 | 6000000 | -0.78 | -1.01 | B>B |
| chr8 | 6000000 | 6100000 | -0.63 | -1.02 | B>B |
| chr8 | 6100000 | 6200000 | -0.37 | -0.72 | B>B |
| chr8 | 6200000 | 6300000 | 0.05 | -0.10 | B>B |
| chr8 | 6300000 | 6400000 | 0.46 | 0.62 | A>A |
| chr8 | 6400000 | 6500000 | 0.39 | 0.54 | A>A |
| chr8 | 6500000 | 6600000 | 0.42 | 0.49 | A>A |
| chr8 | 6600000 | 6700000 | 0.49 | 0.21 | A>A |
| chr8 | 6700000 | 6800000 | 0.25 | -0.14 | A>A |
| chr8 | 6800000 | 6900000 | 0.00 | 0.00 | NO |
| chr8 | 6900000 | 7000000 | 0.00 | 0.00 | NO |
| chr8 | 7000000 | 7100000 | 0.00 | 0.00 | NO |
| chr8 | 7100000 | 7200000 | 0.00 | 0.00 | NO |
| chr8 | 7200000 | 7300000 | 0.00 | 0.00 | NO |
| chr8 | 7300000 | 7400000 | 0.00 | 0.00 | NO |
| chr8 | 7400000 | 7500000 | 0.00 | 0.00 | NO |
| chr8 | 7500000 | 7600000 | 0.00 | 0.00 | NO |
| chr8 | 7600000 | 7700000 | 0.00 | 0.00 | NO |
| chr8 | 7700000 | 7800000 | 0.00 | 0.00 | NO |
| chr8 | 7800000 | 7900000 | 0.00 | 0.00 | NO |
| chr8 | 7900000 | 8000000 | 0.00 | 0.00 | NO |
| chr8 | 8000000 | 8100000 | 0.00 | 0.00 | NO |
| chr8 | 8100000 | 8200000 | 1.00 | 0.98 | A>A |
| chr8 | 8200000 | 8300000 | 1.07 | 1.11 | A>A |
| chr8 | 8300000 | 8400000 | 1.17 | 1.09 | A>A |
| chr8 | 8400000 | 8500000 | 1.12 | 1.07 | A>A |
| chr8 | 8500000 | 8600000 | 0.71 | 0.81 | A>A |
| chr8 | 8600000 | 8700000 | 0.95 | 0.91 | A>A |
| chr8 | 8700000 | 8800000 | 1.11 | 1.13 | A>A |
| chr8 | 8800000 | 8900000 | 0.77 | 0.80 | A>A |
| chr8 | 8900000 | 9000000 | 0.81 | 0.83 | A>A |
| chr8 | 9000000 | 9100000 | 0.71 | 0.53 | A>A |
| chr8 | 9100000 | 9200000 | 0.58 | 0.36 | A>A |
| chr8 | 9200000 | 9300000 | 0.57 | 0.45 | A>A |
| chr8 | 9300000 | 9400000 | 0.63 | 0.54 | A>A |
| chr8 | 9400000 | 9500000 | 0.71 | 0.72 | A>A |
| chr8 | 9500000 | 9600000 | 0.62 | 0.57 | A>A |
| chr8 | 9600000 | 9700000 | 0.50 | 0.40 | A>A |
| chr8 | 9700000 | 9800000 | 0.31 | 0.10 | A>A |
| chr8 | 9800000 | 9900000 | 0.22 | -0.03 | A>A |
| chr8 | 9900000 | 10000000 | 0.36 | 0.21 | A>A |
| chr8 | 10000000 | 10100000 | 0.32 | 0.18 | A>A |
| chr8 | 10100000 | 10200000 | 0.29 | 0.15 | A>A |
| chr8 | 10200000 | 10300000 | 0.24 | 0.17 | A>A |
| chr8 | 10300000 | 10400000 | 0.16 | -0.10 | A>A |
| chr8 | 10400000 | 10500000 | 0.53 | 0.32 | A>A |
| chr8 | 10500000 | 10600000 | 0.85 | 0.57 | A>A |
| chr8 | 10600000 | 10700000 | 0.57 | 0.36 | A>A |
| chr8 | 10700000 | 10800000 | 0.37 | -0.07 | A>A |

| | | | | | |
|---|---|---|---|---|---|
| chr8 | 10800000 | 10900000 | 0.25 | -0.13 | A>A |
| chr8 | 10900000 | 11000000 | 0.12 | -0.34 | B>B |
| chr8 | 11000000 | 11100000 | 0.10 | -0.39 | B>B |
| chr8 | 11100000 | 11200000 | 0.52 | 0.19 | A>A |
| chr8 | 11200000 | 11300000 | 0.72 | 0.47 | A>A |
| chr8 | 11300000 | 11400000 | 0.89 | 0.61 | A>A |
| chr8 | 11400000 | 11500000 | 0.69 | 0.34 | A>A |
| chr8 | 11500000 | 11600000 | 0.81 | 0.41 | A>A |
| chr8 | 11600000 | 11700000 | 0.90 | 0.77 | A>A |
| chr8 | 11700000 | 11800000 | 1.04 | 0.81 | A>A |
| chr8 | 11800000 | 11900000 | 0.63 | 0.30 | A>A |
| chr8 | 11900000 | 12000000 | 0.00 | 0.00 | NO |
| chr8 | 12000000 | 12100000 | 0.00 | 0.00 | NO |
| chr8 | 12100000 | 12200000 | 0.00 | 0.00 | NO |
| chr8 | 12200000 | 12300000 | 0.00 | 0.00 | NO |
| chr8 | 12300000 | 12400000 | 0.00 | 0.00 | NO |
| chr8 | 12400000 | 12500000 | 0.00 | 0.00 | NO |
| chr8 | 12500000 | 12600000 | 0.00 | 0.00 | NO |
| chr8 | 12600000 | 12700000 | 0.04 | -0.55 | A>B |
| chr8 | 12700000 | 12800000 | -0.52 | -1.07 | B>B |
| chr8 | 12800000 | 12900000 | -0.69 | -1.13 | B>B |
| chr8 | 12900000 | 13000000 | -0.80 | -1.03 | B>B |
| chr8 | 13000000 | 13100000 | -1.01 | -1.18 | B>B |
| chr8 | 13100000 | 13200000 | -0.94 | -1.19 | B>B |
| chr8 | 13200000 | 13300000 | -1.07 | -1.16 | B>B |
| chr8 | 13300000 | 13400000 | -1.18 | -1.25 | B>B |
| chr8 | 13400000 | 13500000 | -1.09 | -1.18 | B>B |
| chr8 | 13500000 | 13600000 | -1.02 | -1.10 | B>B |
| chr8 | 13600000 | 13700000 | -1.08 | -1.18 | B>B |
| chr8 | 13700000 | 13800000 | -1.20 | -1.22 | B>B |
| chr8 | 13800000 | 13900000 | -1.21 | -1.20 | B>B |
| chr8 | 13900000 | 14000000 | -1.23 | -1.29 | B>B |
| chr8 | 14000000 | 14100000 | -1.15 | -1.20 | B>B |
| chr8 | 14100000 | 14200000 | -1.11 | -1.17 | B>B |
| chr8 | 14200000 | 14300000 | -1.17 | -1.26 | B>B |
| chr8 | 14300000 | 14400000 | -1.06 | -1.17 | B>B |
| chr8 | 14400000 | 14500000 | -1.13 | -1.17 | B>B |
| chr8 | 14500000 | 14600000 | -1.05 | -1.15 | B>B |
| chr8 | 14600000 | 14700000 | -1.10 | -1.12 | B>B |
| chr8 | 14700000 | 14800000 | -1.00 | -1.12 | B>B |
| chr8 | 14800000 | 14900000 | -1.05 | -1.16 | B>B |
| chr8 | 14900000 | 15000000 | -0.96 | -1.13 | B>B |
| chr8 | 15000000 | 15100000 | -0.81 | -1.09 | B>B |
| chr8 | 15100000 | 15200000 | -0.77 | -1.03 | B>B |
| chr8 | 15200000 | 15300000 | -0.57 | -0.68 | B>B |
| chr8 | 15300000 | 15400000 | -0.37 | -0.45 | B>B |
| chr8 | 15400000 | 15500000 | -0.23 | -0.35 | B>B |
| chr8 | 15500000 | 15600000 | -0.26 | -0.31 | B>B |
| chr8 | 15600000 | 15700000 | -0.54 | -0.67 | B>B |
| chr8 | 15700000 | 15800000 | -0.63 | -0.85 | B>B |
| chr8 | 15800000 | 15900000 | -0.73 | -0.97 | B>B |
| chr8 | 15900000 | 16000000 | -0.60 | -0.92 | B>B |
| chr8 | 16000000 | 16100000 | -0.59 | -0.91 | B>B |
| chr8 | 16100000 | 16200000 | -0.72 | -1.08 | B>B |
| chr8 | 16200000 | 16300000 | -0.61 | -0.93 | B>B |
| chr8 | 16300000 | 16400000 | -0.30 | -0.83 | B>B |

| | | | | | |
|---|---|---|---|---|---|
| chr8 | 16400000 | 16500000 | -0.10 | -0.52 | B>B |
| chr8 | 16500000 | 16600000 | 0.00 | -0.35 | B>B |
| chr8 | 16600000 | 16700000 | 0.07 | -0.22 | B>B |
| chr8 | 16700000 | 16800000 | -0.08 | -0.48 | B>B |
| chr8 | 16800000 | 16900000 | 0.17 | -0.29 | B>B |
| chr8 | 16900000 | 17000000 | 0.29 | 0.07 | A>A |
| chr8 | 17000000 | 17100000 | 0.57 | 0.53 | A>A |
| chr8 | 17100000 | 17200000 | 0.69 | 0.70 | A>A |
| chr8 | 17200000 | 17300000 | 0.56 | 0.60 | A>A |
| chr8 | 17300000 | 17400000 | 0.66 | 0.70 | A>A |
| chr8 | 17400000 | 17500000 | 0.69 | 0.77 | A>A |
| chr8 | 17500000 | 17600000 | 0.82 | 0.98 | A>A |
| chr8 | 17600000 | 17700000 | 1.01 | 1.17 | A>A |
| chr8 | 17700000 | 17800000 | 0.89 | 1.03 | A>A |
| chr8 | 17800000 | 17900000 | 0.81 | 0.97 | A>A |
| chr8 | 17900000 | 18000000 | 0.61 | 0.77 | A>A |
| chr8 | 18000000 | 18100000 | 0.33 | 0.55 | A>A |
| chr8 | 18100000 | 18200000 | 0.17 | 0.36 | A>A |
| chr8 | 18200000 | 18300000 | 0.23 | 0.54 | A>A |
| chr8 | 18300000 | 18400000 | 0.27 | 0.53 | A>A |
| chr8 | 18400000 | 18500000 | 0.41 | 0.64 | A>A |
| chr8 | 18500000 | 18600000 | 0.48 | 0.75 | A>A |
| chr8 | 18600000 | 18700000 | 0.50 | 0.64 | A>A |
| chr8 | 18700000 | 18800000 | 0.61 | 0.68 | A>A |
| chr8 | 18800000 | 18900000 | 0.87 | 0.93 | A>A |
| chr8 | 18900000 | 19000000 | 0.81 | 0.88 | A>A |
| chr8 | 19000000 | 19100000 | 0.80 | 0.84 | A>A |
| chr8 | 19100000 | 19200000 | 0.77 | 0.73 | A>A |
| chr8 | 19200000 | 19300000 | 0.64 | 0.67 | A>A |
| chr8 | 19300000 | 19400000 | 0.87 | 0.83 | A>A |
| chr8 | 19400000 | 19500000 | 0.92 | 0.90 | A>A |
| chr8 | 19500000 | 19600000 | 0.87 | 0.83 | A>A |
| chr8 | 19600000 | 19700000 | 0.72 | 0.49 | A>A |
| chr8 | 19700000 | 19800000 | 0.43 | -0.15 | A>A |
| chr8 | 19800000 | 19900000 | 0.27 | -0.50 | A>B |
| chr8 | 19900000 | 20000000 | 0.30 | -0.32 | A>B |
| chr8 | 20000000 | 20100000 | 0.38 | -0.01 | A>A |
| chr8 | 20100000 | 20200000 | 0.35 | -0.26 | A>B |
| chr8 | 20200000 | 20300000 | 0.16 | -0.44 | A>B |
| chr8 | 20300000 | 20400000 | 0.00 | -0.83 | A>B |
| chr8 | 20400000 | 20500000 | -0.13 | -1.07 | A>B |
| chr8 | 20500000 | 20600000 | -0.23 | -1.09 | A>B |
| chr8 | 20600000 | 20700000 | -0.28 | -1.14 | A>B |
| chr8 | 20700000 | 20800000 | -0.21 | -1.09 | A>B |
| chr8 | 20800000 | 20900000 | -0.01 | -0.78 | A>B |
| chr8 | 20900000 | 21000000 | 0.00 | -0.75 | A>B |
| chr8 | 21000000 | 21100000 | 0.09 | -0.98 | A>B |
| chr8 | 21100000 | 21200000 | 0.09 | -0.90 | A>B |
| chr8 | 21200000 | 21300000 | 0.25 | -0.68 | A>B |
| chr8 | 21300000 | 21400000 | 0.38 | -0.39 | A>B |
| chr8 | 21400000 | 21500000 | 0.53 | -0.14 | A>B |
| chr8 | 21500000 | 21600000 | 0.67 | 0.14 | A>A |
| chr8 | 21600000 | 21700000 | 0.61 | 0.06 | A>A |
| chr8 | 21700000 | 21800000 | 0.54 | 0.24 | A>A |
| chr8 | 21800000 | 21900000 | 0.77 | 0.68 | A>A |
| chr8 | 21900000 | 22000000 | 1.05 | 0.90 | A>A |

| | | | | | |
|---|---|---|---|---|---|
| chr8 | 22000000 | 22100000 | 0.90 | 0.96 | A>A |
| chr8 | 22100000 | 22200000 | 0.92 | 0.96 | A>A |
| chr8 | 22200000 | 22300000 | 0.88 | 0.89 | A>A |
| chr8 | 22300000 | 22400000 | 0.98 | 1.06 | A>A |
| chr8 | 22400000 | 22500000 | 0.79 | 0.88 | A>A |
| chr8 | 22500000 | 22600000 | 0.90 | 0.93 | A>A |
| chr8 | 22600000 | 22700000 | 0.94 | 0.95 | A>A |
| chr8 | 22700000 | 22800000 | 0.80 | 0.89 | A>A |
| chr8 | 22800000 | 22900000 | 0.88 | 0.94 | A>A |
| chr8 | 22900000 | 23000000 | 0.87 | 0.93 | A>A |
| chr8 | 23000000 | 23100000 | 0.95 | 1.01 | A>A |
| chr8 | 23100000 | 23200000 | 0.87 | 0.93 | A>A |
| chr8 | 23200000 | 23300000 | 0.93 | 0.94 | A>A |
| chr8 | 23300000 | 23400000 | 0.81 | 0.78 | A>A |
| chr8 | 23400000 | 23500000 | 0.83 | 0.72 | A>A |
| chr8 | 23500000 | 23600000 | 0.74 | 0.69 | A>A |
| chr8 | 23600000 | 23700000 | 0.57 | 0.54 | A>A |
| chr8 | 23700000 | 23800000 | 0.58 | 0.67 | A>A |
| chr8 | 23800000 | 23900000 | 0.39 | 0.58 | A>A |
| chr8 | 23900000 | 24000000 | 0.47 | 0.59 | A>A |
| chr8 | 24000000 | 24100000 | 0.44 | 0.52 | A>A |
| chr8 | 24100000 | 24200000 | 0.17 | 0.48 | A>A |
| chr8 | 24200000 | 24300000 | 0.11 | 0.37 | A>A |
| chr8 | 24300000 | 24400000 | 0.02 | -0.32 | B>B |
| chr8 | 24400000 | 24500000 | -0.25 | -0.86 | A>B |
| chr8 | 24500000 | 24600000 | -0.31 | -0.97 | A>B |
| chr8 | 24600000 | 24700000 | -0.22 | -0.81 | A>B |
| chr8 | 24700000 | 24800000 | 0.07 | -0.24 | B>B |
| chr8 | 24800000 | 24900000 | 0.41 | 0.05 | A>A |
| chr8 | 24900000 | 25000000 | 0.43 | 0.21 | A>A |
| chr8 | 25000000 | 25100000 | 0.82 | 0.63 | A>A |
| chr8 | 25100000 | 25200000 | 0.86 | 0.84 | A>A |
| chr8 | 25200000 | 25300000 | 0.92 | 0.86 | A>A |
| chr8 | 25300000 | 25400000 | 0.65 | 0.37 | A>A |
| chr8 | 25400000 | 25500000 | 0.19 | -0.16 | A>A |
| chr8 | 25500000 | 25600000 | 0.05 | -0.42 | B>B |
| chr8 | 25600000 | 25700000 | -0.18 | -0.67 | B>B |
| chr8 | 25700000 | 25800000 | 0.06 | -0.54 | A>B |
| chr8 | 25800000 | 25900000 | 0.23 | -0.29 | B>B |
| chr8 | 25900000 | 26000000 | 0.43 | 0.19 | A>A |
| chr8 | 26000000 | 26100000 | 0.78 | 0.64 | A>A |
| chr8 | 26100000 | 26200000 | 0.69 | 0.62 | A>A |
| chr8 | 26200000 | 26300000 | 0.64 | 0.47 | A>A |
| chr8 | 26300000 | 26400000 | 0.76 | 0.62 | A>A |
| chr8 | 26400000 | 26500000 | 0.63 | 0.44 | A>A |
| chr8 | 26500000 | 26600000 | 0.17 | -0.16 | A>A |
| chr8 | 26600000 | 26700000 | -0.14 | -0.69 | B>B |
| chr8 | 26700000 | 26800000 | -0.22 | -0.72 | B>B |
| chr8 | 26800000 | 26900000 | -0.14 | -0.69 | B>B |
| chr8 | 26900000 | 27000000 | -0.19 | -0.57 | B>B |
| chr8 | 27000000 | 27100000 | 0.10 | -0.41 | B>B |
| chr8 | 27100000 | 27200000 | 0.62 | 0.42 | A>A |
| chr8 | 27200000 | 27300000 | 0.85 | 0.70 | A>A |
| chr8 | 27300000 | 27400000 | 0.90 | 0.75 | A>A |
| chr8 | 27400000 | 27500000 | 0.92 | 0.84 | A>A |
| chr8 | 27500000 | 27600000 | 0.97 | 0.96 | A>A |

| | | | | | |
|---|---|---|---|---|---|
| chr8 | 27600000 | 27700000 | 0.92 | 0.90 | A>A |
| chr8 | 27700000 | 27800000 | 1.08 | 0.97 | A>A |
| chr8 | 27800000 | 27900000 | 1.01 | 1.01 | A>A |
| chr8 | 27900000 | 28000000 | 0.84 | 0.84 | A>A |
| chr8 | 28000000 | 28100000 | 0.96 | 0.97 | A>A |
| chr8 | 28100000 | 28200000 | 0.84 | 0.82 | A>A |
| chr8 | 28200000 | 28300000 | 0.88 | 1.00 | A>A |
| chr8 | 28300000 | 28400000 | 0.91 | 0.91 | A>A |
| chr8 | 28400000 | 28500000 | 0.88 | 0.90 | A>A |
| chr8 | 28500000 | 28600000 | 0.90 | 0.90 | A>A |
| chr8 | 28600000 | 28700000 | 0.78 | 0.82 | A>A |
| chr8 | 28700000 | 28800000 | 0.94 | 0.94 | A>A |
| chr8 | 28800000 | 28900000 | 0.83 | 0.95 | A>A |
| chr8 | 28900000 | 29000000 | 0.78 | 0.87 | A>A |
| chr8 | 29000000 | 29100000 | 0.89 | 0.98 | A>A |
| chr8 | 29100000 | 29200000 | 0.78 | 0.83 | A>A |
| chr8 | 29200000 | 29300000 | 0.92 | 0.97 | A>A |
| chr8 | 29300000 | 29400000 | 0.89 | 0.95 | A>A |
| chr8 | 29400000 | 29500000 | 0.89 | 0.88 | A>A |
| chr8 | 29500000 | 29600000 | 0.84 | 0.82 | A>A |
| chr8 | 29600000 | 29700000 | 0.87 | 0.96 | A>A |
| chr8 | 29700000 | 29800000 | 0.94 | 0.91 | A>A |
| chr8 | 29800000 | 29900000 | 0.80 | 0.79 | A>A |
| chr8 | 29900000 | 30000000 | 0.79 | 0.92 | A>A |
| chr8 | 30000000 | 30100000 | 0.77 | 0.85 | A>A |
| chr8 | 30100000 | 30200000 | 0.92 | 1.06 | A>A |
| chr8 | 30200000 | 30300000 | 0.91 | 1.04 | A>A |
| chr8 | 30300000 | 30400000 | 0.97 | 1.12 | A>A |
| chr8 | 30400000 | 30500000 | 0.77 | 0.91 | A>A |
| chr8 | 30500000 | 30600000 | 1.01 | 0.93 | A>A |
| chr8 | 30600000 | 30700000 | 0.83 | 0.81 | A>A |
| chr8 | 30700000 | 30800000 | 0.55 | 0.42 | A>A |
| chr8 | 30800000 | 30900000 | 0.59 | 0.41 | A>A |
| chr8 | 30900000 | 31000000 | 0.66 | 0.56 | A>A |
| chr8 | 31000000 | 31100000 | 0.40 | 0.34 | A>A |
| chr8 | 31100000 | 31200000 | 0.10 | 0.06 | A>A |
| chr8 | 31200000 | 31300000 | -0.13 | -0.17 | B>B |
| chr8 | 31300000 | 31400000 | 0.05 | -0.05 | B>B |
| chr8 | 31400000 | 31500000 | 0.12 | 0.16 | A>A |
| chr8 | 31500000 | 31600000 | 0.36 | 0.47 | A>A |
| chr8 | 31600000 | 31700000 | 0.44 | 0.48 | A>A |
| chr8 | 31700000 | 31800000 | 0.42 | 0.49 | A>A |
| chr8 | 31800000 | 31900000 | 0.30 | 0.48 | A>A |
| chr8 | 31900000 | 32000000 | 0.38 | 0.32 | A>A |
| chr8 | 32000000 | 32100000 | 0.40 | 0.38 | A>A |
| chr8 | 32100000 | 32200000 | 0.67 | 0.56 | A>A |
| chr8 | 32200000 | 32300000 | 0.85 | 0.72 | A>A |
| chr8 | 32300000 | 32400000 | 0.71 | 0.60 | A>A |
| chr8 | 32400000 | 32500000 | 0.54 | 0.61 | A>A |
| chr8 | 32500000 | 32600000 | 0.67 | 0.74 | A>A |
| chr8 | 32600000 | 32700000 | 0.55 | 0.54 | A>A |
| chr8 | 32700000 | 32800000 | 0.61 | 0.49 | A>A |
| chr8 | 32800000 | 32900000 | 0.50 | 0.45 | A>A |
| chr8 | 32900000 | 33000000 | 0.53 | 0.44 | A>A |
| chr8 | 33000000 | 33100000 | 0.69 | 0.40 | A>A |
| chr8 | 33100000 | 33200000 | 0.55 | 0.35 | A>A |

| | | | | | |
|---|---|---|---|---|---|
| chr8 | 33200000 | 33300000 | 0.61 | 0.48 | A>A |
| chr8 | 33300000 | 33400000 | 0.73 | 0.60 | A>A |
| chr8 | 33400000 | 33500000 | 0.58 | 0.32 | A>A |
| chr8 | 33500000 | 33600000 | -0.08 | -0.55 | B>B |
| chr8 | 33600000 | 33700000 | -0.28 | -0.76 | B>B |
| chr8 | 33700000 | 33800000 | -0.43 | -0.75 | B>B |
| chr8 | 33800000 | 33900000 | -0.45 | -0.43 | B>B |
| chr8 | 33900000 | 34000000 | -0.41 | -0.41 | B>B |
| chr8 | 34000000 | 34100000 | -0.49 | -0.78 | B>B |
| chr8 | 34100000 | 34200000 | -0.58 | -0.98 | B>B |
| chr8 | 34200000 | 34300000 | -0.54 | -0.98 | B>B |
| chr8 | 34300000 | 34400000 | -0.43 | -1.03 | A>B |
| chr8 | 34400000 | 34500000 | -0.35 | -1.03 | A>B |
| chr8 | 34500000 | 34600000 | -0.17 | -1.01 | A>B |
| chr8 | 34600000 | 34700000 | -0.05 | -0.96 | A>B |
| chr8 | 34700000 | 34800000 | 0.04 | -0.91 | A>B |
| chr8 | 34800000 | 34900000 | 0.12 | -1.00 | A>B |
| chr8 | 34900000 | 35000000 | 0.10 | -1.03 | A>B |
| chr8 | 35000000 | 35100000 | -0.02 | -1.04 | A>B |
| chr8 | 35100000 | 35200000 | -0.10 | -1.03 | A>B |
| chr8 | 35200000 | 35300000 | -0.25 | -1.14 | A>B |
| chr8 | 35300000 | 35400000 | -0.36 | -1.09 | A>B |
| chr8 | 35400000 | 35500000 | -0.40 | -1.16 | A>B |
| chr8 | 35500000 | 35600000 | -0.48 | -1.16 | A>B |
| chr8 | 35600000 | 35700000 | -0.57 | -1.12 | B>B |
| chr8 | 35700000 | 35800000 | -0.62 | -1.19 | B>B |
| chr8 | 35800000 | 35900000 | -0.81 | -1.15 | B>B |
| chr8 | 35900000 | 36000000 | -0.83 | -1.17 | B>B |
| chr8 | 36000000 | 36100000 | -0.88 | -1.11 | B>B |
| chr8 | 36100000 | 36200000 | -1.05 | -1.24 | B>B |
| chr8 | 36200000 | 36300000 | -1.13 | -1.21 | B>B |
| chr8 | 36300000 | 36400000 | -1.11 | -1.25 | B>B |
| chr8 | 36400000 | 36500000 | -1.18 | -1.25 | B>B |
| chr8 | 36500000 | 36600000 | -1.06 | -1.10 | B>B |
| chr8 | 36600000 | 36700000 | -0.56 | -0.78 | B>B |
| chr8 | 36700000 | 36800000 | -0.36 | -0.60 | B>B |
| chr8 | 36800000 | 36900000 | -0.41 | -0.73 | B>B |
| chr8 | 36900000 | 37000000 | -0.04 | -0.14 | B>B |
| chr8 | 37000000 | 37100000 | 0.13 | 0.25 | A>A |
| chr8 | 37100000 | 37200000 | 0.25 | 0.36 | A>A |
| chr8 | 37200000 | 37300000 | 0.34 | 0.45 | A>A |
| chr8 | 37300000 | 37400000 | 0.44 | 0.49 | A>A |
| chr8 | 37400000 | 37500000 | 0.59 | 0.63 | A>A |
| chr8 | 37500000 | 37600000 | 0.45 | 0.47 | A>A |
| chr8 | 37600000 | 37700000 | 0.73 | 0.65 | A>A |
| chr8 | 37700000 | 37800000 | 0.88 | 0.78 | A>A |
| chr8 | 37800000 | 37900000 | 0.88 | 0.76 | A>A |
| chr8 | 37900000 | 38000000 | 0.96 | 0.90 | A>A |
| chr8 | 38000000 | 38100000 | 0.89 | 0.90 | A>A |
| chr8 | 38100000 | 38200000 | 1.02 | 1.07 | A>A |
| chr8 | 38200000 | 38300000 | 0.91 | 0.91 | A>A |
| chr8 | 38300000 | 38400000 | 0.95 | 0.96 | A>A |
| chr8 | 38400000 | 38500000 | 1.01 | 0.94 | A>A |
| chr8 | 38500000 | 38600000 | 0.82 | 0.83 | A>A |
| chr8 | 38600000 | 38700000 | 0.82 | 0.85 | A>A |
| chr8 | 38700000 | 38800000 | 0.85 | 0.80 | A>A |

| | | | | | |
|---|---|---|---|---|---|
| chr8 | 38800000 | 38900000 | 0.85 | 0.80 | A>A |
| chr8 | 38900000 | 39000000 | 0.77 | 0.57 | A>A |
| chr8 | 39000000 | 39100000 | 0.36 | 0.43 | A>A |
| chr8 | 39100000 | 39200000 | 0.26 | 0.15 | A>A |
| chr8 | 39200000 | 39300000 | 0.00 | 0.00 | NO |
| chr8 | 39300000 | 39400000 | 0.00 | 0.00 | NO |
| chr8 | 39400000 | 39500000 | -0.14 | -0.06 | B>B |
| chr8 | 39500000 | 39600000 | -0.13 | -0.18 | B>B |
| chr8 | 39600000 | 39700000 | 0.06 | -0.10 | B>B |
| chr8 | 39700000 | 39800000 | 0.53 | 0.49 | A>A |
| chr8 | 39800000 | 39900000 | 0.64 | 0.64 | A>A |
| chr8 | 39900000 | 40000000 | 0.92 | 0.93 | A>A |
| chr8 | 40000000 | 40100000 | 0.77 | 0.74 | A>A |
| chr8 | 40100000 | 40200000 | 0.77 | 0.74 | A>A |
| chr8 | 40200000 | 40300000 | 0.80 | 0.77 | A>A |
| chr8 | 40300000 | 40400000 | 0.79 | 0.72 | A>A |
| chr8 | 40400000 | 40500000 | 0.75 | 0.71 | A>A |
| chr8 | 40500000 | 40600000 | 0.76 | 0.72 | A>A |
| chr8 | 40600000 | 40700000 | 0.73 | 0.73 | A>A |
| chr8 | 40700000 | 40800000 | 0.76 | 0.74 | A>A |
| chr8 | 40800000 | 40900000 | 0.58 | 0.56 | A>A |
| chr8 | 40900000 | 41000000 | 0.61 | 0.59 | A>A |
| chr8 | 41000000 | 41100000 | 0.80 | 0.78 | A>A |
| chr8 | 41100000 | 41200000 | 0.83 | 0.77 | A>A |
| chr8 | 41200000 | 41300000 | 0.67 | 0.70 | A>A |
| chr8 | 41300000 | 41400000 | 0.80 | 0.77 | A>A |
| chr8 | 41400000 | 41500000 | 0.89 | 0.89 | A>A |
| chr8 | 41500000 | 41600000 | 0.91 | 0.96 | A>A |
| chr8 | 41600000 | 41700000 | 1.04 | 1.07 | A>A |
| chr8 | 41700000 | 41800000 | 0.86 | 0.82 | A>A |
| chr8 | 41800000 | 41900000 | 0.87 | 0.92 | A>A |
| chr8 | 41900000 | 42000000 | 1.06 | 1.03 | A>A |
| chr8 | 42000000 | 42100000 | 0.94 | 0.95 | A>A |
| chr8 | 42100000 | 42200000 | 0.95 | 0.99 | A>A |
| chr8 | 42200000 | 42300000 | 0.80 | 0.84 | A>A |
| chr8 | 42300000 | 42400000 | 1.05 | 1.01 | A>A |
| chr8 | 42400000 | 42500000 | 0.96 | 1.01 | A>A |
| chr8 | 42500000 | 42600000 | 0.83 | 0.97 | A>A |
| chr8 | 42600000 | 42700000 | 0.93 | 0.97 | A>A |
| chr8 | 42700000 | 42800000 | 0.96 | 1.01 | A>A |
| chr8 | 42800000 | 42900000 | 0.87 | 0.99 | A>A |
| chr8 | 42900000 | 43000000 | 0.57 | 0.95 | A>A |
| chr8 | 43000000 | 43100000 | 0.32 | 0.90 | B>A |
| chr8 | 43100000 | 43200000 | 0.00 | 0.54 | NO |
| chr8 | 43200000 | 43300000 | -0.88 | 0.47 | B>A |
| chr8 | 43300000 | 43400000 | -1.26 | 0.49 | B>A |
| chr8 | 43400000 | 43500000 | -1.38 | 0.45 | B>A |
| chr8 | 43500000 | 43600000 | 0.00 | 0.00 | NO |
| chr8 | 43600000 | 43700000 | -1.60 | 0.32 | B>A |
| chr8 | 43700000 | 43800000 | 0.00 | 0.00 | NO |
| chr8 | 43800000 | 43900000 | 0.00 | 0.00 | NO |
| chr8 | 43900000 | 44000000 | 0.00 | 0.00 | NO |
| chr8 | 44000000 | 44100000 | 0.00 | 0.00 | NO |
| chr8 | 44100000 | 44200000 | 0.00 | 0.00 | NO |
| chr8 | 44200000 | 44300000 | 0.00 | 0.00 | NO |
| chr8 | 44300000 | 44400000 | 0.00 | 0.00 | NO |

| | | | | | |
|---|---|---|---|---|---|
| chr8 | 44400000 | 44500000 | 0.00 | 0.00 | NO |
| chr8 | 44500000 | 44600000 | 0.00 | 0.00 | NO |
| chr8 | 44600000 | 44700000 | 0.00 | 0.00 | NO |
| chr8 | 44700000 | 44800000 | 0.00 | 0.00 | NO |
| chr8 | 44800000 | 44900000 | 0.00 | 0.00 | NO |
| chr8 | 44900000 | 45000000 | 0.00 | 0.00 | NO |
| chr8 | 45000000 | 45100000 | 0.00 | 0.00 | NO |
| chr8 | 45100000 | 45200000 | 0.00 | 0.00 | NO |
| chr8 | 45200000 | 45300000 | 0.00 | 0.00 | NO |
| chr8 | 45300000 | 45400000 | 0.00 | 0.00 | NO |
| chr8 | 45400000 | 45500000 | 0.00 | 0.00 | NO |
| chr8 | 45500000 | 45600000 | 0.00 | 0.00 | NO |
| chr8 | 45600000 | 45700000 | 0.00 | 0.00 | NO |
| chr8 | 45700000 | 45800000 | 0.00 | 0.00 | NO |
| chr8 | 45800000 | 45900000 | 0.00 | 0.00 | NO |
| chr8 | 45900000 | 46000000 | 0.00 | 0.00 | NO |
| chr8 | 46000000 | 46100000 | 0.00 | 0.00 | NO |
| chr8 | 46100000 | 46200000 | 0.00 | 0.00 | NO |
| chr8 | 46200000 | 46300000 | 0.00 | 0.00 | NO |
| chr8 | 46300000 | 46400000 | 0.00 | 0.00 | NO |
| chr8 | 46400000 | 46500000 | 0.00 | 0.00 | NO |
| chr8 | 46500000 | 46600000 | 0.00 | 0.00 | NO |
| chr8 | 46600000 | 46700000 | 0.00 | 0.00 | NO |
| chr8 | 46700000 | 46800000 | 0.00 | 0.00 | NO |
| chr8 | 46800000 | 46900000 | 0.00 | 0.00 | NO |
| chr8 | 46900000 | 47000000 | -1.14 | 0.45 | B>A |
| chr8 | 47000000 | 47100000 | 0.00 | 0.00 | NO |
| chr8 | 47100000 | 47200000 | -1.32 | 0.55 | B>A |
| chr8 | 47200000 | 47300000 | -1.48 | 0.74 | B>A |
| chr8 | 47300000 | 47400000 | -1.45 | 0.68 | B>A |
| chr8 | 47400000 | 47500000 | -1.59 | 0.41 | B>A |
| chr8 | 47500000 | 47600000 | -1.39 | 0.30 | B>A |
| chr8 | 47600000 | 47700000 | 0.00 | 0.00 | NO |
| chr8 | 47700000 | 47800000 | -1.32 | 0.37 | B>A |
| chr8 | 47800000 | 47900000 | -0.81 | 0.49 | B>A |
| chr8 | 47900000 | 48000000 | -0.63 | 0.52 | B>A |
| chr8 | 48000000 | 48100000 | -0.17 | 0.67 | B>A |
| chr8 | 48100000 | 48200000 | 0.29 | 0.70 | A>A |
| chr8 | 48200000 | 48300000 | 0.00 | 0.00 | NO |
| chr8 | 48300000 | 48400000 | 0.42 | 0.81 | A>A |
| chr8 | 48400000 | 48500000 | 0.48 | 0.85 | A>A |
| chr8 | 48500000 | 48600000 | 0.56 | 0.90 | A>A |
| chr8 | 48600000 | 48700000 | 0.49 | 0.86 | A>A |
| chr8 | 48700000 | 48800000 | 0.92 | 0.97 | A>A |
| chr8 | 48800000 | 48900000 | 1.10 | 1.06 | A>A |
| chr8 | 48900000 | 49000000 | 0.70 | 0.67 | A>A |
| chr8 | 49000000 | 49100000 | 0.23 | 0.16 | A>A |
| chr8 | 49100000 | 49200000 | -0.05 | -0.03 | B>B |
| chr8 | 49200000 | 49300000 | 0.04 | 0.23 | A>A |
| chr8 | 49300000 | 49400000 | 0.14 | 0.43 | A>A |
| chr8 | 49400000 | 49500000 | 0.25 | 0.44 | A>A |
| chr8 | 49500000 | 49600000 | 0.12 | 0.44 | A>A |
| chr8 | 49600000 | 49700000 | 0.04 | 0.44 | A>A |
| chr8 | 49700000 | 49800000 | -0.02 | 0.45 | A>A |
| chr8 | 49800000 | 49900000 | -0.26 | -0.10 | B>B |
| chr8 | 49900000 | 50000000 | -0.90 | -0.92 | B>B |

| | | | | | |
|---|---|---|---|---|---|
| chr8 | 50000000 | 50100000 | -1.08 | -1.06 | B>B |
| chr8 | 50100000 | 50200000 | -1.13 | -1.14 | B>B |
| chr8 | 50200000 | 50300000 | -1.15 | -1.14 | B>B |
| chr8 | 50300000 | 50400000 | -1.15 | -1.14 | B>B |
| chr8 | 50400000 | 50500000 | -1.12 | -1.17 | B>B |
| chr8 | 50500000 | 50600000 | -1.09 | -1.16 | B>B |
| chr8 | 50600000 | 50700000 | -1.06 | -1.11 | B>B |
| chr8 | 50700000 | 50800000 | -0.96 | -1.15 | B>B |
| chr8 | 50800000 | 50900000 | -0.94 | -1.15 | B>B |
| chr8 | 50900000 | 51000000 | -0.89 | -1.10 | B>B |
| chr8 | 51000000 | 51100000 | -0.93 | -1.12 | B>B |
| chr8 | 51100000 | 51200000 | -0.98 | -1.16 | B>B |
| chr8 | 51200000 | 51300000 | -1.02 | -1.12 | B>B |
| chr8 | 51300000 | 51400000 | -1.09 | -1.12 | B>B |
| chr8 | 51400000 | 51500000 | -1.12 | -1.14 | B>B |
| chr8 | 51500000 | 51600000 | -1.14 | -1.11 | B>B |
| chr8 | 51600000 | 51700000 | -1.16 | -1.07 | B>B |
| chr8 | 51700000 | 51800000 | -1.17 | -1.12 | B>B |
| chr8 | 51800000 | 51900000 | -1.13 | -1.12 | B>B |
| chr8 | 51900000 | 52000000 | -1.16 | -1.09 | B>B |
| chr8 | 52000000 | 52100000 | -1.20 | -1.09 | B>B |
| chr8 | 52100000 | 52200000 | -1.17 | -1.07 | B>B |
| chr8 | 52200000 | 52300000 | -1.19 | -0.98 | B>B |
| chr8 | 52300000 | 52400000 | -1.12 | -0.91 | B>B |
| chr8 | 52400000 | 52500000 | -0.95 | -0.91 | B>B |
| chr8 | 52500000 | 52600000 | -1.00 | -0.94 | B>B |
| chr8 | 52600000 | 52700000 | -0.78 | -0.75 | B>B |
| chr8 | 52700000 | 52800000 | -0.15 | -0.04 | B>B |
| chr8 | 52800000 | 52900000 | 0.07 | 0.29 | A>A |
| chr8 | 52900000 | 53000000 | -0.11 | -0.10 | B>B |
| chr8 | 53000000 | 53100000 | -0.07 | -0.22 | B>B |
| chr8 | 53100000 | 53200000 | -0.23 | -0.34 | B>B |
| chr8 | 53200000 | 53300000 | -0.05 | -0.15 | B>B |
| chr8 | 53300000 | 53400000 | 0.08 | 0.15 | A>A |
| chr8 | 53400000 | 53500000 | 0.18 | 0.26 | A>A |
| chr8 | 53500000 | 53600000 | 0.41 | 0.53 | A>A |
| chr8 | 53600000 | 53700000 | 0.47 | 0.62 | A>A |
| chr8 | 53700000 | 53800000 | 0.54 | 0.72 | A>A |
| chr8 | 53800000 | 53900000 | 0.29 | 0.48 | A>A |
| chr8 | 53900000 | 54000000 | -0.16 | -0.14 | B>B |
| chr8 | 54000000 | 54100000 | -0.28 | -0.15 | B>B |
| chr8 | 54100000 | 54200000 | -0.09 | 0.03 | B>B |
| chr8 | 54200000 | 54300000 | 0.08 | 0.25 | A>A |
| chr8 | 54300000 | 54400000 | 0.12 | 0.11 | A>A |
| chr8 | 54400000 | 54500000 | 0.19 | 0.02 | A>A |
| chr8 | 54500000 | 54600000 | 0.24 | 0.04 | A>A |
| chr8 | 54600000 | 54700000 | 0.39 | 0.31 | A>A |
| chr8 | 54700000 | 54800000 | 0.47 | 0.44 | A>A |
| chr8 | 54800000 | 54900000 | 0.66 | 0.68 | A>A |
| chr8 | 54900000 | 55000000 | 0.77 | 0.77 | A>A |
| chr8 | 55000000 | 55100000 | 0.53 | 0.34 | A>A |
| chr8 | 55100000 | 55200000 | 0.19 | -0.18 | A>A |
| chr8 | 55200000 | 55300000 | 0.12 | -0.21 | B>B |
| chr8 | 55300000 | 55400000 | 0.11 | -0.13 | B>B |
| chr8 | 55400000 | 55500000 | -0.12 | -0.50 | B>B |
| chr8 | 55500000 | 55600000 | -0.36 | -0.76 | B>B |

| | | | | | |
|---|---|---|---|---|---|
| chr8 | 55600000 | 55700000 | -0.44 | -0.95 | B>B |
| chr8 | 55700000 | 55800000 | -0.59 | -0.99 | B>B |
| chr8 | 55800000 | 55900000 | -0.67 | -1.02 | B>B |
| chr8 | 55900000 | 56000000 | -0.67 | -0.94 | B>B |
| chr8 | 56000000 | 56100000 | -0.49 | -0.90 | B>B |
| chr8 | 56100000 | 56200000 | -0.52 | -1.07 | B>B |
| chr8 | 56200000 | 56300000 | -0.44 | -1.01 | B>B |
| chr8 | 56300000 | 56400000 | -0.23 | -1.01 | A>B |
| chr8 | 56400000 | 56500000 | -0.15 | -0.92 | A>B |
| chr8 | 56500000 | 56600000 | -0.20 | -0.93 | A>B |
| chr8 | 56600000 | 56700000 | 0.27 | -0.22 | A>A |
| chr8 | 56700000 | 56800000 | 0.65 | 0.45 | A>A |
| chr8 | 56800000 | 56900000 | 0.68 | 0.63 | A>A |
| chr8 | 56900000 | 57000000 | 0.53 | 0.37 | A>A |
| chr8 | 57000000 | 57100000 | 0.29 | 0.16 | A>A |
| chr8 | 57100000 | 57200000 | 0.13 | 0.00 | A>A |
| chr8 | 57200000 | 57300000 | -0.09 | -0.08 | B>B |
| chr8 | 57300000 | 57400000 | -0.25 | -0.40 | B>B |
| chr8 | 57400000 | 57500000 | -0.40 | -0.83 | B>B |
| chr8 | 57500000 | 57600000 | -0.65 | -1.01 | B>B |
| chr8 | 57600000 | 57700000 | -0.70 | -0.92 | B>B |
| chr8 | 57700000 | 57800000 | -0.66 | -0.91 | B>B |
| chr8 | 57800000 | 57900000 | -0.19 | -0.41 | B>B |
| chr8 | 57900000 | 58000000 | -0.10 | -0.35 | B>B |
| chr8 | 58000000 | 58100000 | -0.16 | -0.25 | B>B |
| chr8 | 58100000 | 58200000 | -0.05 | 0.06 | A>A |
| chr8 | 58200000 | 58300000 | -0.10 | 0.05 | B>B |
| chr8 | 58300000 | 58400000 | 0.06 | 0.25 | A>A |
| chr8 | 58400000 | 58500000 | 0.27 | 0.48 | A>A |
| chr8 | 58500000 | 58600000 | 0.02 | 0.45 | A>A |
| chr8 | 58600000 | 58700000 | -0.03 | 0.53 | B>A |
| chr8 | 58700000 | 58800000 | -0.23 | 0.53 | B>A |
| chr8 | 58800000 | 58900000 | -0.19 | 0.47 | B>A |
| chr8 | 58900000 | 59000000 | -0.04 | 0.53 | B>A |
| chr8 | 59000000 | 59100000 | -0.15 | 0.60 | B>A |
| chr8 | 59100000 | 59200000 | -0.31 | 0.49 | B>A |
| chr8 | 59200000 | 59300000 | -0.29 | 0.40 | B>A |
| chr8 | 59300000 | 59400000 | 0.13 | 0.46 | A>A |
| chr8 | 59400000 | 59500000 | 0.33 | 0.55 | A>A |
| chr8 | 59500000 | 59600000 | 0.38 | 0.52 | A>A |
| chr8 | 59600000 | 59700000 | 0.03 | 0.14 | A>A |
| chr8 | 59700000 | 59800000 | 0.17 | 0.31 | A>A |
| chr8 | 59800000 | 59900000 | 0.05 | 0.30 | A>A |
| chr8 | 59900000 | 60000000 | -0.02 | 0.16 | A>A |
| chr8 | 60000000 | 60100000 | -0.30 | -0.08 | B>B |
| chr8 | 60100000 | 60200000 | -0.53 | -0.35 | B>B |
| chr8 | 60200000 | 60300000 | -0.59 | -0.45 | B>B |
| chr8 | 60300000 | 60400000 | -0.82 | -0.45 | B>B |
| chr8 | 60400000 | 60500000 | -0.96 | -0.46 | B>B |
| chr8 | 60500000 | 60600000 | -0.99 | -0.51 | B>B |
| chr8 | 60600000 | 60700000 | -1.10 | -0.47 | B>A |
| chr8 | 60700000 | 60800000 | -1.01 | -0.44 | B>A |
| chr8 | 60800000 | 60900000 | -0.75 | -0.50 | B>B |
| chr8 | 60900000 | 61000000 | -0.43 | -0.28 | B>B |
| chr8 | 61000000 | 61100000 | -0.30 | -0.09 | B>B |
| chr8 | 61100000 | 61200000 | 0.11 | 0.17 | A>A |

| | | | | | |
|---|---|---|---|---|---|
| chr8 | 61200000 | 61300000 | 0.10 | 0.05 | A>A |
| chr8 | 61300000 | 61400000 | 0.25 | 0.16 | A>A |
| chr8 | 61400000 | 61500000 | 0.55 | 0.51 | A>A |
| chr8 | 61500000 | 61600000 | 0.48 | 0.39 | A>A |
| chr8 | 61600000 | 61700000 | 0.61 | 0.49 | A>A |
| chr8 | 61700000 | 61800000 | 0.55 | 0.37 | A>A |
| chr8 | 61800000 | 61900000 | 0.48 | 0.20 | A>A |
| chr8 | 61900000 | 62000000 | 0.31 | -0.01 | A>A |
| chr8 | 62000000 | 62100000 | 0.04 | -0.23 | B>B |
| chr8 | 62100000 | 62200000 | -0.08 | -0.47 | B>B |
| chr8 | 62200000 | 62300000 | -0.12 | -0.57 | B>B |
| chr8 | 62300000 | 62400000 | 0.05 | -0.37 | B>B |
| chr8 | 62400000 | 62500000 | 0.27 | 0.39 | A>A |
| chr8 | 62500000 | 62600000 | 0.75 | 0.91 | A>A |
| chr8 | 62600000 | 62700000 | 0.88 | 0.76 | A>A |
| chr8 | 62700000 | 62800000 | 0.49 | 0.25 | A>A |
| chr8 | 62800000 | 62900000 | 0.45 | 0.29 | A>A |
| chr8 | 62900000 | 63000000 | 0.41 | 0.34 | A>A |
| chr8 | 63000000 | 63100000 | 0.35 | 0.21 | A>A |
| chr8 | 63100000 | 63200000 | 0.41 | 0.16 | A>A |
| chr8 | 63200000 | 63300000 | 0.39 | -0.01 | A>A |
| chr8 | 63300000 | 63400000 | 0.18 | -0.20 | B>B |
| chr8 | 63400000 | 63500000 | 0.09 | -0.43 | B>B |
| chr8 | 63500000 | 63600000 | -0.09 | -0.66 | B>B |
| chr8 | 63600000 | 63700000 | -0.18 | -0.70 | B>B |
| chr8 | 63700000 | 63800000 | -0.18 | -0.74 | B>B |
| chr8 | 63800000 | 63900000 | -0.06 | -0.48 | B>B |
| chr8 | 63900000 | 64000000 | 0.24 | -0.03 | A>A |
| chr8 | 64000000 | 64100000 | 0.28 | 0.00 | A>A |
| chr8 | 64100000 | 64200000 | 0.13 | -0.26 | B>B |
| chr8 | 64200000 | 64300000 | -0.34 | -0.80 | B>B |
| chr8 | 64300000 | 64400000 | -0.70 | -0.95 | B>B |
| chr8 | 64400000 | 64500000 | -0.95 | -1.00 | B>B |
| chr8 | 64500000 | 64600000 | -1.05 | -0.74 | B>B |
| chr8 | 64600000 | 64700000 | -1.16 | -0.77 | B>B |
| chr8 | 64700000 | 64800000 | -1.25 | -1.01 | B>B |
| chr8 | 64800000 | 64900000 | -1.29 | -1.06 | B>B |
| chr8 | 64900000 | 65000000 | -1.27 | -1.10 | B>B |
| chr8 | 65000000 | 65100000 | -1.27 | -1.10 | B>B |
| chr8 | 65100000 | 65200000 | -1.23 | -1.11 | B>B |
| chr8 | 65200000 | 65300000 | -0.96 | -1.01 | B>B |
| chr8 | 65300000 | 65400000 | -0.78 | -0.93 | B>B |
| chr8 | 65400000 | 65500000 | -0.54 | -0.87 | B>B |
| chr8 | 65500000 | 65600000 | -0.36 | -0.88 | B>B |
| chr8 | 65600000 | 65700000 | -0.32 | -0.85 | B>B |
| chr8 | 65700000 | 65800000 | -0.45 | -0.97 | B>B |
| chr8 | 65800000 | 65900000 | -0.62 | -0.98 | B>B |
| chr8 | 65900000 | 66000000 | -0.49 | -0.94 | B>B |
| chr8 | 66000000 | 66100000 | -0.30 | -0.92 | A>B |
| chr8 | 66100000 | 66200000 | -0.26 | -0.83 | B>B |
| chr8 | 66200000 | 66300000 | -0.21 | -0.82 | A>B |
| chr8 | 66300000 | 66400000 | -0.07 | -0.87 | A>B |
| chr8 | 66400000 | 66500000 | 0.01 | -0.56 | B>B |
| chr8 | 66500000 | 66600000 | 0.48 | 0.34 | A>A |
| chr8 | 66600000 | 66700000 | 0.62 | 0.53 | A>A |
| chr8 | 66700000 | 66800000 | 0.82 | 0.75 | A>A |

| | | | | | |
|---|---|---|---|---|---|
| chr8 | 66800000 | 66900000 | 0.79 | 0.77 | A>A |
| chr8 | 66900000 | 67000000 | 0.67 | 0.53 | A>A |
| chr8 | 67000000 | 67100000 | 0.87 | 0.45 | A>A |
| chr8 | 67100000 | 67200000 | 0.71 | 0.27 | A>A |
| chr8 | 67200000 | 67300000 | 0.70 | 0.31 | A>A |
| chr8 | 67300000 | 67400000 | 0.64 | 0.49 | A>A |
| chr8 | 67400000 | 67500000 | 0.98 | 0.87 | A>A |
| chr8 | 67500000 | 67600000 | 0.85 | 0.82 | A>A |
| chr8 | 67600000 | 67700000 | 0.83 | 0.86 | A>A |
| chr8 | 67700000 | 67800000 | 0.89 | 0.87 | A>A |
| chr8 | 67800000 | 67900000 | 0.91 | 0.79 | A>A |
| chr8 | 67900000 | 68000000 | 0.83 | 0.77 | A>A |
| chr8 | 68000000 | 68100000 | 0.79 | 0.71 | A>A |
| chr8 | 68100000 | 68200000 | 0.64 | 0.51 | A>A |
| chr8 | 68200000 | 68300000 | 0.48 | 0.19 | A>A |
| chr8 | 68300000 | 68400000 | 0.31 | -0.31 | A>B |
| chr8 | 68400000 | 68500000 | 0.01 | -0.70 | A>B |
| chr8 | 68500000 | 68600000 | -0.17 | -0.98 | A>B |
| chr8 | 68600000 | 68700000 | -0.15 | -1.09 | A>B |
| chr8 | 68700000 | 68800000 | -0.44 | -1.08 | A>B |
| chr8 | 68800000 | 68900000 | -0.61 | -1.17 | B>B |
| chr8 | 68900000 | 69000000 | -0.68 | -1.23 | B>B |
| chr8 | 69000000 | 69100000 | -0.75 | -1.15 | B>B |
| chr8 | 69100000 | 69200000 | -0.68 | -1.18 | B>B |
| chr8 | 69200000 | 69300000 | -0.60 | -1.05 | B>B |
| chr8 | 69300000 | 69400000 | -0.73 | -1.09 | B>B |
| chr8 | 69400000 | 69500000 | -0.64 | -1.05 | B>B |
| chr8 | 69500000 | 69600000 | -0.39 | -1.10 | A>B |
| chr8 | 69600000 | 69700000 | -0.37 | -1.16 | A>B |
| chr8 | 69700000 | 69800000 | -0.57 | -1.18 | A>B |
| chr8 | 69800000 | 69900000 | -0.82 | -1.18 | B>B |
| chr8 | 69900000 | 70000000 | -0.85 | -1.16 | B>B |
| chr8 | 70000000 | 70100000 | -0.94 | -1.11 | B>B |
| chr8 | 70100000 | 70200000 | -0.93 | -1.09 | B>B |
| chr8 | 70200000 | 70300000 | -0.92 | -1.06 | B>B |
| chr8 | 70300000 | 70400000 | -0.76 | -1.13 | B>B |
| chr8 | 70400000 | 70500000 | -0.55 | -1.03 | B>B |
| chr8 | 70500000 | 70600000 | -0.23 | -0.75 | B>B |
| chr8 | 70600000 | 70700000 | 0.02 | -0.85 | A>B |
| chr8 | 70700000 | 70800000 | 0.15 | -0.63 | A>B |
| chr8 | 70800000 | 70900000 | 0.25 | -0.16 | A>A |
| chr8 | 70900000 | 71000000 | 0.56 | 0.09 | A>A |
| chr8 | 71000000 | 71100000 | 0.48 | 0.36 | A>A |
| chr8 | 71100000 | 71200000 | 0.65 | 0.75 | A>A |
| chr8 | 71200000 | 71300000 | 0.72 | 0.79 | A>A |
| chr8 | 71300000 | 71400000 | 0.71 | 0.53 | A>A |
| chr8 | 71400000 | 71500000 | 0.50 | 0.39 | A>A |
| chr8 | 71500000 | 71600000 | 0.38 | 0.38 | A>A |
| chr8 | 71600000 | 71700000 | -0.11 | -0.43 | B>B |
| chr8 | 71700000 | 71800000 | -0.38 | -0.80 | B>B |
| chr8 | 71800000 | 71900000 | -0.45 | -0.91 | B>B |
| chr8 | 71900000 | 72000000 | -0.48 | -0.98 | B>B |
| chr8 | 72000000 | 72100000 | -0.51 | -1.05 | B>B |
| chr8 | 72100000 | 72200000 | -0.41 | -1.03 | A>B |
| chr8 | 72200000 | 72300000 | -0.39 | -0.99 | A>B |
| chr8 | 72300000 | 72400000 | -0.53 | -1.10 | B>B |

| | | | | | |
|---|---|---|---|---|---|
| chr8 | 72400000 | 72500000 | -0.49 | -0.99 | B>B |
| chr8 | 72500000 | 72600000 | -0.38 | -1.04 | A>B |
| chr8 | 72600000 | 72700000 | -0.30 | -0.80 | B>B |
| chr8 | 72700000 | 72800000 | 0.06 | -0.37 | B>B |
| chr8 | 72800000 | 72900000 | 0.14 | -0.44 | B>B |
| chr8 | 72900000 | 73000000 | 0.29 | -0.09 | A>A |
| chr8 | 73000000 | 73100000 | 0.22 | -0.15 | A>A |
| chr8 | 73100000 | 73200000 | -0.25 | -0.65 | B>B |
| chr8 | 73200000 | 73300000 | -0.43 | -0.69 | B>B |
| chr8 | 73300000 | 73400000 | -0.59 | -0.99 | B>B |
| chr8 | 73400000 | 73500000 | -0.42 | -0.97 | B>B |
| chr8 | 73500000 | 73600000 | -0.47 | -1.08 | A>B |
| chr8 | 73600000 | 73700000 | -0.48 | -1.06 | B>B |
| chr8 | 73700000 | 73800000 | -0.30 | -0.86 | B>B |
| chr8 | 73800000 | 73900000 | -0.09 | -0.63 | B>B |
| chr8 | 73900000 | 74000000 | 0.36 | -0.02 | A>A |
| chr8 | 74000000 | 74100000 | 0.32 | -0.08 | A>A |
| chr8 | 74100000 | 74200000 | 0.51 | 0.19 | A>A |
| chr8 | 74200000 | 74300000 | 0.88 | 0.87 | A>A |
| chr8 | 74300000 | 74400000 | 0.91 | 0.80 | A>A |
| chr8 | 74400000 | 74500000 | 0.65 | 0.48 | A>A |
| chr8 | 74500000 | 74600000 | 0.53 | 0.43 | A>A |
| chr8 | 74600000 | 74700000 | 0.67 | 0.60 | A>A |
| chr8 | 74700000 | 74800000 | 0.85 | 0.83 | A>A |
| chr8 | 74800000 | 74900000 | 0.82 | 0.64 | A>A |
| chr8 | 74900000 | 75000000 | 0.34 | 0.09 | A>A |
| chr8 | 75000000 | 75100000 | -0.03 | -0.38 | B>B |
| chr8 | 75100000 | 75200000 | -0.16 | -0.59 | B>B |
| chr8 | 75200000 | 75300000 | -0.15 | -0.42 | B>B |
| chr8 | 75300000 | 75400000 | -0.60 | -0.92 | B>B |
| chr8 | 75400000 | 75500000 | -0.91 | -1.04 | B>B |
| chr8 | 75500000 | 75600000 | -0.97 | -1.06 | B>B |
| chr8 | 75600000 | 75700000 | -0.95 | -1.13 | B>B |
| chr8 | 75700000 | 75800000 | -0.71 | -0.96 | B>B |
| chr8 | 75800000 | 75900000 | -0.94 | -1.01 | B>B |
| chr8 | 75900000 | 76000000 | -1.10 | -1.02 | B>B |
| chr8 | 76000000 | 76100000 | -1.07 | -0.78 | B>B |
| chr8 | 76100000 | 76200000 | -0.91 | -0.47 | B>B |
| chr8 | 76200000 | 76300000 | -0.44 | -0.16 | B>B |
| chr8 | 76300000 | 76400000 | -0.66 | 0.17 | B>A |
| chr8 | 76400000 | 76500000 | -0.97 | -0.24 | B>A |
| chr8 | 76500000 | 76600000 | -1.02 | -0.74 | B>B |
| chr8 | 76600000 | 76700000 | -1.12 | -0.90 | B>B |
| chr8 | 76700000 | 76800000 | -1.18 | -0.99 | B>B |
| chr8 | 76800000 | 76900000 | -1.07 | -0.97 | B>B |
| chr8 | 76900000 | 77000000 | -1.10 | -1.01 | B>B |
| chr8 | 77000000 | 77100000 | -1.10 | -0.95 | B>B |
| chr8 | 77100000 | 77200000 | -1.07 | -0.89 | B>B |
| chr8 | 77200000 | 77300000 | -0.94 | -0.63 | B>B |
| chr8 | 77300000 | 77400000 | -0.93 | -0.28 | B>A |
| chr8 | 77400000 | 77500000 | -0.84 | -0.20 | B>A |
| chr8 | 77500000 | 77600000 | -0.54 | -0.11 | B>B |
| chr8 | 77600000 | 77700000 | -0.31 | -0.03 | B>B |
| chr8 | 77700000 | 77800000 | -0.46 | -0.21 | B>B |
| chr8 | 77800000 | 77900000 | -0.40 | -0.20 | B>B |
| chr8 | 77900000 | 78000000 | -0.58 | -0.33 | B>B |

| | | | | | |
|---|---|---|---|---|---|
| chr8 | 78000000 | 78100000 | -0.85 | -0.53 | B>B |
| chr8 | 78100000 | 78200000 | -0.98 | -0.85 | B>B |
| chr8 | 78200000 | 78300000 | -0.99 | -0.83 | B>B |
| chr8 | 78300000 | 78400000 | -1.09 | -0.88 | B>B |
| chr8 | 78400000 | 78500000 | -1.07 | -0.92 | B>B |
| chr8 | 78500000 | 78600000 | -1.16 | -0.97 | B>B |
| chr8 | 78600000 | 78700000 | -1.17 | -1.03 | B>B |
| chr8 | 78700000 | 78800000 | -1.15 | -1.00 | B>B |
| chr8 | 78800000 | 78900000 | -1.22 | -1.01 | B>B |
| chr8 | 78900000 | 79000000 | -1.24 | -1.05 | B>B |
| chr8 | 79000000 | 79100000 | -1.23 | -1.03 | B>B |
| chr8 | 79100000 | 79200000 | -1.17 | -0.98 | B>B |
| chr8 | 79200000 | 79300000 | -1.23 | -1.07 | B>B |
| chr8 | 79300000 | 79400000 | -1.15 | -1.01 | B>B |
| chr8 | 79400000 | 79500000 | -0.76 | -0.96 | B>B |
| chr8 | 79500000 | 79600000 | -0.39 | -0.69 | B>B |
| chr8 | 79600000 | 79700000 | -0.42 | -0.54 | B>B |
| chr8 | 79700000 | 79800000 | -0.72 | -0.73 | B>B |
| chr8 | 79800000 | 79900000 | -0.94 | -0.68 | B>B |
| chr8 | 79900000 | 80000000 | -1.03 | -0.54 | B>B |
| chr8 | 80000000 | 80100000 | -1.07 | -0.71 | B>B |
| chr8 | 80100000 | 80200000 | -1.07 | -0.83 | B>B |
| chr8 | 80200000 | 80300000 | -0.91 | -0.92 | B>B |
| chr8 | 80300000 | 80400000 | -0.73 | -1.01 | B>B |
| chr8 | 80400000 | 80500000 | -0.57 | -0.89 | B>B |
| chr8 | 80500000 | 80600000 | -0.04 | -0.30 | B>B |
| chr8 | 80600000 | 80700000 | 0.42 | 0.18 | A>A |
| chr8 | 80700000 | 80800000 | 0.62 | 0.38 | A>A |
| chr8 | 80800000 | 80900000 | 0.67 | 0.47 | A>A |
| chr8 | 80900000 | 81000000 | 0.58 | 0.48 | A>A |
| chr8 | 81000000 | 81100000 | 0.92 | 0.81 | A>A |
| chr8 | 81100000 | 81200000 | 0.64 | 0.60 | A>A |
| chr8 | 81200000 | 81300000 | 0.75 | 0.66 | A>A |
| chr8 | 81300000 | 81400000 | 0.69 | 0.73 | A>A |
| chr8 | 81400000 | 81500000 | 0.65 | 0.64 | A>A |
| chr8 | 81500000 | 81600000 | 0.24 | 0.52 | A>A |
| chr8 | 81600000 | 81700000 | 0.08 | 0.66 | B>A |
| chr8 | 81700000 | 81800000 | 0.02 | 0.71 | B>A |
| chr8 | 81800000 | 81900000 | 0.24 | 0.76 | B>A |
| chr8 | 81900000 | 82000000 | 0.16 | 0.45 | A>A |
| chr8 | 82000000 | 82100000 | 0.15 | 0.44 | A>A |
| chr8 | 82100000 | 82200000 | 0.05 | 0.30 | A>A |
| chr8 | 82200000 | 82300000 | -0.21 | -0.23 | B>B |
| chr8 | 82300000 | 82400000 | -0.45 | -0.46 | B>B |
| chr8 | 82400000 | 82500000 | -0.21 | -0.28 | B>B |
| chr8 | 82500000 | 82600000 | 0.26 | 0.21 | A>A |
| chr8 | 82600000 | 82700000 | 0.19 | 0.39 | A>A |
| chr8 | 82700000 | 82800000 | -0.06 | 0.23 | A>A |
| chr8 | 82800000 | 82900000 | -0.61 | -0.01 | B>A |
| chr8 | 82900000 | 83000000 | -0.80 | 0.30 | B>A |
| chr8 | 83000000 | 83100000 | -0.92 | 0.43 | B>A |
| chr8 | 83100000 | 83200000 | -0.94 | 0.42 | B>A |
| chr8 | 83200000 | 83300000 | -0.83 | 0.21 | B>A |
| chr8 | 83300000 | 83400000 | -0.71 | 0.19 | B>A |
| chr8 | 83400000 | 83500000 | -0.47 | 0.17 | B>A |
| chr8 | 83500000 | 83600000 | -0.60 | 0.19 | B>A |

| | | | | | |
|---|---|---|---|---|---|
| chr8 | 83600000 | 83700000 | -0.80 | 0.12 | B>A |
| chr8 | 83700000 | 83800000 | -0.84 | 0.19 | B>A |
| chr8 | 83800000 | 83900000 | -0.87 | 0.14 | B>A |
| chr8 | 83900000 | 84000000 | -0.93 | 0.13 | B>A |
| chr8 | 84000000 | 84100000 | -0.96 | -0.01 | B>A |
| chr8 | 84100000 | 84200000 | -0.97 | -0.10 | B>A |
| chr8 | 84200000 | 84300000 | -1.07 | -0.29 | B>A |
| chr8 | 84300000 | 84400000 | -1.04 | -0.46 | B>A |
| chr8 | 84400000 | 84500000 | -1.09 | -0.58 | B>B |
| chr8 | 84500000 | 84600000 | -1.08 | -0.72 | B>B |
| chr8 | 84600000 | 84700000 | -1.17 | -0.77 | B>B |
| chr8 | 84700000 | 84800000 | -1.09 | -0.82 | B>B |
| chr8 | 84800000 | 84900000 | -1.15 | -0.90 | B>B |
| chr8 | 84900000 | 85000000 | -1.20 | -0.89 | B>B |
| chr8 | 85000000 | 85100000 | -1.10 | -0.84 | B>B |
| chr8 | 85100000 | 85200000 | -1.14 | -0.88 | B>B |
| chr8 | 85200000 | 85300000 | -1.26 | -1.03 | B>B |
| chr8 | 85300000 | 85400000 | -1.22 | -1.08 | B>B |
| chr8 | 85400000 | 85500000 | -1.20 | -0.97 | B>B |
| chr8 | 85500000 | 85600000 | -1.02 | -0.84 | B>B |
| chr8 | 85600000 | 85700000 | -1.26 | -1.22 | B>B |
| chr8 | 85700000 | 85800000 | -1.26 | -1.17 | B>B |
| chr8 | 85800000 | 85900000 | -1.18 | -1.17 | B>B |
| chr8 | 85900000 | 86000000 | -0.67 | -0.91 | B>B |
| chr8 | 86000000 | 86100000 | -0.01 | -0.26 | B>B |
| chr8 | 86100000 | 86200000 | 0.11 | -0.08 | A>A |
| chr8 | 86200000 | 86300000 | -0.34 | -0.70 | B>B |
| chr8 | 86300000 | 86400000 | -0.38 | -0.63 | B>B |
| chr8 | 86400000 | 86500000 | -0.54 | -0.34 | B>B |
| chr8 | 86500000 | 86600000 | 0.00 | 0.00 | NO |
| chr8 | 86600000 | 86700000 | 0.00 | 0.00 | NO |
| chr8 | 86700000 | 86800000 | 0.00 | 0.00 | NO |
| chr8 | 86800000 | 86900000 | 0.00 | -0.04 | NO |
| chr8 | 86900000 | 87000000 | -0.15 | -0.33 | B>B |
| chr8 | 87000000 | 87100000 | 0.21 | -0.15 | A>A |
| chr8 | 87100000 | 87200000 | 0.79 | 0.48 | A>A |
| chr8 | 87200000 | 87300000 | 0.98 | 0.74 | A>A |
| chr8 | 87300000 | 87400000 | 0.86 | 0.75 | A>A |
| chr8 | 87400000 | 87500000 | 0.77 | 0.73 | A>A |
| chr8 | 87500000 | 87600000 | 0.39 | 0.22 | A>A |
| chr8 | 87600000 | 87700000 | -0.30 | -0.53 | B>B |
| chr8 | 87700000 | 87800000 | -0.58 | -0.68 | B>B |
| chr8 | 87800000 | 87900000 | -0.80 | -0.57 | B>B |
| chr8 | 87900000 | 88000000 | -1.02 | -0.36 | B>A |
| chr8 | 88000000 | 88100000 | -1.09 | -0.28 | B>A |
| chr8 | 88100000 | 88200000 | -1.08 | -0.20 | B>A |
| chr8 | 88200000 | 88300000 | -1.10 | -0.06 | B>A |
| chr8 | 88300000 | 88400000 | -1.16 | 0.01 | B>A |
| chr8 | 88400000 | 88500000 | -1.07 | -0.07 | B>A |
| chr8 | 88500000 | 88600000 | -1.02 | -0.19 | B>A |
| chr8 | 88600000 | 88700000 | -1.04 | -0.50 | B>A |
| chr8 | 88700000 | 88800000 | -1.04 | -0.77 | B>B |
| chr8 | 88800000 | 88900000 | -1.04 | -0.88 | B>B |
| chr8 | 88900000 | 89000000 | -1.04 | -0.90 | B>B |
| chr8 | 89000000 | 89100000 | -1.04 | -0.91 | B>B |
| chr8 | 89100000 | 89200000 | -1.01 | -1.00 | B>B |

| | | | | | |
|---|---|---|---|---|---|
| chr8 | 89200000 | 89300000 | -0.93 | -0.86 | B>B |
| chr8 | 89300000 | 89400000 | -0.74 | -0.64 | B>B |
| chr8 | 89400000 | 89500000 | -0.91 | -0.91 | B>B |
| chr8 | 89500000 | 89600000 | -0.90 | -0.93 | B>B |
| chr8 | 89600000 | 89700000 | -0.88 | -0.87 | B>B |
| chr8 | 89700000 | 89800000 | -0.64 | -0.81 | B>B |
| chr8 | 89800000 | 89900000 | -0.66 | -0.69 | B>B |
| chr8 | 89900000 | 90000000 | -0.62 | -0.63 | B>B |
| chr8 | 90000000 | 90100000 | -0.65 | -0.27 | B>B |
| chr8 | 90100000 | 90200000 | -0.46 | 0.05 | B>A |
| chr8 | 90200000 | 90300000 | -0.20 | 0.13 | B>B |
| chr8 | 90300000 | 90400000 | -0.07 | 0.15 | A>A |
| chr8 | 90400000 | 90500000 | 0.07 | 0.21 | A>A |
| chr8 | 90500000 | 90600000 | 0.06 | 0.25 | A>A |
| chr8 | 90600000 | 90700000 | -0.25 | -0.02 | B>B |
| chr8 | 90700000 | 90800000 | 0.41 | 0.44 | A>A |
| chr8 | 90800000 | 90900000 | 0.77 | 0.77 | A>A |
| chr8 | 90900000 | 91000000 | 0.73 | 0.65 | A>A |
| chr8 | 91000000 | 91100000 | 0.01 | -0.14 | B>B |
| chr8 | 91100000 | 91200000 | -0.27 | -0.56 | B>B |
| chr8 | 91200000 | 91300000 | -0.37 | -0.73 | B>B |
| chr8 | 91300000 | 91400000 | -0.35 | -0.63 | B>B |
| chr8 | 91400000 | 91500000 | -0.26 | -0.59 | B>B |
| chr8 | 91500000 | 91600000 | -0.11 | -0.43 | B>B |
| chr8 | 91600000 | 91700000 | 0.20 | -0.05 | A>A |
| chr8 | 91700000 | 91800000 | -0.10 | -0.41 | B>B |
| chr8 | 91800000 | 91900000 | -0.25 | -0.61 | B>B |
| chr8 | 91900000 | 92000000 | -0.07 | -0.33 | B>B |
| chr8 | 92000000 | 92100000 | 0.36 | 0.10 | A>A |
| chr8 | 92100000 | 92200000 | -0.16 | -0.49 | B>B |
| chr8 | 92200000 | 92300000 | -0.45 | -0.85 | B>B |
| chr8 | 92300000 | 92400000 | -0.59 | -0.98 | B>B |
| chr8 | 92400000 | 92500000 | -0.79 | -1.02 | B>B |
| chr8 | 92500000 | 92600000 | -1.18 | -1.07 | B>B |
| chr8 | 92600000 | 92700000 | -1.21 | -1.05 | B>B |
| chr8 | 92700000 | 92800000 | -1.22 | -1.02 | B>B |
| chr8 | 92800000 | 92900000 | -1.09 | -1.02 | B>B |
| chr8 | 92900000 | 93000000 | -0.92 | -1.02 | B>B |
| chr8 | 93000000 | 93100000 | -0.56 | -0.72 | B>B |
| chr8 | 93100000 | 93200000 | -0.61 | -0.72 | B>B |
| chr8 | 93200000 | 93300000 | -0.86 | -0.88 | B>B |
| chr8 | 93300000 | 93400000 | -0.88 | -0.86 | B>B |
| chr8 | 93400000 | 93500000 | -0.79 | -0.81 | B>B |
| chr8 | 93500000 | 93600000 | -0.72 | -0.85 | B>B |
| chr8 | 93600000 | 93700000 | -0.61 | -0.74 | B>B |
| chr8 | 93700000 | 93800000 | -0.55 | -0.63 | B>B |
| chr8 | 93800000 | 93900000 | -0.37 | -0.32 | B>B |
| chr8 | 93900000 | 94000000 | -0.06 | 0.09 | A>A |
| chr8 | 94000000 | 94100000 | -0.19 | -0.14 | B>B |
| chr8 | 94100000 | 94200000 | -0.24 | -0.32 | B>B |
| chr8 | 94200000 | 94300000 | -0.25 | -0.65 | B>B |
| chr8 | 94300000 | 94400000 | -0.16 | -0.67 | B>B |
| chr8 | 94400000 | 94500000 | -0.09 | -0.77 | A>B |
| chr8 | 94500000 | 94600000 | 0.06 | -0.76 | A>B |
| chr8 | 94600000 | 94700000 | 0.07 | -0.30 | B>B |
| chr8 | 94700000 | 94800000 | 0.57 | 0.49 | A>A |

| | | | | | |
|---|---|---|---|---|---|
| chr8 | 94800000 | 94900000 | 0.88 | 0.74 | A>A |
| chr8 | 94900000 | 95000000 | 0.79 | 0.65 | A>A |
| chr8 | 95000000 | 95100000 | 0.89 | 0.83 | A>A |
| chr8 | 95100000 | 95200000 | 1.17 | 1.00 | A>A |
| chr8 | 95200000 | 95300000 | 1.10 | 1.03 | A>A |
| chr8 | 95300000 | 95400000 | 0.92 | 0.86 | A>A |
| chr8 | 95400000 | 95500000 | 1.16 | 1.04 | A>A |
| chr8 | 95500000 | 95600000 | 1.05 | 0.97 | A>A |
| chr8 | 95600000 | 95700000 | 1.10 | 0.91 | A>A |
| chr8 | 95700000 | 95800000 | 1.08 | 0.88 | A>A |
| chr8 | 95800000 | 95900000 | 1.11 | 0.87 | A>A |
| chr8 | 95900000 | 96000000 | 1.00 | 1.05 | A>A |
| chr8 | 96000000 | 96100000 | 0.91 | 0.88 | A>A |
| chr8 | 96100000 | 96200000 | 0.98 | 0.82 | A>A |
| chr8 | 96200000 | 96300000 | 0.66 | 0.43 | A>A |
| chr8 | 96300000 | 96400000 | 0.02 | -0.72 | A>B |
| chr8 | 96400000 | 96500000 | -0.29 | -1.13 | A>B |
| chr8 | 96500000 | 96600000 | -0.57 | -1.14 | B>B |
| chr8 | 96600000 | 96700000 | -0.51 | -1.03 | B>B |
| chr8 | 96700000 | 96800000 | -0.37 | -1.01 | A>B |
| chr8 | 96800000 | 96900000 | -0.21 | -0.94 | A>B |
| chr8 | 96900000 | 97000000 | -0.44 | -1.03 | A>B |
| chr8 | 97000000 | 97100000 | -0.50 | -1.09 | A>B |
| chr8 | 97100000 | 97200000 | -0.10 | -0.62 | B>B |
| chr8 | 97200000 | 97300000 | 0.28 | 0.00 | A>A |
| chr8 | 97300000 | 97400000 | 0.03 | -0.33 | B>B |
| chr8 | 97400000 | 97500000 | -0.23 | -0.89 | A>B |
| chr8 | 97500000 | 97600000 | -0.26 | -0.93 | A>B |
| chr8 | 97600000 | 97700000 | -0.44 | -0.83 | B>B |
| chr8 | 97700000 | 97800000 | -0.72 | -1.04 | B>B |
| chr8 | 97800000 | 97900000 | -0.64 | -1.21 | B>B |
| chr8 | 97900000 | 98000000 | -0.60 | -1.12 | B>B |
| chr8 | 98000000 | 98100000 | -0.60 | -1.23 | A>B |
| chr8 | 98100000 | 98200000 | -0.46 | -1.08 | A>B |
| chr8 | 98200000 | 98300000 | -0.04 | -0.82 | A>B |
| chr8 | 98300000 | 98400000 | 0.15 | -0.88 | A>B |
| chr8 | 98400000 | 98500000 | 0.12 | -0.70 | A>B |
| chr8 | 98500000 | 98600000 | 0.21 | -0.25 | B>B |
| chr8 | 98600000 | 98700000 | 0.77 | 0.60 | A>A |
| chr8 | 98700000 | 98800000 | 0.92 | 0.87 | A>A |
| chr8 | 98800000 | 98900000 | 0.97 | 0.92 | A>A |
| chr8 | 98900000 | 99000000 | 1.02 | 0.90 | A>A |
| chr8 | 99000000 | 99100000 | 1.00 | 0.86 | A>A |
| chr8 | 99100000 | 99200000 | 0.75 | 0.79 | A>A |
| chr8 | 99200000 | 99300000 | 0.66 | 0.67 | A>A |
| chr8 | 99300000 | 99400000 | 0.53 | 0.63 | A>A |
| chr8 | 99400000 | 99500000 | 0.47 | 0.59 | A>A |
| chr8 | 99500000 | 99600000 | 0.46 | 0.66 | A>A |
| chr8 | 99600000 | 99700000 | 0.65 | 0.77 | A>A |
| chr8 | 99700000 | 99800000 | 0.81 | 0.87 | A>A |
| chr8 | 99800000 | 99900000 | 0.77 | 0.91 | A>A |
| chr8 | 99900000 | 100000000 | 0.74 | 0.80 | A>A |
| chr8 | 1E+08 | 100100000 | 0.75 | 0.80 | A>A |
| chr8 | 1E+08 | 100200000 | 0.72 | 0.76 | A>A |
| chr8 | 1E+08 | 100300000 | 0.62 | 0.68 | A>A |
| chr8 | 1E+08 | 100400000 | 0.47 | 0.62 | A>A |

| | | | | | |
|---|---|---|---|---|---|
| chr8 | 1E+08 | 100500000 | 0.47 | 0.61 | A>A |
| chr8 | 1.01E+08 | 100600000 | 0.62 | 0.65 | A>A |
| chr8 | 1.01E+08 | 100700000 | 0.51 | 0.59 | A>A |
| chr8 | 1.01E+08 | 100800000 | 0.47 | 0.57 | A>A |
| chr8 | 1.01E+08 | 100900000 | 0.46 | 0.61 | A>A |
| chr8 | 1.01E+08 | 101000000 | 0.45 | 0.49 | A>A |
| chr8 | 1.01E+08 | 101100000 | 0.50 | 0.50 | A>A |
| chr8 | 1.01E+08 | 101200000 | 0.73 | 0.73 | A>A |
| chr8 | 1.01E+08 | 101300000 | 0.91 | 0.95 | A>A |
| chr8 | 1.01E+08 | 101400000 | 1.02 | 1.10 | A>A |
| chr8 | 1.01E+08 | 101500000 | 1.09 | 1.12 | A>A |
| chr8 | 1.02E+08 | 101600000 | 0.81 | 0.86 | A>A |
| chr8 | 1.02E+08 | 101700000 | 1.00 | 0.96 | A>A |
| chr8 | 1.02E+08 | 101800000 | 1.09 | 1.01 | A>A |
| chr8 | 1.02E+08 | 101900000 | 0.88 | 0.94 | A>A |
| chr8 | 1.02E+08 | 102000000 | 1.04 | 1.10 | A>A |
| chr8 | 1.02E+08 | 102100000 | 1.19 | 1.17 | A>A |
| chr8 | 1.02E+08 | 102200000 | 1.03 | 0.98 | A>A |
| chr8 | 1.02E+08 | 102300000 | 0.84 | 0.98 | A>A |
| chr8 | 1.02E+08 | 102400000 | 0.89 | 0.97 | A>A |
| chr8 | 1.02E+08 | 102500000 | 1.10 | 1.21 | A>A |
| chr8 | 1.03E+08 | 102600000 | 1.03 | 0.95 | A>A |
| chr8 | 1.03E+08 | 102700000 | 0.80 | 0.86 | A>A |
| chr8 | 1.03E+08 | 102800000 | 0.98 | 1.00 | A>A |
| chr8 | 1.03E+08 | 102900000 | 0.96 | 0.97 | A>A |
| chr8 | 1.03E+08 | 103000000 | 0.93 | 1.03 | A>A |
| chr8 | 1.03E+08 | 103100000 | 0.92 | 1.00 | A>A |
| chr8 | 1.03E+08 | 103200000 | 0.86 | 0.93 | A>A |
| chr8 | 1.03E+08 | 103300000 | 0.86 | 0.85 | A>A |
| chr8 | 1.03E+08 | 103400000 | 1.06 | 1.07 | A>A |
| chr8 | 1.03E+08 | 103500000 | 1.17 | 1.12 | A>A |
| chr8 | 1.04E+08 | 103600000 | 1.17 | 1.14 | A>A |
| chr8 | 1.04E+08 | 103700000 | 1.13 | 1.21 | A>A |
| chr8 | 1.04E+08 | 103800000 | 1.37 | 1.29 | A>A |
| chr8 | 1.04E+08 | 103900000 | 1.03 | 0.91 | A>A |
| chr8 | 1.04E+08 | 104000000 | 1.05 | 0.86 | A>A |
| chr8 | 1.04E+08 | 104100000 | 0.85 | 0.60 | A>A |
| chr8 | 1.04E+08 | 104200000 | 0.41 | -0.04 | A>A |
| chr8 | 1.04E+08 | 104300000 | 0.21 | -0.18 | A>A |
| chr8 | 1.04E+08 | 104400000 | 0.46 | 0.18 | A>A |
| chr8 | 1.04E+08 | 104500000 | 0.46 | 0.21 | A>A |
| chr8 | 1.05E+08 | 104600000 | -0.31 | -0.52 | B>B |
| chr8 | 1.05E+08 | 104700000 | -0.68 | -1.06 | B>B |
| chr8 | 1.05E+08 | 104800000 | -0.43 | -0.74 | B>B |
| chr8 | 1.05E+08 | 104900000 | -0.64 | -0.71 | B>B |
| chr8 | 1.05E+08 | 105000000 | -0.84 | -0.77 | B>B |
| chr8 | 1.05E+08 | 105100000 | -0.82 | -0.72 | B>B |
| chr8 | 1.05E+08 | 105200000 | -0.70 | -0.65 | B>B |
| chr8 | 1.05E+08 | 105300000 | -0.49 | -0.67 | B>B |
| chr8 | 1.05E+08 | 105400000 | -0.26 | -0.65 | B>B |
| chr8 | 1.05E+08 | 105500000 | -0.15 | -0.46 | B>B |
| chr8 | 1.06E+08 | 105600000 | 0.21 | -0.09 | A>A |
| chr8 | 1.06E+08 | 105700000 | 0.10 | -0.41 | B>B |
| chr8 | 1.06E+08 | 105800000 | -0.08 | -0.50 | B>B |
| chr8 | 1.06E+08 | 105900000 | -0.63 | -0.84 | B>B |
| chr8 | 1.06E+08 | 106000000 | -0.87 | -1.07 | B>B |

| | | | | | |
|---|---|---|---|---|---|
| chr8 | 1.06E+08 | 106100000 | -0.93 | -1.05 | B>B |
| chr8 | 1.06E+08 | 106200000 | -0.97 | -1.07 | B>B |
| chr8 | 1.06E+08 | 106300000 | -0.93 | -0.98 | B>B |
| chr8 | 1.06E+08 | 106400000 | -0.80 | -0.73 | B>B |
| chr8 | 1.06E+08 | 106500000 | -0.84 | -0.66 | B>B |
| chr8 | 1.07E+08 | 106600000 | -0.80 | -0.65 | B>B |
| chr8 | 1.07E+08 | 106700000 | -0.67 | -0.64 | B>B |
| chr8 | 1.07E+08 | 106800000 | -0.42 | -0.44 | B>B |
| chr8 | 1.07E+08 | 106900000 | 0.02 | 0.04 | A>A |
| chr8 | 1.07E+08 | 107000000 | 0.21 | 0.35 | A>A |
| chr8 | 1.07E+08 | 107100000 | 0.48 | 0.64 | A>A |
| chr8 | 1.07E+08 | 107200000 | 0.37 | 0.53 | A>A |
| chr8 | 1.07E+08 | 107300000 | 0.08 | 0.39 | A>A |
| chr8 | 1.07E+08 | 107400000 | -0.22 | 0.20 | B>B |
| chr8 | 1.07E+08 | 107500000 | -0.31 | 0.22 | B>A |
| chr8 | 1.08E+08 | 107600000 | -0.15 | 0.20 | A>A |
| chr8 | 1.08E+08 | 107700000 | 0.12 | 0.30 | A>A |
| chr8 | 1.08E+08 | 107800000 | 0.33 | 0.21 | A>A |
| chr8 | 1.08E+08 | 107900000 | 0.35 | 0.18 | A>A |
| chr8 | 1.08E+08 | 108000000 | 0.45 | 0.19 | A>A |
| chr8 | 1.08E+08 | 108100000 | 0.49 | 0.15 | A>A |
| chr8 | 1.08E+08 | 108200000 | 0.81 | 0.59 | A>A |
| chr8 | 1.08E+08 | 108300000 | 0.81 | 0.61 | A>A |
| chr8 | 1.08E+08 | 108400000 | 0.66 | 0.50 | A>A |
| chr8 | 1.08E+08 | 108500000 | 0.65 | 0.54 | A>A |
| chr8 | 1.09E+08 | 108600000 | 0.32 | 0.12 | A>A |
| chr8 | 1.09E+08 | 108700000 | -0.05 | -0.70 | A>B |
| chr8 | 1.09E+08 | 108800000 | -0.13 | -0.68 | B>B |
| chr8 | 1.09E+08 | 108900000 | -0.40 | -0.83 | B>B |
| chr8 | 1.09E+08 | 109000000 | -0.44 | -0.90 | B>B |
| chr8 | 1.09E+08 | 109100000 | -0.44 | -0.95 | B>B |
| chr8 | 1.09E+08 | 109200000 | -0.22 | -0.79 | B>B |
| chr8 | 1.09E+08 | 109300000 | 0.09 | -0.34 | B>B |
| chr8 | 1.09E+08 | 109400000 | -0.32 | -0.86 | B>B |
| chr8 | 1.09E+08 | 109500000 | -0.05 | -0.49 | B>B |
| chr8 | 1.1E+08 | 109600000 | -0.30 | -0.75 | B>B |
| chr8 | 1.1E+08 | 109700000 | -0.33 | -0.94 | A>B |
| chr8 | 1.1E+08 | 109800000 | -0.20 | -0.76 | B>B |
| chr8 | 1.1E+08 | 109900000 | -0.15 | -0.57 | B>B |
| chr8 | 1.1E+08 | 110000000 | 0.45 | 0.17 | A>A |
| chr8 | 1.1E+08 | 110100000 | 0.70 | 0.24 | A>A |
| chr8 | 1.1E+08 | 110200000 | 0.51 | 0.12 | A>A |
| chr8 | 1.1E+08 | 110300000 | 0.46 | 0.18 | A>A |
| chr8 | 1.1E+08 | 110400000 | 0.15 | -0.10 | A>A |
| chr8 | 1.1E+08 | 110500000 | -0.24 | -0.51 | B>B |
| chr8 | 1.11E+08 | 110600000 | 0.06 | -0.05 | A>A |
| chr8 | 1.11E+08 | 110700000 | 0.11 | 0.18 | A>A |
| chr8 | 1.11E+08 | 110800000 | -0.52 | -0.05 | B>B |
| chr8 | 1.11E+08 | 110900000 | -0.95 | 0.11 | B>A |
| chr8 | 1.11E+08 | 111000000 | -0.90 | 0.03 | B>A |
| chr8 | 1.11E+08 | 111100000 | -0.96 | -0.35 | B>A |
| chr8 | 1.11E+08 | 111200000 | -1.04 | -0.84 | B>B |
| chr8 | 1.11E+08 | 111300000 | -1.11 | -0.93 | B>B |
| chr8 | 1.11E+08 | 111400000 | -1.06 | -1.00 | B>B |
| chr8 | 1.11E+08 | 111500000 | -1.06 | -1.03 | B>B |
| chr8 | 1.12E+08 | 111600000 | -1.08 | -0.92 | B>B |

| | | | | | |
|---|---|---|---|---|---|
| chr8 | 1.12E+08 | 111700000 | -1.06 | -0.89 | B>B |
| chr8 | 1.12E+08 | 111800000 | -0.92 | -0.74 | B>B |
| chr8 | 1.12E+08 | 111900000 | -0.57 | -0.17 | B>B |
| chr8 | 1.12E+08 | 112000000 | -0.69 | -0.04 | B>A |
| chr8 | 1.12E+08 | 112100000 | -0.81 | 0.03 | B>A |
| chr8 | 1.12E+08 | 112200000 | -0.90 | -0.25 | B>A |
| chr8 | 1.12E+08 | 112300000 | -0.93 | -0.30 | B>A |
| chr8 | 1.12E+08 | 112400000 | -0.97 | -0.42 | B>A |
| chr8 | 1.12E+08 | 112500000 | -0.96 | -0.46 | B>B |
| chr8 | 1.13E+08 | 112600000 | -1.06 | -0.65 | B>B |
| chr8 | 1.13E+08 | 112700000 | -1.07 | -0.58 | B>B |
| chr8 | 1.13E+08 | 112800000 | -1.07 | -0.56 | B>A |
| chr8 | 1.13E+08 | 112900000 | -1.02 | -0.22 | B>A |
| chr8 | 1.13E+08 | 113000000 | -1.07 | -0.12 | B>A |
| chr8 | 1.13E+08 | 113100000 | -1.14 | -0.01 | B>A |
| chr8 | 1.13E+08 | 113200000 | -1.10 | 0.02 | B>A |
| chr8 | 1.13E+08 | 113300000 | -1.12 | 0.01 | B>A |
| chr8 | 1.13E+08 | 113400000 | -1.09 | -0.02 | B>A |
| chr8 | 1.13E+08 | 113500000 | -1.12 | 0.00 | B>A |
| chr8 | 1.14E+08 | 113600000 | -1.12 | 0.02 | B>A |
| chr8 | 1.14E+08 | 113700000 | -1.14 | 0.01 | B>A |
| chr8 | 1.14E+08 | 113800000 | -1.04 | 0.02 | B>A |
| chr8 | 1.14E+08 | 113900000 | -1.03 | 0.00 | B>A |
| chr8 | 1.14E+08 | 114000000 | -1.06 | 0.05 | B>A |
| chr8 | 1.14E+08 | 114100000 | -1.21 | -0.02 | B>A |
| chr8 | 1.14E+08 | 114200000 | -1.15 | -0.03 | B>A |
| chr8 | 1.14E+08 | 114300000 | -1.13 | 0.00 | B>A |
| chr8 | 1.14E+08 | 114400000 | -1.02 | 0.02 | B>A |
| chr8 | 1.14E+08 | 114500000 | -0.89 | 0.00 | B>A |
| chr8 | 1.15E+08 | 114600000 | -1.08 | 0.03 | B>A |
| chr8 | 1.15E+08 | 114700000 | -1.04 | 0.03 | B>A |
| chr8 | 1.15E+08 | 114800000 | -1.12 | 0.08 | B>A |
| chr8 | 1.15E+08 | 114900000 | -1.10 | 0.02 | B>A |
| chr8 | 1.15E+08 | 115000000 | -1.18 | 0.03 | B>A |
| chr8 | 1.15E+08 | 115100000 | -1.07 | 0.10 | B>A |
| chr8 | 1.15E+08 | 115200000 | -0.95 | 0.13 | B>A |
| chr8 | 1.15E+08 | 115300000 | -1.00 | 0.22 | B>A |
| chr8 | 1.15E+08 | 115400000 | -1.06 | 0.41 | B>A |
| chr8 | 1.15E+08 | 115500000 | -0.93 | 0.25 | B>A |
| chr8 | 1.16E+08 | 115600000 | -0.96 | 0.34 | B>A |
| chr8 | 1.16E+08 | 115700000 | -0.94 | 0.36 | B>A |
| chr8 | 1.16E+08 | 115800000 | -0.98 | 0.36 | B>A |
| chr8 | 1.16E+08 | 115900000 | -0.92 | 0.37 | B>A |
| chr8 | 1.16E+08 | 116000000 | -1.05 | 0.59 | B>A |
| chr8 | 1.16E+08 | 116100000 | -0.86 | 0.60 | B>A |
| chr8 | 1.16E+08 | 116200000 | -0.86 | 0.58 | B>A |
| chr8 | 1.16E+08 | 116300000 | -0.85 | 0.70 | B>A |
| chr8 | 1.16E+08 | 116400000 | -0.81 | 0.62 | B>A |
| chr8 | 1.16E+08 | 116500000 | -0.82 | 0.62 | B>A |
| chr8 | 1.17E+08 | 116600000 | -0.80 | 0.76 | B>A |
| chr8 | 1.17E+08 | 116700000 | -0.52 | 0.69 | B>A |
| chr8 | 1.17E+08 | 116800000 | -0.63 | 0.81 | B>A |
| chr8 | 1.17E+08 | 116900000 | -0.74 | 0.78 | B>A |
| chr8 | 1.17E+08 | 117000000 | -0.71 | 0.77 | B>A |
| chr8 | 1.17E+08 | 117100000 | -0.78 | 0.73 | B>A |
| chr8 | 1.17E+08 | 117200000 | -0.70 | 0.78 | B>A |

| | | | | | |
|---|---|---|---|---|---|
| chr8 | 1.17E+08 | 117300000 | -0.64 | 0.69 | B>A |
| chr8 | 1.17E+08 | 117400000 | -0.53 | 0.67 | B>A |
| chr8 | 1.17E+08 | 117500000 | -0.22 | 0.63 | B>A |
| chr8 | 1.18E+08 | 117600000 | 0.09 | 0.61 | B>A |
| chr8 | 1.18E+08 | 117700000 | 0.32 | 0.63 | A>A |
| chr8 | 1.18E+08 | 117800000 | 0.45 | 0.49 | A>A |
| chr8 | 1.18E+08 | 117900000 | 0.34 | 0.26 | A>A |
| chr8 | 1.18E+08 | 118000000 | 0.04 | -0.09 | B>B |
| chr8 | 1.18E+08 | 118100000 | -0.30 | -0.19 | B>B |
| chr8 | 1.18E+08 | 118200000 | -0.57 | -0.57 | B>B |
| chr8 | 1.18E+08 | 118300000 | -0.46 | -0.44 | B>B |
| chr8 | 1.18E+08 | 118400000 | -0.19 | -0.08 | B>B |
| chr8 | 1.18E+08 | 118500000 | -0.06 | 0.07 | A>A |
| chr8 | 1.19E+08 | 118600000 | 0.14 | 0.28 | A>A |
| chr8 | 1.19E+08 | 118700000 | 0.26 | 0.42 | A>A |
| chr8 | 1.19E+08 | 118800000 | 0.35 | 0.41 | A>A |
| chr8 | 1.19E+08 | 118900000 | 0.50 | 0.47 | A>A |
| chr8 | 1.19E+08 | 119000000 | 0.77 | 0.64 | A>A |
| chr8 | 1.19E+08 | 119100000 | 0.79 | 0.69 | A>A |
| chr8 | 1.19E+08 | 119200000 | 0.47 | 0.45 | A>A |
| chr8 | 1.19E+08 | 119300000 | 0.52 | 0.43 | A>A |
| chr8 | 1.19E+08 | 119400000 | 0.57 | 0.45 | A>A |
| chr8 | 1.19E+08 | 119500000 | 0.26 | 0.37 | A>A |
| chr8 | 1.2E+08 | 119600000 | -0.03 | 0.10 | A>A |
| chr8 | 1.2E+08 | 119700000 | -0.20 | -0.06 | B>B |
| chr8 | 1.2E+08 | 119800000 | -0.28 | 0.14 | B>B |
| chr8 | 1.2E+08 | 119900000 | -0.08 | 0.41 | A>A |
| chr8 | 1.2E+08 | 120000000 | -0.07 | 0.42 | A>A |
| chr8 | 1.2E+08 | 120100000 | -0.37 | 0.19 | B>A |
| chr8 | 1.2E+08 | 120200000 | -0.58 | -0.27 | B>B |
| chr8 | 1.2E+08 | 120300000 | -0.53 | -0.27 | B>B |
| chr8 | 1.2E+08 | 120400000 | -0.61 | -0.44 | B>B |
| chr8 | 1.2E+08 | 120500000 | -0.32 | -0.22 | B>B |
| chr8 | 1.21E+08 | 120600000 | -0.18 | -0.21 | B>B |
| chr8 | 1.21E+08 | 120700000 | -0.02 | -0.17 | B>B |
| chr8 | 1.21E+08 | 120800000 | 0.23 | 0.14 | A>A |
| chr8 | 1.21E+08 | 120900000 | 0.54 | 0.39 | A>A |
| chr8 | 1.21E+08 | 121000000 | 0.64 | 0.62 | A>A |
| chr8 | 1.21E+08 | 121100000 | 0.68 | 0.56 | A>A |
| chr8 | 1.21E+08 | 121200000 | 0.13 | 0.01 | A>A |
| chr8 | 1.21E+08 | 121300000 | 0.01 | -0.19 | B>B |
| chr8 | 1.21E+08 | 121400000 | 0.01 | -0.13 | B>B |
| chr8 | 1.21E+08 | 121500000 | 0.31 | 0.25 | A>A |
| chr8 | 1.22E+08 | 121600000 | 0.17 | 0.26 | A>A |
| chr8 | 1.22E+08 | 121700000 | -0.10 | 0.13 | A>A |
| chr8 | 1.22E+08 | 121800000 | 0.05 | 0.27 | A>A |
| chr8 | 1.22E+08 | 121900000 | 0.17 | 0.27 | A>A |
| chr8 | 1.22E+08 | 122000000 | 0.23 | 0.35 | A>A |
| chr8 | 1.22E+08 | 122100000 | 0.29 | 0.69 | A>A |
| chr8 | 1.22E+08 | 122200000 | 0.38 | 0.81 | A>A |
| chr8 | 1.22E+08 | 122300000 | 0.33 | 0.60 | A>A |
| chr8 | 1.22E+08 | 122400000 | 0.49 | 0.67 | A>A |
| chr8 | 1.22E+08 | 122500000 | 0.41 | 0.69 | A>A |
| chr8 | 1.23E+08 | 122600000 | 0.50 | 0.62 | A>A |
| chr8 | 1.23E+08 | 122700000 | 0.51 | 0.60 | A>A |
| chr8 | 1.23E+08 | 122800000 | 0.72 | 0.85 | A>A |

| | | | | | |
|---|---|---|---|---|---|
| chr8 | 1.23E+08 | 122900000 | 0.64 | 0.64 | A>A |
| chr8 | 1.23E+08 | 123000000 | 0.58 | 0.67 | A>A |
| chr8 | 1.23E+08 | 123100000 | 0.54 | 0.68 | A>A |
| chr8 | 1.23E+08 | 123200000 | 0.47 | 0.59 | A>A |
| chr8 | 1.23E+08 | 123300000 | 0.37 | 0.51 | A>A |
| chr8 | 1.23E+08 | 123400000 | 0.19 | 0.47 | A>A |
| chr8 | 1.23E+08 | 123500000 | 0.27 | 0.40 | A>A |
| chr8 | 1.24E+08 | 123600000 | 0.29 | 0.51 | A>A |
| chr8 | 1.24E+08 | 123700000 | 0.47 | 0.52 | A>A |
| chr8 | 1.24E+08 | 123800000 | 0.70 | 0.79 | A>A |
| chr8 | 1.24E+08 | 123900000 | 0.74 | 0.83 | A>A |
| chr8 | 1.24E+08 | 124000000 | 0.71 | 0.80 | A>A |
| chr8 | 1.24E+08 | 124100000 | 0.82 | 0.82 | A>A |
| chr8 | 1.24E+08 | 124200000 | 0.92 | 0.90 | A>A |
| chr8 | 1.24E+08 | 124300000 | 1.00 | 0.96 | A>A |
| chr8 | 1.24E+08 | 124400000 | 1.05 | 0.98 | A>A |
| chr8 | 1.24E+08 | 124500000 | 1.03 | 0.84 | A>A |
| chr8 | 1.25E+08 | 124600000 | 1.01 | 1.05 | A>A |
| chr8 | 1.25E+08 | 124700000 | 1.03 | 0.93 | A>A |
| chr8 | 1.25E+08 | 124800000 | 0.94 | 0.89 | A>A |
| chr8 | 1.25E+08 | 124900000 | 0.87 | 0.83 | A>A |
| chr8 | 1.25E+08 | 125000000 | 1.01 | 0.94 | A>A |
| chr8 | 1.25E+08 | 125100000 | 0.99 | 0.89 | A>A |
| chr8 | 1.25E+08 | 125200000 | 1.03 | 0.97 | A>A |
| chr8 | 1.25E+08 | 125300000 | 1.10 | 0.92 | A>A |
| chr8 | 1.25E+08 | 125400000 | 1.00 | 0.95 | A>A |
| chr8 | 1.25E+08 | 125500000 | 0.77 | 0.79 | A>A |
| chr8 | 1.26E+08 | 125600000 | 0.77 | 0.88 | A>A |
| chr8 | 1.26E+08 | 125700000 | 0.54 | 0.68 | A>A |
| chr8 | 1.26E+08 | 125800000 | 0.40 | 0.73 | A>A |
| chr8 | 1.26E+08 | 125900000 | 0.28 | 0.62 | A>A |
| chr8 | 1.26E+08 | 126000000 | 0.22 | 0.46 | A>A |
| chr8 | 1.26E+08 | 126100000 | 0.52 | 0.54 | A>A |
| chr8 | 1.26E+08 | 126200000 | 0.51 | 0.53 | A>A |
| chr8 | 1.26E+08 | 126300000 | 0.64 | 0.59 | A>A |
| chr8 | 1.26E+08 | 126400000 | 0.64 | 0.64 | A>A |
| chr8 | 1.26E+08 | 126500000 | 0.64 | 0.53 | A>A |
| chr8 | 1.27E+08 | 126600000 | 0.74 | 0.64 | A>A |
| chr8 | 1.27E+08 | 126700000 | 0.52 | 0.44 | A>A |
| chr8 | 1.27E+08 | 126800000 | 0.04 | -0.16 | B>B |
| chr8 | 1.27E+08 | 126900000 | -0.21 | -0.52 | B>B |
| chr8 | 1.27E+08 | 127000000 | -0.38 | -0.62 | B>B |
| chr8 | 1.27E+08 | 127100000 | -0.48 | -0.67 | B>B |
| chr8 | 1.27E+08 | 127200000 | -0.58 | -0.67 | B>B |
| chr8 | 1.27E+08 | 127300000 | -0.47 | -0.38 | B>B |
| chr8 | 1.27E+08 | 127400000 | -0.24 | 0.17 | B>B |
| chr8 | 1.27E+08 | 127500000 | -0.03 | 0.27 | A>A |
| chr8 | 1.28E+08 | 127600000 | 0.05 | 0.27 | A>A |
| chr8 | 1.28E+08 | 127700000 | -0.09 | 0.06 | B>B |
| chr8 | 1.28E+08 | 127800000 | -0.12 | -0.06 | B>B |
| chr8 | 1.28E+08 | 127900000 | 0.12 | 0.11 | A>A |
| chr8 | 1.28E+08 | 128000000 | 0.15 | 0.14 | A>A |
| chr8 | 1.28E+08 | 128100000 | 0.16 | 0.18 | A>A |
| chr8 | 1.28E+08 | 128200000 | 0.28 | 0.32 | A>A |
| chr8 | 1.28E+08 | 128300000 | 0.44 | 0.55 | A>A |
| chr8 | 1.28E+08 | 128400000 | 0.55 | 0.52 | A>A |

| | | | | | |
|---|---|---|---|---|---|
| chr8 | 1.28E+08 | 128500000 | 0.44 | 0.59 | A>A |
| chr8 | 1.29E+08 | 128600000 | 0.48 | 0.55 | A>A |
| chr8 | 1.29E+08 | 128700000 | 0.48 | 0.59 | A>A |
| chr8 | 1.29E+08 | 128800000 | 0.51 | 0.48 | A>A |
| chr8 | 1.29E+08 | 128900000 | 0.67 | 0.55 | A>A |
| chr8 | 1.29E+08 | 129000000 | 0.55 | 0.53 | A>A |
| chr8 | 1.29E+08 | 129100000 | 0.49 | 0.47 | A>A |
| chr8 | 1.29E+08 | 129200000 | 0.53 | 0.51 | A>A |
| chr8 | 1.29E+08 | 129300000 | 0.59 | 0.52 | A>A |
| chr8 | 1.29E+08 | 129400000 | 0.68 | 0.48 | A>A |
| chr8 | 1.29E+08 | 129500000 | 0.65 | 0.50 | A>A |
| chr8 | 1.3E+08 | 129600000 | 0.59 | 0.52 | A>A |
| chr8 | 1.3E+08 | 129700000 | 0.53 | 0.36 | A>A |
| chr8 | 1.3E+08 | 129800000 | 0.42 | 0.05 | A>A |
| chr8 | 1.3E+08 | 129900000 | 0.45 | 0.05 | A>A |
| chr8 | 1.3E+08 | 130000000 | 0.38 | -0.17 | A>A |
| chr8 | 1.3E+08 | 130100000 | 0.32 | -0.20 | A>A |
| chr8 | 1.3E+08 | 130200000 | 0.39 | -0.24 | A>B |
| chr8 | 1.3E+08 | 130300000 | 0.30 | -0.21 | A>A |
| chr8 | 1.3E+08 | 130400000 | 0.29 | -0.23 | A>A |
| chr8 | 1.3E+08 | 130500000 | 0.28 | -0.25 | A>A |
| chr8 | 1.31E+08 | 130600000 | 0.31 | -0.08 | A>A |
| chr8 | 1.31E+08 | 130700000 | 0.33 | -0.04 | A>A |
| chr8 | 1.31E+08 | 130800000 | 0.40 | 0.20 | A>A |
| chr8 | 1.31E+08 | 130900000 | 0.50 | 0.52 | A>A |
| chr8 | 1.31E+08 | 131000000 | 0.67 | 0.75 | A>A |
| chr8 | 1.31E+08 | 131100000 | 0.65 | 0.73 | A>A |
| chr8 | 1.31E+08 | 131200000 | 0.54 | 0.58 | A>A |
| chr8 | 1.31E+08 | 131300000 | 0.53 | 0.68 | A>A |
| chr8 | 1.31E+08 | 131400000 | 0.57 | 0.69 | A>A |
| chr8 | 1.31E+08 | 131500000 | 0.69 | 0.71 | A>A |
| chr8 | 1.32E+08 | 131600000 | 0.64 | 0.70 | A>A |
| chr8 | 1.32E+08 | 131700000 | 0.55 | 0.61 | A>A |
| chr8 | 1.32E+08 | 131800000 | 0.38 | 0.28 | A>A |
| chr8 | 1.32E+08 | 131900000 | 0.31 | -0.03 | A>A |
| chr8 | 1.32E+08 | 132000000 | -0.02 | -0.61 | A>B |
| chr8 | 1.32E+08 | 132100000 | -0.15 | -0.52 | B>B |
| chr8 | 1.32E+08 | 132200000 | -0.40 | -0.76 | B>B |
| chr8 | 1.32E+08 | 132300000 | -0.45 | -0.68 | B>B |
| chr8 | 1.32E+08 | 132400000 | -0.53 | -0.74 | B>B |
| chr8 | 1.32E+08 | 132500000 | -0.53 | -0.72 | B>B |
| chr8 | 1.33E+08 | 132600000 | -0.53 | -0.71 | B>B |
| chr8 | 1.33E+08 | 132700000 | -0.53 | -0.71 | B>B |
| chr8 | 1.33E+08 | 132800000 | -0.34 | -0.59 | B>B |
| chr8 | 1.33E+08 | 132900000 | -0.03 | -0.17 | B>B |
| chr8 | 1.33E+08 | 133000000 | 0.17 | 0.06 | A>A |
| chr8 | 1.33E+08 | 133100000 | 0.01 | -0.30 | B>B |
| chr8 | 1.33E+08 | 133200000 | -0.10 | -0.49 | B>B |
| chr8 | 1.33E+08 | 133300000 | -0.26 | -0.62 | B>B |
| chr8 | 1.33E+08 | 133400000 | -0.17 | -0.56 | B>B |
| chr8 | 1.33E+08 | 133500000 | -0.09 | -0.60 | B>B |
| chr8 | 1.34E+08 | 133600000 | -0.23 | -0.51 | B>B |
| chr8 | 1.34E+08 | 133700000 | -0.14 | -0.31 | B>B |
| chr8 | 1.34E+08 | 133800000 | 0.16 | -0.12 | A>A |
| chr8 | 1.34E+08 | 133900000 | 0.36 | 0.19 | A>A |
| chr8 | 1.34E+08 | 134000000 | 0.56 | 0.19 | A>A |

| | | | | | |
|---|---|---|---|---|---|
| chr8 | 1.34E+08 | 134100000 | 0.68 | 0.28 | A>A |
| chr8 | 1.34E+08 | 134200000 | 0.68 | 0.27 | A>A |
| chr8 | 1.34E+08 | 134300000 | 0.61 | 0.51 | A>A |
| chr8 | 1.34E+08 | 134400000 | 0.80 | 0.73 | A>A |
| chr8 | 1.34E+08 | 134500000 | 0.49 | 0.51 | A>A |
| chr8 | 1.35E+08 | 134600000 | 0.28 | 0.12 | A>A |
| chr8 | 1.35E+08 | 134700000 | 0.22 | -0.03 | A>A |
| chr8 | 1.35E+08 | 134800000 | 0.04 | -0.28 | B>B |
| chr8 | 1.35E+08 | 134900000 | -0.09 | -0.39 | B>B |
| chr8 | 1.35E+08 | 135000000 | -0.35 | -0.51 | B>B |
| chr8 | 1.35E+08 | 135100000 | -0.58 | -0.68 | B>B |
| chr8 | 1.35E+08 | 135200000 | -0.71 | -0.65 | B>B |
| chr8 | 1.35E+08 | 135300000 | -0.62 | -0.71 | B>B |
| chr8 | 1.35E+08 | 135400000 | -0.49 | -0.64 | B>B |
| chr8 | 1.35E+08 | 135500000 | -0.19 | -0.39 | B>B |
| chr8 | 1.36E+08 | 135600000 | 0.17 | 0.20 | A>A |
| chr8 | 1.36E+08 | 135700000 | 0.17 | 0.18 | A>A |
| chr8 | 1.36E+08 | 135800000 | 0.23 | 0.34 | A>A |
| chr8 | 1.36E+08 | 135900000 | 0.16 | 0.14 | A>A |
| chr8 | 1.36E+08 | 136000000 | -0.28 | -0.46 | B>B |
| chr8 | 1.36E+08 | 136100000 | -0.58 | -0.74 | B>B |
| chr8 | 1.36E+08 | 136200000 | -0.68 | -0.80 | B>B |
| chr8 | 1.36E+08 | 136300000 | -0.71 | -0.71 | B>B |
| chr8 | 1.36E+08 | 136400000 | -0.66 | -0.75 | B>B |
| chr8 | 1.36E+08 | 136500000 | -0.37 | -0.61 | B>B |
| chr8 | 1.37E+08 | 136600000 | -0.62 | -0.79 | B>B |
| chr8 | 1.37E+08 | 136700000 | -0.59 | -0.85 | B>B |
| chr8 | 1.37E+08 | 136800000 | -0.50 | -0.75 | B>B |
| chr8 | 1.37E+08 | 136900000 | -0.65 | -0.83 | B>B |
| chr8 | 1.37E+08 | 137000000 | -0.80 | -0.84 | B>B |
| chr8 | 1.37E+08 | 137100000 | -0.80 | -0.90 | B>B |
| chr8 | 1.37E+08 | 137200000 | -0.85 | -0.91 | B>B |
| chr8 | 1.37E+08 | 137300000 | -0.84 | -0.97 | B>B |
| chr8 | 1.37E+08 | 137400000 | -0.89 | -1.00 | B>B |
| chr8 | 1.37E+08 | 137500000 | -0.84 | -0.96 | B>B |
| chr8 | 1.38E+08 | 137600000 | -0.87 | -0.95 | B>B |
| chr8 | 1.38E+08 | 137700000 | -0.82 | -0.89 | B>B |
| chr8 | 1.38E+08 | 137800000 | -0.72 | -0.89 | B>B |
| chr8 | 1.38E+08 | 137900000 | -0.64 | -0.95 | B>B |
| chr8 | 1.38E+08 | 138000000 | -0.51 | -0.95 | B>B |
| chr8 | 1.38E+08 | 138100000 | -0.27 | -0.91 | A>B |
| chr8 | 1.38E+08 | 138200000 | -0.28 | -0.88 | A>B |
| chr8 | 1.38E+08 | 138300000 | -0.34 | -0.92 | B>B |
| chr8 | 1.38E+08 | 138400000 | -0.45 | -0.89 | B>B |
| chr8 | 1.38E+08 | 138500000 | -0.69 | -0.88 | B>B |
| chr8 | 1.39E+08 | 138600000 | -0.81 | -0.97 | B>B |
| chr8 | 1.39E+08 | 138700000 | -1.01 | -1.02 | B>B |
| chr8 | 1.39E+08 | 138800000 | -0.99 | -1.05 | B>B |
| chr8 | 1.39E+08 | 138900000 | -1.05 | -0.92 | B>B |
| chr8 | 1.39E+08 | 139000000 | -0.98 | -0.99 | B>B |
| chr8 | 1.39E+08 | 139100000 | -0.99 | -0.96 | B>B |
| chr8 | 1.39E+08 | 139200000 | -0.74 | -0.90 | B>B |
| chr8 | 1.39E+08 | 139300000 | -0.85 | -0.92 | B>B |
| chr8 | 1.39E+08 | 139400000 | -0.89 | -0.97 | B>B |
| chr8 | 1.39E+08 | 139500000 | -0.91 | -0.99 | B>B |
| chr8 | 1.4E+08 | 139600000 | -0.81 | -0.93 | B>B |

| | | | | | |
|---|---|---|---|---|---|
| chr8 | 1.4E+08 | 139700000 | -0.66 | -0.81 | B>B |
| chr8 | 1.4E+08 | 139800000 | -0.19 | -0.48 | B>B |
| chr8 | 1.4E+08 | 139900000 | 0.04 | -0.72 | A>B |
| chr8 | 1.4E+08 | 140000000 | -0.07 | -0.76 | A>B |
| chr8 | 1.4E+08 | 140100000 | -0.27 | -0.54 | B>B |
| chr8 | 1.4E+08 | 140200000 | -0.28 | -0.49 | B>B |
| chr8 | 1.4E+08 | 140300000 | -0.02 | -0.16 | B>B |
| chr8 | 1.4E+08 | 140400000 | -0.11 | -0.42 | B>B |
| chr8 | 1.4E+08 | 140500000 | -0.09 | -0.62 | B>B |
| chr8 | 1.41E+08 | 140600000 | -0.09 | -0.65 | B>B |
| chr8 | 1.41E+08 | 140700000 | 0.14 | -0.13 | A>A |
| chr8 | 1.41E+08 | 140800000 | 0.43 | 0.40 | A>A |
| chr8 | 1.41E+08 | 140900000 | 0.49 | 0.62 | A>A |
| chr8 | 1.41E+08 | 141000000 | 0.50 | 0.53 | A>A |
| chr8 | 1.41E+08 | 141100000 | 0.42 | 0.53 | A>A |
| chr8 | 1.41E+08 | 141200000 | 0.52 | 0.55 | A>A |
| chr8 | 1.41E+08 | 141300000 | 0.76 | 0.72 | A>A |
| chr8 | 1.41E+08 | 141400000 | 0.84 | 0.70 | A>A |
| chr8 | 1.41E+08 | 141500000 | 0.91 | 0.88 | A>A |
| chr8 | 1.42E+08 | 141600000 | 1.18 | 1.17 | A>A |
| chr8 | 1.42E+08 | 141700000 | 0.97 | 1.13 | A>A |
| chr8 | 1.42E+08 | 141800000 | 1.05 | 1.12 | A>A |
| chr8 | 1.42E+08 | 141900000 | 1.04 | 1.09 | A>A |
| chr8 | 1.42E+08 | 142000000 | 1.12 | 1.17 | A>A |
| chr8 | 1.42E+08 | 142100000 | 1.23 | 1.19 | A>A |
| chr8 | 1.42E+08 | 142200000 | 1.07 | 1.07 | A>A |
| chr8 | 1.42E+08 | 142300000 | 1.23 | 1.15 | A>A |
| chr8 | 1.42E+08 | 142400000 | 1.08 | 1.03 | A>A |
| chr8 | 1.42E+08 | 142500000 | 0.90 | 0.93 | A>A |
| chr8 | 1.43E+08 | 142600000 | 0.82 | 0.86 | A>A |
| chr8 | 1.43E+08 | 142700000 | 0.82 | 0.73 | A>A |
| chr8 | 1.43E+08 | 142800000 | 0.00 | 0.00 | NO |
| chr8 | 1.43E+08 | 142900000 | 0.63 | 0.68 | A>A |
| chr8 | 1.43E+08 | 143000000 | 0.71 | 0.75 | A>A |
| chr8 | 1.43E+08 | 143100000 | 0.85 | 0.79 | A>A |
| chr8 | 1.43E+08 | 143200000 | 0.89 | 0.85 | A>A |
| chr8 | 1.43E+08 | 143300000 | 0.81 | 0.81 | A>A |
| chr8 | 1.43E+08 | 143400000 | 0.75 | 0.72 | A>A |
| chr8 | 1.43E+08 | 143500000 | 0.80 | 0.82 | A>A |
| chr8 | 1.44E+08 | 143600000 | 0.78 | 0.75 | A>A |
| chr8 | 1.44E+08 | 143700000 | 0.89 | 0.86 | A>A |
| chr8 | 1.44E+08 | 143800000 | 0.92 | 0.91 | A>A |
| chr8 | 1.44E+08 | 143900000 | 0.94 | 0.87 | A>A |
| chr8 | 1.44E+08 | 144000000 | 0.79 | 0.82 | A>A |
| chr8 | 1.44E+08 | 144100000 | 0.87 | 0.85 | A>A |
| chr8 | 1.44E+08 | 144200000 | 0.79 | 0.78 | A>A |
| chr8 | 1.44E+08 | 144300000 | 0.82 | 0.85 | A>A |
| chr8 | 1.44E+08 | 144400000 | 0.92 | 0.84 | A>A |
| chr8 | 1.44E+08 | 144500000 | 0.87 | 0.75 | A>A |
| chr8 | 1.45E+08 | 144600000 | 0.94 | 0.75 | A>A |
| chr8 | 1.45E+08 | 144700000 | 0.86 | 0.78 | A>A |
| chr8 | 1.45E+08 | 144800000 | 0.80 | 0.73 | A>A |
| chr8 | 1.45E+08 | 144900000 | 0.77 | 0.65 | A>A |
| chr8 | 1.45E+08 | 145000000 | 0.82 | 0.64 | A>A |
| chr8 | 1.45E+08 | 145100000 | 0.89 | 0.75 | A>A |
| chr8 | 1.45E+08 | 145200000 | 0.77 | 0.69 | A>A |

| | | | | | |
|---|---|---|---|---|---|
| chr8 | 1.45E+08 | 145300000 | 0.76 | 0.73 | A>A |
| chr8 | 1.45E+08 | 145400000 | 0.00 | 0.00 | NO |
| chr8 | 1.45E+08 | 145500000 | 0.00 | 0.00 | NO |
| chr8 | 1.46E+08 | 145600000 | 0.72 | 0.64 | A>A |
| chr8 | 1.46E+08 | 145700000 | 0.81 | 0.71 | A>A |
| chr8 | 1.46E+08 | 145800000 | 0.66 | 0.69 | A>A |
| chr8 | 1.46E+08 | 145900000 | 0.72 | 0.77 | A>A |
| chr8 | 1.46E+08 | 146000000 | 0.74 | 0.79 | A>A |
| chr8 | 1.46E+08 | 146100000 | 0.74 | 0.81 | A>A |
| chr8 | 1.46E+08 | 146200000 | 0.67 | 0.86 | A>A |
| chr8 | 1.46E+08 | 146300000 | 0.65 | 0.66 | A>A |
| chr8 | 1.46E+08 | 146364022 | 0.00 | 0.00 | NO |
| chr9 | 0 | 100000 | 0.00 | 0.00 | NO |
| chr9 | 100000 | 200000 | 0.00 | 0.00 | NO |
| chr9 | 200000 | 300000 | 0.00 | -0.06 | B>B |
| chr9 | 300000 | 400000 | 0.21 | 0.24 | A>A |
| chr9 | 400000 | 500000 | 0.99 | 1.04 | A>A |
| chr9 | 500000 | 600000 | 0.89 | 0.81 | A>A |
| chr9 | 600000 | 700000 | 0.64 | 0.73 | A>A |
| chr9 | 700000 | 800000 | 0.53 | 0.59 | A>A |
| chr9 | 800000 | 900000 | -0.07 | -0.13 | B>B |
| chr9 | 900000 | 1000000 | -0.39 | -0.41 | B>B |
| chr9 | 1000000 | 1100000 | -0.85 | -0.93 | B>B |
| chr9 | 1100000 | 1200000 | -1.39 | -1.44 | B>B |
| chr9 | 1200000 | 1300000 | -1.54 | -1.48 | B>B |
| chr9 | 1300000 | 1400000 | -1.52 | -1.43 | B>B |
| chr9 | 1400000 | 1500000 | -1.57 | -1.42 | B>B |
| chr9 | 1500000 | 1600000 | -1.54 | -1.42 | B>B |
| chr9 | 1600000 | 1700000 | -1.36 | -1.33 | B>B |
| chr9 | 1700000 | 1800000 | -1.26 | -1.37 | B>B |
| chr9 | 1800000 | 1900000 | -0.99 | -1.30 | B>B |
| chr9 | 1900000 | 2000000 | -0.70 | -1.11 | B>B |
| chr9 | 2000000 | 2100000 | -0.24 | -0.46 | B>B |
| chr9 | 2100000 | 2200000 | -0.18 | -0.29 | B>B |
| chr9 | 2200000 | 2300000 | -0.21 | -0.45 | B>B |
| chr9 | 2300000 | 2400000 | -0.54 | -0.85 | B>B |
| chr9 | 2400000 | 2500000 | -0.46 | -0.69 | B>B |
| chr9 | 2500000 | 2600000 | -0.35 | -0.70 | B>B |
| chr9 | 2600000 | 2700000 | -0.02 | -0.39 | B>B |
| chr9 | 2700000 | 2800000 | 0.10 | -0.20 | B>B |
| chr9 | 2800000 | 2900000 | -0.06 | -0.36 | B>B |
| chr9 | 2900000 | 3000000 | -0.72 | -1.16 | B>B |
| chr9 | 3000000 | 3100000 | -0.73 | -1.18 | B>B |
| chr9 | 3100000 | 3200000 | -0.68 | -0.98 | B>B |
| chr9 | 3200000 | 3300000 | -0.33 | -0.48 | B>B |
| chr9 | 3300000 | 3400000 | -0.02 | -0.25 | B>B |
| chr9 | 3400000 | 3500000 | 0.19 | 0.03 | A>A |
| chr9 | 3500000 | 3600000 | 0.22 | -0.06 | A>A |
| chr9 | 3600000 | 3700000 | 0.03 | -0.07 | B>B |
| chr9 | 3700000 | 3800000 | 0.43 | 0.37 | A>A |
| chr9 | 3800000 | 3900000 | 0.68 | 0.60 | A>A |
| chr9 | 3900000 | 4000000 | 0.80 | 0.52 | A>A |
| chr9 | 4000000 | 4100000 | 0.75 | 0.54 | A>A |
| chr9 | 4100000 | 4200000 | 0.76 | 0.64 | A>A |
| chr9 | 4200000 | 4300000 | 0.75 | 0.72 | A>A |
| chr9 | 4300000 | 4400000 | 0.35 | 0.33 | A>A |

| | | | | | |
|---|---|---|---|---|---|
| chr9 | 4400000 | 4500000 | 0.44 | 0.36 | A>A |
| chr9 | 4500000 | 4600000 | 0.67 | 0.53 | A>A |
| chr9 | 4600000 | 4700000 | 0.85 | 0.76 | A>A |
| chr9 | 4700000 | 4800000 | 0.93 | 0.78 | A>A |
| chr9 | 4800000 | 4900000 | 1.14 | 0.99 | A>A |
| chr9 | 4900000 | 5000000 | 0.94 | 0.91 | A>A |
| chr9 | 5000000 | 5100000 | 0.71 | 0.74 | A>A |
| chr9 | 5100000 | 5200000 | 0.58 | 0.65 | A>A |
| chr9 | 5200000 | 5300000 | 0.89 | 0.75 | A>A |
| chr9 | 5300000 | 5400000 | 0.88 | 0.78 | A>A |
| chr9 | 5400000 | 5500000 | 0.76 | 0.77 | A>A |
| chr9 | 5500000 | 5600000 | 0.85 | 0.92 | A>A |
| chr9 | 5600000 | 5700000 | 1.00 | 1.09 | A>A |
| chr9 | 5700000 | 5800000 | 1.05 | 1.15 | A>A |
| chr9 | 5800000 | 5900000 | 1.37 | 1.41 | A>A |
| chr9 | 5900000 | 6000000 | 1.29 | 1.45 | A>A |
| chr9 | 6000000 | 6100000 | 0.94 | 1.12 | A>A |
| chr9 | 6100000 | 6200000 | 0.94 | 1.24 | A>A |
| chr9 | 6200000 | 6300000 | 0.89 | 1.27 | A>A |
| chr9 | 6300000 | 6400000 | 0.78 | 0.88 | A>A |
| chr9 | 6400000 | 6500000 | 1.27 | 1.36 | A>A |
| chr9 | 6500000 | 6600000 | 1.13 | 0.97 | A>A |
| chr9 | 6600000 | 6700000 | 1.16 | 1.06 | A>A |
| chr9 | 6700000 | 6800000 | 0.96 | 0.74 | A>A |
| chr9 | 6800000 | 6900000 | 0.51 | 0.22 | A>A |
| chr9 | 6900000 | 7000000 | 0.21 | -0.06 | A>A |
| chr9 | 7000000 | 7100000 | 0.02 | -0.27 | B>B |
| chr9 | 7100000 | 7200000 | 0.03 | -0.28 | B>B |
| chr9 | 7200000 | 7300000 | -0.30 | -0.73 | B>B |
| chr9 | 7300000 | 7400000 | -0.51 | -0.95 | B>B |
| chr9 | 7400000 | 7500000 | -0.69 | -1.03 | B>B |
| chr9 | 7500000 | 7600000 | -0.72 | -1.04 | B>B |
| chr9 | 7600000 | 7700000 | -0.88 | -1.01 | B>B |
| chr9 | 7700000 | 7800000 | -0.88 | -0.96 | B>B |
| chr9 | 7800000 | 7900000 | -0.93 | -0.97 | B>B |
| chr9 | 7900000 | 8000000 | -1.18 | -1.01 | B>B |
| chr9 | 8000000 | 8100000 | -1.12 | -0.97 | B>B |
| chr9 | 8100000 | 8200000 | -1.04 | -0.96 | B>B |
| chr9 | 8200000 | 8300000 | -1.05 | -0.98 | B>B |
| chr9 | 8300000 | 8400000 | -1.07 | -0.97 | B>B |
| chr9 | 8400000 | 8500000 | -1.12 | -0.94 | B>B |
| chr9 | 8500000 | 8600000 | -1.03 | -0.91 | B>B |
| chr9 | 8600000 | 8700000 | -1.06 | -0.84 | B>B |
| chr9 | 8700000 | 8800000 | -0.99 | -0.81 | B>B |
| chr9 | 8800000 | 8900000 | -0.99 | -0.81 | B>B |
| chr9 | 8900000 | 9000000 | -0.96 | -0.85 | B>B |
| chr9 | 9000000 | 9100000 | -1.03 | -0.92 | B>B |
| chr9 | 9100000 | 9200000 | -1.03 | -0.98 | B>B |
| chr9 | 9200000 | 9300000 | -1.04 | -1.03 | B>B |
| chr9 | 9300000 | 9400000 | -1.10 | -0.99 | B>B |
| chr9 | 9400000 | 9500000 | -1.12 | -1.00 | B>B |
| chr9 | 9500000 | 9600000 | -1.09 | -1.07 | B>B |
| chr9 | 9600000 | 9700000 | -1.06 | -1.06 | B>B |
| chr9 | 9700000 | 9800000 | -1.04 | -1.01 | B>B |
| chr9 | 9800000 | 9900000 | -1.09 | -0.93 | B>B |
| chr9 | 9900000 | 10000000 | -0.99 | -0.99 | B>B |

| | | | | | |
|---|---|---|---|---|---|
| chr9 | 10000000 | 10100000 | -1.03 | -0.97 | B>B |
| chr9 | 10100000 | 10200000 | -1.11 | -1.01 | B>B |
| chr9 | 10200000 | 10300000 | -1.05 | -1.05 | B>B |
| chr9 | 10300000 | 10400000 | -1.03 | -1.01 | B>B |
| chr9 | 10400000 | 10500000 | -1.04 | -1.05 | B>B |
| chr9 | 10500000 | 10600000 | -0.99 | -1.01 | B>B |
| chr9 | 10600000 | 10700000 | -0.97 | -0.99 | B>B |
| chr9 | 10700000 | 10800000 | -1.06 | -1.01 | B>B |
| chr9 | 10800000 | 10900000 | -0.95 | -0.94 | B>B |
| chr9 | 10900000 | 11000000 | -0.96 | -0.99 | B>B |
| chr9 | 11000000 | 11100000 | -0.95 | -1.00 | B>B |
| chr9 | 11100000 | 11200000 | -0.99 | -0.93 | B>B |
| chr9 | 11200000 | 11300000 | -0.97 | -0.97 | B>B |
| chr9 | 11300000 | 11400000 | -0.95 | -0.95 | B>B |
| chr9 | 11400000 | 11500000 | -0.99 | -0.98 | B>B |
| chr9 | 11500000 | 11600000 | -0.98 | -0.94 | B>B |
| chr9 | 11600000 | 11700000 | -0.99 | -0.95 | B>B |
| chr9 | 11700000 | 11800000 | -1.02 | -0.95 | B>B |
| chr9 | 11800000 | 11900000 | -1.00 | -1.00 | B>B |
| chr9 | 11900000 | 12000000 | -1.09 | -0.99 | B>B |
| chr9 | 12000000 | 12100000 | -1.04 | -1.02 | B>B |
| chr9 | 12100000 | 12200000 | -1.10 | -1.03 | B>B |
| chr9 | 12200000 | 12300000 | -1.03 | -0.98 | B>B |
| chr9 | 12300000 | 12400000 | -1.08 | -1.07 | B>B |
| chr9 | 12400000 | 12500000 | -1.06 | -0.98 | B>B |
| chr9 | 12500000 | 12600000 | -1.05 | -1.05 | B>B |
| chr9 | 12600000 | 12700000 | -0.74 | -0.88 | B>B |
| chr9 | 12700000 | 12800000 | 0.15 | -0.12 | A>A |
| chr9 | 12800000 | 12900000 | 0.29 | -0.06 | A>A |
| chr9 | 12900000 | 13000000 | -0.15 | -0.54 | B>B |
| chr9 | 13000000 | 13100000 | -0.19 | -0.44 | B>B |
| chr9 | 13100000 | 13200000 | 0.08 | -0.19 | B>B |
| chr9 | 13200000 | 13300000 | 0.17 | -0.13 | A>A |
| chr9 | 13300000 | 13400000 | -0.29 | -0.51 | B>B |
| chr9 | 13400000 | 13500000 | -0.43 | -0.51 | B>B |
| chr9 | 13500000 | 13600000 | -0.51 | -0.75 | B>B |
| chr9 | 13600000 | 13700000 | -0.58 | -0.75 | B>B |
| chr9 | 13700000 | 13800000 | -0.42 | -0.71 | B>B |
| chr9 | 13800000 | 13900000 | -0.40 | -0.67 | B>B |
| chr9 | 13900000 | 14000000 | -0.16 | -0.58 | B>B |
| chr9 | 14000000 | 14100000 | 0.13 | -0.15 | B>B |
| chr9 | 14100000 | 14200000 | 0.46 | 0.23 | A>A |
| chr9 | 14200000 | 14300000 | 0.54 | 0.22 | A>A |
| chr9 | 14300000 | 14400000 | 0.09 | -0.06 | A>A |
| chr9 | 14400000 | 14500000 | -0.33 | -0.41 | B>B |
| chr9 | 14500000 | 14600000 | -0.42 | -0.50 | B>B |
| chr9 | 14600000 | 14700000 | -0.20 | -0.26 | B>B |
| chr9 | 14700000 | 14800000 | -0.54 | -0.61 | B>B |
| chr9 | 14800000 | 14900000 | -0.73 | -0.81 | B>B |
| chr9 | 14900000 | 15000000 | -0.75 | -0.81 | B>B |
| chr9 | 15000000 | 15100000 | -0.63 | -0.77 | B>B |
| chr9 | 15100000 | 15200000 | -0.22 | -0.44 | B>B |
| chr9 | 15200000 | 15300000 | 0.20 | 0.07 | A>A |
| chr9 | 15300000 | 15400000 | 0.62 | 0.41 | A>A |
| chr9 | 15400000 | 15500000 | 0.51 | 0.43 | A>A |
| chr9 | 15500000 | 15600000 | 0.43 | 0.44 | A>A |

| | | | | | |
|---|---|---|---|---|---|
| chr9 | 15600000 | 15700000 | 0.00 | -0.04 | B>B |
| chr9 | 15700000 | 15800000 | -0.23 | 0.00 | B>B |
| chr9 | 15800000 | 15900000 | -0.18 | 0.14 | B>B |
| chr9 | 15900000 | 16000000 | -0.06 | 0.18 | A>A |
| chr9 | 16000000 | 16100000 | 0.15 | 0.48 | A>A |
| chr9 | 16100000 | 16200000 | 0.24 | 0.52 | A>A |
| chr9 | 16200000 | 16300000 | 0.61 | 0.79 | A>A |
| chr9 | 16300000 | 16400000 | 0.67 | 0.85 | A>A |
| chr9 | 16400000 | 16500000 | 0.68 | 0.91 | A>A |
| chr9 | 16500000 | 16600000 | 0.63 | 0.75 | A>A |
| chr9 | 16600000 | 16700000 | 0.64 | 0.83 | A>A |
| chr9 | 16700000 | 16800000 | 0.60 | 0.71 | A>A |
| chr9 | 16800000 | 16900000 | 0.50 | 0.67 | A>A |
| chr9 | 16900000 | 17000000 | 0.28 | 0.31 | A>A |
| chr9 | 17000000 | 17100000 | 0.06 | 0.11 | A>A |
| chr9 | 17100000 | 17200000 | 0.20 | 0.24 | A>A |
| chr9 | 17200000 | 17300000 | 0.34 | 0.19 | A>A |
| chr9 | 17300000 | 17400000 | 0.14 | 0.08 | A>A |
| chr9 | 17400000 | 17500000 | -0.08 | -0.05 | B>B |
| chr9 | 17500000 | 17600000 | -0.39 | -0.59 | B>B |
| chr9 | 17600000 | 17700000 | -0.74 | -1.02 | B>B |
| chr9 | 17700000 | 17800000 | -0.73 | -0.92 | B>B |
| chr9 | 17800000 | 17900000 | -0.72 | -0.80 | B>B |
| chr9 | 17900000 | 18000000 | -0.77 | -1.01 | B>B |
| chr9 | 18000000 | 18100000 | -0.80 | -0.92 | B>B |
| chr9 | 18100000 | 18200000 | -0.69 | -0.79 | B>B |
| chr9 | 18200000 | 18300000 | -0.36 | -0.33 | B>B |
| chr9 | 18300000 | 18400000 | -0.11 | -0.11 | B>B |
| chr9 | 18400000 | 18500000 | -0.36 | -0.21 | B>B |
| chr9 | 18500000 | 18600000 | -0.69 | -0.39 | B>B |
| chr9 | 18600000 | 18700000 | -0.57 | -0.48 | B>B |
| chr9 | 18700000 | 18800000 | -0.55 | -0.63 | B>B |
| chr9 | 18800000 | 18900000 | -0.59 | -0.66 | B>B |
| chr9 | 18900000 | 19000000 | -0.39 | -0.64 | B>B |
| chr9 | 19000000 | 19100000 | 0.17 | -0.07 | A>A |
| chr9 | 19100000 | 19200000 | 0.87 | 0.51 | A>A |
| chr9 | 19200000 | 19300000 | 0.91 | 0.60 | A>A |
| chr9 | 19300000 | 19400000 | 0.67 | 0.43 | A>A |
| chr9 | 19400000 | 19500000 | -0.02 | -0.26 | B>B |
| chr9 | 19500000 | 19600000 | -0.52 | -0.91 | B>B |
| chr9 | 19600000 | 19700000 | -0.67 | -1.14 | B>B |
| chr9 | 19700000 | 19800000 | -0.84 | -1.15 | B>B |
| chr9 | 19800000 | 19900000 | -0.85 | -1.08 | B>B |
| chr9 | 19900000 | 20000000 | -0.60 | -0.83 | B>B |
| chr9 | 20000000 | 20100000 | -0.76 | -1.01 | B>B |
| chr9 | 20100000 | 20200000 | -0.80 | -1.00 | B>B |
| chr9 | 20200000 | 20300000 | -0.45 | -0.77 | B>B |
| chr9 | 20300000 | 20400000 | -0.03 | -0.14 | B>B |
| chr9 | 20400000 | 20500000 | 0.35 | 0.32 | A>A |
| chr9 | 20500000 | 20600000 | 0.52 | 0.38 | A>A |
| chr9 | 20600000 | 20700000 | 0.41 | 0.29 | A>A |
| chr9 | 20700000 | 20800000 | 0.59 | 0.38 | A>A |
| chr9 | 20800000 | 20900000 | 0.28 | 0.04 | A>A |
| chr9 | 20900000 | 21000000 | 0.09 | -0.12 | B>B |
| chr9 | 21000000 | 21100000 | 0.02 | -0.14 | B>B |
| chr9 | 21100000 | 21200000 | -0.33 | -0.35 | B>B |

| | | | | | |
|---|---|---|---|---|---|
| chr9 | 21200000 | 21300000 | -0.36 | -0.28 | B>B |
| chr9 | 21300000 | 21400000 | -0.11 | -0.04 | B>B |
| chr9 | 21400000 | 21500000 | -0.07 | 0.10 | A>A |
| chr9 | 21500000 | 21600000 | 0.41 | 0.59 | A>A |
| chr9 | 21600000 | 21700000 | 0.85 | 0.58 | A>A |
| chr9 | 21700000 | 21800000 | 0.37 | 0.26 | A>A |
| chr9 | 21800000 | 21900000 | 0.00 | 0.00 | NO |
| chr9 | 21900000 | 22000000 | 0.00 | 0.00 | NO |
| chr9 | 22000000 | 22100000 | 0.00 | 0.00 | NO |
| chr9 | 22100000 | 22200000 | 0.11 | 0.38 | A>A |
| chr9 | 22200000 | 22300000 | -0.13 | 0.04 | B>B |
| chr9 | 22300000 | 22400000 | -0.40 | -0.15 | B>B |
| chr9 | 22400000 | 22500000 | -0.45 | -0.34 | B>B |
| chr9 | 22500000 | 22600000 | -0.90 | -0.78 | B>B |
| chr9 | 22600000 | 22700000 | -0.93 | -0.91 | B>B |
| chr9 | 22700000 | 22800000 | -1.10 | -0.99 | B>B |
| chr9 | 22800000 | 22900000 | -0.98 | -1.03 | B>B |
| chr9 | 22900000 | 23000000 | -1.03 | -1.04 | B>B |
| chr9 | 23000000 | 23100000 | -1.04 | -1.01 | B>B |
| chr9 | 23100000 | 23200000 | -1.08 | -0.96 | B>B |
| chr9 | 23200000 | 23300000 | -1.01 | -1.00 | B>B |
| chr9 | 23300000 | 23400000 | -1.02 | -0.91 | B>B |
| chr9 | 23400000 | 23500000 | -0.98 | -0.95 | B>B |
| chr9 | 23500000 | 23600000 | -0.94 | -1.01 | B>B |
| chr9 | 23600000 | 23700000 | -0.92 | -1.00 | B>B |
| chr9 | 23700000 | 23800000 | -0.85 | -0.98 | B>B |
| chr9 | 23800000 | 23900000 | -0.82 | -0.89 | B>B |
| chr9 | 23900000 | 24000000 | -0.97 | -0.97 | B>B |
| chr9 | 24000000 | 24100000 | -0.98 | -0.98 | B>B |
| chr9 | 24100000 | 24200000 | -1.03 | -0.94 | B>B |
| chr9 | 24200000 | 24300000 | -1.00 | -0.95 | B>B |
| chr9 | 24300000 | 24400000 | -1.00 | -0.95 | B>B |
| chr9 | 24400000 | 24500000 | -0.98 | -1.01 | B>B |
| chr9 | 24500000 | 24600000 | -1.02 | -0.98 | B>B |
| chr9 | 24600000 | 24700000 | -1.05 | -1.02 | B>B |
| chr9 | 24700000 | 24800000 | -0.98 | -0.98 | B>B |
| chr9 | 24800000 | 24900000 | -1.02 | -0.98 | B>B |
| chr9 | 24900000 | 25000000 | -1.07 | -0.99 | B>B |
| chr9 | 25000000 | 25100000 | -0.93 | -0.97 | B>B |
| chr9 | 25100000 | 25200000 | -0.89 | -1.00 | B>B |
| chr9 | 25200000 | 25300000 | -1.00 | -0.98 | B>B |
| chr9 | 25300000 | 25400000 | -0.98 | -0.92 | B>B |
| chr9 | 25400000 | 25500000 | -0.97 | -0.92 | B>B |
| chr9 | 25500000 | 25600000 | -0.95 | -0.95 | B>B |
| chr9 | 25600000 | 25700000 | -0.89 | -0.82 | B>B |
| chr9 | 25700000 | 25800000 | -0.91 | -0.84 | B>B |
| chr9 | 25800000 | 25900000 | -0.92 | -0.82 | B>B |
| chr9 | 25900000 | 26000000 | -0.87 | -0.56 | B>B |
| chr9 | 26000000 | 26100000 | -0.79 | -0.11 | B>A |
| chr9 | 26100000 | 26200000 | -0.82 | 0.03 | B>A |
| chr9 | 26200000 | 26300000 | -0.79 | -0.03 | B>A |
| chr9 | 26300000 | 26400000 | -0.90 | -0.57 | B>B |
| chr9 | 26400000 | 26500000 | -0.96 | -0.94 | B>B |
| chr9 | 26500000 | 26600000 | -0.98 | -1.01 | B>B |
| chr9 | 26600000 | 26700000 | -0.80 | -0.82 | B>B |
| chr9 | 26700000 | 26800000 | -0.35 | -0.66 | B>B |

| | | | | | |
|---|---|---|---|---|---|
| chr9 | 26800000 | 26900000 | 0.08 | -0.33 | B>B |
| chr9 | 26900000 | 27000000 | 0.39 | 0.07 | A>A |
| chr9 | 27000000 | 27100000 | 0.08 | -0.16 | B>B |
| chr9 | 27100000 | 27200000 | -0.22 | -0.27 | B>B |
| chr9 | 27200000 | 27300000 | -0.33 | -0.46 | B>B |
| chr9 | 27300000 | 27400000 | 0.00 | -0.65 | A>B |
| chr9 | 27400000 | 27500000 | 0.23 | -0.76 | A>B |
| chr9 | 27500000 | 27600000 | -0.05 | -0.60 | B>B |
| chr9 | 27600000 | 27700000 | -0.59 | -0.99 | B>B |
| chr9 | 27700000 | 27800000 | -0.77 | -0.94 | B>B |
| chr9 | 27800000 | 27900000 | -0.75 | -1.01 | B>B |
| chr9 | 27900000 | 28000000 | -0.82 | -1.04 | B>B |
| chr9 | 28000000 | 28100000 | -0.76 | -1.00 | B>B |
| chr9 | 28100000 | 28200000 | -0.76 | -0.94 | B>B |
| chr9 | 28200000 | 28300000 | -0.81 | -0.87 | B>B |
| chr9 | 28300000 | 28400000 | -0.73 | -0.84 | B>B |
| chr9 | 28400000 | 28500000 | -0.77 | -0.77 | B>B |
| chr9 | 28500000 | 28600000 | -0.56 | -0.57 | B>B |
| chr9 | 28600000 | 28700000 | -0.40 | -0.38 | B>B |
| chr9 | 28700000 | 28800000 | -0.10 | -0.12 | B>B |
| chr9 | 28800000 | 28900000 | 0.17 | 0.18 | A>A |
| chr9 | 28900000 | 29000000 | 0.18 | 0.18 | A>A |
| chr9 | 29000000 | 29100000 | 0.12 | 0.10 | A>A |
| chr9 | 29100000 | 29200000 | -0.02 | 0.08 | A>A |
| chr9 | 29200000 | 29300000 | -0.32 | -0.05 | B>B |
| chr9 | 29300000 | 29400000 | -0.44 | 0.03 | B>B |
| chr9 | 29400000 | 29500000 | -0.64 | 0.08 | B>A |
| chr9 | 29500000 | 29600000 | -0.77 | 0.14 | B>A |
| chr9 | 29600000 | 29700000 | -0.74 | 0.11 | B>A |
| chr9 | 29700000 | 29800000 | -0.89 | -0.05 | B>A |
| chr9 | 29800000 | 29900000 | -0.92 | -0.16 | B>A |
| chr9 | 29900000 | 30000000 | -0.98 | -0.01 | B>A |
| chr9 | 30000000 | 30100000 | -0.97 | 0.05 | B>A |
| chr9 | 30100000 | 30200000 | -0.94 | 0.05 | B>A |
| chr9 | 30200000 | 30300000 | -0.95 | 0.02 | B>A |
| chr9 | 30300000 | 30400000 | -0.98 | -0.02 | B>A |
| chr9 | 30400000 | 30500000 | -1.00 | -0.14 | B>A |
| chr9 | 30500000 | 30600000 | -1.02 | -0.25 | B>A |
| chr9 | 30600000 | 30700000 | -0.98 | -0.29 | B>A |
| chr9 | 30700000 | 30800000 | -1.06 | -0.46 | B>A |
| chr9 | 30800000 | 30900000 | -1.03 | -0.70 | B>B |
| chr9 | 30900000 | 31000000 | -1.04 | -0.87 | B>B |
| chr9 | 31000000 | 31100000 | -1.02 | -0.93 | B>B |
| chr9 | 31100000 | 31200000 | -1.05 | -1.02 | B>B |
| chr9 | 31200000 | 31300000 | -1.06 | -1.02 | B>B |
| chr9 | 31300000 | 31400000 | -1.06 | -1.00 | B>B |
| chr9 | 31400000 | 31500000 | -1.12 | -1.08 | B>B |
| chr9 | 31500000 | 31600000 | -1.09 | -1.07 | B>B |
| chr9 | 31600000 | 31700000 | -1.12 | -1.09 | B>B |
| chr9 | 31700000 | 31800000 | -1.10 | -1.14 | B>B |
| chr9 | 31800000 | 31900000 | -1.09 | -1.15 | B>B |
| chr9 | 31900000 | 32000000 | -1.18 | -1.16 | B>B |
| chr9 | 32000000 | 32100000 | -1.20 | -1.27 | B>B |
| chr9 | 32100000 | 32200000 | -1.07 | -1.20 | B>B |
| chr9 | 32200000 | 32300000 | -0.95 | -1.01 | B>B |
| chr9 | 32300000 | 32400000 | -0.51 | -0.48 | B>B |

| | | | | | |
|---|---|---|---|---|---|
| chr9 | 32400000 | 32500000 | 0.08 | 0.15 | A>A |
| chr9 | 32500000 | 32600000 | 0.03 | 0.19 | A>A |
| chr9 | 32600000 | 32700000 | -0.15 | 0.43 | B>A |
| chr9 | 32700000 | 32800000 | -0.31 | 0.42 | B>A |
| chr9 | 32800000 | 32900000 | -0.24 | 0.38 | B>A |
| chr9 | 32900000 | 33000000 | 0.05 | 0.40 | A>A |
| chr9 | 33000000 | 33100000 | 1.01 | 0.88 | A>A |
| chr9 | 33100000 | 33200000 | 1.17 | 1.17 | A>A |
| chr9 | 33200000 | 33300000 | 1.12 | 1.25 | A>A |
| chr9 | 33300000 | 33400000 | 1.20 | 1.36 | A>A |
| chr9 | 33400000 | 33500000 | 1.25 | 1.34 | A>A |
| chr9 | 33500000 | 33600000 | 0.00 | 0.00 | NO |
| chr9 | 33600000 | 33700000 | 0.00 | 0.00 | NO |
| chr9 | 33700000 | 33800000 | 0.00 | 0.00 | NO |
| chr9 | 33800000 | 33900000 | 1.33 | 1.41 | A>A |
| chr9 | 33900000 | 34000000 | 1.32 | 1.41 | A>A |
| chr9 | 34000000 | 34100000 | 1.26 | 1.33 | A>A |
| chr9 | 34100000 | 34200000 | 1.44 | 1.39 | A>A |
| chr9 | 34200000 | 34300000 | 1.25 | 1.29 | A>A |
| chr9 | 34300000 | 34400000 | 1.11 | 1.28 | A>A |
| chr9 | 34400000 | 34500000 | 1.25 | 1.36 | A>A |
| chr9 | 34500000 | 34600000 | 1.23 | 1.35 | A>A |
| chr9 | 34600000 | 34700000 | 1.15 | 1.16 | A>A |
| chr9 | 34700000 | 34800000 | 0.99 | 1.08 | A>A |
| chr9 | 34800000 | 34900000 | 0.91 | 1.14 | A>A |
| chr9 | 34900000 | 35000000 | 0.95 | 1.07 | A>A |
| chr9 | 35000000 | 35100000 | 0.95 | 1.12 | A>A |
| chr9 | 35100000 | 35200000 | 0.94 | 1.22 | A>A |
| chr9 | 35200000 | 35300000 | 1.00 | 1.21 | A>A |
| chr9 | 35300000 | 35400000 | 1.04 | 1.14 | A>A |
| chr9 | 35400000 | 35500000 | 0.83 | 1.06 | A>A |
| chr9 | 35500000 | 35600000 | 1.01 | 1.12 | A>A |
| chr9 | 35600000 | 35700000 | 1.09 | 1.17 | A>A |
| chr9 | 35700000 | 35800000 | 1.24 | 1.27 | A>A |
| chr9 | 35800000 | 35900000 | 1.26 | 1.38 | A>A |
| chr9 | 35900000 | 36000000 | 1.24 | 1.23 | A>A |
| chr9 | 36000000 | 36100000 | 1.30 | 1.37 | A>A |
| chr9 | 36100000 | 36200000 | 1.15 | 1.38 | A>A |
| chr9 | 36200000 | 36300000 | 1.32 | 1.34 | A>A |
| chr9 | 36300000 | 36400000 | 1.25 | 1.29 | A>A |
| chr9 | 36400000 | 36500000 | 1.17 | 1.42 | A>A |
| chr9 | 36500000 | 36600000 | 1.30 | 1.35 | A>A |
| chr9 | 36600000 | 36700000 | 1.18 | 1.40 | A>A |
| chr9 | 36700000 | 36800000 | 1.12 | 1.30 | A>A |
| chr9 | 36800000 | 36900000 | 1.13 | 1.28 | A>A |
| chr9 | 36900000 | 37000000 | 1.18 | 1.33 | A>A |
| chr9 | 37000000 | 37100000 | 1.08 | 1.16 | A>A |
| chr9 | 37100000 | 37200000 | 1.24 | 1.37 | A>A |
| chr9 | 37200000 | 37300000 | 1.19 | 1.36 | A>A |
| chr9 | 37300000 | 37400000 | 1.08 | 1.16 | A>A |
| chr9 | 37400000 | 37500000 | 1.14 | 1.22 | A>A |
| chr9 | 37500000 | 37600000 | 1.13 | 1.35 | A>A |
| chr9 | 37600000 | 37700000 | 1.21 | 1.26 | A>A |
| chr9 | 37700000 | 37800000 | 1.16 | 1.17 | A>A |
| chr9 | 37800000 | 37900000 | 1.10 | 1.19 | A>A |
| chr9 | 37900000 | 38000000 | 1.03 | 1.24 | A>A |

| | | | | | |
|---|---|---|---|---|---|
| chr9 | 38000000 | 38100000 | 1.16 | 1.26 | A>A |
| chr9 | 38100000 | 38200000 | 1.12 | 1.35 | A>A |
| chr9 | 38200000 | 38300000 | 1.19 | 1.26 | A>A |
| chr9 | 38300000 | 38400000 | 0.92 | 1.13 | A>A |
| chr9 | 38400000 | 38500000 | 1.21 | 1.42 | A>A |
| chr9 | 38500000 | 38600000 | 0.00 | 0.00 | NO |
| chr9 | 38600000 | 38700000 | 0.00 | 0.00 | NO |
| chr9 | 38700000 | 38800000 | 0.00 | 0.00 | NO |
| chr9 | 38800000 | 38900000 | 0.00 | 0.00 | NO |
| chr9 | 38900000 | 39000000 | 0.00 | 0.00 | NO |
| chr9 | 39000000 | 39100000 | 0.00 | 0.00 | NO |
| chr9 | 39100000 | 39200000 | 0.00 | 0.00 | NO |
| chr9 | 39200000 | 39300000 | 0.00 | 0.00 | NO |
| chr9 | 39300000 | 39400000 | 0.00 | 0.00 | NO |
| chr9 | 39400000 | 39500000 | 0.00 | 0.00 | NO |
| chr9 | 39500000 | 39600000 | 0.00 | 0.00 | NO |
| chr9 | 39600000 | 39700000 | 0.00 | 0.00 | NO |
| chr9 | 39700000 | 39800000 | 0.00 | 0.00 | NO |
| chr9 | 39800000 | 39900000 | 0.00 | 0.00 | NO |
| chr9 | 39900000 | 40000000 | 0.00 | 0.00 | NO |
| chr9 | 40000000 | 40100000 | 0.00 | 0.00 | NO |
| chr9 | 40100000 | 40200000 | 0.00 | 0.00 | NO |
| chr9 | 40200000 | 40300000 | 0.00 | 0.00 | NO |
| chr9 | 40300000 | 40400000 | 0.00 | 0.00 | NO |
| chr9 | 40400000 | 40500000 | 0.00 | 0.00 | NO |
| chr9 | 40500000 | 40600000 | 0.00 | 0.00 | NO |
| chr9 | 40600000 | 40700000 | 0.00 | 0.00 | NO |
| chr9 | 40700000 | 40800000 | 0.00 | 0.00 | NO |
| chr9 | 40800000 | 40900000 | 0.00 | 0.00 | NO |
| chr9 | 40900000 | 41000000 | 0.00 | 0.00 | NO |
| chr9 | 41000000 | 41100000 | 0.00 | 0.00 | NO |
| chr9 | 41100000 | 41200000 | 0.00 | 0.00 | NO |
| chr9 | 41200000 | 41300000 | 0.00 | 0.00 | NO |
| chr9 | 41300000 | 41400000 | 0.00 | 0.00 | NO |
| chr9 | 41400000 | 41500000 | 0.00 | 0.00 | NO |
| chr9 | 41500000 | 41600000 | 0.00 | 0.00 | NO |
| chr9 | 41600000 | 41700000 | 0.00 | 0.00 | NO |
| chr9 | 41700000 | 41800000 | 0.00 | 0.00 | NO |
| chr9 | 41800000 | 41900000 | 0.00 | 0.00 | NO |
| chr9 | 41900000 | 42000000 | 0.00 | 0.00 | NO |
| chr9 | 42000000 | 42100000 | 0.00 | 0.00 | NO |
| chr9 | 42100000 | 42200000 | 0.00 | 0.00 | NO |
| chr9 | 42200000 | 42300000 | 0.00 | 0.00 | NO |
| chr9 | 42300000 | 42400000 | 0.00 | 0.00 | NO |
| chr9 | 42400000 | 42500000 | 0.00 | 0.00 | NO |
| chr9 | 42500000 | 42600000 | 0.00 | 0.00 | NO |
| chr9 | 42600000 | 42700000 | 0.00 | 0.00 | NO |
| chr9 | 42700000 | 42800000 | 0.00 | 0.00 | NO |
| chr9 | 42800000 | 42900000 | 0.00 | 0.00 | NO |
| chr9 | 42900000 | 43000000 | 0.00 | 0.00 | NO |
| chr9 | 43000000 | 43100000 | 0.00 | 0.00 | NO |
| chr9 | 43100000 | 43200000 | 0.00 | 0.00 | NO |
| chr9 | 43200000 | 43300000 | 0.00 | 0.00 | NO |
| chr9 | 43300000 | 43400000 | 0.00 | 0.00 | NO |
| chr9 | 43400000 | 43500000 | 0.00 | 0.00 | NO |
| chr9 | 43500000 | 43600000 | 0.00 | 0.00 | NO |

| | | | | | |
|---|---|---|---|---|---|
| chr9 | 43600000 | 43700000 | 0.00 | 0.00 | NO |
| chr9 | 43700000 | 43800000 | 0.00 | 0.00 | NO |
| chr9 | 43800000 | 43900000 | 0.00 | 0.00 | NO |
| chr9 | 43900000 | 44000000 | 0.00 | 0.00 | NO |
| chr9 | 44000000 | 44100000 | 0.00 | 0.00 | NO |
| chr9 | 44100000 | 44200000 | 0.00 | 0.00 | NO |
| chr9 | 44200000 | 44300000 | 0.00 | 0.00 | NO |
| chr9 | 44300000 | 44400000 | 0.00 | 0.00 | NO |
| chr9 | 44400000 | 44500000 | 0.00 | 0.00 | NO |
| chr9 | 44500000 | 44600000 | 0.00 | 0.00 | NO |
| chr9 | 44600000 | 44700000 | 0.00 | 0.00 | NO |
| chr9 | 44700000 | 44800000 | 0.00 | 0.00 | NO |
| chr9 | 44800000 | 44900000 | 0.00 | 0.00 | NO |
| chr9 | 44900000 | 45000000 | 0.00 | 0.00 | NO |
| chr9 | 45000000 | 45100000 | 0.00 | 0.00 | NO |
| chr9 | 45100000 | 45200000 | 0.00 | 0.00 | NO |
| chr9 | 45200000 | 45300000 | 0.00 | 0.00 | NO |
| chr9 | 45300000 | 45400000 | 0.00 | 0.00 | NO |
| chr9 | 45400000 | 45500000 | 0.00 | 0.00 | NO |
| chr9 | 45500000 | 45600000 | 0.00 | 0.00 | NO |
| chr9 | 45600000 | 45700000 | 0.00 | 0.00 | NO |
| chr9 | 45700000 | 45800000 | 0.00 | 0.00 | NO |
| chr9 | 45800000 | 45900000 | 0.00 | 0.00 | NO |
| chr9 | 45900000 | 46000000 | 0.00 | 0.00 | NO |
| chr9 | 46000000 | 46100000 | 0.00 | 0.00 | NO |
| chr9 | 46100000 | 46200000 | 0.00 | 0.00 | NO |
| chr9 | 46200000 | 46300000 | 0.00 | 0.00 | NO |
| chr9 | 46300000 | 46400000 | 0.00 | 0.00 | NO |
| chr9 | 46400000 | 46500000 | 0.00 | 0.00 | NO |
| chr9 | 46500000 | 46600000 | 0.00 | 0.00 | NO |
| chr9 | 46600000 | 46700000 | 0.00 | 0.00 | NO |
| chr9 | 46700000 | 46800000 | 0.00 | 0.00 | NO |
| chr9 | 46800000 | 46900000 | 0.00 | 0.00 | NO |
| chr9 | 46900000 | 47000000 | 0.00 | 0.00 | NO |
| chr9 | 47000000 | 47100000 | 0.00 | 0.00 | NO |
| chr9 | 47100000 | 47200000 | 0.00 | 0.00 | NO |
| chr9 | 47200000 | 47300000 | 0.00 | 0.00 | NO |
| chr9 | 47300000 | 47400000 | 0.00 | 0.00 | NO |
| chr9 | 47400000 | 47500000 | 0.00 | 0.00 | NO |
| chr9 | 47500000 | 47600000 | 0.00 | 0.00 | NO |
| chr9 | 47600000 | 47700000 | 0.00 | 0.00 | NO |
| chr9 | 47700000 | 47800000 | 0.00 | 0.00 | NO |
| chr9 | 47800000 | 47900000 | 0.00 | 0.00 | NO |
| chr9 | 47900000 | 48000000 | 0.00 | 0.00 | NO |
| chr9 | 48000000 | 48100000 | 0.00 | 0.00 | NO |
| chr9 | 48100000 | 48200000 | 0.00 | 0.00 | NO |
| chr9 | 48200000 | 48300000 | 0.00 | 0.00 | NO |
| chr9 | 48300000 | 48400000 | 0.00 | 0.00 | NO |
| chr9 | 48400000 | 48500000 | 0.00 | 0.00 | NO |
| chr9 | 48500000 | 48600000 | 0.00 | 0.00 | NO |
| chr9 | 48600000 | 48700000 | 0.00 | 0.00 | NO |
| chr9 | 48700000 | 48800000 | 0.00 | 0.00 | NO |
| chr9 | 48800000 | 48900000 | 0.00 | 0.00 | NO |
| chr9 | 48900000 | 49000000 | 0.00 | 0.00 | NO |
| chr9 | 49000000 | 49100000 | 0.00 | 0.00 | NO |
| chr9 | 49100000 | 49200000 | 0.00 | 0.00 | NO |

| | | | | | |
|---|---|---|---|---|---|
| chr9 | 49200000 | 49300000 | 0.00 | 0.00 | NO |
| chr9 | 49300000 | 49400000 | 0.00 | 0.00 | NO |
| chr9 | 49400000 | 49500000 | 0.00 | 0.00 | NO |
| chr9 | 49500000 | 49600000 | 0.00 | 0.00 | NO |
| chr9 | 49600000 | 49700000 | 0.00 | 0.00 | NO |
| chr9 | 49700000 | 49800000 | 0.00 | 0.00 | NO |
| chr9 | 49800000 | 49900000 | 0.00 | 0.00 | NO |
| chr9 | 49900000 | 50000000 | 0.00 | 0.00 | NO |
| chr9 | 50000000 | 50100000 | 0.00 | 0.00 | NO |
| chr9 | 50100000 | 50200000 | 0.00 | 0.00 | NO |
| chr9 | 50200000 | 50300000 | 0.00 | 0.00 | NO |
| chr9 | 50300000 | 50400000 | 0.00 | 0.00 | NO |
| chr9 | 50400000 | 50500000 | 0.00 | 0.00 | NO |
| chr9 | 50500000 | 50600000 | 0.00 | 0.00 | NO |
| chr9 | 50600000 | 50700000 | 0.00 | 0.00 | NO |
| chr9 | 50700000 | 50800000 | 0.00 | 0.00 | NO |
| chr9 | 50800000 | 50900000 | 0.00 | 0.00 | NO |
| chr9 | 50900000 | 51000000 | 0.00 | 0.00 | NO |
| chr9 | 51000000 | 51100000 | 0.00 | 0.00 | NO |
| chr9 | 51100000 | 51200000 | 0.00 | 0.00 | NO |
| chr9 | 51200000 | 51300000 | 0.00 | 0.00 | NO |
| chr9 | 51300000 | 51400000 | 0.00 | 0.00 | NO |
| chr9 | 51400000 | 51500000 | 0.00 | 0.00 | NO |
| chr9 | 51500000 | 51600000 | 0.00 | 0.00 | NO |
| chr9 | 51600000 | 51700000 | 0.00 | 0.00 | NO |
| chr9 | 51700000 | 51800000 | 0.00 | 0.00 | NO |
| chr9 | 51800000 | 51900000 | 0.00 | 0.00 | NO |
| chr9 | 51900000 | 52000000 | 0.00 | 0.00 | NO |
| chr9 | 52000000 | 52100000 | 0.00 | 0.00 | NO |
| chr9 | 52100000 | 52200000 | 0.00 | 0.00 | NO |
| chr9 | 52200000 | 52300000 | 0.00 | 0.00 | NO |
| chr9 | 52300000 | 52400000 | 0.00 | 0.00 | NO |
| chr9 | 52400000 | 52500000 | 0.00 | 0.00 | NO |
| chr9 | 52500000 | 52600000 | 0.00 | 0.00 | NO |
| chr9 | 52600000 | 52700000 | 0.00 | 0.00 | NO |
| chr9 | 52700000 | 52800000 | 0.00 | 0.00 | NO |
| chr9 | 52800000 | 52900000 | 0.00 | 0.00 | NO |
| chr9 | 52900000 | 53000000 | 0.00 | 0.00 | NO |
| chr9 | 53000000 | 53100000 | 0.00 | 0.00 | NO |
| chr9 | 53100000 | 53200000 | 0.00 | 0.00 | NO |
| chr9 | 53200000 | 53300000 | 0.00 | 0.00 | NO |
| chr9 | 53300000 | 53400000 | 0.00 | 0.00 | NO |
| chr9 | 53400000 | 53500000 | 0.00 | 0.00 | NO |
| chr9 | 53500000 | 53600000 | 0.00 | 0.00 | NO |
| chr9 | 53600000 | 53700000 | 0.00 | 0.00 | NO |
| chr9 | 53700000 | 53800000 | 0.00 | 0.00 | NO |
| chr9 | 53800000 | 53900000 | 0.00 | 0.00 | NO |
| chr9 | 53900000 | 54000000 | 0.00 | 0.00 | NO |
| chr9 | 54000000 | 54100000 | 0.00 | 0.00 | NO |
| chr9 | 54100000 | 54200000 | 0.00 | 0.00 | NO |
| chr9 | 54200000 | 54300000 | 0.00 | 0.00 | NO |
| chr9 | 54300000 | 54400000 | 0.00 | 0.00 | NO |
| chr9 | 54400000 | 54500000 | 0.00 | 0.00 | NO |
| chr9 | 54500000 | 54600000 | 0.00 | 0.00 | NO |
| chr9 | 54600000 | 54700000 | 0.00 | 0.00 | NO |
| chr9 | 54700000 | 54800000 | 0.00 | 0.00 | NO |

| | | | | | |
|---|---|---|---|---|---|
| chr9 | 54800000 | 54900000 | 0.00 | 0.00 | NO |
| chr9 | 54900000 | 55000000 | 0.00 | 0.00 | NO |
| chr9 | 55000000 | 55100000 | 0.00 | 0.00 | NO |
| chr9 | 55100000 | 55200000 | 0.00 | 0.00 | NO |
| chr9 | 55200000 | 55300000 | 0.00 | 0.00 | NO |
| chr9 | 55300000 | 55400000 | 0.00 | 0.00 | NO |
| chr9 | 55400000 | 55500000 | 0.00 | 0.00 | NO |
| chr9 | 55500000 | 55600000 | 0.00 | 0.00 | NO |
| chr9 | 55600000 | 55700000 | 0.00 | 0.00 | NO |
| chr9 | 55700000 | 55800000 | 0.00 | 0.00 | NO |
| chr9 | 55800000 | 55900000 | 0.00 | 0.00 | NO |
| chr9 | 55900000 | 56000000 | 0.00 | 0.00 | NO |
| chr9 | 56000000 | 56100000 | 0.00 | 0.00 | NO |
| chr9 | 56100000 | 56200000 | 0.00 | 0.00 | NO |
| chr9 | 56200000 | 56300000 | 0.00 | 0.00 | NO |
| chr9 | 56300000 | 56400000 | 0.00 | 0.00 | NO |
| chr9 | 56400000 | 56500000 | 0.00 | 0.00 | NO |
| chr9 | 56500000 | 56600000 | 0.00 | 0.00 | NO |
| chr9 | 56600000 | 56700000 | 0.00 | 0.00 | NO |
| chr9 | 56700000 | 56800000 | 0.00 | 0.00 | NO |
| chr9 | 56800000 | 56900000 | 0.00 | 0.00 | NO |
| chr9 | 56900000 | 57000000 | 0.00 | 0.00 | NO |
| chr9 | 57000000 | 57100000 | 0.00 | 0.00 | NO |
| chr9 | 57100000 | 57200000 | 0.00 | 0.00 | NO |
| chr9 | 57200000 | 57300000 | 0.00 | 0.00 | NO |
| chr9 | 57300000 | 57400000 | 0.00 | 0.00 | NO |
| chr9 | 57400000 | 57500000 | 0.00 | 0.00 | NO |
| chr9 | 57500000 | 57600000 | 0.00 | 0.00 | NO |
| chr9 | 57600000 | 57700000 | 0.00 | 0.00 | NO |
| chr9 | 57700000 | 57800000 | 0.00 | 0.00 | NO |
| chr9 | 57800000 | 57900000 | 0.00 | 0.00 | NO |
| chr9 | 57900000 | 58000000 | 0.00 | 0.00 | NO |
| chr9 | 58000000 | 58100000 | 0.00 | 0.00 | NO |
| chr9 | 58100000 | 58200000 | 0.00 | 0.00 | NO |
| chr9 | 58200000 | 58300000 | 0.00 | 0.00 | NO |
| chr9 | 58300000 | 58400000 | 0.00 | 0.00 | NO |
| chr9 | 58400000 | 58500000 | 0.00 | 0.00 | NO |
| chr9 | 58500000 | 58600000 | 0.00 | 0.00 | NO |
| chr9 | 58600000 | 58700000 | 0.00 | 0.00 | NO |
| chr9 | 58700000 | 58800000 | 0.00 | 0.00 | NO |
| chr9 | 58800000 | 58900000 | 0.00 | 0.00 | NO |
| chr9 | 58900000 | 59000000 | 0.00 | 0.00 | NO |
| chr9 | 59000000 | 59100000 | 0.00 | 0.00 | NO |
| chr9 | 59100000 | 59200000 | 0.00 | 0.00 | NO |
| chr9 | 59200000 | 59300000 | 0.00 | 0.00 | NO |
| chr9 | 59300000 | 59400000 | 0.00 | 0.00 | NO |
| chr9 | 59400000 | 59500000 | 0.00 | 0.00 | NO |
| chr9 | 59500000 | 59600000 | 0.00 | 0.00 | NO |
| chr9 | 59600000 | 59700000 | 0.00 | 0.00 | NO |
| chr9 | 59700000 | 59800000 | 0.00 | 0.00 | NO |
| chr9 | 59800000 | 59900000 | 0.00 | 0.00 | NO |
| chr9 | 59900000 | 60000000 | 0.00 | 0.00 | NO |
| chr9 | 60000000 | 60100000 | 0.00 | 0.00 | NO |
| chr9 | 60100000 | 60200000 | 0.00 | 0.00 | NO |
| chr9 | 60200000 | 60300000 | 0.00 | 0.00 | NO |
| chr9 | 60300000 | 60400000 | 0.00 | 0.00 | NO |

| | | | | | |
|---|---|---|---|---|---|
| chr9 | 60400000 | 60500000 | 0.00 | 0.00 | NO |
| chr9 | 60500000 | 60600000 | 0.00 | 0.00 | NO |
| chr9 | 60600000 | 60700000 | 0.00 | 0.00 | NO |
| chr9 | 60700000 | 60800000 | 0.00 | 0.00 | NO |
| chr9 | 60800000 | 60900000 | 0.00 | 0.00 | NO |
| chr9 | 60900000 | 61000000 | 0.00 | 0.00 | NO |
| chr9 | 61000000 | 61100000 | 0.00 | 0.00 | NO |
| chr9 | 61100000 | 61200000 | 0.00 | 0.00 | NO |
| chr9 | 61200000 | 61300000 | 0.00 | 0.00 | NO |
| chr9 | 61300000 | 61400000 | 0.00 | 0.00 | NO |
| chr9 | 61400000 | 61500000 | 0.00 | 0.00 | NO |
| chr9 | 61500000 | 61600000 | 0.00 | 0.00 | NO |
| chr9 | 61600000 | 61700000 | 0.00 | 0.00 | NO |
| chr9 | 61700000 | 61800000 | 0.00 | 0.00 | NO |
| chr9 | 61800000 | 61900000 | 0.00 | 0.00 | NO |
| chr9 | 61900000 | 62000000 | 0.00 | 0.00 | NO |
| chr9 | 62000000 | 62100000 | 0.00 | 0.00 | NO |
| chr9 | 62100000 | 62200000 | 0.00 | 0.00 | NO |
| chr9 | 62200000 | 62300000 | 0.00 | 0.00 | NO |
| chr9 | 62300000 | 62400000 | 0.00 | 0.00 | NO |
| chr9 | 62400000 | 62500000 | 0.00 | 0.00 | NO |
| chr9 | 62500000 | 62600000 | 0.00 | 0.00 | NO |
| chr9 | 62600000 | 62700000 | 0.00 | 0.00 | NO |
| chr9 | 62700000 | 62800000 | 0.00 | 0.00 | NO |
| chr9 | 62800000 | 62900000 | 0.00 | 0.00 | NO |
| chr9 | 62900000 | 63000000 | 0.00 | 0.00 | NO |
| chr9 | 63000000 | 63100000 | 0.00 | 0.00 | NO |
| chr9 | 63100000 | 63200000 | 0.00 | 0.00 | NO |
| chr9 | 63200000 | 63300000 | 0.00 | 0.00 | NO |
| chr9 | 63300000 | 63400000 | 0.00 | 0.00 | NO |
| chr9 | 63400000 | 63500000 | 0.00 | 0.00 | NO |
| chr9 | 63500000 | 63600000 | 0.00 | 0.00 | NO |
| chr9 | 63600000 | 63700000 | 0.00 | 0.00 | NO |
| chr9 | 63700000 | 63800000 | 0.00 | 0.00 | NO |
| chr9 | 63800000 | 63900000 | 0.00 | 0.00 | NO |
| chr9 | 63900000 | 64000000 | 0.00 | 0.00 | NO |
| chr9 | 64000000 | 64100000 | 0.00 | 0.00 | NO |
| chr9 | 64100000 | 64200000 | 0.00 | 0.00 | NO |
| chr9 | 64200000 | 64300000 | 0.00 | 0.00 | NO |
| chr9 | 64300000 | 64400000 | 0.00 | 0.00 | NO |
| chr9 | 64400000 | 64500000 | 0.00 | 0.00 | NO |
| chr9 | 64500000 | 64600000 | 0.00 | 0.00 | NO |
| chr9 | 64600000 | 64700000 | 0.00 | 0.00 | NO |
| chr9 | 64700000 | 64800000 | 0.00 | 0.00 | NO |
| chr9 | 64800000 | 64900000 | 0.00 | 0.00 | NO |
| chr9 | 64900000 | 65000000 | 0.00 | 0.00 | NO |
| chr9 | 65000000 | 65100000 | 0.00 | 0.00 | NO |
| chr9 | 65100000 | 65200000 | 0.00 | 0.00 | NO |
| chr9 | 65200000 | 65300000 | 0.00 | 0.00 | NO |
| chr9 | 65300000 | 65400000 | 0.00 | 0.00 | NO |
| chr9 | 65400000 | 65500000 | 0.00 | 0.00 | NO |
| chr9 | 65500000 | 65600000 | 0.00 | 0.00 | NO |
| chr9 | 65600000 | 65700000 | 0.00 | 0.00 | NO |
| chr9 | 65700000 | 65800000 | 0.00 | 0.00 | NO |
| chr9 | 65800000 | 65900000 | 0.00 | 0.00 | NO |
| chr9 | 65900000 | 66000000 | 0.00 | 0.00 | NO |

| | | | | | |
|---|---|---|---|---|---|
| chr9 | 66000000 | 66100000 | 0.00 | 0.00 | NO |
| chr9 | 66100000 | 66200000 | 0.00 | 0.00 | NO |
| chr9 | 66200000 | 66300000 | 0.00 | 0.00 | NO |
| chr9 | 66300000 | 66400000 | 0.00 | 0.00 | NO |
| chr9 | 66400000 | 66500000 | 0.00 | 0.00 | NO |
| chr9 | 66500000 | 66600000 | 0.00 | 0.00 | NO |
| chr9 | 66600000 | 66700000 | 0.00 | 0.00 | NO |
| chr9 | 66700000 | 66800000 | 0.00 | 0.00 | NO |
| chr9 | 66800000 | 66900000 | 0.00 | 0.00 | NO |
| chr9 | 66900000 | 67000000 | 0.00 | 0.00 | NO |
| chr9 | 67000000 | 67100000 | 0.00 | 0.00 | NO |
| chr9 | 67100000 | 67200000 | 0.00 | 0.00 | NO |
| chr9 | 67200000 | 67300000 | 0.00 | 0.00 | NO |
| chr9 | 67300000 | 67400000 | 0.00 | 0.00 | NO |
| chr9 | 67400000 | 67500000 | 0.00 | 0.00 | NO |
| chr9 | 67500000 | 67600000 | 0.00 | 0.00 | NO |
| chr9 | 67600000 | 67700000 | 0.00 | 0.00 | NO |
| chr9 | 67700000 | 67800000 | 0.00 | 0.00 | NO |
| chr9 | 67800000 | 67900000 | 0.00 | 0.00 | NO |
| chr9 | 67900000 | 68000000 | 0.00 | 0.00 | NO |
| chr9 | 68000000 | 68100000 | 0.00 | 0.00 | NO |
| chr9 | 68100000 | 68200000 | 0.00 | 0.00 | NO |
| chr9 | 68200000 | 68300000 | 0.00 | 0.00 | NO |
| chr9 | 68300000 | 68400000 | 0.00 | 0.00 | NO |
| chr9 | 68400000 | 68500000 | 0.00 | 0.00 | NO |
| chr9 | 68500000 | 68600000 | 0.00 | 0.00 | NO |
| chr9 | 68600000 | 68700000 | 0.00 | 0.00 | NO |
| chr9 | 68700000 | 68800000 | 0.00 | 0.00 | NO |
| chr9 | 68800000 | 68900000 | 0.00 | 0.00 | NO |
| chr9 | 68900000 | 69000000 | 0.00 | 0.00 | NO |
| chr9 | 69000000 | 69100000 | 0.00 | 0.00 | NO |
| chr9 | 69100000 | 69200000 | 0.00 | 0.00 | NO |
| chr9 | 69200000 | 69300000 | 0.00 | 0.00 | NO |
| chr9 | 69300000 | 69400000 | 0.00 | 0.00 | NO |
| chr9 | 69400000 | 69500000 | 0.00 | 0.00 | NO |
| chr9 | 69500000 | 69600000 | 0.00 | 0.00 | NO |
| chr9 | 69600000 | 69700000 | 0.00 | 0.00 | NO |
| chr9 | 69700000 | 69800000 | 0.00 | 0.00 | NO |
| chr9 | 69800000 | 69900000 | 0.00 | 0.00 | NO |
| chr9 | 69900000 | 70000000 | 0.00 | 0.00 | NO |
| chr9 | 70000000 | 70100000 | 0.00 | 0.00 | NO |
| chr9 | 70100000 | 70200000 | 0.00 | 0.00 | NO |
| chr9 | 70200000 | 70300000 | 0.00 | 0.00 | NO |
| chr9 | 70300000 | 70400000 | 0.00 | 0.00 | NO |
| chr9 | 70400000 | 70500000 | 0.00 | 0.00 | NO |
| chr9 | 70500000 | 70600000 | 0.00 | 0.00 | NO |
| chr9 | 70600000 | 70700000 | 0.00 | 0.00 | NO |
| chr9 | 70700000 | 70800000 | 0.00 | 0.00 | NO |
| chr9 | 70800000 | 70900000 | 0.00 | 0.00 | NO |
| chr9 | 70900000 | 71000000 | 0.00 | 0.00 | NO |
| chr9 | 71000000 | 71100000 | 0.00 | -0.94 | NO |
| chr9 | 71100000 | 71200000 | -0.39 | -0.96 | B>B |
| chr9 | 71200000 | 71300000 | -0.72 | -1.26 | B>B |
| chr9 | 71300000 | 71400000 | -0.57 | -1.16 | A>B |
| chr9 | 71400000 | 71500000 | -0.58 | -1.20 | A>B |
| chr9 | 71500000 | 71600000 | -0.61 | -1.26 | A>B |

| | | | | | |
|---|---|---|---|---|---|
| chr9 | 71600000 | 71700000 | -0.03 | -0.43 | B>B |
| chr9 | 71700000 | 71800000 | 0.25 | 0.22 | A>A |
| chr9 | 71800000 | 71900000 | 0.04 | -0.04 | B>B |
| chr9 | 71900000 | 72000000 | -0.23 | -0.15 | B>B |
| chr9 | 72000000 | 72100000 | -0.37 | -0.26 | B>B |
| chr9 | 72100000 | 72200000 | -0.42 | -0.28 | B>B |
| chr9 | 72200000 | 72300000 | -0.47 | -0.44 | B>B |
| chr9 | 72300000 | 72400000 | 0.06 | 0.03 | A>A |
| chr9 | 72400000 | 72500000 | 0.09 | -0.01 | A>A |
| chr9 | 72500000 | 72600000 | -0.23 | -0.39 | B>B |
| chr9 | 72600000 | 72700000 | -0.49 | -1.04 | B>B |
| chr9 | 72700000 | 72800000 | -0.57 | -1.07 | B>B |
| chr9 | 72800000 | 72900000 | -0.35 | -0.66 | B>B |
| chr9 | 72900000 | 73000000 | 0.12 | -0.30 | B>B |
| chr9 | 73000000 | 73100000 | 0.32 | -0.46 | A>B |
| chr9 | 73100000 | 73200000 | -0.10 | -0.89 | A>B |
| chr9 | 73200000 | 73300000 | -0.37 | -0.80 | B>B |
| chr9 | 73300000 | 73400000 | -0.87 | -1.14 | B>B |
| chr9 | 73400000 | 73500000 | -0.84 | -1.26 | B>B |
| chr9 | 73500000 | 73600000 | -0.77 | -1.28 | B>B |
| chr9 | 73600000 | 73700000 | -0.92 | -1.32 | B>B |
| chr9 | 73700000 | 73800000 | -0.91 | -1.25 | B>B |
| chr9 | 73800000 | 73900000 | -0.65 | -0.98 | B>B |
| chr9 | 73900000 | 74000000 | -0.73 | -0.97 | B>B |
| chr9 | 74000000 | 74100000 | -0.50 | -0.82 | B>B |
| chr9 | 74100000 | 74200000 | -0.18 | -0.62 | B>B |
| chr9 | 74200000 | 74300000 | 0.38 | 0.01 | A>A |
| chr9 | 74300000 | 74400000 | 0.44 | 0.15 | A>A |
| chr9 | 74400000 | 74500000 | 0.26 | 0.13 | A>A |
| chr9 | 74500000 | 74600000 | 0.35 | 0.36 | A>A |
| chr9 | 74600000 | 74700000 | 0.16 | 0.35 | A>A |
| chr9 | 74700000 | 74800000 | 0.34 | 0.59 | A>A |
| chr9 | 74800000 | 74900000 | 0.31 | 0.49 | A>A |
| chr9 | 74900000 | 75000000 | 0.67 | 0.84 | A>A |
| chr9 | 75000000 | 75100000 | 1.11 | 1.03 | A>A |
| chr9 | 75100000 | 75200000 | 0.87 | 0.95 | A>A |
| chr9 | 75200000 | 75300000 | 0.64 | 0.78 | A>A |
| chr9 | 75300000 | 75400000 | 0.60 | 0.73 | A>A |
| chr9 | 75400000 | 75500000 | 0.35 | 0.47 | A>A |
| chr9 | 75500000 | 75600000 | 0.47 | 0.64 | A>A |
| chr9 | 75600000 | 75700000 | 0.55 | 0.68 | A>A |
| chr9 | 75700000 | 75800000 | 0.55 | 0.51 | A>A |
| chr9 | 75800000 | 75900000 | -0.35 | -0.24 | B>B |
| chr9 | 75900000 | 76000000 | -0.67 | -0.24 | B>B |
| chr9 | 76000000 | 76100000 | -0.67 | -0.15 | B>A |
| chr9 | 76100000 | 76200000 | -0.62 | -0.14 | B>B |
| chr9 | 76200000 | 76300000 | -0.67 | -0.35 | B>B |
| chr9 | 76300000 | 76400000 | -0.71 | -0.74 | B>B |
| chr9 | 76400000 | 76500000 | -0.68 | -1.01 | B>B |
| chr9 | 76500000 | 76600000 | -0.78 | -1.00 | B>B |
| chr9 | 76600000 | 76700000 | -0.84 | -1.09 | B>B |
| chr9 | 76700000 | 76800000 | -0.78 | -1.14 | B>B |
| chr9 | 76800000 | 76900000 | -0.80 | -1.06 | B>B |
| chr9 | 76900000 | 77000000 | -0.84 | -1.06 | B>B |
| chr9 | 77000000 | 77100000 | -0.79 | -1.08 | B>B |
| chr9 | 77100000 | 77200000 | -0.66 | -0.86 | B>B |

| | | | | | |
|---|---|---|---|---|---|
| chr9 | 77200000 | 77300000 | -0.63 | -0.78 | B>B |
| chr9 | 77300000 | 77400000 | -0.40 | -0.51 | B>B |
| chr9 | 77400000 | 77500000 | -0.04 | 0.21 | A>A |
| chr9 | 77500000 | 77600000 | 0.07 | 0.16 | A>A |
| chr9 | 77600000 | 77700000 | 0.03 | -0.10 | B>B |
| chr9 | 77700000 | 77800000 | -0.37 | -0.37 | B>B |
| chr9 | 77800000 | 77900000 | -0.69 | -0.78 | B>B |
| chr9 | 77900000 | 78000000 | -0.89 | -1.08 | B>B |
| chr9 | 78000000 | 78100000 | -0.92 | -1.21 | B>B |
| chr9 | 78100000 | 78200000 | -1.01 | -1.18 | B>B |
| chr9 | 78200000 | 78300000 | -1.02 | -1.18 | B>B |
| chr9 | 78300000 | 78400000 | -1.00 | -1.11 | B>B |
| chr9 | 78400000 | 78500000 | -0.80 | -0.98 | B>B |
| chr9 | 78500000 | 78600000 | -0.62 | -0.81 | B>B |
| chr9 | 78600000 | 78700000 | -0.55 | -0.95 | B>B |
| chr9 | 78700000 | 78800000 | -0.42 | -0.80 | B>B |
| chr9 | 78800000 | 78900000 | -0.21 | -0.73 | B>B |
| chr9 | 78900000 | 79000000 | 0.07 | -0.27 | B>B |
| chr9 | 79000000 | 79100000 | 0.04 | -0.37 | B>B |
| chr9 | 79100000 | 79200000 | -0.20 | -0.91 | A>B |
| chr9 | 79200000 | 79300000 | -0.19 | -0.81 | A>B |
| chr9 | 79300000 | 79400000 | -0.12 | -0.72 | A>B |
| chr9 | 79400000 | 79500000 | 0.10 | -0.71 | A>B |
| chr9 | 79500000 | 79600000 | 0.08 | -0.94 | A>B |
| chr9 | 79600000 | 79700000 | 0.11 | -0.77 | A>B |
| chr9 | 79700000 | 79800000 | 0.14 | -0.54 | A>B |
| chr9 | 79800000 | 79900000 | 0.61 | 0.07 | A>A |
| chr9 | 79900000 | 80000000 | 0.64 | 0.25 | A>A |
| chr9 | 80000000 | 80100000 | 0.49 | 0.37 | A>A |
| chr9 | 80100000 | 80200000 | 0.64 | 0.72 | A>A |
| chr9 | 80200000 | 80300000 | 0.49 | 0.69 | A>A |
| chr9 | 80300000 | 80400000 | 0.42 | 0.55 | A>A |
| chr9 | 80400000 | 80500000 | 0.67 | 0.66 | A>A |
| chr9 | 80500000 | 80600000 | 0.70 | 0.81 | A>A |
| chr9 | 80600000 | 80700000 | 0.57 | 0.67 | A>A |
| chr9 | 80700000 | 80800000 | 0.36 | 0.43 | A>A |
| chr9 | 80800000 | 80900000 | 0.48 | 0.38 | A>A |
| chr9 | 80900000 | 81000000 | 0.33 | 0.31 | A>A |
| chr9 | 81000000 | 81100000 | 0.11 | 0.16 | A>A |
| chr9 | 81100000 | 81200000 | 0.33 | 0.30 | A>A |
| chr9 | 81200000 | 81300000 | 0.37 | 0.30 | A>A |
| chr9 | 81300000 | 81400000 | 0.58 | 0.58 | A>A |
| chr9 | 81400000 | 81500000 | 0.47 | 0.52 | A>A |
| chr9 | 81500000 | 81600000 | 0.50 | 0.50 | A>A |
| chr9 | 81600000 | 81700000 | 0.48 | 0.49 | A>A |
| chr9 | 81700000 | 81800000 | 0.37 | 0.27 | A>A |
| chr9 | 81800000 | 81900000 | 0.38 | 0.24 | A>A |
| chr9 | 81900000 | 82000000 | 0.43 | 0.28 | A>A |
| chr9 | 82000000 | 82100000 | 0.33 | 0.18 | A>A |
| chr9 | 82100000 | 82200000 | 0.17 | 0.13 | A>A |
| chr9 | 82200000 | 82300000 | 0.25 | 0.10 | A>A |
| chr9 | 82300000 | 82400000 | 0.07 | -0.02 | A>A |
| chr9 | 82400000 | 82500000 | -0.03 | -0.05 | B>B |
| chr9 | 82500000 | 82600000 | -0.08 | -0.22 | B>B |
| chr9 | 82600000 | 82700000 | -0.21 | -0.48 | B>B |
| chr9 | 82700000 | 82800000 | -0.29 | -0.50 | B>B |

| | | | | | |
|---|---|---|---|---|---|
| chr9 | 82800000 | 82900000 | -0.38 | -0.69 | B>B |
| chr9 | 82900000 | 83000000 | -0.42 | -0.82 | B>B |
| chr9 | 83000000 | 83100000 | -0.55 | -0.76 | B>B |
| chr9 | 83100000 | 83200000 | -0.37 | -0.64 | B>B |
| chr9 | 83200000 | 83300000 | -0.18 | -0.32 | B>B |
| chr9 | 83300000 | 83400000 | -0.16 | -0.17 | B>B |
| chr9 | 83400000 | 83500000 | -0.05 | -0.03 | B>B |
| chr9 | 83500000 | 83600000 | 0.02 | 0.14 | A>A |
| chr9 | 83600000 | 83700000 | 0.05 | 0.07 | A>A |
| chr9 | 83700000 | 83800000 | 0.12 | 0.12 | A>A |
| chr9 | 83800000 | 83900000 | 0.20 | 0.21 | A>A |
| chr9 | 83900000 | 84000000 | 0.32 | 0.29 | A>A |
| chr9 | 84000000 | 84100000 | 0.40 | 0.41 | A>A |
| chr9 | 84100000 | 84200000 | 0.32 | 0.33 | A>A |
| chr9 | 84200000 | 84300000 | 0.34 | 0.21 | A>A |
| chr9 | 84300000 | 84400000 | 0.37 | 0.15 | A>A |
| chr9 | 84400000 | 84500000 | 0.27 | 0.07 | A>A |
| chr9 | 84500000 | 84600000 | 0.07 | -0.22 | B>B |
| chr9 | 84600000 | 84700000 | -0.06 | -0.37 | B>B |
| chr9 | 84700000 | 84800000 | -0.19 | -0.50 | B>B |
| chr9 | 84800000 | 84900000 | -0.21 | -0.59 | B>B |
| chr9 | 84900000 | 85000000 | -0.26 | -0.54 | B>B |
| chr9 | 85000000 | 85100000 | -0.31 | -0.56 | B>B |
| chr9 | 85100000 | 85200000 | -0.22 | -0.58 | B>B |
| chr9 | 85200000 | 85300000 | -0.26 | -0.53 | B>B |
| chr9 | 85300000 | 85400000 | -0.09 | -0.59 | B>B |
| chr9 | 85400000 | 85500000 | 0.04 | -0.47 | B>B |
| chr9 | 85500000 | 85600000 | 0.09 | -0.16 | B>B |
| chr9 | 85600000 | 85700000 | 0.28 | 0.10 | A>A |
| chr9 | 85700000 | 85800000 | 0.37 | 0.26 | A>A |
| chr9 | 85800000 | 85900000 | 0.34 | 0.17 | A>A |
| chr9 | 85900000 | 86000000 | 0.42 | 0.19 | A>A |
| chr9 | 86000000 | 86100000 | 0.63 | 0.33 | A>A |
| chr9 | 86100000 | 86200000 | 0.49 | 0.21 | A>A |
| chr9 | 86200000 | 86300000 | 0.52 | 0.35 | A>A |
| chr9 | 86300000 | 86400000 | 0.66 | 0.56 | A>A |
| chr9 | 86400000 | 86500000 | 0.00 | 0.00 | NO |
| chr9 | 86500000 | 86600000 | 0.55 | 0.44 | A>A |
| chr9 | 86600000 | 86700000 | 0.57 | 0.30 | A>A |
| chr9 | 86700000 | 86800000 | 0.61 | 0.31 | A>A |
| chr9 | 86800000 | 86900000 | 0.34 | 0.15 | A>A |
| chr9 | 86900000 | 87000000 | 0.05 | 0.06 | A>A |
| chr9 | 87000000 | 87100000 | -0.30 | -0.14 | B>B |
| chr9 | 87100000 | 87200000 | -0.62 | -0.22 | B>B |
| chr9 | 87200000 | 87300000 | -0.61 | -0.23 | B>B |
| chr9 | 87300000 | 87400000 | -0.43 | 0.14 | B>A |
| chr9 | 87400000 | 87500000 | -0.16 | 0.23 | A>A |
| chr9 | 87500000 | 87600000 | -0.18 | 0.27 | A>A |
| chr9 | 87600000 | 87700000 | -0.23 | 0.35 | B>A |
| chr9 | 87700000 | 87800000 | -0.37 | 0.23 | B>A |
| chr9 | 87800000 | 87900000 | -0.44 | 0.06 | B>B |
| chr9 | 87900000 | 88000000 | -0.53 | -0.23 | B>B |
| chr9 | 88000000 | 88100000 | -0.43 | -0.16 | B>B |
| chr9 | 88100000 | 88200000 | -0.08 | -0.09 | B>B |
| chr9 | 88200000 | 88300000 | 0.13 | 0.16 | A>A |
| chr9 | 88300000 | 88400000 | 0.36 | 0.43 | A>A |

| | | | | | |
|---|---|---|---|---|---|
| chr9 | 88400000 | 88500000 | 0.00 | 0.00 | NO |
| chr9 | 88500000 | 88600000 | 0.72 | 0.70 | A>A |
| chr9 | 88600000 | 88700000 | 0.75 | 0.70 | A>A |
| chr9 | 88700000 | 88800000 | 0.87 | 0.95 | A>A |
| chr9 | 88800000 | 88900000 | 0.61 | 0.60 | A>A |
| chr9 | 88900000 | 89000000 | 0.50 | 0.48 | A>A |
| chr9 | 89000000 | 89100000 | 0.06 | -0.02 | A>A |
| chr9 | 89100000 | 89200000 | -0.13 | -0.23 | B>B |
| chr9 | 89200000 | 89300000 | -0.32 | -0.34 | B>B |
| chr9 | 89300000 | 89400000 | -0.39 | -0.36 | B>B |
| chr9 | 89400000 | 89500000 | -0.34 | -0.31 | B>B |
| chr9 | 89500000 | 89600000 | -0.14 | -0.29 | B>B |
| chr9 | 89600000 | 89700000 | -0.11 | -0.20 | B>B |
| chr9 | 89700000 | 89800000 | -0.22 | -0.20 | B>B |
| chr9 | 89800000 | 89900000 | -0.11 | -0.21 | B>B |
| chr9 | 89900000 | 90000000 | 0.26 | 0.21 | A>A |
| chr9 | 90000000 | 90100000 | 0.68 | 0.60 | A>A |
| chr9 | 90100000 | 90200000 | 0.51 | 0.59 | A>A |
| chr9 | 90200000 | 90300000 | 0.39 | 0.53 | A>A |
| chr9 | 90300000 | 90400000 | 0.66 | 0.63 | A>A |
| chr9 | 90400000 | 90500000 | 0.71 | 0.65 | A>A |
| chr9 | 90500000 | 90600000 | 0.67 | 0.44 | A>A |
| chr9 | 90600000 | 90700000 | 0.65 | 0.43 | A>A |
| chr9 | 90700000 | 90800000 | 0.53 | 0.44 | A>A |
| chr9 | 90800000 | 90900000 | 0.80 | 0.64 | A>A |
| chr9 | 90900000 | 91000000 | 0.76 | 0.68 | A>A |
| chr9 | 91000000 | 91100000 | 0.84 | 0.72 | A>A |
| chr9 | 91100000 | 91200000 | 0.65 | 0.48 | A>A |
| chr9 | 91200000 | 91300000 | 0.52 | 0.19 | A>A |
| chr9 | 91300000 | 91400000 | 0.68 | 0.18 | A>A |
| chr9 | 91400000 | 91500000 | 0.81 | 0.18 | A>B |
| chr9 | 91500000 | 91600000 | 0.70 | 0.09 | A>B |
| chr9 | 91600000 | 91700000 | 0.56 | -0.01 | A>A |
| chr9 | 91700000 | 91800000 | 0.60 | -0.03 | A>B |
| chr9 | 91800000 | 91900000 | 0.63 | 0.01 | A>B |
| chr9 | 91900000 | 92000000 | 0.81 | 0.51 | A>A |
| chr9 | 92000000 | 92100000 | 0.84 | 0.57 | A>A |
| chr9 | 92100000 | 92200000 | 0.69 | 0.41 | A>A |
| chr9 | 92200000 | 92300000 | 0.41 | 0.15 | A>A |
| chr9 | 92300000 | 92400000 | 0.00 | 0.00 | NO |
| chr9 | 92400000 | 92500000 | 0.27 | 0.07 | A>A |
| chr9 | 92500000 | 92600000 | 0.00 | 0.00 | NO |
| chr9 | 92600000 | 92700000 | 0.00 | 0.00 | NO |
| chr9 | 92700000 | 92800000 | 0.51 | 0.10 | A>A |
| chr9 | 92800000 | 92900000 | 0.39 | 0.12 | A>A |
| chr9 | 92900000 | 93000000 | 0.06 | 0.18 | A>A |
| chr9 | 93000000 | 93100000 | 0.28 | 0.24 | A>A |
| chr9 | 93100000 | 93200000 | 0.39 | 0.35 | A>A |
| chr9 | 93200000 | 93300000 | 0.25 | 0.21 | A>A |
| chr9 | 93300000 | 93400000 | 0.10 | 0.04 | A>A |
| chr9 | 93400000 | 93500000 | 0.00 | 0.04 | A>A |
| chr9 | 93500000 | 93600000 | 0.18 | 0.10 | A>A |
| chr9 | 93600000 | 93700000 | 0.21 | 0.10 | A>A |
| chr9 | 93700000 | 93800000 | 0.48 | 0.33 | A>A |
| chr9 | 93800000 | 93900000 | 0.66 | 0.57 | A>A |
| chr9 | 93900000 | 94000000 | 0.72 | 0.62 | A>A |

| | | | | | |
|---|---|---|---|---|---|
| chr9 | 94000000 | 94100000 | 0.67 | 0.70 | A>A |
| chr9 | 94100000 | 94200000 | 0.62 | 0.50 | A>A |
| chr9 | 94200000 | 94300000 | 0.51 | 0.23 | A>A |
| chr9 | 94300000 | 94400000 | 0.46 | 0.24 | A>A |
| chr9 | 94400000 | 94500000 | 0.34 | 0.15 | A>A |
| chr9 | 94500000 | 94600000 | 0.55 | 0.26 | A>A |
| chr9 | 94600000 | 94700000 | 0.68 | 0.42 | A>A |
| chr9 | 94700000 | 94800000 | 0.64 | 0.51 | A>A |
| chr9 | 94800000 | 94900000 | 0.62 | 0.64 | A>A |
| chr9 | 94900000 | 95000000 | 0.82 | 0.87 | A>A |
| chr9 | 95000000 | 95100000 | 0.86 | 0.87 | A>A |
| chr9 | 95100000 | 95200000 | 0.83 | 0.84 | A>A |
| chr9 | 95200000 | 95300000 | 0.70 | 0.78 | A>A |
| chr9 | 95300000 | 95400000 | 0.68 | 0.80 | A>A |
| chr9 | 95400000 | 95500000 | 0.78 | 0.90 | A>A |
| chr9 | 95500000 | 95600000 | 0.82 | 1.00 | A>A |
| chr9 | 95600000 | 95700000 | 0.80 | 0.97 | A>A |
| chr9 | 95700000 | 95800000 | 0.87 | 0.97 | A>A |
| chr9 | 95800000 | 95900000 | 0.96 | 1.12 | A>A |
| chr9 | 95900000 | 96000000 | 0.87 | 0.97 | A>A |
| chr9 | 96000000 | 96100000 | 0.87 | 0.93 | A>A |
| chr9 | 96100000 | 96200000 | 0.83 | 0.93 | A>A |
| chr9 | 96200000 | 96300000 | 0.92 | 1.09 | A>A |
| chr9 | 96300000 | 96400000 | 0.70 | 0.88 | A>A |
| chr9 | 96400000 | 96500000 | 0.52 | 0.75 | A>A |
| chr9 | 96500000 | 96600000 | 0.33 | 0.73 | A>A |
| chr9 | 96600000 | 96700000 | 0.57 | 0.78 | A>A |
| chr9 | 96700000 | 96800000 | 0.55 | 0.68 | A>A |
| chr9 | 96800000 | 96900000 | 0.83 | 0.91 | A>A |
| chr9 | 96900000 | 97000000 | 0.75 | 0.91 | A>A |
| chr9 | 97000000 | 97100000 | 0.00 | 0.83 | NO |
| chr9 | 97100000 | 97200000 | 0.00 | 0.00 | NO |
| chr9 | 97200000 | 97300000 | 0.74 | 0.82 | A>A |
| chr9 | 97300000 | 97400000 | 0.58 | 0.68 | A>A |
| chr9 | 97400000 | 97500000 | 0.55 | 0.72 | A>A |
| chr9 | 97500000 | 97600000 | 0.70 | 0.78 | A>A |
| chr9 | 97600000 | 97700000 | 0.92 | 0.88 | A>A |
| chr9 | 97700000 | 97800000 | 0.87 | 0.91 | A>A |
| chr9 | 97800000 | 97900000 | 0.58 | 0.66 | A>A |
| chr9 | 97900000 | 98000000 | 0.62 | 0.68 | A>A |
| chr9 | 98000000 | 98100000 | 0.61 | 0.75 | A>A |
| chr9 | 98100000 | 98200000 | 0.71 | 0.74 | A>A |
| chr9 | 98200000 | 98300000 | 0.58 | 0.81 | A>A |
| chr9 | 98300000 | 98400000 | 0.64 | 0.71 | A>A |
| chr9 | 98400000 | 98500000 | 0.56 | 0.66 | A>A |
| chr9 | 98500000 | 98600000 | 0.64 | 0.70 | A>A |
| chr9 | 98600000 | 98700000 | 0.65 | 0.70 | A>A |
| chr9 | 98700000 | 98800000 | 0.70 | 0.67 | A>A |
| chr9 | 98800000 | 98900000 | 0.69 | 0.83 | A>A |
| chr9 | 98900000 | 99000000 | 0.56 | 0.72 | A>A |
| chr9 | 99000000 | 99100000 | 0.51 | 0.70 | A>A |
| chr9 | 99100000 | 99200000 | 0.80 | 0.96 | A>A |
| chr9 | 99200000 | 99300000 | 0.71 | 0.97 | A>A |
| chr9 | 99300000 | 99400000 | 0.79 | 1.03 | A>A |
| chr9 | 99400000 | 99500000 | 0.50 | 0.85 | A>A |
| chr9 | 99500000 | 99600000 | 0.40 | 0.90 | A>A |

| | | | | | |
|---|---|---|---|---|---|
| chr9 | 99600000 | 99700000 | 0.00 | 0.87 | NO |
| chr9 | 99700000 | 99800000 | 0.00 | 0.00 | NO |
| chr9 | 99800000 | 99900000 | 0.00 | 0.96 | NO |
| chr9 | 99900000 | 100000000 | 0.00 | 0.00 | NO |
| chr9 | 1E+08 | 100100000 | 0.65 | 0.97 | A>A |
| chr9 | 1E+08 | 100200000 | 0.71 | 0.99 | A>A |
| chr9 | 1E+08 | 100300000 | 0.89 | 1.02 | A>A |
| chr9 | 1E+08 | 100400000 | 0.85 | 0.99 | A>A |
| chr9 | 1E+08 | 100500000 | 0.92 | 0.86 | A>A |
| chr9 | 1.01E+08 | 100600000 | 1.00 | 0.86 | A>A |
| chr9 | 1.01E+08 | 100700000 | 0.91 | 0.76 | A>A |
| chr9 | 1.01E+08 | 100800000 | 0.90 | 0.72 | A>A |
| chr9 | 1.01E+08 | 100900000 | 0.82 | 0.67 | A>A |
| chr9 | 1.01E+08 | 101000000 | 0.76 | 0.56 | A>A |
| chr9 | 1.01E+08 | 101100000 | 0.49 | 0.20 | A>A |
| chr9 | 1.01E+08 | 101200000 | 0.19 | -0.21 | B>B |
| chr9 | 1.01E+08 | 101300000 | 0.17 | -0.35 | B>B |
| chr9 | 1.01E+08 | 101400000 | 0.18 | -0.38 | B>B |
| chr9 | 1.01E+08 | 101500000 | 0.29 | -0.25 | A>A |
| chr9 | 1.02E+08 | 101600000 | 0.38 | 0.04 | A>A |
| chr9 | 1.02E+08 | 101700000 | 0.19 | -0.26 | B>B |
| chr9 | 1.02E+08 | 101800000 | 0.19 | -0.29 | B>B |
| chr9 | 1.02E+08 | 101900000 | 0.15 | -0.22 | B>B |
| chr9 | 1.02E+08 | 102000000 | 0.01 | -0.21 | B>B |
| chr9 | 1.02E+08 | 102100000 | -0.23 | -0.63 | B>B |
| chr9 | 1.02E+08 | 102200000 | -0.30 | -0.87 | B>B |
| chr9 | 1.02E+08 | 102300000 | -0.63 | -0.95 | B>B |
| chr9 | 1.02E+08 | 102400000 | -0.72 | -1.03 | B>B |
| chr9 | 1.02E+08 | 102500000 | -0.73 | -1.01 | B>B |
| chr9 | 1.03E+08 | 102600000 | -0.35 | -0.63 | B>B |
| chr9 | 1.03E+08 | 102700000 | -0.06 | -0.21 | B>B |
| chr9 | 1.03E+08 | 102800000 | 0.01 | -0.13 | B>B |
| chr9 | 1.03E+08 | 102900000 | 0.28 | 0.16 | A>A |
| chr9 | 1.03E+08 | 103000000 | 0.14 | 0.07 | A>A |
| chr9 | 1.03E+08 | 103100000 | 0.15 | 0.08 | A>A |
| chr9 | 1.03E+08 | 103200000 | 0.37 | 0.23 | A>A |
| chr9 | 1.03E+08 | 103300000 | 0.25 | 0.00 | A>A |
| chr9 | 1.03E+08 | 103400000 | -0.18 | -0.31 | B>B |
| chr9 | 1.03E+08 | 103500000 | -0.57 | -0.70 | B>B |
| chr9 | 1.04E+08 | 103600000 | -0.86 | -0.88 | B>B |
| chr9 | 1.04E+08 | 103700000 | -0.78 | -0.97 | B>B |
| chr9 | 1.04E+08 | 103800000 | -0.75 | -0.87 | B>B |
| chr9 | 1.04E+08 | 103900000 | -0.62 | -0.75 | B>B |
| chr9 | 1.04E+08 | 104000000 | -0.59 | -0.76 | B>B |
| chr9 | 1.04E+08 | 104100000 | -0.22 | -0.45 | B>B |
| chr9 | 1.04E+08 | 104200000 | 0.09 | -0.12 | B>B |
| chr9 | 1.04E+08 | 104300000 | -0.01 | -0.34 | B>B |
| chr9 | 1.04E+08 | 104400000 | -0.34 | -0.52 | B>B |
| chr9 | 1.04E+08 | 104500000 | -0.63 | -0.84 | B>B |
| chr9 | 1.05E+08 | 104600000 | -0.77 | -0.90 | B>B |
| chr9 | 1.05E+08 | 104700000 | -0.86 | -0.96 | B>B |
| chr9 | 1.05E+08 | 104800000 | -0.80 | -0.93 | B>B |
| chr9 | 1.05E+08 | 104900000 | -0.96 | -0.95 | B>B |
| chr9 | 1.05E+08 | 105000000 | -0.84 | -0.95 | B>B |
| chr9 | 1.05E+08 | 105100000 | -0.89 | -0.93 | B>B |
| chr9 | 1.05E+08 | 105200000 | -0.84 | -0.94 | B>B |

| | | | | | |
|---|---|---|---|---|---|
| chr9 | 1.05E+08 | 105300000 | -0.78 | -0.90 | B>B |
| chr9 | 1.05E+08 | 105400000 | -0.78 | -0.88 | B>B |
| chr9 | 1.05E+08 | 105500000 | -0.83 | -0.88 | B>B |
| chr9 | 1.06E+08 | 105600000 | -0.82 | -0.92 | B>B |
| chr9 | 1.06E+08 | 105700000 | -0.77 | -0.87 | B>B |
| chr9 | 1.06E+08 | 105800000 | -0.75 | -0.89 | B>B |
| chr9 | 1.06E+08 | 105900000 | -0.78 | -0.89 | B>B |
| chr9 | 1.06E+08 | 106000000 | -0.77 | -0.86 | B>B |
| chr9 | 1.06E+08 | 106100000 | -0.79 | -0.81 | B>B |
| chr9 | 1.06E+08 | 106200000 | -0.76 | -0.84 | B>B |
| chr9 | 1.06E+08 | 106300000 | -0.80 | -0.83 | B>B |
| chr9 | 1.06E+08 | 106400000 | -0.74 | -0.77 | B>B |
| chr9 | 1.06E+08 | 106500000 | -0.76 | -0.74 | B>B |
| chr9 | 1.07E+08 | 106600000 | -0.72 | -0.69 | B>B |
| chr9 | 1.07E+08 | 106700000 | -0.72 | -0.70 | B>B |
| chr9 | 1.07E+08 | 106800000 | -0.75 | -0.74 | B>B |
| chr9 | 1.07E+08 | 106900000 | -0.50 | -0.48 | B>B |
| chr9 | 1.07E+08 | 107000000 | -0.51 | -0.39 | B>B |
| chr9 | 1.07E+08 | 107100000 | -0.69 | -0.33 | B>B |
| chr9 | 1.07E+08 | 107200000 | -0.78 | -0.54 | B>B |
| chr9 | 1.07E+08 | 107300000 | -0.82 | -0.90 | B>B |
| chr9 | 1.07E+08 | 107400000 | -0.86 | -0.90 | B>B |
| chr9 | 1.07E+08 | 107500000 | -0.61 | -0.71 | B>B |
| chr9 | 1.08E+08 | 107600000 | 0.06 | 0.07 | A>A |
| chr9 | 1.08E+08 | 107700000 | 0.37 | 0.36 | A>A |
| chr9 | 1.08E+08 | 107800000 | 0.52 | 0.49 | A>A |
| chr9 | 1.08E+08 | 107900000 | 0.62 | 0.49 | A>A |
| chr9 | 1.08E+08 | 108000000 | 0.39 | 0.37 | A>A |
| chr9 | 1.08E+08 | 108100000 | 0.45 | 0.50 | A>A |
| chr9 | 1.08E+08 | 108200000 | 0.29 | 0.33 | A>A |
| chr9 | 1.08E+08 | 108300000 | 0.18 | 0.15 | A>A |
| chr9 | 1.08E+08 | 108400000 | -0.05 | -0.10 | B>B |
| chr9 | 1.08E+08 | 108500000 | -0.10 | -0.20 | B>B |
| chr9 | 1.09E+08 | 108600000 | -0.47 | -0.53 | B>B |
| chr9 | 1.09E+08 | 108700000 | -0.34 | -0.68 | B>B |
| chr9 | 1.09E+08 | 108800000 | -0.47 | -0.96 | B>B |
| chr9 | 1.09E+08 | 108900000 | -0.75 | -0.94 | B>B |
| chr9 | 1.09E+08 | 109000000 | -0.86 | -0.98 | B>B |
| chr9 | 1.09E+08 | 109100000 | -0.83 | -1.03 | B>B |
| chr9 | 1.09E+08 | 109200000 | -0.77 | -0.97 | B>B |
| chr9 | 1.09E+08 | 109300000 | -0.79 | -0.95 | B>B |
| chr9 | 1.09E+08 | 109400000 | -0.77 | -0.92 | B>B |
| chr9 | 1.09E+08 | 109500000 | -0.66 | -0.92 | B>B |
| chr9 | 1.1E+08 | 109600000 | -0.55 | -0.84 | B>B |
| chr9 | 1.1E+08 | 109700000 | -0.11 | -0.41 | B>B |
| chr9 | 1.1E+08 | 109800000 | -0.23 | -0.47 | B>B |
| chr9 | 1.1E+08 | 109900000 | -0.27 | -0.69 | B>B |
| chr9 | 1.1E+08 | 110000000 | 0.01 | -0.32 | B>B |
| chr9 | 1.1E+08 | 110100000 | 0.35 | 0.09 | A>A |
| chr9 | 1.1E+08 | 110200000 | 0.30 | 0.08 | A>A |
| chr9 | 1.1E+08 | 110300000 | 0.39 | 0.30 | A>A |
| chr9 | 1.1E+08 | 110400000 | 0.54 | 0.45 | A>A |
| chr9 | 1.1E+08 | 110500000 | 0.55 | 0.39 | A>A |
| chr9 | 1.11E+08 | 110600000 | 0.47 | 0.40 | A>A |
| chr9 | 1.11E+08 | 110700000 | 0.42 | 0.35 | A>A |
| chr9 | 1.11E+08 | 110800000 | 0.51 | 0.30 | A>A |

| | | | | | |
|---|---|---|---|---|---|
| chr9 | 1.11E+08 | 110900000 | 0.44 | 0.35 | A>A |
| chr9 | 1.11E+08 | 111000000 | 0.38 | 0.30 | A>A |
| chr9 | 1.11E+08 | 111100000 | 0.27 | 0.26 | A>A |
| chr9 | 1.11E+08 | 111200000 | 0.38 | 0.22 | A>A |
| chr9 | 1.11E+08 | 111300000 | 0.37 | 0.20 | A>A |
| chr9 | 1.11E+08 | 111400000 | 0.29 | 0.15 | A>A |
| chr9 | 1.11E+08 | 111500000 | 0.24 | 0.11 | A>A |
| chr9 | 1.12E+08 | 111600000 | 0.20 | 0.11 | A>A |
| chr9 | 1.12E+08 | 111700000 | 0.48 | 0.33 | A>A |
| chr9 | 1.12E+08 | 111800000 | 0.66 | 0.46 | A>A |
| chr9 | 1.12E+08 | 111900000 | 0.70 | 0.59 | A>A |
| chr9 | 1.12E+08 | 112000000 | 0.57 | 0.47 | A>A |
| chr9 | 1.12E+08 | 112100000 | 0.70 | 0.54 | A>A |
| chr9 | 1.12E+08 | 112200000 | 0.45 | 0.35 | A>A |
| chr9 | 1.12E+08 | 112300000 | 0.48 | 0.27 | A>A |
| chr9 | 1.12E+08 | 112400000 | 0.10 | -0.20 | B>B |
| chr9 | 1.12E+08 | 112500000 | -0.03 | -0.38 | B>B |
| chr9 | 1.13E+08 | 112600000 | 0.03 | -0.33 | B>B |
| chr9 | 1.13E+08 | 112700000 | 0.20 | -0.01 | A>A |
| chr9 | 1.13E+08 | 112800000 | 0.40 | 0.26 | A>A |
| chr9 | 1.13E+08 | 112900000 | 0.41 | 0.20 | A>A |
| chr9 | 1.13E+08 | 113000000 | 0.41 | 0.26 | A>A |
| chr9 | 1.13E+08 | 113100000 | 0.52 | 0.34 | A>A |
| chr9 | 1.13E+08 | 113200000 | 0.38 | 0.24 | A>A |
| chr9 | 1.13E+08 | 113300000 | 0.30 | 0.13 | A>A |
| chr9 | 1.13E+08 | 113400000 | 0.38 | 0.21 | A>A |
| chr9 | 1.13E+08 | 113500000 | 0.41 | 0.43 | A>A |
| chr9 | 1.14E+08 | 113600000 | 0.46 | 0.55 | A>A |
| chr9 | 1.14E+08 | 113700000 | 0.49 | 0.54 | A>A |
| chr9 | 1.14E+08 | 113800000 | 0.54 | 0.52 | A>A |
| chr9 | 1.14E+08 | 113900000 | 0.50 | 0.43 | A>A |
| chr9 | 1.14E+08 | 114000000 | 0.43 | 0.42 | A>A |
| chr9 | 1.14E+08 | 114100000 | 0.40 | 0.36 | A>A |
| chr9 | 1.14E+08 | 114200000 | 0.58 | 0.54 | A>A |
| chr9 | 1.14E+08 | 114300000 | 0.65 | 0.85 | A>A |
| chr9 | 1.14E+08 | 114400000 | 0.68 | 0.84 | A>A |
| chr9 | 1.14E+08 | 114500000 | 0.65 | 0.85 | A>A |
| chr9 | 1.15E+08 | 114600000 | 0.62 | 0.74 | A>A |
| chr9 | 1.15E+08 | 114700000 | 0.77 | 0.96 | A>A |
| chr9 | 1.15E+08 | 114800000 | 0.71 | 0.81 | A>A |
| chr9 | 1.15E+08 | 114900000 | 0.44 | 0.63 | A>A |
| chr9 | 1.15E+08 | 115000000 | 0.73 | 0.85 | A>A |
| chr9 | 1.15E+08 | 115100000 | 0.81 | 0.82 | A>A |
| chr9 | 1.15E+08 | 115200000 | 0.54 | 0.66 | A>A |
| chr9 | 1.15E+08 | 115300000 | 0.65 | 0.83 | A>A |
| chr9 | 1.15E+08 | 115400000 | 0.69 | 0.88 | A>A |
| chr9 | 1.15E+08 | 115500000 | 0.55 | 0.78 | A>A |
| chr9 | 1.16E+08 | 115600000 | 0.45 | 0.81 | A>A |
| chr9 | 1.16E+08 | 115700000 | 0.24 | 0.85 | B>A |
| chr9 | 1.16E+08 | 115800000 | 0.03 | 0.89 | B>A |
| chr9 | 1.16E+08 | 115900000 | 0.30 | 0.87 | B>A |
| chr9 | 1.16E+08 | 116000000 | 0.63 | 0.98 | A>A |
| chr9 | 1.16E+08 | 116100000 | 0.72 | 1.05 | A>A |
| chr9 | 1.16E+08 | 116200000 | 0.85 | 0.94 | A>A |
| chr9 | 1.16E+08 | 116300000 | 0.94 | 0.97 | A>A |
| chr9 | 1.16E+08 | 116400000 | 0.82 | 0.95 | A>A |

| | | | | | |
|---|---|---|---|---|---|
| chr9 | 1.16E+08 | 116500000 | 0.61 | 0.71 | A>A |
| chr9 | 1.17E+08 | 116600000 | 0.73 | 0.83 | A>A |
| chr9 | 1.17E+08 | 116700000 | 0.84 | 0.85 | A>A |
| chr9 | 1.17E+08 | 116800000 | 0.70 | 0.94 | A>A |
| chr9 | 1.17E+08 | 116900000 | 0.70 | 0.77 | A>A |
| chr9 | 1.17E+08 | 117000000 | 0.81 | 0.92 | A>A |
| chr9 | 1.17E+08 | 117100000 | 0.64 | 0.77 | A>A |
| chr9 | 1.17E+08 | 117200000 | 0.63 | 0.76 | A>A |
| chr9 | 1.17E+08 | 117300000 | 0.73 | 0.74 | A>A |
| chr9 | 1.17E+08 | 117400000 | 0.74 | 0.77 | A>A |
| chr9 | 1.17E+08 | 117500000 | 0.46 | 0.59 | A>A |
| chr9 | 1.18E+08 | 117600000 | 0.72 | 0.81 | A>A |
| chr9 | 1.18E+08 | 117700000 | 0.72 | 0.67 | A>A |
| chr9 | 1.18E+08 | 117800000 | 0.40 | 0.61 | A>A |
| chr9 | 1.18E+08 | 117900000 | 0.56 | 0.63 | A>A |
| chr9 | 1.18E+08 | 118000000 | 0.56 | 0.56 | A>A |
| chr9 | 1.18E+08 | 118100000 | 0.57 | 0.75 | A>A |
| chr9 | 1.18E+08 | 118200000 | 0.62 | 0.69 | A>A |
| chr9 | 1.18E+08 | 118300000 | 0.45 | 0.61 | A>A |
| chr9 | 1.18E+08 | 118400000 | 0.21 | 0.30 | A>A |
| chr9 | 1.18E+08 | 118500000 | 0.40 | 0.30 | A>A |
| chr9 | 1.19E+08 | 118600000 | 0.56 | 0.29 | A>A |
| chr9 | 1.19E+08 | 118700000 | 0.51 | 0.30 | A>A |
| chr9 | 1.19E+08 | 118800000 | 0.53 | 0.57 | A>A |
| chr9 | 1.19E+08 | 118900000 | 0.64 | 0.52 | A>A |
| chr9 | 1.19E+08 | 119000000 | 0.34 | 0.39 | A>A |
| chr9 | 1.19E+08 | 119100000 | 0.29 | 0.28 | A>A |
| chr9 | 1.19E+08 | 119200000 | 0.29 | 0.32 | A>A |
| chr9 | 1.19E+08 | 119300000 | 0.36 | 0.27 | A>A |
| chr9 | 1.19E+08 | 119400000 | 0.33 | 0.30 | A>A |
| chr9 | 1.19E+08 | 119500000 | 0.12 | 0.12 | A>A |
| chr9 | 1.2E+08 | 119600000 | -0.25 | -0.08 | B>B |
| chr9 | 1.2E+08 | 119700000 | -0.28 | 0.07 | B>B |
| chr9 | 1.2E+08 | 119800000 | -0.41 | -0.04 | B>B |
| chr9 | 1.2E+08 | 119900000 | -0.53 | -0.29 | B>B |
| chr9 | 1.2E+08 | 120000000 | -0.72 | -0.66 | B>B |
| chr9 | 1.2E+08 | 120100000 | -0.81 | -0.88 | B>B |
| chr9 | 1.2E+08 | 120200000 | -0.71 | -0.82 | B>B |
| chr9 | 1.2E+08 | 120300000 | -0.73 | -0.73 | B>B |
| chr9 | 1.2E+08 | 120400000 | -0.76 | -0.86 | B>B |
| chr9 | 1.2E+08 | 120500000 | -0.58 | -0.73 | B>B |
| chr9 | 1.21E+08 | 120600000 | -0.70 | -0.88 | B>B |
| chr9 | 1.21E+08 | 120700000 | -0.81 | -1.01 | B>B |
| chr9 | 1.21E+08 | 120800000 | -0.90 | -1.00 | B>B |
| chr9 | 1.21E+08 | 120900000 | -0.84 | -0.98 | B>B |
| chr9 | 1.21E+08 | 121000000 | -0.73 | -0.71 | B>B |
| chr9 | 1.21E+08 | 121100000 | -0.60 | -0.50 | B>B |
| chr9 | 1.21E+08 | 121200000 | -0.63 | -0.47 | B>B |
| chr9 | 1.21E+08 | 121300000 | -0.53 | -0.45 | B>B |
| chr9 | 1.21E+08 | 121400000 | -0.41 | -0.17 | B>B |
| chr9 | 1.21E+08 | 121500000 | -0.45 | -0.18 | B>B |
| chr9 | 1.22E+08 | 121600000 | -0.60 | -0.30 | B>B |
| chr9 | 1.22E+08 | 121700000 | -0.67 | -0.55 | B>B |
| chr9 | 1.22E+08 | 121800000 | -0.83 | -0.60 | B>B |
| chr9 | 1.22E+08 | 121900000 | -0.77 | -0.76 | B>B |
| chr9 | 1.22E+08 | 122000000 | -0.75 | -0.92 | B>B |

| | | | | | |
|---|---|---|---|---|---|
| chr9 | 1.22E+08 | 122100000 | -0.69 | -0.86 | B>B |
| chr9 | 1.22E+08 | 122200000 | -0.70 | -0.80 | B>B |
| chr9 | 1.22E+08 | 122300000 | -0.78 | -0.93 | B>B |
| chr9 | 1.22E+08 | 122400000 | -0.72 | -1.01 | B>B |
| chr9 | 1.22E+08 | 122500000 | -0.77 | -1.03 | B>B |
| chr9 | 1.23E+08 | 122600000 | -0.70 | -1.05 | B>B |
| chr9 | 1.23E+08 | 122700000 | -0.58 | -1.02 | B>B |
| chr9 | 1.23E+08 | 122800000 | -0.40 | -0.83 | B>B |
| chr9 | 1.23E+08 | 122900000 | -0.54 | -1.04 | B>B |
| chr9 | 1.23E+08 | 123000000 | -0.47 | -1.08 | A>B |
| chr9 | 1.23E+08 | 123100000 | -0.36 | -0.93 | B>B |
| chr9 | 1.23E+08 | 123200000 | 0.12 | -0.25 | B>B |
| chr9 | 1.23E+08 | 123300000 | 0.25 | -0.01 | A>A |
| chr9 | 1.23E+08 | 123400000 | 0.60 | 0.35 | A>A |
| chr9 | 1.23E+08 | 123500000 | 0.77 | 0.70 | A>A |
| chr9 | 1.24E+08 | 123600000 | 0.86 | 0.74 | A>A |
| chr9 | 1.24E+08 | 123700000 | 0.92 | 0.84 | A>A |
| chr9 | 1.24E+08 | 123800000 | 0.88 | 0.83 | A>A |
| chr9 | 1.24E+08 | 123900000 | 0.85 | 0.89 | A>A |
| chr9 | 1.24E+08 | 124000000 | 0.88 | 0.93 | A>A |
| chr9 | 1.24E+08 | 124100000 | 1.10 | 1.04 | A>A |
| chr9 | 1.24E+08 | 124200000 | 0.90 | 0.87 | A>A |
| chr9 | 1.24E+08 | 124300000 | 0.85 | 0.83 | A>A |
| chr9 | 1.24E+08 | 124400000 | 0.77 | 0.93 | A>A |
| chr9 | 1.24E+08 | 124500000 | 0.84 | 0.84 | A>A |
| chr9 | 1.25E+08 | 124600000 | 0.68 | 0.96 | A>A |
| chr9 | 1.25E+08 | 124700000 | 0.74 | 0.92 | A>A |
| chr9 | 1.25E+08 | 124800000 | 0.66 | 0.74 | A>A |
| chr9 | 1.25E+08 | 124900000 | 0.43 | 0.74 | A>A |
| chr9 | 1.25E+08 | 125000000 | 0.70 | 0.91 | A>A |
| chr9 | 1.25E+08 | 125100000 | 0.57 | 0.79 | A>A |
| chr9 | 1.25E+08 | 125200000 | 0.38 | 0.72 | A>A |
| chr9 | 1.25E+08 | 125300000 | 0.01 | 0.52 | B>A |
| chr9 | 1.25E+08 | 125400000 | -0.28 | 0.40 | B>A |
| chr9 | 1.25E+08 | 125500000 | -0.37 | 0.35 | B>A |
| chr9 | 1.26E+08 | 125600000 | 0.00 | 0.45 | A>A |
| chr9 | 1.26E+08 | 125700000 | 0.68 | 0.85 | A>A |
| chr9 | 1.26E+08 | 125800000 | 0.62 | 0.88 | A>A |
| chr9 | 1.26E+08 | 125900000 | 0.60 | 0.76 | A>A |
| chr9 | 1.26E+08 | 126000000 | 0.61 | 0.80 | A>A |
| chr9 | 1.26E+08 | 126100000 | 0.64 | 0.84 | A>A |
| chr9 | 1.26E+08 | 126200000 | 0.52 | 0.70 | A>A |
| chr9 | 1.26E+08 | 126300000 | 0.61 | 0.68 | A>A |
| chr9 | 1.26E+08 | 126400000 | 0.58 | 0.71 | A>A |
| chr9 | 1.26E+08 | 126500000 | 0.66 | 0.75 | A>A |
| chr9 | 1.27E+08 | 126600000 | 0.62 | 0.79 | A>A |
| chr9 | 1.27E+08 | 126700000 | 0.73 | 0.80 | A>A |
| chr9 | 1.27E+08 | 126800000 | 0.61 | 0.68 | A>A |
| chr9 | 1.27E+08 | 126900000 | 0.57 | 0.60 | A>A |
| chr9 | 1.27E+08 | 127000000 | 0.71 | 0.82 | A>A |
| chr9 | 1.27E+08 | 127100000 | 0.73 | 0.83 | A>A |
| chr9 | 1.27E+08 | 127200000 | 0.72 | 0.79 | A>A |
| chr9 | 1.27E+08 | 127300000 | 0.83 | 0.87 | A>A |
| chr9 | 1.27E+08 | 127400000 | 0.80 | 0.86 | A>A |
| chr9 | 1.27E+08 | 127500000 | 0.67 | 0.86 | A>A |
| chr9 | 1.28E+08 | 127600000 | 0.84 | 0.90 | A>A |

| | | | | | |
|---|---|---|---|---|---|
| chr9 | 1.28E+08 | 127700000 | 0.80 | 0.99 | A>A |
| chr9 | 1.28E+08 | 127800000 | 0.81 | 0.86 | A>A |
| chr9 | 1.28E+08 | 127900000 | 0.77 | 0.81 | A>A |
| chr9 | 1.28E+08 | 128000000 | 0.82 | 0.88 | A>A |
| chr9 | 1.28E+08 | 128100000 | 0.89 | 0.90 | A>A |
| chr9 | 1.28E+08 | 128200000 | 0.58 | 0.68 | A>A |
| chr9 | 1.28E+08 | 128300000 | 0.64 | 0.76 | A>A |
| chr9 | 1.28E+08 | 128400000 | 0.60 | 0.73 | A>A |
| chr9 | 1.28E+08 | 128500000 | 0.67 | 0.74 | A>A |
| chr9 | 1.29E+08 | 128600000 | 0.57 | 0.79 | A>A |
| chr9 | 1.29E+08 | 128700000 | 0.56 | 0.69 | A>A |
| chr9 | 1.29E+08 | 128800000 | 0.48 | 0.65 | A>A |
| chr9 | 1.29E+08 | 128900000 | 0.44 | 0.57 | A>A |
| chr9 | 1.29E+08 | 129000000 | 0.35 | 0.59 | A>A |
| chr9 | 1.29E+08 | 129100000 | 0.51 | 0.71 | A>A |
| chr9 | 1.29E+08 | 129200000 | 0.64 | 0.75 | A>A |
| chr9 | 1.29E+08 | 129300000 | 0.66 | 0.77 | A>A |
| chr9 | 1.29E+08 | 129400000 | 0.66 | 0.80 | A>A |
| chr9 | 1.29E+08 | 129500000 | 0.83 | 0.89 | A>A |
| chr9 | 1.3E+08 | 129600000 | 0.73 | 0.78 | A>A |
| chr9 | 1.3E+08 | 129700000 | 0.68 | 0.85 | A>A |
| chr9 | 1.3E+08 | 129800000 | 0.65 | 0.84 | A>A |
| chr9 | 1.3E+08 | 129900000 | 0.86 | 0.91 | A>A |
| chr9 | 1.3E+08 | 130000000 | 0.64 | 0.75 | A>A |
| chr9 | 1.3E+08 | 130100000 | 0.75 | 0.77 | A>A |
| chr9 | 1.3E+08 | 130200000 | 0.74 | 0.81 | A>A |
| chr9 | 1.3E+08 | 130300000 | 0.72 | 0.68 | A>A |
| chr9 | 1.3E+08 | 130400000 | 0.67 | 0.78 | A>A |
| chr9 | 1.3E+08 | 130500000 | 0.64 | 0.73 | A>A |
| chr9 | 1.31E+08 | 130600000 | 0.67 | 0.64 | A>A |
| chr9 | 1.31E+08 | 130700000 | 0.61 | 0.65 | A>A |
| chr9 | 1.31E+08 | 130800000 | 0.77 | 0.79 | A>A |
| chr9 | 1.31E+08 | 130900000 | 0.76 | 0.61 | A>A |
| chr9 | 1.31E+08 | 131000000 | 0.71 | 0.59 | A>A |
| chr9 | 1.31E+08 | 131100000 | 0.61 | 0.55 | A>A |
| chr9 | 1.31E+08 | 131200000 | 0.55 | 0.58 | A>A |
| chr9 | 1.31E+08 | 131300000 | 0.62 | 0.65 | A>A |
| chr9 | 1.31E+08 | 131400000 | 0.65 | 0.60 | A>A |
| chr9 | 1.31E+08 | 131500000 | 0.62 | 0.58 | A>A |
| chr9 | 1.32E+08 | 131600000 | 0.57 | 0.53 | A>A |
| chr9 | 1.32E+08 | 131700000 | 0.52 | 0.56 | A>A |
| chr9 | 1.32E+08 | 131800000 | 0.63 | 0.56 | A>A |
| chr9 | 1.32E+08 | 131900000 | 0.49 | 0.58 | A>A |
| chr9 | 1.32E+08 | 132000000 | 0.57 | 0.60 | A>A |
| chr9 | 1.32E+08 | 132100000 | 0.83 | 0.73 | A>A |
| chr9 | 1.32E+08 | 132200000 | 0.64 | 0.64 | A>A |
| chr9 | 1.32E+08 | 132300000 | 0.62 | 0.64 | A>A |
| chr9 | 1.32E+08 | 132400000 | 0.57 | 0.60 | A>A |
| chr9 | 1.32E+08 | 132500000 | 0.61 | 0.66 | A>A |
| chr9 | 1.33E+08 | 132600000 | 0.53 | 0.63 | A>A |
| chr9 | 1.33E+08 | 132700000 | 0.68 | 0.63 | A>A |
| chr9 | 1.33E+08 | 132800000 | 0.78 | 0.88 | A>A |
| chr9 | 1.33E+08 | 132900000 | 0.57 | 0.64 | A>A |
| chr9 | 1.33E+08 | 133000000 | 0.69 | 0.74 | A>A |
| chr9 | 1.33E+08 | 133100000 | 0.54 | 0.67 | A>A |
| chr9 | 1.33E+08 | 133200000 | 0.00 | 0.00 | NO |

| | | | | | |
|---|---|---|---|---|---|
| chr9 | 1.33E+08 | 133300000 | 0.65 | 0.70 | A>A |
| chr9 | 1.33E+08 | 133400000 | 0.67 | 0.75 | A>A |
| chr9 | 1.33E+08 | 133500000 | 0.70 | 0.90 | A>A |
| chr9 | 1.34E+08 | 133600000 | 0.64 | 0.68 | A>A |
| chr9 | 1.34E+08 | 133700000 | 0.87 | 0.80 | A>A |
| chr9 | 1.34E+08 | 133800000 | 0.74 | 0.80 | A>A |
| chr9 | 1.34E+08 | 133900000 | 0.68 | 0.80 | A>A |
| chr9 | 1.34E+08 | 134000000 | 0.65 | 0.72 | A>A |
| chr9 | 1.34E+08 | 134100000 | 0.75 | 0.76 | A>A |
| chr9 | 1.34E+08 | 134200000 | 0.77 | 0.78 | A>A |
| chr9 | 1.34E+08 | 134300000 | 0.66 | 0.75 | A>A |
| chr9 | 1.34E+08 | 134400000 | 0.62 | 0.75 | A>A |
| chr9 | 1.34E+08 | 134500000 | 0.61 | 0.69 | A>A |
| chr9 | 1.35E+08 | 134600000 | 0.74 | 0.88 | A>A |
| chr9 | 1.35E+08 | 134700000 | 0.45 | 0.68 | A>A |
| chr9 | 1.35E+08 | 134800000 | 0.67 | 0.74 | A>A |
| chr9 | 1.35E+08 | 134900000 | 0.67 | 0.82 | A>A |
| chr9 | 1.35E+08 | 135000000 | 0.63 | 0.80 | A>A |
| chr9 | 1.35E+08 | 135100000 | 0.67 | 0.73 | A>A |
| chr9 | 1.35E+08 | 135200000 | 0.57 | 0.77 | A>A |
| chr9 | 1.35E+08 | 135300000 | 0.74 | 0.77 | A>A |
| chr9 | 1.35E+08 | 135400000 | 0.67 | 0.84 | A>A |
| chr9 | 1.35E+08 | 135500000 | 0.58 | 0.78 | A>A |
| chr9 | 1.36E+08 | 135600000 | 0.75 | 0.89 | A>A |
| chr9 | 1.36E+08 | 135700000 | 0.61 | 0.88 | A>A |
| chr9 | 1.36E+08 | 135800000 | 0.71 | 0.88 | A>A |
| chr9 | 1.36E+08 | 135900000 | 0.68 | 0.81 | A>A |
| chr9 | 1.36E+08 | 136000000 | 0.84 | 0.90 | A>A |
| chr9 | 1.36E+08 | 136100000 | 0.73 | 0.78 | A>A |
| chr9 | 1.36E+08 | 136200000 | 0.65 | 0.67 | A>A |
| chr9 | 1.36E+08 | 136300000 | 0.78 | 0.75 | A>A |
| chr9 | 1.36E+08 | 136400000 | 0.68 | 0.79 | A>A |
| chr9 | 1.36E+08 | 136500000 | 0.67 | 0.82 | A>A |
| chr9 | 1.37E+08 | 136600000 | 0.71 | 0.76 | A>A |
| chr9 | 1.37E+08 | 136700000 | 0.62 | 0.71 | A>A |
| chr9 | 1.37E+08 | 136800000 | 0.86 | 0.89 | A>A |
| chr9 | 1.37E+08 | 136900000 | 0.67 | 0.80 | A>A |
| chr9 | 1.37E+08 | 137000000 | 0.70 | 0.77 | A>A |
| chr9 | 1.37E+08 | 137100000 | 0.00 | 0.67 | NO |
| chr9 | 1.37E+08 | 137200000 | 0.80 | 0.86 | A>A |
| chr9 | 1.37E+08 | 137300000 | 0.88 | 0.90 | A>A |
| chr9 | 1.37E+08 | 137400000 | 0.64 | 0.66 | A>A |
| chr9 | 1.37E+08 | 137500000 | 0.70 | 0.77 | A>A |
| chr9 | 1.38E+08 | 137600000 | 0.61 | 0.75 | A>A |
| chr9 | 1.38E+08 | 137700000 | 0.54 | 0.59 | A>A |
| chr9 | 1.38E+08 | 137800000 | 0.58 | 0.72 | A>A |
| chr9 | 1.38E+08 | 137900000 | 0.53 | 0.69 | A>A |
| chr9 | 1.38E+08 | 138000000 | 0.37 | 0.47 | A>A |
| chr9 | 1.38E+08 | 138100000 | 0.57 | 0.68 | A>A |
| chr9 | 1.38E+08 | 138200000 | 0.60 | 0.82 | A>A |
| chr9 | 1.38E+08 | 138300000 | 0.53 | 0.59 | A>A |
| chr9 | 1.38E+08 | 138400000 | 0.55 | 0.69 | A>A |
| chr9 | 1.38E+08 | 138500000 | 0.61 | 0.74 | A>A |
| chr9 | 1.39E+08 | 138600000 | 0.65 | 0.80 | A>A |
| chr9 | 1.39E+08 | 138700000 | 0.68 | 0.81 | A>A |
| chr9 | 1.39E+08 | 138800000 | 0.65 | 0.86 | A>A |

| | | | | | |
|---|---|---|---|---|---|
| chr9 | 1.39E+08 | 138900000 | 0.71 | 0.77 | A>A |
| chr9 | 1.39E+08 | 139000000 | 0.81 | 0.91 | A>A |
| chr9 | 1.39E+08 | 139100000 | 0.66 | 0.77 | A>A |
| chr9 | 1.39E+08 | 139200000 | 0.69 | 0.80 | A>A |
| chr9 | 1.39E+08 | 139300000 | 0.60 | 0.63 | A>A |
| chr9 | 1.39E+08 | 139400000 | 0.56 | 0.63 | A>A |
| chr9 | 1.39E+08 | 139500000 | 0.79 | 0.92 | A>A |
| chr9 | 1.4E+08 | 139600000 | 0.60 | 0.67 | A>A |
| chr9 | 1.4E+08 | 139700000 | 0.59 | 0.72 | A>A |
| chr9 | 1.4E+08 | 139800000 | 0.73 | 0.75 | A>A |
| chr9 | 1.4E+08 | 139900000 | 0.73 | 0.80 | A>A |
| chr9 | 1.4E+08 | 140000000 | 0.55 | 0.73 | A>A |
| chr9 | 1.4E+08 | 140100000 | 0.66 | 0.78 | A>A |
| chr9 | 1.4E+08 | 140200000 | 0.70 | 0.67 | A>A |
| chr9 | 1.4E+08 | 140300000 | 0.90 | 0.91 | A>A |
| chr9 | 1.4E+08 | 140400000 | 0.75 | 0.85 | A>A |
| chr9 | 1.4E+08 | 140500000 | 0.77 | 0.93 | A>A |
| chr9 | 1.41E+08 | 140600000 | 0.83 | 0.97 | A>A |
| chr9 | 1.41E+08 | 140700000 | 0.80 | 0.85 | A>A |
| chr9 | 1.41E+08 | 140800000 | 0.58 | 0.59 | A>A |
| chr9 | 1.41E+08 | 140900000 | 0.62 | 0.57 | A>A |
| chr9 | 1.41E+08 | 141000000 | 0.51 | 0.53 | A>A |
| chr9 | 1.41E+08 | 141100000 | 0.00 | 0.00 | NO |
| chr9 | 1.41E+08 | 141200000 | 0.00 | 0.00 | NO |
| chr9 | 1.41E+08 | 141213431 | 0.00 | 0.00 | NO |
| chr10 | 0 | 100000 | 0.00 | 0.00 | NO |
| chr10 | 100000 | 200000 | 1.18 | 1.14 | A>A |
| chr10 | 200000 | 300000 | 1.00 | 1.27 | A>A |
| chr10 | 300000 | 400000 | 1.13 | 1.33 | A>A |
| chr10 | 400000 | 500000 | 1.03 | 1.20 | A>A |
| chr10 | 500000 | 600000 | 0.98 | 1.17 | A>A |
| chr10 | 600000 | 700000 | 1.10 | 1.37 | A>A |
| chr10 | 700000 | 800000 | 1.02 | 1.31 | A>A |
| chr10 | 800000 | 900000 | 1.01 | 1.15 | A>A |
| chr10 | 900000 | 1000000 | 1.23 | 1.31 | A>A |
| chr10 | 1000000 | 1100000 | 1.28 | 1.27 | A>A |
| chr10 | 1100000 | 1200000 | 0.55 | 0.44 | A>A |
| chr10 | 1200000 | 1300000 | 0.34 | -0.08 | A>A |
| chr10 | 1300000 | 1400000 | 0.14 | -0.22 | B>B |
| chr10 | 1400000 | 1500000 | -0.08 | -0.29 | B>B |
| chr10 | 1500000 | 1600000 | -0.19 | -0.28 | B>B |
| chr10 | 1600000 | 1700000 | -0.28 | -0.47 | B>B |
| chr10 | 1700000 | 1800000 | -0.43 | -0.50 | B>B |
| chr10 | 1800000 | 1900000 | -0.63 | -0.84 | B>B |
| chr10 | 1900000 | 2000000 | -0.67 | -0.89 | B>B |
| chr10 | 2000000 | 2100000 | -0.70 | -0.94 | B>B |
| chr10 | 2100000 | 2200000 | -0.69 | -0.89 | B>B |
| chr10 | 2200000 | 2300000 | -0.61 | -0.98 | B>B |
| chr10 | 2300000 | 2400000 | -0.69 | -1.01 | B>B |
| chr10 | 2400000 | 2500000 | -0.63 | -1.08 | B>B |
| chr10 | 2500000 | 2600000 | -0.50 | -1.09 | A>B |
| chr10 | 2600000 | 2700000 | -0.51 | -1.00 | B>B |
| chr10 | 2700000 | 2800000 | -0.53 | -0.98 | B>B |
| chr10 | 2800000 | 2900000 | -0.31 | -0.65 | B>B |
| chr10 | 2900000 | 3000000 | 0.13 | -0.24 | B>B |
| chr10 | 3000000 | 3100000 | 0.37 | 0.10 | A>A |

| | | | | | |
|---|---|---|---|---|---|
| chr10 | 3100000 | 3200000 | 0.88 | 0.88 | A>A |
| chr10 | 3200000 | 3300000 | 0.59 | 0.73 | A>A |
| chr10 | 3300000 | 3400000 | 0.79 | 0.90 | A>A |
| chr10 | 3400000 | 3500000 | 0.87 | 0.86 | A>A |
| chr10 | 3500000 | 3600000 | 0.76 | 0.97 | A>A |
| chr10 | 3600000 | 3700000 | 0.82 | 0.87 | A>A |
| chr10 | 3700000 | 3800000 | 0.78 | 0.91 | A>A |
| chr10 | 3800000 | 3900000 | 0.69 | 0.89 | A>A |
| chr10 | 3900000 | 4000000 | 0.83 | 0.89 | A>A |
| chr10 | 4000000 | 4100000 | 0.79 | 0.83 | A>A |
| chr10 | 4100000 | 4200000 | 0.69 | 0.76 | A>A |
| chr10 | 4200000 | 4300000 | 0.59 | 0.79 | A>A |
| chr10 | 4300000 | 4400000 | 0.63 | 0.80 | A>A |
| chr10 | 4400000 | 4500000 | 0.59 | 0.86 | A>A |
| chr10 | 4500000 | 4600000 | 0.58 | 0.76 | A>A |
| chr10 | 4600000 | 4700000 | 0.72 | 0.92 | A>A |
| chr10 | 4700000 | 4800000 | 0.74 | 0.86 | A>A |
| chr10 | 4800000 | 4900000 | 0.57 | 0.65 | A>A |
| chr10 | 4900000 | 5000000 | 1.02 | 0.91 | A>A |
| chr10 | 5000000 | 5100000 | 0.00 | 0.00 | NO |
| chr10 | 5100000 | 5200000 | 0.95 | 0.95 | A>A |
| chr10 | 5200000 | 5300000 | 0.00 | 0.00 | NO |
| chr10 | 5300000 | 5400000 | 1.17 | 1.07 | A>A |
| chr10 | 5400000 | 5500000 | 0.79 | 0.93 | A>A |
| chr10 | 5500000 | 5600000 | 1.08 | 1.29 | A>A |
| chr10 | 5600000 | 5700000 | 1.05 | 1.13 | A>A |
| chr10 | 5700000 | 5800000 | 1.07 | 1.04 | A>A |
| chr10 | 5800000 | 5900000 | 1.09 | 1.17 | A>A |
| chr10 | 5900000 | 6000000 | 1.03 | 1.12 | A>A |
| chr10 | 6000000 | 6100000 | 1.09 | 1.04 | A>A |
| chr10 | 6100000 | 6200000 | 0.98 | 0.91 | A>A |
| chr10 | 6200000 | 6300000 | 1.14 | 1.05 | A>A |
| chr10 | 6300000 | 6400000 | 1.20 | 1.12 | A>A |
| chr10 | 6400000 | 6500000 | 0.78 | 0.74 | A>A |
| chr10 | 6500000 | 6600000 | 0.81 | 0.74 | A>A |
| chr10 | 6600000 | 6700000 | 0.71 | 0.67 | A>A |
| chr10 | 6700000 | 6800000 | 0.60 | 0.67 | A>A |
| chr10 | 6800000 | 6900000 | 0.63 | 0.74 | A>A |
| chr10 | 6900000 | 7000000 | 0.43 | 0.58 | A>A |
| chr10 | 7000000 | 7100000 | 0.41 | 0.63 | A>A |
| chr10 | 7100000 | 7200000 | 0.35 | 0.55 | A>A |
| chr10 | 7200000 | 7300000 | 0.47 | 0.62 | A>A |
| chr10 | 7300000 | 7400000 | 0.72 | 0.81 | A>A |
| chr10 | 7400000 | 7500000 | 0.52 | 0.56 | A>A |
| chr10 | 7500000 | 7600000 | 0.22 | 0.40 | A>A |
| chr10 | 7600000 | 7700000 | 0.03 | 0.33 | A>A |
| chr10 | 7700000 | 7800000 | 0.16 | 0.50 | A>A |
| chr10 | 7800000 | 7900000 | 0.31 | 0.53 | A>A |
| chr10 | 7900000 | 8000000 | 0.15 | 0.60 | A>A |
| chr10 | 8000000 | 8100000 | 0.24 | 0.59 | A>A |
| chr10 | 8100000 | 8200000 | 0.31 | 0.67 | A>A |
| chr10 | 8200000 | 8300000 | 0.31 | 0.79 | A>A |
| chr10 | 8300000 | 8400000 | 0.48 | 0.75 | A>A |
| chr10 | 8400000 | 8500000 | 0.43 | 0.70 | A>A |
| chr10 | 8500000 | 8600000 | 0.33 | 0.69 | A>A |
| chr10 | 8600000 | 8700000 | 0.29 | 0.74 | A>A |

| | | | | | |
|---|---|---|---|---|---|
| chr10 | 8700000 | 8800000 | 0.27 | 0.66 | A>A |
| chr10 | 8800000 | 8900000 | 0.27 | 0.69 | A>A |
| chr10 | 8900000 | 9000000 | -0.05 | 0.53 | B>A |
| chr10 | 9000000 | 9100000 | -0.31 | 0.62 | B>A |
| chr10 | 9100000 | 9200000 | -0.32 | 0.65 | B>A |
| chr10 | 9200000 | 9300000 | -0.31 | 0.46 | B>A |
| chr10 | 9300000 | 9400000 | -0.36 | -0.02 | B>B |
| chr10 | 9400000 | 9500000 | -0.88 | -0.80 | B>B |
| chr10 | 9500000 | 9600000 | -1.01 | -0.97 | B>B |
| chr10 | 9600000 | 9700000 | -0.96 | -1.03 | B>B |
| chr10 | 9700000 | 9800000 | -0.97 | -1.02 | B>B |
| chr10 | 9800000 | 9900000 | -1.03 | -1.05 | B>B |
| chr10 | 9900000 | 10000000 | -0.99 | -1.02 | B>B |
| chr10 | 10000000 | 10100000 | -0.93 | -1.06 | B>B |
| chr10 | 10100000 | 10200000 | -0.94 | -0.98 | B>B |
| chr10 | 10200000 | 10300000 | -0.96 | -1.01 | B>B |
| chr10 | 10300000 | 10400000 | -0.89 | -1.01 | B>B |
| chr10 | 10400000 | 10500000 | -0.95 | -0.94 | B>B |
| chr10 | 10500000 | 10600000 | -0.89 | -0.93 | B>B |
| chr10 | 10600000 | 10700000 | -0.98 | -0.92 | B>B |
| chr10 | 10700000 | 10800000 | -0.93 | -0.94 | B>B |
| chr10 | 10800000 | 10900000 | -0.93 | -0.92 | B>B |
| chr10 | 10900000 | 11000000 | -0.88 | -0.93 | B>B |
| chr10 | 11000000 | 11100000 | -0.74 | -0.91 | B>B |
| chr10 | 11100000 | 11200000 | -0.72 | -0.96 | B>B |
| chr10 | 11200000 | 11300000 | -0.55 | -0.91 | B>B |
| chr10 | 11300000 | 11400000 | -0.20 | -0.79 | A>B |
| chr10 | 11400000 | 11500000 | -0.03 | -0.39 | B>B |
| chr10 | 11500000 | 11600000 | 0.36 | 0.31 | A>A |
| chr10 | 11600000 | 11700000 | 0.79 | 0.59 | A>A |
| chr10 | 11700000 | 11800000 | 0.58 | 0.46 | A>A |
| chr10 | 11800000 | 11900000 | 0.76 | 0.76 | A>A |
| chr10 | 11900000 | 12000000 | 0.92 | 1.05 | A>A |
| chr10 | 12000000 | 12100000 | 1.02 | 0.89 | A>A |
| chr10 | 12100000 | 12200000 | 1.17 | 1.08 | A>A |
| chr10 | 12200000 | 12300000 | 0.96 | 1.02 | A>A |
| chr10 | 12300000 | 12400000 | 0.98 | 0.95 | A>A |
| chr10 | 12400000 | 12500000 | 1.10 | 1.04 | A>A |
| chr10 | 12500000 | 12600000 | 1.07 | 1.01 | A>A |
| chr10 | 12600000 | 12700000 | 1.07 | 0.93 | A>A |
| chr10 | 12700000 | 12800000 | 0.78 | 0.75 | A>A |
| chr10 | 12800000 | 12900000 | 1.06 | 0.85 | A>A |
| chr10 | 12900000 | 13000000 | 0.88 | 0.72 | A>A |
| chr10 | 13000000 | 13100000 | 0.92 | 0.69 | A>A |
| chr10 | 13100000 | 13200000 | 0.85 | 0.81 | A>A |
| chr10 | 13200000 | 13300000 | 1.08 | 1.01 | A>A |
| chr10 | 13300000 | 13400000 | 1.15 | 1.02 | A>A |
| chr10 | 13400000 | 13500000 | 0.88 | 0.91 | A>A |
| chr10 | 13500000 | 13600000 | 1.03 | 0.88 | A>A |
| chr10 | 13600000 | 13700000 | 0.86 | 0.67 | A>A |
| chr10 | 13700000 | 13800000 | 0.98 | 0.66 | A>A |
| chr10 | 13800000 | 13900000 | 0.70 | 0.31 | A>A |
| chr10 | 13900000 | 14000000 | 0.88 | 0.16 | A>B |
| chr10 | 14000000 | 14100000 | 0.67 | 0.11 | A>A |
| chr10 | 14100000 | 14200000 | 0.60 | -0.01 | A>B |
| chr10 | 14200000 | 14300000 | 0.57 | -0.06 | A>B |

| | | | | | |
|---|---|---|---|---|---|
| chr10 | 14300000 | 14400000 | 0.59 | -0.14 | A>B |
| chr10 | 14400000 | 14500000 | 0.67 | 0.12 | A>A |
| chr10 | 14500000 | 14600000 | 0.79 | 0.34 | A>A |
| chr10 | 14600000 | 14700000 | 1.01 | 0.87 | A>A |
| chr10 | 14700000 | 14800000 | 1.22 | 1.00 | A>A |
| chr10 | 14800000 | 14900000 | 0.96 | 0.73 | A>A |
| chr10 | 14900000 | 15000000 | 0.86 | 0.60 | A>A |
| chr10 | 15000000 | 15100000 | 0.00 | 0.56 | NO |
| chr10 | 15100000 | 15200000 | 0.81 | 0.52 | A>A |
| chr10 | 15200000 | 15300000 | 0.69 | 0.28 | A>A |
| chr10 | 15300000 | 15400000 | 0.57 | 0.27 | A>A |
| chr10 | 15400000 | 15500000 | 0.19 | -0.16 | A>A |
| chr10 | 15500000 | 15600000 | -0.25 | -0.74 | B>B |
| chr10 | 15600000 | 15700000 | -0.28 | -0.86 | B>B |
| chr10 | 15700000 | 15800000 | -0.36 | -0.64 | B>B |
| chr10 | 15800000 | 15900000 | -0.01 | -0.21 | B>B |
| chr10 | 15900000 | 16000000 | -0.40 | -0.35 | B>B |
| chr10 | 16000000 | 16100000 | -0.65 | -0.67 | B>B |
| chr10 | 16100000 | 16200000 | -0.56 | -0.61 | B>B |
| chr10 | 16200000 | 16300000 | -0.50 | -0.50 | B>B |
| chr10 | 16300000 | 16400000 | -0.15 | -0.19 | B>B |
| chr10 | 16400000 | 16500000 | 0.16 | 0.17 | A>A |
| chr10 | 16500000 | 16600000 | -0.01 | -0.18 | B>B |
| chr10 | 16600000 | 16700000 | -0.23 | -0.33 | B>B |
| chr10 | 16700000 | 16800000 | -0.27 | -0.29 | B>B |
| chr10 | 16800000 | 16900000 | -0.02 | -0.09 | B>B |
| chr10 | 16900000 | 17000000 | 0.08 | 0.02 | A>A |
| chr10 | 17000000 | 17100000 | 0.32 | 0.06 | A>A |
| chr10 | 17100000 | 17200000 | 0.34 | 0.12 | A>A |
| chr10 | 17200000 | 17300000 | 0.30 | 0.21 | A>A |
| chr10 | 17300000 | 17400000 | -0.13 | -0.20 | B>B |
| chr10 | 17400000 | 17500000 | -0.02 | -0.16 | B>B |
| chr10 | 17500000 | 17600000 | 0.16 | 0.01 | A>A |
| chr10 | 17600000 | 17700000 | 0.34 | 0.07 | A>A |
| chr10 | 17700000 | 17800000 | 0.40 | 0.28 | A>A |
| chr10 | 17800000 | 17900000 | 0.00 | 0.00 | NO |
| chr10 | 17900000 | 18000000 | 0.00 | 0.00 | NO |
| chr10 | 18000000 | 18100000 | 0.00 | 0.00 | NO |
| chr10 | 18100000 | 18200000 | 0.00 | 0.00 | NO |
| chr10 | 18200000 | 18300000 | -0.28 | -0.75 | B>B |
| chr10 | 18300000 | 18400000 | -0.38 | -0.75 | B>B |
| chr10 | 18400000 | 18500000 | -0.31 | -0.61 | B>B |
| chr10 | 18500000 | 18600000 | -0.28 | -0.51 | B>B |
| chr10 | 18600000 | 18700000 | -0.14 | -0.53 | B>B |
| chr10 | 18700000 | 18800000 | -0.12 | -0.50 | B>B |
| chr10 | 18800000 | 18900000 | 0.00 | -0.12 | B>B |
| chr10 | 18900000 | 19000000 | -0.04 | -0.02 | B>B |
| chr10 | 19000000 | 19100000 | -0.51 | -0.29 | B>B |
| chr10 | 19100000 | 19200000 | -0.59 | -0.21 | B>B |
| chr10 | 19200000 | 19300000 | -0.63 | 0.03 | B>A |
| chr10 | 19300000 | 19400000 | -0.50 | 0.08 | B>A |
| chr10 | 19400000 | 19500000 | -0.66 | -0.03 | B>A |
| chr10 | 19500000 | 19600000 | -0.79 | -0.13 | B>A |
| chr10 | 19600000 | 19700000 | -0.76 | -0.45 | B>B |
| chr10 | 19700000 | 19800000 | -0.87 | -0.68 | B>B |
| chr10 | 19800000 | 19900000 | -0.88 | -0.81 | B>B |

| | | | | | |
|---|---|---|---|---|---|
| chr10 | 19900000 | 20000000 | -0.82 | -0.78 | B>B |
| chr10 | 20000000 | 20100000 | -0.72 | -0.67 | B>B |
| chr10 | 20100000 | 20200000 | -0.73 | -0.74 | B>B |
| chr10 | 20200000 | 20300000 | -0.87 | -0.81 | B>B |
| chr10 | 20300000 | 20400000 | -0.81 | -0.92 | B>B |
| chr10 | 20400000 | 20500000 | -0.89 | -0.90 | B>B |
| chr10 | 20500000 | 20600000 | -0.85 | -0.88 | B>B |
| chr10 | 20600000 | 20700000 | -0.84 | -0.83 | B>B |
| chr10 | 20700000 | 20800000 | -0.84 | -0.88 | B>B |
| chr10 | 20800000 | 20900000 | -0.73 | -0.80 | B>B |
| chr10 | 20900000 | 21000000 | -0.59 | -0.84 | B>B |
| chr10 | 21000000 | 21100000 | -0.51 | -0.70 | B>B |
| chr10 | 21100000 | 21200000 | -0.16 | -0.32 | B>B |
| chr10 | 21200000 | 21300000 | 0.04 | -0.15 | B>B |
| chr10 | 21300000 | 21400000 | 0.20 | 0.07 | A>A |
| chr10 | 21400000 | 21500000 | 0.43 | 0.39 | A>A |
| chr10 | 21500000 | 21600000 | 0.44 | 0.35 | A>A |
| chr10 | 21600000 | 21700000 | 0.81 | 0.71 | A>A |
| chr10 | 21700000 | 21800000 | 0.70 | 0.59 | A>A |
| chr10 | 21800000 | 21900000 | 0.71 | 0.69 | A>A |
| chr10 | 21900000 | 22000000 | 0.74 | 0.76 | A>A |
| chr10 | 22000000 | 22100000 | 0.43 | 0.45 | A>A |
| chr10 | 22100000 | 22200000 | 0.26 | 0.46 | A>A |
| chr10 | 22200000 | 22300000 | 0.21 | 0.32 | A>A |
| chr10 | 22300000 | 22400000 | 0.38 | 0.50 | A>A |
| chr10 | 22400000 | 22500000 | 0.51 | 0.40 | A>A |
| chr10 | 22500000 | 22600000 | 0.59 | 0.53 | A>A |
| chr10 | 22600000 | 22700000 | 0.48 | 0.50 | A>A |
| chr10 | 22700000 | 22800000 | 0.40 | 0.45 | A>A |
| chr10 | 22800000 | 22900000 | 0.30 | 0.40 | A>A |
| chr10 | 22900000 | 23000000 | 0.47 | 0.64 | A>A |
| chr10 | 23000000 | 23100000 | 0.30 | 0.59 | A>A |
| chr10 | 23100000 | 23200000 | 0.23 | 0.48 | A>A |
| chr10 | 23200000 | 23300000 | 0.06 | 0.34 | A>A |
| chr10 | 23300000 | 23400000 | -0.03 | 0.12 | A>A |
| chr10 | 23400000 | 23500000 | -0.39 | -0.37 | B>B |
| chr10 | 23500000 | 23600000 | -0.47 | -0.68 | B>B |
| chr10 | 23600000 | 23700000 | -0.39 | -0.61 | B>B |
| chr10 | 23700000 | 23800000 | -0.35 | -0.61 | B>B |
| chr10 | 23800000 | 23900000 | -0.72 | -0.76 | B>B |
| chr10 | 23900000 | 24000000 | -0.93 | -0.83 | B>B |
| chr10 | 24000000 | 24100000 | -1.01 | -0.92 | B>B |
| chr10 | 24100000 | 24200000 | -1.05 | -1.03 | B>B |
| chr10 | 24200000 | 24300000 | -0.80 | -0.78 | B>B |
| chr10 | 24300000 | 24400000 | -0.76 | -0.66 | B>B |
| chr10 | 24400000 | 24500000 | -0.56 | -0.49 | B>B |
| chr10 | 24500000 | 24600000 | -0.30 | -0.06 | B>B |
| chr10 | 24600000 | 24700000 | -0.20 | -0.16 | B>B |
| chr10 | 24700000 | 24800000 | -0.05 | -0.06 | B>B |
| chr10 | 24800000 | 24900000 | -0.01 | -0.02 | B>B |
| chr10 | 24900000 | 25000000 | 0.41 | 0.48 | A>A |
| chr10 | 25000000 | 25100000 | 0.66 | 0.66 | A>A |
| chr10 | 25100000 | 25200000 | 0.61 | 0.77 | A>A |
| chr10 | 25200000 | 25300000 | 0.53 | 0.52 | A>A |
| chr10 | 25300000 | 25400000 | 0.08 | 0.08 | A>A |
| chr10 | 25400000 | 25500000 | -0.32 | -0.29 | B>B |

| | | | | | |
|---|---|---|---|---|---|
| chr10 | 25500000 | 25600000 | -0.51 | -0.15 | B>B |
| chr10 | 25600000 | 25700000 | -0.63 | -0.18 | B>B |
| chr10 | 25700000 | 25800000 | -0.64 | -0.20 | B>B |
| chr10 | 25800000 | 25900000 | -0.63 | -0.39 | B>B |
| chr10 | 25900000 | 26000000 | -0.66 | -0.45 | B>B |
| chr10 | 26000000 | 26100000 | -0.61 | -0.82 | B>B |
| chr10 | 26100000 | 26200000 | -0.66 | -1.00 | B>B |
| chr10 | 26200000 | 26300000 | -0.79 | -1.02 | B>B |
| chr10 | 26300000 | 26400000 | -0.77 | -1.03 | B>B |
| chr10 | 26400000 | 26500000 | -0.64 | -1.08 | B>B |
| chr10 | 26500000 | 26600000 | -0.43 | -0.91 | B>B |
| chr10 | 26600000 | 26700000 | 0.00 | -0.61 | A>B |
| chr10 | 26700000 | 26800000 | 0.19 | -0.53 | A>B |
| chr10 | 26800000 | 26900000 | 0.02 | -0.43 | B>B |
| chr10 | 26900000 | 27000000 | 0.23 | -0.23 | B>B |
| chr10 | 27000000 | 27100000 | 0.42 | 0.27 | A>A |
| chr10 | 27100000 | 27200000 | 0.34 | 0.17 | A>A |
| chr10 | 27200000 | 27300000 | 0.28 | 0.11 | A>A |
| chr10 | 27300000 | 27400000 | 0.60 | 0.44 | A>A |
| chr10 | 27400000 | 27500000 | 0.72 | 0.63 | A>A |
| chr10 | 27500000 | 27600000 | 0.45 | 0.32 | A>A |
| chr10 | 27600000 | 27700000 | 0.05 | -0.10 | B>B |
| chr10 | 27700000 | 27800000 | -0.15 | -0.30 | B>B |
| chr10 | 27800000 | 27900000 | -0.12 | -0.40 | B>B |
| chr10 | 27900000 | 28000000 | -0.40 | -0.71 | B>B |
| chr10 | 28000000 | 28100000 | -0.41 | -0.76 | B>B |
| chr10 | 28100000 | 28200000 | -0.71 | -0.94 | B>B |
| chr10 | 28200000 | 28300000 | -0.58 | -0.91 | B>B |
| chr10 | 28300000 | 28400000 | -0.60 | -0.91 | B>B |
| chr10 | 28400000 | 28500000 | -0.42 | -0.60 | B>B |
| chr10 | 28500000 | 28600000 | -0.17 | -0.32 | B>B |
| chr10 | 28600000 | 28700000 | 0.10 | -0.14 | B>B |
| chr10 | 28700000 | 28800000 | 0.42 | 0.26 | A>A |
| chr10 | 28800000 | 28900000 | 0.79 | 0.67 | A>A |
| chr10 | 28900000 | 29000000 | 0.68 | 0.71 | A>A |
| chr10 | 29000000 | 29100000 | 1.01 | 1.11 | A>A |
| chr10 | 29100000 | 29200000 | 0.92 | 1.02 | A>A |
| chr10 | 29200000 | 29300000 | 0.89 | 0.93 | A>A |
| chr10 | 29300000 | 29400000 | 0.92 | 0.99 | A>A |
| chr10 | 29400000 | 29500000 | 0.89 | 0.84 | A>A |
| chr10 | 29500000 | 29600000 | 0.75 | 0.80 | A>A |
| chr10 | 29600000 | 29700000 | 0.81 | 0.80 | A>A |
| chr10 | 29700000 | 29800000 | 0.78 | 0.88 | A>A |
| chr10 | 29800000 | 29900000 | 0.82 | 0.84 | A>A |
| chr10 | 29900000 | 30000000 | 0.98 | 0.95 | A>A |
| chr10 | 30000000 | 30100000 | 0.94 | 0.94 | A>A |
| chr10 | 30100000 | 30200000 | 0.86 | 0.84 | A>A |
| chr10 | 30200000 | 30300000 | 0.69 | 0.71 | A>A |
| chr10 | 30300000 | 30400000 | 0.88 | 0.79 | A>A |
| chr10 | 30400000 | 30500000 | 0.87 | 0.79 | A>A |
| chr10 | 30500000 | 30600000 | 0.77 | 0.70 | A>A |
| chr10 | 30600000 | 30700000 | 0.78 | 0.76 | A>A |
| chr10 | 30700000 | 30800000 | 1.11 | 1.06 | A>A |
| chr10 | 30800000 | 30900000 | 0.89 | 0.95 | A>A |
| chr10 | 30900000 | 31000000 | 0.70 | 0.78 | A>A |
| chr10 | 31000000 | 31100000 | 0.48 | 0.48 | A>A |

| | | | | | |
|---|---|---|---|---|---|
| chr10 | 31100000 | 31200000 | 0.52 | 0.62 | A>A |
| chr10 | 31200000 | 31300000 | 0.49 | 0.71 | A>A |
| chr10 | 31300000 | 31400000 | 0.47 | 0.54 | A>A |
| chr10 | 31400000 | 31500000 | 0.56 | 0.57 | A>A |
| chr10 | 31500000 | 31600000 | 0.41 | 0.59 | A>A |
| chr10 | 31600000 | 31700000 | 0.54 | 0.69 | A>A |
| chr10 | 31700000 | 31800000 | 0.37 | 0.54 | A>A |
| chr10 | 31800000 | 31900000 | 0.34 | 0.58 | A>A |
| chr10 | 31900000 | 32000000 | 0.36 | 0.56 | A>A |
| chr10 | 32000000 | 32100000 | 0.49 | 0.64 | A>A |
| chr10 | 32100000 | 32200000 | 0.81 | 0.86 | A>A |
| chr10 | 32200000 | 32300000 | 0.83 | 0.88 | A>A |
| chr10 | 32300000 | 32400000 | 0.89 | 0.90 | A>A |
| chr10 | 32400000 | 32500000 | 0.75 | 0.90 | A>A |
| chr10 | 32500000 | 32600000 | 0.79 | 0.87 | A>A |
| chr10 | 32600000 | 32700000 | 0.73 | 0.83 | A>A |
| chr10 | 32700000 | 32800000 | 0.71 | 0.77 | A>A |
| chr10 | 32800000 | 32900000 | 0.62 | 0.70 | A>A |
| chr10 | 32900000 | 33000000 | 0.54 | 0.70 | A>A |
| chr10 | 33000000 | 33100000 | 0.55 | 0.65 | A>A |
| chr10 | 33100000 | 33200000 | 0.75 | 0.86 | A>A |
| chr10 | 33200000 | 33300000 | 0.82 | 0.92 | A>A |
| chr10 | 33300000 | 33400000 | 0.66 | 0.81 | A>A |
| chr10 | 33400000 | 33500000 | 0.50 | 0.70 | A>A |
| chr10 | 33500000 | 33600000 | 0.71 | 0.84 | A>A |
| chr10 | 33600000 | 33700000 | 0.61 | 0.69 | A>A |
| chr10 | 33700000 | 33800000 | 0.71 | 0.74 | A>A |
| chr10 | 33800000 | 33900000 | 0.66 | 0.74 | A>A |
| chr10 | 33900000 | 34000000 | 0.60 | 0.64 | A>A |
| chr10 | 34000000 | 34100000 | 0.60 | 0.68 | A>A |
| chr10 | 34100000 | 34200000 | 0.58 | 0.70 | A>A |
| chr10 | 34200000 | 34300000 | 0.67 | 0.71 | A>A |
| chr10 | 34300000 | 34400000 | 0.60 | 0.69 | A>A |
| chr10 | 34400000 | 34500000 | 0.71 | 0.81 | A>A |
| chr10 | 34500000 | 34600000 | 0.75 | 0.79 | A>A |
| chr10 | 34600000 | 34700000 | 0.74 | 0.86 | A>A |
| chr10 | 34700000 | 34800000 | 0.79 | 0.92 | A>A |
| chr10 | 34800000 | 34900000 | 0.78 | 0.89 | A>A |
| chr10 | 34900000 | 35000000 | 0.63 | 0.72 | A>A |
| chr10 | 35000000 | 35100000 | 0.99 | 1.07 | A>A |
| chr10 | 35100000 | 35200000 | 0.97 | 1.07 | A>A |
| chr10 | 35200000 | 35300000 | 0.77 | 0.84 | A>A |
| chr10 | 35300000 | 35400000 | 0.71 | 0.77 | A>A |
| chr10 | 35400000 | 35500000 | 0.63 | 0.70 | A>A |
| chr10 | 35500000 | 35600000 | 0.44 | 0.49 | A>A |
| chr10 | 35600000 | 35700000 | 0.61 | 0.63 | A>A |
| chr10 | 35700000 | 35800000 | 0.51 | 0.61 | A>A |
| chr10 | 35800000 | 35900000 | 0.28 | 0.26 | A>A |
| chr10 | 35900000 | 36000000 | -0.07 | -0.19 | B>B |
| chr10 | 36000000 | 36100000 | -0.20 | -0.67 | B>B |
| chr10 | 36100000 | 36200000 | -0.25 | -0.74 | B>B |
| chr10 | 36200000 | 36300000 | -0.31 | -0.78 | B>B |
| chr10 | 36300000 | 36400000 | -0.29 | -0.80 | B>B |
| chr10 | 36400000 | 36500000 | -0.34 | -0.82 | B>B |
| chr10 | 36500000 | 36600000 | -0.25 | -0.79 | B>B |
| chr10 | 36600000 | 36700000 | -0.20 | -0.74 | B>B |

| | | | | | |
|---|---|---|---|---|---|
| chr10 | 36700000 | 36800000 | -0.17 | -0.76 | A>B |
| chr10 | 36800000 | 36900000 | -0.09 | -0.68 | A>B |
| chr10 | 36900000 | 37000000 | -0.08 | -0.66 | B>B |
| chr10 | 37000000 | 37100000 | -0.09 | -0.41 | B>B |
| chr10 | 37100000 | 37200000 | -0.23 | -0.03 | B>B |
| chr10 | 37200000 | 37300000 | -0.44 | 0.55 | B>A |
| chr10 | 37300000 | 37400000 | 0.00 | 0.00 | NO |
| chr10 | 37400000 | 37500000 | 0.00 | 0.00 | NO |
| chr10 | 37500000 | 37600000 | -0.46 | 0.55 | B>A |
| chr10 | 37600000 | 37700000 | -0.50 | 0.44 | B>A |
| chr10 | 37700000 | 37800000 | -0.50 | 0.57 | B>A |
| chr10 | 37800000 | 37900000 | -0.50 | 0.67 | B>A |
| chr10 | 37900000 | 38000000 | 0.00 | 0.00 | NO |
| chr10 | 38000000 | 38100000 | -0.61 | 0.48 | B>A |
| chr10 | 38100000 | 38200000 | -0.43 | 0.71 | B>A |
| chr10 | 38200000 | 38300000 | -0.32 | 0.57 | B>A |
| chr10 | 38300000 | 38400000 | 0.00 | 0.00 | NO |
| chr10 | 38400000 | 38500000 | 0.00 | 0.00 | NO |
| chr10 | 38500000 | 38600000 | 0.00 | 0.00 | NO |
| chr10 | 38600000 | 38700000 | 0.00 | 0.00 | NO |
| chr10 | 38700000 | 38800000 | 0.00 | 0.00 | NO |
| chr10 | 38800000 | 38900000 | 0.00 | 0.00 | NO |
| chr10 | 38900000 | 39000000 | 0.00 | 0.00 | NO |
| chr10 | 39000000 | 39100000 | 0.00 | 0.00 | NO |
| chr10 | 39100000 | 39200000 | 0.00 | 0.00 | NO |
| chr10 | 39200000 | 39300000 | 0.00 | 0.00 | NO |
| chr10 | 39300000 | 39400000 | 0.00 | 0.00 | NO |
| chr10 | 39400000 | 39500000 | 0.00 | 0.00 | NO |
| chr10 | 39500000 | 39600000 | 0.00 | 0.00 | NO |
| chr10 | 39600000 | 39700000 | 0.00 | 0.00 | NO |
| chr10 | 39700000 | 39800000 | 0.00 | 0.00 | NO |
| chr10 | 39800000 | 39900000 | 0.00 | 0.00 | NO |
| chr10 | 39900000 | 40000000 | 0.00 | 0.00 | NO |
| chr10 | 40000000 | 40100000 | 0.00 | 0.00 | NO |
| chr10 | 40100000 | 40200000 | 0.00 | 0.00 | NO |
| chr10 | 40200000 | 40300000 | 0.00 | 0.00 | NO |
| chr10 | 40300000 | 40400000 | 0.00 | 0.00 | NO |
| chr10 | 40400000 | 40500000 | 0.00 | 0.00 | NO |
| chr10 | 40500000 | 40600000 | 0.00 | 0.00 | NO |
| chr10 | 40600000 | 40700000 | 0.00 | 0.00 | NO |
| chr10 | 40700000 | 40800000 | 0.00 | 0.00 | NO |
| chr10 | 40800000 | 40900000 | 0.00 | 0.00 | NO |
| chr10 | 40900000 | 41000000 | 0.00 | 0.00 | NO |
| chr10 | 41000000 | 41100000 | 0.00 | 0.00 | NO |
| chr10 | 41100000 | 41200000 | 0.00 | 0.00 | NO |
| chr10 | 41200000 | 41300000 | 0.00 | 0.00 | NO |
| chr10 | 41300000 | 41400000 | 0.00 | 0.00 | NO |
| chr10 | 41400000 | 41500000 | 0.00 | 0.00 | NO |
| chr10 | 41500000 | 41600000 | 0.00 | 0.00 | NO |
| chr10 | 41600000 | 41700000 | 0.00 | 0.00 | NO |
| chr10 | 41700000 | 41800000 | 0.00 | 0.00 | NO |
| chr10 | 41800000 | 41900000 | 0.00 | 0.00 | NO |
| chr10 | 41900000 | 42000000 | 0.00 | 0.00 | NO |
| chr10 | 42000000 | 42100000 | 0.00 | 0.00 | NO |
| chr10 | 42100000 | 42200000 | 0.00 | 0.00 | NO |
| chr10 | 42200000 | 42300000 | 0.00 | 0.00 | NO |

| | | | | | |
|---|---|---|---|---|---|
| chr10 | 42300000 | 42400000 | 0.00 | 0.00 | NO |
| chr10 | 42400000 | 42500000 | 0.00 | 0.00 | NO |
| chr10 | 42500000 | 42600000 | 0.00 | 0.00 | NO |
| chr10 | 42600000 | 42700000 | 0.00 | 0.00 | NO |
| chr10 | 42700000 | 42800000 | 0.00 | 0.00 | NO |
| chr10 | 42800000 | 42900000 | 0.00 | 0.00 | NO |
| chr10 | 42900000 | 43000000 | 0.00 | 0.00 | NO |
| chr10 | 43000000 | 43100000 | 0.00 | 0.00 | NO |
| chr10 | 43100000 | 43200000 | 0.00 | 0.00 | NO |
| chr10 | 43200000 | 43300000 | 0.00 | 0.81 | NO |
| chr10 | 43300000 | 43400000 | 0.00 | 0.00 | NO |
| chr10 | 43400000 | 43500000 | -0.06 | 0.92 | B>A |
| chr10 | 43500000 | 43600000 | 0.13 | 0.92 | B>A |
| chr10 | 43600000 | 43700000 | 0.48 | 0.90 | A>A |
| chr10 | 43700000 | 43800000 | 0.76 | 1.11 | A>A |
| chr10 | 43800000 | 43900000 | 0.75 | 1.16 | A>A |
| chr10 | 43900000 | 44000000 | 0.26 | 0.90 | B>A |
| chr10 | 44000000 | 44100000 | -0.38 | 0.55 | B>A |
| chr10 | 44100000 | 44200000 | -0.35 | -0.01 | B>B |
| chr10 | 44200000 | 44300000 | -0.37 | -0.64 | B>B |
| chr10 | 44300000 | 44400000 | -0.51 | -0.74 | B>B |
| chr10 | 44400000 | 44500000 | -0.50 | -0.80 | B>B |
| chr10 | 44500000 | 44600000 | -0.60 | -0.75 | B>B |
| chr10 | 44600000 | 44700000 | -0.57 | -0.52 | B>B |
| chr10 | 44700000 | 44800000 | -0.39 | -0.36 | B>B |
| chr10 | 44800000 | 44900000 | -0.41 | -0.40 | B>B |
| chr10 | 44900000 | 45000000 | -0.74 | -0.85 | B>B |
| chr10 | 45000000 | 45100000 | -0.81 | -0.97 | B>B |
| chr10 | 45100000 | 45200000 | -0.81 | -0.92 | B>B |
| chr10 | 45200000 | 45300000 | -0.68 | -0.84 | B>B |
| chr10 | 45300000 | 45400000 | -0.33 | -0.40 | B>B |
| chr10 | 45400000 | 45500000 | 0.01 | 0.03 | A>A |
| chr10 | 45500000 | 45600000 | -0.34 | -0.39 | B>B |
| chr10 | 45600000 | 45700000 | 0.00 | 0.00 | NO |
| chr10 | 45700000 | 45800000 | -0.19 | -0.19 | B>B |
| chr10 | 45800000 | 45900000 | 0.00 | -0.05 | NO |
| chr10 | 45900000 | 46000000 | 0.19 | 0.20 | A>A |
| chr10 | 46000000 | 46100000 | 0.45 | 0.39 | A>A |
| chr10 | 46100000 | 46200000 | 0.00 | 0.34 | NO |
| chr10 | 46200000 | 46300000 | 0.00 | 0.00 | NO |
| chr10 | 46300000 | 46400000 | 0.00 | 0.00 | NO |
| chr10 | 46400000 | 46500000 | 0.00 | 0.00 | NO |
| chr10 | 46500000 | 46600000 | 0.00 | 0.00 | NO |
| chr10 | 46600000 | 46700000 | 0.00 | 0.00 | NO |
| chr10 | 46700000 | 46800000 | 0.00 | 0.00 | NO |
| chr10 | 46800000 | 46900000 | 0.00 | 0.00 | NO |
| chr10 | 46900000 | 47000000 | 0.00 | 0.00 | NO |
| chr10 | 47000000 | 47100000 | 1.08 | 1.03 | A>A |
| chr10 | 47100000 | 47200000 | 0.90 | 0.91 | A>A |
| chr10 | 47200000 | 47300000 | 0.00 | 0.00 | NO |
| chr10 | 47300000 | 47400000 | 0.00 | 0.00 | NO |
| chr10 | 47400000 | 47500000 | 0.00 | 0.00 | NO |
| chr10 | 47500000 | 47600000 | 0.00 | 0.00 | NO |
| chr10 | 47600000 | 47700000 | 0.90 | 0.82 | A>A |
| chr10 | 47700000 | 47800000 | 0.00 | 0.00 | NO |
| chr10 | 47800000 | 47900000 | 0.00 | 0.00 | NO |

| | | | | | |
|---|---|---|---|---|---|
| chr10 | 47900000 | 48000000 | 0.00 | 0.00 | NO |
| chr10 | 48000000 | 48100000 | 0.00 | 0.00 | NO |
| chr10 | 48100000 | 48200000 | 0.00 | 0.00 | NO |
| chr10 | 48200000 | 48300000 | 0.00 | 0.00 | NO |
| chr10 | 48300000 | 48400000 | 1.22 | 0.77 | A>A |
| chr10 | 48400000 | 48500000 | 0.88 | 0.16 | A>B |
| chr10 | 48500000 | 48600000 | 0.70 | 0.04 | A>B |
| chr10 | 48600000 | 48700000 | 0.00 | -0.07 | NO |
| chr10 | 48700000 | 48800000 | 0.00 | 0.00 | NO |
| chr10 | 48800000 | 48900000 | 0.00 | 0.00 | NO |
| chr10 | 48900000 | 49000000 | 0.00 | 0.00 | NO |
| chr10 | 49000000 | 49100000 | 0.00 | 0.00 | NO |
| chr10 | 49100000 | 49200000 | 0.00 | 0.00 | NO |
| chr10 | 49200000 | 49300000 | 0.00 | 0.00 | NO |
| chr10 | 49300000 | 49400000 | 0.00 | 0.00 | NO |
| chr10 | 49400000 | 49500000 | 0.81 | 0.25 | A>A |
| chr10 | 49500000 | 49600000 | 0.59 | 0.32 | A>A |
| chr10 | 49600000 | 49700000 | 0.52 | 0.33 | A>A |
| chr10 | 49700000 | 49800000 | 0.37 | 0.10 | A>A |
| chr10 | 49800000 | 49900000 | 0.10 | -0.22 | B>B |
| chr10 | 49900000 | 50000000 | -0.11 | -0.92 | A>B |
| chr10 | 50000000 | 50100000 | -0.19 | -1.11 | A>B |
| chr10 | 50100000 | 50200000 | -0.22 | -1.11 | A>B |
| chr10 | 50200000 | 50300000 | -0.33 | -1.12 | A>B |
| chr10 | 50300000 | 50400000 | -0.25 | -1.13 | A>B |
| chr10 | 50400000 | 50500000 | -0.29 | -1.12 | A>B |
| chr10 | 50500000 | 50600000 | -0.21 | -0.91 | A>B |
| chr10 | 50600000 | 50700000 | -0.08 | -0.40 | B>B |
| chr10 | 50700000 | 50800000 | 0.15 | -0.10 | A>A |
| chr10 | 50800000 | 50900000 | 0.18 | -0.25 | B>B |
| chr10 | 50900000 | 51000000 | 0.34 | -0.13 | A>A |
| chr10 | 51000000 | 51100000 | 0.00 | 0.00 | NO |
| chr10 | 51100000 | 51200000 | 0.00 | 0.00 | NO |
| chr10 | 51200000 | 51300000 | 0.00 | 0.00 | NO |
| chr10 | 51300000 | 51400000 | 0.00 | 0.00 | NO |
| chr10 | 51400000 | 51500000 | 0.00 | 0.00 | NO |
| chr10 | 51500000 | 51600000 | 0.71 | 0.70 | A>A |
| chr10 | 51600000 | 51700000 | 0.00 | 0.00 | NO |
| chr10 | 51700000 | 51800000 | 0.00 | 0.00 | NO |
| chr10 | 51800000 | 51900000 | 0.00 | 0.00 | NO |
| chr10 | 51900000 | 52000000 | 0.00 | 0.00 | NO |
| chr10 | 52000000 | 52100000 | 0.16 | -0.06 | A>A |
| chr10 | 52100000 | 52200000 | 0.47 | 0.26 | A>A |
| chr10 | 52200000 | 52300000 | 0.42 | 0.34 | A>A |
| chr10 | 52300000 | 52400000 | 0.38 | 0.24 | A>A |
| chr10 | 52400000 | 52500000 | 0.00 | 0.00 | NO |
| chr10 | 52500000 | 52600000 | 0.00 | -0.84 | NO |
| chr10 | 52600000 | 52700000 | -0.73 | -0.92 | B>B |
| chr10 | 52700000 | 52800000 | -0.58 | -0.71 | B>B |
| chr10 | 52800000 | 52900000 | -0.64 | -0.81 | B>B |
| chr10 | 52900000 | 53000000 | -0.81 | -0.99 | B>B |
| chr10 | 53000000 | 53100000 | -0.80 | -1.03 | B>B |
| chr10 | 53100000 | 53200000 | -0.72 | -0.98 | B>B |
| chr10 | 53200000 | 53300000 | -0.86 | -0.99 | B>B |
| chr10 | 53300000 | 53400000 | -0.81 | -0.93 | B>B |
| chr10 | 53400000 | 53500000 | -0.50 | -0.78 | B>B |

| | | | | | |
|---|---|---|---|---|---|
| chr10 | 53500000 | 53600000 | -0.70 | -0.96 | B>B |
| chr10 | 53600000 | 53700000 | -0.66 | -0.91 | B>B |
| chr10 | 53700000 | 53800000 | -0.58 | -0.80 | B>B |
| chr10 | 53800000 | 53900000 | -0.62 | -0.87 | B>B |
| chr10 | 53900000 | 54000000 | -0.38 | -0.64 | B>B |
| chr10 | 54000000 | 54100000 | 0.11 | -0.09 | A>A |
| chr10 | 54100000 | 54200000 | 0.65 | 0.45 | A>A |
| chr10 | 54200000 | 54300000 | 0.40 | 0.16 | A>A |
| chr10 | 54300000 | 54400000 | -0.04 | -0.30 | B>B |
| chr10 | 54400000 | 54500000 | -0.40 | -0.65 | B>B |
| chr10 | 54500000 | 54600000 | -0.61 | -0.73 | B>B |
| chr10 | 54600000 | 54700000 | -0.81 | -0.85 | B>B |
| chr10 | 54700000 | 54800000 | -0.93 | -0.94 | B>B |
| chr10 | 54800000 | 54900000 | -0.99 | -0.97 | B>B |
| chr10 | 54900000 | 55000000 | -1.01 | -0.98 | B>B |
| chr10 | 55000000 | 55100000 | -0.99 | -0.98 | B>B |
| chr10 | 55100000 | 55200000 | -0.94 | -0.98 | B>B |
| chr10 | 55200000 | 55300000 | -0.91 | -0.91 | B>B |
| chr10 | 55300000 | 55400000 | -0.82 | -0.82 | B>B |
| chr10 | 55400000 | 55500000 | -0.88 | -0.86 | B>B |
| chr10 | 55500000 | 55600000 | -0.85 | -0.91 | B>B |
| chr10 | 55600000 | 55700000 | -0.82 | -0.91 | B>B |
| chr10 | 55700000 | 55800000 | -0.86 | -0.92 | B>B |
| chr10 | 55800000 | 55900000 | -0.85 | -0.88 | B>B |
| chr10 | 55900000 | 56000000 | -0.85 | -0.92 | B>B |
| chr10 | 56000000 | 56100000 | -0.87 | -0.93 | B>B |
| chr10 | 56100000 | 56200000 | -0.86 | -0.85 | B>B |
| chr10 | 56200000 | 56300000 | -0.84 | -0.92 | B>B |
| chr10 | 56300000 | 56400000 | -0.90 | -0.92 | B>B |
| chr10 | 56400000 | 56500000 | -0.88 | -0.92 | B>B |
| chr10 | 56500000 | 56600000 | -0.90 | -0.90 | B>B |
| chr10 | 56600000 | 56700000 | -0.84 | -0.85 | B>B |
| chr10 | 56700000 | 56800000 | -0.83 | -0.89 | B>B |
| chr10 | 56800000 | 56900000 | -0.81 | -0.82 | B>B |
| chr10 | 56900000 | 57000000 | -0.81 | -0.83 | B>B |
| chr10 | 57000000 | 57100000 | -0.80 | -0.88 | B>B |
| chr10 | 57100000 | 57200000 | -0.88 | -0.86 | B>B |
| chr10 | 57200000 | 57300000 | -0.88 | -0.89 | B>B |
| chr10 | 57300000 | 57400000 | -0.78 | -0.85 | B>B |
| chr10 | 57400000 | 57500000 | -0.78 | -0.85 | B>B |
| chr10 | 57500000 | 57600000 | -0.82 | -0.89 | B>B |
| chr10 | 57600000 | 57700000 | -0.87 | -0.86 | B>B |
| chr10 | 57700000 | 57800000 | -0.86 | -0.84 | B>B |
| chr10 | 57800000 | 57900000 | -0.86 | -0.85 | B>B |
| chr10 | 57900000 | 58000000 | -0.84 | -0.84 | B>B |
| chr10 | 58000000 | 58100000 | -0.83 | -0.81 | B>B |
| chr10 | 58100000 | 58200000 | -0.74 | -0.71 | B>B |
| chr10 | 58200000 | 58300000 | -0.86 | -0.85 | B>B |
| chr10 | 58300000 | 58400000 | -0.86 | -0.80 | B>B |
| chr10 | 58400000 | 58500000 | -0.93 | -0.86 | B>B |
| chr10 | 58500000 | 58600000 | -0.87 | -0.84 | B>B |
| chr10 | 58600000 | 58700000 | -0.90 | -0.88 | B>B |
| chr10 | 58700000 | 58800000 | -0.89 | -0.92 | B>B |
| chr10 | 58800000 | 58900000 | -0.89 | -0.95 | B>B |
| chr10 | 58900000 | 59000000 | -0.91 | -0.89 | B>B |
| chr10 | 59000000 | 59100000 | -0.88 | -0.82 | B>B |

| | | | | | |
|---|---|---|---|---|---|
| chr10 | 59100000 | 59200000 | -0.78 | -0.81 | B>B |
| chr10 | 59200000 | 59300000 | -0.82 | -0.86 | B>B |
| chr10 | 59300000 | 59400000 | -0.40 | -0.39 | B>B |
| chr10 | 59400000 | 59500000 | -0.19 | -0.15 | B>B |
| chr10 | 59500000 | 59600000 | -0.21 | -0.18 | B>B |
| chr10 | 59600000 | 59700000 | -0.02 | -0.01 | B>B |
| chr10 | 59700000 | 59800000 | -0.02 | 0.04 | A>A |
| chr10 | 59800000 | 59900000 | -0.27 | -0.28 | B>B |
| chr10 | 59900000 | 60000000 | 0.00 | 0.02 | A>A |
| chr10 | 60000000 | 60100000 | 0.23 | 0.23 | A>A |
| chr10 | 60100000 | 60200000 | 0.32 | 0.35 | A>A |
| chr10 | 60200000 | 60300000 | 0.45 | 0.48 | A>A |
| chr10 | 60300000 | 60400000 | 0.31 | 0.40 | A>A |
| chr10 | 60400000 | 60500000 | 0.32 | 0.47 | A>A |
| chr10 | 60500000 | 60600000 | 0.24 | 0.27 | A>A |
| chr10 | 60600000 | 60700000 | 0.31 | 0.20 | A>A |
| chr10 | 60700000 | 60800000 | 0.27 | 0.28 | A>A |
| chr10 | 60800000 | 60900000 | 0.32 | 0.30 | A>A |
| chr10 | 60900000 | 61000000 | 0.07 | 0.01 | A>A |
| chr10 | 61000000 | 61100000 | -0.31 | -0.45 | B>B |
| chr10 | 61100000 | 61200000 | -0.54 | -0.63 | B>B |
| chr10 | 61200000 | 61300000 | -0.59 | -0.67 | B>B |
| chr10 | 61300000 | 61400000 | -0.33 | -0.29 | B>B |
| chr10 | 61400000 | 61500000 | -0.25 | -0.03 | B>B |
| chr10 | 61500000 | 61600000 | -0.10 | 0.05 | B>B |
| chr10 | 61600000 | 61700000 | 0.08 | 0.43 | A>A |
| chr10 | 61700000 | 61800000 | -0.18 | 0.47 | B>A |
| chr10 | 61800000 | 61900000 | -0.01 | 0.38 | A>A |
| chr10 | 61900000 | 62000000 | 0.24 | 0.56 | A>A |
| chr10 | 62000000 | 62100000 | 0.24 | 0.64 | A>A |
| chr10 | 62100000 | 62200000 | 0.28 | 0.65 | A>A |
| chr10 | 62200000 | 62300000 | 0.13 | 0.42 | A>A |
| chr10 | 62300000 | 62400000 | 0.09 | 0.44 | A>A |
| chr10 | 62400000 | 62500000 | 0.13 | 0.57 | A>A |
| chr10 | 62500000 | 62600000 | 0.23 | 0.29 | A>A |
| chr10 | 62600000 | 62700000 | 0.42 | 0.21 | A>A |
| chr10 | 62700000 | 62800000 | 0.43 | 0.24 | A>A |
| chr10 | 62800000 | 62900000 | 0.25 | 0.01 | A>A |
| chr10 | 62900000 | 63000000 | 0.05 | -0.17 | B>B |
| chr10 | 63000000 | 63100000 | -0.13 | -0.32 | B>B |
| chr10 | 63100000 | 63200000 | -0.32 | -0.47 | B>B |
| chr10 | 63200000 | 63300000 | -0.31 | -0.49 | B>B |
| chr10 | 63300000 | 63400000 | -0.30 | -0.31 | B>B |
| chr10 | 63400000 | 63500000 | -0.07 | -0.08 | B>B |
| chr10 | 63500000 | 63600000 | 0.14 | 0.19 | A>A |
| chr10 | 63600000 | 63700000 | 0.17 | 0.23 | A>A |
| chr10 | 63700000 | 63800000 | 0.19 | 0.31 | A>A |
| chr10 | 63800000 | 63900000 | 0.33 | 0.37 | A>A |
| chr10 | 63900000 | 64000000 | 0.35 | 0.41 | A>A |
| chr10 | 64000000 | 64100000 | 0.04 | -0.15 | B>B |
| chr10 | 64100000 | 64200000 | 0.09 | -0.22 | B>B |
| chr10 | 64200000 | 64300000 | -0.08 | -0.29 | B>B |
| chr10 | 64300000 | 64400000 | -0.25 | -0.49 | B>B |
| chr10 | 64400000 | 64500000 | -0.38 | -0.79 | B>B |
| chr10 | 64500000 | 64600000 | -0.15 | -0.58 | B>B |
| chr10 | 64600000 | 64700000 | -0.42 | -0.85 | B>B |

| | | | | | |
|---|---|---|---|---|---|
| chr10 | 64700000 | 64800000 | -0.37 | -0.79 | B>B |
| chr10 | 64800000 | 64900000 | -0.12 | -0.52 | B>B |
| chr10 | 64900000 | 65000000 | 0.28 | 0.17 | A>A |
| chr10 | 65000000 | 65100000 | 0.63 | 0.58 | A>A |
| chr10 | 65100000 | 65200000 | 0.64 | 0.57 | A>A |
| chr10 | 65200000 | 65300000 | 0.63 | 0.57 | A>A |
| chr10 | 65300000 | 65400000 | 0.49 | 0.43 | A>A |
| chr10 | 65400000 | 65500000 | 0.28 | 0.21 | A>A |
| chr10 | 65500000 | 65600000 | -0.30 | -0.41 | B>B |
| chr10 | 65600000 | 65700000 | -0.66 | -0.83 | B>B |
| chr10 | 65700000 | 65800000 | -0.70 | -0.96 | B>B |
| chr10 | 65800000 | 65900000 | -0.73 | -1.02 | B>B |
| chr10 | 65900000 | 66000000 | -0.94 | -1.09 | B>B |
| chr10 | 66000000 | 66100000 | -0.97 | -1.07 | B>B |
| chr10 | 66100000 | 66200000 | -0.93 | -1.03 | B>B |
| chr10 | 66200000 | 66300000 | -0.95 | -1.03 | B>B |
| chr10 | 66300000 | 66400000 | -0.85 | -1.01 | B>B |
| chr10 | 66400000 | 66500000 | -0.70 | -0.91 | B>B |
| chr10 | 66500000 | 66600000 | -0.75 | -0.95 | B>B |
| chr10 | 66600000 | 66700000 | -0.81 | -0.97 | B>B |
| chr10 | 66700000 | 66800000 | -0.87 | -0.93 | B>B |
| chr10 | 66800000 | 66900000 | -0.80 | -0.87 | B>B |
| chr10 | 66900000 | 67000000 | -0.80 | -0.85 | B>B |
| chr10 | 67000000 | 67100000 | -0.75 | -0.80 | B>B |
| chr10 | 67100000 | 67200000 | -0.67 | -0.76 | B>B |
| chr10 | 67200000 | 67300000 | -0.66 | -0.73 | B>B |
| chr10 | 67300000 | 67400000 | -0.57 | -0.60 | B>B |
| chr10 | 67400000 | 67500000 | -0.71 | -0.66 | B>B |
| chr10 | 67500000 | 67600000 | -0.71 | -0.74 | B>B |
| chr10 | 67600000 | 67700000 | -0.70 | -0.73 | B>B |
| chr10 | 67700000 | 67800000 | -0.86 | -0.82 | B>B |
| chr10 | 67800000 | 67900000 | -0.89 | -0.89 | B>B |
| chr10 | 67900000 | 68000000 | -0.80 | -0.79 | B>B |
| chr10 | 68000000 | 68100000 | -0.94 | -0.95 | B>B |
| chr10 | 68100000 | 68200000 | -1.01 | -0.99 | B>B |
| chr10 | 68200000 | 68300000 | -1.04 | -1.01 | B>B |
| chr10 | 68300000 | 68400000 | -1.02 | -1.02 | B>B |
| chr10 | 68400000 | 68500000 | -1.06 | -1.02 | B>B |
| chr10 | 68500000 | 68600000 | -1.00 | -0.99 | B>B |
| chr10 | 68600000 | 68700000 | -0.83 | -0.78 | B>B |
| chr10 | 68700000 | 68800000 | -0.87 | -0.58 | B>B |
| chr10 | 68800000 | 68900000 | -0.92 | -0.50 | B>B |
| chr10 | 68900000 | 69000000 | -0.86 | -0.59 | B>B |
| chr10 | 69000000 | 69100000 | -0.78 | -0.65 | B>B |
| chr10 | 69100000 | 69200000 | -0.83 | -0.86 | B>B |
| chr10 | 69200000 | 69300000 | -0.57 | -0.64 | B>B |
| chr10 | 69300000 | 69400000 | -0.50 | -0.77 | B>B |
| chr10 | 69400000 | 69500000 | -0.17 | -0.40 | B>B |
| chr10 | 69500000 | 69600000 | 0.33 | 0.11 | A>A |
| chr10 | 69600000 | 69700000 | 0.58 | 0.56 | A>A |
| chr10 | 69700000 | 69800000 | 0.70 | 0.81 | A>A |
| chr10 | 69800000 | 69900000 | 0.94 | 0.91 | A>A |
| chr10 | 69900000 | 70000000 | 0.80 | 0.80 | A>A |
| chr10 | 70000000 | 70100000 | 0.79 | 0.89 | A>A |
| chr10 | 70100000 | 70200000 | 1.01 | 1.02 | A>A |
| chr10 | 70200000 | 70300000 | 0.98 | 1.06 | A>A |

| | | | | | |
|---|---|---|---|---|---|
| chr10 | 70300000 | 70400000 | 1.16 | 1.08 | A>A |
| chr10 | 70400000 | 70500000 | 1.10 | 1.12 | A>A |
| chr10 | 70500000 | 70600000 | 1.02 | 1.10 | A>A |
| chr10 | 70600000 | 70700000 | 1.06 | 1.06 | A>A |
| chr10 | 70700000 | 70800000 | 1.03 | 1.00 | A>A |
| chr10 | 70800000 | 70900000 | 1.05 | 1.08 | A>A |
| chr10 | 70900000 | 71000000 | 0.93 | 1.04 | A>A |
| chr10 | 71000000 | 71100000 | 1.06 | 0.98 | A>A |
| chr10 | 71100000 | 71200000 | 0.92 | 0.89 | A>A |
| chr10 | 71200000 | 71300000 | 0.92 | 0.91 | A>A |
| chr10 | 71300000 | 71400000 | 0.82 | 0.79 | A>A |
| chr10 | 71400000 | 71500000 | 0.75 | 0.73 | A>A |
| chr10 | 71500000 | 71600000 | 0.88 | 0.81 | A>A |
| chr10 | 71600000 | 71700000 | 0.90 | 0.87 | A>A |
| chr10 | 71700000 | 71800000 | 0.77 | 0.76 | A>A |
| chr10 | 71800000 | 71900000 | 0.91 | 0.84 | A>A |
| chr10 | 71900000 | 72000000 | 0.82 | 0.86 | A>A |
| chr10 | 72000000 | 72100000 | 0.94 | 0.91 | A>A |
| chr10 | 72100000 | 72200000 | 0.86 | 0.78 | A>A |
| chr10 | 72200000 | 72300000 | 0.76 | 0.68 | A>A |
| chr10 | 72300000 | 72400000 | 0.70 | 0.59 | A>A |
| chr10 | 72400000 | 72500000 | 0.77 | 0.69 | A>A |
| chr10 | 72500000 | 72600000 | 0.69 | 0.64 | A>A |
| chr10 | 72600000 | 72700000 | 0.59 | 0.61 | A>A |
| chr10 | 72700000 | 72800000 | 0.65 | 0.64 | A>A |
| chr10 | 72800000 | 72900000 | 0.66 | 0.60 | A>A |
| chr10 | 72900000 | 73000000 | 0.62 | 0.61 | A>A |
| chr10 | 73000000 | 73100000 | 0.52 | 0.56 | A>A |
| chr10 | 73100000 | 73200000 | 0.64 | 0.69 | A>A |
| chr10 | 73200000 | 73300000 | 0.73 | 0.73 | A>A |
| chr10 | 73300000 | 73400000 | 0.56 | 0.64 | A>A |
| chr10 | 73400000 | 73500000 | 0.69 | 0.73 | A>A |
| chr10 | 73500000 | 73600000 | 0.78 | 0.74 | A>A |
| chr10 | 73600000 | 73700000 | 0.73 | 0.67 | A>A |
| chr10 | 73700000 | 73800000 | 0.74 | 0.83 | A>A |
| chr10 | 73800000 | 73900000 | 0.72 | 0.76 | A>A |
| chr10 | 73900000 | 74000000 | 0.80 | 0.80 | A>A |
| chr10 | 74000000 | 74100000 | 0.72 | 0.81 | A>A |
| chr10 | 74100000 | 74200000 | 0.57 | 0.64 | A>A |
| chr10 | 74200000 | 74300000 | 0.81 | 0.80 | A>A |
| chr10 | 74300000 | 74400000 | 0.82 | 0.82 | A>A |
| chr10 | 74400000 | 74500000 | 0.79 | 0.88 | A>A |
| chr10 | 74500000 | 74600000 | 0.72 | 0.84 | A>A |
| chr10 | 74600000 | 74700000 | 0.62 | 0.76 | A>A |
| chr10 | 74700000 | 74800000 | 0.72 | 0.76 | A>A |
| chr10 | 74800000 | 74900000 | 0.83 | 0.88 | A>A |
| chr10 | 74900000 | 75000000 | 0.81 | 0.89 | A>A |
| chr10 | 75000000 | 75100000 | 0.86 | 1.00 | A>A |
| chr10 | 75100000 | 75200000 | 0.89 | 1.00 | A>A |
| chr10 | 75200000 | 75300000 | 0.92 | 1.03 | A>A |
| chr10 | 75300000 | 75400000 | 0.99 | 1.02 | A>A |
| chr10 | 75400000 | 75500000 | 0.00 | 0.00 | NO |
| chr10 | 75500000 | 75600000 | 0.94 | 1.05 | A>A |
| chr10 | 75600000 | 75700000 | 1.00 | 1.13 | A>A |
| chr10 | 75700000 | 75800000 | 1.05 | 1.11 | A>A |
| chr10 | 75800000 | 75900000 | 0.95 | 1.02 | A>A |

| | | | | | |
|---|---|---|---|---|---|
| chr10 | 75900000 | 76000000 | 0.85 | 0.92 | A>A |
| chr10 | 76000000 | 76100000 | 0.55 | 0.68 | A>A |
| chr10 | 76100000 | 76200000 | 0.52 | 0.67 | A>A |
| chr10 | 76200000 | 76300000 | 0.57 | 0.75 | A>A |
| chr10 | 76300000 | 76400000 | 0.52 | 0.71 | A>A |
| chr10 | 76400000 | 76500000 | 0.51 | 0.72 | A>A |
| chr10 | 76500000 | 76600000 | 0.66 | 0.67 | A>A |
| chr10 | 76600000 | 76700000 | 0.81 | 0.92 | A>A |
| chr10 | 76700000 | 76800000 | 0.64 | 0.74 | A>A |
| chr10 | 76800000 | 76900000 | 0.63 | 0.70 | A>A |
| chr10 | 76900000 | 77000000 | 0.79 | 0.78 | A>A |
| chr10 | 77000000 | 77100000 | 0.44 | 0.29 | A>A |
| chr10 | 77100000 | 77200000 | 0.25 | -0.03 | A>A |
| chr10 | 77200000 | 77300000 | 0.18 | -0.20 | B>B |
| chr10 | 77300000 | 77400000 | 0.06 | -0.46 | B>B |
| chr10 | 77400000 | 77500000 | 0.01 | -0.58 | A>B |
| chr10 | 77500000 | 77600000 | 0.07 | -0.58 | A>B |
| chr10 | 77600000 | 77700000 | 0.09 | -0.62 | A>B |
| chr10 | 77700000 | 77800000 | 0.11 | -0.55 | A>B |
| chr10 | 77800000 | 77900000 | -0.07 | -0.58 | B>B |
| chr10 | 77900000 | 78000000 | -0.25 | -0.88 | A>B |
| chr10 | 78000000 | 78100000 | -0.19 | -0.80 | A>B |
| chr10 | 78100000 | 78200000 | -0.31 | -0.90 | A>B |
| chr10 | 78200000 | 78300000 | -0.42 | -0.89 | B>B |
| chr10 | 78300000 | 78400000 | -0.28 | -0.79 | B>B |
| chr10 | 78400000 | 78500000 | -0.40 | -1.04 | A>B |
| chr10 | 78500000 | 78600000 | -0.43 | -1.01 | B>B |
| chr10 | 78600000 | 78700000 | -0.11 | -0.70 | A>B |
| chr10 | 78700000 | 78800000 | 0.06 | -0.34 | B>B |
| chr10 | 78800000 | 78900000 | 0.26 | -0.08 | A>A |
| chr10 | 78900000 | 79000000 | 0.51 | 0.43 | A>A |
| chr10 | 79000000 | 79100000 | 0.66 | 0.59 | A>A |
| chr10 | 79100000 | 79200000 | 0.75 | 0.65 | A>A |
| chr10 | 79200000 | 79300000 | 0.77 | 0.71 | A>A |
| chr10 | 79300000 | 79400000 | 0.79 | 0.67 | A>A |
| chr10 | 79400000 | 79500000 | 0.59 | 0.56 | A>A |
| chr10 | 79500000 | 79600000 | 0.92 | 0.82 | A>A |
| chr10 | 79600000 | 79700000 | 1.07 | 1.14 | A>A |
| chr10 | 79700000 | 79800000 | 0.99 | 1.00 | A>A |
| chr10 | 79800000 | 79900000 | 0.75 | 0.70 | A>A |
| chr10 | 79900000 | 80000000 | 0.66 | 0.77 | A>A |
| chr10 | 80000000 | 80100000 | 0.79 | 0.74 | A>A |
| chr10 | 80100000 | 80200000 | 0.85 | 0.77 | A>A |
| chr10 | 80200000 | 80300000 | 0.77 | 0.85 | A>A |
| chr10 | 80300000 | 80400000 | 0.72 | 0.77 | A>A |
| chr10 | 80400000 | 80500000 | 0.65 | 0.74 | A>A |
| chr10 | 80500000 | 80600000 | 0.82 | 0.91 | A>A |
| chr10 | 80600000 | 80700000 | 0.82 | 0.91 | A>A |
| chr10 | 80700000 | 80800000 | 0.80 | 0.90 | A>A |
| chr10 | 80800000 | 80900000 | 0.89 | 0.98 | A>A |
| chr10 | 80900000 | 81000000 | 0.80 | 0.85 | A>A |
| chr10 | 81000000 | 81100000 | 0.78 | 0.84 | A>A |
| chr10 | 81100000 | 81200000 | 0.92 | 0.78 | A>A |
| chr10 | 81200000 | 81300000 | 0.91 | 0.67 | A>A |
| chr10 | 81300000 | 81400000 | 0.41 | 0.18 | A>A |
| chr10 | 81400000 | 81500000 | 0.00 | 0.00 | NO |

| | | | | | |
|---|---|---|---|---|---|
| chr10 | 81500000 | 81600000 | 0.00 | 0.00 | NO |
| chr10 | 81600000 | 81700000 | 0.00 | 0.00 | NO |
| chr10 | 81700000 | 81800000 | 0.57 | 0.27 | A>A |
| chr10 | 81800000 | 81900000 | 0.63 | 0.48 | A>A |
| chr10 | 81900000 | 82000000 | 0.77 | 0.75 | A>A |
| chr10 | 82000000 | 82100000 | 0.27 | 0.06 | A>A |
| chr10 | 82100000 | 82200000 | 0.19 | 0.07 | A>A |
| chr10 | 82200000 | 82300000 | 0.56 | 0.40 | A>A |
| chr10 | 82300000 | 82400000 | 0.04 | -0.30 | B>B |
| chr10 | 82400000 | 82500000 | -0.40 | -0.60 | B>B |
| chr10 | 82500000 | 82600000 | -0.73 | -1.03 | B>B |
| chr10 | 82600000 | 82700000 | -0.73 | -0.99 | B>B |
| chr10 | 82700000 | 82800000 | -0.78 | -0.89 | B>B |
| chr10 | 82800000 | 82900000 | -0.85 | -0.58 | B>B |
| chr10 | 82900000 | 83000000 | -0.85 | -0.41 | B>B |
| chr10 | 83000000 | 83100000 | -0.97 | -0.38 | B>A |
| chr10 | 83100000 | 83200000 | -1.01 | -0.28 | B>A |
| chr10 | 83200000 | 83300000 | -1.03 | -0.11 | B>A |
| chr10 | 83300000 | 83400000 | -0.98 | 0.00 | B>A |
| chr10 | 83400000 | 83500000 | -1.02 | 0.08 | B>A |
| chr10 | 83500000 | 83600000 | -0.96 | -0.25 | B>A |
| chr10 | 83600000 | 83700000 | -0.99 | -0.72 | B>B |
| chr10 | 83700000 | 83800000 | -1.01 | -0.92 | B>B |
| chr10 | 83800000 | 83900000 | -1.04 | -0.97 | B>B |
| chr10 | 83900000 | 84000000 | -0.99 | -0.87 | B>B |
| chr10 | 84000000 | 84100000 | -0.97 | -0.77 | B>B |
| chr10 | 84100000 | 84200000 | -0.98 | -0.53 | B>B |
| chr10 | 84200000 | 84300000 | -0.88 | -0.46 | B>B |
| chr10 | 84300000 | 84400000 | -0.85 | -0.23 | B>A |
| chr10 | 84400000 | 84500000 | -0.87 | -0.27 | B>A |
| chr10 | 84500000 | 84600000 | -0.96 | -0.54 | B>B |
| chr10 | 84600000 | 84700000 | -0.98 | -0.86 | B>B |
| chr10 | 84700000 | 84800000 | -1.00 | -0.94 | B>B |
| chr10 | 84800000 | 84900000 | -1.08 | -0.99 | B>B |
| chr10 | 84900000 | 85000000 | -1.01 | -1.03 | B>B |
| chr10 | 85000000 | 85100000 | -1.08 | -1.08 | B>B |
| chr10 | 85100000 | 85200000 | -1.11 | -1.10 | B>B |
| chr10 | 85200000 | 85300000 | -1.14 | -1.11 | B>B |
| chr10 | 85300000 | 85400000 | -1.16 | -1.15 | B>B |
| chr10 | 85400000 | 85500000 | -1.09 | -1.12 | B>B |
| chr10 | 85500000 | 85600000 | -1.08 | -1.05 | B>B |
| chr10 | 85600000 | 85700000 | -0.88 | -1.02 | B>B |
| chr10 | 85700000 | 85800000 | -0.77 | -1.13 | B>B |
| chr10 | 85800000 | 85900000 | -0.60 | -0.95 | B>B |
| chr10 | 85900000 | 86000000 | 0.03 | -0.33 | B>B |
| chr10 | 86000000 | 86100000 | 0.11 | -0.07 | A>A |
| chr10 | 86100000 | 86200000 | 0.17 | 0.09 | A>A |
| chr10 | 86200000 | 86300000 | 0.13 | -0.01 | A>A |
| chr10 | 86300000 | 86400000 | -0.42 | -0.73 | B>B |
| chr10 | 86400000 | 86500000 | -0.68 | -1.13 | B>B |
| chr10 | 86500000 | 86600000 | -0.80 | -1.19 | B>B |
| chr10 | 86600000 | 86700000 | -0.97 | -1.18 | B>B |
| chr10 | 86700000 | 86800000 | -1.07 | -1.18 | B>B |
| chr10 | 86800000 | 86900000 | -1.00 | -1.17 | B>B |
| chr10 | 86900000 | 87000000 | -0.86 | -1.10 | B>B |
| chr10 | 87000000 | 87100000 | -0.90 | -1.03 | B>B |

| | | | | | |
|---|---|---|---|---|---|
| chr10 | 87100000 | 87200000 | -0.70 | -0.97 | B>B |
| chr10 | 87200000 | 87300000 | -0.53 | -0.75 | B>B |
| chr10 | 87300000 | 87400000 | -0.13 | -0.29 | B>B |
| chr10 | 87400000 | 87500000 | -0.17 | -0.43 | B>B |
| chr10 | 87500000 | 87600000 | -0.38 | -0.49 | B>B |
| chr10 | 87600000 | 87700000 | -0.40 | -0.62 | B>B |
| chr10 | 87700000 | 87800000 | -0.46 | -0.72 | B>B |
| chr10 | 87800000 | 87900000 | -0.36 | -0.72 | B>B |
| chr10 | 87900000 | 88000000 | -0.18 | -0.55 | B>B |
| chr10 | 88000000 | 88100000 | -0.04 | -0.34 | B>B |
| chr10 | 88100000 | 88200000 | 0.27 | 0.16 | A>A |
| chr10 | 88200000 | 88300000 | 0.86 | 0.66 | A>A |
| chr10 | 88300000 | 88400000 | 0.65 | 0.52 | A>A |
| chr10 | 88400000 | 88500000 | 0.89 | 0.82 | A>A |
| chr10 | 88500000 | 88600000 | 1.10 | 1.02 | A>A |
| chr10 | 88600000 | 88700000 | 0.91 | 0.92 | A>A |
| chr10 | 88700000 | 88800000 | 0.00 | 1.12 | NO |
| chr10 | 88800000 | 88900000 | 0.00 | 0.00 | NO |
| chr10 | 88900000 | 89000000 | 0.00 | 0.00 | NO |
| chr10 | 89000000 | 89100000 | 0.00 | 0.00 | NO |
| chr10 | 89100000 | 89200000 | 0.00 | 0.00 | NO |
| chr10 | 89200000 | 89300000 | 0.00 | 0.00 | NO |
| chr10 | 89300000 | 89400000 | 1.03 | 1.10 | A>A |
| chr10 | 89400000 | 89500000 | 1.06 | 1.12 | A>A |
| chr10 | 89500000 | 89600000 | 0.78 | 0.90 | A>A |
| chr10 | 89600000 | 89700000 | 0.71 | 0.76 | A>A |
| chr10 | 89700000 | 89800000 | 0.63 | 0.80 | A>A |
| chr10 | 89800000 | 89900000 | 0.52 | 0.75 | A>A |
| chr10 | 89900000 | 90000000 | 0.45 | 0.72 | A>A |
| chr10 | 90000000 | 90100000 | 0.37 | 0.71 | A>A |
| chr10 | 90100000 | 90200000 | 0.37 | 0.67 | A>A |
| chr10 | 90200000 | 90300000 | 0.29 | 0.67 | A>A |
| chr10 | 90300000 | 90400000 | 0.14 | 0.39 | A>A |
| chr10 | 90400000 | 90500000 | 0.17 | 0.55 | A>A |
| chr10 | 90500000 | 90600000 | 0.16 | 0.45 | A>A |
| chr10 | 90600000 | 90700000 | 0.44 | 0.50 | A>A |
| chr10 | 90700000 | 90800000 | 0.27 | 0.34 | A>A |
| chr10 | 90800000 | 90900000 | 0.09 | 0.09 | A>A |
| chr10 | 90900000 | 91000000 | 0.12 | 0.12 | A>A |
| chr10 | 91000000 | 91100000 | 0.56 | 0.57 | A>A |
| chr10 | 91100000 | 91200000 | 0.52 | 0.43 | A>A |
| chr10 | 91200000 | 91300000 | 0.36 | 0.25 | A>A |
| chr10 | 91300000 | 91400000 | 0.32 | 0.17 | A>A |
| chr10 | 91400000 | 91500000 | 0.33 | 0.10 | A>A |
| chr10 | 91500000 | 91600000 | -0.27 | -0.40 | B>B |
| chr10 | 91600000 | 91700000 | -0.74 | -0.64 | B>B |
| chr10 | 91700000 | 91800000 | -0.85 | -0.80 | B>B |
| chr10 | 91800000 | 91900000 | -0.80 | -0.96 | B>B |
| chr10 | 91900000 | 92000000 | -0.91 | -1.06 | B>B |
| chr10 | 92000000 | 92100000 | -0.76 | -0.99 | B>B |
| chr10 | 92100000 | 92200000 | -0.83 | -1.18 | B>B |
| chr10 | 92200000 | 92300000 | -0.92 | -1.26 | B>B |
| chr10 | 92300000 | 92400000 | -0.90 | -1.25 | B>B |
| chr10 | 92400000 | 92500000 | -0.69 | -1.14 | B>B |
| chr10 | 92500000 | 92600000 | -0.40 | -0.81 | B>B |
| chr10 | 92600000 | 92700000 | 0.07 | -0.11 | B>B |

| | | | | | |
|---|---|---|---|---|---|
| chr10 | 92700000 | 92800000 | 0.40 | 0.07 | A>A |
| chr10 | 92800000 | 92900000 | 0.31 | -0.13 | A>A |
| chr10 | 92900000 | 93000000 | 0.34 | 0.11 | A>A |
| chr10 | 93000000 | 93100000 | 0.25 | 0.11 | A>A |
| chr10 | 93100000 | 93200000 | 0.37 | 0.22 | A>A |
| chr10 | 93200000 | 93300000 | 0.59 | 0.44 | A>A |
| chr10 | 93300000 | 93400000 | 0.77 | 0.74 | A>A |
| chr10 | 93400000 | 93500000 | 0.64 | 0.51 | A>A |
| chr10 | 93500000 | 93600000 | 0.62 | 0.69 | A>A |
| chr10 | 93600000 | 93700000 | 0.83 | 0.76 | A>A |
| chr10 | 93700000 | 93800000 | 0.88 | 0.86 | A>A |
| chr10 | 93800000 | 93900000 | 0.73 | 0.83 | A>A |
| chr10 | 93900000 | 94000000 | 0.98 | 0.97 | A>A |
| chr10 | 94000000 | 94100000 | 0.93 | 0.90 | A>A |
| chr10 | 94100000 | 94200000 | 0.74 | 0.80 | A>A |
| chr10 | 94200000 | 94300000 | 0.94 | 0.91 | A>A |
| chr10 | 94300000 | 94400000 | 0.96 | 1.00 | A>A |
| chr10 | 94400000 | 94500000 | 0.98 | 1.03 | A>A |
| chr10 | 94500000 | 94600000 | 1.02 | 1.03 | A>A |
| chr10 | 94600000 | 94700000 | 0.73 | 0.85 | A>A |
| chr10 | 94700000 | 94800000 | 0.79 | 0.87 | A>A |
| chr10 | 94800000 | 94900000 | 0.71 | 0.80 | A>A |
| chr10 | 94900000 | 95000000 | 0.90 | 0.97 | A>A |
| chr10 | 95000000 | 95100000 | 0.73 | 0.78 | A>A |
| chr10 | 95100000 | 95200000 | 0.86 | 0.94 | A>A |
| chr10 | 95200000 | 95300000 | 0.88 | 0.93 | A>A |
| chr10 | 95300000 | 95400000 | 0.91 | 0.88 | A>A |
| chr10 | 95400000 | 95500000 | 0.89 | 0.83 | A>A |
| chr10 | 95500000 | 95600000 | 0.81 | 0.81 | A>A |
| chr10 | 95600000 | 95700000 | 0.59 | 0.70 | A>A |
| chr10 | 95700000 | 95800000 | 0.80 | 0.89 | A>A |
| chr10 | 95800000 | 95900000 | 0.74 | 0.84 | A>A |
| chr10 | 95900000 | 96000000 | 0.76 | 0.81 | A>A |
| chr10 | 96000000 | 96100000 | 0.66 | 0.74 | A>A |
| chr10 | 96100000 | 96200000 | 0.80 | 0.85 | A>A |
| chr10 | 96200000 | 96300000 | 0.69 | 0.74 | A>A |
| chr10 | 96300000 | 96400000 | 0.55 | 0.70 | A>A |
| chr10 | 96400000 | 96500000 | 0.24 | 0.72 | A>A |
| chr10 | 96500000 | 96600000 | 0.00 | 0.00 | NO |
| chr10 | 96600000 | 96700000 | 0.00 | 0.00 | NO |
| chr10 | 96700000 | 96800000 | 0.00 | 0.00 | NO |
| chr10 | 96800000 | 96900000 | 0.04 | 0.64 | B>A |
| chr10 | 96900000 | 97000000 | 0.40 | 0.61 | A>A |
| chr10 | 97000000 | 97100000 | 0.65 | 0.83 | A>A |
| chr10 | 97100000 | 97200000 | 0.44 | 0.56 | A>A |
| chr10 | 97200000 | 97300000 | 0.59 | 0.73 | A>A |
| chr10 | 97300000 | 97400000 | 0.47 | 0.60 | A>A |
| chr10 | 97400000 | 97500000 | 0.31 | 0.40 | A>A |
| chr10 | 97500000 | 97600000 | 0.11 | 0.36 | A>A |
| chr10 | 97600000 | 97700000 | 0.20 | 0.28 | A>A |
| chr10 | 97700000 | 97800000 | 0.10 | 0.17 | A>A |
| chr10 | 97800000 | 97900000 | 0.42 | 0.36 | A>A |
| chr10 | 97900000 | 98000000 | 0.26 | 0.18 | A>A |
| chr10 | 98000000 | 98100000 | 0.24 | 0.20 | A>A |
| chr10 | 98100000 | 98200000 | 0.37 | 0.28 | A>A |
| chr10 | 98200000 | 98300000 | 0.55 | 0.41 | A>A |

| | | | | | |
|---|---|---|---|---|---|
| chr10 | 98300000 | 98400000 | 0.61 | 0.66 | A>A |
| chr10 | 98400000 | 98500000 | 0.66 | 0.62 | A>A |
| chr10 | 98500000 | 98600000 | 0.66 | 0.57 | A>A |
| chr10 | 98600000 | 98700000 | 0.77 | 0.83 | A>A |
| chr10 | 98700000 | 98800000 | 0.73 | 0.80 | A>A |
| chr10 | 98800000 | 98900000 | 0.79 | 0.72 | A>A |
| chr10 | 98900000 | 99000000 | 0.87 | 0.77 | A>A |
| chr10 | 99000000 | 99100000 | 0.96 | 0.95 | A>A |
| chr10 | 99100000 | 99200000 | 1.08 | 1.04 | A>A |
| chr10 | 99200000 | 99300000 | 1.13 | 1.13 | A>A |
| chr10 | 99300000 | 99400000 | 1.02 | 1.07 | A>A |
| chr10 | 99400000 | 99500000 | 0.97 | 0.99 | A>A |
| chr10 | 99500000 | 99600000 | 0.80 | 0.68 | A>A |
| chr10 | 99600000 | 99700000 | 0.74 | 0.41 | A>A |
| chr10 | 99700000 | 99800000 | 0.47 | 0.09 | A>A |
| chr10 | 99800000 | 99900000 | 0.39 | 0.03 | A>A |
| chr10 | 99900000 | 100000000 | 0.42 | 0.19 | A>A |
| chr10 | 1E+08 | 100100000 | 0.36 | 0.09 | A>A |
| chr10 | 1E+08 | 100200000 | 0.24 | 0.08 | A>A |
| chr10 | 1E+08 | 100300000 | -0.22 | -0.63 | B>B |
| chr10 | 1E+08 | 100400000 | -0.49 | -1.10 | A>B |
| chr10 | 1E+08 | 100500000 | -0.63 | -1.10 | B>B |
| chr10 | 1.01E+08 | 100600000 | -0.76 | -1.26 | B>B |
| chr10 | 1.01E+08 | 100700000 | -0.76 | -1.30 | B>B |
| chr10 | 1.01E+08 | 100800000 | -0.80 | -1.22 | B>B |
| chr10 | 1.01E+08 | 100900000 | -0.64 | -1.09 | B>B |
| chr10 | 1.01E+08 | 101000000 | -0.39 | -0.91 | B>B |
| chr10 | 1.01E+08 | 101100000 | 0.05 | -0.24 | B>B |
| chr10 | 1.01E+08 | 101200000 | 0.35 | 0.09 | A>A |
| chr10 | 1.01E+08 | 101300000 | 0.61 | 0.26 | A>A |
| chr10 | 1.01E+08 | 101400000 | 0.76 | 0.53 | A>A |
| chr10 | 1.01E+08 | 101500000 | 0.86 | 0.82 | A>A |
| chr10 | 1.02E+08 | 101600000 | 0.97 | 1.09 | A>A |
| chr10 | 1.02E+08 | 101700000 | 0.87 | 0.98 | A>A |
| chr10 | 1.02E+08 | 101800000 | 1.02 | 1.11 | A>A |
| chr10 | 1.02E+08 | 101900000 | 0.90 | 0.98 | A>A |
| chr10 | 1.02E+08 | 102000000 | 0.94 | 1.06 | A>A |
| chr10 | 1.02E+08 | 102100000 | 0.87 | 0.93 | A>A |
| chr10 | 1.02E+08 | 102200000 | 1.04 | 1.07 | A>A |
| chr10 | 1.02E+08 | 102300000 | 0.74 | 0.81 | A>A |
| chr10 | 1.02E+08 | 102400000 | 0.81 | 0.77 | A>A |
| chr10 | 1.02E+08 | 102500000 | 0.89 | 0.95 | A>A |
| chr10 | 1.03E+08 | 102600000 | 0.88 | 0.90 | A>A |
| chr10 | 1.03E+08 | 102700000 | 0.81 | 0.80 | A>A |
| chr10 | 1.03E+08 | 102800000 | 0.95 | 0.94 | A>A |
| chr10 | 1.03E+08 | 102900000 | 0.97 | 0.92 | A>A |
| chr10 | 1.03E+08 | 103000000 | 0.75 | 0.77 | A>A |
| chr10 | 1.03E+08 | 103100000 | 0.78 | 0.86 | A>A |
| chr10 | 1.03E+08 | 103200000 | 0.83 | 0.99 | A>A |
| chr10 | 1.03E+08 | 103300000 | 0.78 | 0.84 | A>A |
| chr10 | 1.03E+08 | 103400000 | 0.76 | 0.81 | A>A |
| chr10 | 1.03E+08 | 103500000 | 0.84 | 0.97 | A>A |
| chr10 | 1.04E+08 | 103600000 | 0.80 | 0.85 | A>A |
| chr10 | 1.04E+08 | 103700000 | 0.92 | 0.94 | A>A |
| chr10 | 1.04E+08 | 103800000 | 0.93 | 0.94 | A>A |
| chr10 | 1.04E+08 | 103900000 | 0.79 | 0.78 | A>A |

| | | | | | |
|---|---|---|---|---|---|
| chr10 | 1.04E+08 | 104000000 | 0.76 | 0.76 | A>A |
| chr10 | 1.04E+08 | 104100000 | 1.06 | 1.02 | A>A |
| chr10 | 1.04E+08 | 104200000 | 0.82 | 0.79 | A>A |
| chr10 | 1.04E+08 | 104300000 | 0.92 | 0.94 | A>A |
| chr10 | 1.04E+08 | 104400000 | 0.97 | 0.94 | A>A |
| chr10 | 1.04E+08 | 104500000 | 0.98 | 0.97 | A>A |
| chr10 | 1.05E+08 | 104600000 | 0.95 | 0.98 | A>A |
| chr10 | 1.05E+08 | 104700000 | 0.82 | 0.92 | A>A |
| chr10 | 1.05E+08 | 104800000 | 1.01 | 1.05 | A>A |
| chr10 | 1.05E+08 | 104900000 | 0.92 | 0.91 | A>A |
| chr10 | 1.05E+08 | 105000000 | 1.15 | 1.20 | A>A |
| chr10 | 1.05E+08 | 105100000 | 0.98 | 1.00 | A>A |
| chr10 | 1.05E+08 | 105200000 | 1.01 | 1.05 | A>A |
| chr10 | 1.05E+08 | 105300000 | 1.18 | 1.11 | A>A |
| chr10 | 1.05E+08 | 105400000 | 1.06 | 1.07 | A>A |
| chr10 | 1.05E+08 | 105500000 | 0.93 | 1.03 | A>A |
| chr10 | 1.06E+08 | 105600000 | 1.06 | 1.08 | A>A |
| chr10 | 1.06E+08 | 105700000 | 0.95 | 0.92 | A>A |
| chr10 | 1.06E+08 | 105800000 | 0.99 | 0.92 | A>A |
| chr10 | 1.06E+08 | 105900000 | 0.87 | 0.80 | A>A |
| chr10 | 1.06E+08 | 106000000 | 0.79 | 0.70 | A>A |
| chr10 | 1.06E+08 | 106100000 | 0.54 | 0.20 | A>A |
| chr10 | 1.06E+08 | 106200000 | -0.08 | -0.68 | A>B |
| chr10 | 1.06E+08 | 106300000 | -0.33 | -1.08 | A>B |
| chr10 | 1.06E+08 | 106400000 | -0.49 | -1.05 | B>B |
| chr10 | 1.06E+08 | 106500000 | -0.58 | -1.11 | B>B |
| chr10 | 1.07E+08 | 106600000 | -0.74 | -1.12 | B>B |
| chr10 | 1.07E+08 | 106700000 | -0.77 | -1.18 | B>B |
| chr10 | 1.07E+08 | 106800000 | -0.93 | -1.11 | B>B |
| chr10 | 1.07E+08 | 106900000 | -0.87 | -1.16 | B>B |
| chr10 | 1.07E+08 | 107000000 | -0.99 | -1.13 | B>B |
| chr10 | 1.07E+08 | 107100000 | -1.06 | -1.13 | B>B |
| chr10 | 1.07E+08 | 107200000 | -1.00 | -1.19 | B>B |
| chr10 | 1.07E+08 | 107300000 | -0.98 | -1.17 | B>B |
| chr10 | 1.07E+08 | 107400000 | -0.91 | -1.14 | B>B |
| chr10 | 1.07E+08 | 107500000 | -0.97 | -1.10 | B>B |
| chr10 | 1.08E+08 | 107600000 | -1.01 | -1.09 | B>B |
| chr10 | 1.08E+08 | 107700000 | -0.93 | -1.06 | B>B |
| chr10 | 1.08E+08 | 107800000 | -0.99 | -1.07 | B>B |
| chr10 | 1.08E+08 | 107900000 | -0.99 | -1.07 | B>B |
| chr10 | 1.08E+08 | 108000000 | -0.99 | -1.07 | B>B |
| chr10 | 1.08E+08 | 108100000 | -1.04 | -1.05 | B>B |
| chr10 | 1.08E+08 | 108200000 | -1.07 | -1.03 | B>B |
| chr10 | 1.08E+08 | 108300000 | -1.02 | -1.05 | B>B |
| chr10 | 1.08E+08 | 108400000 | -0.87 | -0.93 | B>B |
| chr10 | 1.08E+08 | 108500000 | -0.87 | -0.90 | B>B |
| chr10 | 1.09E+08 | 108600000 | -0.82 | -0.91 | B>B |
| chr10 | 1.09E+08 | 108700000 | -0.87 | -0.92 | B>B |
| chr10 | 1.09E+08 | 108800000 | -0.87 | -0.92 | B>B |
| chr10 | 1.09E+08 | 108900000 | -0.88 | -0.95 | B>B |
| chr10 | 1.09E+08 | 109000000 | -0.89 | -0.87 | B>B |
| chr10 | 1.09E+08 | 109100000 | -0.93 | -0.93 | B>B |
| chr10 | 1.09E+08 | 109200000 | -0.93 | -0.91 | B>B |
| chr10 | 1.09E+08 | 109300000 | -0.92 | -0.96 | B>B |
| chr10 | 1.09E+08 | 109400000 | -0.89 | -0.93 | B>B |
| chr10 | 1.09E+08 | 109500000 | -0.96 | -0.86 | B>B |

| | | | | | |
|---|---|---|---|---|---|
| chr10 | 1.1E+08 | 109600000 | -0.88 | -0.88 | B>B |
| chr10 | 1.1E+08 | 109700000 | -0.94 | -0.90 | B>B |
| chr10 | 1.1E+08 | 109800000 | -0.91 | -0.93 | B>B |
| chr10 | 1.1E+08 | 109900000 | -0.92 | -0.97 | B>B |
| chr10 | 1.1E+08 | 110000000 | -0.98 | -0.95 | B>B |
| chr10 | 1.1E+08 | 110100000 | -0.98 | -0.95 | B>B |
| chr10 | 1.1E+08 | 110200000 | -0.90 | -0.86 | B>B |
| chr10 | 1.1E+08 | 110300000 | -0.83 | -0.89 | B>B |
| chr10 | 1.1E+08 | 110400000 | -0.83 | -0.85 | B>B |
| chr10 | 1.1E+08 | 110500000 | -0.90 | -0.92 | B>B |
| chr10 | 1.11E+08 | 110600000 | -1.02 | -0.97 | B>B |
| chr10 | 1.11E+08 | 110700000 | -0.94 | -0.96 | B>B |
| chr10 | 1.11E+08 | 110800000 | -0.95 | -0.98 | B>B |
| chr10 | 1.11E+08 | 110900000 | -0.96 | -1.00 | B>B |
| chr10 | 1.11E+08 | 111000000 | -0.98 | -1.07 | B>B |
| chr10 | 1.11E+08 | 111100000 | -0.95 | -1.06 | B>B |
| chr10 | 1.11E+08 | 111200000 | -1.02 | -1.05 | B>B |
| chr10 | 1.11E+08 | 111300000 | -0.99 | -1.11 | B>B |
| chr10 | 1.11E+08 | 111400000 | -1.03 | -1.13 | B>B |
| chr10 | 1.11E+08 | 111500000 | -0.95 | -1.02 | B>B |
| chr10 | 1.12E+08 | 111600000 | -0.78 | -0.95 | B>B |
| chr10 | 1.12E+08 | 111700000 | -0.14 | -0.18 | B>B |
| chr10 | 1.12E+08 | 111800000 | 0.24 | 0.35 | A>A |
| chr10 | 1.12E+08 | 111900000 | 0.26 | 0.30 | A>A |
| chr10 | 1.12E+08 | 112000000 | 0.42 | 0.42 | A>A |
| chr10 | 1.12E+08 | 112100000 | 0.49 | 0.48 | A>A |
| chr10 | 1.12E+08 | 112200000 | 1.07 | 0.92 | A>A |
| chr10 | 1.12E+08 | 112300000 | 0.99 | 0.95 | A>A |
| chr10 | 1.12E+08 | 112400000 | 0.59 | 0.50 | A>A |
| chr10 | 1.12E+08 | 112500000 | 0.40 | 0.35 | A>A |
| chr10 | 1.13E+08 | 112600000 | 0.31 | 0.36 | A>A |
| chr10 | 1.13E+08 | 112700000 | 0.29 | 0.38 | A>A |
| chr10 | 1.13E+08 | 112800000 | -0.02 | -0.09 | B>B |
| chr10 | 1.13E+08 | 112900000 | -0.38 | -0.58 | B>B |
| chr10 | 1.13E+08 | 113000000 | -0.66 | -1.01 | B>B |
| chr10 | 1.13E+08 | 113100000 | -0.73 | -1.02 | B>B |
| chr10 | 1.13E+08 | 113200000 | -0.89 | -1.10 | B>B |
| chr10 | 1.13E+08 | 113300000 | -0.99 | -1.07 | B>B |
| chr10 | 1.13E+08 | 113400000 | -1.00 | -1.07 | B>B |
| chr10 | 1.13E+08 | 113500000 | -1.09 | -1.09 | B>B |
| chr10 | 1.14E+08 | 113600000 | -1.01 | -1.15 | B>B |
| chr10 | 1.14E+08 | 113700000 | -1.06 | -1.14 | B>B |
| chr10 | 1.14E+08 | 113800000 | -0.91 | -1.14 | B>B |
| chr10 | 1.14E+08 | 113900000 | -0.64 | -0.75 | B>B |
| chr10 | 1.14E+08 | 114000000 | -0.38 | -0.45 | B>B |
| chr10 | 1.14E+08 | 114100000 | -0.31 | -0.41 | B>B |
| chr10 | 1.14E+08 | 114200000 | 0.03 | -0.12 | B>B |
| chr10 | 1.14E+08 | 114300000 | 0.27 | 0.27 | A>A |
| chr10 | 1.14E+08 | 114400000 | 0.49 | 0.57 | A>A |
| chr10 | 1.14E+08 | 114500000 | 0.56 | 0.70 | A>A |
| chr10 | 1.15E+08 | 114600000 | 0.61 | 0.78 | A>A |
| chr10 | 1.15E+08 | 114700000 | 0.69 | 0.76 | A>A |
| chr10 | 1.15E+08 | 114800000 | 0.73 | 0.83 | A>A |
| chr10 | 1.15E+08 | 114900000 | 0.65 | 0.73 | A>A |
| chr10 | 1.15E+08 | 115000000 | 0.57 | 0.46 | A>A |
| chr10 | 1.15E+08 | 115100000 | 0.65 | 0.32 | A>A |

| | | | | | |
|---|---|---|---|---|---|
| chr10 | 1.15E+08 | 115200000 | 0.39 | 0.30 | A>A |
| chr10 | 1.15E+08 | 115300000 | 0.41 | 0.28 | A>A |
| chr10 | 1.15E+08 | 115400000 | 0.53 | 0.36 | A>A |
| chr10 | 1.15E+08 | 115500000 | 0.68 | 0.56 | A>A |
| chr10 | 1.16E+08 | 115600000 | 0.79 | 0.77 | A>A |
| chr10 | 1.16E+08 | 115700000 | 0.88 | 0.87 | A>A |
| chr10 | 1.16E+08 | 115800000 | 0.91 | 0.95 | A>A |
| chr10 | 1.16E+08 | 115900000 | 0.72 | 0.77 | A>A |
| chr10 | 1.16E+08 | 116000000 | 0.78 | 0.79 | A>A |
| chr10 | 1.16E+08 | 116100000 | 0.62 | 0.62 | A>A |
| chr10 | 1.16E+08 | 116200000 | 0.66 | 0.83 | A>A |
| chr10 | 1.16E+08 | 116300000 | 0.56 | 0.62 | A>A |
| chr10 | 1.16E+08 | 116400000 | 0.71 | 0.77 | A>A |
| chr10 | 1.16E+08 | 116500000 | 0.74 | 0.64 | A>A |
| chr10 | 1.17E+08 | 116600000 | 0.54 | 0.52 | A>A |
| chr10 | 1.17E+08 | 116700000 | 0.19 | 0.11 | A>A |
| chr10 | 1.17E+08 | 116800000 | 0.01 | -0.25 | B>B |
| chr10 | 1.17E+08 | 116900000 | -0.27 | -0.56 | B>B |
| chr10 | 1.17E+08 | 117000000 | -0.79 | -0.87 | B>B |
| chr10 | 1.17E+08 | 117100000 | -0.96 | -1.08 | B>B |
| chr10 | 1.17E+08 | 117200000 | -0.85 | -0.92 | B>B |
| chr10 | 1.17E+08 | 117300000 | -0.93 | -1.06 | B>B |
| chr10 | 1.17E+08 | 117400000 | -0.93 | -1.02 | B>B |
| chr10 | 1.17E+08 | 117500000 | -0.63 | -0.93 | B>B |
| chr10 | 1.18E+08 | 117600000 | -0.50 | -0.79 | B>B |
| chr10 | 1.18E+08 | 117700000 | -0.25 | -0.79 | B>B |
| chr10 | 1.18E+08 | 117800000 | -0.28 | -0.79 | B>B |
| chr10 | 1.18E+08 | 117900000 | -0.34 | -0.86 | B>B |
| chr10 | 1.18E+08 | 118000000 | -0.25 | -0.85 | A>B |
| chr10 | 1.18E+08 | 118100000 | -0.40 | -0.80 | B>B |
| chr10 | 1.18E+08 | 118200000 | -0.66 | -0.93 | B>B |
| chr10 | 1.18E+08 | 118300000 | -0.53 | -0.98 | B>B |
| chr10 | 1.18E+08 | 118400000 | -0.32 | -0.75 | B>B |
| chr10 | 1.18E+08 | 118500000 | 0.17 | -0.09 | A>A |
| chr10 | 1.19E+08 | 118600000 | 0.29 | 0.04 | A>A |
| chr10 | 1.19E+08 | 118700000 | 0.29 | 0.24 | A>A |
| chr10 | 1.19E+08 | 118800000 | 0.30 | 0.28 | A>A |
| chr10 | 1.19E+08 | 118900000 | 0.27 | 0.09 | A>A |
| chr10 | 1.19E+08 | 119000000 | 0.27 | 0.09 | A>A |
| chr10 | 1.19E+08 | 119100000 | 0.38 | 0.26 | A>A |
| chr10 | 1.19E+08 | 119200000 | 0.47 | 0.34 | A>A |
| chr10 | 1.19E+08 | 119300000 | 0.13 | -0.23 | B>B |
| chr10 | 1.19E+08 | 119400000 | -0.04 | -0.48 | B>B |
| chr10 | 1.19E+08 | 119500000 | -0.10 | -0.55 | B>B |
| chr10 | 1.2E+08 | 119600000 | -0.29 | -0.85 | B>B |
| chr10 | 1.2E+08 | 119700000 | -0.52 | -0.92 | B>B |
| chr10 | 1.2E+08 | 119800000 | -0.22 | -0.62 | B>B |
| chr10 | 1.2E+08 | 119900000 | -0.35 | -0.79 | B>B |
| chr10 | 1.2E+08 | 120000000 | -0.57 | -0.93 | B>B |
| chr10 | 1.2E+08 | 120100000 | -0.18 | -0.51 | B>B |
| chr10 | 1.2E+08 | 120200000 | -0.26 | -0.70 | B>B |
| chr10 | 1.2E+08 | 120300000 | -0.05 | -0.55 | B>B |
| chr10 | 1.2E+08 | 120400000 | 0.24 | -0.19 | A>A |
| chr10 | 1.2E+08 | 120500000 | 0.45 | 0.10 | A>A |
| chr10 | 1.21E+08 | 120600000 | 0.71 | 0.36 | A>A |
| chr10 | 1.21E+08 | 120700000 | 0.87 | 0.56 | A>A |

| | | | | | |
|---|---|---|---|---|---|
| chr10 | 1.21E+08 | 120800000 | 0.92 | 0.71 | A>A |
| chr10 | 1.21E+08 | 120900000 | 0.98 | 0.87 | A>A |
| chr10 | 1.21E+08 | 121000000 | 0.98 | 1.00 | A>A |
| chr10 | 1.21E+08 | 121100000 | 1.22 | 1.20 | A>A |
| chr10 | 1.21E+08 | 121200000 | 1.05 | 1.04 | A>A |
| chr10 | 1.21E+08 | 121300000 | 0.96 | 1.02 | A>A |
| chr10 | 1.21E+08 | 121400000 | 1.01 | 0.85 | A>A |
| chr10 | 1.21E+08 | 121500000 | 0.82 | 0.66 | A>A |
| chr10 | 1.22E+08 | 121600000 | 0.70 | 0.60 | A>A |
| chr10 | 1.22E+08 | 121700000 | 0.47 | 0.34 | A>A |
| chr10 | 1.22E+08 | 121800000 | -0.26 | -0.69 | B>B |
| chr10 | 1.22E+08 | 121900000 | -0.72 | -1.21 | B>B |
| chr10 | 1.22E+08 | 122000000 | -0.74 | -1.20 | B>B |
| chr10 | 1.22E+08 | 122100000 | -0.85 | -1.24 | B>B |
| chr10 | 1.22E+08 | 122200000 | -0.80 | -1.19 | B>B |
| chr10 | 1.22E+08 | 122300000 | -0.86 | -1.18 | B>B |
| chr10 | 1.22E+08 | 122400000 | -0.92 | -1.23 | B>B |
| chr10 | 1.22E+08 | 122500000 | -0.87 | -1.15 | B>B |
| chr10 | 1.23E+08 | 122600000 | -0.68 | -0.89 | B>B |
| chr10 | 1.23E+08 | 122700000 | -0.20 | 0.00 | B>B |
| chr10 | 1.23E+08 | 122800000 | -0.51 | 0.30 | B>A |
| chr10 | 1.23E+08 | 122900000 | -0.60 | 0.24 | B>A |
| chr10 | 1.23E+08 | 123000000 | -0.66 | 0.17 | B>A |
| chr10 | 1.23E+08 | 123100000 | -0.53 | 0.21 | B>A |
| chr10 | 1.23E+08 | 123200000 | -0.38 | 0.25 | B>A |
| chr10 | 1.23E+08 | 123300000 | -0.28 | 0.25 | B>A |
| chr10 | 1.23E+08 | 123400000 | -0.17 | 0.21 | A>A |
| chr10 | 1.23E+08 | 123500000 | -0.07 | 0.13 | A>A |
| chr10 | 1.24E+08 | 123600000 | 0.31 | 0.44 | A>A |
| chr10 | 1.24E+08 | 123700000 | 0.72 | 0.75 | A>A |
| chr10 | 1.24E+08 | 123800000 | 0.96 | 0.87 | A>A |
| chr10 | 1.24E+08 | 123900000 | 0.95 | 0.99 | A>A |
| chr10 | 1.24E+08 | 124000000 | 0.72 | 0.73 | A>A |
| chr10 | 1.24E+08 | 124100000 | 0.65 | 0.51 | A>A |
| chr10 | 1.24E+08 | 124200000 | 0.89 | 0.60 | A>A |
| chr10 | 1.24E+08 | 124300000 | 0.65 | 0.31 | A>A |
| chr10 | 1.24E+08 | 124400000 | 0.48 | -0.07 | A>A |
| chr10 | 1.24E+08 | 124500000 | 0.29 | -0.29 | B>B |
| chr10 | 1.25E+08 | 124600000 | 0.40 | -0.24 | A>B |
| chr10 | 1.25E+08 | 124700000 | 0.62 | 0.01 | A>B |
| chr10 | 1.25E+08 | 124800000 | 0.52 | 0.06 | A>A |
| chr10 | 1.25E+08 | 124900000 | 0.35 | -0.30 | A>B |
| chr10 | 1.25E+08 | 125000000 | 0.37 | -0.14 | A>A |
| chr10 | 1.25E+08 | 125100000 | 0.24 | -0.53 | A>B |
| chr10 | 1.25E+08 | 125200000 | 0.10 | -0.72 | A>B |
| chr10 | 1.25E+08 | 125300000 | 0.09 | -0.67 | A>B |
| chr10 | 1.25E+08 | 125400000 | 0.00 | -0.45 | B>B |
| chr10 | 1.25E+08 | 125500000 | -0.07 | -0.70 | A>B |
| chr10 | 1.26E+08 | 125600000 | 0.00 | -0.70 | A>B |
| chr10 | 1.26E+08 | 125700000 | 0.06 | -0.49 | B>B |
| chr10 | 1.26E+08 | 125800000 | 0.28 | 0.07 | A>A |
| chr10 | 1.26E+08 | 125900000 | 0.85 | 0.60 | A>A |
| chr10 | 1.26E+08 | 126000000 | 0.78 | 0.33 | A>A |
| chr10 | 1.26E+08 | 126100000 | 0.64 | 0.27 | A>A |
| chr10 | 1.26E+08 | 126200000 | 0.79 | 0.63 | A>A |
| chr10 | 1.26E+08 | 126300000 | 0.87 | 0.77 | A>A |

| | | | | | |
|---|---|---|---|---|---|
| chr10 | 1.26E+08 | 126400000 | 1.00 | 0.94 | A>A |
| chr10 | 1.26E+08 | 126500000 | 1.05 | 0.93 | A>A |
| chr10 | 1.27E+08 | 126600000 | 1.01 | 0.96 | A>A |
| chr10 | 1.27E+08 | 126700000 | 0.81 | 0.89 | A>A |
| chr10 | 1.27E+08 | 126800000 | 0.74 | 0.94 | A>A |
| chr10 | 1.27E+08 | 126900000 | 0.79 | 0.92 | A>A |
| chr10 | 1.27E+08 | 127000000 | 0.71 | 0.80 | A>A |
| chr10 | 1.27E+08 | 127100000 | 0.43 | 0.71 | A>A |
| chr10 | 1.27E+08 | 127200000 | 0.44 | 0.75 | A>A |
| chr10 | 1.27E+08 | 127300000 | 0.49 | 0.75 | A>A |
| chr10 | 1.27E+08 | 127400000 | 0.49 | 0.74 | A>A |
| chr10 | 1.27E+08 | 127500000 | 0.61 | 0.79 | A>A |
| chr10 | 1.28E+08 | 127600000 | 0.50 | 0.65 | A>A |
| chr10 | 1.28E+08 | 127700000 | 0.21 | 0.47 | A>A |
| chr10 | 1.28E+08 | 127800000 | 0.19 | 0.66 | A>A |
| chr10 | 1.28E+08 | 127900000 | 0.20 | 0.55 | A>A |
| chr10 | 1.28E+08 | 128000000 | 0.06 | 0.60 | B>A |
| chr10 | 1.28E+08 | 128100000 | 0.04 | 0.50 | A>A |
| chr10 | 1.28E+08 | 128200000 | -0.01 | 0.37 | A>A |
| chr10 | 1.28E+08 | 128300000 | -0.02 | 0.24 | A>A |
| chr10 | 1.28E+08 | 128400000 | -0.24 | 0.00 | B>B |
| chr10 | 1.28E+08 | 128500000 | -0.26 | 0.01 | B>B |
| chr10 | 1.29E+08 | 128600000 | -0.02 | 0.09 | A>A |
| chr10 | 1.29E+08 | 128700000 | 0.00 | 0.00 | NO |
| chr10 | 1.29E+08 | 128800000 | 0.00 | 0.00 | NO |
| chr10 | 1.29E+08 | 128900000 | 0.26 | 0.33 | A>A |
| chr10 | 1.29E+08 | 129000000 | 0.22 | 0.28 | A>A |
| chr10 | 1.29E+08 | 129100000 | 0.05 | 0.11 | A>A |
| chr10 | 1.29E+08 | 129200000 | -0.05 | 0.00 | B>B |
| chr10 | 1.29E+08 | 129300000 | -0.25 | -0.18 | B>B |
| chr10 | 1.29E+08 | 129400000 | -0.59 | -0.44 | B>B |
| chr10 | 1.29E+08 | 129500000 | -0.79 | -0.59 | B>B |
| chr10 | 1.3E+08 | 129600000 | -0.66 | -0.65 | B>B |
| chr10 | 1.3E+08 | 129700000 | -0.34 | -0.42 | B>B |
| chr10 | 1.3E+08 | 129800000 | 0.08 | -0.09 | B>B |
| chr10 | 1.3E+08 | 129900000 | 0.06 | -0.10 | B>B |
| chr10 | 1.3E+08 | 130000000 | 0.17 | -0.14 | A>A |
| chr10 | 1.3E+08 | 130100000 | -0.33 | -0.62 | B>B |
| chr10 | 1.3E+08 | 130200000 | -0.78 | -1.07 | B>B |
| chr10 | 1.3E+08 | 130300000 | -0.68 | -1.11 | B>B |
| chr10 | 1.3E+08 | 130400000 | -0.62 | -1.06 | B>B |
| chr10 | 1.3E+08 | 130500000 | -0.72 | -1.10 | B>B |
| chr10 | 1.31E+08 | 130600000 | -0.80 | -1.19 | B>B |
| chr10 | 1.31E+08 | 130700000 | -0.85 | -1.08 | B>B |
| chr10 | 1.31E+08 | 130800000 | -0.88 | -1.05 | B>B |
| chr10 | 1.31E+08 | 130900000 | -0.92 | -0.95 | B>B |
| chr10 | 1.31E+08 | 131000000 | -1.09 | -1.13 | B>B |
| chr10 | 1.31E+08 | 131100000 | -1.22 | -1.10 | B>B |
| chr10 | 1.31E+08 | 131200000 | -0.99 | -0.99 | B>B |
| chr10 | 1.31E+08 | 131300000 | -0.84 | -0.85 | B>B |
| chr10 | 1.31E+08 | 131400000 | -0.82 | -0.71 | B>B |
| chr10 | 1.31E+08 | 131500000 | -0.97 | -0.66 | B>B |
| chr10 | 1.32E+08 | 131600000 | -0.87 | -0.55 | B>B |
| chr10 | 1.32E+08 | 131700000 | -0.54 | -0.34 | B>B |
| chr10 | 1.32E+08 | 131800000 | -0.25 | -0.19 | B>B |
| chr10 | 1.32E+08 | 131900000 | -0.17 | -0.20 | B>B |

| | | | | | |
|---|---|---|---|---|---|
| chr10 | 1.32E+08 | 132000000 | -0.02 | 0.06 | A>A |
| chr10 | 1.32E+08 | 132100000 | -0.78 | -0.50 | B>B |
| chr10 | 1.32E+08 | 132200000 | -0.92 | -0.85 | B>B |
| chr10 | 1.32E+08 | 132300000 | -1.10 | -1.00 | B>B |
| chr10 | 1.32E+08 | 132400000 | -1.05 | -1.03 | B>B |
| chr10 | 1.32E+08 | 132500000 | -1.02 | -1.10 | B>B |
| chr10 | 1.33E+08 | 132600000 | -0.92 | -0.99 | B>B |
| chr10 | 1.33E+08 | 132700000 | -1.05 | -1.03 | B>B |
| chr10 | 1.33E+08 | 132800000 | -1.03 | -1.05 | B>B |
| chr10 | 1.33E+08 | 132900000 | -0.89 | -0.83 | B>B |
| chr10 | 1.33E+08 | 133000000 | -0.81 | -0.76 | B>B |
| chr10 | 1.33E+08 | 133100000 | -0.63 | -0.60 | B>B |
| chr10 | 1.33E+08 | 133200000 | -0.58 | -0.38 | B>B |
| chr10 | 1.33E+08 | 133300000 | -0.38 | -0.02 | B>B |
| chr10 | 1.33E+08 | 133400000 | -0.32 | 0.37 | B>A |
| chr10 | 1.33E+08 | 133500000 | -0.12 | 0.54 | B>A |
| chr10 | 1.34E+08 | 133600000 | -0.02 | 0.34 | A>A |
| chr10 | 1.34E+08 | 133700000 | 0.02 | 0.17 | A>A |
| chr10 | 1.34E+08 | 133800000 | 0.89 | 0.96 | A>A |
| chr10 | 1.34E+08 | 133900000 | 1.05 | 1.18 | A>A |
| chr10 | 1.34E+08 | 134000000 | 1.15 | 1.22 | A>A |
| chr10 | 1.34E+08 | 134100000 | 1.36 | 1.29 | A>A |
| chr10 | 1.34E+08 | 134200000 | 1.49 | 1.37 | A>A |
| chr10 | 1.34E+08 | 134300000 | 1.31 | 1.28 | A>A |
| chr10 | 1.34E+08 | 134400000 | 1.50 | 1.37 | A>A |
| chr10 | 1.34E+08 | 134500000 | 1.19 | 1.14 | A>A |
| chr10 | 1.35E+08 | 134600000 | 1.38 | 1.29 | A>A |
| chr10 | 1.35E+08 | 134700000 | 1.22 | 1.17 | A>A |
| chr10 | 1.35E+08 | 134800000 | 1.24 | 1.17 | A>A |
| chr10 | 1.35E+08 | 134900000 | 1.42 | 1.35 | A>A |
| chr10 | 1.35E+08 | 135000000 | 1.27 | 1.24 | A>A |
| chr10 | 1.35E+08 | 135100000 | 1.44 | 1.32 | A>A |
| chr10 | 1.35E+08 | 135200000 | 1.53 | 1.35 | A>A |
| chr10 | 1.35E+08 | 135300000 | 1.20 | 1.13 | A>A |
| chr10 | 1.35E+08 | 135400000 | 0.87 | 0.95 | A>A |
| chr10 | 1.35E+08 | 135500000 | 0.00 | 0.00 | NO |
| chr10 | 1.36E+08 | 135534747 | 0.00 | 0.00 | NO |
| chr11 | 0 | 100000 | 0.00 | 0.00 | NO |
| chr11 | 100000 | 200000 | 0.00 | 0.00 | NO |
| chr11 | 200000 | 300000 | 1.55 | 1.31 | A>A |
| chr11 | 300000 | 400000 | 1.61 | 1.37 | A>A |
| chr11 | 400000 | 500000 | 1.43 | 1.37 | A>A |
| chr11 | 500000 | 600000 | 1.51 | 1.40 | A>A |
| chr11 | 600000 | 700000 | 1.62 | 1.48 | A>A |
| chr11 | 700000 | 800000 | 1.59 | 1.43 | A>A |
| chr11 | 800000 | 900000 | 1.66 | 1.37 | A>A |
| chr11 | 900000 | 1000000 | 1.75 | 1.53 | A>A |
| chr11 | 1000000 | 1100000 | 1.45 | 1.27 | A>A |
| chr11 | 1100000 | 1200000 | 0.00 | 0.00 | NO |
| chr11 | 1200000 | 1300000 | 1.36 | 1.26 | A>A |
| chr11 | 1300000 | 1400000 | 1.57 | 1.52 | A>A |
| chr11 | 1400000 | 1500000 | 1.64 | 1.55 | A>A |
| chr11 | 1500000 | 1600000 | 1.43 | 1.46 | A>A |
| chr11 | 1600000 | 1700000 | 1.20 | 1.36 | A>A |
| chr11 | 1700000 | 1800000 | 1.41 | 1.50 | A>A |
| chr11 | 1800000 | 1900000 | 1.47 | 1.52 | A>A |

| | | | | | |
|---|---|---|---|---|---|
| chr11 | 1900000 | 2000000 | 1.29 | 1.45 | A>A |
| chr11 | 2000000 | 2100000 | 1.30 | 1.35 | A>A |
| chr11 | 2100000 | 2200000 | 1.23 | 1.27 | A>A |
| chr11 | 2200000 | 2300000 | 1.32 | 1.38 | A>A |
| chr11 | 2300000 | 2400000 | 1.47 | 1.47 | A>A |
| chr11 | 2400000 | 2500000 | 1.12 | 1.37 | A>A |
| chr11 | 2500000 | 2600000 | 0.91 | 1.14 | A>A |
| chr11 | 2600000 | 2700000 | 0.79 | 1.37 | B>A |
| chr11 | 2700000 | 2800000 | 1.12 | 1.45 | A>A |
| chr11 | 2800000 | 2900000 | 1.12 | 1.44 | A>A |
| chr11 | 2900000 | 3000000 | 1.18 | 1.41 | A>A |
| chr11 | 3000000 | 3100000 | 1.44 | 1.85 | A>A |
| chr11 | 3100000 | 3200000 | 1.50 | 1.83 | A>A |
| chr11 | 3200000 | 3300000 | 1.19 | 1.71 | B>A |
| chr11 | 3300000 | 3400000 | 0.00 | 0.00 | NO |
| chr11 | 3400000 | 3500000 | 0.00 | 0.00 | NO |
| chr11 | 3500000 | 3600000 | 0.00 | 0.00 | NO |
| chr11 | 3600000 | 3700000 | 1.32 | 1.49 | A>A |
| chr11 | 3700000 | 3800000 | 1.32 | 1.52 | A>A |
| chr11 | 3800000 | 3900000 | 1.02 | 1.15 | A>A |
| chr11 | 3900000 | 4000000 | 0.89 | 0.94 | A>A |
| chr11 | 4000000 | 4100000 | 0.62 | 0.83 | A>A |
| chr11 | 4100000 | 4200000 | 0.33 | 0.31 | A>A |
| chr11 | 4200000 | 4300000 | 0.00 | -0.66 | NO |
| chr11 | 4300000 | 4400000 | 0.00 | 0.00 | NO |
| chr11 | 4400000 | 4500000 | -0.78 | -1.12 | B>B |
| chr11 | 4500000 | 4600000 | -0.67 | -0.80 | B>B |
| chr11 | 4600000 | 4700000 | -0.61 | -0.68 | B>B |
| chr11 | 4700000 | 4800000 | -1.03 | -1.15 | B>B |
| chr11 | 4800000 | 4900000 | -1.11 | -1.34 | B>B |
| chr11 | 4900000 | 5000000 | -1.20 | -1.22 | B>B |
| chr11 | 5000000 | 5100000 | -1.02 | -1.21 | B>B |
| chr11 | 5100000 | 5200000 | -0.78 | -1.03 | B>B |
| chr11 | 5200000 | 5300000 | -0.52 | -0.87 | B>B |
| chr11 | 5300000 | 5400000 | -0.61 | -0.93 | B>B |
| chr11 | 5400000 | 5500000 | -0.75 | -0.97 | B>B |
| chr11 | 5500000 | 5600000 | -0.55 | -0.78 | B>B |
| chr11 | 5600000 | 5700000 | -0.08 | -0.24 | B>B |
| chr11 | 5700000 | 5800000 | -0.44 | -0.69 | B>B |
| chr11 | 5800000 | 5900000 | -0.79 | -0.76 | B>B |
| chr11 | 5900000 | 6000000 | -0.83 | -1.02 | B>B |
| chr11 | 6000000 | 6100000 | -0.84 | -1.09 | B>B |
| chr11 | 6100000 | 6200000 | -0.82 | -1.04 | B>B |
| chr11 | 6200000 | 6300000 | -0.21 | -0.26 | B>B |
| chr11 | 6300000 | 6400000 | -0.09 | -0.02 | B>B |
| chr11 | 6400000 | 6500000 | 0.51 | 0.55 | A>A |
| chr11 | 6500000 | 6600000 | 0.40 | 0.44 | A>A |
| chr11 | 6600000 | 6700000 | 0.48 | 0.37 | A>A |
| chr11 | 6700000 | 6800000 | -0.34 | -0.56 | B>B |
| chr11 | 6800000 | 6900000 | -0.80 | -1.06 | B>B |
| chr11 | 6900000 | 7000000 | -0.79 | -0.97 | B>B |
| chr11 | 7000000 | 7100000 | -0.75 | -0.93 | B>B |
| chr11 | 7100000 | 7200000 | -0.72 | -0.88 | B>B |
| chr11 | 7200000 | 7300000 | -0.70 | -1.12 | B>B |
| chr11 | 7300000 | 7400000 | -0.70 | -1.06 | B>B |
| chr11 | 7400000 | 7500000 | -0.56 | -0.89 | B>B |

| | | | | | |
|---|---|---|---|---|---|
| chr11 | 7500000 | 7600000 | 0.05 | -0.43 | B>B |
| chr11 | 7600000 | 7700000 | 0.24 | -0.11 | A>A |
| chr11 | 7700000 | 7800000 | -0.69 | -0.85 | B>B |
| chr11 | 7800000 | 7900000 | -0.86 | -1.13 | B>B |
| chr11 | 7900000 | 8000000 | -0.51 | -0.87 | B>B |
| chr11 | 8000000 | 8100000 | 0.09 | -0.15 | B>B |
| chr11 | 8100000 | 8200000 | 0.08 | -0.04 | A>A |
| chr11 | 8200000 | 8300000 | 0.27 | 0.19 | A>A |
| chr11 | 8300000 | 8400000 | 0.26 | 0.02 | A>A |
| chr11 | 8400000 | 8500000 | -0.02 | -0.19 | B>B |
| chr11 | 8500000 | 8600000 | 0.15 | -0.04 | A>A |
| chr11 | 8600000 | 8700000 | 0.62 | 0.48 | A>A |
| chr11 | 8700000 | 8800000 | 0.84 | 0.83 | A>A |
| chr11 | 8800000 | 8900000 | 0.88 | 0.79 | A>A |
| chr11 | 8900000 | 9000000 | 0.98 | 0.93 | A>A |
| chr11 | 9000000 | 9100000 | 0.96 | 0.92 | A>A |
| chr11 | 9100000 | 9200000 | 0.92 | 0.97 | A>A |
| chr11 | 9200000 | 9300000 | 1.11 | 1.15 | A>A |
| chr11 | 9300000 | 9400000 | 1.32 | 1.32 | A>A |
| chr11 | 9400000 | 9500000 | 1.16 | 1.14 | A>A |
| chr11 | 9500000 | 9600000 | 0.98 | 1.10 | A>A |
| chr11 | 9600000 | 9700000 | 1.02 | 1.05 | A>A |
| chr11 | 9700000 | 9800000 | 0.90 | 0.98 | A>A |
| chr11 | 9800000 | 9900000 | 0.89 | 1.00 | A>A |
| chr11 | 9900000 | 10000000 | 0.82 | 0.90 | A>A |
| chr11 | 10000000 | 10100000 | 0.83 | 0.90 | A>A |
| chr11 | 10100000 | 10200000 | 0.86 | 0.95 | A>A |
| chr11 | 10200000 | 10300000 | 0.86 | 0.90 | A>A |
| chr11 | 10300000 | 10400000 | 0.72 | 0.73 | A>A |
| chr11 | 10400000 | 10500000 | 0.87 | 0.75 | A>A |
| chr11 | 10500000 | 10600000 | 1.07 | 0.90 | A>A |
| chr11 | 10600000 | 10700000 | 1.01 | 0.84 | A>A |
| chr11 | 10700000 | 10800000 | 0.79 | 0.68 | A>A |
| chr11 | 10800000 | 10900000 | 1.03 | 0.58 | A>A |
| chr11 | 10900000 | 11000000 | 1.11 | 0.52 | A>B |
| chr11 | 11000000 | 11100000 | 0.64 | 0.13 | A>A |
| chr11 | 11100000 | 11200000 | 0.66 | -0.03 | A>B |
| chr11 | 11200000 | 11300000 | 0.64 | -0.05 | A>B |
| chr11 | 11300000 | 11400000 | 0.71 | 0.03 | A>B |
| chr11 | 11400000 | 11500000 | 0.77 | 0.11 | A>B |
| chr11 | 11500000 | 11600000 | 0.85 | 0.09 | A>B |
| chr11 | 11600000 | 11700000 | 0.74 | 0.09 | A>B |
| chr11 | 11700000 | 11800000 | 0.52 | -0.03 | A>A |
| chr11 | 11800000 | 11900000 | 0.66 | 0.37 | A>A |
| chr11 | 11900000 | 12000000 | 0.84 | 0.60 | A>A |
| chr11 | 12000000 | 12100000 | 0.85 | 0.71 | A>A |
| chr11 | 12100000 | 12200000 | 1.01 | 0.88 | A>A |
| chr11 | 12200000 | 12300000 | 0.85 | 0.71 | A>A |
| chr11 | 12300000 | 12400000 | 0.54 | 0.55 | A>A |
| chr11 | 12400000 | 12500000 | 0.68 | 0.80 | A>A |
| chr11 | 12500000 | 12600000 | 0.73 | 0.73 | A>A |
| chr11 | 12600000 | 12700000 | 0.67 | 0.75 | A>A |
| chr11 | 12700000 | 12800000 | 0.71 | 0.90 | A>A |
| chr11 | 12800000 | 12900000 | 0.66 | 0.75 | A>A |
| chr11 | 12900000 | 13000000 | 0.40 | 0.63 | A>A |
| chr11 | 13000000 | 13100000 | 0.86 | 0.95 | A>A |

| | | | | | |
|---|---|---|---|---|---|
| chr11 | 13100000 | 13200000 | 0.73 | 0.97 | A>A |
| chr11 | 13200000 | 13300000 | 0.58 | 0.76 | A>A |
| chr11 | 13300000 | 13400000 | 0.61 | 0.80 | A>A |
| chr11 | 13400000 | 13500000 | 0.29 | 0.56 | A>A |
| chr11 | 13500000 | 13600000 | -0.32 | 0.18 | B>B |
| chr11 | 13600000 | 13700000 | -0.38 | 0.03 | B>B |
| chr11 | 13700000 | 13800000 | -0.08 | 0.21 | A>A |
| chr11 | 13800000 | 13900000 | -0.32 | 0.09 | B>B |
| chr11 | 13900000 | 14000000 | -0.21 | 0.16 | B>B |
| chr11 | 14000000 | 14100000 | -0.16 | 0.02 | B>B |
| chr11 | 14100000 | 14200000 | -0.29 | -0.02 | B>B |
| chr11 | 14200000 | 14300000 | -0.09 | 0.04 | B>B |
| chr11 | 14300000 | 14400000 | 0.52 | 0.53 | A>A |
| chr11 | 14400000 | 14500000 | 0.53 | 0.51 | A>A |
| chr11 | 14500000 | 14600000 | 0.10 | -0.06 | A>A |
| chr11 | 14600000 | 14700000 | -0.26 | -0.44 | B>B |
| chr11 | 14700000 | 14800000 | -0.32 | -0.47 | B>B |
| chr11 | 14800000 | 14900000 | -0.26 | -0.49 | B>B |
| chr11 | 14900000 | 15000000 | -0.43 | -0.56 | B>B |
| chr11 | 15000000 | 15100000 | -0.59 | -0.83 | B>B |
| chr11 | 15100000 | 15200000 | -0.53 | -0.84 | B>B |
| chr11 | 15200000 | 15300000 | -0.63 | -0.98 | B>B |
| chr11 | 15300000 | 15400000 | -0.81 | -1.00 | B>B |
| chr11 | 15400000 | 15500000 | -1.00 | -1.14 | B>B |
| chr11 | 15500000 | 15600000 | -0.70 | -0.98 | B>B |
| chr11 | 15600000 | 15700000 | -0.47 | -0.65 | B>B |
| chr11 | 15700000 | 15800000 | -0.39 | -0.54 | B>B |
| chr11 | 15800000 | 15900000 | -0.15 | -0.16 | B>B |
| chr11 | 15900000 | 16000000 | 0.04 | 0.16 | A>A |
| chr11 | 16000000 | 16100000 | -0.01 | 0.21 | A>A |
| chr11 | 16100000 | 16200000 | 0.05 | 0.22 | A>A |
| chr11 | 16200000 | 16300000 | 0.01 | 0.25 | A>A |
| chr11 | 16300000 | 16400000 | -0.17 | 0.21 | A>A |
| chr11 | 16400000 | 16500000 | -0.33 | 0.17 | B>B |
| chr11 | 16500000 | 16600000 | -0.47 | 0.06 | B>A |
| chr11 | 16600000 | 16700000 | -0.39 | -0.06 | B>B |
| chr11 | 16700000 | 16800000 | -0.06 | 0.14 | A>A |
| chr11 | 16800000 | 16900000 | 0.46 | 0.63 | A>A |
| chr11 | 16900000 | 17000000 | 0.68 | 0.77 | A>A |
| chr11 | 17000000 | 17100000 | 0.78 | 0.85 | A>A |
| chr11 | 17100000 | 17200000 | 0.82 | 0.90 | A>A |
| chr11 | 17200000 | 17300000 | 0.84 | 0.92 | A>A |
| chr11 | 17300000 | 17400000 | 0.91 | 0.83 | A>A |
| chr11 | 17400000 | 17500000 | 0.94 | 0.81 | A>A |
| chr11 | 17500000 | 17600000 | 0.64 | 0.55 | A>A |
| chr11 | 17600000 | 17700000 | 0.39 | 0.17 | A>A |
| chr11 | 17700000 | 17800000 | 0.38 | 0.17 | A>A |
| chr11 | 17800000 | 17900000 | 0.34 | 0.19 | A>A |
| chr11 | 17900000 | 18000000 | 0.33 | 0.30 | A>A |
| chr11 | 18000000 | 18100000 | 0.43 | 0.21 | A>A |
| chr11 | 18100000 | 18200000 | 0.37 | 0.07 | A>A |
| chr11 | 18200000 | 18300000 | 0.39 | 0.04 | A>A |
| chr11 | 18300000 | 18400000 | 0.85 | 0.62 | A>A |
| chr11 | 18400000 | 18500000 | 0.85 | 0.70 | A>A |
| chr11 | 18500000 | 18600000 | 0.69 | 0.65 | A>A |
| chr11 | 18600000 | 18700000 | 0.76 | 0.54 | A>A |

| | | | | | |
|---|---|---|---|---|---|
| chr11 | 18700000 | 18800000 | 0.56 | 0.20 | A>A |
| chr11 | 18800000 | 18900000 | -0.03 | -0.41 | B>B |
| chr11 | 18900000 | 19000000 | -0.39 | -0.65 | B>B |
| chr11 | 19000000 | 19100000 | -0.32 | -0.51 | B>B |
| chr11 | 19100000 | 19200000 | 0.18 | -0.12 | A>A |
| chr11 | 19200000 | 19300000 | 0.20 | -0.21 | B>B |
| chr11 | 19300000 | 19400000 | 0.27 | -0.36 | A>B |
| chr11 | 19400000 | 19500000 | 0.41 | -0.33 | A>B |
| chr11 | 19500000 | 19600000 | 0.26 | -0.24 | A>A |
| chr11 | 19600000 | 19700000 | 0.40 | -0.10 | A>A |
| chr11 | 19700000 | 19800000 | 0.36 | 0.02 | A>A |
| chr11 | 19800000 | 19900000 | 0.26 | -0.17 | A>A |
| chr11 | 19900000 | 20000000 | 0.26 | 0.03 | A>A |
| chr11 | 20000000 | 20100000 | 0.72 | 0.48 | A>A |
| chr11 | 20100000 | 20200000 | 0.28 | -0.04 | A>A |
| chr11 | 20200000 | 20300000 | -0.20 | -0.47 | B>B |
| chr11 | 20300000 | 20400000 | -0.04 | -0.31 | B>B |
| chr11 | 20400000 | 20500000 | 0.22 | 0.08 | A>A |
| chr11 | 20500000 | 20600000 | -0.06 | -0.23 | B>B |
| chr11 | 20600000 | 20700000 | -0.34 | -0.46 | B>B |
| chr11 | 20700000 | 20800000 | -0.76 | -0.83 | B>B |
| chr11 | 20800000 | 20900000 | -0.93 | -0.91 | B>B |
| chr11 | 20900000 | 21000000 | -0.91 | -0.87 | B>B |
| chr11 | 21000000 | 21100000 | -0.98 | -0.85 | B>B |
| chr11 | 21100000 | 21200000 | -0.88 | -0.80 | B>B |
| chr11 | 21200000 | 21300000 | -0.92 | -0.74 | B>B |
| chr11 | 21300000 | 21400000 | -0.84 | -0.60 | B>B |
| chr11 | 21400000 | 21500000 | -0.82 | -0.31 | B>B |
| chr11 | 21500000 | 21600000 | -0.69 | -0.09 | B>A |
| chr11 | 21600000 | 21700000 | -0.68 | -0.04 | B>A |
| chr11 | 21700000 | 21800000 | -0.75 | -0.08 | B>A |
| chr11 | 21800000 | 21900000 | -0.74 | -0.13 | B>A |
| chr11 | 21900000 | 22000000 | -0.41 | -0.06 | B>B |
| chr11 | 22000000 | 22100000 | -0.50 | -0.15 | B>B |
| chr11 | 22100000 | 22200000 | -0.52 | -0.22 | B>B |
| chr11 | 22200000 | 22300000 | -0.17 | -0.24 | B>B |
| chr11 | 22300000 | 22400000 | -0.53 | -0.60 | B>B |
| chr11 | 22400000 | 22500000 | -0.84 | -0.91 | B>B |
| chr11 | 22500000 | 22600000 | -0.81 | -0.91 | B>B |
| chr11 | 22600000 | 22700000 | -0.56 | -0.79 | B>B |
| chr11 | 22700000 | 22800000 | -0.56 | -0.77 | B>B |
| chr11 | 22800000 | 22900000 | -0.51 | -0.54 | B>B |
| chr11 | 22900000 | 23000000 | -0.77 | -0.69 | B>B |
| chr11 | 23000000 | 23100000 | -0.91 | -0.84 | B>B |
| chr11 | 23100000 | 23200000 | -0.88 | -0.85 | B>B |
| chr11 | 23200000 | 23300000 | -0.89 | -0.88 | B>B |
| chr11 | 23300000 | 23400000 | -0.86 | -0.85 | B>B |
| chr11 | 23400000 | 23500000 | -0.79 | -0.79 | B>B |
| chr11 | 23500000 | 23600000 | -0.77 | -0.64 | B>B |
| chr11 | 23600000 | 23700000 | -0.74 | -0.61 | B>B |
| chr11 | 23700000 | 23800000 | -0.81 | -0.59 | B>B |
| chr11 | 23800000 | 23900000 | -0.80 | -0.60 | B>B |
| chr11 | 23900000 | 24000000 | -0.76 | -0.59 | B>B |
| chr11 | 24000000 | 24100000 | -0.78 | -0.62 | B>B |
| chr11 | 24100000 | 24200000 | -0.70 | -0.68 | B>B |
| chr11 | 24200000 | 24300000 | -0.81 | -0.71 | B>B |

| | | | | | |
|---|---|---|---|---|---|
| chr11 | 24300000 | 24400000 | -0.75 | -0.69 | B>B |
| chr11 | 24400000 | 24500000 | -0.79 | -0.63 | B>B |
| chr11 | 24500000 | 24600000 | -0.75 | -0.56 | B>B |
| chr11 | 24600000 | 24700000 | -0.79 | -0.50 | B>B |
| chr11 | 24700000 | 24800000 | -0.74 | -0.56 | B>B |
| chr11 | 24800000 | 24900000 | -0.84 | -0.56 | B>B |
| chr11 | 24900000 | 25000000 | -0.73 | -0.56 | B>B |
| chr11 | 25000000 | 25100000 | -0.78 | -0.50 | B>B |
| chr11 | 25100000 | 25200000 | -0.80 | -0.53 | B>B |
| chr11 | 25200000 | 25300000 | -0.78 | -0.58 | B>B |
| chr11 | 25300000 | 25400000 | -0.73 | -0.64 | B>B |
| chr11 | 25400000 | 25500000 | -0.80 | -0.70 | B>B |
| chr11 | 25500000 | 25600000 | -0.78 | -0.73 | B>B |
| chr11 | 25600000 | 25700000 | -0.85 | -0.76 | B>B |
| chr11 | 25700000 | 25800000 | -0.79 | -0.73 | B>B |
| chr11 | 25800000 | 25900000 | -0.84 | -0.80 | B>B |
| chr11 | 25900000 | 26000000 | -0.81 | -0.76 | B>B |
| chr11 | 26000000 | 26100000 | -0.85 | -0.78 | B>B |
| chr11 | 26100000 | 26200000 | -0.86 | -0.73 | B>B |
| chr11 | 26200000 | 26300000 | -0.77 | -0.57 | B>B |
| chr11 | 26300000 | 26400000 | -0.73 | -0.34 | B>B |
| chr11 | 26400000 | 26500000 | -0.90 | -0.24 | B>A |
| chr11 | 26500000 | 26600000 | -0.79 | -0.02 | B>A |
| chr11 | 26600000 | 26700000 | -0.69 | -0.21 | B>B |
| chr11 | 26700000 | 26800000 | -0.63 | -0.57 | B>B |
| chr11 | 26800000 | 26900000 | -0.80 | -0.85 | B>B |
| chr11 | 26900000 | 27000000 | -0.81 | -0.89 | B>B |
| chr11 | 27000000 | 27100000 | -0.68 | -0.77 | B>B |
| chr11 | 27100000 | 27200000 | -0.48 | -0.49 | B>B |
| chr11 | 27200000 | 27300000 | 0.28 | 0.24 | A>A |
| chr11 | 27300000 | 27400000 | 0.33 | 0.35 | A>A |
| chr11 | 27400000 | 27500000 | 0.27 | 0.45 | A>A |
| chr11 | 27500000 | 27600000 | -0.16 | -0.15 | B>B |
| chr11 | 27600000 | 27700000 | -0.15 | -0.17 | B>B |
| chr11 | 27700000 | 27800000 | 0.13 | 0.31 | A>A |
| chr11 | 27800000 | 27900000 | 0.31 | 0.42 | A>A |
| chr11 | 27900000 | 28000000 | 0.37 | 0.55 | A>A |
| chr11 | 28000000 | 28100000 | 0.35 | 0.53 | A>A |
| chr11 | 28100000 | 28200000 | 0.33 | 0.49 | A>A |
| chr11 | 28200000 | 28300000 | 0.30 | 0.48 | A>A |
| chr11 | 28300000 | 28400000 | 0.25 | 0.42 | A>A |
| chr11 | 28400000 | 28500000 | 0.25 | 0.38 | A>A |
| chr11 | 28500000 | 28600000 | 0.28 | 0.39 | A>A |
| chr11 | 28600000 | 28700000 | 0.32 | 0.46 | A>A |
| chr11 | 28700000 | 28800000 | 0.30 | 0.44 | A>A |
| chr11 | 28800000 | 28900000 | 0.30 | 0.49 | A>A |
| chr11 | 28900000 | 29000000 | 0.37 | 0.51 | A>A |
| chr11 | 29000000 | 29100000 | 0.27 | 0.42 | A>A |
| chr11 | 29100000 | 29200000 | 0.28 | 0.40 | A>A |
| chr11 | 29200000 | 29300000 | 0.20 | 0.19 | A>A |
| chr11 | 29300000 | 29400000 | -0.02 | -0.02 | B>B |
| chr11 | 29400000 | 29500000 | -0.48 | -0.56 | B>B |
| chr11 | 29500000 | 29600000 | -0.70 | -0.75 | B>B |
| chr11 | 29600000 | 29700000 | -0.76 | -0.94 | B>B |
| chr11 | 29700000 | 29800000 | -0.86 | -0.88 | B>B |
| chr11 | 29800000 | 29900000 | -0.88 | -0.93 | B>B |

| | | | | | |
|---|---|---|---|---|---|
| chr11 | 29900000 | 30000000 | -0.84 | -0.92 | B>B |
| chr11 | 30000000 | 30100000 | -0.87 | -0.89 | B>B |
| chr11 | 30100000 | 30200000 | -0.69 | -0.87 | B>B |
| chr11 | 30200000 | 30300000 | -0.69 | -0.80 | B>B |
| chr11 | 30300000 | 30400000 | -0.53 | -0.61 | B>B |
| chr11 | 30400000 | 30500000 | -0.61 | -0.86 | B>B |
| chr11 | 30500000 | 30600000 | -0.76 | -0.91 | B>B |
| chr11 | 30600000 | 30700000 | -0.79 | -0.93 | B>B |
| chr11 | 30700000 | 30800000 | -0.90 | -0.96 | B>B |
| chr11 | 30800000 | 30900000 | -0.81 | -0.89 | B>B |
| chr11 | 30900000 | 31000000 | -0.66 | -0.58 | B>B |
| chr11 | 31000000 | 31100000 | -0.71 | -0.67 | B>B |
| chr11 | 31100000 | 31200000 | -0.86 | -0.80 | B>B |
| chr11 | 31200000 | 31300000 | -0.72 | -0.75 | B>B |
| chr11 | 31300000 | 31400000 | -0.61 | -0.64 | B>B |
| chr11 | 31400000 | 31500000 | -0.12 | -0.17 | B>B |
| chr11 | 31500000 | 31600000 | -0.13 | -0.11 | B>B |
| chr11 | 31600000 | 31700000 | -0.24 | -0.28 | B>B |
| chr11 | 31700000 | 31800000 | -0.38 | -0.38 | B>B |
| chr11 | 31800000 | 31900000 | -0.25 | -0.47 | B>B |
| chr11 | 31900000 | 32000000 | -0.43 | -0.73 | B>B |
| chr11 | 32000000 | 32100000 | -0.21 | -0.57 | B>B |
| chr11 | 32100000 | 32200000 | -0.05 | -0.48 | B>B |
| chr11 | 32200000 | 32300000 | -0.18 | -0.58 | B>B |
| chr11 | 32300000 | 32400000 | 0.14 | -0.28 | B>B |
| chr11 | 32400000 | 32500000 | 0.42 | -0.11 | A>A |
| chr11 | 32500000 | 32600000 | 0.42 | -0.06 | A>A |
| chr11 | 32600000 | 32700000 | 0.39 | 0.14 | A>A |
| chr11 | 32700000 | 32800000 | 0.44 | 0.19 | A>A |
| chr11 | 32800000 | 32900000 | 0.54 | 0.39 | A>A |
| chr11 | 32900000 | 33000000 | 0.70 | 0.67 | A>A |
| chr11 | 33000000 | 33100000 | 0.82 | 0.79 | A>A |
| chr11 | 33100000 | 33200000 | 0.76 | 0.76 | A>A |
| chr11 | 33200000 | 33300000 | 0.89 | 0.80 | A>A |
| chr11 | 33300000 | 33400000 | 0.74 | 0.72 | A>A |
| chr11 | 33400000 | 33500000 | 0.70 | 0.46 | A>A |
| chr11 | 33500000 | 33600000 | 0.81 | 0.48 | A>A |
| chr11 | 33600000 | 33700000 | 0.87 | 0.58 | A>A |
| chr11 | 33700000 | 33800000 | 0.80 | 0.72 | A>A |
| chr11 | 33800000 | 33900000 | 0.82 | 0.70 | A>A |
| chr11 | 33900000 | 34000000 | 0.94 | 0.80 | A>A |
| chr11 | 34000000 | 34100000 | 1.10 | 0.94 | A>A |
| chr11 | 34100000 | 34200000 | 0.89 | 0.85 | A>A |
| chr11 | 34200000 | 34300000 | 0.99 | 1.02 | A>A |
| chr11 | 34300000 | 34400000 | 0.85 | 0.92 | A>A |
| chr11 | 34400000 | 34500000 | 0.79 | 0.82 | A>A |
| chr11 | 34500000 | 34600000 | 0.69 | 0.70 | A>A |
| chr11 | 34600000 | 34700000 | 1.04 | 1.00 | A>A |
| chr11 | 34700000 | 34800000 | 1.15 | 0.94 | A>A |
| chr11 | 34800000 | 34900000 | 1.01 | 0.84 | A>A |
| chr11 | 34900000 | 35000000 | 0.56 | 0.59 | A>A |
| chr11 | 35000000 | 35100000 | 0.74 | 0.81 | A>A |
| chr11 | 35100000 | 35200000 | 0.85 | 0.87 | A>A |
| chr11 | 35200000 | 35300000 | 0.61 | 0.73 | A>A |
| chr11 | 35300000 | 35400000 | 0.69 | 0.77 | A>A |
| chr11 | 35400000 | 35500000 | 0.47 | 0.49 | A>A |

| | | | | | |
|---|---|---|---|---|---|
| chr11 | 35500000 | 35600000 | 0.48 | 0.35 | A>A |
| chr11 | 35600000 | 35700000 | 0.58 | 0.42 | A>A |
| chr11 | 35700000 | 35800000 | 0.48 | 0.51 | A>A |
| chr11 | 35800000 | 35900000 | 0.30 | 0.38 | A>A |
| chr11 | 35900000 | 36000000 | 0.46 | 0.45 | A>A |
| chr11 | 36000000 | 36100000 | 0.39 | 0.39 | A>A |
| chr11 | 36100000 | 36200000 | 0.37 | 0.38 | A>A |
| chr11 | 36200000 | 36300000 | 0.22 | 0.29 | A>A |
| chr11 | 36300000 | 36400000 | 0.32 | 0.20 | A>A |
| chr11 | 36400000 | 36500000 | 0.29 | 0.23 | A>A |
| chr11 | 36500000 | 36600000 | -0.05 | -0.20 | B>B |
| chr11 | 36600000 | 36700000 | -0.35 | -0.44 | B>B |
| chr11 | 36700000 | 36800000 | -0.65 | -0.76 | B>B |
| chr11 | 36800000 | 36900000 | -0.90 | -0.90 | B>B |
| chr11 | 36900000 | 37000000 | -0.94 | -0.93 | B>B |
| chr11 | 37000000 | 37100000 | -0.86 | -0.93 | B>B |
| chr11 | 37100000 | 37200000 | -0.90 | -0.92 | B>B |
| chr11 | 37200000 | 37300000 | -0.89 | -0.91 | B>B |
| chr11 | 37300000 | 37400000 | -0.88 | -0.83 | B>B |
| chr11 | 37400000 | 37500000 | -0.85 | -0.85 | B>B |
| chr11 | 37500000 | 37600000 | -0.86 | -0.85 | B>B |
| chr11 | 37600000 | 37700000 | -0.82 | -0.75 | B>B |
| chr11 | 37700000 | 37800000 | -0.83 | -0.73 | B>B |
| chr11 | 37800000 | 37900000 | -0.83 | -0.71 | B>B |
| chr11 | 37900000 | 38000000 | -0.77 | -0.60 | B>B |
| chr11 | 38000000 | 38100000 | -0.75 | -0.63 | B>B |
| chr11 | 38100000 | 38200000 | -0.75 | -0.64 | B>B |
| chr11 | 38200000 | 38300000 | -0.77 | -0.57 | B>B |
| chr11 | 38300000 | 38400000 | -0.74 | -0.54 | B>B |
| chr11 | 38400000 | 38500000 | -0.77 | -0.57 | B>B |
| chr11 | 38500000 | 38600000 | -0.74 | -0.56 | B>B |
| chr11 | 38600000 | 38700000 | -0.71 | -0.47 | B>B |
| chr11 | 38700000 | 38800000 | -0.71 | -0.47 | B>B |
| chr11 | 38800000 | 38900000 | -0.68 | -0.44 | B>B |
| chr11 | 38900000 | 39000000 | -0.72 | -0.39 | B>B |
| chr11 | 39000000 | 39100000 | -0.72 | -0.41 | B>B |
| chr11 | 39100000 | 39200000 | -0.72 | -0.40 | B>B |
| chr11 | 39200000 | 39300000 | -0.69 | -0.34 | B>B |
| chr11 | 39300000 | 39400000 | -0.65 | -0.36 | B>B |
| chr11 | 39400000 | 39500000 | -0.70 | -0.58 | B>B |
| chr11 | 39500000 | 39600000 | -0.70 | -0.66 | B>B |
| chr11 | 39600000 | 39700000 | -0.71 | -0.77 | B>B |
| chr11 | 39700000 | 39800000 | -0.74 | -0.74 | B>B |
| chr11 | 39800000 | 39900000 | -0.74 | -0.77 | B>B |
| chr11 | 39900000 | 40000000 | -0.71 | -0.75 | B>B |
| chr11 | 40000000 | 40100000 | -0.76 | -0.75 | B>B |
| chr11 | 40100000 | 40200000 | -0.67 | -0.69 | B>B |
| chr11 | 40200000 | 40300000 | -0.57 | -0.61 | B>B |
| chr11 | 40300000 | 40400000 | -0.35 | -0.18 | B>B |
| chr11 | 40400000 | 40500000 | -0.42 | -0.05 | B>B |
| chr11 | 40500000 | 40600000 | -0.59 | -0.13 | B>B |
| chr11 | 40600000 | 40700000 | -0.68 | -0.25 | B>B |
| chr11 | 40700000 | 40800000 | -0.64 | -0.43 | B>B |
| chr11 | 40800000 | 40900000 | -0.76 | -0.72 | B>B |
| chr11 | 40900000 | 41000000 | -0.72 | -0.70 | B>B |
| chr11 | 41000000 | 41100000 | -0.72 | -0.79 | B>B |

| | | | | | |
|---|---|---|---|---|---|
| chr11 | 41100000 | 41200000 | -0.76 | -0.70 | B>B |
| chr11 | 41200000 | 41300000 | -0.72 | -0.59 | B>B |
| chr11 | 41300000 | 41400000 | -0.76 | -0.46 | B>B |
| chr11 | 41400000 | 41500000 | -0.70 | -0.52 | B>B |
| chr11 | 41500000 | 41600000 | -0.74 | -0.65 | B>B |
| chr11 | 41600000 | 41700000 | -0.78 | -0.74 | B>B |
| chr11 | 41700000 | 41800000 | -0.78 | -0.77 | B>B |
| chr11 | 41800000 | 41900000 | -0.81 | -0.77 | B>B |
| chr11 | 41900000 | 42000000 | -0.72 | -0.81 | B>B |
| chr11 | 42000000 | 42100000 | -0.80 | -0.88 | B>B |
| chr11 | 42100000 | 42200000 | -0.82 | -0.88 | B>B |
| chr11 | 42200000 | 42300000 | -0.82 | -0.89 | B>B |
| chr11 | 42300000 | 42400000 | -0.81 | -0.95 | B>B |
| chr11 | 42400000 | 42500000 | -0.89 | -0.94 | B>B |
| chr11 | 42500000 | 42600000 | -0.87 | -0.95 | B>B |
| chr11 | 42600000 | 42700000 | -0.86 | -0.97 | B>B |
| chr11 | 42700000 | 42800000 | -0.89 | -1.00 | B>B |
| chr11 | 42800000 | 42900000 | -0.92 | -1.02 | B>B |
| chr11 | 42900000 | 43000000 | -0.86 | -0.99 | B>B |
| chr11 | 43000000 | 43100000 | -0.92 | -1.09 | B>B |
| chr11 | 43100000 | 43200000 | -0.87 | -1.04 | B>B |
| chr11 | 43200000 | 43300000 | -0.72 | -0.87 | B>B |
| chr11 | 43300000 | 43400000 | -0.16 | -0.17 | B>B |
| chr11 | 43400000 | 43500000 | 0.09 | 0.24 | A>A |
| chr11 | 43500000 | 43600000 | 0.10 | 0.31 | A>A |
| chr11 | 43600000 | 43700000 | 0.30 | 0.45 | A>A |
| chr11 | 43700000 | 43800000 | 0.37 | 0.57 | A>A |
| chr11 | 43800000 | 43900000 | 0.38 | 0.58 | A>A |
| chr11 | 43900000 | 44000000 | 0.63 | 0.74 | A>A |
| chr11 | 44000000 | 44100000 | 0.66 | 0.73 | A>A |
| chr11 | 44100000 | 44200000 | 0.62 | 0.73 | A>A |
| chr11 | 44200000 | 44300000 | 0.70 | 0.73 | A>A |
| chr11 | 44300000 | 44400000 | 0.68 | 0.69 | A>A |
| chr11 | 44400000 | 44500000 | 0.57 | 0.68 | A>A |
| chr11 | 44500000 | 44600000 | 0.58 | 0.63 | A>A |
| chr11 | 44600000 | 44700000 | 0.62 | 0.70 | A>A |
| chr11 | 44700000 | 44800000 | 0.74 | 0.77 | A>A |
| chr11 | 44800000 | 44900000 | 0.86 | 0.82 | A>A |
| chr11 | 44900000 | 45000000 | 0.68 | 0.74 | A>A |
| chr11 | 45000000 | 45100000 | 0.96 | 0.98 | A>A |
| chr11 | 45100000 | 45200000 | 0.76 | 0.85 | A>A |
| chr11 | 45200000 | 45300000 | 0.86 | 0.85 | A>A |
| chr11 | 45300000 | 45400000 | 0.58 | 0.67 | A>A |
| chr11 | 45400000 | 45500000 | 0.68 | 0.54 | A>A |
| chr11 | 45500000 | 45600000 | 0.68 | 0.53 | A>A |
| chr11 | 45600000 | 45700000 | 0.74 | 0.53 | A>A |
| chr11 | 45700000 | 45800000 | 0.73 | 0.57 | A>A |
| chr11 | 45800000 | 45900000 | 0.84 | 0.91 | A>A |
| chr11 | 45900000 | 46000000 | 0.98 | 1.00 | A>A |
| chr11 | 46000000 | 46100000 | 0.95 | 1.03 | A>A |
| chr11 | 46100000 | 46200000 | 1.04 | 1.05 | A>A |
| chr11 | 46200000 | 46300000 | 0.94 | 1.10 | A>A |
| chr11 | 46300000 | 46400000 | 1.01 | 1.04 | A>A |
| chr11 | 46400000 | 46500000 | 1.00 | 1.10 | A>A |
| chr11 | 46500000 | 46600000 | 1.06 | 1.12 | A>A |
| chr11 | 46600000 | 46700000 | 1.07 | 1.16 | A>A |

| | | | | | |
|---|---|---|---|---|---|
| chr11 | 46700000 | 46800000 | 1.16 | 1.18 | A>A |
| chr11 | 46800000 | 46900000 | 1.13 | 1.12 | A>A |
| chr11 | 46900000 | 47000000 | 0.99 | 1.08 | A>A |
| chr11 | 47000000 | 47100000 | 1.10 | 1.13 | A>A |
| chr11 | 47100000 | 47200000 | 0.87 | 1.00 | A>A |
| chr11 | 47200000 | 47300000 | 1.19 | 1.25 | A>A |
| chr11 | 47300000 | 47400000 | 1.10 | 1.11 | A>A |
| chr11 | 47400000 | 47500000 | 1.16 | 1.13 | A>A |
| chr11 | 47500000 | 47600000 | 1.03 | 1.23 | A>A |
| chr11 | 47600000 | 47700000 | 1.00 | 1.10 | A>A |
| chr11 | 47700000 | 47800000 | 0.99 | 1.03 | A>A |
| chr11 | 47800000 | 47900000 | 0.85 | 0.81 | A>A |
| chr11 | 47900000 | 48000000 | 0.73 | 0.71 | A>A |
| chr11 | 48000000 | 48100000 | 0.67 | 0.78 | A>A |
| chr11 | 48100000 | 48200000 | 0.31 | 0.31 | A>A |
| chr11 | 48200000 | 48300000 | -0.59 | -0.89 | B>B |
| chr11 | 48300000 | 48400000 | -1.03 | -1.33 | B>B |
| chr11 | 48400000 | 48500000 | 0.00 | 0.00 | NO |
| chr11 | 48500000 | 48600000 | 0.00 | 0.00 | NO |
| chr11 | 48600000 | 48700000 | 0.00 | 0.00 | NO |
| chr11 | 48700000 | 48800000 | 0.00 | 0.00 | NO |
| chr11 | 48800000 | 48900000 | 0.00 | 0.00 | NO |
| chr11 | 48900000 | 49000000 | 0.00 | 0.00 | NO |
| chr11 | 49000000 | 49100000 | 0.00 | 0.00 | NO |
| chr11 | 49100000 | 49200000 | 0.00 | 0.00 | NO |
| chr11 | 49200000 | 49300000 | 0.00 | 0.00 | NO |
| chr11 | 49300000 | 49400000 | 0.00 | 0.00 | NO |
| chr11 | 49400000 | 49500000 | 0.00 | 0.00 | NO |
| chr11 | 49500000 | 49600000 | 0.00 | 0.00 | NO |
| chr11 | 49600000 | 49700000 | 0.00 | 0.00 | NO |
| chr11 | 49700000 | 49800000 | 0.00 | 0.00 | NO |
| chr11 | 49800000 | 49900000 | 0.00 | 0.00 | NO |
| chr11 | 49900000 | 50000000 | 0.00 | 0.00 | NO |
| chr11 | 50000000 | 50100000 | 0.00 | 0.00 | NO |
| chr11 | 50100000 | 50200000 | 0.00 | 0.00 | NO |
| chr11 | 50200000 | 50300000 | 0.00 | 0.00 | NO |
| chr11 | 50300000 | 50400000 | 0.00 | 0.00 | NO |
| chr11 | 50400000 | 50500000 | 0.00 | 0.00 | NO |
| chr11 | 50500000 | 50600000 | 0.00 | 0.00 | NO |
| chr11 | 50600000 | 50700000 | 0.00 | 0.00 | NO |
| chr11 | 50700000 | 50800000 | 0.00 | 0.00 | NO |
| chr11 | 50800000 | 50900000 | 0.00 | 0.00 | NO |
| chr11 | 50900000 | 51000000 | 0.00 | 0.00 | NO |
| chr11 | 51000000 | 51100000 | 0.00 | 0.00 | NO |
| chr11 | 51100000 | 51200000 | 0.00 | 0.00 | NO |
| chr11 | 51200000 | 51300000 | 0.00 | 0.00 | NO |
| chr11 | 51300000 | 51400000 | 0.00 | 0.00 | NO |
| chr11 | 51400000 | 51500000 | 0.00 | 0.00 | NO |
| chr11 | 51500000 | 51600000 | 0.00 | 0.00 | NO |
| chr11 | 51600000 | 51700000 | 0.00 | 0.00 | NO |
| chr11 | 51700000 | 51800000 | 0.00 | 0.00 | NO |
| chr11 | 51800000 | 51900000 | 0.00 | 0.00 | NO |
| chr11 | 51900000 | 52000000 | 0.00 | 0.00 | NO |
| chr11 | 52000000 | 52100000 | 0.00 | 0.00 | NO |
| chr11 | 52100000 | 52200000 | 0.00 | 0.00 | NO |
| chr11 | 52200000 | 52300000 | 0.00 | 0.00 | NO |

| | | | | | |
|---|---|---|---|---|---|
| chr11 | 52300000 | 52400000 | 0.00 | 0.00 | NO |
| chr11 | 52400000 | 52500000 | 0.00 | 0.00 | NO |
| chr11 | 52500000 | 52600000 | 0.00 | 0.00 | NO |
| chr11 | 52600000 | 52700000 | 0.00 | 0.00 | NO |
| chr11 | 52700000 | 52800000 | 0.00 | 0.00 | NO |
| chr11 | 52800000 | 52900000 | 0.00 | 0.00 | NO |
| chr11 | 52900000 | 53000000 | 0.00 | 0.00 | NO |
| chr11 | 53000000 | 53100000 | 0.00 | 0.00 | NO |
| chr11 | 53100000 | 53200000 | 0.00 | 0.00 | NO |
| chr11 | 53200000 | 53300000 | 0.00 | 0.00 | NO |
| chr11 | 53300000 | 53400000 | 0.00 | 0.00 | NO |
| chr11 | 53400000 | 53500000 | 0.00 | 0.00 | NO |
| chr11 | 53500000 | 53600000 | 0.00 | 0.00 | NO |
| chr11 | 53600000 | 53700000 | 0.00 | 0.00 | NO |
| chr11 | 53700000 | 53800000 | 0.00 | 0.00 | NO |
| chr11 | 53800000 | 53900000 | 0.00 | 0.00 | NO |
| chr11 | 53900000 | 54000000 | 0.00 | 0.00 | NO |
| chr11 | 54000000 | 54100000 | 0.00 | 0.00 | NO |
| chr11 | 54100000 | 54200000 | 0.00 | 0.00 | NO |
| chr11 | 54200000 | 54300000 | 0.00 | 0.00 | NO |
| chr11 | 54300000 | 54400000 | 0.00 | 0.00 | NO |
| chr11 | 54400000 | 54500000 | 0.00 | 0.00 | NO |
| chr11 | 54500000 | 54600000 | 0.00 | 0.00 | NO |
| chr11 | 54600000 | 54700000 | 0.00 | 0.00 | NO |
| chr11 | 54700000 | 54800000 | 0.00 | 0.00 | NO |
| chr11 | 54800000 | 54900000 | 0.00 | 0.00 | NO |
| chr11 | 54900000 | 55000000 | 0.00 | 0.00 | NO |
| chr11 | 55000000 | 55100000 | 0.00 | 0.00 | NO |
| chr11 | 55100000 | 55200000 | -1.15 | -1.24 | B>B |
| chr11 | 55200000 | 55300000 | -1.22 | -1.38 | B>B |
| chr11 | 55300000 | 55400000 | -1.14 | -1.22 | B>B |
| chr11 | 55400000 | 55500000 | -1.09 | -1.15 | B>B |
| chr11 | 55500000 | 55600000 | -1.03 | -1.15 | B>B |
| chr11 | 55600000 | 55700000 | -0.95 | -1.10 | B>B |
| chr11 | 55700000 | 55800000 | -0.85 | -0.91 | B>B |
| chr11 | 55800000 | 55900000 | -0.90 | -1.02 | B>B |
| chr11 | 55900000 | 56000000 | -1.01 | -1.06 | B>B |
| chr11 | 56000000 | 56100000 | -0.91 | -1.06 | B>B |
| chr11 | 56100000 | 56200000 | -0.77 | -1.01 | B>B |
| chr11 | 56200000 | 56300000 | -0.70 | -0.98 | B>B |
| chr11 | 56300000 | 56400000 | -0.55 | -0.91 | B>B |
| chr11 | 56400000 | 56500000 | -0.56 | -0.79 | B>B |
| chr11 | 56500000 | 56600000 | -0.29 | -0.49 | B>B |
| chr11 | 56600000 | 56700000 | 0.40 | 0.33 | A>A |
| chr11 | 56700000 | 56800000 | 0.48 | 0.35 | A>A |
| chr11 | 56800000 | 56900000 | 0.43 | 0.24 | A>A |
| chr11 | 56900000 | 57000000 | 0.33 | 0.09 | A>A |
| chr11 | 57000000 | 57100000 | 0.22 | 0.19 | A>A |
| chr11 | 57100000 | 57200000 | 0.73 | 0.70 | A>A |
| chr11 | 57200000 | 57300000 | 0.74 | 0.67 | A>A |
| chr11 | 57300000 | 57400000 | 0.61 | 0.58 | A>A |
| chr11 | 57400000 | 57500000 | 0.51 | 0.47 | A>A |
| chr11 | 57500000 | 57600000 | 0.42 | 0.27 | A>A |
| chr11 | 57600000 | 57700000 | -0.19 | -0.44 | B>B |
| chr11 | 57700000 | 57800000 | -0.40 | -0.55 | B>B |
| chr11 | 57800000 | 57900000 | -0.63 | -0.80 | B>B |

| | | | | | |
|---|---|---|---|---|---|
| chr11 | 57900000 | 58000000 | -0.69 | -0.90 | B>B |
| chr11 | 58000000 | 58100000 | -0.75 | -0.92 | B>B |
| chr11 | 58100000 | 58200000 | -0.88 | -0.86 | B>B |
| chr11 | 58200000 | 58300000 | -0.53 | -0.54 | B>B |
| chr11 | 58300000 | 58400000 | -0.01 | -0.02 | B>B |
| chr11 | 58400000 | 58500000 | -0.50 | -0.56 | B>B |
| chr11 | 58500000 | 58600000 | -0.74 | -0.97 | B>B |
| chr11 | 58600000 | 58700000 | -0.63 | -0.90 | B>B |
| chr11 | 58700000 | 58800000 | -0.62 | -0.92 | B>B |
| chr11 | 58800000 | 58900000 | -0.27 | -0.43 | B>B |
| chr11 | 58900000 | 59000000 | -0.20 | -0.39 | B>B |
| chr11 | 59000000 | 59100000 | -0.51 | -0.90 | B>B |
| chr11 | 59100000 | 59200000 | -0.68 | -0.96 | B>B |
| chr11 | 59200000 | 59300000 | -0.28 | -0.68 | B>B |
| chr11 | 59300000 | 59400000 | 0.52 | 0.34 | A>A |
| chr11 | 59400000 | 59500000 | 0.36 | 0.18 | A>A |
| chr11 | 59500000 | 59600000 | 0.19 | 0.02 | A>A |
| chr11 | 59600000 | 59700000 | -0.31 | -0.17 | B>B |
| chr11 | 59700000 | 59800000 | -0.65 | -0.80 | B>B |
| chr11 | 59800000 | 59900000 | -0.57 | -0.81 | B>B |
| chr11 | 59900000 | 60000000 | -0.70 | -0.97 | B>B |
| chr11 | 60000000 | 60100000 | -0.69 | -1.09 | B>B |
| chr11 | 60100000 | 60200000 | -0.76 | -1.09 | B>B |
| chr11 | 60200000 | 60300000 | -0.84 | -1.30 | B>B |
| chr11 | 60300000 | 60400000 | -0.82 | -1.25 | B>B |
| chr11 | 60400000 | 60500000 | -0.40 | -1.01 | A>B |
| chr11 | 60500000 | 60600000 | 0.28 | -0.16 | A>A |
| chr11 | 60600000 | 60700000 | 0.76 | 0.54 | A>A |
| chr11 | 60700000 | 60800000 | 0.91 | 0.84 | A>A |
| chr11 | 60800000 | 60900000 | 0.92 | 0.88 | A>A |
| chr11 | 60900000 | 61000000 | 0.84 | 0.83 | A>A |
| chr11 | 61000000 | 61100000 | 1.20 | 1.17 | A>A |
| chr11 | 61100000 | 61200000 | 1.22 | 1.26 | A>A |
| chr11 | 61200000 | 61300000 | 1.17 | 1.26 | A>A |
| chr11 | 61300000 | 61400000 | 1.25 | 1.35 | A>A |
| chr11 | 61400000 | 61500000 | 1.32 | 1.36 | A>A |
| chr11 | 61500000 | 61600000 | 1.18 | 1.19 | A>A |
| chr11 | 61600000 | 61700000 | 1.03 | 1.06 | A>A |
| chr11 | 61700000 | 61800000 | 1.21 | 1.25 | A>A |
| chr11 | 61800000 | 61900000 | 0.96 | 1.03 | A>A |
| chr11 | 61900000 | 62000000 | 0.87 | 1.01 | A>A |
| chr11 | 62000000 | 62100000 | 0.94 | 1.13 | A>A |
| chr11 | 62100000 | 62200000 | 0.78 | 0.95 | A>A |
| chr11 | 62200000 | 62300000 | 0.99 | 1.04 | A>A |
| chr11 | 62300000 | 62400000 | 1.01 | 1.01 | A>A |
| chr11 | 62400000 | 62500000 | 1.01 | 1.09 | A>A |
| chr11 | 62500000 | 62600000 | 0.99 | 1.05 | A>A |
| chr11 | 62600000 | 62700000 | 0.70 | 0.72 | A>A |
| chr11 | 62700000 | 62800000 | 0.48 | 0.43 | A>A |
| chr11 | 62800000 | 62900000 | 0.00 | 0.00 | NO |
| chr11 | 62900000 | 63000000 | 0.01 | 0.07 | A>A |
| chr11 | 63000000 | 63100000 | 0.01 | 0.07 | A>A |
| chr11 | 63100000 | 63200000 | 0.03 | 0.12 | A>A |
| chr11 | 63200000 | 63300000 | 0.38 | 0.43 | A>A |
| chr11 | 63300000 | 63400000 | 0.68 | 0.78 | A>A |
| chr11 | 63400000 | 63500000 | 0.70 | 0.79 | A>A |

| | | | | | |
|---|---|---|---|---|---|
| chr11 | 63500000 | 63600000 | 0.73 | 0.79 | A>A |
| chr11 | 63600000 | 63700000 | 1.02 | 0.97 | A>A |
| chr11 | 63700000 | 63800000 | 0.92 | 0.95 | A>A |
| chr11 | 63800000 | 63900000 | 1.07 | 1.09 | A>A |
| chr11 | 63900000 | 64000000 | 0.99 | 1.04 | A>A |
| chr11 | 64000000 | 64100000 | 0.99 | 1.00 | A>A |
| chr11 | 64100000 | 64200000 | 0.93 | 0.92 | A>A |
| chr11 | 64200000 | 64300000 | 0.84 | 0.95 | A>A |
| chr11 | 64300000 | 64400000 | 0.98 | 1.00 | A>A |
| chr11 | 64400000 | 64500000 | 0.83 | 0.89 | A>A |
| chr11 | 64500000 | 64600000 | 0.76 | 0.86 | A>A |
| chr11 | 64600000 | 64700000 | 0.90 | 0.95 | A>A |
| chr11 | 64700000 | 64800000 | 0.94 | 0.89 | A>A |
| chr11 | 64800000 | 64900000 | 0.75 | 0.77 | A>A |
| chr11 | 64900000 | 65000000 | 0.77 | 0.83 | A>A |
| chr11 | 65000000 | 65100000 | 0.81 | 0.81 | A>A |
| chr11 | 65100000 | 65200000 | 0.77 | 0.71 | A>A |
| chr11 | 65200000 | 65300000 | 0.82 | 0.80 | A>A |
| chr11 | 65300000 | 65400000 | 0.65 | 0.64 | A>A |
| chr11 | 65400000 | 65500000 | 0.65 | 0.72 | A>A |
| chr11 | 65500000 | 65600000 | 0.78 | 0.80 | A>A |
| chr11 | 65600000 | 65700000 | 0.70 | 0.74 | A>A |
| chr11 | 65700000 | 65800000 | 0.64 | 0.70 | A>A |
| chr11 | 65800000 | 65900000 | 0.76 | 0.84 | A>A |
| chr11 | 65900000 | 66000000 | 0.79 | 0.89 | A>A |
| chr11 | 66000000 | 66100000 | 0.78 | 0.82 | A>A |
| chr11 | 66100000 | 66200000 | 0.72 | 0.83 | A>A |
| chr11 | 66200000 | 66300000 | 0.76 | 0.81 | A>A |
| chr11 | 66300000 | 66400000 | 0.81 | 0.81 | A>A |
| chr11 | 66400000 | 66500000 | 0.71 | 0.78 | A>A |
| chr11 | 66500000 | 66600000 | 0.68 | 0.80 | A>A |
| chr11 | 66600000 | 66700000 | 0.73 | 0.81 | A>A |
| chr11 | 66700000 | 66800000 | 0.76 | 0.76 | A>A |
| chr11 | 66800000 | 66900000 | 0.74 | 0.75 | A>A |
| chr11 | 66900000 | 67000000 | 0.78 | 0.82 | A>A |
| chr11 | 67000000 | 67100000 | 0.72 | 0.80 | A>A |
| chr11 | 67100000 | 67200000 | 0.76 | 0.75 | A>A |
| chr11 | 67200000 | 67300000 | 0.78 | 0.84 | A>A |
| chr11 | 67300000 | 67400000 | 0.80 | 0.83 | A>A |
| chr11 | 67400000 | 67500000 | 0.78 | 0.84 | A>A |
| chr11 | 67500000 | 67600000 | 0.00 | 0.00 | NO |
| chr11 | 67600000 | 67700000 | 0.00 | 0.00 | NO |
| chr11 | 67700000 | 67800000 | 0.00 | 0.00 | NO |
| chr11 | 67800000 | 67900000 | 0.82 | 0.86 | A>A |
| chr11 | 67900000 | 68000000 | 0.87 | 0.87 | A>A |
| chr11 | 68000000 | 68100000 | 0.80 | 0.87 | A>A |
| chr11 | 68100000 | 68200000 | 0.80 | 0.90 | A>A |
| chr11 | 68200000 | 68300000 | 0.74 | 0.80 | A>A |
| chr11 | 68300000 | 68400000 | 0.73 | 0.78 | A>A |
| chr11 | 68400000 | 68500000 | 0.72 | 0.79 | A>A |
| chr11 | 68500000 | 68600000 | 0.71 | 0.80 | A>A |
| chr11 | 68600000 | 68700000 | 0.74 | 0.79 | A>A |
| chr11 | 68700000 | 68800000 | 0.73 | 0.79 | A>A |
| chr11 | 68800000 | 68900000 | 0.56 | 0.57 | A>A |
| chr11 | 68900000 | 69000000 | 0.75 | 0.74 | A>A |
| chr11 | 69000000 | 69100000 | 0.70 | 0.76 | A>A |

| | | | | | |
|---|---|---|---|---|---|
| chr11 | 69100000 | 69200000 | 0.73 | 0.75 | A>A |
| chr11 | 69200000 | 69300000 | 0.74 | 0.77 | A>A |
| chr11 | 69300000 | 69400000 | 0.77 | 0.77 | A>A |
| chr11 | 69400000 | 69500000 | 0.56 | 0.66 | A>A |
| chr11 | 69500000 | 69600000 | 0.72 | 0.74 | A>A |
| chr11 | 69600000 | 69700000 | 0.68 | 0.72 | A>A |
| chr11 | 69700000 | 69800000 | 0.00 | 0.00 | NO |
| chr11 | 69800000 | 69900000 | 0.78 | 0.81 | A>A |
| chr11 | 69900000 | 70000000 | 0.82 | 0.85 | A>A |
| chr11 | 70000000 | 70100000 | 0.68 | 0.74 | A>A |
| chr11 | 70100000 | 70200000 | 0.89 | 0.91 | A>A |
| chr11 | 70200000 | 70300000 | 0.77 | 0.85 | A>A |
| chr11 | 70300000 | 70400000 | 0.65 | 0.72 | A>A |
| chr11 | 70400000 | 70500000 | 0.74 | 0.76 | A>A |
| chr11 | 70500000 | 70600000 | 0.62 | 0.72 | A>A |
| chr11 | 70600000 | 70700000 | 0.69 | 0.76 | A>A |
| chr11 | 70700000 | 70800000 | 0.69 | 0.69 | A>A |
| chr11 | 70800000 | 70900000 | 0.68 | 0.73 | A>A |
| chr11 | 70900000 | 71000000 | 0.79 | 0.82 | A>A |
| chr11 | 71000000 | 71100000 | 0.70 | 0.78 | A>A |
| chr11 | 71100000 | 71200000 | 0.81 | 0.82 | A>A |
| chr11 | 71200000 | 71300000 | 1.00 | 0.97 | A>A |
| chr11 | 71300000 | 71400000 | 0.00 | 0.00 | NO |
| chr11 | 71400000 | 71500000 | 0.00 | 0.00 | NO |
| chr11 | 71500000 | 71600000 | 0.00 | 0.00 | NO |
| chr11 | 71600000 | 71700000 | 0.92 | 0.91 | A>A |
| chr11 | 71700000 | 71800000 | 1.00 | 0.97 | A>A |
| chr11 | 71800000 | 71900000 | 0.93 | 1.03 | A>A |
| chr11 | 71900000 | 72000000 | 0.94 | 0.93 | A>A |
| chr11 | 72000000 | 72100000 | 0.93 | 0.99 | A>A |
| chr11 | 72100000 | 72200000 | 0.74 | 0.82 | A>A |
| chr11 | 72200000 | 72300000 | 0.86 | 0.87 | A>A |
| chr11 | 72300000 | 72400000 | 0.91 | 0.90 | A>A |
| chr11 | 72400000 | 72500000 | 0.92 | 0.93 | A>A |
| chr11 | 72500000 | 72600000 | 0.77 | 0.80 | A>A |
| chr11 | 72600000 | 72700000 | 0.80 | 0.79 | A>A |
| chr11 | 72700000 | 72800000 | 0.92 | 0.91 | A>A |
| chr11 | 72800000 | 72900000 | 0.75 | 0.76 | A>A |
| chr11 | 72900000 | 73000000 | 0.89 | 0.82 | A>A |
| chr11 | 73000000 | 73100000 | 0.93 | 0.86 | A>A |
| chr11 | 73100000 | 73200000 | 0.81 | 0.87 | A>A |
| chr11 | 73200000 | 73300000 | 0.94 | 0.82 | A>A |
| chr11 | 73300000 | 73400000 | 0.74 | 0.78 | A>A |
| chr11 | 73400000 | 73500000 | 0.86 | 0.89 | A>A |
| chr11 | 73500000 | 73600000 | 0.97 | 0.90 | A>A |
| chr11 | 73600000 | 73700000 | 0.77 | 0.83 | A>A |
| chr11 | 73700000 | 73800000 | 0.97 | 0.95 | A>A |
| chr11 | 73800000 | 73900000 | 0.90 | 0.85 | A>A |
| chr11 | 73900000 | 74000000 | 0.69 | 0.80 | A>A |
| chr11 | 74000000 | 74100000 | 0.83 | 0.82 | A>A |
| chr11 | 74100000 | 74200000 | 0.76 | 0.82 | A>A |
| chr11 | 74200000 | 74300000 | 0.84 | 0.83 | A>A |
| chr11 | 74300000 | 74400000 | 0.82 | 0.85 | A>A |
| chr11 | 74400000 | 74500000 | 0.85 | 0.86 | A>A |
| chr11 | 74500000 | 74600000 | 0.72 | 0.82 | A>A |
| chr11 | 74600000 | 74700000 | 0.70 | 0.84 | A>A |

| | | | | | |
|---|---|---|---|---|---|
| chr11 | 74700000 | 74800000 | 0.76 | 0.77 | A>A |
| chr11 | 74800000 | 74900000 | 0.84 | 0.81 | A>A |
| chr11 | 74900000 | 75000000 | 0.73 | 0.80 | A>A |
| chr11 | 75000000 | 75100000 | 0.98 | 1.05 | A>A |
| chr11 | 75100000 | 75200000 | 1.05 | 1.06 | A>A |
| chr11 | 75200000 | 75300000 | 0.93 | 0.96 | A>A |
| chr11 | 75300000 | 75400000 | 0.93 | 0.96 | A>A |
| chr11 | 75400000 | 75500000 | 0.86 | 0.88 | A>A |
| chr11 | 75500000 | 75600000 | 0.77 | 0.84 | A>A |
| chr11 | 75600000 | 75700000 | 0.72 | 0.82 | A>A |
| chr11 | 75700000 | 75800000 | 0.86 | 0.89 | A>A |
| chr11 | 75800000 | 75900000 | 0.80 | 0.79 | A>A |
| chr11 | 75900000 | 76000000 | 0.94 | 0.88 | A>A |
| chr11 | 76000000 | 76100000 | 0.81 | 0.83 | A>A |
| chr11 | 76100000 | 76200000 | 0.95 | 0.92 | A>A |
| chr11 | 76200000 | 76300000 | 0.81 | 0.89 | A>A |
| chr11 | 76300000 | 76400000 | 0.83 | 0.90 | A>A |
| chr11 | 76400000 | 76500000 | 0.89 | 0.92 | A>A |
| chr11 | 76500000 | 76600000 | 0.85 | 0.98 | A>A |
| chr11 | 76600000 | 76700000 | 0.74 | 0.90 | A>A |
| chr11 | 76700000 | 76800000 | 0.93 | 0.98 | A>A |
| chr11 | 76800000 | 76900000 | 0.89 | 0.87 | A>A |
| chr11 | 76900000 | 77000000 | 0.76 | 0.94 | A>A |
| chr11 | 77000000 | 77100000 | 0.86 | 0.86 | A>A |
| chr11 | 77100000 | 77200000 | 0.88 | 0.99 | A>A |
| chr11 | 77200000 | 77300000 | 0.74 | 0.86 | A>A |
| chr11 | 77300000 | 77400000 | 0.94 | 0.99 | A>A |
| chr11 | 77400000 | 77500000 | 0.98 | 0.98 | A>A |
| chr11 | 77500000 | 77600000 | 0.97 | 0.98 | A>A |
| chr11 | 77600000 | 77700000 | 0.00 | 0.00 | NO |
| chr11 | 77700000 | 77800000 | 0.98 | 0.94 | A>A |
| chr11 | 77800000 | 77900000 | 0.92 | 0.77 | A>A |
| chr11 | 77900000 | 78000000 | 0.66 | 0.68 | A>A |
| chr11 | 78000000 | 78100000 | 0.49 | 0.54 | A>A |
| chr11 | 78100000 | 78200000 | 0.43 | 0.51 | A>A |
| chr11 | 78200000 | 78300000 | 0.40 | 0.29 | A>A |
| chr11 | 78300000 | 78400000 | -0.26 | -0.63 | B>B |
| chr11 | 78400000 | 78500000 | -0.37 | -0.96 | A>B |
| chr11 | 78500000 | 78600000 | -0.29 | -1.01 | A>B |
| chr11 | 78600000 | 78700000 | -0.17 | -0.97 | A>B |
| chr11 | 78700000 | 78800000 | -0.04 | -0.73 | A>B |
| chr11 | 78800000 | 78900000 | -0.01 | -0.68 | A>B |
| chr11 | 78900000 | 79000000 | -0.14 | -0.75 | A>B |
| chr11 | 79000000 | 79100000 | -0.24 | -0.85 | A>B |
| chr11 | 79100000 | 79200000 | -0.30 | -0.85 | B>B |
| chr11 | 79200000 | 79300000 | -0.69 | -0.92 | B>B |
| chr11 | 79300000 | 79400000 | -0.82 | -0.89 | B>B |
| chr11 | 79400000 | 79500000 | -0.89 | -0.87 | B>B |
| chr11 | 79500000 | 79600000 | -0.97 | -0.85 | B>B |
| chr11 | 79600000 | 79700000 | -0.97 | -0.84 | B>B |
| chr11 | 79700000 | 79800000 | -0.98 | -0.71 | B>B |
| chr11 | 79800000 | 79900000 | -0.92 | -0.72 | B>B |
| chr11 | 79900000 | 80000000 | -0.94 | -0.69 | B>B |
| chr11 | 80000000 | 80100000 | -0.96 | -0.69 | B>B |
| chr11 | 80100000 | 80200000 | -0.95 | -0.64 | B>B |
| chr11 | 80200000 | 80300000 | -0.89 | -0.65 | B>B |

| | | | | | |
|---|---|---|---|---|---|
| chr11 | 80300000 | 80400000 | -0.94 | -0.66 | B>B |
| chr11 | 80400000 | 80500000 | -0.93 | -0.65 | B>B |
| chr11 | 80500000 | 80600000 | -0.95 | -0.71 | B>B |
| chr11 | 80600000 | 80700000 | -0.89 | -0.65 | B>B |
| chr11 | 80700000 | 80800000 | -0.93 | -0.78 | B>B |
| chr11 | 80800000 | 80900000 | -0.92 | -0.95 | B>B |
| chr11 | 80900000 | 81000000 | -0.95 | -0.94 | B>B |
| chr11 | 81000000 | 81100000 | -0.89 | -1.05 | B>B |
| chr11 | 81100000 | 81200000 | -0.94 | -1.04 | B>B |
| chr11 | 81200000 | 81300000 | -0.87 | -1.01 | B>B |
| chr11 | 81300000 | 81400000 | -0.85 | -1.01 | B>B |
| chr11 | 81400000 | 81500000 | -0.93 | -1.04 | B>B |
| chr11 | 81500000 | 81600000 | -0.98 | -1.05 | B>B |
| chr11 | 81600000 | 81700000 | -0.91 | -1.07 | B>B |
| chr11 | 81700000 | 81800000 | -0.95 | -1.11 | B>B |
| chr11 | 81800000 | 81900000 | -0.95 | -1.05 | B>B |
| chr11 | 81900000 | 82000000 | -0.92 | -1.05 | B>B |
| chr11 | 82000000 | 82100000 | -1.00 | -1.08 | B>B |
| chr11 | 82100000 | 82200000 | -1.00 | -1.09 | B>B |
| chr11 | 82200000 | 82300000 | -0.95 | -1.09 | B>B |
| chr11 | 82300000 | 82400000 | -0.70 | -0.91 | B>B |
| chr11 | 82400000 | 82500000 | -0.32 | -0.56 | B>B |
| chr11 | 82500000 | 82600000 | 0.10 | -0.32 | B>B |
| chr11 | 82600000 | 82700000 | 0.37 | 0.14 | A>A |
| chr11 | 82700000 | 82800000 | 0.50 | 0.38 | A>A |
| chr11 | 82800000 | 82900000 | 0.50 | 0.50 | A>A |
| chr11 | 82900000 | 83000000 | 0.11 | -0.05 | A>A |
| chr11 | 83000000 | 83100000 | -0.49 | -0.84 | B>B |
| chr11 | 83100000 | 83200000 | -0.70 | -1.12 | B>B |
| chr11 | 83200000 | 83300000 | -0.77 | -1.09 | B>B |
| chr11 | 83300000 | 83400000 | -0.72 | -1.14 | B>B |
| chr11 | 83400000 | 83500000 | -0.58 | -0.96 | B>B |
| chr11 | 83500000 | 83600000 | -0.55 | -0.75 | B>B |
| chr11 | 83600000 | 83700000 | -0.91 | -1.09 | B>B |
| chr11 | 83700000 | 83800000 | -1.00 | -1.08 | B>B |
| chr11 | 83800000 | 83900000 | -1.00 | -1.14 | B>B |
| chr11 | 83900000 | 84000000 | -0.89 | -1.09 | B>B |
| chr11 | 84000000 | 84100000 | -0.82 | -1.07 | B>B |
| chr11 | 84100000 | 84200000 | -0.72 | -1.04 | B>B |
| chr11 | 84200000 | 84300000 | -0.54 | -0.52 | B>B |
| chr11 | 84300000 | 84400000 | -0.62 | -0.27 | B>B |
| chr11 | 84400000 | 84500000 | -0.56 | -0.32 | B>B |
| chr11 | 84500000 | 84600000 | -0.53 | -0.16 | B>B |
| chr11 | 84600000 | 84700000 | -0.50 | -0.07 | B>B |
| chr11 | 84700000 | 84800000 | -0.26 | -0.02 | B>B |
| chr11 | 84800000 | 84900000 | -0.14 | -0.04 | B>B |
| chr11 | 84900000 | 85000000 | -0.12 | 0.02 | B>B |
| chr11 | 85000000 | 85100000 | -0.18 | 0.01 | B>B |
| chr11 | 85100000 | 85200000 | -0.34 | -0.16 | B>B |
| chr11 | 85200000 | 85300000 | -0.37 | -0.13 | B>B |
| chr11 | 85300000 | 85400000 | 0.30 | 0.36 | A>A |
| chr11 | 85400000 | 85500000 | 0.73 | 0.79 | A>A |
| chr11 | 85500000 | 85600000 | 0.55 | 0.63 | A>A |
| chr11 | 85600000 | 85700000 | 0.58 | 0.68 | A>A |
| chr11 | 85700000 | 85800000 | 0.71 | 0.87 | A>A |
| chr11 | 85800000 | 85900000 | 0.92 | 0.94 | A>A |

| | | | | | |
|---|---|---|---|---|---|
| chr11 | 85900000 | 86000000 | 0.64 | 0.76 | A>A |
| chr11 | 86000000 | 86100000 | 0.55 | 0.61 | A>A |
| chr11 | 86100000 | 86200000 | 0.57 | 0.62 | A>A |
| chr11 | 86200000 | 86300000 | 0.81 | 0.87 | A>A |
| chr11 | 86300000 | 86400000 | 0.74 | 0.80 | A>A |
| chr11 | 86400000 | 86500000 | 0.99 | 0.98 | A>A |
| chr11 | 86500000 | 86600000 | 0.45 | 0.50 | A>A |
| chr11 | 86600000 | 86700000 | 0.21 | 0.26 | A>A |
| chr11 | 86700000 | 86800000 | 0.37 | 0.44 | A>A |
| chr11 | 86800000 | 86900000 | 0.35 | 0.48 | A>A |
| chr11 | 86900000 | 87000000 | 0.23 | 0.31 | A>A |
| chr11 | 87000000 | 87100000 | -0.18 | -0.17 | B>B |
| chr11 | 87100000 | 87200000 | -0.56 | -0.68 | B>B |
| chr11 | 87200000 | 87300000 | -0.86 | -1.06 | B>B |
| chr11 | 87300000 | 87400000 | -0.89 | -1.15 | B>B |
| chr11 | 87400000 | 87500000 | -0.88 | -1.03 | B>B |
| chr11 | 87500000 | 87600000 | -0.77 | -0.85 | B>B |
| chr11 | 87600000 | 87700000 | -0.51 | -0.50 | B>B |
| chr11 | 87700000 | 87800000 | 0.00 | -0.11 | NO |
| chr11 | 87800000 | 87900000 | -0.20 | 0.01 | B>B |
| chr11 | 87900000 | 88000000 | -0.18 | 0.05 | B>B |
| chr11 | 88000000 | 88100000 | -0.04 | 0.16 | A>A |
| chr11 | 88100000 | 88200000 | 0.13 | 0.22 | A>A |
| chr11 | 88200000 | 88300000 | -0.06 | 0.05 | B>B |
| chr11 | 88300000 | 88400000 | -0.31 | -0.04 | B>B |
| chr11 | 88400000 | 88500000 | -0.42 | -0.10 | B>B |
| chr11 | 88500000 | 88600000 | -0.38 | -0.06 | B>B |
| chr11 | 88600000 | 88700000 | 0.00 | 0.00 | NO |
| chr11 | 88700000 | 88800000 | 0.00 | 0.00 | NO |
| chr11 | 88800000 | 88900000 | 0.00 | 0.00 | NO |
| chr11 | 88900000 | 89000000 | -1.13 | -1.05 | B>B |
| chr11 | 89000000 | 89100000 | 0.00 | 0.00 | NO |
| chr11 | 89100000 | 89200000 | 0.00 | 0.00 | NO |
| chr11 | 89200000 | 89300000 | -1.15 | -1.14 | B>B |
| chr11 | 89300000 | 89400000 | 0.00 | 0.00 | NO |
| chr11 | 89400000 | 89500000 | 0.00 | 0.00 | NO |
| chr11 | 89500000 | 89600000 | 0.00 | 0.00 | NO |
| chr11 | 89600000 | 89700000 | 0.00 | 0.00 | NO |
| chr11 | 89700000 | 89800000 | 0.00 | 0.00 | NO |
| chr11 | 89800000 | 89900000 | 0.00 | -0.65 | NO |
| chr11 | 89900000 | 90000000 | -0.27 | -0.37 | B>B |
| chr11 | 90000000 | 90100000 | -0.75 | -0.82 | B>B |
| chr11 | 90100000 | 90200000 | -0.77 | -0.87 | B>B |
| chr11 | 90200000 | 90300000 | -0.69 | -0.84 | B>B |
| chr11 | 90300000 | 90400000 | -0.64 | -0.84 | B>B |
| chr11 | 90400000 | 90500000 | -0.52 | -0.84 | B>B |
| chr11 | 90500000 | 90600000 | -0.35 | -0.78 | B>B |
| chr11 | 90600000 | 90700000 | -0.23 | -0.74 | B>B |
| chr11 | 90700000 | 90800000 | -0.05 | -0.71 | A>B |
| chr11 | 90800000 | 90900000 | 0.04 | -0.60 | A>B |
| chr11 | 90900000 | 91000000 | 0.09 | -0.62 | A>B |
| chr11 | 91000000 | 91100000 | 0.13 | -0.48 | A>B |
| chr11 | 91100000 | 91200000 | 0.17 | -0.25 | B>B |
| chr11 | 91200000 | 91300000 | 0.15 | -0.35 | B>B |
| chr11 | 91300000 | 91400000 | 0.14 | -0.44 | B>B |
| chr11 | 91400000 | 91500000 | 0.09 | -0.43 | B>B |

| | | | | | |
|---|---|---|---|---|---|
| chr11 | 91500000 | 91600000 | 0.01 | -0.41 | B>B |
| chr11 | 91600000 | 91700000 | -0.04 | -0.52 | B>B |
| chr11 | 91700000 | 91800000 | -0.17 | -0.52 | B>B |
| chr11 | 91800000 | 91900000 | -0.15 | -0.48 | B>B |
| chr11 | 91900000 | 92000000 | 0.03 | -0.27 | B>B |
| chr11 | 92000000 | 92100000 | 0.14 | -0.19 | B>B |
| chr11 | 92100000 | 92200000 | -0.03 | -0.32 | B>B |
| chr11 | 92200000 | 92300000 | -0.16 | -0.45 | B>B |
| chr11 | 92300000 | 92400000 | -0.33 | -0.70 | B>B |
| chr11 | 92400000 | 92500000 | -0.25 | -0.76 | B>B |
| chr11 | 92500000 | 92600000 | -0.25 | -0.67 | B>B |
| chr11 | 92600000 | 92700000 | -0.02 | -0.13 | B>B |
| chr11 | 92700000 | 92800000 | 0.31 | 0.36 | A>A |
| chr11 | 92800000 | 92900000 | 0.13 | 0.06 | A>A |
| chr11 | 92900000 | 93000000 | -0.02 | 0.00 | B>B |
| chr11 | 93000000 | 93100000 | -0.33 | -0.10 | B>B |
| chr11 | 93100000 | 93200000 | 0.00 | -0.09 | NO |
| chr11 | 93200000 | 93300000 | 0.04 | 0.14 | A>A |
| chr11 | 93300000 | 93400000 | 0.17 | 0.22 | A>A |
| chr11 | 93400000 | 93500000 | 0.66 | 0.58 | A>A |
| chr11 | 93500000 | 93600000 | 0.66 | 0.28 | A>A |
| chr11 | 93600000 | 93700000 | 0.19 | -0.27 | B>B |
| chr11 | 93700000 | 93800000 | 0.01 | -0.62 | A>B |
| chr11 | 93800000 | 93900000 | 0.21 | -0.34 | B>B |
| chr11 | 93900000 | 94000000 | 0.00 | -0.40 | NO |
| chr11 | 94000000 | 94100000 | 0.00 | -0.51 | NO |
| chr11 | 94100000 | 94200000 | 0.30 | -0.12 | A>A |
| chr11 | 94200000 | 94300000 | 0.48 | 0.26 | A>A |
| chr11 | 94300000 | 94400000 | 0.51 | 0.41 | A>A |
| chr11 | 94400000 | 94500000 | 0.71 | 0.74 | A>A |
| chr11 | 94500000 | 94600000 | 0.63 | 0.62 | A>A |
| chr11 | 94600000 | 94700000 | 0.40 | 0.38 | A>A |
| chr11 | 94700000 | 94800000 | 0.14 | 0.17 | A>A |
| chr11 | 94800000 | 94900000 | -0.01 | 0.12 | A>A |
| chr11 | 94900000 | 95000000 | -0.18 | -0.01 | B>B |
| chr11 | 95000000 | 95100000 | -0.23 | 0.01 | B>B |
| chr11 | 95100000 | 95200000 | -0.23 | -0.16 | B>B |
| chr11 | 95200000 | 95300000 | -0.37 | -0.24 | B>B |
| chr11 | 95300000 | 95400000 | -0.45 | -0.44 | B>B |
| chr11 | 95400000 | 95500000 | -0.27 | -0.15 | B>B |
| chr11 | 95500000 | 95600000 | 0.07 | 0.15 | A>A |
| chr11 | 95600000 | 95700000 | 0.42 | 0.39 | A>A |
| chr11 | 95700000 | 95800000 | 0.42 | 0.49 | A>A |
| chr11 | 95800000 | 95900000 | 0.45 | 0.53 | A>A |
| chr11 | 95900000 | 96000000 | 0.61 | 0.65 | A>A |
| chr11 | 96000000 | 96100000 | 0.47 | 0.56 | A>A |
| chr11 | 96100000 | 96200000 | -0.15 | -0.13 | B>B |
| chr11 | 96200000 | 96300000 | -0.56 | -0.60 | B>B |
| chr11 | 96300000 | 96400000 | 0.00 | 0.00 | NO |
| chr11 | 96400000 | 96500000 | 0.00 | 0.00 | NO |
| chr11 | 96500000 | 96600000 | -0.80 | -0.88 | B>B |
| chr11 | 96600000 | 96700000 | -0.91 | -0.93 | B>B |
| chr11 | 96700000 | 96800000 | -0.99 | -0.94 | B>B |
| chr11 | 96800000 | 96900000 | -0.89 | -0.88 | B>B |
| chr11 | 96900000 | 97000000 | -0.90 | -0.87 | B>B |
| chr11 | 97000000 | 97100000 | -0.81 | -0.88 | B>B |

| | | | | | |
|---|---|---|---|---|---|
| chr11 | 97100000 | 97200000 | -0.92 | -0.88 | B>B |
| chr11 | 97200000 | 97300000 | -0.90 | -0.89 | B>B |
| chr11 | 97300000 | 97400000 | -0.83 | -0.86 | B>B |
| chr11 | 97400000 | 97500000 | -0.81 | -0.82 | B>B |
| chr11 | 97500000 | 97600000 | -0.81 | -0.80 | B>B |
| chr11 | 97600000 | 97700000 | -0.81 | -0.89 | B>B |
| chr11 | 97700000 | 97800000 | -0.81 | -0.84 | B>B |
| chr11 | 97800000 | 97900000 | -0.84 | -0.78 | B>B |
| chr11 | 97900000 | 98000000 | -0.83 | -0.80 | B>B |
| chr11 | 98000000 | 98100000 | -0.80 | -0.78 | B>B |
| chr11 | 98100000 | 98200000 | -0.76 | -0.79 | B>B |
| chr11 | 98200000 | 98300000 | -0.72 | -0.70 | B>B |
| chr11 | 98300000 | 98400000 | -0.79 | -0.76 | B>B |
| chr11 | 98400000 | 98500000 | -0.77 | -0.77 | B>B |
| chr11 | 98500000 | 98600000 | -0.73 | -0.71 | B>B |
| chr11 | 98600000 | 98700000 | -0.81 | -0.78 | B>B |
| chr11 | 98700000 | 98800000 | -0.78 | -0.66 | B>B |
| chr11 | 98800000 | 98900000 | -0.59 | -0.50 | B>B |
| chr11 | 98900000 | 99000000 | -0.49 | -0.42 | B>B |
| chr11 | 99000000 | 99100000 | -0.43 | -0.31 | B>B |
| chr11 | 99100000 | 99200000 | -0.30 | -0.30 | B>B |
| chr11 | 99200000 | 99300000 | -0.33 | -0.36 | B>B |
| chr11 | 99300000 | 99400000 | -0.40 | -0.38 | B>B |
| chr11 | 99400000 | 99500000 | -0.48 | -0.41 | B>B |
| chr11 | 99500000 | 99600000 | -0.41 | -0.34 | B>B |
| chr11 | 99600000 | 99700000 | -0.59 | -0.54 | B>B |
| chr11 | 99700000 | 99800000 | -0.77 | -0.68 | B>B |
| chr11 | 99800000 | 99900000 | -0.87 | -0.79 | B>B |
| chr11 | 99900000 | 100000000 | -0.88 | -0.82 | B>B |
| chr11 | 1E+08 | 100100000 | -0.91 | -0.87 | B>B |
| chr11 | 1E+08 | 100200000 | -0.85 | -0.92 | B>B |
| chr11 | 1E+08 | 100300000 | -0.75 | -0.86 | B>B |
| chr11 | 1E+08 | 100400000 | -0.72 | -0.80 | B>B |
| chr11 | 1E+08 | 100500000 | -0.31 | -0.30 | B>B |
| chr11 | 1.01E+08 | 100600000 | 0.04 | 0.15 | A>A |
| chr11 | 1.01E+08 | 100700000 | 0.32 | 0.39 | A>A |
| chr11 | 1.01E+08 | 100800000 | 0.00 | 0.06 | A>A |
| chr11 | 1.01E+08 | 100900000 | -0.37 | -0.42 | B>B |
| chr11 | 1.01E+08 | 101000000 | -0.82 | -0.92 | B>B |
| chr11 | 1.01E+08 | 101100000 | -0.85 | -1.07 | B>B |
| chr11 | 1.01E+08 | 101200000 | 0.00 | -1.06 | NO |
| chr11 | 1.01E+08 | 101300000 | -0.49 | -0.54 | B>B |
| chr11 | 1.01E+08 | 101400000 | -0.04 | 0.31 | A>A |
| chr11 | 1.01E+08 | 101500000 | 0.11 | 0.38 | A>A |
| chr11 | 1.02E+08 | 101600000 | 0.00 | 0.00 | NO |
| chr11 | 1.02E+08 | 101700000 | 0.15 | 0.37 | A>A |
| chr11 | 1.02E+08 | 101800000 | 0.37 | 0.52 | A>A |
| chr11 | 1.02E+08 | 101900000 | 0.44 | 0.50 | A>A |
| chr11 | 1.02E+08 | 102000000 | 0.57 | 0.68 | A>A |
| chr11 | 1.02E+08 | 102100000 | 0.70 | 0.86 | A>A |
| chr11 | 1.02E+08 | 102200000 | 0.65 | 0.74 | A>A |
| chr11 | 1.02E+08 | 102300000 | 0.71 | 0.82 | A>A |
| chr11 | 1.02E+08 | 102400000 | 1.02 | 0.97 | A>A |
| chr11 | 1.02E+08 | 102500000 | 0.90 | 0.95 | A>A |
| chr11 | 1.03E+08 | 102600000 | 0.55 | 0.43 | A>A |
| chr11 | 1.03E+08 | 102700000 | 0.48 | 0.32 | A>A |

| | | | | | |
|---|---|---|---|---|---|
| chr11 | 1.03E+08 | 102800000 | 0.33 | 0.31 | A>A |
| chr11 | 1.03E+08 | 102900000 | 0.37 | 0.21 | A>A |
| chr11 | 1.03E+08 | 103000000 | 0.42 | 0.37 | A>A |
| chr11 | 1.03E+08 | 103100000 | 0.27 | 0.18 | A>A |
| chr11 | 1.03E+08 | 103200000 | 0.12 | 0.16 | A>A |
| chr11 | 1.03E+08 | 103300000 | 0.14 | 0.06 | A>A |
| chr11 | 1.03E+08 | 103400000 | 0.26 | 0.08 | A>A |
| chr11 | 1.03E+08 | 103500000 | 0.18 | 0.05 | A>A |
| chr11 | 1.04E+08 | 103600000 | 0.10 | 0.01 | A>A |
| chr11 | 1.04E+08 | 103700000 | 0.08 | 0.05 | A>A |
| chr11 | 1.04E+08 | 103800000 | -0.03 | -0.04 | B>B |
| chr11 | 1.04E+08 | 103900000 | -0.02 | 0.07 | A>A |
| chr11 | 1.04E+08 | 104000000 | 0.03 | 0.07 | A>A |
| chr11 | 1.04E+08 | 104100000 | -0.08 | -0.14 | B>B |
| chr11 | 1.04E+08 | 104200000 | -0.20 | -0.38 | B>B |
| chr11 | 1.04E+08 | 104300000 | -0.30 | -0.36 | B>B |
| chr11 | 1.04E+08 | 104400000 | -0.35 | -0.35 | B>B |
| chr11 | 1.04E+08 | 104500000 | -0.45 | -0.40 | B>B |
| chr11 | 1.05E+08 | 104600000 | -0.44 | -0.35 | B>B |
| chr11 | 1.05E+08 | 104700000 | -0.53 | -0.37 | B>B |
| chr11 | 1.05E+08 | 104800000 | -0.59 | -0.36 | B>B |
| chr11 | 1.05E+08 | 104900000 | -0.38 | -0.42 | B>B |
| chr11 | 1.05E+08 | 105000000 | -0.37 | -0.59 | B>B |
| chr11 | 1.05E+08 | 105100000 | -0.50 | -0.69 | B>B |
| chr11 | 1.05E+08 | 105200000 | -0.54 | -0.69 | B>B |
| chr11 | 1.05E+08 | 105300000 | -0.73 | -0.78 | B>B |
| chr11 | 1.05E+08 | 105400000 | -0.84 | -0.83 | B>B |
| chr11 | 1.05E+08 | 105500000 | -0.80 | -0.82 | B>B |
| chr11 | 1.06E+08 | 105600000 | -0.69 | -0.87 | B>B |
| chr11 | 1.06E+08 | 105700000 | -0.80 | -0.74 | B>B |
| chr11 | 1.06E+08 | 105800000 | -0.63 | -0.48 | B>B |
| chr11 | 1.06E+08 | 105900000 | -0.53 | -0.33 | B>B |
| chr11 | 1.06E+08 | 106000000 | -0.33 | -0.09 | B>B |
| chr11 | 1.06E+08 | 106100000 | -0.49 | -0.28 | B>B |
| chr11 | 1.06E+08 | 106200000 | -0.65 | -0.47 | B>B |
| chr11 | 1.06E+08 | 106300000 | -0.45 | -0.42 | B>B |
| chr11 | 1.06E+08 | 106400000 | -0.57 | -0.46 | B>B |
| chr11 | 1.06E+08 | 106500000 | -0.59 | -0.53 | B>B |
| chr11 | 1.07E+08 | 106600000 | -0.66 | -0.68 | B>B |
| chr11 | 1.07E+08 | 106700000 | -0.73 | -0.79 | B>B |
| chr11 | 1.07E+08 | 106800000 | -0.69 | -0.84 | B>B |
| chr11 | 1.07E+08 | 106900000 | -0.49 | -0.65 | B>B |
| chr11 | 1.07E+08 | 107000000 | -0.71 | -0.74 | B>B |
| chr11 | 1.07E+08 | 107100000 | -0.74 | -0.97 | B>B |
| chr11 | 1.07E+08 | 107200000 | -0.54 | -0.73 | B>B |
| chr11 | 1.07E+08 | 107300000 | -0.22 | -0.33 | B>B |
| chr11 | 1.07E+08 | 107400000 | -0.04 | -0.27 | B>B |
| chr11 | 1.07E+08 | 107500000 | 0.16 | -0.24 | B>B |
| chr11 | 1.08E+08 | 107600000 | 0.22 | -0.44 | A>B |
| chr11 | 1.08E+08 | 107700000 | 0.44 | -0.03 | A>A |
| chr11 | 1.08E+08 | 107800000 | 0.66 | 0.28 | A>A |
| chr11 | 1.08E+08 | 107900000 | 0.77 | 0.33 | A>A |
| chr11 | 1.08E+08 | 108000000 | 0.79 | 0.76 | A>A |
| chr11 | 1.08E+08 | 108100000 | 0.75 | 0.90 | A>A |
| chr11 | 1.08E+08 | 108200000 | 0.72 | 0.83 | A>A |
| chr11 | 1.08E+08 | 108300000 | 0.46 | 0.63 | A>A |

| | | | | | |
|---|---|---|---|---|---|
| chr11 | 1.08E+08 | 108400000 | 0.59 | 0.74 | A>A |
| chr11 | 1.08E+08 | 108500000 | 0.64 | 0.70 | A>A |
| chr11 | 1.09E+08 | 108600000 | 0.16 | 0.25 | A>A |
| chr11 | 1.09E+08 | 108700000 | 0.11 | 0.09 | A>A |
| chr11 | 1.09E+08 | 108800000 | 0.20 | 0.16 | A>A |
| chr11 | 1.09E+08 | 108900000 | 0.00 | -0.05 | B>B |
| chr11 | 1.09E+08 | 109000000 | -0.25 | -0.26 | B>B |
| chr11 | 1.09E+08 | 109100000 | -0.43 | -0.34 | B>B |
| chr11 | 1.09E+08 | 109200000 | -0.61 | -0.41 | B>B |
| chr11 | 1.09E+08 | 109300000 | -0.60 | -0.48 | B>B |
| chr11 | 1.09E+08 | 109400000 | -0.69 | -0.58 | B>B |
| chr11 | 1.09E+08 | 109500000 | -0.70 | -0.48 | B>B |
| chr11 | 1.1E+08 | 109600000 | -0.63 | -0.31 | B>B |
| chr11 | 1.1E+08 | 109700000 | -0.55 | -0.09 | B>B |
| chr11 | 1.1E+08 | 109800000 | -0.38 | -0.13 | B>B |
| chr11 | 1.1E+08 | 109900000 | -0.02 | 0.15 | A>A |
| chr11 | 1.1E+08 | 110000000 | 0.18 | 0.33 | A>A |
| chr11 | 1.1E+08 | 110100000 | 0.20 | 0.28 | A>A |
| chr11 | 1.1E+08 | 110200000 | 0.58 | 0.55 | A>A |
| chr11 | 1.1E+08 | 110300000 | 0.32 | 0.25 | A>A |
| chr11 | 1.1E+08 | 110400000 | -0.25 | -0.45 | B>B |
| chr11 | 1.1E+08 | 110500000 | -0.66 | -0.94 | B>B |
| chr11 | 1.11E+08 | 110600000 | -0.75 | -1.05 | B>B |
| chr11 | 1.11E+08 | 110700000 | -0.75 | -1.11 | B>B |
| chr11 | 1.11E+08 | 110800000 | -0.75 | -1.08 | B>B |
| chr11 | 1.11E+08 | 110900000 | -0.68 | -1.06 | B>B |
| chr11 | 1.11E+08 | 111000000 | -0.67 | -1.06 | B>B |
| chr11 | 1.11E+08 | 111100000 | -0.38 | -0.80 | B>B |
| chr11 | 1.11E+08 | 111200000 | 0.03 | -0.14 | B>B |
| chr11 | 1.11E+08 | 111300000 | 0.18 | -0.01 | A>A |
| chr11 | 1.11E+08 | 111400000 | 0.29 | -0.07 | A>A |
| chr11 | 1.11E+08 | 111500000 | 0.52 | 0.42 | A>A |
| chr11 | 1.12E+08 | 111600000 | 0.86 | 0.74 | A>A |
| chr11 | 1.12E+08 | 111700000 | 0.73 | 0.69 | A>A |
| chr11 | 1.12E+08 | 111800000 | 0.77 | 0.71 | A>A |
| chr11 | 1.12E+08 | 111900000 | 1.07 | 0.84 | A>A |
| chr11 | 1.12E+08 | 112000000 | 0.83 | 0.73 | A>A |
| chr11 | 1.12E+08 | 112100000 | 0.57 | 0.46 | A>A |
| chr11 | 1.12E+08 | 112200000 | 0.09 | -0.02 | A>A |
| chr11 | 1.12E+08 | 112300000 | -0.23 | -0.26 | B>B |
| chr11 | 1.12E+08 | 112400000 | -0.36 | -0.37 | B>B |
| chr11 | 1.12E+08 | 112500000 | -0.43 | -0.63 | B>B |
| chr11 | 1.13E+08 | 112600000 | -0.62 | -0.93 | B>B |
| chr11 | 1.13E+08 | 112700000 | -0.56 | -1.04 | B>B |
| chr11 | 1.13E+08 | 112800000 | -0.63 | -1.02 | B>B |
| chr11 | 1.13E+08 | 112900000 | -0.63 | -1.03 | B>B |
| chr11 | 1.13E+08 | 113000000 | -0.57 | -1.14 | B>B |
| chr11 | 1.13E+08 | 113100000 | -0.39 | -1.04 | A>B |
| chr11 | 1.13E+08 | 113200000 | -0.22 | -0.48 | B>B |
| chr11 | 1.13E+08 | 113300000 | -0.03 | -0.34 | B>B |
| chr11 | 1.13E+08 | 113400000 | -0.14 | -0.47 | B>B |
| chr11 | 1.13E+08 | 113500000 | -0.12 | -0.24 | B>B |
| chr11 | 1.14E+08 | 113600000 | 0.03 | -0.01 | A>A |
| chr11 | 1.14E+08 | 113700000 | 0.32 | 0.08 | A>A |
| chr11 | 1.14E+08 | 113800000 | 0.37 | 0.09 | A>A |
| chr11 | 1.14E+08 | 113900000 | 0.12 | -0.15 | B>B |

| | | | | | |
|---|---|---|---|---|---|
| chr11 | 1.14E+08 | 114000000 | 0.32 | -0.07 | A>A |
| chr11 | 1.14E+08 | 114100000 | 0.28 | -0.14 | A>A |
| chr11 | 1.14E+08 | 114200000 | 0.21 | 0.06 | A>A |
| chr11 | 1.14E+08 | 114300000 | 0.17 | 0.06 | A>A |
| chr11 | 1.14E+08 | 114400000 | -0.12 | -0.19 | B>B |
| chr11 | 1.14E+08 | 114500000 | -0.41 | -0.47 | B>B |
| chr11 | 1.15E+08 | 114600000 | -0.56 | -0.90 | B>B |
| chr11 | 1.15E+08 | 114700000 | -0.43 | -0.88 | B>B |
| chr11 | 1.15E+08 | 114800000 | -0.60 | -0.97 | B>B |
| chr11 | 1.15E+08 | 114900000 | -0.65 | -0.95 | B>B |
| chr11 | 1.15E+08 | 115000000 | -0.38 | -0.76 | B>B |
| chr11 | 1.15E+08 | 115100000 | -0.01 | -0.36 | B>B |
| chr11 | 1.15E+08 | 115200000 | 0.05 | -0.20 | B>B |
| chr11 | 1.15E+08 | 115300000 | 0.23 | 0.02 | A>A |
| chr11 | 1.15E+08 | 115400000 | 0.27 | -0.03 | A>A |
| chr11 | 1.15E+08 | 115500000 | 0.09 | -0.30 | B>B |
| chr11 | 1.16E+08 | 115600000 | 0.08 | -0.42 | B>B |
| chr11 | 1.16E+08 | 115700000 | -0.10 | -0.61 | B>B |
| chr11 | 1.16E+08 | 115800000 | -0.22 | -0.80 | B>B |
| chr11 | 1.16E+08 | 115900000 | -0.31 | -0.78 | B>B |
| chr11 | 1.16E+08 | 116000000 | -0.29 | -0.86 | B>B |
| chr11 | 1.16E+08 | 116100000 | -0.21 | -0.69 | B>B |
| chr11 | 1.16E+08 | 116200000 | -0.24 | -0.60 | B>B |
| chr11 | 1.16E+08 | 116300000 | -0.27 | -0.67 | B>B |
| chr11 | 1.16E+08 | 116400000 | -0.12 | -0.63 | B>B |
| chr11 | 1.16E+08 | 116500000 | 0.05 | -0.57 | A>B |
| chr11 | 1.17E+08 | 116600000 | 0.14 | -0.13 | A>A |
| chr11 | 1.17E+08 | 116700000 | 0.91 | 0.73 | A>A |
| chr11 | 1.17E+08 | 116800000 | 1.03 | 0.96 | A>A |
| chr11 | 1.17E+08 | 116900000 | 1.21 | 1.15 | A>A |
| chr11 | 1.17E+08 | 117000000 | 1.15 | 1.10 | A>A |
| chr11 | 1.17E+08 | 117100000 | 1.23 | 1.10 | A>A |
| chr11 | 1.17E+08 | 117200000 | 1.32 | 1.28 | A>A |
| chr11 | 1.17E+08 | 117300000 | 1.15 | 1.18 | A>A |
| chr11 | 1.17E+08 | 117400000 | 0.99 | 1.12 | A>A |
| chr11 | 1.17E+08 | 117500000 | 1.07 | 1.12 | A>A |
| chr11 | 1.18E+08 | 117600000 | 1.27 | 1.34 | A>A |
| chr11 | 1.18E+08 | 117700000 | 0.93 | 1.11 | A>A |
| chr11 | 1.18E+08 | 117800000 | 1.04 | 1.32 | A>A |
| chr11 | 1.18E+08 | 117900000 | 1.15 | 1.23 | A>A |
| chr11 | 1.18E+08 | 118000000 | 0.96 | 1.09 | A>A |
| chr11 | 1.18E+08 | 118100000 | 1.11 | 1.13 | A>A |
| chr11 | 1.18E+08 | 118200000 | 1.27 | 1.36 | A>A |
| chr11 | 1.18E+08 | 118300000 | 1.24 | 1.38 | A>A |
| chr11 | 1.18E+08 | 118400000 | 1.38 | 1.33 | A>A |
| chr11 | 1.18E+08 | 118500000 | 1.35 | 1.39 | A>A |
| chr11 | 1.19E+08 | 118600000 | 1.16 | 1.42 | A>A |
| chr11 | 1.19E+08 | 118700000 | 1.42 | 1.55 | A>A |
| chr11 | 1.19E+08 | 118800000 | 1.59 | 1.75 | A>A |
| chr11 | 1.19E+08 | 118900000 | 1.22 | 1.31 | A>A |
| chr11 | 1.19E+08 | 119000000 | 1.57 | 1.65 | A>A |
| chr11 | 1.19E+08 | 119100000 | 1.60 | 1.63 | A>A |
| chr11 | 1.19E+08 | 119200000 | 1.44 | 1.45 | A>A |
| chr11 | 1.19E+08 | 119300000 | 1.20 | 1.18 | A>A |
| chr11 | 1.19E+08 | 119400000 | 0.97 | 0.91 | A>A |
| chr11 | 1.19E+08 | 119500000 | 1.01 | 0.85 | A>A |

| | | | | | |
|---|---|---|---|---|---|
| chr11 | 1.2E+08 | 119600000 | 0.87 | 0.69 | A>A |
| chr11 | 1.2E+08 | 119700000 | 0.84 | 0.63 | A>A |
| chr11 | 1.2E+08 | 119800000 | 0.71 | 0.43 | A>A |
| chr11 | 1.2E+08 | 119900000 | 0.69 | 0.39 | A>A |
| chr11 | 1.2E+08 | 120000000 | 0.65 | 0.54 | A>A |
| chr11 | 1.2E+08 | 120100000 | 0.73 | 0.70 | A>A |
| chr11 | 1.2E+08 | 120200000 | 0.84 | 0.67 | A>A |
| chr11 | 1.2E+08 | 120300000 | 1.10 | 0.91 | A>A |
| chr11 | 1.2E+08 | 120400000 | 0.64 | 0.45 | A>A |
| chr11 | 1.2E+08 | 120500000 | 0.48 | 0.21 | A>A |
| chr11 | 1.21E+08 | 120600000 | 0.40 | 0.08 | A>A |
| chr11 | 1.21E+08 | 120700000 | 0.35 | 0.04 | A>A |
| chr11 | 1.21E+08 | 120800000 | 0.39 | -0.06 | A>A |
| chr11 | 1.21E+08 | 120900000 | 0.33 | 0.01 | A>A |
| chr11 | 1.21E+08 | 121000000 | 0.35 | 0.14 | A>A |
| chr11 | 1.21E+08 | 121100000 | 0.06 | -0.16 | B>B |
| chr11 | 1.21E+08 | 121200000 | 0.14 | -0.09 | A>A |
| chr11 | 1.21E+08 | 121300000 | 0.14 | 0.05 | A>A |
| chr11 | 1.21E+08 | 121400000 | 0.46 | 0.31 | A>A |
| chr11 | 1.21E+08 | 121500000 | 0.19 | 0.06 | A>A |
| chr11 | 1.22E+08 | 121600000 | -0.33 | -0.54 | B>B |
| chr11 | 1.22E+08 | 121700000 | -0.44 | -0.89 | B>B |
| chr11 | 1.22E+08 | 121800000 | -0.47 | -0.86 | B>B |
| chr11 | 1.22E+08 | 121900000 | -0.26 | -0.68 | B>B |
| chr11 | 1.22E+08 | 122000000 | 0.04 | -0.14 | B>B |
| chr11 | 1.22E+08 | 122100000 | 0.23 | -0.06 | A>A |
| chr11 | 1.22E+08 | 122200000 | 0.02 | -0.58 | A>B |
| chr11 | 1.22E+08 | 122300000 | 0.01 | -0.67 | A>B |
| chr11 | 1.22E+08 | 122400000 | -0.05 | -0.44 | B>B |
| chr11 | 1.22E+08 | 122500000 | 0.31 | 0.04 | A>A |
| chr11 | 1.23E+08 | 122600000 | 0.98 | 0.79 | A>A |
| chr11 | 1.23E+08 | 122700000 | 0.77 | 0.63 | A>A |
| chr11 | 1.23E+08 | 122800000 | 0.87 | 0.64 | A>A |
| chr11 | 1.23E+08 | 122900000 | 0.89 | 0.60 | A>A |
| chr11 | 1.23E+08 | 123000000 | 1.01 | 0.87 | A>A |
| chr11 | 1.23E+08 | 123100000 | 1.28 | 1.11 | A>A |
| chr11 | 1.23E+08 | 123200000 | 0.97 | 0.67 | A>A |
| chr11 | 1.23E+08 | 123300000 | 0.98 | 0.75 | A>A |
| chr11 | 1.23E+08 | 123400000 | 0.86 | 0.78 | A>A |
| chr11 | 1.23E+08 | 123500000 | 0.63 | 0.56 | A>A |
| chr11 | 1.24E+08 | 123600000 | 0.01 | -0.12 | B>B |
| chr11 | 1.24E+08 | 123700000 | -0.64 | -1.05 | B>B |
| chr11 | 1.24E+08 | 123800000 | -0.92 | -1.39 | B>B |
| chr11 | 1.24E+08 | 123900000 | -1.06 | -1.30 | B>B |
| chr11 | 1.24E+08 | 124000000 | -0.64 | -1.12 | B>B |
| chr11 | 1.24E+08 | 124100000 | -0.76 | -1.32 | B>B |
| chr11 | 1.24E+08 | 124200000 | -0.98 | -1.31 | B>B |
| chr11 | 1.24E+08 | 124300000 | -0.72 | -1.09 | B>B |
| chr11 | 1.24E+08 | 124400000 | -0.78 | -1.08 | B>B |
| chr11 | 1.24E+08 | 124500000 | -0.47 | -0.75 | B>B |
| chr11 | 1.25E+08 | 124600000 | 0.23 | 0.03 | A>A |
| chr11 | 1.25E+08 | 124700000 | 0.43 | 0.34 | A>A |
| chr11 | 1.25E+08 | 124800000 | 0.32 | 0.10 | A>A |
| chr11 | 1.25E+08 | 124900000 | 0.21 | 0.09 | A>A |
| chr11 | 1.25E+08 | 125000000 | 0.29 | 0.05 | A>A |
| chr11 | 1.25E+08 | 125100000 | 0.35 | 0.02 | A>A |

| | | | | | |
|---|---|---|---|---|---|
| chr11 | 1.25E+08 | 125200000 | 0.37 | -0.09 | A>A |
| chr11 | 1.25E+08 | 125300000 | 0.33 | -0.02 | A>A |
| chr11 | 1.25E+08 | 125400000 | 0.42 | 0.25 | A>A |
| chr11 | 1.25E+08 | 125500000 | 0.66 | 0.59 | A>A |
| chr11 | 1.26E+08 | 125600000 | 0.33 | 0.27 | A>A |
| chr11 | 1.26E+08 | 125700000 | -0.27 | -0.12 | B>B |
| chr11 | 1.26E+08 | 125800000 | 0.41 | 0.50 | A>A |
| chr11 | 1.26E+08 | 125900000 | 1.03 | 0.95 | A>A |
| chr11 | 1.26E+08 | 126000000 | 1.05 | 1.17 | A>A |
| chr11 | 1.26E+08 | 126100000 | 0.90 | 1.03 | A>A |
| chr11 | 1.26E+08 | 126200000 | 0.89 | 0.84 | A>A |
| chr11 | 1.26E+08 | 126300000 | 0.59 | 0.63 | A>A |
| chr11 | 1.26E+08 | 126400000 | 0.24 | 0.25 | A>A |
| chr11 | 1.26E+08 | 126500000 | 0.19 | 0.04 | A>A |
| chr11 | 1.27E+08 | 126600000 | 0.14 | -0.03 | A>A |
| chr11 | 1.27E+08 | 126700000 | 0.23 | 0.09 | A>A |
| chr11 | 1.27E+08 | 126800000 | 0.27 | 0.17 | A>A |
| chr11 | 1.27E+08 | 126900000 | 0.22 | 0.07 | A>A |
| chr11 | 1.27E+08 | 127000000 | -0.13 | -0.33 | B>B |
| chr11 | 1.27E+08 | 127100000 | -0.55 | -0.83 | B>B |
| chr11 | 1.27E+08 | 127200000 | -0.80 | -1.04 | B>B |
| chr11 | 1.27E+08 | 127300000 | -0.88 | -1.06 | B>B |
| chr11 | 1.27E+08 | 127400000 | -0.91 | -1.18 | B>B |
| chr11 | 1.27E+08 | 127500000 | -0.94 | -1.27 | B>B |
| chr11 | 1.28E+08 | 127600000 | -1.02 | -1.29 | B>B |
| chr11 | 1.28E+08 | 127700000 | -1.02 | -1.30 | B>B |
| chr11 | 1.28E+08 | 127800000 | -0.98 | -1.27 | B>B |
| chr11 | 1.28E+08 | 127900000 | -0.97 | -1.07 | B>B |
| chr11 | 1.28E+08 | 128000000 | -0.76 | -0.81 | B>B |
| chr11 | 1.28E+08 | 128100000 | -0.16 | -0.36 | B>B |
| chr11 | 1.28E+08 | 128200000 | -0.09 | -0.39 | B>B |
| chr11 | 1.28E+08 | 128300000 | -0.01 | -0.39 | B>B |
| chr11 | 1.28E+08 | 128400000 | 0.25 | -0.05 | A>A |
| chr11 | 1.28E+08 | 128500000 | 0.16 | -0.23 | B>B |
| chr11 | 1.29E+08 | 128600000 | -0.16 | -0.54 | B>B |
| chr11 | 1.29E+08 | 128700000 | -0.15 | -0.54 | B>B |
| chr11 | 1.29E+08 | 128800000 | 0.02 | -0.14 | B>B |
| chr11 | 1.29E+08 | 128900000 | 0.13 | -0.05 | A>A |
| chr11 | 1.29E+08 | 129000000 | 0.09 | 0.04 | A>A |
| chr11 | 1.29E+08 | 129100000 | 0.34 | 0.31 | A>A |
| chr11 | 1.29E+08 | 129200000 | 0.37 | 0.32 | A>A |
| chr11 | 1.29E+08 | 129300000 | -0.11 | -0.29 | B>B |
| chr11 | 1.29E+08 | 129400000 | -0.46 | -0.53 | B>B |
| chr11 | 1.29E+08 | 129500000 | -0.38 | -0.53 | B>B |
| chr11 | 1.3E+08 | 129600000 | -0.13 | -0.24 | B>B |
| chr11 | 1.3E+08 | 129700000 | 0.22 | 0.35 | A>A |
| chr11 | 1.3E+08 | 129800000 | 0.67 | 0.72 | A>A |
| chr11 | 1.3E+08 | 129900000 | 0.98 | 0.97 | A>A |
| chr11 | 1.3E+08 | 130000000 | 0.99 | 1.04 | A>A |
| chr11 | 1.3E+08 | 130100000 | 0.70 | 0.76 | A>A |
| chr11 | 1.3E+08 | 130200000 | 0.93 | 0.99 | A>A |
| chr11 | 1.3E+08 | 130300000 | 0.58 | 0.80 | A>A |
| chr11 | 1.3E+08 | 130400000 | 0.79 | 0.88 | A>A |
| chr11 | 1.3E+08 | 130500000 | 0.99 | 1.20 | A>A |
| chr11 | 1.31E+08 | 130600000 | 0.91 | 1.02 | A>A |
| chr11 | 1.31E+08 | 130700000 | 0.85 | 0.92 | A>A |

| | | | | | |
|---|---|---|---|---|---|
| chr11 | 1.31E+08 | 130800000 | 0.25 | 0.45 | A>A |
| chr11 | 1.31E+08 | 130900000 | -0.29 | -0.20 | B>B |
| chr11 | 1.31E+08 | 131000000 | -0.47 | -0.72 | B>B |
| chr11 | 1.31E+08 | 131100000 | -0.63 | -1.01 | B>B |
| chr11 | 1.31E+08 | 131200000 | -0.81 | -1.03 | B>B |
| chr11 | 1.31E+08 | 131300000 | -0.74 | -1.07 | B>B |
| chr11 | 1.31E+08 | 131400000 | -0.71 | -1.00 | B>B |
| chr11 | 1.31E+08 | 131500000 | -0.57 | -1.05 | B>B |
| chr11 | 1.32E+08 | 131600000 | -0.58 | -0.81 | B>B |
| chr11 | 1.32E+08 | 131700000 | -0.92 | -1.06 | B>B |
| chr11 | 1.32E+08 | 131800000 | -1.02 | -1.13 | B>B |
| chr11 | 1.32E+08 | 131900000 | -1.08 | -1.12 | B>B |
| chr11 | 1.32E+08 | 132000000 | -1.10 | -1.06 | B>B |
| chr11 | 1.32E+08 | 132100000 | -1.13 | -1.17 | B>B |
| chr11 | 1.32E+08 | 132200000 | -1.12 | -1.08 | B>B |
| chr11 | 1.32E+08 | 132300000 | -1.14 | -1.18 | B>B |
| chr11 | 1.32E+08 | 132400000 | -1.15 | -1.17 | B>B |
| chr11 | 1.32E+08 | 132500000 | -1.23 | -1.26 | B>B |
| chr11 | 1.33E+08 | 132600000 | -1.14 | -1.28 | B>B |
| chr11 | 1.33E+08 | 132700000 | -1.06 | -1.28 | B>B |
| chr11 | 1.33E+08 | 132800000 | -1.03 | -1.27 | B>B |
| chr11 | 1.33E+08 | 132900000 | -0.93 | -1.17 | B>B |
| chr11 | 1.33E+08 | 133000000 | -0.77 | -1.03 | B>B |
| chr11 | 1.33E+08 | 133100000 | -0.91 | -1.29 | B>B |
| chr11 | 1.33E+08 | 133200000 | -1.16 | -1.34 | B>B |
| chr11 | 1.33E+08 | 133300000 | -1.20 | -1.38 | B>B |
| chr11 | 1.33E+08 | 133400000 | -1.17 | -1.35 | B>B |
| chr11 | 1.33E+08 | 133500000 | -1.18 | -1.36 | B>B |
| chr11 | 1.34E+08 | 133600000 | -1.09 | -1.30 | B>B |
| chr11 | 1.34E+08 | 133700000 | -1.00 | -1.19 | B>B |
| chr11 | 1.34E+08 | 133800000 | -0.48 | -0.88 | B>B |
| chr11 | 1.34E+08 | 133900000 | 0.16 | -0.34 | B>B |
| chr11 | 1.34E+08 | 134000000 | 0.21 | -0.24 | B>B |
| chr11 | 1.34E+08 | 134100000 | 0.56 | 0.23 | A>A |
| chr11 | 1.34E+08 | 134200000 | 0.42 | 0.20 | A>A |
| chr11 | 1.34E+08 | 134300000 | -0.03 | -0.20 | B>B |
| chr11 | 1.34E+08 | 134400000 | -0.14 | -0.47 | B>B |
| chr11 | 1.34E+08 | 134500000 | -0.31 | -0.51 | B>B |
| chr11 | 1.35E+08 | 134600000 | -0.21 | -0.41 | B>B |
| chr11 | 1.35E+08 | 134700000 | -0.40 | -0.31 | B>B |
| chr11 | 1.35E+08 | 134800000 | -0.44 | -0.56 | B>B |
| chr11 | 1.35E+08 | 134900000 | -0.62 | -0.78 | B>B |
| chr11 | 1.35E+08 | 135000000 | 0.00 | 0.00 | NO |
| chr11 | 1.35E+08 | 135006516 | 0.00 | 0.00 | NO |
| chr12 | 0 | 100000 | 0.00 | 0.00 | NO |
| chr12 | 100000 | 200000 | 0.00 | 0.00 | NO |
| chr12 | 200000 | 300000 | 1.21 | 1.09 | A>A |
| chr12 | 300000 | 400000 | 1.20 | 1.28 | A>A |
| chr12 | 400000 | 500000 | 1.23 | 1.28 | A>A |
| chr12 | 500000 | 600000 | 1.24 | 1.19 | A>A |
| chr12 | 600000 | 700000 | 0.99 | 1.16 | A>A |
| chr12 | 700000 | 800000 | 1.42 | 1.36 | A>A |
| chr12 | 800000 | 900000 | 1.29 | 1.24 | A>A |
| chr12 | 900000 | 1000000 | 1.04 | 1.21 | A>A |
| chr12 | 1000000 | 1100000 | 0.94 | 1.03 | A>A |
| chr12 | 1100000 | 1200000 | 0.93 | 0.95 | A>A |

| | | | | | |
|---|---|---|---|---|---|
| chr12 | 1200000 | 1300000 | 0.58 | 0.67 | A>A |
| chr12 | 1300000 | 1400000 | 0.66 | 0.82 | A>A |
| chr12 | 1400000 | 1500000 | 0.52 | 0.61 | A>A |
| chr12 | 1500000 | 1600000 | 0.69 | 0.65 | A>A |
| chr12 | 1600000 | 1700000 | 0.66 | 0.67 | A>A |
| chr12 | 1700000 | 1800000 | 0.64 | 0.49 | A>A |
| chr12 | 1800000 | 1900000 | 0.81 | 0.50 | A>A |
| chr12 | 1900000 | 2000000 | 0.43 | 0.02 | A>A |
| chr12 | 2000000 | 2100000 | 0.33 | -0.05 | A>A |
| chr12 | 2100000 | 2200000 | 0.16 | -0.35 | B>B |
| chr12 | 2200000 | 2300000 | 0.02 | -0.79 | A>B |
| chr12 | 2300000 | 2400000 | -0.11 | -0.98 | A>B |
| chr12 | 2400000 | 2500000 | -0.01 | -1.04 | A>B |
| chr12 | 2500000 | 2600000 | -0.04 | -0.92 | A>B |
| chr12 | 2600000 | 2700000 | 0.03 | -0.76 | A>B |
| chr12 | 2700000 | 2800000 | 0.14 | -0.58 | A>B |
| chr12 | 2800000 | 2900000 | 0.39 | -0.34 | A>B |
| chr12 | 2900000 | 3000000 | 1.03 | 0.44 | A>B |
| chr12 | 3000000 | 3100000 | 1.12 | 0.71 | A>A |
| chr12 | 3100000 | 3200000 | 0.86 | 0.46 | A>A |
| chr12 | 3200000 | 3300000 | 0.59 | 0.13 | A>A |
| chr12 | 3300000 | 3400000 | 0.51 | 0.07 | A>A |
| chr12 | 3400000 | 3500000 | 0.00 | -0.44 | B>B |
| chr12 | 3500000 | 3600000 | -0.16 | -0.78 | A>B |
| chr12 | 3600000 | 3700000 | -0.18 | -0.81 | A>B |
| chr12 | 3700000 | 3800000 | -0.10 | -0.80 | A>B |
| chr12 | 3800000 | 3900000 | -0.01 | -0.62 | A>B |
| chr12 | 3900000 | 4000000 | -0.09 | -0.48 | B>B |
| chr12 | 4000000 | 4100000 | -0.14 | -0.54 | B>B |
| chr12 | 4100000 | 4200000 | -0.14 | -0.51 | B>B |
| chr12 | 4200000 | 4300000 | -0.23 | -0.62 | B>B |
| chr12 | 4300000 | 4400000 | -0.18 | -0.52 | B>B |
| chr12 | 4400000 | 4500000 | 0.13 | -0.34 | B>B |
| chr12 | 4500000 | 4600000 | -0.03 | -0.49 | B>B |
| chr12 | 4600000 | 4700000 | 0.10 | -0.32 | B>B |
| chr12 | 4700000 | 4800000 | 0.08 | -0.43 | B>B |
| chr12 | 4800000 | 4900000 | -0.29 | -0.80 | B>B |
| chr12 | 4900000 | 5000000 | -0.33 | -0.88 | B>B |
| chr12 | 5000000 | 5100000 | -0.36 | -1.03 | A>B |
| chr12 | 5100000 | 5200000 | -0.44 | -0.98 | B>B |
| chr12 | 5200000 | 5300000 | -0.52 | -1.00 | B>B |
| chr12 | 5300000 | 5400000 | -0.65 | -1.12 | B>B |
| chr12 | 5400000 | 5500000 | -0.70 | -1.08 | B>B |
| chr12 | 5500000 | 5600000 | -0.44 | -1.06 | A>B |
| chr12 | 5600000 | 5700000 | -0.47 | -1.06 | A>B |
| chr12 | 5700000 | 5800000 | -0.42 | -1.12 | A>B |
| chr12 | 5800000 | 5900000 | -0.43 | -1.10 | A>B |
| chr12 | 5900000 | 6000000 | -0.32 | -0.95 | A>B |
| chr12 | 6000000 | 6100000 | 0.05 | -0.55 | A>B |
| chr12 | 6100000 | 6200000 | 0.55 | 0.06 | A>A |
| chr12 | 6200000 | 6300000 | 0.95 | 0.50 | A>A |
| chr12 | 6300000 | 6400000 | 1.04 | 0.61 | A>A |
| chr12 | 6400000 | 6500000 | 1.56 | 1.21 | A>A |
| chr12 | 6500000 | 6600000 | 1.48 | 1.32 | A>A |
| chr12 | 6600000 | 6700000 | 1.52 | 1.52 | A>A |
| chr12 | 6700000 | 6800000 | 1.80 | 1.51 | A>A |

| | | | | | |
|---|---|---|---|---|---|
| chr12 | 6800000 | 6900000 | 1.67 | 1.61 | A>A |
| chr12 | 6900000 | 7000000 | 1.58 | 1.55 | A>A |
| chr12 | 7000000 | 7100000 | 1.44 | 1.35 | A>A |
| chr12 | 7100000 | 7200000 | 1.24 | 1.02 | A>A |
| chr12 | 7200000 | 7300000 | 0.92 | 0.66 | A>A |
| chr12 | 7300000 | 7400000 | 0.35 | 0.03 | A>A |
| chr12 | 7400000 | 7500000 | -0.17 | -0.42 | B>B |
| chr12 | 7500000 | 7600000 | -0.04 | -0.44 | B>B |
| chr12 | 7600000 | 7700000 | -0.12 | -0.47 | B>B |
| chr12 | 7700000 | 7800000 | -0.07 | -0.43 | B>B |
| chr12 | 7800000 | 7900000 | 0.37 | -0.16 | A>A |
| chr12 | 7900000 | 8000000 | 0.56 | -0.13 | A>B |
| chr12 | 8000000 | 8100000 | 1.02 | 0.32 | A>B |
| chr12 | 8100000 | 8200000 | 0.96 | 0.45 | A>A |
| chr12 | 8200000 | 8300000 | 0.70 | 0.11 | A>B |
| chr12 | 8300000 | 8400000 | 0.00 | 0.00 | NO |
| chr12 | 8400000 | 8500000 | 0.00 | 0.00 | NO |
| chr12 | 8500000 | 8600000 | 0.00 | 0.00 | NO |
| chr12 | 8600000 | 8700000 | -0.22 | -0.54 | B>B |
| chr12 | 8700000 | 8800000 | 0.06 | -0.36 | B>B |
| chr12 | 8800000 | 8900000 | 0.38 | 0.11 | A>A |
| chr12 | 8900000 | 9000000 | 0.47 | -0.03 | A>A |
| chr12 | 9000000 | 9100000 | 0.37 | 0.00 | A>A |
| chr12 | 9100000 | 9200000 | 0.15 | -0.18 | B>B |
| chr12 | 9200000 | 9300000 | -0.31 | -0.40 | B>B |
| chr12 | 9300000 | 9400000 | -0.39 | -0.53 | B>B |
| chr12 | 9400000 | 9500000 | 0.01 | -0.17 | B>B |
| chr12 | 9500000 | 9600000 | 0.00 | 0.00 | NO |
| chr12 | 9600000 | 9700000 | 0.00 | 0.00 | NO |
| chr12 | 9700000 | 9800000 | 0.00 | 0.11 | NO |
| chr12 | 9800000 | 9900000 | 0.32 | 0.30 | A>A |
| chr12 | 9900000 | 10000000 | -0.06 | 0.00 | B>B |
| chr12 | 10000000 | 10100000 | -0.14 | -0.05 | B>B |
| chr12 | 10100000 | 10200000 | -0.29 | -0.14 | B>B |
| chr12 | 10200000 | 10300000 | -0.24 | -0.04 | B>B |
| chr12 | 10300000 | 10400000 | -0.01 | 0.07 | A>A |
| chr12 | 10400000 | 10500000 | -0.22 | 0.08 | B>B |
| chr12 | 10500000 | 10600000 | 0.00 | 0.48 | NO |
| chr12 | 10600000 | 10700000 | -0.12 | 0.29 | A>A |
| chr12 | 10700000 | 10800000 | 0.00 | 0.12 | A>A |
| chr12 | 10800000 | 10900000 | 0.01 | 0.06 | A>A |
| chr12 | 10900000 | 11000000 | -0.31 | -0.29 | B>B |
| chr12 | 11000000 | 11100000 | -0.29 | -0.23 | B>B |
| chr12 | 11100000 | 11200000 | -0.19 | -0.12 | B>B |
| chr12 | 11200000 | 11300000 | 0.00 | 0.00 | NO |
| chr12 | 11300000 | 11400000 | -0.39 | -0.28 | B>B |
| chr12 | 11400000 | 11500000 | 0.00 | -0.79 | NO |
| chr12 | 11500000 | 11600000 | 0.00 | -0.60 | NO |
| chr12 | 11600000 | 11700000 | -0.30 | -0.20 | B>B |
| chr12 | 11700000 | 11800000 | 0.22 | 0.25 | A>A |
| chr12 | 11800000 | 11900000 | 0.34 | 0.40 | A>A |
| chr12 | 11900000 | 12000000 | 0.36 | 0.47 | A>A |
| chr12 | 12000000 | 12100000 | 0.35 | 0.35 | A>A |
| chr12 | 12100000 | 12200000 | 0.25 | 0.30 | A>A |
| chr12 | 12200000 | 12300000 | 0.63 | 0.63 | A>A |
| chr12 | 12300000 | 12400000 | 0.95 | 0.83 | A>A |

| | | | | | |
|---|---|---|---|---|---|
| chr12 | 12400000 | 12500000 | 0.78 | 0.82 | A>A |
| chr12 | 12500000 | 12600000 | 0.83 | 0.87 | A>A |
| chr12 | 12600000 | 12700000 | 1.09 | 1.03 | A>A |
| chr12 | 12700000 | 12800000 | 1.15 | 0.92 | A>A |
| chr12 | 12800000 | 12900000 | 0.78 | 0.72 | A>A |
| chr12 | 12900000 | 13000000 | 1.03 | 0.96 | A>A |
| chr12 | 13000000 | 13100000 | 1.11 | 0.97 | A>A |
| chr12 | 13100000 | 13200000 | 0.69 | 0.60 | A>A |
| chr12 | 13200000 | 13300000 | 0.81 | 0.53 | A>A |
| chr12 | 13300000 | 13400000 | 0.72 | 0.44 | A>A |
| chr12 | 13400000 | 13500000 | 0.21 | -0.12 | A>A |
| chr12 | 13500000 | 13600000 | 0.21 | 0.02 | A>A |
| chr12 | 13600000 | 13700000 | 0.06 | -0.08 | B>B |
| chr12 | 13700000 | 13800000 | 0.03 | -0.32 | B>B |
| chr12 | 13800000 | 13900000 | -0.07 | -0.39 | B>B |
| chr12 | 13900000 | 14000000 | 0.01 | -0.46 | B>B |
| chr12 | 14000000 | 14100000 | 0.05 | -0.27 | B>B |
| chr12 | 14100000 | 14200000 | -0.11 | -0.32 | B>B |
| chr12 | 14200000 | 14300000 | -0.25 | -0.45 | B>B |
| chr12 | 14300000 | 14400000 | -0.16 | -0.08 | B>B |
| chr12 | 14400000 | 14500000 | -0.07 | 0.55 | B>A |
| chr12 | 14500000 | 14600000 | 0.16 | 0.69 | B>A |
| chr12 | 14600000 | 14700000 | -0.07 | 0.31 | A>A |
| chr12 | 14700000 | 14800000 | -0.18 | 0.17 | B>B |
| chr12 | 14800000 | 14900000 | -0.21 | 0.05 | B>B |
| chr12 | 14900000 | 15000000 | 0.14 | 0.08 | A>A |
| chr12 | 15000000 | 15100000 | -0.04 | -0.15 | B>B |
| chr12 | 15100000 | 15200000 | -0.17 | -0.16 | B>B |
| chr12 | 15200000 | 15300000 | -0.35 | -0.34 | B>B |
| chr12 | 15300000 | 15400000 | -0.15 | 0.00 | B>B |
| chr12 | 15400000 | 15500000 | -0.48 | -0.30 | B>B |
| chr12 | 15500000 | 15600000 | -0.48 | -0.40 | B>B |
| chr12 | 15600000 | 15700000 | -0.32 | -0.24 | B>B |
| chr12 | 15700000 | 15800000 | 0.18 | 0.31 | A>A |
| chr12 | 15800000 | 15900000 | 0.51 | 0.64 | A>A |
| chr12 | 15900000 | 16000000 | 0.33 | 0.49 | A>A |
| chr12 | 16000000 | 16100000 | 0.12 | 0.26 | A>A |
| chr12 | 16100000 | 16200000 | -0.14 | 0.10 | B>B |
| chr12 | 16200000 | 16300000 | -0.43 | 0.02 | B>B |
| chr12 | 16300000 | 16400000 | -0.44 | -0.07 | B>B |
| chr12 | 16400000 | 16500000 | -0.34 | 0.00 | B>B |
| chr12 | 16500000 | 16600000 | -0.25 | -0.07 | B>B |
| chr12 | 16600000 | 16700000 | -0.37 | -0.23 | B>B |
| chr12 | 16700000 | 16800000 | -0.36 | -0.27 | B>B |
| chr12 | 16800000 | 16900000 | -0.42 | -0.43 | B>B |
| chr12 | 16900000 | 17000000 | -0.33 | -0.42 | B>B |
| chr12 | 17000000 | 17100000 | -0.21 | -0.21 | B>B |
| chr12 | 17100000 | 17200000 | -0.03 | 0.15 | A>A |
| chr12 | 17200000 | 17300000 | 0.10 | 0.29 | A>A |
| chr12 | 17300000 | 17400000 | 0.06 | 0.22 | A>A |
| chr12 | 17400000 | 17500000 | 0.13 | 0.32 | A>A |
| chr12 | 17500000 | 17600000 | 0.12 | 0.23 | A>A |
| chr12 | 17600000 | 17700000 | 0.06 | 0.03 | A>A |
| chr12 | 17700000 | 17800000 | -0.08 | 0.04 | B>B |
| chr12 | 17800000 | 17900000 | -0.32 | 0.04 | B>B |
| chr12 | 17900000 | 18000000 | -0.43 | -0.15 | B>B |

| | | | | | |
|---|---|---|---|---|---|
| chr12 | 18000000 | 18100000 | -0.40 | -0.08 | B>B |
| chr12 | 18100000 | 18200000 | -0.40 | 0.00 | B>B |
| chr12 | 18200000 | 18300000 | -0.54 | 0.17 | B>A |
| chr12 | 18300000 | 18400000 | -0.51 | 0.31 | B>A |
| chr12 | 18400000 | 18500000 | -0.60 | 0.31 | B>A |
| chr12 | 18500000 | 18600000 | -0.70 | 0.35 | B>A |
| chr12 | 18600000 | 18700000 | -0.73 | 0.37 | B>A |
| chr12 | 18700000 | 18800000 | -0.73 | 0.35 | B>A |
| chr12 | 18800000 | 18900000 | -0.59 | 0.44 | B>A |
| chr12 | 18900000 | 19000000 | -0.52 | 0.43 | B>A |
| chr12 | 19000000 | 19100000 | -0.22 | 0.55 | B>A |
| chr12 | 19100000 | 19200000 | -0.14 | 0.69 | B>A |
| chr12 | 19200000 | 19300000 | 0.01 | 0.65 | B>A |
| chr12 | 19300000 | 19400000 | -0.12 | 0.50 | B>A |
| chr12 | 19400000 | 19500000 | 0.08 | 0.61 | B>A |
| chr12 | 19500000 | 19600000 | 0.26 | 0.83 | B>A |
| chr12 | 19600000 | 19700000 | 0.37 | 0.77 | A>A |
| chr12 | 19700000 | 19800000 | 0.11 | 0.68 | B>A |
| chr12 | 19800000 | 19900000 | 0.00 | 0.54 | B>A |
| chr12 | 19900000 | 20000000 | -0.03 | 0.69 | B>A |
| chr12 | 20000000 | 20100000 | 0.02 | 0.68 | B>A |
| chr12 | 20100000 | 20200000 | 0.12 | 0.80 | B>A |
| chr12 | 20200000 | 20300000 | 0.04 | 0.78 | B>A |
| chr12 | 20300000 | 20400000 | 0.14 | 0.64 | A>A |
| chr12 | 20400000 | 20500000 | 0.17 | 0.58 | A>A |
| chr12 | 20500000 | 20600000 | 0.15 | 0.51 | A>A |
| chr12 | 20600000 | 20700000 | 0.30 | 0.81 | A>A |
| chr12 | 20700000 | 20800000 | 0.19 | 0.66 | A>A |
| chr12 | 20800000 | 20900000 | -0.11 | 0.44 | B>A |
| chr12 | 20900000 | 21000000 | -0.01 | 0.49 | A>A |
| chr12 | 21000000 | 21100000 | 0.01 | 0.54 | B>A |
| chr12 | 21100000 | 21200000 | -0.23 | 0.39 | B>A |
| chr12 | 21200000 | 21300000 | -0.17 | 0.51 | B>A |
| chr12 | 21300000 | 21400000 | -0.26 | 0.30 | B>A |
| chr12 | 21400000 | 21500000 | -0.26 | 0.21 | B>B |
| chr12 | 21500000 | 21600000 | -0.11 | 0.19 | A>A |
| chr12 | 21600000 | 21700000 | 0.13 | 0.40 | A>A |
| chr12 | 21700000 | 21800000 | 0.04 | 0.34 | A>A |
| chr12 | 21800000 | 21900000 | -0.15 | 0.24 | A>A |
| chr12 | 21900000 | 22000000 | -0.60 | 0.08 | B>A |
| chr12 | 22000000 | 22100000 | -0.63 | 0.03 | B>A |
| chr12 | 22100000 | 22200000 | -0.45 | 0.01 | B>B |
| chr12 | 22200000 | 22300000 | -0.24 | -0.23 | B>B |
| chr12 | 22300000 | 22400000 | -0.25 | -0.35 | B>B |
| chr12 | 22400000 | 22500000 | -0.24 | -0.30 | B>B |
| chr12 | 22500000 | 22600000 | -0.22 | -0.26 | B>B |
| chr12 | 22600000 | 22700000 | 0.13 | 0.03 | A>A |
| chr12 | 22700000 | 22800000 | 0.42 | 0.29 | A>A |
| chr12 | 22800000 | 22900000 | -0.16 | -0.44 | B>B |
| chr12 | 22900000 | 23000000 | -0.48 | -0.85 | B>B |
| chr12 | 23000000 | 23100000 | -0.53 | -0.92 | B>B |
| chr12 | 23100000 | 23200000 | -0.64 | -0.98 | B>B |
| chr12 | 23200000 | 23300000 | -0.65 | -0.92 | B>B |
| chr12 | 23300000 | 23400000 | -0.65 | -0.78 | B>B |
| chr12 | 23400000 | 23500000 | -0.68 | -0.90 | B>B |
| chr12 | 23500000 | 23600000 | -0.68 | -0.88 | B>B |

| | | | | | |
|---|---|---|---|---|---|
| chr12 | 23600000 | 23700000 | -0.67 | -0.88 | B>B |
| chr12 | 23700000 | 23800000 | -0.71 | -0.92 | B>B |
| chr12 | 23800000 | 23900000 | -0.74 | -0.87 | B>B |
| chr12 | 23900000 | 24000000 | -0.67 | -0.90 | B>B |
| chr12 | 24000000 | 24100000 | -0.71 | -0.82 | B>B |
| chr12 | 24100000 | 24200000 | -0.67 | -0.86 | B>B |
| chr12 | 24200000 | 24300000 | -0.70 | -0.87 | B>B |
| chr12 | 24300000 | 24400000 | -0.71 | -0.88 | B>B |
| chr12 | 24400000 | 24500000 | -0.64 | -0.88 | B>B |
| chr12 | 24500000 | 24600000 | -0.67 | -0.91 | B>B |
| chr12 | 24600000 | 24700000 | -0.68 | -0.89 | B>B |
| chr12 | 24700000 | 24800000 | -0.67 | -0.92 | B>B |
| chr12 | 24800000 | 24900000 | -0.59 | -0.89 | B>B |
| chr12 | 24900000 | 25000000 | -0.49 | -0.92 | B>B |
| chr12 | 25000000 | 25100000 | -0.14 | -0.77 | A>B |
| chr12 | 25100000 | 25200000 | -0.25 | -0.66 | B>B |
| chr12 | 25200000 | 25300000 | -0.20 | -0.72 | B>B |
| chr12 | 25300000 | 25400000 | 0.06 | -0.25 | B>B |
| chr12 | 25400000 | 25500000 | 0.22 | -0.30 | B>B |
| chr12 | 25500000 | 25600000 | 0.05 | -0.16 | B>B |
| chr12 | 25600000 | 25700000 | -0.21 | -0.19 | B>B |
| chr12 | 25700000 | 25800000 | -0.20 | -0.24 | B>B |
| chr12 | 25800000 | 25900000 | -0.16 | -0.10 | B>B |
| chr12 | 25900000 | 26000000 | -0.09 | -0.14 | B>B |
| chr12 | 26000000 | 26100000 | 0.05 | 0.05 | A>A |
| chr12 | 26100000 | 26200000 | 0.10 | 0.03 | A>A |
| chr12 | 26200000 | 26300000 | 0.15 | -0.05 | A>A |
| chr12 | 26300000 | 26400000 | 0.35 | 0.07 | A>A |
| chr12 | 26400000 | 26500000 | 0.33 | -0.01 | A>A |
| chr12 | 26500000 | 26600000 | 0.16 | -0.22 | B>B |
| chr12 | 26600000 | 26700000 | 0.05 | -0.17 | B>B |
| chr12 | 26700000 | 26800000 | 0.07 | -0.23 | B>B |
| chr12 | 26800000 | 26900000 | 0.24 | 0.00 | A>A |
| chr12 | 26900000 | 27000000 | 0.44 | 0.24 | A>A |
| chr12 | 27000000 | 27100000 | 0.66 | 0.48 | A>A |
| chr12 | 27100000 | 27200000 | 0.65 | 0.51 | A>A |
| chr12 | 27200000 | 27300000 | 0.00 | 0.44 | NO |
| chr12 | 27300000 | 27400000 | 0.70 | 0.50 | A>A |
| chr12 | 27400000 | 27500000 | 0.62 | 0.41 | A>A |
| chr12 | 27500000 | 27600000 | 0.50 | 0.43 | A>A |
| chr12 | 27600000 | 27700000 | 0.71 | 0.53 | A>A |
| chr12 | 27700000 | 27800000 | 0.70 | 0.58 | A>A |
| chr12 | 27800000 | 27900000 | 0.40 | 0.31 | A>A |
| chr12 | 27900000 | 28000000 | 0.38 | 0.16 | A>A |
| chr12 | 28000000 | 28100000 | 0.21 | -0.01 | A>A |
| chr12 | 28100000 | 28200000 | 0.25 | 0.17 | A>A |
| chr12 | 28200000 | 28300000 | 0.32 | 0.16 | A>A |
| chr12 | 28300000 | 28400000 | 0.16 | 0.05 | A>A |
| chr12 | 28400000 | 28500000 | 0.14 | 0.06 | A>A |
| chr12 | 28500000 | 28600000 | 0.04 | -0.07 | B>B |
| chr12 | 28600000 | 28700000 | -0.11 | -0.15 | B>B |
| chr12 | 28700000 | 28800000 | -0.14 | -0.25 | B>B |
| chr12 | 28800000 | 28900000 | -0.25 | -0.28 | B>B |
| chr12 | 28900000 | 29000000 | -0.22 | -0.33 | B>B |
| chr12 | 29000000 | 29100000 | -0.32 | -0.47 | B>B |
| chr12 | 29100000 | 29200000 | -0.38 | -0.72 | B>B |

| | | | | | |
|---|---|---|---|---|---|
| chr12 | 29200000 | 29300000 | -0.42 | -0.79 | B>B |
| chr12 | 29300000 | 29400000 | -0.53 | -0.93 | B>B |
| chr12 | 29400000 | 29500000 | -0.39 | -0.69 | B>B |
| chr12 | 29500000 | 29600000 | -0.29 | -0.57 | B>B |
| chr12 | 29600000 | 29700000 | -0.44 | -0.73 | B>B |
| chr12 | 29700000 | 29800000 | -0.46 | -0.74 | B>B |
| chr12 | 29800000 | 29900000 | -0.49 | -0.81 | B>B |
| chr12 | 29900000 | 30000000 | -0.57 | -0.84 | B>B |
| chr12 | 30000000 | 30100000 | -0.71 | -0.90 | B>B |
| chr12 | 30100000 | 30200000 | -0.63 | -0.90 | B>B |
| chr12 | 30200000 | 30300000 | -0.50 | -0.85 | B>B |
| chr12 | 30300000 | 30400000 | -0.53 | -0.86 | B>B |
| chr12 | 30400000 | 30500000 | -0.50 | -0.81 | B>B |
| chr12 | 30500000 | 30600000 | -0.39 | -0.65 | B>B |
| chr12 | 30600000 | 30700000 | -0.20 | -0.56 | B>B |
| chr12 | 30700000 | 30800000 | -0.08 | -0.33 | B>B |
| chr12 | 30800000 | 30900000 | 0.15 | -0.09 | A>A |
| chr12 | 30900000 | 31000000 | 0.03 | -0.22 | B>B |
| chr12 | 31000000 | 31100000 | 0.05 | -0.25 | B>B |
| chr12 | 31100000 | 31200000 | 0.19 | -0.13 | A>A |
| chr12 | 31200000 | 31300000 | 0.00 | 0.00 | NO |
| chr12 | 31300000 | 31400000 | 0.00 | 0.00 | NO |
| chr12 | 31400000 | 31500000 | 0.50 | 0.43 | A>A |
| chr12 | 31500000 | 31600000 | 0.69 | 0.54 | A>A |
| chr12 | 31600000 | 31700000 | 0.82 | 0.78 | A>A |
| chr12 | 31700000 | 31800000 | 0.77 | 0.66 | A>A |
| chr12 | 31800000 | 31900000 | 0.62 | 0.68 | A>A |
| chr12 | 31900000 | 32000000 | 0.94 | 0.90 | A>A |
| chr12 | 32000000 | 32100000 | 0.87 | 0.90 | A>A |
| chr12 | 32100000 | 32200000 | 0.72 | 0.73 | A>A |
| chr12 | 32200000 | 32300000 | 0.70 | 0.68 | A>A |
| chr12 | 32300000 | 32400000 | 0.71 | 0.71 | A>A |
| chr12 | 32400000 | 32500000 | 0.52 | 0.56 | A>A |
| chr12 | 32500000 | 32600000 | 0.36 | 0.44 | A>A |
| chr12 | 32600000 | 32700000 | 0.49 | 0.60 | A>A |
| chr12 | 32700000 | 32800000 | 0.49 | 0.57 | A>A |
| chr12 | 32800000 | 32900000 | 0.20 | 0.24 | A>A |
| chr12 | 32900000 | 33000000 | -0.09 | -0.23 | B>B |
| chr12 | 33000000 | 33100000 | -0.10 | -0.26 | B>B |
| chr12 | 33100000 | 33200000 | -0.40 | -0.58 | B>B |
| chr12 | 33200000 | 33300000 | -0.51 | -0.83 | B>B |
| chr12 | 33300000 | 33400000 | -0.56 | -0.91 | B>B |
| chr12 | 33400000 | 33500000 | -0.65 | -0.91 | B>B |
| chr12 | 33500000 | 33600000 | -0.59 | -0.88 | B>B |
| chr12 | 33600000 | 33700000 | -0.66 | -0.90 | B>B |
| chr12 | 33700000 | 33800000 | -0.70 | -0.96 | B>B |
| chr12 | 33800000 | 33900000 | -0.78 | -0.97 | B>B |
| chr12 | 33900000 | 34000000 | -0.89 | -1.06 | B>B |
| chr12 | 34000000 | 34100000 | -0.82 | -0.95 | B>B |
| chr12 | 34100000 | 34200000 | 0.00 | 0.00 | NO |
| chr12 | 34200000 | 34300000 | 0.00 | 0.00 | NO |
| chr12 | 34300000 | 34400000 | 0.00 | -0.26 | NO |
| chr12 | 34400000 | 34500000 | -0.27 | -0.01 | B>B |
| chr12 | 34500000 | 34600000 | -0.58 | 0.20 | B>A |
| chr12 | 34600000 | 34700000 | 0.00 | 0.00 | NO |
| chr12 | 34700000 | 34800000 | 0.00 | -0.05 | NO |

| | | | | | |
|---|---|---|---|---|---|
| chr12 | 34800000 | 34900000 | 0.00 | 0.00 | NO |
| chr12 | 34900000 | 35000000 | 0.00 | 0.00 | NO |
| chr12 | 35000000 | 35100000 | 0.00 | 0.00 | NO |
| chr12 | 35100000 | 35200000 | 0.00 | 0.00 | NO |
| chr12 | 35200000 | 35300000 | 0.00 | 0.00 | NO |
| chr12 | 35300000 | 35400000 | 0.00 | 0.00 | NO |
| chr12 | 35400000 | 35500000 | 0.00 | 0.00 | NO |
| chr12 | 35500000 | 35600000 | 0.00 | 0.00 | NO |
| chr12 | 35600000 | 35700000 | 0.00 | 0.00 | NO |
| chr12 | 35700000 | 35800000 | 0.00 | 0.00 | NO |
| chr12 | 35800000 | 35900000 | 0.00 | 0.00 | NO |
| chr12 | 35900000 | 36000000 | 0.00 | 0.00 | NO |
| chr12 | 36000000 | 36100000 | 0.00 | 0.00 | NO |
| chr12 | 36100000 | 36200000 | 0.00 | 0.00 | NO |
| chr12 | 36200000 | 36300000 | 0.00 | 0.00 | NO |
| chr12 | 36300000 | 36400000 | 0.00 | 0.00 | NO |
| chr12 | 36400000 | 36500000 | 0.00 | 0.00 | NO |
| chr12 | 36500000 | 36600000 | 0.00 | 0.00 | NO |
| chr12 | 36600000 | 36700000 | 0.00 | 0.00 | NO |
| chr12 | 36700000 | 36800000 | 0.00 | 0.00 | NO |
| chr12 | 36800000 | 36900000 | 0.00 | 0.00 | NO |
| chr12 | 36900000 | 37000000 | 0.00 | 0.00 | NO |
| chr12 | 37000000 | 37100000 | 0.00 | 0.00 | NO |
| chr12 | 37100000 | 37200000 | 0.00 | 0.00 | NO |
| chr12 | 37200000 | 37300000 | 0.00 | 0.00 | NO |
| chr12 | 37300000 | 37400000 | 0.00 | 0.00 | NO |
| chr12 | 37400000 | 37500000 | 0.00 | 0.00 | NO |
| chr12 | 37500000 | 37600000 | 0.00 | 0.00 | NO |
| chr12 | 37600000 | 37700000 | 0.00 | 0.00 | NO |
| chr12 | 37700000 | 37800000 | 0.00 | 0.00 | NO |
| chr12 | 37800000 | 37900000 | 0.00 | 0.00 | NO |
| chr12 | 37900000 | 38000000 | 0.00 | 0.00 | NO |
| chr12 | 38000000 | 38100000 | 0.00 | 0.00 | NO |
| chr12 | 38100000 | 38200000 | 0.00 | 0.00 | NO |
| chr12 | 38200000 | 38300000 | 0.00 | 0.00 | NO |
| chr12 | 38300000 | 38400000 | 0.00 | -0.10 | NO |
| chr12 | 38400000 | 38500000 | 0.00 | 0.00 | NO |
| chr12 | 38500000 | 38600000 | 0.00 | 0.12 | NO |
| chr12 | 38600000 | 38700000 | 0.00 | 0.00 | NO |
| chr12 | 38700000 | 38800000 | 0.00 | 0.00 | NO |
| chr12 | 38800000 | 38900000 | -0.74 | -0.29 | B>B |
| chr12 | 38900000 | 39000000 | -0.66 | -0.36 | B>B |
| chr12 | 39000000 | 39100000 | -0.41 | -0.27 | B>B |
| chr12 | 39100000 | 39200000 | -0.10 | -0.03 | B>B |
| chr12 | 39200000 | 39300000 | -0.09 | 0.00 | B>B |
| chr12 | 39300000 | 39400000 | -0.49 | -0.32 | B>B |
| chr12 | 39400000 | 39500000 | -0.54 | -0.40 | B>B |
| chr12 | 39500000 | 39600000 | -0.49 | -0.25 | B>B |
| chr12 | 39600000 | 39700000 | -0.39 | -0.23 | B>B |
| chr12 | 39700000 | 39800000 | -0.01 | 0.06 | A>A |
| chr12 | 39800000 | 39900000 | 0.06 | 0.10 | A>A |
| chr12 | 39900000 | 40000000 | -0.22 | -0.08 | B>B |
| chr12 | 40000000 | 40100000 | -0.42 | -0.09 | B>B |
| chr12 | 40100000 | 40200000 | -0.49 | -0.12 | B>B |
| chr12 | 40200000 | 40300000 | -0.29 | -0.03 | B>B |
| chr12 | 40300000 | 40400000 | -0.20 | -0.05 | B>B |

| | | | | | |
|---|---|---|---|---|---|
| chr12 | 40400000 | 40500000 | -0.01 | 0.14 | A>A |
| chr12 | 40500000 | 40600000 | 0.19 | 0.28 | A>A |
| chr12 | 40600000 | 40700000 | 0.47 | 0.48 | A>A |
| chr12 | 40700000 | 40800000 | 0.08 | 0.04 | A>A |
| chr12 | 40800000 | 40900000 | -0.05 | -0.16 | B>B |
| chr12 | 40900000 | 41000000 | -0.04 | -0.13 | B>B |
| chr12 | 41000000 | 41100000 | 0.26 | 0.23 | A>A |
| chr12 | 41100000 | 41200000 | 0.24 | 0.40 | A>A |
| chr12 | 41200000 | 41300000 | 0.12 | 0.35 | A>A |
| chr12 | 41300000 | 41400000 | -0.08 | 0.11 | A>A |
| chr12 | 41400000 | 41500000 | -0.21 | -0.13 | B>B |
| chr12 | 41500000 | 41600000 | -0.37 | -0.36 | B>B |
| chr12 | 41600000 | 41700000 | -0.47 | -0.43 | B>B |
| chr12 | 41700000 | 41800000 | -0.71 | -0.62 | B>B |
| chr12 | 41800000 | 41900000 | -0.66 | -0.75 | B>B |
| chr12 | 41900000 | 42000000 | -0.66 | -0.72 | B>B |
| chr12 | 42000000 | 42100000 | -0.55 | -0.51 | B>B |
| chr12 | 42100000 | 42200000 | -0.46 | -0.49 | B>B |
| chr12 | 42200000 | 42300000 | -0.47 | -0.55 | B>B |
| chr12 | 42300000 | 42400000 | -0.23 | -0.44 | B>B |
| chr12 | 42400000 | 42500000 | -0.12 | -0.23 | B>B |
| chr12 | 42500000 | 42600000 | 0.38 | 0.24 | A>A |
| chr12 | 42600000 | 42700000 | 0.55 | 0.29 | A>A |
| chr12 | 42700000 | 42800000 | 0.26 | 0.22 | A>A |
| chr12 | 42800000 | 42900000 | 0.02 | 0.12 | A>A |
| chr12 | 42900000 | 43000000 | -0.15 | 0.12 | B>B |
| chr12 | 43000000 | 43100000 | -0.31 | 0.25 | B>A |
| chr12 | 43100000 | 43200000 | -0.30 | 0.30 | B>A |
| chr12 | 43200000 | 43300000 | -0.31 | 0.32 | B>A |
| chr12 | 43300000 | 43400000 | -0.34 | 0.33 | B>A |
| chr12 | 43400000 | 43500000 | -0.42 | 0.33 | B>A |
| chr12 | 43500000 | 43600000 | -0.49 | 0.24 | B>A |
| chr12 | 43600000 | 43700000 | -0.43 | 0.15 | B>A |
| chr12 | 43700000 | 43800000 | -0.62 | -0.30 | B>B |
| chr12 | 43800000 | 43900000 | -0.69 | -0.56 | B>B |
| chr12 | 43900000 | 44000000 | -0.83 | -0.83 | B>B |
| chr12 | 44000000 | 44100000 | -0.68 | -0.83 | B>B |
| chr12 | 44100000 | 44200000 | -0.30 | -0.38 | B>B |
| chr12 | 44200000 | 44300000 | -0.14 | -0.33 | B>B |
| chr12 | 44300000 | 44400000 | -0.54 | -0.71 | B>B |
| chr12 | 44400000 | 44500000 | -0.75 | -0.97 | B>B |
| chr12 | 44500000 | 44600000 | -0.97 | -1.16 | B>B |
| chr12 | 44600000 | 44700000 | -0.87 | -1.21 | B>B |
| chr12 | 44700000 | 44800000 | -0.95 | -1.25 | B>B |
| chr12 | 44800000 | 44900000 | -0.84 | -1.24 | B>B |
| chr12 | 44900000 | 45000000 | -0.87 | -1.18 | B>B |
| chr12 | 45000000 | 45100000 | -0.87 | -1.19 | B>B |
| chr12 | 45100000 | 45200000 | -0.80 | -1.19 | B>B |
| chr12 | 45200000 | 45300000 | -0.80 | -1.04 | B>B |
| chr12 | 45300000 | 45400000 | -0.58 | -0.84 | B>B |
| chr12 | 45400000 | 45500000 | -0.64 | -0.92 | B>B |
| chr12 | 45500000 | 45600000 | -0.19 | -0.32 | B>B |
| chr12 | 45600000 | 45700000 | 0.25 | 0.14 | A>A |
| chr12 | 45700000 | 45800000 | 0.14 | 0.00 | A>A |
| chr12 | 45800000 | 45900000 | -0.10 | -0.10 | B>B |
| chr12 | 45900000 | 46000000 | 0.05 | -0.01 | A>A |

| | | | | | |
|---|---|---|---|---|---|
| chr12 | 46000000 | 46100000 | 0.21 | 0.14 | A>A |
| chr12 | 46100000 | 46200000 | 0.35 | 0.31 | A>A |
| chr12 | 46200000 | 46300000 | 0.38 | 0.38 | A>A |
| chr12 | 46300000 | 46400000 | 0.36 | 0.50 | A>A |
| chr12 | 46400000 | 46500000 | 0.26 | 0.32 | A>A |
| chr12 | 46500000 | 46600000 | 0.27 | 0.40 | A>A |
| chr12 | 46600000 | 46700000 | 0.49 | 0.49 | A>A |
| chr12 | 46700000 | 46800000 | 0.62 | 0.57 | A>A |
| chr12 | 46800000 | 46900000 | 0.85 | 0.78 | A>A |
| chr12 | 46900000 | 47000000 | 0.67 | 0.77 | A>A |
| chr12 | 47000000 | 47100000 | 0.44 | 0.50 | A>A |
| chr12 | 47100000 | 47200000 | 0.27 | 0.28 | A>A |
| chr12 | 47200000 | 47300000 | 0.18 | 0.30 | A>A |
| chr12 | 47300000 | 47400000 | 0.24 | 0.34 | A>A |
| chr12 | 47400000 | 47500000 | 0.59 | 0.64 | A>A |
| chr12 | 47500000 | 47600000 | 0.99 | 0.87 | A>A |
| chr12 | 47600000 | 47700000 | 0.94 | 0.87 | A>A |
| chr12 | 47700000 | 47800000 | 0.80 | 0.69 | A>A |
| chr12 | 47800000 | 47900000 | 0.45 | 0.47 | A>A |
| chr12 | 47900000 | 48000000 | 0.57 | 0.44 | A>A |
| chr12 | 48000000 | 48100000 | 0.58 | 0.59 | A>A |
| chr12 | 48100000 | 48200000 | 1.02 | 0.93 | A>A |
| chr12 | 48200000 | 48300000 | 1.02 | 0.95 | A>A |
| chr12 | 48300000 | 48400000 | 1.07 | 0.81 | A>A |
| chr12 | 48400000 | 48500000 | 0.99 | 0.91 | A>A |
| chr12 | 48500000 | 48600000 | 0.79 | 0.73 | A>A |
| chr12 | 48600000 | 48700000 | 0.53 | 0.51 | A>A |
| chr12 | 48700000 | 48800000 | 0.44 | 0.51 | A>A |
| chr12 | 48800000 | 48900000 | 0.43 | 0.45 | A>A |
| chr12 | 48900000 | 49000000 | 0.48 | 0.51 | A>A |
| chr12 | 49000000 | 49100000 | 1.02 | 0.86 | A>A |
| chr12 | 49100000 | 49200000 | 1.27 | 1.19 | A>A |
| chr12 | 49200000 | 49300000 | 1.45 | 1.29 | A>A |
| chr12 | 49300000 | 49400000 | 1.47 | 1.23 | A>A |
| chr12 | 49400000 | 49500000 | 1.55 | 1.24 | A>A |
| chr12 | 49500000 | 49600000 | 1.52 | 1.25 | A>A |
| chr12 | 49600000 | 49700000 | 1.35 | 1.06 | A>A |
| chr12 | 49700000 | 49800000 | 1.47 | 1.13 | A>A |
| chr12 | 49800000 | 49900000 | 1.36 | 1.17 | A>A |
| chr12 | 49900000 | 50000000 | 1.28 | 1.09 | A>A |
| chr12 | 50000000 | 50100000 | 1.30 | 1.04 | A>A |
| chr12 | 50100000 | 50200000 | 1.31 | 1.01 | A>A |
| chr12 | 50200000 | 50300000 | 1.28 | 1.09 | A>A |
| chr12 | 50300000 | 50400000 | 1.31 | 1.05 | A>A |
| chr12 | 50400000 | 50500000 | 1.19 | 0.92 | A>A |
| chr12 | 50500000 | 50600000 | 0.99 | 1.01 | A>A |
| chr12 | 50600000 | 50700000 | 1.20 | 1.11 | A>A |
| chr12 | 50700000 | 50800000 | 1.08 | 0.93 | A>A |
| chr12 | 50800000 | 50900000 | 0.95 | 0.93 | A>A |
| chr12 | 50900000 | 51000000 | 1.13 | 0.98 | A>A |
| chr12 | 51000000 | 51100000 | 1.08 | 0.93 | A>A |
| chr12 | 51100000 | 51200000 | 0.92 | 0.85 | A>A |
| chr12 | 51200000 | 51300000 | 0.90 | 0.88 | A>A |
| chr12 | 51300000 | 51400000 | 0.98 | 0.93 | A>A |
| chr12 | 51400000 | 51500000 | 0.97 | 0.91 | A>A |
| chr12 | 51500000 | 51600000 | 1.25 | 0.94 | A>A |

| | | | | | |
|---|---|---|---|---|---|
| chr12 | 51600000 | 51700000 | 1.20 | 0.92 | A>A |
| chr12 | 51700000 | 51800000 | 0.92 | 0.76 | A>A |
| chr12 | 51800000 | 51900000 | 0.92 | 0.82 | A>A |
| chr12 | 51900000 | 52000000 | 0.90 | 0.79 | A>A |
| chr12 | 52000000 | 52100000 | 1.06 | 0.84 | A>A |
| chr12 | 52100000 | 52200000 | 0.96 | 0.82 | A>A |
| chr12 | 52200000 | 52300000 | 1.00 | 0.76 | A>A |
| chr12 | 52300000 | 52400000 | 1.07 | 0.93 | A>A |
| chr12 | 52400000 | 52500000 | 0.96 | 0.89 | A>A |
| chr12 | 52500000 | 52600000 | 0.74 | 0.88 | A>A |
| chr12 | 52600000 | 52700000 | 1.01 | 1.05 | A>A |
| chr12 | 52700000 | 52800000 | 0.77 | 0.75 | A>A |
| chr12 | 52800000 | 52900000 | 0.84 | 0.79 | A>A |
| chr12 | 52900000 | 53000000 | 0.69 | 0.66 | A>A |
| chr12 | 53000000 | 53100000 | 0.80 | 0.78 | A>A |
| chr12 | 53100000 | 53200000 | 1.00 | 0.83 | A>A |
| chr12 | 53200000 | 53300000 | 0.80 | 0.71 | A>A |
| chr12 | 53300000 | 53400000 | 0.88 | 1.01 | A>A |
| chr12 | 53400000 | 53500000 | 1.38 | 1.13 | A>A |
| chr12 | 53500000 | 53600000 | 1.44 | 1.27 | A>A |
| chr12 | 53600000 | 53700000 | 1.41 | 1.27 | A>A |
| chr12 | 53700000 | 53800000 | 1.57 | 1.30 | A>A |
| chr12 | 53800000 | 53900000 | 1.45 | 1.32 | A>A |
| chr12 | 53900000 | 54000000 | 1.35 | 1.27 | A>A |
| chr12 | 54000000 | 54100000 | 1.41 | 1.20 | A>A |
| chr12 | 54100000 | 54200000 | 1.12 | 0.94 | A>A |
| chr12 | 54200000 | 54300000 | 1.37 | 1.23 | A>A |
| chr12 | 54300000 | 54400000 | 1.09 | 0.97 | A>A |
| chr12 | 54400000 | 54500000 | 1.19 | 1.11 | A>A |
| chr12 | 54500000 | 54600000 | 1.19 | 1.06 | A>A |
| chr12 | 54600000 | 54700000 | 1.18 | 1.04 | A>A |
| chr12 | 54700000 | 54800000 | 1.13 | 0.81 | A>A |
| chr12 | 54800000 | 54900000 | 0.81 | 0.23 | A>A |
| chr12 | 54900000 | 55000000 | 0.45 | -0.27 | A>B |
| chr12 | 55000000 | 55100000 | -0.19 | -0.80 | A>B |
| chr12 | 55100000 | 55200000 | -0.44 | -0.80 | B>B |
| chr12 | 55200000 | 55300000 | -0.47 | -0.85 | B>B |
| chr12 | 55300000 | 55400000 | -0.44 | -0.68 | B>B |
| chr12 | 55400000 | 55500000 | -0.52 | -0.92 | B>B |
| chr12 | 55500000 | 55600000 | -0.61 | -1.02 | B>B |
| chr12 | 55600000 | 55700000 | -0.78 | -1.21 | B>B |
| chr12 | 55700000 | 55800000 | -0.68 | -1.28 | A>B |
| chr12 | 55800000 | 55900000 | -0.70 | -1.25 | B>B |
| chr12 | 55900000 | 56000000 | -0.70 | -1.16 | B>B |
| chr12 | 56000000 | 56100000 | -0.04 | -0.32 | B>B |
| chr12 | 56100000 | 56200000 | 0.78 | 0.61 | A>A |
| chr12 | 56200000 | 56300000 | 1.15 | 0.95 | A>A |
| chr12 | 56300000 | 56400000 | 1.18 | 1.12 | A>A |
| chr12 | 56400000 | 56500000 | 1.30 | 1.21 | A>A |
| chr12 | 56500000 | 56600000 | 1.45 | 1.38 | A>A |
| chr12 | 56600000 | 56700000 | 1.47 | 1.23 | A>A |
| chr12 | 56700000 | 56800000 | 1.55 | 1.47 | A>A |
| chr12 | 56800000 | 56900000 | 1.43 | 1.34 | A>A |
| chr12 | 56900000 | 57000000 | 1.56 | 1.31 | A>A |
| chr12 | 57000000 | 57100000 | 1.50 | 1.39 | A>A |
| chr12 | 57100000 | 57200000 | 1.24 | 1.09 | A>A |

| | | | | | |
|---|---|---|---|---|---|
| chr12 | 57200000 | 57300000 | 1.45 | 1.24 | A>A |
| chr12 | 57300000 | 57400000 | 1.50 | 1.23 | A>A |
| chr12 | 57400000 | 57500000 | 1.39 | 1.27 | A>A |
| chr12 | 57500000 | 57600000 | 1.42 | 1.26 | A>A |
| chr12 | 57600000 | 57700000 | 1.52 | 1.30 | A>A |
| chr12 | 57700000 | 57800000 | 1.65 | 1.48 | A>A |
| chr12 | 57800000 | 57900000 | 1.48 | 1.36 | A>A |
| chr12 | 57900000 | 58000000 | 1.60 | 1.37 | A>A |
| chr12 | 58000000 | 58100000 | 1.60 | 1.42 | A>A |
| chr12 | 58100000 | 58200000 | 1.33 | 1.29 | A>A |
| chr12 | 58200000 | 58300000 | 0.86 | 1.02 | A>A |
| chr12 | 58300000 | 58400000 | 0.05 | 0.44 | A>A |
| chr12 | 58400000 | 58500000 | -0.41 | 0.23 | B>A |
| chr12 | 58500000 | 58600000 | -0.31 | 0.30 | B>A |
| chr12 | 58600000 | 58700000 | -0.26 | 0.34 | B>A |
| chr12 | 58700000 | 58800000 | -0.11 | 0.23 | A>A |
| chr12 | 58800000 | 58900000 | -0.02 | 0.40 | A>A |
| chr12 | 58900000 | 59000000 | -0.20 | 0.25 | A>A |
| chr12 | 59000000 | 59100000 | -0.12 | 0.32 | A>A |
| chr12 | 59100000 | 59200000 | -0.10 | 0.26 | A>A |
| chr12 | 59200000 | 59300000 | -0.08 | 0.25 | A>A |
| chr12 | 59300000 | 59400000 | -0.09 | 0.36 | A>A |
| chr12 | 59400000 | 59500000 | -0.18 | 0.25 | A>A |
| chr12 | 59500000 | 59600000 | -0.45 | -0.07 | B>B |
| chr12 | 59600000 | 59700000 | -0.85 | -0.23 | B>A |
| chr12 | 59700000 | 59800000 | -0.87 | -0.24 | B>A |
| chr12 | 59800000 | 59900000 | -0.85 | -0.32 | B>A |
| chr12 | 59900000 | 60000000 | -0.81 | -0.17 | B>A |
| chr12 | 60000000 | 60100000 | -0.83 | -0.19 | B>A |
| chr12 | 60100000 | 60200000 | -0.81 | -0.20 | B>A |
| chr12 | 60200000 | 60300000 | -0.73 | -0.28 | B>B |
| chr12 | 60300000 | 60400000 | -0.76 | -0.33 | B>B |
| chr12 | 60400000 | 60500000 | -0.67 | -0.36 | B>B |
| chr12 | 60500000 | 60600000 | -0.45 | -0.25 | B>B |
| chr12 | 60600000 | 60700000 | -0.40 | -0.27 | B>B |
| chr12 | 60700000 | 60800000 | -0.45 | -0.31 | B>B |
| chr12 | 60800000 | 60900000 | -0.61 | -0.38 | B>B |
| chr12 | 60900000 | 61000000 | -0.71 | -0.68 | B>B |
| chr12 | 61000000 | 61100000 | -0.80 | -0.93 | B>B |
| chr12 | 61100000 | 61200000 | -0.82 | -0.96 | B>B |
| chr12 | 61200000 | 61300000 | -0.80 | -0.96 | B>B |
| chr12 | 61300000 | 61400000 | -0.84 | -0.93 | B>B |
| chr12 | 61400000 | 61500000 | -0.84 | -0.95 | B>B |
| chr12 | 61500000 | 61600000 | -0.85 | -0.96 | B>B |
| chr12 | 61600000 | 61700000 | -0.83 | -0.95 | B>B |
| chr12 | 61700000 | 61800000 | -0.85 | -1.02 | B>B |
| chr12 | 61800000 | 61900000 | -0.90 | -1.05 | B>B |
| chr12 | 61900000 | 62000000 | -0.90 | -1.07 | B>B |
| chr12 | 62000000 | 62100000 | -0.91 | -1.10 | B>B |
| chr12 | 62100000 | 62200000 | -0.91 | -1.07 | B>B |
| chr12 | 62200000 | 62300000 | -0.88 | -1.12 | B>B |
| chr12 | 62300000 | 62400000 | -0.85 | -0.99 | B>B |
| chr12 | 62400000 | 62500000 | -0.79 | -1.00 | B>B |
| chr12 | 62500000 | 62600000 | -0.63 | -0.70 | B>B |
| chr12 | 62600000 | 62700000 | -0.17 | -0.13 | B>B |
| chr12 | 62700000 | 62800000 | 0.11 | 0.26 | A>A |

| | | | | | |
|---|---|---|---|---|---|
| chr12 | 62800000 | 62900000 | 0.37 | 0.46 | A>A |
| chr12 | 62900000 | 63000000 | 0.60 | 0.57 | A>A |
| chr12 | 63000000 | 63100000 | 0.85 | 0.77 | A>A |
| chr12 | 63100000 | 63200000 | 0.80 | 0.84 | A>A |
| chr12 | 63200000 | 63300000 | 0.25 | 0.48 | A>A |
| chr12 | 63300000 | 63400000 | 0.20 | 0.49 | A>A |
| chr12 | 63400000 | 63500000 | -0.03 | 0.39 | A>A |
| chr12 | 63500000 | 63600000 | -0.32 | 0.60 | B>A |
| chr12 | 63600000 | 63700000 | -0.16 | 0.63 | B>A |
| chr12 | 63700000 | 63800000 | -0.19 | 0.59 | B>A |
| chr12 | 63800000 | 63900000 | -0.25 | 0.55 | B>A |
| chr12 | 63900000 | 64000000 | 0.00 | 0.00 | NO |
| chr12 | 64000000 | 64100000 | 0.00 | 0.00 | NO |
| chr12 | 64100000 | 64200000 | 0.12 | 0.60 | A>A |
| chr12 | 64200000 | 64300000 | 0.50 | 0.83 | A>A |
| chr12 | 64300000 | 64400000 | 0.61 | 0.90 | A>A |
| chr12 | 64400000 | 64500000 | 0.63 | 0.88 | A>A |
| chr12 | 64500000 | 64600000 | 0.59 | 0.72 | A>A |
| chr12 | 64600000 | 64700000 | 0.41 | 0.72 | A>A |
| chr12 | 64700000 | 64800000 | 0.62 | 0.85 | A>A |
| chr12 | 64800000 | 64900000 | 0.98 | 0.97 | A>A |
| chr12 | 64900000 | 65000000 | 0.98 | 0.98 | A>A |
| chr12 | 65000000 | 65100000 | 1.13 | 1.08 | A>A |
| chr12 | 65100000 | 65200000 | 0.62 | 0.63 | A>A |
| chr12 | 65200000 | 65300000 | 0.03 | 0.20 | A>A |
| chr12 | 65300000 | 65400000 | -0.18 | 0.03 | B>B |
| chr12 | 65400000 | 65500000 | -0.22 | 0.05 | B>B |
| chr12 | 65500000 | 65600000 | 0.14 | 0.23 | A>A |
| chr12 | 65600000 | 65700000 | 0.19 | 0.30 | A>A |
| chr12 | 65700000 | 65800000 | 0.12 | 0.32 | A>A |
| chr12 | 65800000 | 65900000 | 0.15 | 0.43 | A>A |
| chr12 | 65900000 | 66000000 | 0.17 | 0.33 | A>A |
| chr12 | 66000000 | 66100000 | 0.43 | 0.47 | A>A |
| chr12 | 66100000 | 66200000 | 0.50 | 0.45 | A>A |
| chr12 | 66200000 | 66300000 | 0.51 | 0.52 | A>A |
| chr12 | 66300000 | 66400000 | 0.44 | 0.46 | A>A |
| chr12 | 66400000 | 66500000 | 0.19 | 0.28 | A>A |
| chr12 | 66500000 | 66600000 | 0.08 | 0.01 | A>A |
| chr12 | 66600000 | 66700000 | -0.10 | -0.24 | B>B |
| chr12 | 66700000 | 66800000 | -0.24 | -0.37 | B>B |
| chr12 | 66800000 | 66900000 | -0.56 | -0.53 | B>B |
| chr12 | 66900000 | 67000000 | -0.65 | -0.51 | B>B |
| chr12 | 67000000 | 67100000 | -0.56 | -0.50 | B>B |
| chr12 | 67100000 | 67200000 | -0.62 | -0.57 | B>B |
| chr12 | 67200000 | 67300000 | -0.59 | -0.51 | B>B |
| chr12 | 67300000 | 67400000 | -0.46 | -0.42 | B>B |
| chr12 | 67400000 | 67500000 | -0.35 | -0.34 | B>B |
| chr12 | 67500000 | 67600000 | -0.56 | -0.56 | B>B |
| chr12 | 67600000 | 67700000 | -0.30 | -0.33 | B>B |
| chr12 | 67700000 | 67800000 | -0.36 | -0.26 | B>B |
| chr12 | 67800000 | 67900000 | -0.53 | -0.35 | B>B |
| chr12 | 67900000 | 68000000 | -0.57 | -0.37 | B>B |
| chr12 | 68000000 | 68100000 | -0.36 | -0.16 | B>B |
| chr12 | 68100000 | 68200000 | -0.46 | -0.23 | B>B |
| chr12 | 68200000 | 68300000 | -0.62 | -0.27 | B>B |
| chr12 | 68300000 | 68400000 | -0.65 | -0.44 | B>B |

| | | | | | |
|---|---|---|---|---|---|
| chr12 | 68400000 | 68500000 | -0.74 | -0.72 | B>B |
| chr12 | 68500000 | 68600000 | -0.74 | -0.82 | B>B |
| chr12 | 68600000 | 68700000 | -0.36 | -0.57 | B>B |
| chr12 | 68700000 | 68800000 | 0.05 | -0.19 | B>B |
| chr12 | 68800000 | 68900000 | 0.05 | -0.20 | B>B |
| chr12 | 68900000 | 69000000 | 0.23 | -0.15 | A>A |
| chr12 | 69000000 | 69100000 | 0.44 | 0.22 | A>A |
| chr12 | 69100000 | 69200000 | 0.68 | 0.53 | A>A |
| chr12 | 69200000 | 69300000 | 0.67 | 0.61 | A>A |
| chr12 | 69300000 | 69400000 | 0.86 | 0.81 | A>A |
| chr12 | 69400000 | 69500000 | 0.81 | 0.77 | A>A |
| chr12 | 69500000 | 69600000 | 0.67 | 0.67 | A>A |
| chr12 | 69600000 | 69700000 | 0.74 | 0.73 | A>A |
| chr12 | 69700000 | 69800000 | 0.66 | 0.64 | A>A |
| chr12 | 69800000 | 69900000 | 0.77 | 0.70 | A>A |
| chr12 | 69900000 | 70000000 | 0.70 | 0.65 | A>A |
| chr12 | 70000000 | 70100000 | 0.43 | 0.37 | A>A |
| chr12 | 70100000 | 70200000 | 0.04 | 0.18 | A>A |
| chr12 | 70200000 | 70300000 | -0.20 | -0.06 | B>B |
| chr12 | 70300000 | 70400000 | -0.32 | -0.06 | B>B |
| chr12 | 70400000 | 70500000 | -0.38 | -0.09 | B>B |
| chr12 | 70500000 | 70600000 | -0.44 | -0.11 | B>B |
| chr12 | 70600000 | 70700000 | -0.12 | 0.11 | B>B |
| chr12 | 70700000 | 70800000 | -0.06 | 0.06 | A>A |
| chr12 | 70800000 | 70900000 | -0.34 | -0.06 | B>B |
| chr12 | 70900000 | 71000000 | -0.29 | -0.09 | B>B |
| chr12 | 71000000 | 71100000 | -0.28 | -0.05 | B>B |
| chr12 | 71100000 | 71200000 | -0.56 | -0.36 | B>B |
| chr12 | 71200000 | 71300000 | -0.81 | -0.73 | B>B |
| chr12 | 71300000 | 71400000 | -0.87 | -0.96 | B>B |
| chr12 | 71400000 | 71500000 | -0.88 | -1.00 | B>B |
| chr12 | 71500000 | 71600000 | -0.94 | -0.94 | B>B |
| chr12 | 71600000 | 71700000 | -1.01 | -1.05 | B>B |
| chr12 | 71700000 | 71800000 | -0.84 | -0.96 | B>B |
| chr12 | 71800000 | 71900000 | -0.62 | -0.92 | B>B |
| chr12 | 71900000 | 72000000 | -0.31 | -0.76 | B>B |
| chr12 | 72000000 | 72100000 | 0.13 | -0.20 | B>B |
| chr12 | 72100000 | 72200000 | -0.06 | -0.28 | B>B |
| chr12 | 72200000 | 72300000 | -0.07 | -0.38 | B>B |
| chr12 | 72300000 | 72400000 | -0.54 | -0.90 | B>B |
| chr12 | 72400000 | 72500000 | -0.87 | -1.06 | B>B |
| chr12 | 72500000 | 72600000 | -0.93 | -1.19 | B>B |
| chr12 | 72600000 | 72700000 | -0.87 | -1.02 | B>B |
| chr12 | 72700000 | 72800000 | -0.81 | -0.98 | B>B |
| chr12 | 72800000 | 72900000 | -0.90 | -1.01 | B>B |
| chr12 | 72900000 | 73000000 | -0.90 | -1.00 | B>B |
| chr12 | 73000000 | 73100000 | -0.87 | -0.97 | B>B |
| chr12 | 73100000 | 73200000 | -0.86 | -0.97 | B>B |
| chr12 | 73200000 | 73300000 | -0.87 | -1.03 | B>B |
| chr12 | 73300000 | 73400000 | -0.85 | -0.98 | B>B |
| chr12 | 73400000 | 73500000 | -0.83 | -0.90 | B>B |
| chr12 | 73500000 | 73600000 | -0.85 | -0.89 | B>B |
| chr12 | 73600000 | 73700000 | -0.79 | -0.85 | B>B |
| chr12 | 73700000 | 73800000 | -0.80 | -0.73 | B>B |
| chr12 | 73800000 | 73900000 | -0.79 | -0.73 | B>B |
| chr12 | 73900000 | 74000000 | -0.73 | -0.64 | B>B |

| | | | | | |
|---|---|---|---|---|---|
| chr12 | 74000000 | 74100000 | -0.62 | -0.56 | B>B |
| chr12 | 74100000 | 74200000 | -0.51 | -0.44 | B>B |
| chr12 | 74200000 | 74300000 | -0.37 | -0.37 | B>B |
| chr12 | 74300000 | 74400000 | -0.25 | -0.26 | B>B |
| chr12 | 74400000 | 74500000 | -0.14 | -0.21 | B>B |
| chr12 | 74500000 | 74600000 | -0.16 | -0.27 | B>B |
| chr12 | 74600000 | 74700000 | -0.26 | -0.26 | B>B |
| chr12 | 74700000 | 74800000 | -0.27 | -0.14 | B>B |
| chr12 | 74800000 | 74900000 | -0.34 | -0.22 | B>B |
| chr12 | 74900000 | 75000000 | -0.44 | -0.13 | B>B |
| chr12 | 75000000 | 75100000 | -0.55 | -0.07 | B>B |
| chr12 | 75100000 | 75200000 | -0.35 | 0.37 | B>A |
| chr12 | 75200000 | 75300000 | -0.08 | 0.71 | B>A |
| chr12 | 75300000 | 75400000 | -0.14 | 0.70 | B>A |
| chr12 | 75400000 | 75500000 | -0.29 | 0.50 | B>A |
| chr12 | 75500000 | 75600000 | -0.37 | 0.44 | B>A |
| chr12 | 75600000 | 75700000 | -0.34 | 0.34 | B>A |
| chr12 | 75700000 | 75800000 | -0.10 | 0.34 | A>A |
| chr12 | 75800000 | 75900000 | 0.03 | 0.37 | A>A |
| chr12 | 75900000 | 76000000 | 0.53 | 0.48 | A>A |
| chr12 | 76000000 | 76100000 | 0.99 | 0.77 | A>A |
| chr12 | 76100000 | 76200000 | 0.98 | 0.83 | A>A |
| chr12 | 76200000 | 76300000 | 1.06 | 0.81 | A>A |
| chr12 | 76300000 | 76400000 | 1.02 | 0.87 | A>A |
| chr12 | 76400000 | 76500000 | 0.52 | 0.52 | A>A |
| chr12 | 76500000 | 76600000 | 0.54 | 0.32 | A>A |
| chr12 | 76600000 | 76700000 | 0.41 | 0.27 | A>A |
| chr12 | 76700000 | 76800000 | 0.44 | 0.28 | A>A |
| chr12 | 76800000 | 76900000 | 0.24 | 0.15 | A>A |
| chr12 | 76900000 | 77000000 | 0.20 | -0.02 | A>A |
| chr12 | 77000000 | 77100000 | -0.02 | -0.37 | B>B |
| chr12 | 77100000 | 77200000 | 0.10 | -0.28 | B>B |
| chr12 | 77200000 | 77300000 | 0.00 | -0.44 | B>B |
| chr12 | 77300000 | 77400000 | -0.45 | -0.87 | B>B |
| chr12 | 77400000 | 77500000 | -0.49 | -0.75 | B>B |
| chr12 | 77500000 | 77600000 | -0.88 | -1.15 | B>B |
| chr12 | 77600000 | 77700000 | -0.90 | -1.24 | B>B |
| chr12 | 77700000 | 77800000 | -0.79 | -1.17 | B>B |
| chr12 | 77800000 | 77900000 | -0.97 | -1.26 | B>B |
| chr12 | 77900000 | 78000000 | -0.89 | -1.18 | B>B |
| chr12 | 78000000 | 78100000 | -0.72 | -0.96 | B>B |
| chr12 | 78100000 | 78200000 | -0.44 | -0.36 | B>B |
| chr12 | 78200000 | 78300000 | -0.25 | -0.23 | B>B |
| chr12 | 78300000 | 78400000 | -0.20 | -0.24 | B>B |
| chr12 | 78400000 | 78500000 | -0.19 | -0.20 | B>B |
| chr12 | 78500000 | 78600000 | -0.17 | -0.09 | B>B |
| chr12 | 78600000 | 78700000 | -0.16 | 0.05 | B>B |
| chr12 | 78700000 | 78800000 | -0.15 | 0.10 | B>B |
| chr12 | 78800000 | 78900000 | -0.15 | 0.01 | B>B |
| chr12 | 78900000 | 79000000 | -0.14 | -0.03 | B>B |
| chr12 | 79000000 | 79100000 | 0.04 | 0.09 | A>A |
| chr12 | 79100000 | 79200000 | 0.10 | 0.14 | A>A |
| chr12 | 79200000 | 79300000 | 0.14 | 0.24 | A>A |
| chr12 | 79300000 | 79400000 | 0.05 | 0.19 | A>A |
| chr12 | 79400000 | 79500000 | 0.11 | 0.19 | A>A |
| chr12 | 79500000 | 79600000 | 0.14 | 0.14 | A>A |

| | | | | | |
|---|---|---|---|---|---|
| chr12 | 79600000 | 79700000 | 0.22 | 0.17 | A>A |
| chr12 | 79700000 | 79800000 | 0.05 | 0.03 | A>A |
| chr12 | 79800000 | 79900000 | 0.13 | 0.01 | A>A |
| chr12 | 79900000 | 80000000 | 0.20 | 0.13 | A>A |
| chr12 | 80000000 | 80100000 | 0.46 | 0.31 | A>A |
| chr12 | 80100000 | 80200000 | 0.20 | 0.05 | A>A |
| chr12 | 80200000 | 80300000 | 0.28 | -0.06 | A>A |
| chr12 | 80300000 | 80400000 | 0.02 | -0.39 | B>B |
| chr12 | 80400000 | 80500000 | -0.25 | -0.71 | B>B |
| chr12 | 80500000 | 80600000 | -0.32 | -0.68 | B>B |
| chr12 | 80600000 | 80700000 | -0.49 | -0.73 | B>B |
| chr12 | 80700000 | 80800000 | -0.68 | -0.97 | B>B |
| chr12 | 80800000 | 80900000 | -0.73 | -0.75 | B>B |
| chr12 | 80900000 | 81000000 | -0.73 | -0.65 | B>B |
| chr12 | 81000000 | 81100000 | -0.85 | -0.91 | B>B |
| chr12 | 81100000 | 81200000 | -0.80 | -0.95 | B>B |
| chr12 | 81200000 | 81300000 | -0.88 | -1.10 | B>B |
| chr12 | 81300000 | 81400000 | -0.95 | -1.12 | B>B |
| chr12 | 81400000 | 81500000 | -1.01 | -1.14 | B>B |
| chr12 | 81500000 | 81600000 | -0.93 | -1.09 | B>B |
| chr12 | 81600000 | 81700000 | -0.89 | -1.06 | B>B |
| chr12 | 81700000 | 81800000 | -0.99 | -1.13 | B>B |
| chr12 | 81800000 | 81900000 | -1.02 | -1.11 | B>B |
| chr12 | 81900000 | 82000000 | -0.94 | -1.14 | B>B |
| chr12 | 82000000 | 82100000 | -0.85 | -1.05 | B>B |
| chr12 | 82100000 | 82200000 | -0.96 | -1.20 | B>B |
| chr12 | 82200000 | 82300000 | -0.99 | -1.27 | B>B |
| chr12 | 82300000 | 82400000 | -0.99 | -1.30 | B>B |
| chr12 | 82400000 | 82500000 | -1.09 | -1.24 | B>B |
| chr12 | 82500000 | 82600000 | -1.02 | -1.23 | B>B |
| chr12 | 82600000 | 82700000 | -0.99 | -1.22 | B>B |
| chr12 | 82700000 | 82800000 | -0.57 | -0.68 | B>B |
| chr12 | 82800000 | 82900000 | -0.46 | -0.42 | B>B |
| chr12 | 82900000 | 83000000 | -0.39 | -0.40 | B>B |
| chr12 | 83000000 | 83100000 | -0.28 | -0.23 | B>B |
| chr12 | 83100000 | 83200000 | -0.18 | -0.11 | B>B |
| chr12 | 83200000 | 83300000 | -0.18 | -0.13 | B>B |
| chr12 | 83300000 | 83400000 | -0.33 | -0.21 | B>B |
| chr12 | 83400000 | 83500000 | -0.45 | -0.28 | B>B |
| chr12 | 83500000 | 83600000 | -0.60 | -0.34 | B>B |
| chr12 | 83600000 | 83700000 | -0.74 | -0.36 | B>B |
| chr12 | 83700000 | 83800000 | -0.76 | -0.29 | B>B |
| chr12 | 83800000 | 83900000 | -0.76 | -0.15 | B>A |
| chr12 | 83900000 | 84000000 | -0.75 | -0.15 | B>A |
| chr12 | 84000000 | 84100000 | -0.74 | -0.07 | B>A |
| chr12 | 84100000 | 84200000 | -0.62 | -0.07 | B>A |
| chr12 | 84200000 | 84300000 | -0.61 | -0.14 | B>B |
| chr12 | 84300000 | 84400000 | -0.63 | -0.13 | B>B |
| chr12 | 84400000 | 84500000 | -0.77 | -0.28 | B>B |
| chr12 | 84500000 | 84600000 | -0.85 | -0.23 | B>A |
| chr12 | 84600000 | 84700000 | -0.83 | -0.29 | B>A |
| chr12 | 84700000 | 84800000 | -0.85 | -0.32 | B>A |
| chr12 | 84800000 | 84900000 | -0.91 | -0.56 | B>B |
| chr12 | 84900000 | 85000000 | -0.96 | -0.88 | B>B |
| chr12 | 85000000 | 85100000 | -1.05 | -1.06 | B>B |
| chr12 | 85100000 | 85200000 | -1.01 | -1.12 | B>B |

| | | | | | |
|---|---|---|---|---|---|
| chr12 | 85200000 | 85300000 | -0.80 | -0.72 | B>B |
| chr12 | 85300000 | 85400000 | -0.92 | -0.84 | B>B |
| chr12 | 85400000 | 85500000 | -0.67 | -0.41 | B>B |
| chr12 | 85500000 | 85600000 | -0.69 | -0.04 | B>A |
| chr12 | 85600000 | 85700000 | -0.59 | 0.08 | B>A |
| chr12 | 85700000 | 85800000 | -0.63 | 0.01 | B>A |
| chr12 | 85800000 | 85900000 | -0.41 | 0.16 | B>A |
| chr12 | 85900000 | 86000000 | -0.36 | 0.35 | B>A |
| chr12 | 86000000 | 86100000 | -0.29 | 0.42 | B>A |
| chr12 | 86100000 | 86200000 | -0.20 | 0.43 | B>A |
| chr12 | 86200000 | 86300000 | -0.18 | 0.35 | B>A |
| chr12 | 86300000 | 86400000 | -0.24 | 0.26 | A>A |
| chr12 | 86400000 | 86500000 | -0.26 | 0.38 | B>A |
| chr12 | 86500000 | 86600000 | -0.13 | 0.42 | B>A |
| chr12 | 86600000 | 86700000 | -0.09 | 0.44 | B>A |
| chr12 | 86700000 | 86800000 | -0.02 | 0.58 | B>A |
| chr12 | 86800000 | 86900000 | -0.02 | 0.54 | B>A |
| chr12 | 86900000 | 87000000 | -0.04 | 0.56 | B>A |
| chr12 | 87000000 | 87100000 | 0.04 | 0.53 | A>A |
| chr12 | 87100000 | 87200000 | -0.02 | 0.52 | B>A |
| chr12 | 87200000 | 87300000 | -0.01 | 0.49 | A>A |
| chr12 | 87300000 | 87400000 | -0.16 | 0.55 | B>A |
| chr12 | 87400000 | 87500000 | -0.20 | 0.68 | B>A |
| chr12 | 87500000 | 87600000 | -0.20 | 0.68 | B>A |
| chr12 | 87600000 | 87700000 | -0.10 | 0.74 | B>A |
| chr12 | 87700000 | 87800000 | -0.05 | 0.73 | B>A |
| chr12 | 87800000 | 87900000 | -0.06 | 0.75 | B>A |
| chr12 | 87900000 | 88000000 | -0.04 | 0.90 | B>A |
| chr12 | 88000000 | 88100000 | -0.05 | 0.76 | B>A |
| chr12 | 88100000 | 88200000 | 0.03 | 0.78 | B>A |
| chr12 | 88200000 | 88300000 | -0.09 | 0.60 | B>A |
| chr12 | 88300000 | 88400000 | -0.26 | 0.44 | B>A |
| chr12 | 88400000 | 88500000 | 0.08 | 0.55 | A>A |
| chr12 | 88500000 | 88600000 | 0.30 | 0.55 | A>A |
| chr12 | 88600000 | 88700000 | 0.38 | 0.70 | A>A |
| chr12 | 88700000 | 88800000 | 0.28 | 0.60 | A>A |
| chr12 | 88800000 | 88900000 | 0.25 | 0.58 | A>A |
| chr12 | 88900000 | 89000000 | 0.23 | 0.62 | A>A |
| chr12 | 89000000 | 89100000 | 0.19 | 0.64 | A>A |
| chr12 | 89100000 | 89200000 | 0.20 | 0.69 | A>A |
| chr12 | 89200000 | 89300000 | 0.21 | 0.72 | A>A |
| chr12 | 89300000 | 89400000 | 0.13 | 0.53 | A>A |
| chr12 | 89400000 | 89500000 | 0.14 | 0.37 | A>A |
| chr12 | 89500000 | 89600000 | 0.34 | 0.54 | A>A |
| chr12 | 89600000 | 89700000 | 0.43 | 0.58 | A>A |
| chr12 | 89700000 | 89800000 | 0.36 | 0.53 | A>A |
| chr12 | 89800000 | 89900000 | 0.25 | 0.39 | A>A |
| chr12 | 89900000 | 90000000 | 0.16 | 0.24 | A>A |
| chr12 | 90000000 | 90100000 | 0.28 | 0.36 | A>A |
| chr12 | 90100000 | 90200000 | 0.36 | 0.43 | A>A |
| chr12 | 90200000 | 90300000 | 0.45 | 0.45 | A>A |
| chr12 | 90300000 | 90400000 | 0.30 | 0.39 | A>A |
| chr12 | 90400000 | 90500000 | 0.29 | 0.32 | A>A |
| chr12 | 90500000 | 90600000 | 0.14 | 0.27 | A>A |
| chr12 | 90600000 | 90700000 | 0.01 | 0.24 | A>A |
| chr12 | 90700000 | 90800000 | -0.06 | 0.15 | A>A |

| | | | | | |
|---|---|---|---|---|---|
| chr12 | 90800000 | 90900000 | -0.18 | 0.07 | B>B |
| chr12 | 90900000 | 91000000 | -0.07 | 0.08 | A>A |
| chr12 | 91000000 | 91100000 | -0.18 | -0.08 | B>B |
| chr12 | 91100000 | 91200000 | -0.19 | -0.08 | B>B |
| chr12 | 91200000 | 91300000 | -0.18 | -0.03 | B>B |
| chr12 | 91300000 | 91400000 | -0.39 | -0.26 | B>B |
| chr12 | 91400000 | 91500000 | -0.87 | -0.79 | B>B |
| chr12 | 91500000 | 91600000 | -0.91 | -0.89 | B>B |
| chr12 | 91600000 | 91700000 | -0.92 | -0.81 | B>B |
| chr12 | 91700000 | 91800000 | -0.43 | -0.29 | B>B |
| chr12 | 91800000 | 91900000 | -0.35 | -0.04 | B>B |
| chr12 | 91900000 | 92000000 | -0.23 | 0.06 | B>B |
| chr12 | 92000000 | 92100000 | -0.17 | 0.13 | B>B |
| chr12 | 92100000 | 92200000 | -0.12 | 0.13 | A>A |
| chr12 | 92200000 | 92300000 | 0.12 | 0.33 | A>A |
| chr12 | 92300000 | 92400000 | 0.17 | 0.36 | A>A |
| chr12 | 92400000 | 92500000 | 0.21 | 0.34 | A>A |
| chr12 | 92500000 | 92600000 | 0.14 | 0.15 | A>A |
| chr12 | 92600000 | 92700000 | -0.08 | 0.05 | B>B |
| chr12 | 92700000 | 92800000 | -0.18 | -0.05 | B>B |
| chr12 | 92800000 | 92900000 | -0.27 | -0.05 | B>B |
| chr12 | 92900000 | 93000000 | -0.18 | -0.02 | B>B |
| chr12 | 93000000 | 93100000 | -0.26 | 0.00 | B>B |
| chr12 | 93100000 | 93200000 | -0.13 | 0.14 | A>A |
| chr12 | 93200000 | 93300000 | 0.28 | 0.33 | A>A |
| chr12 | 93300000 | 93400000 | 0.23 | 0.37 | A>A |
| chr12 | 93400000 | 93500000 | 0.07 | 0.52 | A>A |
| chr12 | 93500000 | 93600000 | 0.09 | 0.56 | A>A |
| chr12 | 93600000 | 93700000 | 0.14 | 0.47 | A>A |
| chr12 | 93700000 | 93800000 | 0.25 | 0.44 | A>A |
| chr12 | 93800000 | 93900000 | 0.47 | 0.41 | A>A |
| chr12 | 93900000 | 94000000 | 0.51 | 0.39 | A>A |
| chr12 | 94000000 | 94100000 | 0.68 | 0.34 | A>A |
| chr12 | 94100000 | 94200000 | 0.49 | 0.30 | A>A |
| chr12 | 94200000 | 94300000 | 0.36 | 0.19 | A>A |
| chr12 | 94300000 | 94400000 | 0.25 | 0.06 | A>A |
| chr12 | 94400000 | 94500000 | 0.24 | 0.05 | A>A |
| chr12 | 94500000 | 94600000 | 0.25 | 0.01 | A>A |
| chr12 | 94600000 | 94700000 | 0.29 | 0.01 | A>A |
| chr12 | 94700000 | 94800000 | 0.31 | 0.11 | A>A |
| chr12 | 94800000 | 94900000 | 0.26 | 0.14 | A>A |
| chr12 | 94900000 | 95000000 | 0.34 | 0.17 | A>A |
| chr12 | 95000000 | 95100000 | 0.59 | 0.33 | A>A |
| chr12 | 95100000 | 95200000 | 0.49 | 0.13 | A>A |
| chr12 | 95200000 | 95300000 | 0.51 | 0.10 | A>A |
| chr12 | 95300000 | 95400000 | 0.65 | 0.27 | A>A |
| chr12 | 95400000 | 95500000 | 0.86 | 0.61 | A>A |
| chr12 | 95500000 | 95600000 | 1.12 | 0.98 | A>A |
| chr12 | 95600000 | 95700000 | 1.13 | 0.86 | A>A |
| chr12 | 95700000 | 95800000 | 0.82 | 0.53 | A>A |
| chr12 | 95800000 | 95900000 | 1.02 | 0.60 | A>A |
| chr12 | 95900000 | 96000000 | 0.70 | 0.55 | A>A |
| chr12 | 96000000 | 96100000 | 1.02 | 0.81 | A>A |
| chr12 | 96100000 | 96200000 | 1.01 | 0.83 | A>A |
| chr12 | 96200000 | 96300000 | 0.83 | 0.65 | A>A |
| chr12 | 96300000 | 96400000 | 0.72 | 0.52 | A>A |

| | | | | | |
|---|---|---|---|---|---|
| chr12 | 96400000 | 96500000 | 0.57 | 0.35 | A>A |
| chr12 | 96500000 | 96600000 | 0.56 | 0.37 | A>A |
| chr12 | 96600000 | 96700000 | 0.44 | 0.31 | A>A |
| chr12 | 96700000 | 96800000 | 0.51 | 0.32 | A>A |
| chr12 | 96800000 | 96900000 | 0.31 | 0.13 | A>A |
| chr12 | 96900000 | 97000000 | -0.31 | -0.35 | B>B |
| chr12 | 97000000 | 97100000 | -0.66 | -0.80 | B>B |
| chr12 | 97100000 | 97200000 | -0.56 | -0.88 | B>B |
| chr12 | 97200000 | 97300000 | -0.49 | -0.84 | B>B |
| chr12 | 97300000 | 97400000 | -0.44 | -0.64 | B>B |
| chr12 | 97400000 | 97500000 | -0.87 | -1.12 | B>B |
| chr12 | 97500000 | 97600000 | -0.91 | -1.20 | B>B |
| chr12 | 97600000 | 97700000 | -0.95 | -1.03 | B>B |
| chr12 | 97700000 | 97800000 | -0.81 | -0.68 | B>B |
| chr12 | 97800000 | 97900000 | -0.83 | -0.35 | B>B |
| chr12 | 97900000 | 98000000 | -0.86 | -0.17 | B>A |
| chr12 | 98000000 | 98100000 | -0.86 | -0.25 | B>A |
| chr12 | 98100000 | 98200000 | -0.95 | -0.37 | B>A |
| chr12 | 98200000 | 98300000 | -1.00 | -0.45 | B>A |
| chr12 | 98300000 | 98400000 | -0.89 | -0.56 | B>B |
| chr12 | 98400000 | 98500000 | -0.89 | -1.01 | B>B |
| chr12 | 98500000 | 98600000 | -0.90 | -1.20 | B>B |
| chr12 | 98600000 | 98700000 | -0.75 | -1.08 | B>B |
| chr12 | 98700000 | 98800000 | -0.48 | -0.72 | B>B |
| chr12 | 98800000 | 98900000 | 0.11 | -0.07 | A>A |
| chr12 | 98900000 | 99000000 | 0.42 | 0.25 | A>A |
| chr12 | 99000000 | 99100000 | 0.19 | 0.06 | A>A |
| chr12 | 99100000 | 99200000 | 0.07 | -0.30 | B>B |
| chr12 | 99200000 | 99300000 | -0.17 | -0.51 | B>B |
| chr12 | 99300000 | 99400000 | -0.28 | -0.42 | B>B |
| chr12 | 99400000 | 99500000 | -0.34 | -0.55 | B>B |
| chr12 | 99500000 | 99600000 | -0.60 | -0.66 | B>B |
| chr12 | 99600000 | 99700000 | -0.50 | -0.75 | B>B |
| chr12 | 99700000 | 99800000 | -0.34 | -0.72 | B>B |
| chr12 | 99800000 | 99900000 | -0.16 | -0.60 | B>B |
| chr12 | 99900000 | 100000000 | -0.21 | -0.59 | B>B |
| chr12 | 1E+08 | 100100000 | -0.32 | -0.54 | B>B |
| chr12 | 1E+08 | 100200000 | -0.23 | -0.35 | B>B |
| chr12 | 1E+08 | 100300000 | 0.05 | -0.02 | A>A |
| chr12 | 1E+08 | 100400000 | 0.27 | 0.11 | A>A |
| chr12 | 1E+08 | 100500000 | 0.22 | 0.10 | A>A |
| chr12 | 1.01E+08 | 100600000 | 0.42 | 0.27 | A>A |
| chr12 | 1.01E+08 | 100700000 | 0.21 | 0.18 | A>A |
| chr12 | 1.01E+08 | 100800000 | -0.05 | 0.09 | A>A |
| chr12 | 1.01E+08 | 100900000 | -0.07 | 0.30 | A>A |
| chr12 | 1.01E+08 | 101000000 | -0.11 | 0.08 | B>B |
| chr12 | 1.01E+08 | 101100000 | -0.41 | -0.42 | B>B |
| chr12 | 1.01E+08 | 101200000 | -0.71 | -0.82 | B>B |
| chr12 | 1.01E+08 | 101300000 | -0.94 | -1.00 | B>B |
| chr12 | 1.01E+08 | 101400000 | -1.00 | -0.93 | B>B |
| chr12 | 1.01E+08 | 101500000 | -0.95 | -0.76 | B>B |
| chr12 | 1.02E+08 | 101600000 | -0.94 | -0.96 | B>B |
| chr12 | 1.02E+08 | 101700000 | -0.67 | -0.82 | B>B |
| chr12 | 1.02E+08 | 101800000 | -0.09 | -0.30 | B>B |
| chr12 | 1.02E+08 | 101900000 | 0.18 | -0.25 | B>B |
| chr12 | 1.02E+08 | 102000000 | 0.02 | -0.40 | B>B |

| | | | | | |
|---|---|---|---|---|---|
| chr12 | 1.02E+08 | 102100000 | 0.14 | -0.32 | B>B |
| chr12 | 1.02E+08 | 102200000 | 0.12 | -0.15 | B>B |
| chr12 | 1.02E+08 | 102300000 | 0.18 | -0.11 | A>A |
| chr12 | 1.02E+08 | 102400000 | -0.19 | -0.57 | B>B |
| chr12 | 1.02E+08 | 102500000 | -0.08 | -0.39 | B>B |
| chr12 | 1.03E+08 | 102600000 | -0.13 | -0.52 | B>B |
| chr12 | 1.03E+08 | 102700000 | -0.45 | -0.81 | B>B |
| chr12 | 1.03E+08 | 102800000 | -0.58 | -1.14 | B>B |
| chr12 | 1.03E+08 | 102900000 | -0.43 | -0.93 | B>B |
| chr12 | 1.03E+08 | 103000000 | -0.23 | -0.64 | B>B |
| chr12 | 1.03E+08 | 103100000 | -0.32 | -0.73 | B>B |
| chr12 | 1.03E+08 | 103200000 | -0.42 | -0.83 | B>B |
| chr12 | 1.03E+08 | 103300000 | -0.52 | -0.90 | B>B |
| chr12 | 1.03E+08 | 103400000 | -0.69 | -1.08 | B>B |
| chr12 | 1.03E+08 | 103500000 | -1.00 | -1.40 | B>B |
| chr12 | 1.04E+08 | 103600000 | -0.95 | -1.38 | B>B |
| chr12 | 1.04E+08 | 103700000 | -0.93 | -1.40 | B>B |
| chr12 | 1.04E+08 | 103800000 | -0.88 | -1.21 | B>B |
| chr12 | 1.04E+08 | 103900000 | -0.65 | -0.91 | B>B |
| chr12 | 1.04E+08 | 104000000 | -0.29 | -0.56 | B>B |
| chr12 | 1.04E+08 | 104100000 | 0.09 | -0.08 | A>A |
| chr12 | 1.04E+08 | 104200000 | 0.33 | 0.17 | A>A |
| chr12 | 1.04E+08 | 104300000 | 0.51 | 0.38 | A>A |
| chr12 | 1.04E+08 | 104400000 | 0.73 | 0.59 | A>A |
| chr12 | 1.04E+08 | 104500000 | 0.74 | 0.73 | A>A |
| chr12 | 1.05E+08 | 104600000 | 0.81 | 0.73 | A>A |
| chr12 | 1.05E+08 | 104700000 | 1.19 | 1.11 | A>A |
| chr12 | 1.05E+08 | 104800000 | 0.69 | 0.75 | A>A |
| chr12 | 1.05E+08 | 104900000 | 0.63 | 0.75 | A>A |
| chr12 | 1.05E+08 | 105000000 | 0.64 | 0.77 | A>A |
| chr12 | 1.05E+08 | 105100000 | 0.59 | 0.72 | A>A |
| chr12 | 1.05E+08 | 105200000 | 0.59 | 0.67 | A>A |
| chr12 | 1.05E+08 | 105300000 | 0.54 | 0.64 | A>A |
| chr12 | 1.05E+08 | 105400000 | 0.75 | 0.80 | A>A |
| chr12 | 1.05E+08 | 105500000 | 0.62 | 0.67 | A>A |
| chr12 | 1.06E+08 | 105600000 | 0.84 | 0.73 | A>A |
| chr12 | 1.06E+08 | 105700000 | 1.06 | 0.92 | A>A |
| chr12 | 1.06E+08 | 105800000 | 0.77 | 0.77 | A>A |
| chr12 | 1.06E+08 | 105900000 | 0.46 | 0.40 | A>A |
| chr12 | 1.06E+08 | 106000000 | 0.51 | 0.43 | A>A |
| chr12 | 1.06E+08 | 106100000 | 0.38 | 0.35 | A>A |
| chr12 | 1.06E+08 | 106200000 | 0.53 | 0.35 | A>A |
| chr12 | 1.06E+08 | 106300000 | 0.50 | 0.34 | A>A |
| chr12 | 1.06E+08 | 106400000 | 0.62 | 0.44 | A>A |
| chr12 | 1.06E+08 | 106500000 | 0.70 | 0.50 | A>A |
| chr12 | 1.07E+08 | 106600000 | 0.52 | 0.43 | A>A |
| chr12 | 1.07E+08 | 106700000 | 0.59 | 0.55 | A>A |
| chr12 | 1.07E+08 | 106800000 | 0.54 | 0.46 | A>A |
| chr12 | 1.07E+08 | 106900000 | 0.47 | 0.26 | A>A |
| chr12 | 1.07E+08 | 107000000 | 0.35 | 0.11 | A>A |
| chr12 | 1.07E+08 | 107100000 | 0.41 | 0.21 | A>A |
| chr12 | 1.07E+08 | 107200000 | 0.59 | 0.30 | A>A |
| chr12 | 1.07E+08 | 107300000 | 0.50 | 0.38 | A>A |
| chr12 | 1.07E+08 | 107400000 | 0.55 | 0.41 | A>A |
| chr12 | 1.07E+08 | 107500000 | 0.49 | 0.47 | A>A |
| chr12 | 1.08E+08 | 107600000 | 0.33 | 0.12 | A>A |

| | | | | | |
|---|---|---|---|---|---|
| chr12 | 1.08E+08 | 107700000 | 0.41 | 0.16 | A>A |
| chr12 | 1.08E+08 | 107800000 | 0.51 | 0.41 | A>A |
| chr12 | 1.08E+08 | 107900000 | 0.29 | 0.30 | A>A |
| chr12 | 1.08E+08 | 108000000 | 0.20 | 0.21 | A>A |
| chr12 | 1.08E+08 | 108100000 | 0.27 | 0.20 | A>A |
| chr12 | 1.08E+08 | 108200000 | 0.26 | -0.14 | A>A |
| chr12 | 1.08E+08 | 108300000 | 0.01 | -0.42 | B>B |
| chr12 | 1.08E+08 | 108400000 | -0.24 | -0.52 | B>B |
| chr12 | 1.08E+08 | 108500000 | -0.30 | -0.64 | B>B |
| chr12 | 1.09E+08 | 108600000 | -0.16 | -0.50 | B>B |
| chr12 | 1.09E+08 | 108700000 | -0.12 | -0.37 | B>B |
| chr12 | 1.09E+08 | 108800000 | -0.02 | -0.25 | B>B |
| chr12 | 1.09E+08 | 108900000 | 0.06 | -0.28 | B>B |
| chr12 | 1.09E+08 | 109000000 | 0.54 | 0.27 | A>A |
| chr12 | 1.09E+08 | 109100000 | 0.90 | 0.63 | A>A |
| chr12 | 1.09E+08 | 109200000 | 1.02 | 0.72 | A>A |
| chr12 | 1.09E+08 | 109300000 | 0.95 | 0.61 | A>A |
| chr12 | 1.09E+08 | 109400000 | 0.75 | 0.45 | A>A |
| chr12 | 1.09E+08 | 109500000 | 0.75 | 0.43 | A>A |
| chr12 | 1.1E+08 | 109600000 | 0.98 | 0.70 | A>A |
| chr12 | 1.1E+08 | 109700000 | 1.05 | 0.80 | A>A |
| chr12 | 1.1E+08 | 109800000 | 1.00 | 0.78 | A>A |
| chr12 | 1.1E+08 | 109900000 | 0.88 | 0.79 | A>A |
| chr12 | 1.1E+08 | 110000000 | 0.98 | 0.95 | A>A |
| chr12 | 1.1E+08 | 110100000 | 1.08 | 0.91 | A>A |
| chr12 | 1.1E+08 | 110200000 | 1.13 | 0.96 | A>A |
| chr12 | 1.1E+08 | 110300000 | 0.92 | 1.02 | A>A |
| chr12 | 1.1E+08 | 110400000 | 1.11 | 1.06 | A>A |
| chr12 | 1.1E+08 | 110500000 | 1.21 | 1.14 | A>A |
| chr12 | 1.11E+08 | 110600000 | 1.16 | 1.13 | A>A |
| chr12 | 1.11E+08 | 110700000 | 1.22 | 1.22 | A>A |
| chr12 | 1.11E+08 | 110800000 | 1.27 | 1.29 | A>A |
| chr12 | 1.11E+08 | 110900000 | 1.00 | 1.04 | A>A |
| chr12 | 1.11E+08 | 111000000 | 1.11 | 1.10 | A>A |
| chr12 | 1.11E+08 | 111100000 | 0.90 | 0.98 | A>A |
| chr12 | 1.11E+08 | 111200000 | 1.05 | 0.93 | A>A |
| chr12 | 1.11E+08 | 111300000 | 0.96 | 0.84 | A>A |
| chr12 | 1.11E+08 | 111400000 | 0.69 | 0.67 | A>A |
| chr12 | 1.11E+08 | 111500000 | 0.73 | 0.71 | A>A |
| chr12 | 1.12E+08 | 111600000 | 0.88 | 0.64 | A>A |
| chr12 | 1.12E+08 | 111700000 | 0.77 | 0.65 | A>A |
| chr12 | 1.12E+08 | 111800000 | 0.72 | 0.71 | A>A |
| chr12 | 1.12E+08 | 111900000 | 0.95 | 0.97 | A>A |
| chr12 | 1.12E+08 | 112000000 | 1.06 | 1.01 | A>A |
| chr12 | 1.12E+08 | 112100000 | 1.14 | 1.17 | A>A |
| chr12 | 1.12E+08 | 112200000 | 1.07 | 1.10 | A>A |
| chr12 | 1.12E+08 | 112300000 | 1.20 | 1.12 | A>A |
| chr12 | 1.12E+08 | 112400000 | 1.01 | 1.08 | A>A |
| chr12 | 1.12E+08 | 112500000 | 1.15 | 1.12 | A>A |
| chr12 | 1.13E+08 | 112600000 | 1.22 | 1.25 | A>A |
| chr12 | 1.13E+08 | 112700000 | 1.11 | 1.03 | A>A |
| chr12 | 1.13E+08 | 112800000 | 1.11 | 1.08 | A>A |
| chr12 | 1.13E+08 | 112900000 | 0.97 | 0.97 | A>A |
| chr12 | 1.13E+08 | 113000000 | 0.69 | 0.71 | A>A |
| chr12 | 1.13E+08 | 113100000 | 0.64 | 0.47 | A>A |
| chr12 | 1.13E+08 | 113200000 | 0.60 | 0.52 | A>A |

| | | | | | |
|---|---|---|---|---|---|
| chr12 | 1.13E+08 | 113300000 | 0.80 | 0.71 | A>A |
| chr12 | 1.13E+08 | 113400000 | 0.76 | 0.72 | A>A |
| chr12 | 1.13E+08 | 113500000 | 0.73 | 0.77 | A>A |
| chr12 | 1.14E+08 | 113600000 | 0.99 | 0.92 | A>A |
| chr12 | 1.14E+08 | 113700000 | 1.00 | 0.89 | A>A |
| chr12 | 1.14E+08 | 113800000 | 0.98 | 0.79 | A>A |
| chr12 | 1.14E+08 | 113900000 | 0.63 | 0.52 | A>A |
| chr12 | 1.14E+08 | 114000000 | 0.32 | 0.06 | A>A |
| chr12 | 1.14E+08 | 114100000 | 0.30 | 0.04 | A>A |
| chr12 | 1.14E+08 | 114200000 | 0.40 | 0.09 | A>A |
| chr12 | 1.14E+08 | 114300000 | 0.43 | 0.16 | A>A |
| chr12 | 1.14E+08 | 114400000 | 0.28 | -0.01 | A>A |
| chr12 | 1.14E+08 | 114500000 | -0.16 | -0.81 | A>B |
| chr12 | 1.15E+08 | 114600000 | -0.40 | -1.30 | A>B |
| chr12 | 1.15E+08 | 114700000 | -0.38 | -1.27 | A>B |
| chr12 | 1.15E+08 | 114800000 | -0.34 | -1.29 | A>B |
| chr12 | 1.15E+08 | 114900000 | -0.21 | -1.23 | A>B |
| chr12 | 1.15E+08 | 115000000 | -0.12 | -1.07 | A>B |
| chr12 | 1.15E+08 | 115100000 | 0.05 | -0.84 | A>B |
| chr12 | 1.15E+08 | 115200000 | 0.23 | -0.66 | A>B |
| chr12 | 1.15E+08 | 115300000 | 0.38 | -0.97 | A>B |
| chr12 | 1.15E+08 | 115400000 | 0.36 | -1.04 | A>B |
| chr12 | 1.15E+08 | 115500000 | 0.25 | -1.04 | A>B |
| chr12 | 1.16E+08 | 115600000 | 0.20 | -1.12 | A>B |
| chr12 | 1.16E+08 | 115700000 | 0.21 | -1.09 | A>B |
| chr12 | 1.16E+08 | 115800000 | 0.29 | -1.07 | A>B |
| chr12 | 1.16E+08 | 115900000 | 0.27 | -1.08 | A>B |
| chr12 | 1.16E+08 | 116000000 | 0.13 | -1.16 | A>B |
| chr12 | 1.16E+08 | 116100000 | 0.09 | -1.21 | A>B |
| chr12 | 1.16E+08 | 116200000 | 0.06 | -1.09 | A>B |
| chr12 | 1.16E+08 | 116300000 | 0.21 | -1.03 | A>B |
| chr12 | 1.16E+08 | 116400000 | 0.18 | -0.57 | A>B |
| chr12 | 1.16E+08 | 116500000 | 0.51 | 0.21 | A>A |
| chr12 | 1.17E+08 | 116600000 | 0.69 | 0.44 | A>A |
| chr12 | 1.17E+08 | 116700000 | 0.68 | 0.44 | A>A |
| chr12 | 1.17E+08 | 116800000 | 0.80 | 0.59 | A>A |
| chr12 | 1.17E+08 | 116900000 | 0.81 | 0.65 | A>A |
| chr12 | 1.17E+08 | 117000000 | 0.75 | 0.57 | A>A |
| chr12 | 1.17E+08 | 117100000 | 0.60 | 0.39 | A>A |
| chr12 | 1.17E+08 | 117200000 | 0.50 | 0.35 | A>A |
| chr12 | 1.17E+08 | 117300000 | 0.67 | 0.49 | A>A |
| chr12 | 1.17E+08 | 117400000 | 0.61 | 0.52 | A>A |
| chr12 | 1.17E+08 | 117500000 | 0.50 | 0.53 | A>A |
| chr12 | 1.18E+08 | 117600000 | 0.66 | 0.72 | A>A |
| chr12 | 1.18E+08 | 117700000 | 0.58 | 0.53 | A>A |
| chr12 | 1.18E+08 | 117800000 | 0.56 | 0.57 | A>A |
| chr12 | 1.18E+08 | 117900000 | 0.56 | 0.54 | A>A |
| chr12 | 1.18E+08 | 118000000 | 0.67 | 0.62 | A>A |
| chr12 | 1.18E+08 | 118100000 | 0.40 | 0.51 | A>A |
| chr12 | 1.18E+08 | 118200000 | 0.60 | 0.65 | A>A |
| chr12 | 1.18E+08 | 118300000 | 0.48 | 0.47 | A>A |
| chr12 | 1.18E+08 | 118400000 | 0.58 | 0.53 | A>A |
| chr12 | 1.18E+08 | 118500000 | 0.70 | 0.69 | A>A |
| chr12 | 1.19E+08 | 118600000 | 0.53 | 0.42 | A>A |
| chr12 | 1.19E+08 | 118700000 | 0.39 | 0.14 | A>A |
| chr12 | 1.19E+08 | 118800000 | 0.34 | 0.20 | A>A |

| | | | | | |
|---|---|---|---|---|---|
| chr12 | 1.19E+08 | 118900000 | -0.17 | -0.51 | B>B |
| chr12 | 1.19E+08 | 119000000 | -0.64 | -1.10 | B>B |
| chr12 | 1.19E+08 | 119100000 | -0.68 | -1.28 | A>B |
| chr12 | 1.19E+08 | 119200000 | -0.82 | -1.27 | B>B |
| chr12 | 1.19E+08 | 119300000 | -0.89 | -1.30 | B>B |
| chr12 | 1.19E+08 | 119400000 | -0.84 | -1.24 | B>B |
| chr12 | 1.19E+08 | 119500000 | -0.72 | -1.12 | B>B |
| chr12 | 1.2E+08 | 119600000 | -0.58 | -0.88 | B>B |
| chr12 | 1.2E+08 | 119700000 | -0.47 | -0.70 | B>B |
| chr12 | 1.2E+08 | 119800000 | -0.63 | -1.11 | B>B |
| chr12 | 1.2E+08 | 119900000 | -0.61 | -1.02 | B>B |
| chr12 | 1.2E+08 | 120000000 | -0.57 | -0.98 | B>B |
| chr12 | 1.2E+08 | 120100000 | -0.26 | -0.58 | B>B |
| chr12 | 1.2E+08 | 120200000 | 0.49 | 0.39 | A>A |
| chr12 | 1.2E+08 | 120300000 | 0.84 | 0.73 | A>A |
| chr12 | 1.2E+08 | 120400000 | 0.96 | 0.88 | A>A |
| chr12 | 1.2E+08 | 120500000 | 1.09 | 1.15 | A>A |
| chr12 | 1.21E+08 | 120600000 | 1.47 | 1.34 | A>A |
| chr12 | 1.21E+08 | 120700000 | 1.61 | 1.48 | A>A |
| chr12 | 1.21E+08 | 120800000 | 1.24 | 1.33 | A>A |
| chr12 | 1.21E+08 | 120900000 | 1.32 | 1.31 | A>A |
| chr12 | 1.21E+08 | 121000000 | 1.41 | 1.27 | A>A |
| chr12 | 1.21E+08 | 121100000 | 1.10 | 1.07 | A>A |
| chr12 | 1.21E+08 | 121200000 | 1.13 | 1.10 | A>A |
| chr12 | 1.21E+08 | 121300000 | 1.14 | 1.11 | A>A |
| chr12 | 1.21E+08 | 121400000 | 1.02 | 0.98 | A>A |
| chr12 | 1.21E+08 | 121500000 | 0.91 | 0.97 | A>A |
| chr12 | 1.22E+08 | 121600000 | 1.13 | 1.02 | A>A |
| chr12 | 1.22E+08 | 121700000 | 1.19 | 1.07 | A>A |
| chr12 | 1.22E+08 | 121800000 | 1.40 | 1.20 | A>A |
| chr12 | 1.22E+08 | 121900000 | 1.28 | 1.19 | A>A |
| chr12 | 1.22E+08 | 122000000 | 1.27 | 1.15 | A>A |
| chr12 | 1.22E+08 | 122100000 | 1.35 | 1.27 | A>A |
| chr12 | 1.22E+08 | 122200000 | 1.26 | 1.17 | A>A |
| chr12 | 1.22E+08 | 122300000 | 1.49 | 1.32 | A>A |
| chr12 | 1.22E+08 | 122400000 | 1.31 | 1.25 | A>A |
| chr12 | 1.22E+08 | 122500000 | 1.16 | 1.11 | A>A |
| chr12 | 1.23E+08 | 122600000 | 0.00 | 0.00 | NO |
| chr12 | 1.23E+08 | 122700000 | 1.27 | 1.17 | A>A |
| chr12 | 1.23E+08 | 122800000 | 1.38 | 1.18 | A>A |
| chr12 | 1.23E+08 | 122900000 | 1.27 | 1.19 | A>A |
| chr12 | 1.23E+08 | 123000000 | 1.47 | 1.32 | A>A |
| chr12 | 1.23E+08 | 123100000 | 1.38 | 1.12 | A>A |
| chr12 | 1.23E+08 | 123200000 | 1.30 | 1.22 | A>A |
| chr12 | 1.23E+08 | 123300000 | 1.25 | 1.12 | A>A |
| chr12 | 1.23E+08 | 123400000 | 1.21 | 1.23 | A>A |
| chr12 | 1.23E+08 | 123500000 | 1.25 | 1.14 | A>A |
| chr12 | 1.24E+08 | 123600000 | 1.34 | 1.18 | A>A |
| chr12 | 1.24E+08 | 123700000 | 1.25 | 1.17 | A>A |
| chr12 | 1.24E+08 | 123800000 | 1.14 | 1.06 | A>A |
| chr12 | 1.24E+08 | 123900000 | 1.24 | 1.23 | A>A |
| chr12 | 1.24E+08 | 124000000 | 1.27 | 1.19 | A>A |
| chr12 | 1.24E+08 | 124100000 | 1.25 | 1.05 | A>A |
| chr12 | 1.24E+08 | 124200000 | 1.18 | 1.11 | A>A |
| chr12 | 1.24E+08 | 124300000 | 1.06 | 0.90 | A>A |
| chr12 | 1.24E+08 | 124400000 | 1.04 | 0.93 | A>A |

| | | | | | |
|---|---|---|---|---|---|
| chr12 | 1.24E+08 | 124500000 | 0.91 | 0.87 | A>A |
| chr12 | 1.25E+08 | 124600000 | 0.78 | 0.79 | A>A |
| chr12 | 1.25E+08 | 124700000 | 0.96 | 0.95 | A>A |
| chr12 | 1.25E+08 | 124800000 | 0.85 | 0.85 | A>A |
| chr12 | 1.25E+08 | 124900000 | 0.82 | 0.97 | A>A |
| chr12 | 1.25E+08 | 125000000 | 0.95 | 0.95 | A>A |
| chr12 | 1.25E+08 | 125100000 | 0.97 | 1.06 | A>A |
| chr12 | 1.25E+08 | 125200000 | 1.09 | 1.17 | A>A |
| chr12 | 1.25E+08 | 125300000 | 0.81 | 0.87 | A>A |
| chr12 | 1.25E+08 | 125400000 | 1.06 | 0.98 | A>A |
| chr12 | 1.25E+08 | 125500000 | 1.13 | 0.78 | A>A |
| chr12 | 1.26E+08 | 125600000 | 0.53 | 0.38 | A>A |
| chr12 | 1.26E+08 | 125700000 | 0.11 | -0.10 | A>A |
| chr12 | 1.26E+08 | 125800000 | -0.23 | -0.67 | B>B |
| chr12 | 1.26E+08 | 125900000 | -0.42 | -0.74 | B>B |
| chr12 | 1.26E+08 | 126000000 | -0.70 | -0.89 | B>B |
| chr12 | 1.26E+08 | 126100000 | -0.83 | -0.85 | B>B |
| chr12 | 1.26E+08 | 126200000 | -0.98 | -0.91 | B>B |
| chr12 | 1.26E+08 | 126300000 | -0.92 | -0.77 | B>B |
| chr12 | 1.26E+08 | 126400000 | -0.91 | -0.75 | B>B |
| chr12 | 1.26E+08 | 126500000 | -0.92 | -0.67 | B>B |
| chr12 | 1.27E+08 | 126600000 | -0.72 | -0.71 | B>B |
| chr12 | 1.27E+08 | 126700000 | -0.85 | -0.89 | B>B |
| chr12 | 1.27E+08 | 126800000 | -0.86 | -1.06 | B>B |
| chr12 | 1.27E+08 | 126900000 | -0.87 | -1.01 | B>B |
| chr12 | 1.27E+08 | 127000000 | -0.88 | -0.96 | B>B |
| chr12 | 1.27E+08 | 127100000 | -0.80 | -0.99 | B>B |
| chr12 | 1.27E+08 | 127200000 | -0.82 | -0.91 | B>B |
| chr12 | 1.27E+08 | 127300000 | -0.72 | -0.71 | B>B |
| chr12 | 1.27E+08 | 127400000 | -0.72 | -0.78 | B>B |
| chr12 | 1.27E+08 | 127500000 | -0.62 | -0.77 | B>B |
| chr12 | 1.28E+08 | 127600000 | -0.48 | -0.61 | B>B |
| chr12 | 1.28E+08 | 127700000 | -0.39 | -0.62 | B>B |
| chr12 | 1.28E+08 | 127800000 | -0.73 | -0.94 | B>B |
| chr12 | 1.28E+08 | 127900000 | -0.68 | -0.81 | B>B |
| chr12 | 1.28E+08 | 128000000 | -0.80 | -0.91 | B>B |
| chr12 | 1.28E+08 | 128100000 | -0.82 | -1.02 | B>B |
| chr12 | 1.28E+08 | 128200000 | -0.79 | -1.03 | B>B |
| chr12 | 1.28E+08 | 128300000 | -0.82 | -1.03 | B>B |
| chr12 | 1.28E+08 | 128400000 | -0.84 | -1.07 | B>B |
| chr12 | 1.28E+08 | 128500000 | -0.86 | -1.05 | B>B |
| chr12 | 1.29E+08 | 128600000 | -0.72 | -0.92 | B>B |
| chr12 | 1.29E+08 | 128700000 | -0.76 | -1.02 | B>B |
| chr12 | 1.29E+08 | 128800000 | -0.81 | -1.07 | B>B |
| chr12 | 1.29E+08 | 128900000 | -0.77 | -1.06 | B>B |
| chr12 | 1.29E+08 | 129000000 | -0.73 | -1.08 | B>B |
| chr12 | 1.29E+08 | 129100000 | -0.70 | -1.04 | B>B |
| chr12 | 1.29E+08 | 129200000 | -0.69 | -0.98 | B>B |
| chr12 | 1.29E+08 | 129300000 | -0.43 | -0.71 | B>B |
| chr12 | 1.29E+08 | 129400000 | -0.44 | -0.76 | B>B |
| chr12 | 1.29E+08 | 129500000 | -0.56 | -0.94 | B>B |
| chr12 | 1.3E+08 | 129600000 | -0.77 | -1.05 | B>B |
| chr12 | 1.3E+08 | 129700000 | -0.80 | -1.04 | B>B |
| chr12 | 1.3E+08 | 129800000 | -0.82 | -1.09 | B>B |
| chr12 | 1.3E+08 | 129900000 | -0.83 | -1.12 | B>B |
| chr12 | 1.3E+08 | 130000000 | -0.79 | -1.20 | B>B |

| | | | | | |
|---|---|---|---|---|---|
| chr12 | 1.3E+08 | 130100000 | -0.91 | -1.22 | B>B |
| chr12 | 1.3E+08 | 130200000 | -0.87 | -1.21 | B>B |
| chr12 | 1.3E+08 | 130300000 | -0.90 | -1.13 | B>B |
| chr12 | 1.3E+08 | 130400000 | -0.81 | -1.05 | B>B |
| chr12 | 1.3E+08 | 130500000 | -0.70 | -0.89 | B>B |
| chr12 | 1.31E+08 | 130600000 | -0.33 | -0.07 | B>B |
| chr12 | 1.31E+08 | 130700000 | -0.28 | -0.03 | B>B |
| chr12 | 1.31E+08 | 130800000 | -0.44 | -0.34 | B>B |
| chr12 | 1.31E+08 | 130900000 | -0.32 | -0.35 | B>B |
| chr12 | 1.31E+08 | 131000000 | -0.13 | -0.39 | B>B |
| chr12 | 1.31E+08 | 131100000 | -0.18 | -0.50 | B>B |
| chr12 | 1.31E+08 | 131200000 | 0.07 | -0.35 | B>B |
| chr12 | 1.31E+08 | 131300000 | 0.39 | 0.15 | A>A |
| chr12 | 1.31E+08 | 131400000 | 0.78 | 0.72 | A>A |
| chr12 | 1.31E+08 | 131500000 | 0.78 | 0.58 | A>A |
| chr12 | 1.32E+08 | 131600000 | 0.76 | 0.67 | A>A |
| chr12 | 1.32E+08 | 131700000 | 0.69 | 0.73 | A>A |
| chr12 | 1.32E+08 | 131800000 | 1.10 | 1.00 | A>A |
| chr12 | 1.32E+08 | 131900000 | 0.92 | 0.82 | A>A |
| chr12 | 1.32E+08 | 132000000 | 0.78 | 0.73 | A>A |
| chr12 | 1.32E+08 | 132100000 | 0.74 | 0.58 | A>A |
| chr12 | 1.32E+08 | 132200000 | 1.01 | 0.73 | A>A |
| chr12 | 1.32E+08 | 132300000 | 1.10 | 0.95 | A>A |
| chr12 | 1.32E+08 | 132400000 | 1.39 | 1.35 | A>A |
| chr12 | 1.32E+08 | 132500000 | 1.59 | 1.39 | A>A |
| chr12 | 1.33E+08 | 132600000 | 1.49 | 1.46 | A>A |
| chr12 | 1.33E+08 | 132700000 | 1.25 | 1.26 | A>A |
| chr12 | 1.33E+08 | 132800000 | 0.00 | 0.00 | NO |
| chr12 | 1.33E+08 | 132900000 | 1.29 | 1.34 | A>A |
| chr12 | 1.33E+08 | 133000000 | 1.44 | 1.31 | A>A |
| chr12 | 1.33E+08 | 133100000 | 1.66 | 1.55 | A>A |
| chr12 | 1.33E+08 | 133200000 | 1.54 | 1.40 | A>A |
| chr12 | 1.33E+08 | 133300000 | 1.77 | 1.52 | A>A |
| chr12 | 1.33E+08 | 133400000 | 1.79 | 1.75 | A>A |
| chr12 | 1.33E+08 | 133500000 | 1.16 | 1.57 | A>A |
| chr12 | 1.34E+08 | 133600000 | 0.00 | 0.00 | NO |
| chr12 | 1.34E+08 | 133700000 | 0.00 | 0.00 | NO |
| chr12 | 1.34E+08 | 133800000 | 0.33 | 1.25 | B>A |
| chr12 | 1.34E+08 | 133851895 | 0.00 | 0.00 | NO |
| chr13 | 0 | 100000 | 0.00 | 0.00 | NO |
| chr13 | 100000 | 200000 | 0.00 | 0.00 | NO |
| chr13 | 200000 | 300000 | 0.00 | 0.00 | NO |
| chr13 | 300000 | 400000 | 0.00 | 0.00 | NO |
| chr13 | 400000 | 500000 | 0.00 | 0.00 | NO |
| chr13 | 500000 | 600000 | 0.00 | 0.00 | NO |
| chr13 | 600000 | 700000 | 0.00 | 0.00 | NO |
| chr13 | 700000 | 800000 | 0.00 | 0.00 | NO |
| chr13 | 800000 | 900000 | 0.00 | 0.00 | NO |
| chr13 | 900000 | 1000000 | 0.00 | 0.00 | NO |
| chr13 | 1000000 | 1100000 | 0.00 | 0.00 | NO |
| chr13 | 1100000 | 1200000 | 0.00 | 0.00 | NO |
| chr13 | 1200000 | 1300000 | 0.00 | 0.00 | NO |
| chr13 | 1300000 | 1400000 | 0.00 | 0.00 | NO |
| chr13 | 1400000 | 1500000 | 0.00 | 0.00 | NO |
| chr13 | 1500000 | 1600000 | 0.00 | 0.00 | NO |
| chr13 | 1600000 | 1700000 | 0.00 | 0.00 | NO |

| | | | | | |
|---|---|---|---|---|---|
| chr13 | 1700000 | 1800000 | 0.00 | 0.00 | NO |
| chr13 | 1800000 | 1900000 | 0.00 | 0.00 | NO |
| chr13 | 1900000 | 2000000 | 0.00 | 0.00 | NO |
| chr13 | 2000000 | 2100000 | 0.00 | 0.00 | NO |
| chr13 | 2100000 | 2200000 | 0.00 | 0.00 | NO |
| chr13 | 2200000 | 2300000 | 0.00 | 0.00 | NO |
| chr13 | 2300000 | 2400000 | 0.00 | 0.00 | NO |
| chr13 | 2400000 | 2500000 | 0.00 | 0.00 | NO |
| chr13 | 2500000 | 2600000 | 0.00 | 0.00 | NO |
| chr13 | 2600000 | 2700000 | 0.00 | 0.00 | NO |
| chr13 | 2700000 | 2800000 | 0.00 | 0.00 | NO |
| chr13 | 2800000 | 2900000 | 0.00 | 0.00 | NO |
| chr13 | 2900000 | 3000000 | 0.00 | 0.00 | NO |
| chr13 | 3000000 | 3100000 | 0.00 | 0.00 | NO |
| chr13 | 3100000 | 3200000 | 0.00 | 0.00 | NO |
| chr13 | 3200000 | 3300000 | 0.00 | 0.00 | NO |
| chr13 | 3300000 | 3400000 | 0.00 | 0.00 | NO |
| chr13 | 3400000 | 3500000 | 0.00 | 0.00 | NO |
| chr13 | 3500000 | 3600000 | 0.00 | 0.00 | NO |
| chr13 | 3600000 | 3700000 | 0.00 | 0.00 | NO |
| chr13 | 3700000 | 3800000 | 0.00 | 0.00 | NO |
| chr13 | 3800000 | 3900000 | 0.00 | 0.00 | NO |
| chr13 | 3900000 | 4000000 | 0.00 | 0.00 | NO |
| chr13 | 4000000 | 4100000 | 0.00 | 0.00 | NO |
| chr13 | 4100000 | 4200000 | 0.00 | 0.00 | NO |
| chr13 | 4200000 | 4300000 | 0.00 | 0.00 | NO |
| chr13 | 4300000 | 4400000 | 0.00 | 0.00 | NO |
| chr13 | 4400000 | 4500000 | 0.00 | 0.00 | NO |
| chr13 | 4500000 | 4600000 | 0.00 | 0.00 | NO |
| chr13 | 4600000 | 4700000 | 0.00 | 0.00 | NO |
| chr13 | 4700000 | 4800000 | 0.00 | 0.00 | NO |
| chr13 | 4800000 | 4900000 | 0.00 | 0.00 | NO |
| chr13 | 4900000 | 5000000 | 0.00 | 0.00 | NO |
| chr13 | 5000000 | 5100000 | 0.00 | 0.00 | NO |
| chr13 | 5100000 | 5200000 | 0.00 | 0.00 | NO |
| chr13 | 5200000 | 5300000 | 0.00 | 0.00 | NO |
| chr13 | 5300000 | 5400000 | 0.00 | 0.00 | NO |
| chr13 | 5400000 | 5500000 | 0.00 | 0.00 | NO |
| chr13 | 5500000 | 5600000 | 0.00 | 0.00 | NO |
| chr13 | 5600000 | 5700000 | 0.00 | 0.00 | NO |
| chr13 | 5700000 | 5800000 | 0.00 | 0.00 | NO |
| chr13 | 5800000 | 5900000 | 0.00 | 0.00 | NO |
| chr13 | 5900000 | 6000000 | 0.00 | 0.00 | NO |
| chr13 | 6000000 | 6100000 | 0.00 | 0.00 | NO |
| chr13 | 6100000 | 6200000 | 0.00 | 0.00 | NO |
| chr13 | 6200000 | 6300000 | 0.00 | 0.00 | NO |
| chr13 | 6300000 | 6400000 | 0.00 | 0.00 | NO |
| chr13 | 6400000 | 6500000 | 0.00 | 0.00 | NO |
| chr13 | 6500000 | 6600000 | 0.00 | 0.00 | NO |
| chr13 | 6600000 | 6700000 | 0.00 | 0.00 | NO |
| chr13 | 6700000 | 6800000 | 0.00 | 0.00 | NO |
| chr13 | 6800000 | 6900000 | 0.00 | 0.00 | NO |
| chr13 | 6900000 | 7000000 | 0.00 | 0.00 | NO |
| chr13 | 7000000 | 7100000 | 0.00 | 0.00 | NO |
| chr13 | 7100000 | 7200000 | 0.00 | 0.00 | NO |
| chr13 | 7200000 | 7300000 | 0.00 | 0.00 | NO |

| | | | | | |
|---|---|---|---|---|---|
| chr13 | 7300000 | 7400000 | 0.00 | 0.00 | NO |
| chr13 | 7400000 | 7500000 | 0.00 | 0.00 | NO |
| chr13 | 7500000 | 7600000 | 0.00 | 0.00 | NO |
| chr13 | 7600000 | 7700000 | 0.00 | 0.00 | NO |
| chr13 | 7700000 | 7800000 | 0.00 | 0.00 | NO |
| chr13 | 7800000 | 7900000 | 0.00 | 0.00 | NO |
| chr13 | 7900000 | 8000000 | 0.00 | 0.00 | NO |
| chr13 | 8000000 | 8100000 | 0.00 | 0.00 | NO |
| chr13 | 8100000 | 8200000 | 0.00 | 0.00 | NO |
| chr13 | 8200000 | 8300000 | 0.00 | 0.00 | NO |
| chr13 | 8300000 | 8400000 | 0.00 | 0.00 | NO |
| chr13 | 8400000 | 8500000 | 0.00 | 0.00 | NO |
| chr13 | 8500000 | 8600000 | 0.00 | 0.00 | NO |
| chr13 | 8600000 | 8700000 | 0.00 | 0.00 | NO |
| chr13 | 8700000 | 8800000 | 0.00 | 0.00 | NO |
| chr13 | 8800000 | 8900000 | 0.00 | 0.00 | NO |
| chr13 | 8900000 | 9000000 | 0.00 | 0.00 | NO |
| chr13 | 9000000 | 9100000 | 0.00 | 0.00 | NO |
| chr13 | 9100000 | 9200000 | 0.00 | 0.00 | NO |
| chr13 | 9200000 | 9300000 | 0.00 | 0.00 | NO |
| chr13 | 9300000 | 9400000 | 0.00 | 0.00 | NO |
| chr13 | 9400000 | 9500000 | 0.00 | 0.00 | NO |
| chr13 | 9500000 | 9600000 | 0.00 | 0.00 | NO |
| chr13 | 9600000 | 9700000 | 0.00 | 0.00 | NO |
| chr13 | 9700000 | 9800000 | 0.00 | 0.00 | NO |
| chr13 | 9800000 | 9900000 | 0.00 | 0.00 | NO |
| chr13 | 9900000 | 10000000 | 0.00 | 0.00 | NO |
| chr13 | 10000000 | 10100000 | 0.00 | 0.00 | NO |
| chr13 | 10100000 | 10200000 | 0.00 | 0.00 | NO |
| chr13 | 10200000 | 10300000 | 0.00 | 0.00 | NO |
| chr13 | 10300000 | 10400000 | 0.00 | 0.00 | NO |
| chr13 | 10400000 | 10500000 | 0.00 | 0.00 | NO |
| chr13 | 10500000 | 10600000 | 0.00 | 0.00 | NO |
| chr13 | 10600000 | 10700000 | 0.00 | 0.00 | NO |
| chr13 | 10700000 | 10800000 | 0.00 | 0.00 | NO |
| chr13 | 10800000 | 10900000 | 0.00 | 0.00 | NO |
| chr13 | 10900000 | 11000000 | 0.00 | 0.00 | NO |
| chr13 | 11000000 | 11100000 | 0.00 | 0.00 | NO |
| chr13 | 11100000 | 11200000 | 0.00 | 0.00 | NO |
| chr13 | 11200000 | 11300000 | 0.00 | 0.00 | NO |
| chr13 | 11300000 | 11400000 | 0.00 | 0.00 | NO |
| chr13 | 11400000 | 11500000 | 0.00 | 0.00 | NO |
| chr13 | 11500000 | 11600000 | 0.00 | 0.00 | NO |
| chr13 | 11600000 | 11700000 | 0.00 | 0.00 | NO |
| chr13 | 11700000 | 11800000 | 0.00 | 0.00 | NO |
| chr13 | 11800000 | 11900000 | 0.00 | 0.00 | NO |
| chr13 | 11900000 | 12000000 | 0.00 | 0.00 | NO |
| chr13 | 12000000 | 12100000 | 0.00 | 0.00 | NO |
| chr13 | 12100000 | 12200000 | 0.00 | 0.00 | NO |
| chr13 | 12200000 | 12300000 | 0.00 | 0.00 | NO |
| chr13 | 12300000 | 12400000 | 0.00 | 0.00 | NO |
| chr13 | 12400000 | 12500000 | 0.00 | 0.00 | NO |
| chr13 | 12500000 | 12600000 | 0.00 | 0.00 | NO |
| chr13 | 12600000 | 12700000 | 0.00 | 0.00 | NO |
| chr13 | 12700000 | 12800000 | 0.00 | 0.00 | NO |
| chr13 | 12800000 | 12900000 | 0.00 | 0.00 | NO |

| | | | | | |
|---|---|---|---|---|---|
| chr13 | 12900000 | 13000000 | 0.00 | 0.00 | NO |
| chr13 | 13000000 | 13100000 | 0.00 | 0.00 | NO |
| chr13 | 13100000 | 13200000 | 0.00 | 0.00 | NO |
| chr13 | 13200000 | 13300000 | 0.00 | 0.00 | NO |
| chr13 | 13300000 | 13400000 | 0.00 | 0.00 | NO |
| chr13 | 13400000 | 13500000 | 0.00 | 0.00 | NO |
| chr13 | 13500000 | 13600000 | 0.00 | 0.00 | NO |
| chr13 | 13600000 | 13700000 | 0.00 | 0.00 | NO |
| chr13 | 13700000 | 13800000 | 0.00 | 0.00 | NO |
| chr13 | 13800000 | 13900000 | 0.00 | 0.00 | NO |
| chr13 | 13900000 | 14000000 | 0.00 | 0.00 | NO |
| chr13 | 14000000 | 14100000 | 0.00 | 0.00 | NO |
| chr13 | 14100000 | 14200000 | 0.00 | 0.00 | NO |
| chr13 | 14200000 | 14300000 | 0.00 | 0.00 | NO |
| chr13 | 14300000 | 14400000 | 0.00 | 0.00 | NO |
| chr13 | 14400000 | 14500000 | 0.00 | 0.00 | NO |
| chr13 | 14500000 | 14600000 | 0.00 | 0.00 | NO |
| chr13 | 14600000 | 14700000 | 0.00 | 0.00 | NO |
| chr13 | 14700000 | 14800000 | 0.00 | 0.00 | NO |
| chr13 | 14800000 | 14900000 | 0.00 | 0.00 | NO |
| chr13 | 14900000 | 15000000 | 0.00 | 0.00 | NO |
| chr13 | 15000000 | 15100000 | 0.00 | 0.00 | NO |
| chr13 | 15100000 | 15200000 | 0.00 | 0.00 | NO |
| chr13 | 15200000 | 15300000 | 0.00 | 0.00 | NO |
| chr13 | 15300000 | 15400000 | 0.00 | 0.00 | NO |
| chr13 | 15400000 | 15500000 | 0.00 | 0.00 | NO |
| chr13 | 15500000 | 15600000 | 0.00 | 0.00 | NO |
| chr13 | 15600000 | 15700000 | 0.00 | 0.00 | NO |
| chr13 | 15700000 | 15800000 | 0.00 | 0.00 | NO |
| chr13 | 15800000 | 15900000 | 0.00 | 0.00 | NO |
| chr13 | 15900000 | 16000000 | 0.00 | 0.00 | NO |
| chr13 | 16000000 | 16100000 | 0.00 | 0.00 | NO |
| chr13 | 16100000 | 16200000 | 0.00 | 0.00 | NO |
| chr13 | 16200000 | 16300000 | 0.00 | 0.00 | NO |
| chr13 | 16300000 | 16400000 | 0.00 | 0.00 | NO |
| chr13 | 16400000 | 16500000 | 0.00 | 0.00 | NO |
| chr13 | 16500000 | 16600000 | 0.00 | 0.00 | NO |
| chr13 | 16600000 | 16700000 | 0.00 | 0.00 | NO |
| chr13 | 16700000 | 16800000 | 0.00 | 0.00 | NO |
| chr13 | 16800000 | 16900000 | 0.00 | 0.00 | NO |
| chr13 | 16900000 | 17000000 | 0.00 | 0.00 | NO |
| chr13 | 17000000 | 17100000 | 0.00 | 0.00 | NO |
| chr13 | 17100000 | 17200000 | 0.00 | 0.00 | NO |
| chr13 | 17200000 | 17300000 | 0.00 | 0.00 | NO |
| chr13 | 17300000 | 17400000 | 0.00 | 0.00 | NO |
| chr13 | 17400000 | 17500000 | 0.00 | 0.00 | NO |
| chr13 | 17500000 | 17600000 | 0.00 | 0.00 | NO |
| chr13 | 17600000 | 17700000 | 0.00 | 0.00 | NO |
| chr13 | 17700000 | 17800000 | 0.00 | 0.00 | NO |
| chr13 | 17800000 | 17900000 | 0.00 | 0.00 | NO |
| chr13 | 17900000 | 18000000 | 0.00 | 0.00 | NO |
| chr13 | 18000000 | 18100000 | 0.00 | 0.00 | NO |
| chr13 | 18100000 | 18200000 | 0.00 | 0.00 | NO |
| chr13 | 18200000 | 18300000 | 0.00 | 0.00 | NO |
| chr13 | 18300000 | 18400000 | 0.00 | 0.00 | NO |
| chr13 | 18400000 | 18500000 | 0.00 | 0.00 | NO |

| | | | | | |
|---|---|---|---|---|---|
| chr13 | 18500000 | 18600000 | 0.00 | 0.00 | NO |
| chr13 | 18600000 | 18700000 | 0.00 | 0.00 | NO |
| chr13 | 18700000 | 18800000 | 0.00 | 0.00 | NO |
| chr13 | 18800000 | 18900000 | 0.00 | 0.00 | NO |
| chr13 | 18900000 | 19000000 | 0.00 | 0.00 | NO |
| chr13 | 19000000 | 19100000 | 0.00 | 0.00 | NO |
| chr13 | 19100000 | 19200000 | 0.00 | 0.00 | NO |
| chr13 | 19200000 | 19300000 | 0.00 | 0.00 | NO |
| chr13 | 19300000 | 19400000 | 0.00 | 0.00 | NO |
| chr13 | 19400000 | 19500000 | 0.00 | 0.00 | NO |
| chr13 | 19500000 | 19600000 | 0.00 | -1.68 | NO |
| chr13 | 19600000 | 19700000 | 0.00 | -1.34 | NO |
| chr13 | 19700000 | 19800000 | 0.00 | 0.00 | NO |
| chr13 | 19800000 | 19900000 | 0.00 | 0.00 | NO |
| chr13 | 19900000 | 20000000 | 0.00 | 0.00 | NO |
| chr13 | 20000000 | 20100000 | 0.00 | 0.00 | NO |
| chr13 | 20100000 | 20200000 | 0.00 | 0.00 | NO |
| chr13 | 20200000 | 20300000 | 0.80 | 0.59 | A>A |
| chr13 | 20300000 | 20400000 | 0.92 | 0.80 | A>A |
| chr13 | 20400000 | 20500000 | 0.86 | 0.84 | A>A |
| chr13 | 20500000 | 20600000 | 0.95 | 0.77 | A>A |
| chr13 | 20600000 | 20700000 | 0.98 | 0.86 | A>A |
| chr13 | 20700000 | 20800000 | 0.75 | 0.80 | A>A |
| chr13 | 20800000 | 20900000 | 0.77 | 0.80 | A>A |
| chr13 | 20900000 | 21000000 | 0.74 | 0.77 | A>A |
| chr13 | 21000000 | 21100000 | 0.60 | 0.74 | A>A |
| chr13 | 21100000 | 21200000 | 0.89 | 0.81 | A>A |
| chr13 | 21200000 | 21300000 | 0.74 | 0.82 | A>A |
| chr13 | 21300000 | 21400000 | 0.93 | 0.82 | A>A |
| chr13 | 21400000 | 21500000 | 0.83 | 0.76 | A>A |
| chr13 | 21500000 | 21600000 | 0.84 | 0.74 | A>A |
| chr13 | 21600000 | 21700000 | 0.98 | 0.91 | A>A |
| chr13 | 21700000 | 21800000 | 0.76 | 0.70 | A>A |
| chr13 | 21800000 | 21900000 | 0.00 | 0.66 | NO |
| chr13 | 21900000 | 22000000 | 0.76 | 0.71 | A>A |
| chr13 | 22000000 | 22100000 | 0.63 | 0.59 | A>A |
| chr13 | 22100000 | 22200000 | 0.34 | 0.48 | A>A |
| chr13 | 22200000 | 22300000 | 0.02 | 0.06 | A>A |
| chr13 | 22300000 | 22400000 | -0.31 | -0.25 | B>B |
| chr13 | 22400000 | 22500000 | -0.58 | -0.26 | B>B |
| chr13 | 22500000 | 22600000 | -0.73 | -0.31 | B>B |
| chr13 | 22600000 | 22700000 | -0.85 | -0.44 | B>B |
| chr13 | 22700000 | 22800000 | -0.94 | -0.45 | B>B |
| chr13 | 22800000 | 22900000 | -0.94 | -0.51 | B>B |
| chr13 | 22900000 | 23000000 | -0.99 | -0.57 | B>B |
| chr13 | 23000000 | 23100000 | -1.08 | -0.45 | B>A |
| chr13 | 23100000 | 23200000 | -1.02 | -0.37 | B>A |
| chr13 | 23200000 | 23300000 | -0.90 | -0.25 | B>A |
| chr13 | 23300000 | 23400000 | -0.79 | 0.17 | B>A |
| chr13 | 23400000 | 23500000 | -0.75 | 0.29 | B>A |
| chr13 | 23500000 | 23600000 | -0.64 | 0.29 | B>A |
| chr13 | 23600000 | 23700000 | -0.54 | 0.18 | B>A |
| chr13 | 23700000 | 23800000 | -0.27 | 0.17 | B>B |
| chr13 | 23800000 | 23900000 | -0.05 | 0.19 | A>A |
| chr13 | 23900000 | 24000000 | 0.49 | 0.54 | A>A |
| chr13 | 24000000 | 24100000 | 0.54 | 0.61 | A>A |

| | | | | | |
|---|---|---|---|---|---|
| chr13 | 24100000 | 24200000 | 0.65 | 0.59 | A>A |
| chr13 | 24200000 | 24300000 | 0.62 | 0.67 | A>A |
| chr13 | 24300000 | 24400000 | 0.79 | 0.83 | A>A |
| chr13 | 24400000 | 24500000 | 0.63 | 0.69 | A>A |
| chr13 | 24500000 | 24600000 | 0.61 | 0.77 | A>A |
| chr13 | 24600000 | 24700000 | 0.94 | 0.84 | A>A |
| chr13 | 24700000 | 24800000 | 0.78 | 0.87 | A>A |
| chr13 | 24800000 | 24900000 | 0.69 | 0.65 | A>A |
| chr13 | 24900000 | 25000000 | 0.00 | 0.00 | NO |
| chr13 | 25000000 | 25100000 | 0.84 | 0.82 | A>A |
| chr13 | 25100000 | 25200000 | 0.00 | 0.00 | NO |
| chr13 | 25200000 | 25300000 | 0.73 | 0.50 | A>A |
| chr13 | 25300000 | 25400000 | 0.74 | 0.46 | A>A |
| chr13 | 25400000 | 25500000 | 0.88 | 0.49 | A>A |
| chr13 | 25500000 | 25600000 | 0.00 | 0.00 | NO |
| chr13 | 25600000 | 25700000 | 0.65 | 0.18 | A>A |
| chr13 | 25700000 | 25800000 | 0.57 | 0.17 | A>A |
| chr13 | 25800000 | 25900000 | 0.74 | 0.43 | A>A |
| chr13 | 25900000 | 26000000 | 0.42 | -0.09 | A>A |
| chr13 | 26000000 | 26100000 | 0.05 | -0.46 | B>B |
| chr13 | 26100000 | 26200000 | -0.16 | -0.90 | A>B |
| chr13 | 26200000 | 26300000 | -0.21 | -1.39 | A>B |
| chr13 | 26300000 | 26400000 | -0.16 | -1.31 | A>B |
| chr13 | 26400000 | 26500000 | -0.09 | -1.32 | A>B |
| chr13 | 26500000 | 26600000 | 0.01 | -1.17 | A>B |
| chr13 | 26600000 | 26700000 | 0.05 | -0.58 | A>B |
| chr13 | 26700000 | 26800000 | 0.26 | -0.04 | A>A |
| chr13 | 26800000 | 26900000 | 0.53 | 0.33 | A>A |
| chr13 | 26900000 | 27000000 | 0.38 | 0.13 | A>A |
| chr13 | 27000000 | 27100000 | 0.22 | -0.05 | A>A |
| chr13 | 27100000 | 27200000 | 0.08 | -0.10 | B>B |
| chr13 | 27200000 | 27300000 | 0.08 | -0.11 | B>B |
| chr13 | 27300000 | 27400000 | 0.21 | -0.14 | A>A |
| chr13 | 27400000 | 27500000 | 0.30 | -0.01 | A>A |
| chr13 | 27500000 | 27600000 | 0.62 | 0.41 | A>A |
| chr13 | 27600000 | 27700000 | 0.89 | 0.78 | A>A |
| chr13 | 27700000 | 27800000 | 0.93 | 0.78 | A>A |
| chr13 | 27800000 | 27900000 | 0.83 | 0.54 | A>A |
| chr13 | 27900000 | 28000000 | 0.54 | 0.52 | A>A |
| chr13 | 28000000 | 28100000 | 0.63 | 0.51 | A>A |
| chr13 | 28100000 | 28200000 | 0.61 | 0.51 | A>A |
| chr13 | 28200000 | 28300000 | 0.57 | 0.36 | A>A |
| chr13 | 28300000 | 28400000 | 0.45 | 0.25 | A>A |
| chr13 | 28400000 | 28500000 | 0.41 | 0.33 | A>A |
| chr13 | 28500000 | 28600000 | 0.48 | 0.32 | A>A |
| chr13 | 28600000 | 28700000 | 0.45 | 0.44 | A>A |
| chr13 | 28700000 | 28800000 | 0.67 | 0.54 | A>A |
| chr13 | 28800000 | 28900000 | 0.56 | 0.32 | A>A |
| chr13 | 28900000 | 29000000 | 0.27 | -0.11 | A>A |
| chr13 | 29000000 | 29100000 | 0.27 | -0.16 | A>A |
| chr13 | 29100000 | 29200000 | 0.38 | -0.10 | A>A |
| chr13 | 29200000 | 29300000 | 0.50 | 0.22 | A>A |
| chr13 | 29300000 | 29400000 | 0.36 | 0.09 | A>A |
| chr13 | 29400000 | 29500000 | -0.18 | -0.69 | B>B |
| chr13 | 29500000 | 29600000 | -0.38 | -1.20 | A>B |
| chr13 | 29600000 | 29700000 | -0.49 | -1.29 | A>B |

| | | | | | |
|---|---|---|---|---|---|
| chr13 | 29700000 | 29800000 | -0.39 | -1.33 | A>B |
| chr13 | 29800000 | 29900000 | -0.28 | -1.04 | A>B |
| chr13 | 29900000 | 30000000 | 0.00 | -0.32 | B>B |
| chr13 | 30000000 | 30100000 | 0.33 | 0.12 | A>A |
| chr13 | 30100000 | 30200000 | 0.55 | 0.29 | A>A |
| chr13 | 30200000 | 30300000 | 0.38 | 0.12 | A>A |
| chr13 | 30300000 | 30400000 | 0.61 | 0.42 | A>A |
| chr13 | 30400000 | 30500000 | 0.70 | 0.67 | A>A |
| chr13 | 30500000 | 30600000 | 0.63 | 0.44 | A>A |
| chr13 | 30600000 | 30700000 | 0.57 | 0.39 | A>A |
| chr13 | 30700000 | 30800000 | 0.69 | 0.54 | A>A |
| chr13 | 30800000 | 30900000 | 0.76 | 0.76 | A>A |
| chr13 | 30900000 | 31000000 | 0.79 | 0.72 | A>A |
| chr13 | 31000000 | 31100000 | 0.73 | 0.67 | A>A |
| chr13 | 31100000 | 31200000 | 0.73 | 0.66 | A>A |
| chr13 | 31200000 | 31300000 | 0.76 | 0.66 | A>A |
| chr13 | 31300000 | 31400000 | 0.71 | 0.62 | A>A |
| chr13 | 31400000 | 31500000 | 0.71 | 0.48 | A>A |
| chr13 | 31500000 | 31600000 | 0.68 | 0.36 | A>A |
| chr13 | 31600000 | 31700000 | 0.66 | 0.37 | A>A |
| chr13 | 31700000 | 31800000 | 0.58 | 0.35 | A>A |
| chr13 | 31800000 | 31900000 | 0.10 | -0.09 | A>A |
| chr13 | 31900000 | 32000000 | -0.06 | -0.30 | B>B |
| chr13 | 32000000 | 32100000 | -0.32 | -0.56 | B>B |
| chr13 | 32100000 | 32200000 | -0.44 | -0.85 | B>B |
| chr13 | 32200000 | 32300000 | -0.32 | -0.61 | B>B |
| chr13 | 32300000 | 32400000 | -0.07 | -0.17 | B>B |
| chr13 | 32400000 | 32500000 | 0.20 | 0.15 | A>A |
| chr13 | 32500000 | 32600000 | 0.28 | 0.44 | A>A |
| chr13 | 32600000 | 32700000 | 0.28 | 0.43 | A>A |
| chr13 | 32700000 | 32800000 | 0.35 | 0.37 | A>A |
| chr13 | 32800000 | 32900000 | 0.45 | 0.31 | A>A |
| chr13 | 32900000 | 33000000 | 0.66 | 0.60 | A>A |
| chr13 | 33000000 | 33100000 | 0.82 | 0.66 | A>A |
| chr13 | 33100000 | 33200000 | 0.75 | 0.74 | A>A |
| chr13 | 33200000 | 33300000 | 0.71 | 0.71 | A>A |
| chr13 | 33300000 | 33400000 | 0.80 | 0.76 | A>A |
| chr13 | 33400000 | 33500000 | 0.80 | 0.61 | A>A |
| chr13 | 33500000 | 33600000 | 0.66 | 0.60 | A>A |
| chr13 | 33600000 | 33700000 | 0.68 | 0.62 | A>A |
| chr13 | 33700000 | 33800000 | 0.71 | 0.69 | A>A |
| chr13 | 33800000 | 33900000 | 0.65 | 0.68 | A>A |
| chr13 | 33900000 | 34000000 | 0.72 | 0.60 | A>A |
| chr13 | 34000000 | 34100000 | 0.71 | 0.65 | A>A |
| chr13 | 34100000 | 34200000 | 0.60 | 0.63 | A>A |
| chr13 | 34200000 | 34300000 | 0.71 | 0.70 | A>A |
| chr13 | 34300000 | 34400000 | 0.61 | 0.61 | A>A |
| chr13 | 34400000 | 34500000 | 0.61 | 0.66 | A>A |
| chr13 | 34500000 | 34600000 | 0.67 | 0.72 | A>A |
| chr13 | 34600000 | 34700000 | 0.60 | 0.72 | A>A |
| chr13 | 34700000 | 34800000 | 0.72 | 0.66 | A>A |
| chr13 | 34800000 | 34900000 | 0.70 | 0.76 | A>A |
| chr13 | 34900000 | 35000000 | 0.60 | 0.64 | A>A |
| chr13 | 35000000 | 35100000 | 0.66 | 0.73 | A>A |
| chr13 | 35100000 | 35200000 | 0.66 | 0.67 | A>A |
| chr13 | 35200000 | 35300000 | 0.63 | 0.68 | A>A |

| | | | | | |
|---|---|---|---|---|---|
| chr13 | 35300000 | 35400000 | 0.78 | 0.77 | A>A |
| chr13 | 35400000 | 35500000 | 0.72 | 0.76 | A>A |
| chr13 | 35500000 | 35600000 | 0.64 | 0.71 | A>A |
| chr13 | 35600000 | 35700000 | 0.00 | 0.77 | NO |
| chr13 | 35700000 | 35800000 | 0.46 | 0.57 | A>A |
| chr13 | 35800000 | 35900000 | 0.33 | 0.64 | A>A |
| chr13 | 35900000 | 36000000 | 0.26 | 0.55 | A>A |
| chr13 | 36000000 | 36100000 | 0.34 | 0.59 | A>A |
| chr13 | 36100000 | 36200000 | 0.39 | 0.61 | A>A |
| chr13 | 36200000 | 36300000 | 0.42 | 0.51 | A>A |
| chr13 | 36300000 | 36400000 | 0.36 | 0.52 | A>A |
| chr13 | 36400000 | 36500000 | 0.24 | 0.43 | A>A |
| chr13 | 36500000 | 36600000 | -0.02 | 0.03 | A>A |
| chr13 | 36600000 | 36700000 | -0.27 | -0.53 | B>B |
| chr13 | 36700000 | 36800000 | -0.27 | -0.51 | B>B |
| chr13 | 36800000 | 36900000 | 0.08 | -0.06 | A>A |
| chr13 | 36900000 | 37000000 | 0.21 | -0.26 | B>B |
| chr13 | 37000000 | 37100000 | -0.34 | -0.81 | B>B |
| chr13 | 37100000 | 37200000 | -0.43 | -1.15 | A>B |
| chr13 | 37200000 | 37300000 | -0.29 | -0.99 | A>B |
| chr13 | 37300000 | 37400000 | 0.05 | -0.55 | A>B |
| chr13 | 37400000 | 37500000 | 0.34 | -0.14 | A>A |
| chr13 | 37500000 | 37600000 | 0.53 | 0.19 | A>A |
| chr13 | 37600000 | 37700000 | 0.06 | -0.48 | B>B |
| chr13 | 37700000 | 37800000 | -0.44 | -1.16 | A>B |
| chr13 | 37800000 | 37900000 | -0.66 | -1.25 | A>B |
| chr13 | 37900000 | 38000000 | -0.53 | -1.26 | A>B |
| chr13 | 38000000 | 38100000 | -0.60 | -1.28 | A>B |
| chr13 | 38100000 | 38200000 | -0.52 | -1.28 | A>B |
| chr13 | 38200000 | 38300000 | -0.47 | -1.20 | A>B |
| chr13 | 38300000 | 38400000 | -0.61 | -1.27 | A>B |
| chr13 | 38400000 | 38500000 | -0.63 | -1.25 | A>B |
| chr13 | 38500000 | 38600000 | -0.75 | -1.23 | B>B |
| chr13 | 38600000 | 38700000 | -0.86 | -1.12 | B>B |
| chr13 | 38700000 | 38800000 | -0.91 | -1.22 | B>B |
| chr13 | 38800000 | 38900000 | -0.63 | -1.06 | B>B |
| chr13 | 38900000 | 39000000 | -0.22 | -0.72 | B>B |
| chr13 | 39000000 | 39100000 | -0.50 | -1.12 | A>B |
| chr13 | 39100000 | 39200000 | -0.44 | -1.10 | A>B |
| chr13 | 39200000 | 39300000 | -0.36 | -1.18 | A>B |
| chr13 | 39300000 | 39400000 | -0.53 | -1.12 | A>B |
| chr13 | 39400000 | 39500000 | -0.30 | -0.82 | B>B |
| chr13 | 39500000 | 39600000 | 0.02 | -0.34 | B>B |
| chr13 | 39600000 | 39700000 | 0.14 | 0.05 | A>A |
| chr13 | 39700000 | 39800000 | -0.09 | -0.08 | B>B |
| chr13 | 39800000 | 39900000 | 0.01 | 0.16 | A>A |
| chr13 | 39900000 | 40000000 | 0.16 | 0.23 | A>A |
| chr13 | 40000000 | 40100000 | 0.19 | 0.32 | A>A |
| chr13 | 40100000 | 40200000 | 0.36 | 0.32 | A>A |
| chr13 | 40200000 | 40300000 | 0.36 | 0.47 | A>A |
| chr13 | 40300000 | 40400000 | 0.20 | 0.49 | A>A |
| chr13 | 40400000 | 40500000 | 0.15 | 0.46 | A>A |
| chr13 | 40500000 | 40600000 | 0.34 | 0.55 | A>A |
| chr13 | 40600000 | 40700000 | 0.58 | 0.63 | A>A |
| chr13 | 40700000 | 40800000 | 0.53 | 0.64 | A>A |
| chr13 | 40800000 | 40900000 | 0.52 | 0.59 | A>A |

| | | | | | |
|---|---|---|---|---|---|
| chr13 | 40900000 | 41000000 | 0.57 | 0.65 | A>A |
| chr13 | 41000000 | 41100000 | 0.62 | 0.73 | A>A |
| chr13 | 41100000 | 41200000 | 0.71 | 0.67 | A>A |
| chr13 | 41200000 | 41300000 | 0.49 | 0.65 | A>A |
| chr13 | 41300000 | 41400000 | 0.63 | 0.64 | A>A |
| chr13 | 41400000 | 41500000 | 0.00 | 0.00 | NO |
| chr13 | 41500000 | 41600000 | 0.94 | 0.95 | A>A |
| chr13 | 41600000 | 41700000 | 0.88 | 0.91 | A>A |
| chr13 | 41700000 | 41800000 | 0.97 | 0.96 | A>A |
| chr13 | 41800000 | 41900000 | 0.96 | 0.85 | A>A |
| chr13 | 41900000 | 42000000 | 0.79 | 0.69 | A>A |
| chr13 | 42000000 | 42100000 | 0.96 | 0.85 | A>A |
| chr13 | 42100000 | 42200000 | 0.82 | 0.82 | A>A |
| chr13 | 42200000 | 42300000 | 0.74 | 0.75 | A>A |
| chr13 | 42300000 | 42400000 | 0.91 | 0.85 | A>A |
| chr13 | 42400000 | 42500000 | 0.82 | 0.74 | A>A |
| chr13 | 42500000 | 42600000 | 0.81 | 0.80 | A>A |
| chr13 | 42600000 | 42700000 | 0.74 | 0.83 | A>A |
| chr13 | 42700000 | 42800000 | 0.68 | 0.70 | A>A |
| chr13 | 42800000 | 42900000 | 0.68 | 0.72 | A>A |
| chr13 | 42900000 | 43000000 | 0.40 | 0.48 | A>A |
| chr13 | 43000000 | 43100000 | 0.37 | 0.45 | A>A |
| chr13 | 43100000 | 43200000 | 0.22 | 0.50 | A>A |
| chr13 | 43200000 | 43300000 | 0.32 | 0.45 | A>A |
| chr13 | 43300000 | 43400000 | 0.33 | 0.44 | A>A |
| chr13 | 43400000 | 43500000 | 0.27 | 0.53 | A>A |
| chr13 | 43500000 | 43600000 | 0.50 | 0.60 | A>A |
| chr13 | 43600000 | 43700000 | 0.43 | 0.59 | A>A |
| chr13 | 43700000 | 43800000 | 0.51 | 0.58 | A>A |
| chr13 | 43800000 | 43900000 | 0.54 | 0.55 | A>A |
| chr13 | 43900000 | 44000000 | 0.32 | 0.59 | A>A |
| chr13 | 44000000 | 44100000 | 0.34 | 0.60 | A>A |
| chr13 | 44100000 | 44200000 | 0.34 | 0.65 | A>A |
| chr13 | 44200000 | 44300000 | 0.32 | 0.62 | A>A |
| chr13 | 44300000 | 44400000 | 0.43 | 0.58 | A>A |
| chr13 | 44400000 | 44500000 | 0.54 | 0.54 | A>A |
| chr13 | 44500000 | 44600000 | 0.74 | 0.66 | A>A |
| chr13 | 44600000 | 44700000 | 0.82 | 0.70 | A>A |
| chr13 | 44700000 | 44800000 | 0.76 | 0.73 | A>A |
| chr13 | 44800000 | 44900000 | 0.77 | 0.75 | A>A |
| chr13 | 44900000 | 45000000 | 0.88 | 0.74 | A>A |
| chr13 | 45000000 | 45100000 | 0.59 | 0.60 | A>A |
| chr13 | 45100000 | 45200000 | 0.76 | 0.73 | A>A |
| chr13 | 45200000 | 45300000 | 0.82 | 0.72 | A>A |
| chr13 | 45300000 | 45400000 | 0.83 | 0.78 | A>A |
| chr13 | 45400000 | 45500000 | 0.74 | 0.68 | A>A |
| chr13 | 45500000 | 45600000 | 0.81 | 0.70 | A>A |
| chr13 | 45600000 | 45700000 | 0.97 | 0.78 | A>A |
| chr13 | 45700000 | 45800000 | 1.02 | 0.89 | A>A |
| chr13 | 45800000 | 45900000 | 0.85 | 0.79 | A>A |
| chr13 | 45900000 | 46000000 | 0.91 | 0.83 | A>A |
| chr13 | 46000000 | 46100000 | 0.93 | 0.72 | A>A |
| chr13 | 46100000 | 46200000 | 0.74 | 0.48 | A>A |
| chr13 | 46200000 | 46300000 | 0.57 | 0.30 | A>A |
| chr13 | 46300000 | 46400000 | 0.46 | 0.24 | A>A |
| chr13 | 46400000 | 46500000 | 0.61 | 0.28 | A>A |

| | | | | | |
|---|---|---|---|---|---|
| chr13 | 46500000 | 46600000 | 0.76 | 0.45 | A>A |
| chr13 | 46600000 | 46700000 | 0.77 | 0.49 | A>A |
| chr13 | 46700000 | 46800000 | 0.51 | 0.38 | A>A |
| chr13 | 46800000 | 46900000 | 0.48 | 0.22 | A>A |
| chr13 | 46900000 | 47000000 | 0.52 | 0.28 | A>A |
| chr13 | 47000000 | 47100000 | 0.00 | 0.39 | NO |
| chr13 | 47100000 | 47200000 | 0.82 | 0.69 | A>A |
| chr13 | 47200000 | 47300000 | 0.76 | 0.74 | A>A |
| chr13 | 47300000 | 47400000 | 0.34 | 0.20 | A>A |
| chr13 | 47400000 | 47500000 | -0.05 | -0.51 | B>B |
| chr13 | 47500000 | 47600000 | -0.38 | -0.84 | B>B |
| chr13 | 47600000 | 47700000 | -0.56 | -1.08 | B>B |
| chr13 | 47700000 | 47800000 | -0.55 | -1.10 | B>B |
| chr13 | 47800000 | 47900000 | -0.71 | -1.12 | B>B |
| chr13 | 47900000 | 48000000 | -0.84 | -1.14 | B>B |
| chr13 | 48000000 | 48100000 | -0.81 | -1.18 | B>B |
| chr13 | 48100000 | 48200000 | -0.65 | -1.03 | B>B |
| chr13 | 48200000 | 48300000 | -0.47 | -0.91 | B>B |
| chr13 | 48300000 | 48400000 | -0.27 | -0.49 | B>B |
| chr13 | 48400000 | 48500000 | 0.11 | -0.06 | A>A |
| chr13 | 48500000 | 48600000 | 0.41 | 0.37 | A>A |
| chr13 | 48600000 | 48700000 | 0.53 | 0.49 | A>A |
| chr13 | 48700000 | 48800000 | 0.59 | 0.74 | A>A |
| chr13 | 48800000 | 48900000 | 0.64 | 0.73 | A>A |
| chr13 | 48900000 | 49000000 | 0.50 | 0.67 | A>A |
| chr13 | 49000000 | 49100000 | 0.49 | 0.65 | A>A |
| chr13 | 49100000 | 49200000 | 0.34 | 0.48 | A>A |
| chr13 | 49200000 | 49300000 | 0.14 | 0.36 | A>A |
| chr13 | 49300000 | 49400000 | 0.32 | 0.54 | A>A |
| chr13 | 49400000 | 49500000 | 0.55 | 0.69 | A>A |
| chr13 | 49500000 | 49600000 | 0.70 | 0.72 | A>A |
| chr13 | 49600000 | 49700000 | 0.75 | 0.79 | A>A |
| chr13 | 49700000 | 49800000 | 0.76 | 0.76 | A>A |
| chr13 | 49800000 | 49900000 | 0.76 | 0.79 | A>A |
| chr13 | 49900000 | 50000000 | 0.87 | 0.86 | A>A |
| chr13 | 50000000 | 50100000 | 0.98 | 0.86 | A>A |
| chr13 | 50100000 | 50200000 | 0.86 | 0.82 | A>A |
| chr13 | 50200000 | 50300000 | 0.88 | 0.84 | A>A |
| chr13 | 50300000 | 50400000 | 0.93 | 0.99 | A>A |
| chr13 | 50400000 | 50500000 | 0.89 | 0.84 | A>A |
| chr13 | 50500000 | 50600000 | 0.77 | 0.84 | A>A |
| chr13 | 50600000 | 50700000 | 0.76 | 0.86 | A>A |
| chr13 | 50700000 | 50800000 | 0.73 | 0.82 | A>A |
| chr13 | 50800000 | 50900000 | 0.80 | 0.94 | A>A |
| chr13 | 50900000 | 51000000 | 0.69 | 0.88 | A>A |
| chr13 | 51000000 | 51100000 | 0.70 | 0.89 | A>A |
| chr13 | 51100000 | 51200000 | 0.51 | 0.84 | A>A |
| chr13 | 51200000 | 51300000 | 0.52 | 0.80 | A>A |
| chr13 | 51300000 | 51400000 | 0.45 | 0.73 | A>A |
| chr13 | 51400000 | 51500000 | 0.56 | 0.71 | A>A |
| chr13 | 51500000 | 51600000 | 0.75 | 0.88 | A>A |
| chr13 | 51600000 | 51700000 | 0.79 | 0.75 | A>A |
| chr13 | 51700000 | 51800000 | 0.76 | 0.78 | A>A |
| chr13 | 51800000 | 51900000 | 0.88 | 0.83 | A>A |
| chr13 | 51900000 | 52000000 | 0.98 | 0.81 | A>A |
| chr13 | 52000000 | 52100000 | 1.02 | 0.95 | A>A |

| | | | | | |
|---|---|---|---|---|---|
| chr13 | 52100000 | 52200000 | 1.07 | 0.97 | A>A |
| chr13 | 52200000 | 52300000 | 1.11 | 0.94 | A>A |
| chr13 | 52300000 | 52400000 | 0.84 | 0.77 | A>A |
| chr13 | 52400000 | 52500000 | 0.84 | 0.81 | A>A |
| chr13 | 52500000 | 52600000 | 0.88 | 0.82 | A>A |
| chr13 | 52600000 | 52700000 | 0.80 | 0.75 | A>A |
| chr13 | 52700000 | 52800000 | 0.90 | 0.80 | A>A |
| chr13 | 52800000 | 52900000 | 0.00 | 0.00 | NO |
| chr13 | 52900000 | 53000000 | 0.94 | 0.71 | A>A |
| chr13 | 53000000 | 53100000 | 0.00 | 0.00 | NO |
| chr13 | 53100000 | 53200000 | 0.00 | 0.00 | NO |
| chr13 | 53200000 | 53300000 | 0.63 | 0.11 | A>A |
| chr13 | 53300000 | 53400000 | 0.28 | -0.33 | A>B |
| chr13 | 53400000 | 53500000 | 0.02 | -0.59 | A>B |
| chr13 | 53500000 | 53600000 | -0.29 | -0.89 | A>B |
| chr13 | 53600000 | 53700000 | -0.55 | -1.19 | A>B |
| chr13 | 53700000 | 53800000 | -0.66 | -1.10 | B>B |
| chr13 | 53800000 | 53900000 | -0.91 | -1.04 | B>B |
| chr13 | 53900000 | 54000000 | -1.11 | -1.18 | B>B |
| chr13 | 54000000 | 54100000 | -1.18 | -1.11 | B>B |
| chr13 | 54100000 | 54200000 | -1.11 | -1.17 | B>B |
| chr13 | 54200000 | 54300000 | -1.08 | -1.11 | B>B |
| chr13 | 54300000 | 54400000 | -0.95 | -1.07 | B>B |
| chr13 | 54400000 | 54500000 | -1.01 | -1.07 | B>B |
| chr13 | 54500000 | 54600000 | -1.06 | -1.09 | B>B |
| chr13 | 54600000 | 54700000 | -0.99 | -1.07 | B>B |
| chr13 | 54700000 | 54800000 | -1.00 | -1.08 | B>B |
| chr13 | 54800000 | 54900000 | -0.96 | -1.00 | B>B |
| chr13 | 54900000 | 55000000 | -0.97 | -0.96 | B>B |
| chr13 | 55000000 | 55100000 | -0.91 | -1.05 | B>B |
| chr13 | 55100000 | 55200000 | -0.95 | -1.00 | B>B |
| chr13 | 55200000 | 55300000 | -1.00 | -1.03 | B>B |
| chr13 | 55300000 | 55400000 | -0.91 | -1.01 | B>B |
| chr13 | 55400000 | 55500000 | -0.94 | -0.98 | B>B |
| chr13 | 55500000 | 55600000 | -0.96 | -1.02 | B>B |
| chr13 | 55600000 | 55700000 | -0.93 | -1.02 | B>B |
| chr13 | 55700000 | 55800000 | -0.96 | -1.06 | B>B |
| chr13 | 55800000 | 55900000 | -0.94 | -1.05 | B>B |
| chr13 | 55900000 | 56000000 | -0.95 | -1.01 | B>B |
| chr13 | 56000000 | 56100000 | -0.96 | -1.00 | B>B |
| chr13 | 56100000 | 56200000 | -0.92 | -1.01 | B>B |
| chr13 | 56200000 | 56300000 | -0.96 | -1.03 | B>B |
| chr13 | 56300000 | 56400000 | -0.94 | -1.03 | B>B |
| chr13 | 56400000 | 56500000 | -0.92 | -1.07 | B>B |
| chr13 | 56500000 | 56600000 | -0.94 | -1.00 | B>B |
| chr13 | 56600000 | 56700000 | -0.94 | -1.05 | B>B |
| chr13 | 56700000 | 56800000 | -0.95 | -1.07 | B>B |
| chr13 | 56800000 | 56900000 | -0.97 | -1.04 | B>B |
| chr13 | 56900000 | 57000000 | -0.90 | -1.05 | B>B |
| chr13 | 57000000 | 57100000 | -0.96 | -1.05 | B>B |
| chr13 | 57100000 | 57200000 | -0.94 | -1.01 | B>B |
| chr13 | 57200000 | 57300000 | -0.90 | -1.07 | B>B |
| chr13 | 57300000 | 57400000 | -0.95 | -1.09 | B>B |
| chr13 | 57400000 | 57500000 | -0.98 | -1.11 | B>B |
| chr13 | 57500000 | 57600000 | -0.97 | -1.06 | B>B |
| chr13 | 57600000 | 57700000 | -0.97 | -1.10 | B>B |

| | | | | | |
|---|---|---|---|---|---|
| chr13 | 57700000 | 57800000 | -1.02 | -1.07 | B>B |
| chr13 | 57800000 | 57900000 | -0.99 | -1.10 | B>B |
| chr13 | 57900000 | 58000000 | -1.08 | -1.07 | B>B |
| chr13 | 58000000 | 58100000 | -1.04 | -1.05 | B>B |
| chr13 | 58100000 | 58200000 | -0.95 | -1.01 | B>B |
| chr13 | 58200000 | 58300000 | -0.96 | -0.98 | B>B |
| chr13 | 58300000 | 58400000 | -1.03 | -1.01 | B>B |
| chr13 | 58400000 | 58500000 | -1.06 | -1.07 | B>B |
| chr13 | 58500000 | 58600000 | -1.05 | -1.05 | B>B |
| chr13 | 58600000 | 58700000 | -1.04 | -1.00 | B>B |
| chr13 | 58700000 | 58800000 | -1.05 | -1.00 | B>B |
| chr13 | 58800000 | 58900000 | -1.01 | -0.99 | B>B |
| chr13 | 58900000 | 59000000 | -1.02 | -1.00 | B>B |
| chr13 | 59000000 | 59100000 | -0.97 | -0.93 | B>B |
| chr13 | 59100000 | 59200000 | -1.07 | -0.98 | B>B |
| chr13 | 59200000 | 59300000 | -1.03 | -1.02 | B>B |
| chr13 | 59300000 | 59400000 | -1.01 | -1.00 | B>B |
| chr13 | 59400000 | 59500000 | -1.04 | -0.96 | B>B |
| chr13 | 59500000 | 59600000 | -0.90 | -0.99 | B>B |
| chr13 | 59600000 | 59700000 | -0.91 | -0.96 | B>B |
| chr13 | 59700000 | 59800000 | -0.77 | -0.99 | B>B |
| chr13 | 59800000 | 59900000 | -0.67 | -0.83 | B>B |
| chr13 | 59900000 | 60000000 | -0.51 | -0.78 | B>B |
| chr13 | 60000000 | 60100000 | -0.36 | -0.61 | B>B |
| chr13 | 60100000 | 60200000 | -0.23 | -0.48 | B>B |
| chr13 | 60200000 | 60300000 | 0.05 | -0.16 | B>B |
| chr13 | 60300000 | 60400000 | 0.33 | 0.12 | A>A |
| chr13 | 60400000 | 60500000 | 0.47 | 0.27 | A>A |
| chr13 | 60500000 | 60600000 | 0.59 | 0.30 | A>A |
| chr13 | 60600000 | 60700000 | 0.58 | 0.42 | A>A |
| chr13 | 60700000 | 60800000 | 0.43 | 0.02 | A>A |
| chr13 | 60800000 | 60900000 | 0.15 | -0.26 | B>B |
| chr13 | 60900000 | 61000000 | 0.24 | -0.04 | A>A |
| chr13 | 61000000 | 61100000 | 0.03 | -0.05 | B>B |
| chr13 | 61100000 | 61200000 | -0.03 | -0.14 | B>B |
| chr13 | 61200000 | 61300000 | -0.63 | -0.57 | B>B |
| chr13 | 61300000 | 61400000 | -0.88 | -0.46 | B>B |
| chr13 | 61400000 | 61500000 | -0.97 | -0.34 | B>A |
| chr13 | 61500000 | 61600000 | -0.90 | -0.33 | B>A |
| chr13 | 61600000 | 61700000 | -1.01 | -0.77 | B>B |
| chr13 | 61700000 | 61800000 | -1.05 | -0.95 | B>B |
| chr13 | 61800000 | 61900000 | -1.04 | -0.95 | B>B |
| chr13 | 61900000 | 62000000 | -0.89 | -0.97 | B>B |
| chr13 | 62000000 | 62100000 | -0.88 | -1.06 | B>B |
| chr13 | 62100000 | 62200000 | -0.87 | -1.15 | B>B |
| chr13 | 62200000 | 62300000 | -1.03 | -1.18 | B>B |
| chr13 | 62300000 | 62400000 | -1.01 | -1.18 | B>B |
| chr13 | 62400000 | 62500000 | -1.00 | -1.17 | B>B |
| chr13 | 62500000 | 62600000 | -0.77 | -1.09 | B>B |
| chr13 | 62600000 | 62700000 | -0.58 | -1.03 | B>B |
| chr13 | 62700000 | 62800000 | -0.51 | -1.02 | B>B |
| chr13 | 62800000 | 62900000 | -0.47 | -0.81 | B>B |
| chr13 | 62900000 | 63000000 | -0.51 | -0.36 | B>B |
| chr13 | 63000000 | 63100000 | -0.47 | 0.03 | B>B |
| chr13 | 63100000 | 63200000 | -0.49 | 0.21 | B>A |
| chr13 | 63200000 | 63300000 | -0.39 | 0.38 | B>A |

| | | | | | |
|---|---|---|---|---|---|
| chr13 | 63300000 | 63400000 | -0.45 | 0.32 | B>A |
| chr13 | 63400000 | 63500000 | -0.51 | 0.21 | B>A |
| chr13 | 63500000 | 63600000 | -0.51 | 0.17 | B>A |
| chr13 | 63600000 | 63700000 | -0.41 | -0.07 | B>B |
| chr13 | 63700000 | 63800000 | -0.36 | -0.09 | B>B |
| chr13 | 63800000 | 63900000 | -0.42 | -0.30 | B>B |
| chr13 | 63900000 | 64000000 | -0.42 | -0.44 | B>B |
| chr13 | 64000000 | 64100000 | -0.39 | -0.50 | B>B |
| chr13 | 64100000 | 64200000 | -0.43 | -0.60 | B>B |
| chr13 | 64200000 | 64300000 | -0.49 | -0.57 | B>B |
| chr13 | 64300000 | 64400000 | 0.00 | 0.00 | NO |
| chr13 | 64400000 | 64500000 | -0.47 | -0.71 | B>B |
| chr13 | 64500000 | 64600000 | -0.55 | -0.82 | B>B |
| chr13 | 64600000 | 64700000 | -0.67 | -0.86 | B>B |
| chr13 | 64700000 | 64800000 | -0.64 | -0.84 | B>B |
| chr13 | 64800000 | 64900000 | -0.56 | -0.77 | B>B |
| chr13 | 64900000 | 65000000 | -0.35 | -0.76 | B>B |
| chr13 | 65000000 | 65100000 | -0.01 | -0.49 | B>B |
| chr13 | 65100000 | 65200000 | 0.23 | -0.32 | B>B |
| chr13 | 65200000 | 65300000 | 0.36 | -0.24 | A>B |
| chr13 | 65300000 | 65400000 | 0.29 | -0.27 | A>A |
| chr13 | 65400000 | 65500000 | 0.25 | -0.34 | A>B |
| chr13 | 65500000 | 65600000 | 0.14 | -0.41 | B>B |
| chr13 | 65600000 | 65700000 | 0.12 | -0.54 | A>B |
| chr13 | 65700000 | 65800000 | 0.06 | -0.68 | A>B |
| chr13 | 65800000 | 65900000 | -0.02 | -0.68 | A>B |
| chr13 | 65900000 | 66000000 | -0.11 | -0.73 | A>B |
| chr13 | 66000000 | 66100000 | 0.03 | -0.69 | A>B |
| chr13 | 66100000 | 66200000 | 0.07 | -0.59 | A>B |
| chr13 | 66200000 | 66300000 | 0.04 | -0.33 | B>B |
| chr13 | 66300000 | 66400000 | 0.18 | -0.23 | B>B |
| chr13 | 66400000 | 66500000 | 0.21 | -0.03 | A>A |
| chr13 | 66500000 | 66600000 | 0.39 | 0.27 | A>A |
| chr13 | 66600000 | 66700000 | 0.47 | 0.46 | A>A |
| chr13 | 66700000 | 66800000 | 0.52 | 0.48 | A>A |
| chr13 | 66800000 | 66900000 | 0.51 | 0.54 | A>A |
| chr13 | 66900000 | 67000000 | 0.61 | 0.59 | A>A |
| chr13 | 67000000 | 67100000 | 0.58 | 0.59 | A>A |
| chr13 | 67100000 | 67200000 | 0.57 | 0.54 | A>A |
| chr13 | 67200000 | 67300000 | 0.46 | 0.48 | A>A |
| chr13 | 67300000 | 67400000 | 0.66 | 0.60 | A>A |
| chr13 | 67400000 | 67500000 | 0.69 | 0.76 | A>A |
| chr13 | 67500000 | 67600000 | 0.68 | 0.64 | A>A |
| chr13 | 67600000 | 67700000 | 0.77 | 0.61 | A>A |
| chr13 | 67700000 | 67800000 | 0.70 | 0.55 | A>A |
| chr13 | 67800000 | 67900000 | -0.14 | -0.38 | B>B |
| chr13 | 67900000 | 68000000 | -0.29 | -0.72 | B>B |
| chr13 | 68000000 | 68100000 | -0.56 | -0.85 | B>B |
| chr13 | 68100000 | 68200000 | -0.79 | -0.88 | B>B |
| chr13 | 68200000 | 68300000 | -0.78 | -0.73 | B>B |
| chr13 | 68300000 | 68400000 | -0.63 | -0.50 | B>B |
| chr13 | 68400000 | 68500000 | -0.56 | -0.32 | B>B |
| chr13 | 68500000 | 68600000 | -0.33 | -0.10 | B>B |
| chr13 | 68600000 | 68700000 | -0.07 | 0.10 | A>A |
| chr13 | 68700000 | 68800000 | -0.31 | 0.00 | B>B |
| chr13 | 68800000 | 68900000 | -0.44 | -0.47 | B>B |

| | | | | | |
|---|---|---|---|---|---|
| chr13 | 68900000 | 69000000 | -0.48 | -0.75 | B>B |
| chr13 | 69000000 | 69100000 | -0.53 | -0.86 | B>B |
| chr13 | 69100000 | 69200000 | -0.53 | -0.90 | B>B |
| chr13 | 69200000 | 69300000 | -0.50 | -0.95 | B>B |
| chr13 | 69300000 | 69400000 | -0.67 | -0.95 | B>B |
| chr13 | 69400000 | 69500000 | -0.69 | -1.00 | B>B |
| chr13 | 69500000 | 69600000 | -0.70 | -1.01 | B>B |
| chr13 | 69600000 | 69700000 | -0.77 | -1.03 | B>B |
| chr13 | 69700000 | 69800000 | -0.71 | -0.99 | B>B |
| chr13 | 69800000 | 69900000 | -0.74 | -0.93 | B>B |
| chr13 | 69900000 | 70000000 | -1.00 | -1.03 | B>B |
| chr13 | 70000000 | 70100000 | -1.08 | -1.08 | B>B |
| chr13 | 70100000 | 70200000 | -1.07 | -1.01 | B>B |
| chr13 | 70200000 | 70300000 | -1.18 | -1.15 | B>B |
| chr13 | 70300000 | 70400000 | -1.14 | -1.16 | B>B |
| chr13 | 70400000 | 70500000 | -1.10 | -1.18 | B>B |
| chr13 | 70500000 | 70600000 | -1.23 | -1.21 | B>B |
| chr13 | 70600000 | 70700000 | -1.15 | -1.18 | B>B |
| chr13 | 70700000 | 70800000 | -1.16 | -1.20 | B>B |
| chr13 | 70800000 | 70900000 | -1.24 | -1.24 | B>B |
| chr13 | 70900000 | 71000000 | -1.20 | -1.21 | B>B |
| chr13 | 71000000 | 71100000 | -1.15 | -1.13 | B>B |
| chr13 | 71100000 | 71200000 | -1.01 | -0.96 | B>B |
| chr13 | 71200000 | 71300000 | -1.12 | -1.04 | B>B |
| chr13 | 71300000 | 71400000 | -1.20 | -1.15 | B>B |
| chr13 | 71400000 | 71500000 | -1.13 | -1.06 | B>B |
| chr13 | 71500000 | 71600000 | -1.08 | -1.03 | B>B |
| chr13 | 71600000 | 71700000 | -1.07 | -0.98 | B>B |
| chr13 | 71700000 | 71800000 | -1.11 | -0.83 | B>B |
| chr13 | 71800000 | 71900000 | -0.98 | -0.40 | B>A |
| chr13 | 71900000 | 72000000 | -0.98 | -0.28 | B>A |
| chr13 | 72000000 | 72100000 | -1.01 | -0.31 | B>A |
| chr13 | 72100000 | 72200000 | -1.02 | -0.62 | B>B |
| chr13 | 72200000 | 72300000 | -1.00 | -0.78 | B>B |
| chr13 | 72300000 | 72400000 | -0.87 | -0.79 | B>B |
| chr13 | 72400000 | 72500000 | -0.75 | -0.76 | B>B |
| chr13 | 72500000 | 72600000 | -0.88 | -0.96 | B>B |
| chr13 | 72600000 | 72700000 | -0.89 | -0.99 | B>B |
| chr13 | 72700000 | 72800000 | -0.75 | -1.02 | B>B |
| chr13 | 72800000 | 72900000 | -0.74 | -1.04 | B>B |
| chr13 | 72900000 | 73000000 | -0.61 | -1.07 | B>B |
| chr13 | 73000000 | 73100000 | -0.53 | -1.05 | B>B |
| chr13 | 73100000 | 73200000 | -0.45 | -0.95 | B>B |
| chr13 | 73200000 | 73300000 | -0.09 | -0.54 | B>B |
| chr13 | 73300000 | 73400000 | 0.76 | 0.39 | A>A |
| chr13 | 73400000 | 73500000 | 0.80 | 0.62 | A>A |
| chr13 | 73500000 | 73600000 | 0.90 | 0.72 | A>A |
| chr13 | 73600000 | 73700000 | 0.94 | 0.70 | A>A |
| chr13 | 73700000 | 73800000 | 1.03 | 0.77 | A>A |
| chr13 | 73800000 | 73900000 | 1.04 | 0.88 | A>A |
| chr13 | 73900000 | 74000000 | 1.00 | 0.77 | A>A |
| chr13 | 74000000 | 74100000 | 0.93 | 0.75 | A>A |
| chr13 | 74100000 | 74200000 | 0.83 | 0.78 | A>A |
| chr13 | 74200000 | 74300000 | 0.70 | 0.50 | A>A |
| chr13 | 74300000 | 74400000 | 0.41 | -0.06 | A>A |
| chr13 | 74400000 | 74500000 | 0.44 | -0.07 | A>A |

| | | | | | |
|---|---|---|---|---|---|
| chr13 | 74500000 | 74600000 | 0.55 | -0.04 | A>B |
| chr13 | 74600000 | 74700000 | 0.58 | 0.03 | A>A |
| chr13 | 74700000 | 74800000 | 0.44 | -0.25 | A>B |
| chr13 | 74800000 | 74900000 | 0.21 | -0.66 | A>B |
| chr13 | 74900000 | 75000000 | 0.13 | -0.92 | A>B |
| chr13 | 75000000 | 75100000 | 0.09 | -0.98 | A>B |
| chr13 | 75100000 | 75200000 | 0.09 | -0.97 | A>B |
| chr13 | 75200000 | 75300000 | 0.05 | -0.98 | A>B |
| chr13 | 75300000 | 75400000 | 0.18 | -1.02 | A>B |
| chr13 | 75400000 | 75500000 | 0.27 | -1.03 | A>B |
| chr13 | 75500000 | 75600000 | 0.23 | -1.02 | A>B |
| chr13 | 75600000 | 75700000 | 0.19 | -1.03 | A>B |
| chr13 | 75700000 | 75800000 | 0.27 | -0.77 | A>B |
| chr13 | 75800000 | 75900000 | 0.43 | -0.03 | A>A |
| chr13 | 75900000 | 76000000 | 0.82 | 0.52 | A>A |
| chr13 | 76000000 | 76100000 | 0.98 | 0.64 | A>A |
| chr13 | 76100000 | 76200000 | 0.69 | 0.45 | A>A |
| chr13 | 76200000 | 76300000 | 0.58 | 0.43 | A>A |
| chr13 | 76300000 | 76400000 | 0.81 | 0.67 | A>A |
| chr13 | 76400000 | 76500000 | 0.31 | 0.09 | A>A |
| chr13 | 76500000 | 76600000 | 0.23 | -0.06 | A>A |
| chr13 | 76600000 | 76700000 | -0.10 | -0.25 | B>B |
| chr13 | 76700000 | 76800000 | -0.22 | -0.30 | B>B |
| chr13 | 76800000 | 76900000 | -0.20 | -0.12 | B>B |
| chr13 | 76900000 | 77000000 | -0.23 | -0.14 | B>B |
| chr13 | 77000000 | 77100000 | -0.16 | -0.29 | B>B |
| chr13 | 77100000 | 77200000 | -0.45 | -0.78 | B>B |
| chr13 | 77200000 | 77300000 | -0.38 | -0.87 | B>B |
| chr13 | 77300000 | 77400000 | -0.23 | -0.80 | B>B |
| chr13 | 77400000 | 77500000 | -0.03 | -0.35 | B>B |
| chr13 | 77500000 | 77600000 | 0.33 | 0.07 | A>A |
| chr13 | 77600000 | 77700000 | 0.72 | 0.40 | A>A |
| chr13 | 77700000 | 77800000 | 0.71 | 0.46 | A>A |
| chr13 | 77800000 | 77900000 | 0.69 | 0.35 | A>A |
| chr13 | 77900000 | 78000000 | 0.35 | -0.17 | A>A |
| chr13 | 78000000 | 78100000 | 0.42 | 0.05 | A>A |
| chr13 | 78100000 | 78200000 | 0.28 | 0.08 | A>A |
| chr13 | 78200000 | 78300000 | -0.01 | -0.14 | B>B |
| chr13 | 78300000 | 78400000 | -0.26 | -0.48 | B>B |
| chr13 | 78400000 | 78500000 | -0.50 | -1.08 | A>B |
| chr13 | 78500000 | 78600000 | -0.56 | -1.15 | A>B |
| chr13 | 78600000 | 78700000 | -0.54 | -1.01 | B>B |
| chr13 | 78700000 | 78800000 | -0.54 | -0.95 | B>B |
| chr13 | 78800000 | 78900000 | -0.58 | -1.16 | B>B |
| chr13 | 78900000 | 79000000 | -0.58 | -1.08 | B>B |
| chr13 | 79000000 | 79100000 | -0.52 | -1.01 | B>B |
| chr13 | 79100000 | 79200000 | -0.13 | -0.59 | B>B |
| chr13 | 79200000 | 79300000 | -0.03 | -0.53 | B>B |
| chr13 | 79300000 | 79400000 | -0.38 | -0.96 | A>B |
| chr13 | 79400000 | 79500000 | -0.53 | -1.03 | B>B |
| chr13 | 79500000 | 79600000 | -0.48 | -0.96 | B>B |
| chr13 | 79600000 | 79700000 | -0.49 | -1.09 | A>B |
| chr13 | 79700000 | 79800000 | -0.43 | -1.09 | A>B |
| chr13 | 79800000 | 79900000 | 0.03 | -0.51 | B>B |
| chr13 | 79900000 | 80000000 | 0.55 | 0.36 | A>A |
| chr13 | 80000000 | 80100000 | 0.59 | 0.64 | A>A |

| | | | | | |
|---|---|---|---|---|---|
| chr13 | 80100000 | 80200000 | 0.26 | 0.51 | A>A |
| chr13 | 80200000 | 80300000 | 0.34 | 0.61 | A>A |
| chr13 | 80300000 | 80400000 | 0.44 | 0.74 | A>A |
| chr13 | 80400000 | 80500000 | 0.58 | 0.87 | A>A |
| chr13 | 80500000 | 80600000 | 0.52 | 0.77 | A>A |
| chr13 | 80600000 | 80700000 | 0.66 | 0.75 | A>A |
| chr13 | 80700000 | 80800000 | 0.68 | 0.85 | A>A |
| chr13 | 80800000 | 80900000 | 0.60 | 0.76 | A>A |
| chr13 | 80900000 | 81000000 | 0.13 | 0.48 | A>A |
| chr13 | 81000000 | 81100000 | 0.07 | 0.38 | A>A |
| chr13 | 81100000 | 81200000 | -0.16 | 0.40 | B>A |
| chr13 | 81200000 | 81300000 | -0.42 | 0.26 | B>A |
| chr13 | 81300000 | 81400000 | -0.59 | 0.33 | B>A |
| chr13 | 81400000 | 81500000 | -0.77 | 0.31 | B>A |
| chr13 | 81500000 | 81600000 | -0.64 | 0.40 | B>A |
| chr13 | 81600000 | 81700000 | -0.60 | 0.47 | B>A |
| chr13 | 81700000 | 81800000 | -0.54 | 0.56 | B>A |
| chr13 | 81800000 | 81900000 | -0.51 | 0.64 | B>A |
| chr13 | 81900000 | 82000000 | -0.61 | 0.63 | B>A |
| chr13 | 82000000 | 82100000 | -0.69 | 0.54 | B>A |
| chr13 | 82100000 | 82200000 | -0.79 | 0.52 | B>A |
| chr13 | 82200000 | 82300000 | -0.73 | 0.59 | B>A |
| chr13 | 82300000 | 82400000 | -0.87 | 0.59 | B>A |
| chr13 | 82400000 | 82500000 | -0.85 | 0.53 | B>A |
| chr13 | 82500000 | 82600000 | -0.84 | 0.65 | B>A |
| chr13 | 82600000 | 82700000 | -0.71 | 0.68 | B>A |
| chr13 | 82700000 | 82800000 | -0.64 | 0.66 | B>A |
| chr13 | 82800000 | 82900000 | -0.59 | 0.58 | B>A |
| chr13 | 82900000 | 83000000 | -0.65 | 0.51 | B>A |
| chr13 | 83000000 | 83100000 | -0.69 | 0.59 | B>A |
| chr13 | 83100000 | 83200000 | -0.62 | 0.65 | B>A |
| chr13 | 83200000 | 83300000 | -0.53 | 0.60 | B>A |
| chr13 | 83300000 | 83400000 | -0.54 | 0.57 | B>A |
| chr13 | 83400000 | 83500000 | -0.46 | 0.65 | B>A |
| chr13 | 83500000 | 83600000 | -0.43 | 0.66 | B>A |
| chr13 | 83600000 | 83700000 | -0.40 | 0.62 | B>A |
| chr13 | 83700000 | 83800000 | -0.41 | 0.64 | B>A |
| chr13 | 83800000 | 83900000 | -0.44 | 0.60 | B>A |
| chr13 | 83900000 | 84000000 | -0.46 | 0.52 | B>A |
| chr13 | 84000000 | 84100000 | -0.47 | 0.67 | B>A |
| chr13 | 84100000 | 84200000 | -0.45 | 0.63 | B>A |
| chr13 | 84200000 | 84300000 | -0.38 | 0.56 | B>A |
| chr13 | 84300000 | 84400000 | -0.50 | 0.51 | B>A |
| chr13 | 84400000 | 84500000 | -0.54 | 0.52 | B>A |
| chr13 | 84500000 | 84600000 | -0.69 | 0.44 | B>A |
| chr13 | 84600000 | 84700000 | -0.83 | 0.52 | B>A |
| chr13 | 84700000 | 84800000 | -0.95 | 0.48 | B>A |
| chr13 | 84800000 | 84900000 | -1.01 | 0.51 | B>A |
| chr13 | 84900000 | 85000000 | -1.09 | 0.47 | B>A |
| chr13 | 85000000 | 85100000 | -1.16 | 0.41 | B>A |
| chr13 | 85100000 | 85200000 | -1.19 | 0.41 | B>A |
| chr13 | 85200000 | 85300000 | -1.14 | 0.31 | B>A |
| chr13 | 85300000 | 85400000 | -1.22 | -0.01 | B>A |
| chr13 | 85400000 | 85500000 | -0.80 | -0.39 | B>B |
| chr13 | 85500000 | 85600000 | -0.65 | -0.85 | B>B |
| chr13 | 85600000 | 85700000 | -0.57 | -0.98 | B>B |

| | | | | | |
|---|---|---|---|---|---|
| chr13 | 85700000 | 85800000 | -0.39 | -0.99 | A>B |
| chr13 | 85800000 | 85900000 | -0.27 | -0.99 | A>B |
| chr13 | 85900000 | 86000000 | -0.18 | -0.95 | A>B |
| chr13 | 86000000 | 86100000 | -0.12 | -0.96 | A>B |
| chr13 | 86100000 | 86200000 | -0.08 | -0.97 | A>B |
| chr13 | 86200000 | 86300000 | -0.02 | -0.94 | A>B |
| chr13 | 86300000 | 86400000 | -0.27 | -0.80 | B>B |
| chr13 | 86400000 | 86500000 | -1.02 | -1.06 | B>B |
| chr13 | 86500000 | 86600000 | -1.17 | -1.19 | B>B |
| chr13 | 86600000 | 86700000 | -1.18 | -1.22 | B>B |
| chr13 | 86700000 | 86800000 | -1.10 | -1.14 | B>B |
| chr13 | 86800000 | 86900000 | 0.00 | 0.00 | NO |
| chr13 | 86900000 | 87000000 | -1.10 | -1.10 | B>B |
| chr13 | 87000000 | 87100000 | -1.16 | -1.17 | B>B |
| chr13 | 87100000 | 87200000 | -1.11 | -1.15 | B>B |
| chr13 | 87200000 | 87300000 | -1.10 | -1.14 | B>B |
| chr13 | 87300000 | 87400000 | -1.09 | -1.06 | B>B |
| chr13 | 87400000 | 87500000 | -1.02 | -1.00 | B>B |
| chr13 | 87500000 | 87600000 | -0.90 | -0.90 | B>B |
| chr13 | 87600000 | 87700000 | -0.96 | -0.99 | B>B |
| chr13 | 87700000 | 87800000 | -1.01 | -0.99 | B>B |
| chr13 | 87800000 | 87900000 | -0.91 | -0.92 | B>B |
| chr13 | 87900000 | 88000000 | -0.83 | -0.84 | B>B |
| chr13 | 88000000 | 88100000 | -0.90 | -0.87 | B>B |
| chr13 | 88100000 | 88200000 | -0.85 | -0.93 | B>B |
| chr13 | 88200000 | 88300000 | -0.66 | -0.43 | B>B |
| chr13 | 88300000 | 88400000 | -0.06 | 0.03 | B>B |
| chr13 | 88400000 | 88500000 | -0.04 | 0.04 | A>A |
| chr13 | 88500000 | 88600000 | -0.55 | -0.18 | B>B |
| chr13 | 88600000 | 88700000 | -0.80 | -0.25 | B>A |
| chr13 | 88700000 | 88800000 | -0.80 | -0.26 | B>A |
| chr13 | 88800000 | 88900000 | -0.82 | -0.19 | B>A |
| chr13 | 88900000 | 89000000 | -0.81 | -0.20 | B>A |
| chr13 | 89000000 | 89100000 | -0.79 | -0.13 | B>A |
| chr13 | 89100000 | 89200000 | -0.76 | 0.01 | B>A |
| chr13 | 89200000 | 89300000 | -0.77 | 0.07 | B>A |
| chr13 | 89300000 | 89400000 | -0.81 | 0.11 | B>A |
| chr13 | 89400000 | 89500000 | -0.77 | 0.07 | B>A |
| chr13 | 89500000 | 89600000 | -0.67 | 0.17 | B>A |
| chr13 | 89600000 | 89700000 | -0.49 | 0.12 | B>A |
| chr13 | 89700000 | 89800000 | -0.24 | 0.24 | A>A |
| chr13 | 89800000 | 89900000 | 0.03 | 0.41 | A>A |
| chr13 | 89900000 | 90000000 | 0.17 | 0.44 | A>A |
| chr13 | 90000000 | 90100000 | 0.43 | 0.51 | A>A |
| chr13 | 90100000 | 90200000 | 0.37 | 0.49 | A>A |
| chr13 | 90200000 | 90300000 | 0.38 | 0.39 | A>A |
| chr13 | 90300000 | 90400000 | 0.04 | 0.27 | A>A |
| chr13 | 90400000 | 90500000 | -0.03 | 0.38 | A>A |
| chr13 | 90500000 | 90600000 | -0.05 | 0.30 | A>A |
| chr13 | 90600000 | 90700000 | -0.14 | 0.33 | A>A |
| chr13 | 90700000 | 90800000 | -0.09 | 0.29 | A>A |
| chr13 | 90800000 | 90900000 | -0.04 | 0.36 | A>A |
| chr13 | 90900000 | 91000000 | 0.08 | 0.38 | A>A |
| chr13 | 91000000 | 91100000 | 0.33 | 0.39 | A>A |
| chr13 | 91100000 | 91200000 | 0.17 | 0.43 | A>A |
| chr13 | 91200000 | 91300000 | 0.17 | 0.48 | A>A |

| | | | | | |
|---|---|---|---|---|---|
| chr13 | 91300000 | 91400000 | 0.18 | 0.48 | A>A |
| chr13 | 91400000 | 91500000 | 0.17 | 0.45 | A>A |
| chr13 | 91500000 | 91600000 | 0.24 | 0.33 | A>A |
| chr13 | 91600000 | 91700000 | 0.02 | 0.12 | A>A |
| chr13 | 91700000 | 91800000 | -0.46 | -0.30 | B>B |
| chr13 | 91800000 | 91900000 | -0.52 | -0.46 | B>B |
| chr13 | 91900000 | 92000000 | -0.40 | -0.66 | B>B |
| chr13 | 92000000 | 92100000 | -0.23 | -0.59 | B>B |
| chr13 | 92100000 | 92200000 | -0.75 | -1.29 | B>B |
| chr13 | 92200000 | 92300000 | -0.92 | -1.28 | B>B |
| chr13 | 92300000 | 92400000 | -0.93 | -1.23 | B>B |
| chr13 | 92400000 | 92500000 | -1.00 | -1.21 | B>B |
| chr13 | 92500000 | 92600000 | -1.09 | -1.26 | B>B |
| chr13 | 92600000 | 92700000 | -1.07 | -1.26 | B>B |
| chr13 | 92700000 | 92800000 | -0.99 | -1.23 | B>B |
| chr13 | 92800000 | 92900000 | -0.96 | -1.15 | B>B |
| chr13 | 92900000 | 93000000 | -1.05 | -1.17 | B>B |
| chr13 | 93000000 | 93100000 | -0.96 | -1.18 | B>B |
| chr13 | 93100000 | 93200000 | -1.02 | -1.18 | B>B |
| chr13 | 93200000 | 93300000 | -0.89 | -1.11 | B>B |
| chr13 | 93300000 | 93400000 | -0.85 | -1.11 | B>B |
| chr13 | 93400000 | 93500000 | -0.78 | -0.99 | B>B |
| chr13 | 93500000 | 93600000 | -0.68 | -0.99 | B>B |
| chr13 | 93600000 | 93700000 | -0.49 | -0.93 | B>B |
| chr13 | 93700000 | 93800000 | -0.37 | -0.88 | B>B |
| chr13 | 93800000 | 93900000 | -0.30 | -0.77 | B>B |
| chr13 | 93900000 | 94000000 | -0.31 | -0.58 | B>B |
| chr13 | 94000000 | 94100000 | -0.32 | -0.55 | B>B |
| chr13 | 94100000 | 94200000 | -0.41 | -0.50 | B>B |
| chr13 | 94200000 | 94300000 | -0.38 | -0.47 | B>B |
| chr13 | 94300000 | 94400000 | -0.42 | -0.45 | B>B |
| chr13 | 94400000 | 94500000 | -0.39 | -0.51 | B>B |
| chr13 | 94500000 | 94600000 | -0.32 | -0.46 | B>B |
| chr13 | 94600000 | 94700000 | -0.27 | -0.34 | B>B |
| chr13 | 94700000 | 94800000 | -0.39 | -0.62 | B>B |
| chr13 | 94800000 | 94900000 | -0.33 | -0.61 | B>B |
| chr13 | 94900000 | 95000000 | -0.32 | -0.65 | B>B |
| chr13 | 95000000 | 95100000 | -0.07 | -0.53 | B>B |
| chr13 | 95100000 | 95200000 | 0.28 | -0.42 | A>B |
| chr13 | 95200000 | 95300000 | 0.57 | 0.18 | A>A |
| chr13 | 95300000 | 95400000 | 0.64 | 0.46 | A>A |
| chr13 | 95400000 | 95500000 | 0.88 | 0.50 | A>A |
| chr13 | 95500000 | 95600000 | 0.94 | 0.61 | A>A |
| chr13 | 95600000 | 95700000 | 0.83 | 0.72 | A>A |
| chr13 | 95700000 | 95800000 | 0.87 | 0.82 | A>A |
| chr13 | 95800000 | 95900000 | 0.90 | 0.81 | A>A |
| chr13 | 95900000 | 96000000 | 1.22 | 0.92 | A>A |
| chr13 | 96000000 | 96100000 | 0.85 | 0.67 | A>A |
| chr13 | 96100000 | 96200000 | 0.80 | 0.53 | A>A |
| chr13 | 96200000 | 96300000 | 0.78 | 0.66 | A>A |
| chr13 | 96300000 | 96400000 | 0.85 | 0.66 | A>A |
| chr13 | 96400000 | 96500000 | 0.64 | 0.51 | A>A |
| chr13 | 96500000 | 96600000 | 0.47 | 0.30 | A>A |
| chr13 | 96600000 | 96700000 | 0.24 | -0.04 | A>A |
| chr13 | 96700000 | 96800000 | 0.00 | -0.70 | A>B |
| chr13 | 96800000 | 96900000 | -0.39 | -1.08 | A>B |

| | | | | | |
|---|---|---|---|---|---|
| chr13 | 96900000 | 97000000 | -0.62 | -1.32 | A>B |
| chr13 | 97000000 | 97100000 | -0.63 | -1.28 | A>B |
| chr13 | 97100000 | 97200000 | -0.66 | -1.32 | A>B |
| chr13 | 97200000 | 97300000 | -0.78 | -1.36 | B>B |
| chr13 | 97300000 | 97400000 | -0.75 | -1.31 | B>B |
| chr13 | 97400000 | 97500000 | -0.53 | -1.01 | B>B |
| chr13 | 97500000 | 97600000 | -0.44 | -0.81 | B>B |
| chr13 | 97600000 | 97700000 | -0.03 | -0.10 | B>B |
| chr13 | 97700000 | 97800000 | 0.46 | 0.48 | A>A |
| chr13 | 97800000 | 97900000 | 0.58 | 0.60 | A>A |
| chr13 | 97900000 | 98000000 | 0.69 | 0.60 | A>A |
| chr13 | 98000000 | 98100000 | 0.35 | 0.45 | A>A |
| chr13 | 98100000 | 98200000 | 0.15 | 0.38 | A>A |
| chr13 | 98200000 | 98300000 | 0.17 | 0.34 | A>A |
| chr13 | 98300000 | 98400000 | 0.13 | 0.23 | A>A |
| chr13 | 98400000 | 98500000 | 0.14 | 0.28 | A>A |
| chr13 | 98500000 | 98600000 | 0.55 | 0.53 | A>A |
| chr13 | 98600000 | 98700000 | 0.75 | 0.68 | A>A |
| chr13 | 98700000 | 98800000 | 0.78 | 0.78 | A>A |
| chr13 | 98800000 | 98900000 | 0.89 | 0.87 | A>A |
| chr13 | 98900000 | 99000000 | 0.80 | 0.92 | A>A |
| chr13 | 99000000 | 99100000 | 0.73 | 0.79 | A>A |
| chr13 | 99100000 | 99200000 | 1.00 | 0.97 | A>A |
| chr13 | 99200000 | 99300000 | 1.03 | 0.94 | A>A |
| chr13 | 99300000 | 99400000 | 0.81 | 0.83 | A>A |
| chr13 | 99400000 | 99500000 | 1.01 | 0.88 | A>A |
| chr13 | 99500000 | 99600000 | 0.93 | 0.96 | A>A |
| chr13 | 99600000 | 99700000 | 1.03 | 1.03 | A>A |
| chr13 | 99700000 | 99800000 | 1.07 | 0.98 | A>A |
| chr13 | 99800000 | 99900000 | 1.06 | 0.98 | A>A |
| chr13 | 99900000 | 100000000 | 0.90 | 0.98 | A>A |
| chr13 | 1E+08 | 100100000 | 0.96 | 0.92 | A>A |
| chr13 | 1E+08 | 100200000 | 0.95 | 0.88 | A>A |
| chr13 | 1E+08 | 100300000 | 0.79 | 0.72 | A>A |
| chr13 | 1E+08 | 100400000 | 0.61 | 0.51 | A>A |
| chr13 | 1E+08 | 100500000 | 0.60 | 0.48 | A>A |
| chr13 | 1.01E+08 | 100600000 | 0.53 | 0.56 | A>A |
| chr13 | 1.01E+08 | 100700000 | 0.94 | 0.80 | A>A |
| chr13 | 1.01E+08 | 100800000 | 0.74 | 0.81 | A>A |
| chr13 | 1.01E+08 | 100900000 | 0.73 | 0.70 | A>A |
| chr13 | 1.01E+08 | 101000000 | 0.70 | 0.73 | A>A |
| chr13 | 1.01E+08 | 101100000 | 0.89 | 0.80 | A>A |
| chr13 | 1.01E+08 | 101200000 | 0.63 | 0.70 | A>A |
| chr13 | 1.01E+08 | 101300000 | 0.48 | 0.43 | A>A |
| chr13 | 1.01E+08 | 101400000 | 0.30 | -0.01 | A>A |
| chr13 | 1.01E+08 | 101500000 | -0.01 | -0.38 | B>B |
| chr13 | 1.02E+08 | 101600000 | -0.10 | -0.58 | B>B |
| chr13 | 1.02E+08 | 101700000 | -0.17 | -0.50 | B>B |
| chr13 | 1.02E+08 | 101800000 | 0.04 | -0.11 | B>B |
| chr13 | 1.02E+08 | 101900000 | 0.28 | 0.09 | A>A |
| chr13 | 1.02E+08 | 102000000 | 0.33 | 0.13 | A>A |
| chr13 | 1.02E+08 | 102100000 | 0.41 | 0.29 | A>A |
| chr13 | 1.02E+08 | 102200000 | 0.29 | 0.22 | A>A |
| chr13 | 1.02E+08 | 102300000 | 0.39 | 0.38 | A>A |
| chr13 | 1.02E+08 | 102400000 | 0.36 | 0.33 | A>A |
| chr13 | 1.02E+08 | 102500000 | -0.07 | -0.17 | B>B |

| | | | | | |
|---|---|---|---|---|---|
| chr13 | 1.03E+08 | 102600000 | -0.30 | -0.39 | B>B |
| chr13 | 1.03E+08 | 102700000 | -0.60 | -0.91 | B>B |
| chr13 | 1.03E+08 | 102800000 | -0.57 | -1.09 | B>B |
| chr13 | 1.03E+08 | 102900000 | -0.50 | -1.14 | A>B |
| chr13 | 1.03E+08 | 103000000 | -0.45 | -1.18 | A>B |
| chr13 | 1.03E+08 | 103100000 | -0.30 | -0.83 | B>B |
| chr13 | 1.03E+08 | 103200000 | -0.45 | -0.85 | B>B |
| chr13 | 1.03E+08 | 103300000 | 0.26 | -0.07 | A>A |
| chr13 | 1.03E+08 | 103400000 | 0.36 | 0.16 | A>A |
| chr13 | 1.03E+08 | 103500000 | 0.42 | 0.16 | A>A |
| chr13 | 1.04E+08 | 103600000 | -0.14 | -0.17 | B>B |
| chr13 | 1.04E+08 | 103700000 | -0.56 | -0.72 | B>B |
| chr13 | 1.04E+08 | 103800000 | -0.61 | -0.82 | B>B |
| chr13 | 1.04E+08 | 103900000 | -0.70 | -0.64 | B>B |
| chr13 | 1.04E+08 | 104000000 | -0.81 | -0.82 | B>B |
| chr13 | 1.04E+08 | 104100000 | -0.61 | -0.71 | B>B |
| chr13 | 1.04E+08 | 104200000 | -0.05 | -0.12 | B>B |
| chr13 | 1.04E+08 | 104300000 | -0.47 | -0.40 | B>B |
| chr13 | 1.04E+08 | 104400000 | -0.79 | -0.56 | B>B |
| chr13 | 1.04E+08 | 104500000 | -0.89 | -0.32 | B>A |
| chr13 | 1.05E+08 | 104600000 | -0.87 | 0.05 | B>A |
| chr13 | 1.05E+08 | 104700000 | -1.05 | 0.08 | B>A |
| chr13 | 1.05E+08 | 104800000 | -1.06 | 0.12 | B>A |
| chr13 | 1.05E+08 | 104900000 | -1.05 | 0.08 | B>A |
| chr13 | 1.05E+08 | 105000000 | -0.97 | 0.07 | B>A |
| chr13 | 1.05E+08 | 105100000 | -1.04 | 0.09 | B>A |
| chr13 | 1.05E+08 | 105200000 | -0.98 | 0.20 | B>A |
| chr13 | 1.05E+08 | 105300000 | -0.89 | 0.16 | B>A |
| chr13 | 1.05E+08 | 105400000 | -0.92 | 0.26 | B>A |
| chr13 | 1.05E+08 | 105500000 | -0.73 | 0.23 | B>A |
| chr13 | 1.06E+08 | 105600000 | -0.85 | 0.36 | B>A |
| chr13 | 1.06E+08 | 105700000 | -0.62 | 0.37 | B>A |
| chr13 | 1.06E+08 | 105800000 | -0.43 | 0.37 | B>A |
| chr13 | 1.06E+08 | 105900000 | -0.36 | 0.37 | B>A |
| chr13 | 1.06E+08 | 106000000 | -0.33 | 0.44 | B>A |
| chr13 | 1.06E+08 | 106100000 | -0.26 | 0.44 | B>A |
| chr13 | 1.06E+08 | 106200000 | -0.07 | 0.50 | B>A |
| chr13 | 1.06E+08 | 106300000 | -0.05 | 0.53 | B>A |
| chr13 | 1.06E+08 | 106400000 | -0.01 | 0.52 | B>A |
| chr13 | 1.06E+08 | 106500000 | 0.12 | 0.53 | A>A |
| chr13 | 1.07E+08 | 106600000 | 0.27 | 0.55 | A>A |
| chr13 | 1.07E+08 | 106700000 | 0.39 | 0.53 | A>A |
| chr13 | 1.07E+08 | 106800000 | 0.40 | 0.48 | A>A |
| chr13 | 1.07E+08 | 106900000 | 0.40 | 0.51 | A>A |
| chr13 | 1.07E+08 | 107000000 | 0.29 | 0.46 | A>A |
| chr13 | 1.07E+08 | 107100000 | 0.23 | 0.47 | A>A |
| chr13 | 1.07E+08 | 107200000 | 0.13 | 0.31 | A>A |
| chr13 | 1.07E+08 | 107300000 | -0.11 | -0.27 | B>B |
| chr13 | 1.07E+08 | 107400000 | -0.50 | -0.93 | B>B |
| chr13 | 1.07E+08 | 107500000 | -0.68 | -1.19 | B>B |
| chr13 | 1.08E+08 | 107600000 | -0.53 | -0.87 | B>B |
| chr13 | 1.08E+08 | 107700000 | -0.99 | -1.33 | B>B |
| chr13 | 1.08E+08 | 107800000 | -1.09 | -1.30 | B>B |
| chr13 | 1.08E+08 | 107900000 | -0.92 | -1.14 | B>B |
| chr13 | 1.08E+08 | 108000000 | -1.25 | -1.45 | B>B |
| chr13 | 1.08E+08 | 108100000 | -1.52 | -1.54 | B>B |

| | | | | | |
|---|---|---|---|---|---|
| chr13 | 1.08E+08 | 108200000 | -1.48 | -1.54 | B>B |
| chr13 | 1.08E+08 | 108300000 | -1.35 | -1.52 | B>B |
| chr13 | 1.08E+08 | 108400000 | -1.22 | -1.54 | B>B |
| chr13 | 1.08E+08 | 108500000 | -0.98 | -1.43 | B>B |
| chr13 | 1.09E+08 | 108600000 | -0.97 | -1.28 | B>B |
| chr13 | 1.09E+08 | 108700000 | -1.21 | -1.42 | B>B |
| chr13 | 1.09E+08 | 108800000 | -0.86 | -1.19 | B>B |
| chr13 | 1.09E+08 | 108900000 | -0.08 | -0.22 | B>B |
| chr13 | 1.09E+08 | 109000000 | 0.36 | 0.38 | A>A |
| chr13 | 1.09E+08 | 109100000 | 0.51 | 0.40 | A>A |
| chr13 | 1.09E+08 | 109200000 | 0.45 | 0.54 | A>A |
| chr13 | 1.09E+08 | 109300000 | 0.61 | 0.61 | A>A |
| chr13 | 1.09E+08 | 109400000 | 0.73 | 0.79 | A>A |
| chr13 | 1.09E+08 | 109500000 | 0.74 | 0.71 | A>A |
| chr13 | 1.1E+08 | 109600000 | 0.59 | 0.65 | A>A |
| chr13 | 1.1E+08 | 109700000 | 0.76 | 0.70 | A>A |
| chr13 | 1.1E+08 | 109800000 | 0.76 | 0.79 | A>A |
| chr13 | 1.1E+08 | 109900000 | 0.81 | 0.93 | A>A |
| chr13 | 1.1E+08 | 110000000 | 0.88 | 0.83 | A>A |
| chr13 | 1.1E+08 | 110100000 | 0.89 | 0.88 | A>A |
| chr13 | 1.1E+08 | 110200000 | 0.86 | 0.82 | A>A |
| chr13 | 1.1E+08 | 110300000 | 0.87 | 0.96 | A>A |
| chr13 | 1.1E+08 | 110400000 | 0.90 | 0.94 | A>A |
| chr13 | 1.1E+08 | 110500000 | 0.71 | 0.80 | A>A |
| chr13 | 1.11E+08 | 110600000 | 0.90 | 0.94 | A>A |
| chr13 | 1.11E+08 | 110700000 | 0.96 | 0.95 | A>A |
| chr13 | 1.11E+08 | 110800000 | 0.76 | 0.79 | A>A |
| chr13 | 1.11E+08 | 110900000 | 0.88 | 0.89 | A>A |
| chr13 | 1.11E+08 | 111000000 | 1.20 | 1.04 | A>A |
| chr13 | 1.11E+08 | 111100000 | 1.14 | 0.92 | A>A |
| chr13 | 1.11E+08 | 111200000 | 1.16 | 0.98 | A>A |
| chr13 | 1.11E+08 | 111300000 | 0.98 | 0.97 | A>A |
| chr13 | 1.11E+08 | 111400000 | 1.18 | 1.01 | A>A |
| chr13 | 1.11E+08 | 111500000 | 1.10 | 0.93 | A>A |
| chr13 | 1.12E+08 | 111600000 | 1.03 | 0.93 | A>A |
| chr13 | 1.12E+08 | 111700000 | 0.99 | 0.92 | A>A |
| chr13 | 1.12E+08 | 111800000 | 0.96 | 0.76 | A>A |
| chr13 | 1.12E+08 | 111900000 | 0.94 | 0.80 | A>A |
| chr13 | 1.12E+08 | 112000000 | 0.78 | 0.58 | A>A |
| chr13 | 1.12E+08 | 112100000 | 0.52 | 0.36 | A>A |
| chr13 | 1.12E+08 | 112200000 | 0.27 | 0.17 | A>A |
| chr13 | 1.12E+08 | 112300000 | 0.17 | 0.04 | A>A |
| chr13 | 1.12E+08 | 112400000 | 0.00 | 0.02 | NO |
| chr13 | 1.12E+08 | 112500000 | 0.00 | 0.00 | NO |
| chr13 | 1.13E+08 | 112600000 | 0.06 | 0.05 | A>A |
| chr13 | 1.13E+08 | 112700000 | -0.06 | -0.08 | B>B |
| chr13 | 1.13E+08 | 112800000 | 0.18 | -0.04 | A>A |
| chr13 | 1.13E+08 | 112900000 | 0.12 | -0.11 | A>A |
| chr13 | 1.13E+08 | 113000000 | -0.21 | -0.16 | B>B |
| chr13 | 1.13E+08 | 113100000 | 0.09 | 0.04 | A>A |
| chr13 | 1.13E+08 | 113200000 | 0.59 | 0.50 | A>A |
| chr13 | 1.13E+08 | 113300000 | 0.84 | 0.66 | A>A |
| chr13 | 1.13E+08 | 113400000 | 0.97 | 0.84 | A>A |
| chr13 | 1.13E+08 | 113500000 | 0.99 | 0.91 | A>A |
| chr13 | 1.14E+08 | 113600000 | 0.87 | 0.86 | A>A |
| chr13 | 1.14E+08 | 113700000 | 1.03 | 0.86 | A>A |

| | | | | | |
|---|---|---|---|---|---|
| chr13 | 1.14E+08 | 113800000 | 1.05 | 0.87 | A>A |
| chr13 | 1.14E+08 | 113900000 | 1.22 | 0.97 | A>A |
| chr13 | 1.14E+08 | 114000000 | 1.14 | 0.86 | A>A |
| chr13 | 1.14E+08 | 114100000 | 1.14 | 0.96 | A>A |
| chr13 | 1.14E+08 | 114200000 | 1.15 | 0.96 | A>A |
| chr13 | 1.14E+08 | 114300000 | 0.93 | 0.91 | A>A |
| chr13 | 1.14E+08 | 114400000 | 0.00 | 0.00 | NO |
| chr13 | 1.14E+08 | 114500000 | 0.86 | 0.78 | A>A |
| chr13 | 1.15E+08 | 114600000 | 0.96 | 0.76 | A>A |
| chr13 | 1.15E+08 | 114700000 | 0.00 | 0.00 | NO |
| chr13 | 1.15E+08 | 114800000 | 0.00 | 0.00 | NO |
| chr13 | 1.15E+08 | 114900000 | 1.15 | 0.94 | A>A |
| chr13 | 1.15E+08 | 115000000 | 1.06 | 0.84 | A>A |
| chr13 | 1.15E+08 | 115100000 | 1.06 | 0.82 | A>A |
| chr13 | 1.15E+08 | 115169878 | 0.00 | 0.00 | NO |
| chr14 | 0 | 100000 | 0.00 | 0.00 | NO |
| chr14 | 100000 | 200000 | 0.00 | 0.00 | NO |
| chr14 | 200000 | 300000 | 0.00 | 0.00 | NO |
| chr14 | 300000 | 400000 | 0.00 | 0.00 | NO |
| chr14 | 400000 | 500000 | 0.00 | 0.00 | NO |
| chr14 | 500000 | 600000 | 0.00 | 0.00 | NO |
| chr14 | 600000 | 700000 | 0.00 | 0.00 | NO |
| chr14 | 700000 | 800000 | 0.00 | 0.00 | NO |
| chr14 | 800000 | 900000 | 0.00 | 0.00 | NO |
| chr14 | 900000 | 1000000 | 0.00 | 0.00 | NO |
| chr14 | 1000000 | 1100000 | 0.00 | 0.00 | NO |
| chr14 | 1100000 | 1200000 | 0.00 | 0.00 | NO |
| chr14 | 1200000 | 1300000 | 0.00 | 0.00 | NO |
| chr14 | 1300000 | 1400000 | 0.00 | 0.00 | NO |
| chr14 | 1400000 | 1500000 | 0.00 | 0.00 | NO |
| chr14 | 1500000 | 1600000 | 0.00 | 0.00 | NO |
| chr14 | 1600000 | 1700000 | 0.00 | 0.00 | NO |
| chr14 | 1700000 | 1800000 | 0.00 | 0.00 | NO |
| chr14 | 1800000 | 1900000 | 0.00 | 0.00 | NO |
| chr14 | 1900000 | 2000000 | 0.00 | 0.00 | NO |
| chr14 | 2000000 | 2100000 | 0.00 | 0.00 | NO |
| chr14 | 2100000 | 2200000 | 0.00 | 0.00 | NO |
| chr14 | 2200000 | 2300000 | 0.00 | 0.00 | NO |
| chr14 | 2300000 | 2400000 | 0.00 | 0.00 | NO |
| chr14 | 2400000 | 2500000 | 0.00 | 0.00 | NO |
| chr14 | 2500000 | 2600000 | 0.00 | 0.00 | NO |
| chr14 | 2600000 | 2700000 | 0.00 | 0.00 | NO |
| chr14 | 2700000 | 2800000 | 0.00 | 0.00 | NO |
| chr14 | 2800000 | 2900000 | 0.00 | 0.00 | NO |
| chr14 | 2900000 | 3000000 | 0.00 | 0.00 | NO |
| chr14 | 3000000 | 3100000 | 0.00 | 0.00 | NO |
| chr14 | 3100000 | 3200000 | 0.00 | 0.00 | NO |
| chr14 | 3200000 | 3300000 | 0.00 | 0.00 | NO |
| chr14 | 3300000 | 3400000 | 0.00 | 0.00 | NO |
| chr14 | 3400000 | 3500000 | 0.00 | 0.00 | NO |
| chr14 | 3500000 | 3600000 | 0.00 | 0.00 | NO |
| chr14 | 3600000 | 3700000 | 0.00 | 0.00 | NO |
| chr14 | 3700000 | 3800000 | 0.00 | 0.00 | NO |
| chr14 | 3800000 | 3900000 | 0.00 | 0.00 | NO |
| chr14 | 3900000 | 4000000 | 0.00 | 0.00 | NO |
| chr14 | 4000000 | 4100000 | 0.00 | 0.00 | NO |

| | | | | | |
|---|---|---|---|---|---|
| chr14 | 4100000 | 4200000 | 0.00 | 0.00 | NO |
| chr14 | 4200000 | 4300000 | 0.00 | 0.00 | NO |
| chr14 | 4300000 | 4400000 | 0.00 | 0.00 | NO |
| chr14 | 4400000 | 4500000 | 0.00 | 0.00 | NO |
| chr14 | 4500000 | 4600000 | 0.00 | 0.00 | NO |
| chr14 | 4600000 | 4700000 | 0.00 | 0.00 | NO |
| chr14 | 4700000 | 4800000 | 0.00 | 0.00 | NO |
| chr14 | 4800000 | 4900000 | 0.00 | 0.00 | NO |
| chr14 | 4900000 | 5000000 | 0.00 | 0.00 | NO |
| chr14 | 5000000 | 5100000 | 0.00 | 0.00 | NO |
| chr14 | 5100000 | 5200000 | 0.00 | 0.00 | NO |
| chr14 | 5200000 | 5300000 | 0.00 | 0.00 | NO |
| chr14 | 5300000 | 5400000 | 0.00 | 0.00 | NO |
| chr14 | 5400000 | 5500000 | 0.00 | 0.00 | NO |
| chr14 | 5500000 | 5600000 | 0.00 | 0.00 | NO |
| chr14 | 5600000 | 5700000 | 0.00 | 0.00 | NO |
| chr14 | 5700000 | 5800000 | 0.00 | 0.00 | NO |
| chr14 | 5800000 | 5900000 | 0.00 | 0.00 | NO |
| chr14 | 5900000 | 6000000 | 0.00 | 0.00 | NO |
| chr14 | 6000000 | 6100000 | 0.00 | 0.00 | NO |
| chr14 | 6100000 | 6200000 | 0.00 | 0.00 | NO |
| chr14 | 6200000 | 6300000 | 0.00 | 0.00 | NO |
| chr14 | 6300000 | 6400000 | 0.00 | 0.00 | NO |
| chr14 | 6400000 | 6500000 | 0.00 | 0.00 | NO |
| chr14 | 6500000 | 6600000 | 0.00 | 0.00 | NO |
| chr14 | 6600000 | 6700000 | 0.00 | 0.00 | NO |
| chr14 | 6700000 | 6800000 | 0.00 | 0.00 | NO |
| chr14 | 6800000 | 6900000 | 0.00 | 0.00 | NO |
| chr14 | 6900000 | 7000000 | 0.00 | 0.00 | NO |
| chr14 | 7000000 | 7100000 | 0.00 | 0.00 | NO |
| chr14 | 7100000 | 7200000 | 0.00 | 0.00 | NO |
| chr14 | 7200000 | 7300000 | 0.00 | 0.00 | NO |
| chr14 | 7300000 | 7400000 | 0.00 | 0.00 | NO |
| chr14 | 7400000 | 7500000 | 0.00 | 0.00 | NO |
| chr14 | 7500000 | 7600000 | 0.00 | 0.00 | NO |
| chr14 | 7600000 | 7700000 | 0.00 | 0.00 | NO |
| chr14 | 7700000 | 7800000 | 0.00 | 0.00 | NO |
| chr14 | 7800000 | 7900000 | 0.00 | 0.00 | NO |
| chr14 | 7900000 | 8000000 | 0.00 | 0.00 | NO |
| chr14 | 8000000 | 8100000 | 0.00 | 0.00 | NO |
| chr14 | 8100000 | 8200000 | 0.00 | 0.00 | NO |
| chr14 | 8200000 | 8300000 | 0.00 | 0.00 | NO |
| chr14 | 8300000 | 8400000 | 0.00 | 0.00 | NO |
| chr14 | 8400000 | 8500000 | 0.00 | 0.00 | NO |
| chr14 | 8500000 | 8600000 | 0.00 | 0.00 | NO |
| chr14 | 8600000 | 8700000 | 0.00 | 0.00 | NO |
| chr14 | 8700000 | 8800000 | 0.00 | 0.00 | NO |
| chr14 | 8800000 | 8900000 | 0.00 | 0.00 | NO |
| chr14 | 8900000 | 9000000 | 0.00 | 0.00 | NO |
| chr14 | 9000000 | 9100000 | 0.00 | 0.00 | NO |
| chr14 | 9100000 | 9200000 | 0.00 | 0.00 | NO |
| chr14 | 9200000 | 9300000 | 0.00 | 0.00 | NO |
| chr14 | 9300000 | 9400000 | 0.00 | 0.00 | NO |
| chr14 | 9400000 | 9500000 | 0.00 | 0.00 | NO |
| chr14 | 9500000 | 9600000 | 0.00 | 0.00 | NO |
| chr14 | 9600000 | 9700000 | 0.00 | 0.00 | NO |

| | | | | | |
|---|---|---|---|---|---|
| chr14 | 9700000 | 9800000 | 0.00 | 0.00 | NO |
| chr14 | 9800000 | 9900000 | 0.00 | 0.00 | NO |
| chr14 | 9900000 | 10000000 | 0.00 | 0.00 | NO |
| chr14 | 10000000 | 10100000 | 0.00 | 0.00 | NO |
| chr14 | 10100000 | 10200000 | 0.00 | 0.00 | NO |
| chr14 | 10200000 | 10300000 | 0.00 | 0.00 | NO |
| chr14 | 10300000 | 10400000 | 0.00 | 0.00 | NO |
| chr14 | 10400000 | 10500000 | 0.00 | 0.00 | NO |
| chr14 | 10500000 | 10600000 | 0.00 | 0.00 | NO |
| chr14 | 10600000 | 10700000 | 0.00 | 0.00 | NO |
| chr14 | 10700000 | 10800000 | 0.00 | 0.00 | NO |
| chr14 | 10800000 | 10900000 | 0.00 | 0.00 | NO |
| chr14 | 10900000 | 11000000 | 0.00 | 0.00 | NO |
| chr14 | 11000000 | 11100000 | 0.00 | 0.00 | NO |
| chr14 | 11100000 | 11200000 | 0.00 | 0.00 | NO |
| chr14 | 11200000 | 11300000 | 0.00 | 0.00 | NO |
| chr14 | 11300000 | 11400000 | 0.00 | 0.00 | NO |
| chr14 | 11400000 | 11500000 | 0.00 | 0.00 | NO |
| chr14 | 11500000 | 11600000 | 0.00 | 0.00 | NO |
| chr14 | 11600000 | 11700000 | 0.00 | 0.00 | NO |
| chr14 | 11700000 | 11800000 | 0.00 | 0.00 | NO |
| chr14 | 11800000 | 11900000 | 0.00 | 0.00 | NO |
| chr14 | 11900000 | 12000000 | 0.00 | 0.00 | NO |
| chr14 | 12000000 | 12100000 | 0.00 | 0.00 | NO |
| chr14 | 12100000 | 12200000 | 0.00 | 0.00 | NO |
| chr14 | 12200000 | 12300000 | 0.00 | 0.00 | NO |
| chr14 | 12300000 | 12400000 | 0.00 | 0.00 | NO |
| chr14 | 12400000 | 12500000 | 0.00 | 0.00 | NO |
| chr14 | 12500000 | 12600000 | 0.00 | 0.00 | NO |
| chr14 | 12600000 | 12700000 | 0.00 | 0.00 | NO |
| chr14 | 12700000 | 12800000 | 0.00 | 0.00 | NO |
| chr14 | 12800000 | 12900000 | 0.00 | 0.00 | NO |
| chr14 | 12900000 | 13000000 | 0.00 | 0.00 | NO |
| chr14 | 13000000 | 13100000 | 0.00 | 0.00 | NO |
| chr14 | 13100000 | 13200000 | 0.00 | 0.00 | NO |
| chr14 | 13200000 | 13300000 | 0.00 | 0.00 | NO |
| chr14 | 13300000 | 13400000 | 0.00 | 0.00 | NO |
| chr14 | 13400000 | 13500000 | 0.00 | 0.00 | NO |
| chr14 | 13500000 | 13600000 | 0.00 | 0.00 | NO |
| chr14 | 13600000 | 13700000 | 0.00 | 0.00 | NO |
| chr14 | 13700000 | 13800000 | 0.00 | 0.00 | NO |
| chr14 | 13800000 | 13900000 | 0.00 | 0.00 | NO |
| chr14 | 13900000 | 14000000 | 0.00 | 0.00 | NO |
| chr14 | 14000000 | 14100000 | 0.00 | 0.00 | NO |
| chr14 | 14100000 | 14200000 | 0.00 | 0.00 | NO |
| chr14 | 14200000 | 14300000 | 0.00 | 0.00 | NO |
| chr14 | 14300000 | 14400000 | 0.00 | 0.00 | NO |
| chr14 | 14400000 | 14500000 | 0.00 | 0.00 | NO |
| chr14 | 14500000 | 14600000 | 0.00 | 0.00 | NO |
| chr14 | 14600000 | 14700000 | 0.00 | 0.00 | NO |
| chr14 | 14700000 | 14800000 | 0.00 | 0.00 | NO |
| chr14 | 14800000 | 14900000 | 0.00 | 0.00 | NO |
| chr14 | 14900000 | 15000000 | 0.00 | 0.00 | NO |
| chr14 | 15000000 | 15100000 | 0.00 | 0.00 | NO |
| chr14 | 15100000 | 15200000 | 0.00 | 0.00 | NO |
| chr14 | 15200000 | 15300000 | 0.00 | 0.00 | NO |

| | | | | | |
|---|---|---|---|---|---|
| chr14 | 15300000 | 15400000 | 0.00 | 0.00 | NO |
| chr14 | 15400000 | 15500000 | 0.00 | 0.00 | NO |
| chr14 | 15500000 | 15600000 | 0.00 | 0.00 | NO |
| chr14 | 15600000 | 15700000 | 0.00 | 0.00 | NO |
| chr14 | 15700000 | 15800000 | 0.00 | 0.00 | NO |
| chr14 | 15800000 | 15900000 | 0.00 | 0.00 | NO |
| chr14 | 15900000 | 16000000 | 0.00 | 0.00 | NO |
| chr14 | 16000000 | 16100000 | 0.00 | 0.00 | NO |
| chr14 | 16100000 | 16200000 | 0.00 | 0.00 | NO |
| chr14 | 16200000 | 16300000 | 0.00 | 0.00 | NO |
| chr14 | 16300000 | 16400000 | 0.00 | 0.00 | NO |
| chr14 | 16400000 | 16500000 | 0.00 | 0.00 | NO |
| chr14 | 16500000 | 16600000 | 0.00 | 0.00 | NO |
| chr14 | 16600000 | 16700000 | 0.00 | 0.00 | NO |
| chr14 | 16700000 | 16800000 | 0.00 | 0.00 | NO |
| chr14 | 16800000 | 16900000 | 0.00 | 0.00 | NO |
| chr14 | 16900000 | 17000000 | 0.00 | 0.00 | NO |
| chr14 | 17000000 | 17100000 | 0.00 | 0.00 | NO |
| chr14 | 17100000 | 17200000 | 0.00 | 0.00 | NO |
| chr14 | 17200000 | 17300000 | 0.00 | 0.00 | NO |
| chr14 | 17300000 | 17400000 | 0.00 | 0.00 | NO |
| chr14 | 17400000 | 17500000 | 0.00 | 0.00 | NO |
| chr14 | 17500000 | 17600000 | 0.00 | 0.00 | NO |
| chr14 | 17600000 | 17700000 | 0.00 | 0.00 | NO |
| chr14 | 17700000 | 17800000 | 0.00 | 0.00 | NO |
| chr14 | 17800000 | 17900000 | 0.00 | 0.00 | NO |
| chr14 | 17900000 | 18000000 | 0.00 | 0.00 | NO |
| chr14 | 18000000 | 18100000 | 0.00 | 0.00 | NO |
| chr14 | 18100000 | 18200000 | 0.00 | 0.00 | NO |
| chr14 | 18200000 | 18300000 | 0.00 | 0.00 | NO |
| chr14 | 18300000 | 18400000 | 0.00 | 0.00 | NO |
| chr14 | 18400000 | 18500000 | 0.00 | 0.00 | NO |
| chr14 | 18500000 | 18600000 | 0.00 | 0.00 | NO |
| chr14 | 18600000 | 18700000 | 0.00 | 0.00 | NO |
| chr14 | 18700000 | 18800000 | 0.00 | 0.00 | NO |
| chr14 | 18800000 | 18900000 | 0.00 | 0.00 | NO |
| chr14 | 18900000 | 19000000 | 0.00 | 0.00 | NO |
| chr14 | 19000000 | 19100000 | 0.00 | 0.00 | NO |
| chr14 | 19100000 | 19200000 | 0.00 | 0.00 | NO |
| chr14 | 19200000 | 19300000 | 0.00 | 0.00 | NO |
| chr14 | 19300000 | 19400000 | 0.00 | 0.00 | NO |
| chr14 | 19400000 | 19500000 | 0.00 | 0.00 | NO |
| chr14 | 19500000 | 19600000 | 0.00 | 0.00 | NO |
| chr14 | 19600000 | 19700000 | 0.00 | 0.00 | NO |
| chr14 | 19700000 | 19800000 | 0.00 | 0.00 | NO |
| chr14 | 19800000 | 19900000 | 0.00 | 0.00 | NO |
| chr14 | 19900000 | 20000000 | 0.00 | 0.00 | NO |
| chr14 | 20000000 | 20100000 | 0.00 | 0.00 | NO |
| chr14 | 20100000 | 20200000 | 0.00 | 0.00 | NO |
| chr14 | 20200000 | 20300000 | 0.00 | -1.57 | NO |
| chr14 | 20300000 | 20400000 | -1.02 | -1.50 | B>B |
| chr14 | 20400000 | 20500000 | 0.00 | -1.35 | NO |
| chr14 | 20500000 | 20600000 | 0.00 | -1.27 | NO |
| chr14 | 20600000 | 20700000 | -0.37 | -0.82 | B>B |
| chr14 | 20700000 | 20800000 | 0.25 | -0.04 | A>A |
| chr14 | 20800000 | 20900000 | 0.90 | 0.58 | A>A |

| | | | | | |
|---|---|---|---|---|---|
| chr14 | 20900000 | 21000000 | 0.77 | 0.48 | A>A |
| chr14 | 21000000 | 21100000 | 0.63 | 0.30 | A>A |
| chr14 | 21100000 | 21200000 | 0.69 | 0.32 | A>A |
| chr14 | 21200000 | 21300000 | 0.51 | 0.02 | A>A |
| chr14 | 21300000 | 21400000 | 0.00 | 0.00 | NO |
| chr14 | 21400000 | 21500000 | 0.82 | 0.39 | A>A |
| chr14 | 21500000 | 21600000 | 0.96 | 0.68 | A>A |
| chr14 | 21600000 | 21700000 | 0.93 | 0.74 | A>A |
| chr14 | 21700000 | 21800000 | 1.09 | 0.95 | A>A |
| chr14 | 21800000 | 21900000 | 0.96 | 0.90 | A>A |
| chr14 | 21900000 | 22000000 | 0.90 | 0.59 | A>A |
| chr14 | 22000000 | 22100000 | 0.37 | 0.12 | A>A |
| chr14 | 22100000 | 22200000 | 0.02 | -0.24 | B>B |
| chr14 | 22200000 | 22300000 | -0.12 | -0.35 | B>B |
| chr14 | 22300000 | 22400000 | 0.03 | -0.12 | B>B |
| chr14 | 22400000 | 22500000 | 0.13 | -0.15 | B>B |
| chr14 | 22500000 | 22600000 | 0.04 | -0.13 | B>B |
| chr14 | 22600000 | 22700000 | 0.03 | -0.01 | A>A |
| chr14 | 22700000 | 22800000 | 0.39 | 0.42 | A>A |
| chr14 | 22800000 | 22900000 | 0.44 | 0.56 | A>A |
| chr14 | 22900000 | 23000000 | 0.57 | 0.53 | A>A |
| chr14 | 23000000 | 23100000 | 0.72 | 0.67 | A>A |
| chr14 | 23100000 | 23200000 | 0.00 | 0.72 | NO |
| chr14 | 23200000 | 23300000 | 0.81 | 0.74 | A>A |
| chr14 | 23300000 | 23400000 | 0.96 | 0.84 | A>A |
| chr14 | 23400000 | 23500000 | 1.15 | 1.03 | A>A |
| chr14 | 23500000 | 23600000 | 1.03 | 0.94 | A>A |
| chr14 | 23600000 | 23700000 | 1.00 | 0.88 | A>A |
| chr14 | 23700000 | 23800000 | 0.96 | 0.91 | A>A |
| chr14 | 23800000 | 23900000 | 0.96 | 0.84 | A>A |
| chr14 | 23900000 | 24000000 | 0.89 | 0.74 | A>A |
| chr14 | 24000000 | 24100000 | 0.79 | 0.71 | A>A |
| chr14 | 24100000 | 24200000 | 0.55 | 0.56 | A>A |
| chr14 | 24200000 | 24300000 | 0.70 | 0.56 | A>A |
| chr14 | 24300000 | 24400000 | 0.53 | 0.54 | A>A |
| chr14 | 24400000 | 24500000 | 0.00 | 0.00 | NO |
| chr14 | 24500000 | 24600000 | 0.99 | 0.96 | A>A |
| chr14 | 24600000 | 24700000 | 1.16 | 1.14 | A>A |
| chr14 | 24700000 | 24800000 | 0.99 | 0.81 | A>A |
| chr14 | 24800000 | 24900000 | 0.94 | 0.58 | A>A |
| chr14 | 24900000 | 25000000 | 0.25 | -0.30 | B>B |
| chr14 | 25000000 | 25100000 | -0.54 | -1.19 | A>B |
| chr14 | 25100000 | 25200000 | -0.60 | -1.02 | B>B |
| chr14 | 25200000 | 25300000 | -0.72 | -0.97 | B>B |
| chr14 | 25300000 | 25400000 | -0.63 | -0.79 | B>B |
| chr14 | 25400000 | 25500000 | -0.50 | -0.75 | B>B |
| chr14 | 25500000 | 25600000 | -0.56 | -0.92 | B>B |
| chr14 | 25600000 | 25700000 | -0.85 | -1.07 | B>B |
| chr14 | 25700000 | 25800000 | -1.10 | -1.19 | B>B |
| chr14 | 25800000 | 25900000 | -1.03 | -1.06 | B>B |
| chr14 | 25900000 | 26000000 | -1.10 | -1.02 | B>B |
| chr14 | 26000000 | 26100000 | -1.16 | -0.97 | B>B |
| chr14 | 26100000 | 26200000 | -1.09 | -1.07 | B>B |
| chr14 | 26200000 | 26300000 | -1.05 | -1.01 | B>B |
| chr14 | 26300000 | 26400000 | -0.94 | -0.93 | B>B |
| chr14 | 26400000 | 26500000 | -0.90 | -0.85 | B>B |

| | | | | | |
|---|---|---|---|---|---|
| chr14 | 26500000 | 26600000 | -0.85 | -0.79 | B>B |
| chr14 | 26600000 | 26700000 | -0.69 | -0.40 | B>B |
| chr14 | 26700000 | 26800000 | -0.59 | -0.03 | B>A |
| chr14 | 26800000 | 26900000 | -0.42 | 0.07 | B>B |
| chr14 | 26900000 | 27000000 | -0.10 | 0.20 | A>A |
| chr14 | 27000000 | 27100000 | -0.01 | 0.24 | A>A |
| chr14 | 27100000 | 27200000 | -0.05 | 0.17 | A>A |
| chr14 | 27200000 | 27300000 | 0.10 | 0.24 | A>A |
| chr14 | 27300000 | 27400000 | 0.06 | 0.20 | A>A |
| chr14 | 27400000 | 27500000 | -0.22 | 0.10 | B>B |
| chr14 | 27500000 | 27600000 | -0.62 | -0.04 | B>A |
| chr14 | 27600000 | 27700000 | -0.74 | -0.22 | B>A |
| chr14 | 27700000 | 27800000 | -0.86 | -0.31 | B>A |
| chr14 | 27800000 | 27900000 | -0.86 | -0.33 | B>A |
| chr14 | 27900000 | 28000000 | -0.83 | -0.32 | B>A |
| chr14 | 28000000 | 28100000 | -0.84 | -0.58 | B>B |
| chr14 | 28100000 | 28200000 | -0.82 | -0.70 | B>B |
| chr14 | 28200000 | 28300000 | 0.00 | 0.00 | NO |
| chr14 | 28300000 | 28400000 | -0.67 | -0.70 | B>B |
| chr14 | 28400000 | 28500000 | -0.57 | -0.72 | B>B |
| chr14 | 28500000 | 28600000 | -0.41 | -0.53 | B>B |
| chr14 | 28600000 | 28700000 | -0.20 | -0.38 | B>B |
| chr14 | 28700000 | 28800000 | -0.10 | -0.27 | B>B |
| chr14 | 28800000 | 28900000 | -0.09 | -0.25 | B>B |
| chr14 | 28900000 | 29000000 | -0.14 | -0.30 | B>B |
| chr14 | 29000000 | 29100000 | -0.22 | -0.30 | B>B |
| chr14 | 29100000 | 29200000 | -0.34 | -0.38 | B>B |
| chr14 | 29200000 | 29300000 | -0.52 | -0.57 | B>B |
| chr14 | 29300000 | 29400000 | -0.79 | -0.97 | B>B |
| chr14 | 29400000 | 29500000 | -0.80 | -1.02 | B>B |
| chr14 | 29500000 | 29600000 | -0.80 | -1.07 | B>B |
| chr14 | 29600000 | 29700000 | -0.77 | -1.14 | B>B |
| chr14 | 29700000 | 29800000 | -0.75 | -1.08 | B>B |
| chr14 | 29800000 | 29900000 | -0.84 | -1.09 | B>B |
| chr14 | 29900000 | 30000000 | -0.60 | -0.86 | B>B |
| chr14 | 30000000 | 30100000 | -0.16 | -0.39 | B>B |
| chr14 | 30100000 | 30200000 | 0.03 | -0.30 | B>B |
| chr14 | 30200000 | 30300000 | -0.07 | -0.31 | B>B |
| chr14 | 30300000 | 30400000 | -0.01 | -0.21 | B>B |
| chr14 | 30400000 | 30500000 | -0.26 | -0.42 | B>B |
| chr14 | 30500000 | 30600000 | -0.63 | -0.74 | B>B |
| chr14 | 30600000 | 30700000 | -0.68 | -0.78 | B>B |
| chr14 | 30700000 | 30800000 | -0.53 | -0.74 | B>B |
| chr14 | 30800000 | 30900000 | -0.64 | -0.96 | B>B |
| chr14 | 30900000 | 31000000 | -0.39 | -0.81 | B>B |
| chr14 | 31000000 | 31100000 | 0.30 | -0.01 | A>A |
| chr14 | 31100000 | 31200000 | 0.49 | 0.40 | A>A |
| chr14 | 31200000 | 31300000 | 0.43 | 0.27 | A>A |
| chr14 | 31300000 | 31400000 | 0.69 | 0.57 | A>A |
| chr14 | 31400000 | 31500000 | 0.96 | 0.93 | A>A |
| chr14 | 31500000 | 31600000 | 0.93 | 1.03 | A>A |
| chr14 | 31600000 | 31700000 | 1.04 | 0.99 | A>A |
| chr14 | 31700000 | 31800000 | 1.03 | 0.99 | A>A |
| chr14 | 31800000 | 31900000 | 0.73 | 0.79 | A>A |
| chr14 | 31900000 | 32000000 | 0.39 | 0.59 | A>A |
| chr14 | 32000000 | 32100000 | 0.31 | 0.45 | A>A |

| | | | | | |
|---|---|---|---|---|---|
| chr14 | 32100000 | 32200000 | 0.16 | 0.42 | A>A |
| chr14 | 32200000 | 32300000 | 0.28 | 0.39 | A>A |
| chr14 | 32300000 | 32400000 | 0.35 | 0.47 | A>A |
| chr14 | 32400000 | 32500000 | 0.65 | 0.67 | A>A |
| chr14 | 32500000 | 32600000 | 0.65 | 0.60 | A>A |
| chr14 | 32600000 | 32700000 | 0.52 | 0.61 | A>A |
| chr14 | 32700000 | 32800000 | 0.31 | 0.46 | A>A |
| chr14 | 32800000 | 32900000 | 0.26 | 0.51 | A>A |
| chr14 | 32900000 | 33000000 | 0.17 | 0.39 | A>A |
| chr14 | 33000000 | 33100000 | -0.13 | -0.16 | B>B |
| chr14 | 33100000 | 33200000 | -0.32 | -0.68 | B>B |
| chr14 | 33200000 | 33300000 | -0.64 | -0.93 | B>B |
| chr14 | 33300000 | 33400000 | -0.48 | -0.87 | B>B |
| chr14 | 33400000 | 33500000 | -0.44 | -0.86 | B>B |
| chr14 | 33500000 | 33600000 | -0.64 | -1.06 | B>B |
| chr14 | 33600000 | 33700000 | -0.61 | -1.14 | B>B |
| chr14 | 33700000 | 33800000 | -0.63 | -1.12 | B>B |
| chr14 | 33800000 | 33900000 | -0.60 | -1.10 | B>B |
| chr14 | 33900000 | 34000000 | -0.48 | -1.03 | B>B |
| chr14 | 34000000 | 34100000 | -0.60 | -1.08 | B>B |
| chr14 | 34100000 | 34200000 | -0.44 | -1.03 | A>B |
| chr14 | 34200000 | 34300000 | -0.25 | -0.63 | B>B |
| chr14 | 34300000 | 34400000 | 0.27 | 0.05 | A>A |
| chr14 | 34400000 | 34500000 | 0.46 | 0.25 | A>A |
| chr14 | 34500000 | 34600000 | 0.47 | 0.25 | A>A |
| chr14 | 34600000 | 34700000 | 0.32 | 0.18 | A>A |
| chr14 | 34700000 | 34800000 | 0.13 | -0.02 | A>A |
| chr14 | 34800000 | 34900000 | 0.54 | 0.39 | A>A |
| chr14 | 34900000 | 35000000 | 0.63 | 0.48 | A>A |
| chr14 | 35000000 | 35100000 | 0.98 | 0.83 | A>A |
| chr14 | 35100000 | 35200000 | 1.07 | 0.95 | A>A |
| chr14 | 35200000 | 35300000 | 1.01 | 0.98 | A>A |
| chr14 | 35300000 | 35400000 | 1.07 | 1.08 | A>A |
| chr14 | 35400000 | 35500000 | 0.93 | 0.91 | A>A |
| chr14 | 35500000 | 35600000 | 1.04 | 0.94 | A>A |
| chr14 | 35600000 | 35700000 | 0.76 | 0.96 | A>A |
| chr14 | 35700000 | 35800000 | 1.07 | 1.02 | A>A |
| chr14 | 35800000 | 35900000 | 1.00 | 1.02 | A>A |
| chr14 | 35900000 | 36000000 | 0.60 | 0.71 | A>A |
| chr14 | 36000000 | 36100000 | 0.62 | 0.73 | A>A |
| chr14 | 36100000 | 36200000 | 0.63 | 0.74 | A>A |
| chr14 | 36200000 | 36300000 | 0.65 | 0.77 | A>A |
| chr14 | 36300000 | 36400000 | 0.34 | 0.55 | A>A |
| chr14 | 36400000 | 36500000 | 0.17 | 0.48 | A>A |
| chr14 | 36500000 | 36600000 | 0.09 | 0.26 | A>A |
| chr14 | 36600000 | 36700000 | -0.18 | 0.03 | B>B |
| chr14 | 36700000 | 36800000 | 0.07 | 0.07 | A>A |
| chr14 | 36800000 | 36900000 | -0.20 | 0.02 | B>B |
| chr14 | 36900000 | 37000000 | -0.05 | 0.04 | B>B |
| chr14 | 37000000 | 37100000 | 0.07 | 0.25 | A>A |
| chr14 | 37100000 | 37200000 | 0.05 | 0.33 | A>A |
| chr14 | 37200000 | 37300000 | 0.06 | 0.23 | A>A |
| chr14 | 37300000 | 37400000 | -0.02 | 0.29 | A>A |
| chr14 | 37400000 | 37500000 | 0.25 | 0.44 | A>A |
| chr14 | 37500000 | 37600000 | 0.24 | 0.40 | A>A |
| chr14 | 37600000 | 37700000 | 0.22 | 0.27 | A>A |

| | | | | | |
|---|---|---|---|---|---|
| chr14 | 37700000 | 37800000 | 0.14 | 0.38 | A>A |
| chr14 | 37800000 | 37900000 | 0.32 | 0.51 | A>A |
| chr14 | 37900000 | 38000000 | 0.14 | 0.48 | A>A |
| chr14 | 38000000 | 38100000 | 0.16 | 0.38 | A>A |
| chr14 | 38100000 | 38200000 | -0.13 | 0.17 | A>A |
| chr14 | 38200000 | 38300000 | -0.44 | -0.02 | B>B |
| chr14 | 38300000 | 38400000 | -0.37 | -0.14 | B>B |
| chr14 | 38400000 | 38500000 | -0.71 | -0.51 | B>B |
| chr14 | 38500000 | 38600000 | -0.86 | -0.35 | B>B |
| chr14 | 38600000 | 38700000 | -0.75 | -0.57 | B>B |
| chr14 | 38700000 | 38800000 | -0.91 | -0.96 | B>B |
| chr14 | 38800000 | 38900000 | -0.92 | -1.14 | B>B |
| chr14 | 38900000 | 39000000 | -0.88 | -1.17 | B>B |
| chr14 | 39000000 | 39100000 | -0.76 | -1.10 | B>B |
| chr14 | 39100000 | 39200000 | -0.64 | -1.07 | B>B |
| chr14 | 39200000 | 39300000 | -0.59 | -0.93 | B>B |
| chr14 | 39300000 | 39400000 | -0.48 | -0.68 | B>B |
| chr14 | 39400000 | 39500000 | -0.22 | -0.33 | B>B |
| chr14 | 39500000 | 39600000 | 0.42 | 0.25 | A>A |
| chr14 | 39600000 | 39700000 | 0.84 | 0.74 | A>A |
| chr14 | 39700000 | 39800000 | 0.77 | 0.66 | A>A |
| chr14 | 39800000 | 39900000 | 0.37 | 0.26 | A>A |
| chr14 | 39900000 | 40000000 | -0.51 | -0.58 | B>B |
| chr14 | 40000000 | 40100000 | -0.76 | -1.02 | B>B |
| chr14 | 40100000 | 40200000 | -0.85 | -1.20 | B>B |
| chr14 | 40200000 | 40300000 | -0.87 | -1.17 | B>B |
| chr14 | 40300000 | 40400000 | -0.95 | -1.13 | B>B |
| chr14 | 40400000 | 40500000 | -0.92 | -1.08 | B>B |
| chr14 | 40500000 | 40600000 | -1.03 | -1.12 | B>B |
| chr14 | 40600000 | 40700000 | -0.97 | -1.10 | B>B |
| chr14 | 40700000 | 40800000 | -0.98 | -1.09 | B>B |
| chr14 | 40800000 | 40900000 | -1.00 | -1.08 | B>B |
| chr14 | 40900000 | 41000000 | -1.01 | -1.06 | B>B |
| chr14 | 41000000 | 41100000 | -1.05 | -1.05 | B>B |
| chr14 | 41100000 | 41200000 | -0.98 | -1.05 | B>B |
| chr14 | 41200000 | 41300000 | -0.96 | -1.07 | B>B |
| chr14 | 41300000 | 41400000 | -1.01 | -1.05 | B>B |
| chr14 | 41400000 | 41500000 | -0.96 | -1.00 | B>B |
| chr14 | 41500000 | 41600000 | -0.93 | -0.92 | B>B |
| chr14 | 41600000 | 41700000 | 0.00 | 0.00 | NO |
| chr14 | 41700000 | 41800000 | -0.86 | -0.99 | B>B |
| chr14 | 41800000 | 41900000 | -0.96 | -1.02 | B>B |
| chr14 | 41900000 | 42000000 | -0.94 | -1.00 | B>B |
| chr14 | 42000000 | 42100000 | -0.68 | -0.88 | B>B |
| chr14 | 42100000 | 42200000 | -0.75 | -0.88 | B>B |
| chr14 | 42200000 | 42300000 | -0.65 | -0.79 | B>B |
| chr14 | 42300000 | 42400000 | -0.60 | -0.76 | B>B |
| chr14 | 42400000 | 42500000 | -0.36 | -0.61 | B>B |
| chr14 | 42500000 | 42600000 | -0.16 | -0.40 | B>B |
| chr14 | 42600000 | 42700000 | -0.02 | -0.08 | B>B |
| chr14 | 42700000 | 42800000 | -0.11 | -0.23 | B>B |
| chr14 | 42800000 | 42900000 | -0.24 | -0.38 | B>B |
| chr14 | 42900000 | 43000000 | -0.22 | -0.33 | B>B |
| chr14 | 43000000 | 43100000 | -0.34 | -0.33 | B>B |
| chr14 | 43100000 | 43200000 | -0.33 | -0.28 | B>B |
| chr14 | 43200000 | 43300000 | -0.31 | -0.24 | B>B |

| | | | | | |
|---|---|---|---|---|---|
| chr14 | 43300000 | 43400000 | -0.22 | -0.07 | B>B |
| chr14 | 43400000 | 43500000 | -0.19 | 0.00 | B>B |
| chr14 | 43500000 | 43600000 | -0.22 | -0.02 | B>B |
| chr14 | 43600000 | 43700000 | -0.24 | -0.06 | B>B |
| chr14 | 43700000 | 43800000 | -0.37 | -0.26 | B>B |
| chr14 | 43800000 | 43900000 | -0.39 | -0.24 | B>B |
| chr14 | 43900000 | 44000000 | -0.45 | -0.33 | B>B |
| chr14 | 44000000 | 44100000 | -0.53 | -0.42 | B>B |
| chr14 | 44100000 | 44200000 | -0.53 | -0.47 | B>B |
| chr14 | 44200000 | 44300000 | -0.59 | -0.51 | B>B |
| chr14 | 44300000 | 44400000 | -0.64 | -0.46 | B>B |
| chr14 | 44400000 | 44500000 | -0.73 | -0.57 | B>B |
| chr14 | 44500000 | 44600000 | -0.81 | -0.55 | B>B |
| chr14 | 44600000 | 44700000 | -0.79 | -0.45 | B>B |
| chr14 | 44700000 | 44800000 | -0.72 | -0.41 | B>B |
| chr14 | 44800000 | 44900000 | -0.73 | -0.49 | B>B |
| chr14 | 44900000 | 45000000 | -0.83 | -0.58 | B>B |
| chr14 | 45000000 | 45100000 | -0.86 | -0.60 | B>B |
| chr14 | 45100000 | 45200000 | -0.80 | -0.65 | B>B |
| chr14 | 45200000 | 45300000 | -0.69 | -0.56 | B>B |
| chr14 | 45300000 | 45400000 | -0.18 | -0.24 | B>B |
| chr14 | 45400000 | 45500000 | 0.59 | 0.49 | A>A |
| chr14 | 45500000 | 45600000 | 0.72 | 0.60 | A>A |
| chr14 | 45600000 | 45700000 | 0.48 | 0.36 | A>A |
| chr14 | 45700000 | 45800000 | -0.17 | -0.53 | B>B |
| chr14 | 45800000 | 45900000 | -0.82 | -1.15 | B>B |
| chr14 | 45900000 | 46000000 | -0.91 | -1.19 | B>B |
| chr14 | 46000000 | 46100000 | -1.02 | -1.25 | B>B |
| chr14 | 46100000 | 46200000 | -1.09 | -1.20 | B>B |
| chr14 | 46200000 | 46300000 | -1.05 | -1.24 | B>B |
| chr14 | 46300000 | 46400000 | -1.05 | -1.23 | B>B |
| chr14 | 46400000 | 46500000 | -1.01 | -1.24 | B>B |
| chr14 | 46500000 | 46600000 | -1.00 | -1.20 | B>B |
| chr14 | 46600000 | 46700000 | -1.02 | -1.15 | B>B |
| chr14 | 46700000 | 46800000 | -1.01 | -1.13 | B>B |
| chr14 | 46800000 | 46900000 | -1.01 | -1.11 | B>B |
| chr14 | 46900000 | 47000000 | -0.97 | -1.00 | B>B |
| chr14 | 47000000 | 47100000 | -0.87 | -0.81 | B>B |
| chr14 | 47100000 | 47200000 | -0.95 | -0.90 | B>B |
| chr14 | 47200000 | 47300000 | -1.01 | -1.00 | B>B |
| chr14 | 47300000 | 47400000 | -0.98 | -1.01 | B>B |
| chr14 | 47400000 | 47500000 | -0.93 | -1.02 | B>B |
| chr14 | 47500000 | 47600000 | -0.99 | -1.06 | B>B |
| chr14 | 47600000 | 47700000 | -0.96 | -1.01 | B>B |
| chr14 | 47700000 | 47800000 | -0.95 | -1.02 | B>B |
| chr14 | 47800000 | 47900000 | -0.92 | -1.09 | B>B |
| chr14 | 47900000 | 48000000 | -0.97 | -1.04 | B>B |
| chr14 | 48000000 | 48100000 | -0.94 | -0.99 | B>B |
| chr14 | 48100000 | 48200000 | -0.96 | -0.87 | B>B |
| chr14 | 48200000 | 48300000 | -0.98 | -0.72 | B>B |
| chr14 | 48300000 | 48400000 | -0.99 | -0.51 | B>B |
| chr14 | 48400000 | 48500000 | -1.01 | -0.38 | B>A |
| chr14 | 48500000 | 48600000 | -0.96 | -0.28 | B>A |
| chr14 | 48600000 | 48700000 | -1.02 | -0.17 | B>A |
| chr14 | 48700000 | 48800000 | -1.05 | -0.11 | B>A |
| chr14 | 48800000 | 48900000 | -0.96 | -0.18 | B>A |

| | | | | | |
|---|---|---|---|---|---|
| chr14 | 48900000 | 49000000 | -1.04 | -0.05 | B>A |
| chr14 | 49000000 | 49100000 | -1.09 | -0.03 | B>A |
| chr14 | 49100000 | 49200000 | -1.12 | -0.19 | B>A |
| chr14 | 49200000 | 49300000 | -1.03 | -0.18 | B>A |
| chr14 | 49300000 | 49400000 | -1.07 | -0.33 | B>A |
| chr14 | 49400000 | 49500000 | -1.10 | -0.47 | B>A |
| chr14 | 49500000 | 49600000 | -1.06 | -0.67 | B>B |
| chr14 | 49600000 | 49700000 | -0.96 | -0.85 | B>B |
| chr14 | 49700000 | 49800000 | -0.97 | -1.20 | B>B |
| chr14 | 49800000 | 49900000 | -0.83 | -1.19 | B>B |
| chr14 | 49900000 | 50000000 | -0.40 | -0.70 | B>B |
| chr14 | 50000000 | 50100000 | 0.38 | 0.19 | A>A |
| chr14 | 50100000 | 50200000 | 0.95 | 0.82 | A>A |
| chr14 | 50200000 | 50300000 | 0.96 | 0.82 | A>A |
| chr14 | 50300000 | 50400000 | 1.09 | 0.97 | A>A |
| chr14 | 50400000 | 50500000 | 1.26 | 1.17 | A>A |
| chr14 | 50500000 | 50600000 | 1.03 | 1.03 | A>A |
| chr14 | 50600000 | 50700000 | 1.03 | 0.99 | A>A |
| chr14 | 50700000 | 50800000 | 0.97 | 0.96 | A>A |
| chr14 | 50800000 | 50900000 | 1.01 | 0.95 | A>A |
| chr14 | 50900000 | 51000000 | 0.91 | 0.96 | A>A |
| chr14 | 51000000 | 51100000 | 0.81 | 0.83 | A>A |
| chr14 | 51100000 | 51200000 | 0.94 | 0.93 | A>A |
| chr14 | 51200000 | 51300000 | 0.92 | 0.89 | A>A |
| chr14 | 51300000 | 51400000 | 0.86 | 0.74 | A>A |
| chr14 | 51400000 | 51500000 | 0.89 | 0.70 | A>A |
| chr14 | 51500000 | 51600000 | 0.89 | 0.71 | A>A |
| chr14 | 51600000 | 51700000 | 0.72 | 0.60 | A>A |
| chr14 | 51700000 | 51800000 | 0.56 | 0.52 | A>A |
| chr14 | 51800000 | 51900000 | 0.51 | 0.48 | A>A |
| chr14 | 51900000 | 52000000 | 0.53 | 0.47 | A>A |
| chr14 | 52000000 | 52100000 | 0.58 | 0.48 | A>A |
| chr14 | 52100000 | 52200000 | 0.56 | 0.51 | A>A |
| chr14 | 52200000 | 52300000 | 0.37 | 0.41 | A>A |
| chr14 | 52300000 | 52400000 | 0.40 | 0.47 | A>A |
| chr14 | 52400000 | 52500000 | 0.38 | 0.44 | A>A |
| chr14 | 52500000 | 52600000 | 0.31 | 0.47 | A>A |
| chr14 | 52600000 | 52700000 | 0.03 | 0.38 | A>A |
| chr14 | 52700000 | 52800000 | 0.21 | 0.42 | A>A |
| chr14 | 52800000 | 52900000 | 0.39 | 0.57 | A>A |
| chr14 | 52900000 | 53000000 | 0.31 | 0.42 | A>A |
| chr14 | 53000000 | 53100000 | 0.41 | 0.51 | A>A |
| chr14 | 53100000 | 53200000 | 0.69 | 0.74 | A>A |
| chr14 | 53200000 | 53300000 | 0.62 | 0.73 | A>A |
| chr14 | 53300000 | 53400000 | 0.76 | 0.73 | A>A |
| chr14 | 53400000 | 53500000 | 0.64 | 0.63 | A>A |
| chr14 | 53500000 | 53600000 | 0.49 | 0.58 | A>A |
| chr14 | 53600000 | 53700000 | 0.49 | 0.63 | A>A |
| chr14 | 53700000 | 53800000 | 0.30 | 0.56 | A>A |
| chr14 | 53800000 | 53900000 | 0.29 | 0.50 | A>A |
| chr14 | 53900000 | 54000000 | 0.31 | 0.46 | A>A |
| chr14 | 54000000 | 54100000 | 0.44 | 0.57 | A>A |
| chr14 | 54100000 | 54200000 | 0.45 | 0.58 | A>A |
| chr14 | 54200000 | 54300000 | 0.47 | 0.60 | A>A |
| chr14 | 54300000 | 54400000 | 0.52 | 0.68 | A>A |
| chr14 | 54400000 | 54500000 | 0.40 | 0.57 | A>A |

| | | | | | |
|---|---|---|---|---|---|
| chr14 | 54500000 | 54600000 | 0.38 | 0.59 | A>A |
| chr14 | 54600000 | 54700000 | 0.34 | 0.57 | A>A |
| chr14 | 54700000 | 54800000 | 0.38 | 0.56 | A>A |
| chr14 | 54800000 | 54900000 | 0.56 | 0.69 | A>A |
| chr14 | 54900000 | 55000000 | 0.90 | 0.84 | A>A |
| chr14 | 55000000 | 55100000 | 0.83 | 0.82 | A>A |
| chr14 | 55100000 | 55200000 | 0.91 | 0.90 | A>A |
| chr14 | 55200000 | 55300000 | 0.83 | 0.81 | A>A |
| chr14 | 55300000 | 55400000 | 0.95 | 0.91 | A>A |
| chr14 | 55400000 | 55500000 | 1.07 | 0.97 | A>A |
| chr14 | 55500000 | 55600000 | 1.13 | 0.98 | A>A |
| chr14 | 55600000 | 55700000 | 0.98 | 0.86 | A>A |
| chr14 | 55700000 | 55800000 | 0.88 | 0.74 | A>A |
| chr14 | 55800000 | 55900000 | 0.69 | 0.68 | A>A |
| chr14 | 55900000 | 56000000 | 0.56 | 0.61 | A>A |
| chr14 | 56000000 | 56100000 | 0.76 | 0.70 | A>A |
| chr14 | 56100000 | 56200000 | 0.50 | 0.57 | A>A |
| chr14 | 56200000 | 56300000 | 0.56 | 0.63 | A>A |
| chr14 | 56300000 | 56400000 | 0.49 | 0.54 | A>A |
| chr14 | 56400000 | 56500000 | 0.46 | 0.55 | A>A |
| chr14 | 56500000 | 56600000 | 0.38 | 0.48 | A>A |
| chr14 | 56600000 | 56700000 | 0.39 | 0.55 | A>A |
| chr14 | 56700000 | 56800000 | 0.26 | 0.42 | A>A |
| chr14 | 56800000 | 56900000 | 0.13 | 0.31 | A>A |
| chr14 | 56900000 | 57000000 | 0.05 | 0.12 | A>A |
| chr14 | 57000000 | 57100000 | 0.08 | -0.05 | A>A |
| chr14 | 57100000 | 57200000 | -0.23 | -0.52 | B>B |
| chr14 | 57200000 | 57300000 | -0.37 | -0.82 | B>B |
| chr14 | 57300000 | 57400000 | -0.45 | -0.90 | B>B |
| chr14 | 57400000 | 57500000 | -0.48 | -0.92 | B>B |
| chr14 | 57500000 | 57600000 | -0.40 | -0.68 | B>B |
| chr14 | 57600000 | 57700000 | -0.06 | -0.34 | B>B |
| chr14 | 57700000 | 57800000 | 0.25 | 0.12 | A>A |
| chr14 | 57800000 | 57900000 | 0.10 | 0.00 | A>A |
| chr14 | 57900000 | 58000000 | -0.18 | -0.15 | B>B |
| chr14 | 58000000 | 58100000 | -0.51 | -0.46 | B>B |
| chr14 | 58100000 | 58200000 | -0.66 | -0.47 | B>B |
| chr14 | 58200000 | 58300000 | -0.64 | -0.36 | B>B |
| chr14 | 58300000 | 58400000 | -0.60 | -0.54 | B>B |
| chr14 | 58400000 | 58500000 | -0.60 | -0.67 | B>B |
| chr14 | 58500000 | 58600000 | -0.02 | -0.12 | B>B |
| chr14 | 58600000 | 58700000 | 0.46 | 0.38 | A>A |
| chr14 | 58700000 | 58800000 | 0.76 | 0.65 | A>A |
| chr14 | 58800000 | 58900000 | 0.69 | 0.57 | A>A |
| chr14 | 58900000 | 59000000 | 0.36 | 0.12 | A>A |
| chr14 | 59000000 | 59100000 | -0.02 | -0.33 | B>B |
| chr14 | 59100000 | 59200000 | 0.04 | -0.25 | B>B |
| chr14 | 59200000 | 59300000 | 0.15 | -0.11 | A>A |
| chr14 | 59300000 | 59400000 | -0.17 | -0.52 | B>B |
| chr14 | 59400000 | 59500000 | -0.32 | -0.53 | B>B |
| chr14 | 59500000 | 59600000 | -0.20 | -0.38 | B>B |
| chr14 | 59600000 | 59700000 | 0.11 | 0.06 | A>A |
| chr14 | 59700000 | 59800000 | 0.19 | 0.15 | A>A |
| chr14 | 59800000 | 59900000 | -0.02 | 0.07 | A>A |
| chr14 | 59900000 | 60000000 | 0.05 | -0.16 | B>B |
| chr14 | 60000000 | 60100000 | -0.52 | -0.84 | B>B |

| | | | | | |
|---|---|---|---|---|---|
| chr14 | 60100000 | 60200000 | -0.90 | -1.26 | B>B |
| chr14 | 60200000 | 60300000 | -0.92 | -1.31 | B>B |
| chr14 | 60300000 | 60400000 | -0.73 | -0.95 | B>B |
| chr14 | 60400000 | 60500000 | -0.63 | -0.27 | B>B |
| chr14 | 60500000 | 60600000 | -0.21 | -0.08 | B>B |
| chr14 | 60600000 | 60700000 | -0.19 | -0.23 | B>B |
| chr14 | 60700000 | 60800000 | -0.04 | -0.26 | B>B |
| chr14 | 60800000 | 60900000 | -0.60 | -0.78 | B>B |
| chr14 | 60900000 | 61000000 | -0.64 | -0.84 | B>B |
| chr14 | 61000000 | 61100000 | -0.26 | -0.40 | B>B |
| chr14 | 61100000 | 61200000 | 0.36 | 0.18 | A>A |
| chr14 | 61200000 | 61300000 | 0.40 | 0.24 | A>A |
| chr14 | 61300000 | 61400000 | 0.32 | 0.16 | A>A |
| chr14 | 61400000 | 61500000 | 0.31 | 0.13 | A>A |
| chr14 | 61500000 | 61600000 | 0.40 | 0.14 | A>A |
| chr14 | 61600000 | 61700000 | 0.35 | 0.00 | A>A |
| chr14 | 61700000 | 61800000 | 0.08 | -0.30 | B>B |
| chr14 | 61800000 | 61900000 | 0.09 | -0.57 | A>B |
| chr14 | 61900000 | 62000000 | 0.39 | 0.13 | A>A |
| chr14 | 62000000 | 62100000 | 0.69 | 0.47 | A>A |
| chr14 | 62100000 | 62200000 | 0.63 | 0.40 | A>A |
| chr14 | 62200000 | 62300000 | 0.30 | 0.00 | A>A |
| chr14 | 62300000 | 62400000 | -0.34 | -0.71 | B>B |
| chr14 | 62400000 | 62500000 | -0.85 | -1.23 | B>B |
| chr14 | 62500000 | 62600000 | -0.71 | -1.11 | B>B |
| chr14 | 62600000 | 62700000 | -0.64 | -0.88 | B>B |
| chr14 | 62700000 | 62800000 | -0.91 | -1.08 | B>B |
| chr14 | 62800000 | 62900000 | -0.88 | -1.06 | B>B |
| chr14 | 62900000 | 63000000 | -0.83 | -0.92 | B>B |
| chr14 | 63000000 | 63100000 | -0.50 | -0.78 | B>B |
| chr14 | 63100000 | 63200000 | -0.27 | -0.48 | B>B |
| chr14 | 63200000 | 63300000 | -0.42 | -0.63 | B>B |
| chr14 | 63300000 | 63400000 | -0.65 | -0.92 | B>B |
| chr14 | 63400000 | 63500000 | -0.66 | -0.94 | B>B |
| chr14 | 63500000 | 63600000 | -0.68 | -0.97 | B>B |
| chr14 | 63600000 | 63700000 | -0.68 | -1.15 | B>B |
| chr14 | 63700000 | 63800000 | -0.39 | -0.83 | B>B |
| chr14 | 63800000 | 63900000 | 0.19 | 0.03 | A>A |
| chr14 | 63900000 | 64000000 | 0.54 | 0.48 | A>A |
| chr14 | 64000000 | 64100000 | 0.51 | 0.44 | A>A |
| chr14 | 64100000 | 64200000 | 0.68 | 0.62 | A>A |
| chr14 | 64200000 | 64300000 | 0.55 | 0.58 | A>A |
| chr14 | 64300000 | 64400000 | 0.72 | 0.80 | A>A |
| chr14 | 64400000 | 64500000 | 0.61 | 0.76 | A>A |
| chr14 | 64500000 | 64600000 | 0.67 | 0.76 | A>A |
| chr14 | 64600000 | 64700000 | 0.71 | 0.78 | A>A |
| chr14 | 64700000 | 64800000 | 0.60 | 0.61 | A>A |
| chr14 | 64800000 | 64900000 | 0.89 | 0.89 | A>A |
| chr14 | 64900000 | 65000000 | 0.90 | 0.94 | A>A |
| chr14 | 65000000 | 65100000 | 1.05 | 0.96 | A>A |
| chr14 | 65100000 | 65200000 | 1.07 | 0.98 | A>A |
| chr14 | 65200000 | 65300000 | 1.04 | 0.95 | A>A |
| chr14 | 65300000 | 65400000 | 0.95 | 0.82 | A>A |
| chr14 | 65400000 | 65500000 | 0.77 | 0.70 | A>A |
| chr14 | 65500000 | 65600000 | 0.55 | 0.35 | A>A |
| chr14 | 65600000 | 65700000 | 0.29 | 0.10 | A>A |

| | | | | | |
|---|---|---|---|---|---|
| chr14 | 65700000 | 65800000 | 0.39 | 0.30 | A>A |
| chr14 | 65800000 | 65900000 | 0.19 | 0.14 | A>A |
| chr14 | 65900000 | 66000000 | 0.06 | 0.05 | A>A |
| chr14 | 66000000 | 66100000 | 0.05 | 0.01 | A>A |
| chr14 | 66100000 | 66200000 | -0.14 | -0.16 | B>B |
| chr14 | 66200000 | 66300000 | -0.23 | -0.36 | B>B |
| chr14 | 66300000 | 66400000 | -0.33 | -0.68 | B>B |
| chr14 | 66400000 | 66500000 | -0.52 | -0.85 | B>B |
| chr14 | 66500000 | 66600000 | -0.57 | -1.01 | B>B |
| chr14 | 66600000 | 66700000 | -0.84 | -1.19 | B>B |
| chr14 | 66700000 | 66800000 | -0.79 | -1.11 | B>B |
| chr14 | 66800000 | 66900000 | -0.54 | -0.66 | B>B |
| chr14 | 66900000 | 67000000 | -0.22 | -0.37 | B>B |
| chr14 | 67000000 | 67100000 | -0.18 | -0.10 | B>B |
| chr14 | 67100000 | 67200000 | -0.24 | -0.17 | B>B |
| chr14 | 67200000 | 67300000 | -0.24 | -0.18 | B>B |
| chr14 | 67300000 | 67400000 | -0.31 | -0.19 | B>B |
| chr14 | 67400000 | 67500000 | -0.20 | -0.09 | B>B |
| chr14 | 67500000 | 67600000 | -0.18 | -0.11 | B>B |
| chr14 | 67600000 | 67700000 | 0.32 | 0.24 | A>A |
| chr14 | 67700000 | 67800000 | 0.49 | 0.39 | A>A |
| chr14 | 67800000 | 67900000 | 0.64 | 0.62 | A>A |
| chr14 | 67900000 | 68000000 | 0.78 | 0.76 | A>A |
| chr14 | 68000000 | 68100000 | 0.85 | 0.79 | A>A |
| chr14 | 68100000 | 68200000 | 0.85 | 0.82 | A>A |
| chr14 | 68200000 | 68300000 | 0.94 | 0.84 | A>A |
| chr14 | 68300000 | 68400000 | 0.69 | 0.66 | A>A |
| chr14 | 68400000 | 68500000 | 0.72 | 0.74 | A>A |
| chr14 | 68500000 | 68600000 | 0.82 | 0.85 | A>A |
| chr14 | 68600000 | 68700000 | 0.73 | 0.78 | A>A |
| chr14 | 68700000 | 68800000 | 0.71 | 0.68 | A>A |
| chr14 | 68800000 | 68900000 | 0.81 | 0.77 | A>A |
| chr14 | 68900000 | 69000000 | 0.74 | 0.75 | A>A |
| chr14 | 69000000 | 69100000 | 0.75 | 0.76 | A>A |
| chr14 | 69100000 | 69200000 | 0.79 | 0.79 | A>A |
| chr14 | 69200000 | 69300000 | 0.65 | 0.69 | A>A |
| chr14 | 69300000 | 69400000 | 0.73 | 0.73 | A>A |
| chr14 | 69400000 | 69500000 | 1.01 | 0.93 | A>A |
| chr14 | 69500000 | 69600000 | 0.79 | 0.79 | A>A |
| chr14 | 69600000 | 69700000 | 0.92 | 0.87 | A>A |
| chr14 | 69700000 | 69800000 | 0.93 | 0.87 | A>A |
| chr14 | 69800000 | 69900000 | 0.97 | 0.90 | A>A |
| chr14 | 69900000 | 70000000 | 0.86 | 0.79 | A>A |
| chr14 | 70000000 | 70100000 | 0.83 | 0.80 | A>A |
| chr14 | 70100000 | 70200000 | 0.88 | 0.80 | A>A |
| chr14 | 70200000 | 70300000 | 0.58 | 0.60 | A>A |
| chr14 | 70300000 | 70400000 | 0.56 | 0.52 | A>A |
| chr14 | 70400000 | 70500000 | 0.56 | 0.55 | A>A |
| chr14 | 70500000 | 70600000 | 0.36 | 0.38 | A>A |
| chr14 | 70600000 | 70700000 | 0.27 | 0.36 | A>A |
| chr14 | 70700000 | 70800000 | 0.30 | 0.46 | A>A |
| chr14 | 70800000 | 70900000 | 0.32 | 0.48 | A>A |
| chr14 | 70900000 | 71000000 | 0.20 | 0.35 | A>A |
| chr14 | 71000000 | 71100000 | 0.41 | 0.54 | A>A |
| chr14 | 71100000 | 71200000 | 0.61 | 0.68 | A>A |
| chr14 | 71200000 | 71300000 | 0.67 | 0.67 | A>A |

| | | | | | |
|---|---|---|---|---|---|
| chr14 | 71300000 | 71400000 | 0.60 | 0.63 | A>A |
| chr14 | 71400000 | 71500000 | 0.48 | 0.54 | A>A |
| chr14 | 71500000 | 71600000 | 0.23 | 0.23 | A>A |
| chr14 | 71600000 | 71700000 | 0.01 | 0.00 | A>A |
| chr14 | 71700000 | 71800000 | 0.01 | 0.08 | A>A |
| chr14 | 71800000 | 71900000 | 0.13 | 0.12 | A>A |
| chr14 | 71900000 | 72000000 | 0.14 | 0.08 | A>A |
| chr14 | 72000000 | 72100000 | 0.02 | 0.02 | A>A |
| chr14 | 72100000 | 72200000 | -0.04 | -0.09 | B>B |
| chr14 | 72200000 | 72300000 | -0.18 | -0.19 | B>B |
| chr14 | 72300000 | 72400000 | -0.46 | -0.71 | B>B |
| chr14 | 72400000 | 72500000 | -0.64 | -1.22 | B>B |
| chr14 | 72500000 | 72600000 | -0.75 | -1.38 | A>B |
| chr14 | 72600000 | 72700000 | -0.81 | -1.38 | B>B |
| chr14 | 72700000 | 72800000 | -0.75 | -1.25 | B>B |
| chr14 | 72800000 | 72900000 | -0.54 | -1.04 | B>B |
| chr14 | 72900000 | 73000000 | -0.01 | -0.63 | A>B |
| chr14 | 73000000 | 73100000 | 0.28 | -0.69 | A>B |
| chr14 | 73100000 | 73200000 | 0.28 | -0.63 | A>B |
| chr14 | 73200000 | 73300000 | 0.21 | -0.55 | A>B |
| chr14 | 73300000 | 73400000 | 0.35 | -0.22 | A>A |
| chr14 | 73400000 | 73500000 | 0.67 | 0.38 | A>A |
| chr14 | 73500000 | 73600000 | 0.87 | 0.81 | A>A |
| chr14 | 73600000 | 73700000 | 0.87 | 0.84 | A>A |
| chr14 | 73700000 | 73800000 | 0.84 | 0.86 | A>A |
| chr14 | 73800000 | 73900000 | 0.90 | 0.88 | A>A |
| chr14 | 73900000 | 74000000 | 0.79 | 0.89 | A>A |
| chr14 | 74000000 | 74100000 | 0.93 | 0.90 | A>A |
| chr14 | 74100000 | 74200000 | 0.88 | 0.95 | A>A |
| chr14 | 74200000 | 74300000 | 1.02 | 1.05 | A>A |
| chr14 | 74300000 | 74400000 | 0.94 | 0.95 | A>A |
| chr14 | 74400000 | 74500000 | 0.89 | 0.97 | A>A |
| chr14 | 74500000 | 74600000 | 0.86 | 0.86 | A>A |
| chr14 | 74600000 | 74700000 | 0.90 | 0.93 | A>A |
| chr14 | 74700000 | 74800000 | 0.88 | 0.86 | A>A |
| chr14 | 74800000 | 74900000 | 0.87 | 0.82 | A>A |
| chr14 | 74900000 | 75000000 | 0.86 | 0.83 | A>A |
| chr14 | 75000000 | 75100000 | 0.89 | 0.89 | A>A |
| chr14 | 75100000 | 75200000 | 1.00 | 0.97 | A>A |
| chr14 | 75200000 | 75300000 | 0.94 | 0.89 | A>A |
| chr14 | 75300000 | 75400000 | 0.88 | 0.84 | A>A |
| chr14 | 75400000 | 75500000 | 0.94 | 0.93 | A>A |
| chr14 | 75500000 | 75600000 | 0.75 | 0.85 | A>A |
| chr14 | 75600000 | 75700000 | 0.86 | 0.88 | A>A |
| chr14 | 75700000 | 75800000 | 0.79 | 0.83 | A>A |
| chr14 | 75800000 | 75900000 | 0.78 | 0.77 | A>A |
| chr14 | 75900000 | 76000000 | 0.79 | 0.76 | A>A |
| chr14 | 76000000 | 76100000 | 0.73 | 0.69 | A>A |
| chr14 | 76100000 | 76200000 | 0.66 | 0.69 | A>A |
| chr14 | 76200000 | 76300000 | 0.69 | 0.69 | A>A |
| chr14 | 76300000 | 76400000 | 0.70 | 0.69 | A>A |
| chr14 | 76400000 | 76500000 | 0.62 | 0.62 | A>A |
| chr14 | 76500000 | 76600000 | 0.48 | 0.54 | A>A |
| chr14 | 76600000 | 76700000 | 0.59 | 0.61 | A>A |
| chr14 | 76700000 | 76800000 | 0.48 | 0.45 | A>A |
| chr14 | 76800000 | 76900000 | 0.44 | 0.43 | A>A |

| | | | | | |
|---|---|---|---|---|---|
| chr14 | 76900000 | 77000000 | 0.52 | 0.42 | A>A |
| chr14 | 77000000 | 77100000 | 0.55 | 0.46 | A>A |
| chr14 | 77100000 | 77200000 | 0.63 | 0.55 | A>A |
| chr14 | 77200000 | 77300000 | 0.86 | 0.78 | A>A |
| chr14 | 77300000 | 77400000 | 0.86 | 0.86 | A>A |
| chr14 | 77400000 | 77500000 | 1.24 | 1.12 | A>A |
| chr14 | 77500000 | 77600000 | 0.82 | 0.84 | A>A |
| chr14 | 77600000 | 77700000 | 0.83 | 0.77 | A>A |
| chr14 | 77700000 | 77800000 | 0.95 | 0.86 | A>A |
| chr14 | 77800000 | 77900000 | 0.94 | 0.82 | A>A |
| chr14 | 77900000 | 78000000 | 0.84 | 0.82 | A>A |
| chr14 | 78000000 | 78100000 | 0.87 | 0.72 | A>A |
| chr14 | 78100000 | 78200000 | 0.66 | 0.51 | A>A |
| chr14 | 78200000 | 78300000 | 0.50 | 0.24 | A>A |
| chr14 | 78300000 | 78400000 | 0.08 | -0.25 | B>B |
| chr14 | 78400000 | 78500000 | -0.07 | -0.59 | B>B |
| chr14 | 78500000 | 78600000 | -0.09 | -1.05 | A>B |
| chr14 | 78600000 | 78700000 | -0.20 | -1.06 | A>B |
| chr14 | 78700000 | 78800000 | -0.52 | -1.20 | A>B |
| chr14 | 78800000 | 78900000 | -0.78 | -1.27 | B>B |
| chr14 | 78900000 | 79000000 | -0.80 | -1.23 | B>B |
| chr14 | 79000000 | 79100000 | -0.73 | -1.17 | B>B |
| chr14 | 79100000 | 79200000 | -0.93 | -1.39 | B>B |
| chr14 | 79200000 | 79300000 | -0.96 | -1.42 | B>B |
| chr14 | 79300000 | 79400000 | -0.77 | -1.23 | B>B |
| chr14 | 79400000 | 79500000 | -0.44 | -0.90 | B>B |
| chr14 | 79500000 | 79600000 | -0.53 | -1.08 | B>B |
| chr14 | 79600000 | 79700000 | -0.63 | -1.10 | B>B |
| chr14 | 79700000 | 79800000 | -0.39 | -0.96 | B>B |
| chr14 | 79800000 | 79900000 | -0.42 | -0.96 | B>B |
| chr14 | 79900000 | 80000000 | -0.31 | -0.90 | A>B |
| chr14 | 80000000 | 80100000 | -0.30 | -0.92 | A>B |
| chr14 | 80100000 | 80200000 | -0.27 | -0.93 | A>B |
| chr14 | 80200000 | 80300000 | -0.27 | -0.95 | A>B |
| chr14 | 80300000 | 80400000 | -0.30 | -0.95 | A>B |
| chr14 | 80400000 | 80500000 | -0.31 | -0.91 | A>B |
| chr14 | 80500000 | 80600000 | -0.01 | -0.67 | A>B |
| chr14 | 80600000 | 80700000 | 0.17 | -0.61 | A>B |
| chr14 | 80700000 | 80800000 | 0.21 | -0.50 | A>B |
| chr14 | 80800000 | 80900000 | 0.16 | -0.53 | A>B |
| chr14 | 80900000 | 81000000 | 0.19 | -0.33 | B>B |
| chr14 | 81000000 | 81100000 | 0.27 | -0.09 | A>A |
| chr14 | 81100000 | 81200000 | 0.28 | -0.07 | A>A |
| chr14 | 81200000 | 81300000 | 0.24 | -0.23 | A>A |
| chr14 | 81300000 | 81400000 | 0.07 | -0.39 | B>B |
| chr14 | 81400000 | 81500000 | -0.23 | -0.69 | B>B |
| chr14 | 81500000 | 81600000 | -0.52 | -1.05 | B>B |
| chr14 | 81600000 | 81700000 | 0.00 | -0.39 | B>B |
| chr14 | 81700000 | 81800000 | -0.21 | -0.09 | B>B |
| chr14 | 81800000 | 81900000 | -0.36 | -0.16 | B>B |
| chr14 | 81900000 | 82000000 | -0.08 | -0.17 | B>B |
| chr14 | 82000000 | 82100000 | -0.63 | -1.06 | B>B |
| chr14 | 82100000 | 82200000 | -1.01 | -1.22 | B>B |
| chr14 | 82200000 | 82300000 | -1.17 | -1.34 | B>B |
| chr14 | 82300000 | 82400000 | -1.15 | -1.23 | B>B |
| chr14 | 82400000 | 82500000 | -1.08 | -1.17 | B>B |

| | | | | | |
|---|---|---|---|---|---|
| chr14 | 82500000 | 82600000 | -1.17 | -1.18 | B>B |
| chr14 | 82600000 | 82700000 | -1.11 | -1.18 | B>B |
| chr14 | 82700000 | 82800000 | -1.18 | -1.25 | B>B |
| chr14 | 82800000 | 82900000 | -1.13 | -1.15 | B>B |
| chr14 | 82900000 | 83000000 | -1.16 | -1.14 | B>B |
| chr14 | 83000000 | 83100000 | -1.11 | -1.14 | B>B |
| chr14 | 83100000 | 83200000 | -1.00 | -1.02 | B>B |
| chr14 | 83200000 | 83300000 | -1.02 | -1.04 | B>B |
| chr14 | 83300000 | 83400000 | -1.03 | -1.07 | B>B |
| chr14 | 83400000 | 83500000 | -1.05 | -1.03 | B>B |
| chr14 | 83500000 | 83600000 | -1.02 | -1.00 | B>B |
| chr14 | 83600000 | 83700000 | -0.98 | -0.97 | B>B |
| chr14 | 83700000 | 83800000 | -0.97 | -0.89 | B>B |
| chr14 | 83800000 | 83900000 | -1.02 | -0.85 | B>B |
| chr14 | 83900000 | 84000000 | -0.94 | -0.85 | B>B |
| chr14 | 84000000 | 84100000 | -0.95 | -0.87 | B>B |
| chr14 | 84100000 | 84200000 | -0.90 | -0.87 | B>B |
| chr14 | 84200000 | 84300000 | -0.97 | -0.84 | B>B |
| chr14 | 84300000 | 84400000 | -0.90 | -0.82 | B>B |
| chr14 | 84400000 | 84500000 | -0.89 | -0.78 | B>B |
| chr14 | 84500000 | 84600000 | -0.86 | -0.82 | B>B |
| chr14 | 84600000 | 84700000 | -0.85 | -0.76 | B>B |
| chr14 | 84700000 | 84800000 | -0.84 | -0.78 | B>B |
| chr14 | 84800000 | 84900000 | -0.88 | -0.78 | B>B |
| chr14 | 84900000 | 85000000 | -0.79 | -0.74 | B>B |
| chr14 | 85000000 | 85100000 | -0.79 | -0.76 | B>B |
| chr14 | 85100000 | 85200000 | -0.79 | -0.72 | B>B |
| chr14 | 85200000 | 85300000 | -0.82 | -0.69 | B>B |
| chr14 | 85300000 | 85400000 | -0.77 | -0.64 | B>B |
| chr14 | 85400000 | 85500000 | -0.66 | -0.54 | B>B |
| chr14 | 85500000 | 85600000 | -0.58 | -0.28 | B>B |
| chr14 | 85600000 | 85700000 | -0.29 | -0.02 | B>B |
| chr14 | 85700000 | 85800000 | -0.14 | 0.12 | B>B |
| chr14 | 85800000 | 85900000 | 0.00 | 0.09 | A>A |
| chr14 | 85900000 | 86000000 | 0.06 | 0.09 | A>A |
| chr14 | 86000000 | 86100000 | -0.45 | -0.57 | B>B |
| chr14 | 86100000 | 86200000 | -0.73 | -0.87 | B>B |
| chr14 | 86200000 | 86300000 | -0.85 | -1.00 | B>B |
| chr14 | 86300000 | 86400000 | -0.88 | -1.02 | B>B |
| chr14 | 86400000 | 86500000 | -0.94 | -0.99 | B>B |
| chr14 | 86500000 | 86600000 | -0.97 | -1.11 | B>B |
| chr14 | 86600000 | 86700000 | -1.01 | -1.10 | B>B |
| chr14 | 86700000 | 86800000 | -0.99 | -1.20 | B>B |
| chr14 | 86800000 | 86900000 | -0.95 | -1.22 | B>B |
| chr14 | 86900000 | 87000000 | -0.94 | -1.24 | B>B |
| chr14 | 87000000 | 87100000 | -1.04 | -1.25 | B>B |
| chr14 | 87100000 | 87200000 | -1.08 | -1.24 | B>B |
| chr14 | 87200000 | 87300000 | -1.07 | -1.26 | B>B |
| chr14 | 87300000 | 87400000 | -1.10 | -1.25 | B>B |
| chr14 | 87400000 | 87500000 | -1.20 | -1.25 | B>B |
| chr14 | 87500000 | 87600000 | -1.19 | -1.39 | B>B |
| chr14 | 87600000 | 87700000 | -1.18 | -1.41 | B>B |
| chr14 | 87700000 | 87800000 | -1.23 | -1.43 | B>B |
| chr14 | 87800000 | 87900000 | -1.25 | -1.48 | B>B |
| chr14 | 87900000 | 88000000 | -1.28 | -1.38 | B>B |
| chr14 | 88000000 | 88100000 | -1.22 | -1.46 | B>B |

| | | | | | |
|---|---|---|---|---|---|
| chr14 | 88100000 | 88200000 | -1.29 | -1.46 | B>B |
| chr14 | 88200000 | 88300000 | -1.06 | -1.38 | B>B |
| chr14 | 88300000 | 88400000 | -0.83 | -1.18 | B>B |
| chr14 | 88400000 | 88500000 | -0.02 | -0.29 | B>B |
| chr14 | 88500000 | 88600000 | 0.00 | -0.35 | B>B |
| chr14 | 88600000 | 88700000 | -0.03 | -0.45 | B>B |
| chr14 | 88700000 | 88800000 | -0.06 | -0.70 | A>B |
| chr14 | 88800000 | 88900000 | 0.06 | -0.33 | B>B |
| chr14 | 88900000 | 89000000 | 0.32 | 0.20 | A>A |
| chr14 | 89000000 | 89100000 | 0.61 | 0.41 | A>A |
| chr14 | 89100000 | 89200000 | 0.41 | 0.19 | A>A |
| chr14 | 89200000 | 89300000 | 0.49 | 0.28 | A>A |
| chr14 | 89300000 | 89400000 | 0.38 | 0.22 | A>A |
| chr14 | 89400000 | 89500000 | 0.25 | 0.23 | A>A |
| chr14 | 89500000 | 89600000 | 0.18 | 0.17 | A>A |
| chr14 | 89600000 | 89700000 | 0.25 | 0.23 | A>A |
| chr14 | 89700000 | 89800000 | 0.29 | 0.15 | A>A |
| chr14 | 89800000 | 89900000 | 0.35 | 0.23 | A>A |
| chr14 | 89900000 | 90000000 | 0.19 | -0.02 | A>A |
| chr14 | 90000000 | 90100000 | 0.08 | -0.10 | B>B |
| chr14 | 90100000 | 90200000 | 0.07 | -0.04 | A>A |
| chr14 | 90200000 | 90300000 | 0.10 | -0.08 | A>A |
| chr14 | 90300000 | 90400000 | 0.10 | 0.01 | A>A |
| chr14 | 90400000 | 90500000 | 0.30 | 0.09 | A>A |
| chr14 | 90500000 | 90600000 | 0.31 | -0.12 | A>A |
| chr14 | 90600000 | 90700000 | 0.30 | -0.12 | A>A |
| chr14 | 90700000 | 90800000 | 0.64 | 0.49 | A>A |
| chr14 | 90800000 | 90900000 | 0.82 | 0.71 | A>A |
| chr14 | 90900000 | 91000000 | 0.70 | 0.71 | A>A |
| chr14 | 91000000 | 91100000 | 0.61 | 0.66 | A>A |
| chr14 | 91100000 | 91200000 | 0.63 | 0.63 | A>A |
| chr14 | 91200000 | 91300000 | 0.68 | 0.68 | A>A |
| chr14 | 91300000 | 91400000 | 0.68 | 0.58 | A>A |
| chr14 | 91400000 | 91500000 | 0.77 | 0.58 | A>A |
| chr14 | 91500000 | 91600000 | 0.79 | 0.70 | A>A |
| chr14 | 91600000 | 91700000 | 0.78 | 0.76 | A>A |
| chr14 | 91700000 | 91800000 | 0.85 | 0.84 | A>A |
| chr14 | 91800000 | 91900000 | 0.87 | 0.93 | A>A |
| chr14 | 91900000 | 92000000 | 0.86 | 0.90 | A>A |
| chr14 | 92000000 | 92100000 | 0.66 | 0.78 | A>A |
| chr14 | 92100000 | 92200000 | 0.49 | 0.90 | A>A |
| chr14 | 92200000 | 92300000 | 0.63 | 0.96 | A>A |
| chr14 | 92300000 | 92400000 | 0.69 | 0.91 | A>A |
| chr14 | 92400000 | 92500000 | 0.74 | 0.90 | A>A |
| chr14 | 92500000 | 92600000 | 0.91 | 0.96 | A>A |
| chr14 | 92600000 | 92700000 | 0.86 | 0.76 | A>A |
| chr14 | 92700000 | 92800000 | 0.81 | 0.74 | A>A |
| chr14 | 92800000 | 92900000 | 0.81 | 0.75 | A>A |
| chr14 | 92900000 | 93000000 | 0.89 | 0.89 | A>A |
| chr14 | 93000000 | 93100000 | 0.90 | 0.92 | A>A |
| chr14 | 93100000 | 93200000 | 0.79 | 0.83 | A>A |
| chr14 | 93200000 | 93300000 | 0.81 | 0.81 | A>A |
| chr14 | 93300000 | 93400000 | 0.68 | 0.62 | A>A |
| chr14 | 93400000 | 93500000 | 0.83 | 0.67 | A>A |
| chr14 | 93500000 | 93600000 | 0.79 | 0.72 | A>A |
| chr14 | 93600000 | 93700000 | 0.47 | 0.36 | A>A |

| | | | | | |
|---|---|---|---|---|---|
| chr14 | 93700000 | 93800000 | 0.39 | 0.08 | A>A |
| chr14 | 93800000 | 93900000 | -0.07 | -0.54 | B>B |
| chr14 | 93900000 | 94000000 | -0.25 | -0.68 | B>B |
| chr14 | 94000000 | 94100000 | -0.17 | -0.63 | B>B |
| chr14 | 94100000 | 94200000 | -0.13 | -0.63 | B>B |
| chr14 | 94200000 | 94300000 | 0.02 | -0.57 | A>B |
| chr14 | 94300000 | 94400000 | 0.09 | -0.35 | B>B |
| chr14 | 94400000 | 94500000 | 0.41 | 0.15 | A>A |
| chr14 | 94500000 | 94600000 | 0.39 | 0.25 | A>A |
| chr14 | 94600000 | 94700000 | -0.06 | -0.15 | B>B |
| chr14 | 94700000 | 94800000 | -0.21 | -0.09 | B>B |
| chr14 | 94800000 | 94900000 | 0.04 | 0.21 | A>A |
| chr14 | 94900000 | 95000000 | 0.10 | 0.12 | A>A |
| chr14 | 95000000 | 95100000 | 0.38 | 0.47 | A>A |
| chr14 | 95100000 | 95200000 | 0.32 | 0.27 | A>A |
| chr14 | 95200000 | 95300000 | 0.32 | 0.08 | A>A |
| chr14 | 95300000 | 95400000 | 0.24 | -0.15 | A>A |
| chr14 | 95400000 | 95500000 | 0.28 | -0.12 | A>A |
| chr14 | 95500000 | 95600000 | 0.47 | 0.20 | A>A |
| chr14 | 95600000 | 95700000 | 0.75 | 0.56 | A>A |
| chr14 | 95700000 | 95800000 | 0.88 | 0.78 | A>A |
| chr14 | 95800000 | 95900000 | 0.64 | 0.49 | A>A |
| chr14 | 95900000 | 96000000 | 0.58 | 0.32 | A>A |
| chr14 | 96000000 | 96100000 | 0.13 | -0.20 | B>B |
| chr14 | 96100000 | 96200000 | -0.07 | -0.54 | B>B |
| chr14 | 96200000 | 96300000 | -0.22 | -0.68 | B>B |
| chr14 | 96300000 | 96400000 | -0.21 | -0.66 | B>B |
| chr14 | 96400000 | 96500000 | 0.07 | -0.30 | B>B |
| chr14 | 96500000 | 96600000 | 0.42 | 0.24 | A>A |
| chr14 | 96600000 | 96700000 | 0.69 | 0.53 | A>A |
| chr14 | 96700000 | 96800000 | 0.60 | 0.45 | A>A |
| chr14 | 96800000 | 96900000 | 0.84 | 0.57 | A>A |
| chr14 | 96900000 | 97000000 | 0.74 | 0.45 | A>A |
| chr14 | 97000000 | 97100000 | 0.52 | 0.01 | A>A |
| chr14 | 97100000 | 97200000 | 0.36 | -0.40 | A>B |
| chr14 | 97200000 | 97300000 | 0.44 | -0.12 | A>A |
| chr14 | 97300000 | 97400000 | 0.14 | -0.25 | B>B |
| chr14 | 97400000 | 97500000 | -0.14 | -0.71 | B>B |
| chr14 | 97500000 | 97600000 | -0.12 | -0.73 | A>B |
| chr14 | 97600000 | 97700000 | -0.30 | -0.71 | B>B |
| chr14 | 97700000 | 97800000 | -0.49 | -0.96 | B>B |
| chr14 | 97800000 | 97900000 | -0.56 | -0.94 | B>B |
| chr14 | 97900000 | 98000000 | -0.78 | -1.13 | B>B |
| chr14 | 98000000 | 98100000 | -0.97 | -1.11 | B>B |
| chr14 | 98100000 | 98200000 | -0.97 | -1.18 | B>B |
| chr14 | 98200000 | 98300000 | -1.05 | -1.15 | B>B |
| chr14 | 98300000 | 98400000 | -1.03 | -1.08 | B>B |
| chr14 | 98400000 | 98500000 | -0.99 | -1.08 | B>B |
| chr14 | 98500000 | 98600000 | -1.06 | -1.05 | B>B |
| chr14 | 98600000 | 98700000 | -0.90 | -1.06 | B>B |
| chr14 | 98700000 | 98800000 | -1.03 | -1.01 | B>B |
| chr14 | 98800000 | 98900000 | -1.01 | -1.07 | B>B |
| chr14 | 98900000 | 99000000 | -1.00 | -1.08 | B>B |
| chr14 | 99000000 | 99100000 | -0.91 | -1.01 | B>B |
| chr14 | 99100000 | 99200000 | -0.79 | -0.89 | B>B |
| chr14 | 99200000 | 99300000 | -0.57 | -0.72 | B>B |

| | | | | | |
|---|---|---|---|---|---|
| chr14 | 99300000 | 99400000 | -0.21 | -0.38 | B>B |
| chr14 | 99400000 | 99500000 | -0.09 | -0.29 | B>B |
| chr14 | 99500000 | 99600000 | -0.17 | -0.43 | B>B |
| chr14 | 99600000 | 99700000 | -0.14 | -0.32 | B>B |
| chr14 | 99700000 | 99800000 | 0.02 | -0.05 | B>B |
| chr14 | 99800000 | 99900000 | 0.43 | 0.40 | A>A |
| chr14 | 99900000 | 100000000 | 1.00 | 0.93 | A>A |
| chr14 | 1E+08 | 100100000 | 1.22 | 1.18 | A>A |
| chr14 | 1E+08 | 100200000 | 1.07 | 1.03 | A>A |
| chr14 | 1E+08 | 100300000 | 1.36 | 1.20 | A>A |
| chr14 | 1E+08 | 100400000 | 1.05 | 1.07 | A>A |
| chr14 | 1E+08 | 100500000 | 1.17 | 1.11 | A>A |
| chr14 | 1.01E+08 | 100600000 | 1.21 | 1.18 | A>A |
| chr14 | 1.01E+08 | 100700000 | 1.30 | 1.23 | A>A |
| chr14 | 1.01E+08 | 100800000 | 1.20 | 1.19 | A>A |
| chr14 | 1.01E+08 | 100900000 | 0.94 | 0.98 | A>A |
| chr14 | 1.01E+08 | 101000000 | 0.93 | 0.92 | A>A |
| chr14 | 1.01E+08 | 101100000 | 0.89 | 0.96 | A>A |
| chr14 | 1.01E+08 | 101200000 | 0.70 | 0.78 | A>A |
| chr14 | 1.01E+08 | 101300000 | 0.71 | 0.75 | A>A |
| chr14 | 1.01E+08 | 101400000 | 0.58 | 0.63 | A>A |
| chr14 | 1.01E+08 | 101500000 | 0.73 | 0.76 | A>A |
| chr14 | 1.02E+08 | 101600000 | 0.70 | 0.72 | A>A |
| chr14 | 1.02E+08 | 101700000 | 0.61 | 0.70 | A>A |
| chr14 | 1.02E+08 | 101800000 | 0.62 | 0.68 | A>A |
| chr14 | 1.02E+08 | 101900000 | 0.78 | 0.79 | A>A |
| chr14 | 1.02E+08 | 102000000 | 0.82 | 0.85 | A>A |
| chr14 | 1.02E+08 | 102100000 | 0.72 | 0.73 | A>A |
| chr14 | 1.02E+08 | 102200000 | 0.55 | 0.66 | A>A |
| chr14 | 1.02E+08 | 102300000 | 0.92 | 0.90 | A>A |
| chr14 | 1.02E+08 | 102400000 | 0.97 | 0.89 | A>A |
| chr14 | 1.02E+08 | 102500000 | 0.86 | 0.88 | A>A |
| chr14 | 1.03E+08 | 102600000 | 0.81 | 0.85 | A>A |
| chr14 | 1.03E+08 | 102700000 | 0.88 | 0.87 | A>A |
| chr14 | 1.03E+08 | 102800000 | 0.92 | 0.91 | A>A |
| chr14 | 1.03E+08 | 102900000 | 0.84 | 0.87 | A>A |
| chr14 | 1.03E+08 | 103000000 | 0.82 | 0.83 | A>A |
| chr14 | 1.03E+08 | 103100000 | 0.83 | 0.83 | A>A |
| chr14 | 1.03E+08 | 103200000 | 0.83 | 0.80 | A>A |
| chr14 | 1.03E+08 | 103300000 | 0.83 | 0.84 | A>A |
| chr14 | 1.03E+08 | 103400000 | 0.81 | 0.78 | A>A |
| chr14 | 1.03E+08 | 103500000 | 0.75 | 0.79 | A>A |
| chr14 | 1.04E+08 | 103600000 | 0.72 | 0.75 | A>A |
| chr14 | 1.04E+08 | 103700000 | 0.93 | 0.81 | A>A |
| chr14 | 1.04E+08 | 103800000 | 0.81 | 0.78 | A>A |
| chr14 | 1.04E+08 | 103900000 | 0.84 | 0.85 | A>A |
| chr14 | 1.04E+08 | 104000000 | 0.94 | 0.86 | A>A |
| chr14 | 1.04E+08 | 104100000 | 0.96 | 0.89 | A>A |
| chr14 | 1.04E+08 | 104200000 | 0.99 | 0.89 | A>A |
| chr14 | 1.04E+08 | 104300000 | 0.78 | 0.84 | A>A |
| chr14 | 1.04E+08 | 104400000 | 0.91 | 0.86 | A>A |
| chr14 | 1.04E+08 | 104500000 | 0.72 | 0.77 | A>A |
| chr14 | 1.05E+08 | 104600000 | 0.64 | 0.68 | A>A |
| chr14 | 1.05E+08 | 104700000 | 0.60 | 0.70 | A>A |
| chr14 | 1.05E+08 | 104800000 | 0.64 | 0.74 | A>A |
| chr14 | 1.05E+08 | 104900000 | 0.64 | 0.67 | A>A |

| | | | | | |
|---|---|---|---|---|---|
| chr14 | 1.05E+08 | 105000000 | 0.60 | 0.70 | A>A |
| chr14 | 1.05E+08 | 105100000 | 0.60 | 0.67 | A>A |
| chr14 | 1.05E+08 | 105200000 | 0.95 | 0.87 | A>A |
| chr14 | 1.05E+08 | 105300000 | 1.10 | 1.03 | A>A |
| chr14 | 1.05E+08 | 105400000 | 0.97 | 0.97 | A>A |
| chr14 | 1.05E+08 | 105500000 | 1.18 | 1.13 | A>A |
| chr14 | 1.06E+08 | 105600000 | 1.16 | 1.16 | A>A |
| chr14 | 1.06E+08 | 105700000 | 1.28 | 1.15 | A>A |
| chr14 | 1.06E+08 | 105800000 | 1.22 | 1.23 | A>A |
| chr14 | 1.06E+08 | 105900000 | 1.24 | 1.17 | A>A |
| chr14 | 1.06E+08 | 106000000 | 1.13 | 1.10 | A>A |
| chr14 | 1.06E+08 | 106100000 | 0.00 | 0.00 | NO |
| chr14 | 1.06E+08 | 106200000 | 0.00 | 0.00 | NO |
| chr14 | 1.06E+08 | 106300000 | -0.03 | -0.32 | B>B |
| chr14 | 1.06E+08 | 106400000 | 0.20 | -0.23 | B>B |
| chr14 | 1.06E+08 | 106500000 | 0.00 | 0.00 | NO |
| chr14 | 1.07E+08 | 106600000 | 0.00 | -0.81 | NO |
| chr14 | 1.07E+08 | 106700000 | 0.00 | 0.00 | NO |
| chr14 | 1.07E+08 | 106800000 | -1.02 | -1.69 | A>B |
| chr14 | 1.07E+08 | 106900000 | -1.33 | -1.78 | B>B |
| chr14 | 1.07E+08 | 107000000 | -1.77 | -1.77 | B>B |
| chr14 | 1.07E+08 | 107100000 | -1.64 | -1.49 | B>B |
| chr14 | 1.07E+08 | 107200000 | -1.27 | -0.99 | B>B |
| chr14 | 1.07E+08 | 107300000 | -0.50 | -0.23 | B>B |
| chr14 | 1.07E+08 | 107349540 | 0.00 | 0.00 | NO |
| chr15 | 0 | 100000 | 0.00 | 0.00 | NO |
| chr15 | 100000 | 200000 | 0.00 | 0.00 | NO |
| chr15 | 200000 | 300000 | 0.00 | 0.00 | NO |
| chr15 | 300000 | 400000 | 0.00 | 0.00 | NO |
| chr15 | 400000 | 500000 | 0.00 | 0.00 | NO |
| chr15 | 500000 | 600000 | 0.00 | 0.00 | NO |
| chr15 | 600000 | 700000 | 0.00 | 0.00 | NO |
| chr15 | 700000 | 800000 | 0.00 | 0.00 | NO |
| chr15 | 800000 | 900000 | 0.00 | 0.00 | NO |
| chr15 | 900000 | 1000000 | 0.00 | 0.00 | NO |
| chr15 | 1000000 | 1100000 | 0.00 | 0.00 | NO |
| chr15 | 1100000 | 1200000 | 0.00 | 0.00 | NO |
| chr15 | 1200000 | 1300000 | 0.00 | 0.00 | NO |
| chr15 | 1300000 | 1400000 | 0.00 | 0.00 | NO |
| chr15 | 1400000 | 1500000 | 0.00 | 0.00 | NO |
| chr15 | 1500000 | 1600000 | 0.00 | 0.00 | NO |
| chr15 | 1600000 | 1700000 | 0.00 | 0.00 | NO |
| chr15 | 1700000 | 1800000 | 0.00 | 0.00 | NO |
| chr15 | 1800000 | 1900000 | 0.00 | 0.00 | NO |
| chr15 | 1900000 | 2000000 | 0.00 | 0.00 | NO |
| chr15 | 2000000 | 2100000 | 0.00 | 0.00 | NO |
| chr15 | 2100000 | 2200000 | 0.00 | 0.00 | NO |
| chr15 | 2200000 | 2300000 | 0.00 | 0.00 | NO |
| chr15 | 2300000 | 2400000 | 0.00 | 0.00 | NO |
| chr15 | 2400000 | 2500000 | 0.00 | 0.00 | NO |
| chr15 | 2500000 | 2600000 | 0.00 | 0.00 | NO |
| chr15 | 2600000 | 2700000 | 0.00 | 0.00 | NO |
| chr15 | 2700000 | 2800000 | 0.00 | 0.00 | NO |
| chr15 | 2800000 | 2900000 | 0.00 | 0.00 | NO |
| chr15 | 2900000 | 3000000 | 0.00 | 0.00 | NO |
| chr15 | 3000000 | 3100000 | 0.00 | 0.00 | NO |

| | | | | | |
|---|---|---|---|---|---|
| chr15 | 3100000 | 3200000 | 0.00 | 0.00 | NO |
| chr15 | 3200000 | 3300000 | 0.00 | 0.00 | NO |
| chr15 | 3300000 | 3400000 | 0.00 | 0.00 | NO |
| chr15 | 3400000 | 3500000 | 0.00 | 0.00 | NO |
| chr15 | 3500000 | 3600000 | 0.00 | 0.00 | NO |
| chr15 | 3600000 | 3700000 | 0.00 | 0.00 | NO |
| chr15 | 3700000 | 3800000 | 0.00 | 0.00 | NO |
| chr15 | 3800000 | 3900000 | 0.00 | 0.00 | NO |
| chr15 | 3900000 | 4000000 | 0.00 | 0.00 | NO |
| chr15 | 4000000 | 4100000 | 0.00 | 0.00 | NO |
| chr15 | 4100000 | 4200000 | 0.00 | 0.00 | NO |
| chr15 | 4200000 | 4300000 | 0.00 | 0.00 | NO |
| chr15 | 4300000 | 4400000 | 0.00 | 0.00 | NO |
| chr15 | 4400000 | 4500000 | 0.00 | 0.00 | NO |
| chr15 | 4500000 | 4600000 | 0.00 | 0.00 | NO |
| chr15 | 4600000 | 4700000 | 0.00 | 0.00 | NO |
| chr15 | 4700000 | 4800000 | 0.00 | 0.00 | NO |
| chr15 | 4800000 | 4900000 | 0.00 | 0.00 | NO |
| chr15 | 4900000 | 5000000 | 0.00 | 0.00 | NO |
| chr15 | 5000000 | 5100000 | 0.00 | 0.00 | NO |
| chr15 | 5100000 | 5200000 | 0.00 | 0.00 | NO |
| chr15 | 5200000 | 5300000 | 0.00 | 0.00 | NO |
| chr15 | 5300000 | 5400000 | 0.00 | 0.00 | NO |
| chr15 | 5400000 | 5500000 | 0.00 | 0.00 | NO |
| chr15 | 5500000 | 5600000 | 0.00 | 0.00 | NO |
| chr15 | 5600000 | 5700000 | 0.00 | 0.00 | NO |
| chr15 | 5700000 | 5800000 | 0.00 | 0.00 | NO |
| chr15 | 5800000 | 5900000 | 0.00 | 0.00 | NO |
| chr15 | 5900000 | 6000000 | 0.00 | 0.00 | NO |
| chr15 | 6000000 | 6100000 | 0.00 | 0.00 | NO |
| chr15 | 6100000 | 6200000 | 0.00 | 0.00 | NO |
| chr15 | 6200000 | 6300000 | 0.00 | 0.00 | NO |
| chr15 | 6300000 | 6400000 | 0.00 | 0.00 | NO |
| chr15 | 6400000 | 6500000 | 0.00 | 0.00 | NO |
| chr15 | 6500000 | 6600000 | 0.00 | 0.00 | NO |
| chr15 | 6600000 | 6700000 | 0.00 | 0.00 | NO |
| chr15 | 6700000 | 6800000 | 0.00 | 0.00 | NO |
| chr15 | 6800000 | 6900000 | 0.00 | 0.00 | NO |
| chr15 | 6900000 | 7000000 | 0.00 | 0.00 | NO |
| chr15 | 7000000 | 7100000 | 0.00 | 0.00 | NO |
| chr15 | 7100000 | 7200000 | 0.00 | 0.00 | NO |
| chr15 | 7200000 | 7300000 | 0.00 | 0.00 | NO |
| chr15 | 7300000 | 7400000 | 0.00 | 0.00 | NO |
| chr15 | 7400000 | 7500000 | 0.00 | 0.00 | NO |
| chr15 | 7500000 | 7600000 | 0.00 | 0.00 | NO |
| chr15 | 7600000 | 7700000 | 0.00 | 0.00 | NO |
| chr15 | 7700000 | 7800000 | 0.00 | 0.00 | NO |
| chr15 | 7800000 | 7900000 | 0.00 | 0.00 | NO |
| chr15 | 7900000 | 8000000 | 0.00 | 0.00 | NO |
| chr15 | 8000000 | 8100000 | 0.00 | 0.00 | NO |
| chr15 | 8100000 | 8200000 | 0.00 | 0.00 | NO |
| chr15 | 8200000 | 8300000 | 0.00 | 0.00 | NO |
| chr15 | 8300000 | 8400000 | 0.00 | 0.00 | NO |
| chr15 | 8400000 | 8500000 | 0.00 | 0.00 | NO |
| chr15 | 8500000 | 8600000 | 0.00 | 0.00 | NO |
| chr15 | 8600000 | 8700000 | 0.00 | 0.00 | NO |

| | | | | | |
|---|---|---|---|---|---|
| chr15 | 8700000 | 8800000 | 0.00 | 0.00 | NO |
| chr15 | 8800000 | 8900000 | 0.00 | 0.00 | NO |
| chr15 | 8900000 | 9000000 | 0.00 | 0.00 | NO |
| chr15 | 9000000 | 9100000 | 0.00 | 0.00 | NO |
| chr15 | 9100000 | 9200000 | 0.00 | 0.00 | NO |
| chr15 | 9200000 | 9300000 | 0.00 | 0.00 | NO |
| chr15 | 9300000 | 9400000 | 0.00 | 0.00 | NO |
| chr15 | 9400000 | 9500000 | 0.00 | 0.00 | NO |
| chr15 | 9500000 | 9600000 | 0.00 | 0.00 | NO |
| chr15 | 9600000 | 9700000 | 0.00 | 0.00 | NO |
| chr15 | 9700000 | 9800000 | 0.00 | 0.00 | NO |
| chr15 | 9800000 | 9900000 | 0.00 | 0.00 | NO |
| chr15 | 9900000 | 10000000 | 0.00 | 0.00 | NO |
| chr15 | 10000000 | 10100000 | 0.00 | 0.00 | NO |
| chr15 | 10100000 | 10200000 | 0.00 | 0.00 | NO |
| chr15 | 10200000 | 10300000 | 0.00 | 0.00 | NO |
| chr15 | 10300000 | 10400000 | 0.00 | 0.00 | NO |
| chr15 | 10400000 | 10500000 | 0.00 | 0.00 | NO |
| chr15 | 10500000 | 10600000 | 0.00 | 0.00 | NO |
| chr15 | 10600000 | 10700000 | 0.00 | 0.00 | NO |
| chr15 | 10700000 | 10800000 | 0.00 | 0.00 | NO |
| chr15 | 10800000 | 10900000 | 0.00 | 0.00 | NO |
| chr15 | 10900000 | 11000000 | 0.00 | 0.00 | NO |
| chr15 | 11000000 | 11100000 | 0.00 | 0.00 | NO |
| chr15 | 11100000 | 11200000 | 0.00 | 0.00 | NO |
| chr15 | 11200000 | 11300000 | 0.00 | 0.00 | NO |
| chr15 | 11300000 | 11400000 | 0.00 | 0.00 | NO |
| chr15 | 11400000 | 11500000 | 0.00 | 0.00 | NO |
| chr15 | 11500000 | 11600000 | 0.00 | 0.00 | NO |
| chr15 | 11600000 | 11700000 | 0.00 | 0.00 | NO |
| chr15 | 11700000 | 11800000 | 0.00 | 0.00 | NO |
| chr15 | 11800000 | 11900000 | 0.00 | 0.00 | NO |
| chr15 | 11900000 | 12000000 | 0.00 | 0.00 | NO |
| chr15 | 12000000 | 12100000 | 0.00 | 0.00 | NO |
| chr15 | 12100000 | 12200000 | 0.00 | 0.00 | NO |
| chr15 | 12200000 | 12300000 | 0.00 | 0.00 | NO |
| chr15 | 12300000 | 12400000 | 0.00 | 0.00 | NO |
| chr15 | 12400000 | 12500000 | 0.00 | 0.00 | NO |
| chr15 | 12500000 | 12600000 | 0.00 | 0.00 | NO |
| chr15 | 12600000 | 12700000 | 0.00 | 0.00 | NO |
| chr15 | 12700000 | 12800000 | 0.00 | 0.00 | NO |
| chr15 | 12800000 | 12900000 | 0.00 | 0.00 | NO |
| chr15 | 12900000 | 13000000 | 0.00 | 0.00 | NO |
| chr15 | 13000000 | 13100000 | 0.00 | 0.00 | NO |
| chr15 | 13100000 | 13200000 | 0.00 | 0.00 | NO |
| chr15 | 13200000 | 13300000 | 0.00 | 0.00 | NO |
| chr15 | 13300000 | 13400000 | 0.00 | 0.00 | NO |
| chr15 | 13400000 | 13500000 | 0.00 | 0.00 | NO |
| chr15 | 13500000 | 13600000 | 0.00 | 0.00 | NO |
| chr15 | 13600000 | 13700000 | 0.00 | 0.00 | NO |
| chr15 | 13700000 | 13800000 | 0.00 | 0.00 | NO |
| chr15 | 13800000 | 13900000 | 0.00 | 0.00 | NO |
| chr15 | 13900000 | 14000000 | 0.00 | 0.00 | NO |
| chr15 | 14000000 | 14100000 | 0.00 | 0.00 | NO |
| chr15 | 14100000 | 14200000 | 0.00 | 0.00 | NO |
| chr15 | 14200000 | 14300000 | 0.00 | 0.00 | NO |

| | | | | | |
|---|---|---|---|---|---|
| chr15 | 14300000 | 14400000 | 0.00 | 0.00 | NO |
| chr15 | 14400000 | 14500000 | 0.00 | 0.00 | NO |
| chr15 | 14500000 | 14600000 | 0.00 | 0.00 | NO |
| chr15 | 14600000 | 14700000 | 0.00 | 0.00 | NO |
| chr15 | 14700000 | 14800000 | 0.00 | 0.00 | NO |
| chr15 | 14800000 | 14900000 | 0.00 | 0.00 | NO |
| chr15 | 14900000 | 15000000 | 0.00 | 0.00 | NO |
| chr15 | 15000000 | 15100000 | 0.00 | 0.00 | NO |
| chr15 | 15100000 | 15200000 | 0.00 | 0.00 | NO |
| chr15 | 15200000 | 15300000 | 0.00 | 0.00 | NO |
| chr15 | 15300000 | 15400000 | 0.00 | 0.00 | NO |
| chr15 | 15400000 | 15500000 | 0.00 | 0.00 | NO |
| chr15 | 15500000 | 15600000 | 0.00 | 0.00 | NO |
| chr15 | 15600000 | 15700000 | 0.00 | 0.00 | NO |
| chr15 | 15700000 | 15800000 | 0.00 | 0.00 | NO |
| chr15 | 15800000 | 15900000 | 0.00 | 0.00 | NO |
| chr15 | 15900000 | 16000000 | 0.00 | 0.00 | NO |
| chr15 | 16000000 | 16100000 | 0.00 | 0.00 | NO |
| chr15 | 16100000 | 16200000 | 0.00 | 0.00 | NO |
| chr15 | 16200000 | 16300000 | 0.00 | 0.00 | NO |
| chr15 | 16300000 | 16400000 | 0.00 | 0.00 | NO |
| chr15 | 16400000 | 16500000 | 0.00 | 0.00 | NO |
| chr15 | 16500000 | 16600000 | 0.00 | 0.00 | NO |
| chr15 | 16600000 | 16700000 | 0.00 | 0.00 | NO |
| chr15 | 16700000 | 16800000 | 0.00 | 0.00 | NO |
| chr15 | 16800000 | 16900000 | 0.00 | 0.00 | NO |
| chr15 | 16900000 | 17000000 | 0.00 | 0.00 | NO |
| chr15 | 17000000 | 17100000 | 0.00 | 0.00 | NO |
| chr15 | 17100000 | 17200000 | 0.00 | 0.00 | NO |
| chr15 | 17200000 | 17300000 | 0.00 | 0.00 | NO |
| chr15 | 17300000 | 17400000 | 0.00 | 0.00 | NO |
| chr15 | 17400000 | 17500000 | 0.00 | 0.00 | NO |
| chr15 | 17500000 | 17600000 | 0.00 | 0.00 | NO |
| chr15 | 17600000 | 17700000 | 0.00 | 0.00 | NO |
| chr15 | 17700000 | 17800000 | 0.00 | 0.00 | NO |
| chr15 | 17800000 | 17900000 | 0.00 | 0.00 | NO |
| chr15 | 17900000 | 18000000 | 0.00 | 0.00 | NO |
| chr15 | 18000000 | 18100000 | 0.00 | 0.00 | NO |
| chr15 | 18100000 | 18200000 | 0.00 | 0.00 | NO |
| chr15 | 18200000 | 18300000 | 0.00 | 0.00 | NO |
| chr15 | 18300000 | 18400000 | 0.00 | 0.00 | NO |
| chr15 | 18400000 | 18500000 | 0.00 | 0.00 | NO |
| chr15 | 18500000 | 18600000 | 0.00 | 0.00 | NO |
| chr15 | 18600000 | 18700000 | 0.00 | 0.00 | NO |
| chr15 | 18700000 | 18800000 | 0.00 | 0.00 | NO |
| chr15 | 18800000 | 18900000 | 0.00 | 0.00 | NO |
| chr15 | 18900000 | 19000000 | 0.00 | 0.00 | NO |
| chr15 | 19000000 | 19100000 | 0.00 | 0.00 | NO |
| chr15 | 19100000 | 19200000 | 0.00 | 0.00 | NO |
| chr15 | 19200000 | 19300000 | 0.00 | 0.00 | NO |
| chr15 | 19300000 | 19400000 | 0.00 | 0.00 | NO |
| chr15 | 19400000 | 19500000 | 0.00 | 0.00 | NO |
| chr15 | 19500000 | 19600000 | 0.00 | 0.00 | NO |
| chr15 | 19600000 | 19700000 | 0.00 | 0.00 | NO |
| chr15 | 19700000 | 19800000 | 0.00 | 0.00 | NO |
| chr15 | 19800000 | 19900000 | 0.00 | 0.00 | NO |

| | | | | | |
|---|---|---|---|---|---|
| chr15 | 19900000 | 20000000 | 0.00 | 0.00 | NO |
| chr15 | 20000000 | 20100000 | 0.00 | 0.00 | NO |
| chr15 | 20100000 | 20200000 | 0.00 | 0.00 | NO |
| chr15 | 20200000 | 20300000 | 0.00 | 0.00 | NO |
| chr15 | 20300000 | 20400000 | 0.00 | 0.00 | NO |
| chr15 | 20400000 | 20500000 | 0.00 | 0.00 | NO |
| chr15 | 20500000 | 20600000 | 0.00 | -0.98 | NO |
| chr15 | 20600000 | 20700000 | 0.00 | 0.00 | NO |
| chr15 | 20700000 | 20800000 | 0.00 | 0.00 | NO |
| chr15 | 20800000 | 20900000 | 0.00 | 0.00 | NO |
| chr15 | 20900000 | 21000000 | 0.00 | 0.00 | NO |
| chr15 | 21000000 | 21100000 | 0.00 | 0.00 | NO |
| chr15 | 21100000 | 21200000 | 0.00 | 0.00 | NO |
| chr15 | 21200000 | 21300000 | 0.00 | 0.00 | NO |
| chr15 | 21300000 | 21400000 | 0.00 | 0.00 | NO |
| chr15 | 21400000 | 21500000 | 0.00 | 0.00 | NO |
| chr15 | 21500000 | 21600000 | 0.00 | 0.00 | NO |
| chr15 | 21600000 | 21700000 | 0.00 | 0.00 | NO |
| chr15 | 21700000 | 21800000 | 0.00 | 0.00 | NO |
| chr15 | 21800000 | 21900000 | 0.00 | 0.00 | NO |
| chr15 | 21900000 | 22000000 | 0.00 | 0.00 | NO |
| chr15 | 22000000 | 22100000 | 0.00 | 0.00 | NO |
| chr15 | 22100000 | 22200000 | 0.00 | 0.00 | NO |
| chr15 | 22200000 | 22300000 | 0.00 | 0.00 | NO |
| chr15 | 22300000 | 22400000 | -0.48 | -0.96 | B>B |
| chr15 | 22400000 | 22500000 | -0.52 | -1.02 | B>B |
| chr15 | 22500000 | 22600000 | -0.36 | -0.66 | B>B |
| chr15 | 22600000 | 22700000 | 0.00 | 0.00 | NO |
| chr15 | 22700000 | 22800000 | 0.00 | 0.00 | NO |
| chr15 | 22800000 | 22900000 | 0.45 | 0.40 | A>A |
| chr15 | 22900000 | 23000000 | 0.71 | 0.67 | A>A |
| chr15 | 23000000 | 23100000 | 0.75 | 0.66 | A>A |
| chr15 | 23100000 | 23200000 | 0.00 | 0.00 | NO |
| chr15 | 23200000 | 23300000 | 0.00 | 0.00 | NO |
| chr15 | 23300000 | 23400000 | 0.00 | 0.00 | NO |
| chr15 | 23400000 | 23500000 | 0.00 | 0.00 | NO |
| chr15 | 23500000 | 23600000 | 0.00 | 0.00 | NO |
| chr15 | 23600000 | 23700000 | 0.00 | 0.00 | NO |
| chr15 | 23700000 | 23800000 | -0.22 | -0.35 | B>B |
| chr15 | 23800000 | 23900000 | -0.40 | -0.56 | B>B |
| chr15 | 23900000 | 24000000 | 0.00 | 0.00 | NO |
| chr15 | 24000000 | 24100000 | -0.49 | -0.92 | B>B |
| chr15 | 24100000 | 24200000 | -0.51 | -1.03 | B>B |
| chr15 | 24200000 | 24300000 | -0.60 | -1.30 | A>B |
| chr15 | 24300000 | 24400000 | 0.00 | 0.00 | NO |
| chr15 | 24400000 | 24500000 | 0.00 | 0.00 | NO |
| chr15 | 24500000 | 24600000 | 0.00 | 0.00 | NO |
| chr15 | 24600000 | 24700000 | 0.00 | 0.00 | NO |
| chr15 | 24700000 | 24800000 | 0.00 | 0.00 | NO |
| chr15 | 24800000 | 24900000 | -0.61 | -1.25 | A>B |
| chr15 | 24900000 | 25000000 | -0.48 | -0.58 | B>B |
| chr15 | 25000000 | 25100000 | 0.00 | 0.00 | NO |
| chr15 | 25100000 | 25200000 | 0.00 | 0.00 | NO |
| chr15 | 25200000 | 25300000 | 0.00 | 0.00 | NO |
| chr15 | 25300000 | 25400000 | 0.00 | 0.00 | NO |
| chr15 | 25400000 | 25500000 | -0.37 | -0.61 | B>B |

| | | | | | |
|---|---|---|---|---|---|
| chr15 | 25500000 | 25600000 | 0.00 | 0.00 | NO |
| chr15 | 25600000 | 25700000 | 0.00 | 0.03 | NO |
| chr15 | 25700000 | 25800000 | 0.00 | -0.32 | NO |
| chr15 | 25800000 | 25900000 | 0.00 | -0.44 | NO |
| chr15 | 25900000 | 26000000 | 0.13 | -0.10 | A>A |
| chr15 | 26000000 | 26100000 | 0.00 | 0.00 | NO |
| chr15 | 26100000 | 26200000 | 0.00 | 0.02 | NO |
| chr15 | 26200000 | 26300000 | 0.00 | -0.38 | NO |
| chr15 | 26300000 | 26400000 | 0.00 | -0.45 | NO |
| chr15 | 26400000 | 26500000 | 0.00 | -0.63 | NO |
| chr15 | 26500000 | 26600000 | 0.00 | -0.68 | NO |
| chr15 | 26600000 | 26700000 | 0.00 | -0.76 | NO |
| chr15 | 26700000 | 26800000 | 0.51 | -0.07 | A>A |
| chr15 | 26800000 | 26900000 | 0.50 | 0.02 | A>A |
| chr15 | 26900000 | 27000000 | 0.52 | -0.06 | A>B |
| chr15 | 27000000 | 27100000 | 0.47 | -0.02 | A>A |
| chr15 | 27100000 | 27200000 | 0.05 | -0.51 | B>B |
| chr15 | 27200000 | 27300000 | -0.17 | -1.02 | A>B |
| chr15 | 27300000 | 27400000 | -0.60 | -1.67 | A>B |
| chr15 | 27400000 | 27500000 | -0.65 | -1.76 | A>B |
| chr15 | 27500000 | 27600000 | -0.67 | -1.74 | A>B |
| chr15 | 27600000 | 27700000 | -0.73 | -1.71 | A>B |
| chr15 | 27700000 | 27800000 | -0.83 | -1.76 | A>B |
| chr15 | 27800000 | 27900000 | -0.89 | -1.75 | A>B |
| chr15 | 27900000 | 28000000 | -0.77 | -1.71 | A>B |
| chr15 | 28000000 | 28100000 | -0.94 | -1.79 | A>B |
| chr15 | 28100000 | 28200000 | -0.90 | -1.76 | A>B |
| chr15 | 28200000 | 28300000 | -0.63 | -1.57 | A>B |
| chr15 | 28300000 | 28400000 | 0.09 | -0.47 | B>B |
| chr15 | 28400000 | 28500000 | 0.00 | 0.00 | NO |
| chr15 | 28500000 | 28600000 | 0.00 | 0.00 | NO |
| chr15 | 28600000 | 28700000 | 0.00 | 0.00 | NO |
| chr15 | 28700000 | 28800000 | 0.00 | 0.00 | NO |
| chr15 | 28800000 | 28900000 | 0.00 | 0.00 | NO |
| chr15 | 28900000 | 29000000 | 0.00 | 0.00 | NO |
| chr15 | 29000000 | 29100000 | 0.00 | 0.00 | NO |
| chr15 | 29100000 | 29200000 | 0.00 | 0.64 | NO |
| chr15 | 29200000 | 29300000 | 0.53 | 0.58 | A>A |
| chr15 | 29300000 | 29400000 | 0.36 | 0.48 | A>A |
| chr15 | 29400000 | 29500000 | 0.12 | 0.36 | A>A |
| chr15 | 29500000 | 29600000 | 0.04 | 0.38 | A>A |
| chr15 | 29600000 | 29700000 | -0.01 | 0.36 | A>A |
| chr15 | 29700000 | 29800000 | 0.04 | 0.40 | A>A |
| chr15 | 29800000 | 29900000 | 0.23 | 0.49 | A>A |
| chr15 | 29900000 | 30000000 | 0.54 | 0.62 | A>A |
| chr15 | 30000000 | 30100000 | 0.77 | 0.75 | A>A |
| chr15 | 30100000 | 30200000 | 0.96 | 0.83 | A>A |
| chr15 | 30200000 | 30300000 | 0.76 | 0.69 | A>A |
| chr15 | 30300000 | 30400000 | 0.67 | 0.57 | A>A |
| chr15 | 30400000 | 30500000 | 0.00 | 0.00 | NO |
| chr15 | 30500000 | 30600000 | 0.00 | 0.00 | NO |
| chr15 | 30600000 | 30700000 | 0.00 | 0.00 | NO |
| chr15 | 30700000 | 30800000 | 0.00 | 0.00 | NO |
| chr15 | 30800000 | 30900000 | 0.00 | 0.00 | NO |
| chr15 | 30900000 | 31000000 | 0.00 | 0.00 | NO |
| chr15 | 31000000 | 31100000 | 0.00 | 0.00 | NO |

| | | | | | |
|---|---|---|---|---|---|
| chr15 | 31100000 | 31200000 | 0.56 | 0.31 | A>A |
| chr15 | 31200000 | 31300000 | 0.75 | 0.64 | A>A |
| chr15 | 31300000 | 31400000 | 0.69 | 0.59 | A>A |
| chr15 | 31400000 | 31500000 | 0.54 | 0.35 | A>A |
| chr15 | 31500000 | 31600000 | 0.80 | 0.51 | A>A |
| chr15 | 31600000 | 31700000 | 0.79 | 0.43 | A>A |
| chr15 | 31700000 | 31800000 | 0.52 | 0.06 | A>A |
| chr15 | 31800000 | 31900000 | -0.01 | -1.11 | A>B |
| chr15 | 31900000 | 32000000 | 0.00 | 0.00 | NO |
| chr15 | 32000000 | 32100000 | -0.28 | -1.72 | A>B |
| chr15 | 32100000 | 32200000 | -0.30 | -1.68 | A>B |
| chr15 | 32200000 | 32300000 | -0.40 | -1.58 | A>B |
| chr15 | 32300000 | 32400000 | -0.51 | -1.80 | A>B |
| chr15 | 32400000 | 32500000 | 0.00 | 0.00 | NO |
| chr15 | 32500000 | 32600000 | 0.00 | 0.00 | NO |
| chr15 | 32600000 | 32700000 | 0.00 | 0.00 | NO |
| chr15 | 32700000 | 32800000 | 0.00 | 0.00 | NO |
| chr15 | 32800000 | 32900000 | 0.00 | 0.00 | NO |
| chr15 | 32900000 | 33000000 | 0.65 | 0.51 | A>A |
| chr15 | 33000000 | 33100000 | 0.93 | 0.55 | A>A |
| chr15 | 33100000 | 33200000 | 0.91 | 0.55 | A>A |
| chr15 | 33200000 | 33300000 | 0.75 | 0.46 | A>A |
| chr15 | 33300000 | 33400000 | 0.54 | 0.19 | A>A |
| chr15 | 33400000 | 33500000 | 0.38 | 0.05 | A>A |
| chr15 | 33500000 | 33600000 | -0.06 | -0.30 | B>B |
| chr15 | 33600000 | 33700000 | -0.49 | -0.90 | B>B |
| chr15 | 33700000 | 33800000 | -0.63 | -0.96 | B>B |
| chr15 | 33800000 | 33900000 | -0.70 | -0.98 | B>B |
| chr15 | 33900000 | 34000000 | -0.43 | -0.78 | B>B |
| chr15 | 34000000 | 34100000 | -0.20 | -0.60 | B>B |
| chr15 | 34100000 | 34200000 | -0.12 | -0.57 | B>B |
| chr15 | 34200000 | 34300000 | 0.06 | -0.18 | B>B |
| chr15 | 34300000 | 34400000 | 0.44 | 0.33 | A>A |
| chr15 | 34400000 | 34500000 | 0.61 | 0.84 | A>A |
| chr15 | 34500000 | 34600000 | 0.64 | 0.58 | A>A |
| chr15 | 34600000 | 34700000 | 0.62 | 0.45 | A>A |
| chr15 | 34700000 | 34800000 | 0.00 | 0.00 | NO |
| chr15 | 34800000 | 34900000 | 0.00 | 0.00 | NO |
| chr15 | 34900000 | 35000000 | -0.18 | -0.38 | B>B |
| chr15 | 35000000 | 35100000 | -0.13 | -0.56 | B>B |
| chr15 | 35100000 | 35200000 | 0.32 | -0.10 | A>A |
| chr15 | 35200000 | 35300000 | 0.64 | 0.41 | A>A |
| chr15 | 35300000 | 35400000 | 0.34 | 0.18 | A>A |
| chr15 | 35400000 | 35500000 | -0.17 | -0.09 | B>B |
| chr15 | 35500000 | 35600000 | -0.02 | 0.05 | A>A |
| chr15 | 35600000 | 35700000 | 0.05 | 0.12 | A>A |
| chr15 | 35700000 | 35800000 | 0.00 | 0.27 | NO |
| chr15 | 35800000 | 35900000 | 0.00 | 0.31 | NO |
| chr15 | 35900000 | 36000000 | 0.00 | 0.00 | NO |
| chr15 | 36000000 | 36100000 | 0.00 | 0.29 | NO |
| chr15 | 36100000 | 36200000 | 0.00 | 0.32 | NO |
| chr15 | 36200000 | 36300000 | 0.00 | 0.31 | NO |
| chr15 | 36300000 | 36400000 | 0.00 | 0.34 | NO |
| chr15 | 36400000 | 36500000 | 0.00 | 0.00 | NO |
| chr15 | 36500000 | 36600000 | 0.00 | 0.00 | NO |
| chr15 | 36600000 | 36700000 | 0.00 | 0.39 | NO |

| | | | | | |
|---|---|---|---|---|---|
| chr15 | 36700000 | 36800000 | 0.00 | 0.40 | NO |
| chr15 | 36800000 | 36900000 | 0.00 | 0.32 | NO |
| chr15 | 36900000 | 37000000 | 0.00 | 0.00 | NO |
| chr15 | 37000000 | 37100000 | 0.00 | 0.38 | NO |
| chr15 | 37100000 | 37200000 | 0.19 | 0.35 | A>A |
| chr15 | 37200000 | 37300000 | 0.00 | 0.33 | NO |
| chr15 | 37300000 | 37400000 | 0.21 | 0.35 | A>A |
| chr15 | 37400000 | 37500000 | 0.00 | 0.17 | NO |
| chr15 | 37500000 | 37600000 | 0.00 | 0.00 | NO |
| chr15 | 37600000 | 37700000 | 0.00 | 0.00 | NO |
| chr15 | 37700000 | 37800000 | 0.00 | 0.00 | NO |
| chr15 | 37800000 | 37900000 | 0.00 | 0.00 | NO |
| chr15 | 37900000 | 38000000 | 0.00 | 0.00 | NO |
| chr15 | 38000000 | 38100000 | 0.00 | 0.00 | NO |
| chr15 | 38100000 | 38200000 | 0.00 | -0.06 | NO |
| chr15 | 38200000 | 38300000 | 0.00 | 0.00 | NO |
| chr15 | 38300000 | 38400000 | 0.00 | 0.27 | NO |
| chr15 | 38400000 | 38500000 | 0.00 | 0.00 | NO |
| chr15 | 38500000 | 38600000 | 0.00 | 0.00 | NO |
| chr15 | 38600000 | 38700000 | 0.00 | 0.32 | NO |
| chr15 | 38700000 | 38800000 | 0.00 | 0.29 | NO |
| chr15 | 38800000 | 38900000 | 0.00 | 0.00 | NO |
| chr15 | 38900000 | 39000000 | 0.00 | 0.27 | NO |
| chr15 | 39000000 | 39100000 | 0.00 | 0.00 | NO |
| chr15 | 39100000 | 39200000 | 0.00 | 0.00 | NO |
| chr15 | 39200000 | 39300000 | 0.00 | 0.00 | NO |
| chr15 | 39300000 | 39400000 | 0.00 | 0.00 | NO |
| chr15 | 39400000 | 39500000 | 0.00 | 0.56 | NO |
| chr15 | 39500000 | 39600000 | 0.00 | 0.00 | NO |
| chr15 | 39600000 | 39700000 | 0.00 | 0.00 | NO |
| chr15 | 39700000 | 39800000 | 0.00 | 0.00 | NO |
| chr15 | 39800000 | 39900000 | 0.00 | 0.54 | NO |
| chr15 | 39900000 | 40000000 | 0.00 | 0.00 | NO |
| chr15 | 40000000 | 40100000 | 0.00 | 0.59 | NO |
| chr15 | 40100000 | 40200000 | 0.00 | 0.00 | NO |
| chr15 | 40200000 | 40300000 | 0.00 | 0.00 | NO |
| chr15 | 40300000 | 40400000 | 0.00 | 0.77 | NO |
| chr15 | 40400000 | 40500000 | 0.00 | 0.00 | NO |
| chr15 | 40500000 | 40600000 | 0.00 | 0.00 | NO |
| chr15 | 40600000 | 40700000 | 0.00 | 0.00 | NO |
| chr15 | 40700000 | 40800000 | 0.00 | 0.00 | NO |
| chr15 | 40800000 | 40900000 | 0.00 | 0.00 | NO |
| chr15 | 40900000 | 41000000 | 0.00 | 0.00 | NO |
| chr15 | 41000000 | 41100000 | 0.00 | 0.00 | NO |
| chr15 | 41100000 | 41200000 | 0.00 | 0.00 | NO |
| chr15 | 41200000 | 41300000 | 0.00 | 0.00 | NO |
| chr15 | 41300000 | 41400000 | 0.00 | 0.00 | NO |
| chr15 | 41400000 | 41500000 | 0.00 | 0.00 | NO |
| chr15 | 41500000 | 41600000 | 0.00 | 0.00 | NO |
| chr15 | 41600000 | 41700000 | 0.00 | 0.00 | NO |
| chr15 | 41700000 | 41800000 | 0.00 | 0.00 | NO |
| chr15 | 41800000 | 41900000 | 0.00 | 0.00 | NO |
| chr15 | 41900000 | 42000000 | 0.00 | 0.00 | NO |
| chr15 | 42000000 | 42100000 | 0.00 | 0.00 | NO |
| chr15 | 42100000 | 42200000 | 0.00 | 0.72 | NO |
| chr15 | 42200000 | 42300000 | 0.00 | 0.78 | NO |

| | | | | | |
|---|---|---|---|---|---|
| chr15 | 42300000 | 42400000 | 0.95 | 0.71 | A>A |
| chr15 | 42400000 | 42500000 | 0.00 | 0.00 | NO |
| chr15 | 42500000 | 42600000 | 0.00 | 0.00 | NO |
| chr15 | 42600000 | 42700000 | 0.00 | 0.00 | NO |
| chr15 | 42700000 | 42800000 | 0.00 | 0.00 | NO |
| chr15 | 42800000 | 42900000 | 0.00 | 0.00 | NO |
| chr15 | 42900000 | 43000000 | 0.00 | 0.00 | NO |
| chr15 | 43000000 | 43100000 | 0.00 | 0.00 | NO |
| chr15 | 43100000 | 43200000 | 0.00 | 0.00 | NO |
| chr15 | 43200000 | 43300000 | 0.00 | 0.00 | NO |
| chr15 | 43300000 | 43400000 | 0.00 | 0.00 | NO |
| chr15 | 43400000 | 43500000 | 0.00 | 0.00 | NO |
| chr15 | 43500000 | 43600000 | 0.00 | 0.00 | NO |
| chr15 | 43600000 | 43700000 | 0.00 | 0.00 | NO |
| chr15 | 43700000 | 43800000 | 0.00 | 0.00 | NO |
| chr15 | 43800000 | 43900000 | 0.00 | 0.00 | NO |
| chr15 | 43900000 | 44000000 | 0.00 | 0.00 | NO |
| chr15 | 44000000 | 44100000 | 0.00 | 0.00 | NO |
| chr15 | 44100000 | 44200000 | 0.00 | 0.00 | NO |
| chr15 | 44200000 | 44300000 | 0.00 | 0.00 | NO |
| chr15 | 44300000 | 44400000 | 0.00 | 0.00 | NO |
| chr15 | 44400000 | 44500000 | 0.00 | 0.00 | NO |
| chr15 | 44500000 | 44600000 | 0.00 | 0.00 | NO |
| chr15 | 44600000 | 44700000 | 0.00 | 0.00 | NO |
| chr15 | 44700000 | 44800000 | 0.00 | 0.00 | NO |
| chr15 | 44800000 | 44900000 | 0.00 | 0.00 | NO |
| chr15 | 44900000 | 45000000 | 0.00 | 0.00 | NO |
| chr15 | 45000000 | 45100000 | 0.00 | 0.00 | NO |
| chr15 | 45100000 | 45200000 | 0.00 | 0.00 | NO |
| chr15 | 45200000 | 45300000 | 0.00 | 0.00 | NO |
| chr15 | 45300000 | 45400000 | 0.00 | 0.00 | NO |
| chr15 | 45400000 | 45500000 | 0.00 | 0.92 | NO |
| chr15 | 45500000 | 45600000 | 0.00 | 0.00 | NO |
| chr15 | 45600000 | 45700000 | 0.00 | 0.00 | NO |
| chr15 | 45700000 | 45800000 | 0.00 | 0.00 | NO |
| chr15 | 45800000 | 45900000 | 0.00 | 0.00 | NO |
| chr15 | 45900000 | 46000000 | 0.00 | 0.02 | NO |
| chr15 | 46000000 | 46100000 | 0.00 | -0.67 | NO |
| chr15 | 46100000 | 46200000 | 0.00 | -0.76 | NO |
| chr15 | 46200000 | 46300000 | 0.00 | -0.85 | NO |
| chr15 | 46300000 | 46400000 | 0.00 | -0.91 | NO |
| chr15 | 46400000 | 46500000 | 0.00 | -1.01 | NO |
| chr15 | 46500000 | 46600000 | 0.00 | -0.98 | NO |
| chr15 | 46600000 | 46700000 | 0.00 | -0.99 | NO |
| chr15 | 46700000 | 46800000 | 0.00 | -1.06 | NO |
| chr15 | 46800000 | 46900000 | 0.00 | -1.00 | NO |
| chr15 | 46900000 | 47000000 | 0.00 | -1.00 | NO |
| chr15 | 47000000 | 47100000 | 0.00 | 0.00 | NO |
| chr15 | 47100000 | 47200000 | 0.00 | -1.01 | NO |
| chr15 | 47200000 | 47300000 | 0.00 | -0.97 | NO |
| chr15 | 47300000 | 47400000 | 0.00 | 0.00 | NO |
| chr15 | 47400000 | 47500000 | 0.00 | -0.87 | NO |
| chr15 | 47500000 | 47600000 | 0.00 | -0.90 | NO |
| chr15 | 47600000 | 47700000 | 0.00 | -1.02 | NO |
| chr15 | 47700000 | 47800000 | 0.00 | -0.95 | NO |
| chr15 | 47800000 | 47900000 | 0.00 | -0.93 | NO |

| | | | | | |
|---|---|---|---|---|---|
| chr15 | 47900000 | 48000000 | 0.00 | -0.90 | NO |
| chr15 | 48000000 | 48100000 | -0.51 | -0.81 | B>B |
| chr15 | 48100000 | 48200000 | 0.00 | -0.85 | NO |
| chr15 | 48200000 | 48300000 | -0.50 | -0.95 | B>B |
| chr15 | 48300000 | 48400000 | 0.00 | -0.87 | NO |
| chr15 | 48400000 | 48500000 | 0.00 | -0.53 | NO |
| chr15 | 48500000 | 48600000 | -0.23 | -0.58 | B>B |
| chr15 | 48600000 | 48700000 | 0.00 | -0.11 | NO |
| chr15 | 48700000 | 48800000 | 0.00 | -0.03 | NO |
| chr15 | 48800000 | 48900000 | 0.00 | -0.02 | NO |
| chr15 | 48900000 | 49000000 | 0.00 | 0.07 | NO |
| chr15 | 49000000 | 49100000 | 0.00 | 0.24 | NO |
| chr15 | 49100000 | 49200000 | 0.00 | 0.16 | NO |
| chr15 | 49200000 | 49300000 | 0.00 | 0.00 | NO |
| chr15 | 49300000 | 49400000 | 0.00 | 0.00 | NO |
| chr15 | 49400000 | 49500000 | 0.00 | 0.00 | NO |
| chr15 | 49500000 | 49600000 | 0.00 | 0.00 | NO |
| chr15 | 49600000 | 49700000 | 0.00 | 0.00 | NO |
| chr15 | 49700000 | 49800000 | 0.00 | 0.00 | NO |
| chr15 | 49800000 | 49900000 | 0.00 | 0.00 | NO |
| chr15 | 49900000 | 50000000 | 0.00 | 0.00 | NO |
| chr15 | 50000000 | 50100000 | 0.00 | 0.00 | NO |
| chr15 | 50100000 | 50200000 | 0.14 | -0.10 | A>A |
| chr15 | 50200000 | 50300000 | 0.05 | -0.34 | B>B |
| chr15 | 50300000 | 50400000 | 0.04 | -0.41 | B>B |
| chr15 | 50400000 | 50500000 | 0.25 | -0.12 | A>A |
| chr15 | 50500000 | 50600000 | 0.43 | 0.08 | A>A |
| chr15 | 50600000 | 50700000 | 0.69 | 0.38 | A>A |
| chr15 | 50700000 | 50800000 | 0.84 | 0.70 | A>A |
| chr15 | 50800000 | 50900000 | 0.83 | 0.75 | A>A |
| chr15 | 50900000 | 51000000 | 0.84 | 0.77 | A>A |
| chr15 | 51000000 | 51100000 | 0.89 | 0.68 | A>A |
| chr15 | 51100000 | 51200000 | 0.72 | 0.62 | A>A |
| chr15 | 51200000 | 51300000 | 0.64 | 0.44 | A>A |
| chr15 | 51300000 | 51400000 | 0.60 | 0.20 | A>A |
| chr15 | 51400000 | 51500000 | 0.40 | 0.03 | A>A |
| chr15 | 51500000 | 51600000 | 0.44 | -0.11 | A>A |
| chr15 | 51600000 | 51700000 | 0.49 | -0.01 | A>A |
| chr15 | 51700000 | 51800000 | 0.58 | 0.35 | A>A |
| chr15 | 51800000 | 51900000 | 0.65 | 0.44 | A>A |
| chr15 | 51900000 | 52000000 | 0.64 | 0.51 | A>A |
| chr15 | 52000000 | 52100000 | 0.72 | 0.61 | A>A |
| chr15 | 52100000 | 52200000 | 0.81 | 0.74 | A>A |
| chr15 | 52200000 | 52300000 | 0.73 | 0.73 | A>A |
| chr15 | 52300000 | 52400000 | 0.81 | 0.84 | A>A |
| chr15 | 52400000 | 52500000 | 0.69 | 0.77 | A>A |
| chr15 | 52500000 | 52600000 | 0.54 | 0.60 | A>A |
| chr15 | 52600000 | 52700000 | 0.45 | 0.44 | A>A |
| chr15 | 52700000 | 52800000 | 0.38 | 0.36 | A>A |
| chr15 | 52800000 | 52900000 | 0.47 | 0.47 | A>A |
| chr15 | 52900000 | 53000000 | 0.46 | 0.44 | A>A |
| chr15 | 53000000 | 53100000 | 0.13 | -0.04 | A>A |
| chr15 | 53100000 | 53200000 | -0.57 | -0.97 | B>B |
| chr15 | 53200000 | 53300000 | -0.79 | -1.17 | B>B |
| chr15 | 53300000 | 53400000 | -0.85 | -1.11 | B>B |
| chr15 | 53400000 | 53500000 | -0.63 | -0.89 | B>B |

| | | | | | |
|---|---|---|---|---|---|
| chr15 | 53500000 | 53600000 | -0.30 | -0.58 | B>B |
| chr15 | 53600000 | 53700000 | -0.15 | -0.30 | B>B |
| chr15 | 53700000 | 53800000 | 0.00 | -0.10 | B>B |
| chr15 | 53800000 | 53900000 | 0.09 | 0.01 | A>A |
| chr15 | 53900000 | 54000000 | 0.10 | -0.05 | A>A |
| chr15 | 54000000 | 54100000 | -0.49 | -0.64 | B>B |
| chr15 | 54100000 | 54200000 | 0.00 | -0.93 | NO |
| chr15 | 54200000 | 54300000 | 0.00 | -1.14 | NO |
| chr15 | 54300000 | 54400000 | 0.00 | -1.19 | NO |
| chr15 | 54400000 | 54500000 | 0.00 | -1.23 | NO |
| chr15 | 54500000 | 54600000 | 0.00 | -1.31 | NO |
| chr15 | 54600000 | 54700000 | 0.00 | -1.19 | NO |
| chr15 | 54700000 | 54800000 | 0.00 | 0.00 | NO |
| chr15 | 54800000 | 54900000 | 0.00 | -1.39 | NO |
| chr15 | 54900000 | 55000000 | 0.00 | -1.44 | NO |
| chr15 | 55000000 | 55100000 | 0.00 | 0.00 | NO |
| chr15 | 55100000 | 55200000 | 0.00 | 0.00 | NO |
| chr15 | 55200000 | 55300000 | 0.00 | 0.00 | NO |
| chr15 | 55300000 | 55400000 | -0.97 | -1.37 | B>B |
| chr15 | 55400000 | 55500000 | -0.38 | -0.79 | B>B |
| chr15 | 55500000 | 55600000 | -0.04 | -0.09 | B>B |
| chr15 | 55600000 | 55700000 | 0.20 | 0.23 | A>A |
| chr15 | 55700000 | 55800000 | 0.22 | 0.17 | A>A |
| chr15 | 55800000 | 55900000 | 0.25 | 0.27 | A>A |
| chr15 | 55900000 | 56000000 | 0.33 | 0.29 | A>A |
| chr15 | 56000000 | 56100000 | 0.42 | 0.34 | A>A |
| chr15 | 56100000 | 56200000 | 0.37 | 0.39 | A>A |
| chr15 | 56200000 | 56300000 | 0.37 | 0.39 | A>A |
| chr15 | 56300000 | 56400000 | 0.17 | 0.25 | A>A |
| chr15 | 56400000 | 56500000 | 0.01 | 0.21 | A>A |
| chr15 | 56500000 | 56600000 | -0.17 | 0.04 | B>B |
| chr15 | 56600000 | 56700000 | -0.23 | -0.06 | B>B |
| chr15 | 56700000 | 56800000 | -0.09 | 0.16 | A>A |
| chr15 | 56800000 | 56900000 | -0.24 | 0.21 | B>B |
| chr15 | 56900000 | 57000000 | 0.08 | 0.35 | A>A |
| chr15 | 57000000 | 57100000 | 0.19 | 0.50 | A>A |
| chr15 | 57100000 | 57200000 | 0.29 | 0.48 | A>A |
| chr15 | 57200000 | 57300000 | 0.40 | 0.55 | A>A |
| chr15 | 57300000 | 57400000 | 0.38 | 0.56 | A>A |
| chr15 | 57400000 | 57500000 | 0.34 | 0.49 | A>A |
| chr15 | 57500000 | 57600000 | 0.31 | 0.45 | A>A |
| chr15 | 57600000 | 57700000 | 0.35 | 0.33 | A>A |
| chr15 | 57700000 | 57800000 | 0.41 | 0.31 | A>A |
| chr15 | 57800000 | 57900000 | 0.27 | 0.20 | A>A |
| chr15 | 57900000 | 58000000 | 0.19 | -0.01 | A>A |
| chr15 | 58000000 | 58100000 | -0.02 | -0.43 | B>B |
| chr15 | 58100000 | 58200000 | -0.22 | -0.64 | B>B |
| chr15 | 58200000 | 58300000 | -0.32 | -0.75 | B>B |
| chr15 | 58300000 | 58400000 | -0.18 | -0.47 | B>B |
| chr15 | 58400000 | 58500000 | -0.01 | -0.35 | B>B |
| chr15 | 58500000 | 58600000 | 0.43 | 0.33 | A>A |
| chr15 | 58600000 | 58700000 | 0.59 | 0.53 | A>A |
| chr15 | 58700000 | 58800000 | 0.58 | 0.56 | A>A |
| chr15 | 58800000 | 58900000 | 0.56 | 0.54 | A>A |
| chr15 | 58900000 | 59000000 | 0.46 | 0.48 | A>A |
| chr15 | 59000000 | 59100000 | 0.52 | 0.48 | A>A |

| | | | | | |
|---|---|---|---|---|---|
| chr15 | 59100000 | 59200000 | 0.51 | 0.51 | A>A |
| chr15 | 59200000 | 59300000 | 0.50 | 0.55 | A>A |
| chr15 | 59300000 | 59400000 | 0.57 | 0.60 | A>A |
| chr15 | 59400000 | 59500000 | 0.47 | 0.58 | A>A |
| chr15 | 59500000 | 59600000 | 0.54 | 0.65 | A>A |
| chr15 | 59600000 | 59700000 | 0.57 | 0.69 | A>A |
| chr15 | 59700000 | 59800000 | 0.48 | 0.59 | A>A |
| chr15 | 59800000 | 59900000 | 0.58 | 0.58 | A>A |
| chr15 | 59900000 | 60000000 | 0.42 | 0.37 | A>A |
| chr15 | 60000000 | 60100000 | -0.24 | -0.36 | B>B |
| chr15 | 60100000 | 60200000 | -0.40 | -0.62 | B>B |
| chr15 | 60200000 | 60300000 | -0.45 | -0.57 | B>B |
| chr15 | 60300000 | 60400000 | -0.39 | -0.68 | B>B |
| chr15 | 60400000 | 60500000 | -0.41 | -0.56 | B>B |
| chr15 | 60500000 | 60600000 | -0.35 | -0.60 | B>B |
| chr15 | 60600000 | 60700000 | 0.18 | 0.18 | A>A |
| chr15 | 60700000 | 60800000 | 0.35 | 0.20 | A>A |
| chr15 | 60800000 | 60900000 | -0.18 | -0.37 | B>B |
| chr15 | 60900000 | 61000000 | -0.52 | -0.83 | B>B |
| chr15 | 61000000 | 61100000 | -0.64 | -0.98 | B>B |
| chr15 | 61100000 | 61200000 | -0.61 | -0.99 | B>B |
| chr15 | 61200000 | 61300000 | -0.56 | -0.86 | B>B |
| chr15 | 61300000 | 61400000 | -0.55 | -0.67 | B>B |
| chr15 | 61400000 | 61500000 | -0.48 | -0.59 | B>B |
| chr15 | 61500000 | 61600000 | -0.66 | -0.92 | B>B |
| chr15 | 61600000 | 61700000 | -0.77 | -1.36 | A>B |
| chr15 | 61700000 | 61800000 | -0.69 | -1.45 | A>B |
| chr15 | 61800000 | 61900000 | -0.79 | -1.44 | A>B |
| chr15 | 61900000 | 62000000 | -0.76 | -1.53 | A>B |
| chr15 | 62000000 | 62100000 | -0.64 | -1.19 | B>B |
| chr15 | 62100000 | 62200000 | -0.15 | -0.28 | B>B |
| chr15 | 62200000 | 62300000 | 0.10 | 0.16 | A>A |
| chr15 | 62300000 | 62400000 | 0.21 | 0.31 | A>A |
| chr15 | 62400000 | 62500000 | 0.11 | 0.14 | A>A |
| chr15 | 62500000 | 62600000 | 0.15 | 0.12 | A>A |
| chr15 | 62600000 | 62700000 | 0.32 | 0.37 | A>A |
| chr15 | 62700000 | 62800000 | 0.47 | 0.46 | A>A |
| chr15 | 62800000 | 62900000 | 0.47 | 0.49 | A>A |
| chr15 | 62900000 | 63000000 | 0.47 | 0.47 | A>A |
| chr15 | 63000000 | 63100000 | 0.52 | 0.52 | A>A |
| chr15 | 63100000 | 63200000 | 0.61 | 0.64 | A>A |
| chr15 | 63200000 | 63300000 | 0.64 | 0.63 | A>A |
| chr15 | 63300000 | 63400000 | 0.57 | 0.61 | A>A |
| chr15 | 63400000 | 63500000 | 0.62 | 0.63 | A>A |
| chr15 | 63500000 | 63600000 | 0.57 | 0.62 | A>A |
| chr15 | 63600000 | 63700000 | 0.57 | 0.63 | A>A |
| chr15 | 63700000 | 63800000 | 0.58 | 0.64 | A>A |
| chr15 | 63800000 | 63900000 | 0.61 | 0.65 | A>A |
| chr15 | 63900000 | 64000000 | 0.58 | 0.62 | A>A |
| chr15 | 64000000 | 64100000 | 0.55 | 0.58 | A>A |
| chr15 | 64100000 | 64200000 | 0.55 | 0.58 | A>A |
| chr15 | 64200000 | 64300000 | 0.58 | 0.60 | A>A |
| chr15 | 64300000 | 64400000 | 0.54 | 0.58 | A>A |
| chr15 | 64400000 | 64500000 | 0.54 | 0.57 | A>A |
| chr15 | 64500000 | 64600000 | 0.60 | 0.59 | A>A |
| chr15 | 64600000 | 64700000 | 0.56 | 0.57 | A>A |

| | | | | | |
|---|---|---|---|---|---|
| chr15 | 64700000 | 64800000 | 0.55 | 0.56 | A>A |
| chr15 | 64800000 | 64900000 | 0.55 | 0.60 | A>A |
| chr15 | 64900000 | 65000000 | 0.49 | 0.54 | A>A |
| chr15 | 65000000 | 65100000 | 0.52 | 0.51 | A>A |
| chr15 | 65100000 | 65200000 | 0.56 | 0.60 | A>A |
| chr15 | 65200000 | 65300000 | 0.55 | 0.55 | A>A |
| chr15 | 65300000 | 65400000 | 0.56 | 0.59 | A>A |
| chr15 | 65400000 | 65500000 | 0.54 | 0.60 | A>A |
| chr15 | 65500000 | 65600000 | 0.58 | 0.58 | A>A |
| chr15 | 65600000 | 65700000 | 0.56 | 0.56 | A>A |
| chr15 | 65700000 | 65800000 | 0.56 | 0.56 | A>A |
| chr15 | 65800000 | 65900000 | 0.50 | 0.51 | A>A |
| chr15 | 65900000 | 66000000 | 0.46 | 0.47 | A>A |
| chr15 | 66000000 | 66100000 | 0.52 | 0.55 | A>A |
| chr15 | 66100000 | 66200000 | 0.48 | 0.49 | A>A |
| chr15 | 66200000 | 66300000 | 0.42 | 0.47 | A>A |
| chr15 | 66300000 | 66400000 | 0.44 | 0.48 | A>A |
| chr15 | 66400000 | 66500000 | 0.43 | 0.49 | A>A |
| chr15 | 66500000 | 66600000 | 0.43 | 0.48 | A>A |
| chr15 | 66600000 | 66700000 | 0.45 | 0.50 | A>A |
| chr15 | 66700000 | 66800000 | 0.48 | 0.53 | A>A |
| chr15 | 66800000 | 66900000 | 0.44 | 0.50 | A>A |
| chr15 | 66900000 | 67000000 | 0.48 | 0.50 | A>A |
| chr15 | 67000000 | 67100000 | 0.44 | 0.54 | A>A |
| chr15 | 67100000 | 67200000 | 0.44 | 0.49 | A>A |
| chr15 | 67200000 | 67300000 | 0.37 | 0.46 | A>A |
| chr15 | 67300000 | 67400000 | 0.47 | 0.50 | A>A |
| chr15 | 67400000 | 67500000 | 0.41 | 0.47 | A>A |
| chr15 | 67500000 | 67600000 | 0.35 | 0.46 | A>A |
| chr15 | 67600000 | 67700000 | 0.36 | 0.46 | A>A |
| chr15 | 67700000 | 67800000 | 0.36 | 0.46 | A>A |
| chr15 | 67800000 | 67900000 | 0.41 | 0.46 | A>A |
| chr15 | 67900000 | 68000000 | 0.39 | 0.43 | A>A |
| chr15 | 68000000 | 68100000 | 0.40 | 0.45 | A>A |
| chr15 | 68100000 | 68200000 | 0.39 | 0.45 | A>A |
| chr15 | 68200000 | 68300000 | 0.49 | 0.52 | A>A |
| chr15 | 68300000 | 68400000 | 0.45 | 0.52 | A>A |
| chr15 | 68400000 | 68500000 | 0.45 | 0.49 | A>A |
| chr15 | 68500000 | 68600000 | 0.46 | 0.49 | A>A |
| chr15 | 68600000 | 68700000 | 0.46 | 0.48 | A>A |
| chr15 | 68700000 | 68800000 | 0.40 | 0.44 | A>A |
| chr15 | 68800000 | 68900000 | 0.42 | 0.43 | A>A |
| chr15 | 68900000 | 69000000 | 0.40 | 0.42 | A>A |
| chr15 | 69000000 | 69100000 | 0.44 | 0.41 | A>A |
| chr15 | 69100000 | 69200000 | 0.40 | 0.41 | A>A |
| chr15 | 69200000 | 69300000 | 0.40 | 0.38 | A>A |
| chr15 | 69300000 | 69400000 | 0.44 | 0.46 | A>A |
| chr15 | 69400000 | 69500000 | 0.48 | 0.52 | A>A |
| chr15 | 69500000 | 69600000 | 0.50 | 0.50 | A>A |
| chr15 | 69600000 | 69700000 | 0.49 | 0.54 | A>A |
| chr15 | 69700000 | 69800000 | 0.48 | 0.50 | A>A |
| chr15 | 69800000 | 69900000 | 0.42 | 0.46 | A>A |
| chr15 | 69900000 | 70000000 | 0.39 | 0.47 | A>A |
| chr15 | 70000000 | 70100000 | 0.36 | 0.44 | A>A |
| chr15 | 70100000 | 70200000 | 0.36 | 0.44 | A>A |
| chr15 | 70200000 | 70300000 | 0.33 | 0.42 | A>A |

| | | | | | |
|---|---|---|---|---|---|
| chr15 | 70300000 | 70400000 | 0.40 | 0.43 | A>A |
| chr15 | 70400000 | 70500000 | 0.42 | 0.48 | A>A |
| chr15 | 70500000 | 70600000 | 0.40 | 0.47 | A>A |
| chr15 | 70600000 | 70700000 | 0.40 | 0.48 | A>A |
| chr15 | 70700000 | 70800000 | 0.42 | 0.48 | A>A |
| chr15 | 70800000 | 70900000 | 0.42 | 0.49 | A>A |
| chr15 | 70900000 | 71000000 | 0.42 | 0.49 | A>A |
| chr15 | 71000000 | 71100000 | 0.54 | 0.59 | A>A |
| chr15 | 71100000 | 71200000 | 0.45 | 0.54 | A>A |
| chr15 | 71200000 | 71300000 | 0.42 | 0.46 | A>A |
| chr15 | 71300000 | 71400000 | 0.39 | 0.46 | A>A |
| chr15 | 71400000 | 71500000 | 0.38 | 0.46 | A>A |
| chr15 | 71500000 | 71600000 | 0.40 | 0.47 | A>A |
| chr15 | 71600000 | 71700000 | 0.34 | 0.45 | A>A |
| chr15 | 71700000 | 71800000 | 0.38 | 0.46 | A>A |
| chr15 | 71800000 | 71900000 | 0.37 | 0.50 | A>A |
| chr15 | 71900000 | 72000000 | 0.41 | 0.46 | A>A |
| chr15 | 72000000 | 72100000 | 0.43 | 0.47 | A>A |
| chr15 | 72100000 | 72200000 | 0.50 | 0.49 | A>A |
| chr15 | 72200000 | 72300000 | 0.53 | 0.56 | A>A |
| chr15 | 72300000 | 72400000 | 0.54 | 0.57 | A>A |
| chr15 | 72400000 | 72500000 | 0.51 | 0.58 | A>A |
| chr15 | 72500000 | 72600000 | 0.69 | 0.68 | A>A |
| chr15 | 72600000 | 72700000 | 0.61 | 0.64 | A>A |
| chr15 | 72700000 | 72800000 | 0.67 | 0.68 | A>A |
| chr15 | 72800000 | 72900000 | 0.67 | 0.64 | A>A |
| chr15 | 72900000 | 73000000 | 0.00 | 0.00 | NO |
| chr15 | 73000000 | 73100000 | 0.50 | 0.40 | A>A |
| chr15 | 73100000 | 73200000 | 0.25 | -0.14 | A>A |
| chr15 | 73200000 | 73300000 | 0.16 | -0.22 | B>B |
| chr15 | 73300000 | 73400000 | 0.37 | 0.08 | A>A |
| chr15 | 73400000 | 73500000 | 0.34 | 0.26 | A>A |
| chr15 | 73500000 | 73600000 | 0.31 | 0.23 | A>A |
| chr15 | 73600000 | 73700000 | 0.30 | 0.18 | A>A |
| chr15 | 73700000 | 73800000 | 0.11 | 0.16 | A>A |
| chr15 | 73800000 | 73900000 | 0.34 | 0.28 | A>A |
| chr15 | 73900000 | 74000000 | 0.42 | 0.29 | A>A |
| chr15 | 74000000 | 74100000 | 0.50 | 0.49 | A>A |
| chr15 | 74100000 | 74200000 | 0.00 | 0.51 | NO |
| chr15 | 74200000 | 74300000 | 0.00 | 0.54 | NO |
| chr15 | 74300000 | 74400000 | 0.00 | 0.00 | NO |
| chr15 | 74400000 | 74500000 | 0.00 | 0.00 | NO |
| chr15 | 74500000 | 74600000 | 0.00 | 0.63 | NO |
| chr15 | 74600000 | 74700000 | 0.00 | 0.58 | NO |
| chr15 | 74700000 | 74800000 | 0.00 | 0.00 | NO |
| chr15 | 74800000 | 74900000 | 0.00 | 0.56 | NO |
| chr15 | 74900000 | 75000000 | 0.00 | 0.00 | NO |
| chr15 | 75000000 | 75100000 | 0.23 | 0.14 | A>A |
| chr15 | 75100000 | 75200000 | 0.25 | 0.08 | A>A |
| chr15 | 75200000 | 75300000 | 0.53 | 0.46 | A>A |
| chr15 | 75300000 | 75400000 | 0.68 | 0.69 | A>A |
| chr15 | 75400000 | 75500000 | 0.73 | 0.71 | A>A |
| chr15 | 75500000 | 75600000 | 0.00 | 0.00 | NO |
| chr15 | 75600000 | 75700000 | 0.68 | 0.67 | A>A |
| chr15 | 75700000 | 75800000 | 0.75 | 0.71 | A>A |
| chr15 | 75800000 | 75900000 | 0.75 | 0.72 | A>A |

| | | | | | |
|---|---|---|---|---|---|
| chr15 | 75900000 | 76000000 | 0.70 | 0.69 | A>A |
| chr15 | 76000000 | 76100000 | 0.00 | 0.69 | NO |
| chr15 | 76100000 | 76200000 | 0.61 | 0.63 | A>A |
| chr15 | 76200000 | 76300000 | 0.37 | 0.49 | A>A |
| chr15 | 76300000 | 76400000 | 0.29 | 0.44 | A>A |
| chr15 | 76400000 | 76500000 | 0.26 | 0.45 | A>A |
| chr15 | 76500000 | 76600000 | 0.34 | 0.51 | A>A |
| chr15 | 76600000 | 76700000 | 0.20 | 0.30 | A>A |
| chr15 | 76700000 | 76800000 | -0.07 | 0.16 | A>A |
| chr15 | 76800000 | 76900000 | -0.17 | 0.15 | B>B |
| chr15 | 76900000 | 77000000 | -0.12 | 0.16 | A>A |
| chr15 | 77000000 | 77100000 | -0.01 | 0.20 | A>A |
| chr15 | 77100000 | 77200000 | 0.15 | 0.33 | A>A |
| chr15 | 77200000 | 77300000 | 0.38 | 0.46 | A>A |
| chr15 | 77300000 | 77400000 | 0.45 | 0.49 | A>A |
| chr15 | 77400000 | 77500000 | 0.26 | 0.37 | A>A |
| chr15 | 77500000 | 77600000 | 0.23 | 0.34 | A>A |
| chr15 | 77600000 | 77700000 | 0.32 | 0.37 | A>A |
| chr15 | 77700000 | 77800000 | 0.36 | 0.44 | A>A |
| chr15 | 77800000 | 77900000 | 0.46 | 0.44 | A>A |
| chr15 | 77900000 | 78000000 | 0.56 | 0.52 | A>A |
| chr15 | 78000000 | 78100000 | 0.60 | 0.51 | A>A |
| chr15 | 78100000 | 78200000 | 0.57 | 0.56 | A>A |
| chr15 | 78200000 | 78300000 | 0.00 | 0.00 | NO |
| chr15 | 78300000 | 78400000 | 0.74 | 0.70 | A>A |
| chr15 | 78400000 | 78500000 | 0.78 | 0.72 | A>A |
| chr15 | 78500000 | 78600000 | 0.72 | 0.69 | A>A |
| chr15 | 78600000 | 78700000 | 0.76 | 0.72 | A>A |
| chr15 | 78700000 | 78800000 | 0.80 | 0.75 | A>A |
| chr15 | 78800000 | 78900000 | 0.79 | 0.77 | A>A |
| chr15 | 78900000 | 79000000 | 0.74 | 0.67 | A>A |
| chr15 | 79000000 | 79100000 | 0.00 | 0.00 | NO |
| chr15 | 79100000 | 79200000 | 0.72 | 0.65 | A>A |
| chr15 | 79200000 | 79300000 | 0.47 | 0.37 | A>A |
| chr15 | 79300000 | 79400000 | 0.09 | -0.17 | B>B |
| chr15 | 79400000 | 79500000 | -0.10 | -0.31 | B>B |
| chr15 | 79500000 | 79600000 | -0.20 | -0.40 | B>B |
| chr15 | 79600000 | 79700000 | -0.07 | -0.07 | B>B |
| chr15 | 79700000 | 79800000 | -0.26 | -0.31 | B>B |
| chr15 | 79800000 | 79900000 | -0.23 | -0.35 | B>B |
| chr15 | 79900000 | 80000000 | -0.11 | -0.06 | B>B |
| chr15 | 80000000 | 80100000 | -0.16 | -0.12 | B>B |
| chr15 | 80100000 | 80200000 | 0.21 | 0.22 | A>A |
| chr15 | 80200000 | 80300000 | 0.49 | 0.38 | A>A |
| chr15 | 80300000 | 80400000 | 0.47 | 0.49 | A>A |
| chr15 | 80400000 | 80500000 | 0.29 | 0.20 | A>A |
| chr15 | 80500000 | 80600000 | -0.03 | -0.27 | B>B |
| chr15 | 80600000 | 80700000 | 0.02 | -0.19 | B>B |
| chr15 | 80700000 | 80800000 | 0.16 | 0.19 | A>A |
| chr15 | 80800000 | 80900000 | 0.17 | 0.29 | A>A |
| chr15 | 80900000 | 81000000 | 0.25 | 0.41 | A>A |
| chr15 | 81000000 | 81100000 | 0.27 | 0.37 | A>A |
| chr15 | 81100000 | 81200000 | 0.13 | 0.21 | A>A |
| chr15 | 81200000 | 81300000 | 0.03 | 0.05 | A>A |
| chr15 | 81300000 | 81400000 | -0.48 | -0.79 | B>B |
| chr15 | 81400000 | 81500000 | -0.93 | -1.25 | B>B |

| | | | | | |
|---|---|---|---|---|---|
| chr15 | 81500000 | 81600000 | -0.96 | -1.23 | B>B |
| chr15 | 81600000 | 81700000 | -0.93 | -1.30 | B>B |
| chr15 | 81700000 | 81800000 | -1.22 | -1.49 | B>B |
| chr15 | 81800000 | 81900000 | -1.52 | -1.85 | B>B |
| chr15 | 81900000 | 82000000 | -1.57 | -1.88 | B>B |
| chr15 | 82000000 | 82100000 | -1.36 | -1.85 | B>B |
| chr15 | 82100000 | 82200000 | -1.28 | -1.78 | B>B |
| chr15 | 82200000 | 82300000 | -1.05 | -1.58 | B>B |
| chr15 | 82300000 | 82400000 | -0.62 | -0.94 | B>B |
| chr15 | 82400000 | 82500000 | -0.32 | -0.46 | B>B |
| chr15 | 82500000 | 82600000 | 0.00 | 0.00 | NO |
| chr15 | 82600000 | 82700000 | 0.00 | 0.00 | NO |
| chr15 | 82700000 | 82800000 | 0.00 | 0.00 | NO |
| chr15 | 82800000 | 82900000 | 0.00 | 0.00 | NO |
| chr15 | 82900000 | 83000000 | 0.00 | 0.00 | NO |
| chr15 | 83000000 | 83100000 | 0.00 | 0.00 | NO |
| chr15 | 83100000 | 83200000 | 0.00 | 0.00 | NO |
| chr15 | 83200000 | 83300000 | 0.20 | 0.09 | A>A |
| chr15 | 83300000 | 83400000 | 0.21 | 0.12 | A>A |
| chr15 | 83400000 | 83500000 | 0.32 | 0.32 | A>A |
| chr15 | 83500000 | 83600000 | 0.17 | 0.41 | A>A |
| chr15 | 83600000 | 83700000 | 0.25 | 0.38 | A>A |
| chr15 | 83700000 | 83800000 | 0.10 | 0.12 | A>A |
| chr15 | 83800000 | 83900000 | -0.09 | -0.05 | B>B |
| chr15 | 83900000 | 84000000 | -0.39 | -0.34 | B>B |
| chr15 | 84000000 | 84100000 | -0.56 | -0.53 | B>B |
| chr15 | 84100000 | 84200000 | -0.88 | -1.08 | B>B |
| chr15 | 84200000 | 84300000 | -0.88 | -1.16 | B>B |
| chr15 | 84300000 | 84400000 | -1.15 | -1.33 | B>B |
| chr15 | 84400000 | 84500000 | -1.24 | -1.43 | B>B |
| chr15 | 84500000 | 84600000 | -1.26 | -1.52 | B>B |
| chr15 | 84600000 | 84700000 | -1.36 | -1.52 | B>B |
| chr15 | 84700000 | 84800000 | -1.03 | -1.22 | B>B |
| chr15 | 84800000 | 84900000 | 0.00 | 0.00 | NO |
| chr15 | 84900000 | 85000000 | 0.00 | 0.00 | NO |
| chr15 | 85000000 | 85100000 | 0.00 | 0.00 | NO |
| chr15 | 85100000 | 85200000 | 0.65 | 0.77 | A>A |
| chr15 | 85200000 | 85300000 | 0.74 | 0.84 | A>A |
| chr15 | 85300000 | 85400000 | 0.72 | 0.80 | A>A |
| chr15 | 85400000 | 85500000 | 0.68 | 0.81 | A>A |
| chr15 | 85500000 | 85600000 | 0.73 | 0.84 | A>A |
| chr15 | 85600000 | 85700000 | 0.63 | 0.77 | A>A |
| chr15 | 85700000 | 85800000 | 0.00 | 0.00 | NO |
| chr15 | 85800000 | 85900000 | 0.46 | 0.61 | A>A |
| chr15 | 85900000 | 86000000 | 0.55 | 0.67 | A>A |
| chr15 | 86000000 | 86100000 | 0.39 | 0.60 | A>A |
| chr15 | 86100000 | 86200000 | 0.36 | 0.59 | A>A |
| chr15 | 86200000 | 86300000 | 0.12 | 0.43 | A>A |
| chr15 | 86300000 | 86400000 | -0.10 | 0.04 | B>B |
| chr15 | 86400000 | 86500000 | -0.84 | -1.02 | B>B |
| chr15 | 86500000 | 86600000 | -1.06 | -1.33 | B>B |
| chr15 | 86600000 | 86700000 | -1.14 | -1.31 | B>B |
| chr15 | 86700000 | 86800000 | -1.09 | -1.08 | B>B |
| chr15 | 86800000 | 86900000 | -1.21 | -1.01 | B>B |
| chr15 | 86900000 | 87000000 | -1.18 | -0.75 | B>B |
| chr15 | 87000000 | 87100000 | -1.01 | -0.46 | B>A |

| | | | | | |
|---|---|---|---|---|---|
| chr15 | 87100000 | 87200000 | -0.77 | -0.38 | B>B |
| chr15 | 87200000 | 87300000 | -1.02 | -0.70 | B>B |
| chr15 | 87300000 | 87400000 | -1.08 | -1.02 | B>B |
| chr15 | 87400000 | 87500000 | -1.07 | -1.08 | B>B |
| chr15 | 87500000 | 87600000 | -1.19 | -1.13 | B>B |
| chr15 | 87600000 | 87700000 | -1.23 | -1.14 | B>B |
| chr15 | 87700000 | 87800000 | -1.21 | -1.12 | B>B |
| chr15 | 87800000 | 87900000 | -1.14 | -1.10 | B>B |
| chr15 | 87900000 | 88000000 | -1.06 | -1.13 | B>B |
| chr15 | 88000000 | 88100000 | -1.04 | -1.16 | B>B |
| chr15 | 88100000 | 88200000 | -0.89 | -1.19 | B>B |
| chr15 | 88200000 | 88300000 | -0.78 | -1.17 | B>B |
| chr15 | 88300000 | 88400000 | -0.79 | -1.23 | B>B |
| chr15 | 88400000 | 88500000 | -0.98 | -1.19 | B>B |
| chr15 | 88500000 | 88600000 | -0.98 | -1.22 | B>B |
| chr15 | 88600000 | 88700000 | -0.99 | -1.25 | B>B |
| chr15 | 88700000 | 88800000 | -0.80 | -1.06 | B>B |
| chr15 | 88800000 | 88900000 | -0.60 | -1.16 | B>B |
| chr15 | 88900000 | 89000000 | -0.58 | -1.08 | B>B |
| chr15 | 89000000 | 89100000 | -0.04 | -0.29 | B>B |
| chr15 | 89100000 | 89200000 | 0.15 | -0.19 | B>B |
| chr15 | 89200000 | 89300000 | 0.10 | -0.55 | A>B |
| chr15 | 89300000 | 89400000 | 0.14 | -0.50 | A>B |
| chr15 | 89400000 | 89500000 | 0.50 | 0.26 | A>A |
| chr15 | 89500000 | 89600000 | 0.54 | 0.44 | A>A |
| chr15 | 89600000 | 89700000 | 0.68 | 0.69 | A>A |
| chr15 | 89700000 | 89800000 | 0.63 | 0.66 | A>A |
| chr15 | 89800000 | 89900000 | 0.68 | 0.72 | A>A |
| chr15 | 89900000 | 90000000 | 0.74 | 0.81 | A>A |
| chr15 | 90000000 | 90100000 | 0.71 | 0.79 | A>A |
| chr15 | 90100000 | 90200000 | 0.72 | 0.81 | A>A |
| chr15 | 90200000 | 90300000 | 0.68 | 0.77 | A>A |
| chr15 | 90300000 | 90400000 | 0.78 | 0.83 | A>A |
| chr15 | 90400000 | 90500000 | 0.69 | 0.77 | A>A |
| chr15 | 90500000 | 90600000 | 0.76 | 0.81 | A>A |
| chr15 | 90600000 | 90700000 | 0.65 | 0.74 | A>A |
| chr15 | 90700000 | 90800000 | 0.77 | 0.78 | A>A |
| chr15 | 90800000 | 90900000 | 0.61 | 0.69 | A>A |
| chr15 | 90900000 | 91000000 | 0.75 | 0.78 | A>A |
| chr15 | 91000000 | 91100000 | 0.58 | 0.72 | A>A |
| chr15 | 91100000 | 91200000 | 0.64 | 0.70 | A>A |
| chr15 | 91200000 | 91300000 | 0.59 | 0.66 | A>A |
| chr15 | 91300000 | 91400000 | 0.63 | 0.68 | A>A |
| chr15 | 91400000 | 91500000 | 0.59 | 0.58 | A>A |
| chr15 | 91500000 | 91600000 | 0.25 | 0.13 | A>A |
| chr15 | 91600000 | 91700000 | -0.62 | -0.79 | B>B |
| chr15 | 91700000 | 91800000 | -0.66 | -0.91 | B>B |
| chr15 | 91800000 | 91900000 | -0.54 | -0.75 | B>B |
| chr15 | 91900000 | 92000000 | -0.84 | -1.02 | B>B |
| chr15 | 92000000 | 92100000 | -0.66 | -1.03 | B>B |
| chr15 | 92100000 | 92200000 | -0.68 | -1.10 | B>B |
| chr15 | 92200000 | 92300000 | -0.72 | -1.04 | B>B |
| chr15 | 92300000 | 92400000 | -0.58 | -0.81 | B>B |
| chr15 | 92400000 | 92500000 | -0.13 | -0.09 | B>B |
| chr15 | 92500000 | 92600000 | -0.20 | -0.17 | B>B |
| chr15 | 92600000 | 92700000 | -0.47 | -0.40 | B>B |

| | | | | | |
|---|---|---|---|---|---|
| chr15 | 92700000 | 92800000 | -0.96 | -0.90 | B>B |
| chr15 | 92800000 | 92900000 | -0.88 | -1.07 | B>B |
| chr15 | 92900000 | 93000000 | -0.54 | -1.01 | B>B |
| chr15 | 93000000 | 93100000 | -0.56 | -0.87 | B>B |
| chr15 | 93100000 | 93200000 | -0.35 | -0.50 | B>B |
| chr15 | 93200000 | 93300000 | -0.11 | -0.25 | B>B |
| chr15 | 93300000 | 93400000 | 0.38 | 0.36 | A>A |
| chr15 | 93400000 | 93500000 | 0.50 | 0.54 | A>A |
| chr15 | 93500000 | 93600000 | -0.13 | -0.11 | B>B |
| chr15 | 93600000 | 93700000 | -0.60 | -0.75 | B>B |
| chr15 | 93700000 | 93800000 | -0.70 | -0.97 | B>B |
| chr15 | 93800000 | 93900000 | -0.77 | -1.07 | B>B |
| chr15 | 93900000 | 94000000 | -0.95 | -1.19 | B>B |
| chr15 | 94000000 | 94100000 | -1.09 | -1.25 | B>B |
| chr15 | 94100000 | 94200000 | -1.23 | -1.25 | B>B |
| chr15 | 94200000 | 94300000 | -1.23 | -1.25 | B>B |
| chr15 | 94300000 | 94400000 | -1.24 | -1.28 | B>B |
| chr15 | 94400000 | 94500000 | -1.19 | -1.29 | B>B |
| chr15 | 94500000 | 94600000 | -1.15 | -1.33 | B>B |
| chr15 | 94600000 | 94700000 | -1.18 | -1.21 | B>B |
| chr15 | 94700000 | 94800000 | -1.24 | -1.26 | B>B |
| chr15 | 94800000 | 94900000 | -1.26 | -1.26 | B>B |
| chr15 | 94900000 | 95000000 | -1.30 | -1.34 | B>B |
| chr15 | 95000000 | 95100000 | -1.34 | -1.35 | B>B |
| chr15 | 95100000 | 95200000 | -1.29 | -1.21 | B>B |
| chr15 | 95200000 | 95300000 | -1.20 | -1.09 | B>B |
| chr15 | 95300000 | 95400000 | -1.22 | -1.09 | B>B |
| chr15 | 95400000 | 95500000 | -1.20 | -1.07 | B>B |
| chr15 | 95500000 | 95600000 | -1.11 | -0.84 | B>B |
| chr15 | 95600000 | 95700000 | -1.12 | -0.83 | B>B |
| chr15 | 95700000 | 95800000 | -0.89 | -0.50 | B>B |
| chr15 | 95800000 | 95900000 | -0.75 | -0.24 | B>A |
| chr15 | 95900000 | 96000000 | -0.90 | -0.19 | B>A |
| chr15 | 96000000 | 96100000 | -0.83 | -0.11 | B>A |
| chr15 | 96100000 | 96200000 | -0.76 | -0.09 | B>A |
| chr15 | 96200000 | 96300000 | -0.66 | -0.09 | B>A |
| chr15 | 96300000 | 96400000 | -0.50 | -0.04 | B>B |
| chr15 | 96400000 | 96500000 | -0.40 | -0.10 | B>B |
| chr15 | 96500000 | 96600000 | -0.24 | -0.07 | B>B |
| chr15 | 96600000 | 96700000 | -0.17 | 0.05 | B>B |
| chr15 | 96700000 | 96800000 | -0.14 | 0.02 | B>B |
| chr15 | 96800000 | 96900000 | -0.24 | 0.01 | B>B |
| chr15 | 96900000 | 97000000 | -0.69 | -0.55 | B>B |
| chr15 | 97000000 | 97100000 | -0.94 | -0.90 | B>B |
| chr15 | 97100000 | 97200000 | -1.03 | -1.02 | B>B |
| chr15 | 97200000 | 97300000 | -1.08 | -1.04 | B>B |
| chr15 | 97300000 | 97400000 | -1.13 | -1.08 | B>B |
| chr15 | 97400000 | 97500000 | -1.12 | -1.00 | B>B |
| chr15 | 97500000 | 97600000 | -1.17 | -1.17 | B>B |
| chr15 | 97600000 | 97700000 | -1.16 | -1.16 | B>B |
| chr15 | 97700000 | 97800000 | -1.13 | -0.94 | B>B |
| chr15 | 97800000 | 97900000 | -1.18 | -1.20 | B>B |
| chr15 | 97900000 | 98000000 | -1.28 | -1.36 | B>B |
| chr15 | 98000000 | 98100000 | -1.40 | -1.44 | B>B |
| chr15 | 98100000 | 98200000 | -1.41 | -1.37 | B>B |
| chr15 | 98200000 | 98300000 | -1.35 | -1.14 | B>B |

| | | | | | |
|---|---|---|---|---|---|
| chr15 | 98300000 | 98400000 | -1.40 | -1.11 | B>B |
| chr15 | 98400000 | 98500000 | -1.29 | -1.06 | B>B |
| chr15 | 98500000 | 98600000 | -1.41 | -1.29 | B>B |
| chr15 | 98600000 | 98700000 | -1.36 | -1.47 | B>B |
| chr15 | 98700000 | 98800000 | -1.34 | -1.55 | B>B |
| chr15 | 98800000 | 98900000 | -1.10 | -1.06 | B>B |
| chr15 | 98900000 | 99000000 | -0.74 | -0.74 | B>B |
| chr15 | 99000000 | 99100000 | -0.07 | 0.08 | A>A |
| chr15 | 99100000 | 99200000 | 0.23 | 0.34 | A>A |
| chr15 | 99200000 | 99300000 | 0.39 | 0.56 | A>A |
| chr15 | 99300000 | 99400000 | 0.31 | 0.59 | A>A |
| chr15 | 99400000 | 99500000 | 0.40 | 0.54 | A>A |
| chr15 | 99500000 | 99600000 | 0.29 | 0.37 | A>A |
| chr15 | 99600000 | 99700000 | 0.12 | 0.21 | A>A |
| chr15 | 99700000 | 99800000 | 0.10 | 0.17 | A>A |
| chr15 | 99800000 | 99900000 | 0.12 | 0.10 | A>A |
| chr15 | 99900000 | 100000000 | 0.11 | -0.02 | A>A |
| chr15 | 1E+08 | 100100000 | 0.16 | -0.10 | A>A |
| chr15 | 1E+08 | 100200000 | 0.05 | 0.03 | A>A |
| chr15 | 1E+08 | 100300000 | -0.08 | -0.07 | B>B |
| chr15 | 1E+08 | 100400000 | 0.00 | -0.69 | NO |
| chr15 | 1E+08 | 100500000 | -0.87 | -0.78 | B>B |
| chr15 | 1.01E+08 | 100600000 | -0.80 | -0.79 | B>B |
| chr15 | 1.01E+08 | 100700000 | -0.92 | -0.95 | B>B |
| chr15 | 1.01E+08 | 100800000 | -0.91 | -1.06 | B>B |
| chr15 | 1.01E+08 | 100900000 | -0.77 | -0.87 | B>B |
| chr15 | 1.01E+08 | 101000000 | -1.01 | -1.03 | B>B |
| chr15 | 1.01E+08 | 101100000 | -1.07 | -1.04 | B>B |
| chr15 | 1.01E+08 | 101200000 | -0.14 | -0.06 | B>B |
| chr15 | 1.01E+08 | 101300000 | 0.41 | 0.52 | A>A |
| chr15 | 1.01E+08 | 101400000 | 0.42 | 0.53 | A>A |
| chr15 | 1.01E+08 | 101500000 | 0.29 | 0.50 | A>A |
| chr15 | 1.02E+08 | 101600000 | 0.33 | 0.50 | A>A |
| chr15 | 1.02E+08 | 101700000 | 0.33 | 0.51 | A>A |
| chr15 | 1.02E+08 | 101800000 | 0.33 | 0.53 | A>A |
| chr15 | 1.02E+08 | 101900000 | 0.11 | 0.44 | A>A |
| chr15 | 1.02E+08 | 102000000 | 0.11 | 0.44 | A>A |
| chr15 | 1.02E+08 | 102100000 | 0.25 | 0.52 | A>A |
| chr15 | 1.02E+08 | 102200000 | 0.12 | 0.41 | A>A |
| chr15 | 1.02E+08 | 102300000 | -0.30 | -0.13 | B>B |
| chr15 | 1.02E+08 | 102400000 | -1.41 | -1.39 | B>B |
| chr15 | 1.02E+08 | 102500000 | 0.00 | 0.00 | NO |
| chr15 | 1.03E+08 | 102531392 | 0.00 | 0.00 | NO |
| chr16 | 0 | 100000 | 0.00 | 0.00 | NO |
| chr16 | 100000 | 200000 | 0.57 | 0.66 | A>A |
| chr16 | 200000 | 300000 | 0.70 | 0.72 | A>A |
| chr16 | 300000 | 400000 | 0.72 | 0.77 | A>A |
| chr16 | 400000 | 500000 | 0.59 | 0.65 | A>A |
| chr16 | 500000 | 600000 | 0.67 | 0.80 | A>A |
| chr16 | 600000 | 700000 | 0.67 | 0.75 | A>A |
| chr16 | 700000 | 800000 | 0.55 | 0.67 | A>A |
| chr16 | 800000 | 900000 | 0.46 | 0.55 | A>A |
| chr16 | 900000 | 1000000 | 0.50 | 0.61 | A>A |
| chr16 | 1000000 | 1100000 | 0.55 | 0.64 | A>A |
| chr16 | 1100000 | 1200000 | 0.45 | 0.53 | A>A |
| chr16 | 1200000 | 1300000 | 0.55 | 0.59 | A>A |

| | | | | | |
|---|---|---|---|---|---|
| chr16 | 1300000 | 1400000 | 0.58 | 0.62 | A>A |
| chr16 | 1400000 | 1500000 | 0.64 | 0.66 | A>A |
| chr16 | 1500000 | 1600000 | 0.55 | 0.57 | A>A |
| chr16 | 1600000 | 1700000 | 0.54 | 0.61 | A>A |
| chr16 | 1700000 | 1800000 | 0.59 | 0.65 | A>A |
| chr16 | 1800000 | 1900000 | 0.47 | 0.60 | A>A |
| chr16 | 1900000 | 2000000 | 0.68 | 0.70 | A>A |
| chr16 | 2000000 | 2100000 | 0.65 | 0.60 | A>A |
| chr16 | 2100000 | 2200000 | 0.00 | 0.00 | NO |
| chr16 | 2200000 | 2300000 | 0.65 | 0.73 | A>A |
| chr16 | 2300000 | 2400000 | 0.75 | 0.84 | A>A |
| chr16 | 2400000 | 2500000 | 0.89 | 0.85 | A>A |
| chr16 | 2500000 | 2600000 | 0.75 | 0.79 | A>A |
| chr16 | 2600000 | 2700000 | 0.00 | 0.00 | NO |
| chr16 | 2700000 | 2800000 | 0.82 | 0.89 | A>A |
| chr16 | 2800000 | 2900000 | 0.97 | 0.96 | A>A |
| chr16 | 2900000 | 3000000 | 0.94 | 0.99 | A>A |
| chr16 | 3000000 | 3100000 | 1.22 | 0.99 | A>A |
| chr16 | 3100000 | 3200000 | 1.07 | 1.13 | A>A |
| chr16 | 3200000 | 3300000 | 0.87 | 1.04 | A>A |
| chr16 | 3300000 | 3400000 | 0.82 | 1.10 | A>A |
| chr16 | 3400000 | 3500000 | 0.74 | 1.09 | A>A |
| chr16 | 3500000 | 3600000 | 0.90 | 0.97 | A>A |
| chr16 | 3600000 | 3700000 | 0.88 | 1.06 | A>A |
| chr16 | 3700000 | 3800000 | 0.98 | 1.11 | A>A |
| chr16 | 3800000 | 3900000 | 1.10 | 1.15 | A>A |
| chr16 | 3900000 | 4000000 | 0.93 | 1.04 | A>A |
| chr16 | 4000000 | 4100000 | 0.94 | 1.11 | A>A |
| chr16 | 4100000 | 4200000 | 1.16 | 1.22 | A>A |
| chr16 | 4200000 | 4300000 | 1.05 | 1.12 | A>A |
| chr16 | 4300000 | 4400000 | 1.23 | 1.32 | A>A |
| chr16 | 4400000 | 4500000 | 1.37 | 1.38 | A>A |
| chr16 | 4500000 | 4600000 | 1.38 | 1.47 | A>A |
| chr16 | 4600000 | 4700000 | 1.45 | 1.43 | A>A |
| chr16 | 4700000 | 4800000 | 1.36 | 1.26 | A>A |
| chr16 | 4800000 | 4900000 | 1.40 | 1.46 | A>A |
| chr16 | 4900000 | 5000000 | 1.41 | 1.32 | A>A |
| chr16 | 5000000 | 5100000 | 1.00 | 0.83 | A>A |
| chr16 | 5100000 | 5200000 | 0.00 | 0.00 | NO |
| chr16 | 5200000 | 5300000 | -0.19 | -0.90 | A>B |
| chr16 | 5300000 | 5400000 | -0.83 | -1.25 | B>B |
| chr16 | 5400000 | 5500000 | -0.75 | -1.26 | B>B |
| chr16 | 5500000 | 5600000 | -0.67 | -1.23 | B>B |
| chr16 | 5600000 | 5700000 | -0.75 | -1.20 | B>B |
| chr16 | 5700000 | 5800000 | -1.01 | -1.12 | B>B |
| chr16 | 5800000 | 5900000 | -0.96 | -1.21 | B>B |
| chr16 | 5900000 | 6000000 | -0.92 | -1.13 | B>B |
| chr16 | 6000000 | 6100000 | -0.88 | -1.17 | B>B |
| chr16 | 6100000 | 6200000 | -0.92 | -1.10 | B>B |
| chr16 | 6200000 | 6300000 | -0.95 | -1.06 | B>B |
| chr16 | 6300000 | 6400000 | -0.92 | -1.05 | B>B |
| chr16 | 6400000 | 6500000 | -0.93 | -1.12 | B>B |
| chr16 | 6500000 | 6600000 | -0.96 | -1.09 | B>B |
| chr16 | 6600000 | 6700000 | -0.78 | -1.00 | B>B |
| chr16 | 6700000 | 6800000 | -0.81 | -0.91 | B>B |
| chr16 | 6800000 | 6900000 | -0.86 | -0.97 | B>B |

| | | | | | |
|---|---|---|---|---|---|
| chr16 | 6900000 | 7000000 | -0.84 | -1.04 | B>B |
| chr16 | 7000000 | 7100000 | -0.74 | -0.97 | B>B |
| chr16 | 7100000 | 7200000 | -0.81 | -1.00 | B>B |
| chr16 | 7200000 | 7300000 | -0.77 | -0.97 | B>B |
| chr16 | 7300000 | 7400000 | -0.77 | -0.95 | B>B |
| chr16 | 7400000 | 7500000 | -0.76 | -0.96 | B>B |
| chr16 | 7500000 | 7600000 | -0.71 | -1.01 | B>B |
| chr16 | 7600000 | 7700000 | -0.89 | -1.07 | B>B |
| chr16 | 7700000 | 7800000 | -0.90 | -1.03 | B>B |
| chr16 | 7800000 | 7900000 | -0.85 | -1.00 | B>B |
| chr16 | 7900000 | 8000000 | -0.91 | -1.06 | B>B |
| chr16 | 8000000 | 8100000 | -0.92 | -1.08 | B>B |
| chr16 | 8100000 | 8200000 | -0.80 | -1.07 | B>B |
| chr16 | 8200000 | 8300000 | -0.89 | -1.07 | B>B |
| chr16 | 8300000 | 8400000 | -0.91 | -1.13 | B>B |
| chr16 | 8400000 | 8500000 | -0.88 | -1.15 | B>B |
| chr16 | 8500000 | 8600000 | -0.71 | -1.01 | B>B |
| chr16 | 8600000 | 8700000 | 0.00 | 0.00 | NO |
| chr16 | 8700000 | 8800000 | 0.05 | -0.05 | B>B |
| chr16 | 8800000 | 8900000 | 0.21 | 0.00 | A>A |
| chr16 | 8900000 | 9000000 | 0.37 | 0.41 | A>A |
| chr16 | 9000000 | 9100000 | 0.69 | 0.63 | A>A |
| chr16 | 9100000 | 9200000 | 0.73 | 0.56 | A>A |
| chr16 | 9200000 | 9300000 | 0.04 | -0.17 | B>B |
| chr16 | 9300000 | 9400000 | -0.37 | -0.58 | B>B |
| chr16 | 9400000 | 9500000 | -0.55 | -0.75 | B>B |
| chr16 | 9500000 | 9600000 | -0.63 | -0.84 | B>B |
| chr16 | 9600000 | 9700000 | -0.65 | -0.92 | B>B |
| chr16 | 9700000 | 9800000 | -0.65 | -0.97 | B>B |
| chr16 | 9800000 | 9900000 | -0.63 | -1.05 | B>B |
| chr16 | 9900000 | 10000000 | -0.61 | -1.04 | B>B |
| chr16 | 10000000 | 10100000 | -0.67 | -1.06 | B>B |
| chr16 | 10100000 | 10200000 | -0.62 | -1.03 | B>B |
| chr16 | 10200000 | 10300000 | -0.45 | -0.73 | B>B |
| chr16 | 10300000 | 10400000 | -0.10 | -0.20 | B>B |
| chr16 | 10400000 | 10500000 | 0.08 | 0.26 | A>A |
| chr16 | 10500000 | 10600000 | 0.07 | 0.19 | A>A |
| chr16 | 10600000 | 10700000 | 0.43 | 0.35 | A>A |
| chr16 | 10700000 | 10800000 | 0.49 | 0.53 | A>A |
| chr16 | 10800000 | 10900000 | 0.29 | 0.24 | A>A |
| chr16 | 10900000 | 11000000 | 0.28 | 0.22 | A>A |
| chr16 | 11000000 | 11100000 | 0.52 | 0.51 | A>A |
| chr16 | 11100000 | 11200000 | 0.59 | 0.67 | A>A |
| chr16 | 11200000 | 11300000 | 0.56 | 0.65 | A>A |
| chr16 | 11300000 | 11400000 | 0.68 | 0.74 | A>A |
| chr16 | 11400000 | 11500000 | 0.76 | 0.73 | A>A |
| chr16 | 11500000 | 11600000 | 0.99 | 0.96 | A>A |
| chr16 | 11600000 | 11700000 | 1.11 | 1.07 | A>A |
| chr16 | 11700000 | 11800000 | 1.07 | 1.10 | A>A |
| chr16 | 11800000 | 11900000 | 1.04 | 1.08 | A>A |
| chr16 | 11900000 | 12000000 | 0.85 | 0.92 | A>A |
| chr16 | 12000000 | 12100000 | 0.85 | 0.72 | A>A |
| chr16 | 12100000 | 12200000 | 0.53 | 0.50 | A>A |
| chr16 | 12200000 | 12300000 | 0.29 | 0.22 | A>A |
| chr16 | 12300000 | 12400000 | 0.21 | 0.12 | A>A |
| chr16 | 12400000 | 12500000 | 0.15 | -0.03 | A>A |

| | | | | | |
|---|---|---|---|---|---|
| chr16 | 12500000 | 12600000 | 0.11 | -0.12 | B>B |
| chr16 | 12600000 | 12700000 | 0.16 | -0.11 | A>A |
| chr16 | 12700000 | 12800000 | 0.36 | 0.14 | A>A |
| chr16 | 12800000 | 12900000 | 0.20 | 0.09 | A>A |
| chr16 | 12900000 | 13000000 | 0.11 | -0.24 | B>B |
| chr16 | 13000000 | 13100000 | -0.05 | -0.74 | A>B |
| chr16 | 13100000 | 13200000 | -0.23 | -0.95 | A>B |
| chr16 | 13200000 | 13300000 | -0.27 | -1.05 | A>B |
| chr16 | 13300000 | 13400000 | -0.51 | -1.14 | A>B |
| chr16 | 13400000 | 13500000 | -0.62 | -1.23 | A>B |
| chr16 | 13500000 | 13600000 | -0.58 | -1.12 | B>B |
| chr16 | 13600000 | 13700000 | -0.61 | -1.11 | B>B |
| chr16 | 13700000 | 13800000 | -0.73 | -1.13 | B>B |
| chr16 | 13800000 | 13900000 | -0.52 | -0.98 | B>B |
| chr16 | 13900000 | 14000000 | -0.21 | -0.60 | B>B |
| chr16 | 14000000 | 14100000 | 0.43 | 0.27 | A>A |
| chr16 | 14100000 | 14200000 | 0.59 | 0.38 | A>A |
| chr16 | 14200000 | 14300000 | 0.45 | 0.30 | A>A |
| chr16 | 14300000 | 14400000 | 0.34 | 0.27 | A>A |
| chr16 | 14400000 | 14500000 | 0.72 | 0.68 | A>A |
| chr16 | 14500000 | 14600000 | 0.82 | 0.71 | A>A |
| chr16 | 14600000 | 14700000 | 0.73 | 0.65 | A>A |
| chr16 | 14700000 | 14800000 | 0.85 | 0.92 | A>A |
| chr16 | 14800000 | 14900000 | 0.00 | 0.00 | NO |
| chr16 | 14900000 | 15000000 | 0.00 | 0.00 | NO |
| chr16 | 15000000 | 15100000 | 0.00 | 0.00 | NO |
| chr16 | 15100000 | 15200000 | 0.00 | 0.00 | NO |
| chr16 | 15200000 | 15300000 | 0.00 | 0.00 | NO |
| chr16 | 15300000 | 15400000 | 0.00 | 0.00 | NO |
| chr16 | 15400000 | 15500000 | 0.00 | 0.00 | NO |
| chr16 | 15500000 | 15600000 | 1.21 | 1.23 | A>A |
| chr16 | 15600000 | 15700000 | 1.38 | 1.39 | A>A |
| chr16 | 15700000 | 15800000 | 1.17 | 1.25 | A>A |
| chr16 | 15800000 | 15900000 | 0.87 | 0.96 | A>A |
| chr16 | 15900000 | 16000000 | 0.97 | 0.95 | A>A |
| chr16 | 16000000 | 16100000 | 1.05 | 1.18 | A>A |
| chr16 | 16100000 | 16200000 | 1.00 | 1.03 | A>A |
| chr16 | 16200000 | 16300000 | 0.79 | 0.85 | A>A |
| chr16 | 16300000 | 16400000 | 0.00 | 0.00 | NO |
| chr16 | 16400000 | 16500000 | 0.00 | 0.00 | NO |
| chr16 | 16500000 | 16600000 | 0.00 | 0.00 | NO |
| chr16 | 16600000 | 16700000 | 0.00 | 0.00 | NO |
| chr16 | 16700000 | 16800000 | 0.00 | 0.00 | NO |
| chr16 | 16800000 | 16900000 | 0.00 | 0.00 | NO |
| chr16 | 16900000 | 17000000 | -0.18 | -1.15 | A>B |
| chr16 | 17000000 | 17100000 | -0.20 | -1.32 | A>B |
| chr16 | 17100000 | 17200000 | -0.23 | -1.23 | A>B |
| chr16 | 17200000 | 17300000 | -0.38 | -1.24 | A>B |
| chr16 | 17300000 | 17400000 | -0.26 | -1.17 | A>B |
| chr16 | 17400000 | 17500000 | -0.20 | -1.13 | A>B |
| chr16 | 17500000 | 17600000 | -0.23 | -1.00 | A>B |
| chr16 | 17600000 | 17700000 | -0.51 | -1.11 | A>B |
| chr16 | 17700000 | 17800000 | -0.66 | -1.15 | B>B |
| chr16 | 17800000 | 17900000 | -0.65 | -1.25 | A>B |
| chr16 | 17900000 | 18000000 | -0.57 | -1.24 | A>B |
| chr16 | 18000000 | 18100000 | -0.55 | -1.25 | A>B |

| | | | | | |
|---|---|---|---|---|---|
| chr16 | 18100000 | 18200000 | 0.00 | -1.16 | NO |
| chr16 | 18200000 | 18300000 | 0.00 | 0.00 | NO |
| chr16 | 18300000 | 18400000 | 0.00 | 0.00 | NO |
| chr16 | 18400000 | 18500000 | 0.00 | 0.00 | NO |
| chr16 | 18500000 | 18600000 | 0.00 | 0.00 | NO |
| chr16 | 18600000 | 18700000 | 0.00 | 0.00 | NO |
| chr16 | 18700000 | 18800000 | 0.00 | 0.00 | NO |
| chr16 | 18800000 | 18900000 | 0.00 | 0.00 | NO |
| chr16 | 18900000 | 19000000 | 0.00 | 0.00 | NO |
| chr16 | 19000000 | 19100000 | 0.96 | 0.90 | A>A |
| chr16 | 19100000 | 19200000 | 0.94 | 0.94 | A>A |
| chr16 | 19200000 | 19300000 | 0.77 | 0.68 | A>A |
| chr16 | 19300000 | 19400000 | 0.70 | 0.73 | A>A |
| chr16 | 19400000 | 19500000 | 0.83 | 0.88 | A>A |
| chr16 | 19500000 | 19600000 | 0.61 | 0.57 | A>A |
| chr16 | 19600000 | 19700000 | 0.30 | 0.26 | A>A |
| chr16 | 19700000 | 19800000 | 0.40 | 0.29 | A>A |
| chr16 | 19800000 | 19900000 | 0.37 | 0.23 | A>A |
| chr16 | 19900000 | 20000000 | -0.14 | -0.42 | B>B |
| chr16 | 20000000 | 20100000 | -0.33 | -0.68 | B>B |
| chr16 | 20100000 | 20200000 | -0.37 | -0.68 | B>B |
| chr16 | 20200000 | 20300000 | -0.46 | -0.72 | B>B |
| chr16 | 20300000 | 20400000 | -0.30 | -0.49 | B>B |
| chr16 | 20400000 | 20500000 | 0.00 | 0.00 | NO |
| chr16 | 20500000 | 20600000 | 0.00 | 0.00 | NO |
| chr16 | 20600000 | 20700000 | -0.38 | -0.58 | B>B |
| chr16 | 20700000 | 20800000 | 0.08 | -0.04 | A>A |
| chr16 | 20800000 | 20900000 | 0.41 | 0.41 | A>A |
| chr16 | 20900000 | 21000000 | 0.60 | 0.49 | A>A |
| chr16 | 21000000 | 21100000 | 0.44 | 0.43 | A>A |
| chr16 | 21100000 | 21200000 | 0.52 | 0.57 | A>A |
| chr16 | 21200000 | 21300000 | 0.81 | 0.83 | A>A |
| chr16 | 21300000 | 21400000 | 0.00 | 0.00 | NO |
| chr16 | 21400000 | 21500000 | 0.00 | 0.00 | NO |
| chr16 | 21500000 | 21600000 | 0.00 | 0.00 | NO |
| chr16 | 21600000 | 21700000 | 0.81 | 0.63 | A>A |
| chr16 | 21700000 | 21800000 | 0.00 | 0.00 | NO |
| chr16 | 21800000 | 21900000 | 0.00 | 0.00 | NO |
| chr16 | 21900000 | 22000000 | 0.00 | 0.00 | NO |
| chr16 | 22000000 | 22100000 | 0.96 | 0.96 | A>A |
| chr16 | 22100000 | 22200000 | 0.95 | 0.91 | A>A |
| chr16 | 22200000 | 22300000 | 1.09 | 1.10 | A>A |
| chr16 | 22300000 | 22400000 | 0.83 | 0.85 | A>A |
| chr16 | 22400000 | 22500000 | 0.00 | 0.00 | NO |
| chr16 | 22500000 | 22600000 | 0.00 | 0.00 | NO |
| chr16 | 22600000 | 22700000 | 0.25 | -0.11 | A>A |
| chr16 | 22700000 | 22800000 | 0.04 | -0.25 | B>B |
| chr16 | 22800000 | 22900000 | 0.07 | -0.29 | B>B |
| chr16 | 22900000 | 23000000 | 0.07 | -0.22 | B>B |
| chr16 | 23000000 | 23100000 | 0.26 | -0.01 | A>A |
| chr16 | 23100000 | 23200000 | 0.55 | 0.36 | A>A |
| chr16 | 23200000 | 23300000 | 0.76 | 0.41 | A>A |
| chr16 | 23300000 | 23400000 | 0.88 | 0.63 | A>A |
| chr16 | 23400000 | 23500000 | 0.70 | 0.59 | A>A |
| chr16 | 23500000 | 23600000 | 0.94 | 0.76 | A>A |
| chr16 | 23600000 | 23700000 | 0.77 | 0.40 | A>A |

| | | | | | |
|---|---|---|---|---|---|
| chr16 | 23700000 | 23800000 | 0.39 | -0.20 | A>B |
| chr16 | 23800000 | 23900000 | 0.18 | -0.63 | A>B |
| chr16 | 23900000 | 24000000 | 0.14 | -0.84 | A>B |
| chr16 | 24000000 | 24100000 | -0.02 | -0.54 | B>B |
| chr16 | 24100000 | 24200000 | -0.18 | -0.83 | A>B |
| chr16 | 24200000 | 24300000 | -0.17 | -0.91 | A>B |
| chr16 | 24300000 | 24400000 | -0.04 | -0.76 | A>B |
| chr16 | 24400000 | 24500000 | 0.03 | -0.59 | A>B |
| chr16 | 24500000 | 24600000 | 0.35 | 0.11 | A>A |
| chr16 | 24600000 | 24700000 | 0.82 | 0.61 | A>A |
| chr16 | 24700000 | 24800000 | 0.89 | 0.59 | A>A |
| chr16 | 24800000 | 24900000 | 0.59 | 0.55 | A>A |
| chr16 | 24900000 | 25000000 | 0.67 | 0.59 | A>A |
| chr16 | 25000000 | 25100000 | 0.66 | 0.53 | A>A |
| chr16 | 25100000 | 25200000 | 0.39 | 0.32 | A>A |
| chr16 | 25200000 | 25300000 | -0.01 | -0.10 | B>B |
| chr16 | 25300000 | 25400000 | -0.56 | -0.83 | B>B |
| chr16 | 25400000 | 25500000 | -0.74 | -1.11 | B>B |
| chr16 | 25500000 | 25600000 | -0.61 | -1.13 | B>B |
| chr16 | 25600000 | 25700000 | -0.45 | -1.00 | B>B |
| chr16 | 25700000 | 25800000 | -0.57 | -0.90 | B>B |
| chr16 | 25800000 | 25900000 | -0.64 | -0.80 | B>B |
| chr16 | 25900000 | 26000000 | -0.69 | -0.79 | B>B |
| chr16 | 26000000 | 26100000 | -0.82 | -0.86 | B>B |
| chr16 | 26100000 | 26200000 | -0.81 | -0.96 | B>B |
| chr16 | 26200000 | 26300000 | -0.90 | -1.00 | B>B |
| chr16 | 26300000 | 26400000 | -0.88 | -1.02 | B>B |
| chr16 | 26400000 | 26500000 | -0.89 | -1.02 | B>B |
| chr16 | 26500000 | 26600000 | -0.90 | -1.07 | B>B |
| chr16 | 26600000 | 26700000 | -0.83 | -1.07 | B>B |
| chr16 | 26700000 | 26800000 | -0.70 | -1.10 | B>B |
| chr16 | 26800000 | 26900000 | -0.65 | -1.12 | B>B |
| chr16 | 26900000 | 27000000 | -0.63 | -1.06 | B>B |
| chr16 | 27000000 | 27100000 | -0.46 | -0.84 | B>B |
| chr16 | 27100000 | 27200000 | -0.14 | -0.29 | B>B |
| chr16 | 27200000 | 27300000 | 0.42 | 0.40 | A>A |
| chr16 | 27300000 | 27400000 | 0.41 | 0.49 | A>A |
| chr16 | 27400000 | 27500000 | 0.37 | 0.41 | A>A |
| chr16 | 27500000 | 27600000 | 0.69 | 0.81 | A>A |
| chr16 | 27600000 | 27700000 | 0.53 | 0.72 | A>A |
| chr16 | 27700000 | 27800000 | 0.58 | 0.79 | A>A |
| chr16 | 27800000 | 27900000 | 0.42 | 0.74 | A>A |
| chr16 | 27900000 | 28000000 | 0.61 | 0.73 | A>A |
| chr16 | 28000000 | 28100000 | 0.65 | 0.80 | A>A |
| chr16 | 28100000 | 28200000 | 1.10 | 1.10 | A>A |
| chr16 | 28200000 | 28300000 | 1.24 | 1.22 | A>A |
| chr16 | 28300000 | 28400000 | 0.00 | 0.00 | NO |
| chr16 | 28400000 | 28500000 | 0.00 | 0.00 | NO |
| chr16 | 28500000 | 28600000 | 1.38 | 1.41 | A>A |
| chr16 | 28600000 | 28700000 | 0.00 | 0.00 | NO |
| chr16 | 28700000 | 28800000 | 0.00 | 0.00 | NO |
| chr16 | 28800000 | 28900000 | 0.00 | 0.00 | NO |
| chr16 | 28900000 | 29000000 | 1.61 | 1.53 | A>A |
| chr16 | 29000000 | 29100000 | 0.00 | 0.00 | NO |
| chr16 | 29100000 | 29200000 | 0.00 | 0.00 | NO |
| chr16 | 29200000 | 29300000 | 0.00 | 0.00 | NO |

| | | | | | |
|---|---|---|---|---|---|
| chr16 | 29300000 | 29400000 | 0.00 | 0.00 | NO |
| chr16 | 29400000 | 29500000 | 0.00 | 0.00 | NO |
| chr16 | 29500000 | 29600000 | 0.00 | 0.00 | NO |
| chr16 | 29600000 | 29700000 | 0.00 | 0.00 | NO |
| chr16 | 29700000 | 29800000 | 1.11 | 1.07 | A>A |
| chr16 | 29800000 | 29900000 | 1.21 | 1.12 | A>A |
| chr16 | 29900000 | 30000000 | 1.36 | 1.27 | A>A |
| chr16 | 30000000 | 30100000 | 1.02 | 1.02 | A>A |
| chr16 | 30100000 | 30200000 | 1.19 | 1.11 | A>A |
| chr16 | 30200000 | 30300000 | 0.00 | 0.00 | NO |
| chr16 | 30300000 | 30400000 | 0.00 | 0.00 | NO |
| chr16 | 30400000 | 30500000 | 1.16 | 1.11 | A>A |
| chr16 | 30500000 | 30600000 | 1.47 | 1.29 | A>A |
| chr16 | 30600000 | 30700000 | 1.20 | 1.07 | A>A |
| chr16 | 30700000 | 30800000 | 1.39 | 1.24 | A>A |
| chr16 | 30800000 | 30900000 | 1.29 | 1.31 | A>A |
| chr16 | 30900000 | 31000000 | 1.10 | 1.12 | A>A |
| chr16 | 31000000 | 31100000 | 1.04 | 1.17 | A>A |
| chr16 | 31100000 | 31200000 | 0.95 | 1.02 | A>A |
| chr16 | 31200000 | 31300000 | 0.87 | 0.98 | A>A |
| chr16 | 31300000 | 31400000 | 0.69 | 0.94 | A>A |
| chr16 | 31400000 | 31500000 | 0.49 | 0.77 | A>A |
| chr16 | 31500000 | 31600000 | -0.45 | 0.10 | B>A |
| chr16 | 31600000 | 31700000 | 0.00 | 0.00 | NO |
| chr16 | 31700000 | 31800000 | 0.00 | 0.18 | NO |
| chr16 | 31800000 | 31900000 | 0.00 | 0.12 | NO |
| chr16 | 31900000 | 32000000 | 0.00 | 0.00 | NO |
| chr16 | 32000000 | 32100000 | 0.00 | 0.00 | NO |
| chr16 | 32100000 | 32200000 | 0.00 | 0.00 | NO |
| chr16 | 32200000 | 32300000 | 0.00 | 0.00 | NO |
| chr16 | 32300000 | 32400000 | 0.00 | 0.00 | NO |
| chr16 | 32400000 | 32500000 | 0.00 | 0.00 | NO |
| chr16 | 32500000 | 32600000 | 0.00 | -1.37 | NO |
| chr16 | 32600000 | 32700000 | 0.00 | 0.00 | NO |
| chr16 | 32700000 | 32800000 | 0.00 | 0.00 | NO |
| chr16 | 32800000 | 32900000 | 0.00 | 0.00 | NO |
| chr16 | 32900000 | 33000000 | 0.00 | 0.00 | NO |
| chr16 | 33000000 | 33100000 | 0.00 | 0.00 | NO |
| chr16 | 33100000 | 33200000 | 0.00 | 0.00 | NO |
| chr16 | 33200000 | 33300000 | 0.00 | 0.00 | NO |
| chr16 | 33300000 | 33400000 | 0.00 | 0.00 | NO |
| chr16 | 33400000 | 33500000 | 0.00 | 0.00 | NO |
| chr16 | 33500000 | 33600000 | 0.00 | 0.00 | NO |
| chr16 | 33600000 | 33700000 | 0.00 | 0.00 | NO |
| chr16 | 33700000 | 33800000 | 0.00 | 0.00 | NO |
| chr16 | 33800000 | 33900000 | 0.00 | 0.00 | NO |
| chr16 | 33900000 | 34000000 | 0.00 | 0.00 | NO |
| chr16 | 34000000 | 34100000 | 0.00 | 0.00 | NO |
| chr16 | 34100000 | 34200000 | 0.00 | 0.00 | NO |
| chr16 | 34200000 | 34300000 | -1.45 | -1.45 | B>B |
| chr16 | 34300000 | 34400000 | 0.00 | 0.00 | NO |
| chr16 | 34400000 | 34500000 | 0.00 | -1.50 | NO |
| chr16 | 34500000 | 34600000 | 0.00 | -1.58 | NO |
| chr16 | 34600000 | 34700000 | -1.40 | -1.28 | B>B |
| chr16 | 34700000 | 34800000 | 0.00 | -0.64 | NO |
| chr16 | 34800000 | 34900000 | 0.00 | -0.29 | NO |

| | | | | | |
|---|---|---|---|---|---|
| chr16 | 34900000 | 35000000 | 0.00 | -0.35 | NO |
| chr16 | 35000000 | 35100000 | -1.40 | -0.24 | B>A |
| chr16 | 35100000 | 35200000 | 0.00 | 0.00 | NO |
| chr16 | 35200000 | 35300000 | 0.00 | 0.00 | NO |
| chr16 | 35300000 | 35400000 | 0.00 | 0.00 | NO |
| chr16 | 35400000 | 35500000 | 0.00 | 0.00 | NO |
| chr16 | 35500000 | 35600000 | 0.00 | 0.00 | NO |
| chr16 | 35600000 | 35700000 | 0.00 | 0.00 | NO |
| chr16 | 35700000 | 35800000 | 0.00 | 0.00 | NO |
| chr16 | 35800000 | 35900000 | 0.00 | 0.00 | NO |
| chr16 | 35900000 | 36000000 | 0.00 | 0.00 | NO |
| chr16 | 36000000 | 36100000 | 0.00 | 0.00 | NO |
| chr16 | 36100000 | 36200000 | 0.00 | 0.00 | NO |
| chr16 | 36200000 | 36300000 | 0.00 | 0.00 | NO |
| chr16 | 36300000 | 36400000 | 0.00 | 0.00 | NO |
| chr16 | 36400000 | 36500000 | 0.00 | 0.00 | NO |
| chr16 | 36500000 | 36600000 | 0.00 | 0.00 | NO |
| chr16 | 36600000 | 36700000 | 0.00 | 0.00 | NO |
| chr16 | 36700000 | 36800000 | 0.00 | 0.00 | NO |
| chr16 | 36800000 | 36900000 | 0.00 | 0.00 | NO |
| chr16 | 36900000 | 37000000 | 0.00 | 0.00 | NO |
| chr16 | 37000000 | 37100000 | 0.00 | 0.00 | NO |
| chr16 | 37100000 | 37200000 | 0.00 | 0.00 | NO |
| chr16 | 37200000 | 37300000 | 0.00 | 0.00 | NO |
| chr16 | 37300000 | 37400000 | 0.00 | 0.00 | NO |
| chr16 | 37400000 | 37500000 | 0.00 | 0.00 | NO |
| chr16 | 37500000 | 37600000 | 0.00 | 0.00 | NO |
| chr16 | 37600000 | 37700000 | 0.00 | 0.00 | NO |
| chr16 | 37700000 | 37800000 | 0.00 | 0.00 | NO |
| chr16 | 37800000 | 37900000 | 0.00 | 0.00 | NO |
| chr16 | 37900000 | 38000000 | 0.00 | 0.00 | NO |
| chr16 | 38000000 | 38100000 | 0.00 | 0.00 | NO |
| chr16 | 38100000 | 38200000 | 0.00 | 0.00 | NO |
| chr16 | 38200000 | 38300000 | 0.00 | 0.00 | NO |
| chr16 | 38300000 | 38400000 | 0.00 | 0.00 | NO |
| chr16 | 38400000 | 38500000 | 0.00 | 0.00 | NO |
| chr16 | 38500000 | 38600000 | 0.00 | 0.00 | NO |
| chr16 | 38600000 | 38700000 | 0.00 | 0.00 | NO |
| chr16 | 38700000 | 38800000 | 0.00 | 0.00 | NO |
| chr16 | 38800000 | 38900000 | 0.00 | 0.00 | NO |
| chr16 | 38900000 | 39000000 | 0.00 | 0.00 | NO |
| chr16 | 39000000 | 39100000 | 0.00 | 0.00 | NO |
| chr16 | 39100000 | 39200000 | 0.00 | 0.00 | NO |
| chr16 | 39200000 | 39300000 | 0.00 | 0.00 | NO |
| chr16 | 39300000 | 39400000 | 0.00 | 0.00 | NO |
| chr16 | 39400000 | 39500000 | 0.00 | 0.00 | NO |
| chr16 | 39500000 | 39600000 | 0.00 | 0.00 | NO |
| chr16 | 39600000 | 39700000 | 0.00 | 0.00 | NO |
| chr16 | 39700000 | 39800000 | 0.00 | 0.00 | NO |
| chr16 | 39800000 | 39900000 | 0.00 | 0.00 | NO |
| chr16 | 39900000 | 40000000 | 0.00 | 0.00 | NO |
| chr16 | 40000000 | 40100000 | 0.00 | 0.00 | NO |
| chr16 | 40100000 | 40200000 | 0.00 | 0.00 | NO |
| chr16 | 40200000 | 40300000 | 0.00 | 0.00 | NO |
| chr16 | 40300000 | 40400000 | 0.00 | 0.00 | NO |
| chr16 | 40400000 | 40500000 | 0.00 | 0.00 | NO |

| | | | | | |
|---|---|---|---|---|---|
| chr16 | 40500000 | 40600000 | 0.00 | 0.00 | NO |
| chr16 | 40600000 | 40700000 | 0.00 | 0.00 | NO |
| chr16 | 40700000 | 40800000 | 0.00 | 0.00 | NO |
| chr16 | 40800000 | 40900000 | 0.00 | 0.00 | NO |
| chr16 | 40900000 | 41000000 | 0.00 | 0.00 | NO |
| chr16 | 41000000 | 41100000 | 0.00 | 0.00 | NO |
| chr16 | 41100000 | 41200000 | 0.00 | 0.00 | NO |
| chr16 | 41200000 | 41300000 | 0.00 | 0.00 | NO |
| chr16 | 41300000 | 41400000 | 0.00 | 0.00 | NO |
| chr16 | 41400000 | 41500000 | 0.00 | 0.00 | NO |
| chr16 | 41500000 | 41600000 | 0.00 | 0.00 | NO |
| chr16 | 41600000 | 41700000 | 0.00 | 0.00 | NO |
| chr16 | 41700000 | 41800000 | 0.00 | 0.00 | NO |
| chr16 | 41800000 | 41900000 | 0.00 | 0.00 | NO |
| chr16 | 41900000 | 42000000 | 0.00 | 0.00 | NO |
| chr16 | 42000000 | 42100000 | 0.00 | 0.00 | NO |
| chr16 | 42100000 | 42200000 | 0.00 | 0.00 | NO |
| chr16 | 42200000 | 42300000 | 0.00 | 0.00 | NO |
| chr16 | 42300000 | 42400000 | 0.00 | 0.00 | NO |
| chr16 | 42400000 | 42500000 | 0.00 | 0.00 | NO |
| chr16 | 42500000 | 42600000 | 0.00 | 0.00 | NO |
| chr16 | 42600000 | 42700000 | 0.00 | 0.00 | NO |
| chr16 | 42700000 | 42800000 | 0.00 | 0.00 | NO |
| chr16 | 42800000 | 42900000 | 0.00 | 0.00 | NO |
| chr16 | 42900000 | 43000000 | 0.00 | 0.00 | NO |
| chr16 | 43000000 | 43100000 | 0.00 | 0.00 | NO |
| chr16 | 43100000 | 43200000 | 0.00 | 0.00 | NO |
| chr16 | 43200000 | 43300000 | 0.00 | 0.00 | NO |
| chr16 | 43300000 | 43400000 | 0.00 | 0.00 | NO |
| chr16 | 43400000 | 43500000 | 0.00 | 0.00 | NO |
| chr16 | 43500000 | 43600000 | 0.00 | 0.00 | NO |
| chr16 | 43600000 | 43700000 | 0.00 | 0.00 | NO |
| chr16 | 43700000 | 43800000 | 0.00 | 0.00 | NO |
| chr16 | 43800000 | 43900000 | 0.00 | 0.00 | NO |
| chr16 | 43900000 | 44000000 | 0.00 | 0.00 | NO |
| chr16 | 44000000 | 44100000 | 0.00 | 0.00 | NO |
| chr16 | 44100000 | 44200000 | 0.00 | 0.00 | NO |
| chr16 | 44200000 | 44300000 | 0.00 | 0.00 | NO |
| chr16 | 44300000 | 44400000 | 0.00 | 0.00 | NO |
| chr16 | 44400000 | 44500000 | 0.00 | 0.00 | NO |
| chr16 | 44500000 | 44600000 | 0.00 | 0.00 | NO |
| chr16 | 44600000 | 44700000 | 0.00 | 0.00 | NO |
| chr16 | 44700000 | 44800000 | 0.00 | 0.00 | NO |
| chr16 | 44800000 | 44900000 | 0.00 | 0.00 | NO |
| chr16 | 44900000 | 45000000 | 0.00 | 0.00 | NO |
| chr16 | 45000000 | 45100000 | 0.00 | 0.00 | NO |
| chr16 | 45100000 | 45200000 | 0.00 | 0.00 | NO |
| chr16 | 45200000 | 45300000 | 0.00 | 0.00 | NO |
| chr16 | 45300000 | 45400000 | 0.00 | 0.00 | NO |
| chr16 | 45400000 | 45500000 | 0.00 | 0.00 | NO |
| chr16 | 45500000 | 45600000 | 0.00 | 0.00 | NO |
| chr16 | 45600000 | 45700000 | 0.00 | 0.00 | NO |
| chr16 | 45700000 | 45800000 | 0.00 | 0.00 | NO |
| chr16 | 45800000 | 45900000 | 0.00 | 0.00 | NO |
| chr16 | 45900000 | 46000000 | 0.00 | 0.00 | NO |
| chr16 | 46000000 | 46100000 | 0.00 | 0.00 | NO |

| | | | | | |
|---|---|---|---|---|---|
| chr16 | 46100000 | 46200000 | 0.00 | 0.00 | NO |
| chr16 | 46200000 | 46300000 | 0.00 | 0.00 | NO |
| chr16 | 46300000 | 46400000 | 0.00 | 0.00 | NO |
| chr16 | 46400000 | 46500000 | 0.00 | 0.00 | NO |
| chr16 | 46500000 | 46600000 | -1.29 | -1.57 | B>B |
| chr16 | 46600000 | 46700000 | -0.27 | -0.40 | B>B |
| chr16 | 46700000 | 46800000 | 0.57 | 0.35 | A>A |
| chr16 | 46800000 | 46900000 | 0.66 | 0.53 | A>A |
| chr16 | 46900000 | 47000000 | 0.63 | 0.59 | A>A |
| chr16 | 47000000 | 47100000 | 0.50 | 0.52 | A>A |
| chr16 | 47100000 | 47200000 | 0.24 | 0.37 | A>A |
| chr16 | 47200000 | 47300000 | -0.02 | 0.15 | A>A |
| chr16 | 47300000 | 47400000 | 0.01 | 0.17 | A>A |
| chr16 | 47400000 | 47500000 | 0.16 | 0.27 | A>A |
| chr16 | 47500000 | 47600000 | 0.23 | 0.34 | A>A |
| chr16 | 47600000 | 47700000 | 0.26 | 0.30 | A>A |
| chr16 | 47700000 | 47800000 | 0.23 | 0.22 | A>A |
| chr16 | 47800000 | 47900000 | 0.15 | 0.26 | A>A |
| chr16 | 47900000 | 48000000 | 0.29 | 0.19 | A>A |
| chr16 | 48000000 | 48100000 | 0.39 | 0.22 | A>A |
| chr16 | 48100000 | 48200000 | 0.38 | 0.33 | A>A |
| chr16 | 48200000 | 48300000 | 0.36 | 0.33 | A>A |
| chr16 | 48300000 | 48400000 | 0.32 | 0.34 | A>A |
| chr16 | 48400000 | 48500000 | 0.56 | 0.61 | A>A |
| chr16 | 48500000 | 48600000 | 0.37 | 0.40 | A>A |
| chr16 | 48600000 | 48700000 | -0.11 | -0.11 | B>B |
| chr16 | 48700000 | 48800000 | -0.77 | -1.08 | B>B |
| chr16 | 48800000 | 48900000 | -0.96 | -1.45 | B>B |
| chr16 | 48900000 | 49000000 | -0.84 | -1.50 | A>B |
| chr16 | 49000000 | 49100000 | -0.88 | -1.48 | A>B |
| chr16 | 49100000 | 49200000 | -1.05 | -1.54 | B>B |
| chr16 | 49200000 | 49300000 | -1.06 | -1.44 | B>B |
| chr16 | 49300000 | 49400000 | -0.83 | -1.06 | B>B |
| chr16 | 49400000 | 49500000 | -0.51 | -0.70 | B>B |
| chr16 | 49500000 | 49600000 | -0.20 | -0.41 | B>B |
| chr16 | 49600000 | 49700000 | -0.15 | -0.30 | B>B |
| chr16 | 49700000 | 49800000 | 0.00 | -0.26 | B>B |
| chr16 | 49800000 | 49900000 | 0.14 | -0.15 | B>B |
| chr16 | 49900000 | 50000000 | 0.10 | -0.03 | A>A |
| chr16 | 50000000 | 50100000 | 0.37 | 0.23 | A>A |
| chr16 | 50100000 | 50200000 | 0.81 | 0.57 | A>A |
| chr16 | 50200000 | 50300000 | 0.69 | 0.64 | A>A |
| chr16 | 50300000 | 50400000 | 0.49 | 0.45 | A>A |
| chr16 | 50400000 | 50500000 | 0.12 | 0.17 | A>A |
| chr16 | 50500000 | 50600000 | 0.21 | 0.09 | A>A |
| chr16 | 50600000 | 50700000 | 0.15 | 0.15 | A>A |
| chr16 | 50700000 | 50800000 | 0.12 | 0.22 | A>A |
| chr16 | 50800000 | 50900000 | -0.10 | 0.06 | B>B |
| chr16 | 50900000 | 51000000 | -0.19 | -0.04 | B>B |
| chr16 | 51000000 | 51100000 | -0.17 | -0.05 | B>B |
| chr16 | 51100000 | 51200000 | -0.23 | 0.05 | B>B |
| chr16 | 51200000 | 51300000 | -0.22 | 0.13 | B>B |
| chr16 | 51300000 | 51400000 | -0.27 | 0.19 | B>B |
| chr16 | 51400000 | 51500000 | -0.16 | 0.29 | A>A |
| chr16 | 51500000 | 51600000 | -0.19 | 0.31 | A>A |
| chr16 | 51600000 | 51700000 | -0.11 | 0.32 | A>A |

| | | | | | |
|---|---|---|---|---|---|
| chr16 | 51700000 | 51800000 | -0.26 | 0.20 | B>B |
| chr16 | 51800000 | 51900000 | -0.26 | 0.21 | B>B |
| chr16 | 51900000 | 52000000 | -0.37 | 0.21 | B>A |
| chr16 | 52000000 | 52100000 | -0.59 | 0.11 | B>A |
| chr16 | 52100000 | 52200000 | -0.80 | 0.21 | B>A |
| chr16 | 52200000 | 52300000 | -0.61 | 0.29 | B>A |
| chr16 | 52300000 | 52400000 | -0.58 | 0.39 | B>A |
| chr16 | 52400000 | 52500000 | -0.43 | 0.40 | B>A |
| chr16 | 52500000 | 52600000 | -0.20 | 0.41 | B>A |
| chr16 | 52600000 | 52700000 | -0.12 | 0.51 | B>A |
| chr16 | 52700000 | 52800000 | -0.39 | 0.44 | B>A |
| chr16 | 52800000 | 52900000 | -0.48 | 0.35 | B>A |
| chr16 | 52900000 | 53000000 | -0.52 | 0.26 | B>A |
| chr16 | 53000000 | 53100000 | -0.18 | 0.33 | B>A |
| chr16 | 53100000 | 53200000 | 0.26 | 0.67 | A>A |
| chr16 | 53200000 | 53300000 | 0.33 | 0.67 | A>A |
| chr16 | 53300000 | 53400000 | 0.26 | 0.48 | A>A |
| chr16 | 53400000 | 53500000 | 0.42 | 0.56 | A>A |
| chr16 | 53500000 | 53600000 | 0.25 | 0.34 | A>A |
| chr16 | 53600000 | 53700000 | 0.19 | 0.28 | A>A |
| chr16 | 53700000 | 53800000 | 0.35 | 0.28 | A>A |
| chr16 | 53800000 | 53900000 | 0.26 | 0.23 | A>A |
| chr16 | 53900000 | 54000000 | 0.11 | 0.03 | A>A |
| chr16 | 54000000 | 54100000 | -0.04 | -0.08 | B>B |
| chr16 | 54100000 | 54200000 | -0.05 | -0.11 | B>B |
| chr16 | 54200000 | 54300000 | 0.01 | -0.06 | B>B |
| chr16 | 54300000 | 54400000 | 0.00 | -0.09 | B>B |
| chr16 | 54400000 | 54500000 | -0.01 | -0.16 | B>B |
| chr16 | 54500000 | 54600000 | -0.19 | -0.27 | B>B |
| chr16 | 54600000 | 54700000 | -0.25 | -0.27 | B>B |
| chr16 | 54700000 | 54800000 | -0.40 | -0.41 | B>B |
| chr16 | 54800000 | 54900000 | -0.22 | -0.31 | B>B |
| chr16 | 54900000 | 55000000 | -0.08 | -0.14 | B>B |
| chr16 | 55000000 | 55100000 | -0.20 | -0.38 | B>B |
| chr16 | 55100000 | 55200000 | -0.50 | -0.84 | B>B |
| chr16 | 55200000 | 55300000 | -0.60 | -0.97 | B>B |
| chr16 | 55300000 | 55400000 | -0.58 | -0.84 | B>B |
| chr16 | 55400000 | 55500000 | -0.54 | -0.66 | B>B |
| chr16 | 55500000 | 55600000 | -0.37 | -0.16 | B>B |
| chr16 | 55600000 | 55700000 | -0.46 | -0.37 | B>B |
| chr16 | 55700000 | 55800000 | -0.41 | -0.14 | B>B |
| chr16 | 55800000 | 55900000 | -0.42 | -0.22 | B>B |
| chr16 | 55900000 | 56000000 | -0.78 | -0.65 | B>B |
| chr16 | 56000000 | 56100000 | -0.80 | -0.91 | B>B |
| chr16 | 56100000 | 56200000 | -0.72 | -0.91 | B>B |
| chr16 | 56200000 | 56300000 | -0.36 | -0.59 | B>B |
| chr16 | 56300000 | 56400000 | -0.10 | -0.22 | B>B |
| chr16 | 56400000 | 56500000 | 0.59 | 0.44 | A>A |
| chr16 | 56500000 | 56600000 | 0.81 | 0.72 | A>A |
| chr16 | 56600000 | 56700000 | 0.98 | 0.90 | A>A |
| chr16 | 56700000 | 56800000 | 0.85 | 0.85 | A>A |
| chr16 | 56800000 | 56900000 | 0.85 | 0.93 | A>A |
| chr16 | 56900000 | 57000000 | 1.05 | 1.01 | A>A |
| chr16 | 57000000 | 57100000 | 1.10 | 1.11 | A>A |
| chr16 | 57100000 | 57200000 | 1.07 | 1.05 | A>A |
| chr16 | 57200000 | 57300000 | 1.11 | 1.15 | A>A |

| | | | | | |
|---|---|---|---|---|---|
| chr16 | 57300000 | 57400000 | 1.14 | 1.15 | A>A |
| chr16 | 57400000 | 57500000 | 1.10 | 1.18 | A>A |
| chr16 | 57500000 | 57600000 | 1.23 | 1.15 | A>A |
| chr16 | 57600000 | 57700000 | 1.09 | 1.11 | A>A |
| chr16 | 57700000 | 57800000 | 1.05 | 1.04 | A>A |
| chr16 | 57800000 | 57900000 | 1.00 | 1.10 | A>A |
| chr16 | 57900000 | 58000000 | 1.00 | 1.02 | A>A |
| chr16 | 58000000 | 58100000 | 0.88 | 0.99 | A>A |
| chr16 | 58100000 | 58200000 | 0.82 | 0.92 | A>A |
| chr16 | 58200000 | 58300000 | 0.89 | 0.92 | A>A |
| chr16 | 58300000 | 58400000 | 0.69 | 0.71 | A>A |
| chr16 | 58400000 | 58500000 | 0.84 | 0.83 | A>A |
| chr16 | 58500000 | 58600000 | 0.72 | 0.80 | A>A |
| chr16 | 58600000 | 58700000 | 0.80 | 0.75 | A>A |
| chr16 | 58700000 | 58800000 | 0.31 | 0.32 | A>A |
| chr16 | 58800000 | 58900000 | -0.52 | -0.52 | B>B |
| chr16 | 58900000 | 59000000 | -0.56 | -0.64 | B>B |
| chr16 | 59000000 | 59100000 | -0.42 | -0.56 | B>B |
| chr16 | 59100000 | 59200000 | -0.35 | -0.44 | B>B |
| chr16 | 59200000 | 59300000 | -0.23 | -0.35 | B>B |
| chr16 | 59300000 | 59400000 | -0.21 | -0.35 | B>B |
| chr16 | 59400000 | 59500000 | -0.24 | -0.45 | B>B |
| chr16 | 59500000 | 59600000 | -0.20 | -0.33 | B>B |
| chr16 | 59600000 | 59700000 | -0.25 | -0.41 | B>B |
| chr16 | 59700000 | 59800000 | -0.33 | -0.47 | B>B |
| chr16 | 59800000 | 59900000 | -0.33 | -0.43 | B>B |
| chr16 | 59900000 | 60000000 | -0.25 | -0.40 | B>B |
| chr16 | 60000000 | 60100000 | -0.30 | -0.41 | B>B |
| chr16 | 60100000 | 60200000 | -0.40 | -0.47 | B>B |
| chr16 | 60200000 | 60300000 | -0.51 | -0.55 | B>B |
| chr16 | 60300000 | 60400000 | -0.72 | -0.70 | B>B |
| chr16 | 60400000 | 60500000 | -0.82 | -0.83 | B>B |
| chr16 | 60500000 | 60600000 | -0.80 | -0.84 | B>B |
| chr16 | 60600000 | 60700000 | -0.82 | -0.89 | B>B |
| chr16 | 60700000 | 60800000 | -0.82 | -0.91 | B>B |
| chr16 | 60800000 | 60900000 | -0.86 | -0.92 | B>B |
| chr16 | 60900000 | 61000000 | -0.84 | -0.93 | B>B |
| chr16 | 61000000 | 61100000 | -0.81 | -0.88 | B>B |
| chr16 | 61100000 | 61200000 | -0.84 | -0.90 | B>B |
| chr16 | 61200000 | 61300000 | -0.85 | -0.92 | B>B |
| chr16 | 61300000 | 61400000 | -0.88 | -0.85 | B>B |
| chr16 | 61400000 | 61500000 | -0.87 | -0.87 | B>B |
| chr16 | 61500000 | 61600000 | -0.90 | -0.89 | B>B |
| chr16 | 61600000 | 61700000 | -0.82 | -0.87 | B>B |
| chr16 | 61700000 | 61800000 | -0.86 | -0.89 | B>B |
| chr16 | 61800000 | 61900000 | -0.84 | -0.85 | B>B |
| chr16 | 61900000 | 62000000 | -0.89 | -0.85 | B>B |
| chr16 | 62000000 | 62100000 | -0.82 | -0.83 | B>B |
| chr16 | 62100000 | 62200000 | -0.77 | -0.75 | B>B |
| chr16 | 62200000 | 62300000 | -0.79 | -0.78 | B>B |
| chr16 | 62300000 | 62400000 | -0.87 | -0.88 | B>B |
| chr16 | 62400000 | 62500000 | -0.89 | -0.90 | B>B |
| chr16 | 62500000 | 62600000 | -0.92 | -0.95 | B>B |
| chr16 | 62600000 | 62700000 | -0.89 | -0.91 | B>B |
| chr16 | 62700000 | 62800000 | -0.89 | -0.92 | B>B |
| chr16 | 62800000 | 62900000 | -0.87 | -0.88 | B>B |

| | | | | | |
|---|---|---|---|---|---|
| chr16 | 62900000 | 63000000 | -0.87 | -0.86 | B>B |
| chr16 | 63000000 | 63100000 | -0.84 | -0.81 | B>B |
| chr16 | 63100000 | 63200000 | -0.79 | -0.56 | B>B |
| chr16 | 63200000 | 63300000 | -0.75 | -0.45 | B>B |
| chr16 | 63300000 | 63400000 | -0.63 | -0.39 | B>B |
| chr16 | 63400000 | 63500000 | -0.58 | -0.15 | B>B |
| chr16 | 63500000 | 63600000 | -0.63 | -0.11 | B>A |
| chr16 | 63600000 | 63700000 | -0.51 | 0.01 | B>A |
| chr16 | 63700000 | 63800000 | -0.61 | -0.14 | B>B |
| chr16 | 63800000 | 63900000 | -0.74 | -0.36 | B>B |
| chr16 | 63900000 | 64000000 | -0.77 | -0.44 | B>B |
| chr16 | 64000000 | 64100000 | -0.81 | -0.35 | B>B |
| chr16 | 64100000 | 64200000 | -0.80 | -0.32 | B>B |
| chr16 | 64200000 | 64300000 | -0.81 | -0.21 | B>A |
| chr16 | 64300000 | 64400000 | -0.78 | -0.18 | B>A |
| chr16 | 64400000 | 64500000 | -0.76 | -0.15 | B>A |
| chr16 | 64500000 | 64600000 | -0.81 | -0.52 | B>B |
| chr16 | 64600000 | 64700000 | -0.83 | -0.72 | B>B |
| chr16 | 64700000 | 64800000 | -0.88 | -0.82 | B>B |
| chr16 | 64800000 | 64900000 | -0.87 | -0.88 | B>B |
| chr16 | 64900000 | 65000000 | -0.78 | -0.93 | B>B |
| chr16 | 65000000 | 65100000 | -0.75 | -0.88 | B>B |
| chr16 | 65100000 | 65200000 | -0.78 | -0.76 | B>B |
| chr16 | 65200000 | 65300000 | -0.93 | -1.04 | B>B |
| chr16 | 65300000 | 65400000 | -1.01 | -1.13 | B>B |
| chr16 | 65400000 | 65500000 | -1.07 | -1.21 | B>B |
| chr16 | 65500000 | 65600000 | -1.07 | -1.20 | B>B |
| chr16 | 65600000 | 65700000 | -1.01 | -1.18 | B>B |
| chr16 | 65700000 | 65800000 | -0.92 | -1.17 | B>B |
| chr16 | 65800000 | 65900000 | -0.83 | -1.23 | B>B |
| chr16 | 65900000 | 66000000 | -0.80 | -1.15 | B>B |
| chr16 | 66000000 | 66100000 | -0.76 | -1.15 | B>B |
| chr16 | 66100000 | 66200000 | -0.62 | -1.20 | B>B |
| chr16 | 66200000 | 66300000 | -0.40 | -0.94 | B>B |
| chr16 | 66300000 | 66400000 | 0.03 | -0.12 | B>B |
| chr16 | 66400000 | 66500000 | 0.68 | 0.64 | A>A |
| chr16 | 66500000 | 66600000 | 0.98 | 0.84 | A>A |
| chr16 | 66600000 | 66700000 | 1.01 | 1.02 | A>A |
| chr16 | 66700000 | 66800000 | 1.27 | 1.21 | A>A |
| chr16 | 66800000 | 66900000 | 1.40 | 1.29 | A>A |
| chr16 | 66900000 | 67000000 | 1.27 | 1.33 | A>A |
| chr16 | 67000000 | 67100000 | 1.41 | 1.30 | A>A |
| chr16 | 67100000 | 67200000 | 1.32 | 1.32 | A>A |
| chr16 | 67200000 | 67300000 | 1.42 | 1.34 | A>A |
| chr16 | 67300000 | 67400000 | 1.38 | 1.38 | A>A |
| chr16 | 67400000 | 67500000 | 1.23 | 1.23 | A>A |
| chr16 | 67500000 | 67600000 | 1.10 | 1.17 | A>A |
| chr16 | 67600000 | 67700000 | 1.18 | 1.21 | A>A |
| chr16 | 67700000 | 67800000 | 1.12 | 1.16 | A>A |
| chr16 | 67800000 | 67900000 | 1.23 | 1.13 | A>A |
| chr16 | 67900000 | 68000000 | 1.20 | 1.15 | A>A |
| chr16 | 68000000 | 68100000 | 1.02 | 0.99 | A>A |
| chr16 | 68100000 | 68200000 | 0.95 | 1.00 | A>A |
| chr16 | 68200000 | 68300000 | 0.88 | 0.90 | A>A |
| chr16 | 68300000 | 68400000 | 0.93 | 0.93 | A>A |
| chr16 | 68400000 | 68500000 | 0.90 | 0.87 | A>A |

| | | | | | |
|---|---|---|---|---|---|
| chr16 | 68500000 | 68600000 | 0.72 | 0.83 | A>A |
| chr16 | 68600000 | 68700000 | 0.68 | 0.75 | A>A |
| chr16 | 68700000 | 68800000 | 0.83 | 0.81 | A>A |
| chr16 | 68800000 | 68900000 | 0.73 | 0.71 | A>A |
| chr16 | 68900000 | 69000000 | 0.64 | 0.70 | A>A |
| chr16 | 69000000 | 69100000 | 0.87 | 0.79 | A>A |
| chr16 | 69100000 | 69200000 | 0.69 | 0.69 | A>A |
| chr16 | 69200000 | 69300000 | 0.84 | 0.79 | A>A |
| chr16 | 69300000 | 69400000 | 0.85 | 0.80 | A>A |
| chr16 | 69400000 | 69500000 | 0.75 | 0.72 | A>A |
| chr16 | 69500000 | 69600000 | 0.89 | 0.88 | A>A |
| chr16 | 69600000 | 69700000 | 0.79 | 0.73 | A>A |
| chr16 | 69700000 | 69800000 | 0.64 | 0.77 | A>A |
| chr16 | 69800000 | 69900000 | 0.91 | 0.89 | A>A |
| chr16 | 69900000 | 70000000 | 0.93 | 0.89 | A>A |
| chr16 | 70000000 | 70100000 | 0.00 | 0.00 | NO |
| chr16 | 70100000 | 70200000 | 0.00 | 0.00 | NO |
| chr16 | 70200000 | 70300000 | 0.00 | 0.00 | NO |
| chr16 | 70300000 | 70400000 | 1.06 | 1.04 | A>A |
| chr16 | 70400000 | 70500000 | 1.11 | 1.04 | A>A |
| chr16 | 70500000 | 70600000 | 0.99 | 0.92 | A>A |
| chr16 | 70600000 | 70700000 | 0.78 | 0.70 | A>A |
| chr16 | 70700000 | 70800000 | 0.65 | 0.57 | A>A |
| chr16 | 70800000 | 70900000 | 0.00 | 0.00 | NO |
| chr16 | 70900000 | 71000000 | 0.00 | 0.00 | NO |
| chr16 | 71000000 | 71100000 | 0.00 | 0.00 | NO |
| chr16 | 71100000 | 71200000 | 0.00 | 0.00 | NO |
| chr16 | 71200000 | 71300000 | -0.46 | -0.41 | B>B |
| chr16 | 71300000 | 71400000 | -0.13 | -0.24 | B>B |
| chr16 | 71400000 | 71500000 | 0.13 | 0.20 | A>A |
| chr16 | 71500000 | 71600000 | 0.30 | 0.43 | A>A |
| chr16 | 71600000 | 71700000 | 0.45 | 0.55 | A>A |
| chr16 | 71700000 | 71800000 | 0.68 | 0.72 | A>A |
| chr16 | 71800000 | 71900000 | 0.88 | 0.82 | A>A |
| chr16 | 71900000 | 72000000 | 0.70 | 0.72 | A>A |
| chr16 | 72000000 | 72100000 | 0.34 | 0.37 | A>A |
| chr16 | 72100000 | 72200000 | -0.11 | -0.02 | B>B |
| chr16 | 72200000 | 72300000 | -0.48 | -0.22 | B>B |
| chr16 | 72300000 | 72400000 | -0.58 | -0.12 | B>B |
| chr16 | 72400000 | 72500000 | -0.59 | -0.12 | B>B |
| chr16 | 72500000 | 72600000 | -0.35 | -0.17 | B>B |
| chr16 | 72600000 | 72700000 | -0.19 | -0.09 | B>B |
| chr16 | 72700000 | 72800000 | -0.27 | -0.12 | B>B |
| chr16 | 72800000 | 72900000 | -0.14 | 0.01 | B>B |
| chr16 | 72900000 | 73000000 | 0.16 | 0.19 | A>A |
| chr16 | 73000000 | 73100000 | 0.15 | 0.25 | A>A |
| chr16 | 73100000 | 73200000 | 0.22 | 0.18 | A>A |
| chr16 | 73200000 | 73300000 | -0.11 | -0.02 | B>B |
| chr16 | 73300000 | 73400000 | -0.30 | -0.30 | B>B |
| chr16 | 73400000 | 73500000 | -0.38 | -0.42 | B>B |
| chr16 | 73500000 | 73600000 | -0.48 | -0.47 | B>B |
| chr16 | 73600000 | 73700000 | -0.56 | -0.48 | B>B |
| chr16 | 73700000 | 73800000 | -0.71 | -0.59 | B>B |
| chr16 | 73800000 | 73900000 | -0.77 | -0.66 | B>B |
| chr16 | 73900000 | 74000000 | -0.62 | -0.59 | B>B |
| chr16 | 74000000 | 74100000 | -0.72 | -0.68 | B>B |

| | | | | | |
|---|---|---|---|---|---|
| chr16 | 74100000 | 74200000 | -0.70 | -0.62 | B>B |
| chr16 | 74200000 | 74300000 | -0.50 | -0.57 | B>B |
| chr16 | 74300000 | 74400000 | 0.05 | 0.02 | A>A |
| chr16 | 74400000 | 74500000 | 0.00 | 0.00 | NO |
| chr16 | 74500000 | 74600000 | 0.80 | 0.75 | A>A |
| chr16 | 74600000 | 74700000 | 0.79 | 0.81 | A>A |
| chr16 | 74700000 | 74800000 | 0.97 | 0.96 | A>A |
| chr16 | 74800000 | 74900000 | 0.92 | 0.97 | A>A |
| chr16 | 74900000 | 75000000 | 0.89 | 0.90 | A>A |
| chr16 | 75000000 | 75100000 | 0.97 | 1.00 | A>A |
| chr16 | 75100000 | 75200000 | 0.80 | 0.97 | A>A |
| chr16 | 75200000 | 75300000 | 0.56 | 0.78 | A>A |
| chr16 | 75300000 | 75400000 | 0.71 | 0.87 | A>A |
| chr16 | 75400000 | 75500000 | 0.50 | 0.70 | A>A |
| chr16 | 75500000 | 75600000 | 0.46 | 0.50 | A>A |
| chr16 | 75600000 | 75700000 | 0.19 | 0.25 | A>A |
| chr16 | 75700000 | 75800000 | -0.37 | -0.27 | B>B |
| chr16 | 75800000 | 75900000 | -0.69 | -0.57 | B>B |
| chr16 | 75900000 | 76000000 | -0.94 | -0.94 | B>B |
| chr16 | 76000000 | 76100000 | -1.08 | -0.94 | B>B |
| chr16 | 76100000 | 76200000 | -1.09 | -0.88 | B>B |
| chr16 | 76200000 | 76300000 | -1.03 | -0.87 | B>B |
| chr16 | 76300000 | 76400000 | -0.94 | -0.86 | B>B |
| chr16 | 76400000 | 76500000 | -0.97 | -0.81 | B>B |
| chr16 | 76500000 | 76600000 | -0.92 | -0.85 | B>B |
| chr16 | 76600000 | 76700000 | -0.80 | -0.60 | B>B |
| chr16 | 76700000 | 76800000 | -0.69 | -0.53 | B>B |
| chr16 | 76800000 | 76900000 | -0.57 | -0.42 | B>B |
| chr16 | 76900000 | 77000000 | -0.31 | -0.27 | B>B |
| chr16 | 77000000 | 77100000 | -0.55 | -0.42 | B>B |
| chr16 | 77100000 | 77200000 | -0.59 | -0.54 | B>B |
| chr16 | 77200000 | 77300000 | -0.21 | -0.18 | B>B |
| chr16 | 77300000 | 77400000 | 0.12 | 0.10 | A>A |
| chr16 | 77400000 | 77500000 | -0.02 | -0.09 | B>B |
| chr16 | 77500000 | 77600000 | -0.16 | -0.22 | B>B |
| chr16 | 77600000 | 77700000 | -0.26 | -0.30 | B>B |
| chr16 | 77700000 | 77800000 | -0.25 | -0.41 | B>B |
| chr16 | 77800000 | 77900000 | -0.28 | -0.49 | B>B |
| chr16 | 77900000 | 78000000 | -0.24 | -0.30 | B>B |
| chr16 | 78000000 | 78100000 | -0.41 | -0.48 | B>B |
| chr16 | 78100000 | 78200000 | -0.36 | -0.28 | B>B |
| chr16 | 78200000 | 78300000 | -0.25 | -0.05 | B>B |
| chr16 | 78300000 | 78400000 | -0.42 | -0.16 | B>B |
| chr16 | 78400000 | 78500000 | -0.47 | -0.27 | B>B |
| chr16 | 78500000 | 78600000 | -0.62 | -0.31 | B>B |
| chr16 | 78600000 | 78700000 | -0.65 | -0.33 | B>B |
| chr16 | 78700000 | 78800000 | -0.75 | -0.40 | B>B |
| chr16 | 78800000 | 78900000 | -0.81 | -0.70 | B>B |
| chr16 | 78900000 | 79000000 | -0.81 | -0.92 | B>B |
| chr16 | 79000000 | 79100000 | -0.76 | -0.92 | B>B |
| chr16 | 79100000 | 79200000 | -0.78 | -0.95 | B>B |
| chr16 | 79200000 | 79300000 | -0.74 | -0.94 | B>B |
| chr16 | 79300000 | 79400000 | -0.65 | -0.84 | B>B |
| chr16 | 79400000 | 79500000 | -0.63 | -0.80 | B>B |
| chr16 | 79500000 | 79600000 | -0.59 | -0.66 | B>B |
| chr16 | 79600000 | 79700000 | -0.42 | -0.31 | B>B |

| | | | | | |
|---|---|---|---|---|---|
| chr16 | 79700000 | 79800000 | -0.39 | -0.23 | B>B |
| chr16 | 79800000 | 79900000 | -0.29 | -0.25 | B>B |
| chr16 | 79900000 | 80000000 | -0.12 | -0.18 | B>B |
| chr16 | 80000000 | 80100000 | -0.07 | -0.05 | B>B |
| chr16 | 80100000 | 80200000 | -0.16 | -0.23 | B>B |
| chr16 | 80200000 | 80300000 | -0.09 | -0.26 | B>B |
| chr16 | 80300000 | 80400000 | -0.23 | -0.36 | B>B |
| chr16 | 80400000 | 80500000 | -0.48 | -0.59 | B>B |
| chr16 | 80500000 | 80600000 | -0.57 | -0.63 | B>B |
| chr16 | 80600000 | 80700000 | -0.76 | -1.02 | B>B |
| chr16 | 80700000 | 80800000 | -0.82 | -1.05 | B>B |
| chr16 | 80800000 | 80900000 | -0.58 | -0.71 | B>B |
| chr16 | 80900000 | 81000000 | -0.13 | -0.26 | B>B |
| chr16 | 81000000 | 81100000 | 0.54 | 0.40 | A>A |
| chr16 | 81100000 | 81200000 | 0.89 | 0.71 | A>A |
| chr16 | 81200000 | 81300000 | 0.83 | 0.67 | A>A |
| chr16 | 81300000 | 81400000 | 0.62 | 0.59 | A>A |
| chr16 | 81400000 | 81500000 | 0.59 | 0.69 | A>A |
| chr16 | 81500000 | 81600000 | 0.57 | 0.79 | A>A |
| chr16 | 81600000 | 81700000 | 0.41 | 0.63 | A>A |
| chr16 | 81700000 | 81800000 | 0.37 | 0.49 | A>A |
| chr16 | 81800000 | 81900000 | 0.26 | 0.29 | A>A |
| chr16 | 81900000 | 82000000 | -0.14 | -0.21 | B>B |
| chr16 | 82000000 | 82100000 | -0.34 | -0.47 | B>B |
| chr16 | 82100000 | 82200000 | -0.37 | -0.49 | B>B |
| chr16 | 82200000 | 82300000 | -0.55 | -0.66 | B>B |
| chr16 | 82300000 | 82400000 | -0.87 | -1.13 | B>B |
| chr16 | 82400000 | 82500000 | -0.81 | -1.26 | B>B |
| chr16 | 82500000 | 82600000 | -0.82 | -1.28 | B>B |
| chr16 | 82600000 | 82700000 | -0.69 | -1.17 | B>B |
| chr16 | 82700000 | 82800000 | -0.86 | -1.21 | B>B |
| chr16 | 82800000 | 82900000 | -1.03 | -1.32 | B>B |
| chr16 | 82900000 | 83000000 | -1.03 | -1.21 | B>B |
| chr16 | 83000000 | 83100000 | -1.01 | -1.28 | B>B |
| chr16 | 83100000 | 83200000 | -0.92 | -1.20 | B>B |
| chr16 | 83200000 | 83300000 | -0.87 | -1.21 | B>B |
| chr16 | 83300000 | 83400000 | -0.87 | -1.30 | B>B |
| chr16 | 83400000 | 83500000 | -0.93 | -1.41 | B>B |
| chr16 | 83500000 | 83600000 | -0.87 | -1.42 | B>B |
| chr16 | 83600000 | 83700000 | -0.84 | -1.41 | B>B |
| chr16 | 83700000 | 83800000 | -0.64 | -1.25 | A>B |
| chr16 | 83800000 | 83900000 | -0.13 | -0.58 | B>B |
| chr16 | 83900000 | 84000000 | 0.44 | 0.14 | A>A |
| chr16 | 84000000 | 84100000 | 0.72 | 0.76 | A>A |
| chr16 | 84100000 | 84200000 | 0.92 | 1.02 | A>A |
| chr16 | 84200000 | 84300000 | 0.94 | 1.03 | A>A |
| chr16 | 84300000 | 84400000 | 0.87 | 1.00 | A>A |
| chr16 | 84400000 | 84500000 | 0.93 | 1.01 | A>A |
| chr16 | 84500000 | 84600000 | 0.89 | 1.01 | A>A |
| chr16 | 84600000 | 84700000 | 1.08 | 1.14 | A>A |
| chr16 | 84700000 | 84800000 | 1.09 | 1.09 | A>A |
| chr16 | 84800000 | 84900000 | 0.99 | 1.13 | A>A |
| chr16 | 84900000 | 85000000 | 1.05 | 1.05 | A>A |
| chr16 | 85000000 | 85100000 | 1.09 | 1.06 | A>A |
| chr16 | 85100000 | 85200000 | 0.97 | 1.07 | A>A |
| chr16 | 85200000 | 85300000 | 1.03 | 1.00 | A>A |

| | | | | | |
|---|---|---|---|---|---|
| chr16 | 85300000 | 85400000 | 0.88 | 0.88 | A>A |
| chr16 | 85400000 | 85500000 | 0.82 | 0.86 | A>A |
| chr16 | 85500000 | 85600000 | 0.99 | 1.08 | A>A |
| chr16 | 85600000 | 85700000 | 0.89 | 0.87 | A>A |
| chr16 | 85700000 | 85800000 | 0.76 | 0.74 | A>A |
| chr16 | 85800000 | 85900000 | 0.60 | 0.40 | A>A |
| chr16 | 85900000 | 86000000 | 0.51 | 0.30 | A>A |
| chr16 | 86000000 | 86100000 | 0.32 | 0.18 | A>A |
| chr16 | 86100000 | 86200000 | 0.24 | 0.06 | A>A |
| chr16 | 86200000 | 86300000 | 0.19 | 0.05 | A>A |
| chr16 | 86300000 | 86400000 | 0.23 | 0.05 | A>A |
| chr16 | 86400000 | 86500000 | 0.23 | 0.04 | A>A |
| chr16 | 86500000 | 86600000 | 0.21 | 0.16 | A>A |
| chr16 | 86600000 | 86700000 | 0.41 | 0.44 | A>A |
| chr16 | 86700000 | 86800000 | 0.49 | 0.47 | A>A |
| chr16 | 86800000 | 86900000 | 0.42 | 0.37 | A>A |
| chr16 | 86900000 | 87000000 | 0.40 | 0.37 | A>A |
| chr16 | 87000000 | 87100000 | 0.31 | 0.36 | A>A |
| chr16 | 87100000 | 87200000 | 0.35 | 0.30 | A>A |
| chr16 | 87200000 | 87300000 | 0.48 | 0.41 | A>A |
| chr16 | 87300000 | 87400000 | 0.56 | 0.63 | A>A |
| chr16 | 87400000 | 87500000 | 0.67 | 0.72 | A>A |
| chr16 | 87500000 | 87600000 | 0.68 | 0.77 | A>A |
| chr16 | 87600000 | 87700000 | 0.64 | 0.77 | A>A |
| chr16 | 87700000 | 87800000 | 0.66 | 0.72 | A>A |
| chr16 | 87800000 | 87900000 | 0.88 | 0.90 | A>A |
| chr16 | 87900000 | 88000000 | 0.69 | 0.72 | A>A |
| chr16 | 88000000 | 88100000 | 0.65 | 0.66 | A>A |
| chr16 | 88100000 | 88200000 | 0.00 | 0.00 | NO |
| chr16 | 88200000 | 88300000 | 0.00 | 0.00 | NO |
| chr16 | 88300000 | 88400000 | 0.75 | 0.76 | A>A |
| chr16 | 88400000 | 88500000 | 0.63 | 0.71 | A>A |
| chr16 | 88500000 | 88600000 | 0.61 | 0.80 | A>A |
| chr16 | 88600000 | 88700000 | 0.82 | 0.85 | A>A |
| chr16 | 88700000 | 88800000 | 1.03 | 0.91 | A>A |
| chr16 | 88800000 | 88900000 | 0.84 | 0.90 | A>A |
| chr16 | 88900000 | 89000000 | 0.78 | 0.76 | A>A |
| chr16 | 89000000 | 89100000 | 0.78 | 0.70 | A>A |
| chr16 | 89100000 | 89200000 | 0.75 | 0.72 | A>A |
| chr16 | 89200000 | 89300000 | 0.58 | 0.64 | A>A |
| chr16 | 89300000 | 89400000 | 0.66 | 0.75 | A>A |
| chr16 | 89400000 | 89500000 | 0.77 | 0.87 | A>A |
| chr16 | 89500000 | 89600000 | 0.88 | 0.83 | A>A |
| chr16 | 89600000 | 89700000 | 0.88 | 0.86 | A>A |
| chr16 | 89700000 | 89800000 | 1.01 | 0.98 | A>A |
| chr16 | 89800000 | 89900000 | 1.06 | 1.02 | A>A |
| chr16 | 89900000 | 90000000 | 1.06 | 1.07 | A>A |
| chr16 | 90000000 | 90100000 | 1.03 | 1.03 | A>A |
| chr16 | 90100000 | 90200000 | 0.00 | 0.00 | NO |
| chr16 | 90200000 | 90300000 | 0.00 | 0.00 | NO |
| chr16 | 90300000 | 90354753 | 0.00 | 0.00 | NO |
| chr17 | 0 | 100000 | 0.34 | 0.32 | A>A |
| chr17 | 100000 | 200000 | 0.61 | 0.54 | A>A |
| chr17 | 200000 | 300000 | 0.38 | 0.38 | A>A |
| chr17 | 300000 | 400000 | 0.00 | 0.00 | NO |
| chr17 | 400000 | 500000 | 0.36 | 0.33 | A>A |

| | | | | | |
|---|---|---|---|---|---|
| chr17 | 500000 | 600000 | 0.31 | 0.38 | A>A |
| chr17 | 600000 | 700000 | 0.47 | 0.52 | A>A |
| chr17 | 700000 | 800000 | 0.60 | 0.60 | A>A |
| chr17 | 800000 | 900000 | 0.57 | 0.65 | A>A |
| chr17 | 900000 | 1000000 | 0.73 | 0.77 | A>A |
| chr17 | 1000000 | 1100000 | 0.85 | 0.80 | A>A |
| chr17 | 1100000 | 1200000 | 0.74 | 0.84 | A>A |
| chr17 | 1200000 | 1300000 | 0.89 | 0.91 | A>A |
| chr17 | 1300000 | 1400000 | 1.09 | 1.04 | A>A |
| chr17 | 1400000 | 1500000 | 0.96 | 0.98 | A>A |
| chr17 | 1500000 | 1600000 | 0.97 | 1.09 | A>A |
| chr17 | 1600000 | 1700000 | 1.05 | 1.09 | A>A |
| chr17 | 1700000 | 1800000 | 0.84 | 0.90 | A>A |
| chr17 | 1800000 | 1900000 | 0.88 | 0.88 | A>A |
| chr17 | 1900000 | 2000000 | 0.73 | 0.75 | A>A |
| chr17 | 2000000 | 2100000 | 0.79 | 0.78 | A>A |
| chr17 | 2100000 | 2200000 | 0.62 | 0.76 | A>A |
| chr17 | 2200000 | 2300000 | 0.88 | 0.97 | A>A |
| chr17 | 2300000 | 2400000 | 0.73 | 0.78 | A>A |
| chr17 | 2400000 | 2500000 | 0.64 | 0.74 | A>A |
| chr17 | 2500000 | 2600000 | 0.64 | 0.62 | A>A |
| chr17 | 2600000 | 2700000 | 0.47 | 0.39 | A>A |
| chr17 | 2700000 | 2800000 | 0.25 | 0.25 | A>A |
| chr17 | 2800000 | 2900000 | 0.37 | 0.29 | A>A |
| chr17 | 2900000 | 3000000 | -0.05 | -0.11 | B>B |
| chr17 | 3000000 | 3100000 | -0.32 | -0.52 | B>B |
| chr17 | 3100000 | 3200000 | 0.00 | -0.55 | NO |
| chr17 | 3200000 | 3300000 | -0.33 | -0.53 | B>B |
| chr17 | 3300000 | 3400000 | -0.16 | -0.45 | B>B |
| chr17 | 3400000 | 3500000 | 0.09 | -0.20 | B>B |
| chr17 | 3500000 | 3600000 | 0.40 | 0.27 | A>A |
| chr17 | 3600000 | 3700000 | 0.67 | 0.59 | A>A |
| chr17 | 3700000 | 3800000 | 0.71 | 0.75 | A>A |
| chr17 | 3800000 | 3900000 | 0.86 | 0.84 | A>A |
| chr17 | 3900000 | 4000000 | 0.74 | 0.74 | A>A |
| chr17 | 4000000 | 4100000 | 0.76 | 0.86 | A>A |
| chr17 | 4100000 | 4200000 | 0.80 | 0.75 | A>A |
| chr17 | 4200000 | 4300000 | 0.81 | 0.90 | A>A |
| chr17 | 4300000 | 4400000 | 0.73 | 0.74 | A>A |
| chr17 | 4400000 | 4500000 | 0.99 | 0.95 | A>A |
| chr17 | 4500000 | 4600000 | 0.92 | 0.92 | A>A |
| chr17 | 4600000 | 4700000 | 1.01 | 1.01 | A>A |
| chr17 | 4700000 | 4800000 | 0.99 | 1.02 | A>A |
| chr17 | 4800000 | 4900000 | 0.90 | 1.01 | A>A |
| chr17 | 4900000 | 5000000 | 0.68 | 0.93 | A>A |
| chr17 | 5000000 | 5100000 | 0.00 | 0.85 | NO |
| chr17 | 5100000 | 5200000 | 0.61 | 0.71 | A>A |
| chr17 | 5200000 | 5300000 | 0.73 | 0.70 | A>A |
| chr17 | 5300000 | 5400000 | 0.70 | 0.59 | A>A |
| chr17 | 5400000 | 5500000 | 0.23 | 0.03 | A>A |
| chr17 | 5500000 | 5600000 | 0.09 | -0.18 | B>B |
| chr17 | 5600000 | 5700000 | 0.01 | -0.13 | B>B |
| chr17 | 5700000 | 5800000 | 0.11 | -0.05 | A>A |
| chr17 | 5800000 | 5900000 | 0.06 | -0.14 | B>B |
| chr17 | 5900000 | 6000000 | 0.05 | -0.10 | B>B |
| chr17 | 6000000 | 6100000 | 0.12 | -0.18 | B>B |

| | | | | | |
|---|---|---|---|---|---|
| chr17 | 6100000 | 6200000 | 0.12 | -0.16 | B>B |
| chr17 | 6200000 | 6300000 | 0.16 | -0.05 | A>A |
| chr17 | 6300000 | 6400000 | 0.26 | 0.12 | A>A |
| chr17 | 6400000 | 6500000 | 0.62 | 0.43 | A>A |
| chr17 | 6500000 | 6600000 | 0.47 | 0.40 | A>A |
| chr17 | 6600000 | 6700000 | 0.41 | 0.29 | A>A |
| chr17 | 6700000 | 6800000 | 0.48 | 0.44 | A>A |
| chr17 | 6800000 | 6900000 | 0.49 | 0.43 | A>A |
| chr17 | 6900000 | 7000000 | 0.75 | 0.77 | A>A |
| chr17 | 7000000 | 7100000 | 0.88 | 0.92 | A>A |
| chr17 | 7100000 | 7200000 | 1.14 | 1.14 | A>A |
| chr17 | 7200000 | 7300000 | 1.19 | 1.15 | A>A |
| chr17 | 7300000 | 7400000 | 1.40 | 1.25 | A>A |
| chr17 | 7400000 | 7500000 | 1.19 | 1.19 | A>A |
| chr17 | 7500000 | 7600000 | 1.17 | 1.26 | A>A |
| chr17 | 7600000 | 7700000 | 1.12 | 1.09 | A>A |
| chr17 | 7700000 | 7800000 | 1.16 | 1.21 | A>A |
| chr17 | 7800000 | 7900000 | 1.25 | 1.28 | A>A |
| chr17 | 7900000 | 8000000 | 1.02 | 1.07 | A>A |
| chr17 | 8000000 | 8100000 | 1.34 | 1.32 | A>A |
| chr17 | 8100000 | 8200000 | 1.35 | 1.33 | A>A |
| chr17 | 8200000 | 8300000 | 0.96 | 1.02 | A>A |
| chr17 | 8300000 | 8400000 | 0.61 | 0.63 | A>A |
| chr17 | 8400000 | 8500000 | 0.55 | 0.48 | A>A |
| chr17 | 8500000 | 8600000 | 0.53 | 0.42 | A>A |
| chr17 | 8600000 | 8700000 | 0.34 | 0.43 | A>A |
| chr17 | 8700000 | 8800000 | 0.26 | 0.38 | A>A |
| chr17 | 8800000 | 8900000 | 0.09 | 0.14 | A>A |
| chr17 | 8900000 | 9000000 | 0.14 | 0.19 | A>A |
| chr17 | 9000000 | 9100000 | 0.14 | 0.10 | A>A |
| chr17 | 9100000 | 9200000 | 0.13 | 0.18 | A>A |
| chr17 | 9200000 | 9300000 | 0.05 | 0.17 | A>A |
| chr17 | 9300000 | 9400000 | 0.12 | 0.12 | A>A |
| chr17 | 9400000 | 9500000 | 0.06 | 0.11 | A>A |
| chr17 | 9500000 | 9600000 | 0.09 | 0.13 | A>A |
| chr17 | 9600000 | 9700000 | 0.07 | -0.03 | A>A |
| chr17 | 9700000 | 9800000 | 0.22 | -0.02 | A>A |
| chr17 | 9800000 | 9900000 | -0.16 | -0.40 | B>B |
| chr17 | 9900000 | 10000000 | -0.27 | -0.52 | B>B |
| chr17 | 10000000 | 10100000 | -0.31 | -0.52 | B>B |
| chr17 | 10100000 | 10200000 | -0.41 | -0.71 | B>B |
| chr17 | 10200000 | 10300000 | -0.61 | -1.03 | B>B |
| chr17 | 10300000 | 10400000 | -0.84 | -1.27 | B>B |
| chr17 | 10400000 | 10500000 | -0.78 | -1.15 | B>B |
| chr17 | 10500000 | 10600000 | -0.45 | -0.75 | B>B |
| chr17 | 10600000 | 10700000 | -0.50 | -0.83 | B>B |
| chr17 | 10700000 | 10800000 | -0.66 | -1.01 | B>B |
| chr17 | 10800000 | 10900000 | -0.86 | -1.06 | B>B |
| chr17 | 10900000 | 11000000 | -0.89 | -1.13 | B>B |
| chr17 | 11000000 | 11100000 | -0.82 | -1.06 | B>B |
| chr17 | 11100000 | 11200000 | -0.82 | -1.06 | B>B |
| chr17 | 11200000 | 11300000 | -0.80 | -1.03 | B>B |
| chr17 | 11300000 | 11400000 | -0.84 | -1.02 | B>B |
| chr17 | 11400000 | 11500000 | -0.85 | -1.04 | B>B |
| chr17 | 11500000 | 11600000 | -0.74 | -0.98 | B>B |
| chr17 | 11600000 | 11700000 | -0.75 | -1.04 | B>B |

| | | | | | |
|---|---|---|---|---|---|
| chr17 | 11700000 | 11800000 | -0.74 | -0.99 | B>B |
| chr17 | 11800000 | 11900000 | -0.63 | -0.86 | B>B |
| chr17 | 11900000 | 12000000 | -0.37 | -0.46 | B>B |
| chr17 | 12000000 | 12100000 | -0.49 | -0.73 | B>B |
| chr17 | 12100000 | 12200000 | -0.54 | -0.96 | B>B |
| chr17 | 12200000 | 12300000 | -0.55 | -1.02 | B>B |
| chr17 | 12300000 | 12400000 | -0.46 | -1.04 | B>B |
| chr17 | 12400000 | 12500000 | -0.37 | -1.00 | A>B |
| chr17 | 12500000 | 12600000 | -0.40 | -1.01 | A>B |
| chr17 | 12600000 | 12700000 | -0.57 | -1.01 | B>B |
| chr17 | 12700000 | 12800000 | -0.61 | -1.01 | B>B |
| chr17 | 12800000 | 12900000 | -0.62 | -0.97 | B>B |
| chr17 | 12900000 | 13000000 | -0.60 | -0.89 | B>B |
| chr17 | 13000000 | 13100000 | -0.76 | -1.00 | B>B |
| chr17 | 13100000 | 13200000 | -0.76 | -0.96 | B>B |
| chr17 | 13200000 | 13300000 | -0.75 | -0.72 | B>B |
| chr17 | 13300000 | 13400000 | -0.76 | -0.67 | B>B |
| chr17 | 13400000 | 13500000 | -0.70 | -0.70 | B>B |
| chr17 | 13500000 | 13600000 | -0.70 | -0.78 | B>B |
| chr17 | 13600000 | 13700000 | -0.78 | -0.96 | B>B |
| chr17 | 13700000 | 13800000 | -0.74 | -0.93 | B>B |
| chr17 | 13800000 | 13900000 | -0.70 | -0.90 | B>B |
| chr17 | 13900000 | 14000000 | -0.50 | -0.61 | B>B |
| chr17 | 14000000 | 14100000 | -0.41 | -0.58 | B>B |
| chr17 | 14100000 | 14200000 | -0.46 | -0.60 | B>B |
| chr17 | 14200000 | 14300000 | -0.38 | -0.51 | B>B |
| chr17 | 14300000 | 14400000 | -0.37 | -0.50 | B>B |
| chr17 | 14400000 | 14500000 | -0.37 | -0.47 | B>B |
| chr17 | 14500000 | 14600000 | -0.31 | -0.46 | B>B |
| chr17 | 14600000 | 14700000 | -0.34 | -0.49 | B>B |
| chr17 | 14700000 | 14800000 | -0.46 | -0.67 | B>B |
| chr17 | 14800000 | 14900000 | -0.65 | -0.85 | B>B |
| chr17 | 14900000 | 15000000 | -0.67 | -1.04 | B>B |
| chr17 | 15000000 | 15100000 | -0.67 | -0.93 | B>B |
| chr17 | 15100000 | 15200000 | -0.31 | -0.47 | B>B |
| chr17 | 15200000 | 15300000 | 0.01 | -0.19 | B>B |
| chr17 | 15300000 | 15400000 | -0.22 | -0.39 | B>B |
| chr17 | 15400000 | 15500000 | 0.00 | 0.00 | NO |
| chr17 | 15500000 | 15600000 | 0.00 | 0.00 | NO |
| chr17 | 15600000 | 15700000 | 0.00 | 0.00 | NO |
| chr17 | 15700000 | 15800000 | 0.00 | 0.00 | NO |
| chr17 | 15800000 | 15900000 | 0.51 | 0.55 | A>A |
| chr17 | 15900000 | 16000000 | 0.45 | 0.52 | A>A |
| chr17 | 16000000 | 16100000 | 0.61 | 0.69 | A>A |
| chr17 | 16100000 | 16200000 | 0.60 | 0.75 | A>A |
| chr17 | 16200000 | 16300000 | 0.68 | 0.80 | A>A |
| chr17 | 16300000 | 16400000 | 0.57 | 0.82 | A>A |
| chr17 | 16400000 | 16500000 | 0.38 | 0.68 | A>A |
| chr17 | 16500000 | 16600000 | 0.42 | 0.63 | A>A |
| chr17 | 16600000 | 16700000 | 0.00 | 0.00 | NO |
| chr17 | 16700000 | 16800000 | 0.00 | 0.00 | NO |
| chr17 | 16800000 | 16900000 | 0.38 | 0.58 | A>A |
| chr17 | 16900000 | 17000000 | 0.65 | 0.82 | A>A |
| chr17 | 17000000 | 17100000 | 0.55 | 0.61 | A>A |
| chr17 | 17100000 | 17200000 | 0.66 | 0.76 | A>A |
| chr17 | 17200000 | 17300000 | 0.52 | 0.60 | A>A |

| | | | | | |
|---|---|---|---|---|---|
| chr17 | 17300000 | 17400000 | 0.80 | 0.75 | A>A |
| chr17 | 17400000 | 17500000 | 0.68 | 0.66 | A>A |
| chr17 | 17500000 | 17600000 | 0.53 | 0.61 | A>A |
| chr17 | 17600000 | 17700000 | 0.75 | 0.83 | A>A |
| chr17 | 17700000 | 17800000 | 0.74 | 0.83 | A>A |
| chr17 | 17800000 | 17900000 | 0.97 | 0.90 | A>A |
| chr17 | 17900000 | 18000000 | 0.92 | 0.89 | A>A |
| chr17 | 18000000 | 18100000 | 1.28 | 1.20 | A>A |
| chr17 | 18100000 | 18200000 | 1.48 | 1.42 | A>A |
| chr17 | 18200000 | 18300000 | 1.22 | 1.25 | A>A |
| chr17 | 18300000 | 18400000 | 0.00 | 0.00 | NO |
| chr17 | 18400000 | 18500000 | 0.00 | 0.00 | NO |
| chr17 | 18500000 | 18600000 | 0.00 | 0.00 | NO |
| chr17 | 18600000 | 18700000 | 0.00 | 0.00 | NO |
| chr17 | 18700000 | 18800000 | 0.00 | 0.00 | NO |
| chr17 | 18800000 | 18900000 | 1.09 | 1.12 | A>A |
| chr17 | 18900000 | 19000000 | 0.00 | 0.00 | NO |
| chr17 | 19000000 | 19100000 | 0.00 | 0.00 | NO |
| chr17 | 19100000 | 19200000 | 0.00 | 0.88 | NO |
| chr17 | 19200000 | 19300000 | 0.89 | 0.92 | A>A |
| chr17 | 19300000 | 19400000 | 0.77 | 0.74 | A>A |
| chr17 | 19400000 | 19500000 | 0.76 | 0.71 | A>A |
| chr17 | 19500000 | 19600000 | 0.83 | 0.80 | A>A |
| chr17 | 19600000 | 19700000 | 0.76 | 0.81 | A>A |
| chr17 | 19700000 | 19800000 | 0.64 | 0.64 | A>A |
| chr17 | 19800000 | 19900000 | 0.55 | 0.55 | A>A |
| chr17 | 19900000 | 20000000 | 0.52 | 0.43 | A>A |
| chr17 | 20000000 | 20100000 | 0.30 | 0.37 | A>A |
| chr17 | 20100000 | 20200000 | 0.16 | 0.22 | A>A |
| chr17 | 20200000 | 20300000 | 0.00 | 0.00 | NO |
| chr17 | 20300000 | 20400000 | 0.00 | 0.00 | NO |
| chr17 | 20400000 | 20500000 | 0.00 | 0.00 | NO |
| chr17 | 20500000 | 20600000 | 0.00 | 0.00 | NO |
| chr17 | 20600000 | 20700000 | 0.00 | 0.00 | NO |
| chr17 | 20700000 | 20800000 | 0.00 | 0.00 | NO |
| chr17 | 20800000 | 20900000 | 0.14 | -0.01 | A>A |
| chr17 | 20900000 | 21000000 | 0.14 | 0.13 | A>A |
| chr17 | 21000000 | 21100000 | -0.10 | -0.14 | B>B |
| chr17 | 21100000 | 21200000 | -0.16 | -0.22 | B>B |
| chr17 | 21200000 | 21300000 | -0.41 | -0.70 | B>B |
| chr17 | 21300000 | 21400000 | -0.46 | -0.82 | B>B |
| chr17 | 21400000 | 21500000 | 0.00 | -0.78 | NO |
| chr17 | 21500000 | 21600000 | 0.00 | 0.00 | NO |
| chr17 | 21600000 | 21700000 | 0.00 | 0.00 | NO |
| chr17 | 21700000 | 21800000 | 0.00 | -1.70 | NO |
| chr17 | 21800000 | 21900000 | 0.00 | 0.00 | NO |
| chr17 | 21900000 | 22000000 | 0.00 | 0.00 | NO |
| chr17 | 22000000 | 22100000 | 0.00 | 0.00 | NO |
| chr17 | 22100000 | 22200000 | 0.00 | 0.00 | NO |
| chr17 | 22200000 | 22300000 | 0.00 | 0.00 | NO |
| chr17 | 22300000 | 22400000 | 0.00 | 0.00 | NO |
| chr17 | 22400000 | 22500000 | 0.00 | 0.00 | NO |
| chr17 | 22500000 | 22600000 | 0.00 | 0.00 | NO |
| chr17 | 22600000 | 22700000 | 0.00 | 0.00 | NO |
| chr17 | 22700000 | 22800000 | 0.00 | 0.00 | NO |
| chr17 | 22800000 | 22900000 | 0.00 | 0.00 | NO |

| | | | | | |
|---|---|---|---|---|---|
| chr17 | 22900000 | 23000000 | 0.00 | 0.00 | NO |
| chr17 | 23000000 | 23100000 | 0.00 | 0.00 | NO |
| chr17 | 23100000 | 23200000 | 0.00 | 0.00 | NO |
| chr17 | 23200000 | 23300000 | 0.00 | 0.00 | NO |
| chr17 | 23300000 | 23400000 | 0.00 | 0.00 | NO |
| chr17 | 23400000 | 23500000 | 0.00 | 0.00 | NO |
| chr17 | 23500000 | 23600000 | 0.00 | 0.00 | NO |
| chr17 | 23600000 | 23700000 | 0.00 | 0.00 | NO |
| chr17 | 23700000 | 23800000 | 0.00 | 0.00 | NO |
| chr17 | 23800000 | 23900000 | 0.00 | 0.00 | NO |
| chr17 | 23900000 | 24000000 | 0.00 | 0.00 | NO |
| chr17 | 24000000 | 24100000 | 0.00 | 0.00 | NO |
| chr17 | 24100000 | 24200000 | 0.00 | 0.00 | NO |
| chr17 | 24200000 | 24300000 | 0.00 | 0.00 | NO |
| chr17 | 24300000 | 24400000 | 0.00 | 0.00 | NO |
| chr17 | 24400000 | 24500000 | 0.00 | 0.00 | NO |
| chr17 | 24500000 | 24600000 | 0.00 | 0.00 | NO |
| chr17 | 24600000 | 24700000 | 0.00 | 0.00 | NO |
| chr17 | 24700000 | 24800000 | 0.00 | 0.00 | NO |
| chr17 | 24800000 | 24900000 | 0.00 | 0.00 | NO |
| chr17 | 24900000 | 25000000 | 0.00 | 0.00 | NO |
| chr17 | 25000000 | 25100000 | 0.00 | 0.00 | NO |
| chr17 | 25100000 | 25200000 | 0.00 | 0.00 | NO |
| chr17 | 25200000 | 25300000 | 0.00 | 0.00 | NO |
| chr17 | 25300000 | 25400000 | 0.00 | 0.00 | NO |
| chr17 | 25400000 | 25500000 | 0.00 | -1.59 | NO |
| chr17 | 25500000 | 25600000 | -0.30 | -0.64 | B>B |
| chr17 | 25600000 | 25700000 | -0.13 | -0.30 | B>B |
| chr17 | 25700000 | 25800000 | -0.11 | -0.22 | B>B |
| chr17 | 25800000 | 25900000 | 0.01 | 0.00 | A>A |
| chr17 | 25900000 | 26000000 | -0.21 | -0.30 | B>B |
| chr17 | 26000000 | 26100000 | -0.19 | -0.47 | B>B |
| chr17 | 26100000 | 26200000 | -0.06 | -0.32 | B>B |
| chr17 | 26200000 | 26300000 | -0.02 | -0.20 | B>B |
| chr17 | 26300000 | 26400000 | 0.14 | -0.02 | A>A |
| chr17 | 26400000 | 26500000 | 0.26 | 0.17 | A>A |
| chr17 | 26500000 | 26600000 | 0.35 | 0.33 | A>A |
| chr17 | 26600000 | 26700000 | 0.78 | 0.67 | A>A |
| chr17 | 26700000 | 26800000 | 0.83 | 0.84 | A>A |
| chr17 | 26800000 | 26900000 | 1.18 | 1.07 | A>A |
| chr17 | 26900000 | 27000000 | 1.39 | 1.40 | A>A |
| chr17 | 27000000 | 27100000 | 1.40 | 1.43 | A>A |
| chr17 | 27100000 | 27200000 | 1.26 | 1.33 | A>A |
| chr17 | 27200000 | 27300000 | 1.04 | 1.21 | A>A |
| chr17 | 27300000 | 27400000 | 1.06 | 1.25 | A>A |
| chr17 | 27400000 | 27500000 | 1.06 | 1.20 | A>A |
| chr17 | 27500000 | 27600000 | 1.00 | 1.05 | A>A |
| chr17 | 27600000 | 27700000 | 0.86 | 0.89 | A>A |
| chr17 | 27700000 | 27800000 | 0.85 | 0.99 | A>A |
| chr17 | 27800000 | 27900000 | 0.90 | 0.95 | A>A |
| chr17 | 27900000 | 28000000 | 0.74 | 0.76 | A>A |
| chr17 | 28000000 | 28100000 | 0.69 | 0.69 | A>A |
| chr17 | 28100000 | 28200000 | 0.51 | 0.53 | A>A |
| chr17 | 28200000 | 28300000 | 0.53 | 0.61 | A>A |
| chr17 | 28300000 | 28400000 | 0.55 | 0.51 | A>A |
| chr17 | 28400000 | 28500000 | 0.63 | 0.65 | A>A |

| | | | | | |
|---|---|---|---|---|---|
| chr17 | 28500000 | 28600000 | 0.76 | 0.68 | A>A |
| chr17 | 28600000 | 28700000 | 0.73 | 0.70 | A>A |
| chr17 | 28700000 | 28800000 | 0.83 | 0.84 | A>A |
| chr17 | 28800000 | 28900000 | 0.75 | 0.79 | A>A |
| chr17 | 28900000 | 29000000 | 0.00 | 0.00 | NO |
| chr17 | 29000000 | 29100000 | 0.00 | 0.00 | NO |
| chr17 | 29100000 | 29200000 | 0.74 | 0.76 | A>A |
| chr17 | 29200000 | 29300000 | 0.74 | 0.77 | A>A |
| chr17 | 29300000 | 29400000 | 0.58 | 0.60 | A>A |
| chr17 | 29400000 | 29500000 | 0.72 | 0.66 | A>A |
| chr17 | 29500000 | 29600000 | 0.47 | 0.58 | A>A |
| chr17 | 29600000 | 29700000 | 0.49 | 0.55 | A>A |
| chr17 | 29700000 | 29800000 | 0.38 | 0.46 | A>A |
| chr17 | 29800000 | 29900000 | 0.43 | 0.53 | A>A |
| chr17 | 29900000 | 30000000 | 0.43 | 0.45 | A>A |
| chr17 | 30000000 | 30100000 | 0.47 | 0.49 | A>A |
| chr17 | 30100000 | 30200000 | 0.44 | 0.49 | A>A |
| chr17 | 30200000 | 30300000 | 0.67 | 0.66 | A>A |
| chr17 | 30300000 | 30400000 | 0.00 | 0.00 | NO |
| chr17 | 30400000 | 30500000 | 0.57 | 0.62 | A>A |
| chr17 | 30500000 | 30600000 | 0.51 | 0.52 | A>A |
| chr17 | 30600000 | 30700000 | 0.58 | 0.49 | A>A |
| chr17 | 30700000 | 30800000 | 0.28 | 0.17 | A>A |
| chr17 | 30800000 | 30900000 | -0.26 | -0.53 | B>B |
| chr17 | 30900000 | 31000000 | -0.53 | -0.66 | B>B |
| chr17 | 31000000 | 31100000 | -0.53 | -0.62 | B>B |
| chr17 | 31100000 | 31200000 | -0.45 | -0.61 | B>B |
| chr17 | 31200000 | 31300000 | -0.34 | -0.74 | B>B |
| chr17 | 31300000 | 31400000 | -0.48 | -1.07 | A>B |
| chr17 | 31400000 | 31500000 | -0.49 | -1.17 | A>B |
| chr17 | 31500000 | 31600000 | -0.49 | -1.09 | A>B |
| chr17 | 31600000 | 31700000 | -0.58 | -1.08 | B>B |
| chr17 | 31700000 | 31800000 | -0.71 | -1.20 | B>B |
| chr17 | 31800000 | 31900000 | -0.70 | -1.22 | B>B |
| chr17 | 31900000 | 32000000 | -0.70 | -1.21 | B>B |
| chr17 | 32000000 | 32100000 | -0.67 | -1.09 | B>B |
| chr17 | 32100000 | 32200000 | -0.69 | -1.28 | A>B |
| chr17 | 32200000 | 32300000 | -0.74 | -1.33 | A>B |
| chr17 | 32300000 | 32400000 | -0.77 | -1.32 | B>B |
| chr17 | 32400000 | 32500000 | -0.69 | -1.21 | B>B |
| chr17 | 32500000 | 32600000 | -0.43 | -0.89 | B>B |
| chr17 | 32600000 | 32700000 | -0.64 | -0.95 | B>B |
| chr17 | 32700000 | 32800000 | -0.52 | -0.88 | B>B |
| chr17 | 32800000 | 32900000 | -0.52 | -0.94 | B>B |
| chr17 | 32900000 | 33000000 | -0.44 | -0.93 | B>B |
| chr17 | 33000000 | 33100000 | -0.54 | -1.09 | B>B |
| chr17 | 33100000 | 33200000 | -0.51 | -0.97 | B>B |
| chr17 | 33200000 | 33300000 | -0.39 | -0.79 | B>B |
| chr17 | 33300000 | 33400000 | 0.09 | -0.25 | B>B |
| chr17 | 33400000 | 33500000 | 0.01 | -0.38 | B>B |
| chr17 | 33500000 | 33600000 | 0.03 | -0.44 | B>B |
| chr17 | 33600000 | 33700000 | -0.07 | -0.50 | B>B |
| chr17 | 33700000 | 33800000 | -0.09 | -0.45 | B>B |
| chr17 | 33800000 | 33900000 | -0.08 | -0.39 | B>B |
| chr17 | 33900000 | 34000000 | 0.36 | 0.07 | A>A |
| chr17 | 34000000 | 34100000 | 0.30 | 0.11 | A>A |

| | | | | | |
|---|---|---|---|---|---|
| chr17 | 34100000 | 34200000 | 0.42 | 0.15 | A>A |
| chr17 | 34200000 | 34300000 | 0.31 | -0.06 | A>A |
| chr17 | 34300000 | 34400000 | 0.08 | -0.31 | B>B |
| chr17 | 34400000 | 34500000 | 0.00 | -0.38 | NO |
| chr17 | 34500000 | 34600000 | 0.00 | 0.00 | NO |
| chr17 | 34600000 | 34700000 | 0.00 | 0.00 | NO |
| chr17 | 34700000 | 34800000 | 0.00 | 0.00 | NO |
| chr17 | 34800000 | 34900000 | 0.67 | 0.32 | A>A |
| chr17 | 34900000 | 35000000 | 0.57 | 0.55 | A>A |
| chr17 | 35000000 | 35100000 | 0.39 | 0.40 | A>A |
| chr17 | 35100000 | 35200000 | 0.39 | 0.39 | A>A |
| chr17 | 35200000 | 35300000 | 0.54 | 0.57 | A>A |
| chr17 | 35300000 | 35400000 | 0.45 | 0.47 | A>A |
| chr17 | 35400000 | 35500000 | 0.29 | 0.37 | A>A |
| chr17 | 35500000 | 35600000 | 0.52 | 0.53 | A>A |
| chr17 | 35600000 | 35700000 | 0.67 | 0.75 | A>A |
| chr17 | 35700000 | 35800000 | 0.75 | 0.85 | A>A |
| chr17 | 35800000 | 35900000 | 0.78 | 0.93 | A>A |
| chr17 | 35900000 | 36000000 | 0.79 | 0.87 | A>A |
| chr17 | 36000000 | 36100000 | 0.89 | 0.97 | A>A |
| chr17 | 36100000 | 36200000 | 0.83 | 0.87 | A>A |
| chr17 | 36200000 | 36300000 | 0.00 | 0.00 | NO |
| chr17 | 36300000 | 36400000 | 0.00 | 0.00 | NO |
| chr17 | 36400000 | 36500000 | 0.00 | 0.00 | NO |
| chr17 | 36500000 | 36600000 | 0.95 | 0.97 | A>A |
| chr17 | 36600000 | 36700000 | 0.00 | 0.99 | NO |
| chr17 | 36700000 | 36800000 | 1.03 | 1.04 | A>A |
| chr17 | 36800000 | 36900000 | 1.11 | 1.03 | A>A |
| chr17 | 36900000 | 37000000 | 1.04 | 1.04 | A>A |
| chr17 | 37000000 | 37100000 | 0.82 | 0.84 | A>A |
| chr17 | 37100000 | 37200000 | 0.65 | 0.69 | A>A |
| chr17 | 37200000 | 37300000 | 0.77 | 0.77 | A>A |
| chr17 | 37300000 | 37400000 | 0.67 | 0.71 | A>A |
| chr17 | 37400000 | 37500000 | 0.67 | 0.67 | A>A |
| chr17 | 37500000 | 37600000 | 0.68 | 0.67 | A>A |
| chr17 | 37600000 | 37700000 | 0.64 | 0.70 | A>A |
| chr17 | 37700000 | 37800000 | 0.68 | 0.66 | A>A |
| chr17 | 37800000 | 37900000 | 0.90 | 0.84 | A>A |
| chr17 | 37900000 | 38000000 | 0.65 | 0.64 | A>A |
| chr17 | 38000000 | 38100000 | 0.74 | 0.70 | A>A |
| chr17 | 38100000 | 38200000 | 0.70 | 0.68 | A>A |
| chr17 | 38200000 | 38300000 | 0.90 | 0.83 | A>A |
| chr17 | 38300000 | 38400000 | 0.74 | 0.64 | A>A |
| chr17 | 38400000 | 38500000 | 0.67 | 0.66 | A>A |
| chr17 | 38500000 | 38600000 | 0.60 | 0.56 | A>A |
| chr17 | 38600000 | 38700000 | 0.66 | 0.64 | A>A |
| chr17 | 38700000 | 38800000 | 0.29 | 0.31 | A>A |
| chr17 | 38800000 | 38900000 | 0.48 | 0.37 | A>A |
| chr17 | 38900000 | 39000000 | 0.25 | 0.26 | A>A |
| chr17 | 39000000 | 39100000 | 0.21 | 0.26 | A>A |
| chr17 | 39100000 | 39200000 | 0.24 | 0.30 | A>A |
| chr17 | 39200000 | 39300000 | 0.31 | 0.25 | A>A |
| chr17 | 39300000 | 39400000 | 0.28 | 0.25 | A>A |
| chr17 | 39400000 | 39500000 | 0.23 | 0.20 | A>A |
| chr17 | 39500000 | 39600000 | 0.30 | 0.27 | A>A |
| chr17 | 39600000 | 39700000 | 0.32 | 0.30 | A>A |

| | | | | | |
|---|---|---|---|---|---|
| chr17 | 39700000 | 39800000 | 0.32 | 0.27 | A>A |
| chr17 | 39800000 | 39900000 | 0.42 | 0.46 | A>A |
| chr17 | 39900000 | 40000000 | 0.82 | 0.75 | A>A |
| chr17 | 40000000 | 40100000 | 0.79 | 0.72 | A>A |
| chr17 | 40100000 | 40200000 | 0.70 | 0.68 | A>A |
| chr17 | 40200000 | 40300000 | 0.72 | 0.72 | A>A |
| chr17 | 40300000 | 40400000 | 0.69 | 0.65 | A>A |
| chr17 | 40400000 | 40500000 | 0.71 | 0.70 | A>A |
| chr17 | 40500000 | 40600000 | 0.71 | 0.73 | A>A |
| chr17 | 40600000 | 40700000 | 0.73 | 0.69 | A>A |
| chr17 | 40700000 | 40800000 | 0.79 | 0.83 | A>A |
| chr17 | 40800000 | 40900000 | 0.89 | 0.85 | A>A |
| chr17 | 40900000 | 41000000 | 0.81 | 0.80 | A>A |
| chr17 | 41000000 | 41100000 | 0.81 | 0.78 | A>A |
| chr17 | 41100000 | 41200000 | 0.72 | 0.73 | A>A |
| chr17 | 41200000 | 41300000 | 0.71 | 0.74 | A>A |
| chr17 | 41300000 | 41400000 | 0.62 | 0.74 | A>A |
| chr17 | 41400000 | 41500000 | 0.46 | 0.64 | A>A |
| chr17 | 41500000 | 41600000 | 0.61 | 0.68 | A>A |
| chr17 | 41600000 | 41700000 | 0.66 | 0.67 | A>A |
| chr17 | 41700000 | 41800000 | 0.76 | 0.73 | A>A |
| chr17 | 41800000 | 41900000 | 0.80 | 0.83 | A>A |
| chr17 | 41900000 | 42000000 | 0.93 | 0.83 | A>A |
| chr17 | 42000000 | 42100000 | 0.84 | 0.82 | A>A |
| chr17 | 42100000 | 42200000 | 0.96 | 0.85 | A>A |
| chr17 | 42200000 | 42300000 | 0.84 | 0.88 | A>A |
| chr17 | 42300000 | 42400000 | 1.00 | 0.90 | A>A |
| chr17 | 42400000 | 42500000 | 0.73 | 0.71 | A>A |
| chr17 | 42500000 | 42600000 | 0.66 | 0.70 | A>A |
| chr17 | 42600000 | 42700000 | 0.86 | 0.72 | A>A |
| chr17 | 42700000 | 42800000 | 0.71 | 0.69 | A>A |
| chr17 | 42800000 | 42900000 | 0.86 | 0.78 | A>A |
| chr17 | 42900000 | 43000000 | 0.82 | 0.74 | A>A |
| chr17 | 43000000 | 43100000 | 1.02 | 0.86 | A>A |
| chr17 | 43100000 | 43200000 | 0.92 | 0.86 | A>A |
| chr17 | 43200000 | 43300000 | 0.96 | 0.88 | A>A |
| chr17 | 43300000 | 43400000 | 0.78 | 0.81 | A>A |
| chr17 | 43400000 | 43500000 | 0.00 | 0.00 | NO |
| chr17 | 43500000 | 43600000 | 0.00 | 0.00 | NO |
| chr17 | 43600000 | 43700000 | 0.00 | 0.00 | NO |
| chr17 | 43700000 | 43800000 | 0.00 | 0.00 | NO |
| chr17 | 43800000 | 43900000 | 0.00 | 0.00 | NO |
| chr17 | 43900000 | 44000000 | 0.00 | 0.00 | NO |
| chr17 | 44000000 | 44100000 | 0.00 | 0.00 | NO |
| chr17 | 44100000 | 44200000 | 0.00 | 0.00 | NO |
| chr17 | 44200000 | 44300000 | 0.00 | 0.00 | NO |
| chr17 | 44300000 | 44400000 | 0.00 | 0.00 | NO |
| chr17 | 44400000 | 44500000 | 0.00 | 0.00 | NO |
| chr17 | 44500000 | 44600000 | 0.00 | 0.00 | NO |
| chr17 | 44600000 | 44700000 | 0.00 | 0.00 | NO |
| chr17 | 44700000 | 44800000 | 0.00 | 0.00 | NO |
| chr17 | 44800000 | 44900000 | 0.54 | 0.51 | A>A |
| chr17 | 44900000 | 45000000 | 0.63 | 0.53 | A>A |
| chr17 | 45000000 | 45100000 | 0.41 | 0.49 | A>A |
| chr17 | 45100000 | 45200000 | 0.00 | 0.00 | NO |
| chr17 | 45200000 | 45300000 | 0.49 | 0.50 | A>A |

| | | | | | |
|---|---|---|---|---|---|
| chr17 | 45300000 | 45400000 | 0.61 | 0.62 | A>A |
| chr17 | 45400000 | 45500000 | 0.53 | 0.54 | A>A |
| chr17 | 45500000 | 45600000 | 0.00 | 0.00 | NO |
| chr17 | 45600000 | 45700000 | 0.00 | 0.00 | NO |
| chr17 | 45700000 | 45800000 | 0.93 | 0.84 | A>A |
| chr17 | 45800000 | 45900000 | 0.99 | 0.88 | A>A |
| chr17 | 45900000 | 46000000 | 0.98 | 0.92 | A>A |
| chr17 | 46000000 | 46100000 | 0.99 | 0.91 | A>A |
| chr17 | 46100000 | 46200000 | 0.79 | 0.79 | A>A |
| chr17 | 46200000 | 46300000 | 0.61 | 0.60 | A>A |
| chr17 | 46300000 | 46400000 | 0.75 | 0.77 | A>A |
| chr17 | 46400000 | 46500000 | 0.61 | 0.55 | A>A |
| chr17 | 46500000 | 46600000 | 0.77 | 0.69 | A>A |
| chr17 | 46600000 | 46700000 | 0.77 | 0.67 | A>A |
| chr17 | 46700000 | 46800000 | 0.57 | 0.65 | A>A |
| chr17 | 46800000 | 46900000 | 0.86 | 0.83 | A>A |
| chr17 | 46900000 | 47000000 | 0.80 | 0.77 | A>A |
| chr17 | 47000000 | 47100000 | 1.00 | 0.91 | A>A |
| chr17 | 47100000 | 47200000 | 0.80 | 0.76 | A>A |
| chr17 | 47200000 | 47300000 | 0.79 | 0.74 | A>A |
| chr17 | 47300000 | 47400000 | 0.76 | 0.69 | A>A |
| chr17 | 47400000 | 47500000 | 0.80 | 0.75 | A>A |
| chr17 | 47500000 | 47600000 | 0.91 | 0.78 | A>A |
| chr17 | 47600000 | 47700000 | 0.72 | 0.80 | A>A |
| chr17 | 47700000 | 47800000 | 0.88 | 0.83 | A>A |
| chr17 | 47800000 | 47900000 | 1.01 | 0.93 | A>A |
| chr17 | 47900000 | 48000000 | 0.94 | 0.92 | A>A |
| chr17 | 48000000 | 48100000 | 1.01 | 0.97 | A>A |
| chr17 | 48100000 | 48200000 | 1.16 | 1.18 | A>A |
| chr17 | 48200000 | 48300000 | 1.14 | 1.09 | A>A |
| chr17 | 48300000 | 48400000 | 1.06 | 1.10 | A>A |
| chr17 | 48400000 | 48500000 | 1.05 | 1.07 | A>A |
| chr17 | 48500000 | 48600000 | 0.90 | 0.94 | A>A |
| chr17 | 48600000 | 48700000 | 0.92 | 0.94 | A>A |
| chr17 | 48700000 | 48800000 | 0.82 | 0.96 | A>A |
| chr17 | 48800000 | 48900000 | 0.58 | 0.61 | A>A |
| chr17 | 48900000 | 49000000 | 0.70 | 0.82 | A>A |
| chr17 | 49000000 | 49100000 | 0.51 | 0.58 | A>A |
| chr17 | 49100000 | 49200000 | 0.39 | 0.48 | A>A |
| chr17 | 49200000 | 49300000 | 0.17 | 0.22 | A>A |
| chr17 | 49300000 | 49400000 | -0.26 | -0.36 | B>B |
| chr17 | 49400000 | 49500000 | -0.74 | -0.86 | B>B |
| chr17 | 49500000 | 49600000 | -0.73 | -0.92 | B>B |
| chr17 | 49600000 | 49700000 | -0.84 | -1.07 | B>B |
| chr17 | 49700000 | 49800000 | -0.89 | -1.20 | B>B |
| chr17 | 49800000 | 49900000 | -0.98 | -1.13 | B>B |
| chr17 | 49900000 | 50000000 | -1.07 | -1.24 | B>B |
| chr17 | 50000000 | 50100000 | -1.09 | -1.16 | B>B |
| chr17 | 50100000 | 50200000 | -1.11 | -1.14 | B>B |
| chr17 | 50200000 | 50300000 | -1.06 | -1.11 | B>B |
| chr17 | 50300000 | 50400000 | -1.05 | -1.10 | B>B |
| chr17 | 50400000 | 50500000 | -1.05 | -1.07 | B>B |
| chr17 | 50500000 | 50600000 | -1.00 | -1.09 | B>B |
| chr17 | 50600000 | 50700000 | -0.98 | -1.09 | B>B |
| chr17 | 50700000 | 50800000 | -0.95 | -1.06 | B>B |
| chr17 | 50800000 | 50900000 | -0.94 | -1.02 | B>B |

| | | | | | |
|---|---|---|---|---|---|
| chr17 | 50900000 | 51000000 | -0.96 | -1.02 | B>B |
| chr17 | 51000000 | 51100000 | -0.91 | -0.97 | B>B |
| chr17 | 51100000 | 51200000 | -0.90 | -0.94 | B>B |
| chr17 | 51200000 | 51300000 | -0.91 | -0.86 | B>B |
| chr17 | 51300000 | 51400000 | -0.92 | -0.87 | B>B |
| chr17 | 51400000 | 51500000 | -0.89 | -0.80 | B>B |
| chr17 | 51500000 | 51600000 | -0.92 | -0.72 | B>B |
| chr17 | 51600000 | 51700000 | -0.89 | -0.68 | B>B |
| chr17 | 51700000 | 51800000 | -0.88 | -0.63 | B>B |
| chr17 | 51800000 | 51900000 | -0.90 | -0.65 | B>B |
| chr17 | 51900000 | 52000000 | -0.91 | -0.74 | B>B |
| chr17 | 52000000 | 52100000 | -0.91 | -0.78 | B>B |
| chr17 | 52100000 | 52200000 | -0.94 | -0.71 | B>B |
| chr17 | 52200000 | 52300000 | -0.94 | -0.66 | B>B |
| chr17 | 52300000 | 52400000 | -0.98 | -0.67 | B>B |
| chr17 | 52400000 | 52500000 | -1.05 | -0.71 | B>B |
| chr17 | 52500000 | 52600000 | -1.04 | -0.96 | B>B |
| chr17 | 52600000 | 52700000 | -1.04 | -1.13 | B>B |
| chr17 | 52700000 | 52800000 | -1.08 | -1.19 | B>B |
| chr17 | 52800000 | 52900000 | -1.08 | -1.16 | B>B |
| chr17 | 52900000 | 53000000 | -0.99 | -0.97 | B>B |
| chr17 | 53000000 | 53100000 | -0.62 | -0.59 | B>B |
| chr17 | 53100000 | 53200000 | -0.59 | -0.68 | B>B |
| chr17 | 53200000 | 53300000 | -0.55 | -0.79 | B>B |
| chr17 | 53300000 | 53400000 | -0.46 | -0.71 | B>B |
| chr17 | 53400000 | 53500000 | -0.37 | -0.51 | B>B |
| chr17 | 53500000 | 53600000 | -0.32 | -0.42 | B>B |
| chr17 | 53600000 | 53700000 | -0.65 | -0.66 | B>B |
| chr17 | 53700000 | 53800000 | -0.73 | -0.89 | B>B |
| chr17 | 53800000 | 53900000 | -0.79 | -0.84 | B>B |
| chr17 | 53900000 | 54000000 | -0.94 | -1.27 | B>B |
| chr17 | 54000000 | 54100000 | -1.08 | -1.34 | B>B |
| chr17 | 54100000 | 54200000 | -1.08 | -1.30 | B>B |
| chr17 | 54200000 | 54300000 | -1.04 | -1.38 | B>B |
| chr17 | 54300000 | 54400000 | -1.06 | -1.41 | B>B |
| chr17 | 54400000 | 54500000 | -1.07 | -1.40 | B>B |
| chr17 | 54500000 | 54600000 | -1.01 | -1.39 | B>B |
| chr17 | 54600000 | 54700000 | -0.83 | -1.18 | B>B |
| chr17 | 54700000 | 54800000 | -0.73 | -1.03 | B>B |
| chr17 | 54800000 | 54900000 | -0.60 | -0.79 | B>B |
| chr17 | 54900000 | 55000000 | -0.21 | -0.15 | B>B |
| chr17 | 55000000 | 55100000 | -0.06 | -0.05 | B>B |
| chr17 | 55100000 | 55200000 | 0.04 | 0.11 | A>A |
| chr17 | 55200000 | 55300000 | -0.07 | 0.00 | B>B |
| chr17 | 55300000 | 55400000 | 0.04 | 0.11 | A>A |
| chr17 | 55400000 | 55500000 | 0.10 | 0.18 | A>A |
| chr17 | 55500000 | 55600000 | 0.02 | 0.09 | A>A |
| chr17 | 55600000 | 55700000 | -0.03 | 0.05 | A>A |
| chr17 | 55700000 | 55800000 | -0.04 | 0.01 | B>B |
| chr17 | 55800000 | 55900000 | 0.00 | 0.05 | A>A |
| chr17 | 55900000 | 56000000 | 0.43 | 0.49 | A>A |
| chr17 | 56000000 | 56100000 | 0.55 | 0.56 | A>A |
| chr17 | 56100000 | 56200000 | 0.36 | 0.26 | A>A |
| chr17 | 56200000 | 56300000 | 0.32 | 0.19 | A>A |
| chr17 | 56300000 | 56400000 | 0.40 | 0.26 | A>A |
| chr17 | 56400000 | 56500000 | 0.65 | 0.45 | A>A |

| | | | | | |
|---|---|---|---|---|---|
| chr17 | 56500000 | 56600000 | 0.37 | 0.26 | A>A |
| chr17 | 56600000 | 56700000 | 0.44 | 0.34 | A>A |
| chr17 | 56700000 | 56800000 | 0.55 | 0.36 | A>A |
| chr17 | 56800000 | 56900000 | 0.23 | 0.12 | A>A |
| chr17 | 56900000 | 57000000 | 0.19 | 0.11 | A>A |
| chr17 | 57000000 | 57100000 | 0.19 | 0.13 | A>A |
| chr17 | 57100000 | 57200000 | 0.59 | 0.49 | A>A |
| chr17 | 57200000 | 57300000 | 0.59 | 0.67 | A>A |
| chr17 | 57300000 | 57400000 | 0.50 | 0.53 | A>A |
| chr17 | 57400000 | 57500000 | 0.65 | 0.68 | A>A |
| chr17 | 57500000 | 57600000 | 0.50 | 0.52 | A>A |
| chr17 | 57600000 | 57700000 | 0.67 | 0.69 | A>A |
| chr17 | 57700000 | 57800000 | 0.71 | 0.83 | A>A |
| chr17 | 57800000 | 57900000 | 0.97 | 1.13 | A>A |
| chr17 | 57900000 | 58000000 | 0.87 | 0.91 | A>A |
| chr17 | 58000000 | 58100000 | 0.00 | 0.00 | NO |
| chr17 | 58100000 | 58200000 | 0.49 | 0.61 | A>A |
| chr17 | 58200000 | 58300000 | 0.43 | 0.48 | A>A |
| chr17 | 58300000 | 58400000 | 0.44 | 0.43 | A>A |
| chr17 | 58400000 | 58500000 | 0.48 | 0.47 | A>A |
| chr17 | 58500000 | 58600000 | 0.33 | 0.38 | A>A |
| chr17 | 58600000 | 58700000 | 0.44 | 0.48 | A>A |
| chr17 | 58700000 | 58800000 | 0.09 | 0.25 | A>A |
| chr17 | 58800000 | 58900000 | -0.06 | 0.07 | A>A |
| chr17 | 58900000 | 59000000 | 0.02 | 0.09 | A>A |
| chr17 | 59000000 | 59100000 | -0.12 | 0.01 | B>B |
| chr17 | 59100000 | 59200000 | -0.15 | -0.02 | B>B |
| chr17 | 59200000 | 59300000 | -0.11 | 0.05 | B>B |
| chr17 | 59300000 | 59400000 | -0.06 | 0.00 | B>B |
| chr17 | 59400000 | 59500000 | -0.11 | 0.02 | B>B |
| chr17 | 59500000 | 59600000 | 0.19 | 0.21 | A>A |
| chr17 | 59600000 | 59700000 | 0.00 | 0.12 | A>A |
| chr17 | 59700000 | 59800000 | -0.08 | 0.08 | A>A |
| chr17 | 59800000 | 59900000 | 0.05 | 0.12 | A>A |
| chr17 | 59900000 | 60000000 | 0.18 | 0.21 | A>A |
| chr17 | 60000000 | 60100000 | 0.47 | 0.47 | A>A |
| chr17 | 60100000 | 60200000 | 0.41 | 0.38 | A>A |
| chr17 | 60200000 | 60300000 | 0.36 | 0.32 | A>A |
| chr17 | 60300000 | 60400000 | 0.00 | 0.00 | NO |
| chr17 | 60400000 | 60500000 | 0.13 | 0.03 | A>A |
| chr17 | 60500000 | 60600000 | 0.00 | 0.00 | NO |
| chr17 | 60600000 | 60700000 | 0.43 | 0.42 | A>A |
| chr17 | 60700000 | 60800000 | 0.35 | 0.35 | A>A |
| chr17 | 60800000 | 60900000 | 0.39 | 0.40 | A>A |
| chr17 | 60900000 | 61000000 | 0.40 | 0.45 | A>A |
| chr17 | 61000000 | 61100000 | 0.20 | 0.28 | A>A |
| chr17 | 61100000 | 61200000 | 0.16 | 0.22 | A>A |
| chr17 | 61200000 | 61300000 | 0.13 | 0.19 | A>A |
| chr17 | 61300000 | 61400000 | 0.14 | 0.22 | A>A |
| chr17 | 61400000 | 61500000 | 0.10 | 0.22 | A>A |
| chr17 | 61500000 | 61600000 | 0.57 | 0.55 | A>A |
| chr17 | 61600000 | 61700000 | 0.65 | 0.61 | A>A |
| chr17 | 61700000 | 61800000 | 0.79 | 0.82 | A>A |
| chr17 | 61800000 | 61900000 | 1.16 | 1.09 | A>A |
| chr17 | 61900000 | 62000000 | 0.00 | 1.03 | NO |
| chr17 | 62000000 | 62100000 | 0.94 | 0.88 | A>A |

| | | | | | |
|---|---|---|---|---|---|
| chr17 | 62100000 | 62200000 | 1.11 | 0.89 | A>A |
| chr17 | 62200000 | 62300000 | 0.97 | 0.87 | A>A |
| chr17 | 62300000 | 62400000 | 0.72 | 0.68 | A>A |
| chr17 | 62400000 | 62500000 | 0.00 | 0.00 | NO |
| chr17 | 62500000 | 62600000 | 0.86 | 0.84 | A>A |
| chr17 | 62600000 | 62700000 | 0.85 | 0.84 | A>A |
| chr17 | 62700000 | 62800000 | 0.00 | 0.75 | NO |
| chr17 | 62800000 | 62900000 | 0.00 | 0.00 | NO |
| chr17 | 62900000 | 63000000 | 0.00 | 0.00 | NO |
| chr17 | 63000000 | 63100000 | 0.63 | 0.56 | A>A |
| chr17 | 63100000 | 63200000 | 0.42 | 0.33 | A>A |
| chr17 | 63200000 | 63300000 | -0.13 | -0.10 | B>B |
| chr17 | 63300000 | 63400000 | -0.22 | -0.17 | B>B |
| chr17 | 63400000 | 63500000 | -0.11 | -0.12 | B>B |
| chr17 | 63500000 | 63600000 | -0.12 | 0.00 | B>B |
| chr17 | 63600000 | 63700000 | -0.06 | 0.01 | B>B |
| chr17 | 63700000 | 63800000 | -0.07 | 0.02 | B>B |
| chr17 | 63800000 | 63900000 | -0.13 | -0.03 | B>B |
| chr17 | 63900000 | 64000000 | -0.18 | -0.11 | B>B |
| chr17 | 64000000 | 64100000 | -0.16 | -0.06 | B>B |
| chr17 | 64100000 | 64200000 | -0.19 | -0.04 | B>B |
| chr17 | 64200000 | 64300000 | 0.06 | 0.14 | A>A |
| chr17 | 64300000 | 64400000 | 0.23 | 0.28 | A>A |
| chr17 | 64400000 | 64500000 | 0.18 | 0.31 | A>A |
| chr17 | 64500000 | 64600000 | 0.15 | 0.24 | A>A |
| chr17 | 64600000 | 64700000 | -0.05 | 0.07 | A>A |
| chr17 | 64700000 | 64800000 | -0.06 | 0.07 | A>A |
| chr17 | 64800000 | 64900000 | 0.02 | 0.05 | A>A |
| chr17 | 64900000 | 65000000 | 0.08 | 0.17 | A>A |
| chr17 | 65000000 | 65100000 | 0.19 | 0.29 | A>A |
| chr17 | 65100000 | 65200000 | 0.26 | 0.32 | A>A |
| chr17 | 65200000 | 65300000 | 0.33 | 0.38 | A>A |
| chr17 | 65300000 | 65400000 | 0.28 | 0.35 | A>A |
| chr17 | 65400000 | 65500000 | 0.29 | 0.28 | A>A |
| chr17 | 65500000 | 65600000 | 0.27 | 0.22 | A>A |
| chr17 | 65600000 | 65700000 | 0.21 | 0.18 | A>A |
| chr17 | 65700000 | 65800000 | 0.45 | 0.41 | A>A |
| chr17 | 65800000 | 65900000 | 0.43 | 0.45 | A>A |
| chr17 | 65900000 | 66000000 | 0.34 | 0.36 | A>A |
| chr17 | 66000000 | 66100000 | 0.39 | 0.31 | A>A |
| chr17 | 66100000 | 66200000 | 0.05 | 0.02 | A>A |
| chr17 | 66200000 | 66300000 | 0.02 | -0.13 | B>B |
| chr17 | 66300000 | 66400000 | 0.12 | -0.11 | A>A |
| chr17 | 66400000 | 66500000 | -0.15 | -0.24 | B>B |
| chr17 | 66500000 | 66600000 | -0.43 | -0.77 | B>B |
| chr17 | 66600000 | 66700000 | -0.67 | -1.14 | B>B |
| chr17 | 66700000 | 66800000 | -0.63 | -0.85 | B>B |
| chr17 | 66800000 | 66900000 | -0.85 | -0.92 | B>B |
| chr17 | 66900000 | 67000000 | -0.85 | -0.94 | B>B |
| chr17 | 67000000 | 67100000 | -0.89 | -1.13 | B>B |
| chr17 | 67100000 | 67200000 | -0.76 | -0.92 | B>B |
| chr17 | 67200000 | 67300000 | -0.64 | -0.68 | B>B |
| chr17 | 67300000 | 67400000 | -0.54 | -0.48 | B>B |
| chr17 | 67400000 | 67500000 | -0.46 | -0.31 | B>B |
| chr17 | 67500000 | 67600000 | -0.63 | -0.58 | B>B |
| chr17 | 67600000 | 67700000 | -0.49 | -0.74 | B>B |

| | | | | | |
|---|---|---|---|---|---|
| chr17 | 67700000 | 67800000 | -0.45 | -0.70 | B>B |
| chr17 | 67800000 | 67900000 | -0.45 | -0.78 | B>B |
| chr17 | 67900000 | 68000000 | -0.50 | -0.87 | B>B |
| chr17 | 68000000 | 68100000 | -0.53 | -0.84 | B>B |
| chr17 | 68100000 | 68200000 | -0.62 | -0.75 | B>B |
| chr17 | 68200000 | 68300000 | -0.87 | -0.91 | B>B |
| chr17 | 68300000 | 68400000 | -0.94 | -1.02 | B>B |
| chr17 | 68400000 | 68500000 | -0.94 | -0.99 | B>B |
| chr17 | 68500000 | 68600000 | -0.94 | -0.83 | B>B |
| chr17 | 68600000 | 68700000 | -0.91 | -0.62 | B>B |
| chr17 | 68700000 | 68800000 | -0.98 | -0.66 | B>B |
| chr17 | 68800000 | 68900000 | -1.08 | -0.75 | B>B |
| chr17 | 68900000 | 69000000 | -1.05 | -0.82 | B>B |
| chr17 | 69000000 | 69100000 | -1.00 | -0.73 | B>B |
| chr17 | 69100000 | 69200000 | -1.00 | -0.65 | B>B |
| chr17 | 69200000 | 69300000 | -0.92 | -0.71 | B>B |
| chr17 | 69300000 | 69400000 | -0.87 | -0.69 | B>B |
| chr17 | 69400000 | 69500000 | -0.85 | -0.60 | B>B |
| chr17 | 69500000 | 69600000 | -0.95 | -0.77 | B>B |
| chr17 | 69600000 | 69700000 | -1.02 | -0.87 | B>B |
| chr17 | 69700000 | 69800000 | -0.89 | -0.78 | B>B |
| chr17 | 69800000 | 69900000 | -0.87 | -0.71 | B>B |
| chr17 | 69900000 | 70000000 | -0.72 | -0.63 | B>B |
| chr17 | 70000000 | 70100000 | -0.70 | -0.56 | B>B |
| chr17 | 70100000 | 70200000 | -0.49 | -0.26 | B>B |
| chr17 | 70200000 | 70300000 | -0.47 | -0.25 | B>B |
| chr17 | 70300000 | 70400000 | -0.46 | -0.23 | B>B |
| chr17 | 70400000 | 70500000 | -0.43 | -0.21 | B>B |
| chr17 | 70500000 | 70600000 | -0.40 | -0.14 | B>B |
| chr17 | 70600000 | 70700000 | -0.06 | 0.08 | A>A |
| chr17 | 70700000 | 70800000 | -0.26 | -0.01 | B>B |
| chr17 | 70800000 | 70900000 | -0.24 | 0.01 | B>B |
| chr17 | 70900000 | 71000000 | -0.19 | 0.10 | B>B |
| chr17 | 71000000 | 71100000 | -0.10 | 0.14 | A>A |
| chr17 | 71100000 | 71200000 | -0.09 | 0.09 | A>A |
| chr17 | 71200000 | 71300000 | 0.26 | 0.29 | A>A |
| chr17 | 71300000 | 71400000 | 0.34 | 0.23 | A>A |
| chr17 | 71400000 | 71500000 | 0.14 | 0.00 | A>A |
| chr17 | 71500000 | 71600000 | 0.17 | -0.03 | A>A |
| chr17 | 71600000 | 71700000 | 0.19 | -0.03 | A>A |
| chr17 | 71700000 | 71800000 | 0.41 | 0.01 | A>A |
| chr17 | 71800000 | 71900000 | 0.35 | 0.01 | A>A |
| chr17 | 71900000 | 72000000 | 0.15 | -0.09 | A>A |
| chr17 | 72000000 | 72100000 | 0.29 | -0.04 | A>A |
| chr17 | 72100000 | 72200000 | 0.27 | 0.10 | A>A |
| chr17 | 72200000 | 72300000 | 0.66 | 0.44 | A>A |
| chr17 | 72300000 | 72400000 | 0.78 | 0.52 | A>A |
| chr17 | 72400000 | 72500000 | 0.90 | 0.68 | A>A |
| chr17 | 72500000 | 72600000 | 0.77 | 0.41 | A>A |
| chr17 | 72600000 | 72700000 | 0.87 | 0.56 | A>A |
| chr17 | 72700000 | 72800000 | 1.10 | 0.97 | A>A |
| chr17 | 72800000 | 72900000 | 1.15 | 1.12 | A>A |
| chr17 | 72900000 | 73000000 | 1.27 | 1.30 | A>A |
| chr17 | 73000000 | 73100000 | 1.45 | 1.51 | A>A |
| chr17 | 73100000 | 73200000 | 1.37 | 1.35 | A>A |
| chr17 | 73200000 | 73300000 | 1.29 | 1.33 | A>A |

| | | | | | |
|---|---|---|---|---|---|
| chr17 | 73300000 | 73400000 | 1.30 | 1.39 | A>A |
| chr17 | 73400000 | 73500000 | 1.32 | 1.36 | A>A |
| chr17 | 73500000 | 73600000 | 1.43 | 1.35 | A>A |
| chr17 | 73600000 | 73700000 | 1.36 | 1.36 | A>A |
| chr17 | 73700000 | 73800000 | 1.43 | 1.43 | A>A |
| chr17 | 73800000 | 73900000 | 1.39 | 1.41 | A>A |
| chr17 | 73900000 | 74000000 | 1.40 | 1.22 | A>A |
| chr17 | 74000000 | 74100000 | 1.22 | 1.25 | A>A |
| chr17 | 74100000 | 74200000 | 1.23 | 1.22 | A>A |
| chr17 | 74200000 | 74300000 | 1.02 | 1.05 | A>A |
| chr17 | 74300000 | 74400000 | 1.04 | 0.98 | A>A |
| chr17 | 74400000 | 74500000 | 1.10 | 0.95 | A>A |
| chr17 | 74500000 | 74600000 | 0.99 | 0.93 | A>A |
| chr17 | 74600000 | 74700000 | 1.06 | 1.02 | A>A |
| chr17 | 74700000 | 74800000 | 0.85 | 0.85 | A>A |
| chr17 | 74800000 | 74900000 | 0.68 | 0.63 | A>A |
| chr17 | 74900000 | 75000000 | 0.80 | 0.61 | A>A |
| chr17 | 75000000 | 75100000 | 0.56 | 0.51 | A>A |
| chr17 | 75100000 | 75200000 | 0.65 | 0.73 | A>A |
| chr17 | 75200000 | 75300000 | 0.57 | 0.57 | A>A |
| chr17 | 75300000 | 75400000 | 0.54 | 0.56 | A>A |
| chr17 | 75400000 | 75500000 | 0.40 | 0.42 | A>A |
| chr17 | 75500000 | 75600000 | 0.17 | 0.26 | A>A |
| chr17 | 75600000 | 75700000 | 0.19 | 0.32 | A>A |
| chr17 | 75700000 | 75800000 | 0.19 | 0.26 | A>A |
| chr17 | 75800000 | 75900000 | 0.31 | 0.27 | A>A |
| chr17 | 75900000 | 76000000 | 0.27 | 0.34 | A>A |
| chr17 | 76000000 | 76100000 | 0.39 | 0.45 | A>A |
| chr17 | 76100000 | 76200000 | 0.72 | 0.79 | A>A |
| chr17 | 76200000 | 76300000 | 0.77 | 0.66 | A>A |
| chr17 | 76300000 | 76400000 | 0.80 | 0.83 | A>A |
| chr17 | 76400000 | 76500000 | 0.46 | 0.51 | A>A |
| chr17 | 76500000 | 76600000 | 0.63 | 0.66 | A>A |
| chr17 | 76600000 | 76700000 | 0.68 | 0.67 | A>A |
| chr17 | 76700000 | 76800000 | 0.47 | 0.56 | A>A |
| chr17 | 76800000 | 76900000 | 0.61 | 0.75 | A>A |
| chr17 | 76900000 | 77000000 | 0.55 | 0.79 | A>A |
| chr17 | 77000000 | 77100000 | 0.24 | 0.52 | A>A |
| chr17 | 77100000 | 77200000 | 0.03 | 0.37 | A>A |
| chr17 | 77200000 | 77300000 | -0.06 | 0.39 | A>A |
| chr17 | 77300000 | 77400000 | -0.02 | 0.34 | A>A |
| chr17 | 77400000 | 77500000 | 0.17 | 0.40 | A>A |
| chr17 | 77500000 | 77600000 | 0.00 | 0.00 | NO |
| chr17 | 77600000 | 77700000 | 0.11 | 0.31 | A>A |
| chr17 | 77700000 | 77800000 | 0.53 | 0.62 | A>A |
| chr17 | 77800000 | 77900000 | 0.71 | 0.82 | A>A |
| chr17 | 77900000 | 78000000 | 0.57 | 0.69 | A>A |
| chr17 | 78000000 | 78100000 | 0.63 | 0.78 | A>A |
| chr17 | 78100000 | 78200000 | 0.74 | 0.90 | A>A |
| chr17 | 78200000 | 78300000 | 0.82 | 0.87 | A>A |
| chr17 | 78300000 | 78400000 | 0.65 | 0.67 | A>A |
| chr17 | 78400000 | 78500000 | 0.61 | 0.67 | A>A |
| chr17 | 78500000 | 78600000 | 0.52 | 0.62 | A>A |
| chr17 | 78600000 | 78700000 | 0.50 | 0.59 | A>A |
| chr17 | 78700000 | 78800000 | 0.62 | 0.64 | A>A |
| chr17 | 78800000 | 78900000 | 0.46 | 0.61 | A>A |

| | | | | | |
|---|---|---|---|---|---|
| chr17 | 78900000 | 79000000 | 0.50 | 0.61 | A>A |
| chr17 | 79000000 | 79100000 | 0.86 | 0.88 | A>A |
| chr17 | 79100000 | 79200000 | 0.87 | 0.86 | A>A |
| chr17 | 79200000 | 79300000 | 0.80 | 0.86 | A>A |
| chr17 | 79300000 | 79400000 | 1.17 | 1.10 | A>A |
| chr17 | 79400000 | 79500000 | 0.84 | 0.94 | A>A |
| chr17 | 79500000 | 79600000 | 0.95 | 1.00 | A>A |
| chr17 | 79600000 | 79700000 | 1.06 | 1.05 | A>A |
| chr17 | 79700000 | 79800000 | 0.00 | 0.00 | NO |
| chr17 | 79800000 | 79900000 | 1.23 | 1.13 | A>A |
| chr17 | 79900000 | 80000000 | 1.15 | 1.10 | A>A |
| chr17 | 80000000 | 80100000 | 1.11 | 1.08 | A>A |
| chr17 | 80100000 | 80200000 | 1.00 | 0.98 | A>A |
| chr17 | 80200000 | 80300000 | 1.03 | 1.03 | A>A |
| chr17 | 80300000 | 80400000 | 0.88 | 0.91 | A>A |
| chr17 | 80400000 | 80500000 | 0.96 | 0.97 | A>A |
| chr17 | 80500000 | 80600000 | 0.77 | 0.83 | A>A |
| chr17 | 80600000 | 80700000 | 0.88 | 0.86 | A>A |
| chr17 | 80700000 | 80800000 | 0.61 | 0.61 | A>A |
| chr17 | 80800000 | 80900000 | 0.66 | 0.57 | A>A |
| chr17 | 80900000 | 81000000 | 0.66 | 0.58 | A>A |
| chr17 | 81000000 | 81100000 | 0.68 | 0.64 | A>A |
| chr17 | 81100000 | 81195210 | 0.00 | 0.00 | NO |
| chr18 | 0 | 100000 | 0.00 | 0.00 | NO |
| chr18 | 100000 | 200000 | 0.00 | 0.00 | NO |
| chr18 | 200000 | 300000 | 0.82 | 0.52 | A>A |
| chr18 | 300000 | 400000 | 0.57 | -0.12 | A>B |
| chr18 | 400000 | 500000 | 0.70 | -0.10 | A>B |
| chr18 | 500000 | 600000 | 0.67 | 0.09 | A>A |
| chr18 | 600000 | 700000 | 0.79 | 0.58 | A>A |
| chr18 | 700000 | 800000 | 0.76 | 0.72 | A>A |
| chr18 | 800000 | 900000 | 0.33 | 0.28 | A>A |
| chr18 | 900000 | 1000000 | -0.01 | -0.34 | B>B |
| chr18 | 1000000 | 1100000 | -0.41 | -0.69 | B>B |
| chr18 | 1100000 | 1200000 | -0.50 | -0.67 | B>B |
| chr18 | 1200000 | 1300000 | -0.47 | -0.47 | B>B |
| chr18 | 1300000 | 1400000 | -0.22 | -0.33 | B>B |
| chr18 | 1400000 | 1500000 | -0.40 | -0.47 | B>B |
| chr18 | 1500000 | 1600000 | -0.30 | -0.41 | B>B |
| chr18 | 1600000 | 1700000 | -0.45 | -0.49 | B>B |
| chr18 | 1700000 | 1800000 | -0.50 | -0.58 | B>B |
| chr18 | 1800000 | 1900000 | -0.57 | -0.34 | B>B |
| chr18 | 1900000 | 2000000 | -0.40 | -0.07 | B>B |
| chr18 | 2000000 | 2100000 | -0.24 | 0.01 | B>B |
| chr18 | 2100000 | 2200000 | -0.05 | 0.03 | B>B |
| chr18 | 2200000 | 2300000 | 0.15 | 0.22 | A>A |
| chr18 | 2300000 | 2400000 | 0.27 | 0.39 | A>A |
| chr18 | 2400000 | 2500000 | 0.25 | 0.37 | A>A |
| chr18 | 2500000 | 2600000 | 0.36 | 0.46 | A>A |
| chr18 | 2600000 | 2700000 | 0.82 | 0.79 | A>A |
| chr18 | 2700000 | 2800000 | 0.80 | 0.83 | A>A |
| chr18 | 2800000 | 2900000 | 0.81 | 0.78 | A>A |
| chr18 | 2900000 | 3000000 | 0.76 | 0.84 | A>A |
| chr18 | 3000000 | 3100000 | 0.99 | 0.95 | A>A |
| chr18 | 3100000 | 3200000 | 0.70 | 0.89 | A>A |
| chr18 | 3200000 | 3300000 | 0.66 | 0.68 | A>A |

| | | | | | |
|---|---|---|---|---|---|
| chr18 | 3300000 | 3400000 | 0.85 | 0.82 | A>A |
| chr18 | 3400000 | 3500000 | 0.78 | 0.87 | A>A |
| chr18 | 3500000 | 3600000 | 0.79 | 0.92 | A>A |
| chr18 | 3600000 | 3700000 | 0.75 | 0.70 | A>A |
| chr18 | 3700000 | 3800000 | 0.63 | 0.63 | A>A |
| chr18 | 3800000 | 3900000 | 0.66 | 0.62 | A>A |
| chr18 | 3900000 | 4000000 | 0.56 | 0.54 | A>A |
| chr18 | 4000000 | 4100000 | 0.28 | 0.11 | A>A |
| chr18 | 4100000 | 4200000 | -0.21 | -0.68 | B>B |
| chr18 | 4200000 | 4300000 | -0.42 | -1.15 | A>B |
| chr18 | 4300000 | 4400000 | -0.35 | -1.07 | A>B |
| chr18 | 4400000 | 4500000 | -0.51 | -1.11 | A>B |
| chr18 | 4500000 | 4600000 | -0.79 | -1.06 | B>B |
| chr18 | 4600000 | 4700000 | -1.01 | -1.06 | B>B |
| chr18 | 4700000 | 4800000 | -0.96 | -1.00 | B>B |
| chr18 | 4800000 | 4900000 | -0.95 | -1.00 | B>B |
| chr18 | 4900000 | 5000000 | -0.87 | -0.89 | B>B |
| chr18 | 5000000 | 5100000 | -0.46 | -0.63 | B>B |
| chr18 | 5100000 | 5200000 | -0.38 | -0.73 | B>B |
| chr18 | 5200000 | 5300000 | -0.06 | -0.31 | B>B |
| chr18 | 5300000 | 5400000 | -0.19 | -0.67 | B>B |
| chr18 | 5400000 | 5500000 | -0.18 | -0.79 | A>B |
| chr18 | 5500000 | 5600000 | -0.23 | -0.80 | B>B |
| chr18 | 5600000 | 5700000 | -0.26 | -0.74 | B>B |
| chr18 | 5700000 | 5800000 | -0.37 | -0.84 | B>B |
| chr18 | 5800000 | 5900000 | -0.21 | -0.58 | B>B |
| chr18 | 5900000 | 6000000 | -0.14 | -0.57 | B>B |
| chr18 | 6000000 | 6100000 | -0.25 | -0.72 | B>B |
| chr18 | 6100000 | 6200000 | -0.42 | -0.73 | B>B |
| chr18 | 6200000 | 6300000 | -0.39 | -0.73 | B>B |
| chr18 | 6300000 | 6400000 | -0.38 | -0.68 | B>B |
| chr18 | 6400000 | 6500000 | -0.27 | -0.72 | B>B |
| chr18 | 6500000 | 6600000 | -0.26 | -0.76 | B>B |
| chr18 | 6600000 | 6700000 | -0.34 | -0.91 | B>B |
| chr18 | 6700000 | 6800000 | -0.24 | -0.85 | A>B |
| chr18 | 6800000 | 6900000 | -0.10 | -0.62 | B>B |
| chr18 | 6900000 | 7000000 | 0.13 | -0.38 | B>B |
| chr18 | 7000000 | 7100000 | 0.30 | -0.07 | A>A |
| chr18 | 7100000 | 7200000 | 0.39 | 0.10 | A>A |
| chr18 | 7200000 | 7300000 | 0.46 | 0.51 | A>A |
| chr18 | 7300000 | 7400000 | 0.44 | 0.62 | A>A |
| chr18 | 7400000 | 7500000 | 0.67 | 0.61 | A>A |
| chr18 | 7500000 | 7600000 | 0.57 | 0.62 | A>A |
| chr18 | 7600000 | 7700000 | 0.56 | 0.72 | A>A |
| chr18 | 7700000 | 7800000 | 0.53 | 0.67 | A>A |
| chr18 | 7800000 | 7900000 | 0.59 | 0.68 | A>A |
| chr18 | 7900000 | 8000000 | 0.56 | 0.72 | A>A |
| chr18 | 8000000 | 8100000 | 0.49 | 0.64 | A>A |
| chr18 | 8100000 | 8200000 | 0.60 | 0.62 | A>A |
| chr18 | 8200000 | 8300000 | 0.58 | 0.67 | A>A |
| chr18 | 8300000 | 8400000 | 0.56 | 0.61 | A>A |
| chr18 | 8400000 | 8500000 | 0.57 | 0.61 | A>A |
| chr18 | 8500000 | 8600000 | 0.62 | 0.67 | A>A |
| chr18 | 8600000 | 8700000 | 0.65 | 0.73 | A>A |
| chr18 | 8700000 | 8800000 | 0.87 | 0.92 | A>A |
| chr18 | 8800000 | 8900000 | 0.77 | 0.81 | A>A |

| | | | | | |
|---|---|---|---|---|---|
| chr18 | 8900000 | 9000000 | 0.72 | 0.76 | A>A |
| chr18 | 9000000 | 9100000 | 0.67 | 0.73 | A>A |
| chr18 | 9100000 | 9200000 | 0.74 | 0.74 | A>A |
| chr18 | 9200000 | 9300000 | 0.68 | 0.77 | A>A |
| chr18 | 9300000 | 9400000 | 0.63 | 0.71 | A>A |
| chr18 | 9400000 | 9500000 | 0.71 | 0.74 | A>A |
| chr18 | 9500000 | 9600000 | 0.66 | 0.63 | A>A |
| chr18 | 9600000 | 9700000 | 0.73 | 0.69 | A>A |
| chr18 | 9700000 | 9800000 | 0.62 | 0.70 | A>A |
| chr18 | 9800000 | 9900000 | 0.72 | 0.66 | A>A |
| chr18 | 9900000 | 10000000 | 0.54 | 0.52 | A>A |
| chr18 | 10000000 | 10100000 | 0.52 | 0.37 | A>A |
| chr18 | 10100000 | 10200000 | 0.55 | 0.38 | A>A |
| chr18 | 10200000 | 10300000 | 0.48 | 0.38 | A>A |
| chr18 | 10300000 | 10400000 | 0.39 | 0.27 | A>A |
| chr18 | 10400000 | 10500000 | 0.48 | 0.25 | A>A |
| chr18 | 10500000 | 10600000 | 0.49 | 0.40 | A>A |
| chr18 | 10600000 | 10700000 | 0.00 | 0.00 | NO |
| chr18 | 10700000 | 10800000 | 0.44 | 0.30 | A>A |
| chr18 | 10800000 | 10900000 | 0.42 | 0.31 | A>A |
| chr18 | 10900000 | 11000000 | 0.38 | 0.25 | A>A |
| chr18 | 11000000 | 11100000 | 0.38 | 0.23 | A>A |
| chr18 | 11100000 | 11200000 | 0.40 | 0.26 | A>A |
| chr18 | 11200000 | 11300000 | 0.46 | 0.27 | A>A |
| chr18 | 11300000 | 11400000 | 0.54 | 0.40 | A>A |
| chr18 | 11400000 | 11500000 | 0.55 | 0.42 | A>A |
| chr18 | 11500000 | 11600000 | 0.44 | 0.37 | A>A |
| chr18 | 11600000 | 11700000 | 0.00 | 0.48 | NO |
| chr18 | 11700000 | 11800000 | 0.74 | 0.54 | A>A |
| chr18 | 11800000 | 11900000 | 0.80 | 0.64 | A>A |
| chr18 | 11900000 | 12000000 | 0.83 | 0.74 | A>A |
| chr18 | 12000000 | 12100000 | 0.63 | 0.57 | A>A |
| chr18 | 12100000 | 12200000 | 0.00 | 0.00 | NO |
| chr18 | 12200000 | 12300000 | 0.69 | 0.66 | A>A |
| chr18 | 12300000 | 12400000 | 0.75 | 0.70 | A>A |
| chr18 | 12400000 | 12500000 | 0.76 | 0.71 | A>A |
| chr18 | 12500000 | 12600000 | 0.76 | 0.77 | A>A |
| chr18 | 12600000 | 12700000 | 0.74 | 0.72 | A>A |
| chr18 | 12700000 | 12800000 | 0.87 | 0.77 | A>A |
| chr18 | 12800000 | 12900000 | 0.89 | 0.81 | A>A |
| chr18 | 12900000 | 13000000 | 0.81 | 0.75 | A>A |
| chr18 | 13000000 | 13100000 | 0.91 | 0.80 | A>A |
| chr18 | 13100000 | 13200000 | 0.65 | 0.64 | A>A |
| chr18 | 13200000 | 13300000 | 0.38 | 0.49 | A>A |
| chr18 | 13300000 | 13400000 | 0.29 | 0.44 | A>A |
| chr18 | 13400000 | 13500000 | 0.44 | 0.51 | A>A |
| chr18 | 13500000 | 13600000 | 0.43 | 0.51 | A>A |
| chr18 | 13600000 | 13700000 | 0.47 | 0.63 | A>A |
| chr18 | 13700000 | 13800000 | 0.61 | 0.58 | A>A |
| chr18 | 13800000 | 13900000 | 0.01 | 0.16 | A>A |
| chr18 | 13900000 | 14000000 | -0.38 | -0.26 | B>B |
| chr18 | 14000000 | 14100000 | -0.31 | 0.01 | B>B |
| chr18 | 14100000 | 14200000 | 0.00 | 0.00 | NO |
| chr18 | 14200000 | 14300000 | 0.00 | 0.00 | NO |
| chr18 | 14300000 | 14400000 | 0.00 | 0.00 | NO |
| chr18 | 14400000 | 14500000 | 0.00 | 0.00 | NO |

| | | | | | |
|---|---|---|---|---|---|
| chr18 | 14500000 | 14600000 | 0.00 | 0.00 | NO |
| chr18 | 14600000 | 14700000 | 0.00 | 0.00 | NO |
| chr18 | 14700000 | 14800000 | 0.00 | 0.00 | NO |
| chr18 | 14800000 | 14900000 | 0.00 | 0.00 | NO |
| chr18 | 14900000 | 15000000 | 0.00 | 0.00 | NO |
| chr18 | 15000000 | 15100000 | 0.00 | 0.00 | NO |
| chr18 | 15100000 | 15200000 | 0.00 | 0.00 | NO |
| chr18 | 15200000 | 15300000 | 0.00 | 0.00 | NO |
| chr18 | 15300000 | 15400000 | 0.00 | 0.00 | NO |
| chr18 | 15400000 | 15500000 | 0.00 | 0.00 | NO |
| chr18 | 15500000 | 15600000 | 0.00 | 0.00 | NO |
| chr18 | 15600000 | 15700000 | 0.00 | 0.00 | NO |
| chr18 | 15700000 | 15800000 | 0.00 | 0.00 | NO |
| chr18 | 15800000 | 15900000 | 0.00 | 0.00 | NO |
| chr18 | 15900000 | 16000000 | 0.00 | 0.00 | NO |
| chr18 | 16000000 | 16100000 | 0.00 | 0.00 | NO |
| chr18 | 16100000 | 16200000 | 0.00 | 0.00 | NO |
| chr18 | 16200000 | 16300000 | 0.00 | 0.00 | NO |
| chr18 | 16300000 | 16400000 | 0.00 | 0.00 | NO |
| chr18 | 16400000 | 16500000 | 0.00 | 0.00 | NO |
| chr18 | 16500000 | 16600000 | 0.00 | 0.00 | NO |
| chr18 | 16600000 | 16700000 | 0.00 | 0.00 | NO |
| chr18 | 16700000 | 16800000 | 0.00 | 0.00 | NO |
| chr18 | 16800000 | 16900000 | 0.00 | 0.00 | NO |
| chr18 | 16900000 | 17000000 | 0.00 | 0.00 | NO |
| chr18 | 17000000 | 17100000 | 0.00 | 0.00 | NO |
| chr18 | 17100000 | 17200000 | 0.00 | 0.00 | NO |
| chr18 | 17200000 | 17300000 | 0.00 | 0.00 | NO |
| chr18 | 17300000 | 17400000 | 0.00 | 0.00 | NO |
| chr18 | 17400000 | 17500000 | 0.00 | 0.00 | NO |
| chr18 | 17500000 | 17600000 | 0.00 | 0.00 | NO |
| chr18 | 17600000 | 17700000 | 0.00 | 0.00 | NO |
| chr18 | 17700000 | 17800000 | 0.00 | 0.00 | NO |
| chr18 | 17800000 | 17900000 | 0.00 | 0.00 | NO |
| chr18 | 17900000 | 18000000 | 0.00 | 0.00 | NO |
| chr18 | 18000000 | 18100000 | 0.00 | 0.00 | NO |
| chr18 | 18100000 | 18200000 | 0.00 | 0.00 | NO |
| chr18 | 18200000 | 18300000 | 0.00 | 0.00 | NO |
| chr18 | 18300000 | 18400000 | 0.00 | 0.00 | NO |
| chr18 | 18400000 | 18500000 | 0.00 | 0.00 | NO |
| chr18 | 18500000 | 18600000 | 0.00 | 0.00 | NO |
| chr18 | 18600000 | 18700000 | 0.78 | 0.78 | A>A |
| chr18 | 18700000 | 18800000 | 0.47 | 0.70 | A>A |
| chr18 | 18800000 | 18900000 | 0.69 | 0.69 | A>A |
| chr18 | 18900000 | 19000000 | 0.59 | 0.75 | A>A |
| chr18 | 19000000 | 19100000 | 0.65 | 0.73 | A>A |
| chr18 | 19100000 | 19200000 | 0.86 | 0.81 | A>A |
| chr18 | 19200000 | 19300000 | 0.83 | 0.82 | A>A |
| chr18 | 19300000 | 19400000 | 0.81 | 0.75 | A>A |
| chr18 | 19400000 | 19500000 | 0.64 | 0.71 | A>A |
| chr18 | 19500000 | 19600000 | 0.65 | 0.69 | A>A |
| chr18 | 19600000 | 19700000 | 0.69 | 0.71 | A>A |
| chr18 | 19700000 | 19800000 | 0.62 | 0.62 | A>A |
| chr18 | 19800000 | 19900000 | 0.66 | 0.71 | A>A |
| chr18 | 19900000 | 20000000 | 0.57 | 0.64 | A>A |
| chr18 | 20000000 | 20100000 | 0.65 | 0.69 | A>A |

| | | | | | |
|---|---|---|---|---|---|
| chr18 | 20100000 | 20200000 | 0.62 | 0.65 | A>A |
| chr18 | 20200000 | 20300000 | 0.68 | 0.73 | A>A |
| chr18 | 20300000 | 20400000 | 0.66 | 0.66 | A>A |
| chr18 | 20400000 | 20500000 | 0.66 | 0.71 | A>A |
| chr18 | 20500000 | 20600000 | 0.69 | 0.72 | A>A |
| chr18 | 20600000 | 20700000 | 0.73 | 0.69 | A>A |
| chr18 | 20700000 | 20800000 | 0.80 | 0.78 | A>A |
| chr18 | 20800000 | 20900000 | 0.71 | 0.76 | A>A |
| chr18 | 20900000 | 21000000 | 0.66 | 0.73 | A>A |
| chr18 | 21000000 | 21100000 | 0.73 | 0.76 | A>A |
| chr18 | 21100000 | 21200000 | 0.82 | 0.85 | A>A |
| chr18 | 21200000 | 21300000 | 0.92 | 0.90 | A>A |
| chr18 | 21300000 | 21400000 | 0.80 | 0.84 | A>A |
| chr18 | 21400000 | 21500000 | 0.76 | 0.85 | A>A |
| chr18 | 21500000 | 21600000 | 0.80 | 0.91 | A>A |
| chr18 | 21600000 | 21700000 | 0.90 | 0.89 | A>A |
| chr18 | 21700000 | 21800000 | 0.81 | 0.78 | A>A |
| chr18 | 21800000 | 21900000 | 0.75 | 0.76 | A>A |
| chr18 | 21900000 | 22000000 | 0.70 | 0.58 | A>A |
| chr18 | 22000000 | 22100000 | 0.59 | 0.12 | A>A |
| chr18 | 22100000 | 22200000 | -0.10 | -0.58 | B>B |
| chr18 | 22200000 | 22300000 | -0.37 | -1.13 | A>B |
| chr18 | 22300000 | 22400000 | -0.41 | -1.17 | A>B |
| chr18 | 22400000 | 22500000 | -0.44 | -1.32 | A>B |
| chr18 | 22500000 | 22600000 | -0.63 | -1.37 | A>B |
| chr18 | 22600000 | 22700000 | -0.46 | -1.39 | A>B |
| chr18 | 22700000 | 22800000 | -0.59 | -1.34 | A>B |
| chr18 | 22800000 | 22900000 | -0.52 | -1.21 | A>B |
| chr18 | 22900000 | 23000000 | -0.49 | -0.95 | B>B |
| chr18 | 23000000 | 23100000 | -0.62 | -1.23 | A>B |
| chr18 | 23100000 | 23200000 | -0.60 | -1.21 | A>B |
| chr18 | 23200000 | 23300000 | -0.59 | -1.22 | A>B |
| chr18 | 23300000 | 23400000 | -0.51 | -1.18 | A>B |
| chr18 | 23400000 | 23500000 | -0.22 | -0.70 | B>B |
| chr18 | 23500000 | 23600000 | 0.28 | -0.12 | A>A |
| chr18 | 23600000 | 23700000 | 0.53 | 0.24 | A>A |
| chr18 | 23700000 | 23800000 | 0.41 | 0.11 | A>A |
| chr18 | 23800000 | 23900000 | 0.55 | 0.29 | A>A |
| chr18 | 23900000 | 24000000 | 0.57 | 0.43 | A>A |
| chr18 | 24000000 | 24100000 | 0.68 | 0.70 | A>A |
| chr18 | 24100000 | 24200000 | 0.72 | 0.67 | A>A |
| chr18 | 24200000 | 24300000 | 0.66 | 0.51 | A>A |
| chr18 | 24300000 | 24400000 | 0.56 | 0.42 | A>A |
| chr18 | 24400000 | 24500000 | 0.50 | 0.20 | A>A |
| chr18 | 24500000 | 24600000 | 0.28 | 0.04 | A>A |
| chr18 | 24600000 | 24700000 | -0.24 | -0.20 | B>B |
| chr18 | 24700000 | 24800000 | -0.26 | -0.15 | B>B |
| chr18 | 24800000 | 24900000 | -0.52 | -0.58 | B>B |
| chr18 | 24900000 | 25000000 | -0.62 | -0.87 | B>B |
| chr18 | 25000000 | 25100000 | -0.66 | -0.96 | B>B |
| chr18 | 25100000 | 25200000 | -0.53 | -0.51 | B>B |
| chr18 | 25200000 | 25300000 | -0.53 | -0.13 | B>B |
| chr18 | 25300000 | 25400000 | -0.44 | -0.03 | B>B |
| chr18 | 25400000 | 25500000 | -0.30 | 0.09 | B>B |
| chr18 | 25500000 | 25600000 | -0.04 | 0.14 | A>A |
| chr18 | 25600000 | 25700000 | 0.13 | 0.29 | A>A |

| | | | | | |
|---|---|---|---|---|---|
| chr18 | 25700000 | 25800000 | 0.00 | 0.11 | A>A |
| chr18 | 25800000 | 25900000 | -0.66 | -0.65 | B>B |
| chr18 | 25900000 | 26000000 | -0.89 | -0.95 | B>B |
| chr18 | 26000000 | 26100000 | -1.05 | -0.95 | B>B |
| chr18 | 26100000 | 26200000 | -1.09 | -0.98 | B>B |
| chr18 | 26200000 | 26300000 | -1.08 | -0.98 | B>B |
| chr18 | 26300000 | 26400000 | -1.02 | -0.96 | B>B |
| chr18 | 26400000 | 26500000 | -0.96 | -0.85 | B>B |
| chr18 | 26500000 | 26600000 | -0.76 | -0.64 | B>B |
| chr18 | 26600000 | 26700000 | -0.51 | -0.35 | B>B |
| chr18 | 26700000 | 26800000 | -0.39 | -0.07 | B>B |
| chr18 | 26800000 | 26900000 | -0.68 | -0.30 | B>B |
| chr18 | 26900000 | 27000000 | -0.81 | -0.33 | B>B |
| chr18 | 27000000 | 27100000 | -0.95 | -0.54 | B>B |
| chr18 | 27100000 | 27200000 | -0.92 | -0.73 | B>B |
| chr18 | 27200000 | 27300000 | -0.99 | -0.77 | B>B |
| chr18 | 27300000 | 27400000 | -1.04 | -0.76 | B>B |
| chr18 | 27400000 | 27500000 | -1.05 | -0.48 | B>A |
| chr18 | 27500000 | 27600000 | -1.10 | -0.35 | B>A |
| chr18 | 27600000 | 27700000 | -1.08 | -0.28 | B>A |
| chr18 | 27700000 | 27800000 | -1.14 | -0.30 | B>A |
| chr18 | 27800000 | 27900000 | -1.11 | -0.45 | B>A |
| chr18 | 27900000 | 28000000 | -1.16 | -0.68 | B>B |
| chr18 | 28000000 | 28100000 | -1.18 | -0.84 | B>B |
| chr18 | 28100000 | 28200000 | -1.22 | -0.87 | B>B |
| chr18 | 28200000 | 28300000 | -1.20 | -0.89 | B>B |
| chr18 | 28300000 | 28400000 | -1.08 | -0.72 | B>B |
| chr18 | 28400000 | 28500000 | -0.68 | -0.24 | B>B |
| chr18 | 28500000 | 28600000 | 0.01 | 0.31 | A>A |
| chr18 | 28600000 | 28700000 | 0.41 | 0.66 | A>A |
| chr18 | 28700000 | 28800000 | 0.59 | 0.74 | A>A |
| chr18 | 28800000 | 28900000 | 0.57 | 0.73 | A>A |
| chr18 | 28900000 | 29000000 | 0.55 | 0.74 | A>A |
| chr18 | 29000000 | 29100000 | 0.50 | 0.70 | A>A |
| chr18 | 29100000 | 29200000 | 0.49 | 0.58 | A>A |
| chr18 | 29200000 | 29300000 | 0.55 | 0.68 | A>A |
| chr18 | 29300000 | 29400000 | 0.43 | 0.65 | A>A |
| chr18 | 29400000 | 29500000 | 0.68 | 0.73 | A>A |
| chr18 | 29500000 | 29600000 | 0.61 | 0.68 | A>A |
| chr18 | 29600000 | 29700000 | 0.61 | 0.56 | A>A |
| chr18 | 29700000 | 29800000 | 0.04 | -0.15 | B>B |
| chr18 | 29800000 | 29900000 | 0.29 | 0.10 | A>A |
| chr18 | 29900000 | 30000000 | 0.46 | 0.41 | A>A |
| chr18 | 30000000 | 30100000 | 0.33 | 0.19 | A>A |
| chr18 | 30100000 | 30200000 | -0.14 | -0.35 | B>B |
| chr18 | 30200000 | 30300000 | -0.55 | -0.67 | B>B |
| chr18 | 30300000 | 30400000 | -0.70 | -0.60 | B>B |
| chr18 | 30400000 | 30500000 | -0.84 | -0.67 | B>B |
| chr18 | 30500000 | 30600000 | -0.93 | -0.77 | B>B |
| chr18 | 30600000 | 30700000 | -1.18 | -1.14 | B>B |
| chr18 | 30700000 | 30800000 | -1.12 | -1.14 | B>B |
| chr18 | 30800000 | 30900000 | -1.10 | -1.13 | B>B |
| chr18 | 30900000 | 31000000 | -1.23 | -1.23 | B>B |
| chr18 | 31000000 | 31100000 | -1.04 | -1.18 | B>B |
| chr18 | 31100000 | 31200000 | -0.93 | -1.13 | B>B |
| chr18 | 31200000 | 31300000 | -1.01 | -1.25 | B>B |

| | | | | | |
|---|---|---|---|---|---|
| chr18 | 31300000 | 31400000 | -1.05 | -1.16 | B>B |
| chr18 | 31400000 | 31500000 | -1.26 | -1.25 | B>B |
| chr18 | 31500000 | 31600000 | -1.34 | -1.24 | B>B |
| chr18 | 31600000 | 31700000 | -1.18 | -1.26 | B>B |
| chr18 | 31700000 | 31800000 | -1.08 | -1.14 | B>B |
| chr18 | 31800000 | 31900000 | -1.11 | -1.11 | B>B |
| chr18 | 31900000 | 32000000 | -1.05 | -1.15 | B>B |
| chr18 | 32000000 | 32100000 | -0.96 | -1.15 | B>B |
| chr18 | 32100000 | 32200000 | -0.79 | -1.01 | B>B |
| chr18 | 32200000 | 32300000 | -0.48 | -0.61 | B>B |
| chr18 | 32300000 | 32400000 | -0.28 | -0.47 | B>B |
| chr18 | 32400000 | 32500000 | 0.27 | 0.08 | A>A |
| chr18 | 32500000 | 32600000 | 0.67 | 0.56 | A>A |
| chr18 | 32600000 | 32700000 | 0.41 | 0.48 | A>A |
| chr18 | 32700000 | 32800000 | 0.27 | 0.39 | A>A |
| chr18 | 32800000 | 32900000 | 0.30 | 0.63 | A>A |
| chr18 | 32900000 | 33000000 | 0.45 | 0.64 | A>A |
| chr18 | 33000000 | 33100000 | 0.68 | 0.84 | A>A |
| chr18 | 33100000 | 33200000 | 0.61 | 0.76 | A>A |
| chr18 | 33200000 | 33300000 | 0.41 | 0.59 | A>A |
| chr18 | 33300000 | 33400000 | 0.26 | 0.35 | A>A |
| chr18 | 33400000 | 33500000 | 0.16 | 0.24 | A>A |
| chr18 | 33500000 | 33600000 | 0.46 | 0.49 | A>A |
| chr18 | 33600000 | 33700000 | 0.68 | 0.60 | A>A |
| chr18 | 33700000 | 33800000 | 0.75 | 0.72 | A>A |
| chr18 | 33800000 | 33900000 | 0.49 | 0.47 | A>A |
| chr18 | 33900000 | 34000000 | 0.54 | 0.33 | A>A |
| chr18 | 34000000 | 34100000 | 0.43 | 0.15 | A>A |
| chr18 | 34100000 | 34200000 | 0.40 | 0.12 | A>A |
| chr18 | 34200000 | 34300000 | 0.41 | 0.13 | A>A |
| chr18 | 34300000 | 34400000 | 0.40 | 0.29 | A>A |
| chr18 | 34400000 | 34500000 | 0.26 | 0.35 | A>A |
| chr18 | 34500000 | 34600000 | -0.19 | 0.00 | B>B |
| chr18 | 34600000 | 34700000 | -0.46 | -0.15 | B>B |
| chr18 | 34700000 | 34800000 | -0.20 | -0.04 | B>B |
| chr18 | 34800000 | 34900000 | 0.21 | 0.20 | A>A |
| chr18 | 34900000 | 35000000 | 0.22 | -0.03 | A>A |
| chr18 | 35000000 | 35100000 | 0.17 | -0.13 | A>A |
| chr18 | 35100000 | 35200000 | -0.16 | -0.46 | B>B |
| chr18 | 35200000 | 35300000 | -0.58 | -0.99 | B>B |
| chr18 | 35300000 | 35400000 | -0.76 | -1.03 | B>B |
| chr18 | 35400000 | 35500000 | -0.99 | -1.11 | B>B |
| chr18 | 35500000 | 35600000 | -1.07 | -1.10 | B>B |
| chr18 | 35600000 | 35700000 | -1.12 | -1.08 | B>B |
| chr18 | 35700000 | 35800000 | -1.13 | -1.12 | B>B |
| chr18 | 35800000 | 35900000 | -1.12 | -1.08 | B>B |
| chr18 | 35900000 | 36000000 | -1.06 | -1.04 | B>B |
| chr18 | 36000000 | 36100000 | -1.04 | -0.98 | B>B |
| chr18 | 36100000 | 36200000 | -1.05 | -0.98 | B>B |
| chr18 | 36200000 | 36300000 | -1.05 | -0.95 | B>B |
| chr18 | 36300000 | 36400000 | -0.99 | -0.90 | B>B |
| chr18 | 36400000 | 36500000 | -0.94 | -0.82 | B>B |
| chr18 | 36500000 | 36600000 | -0.84 | -0.70 | B>B |
| chr18 | 36600000 | 36700000 | -0.88 | -0.81 | B>B |
| chr18 | 36700000 | 36800000 | -0.81 | -0.76 | B>B |
| chr18 | 36800000 | 36900000 | -0.63 | -0.76 | B>B |

| | | | | | |
|---|---|---|---|---|---|
| chr18 | 36900000 | 37000000 | -0.42 | -0.45 | B>B |
| chr18 | 37000000 | 37100000 | -0.08 | -0.32 | B>B |
| chr18 | 37100000 | 37200000 | -0.05 | -0.13 | B>B |
| chr18 | 37200000 | 37300000 | 0.07 | 0.14 | A>A |
| chr18 | 37300000 | 37400000 | 0.12 | 0.11 | A>A |
| chr18 | 37400000 | 37500000 | -0.39 | -0.28 | B>B |
| chr18 | 37500000 | 37600000 | -0.49 | -0.22 | B>B |
| chr18 | 37600000 | 37700000 | -0.67 | -0.56 | B>B |
| chr18 | 37700000 | 37800000 | -0.85 | -0.66 | B>B |
| chr18 | 37800000 | 37900000 | -0.99 | -0.95 | B>B |
| chr18 | 37900000 | 38000000 | -0.99 | -1.03 | B>B |
| chr18 | 38000000 | 38100000 | -1.07 | -1.11 | B>B |
| chr18 | 38100000 | 38200000 | -1.01 | -1.11 | B>B |
| chr18 | 38200000 | 38300000 | -0.98 | -1.20 | B>B |
| chr18 | 38300000 | 38400000 | -0.88 | -1.16 | B>B |
| chr18 | 38400000 | 38500000 | -0.77 | -1.16 | B>B |
| chr18 | 38500000 | 38600000 | -0.83 | -1.14 | B>B |
| chr18 | 38600000 | 38700000 | -1.03 | -1.25 | B>B |
| chr18 | 38700000 | 38800000 | -1.02 | -1.19 | B>B |
| chr18 | 38800000 | 38900000 | -1.00 | -1.21 | B>B |
| chr18 | 38900000 | 39000000 | -0.94 | -1.21 | B>B |
| chr18 | 39000000 | 39100000 | -0.99 | -1.21 | B>B |
| chr18 | 39100000 | 39200000 | -0.98 | -1.18 | B>B |
| chr18 | 39200000 | 39300000 | -1.01 | -1.31 | B>B |
| chr18 | 39300000 | 39400000 | -0.75 | -1.20 | B>B |
| chr18 | 39400000 | 39500000 | -0.63 | -1.03 | B>B |
| chr18 | 39500000 | 39600000 | -0.04 | -0.28 | B>B |
| chr18 | 39600000 | 39700000 | -0.26 | -0.42 | B>B |
| chr18 | 39700000 | 39800000 | -0.61 | -1.14 | B>B |
| chr18 | 39800000 | 39900000 | -0.81 | -1.20 | B>B |
| chr18 | 39900000 | 40000000 | -1.01 | -1.24 | B>B |
| chr18 | 40000000 | 40100000 | -1.11 | -1.26 | B>B |
| chr18 | 40100000 | 40200000 | -1.05 | -1.23 | B>B |
| chr18 | 40200000 | 40300000 | -1.15 | -1.24 | B>B |
| chr18 | 40300000 | 40400000 | -1.20 | -1.24 | B>B |
| chr18 | 40400000 | 40500000 | -1.22 | -1.29 | B>B |
| chr18 | 40500000 | 40600000 | -1.25 | -1.30 | B>B |
| chr18 | 40600000 | 40700000 | -1.20 | -1.30 | B>B |
| chr18 | 40700000 | 40800000 | -1.18 | -1.29 | B>B |
| chr18 | 40800000 | 40900000 | -1.09 | -1.23 | B>B |
| chr18 | 40900000 | 41000000 | -1.24 | -1.24 | B>B |
| chr18 | 41000000 | 41100000 | -1.08 | -1.27 | B>B |
| chr18 | 41100000 | 41200000 | -0.86 | -1.02 | B>B |
| chr18 | 41200000 | 41300000 | -1.00 | -1.00 | B>B |
| chr18 | 41300000 | 41400000 | -1.28 | -1.11 | B>B |
| chr18 | 41400000 | 41500000 | -1.26 | -1.14 | B>B |
| chr18 | 41500000 | 41600000 | -1.16 | -1.07 | B>B |
| chr18 | 41600000 | 41700000 | -1.03 | -0.90 | B>B |
| chr18 | 41700000 | 41800000 | -0.94 | -0.79 | B>B |
| chr18 | 41800000 | 41900000 | -0.94 | -0.81 | B>B |
| chr18 | 41900000 | 42000000 | -0.66 | -0.74 | B>B |
| chr18 | 42000000 | 42100000 | -0.51 | -0.59 | B>B |
| chr18 | 42100000 | 42200000 | -0.16 | -0.26 | B>B |
| chr18 | 42200000 | 42300000 | 0.11 | 0.05 | A>A |
| chr18 | 42300000 | 42400000 | 0.31 | 0.27 | A>A |
| chr18 | 42400000 | 42500000 | 0.16 | 0.12 | A>A |

| | | | | | |
|---|---|---|---|---|---|
| chr18 | 42500000 | 42600000 | -0.01 | -0.10 | B>B |
| chr18 | 42600000 | 42700000 | -0.06 | -0.39 | B>B |
| chr18 | 42700000 | 42800000 | -0.19 | -0.61 | B>B |
| chr18 | 42800000 | 42900000 | -0.34 | -0.80 | B>B |
| chr18 | 42900000 | 43000000 | -0.38 | -1.04 | A>B |
| chr18 | 43000000 | 43100000 | -0.33 | -1.08 | A>B |
| chr18 | 43100000 | 43200000 | 0.04 | -0.79 | A>B |
| chr18 | 43200000 | 43300000 | 0.42 | 0.04 | A>A |
| chr18 | 43300000 | 43400000 | 0.78 | 0.52 | A>A |
| chr18 | 43400000 | 43500000 | 0.60 | 0.40 | A>A |
| chr18 | 43500000 | 43600000 | 0.64 | 0.34 | A>A |
| chr18 | 43600000 | 43700000 | 0.78 | 0.46 | A>A |
| chr18 | 43700000 | 43800000 | 0.80 | 0.58 | A>A |
| chr18 | 43800000 | 43900000 | 0.66 | 0.38 | A>A |
| chr18 | 43900000 | 44000000 | 0.57 | 0.16 | A>A |
| chr18 | 44000000 | 44100000 | 0.47 | 0.04 | A>A |
| chr18 | 44100000 | 44200000 | 0.30 | -0.03 | A>A |
| chr18 | 44200000 | 44300000 | 0.39 | 0.19 | A>A |
| chr18 | 44300000 | 44400000 | 0.47 | 0.46 | A>A |
| chr18 | 44400000 | 44500000 | 0.67 | 0.59 | A>A |
| chr18 | 44500000 | 44600000 | 0.42 | 0.36 | A>A |
| chr18 | 44600000 | 44700000 | 0.45 | 0.31 | A>A |
| chr18 | 44700000 | 44800000 | 0.30 | -0.03 | A>A |
| chr18 | 44800000 | 44900000 | 0.00 | -0.51 | B>B |
| chr18 | 44900000 | 45000000 | -0.01 | -0.65 | A>B |
| chr18 | 45000000 | 45100000 | 0.05 | -0.61 | A>B |
| chr18 | 45100000 | 45200000 | 0.07 | -0.44 | B>B |
| chr18 | 45200000 | 45300000 | 0.25 | -0.04 | A>A |
| chr18 | 45300000 | 45400000 | 0.60 | 0.57 | A>A |
| chr18 | 45400000 | 45500000 | 0.80 | 0.76 | A>A |
| chr18 | 45500000 | 45600000 | 0.63 | 0.63 | A>A |
| chr18 | 45600000 | 45700000 | 0.59 | 0.68 | A>A |
| chr18 | 45700000 | 45800000 | 0.59 | 0.67 | A>A |
| chr18 | 45800000 | 45900000 | 0.62 | 0.68 | A>A |
| chr18 | 45900000 | 46000000 | 0.57 | 0.63 | A>A |
| chr18 | 46000000 | 46100000 | 0.63 | 0.70 | A>A |
| chr18 | 46100000 | 46200000 | 0.59 | 0.68 | A>A |
| chr18 | 46200000 | 46300000 | 0.49 | 0.61 | A>A |
| chr18 | 46300000 | 46400000 | 0.60 | 0.75 | A>A |
| chr18 | 46400000 | 46500000 | 0.66 | 0.79 | A>A |
| chr18 | 46500000 | 46600000 | 0.64 | 0.70 | A>A |
| chr18 | 46600000 | 46700000 | 0.56 | 0.66 | A>A |
| chr18 | 46700000 | 46800000 | 0.55 | 0.62 | A>A |
| chr18 | 46800000 | 46900000 | 0.65 | 0.71 | A>A |
| chr18 | 46900000 | 47000000 | 0.60 | 0.62 | A>A |
| chr18 | 47000000 | 47100000 | 0.58 | 0.56 | A>A |
| chr18 | 47100000 | 47200000 | 0.58 | 0.53 | A>A |
| chr18 | 47200000 | 47300000 | 0.54 | 0.53 | A>A |
| chr18 | 47300000 | 47400000 | 0.57 | 0.52 | A>A |
| chr18 | 47400000 | 47500000 | 0.35 | 0.27 | A>A |
| chr18 | 47500000 | 47600000 | 0.22 | 0.08 | A>A |
| chr18 | 47600000 | 47700000 | 0.20 | 0.02 | A>A |
| chr18 | 47700000 | 47800000 | 0.39 | 0.26 | A>A |
| chr18 | 47800000 | 47900000 | 0.52 | 0.44 | A>A |
| chr18 | 47900000 | 48000000 | 0.29 | 0.15 | A>A |
| chr18 | 48000000 | 48100000 | 0.06 | -0.11 | B>B |

| | | | | | |
|---|---|---|---|---|---|
| chr18 | 48100000 | 48200000 | 0.15 | -0.08 | A>A |
| chr18 | 48200000 | 48300000 | 0.29 | -0.10 | A>A |
| chr18 | 48300000 | 48400000 | 0.47 | 0.24 | A>A |
| chr18 | 48400000 | 48500000 | 0.66 | 0.56 | A>A |
| chr18 | 48500000 | 48600000 | 0.57 | 0.50 | A>A |
| chr18 | 48600000 | 48700000 | 0.36 | 0.07 | A>A |
| chr18 | 48700000 | 48800000 | 0.46 | -0.03 | A>A |
| chr18 | 48800000 | 48900000 | 0.34 | -0.47 | A>B |
| chr18 | 48900000 | 49000000 | 0.33 | -0.61 | A>B |
| chr18 | 49000000 | 49100000 | 0.20 | -0.62 | A>B |
| chr18 | 49100000 | 49200000 | 0.01 | -0.62 | A>B |
| chr18 | 49200000 | 49300000 | -0.36 | -0.86 | B>B |
| chr18 | 49300000 | 49400000 | -0.53 | -1.03 | B>B |
| chr18 | 49400000 | 49500000 | -0.67 | -1.01 | B>B |
| chr18 | 49500000 | 49600000 | -0.76 | -0.96 | B>B |
| chr18 | 49600000 | 49700000 | -0.70 | -0.89 | B>B |
| chr18 | 49700000 | 49800000 | -0.44 | -0.78 | B>B |
| chr18 | 49800000 | 49900000 | -0.04 | -0.31 | B>B |
| chr18 | 49900000 | 50000000 | 0.18 | -0.05 | A>A |
| chr18 | 50000000 | 50100000 | 0.34 | 0.17 | A>A |
| chr18 | 50100000 | 50200000 | 0.23 | 0.00 | A>A |
| chr18 | 50200000 | 50300000 | -0.21 | -0.65 | B>B |
| chr18 | 50300000 | 50400000 | -0.64 | -1.12 | B>B |
| chr18 | 50400000 | 50500000 | -0.71 | -1.18 | B>B |
| chr18 | 50500000 | 50600000 | -1.14 | -1.29 | B>B |
| chr18 | 50600000 | 50700000 | -1.18 | -1.28 | B>B |
| chr18 | 50700000 | 50800000 | -1.37 | -1.34 | B>B |
| chr18 | 50800000 | 50900000 | -1.28 | -1.29 | B>B |
| chr18 | 50900000 | 51000000 | -1.08 | -1.30 | B>B |
| chr18 | 51000000 | 51100000 | -1.19 | -1.36 | B>B |
| chr18 | 51100000 | 51200000 | -1.21 | -1.34 | B>B |
| chr18 | 51200000 | 51300000 | -1.14 | -1.31 | B>B |
| chr18 | 51300000 | 51400000 | -1.03 | -1.30 | B>B |
| chr18 | 51400000 | 51500000 | -0.78 | -1.31 | B>B |
| chr18 | 51500000 | 51600000 | -0.67 | -1.19 | B>B |
| chr18 | 51600000 | 51700000 | -0.28 | -0.42 | B>B |
| chr18 | 51700000 | 51800000 | 0.49 | 0.39 | A>A |
| chr18 | 51800000 | 51900000 | 0.38 | 0.42 | A>A |
| chr18 | 51900000 | 52000000 | 0.18 | 0.37 | A>A |
| chr18 | 52000000 | 52100000 | 0.00 | 0.29 | NO |
| chr18 | 52100000 | 52200000 | 0.00 | 0.00 | NO |
| chr18 | 52200000 | 52300000 | 0.17 | 0.34 | A>A |
| chr18 | 52300000 | 52400000 | 0.43 | 0.54 | A>A |
| chr18 | 52400000 | 52500000 | 0.67 | 0.74 | A>A |
| chr18 | 52500000 | 52600000 | 0.60 | 0.61 | A>A |
| chr18 | 52600000 | 52700000 | 0.50 | 0.63 | A>A |
| chr18 | 52700000 | 52800000 | 0.48 | 0.67 | A>A |
| chr18 | 52800000 | 52900000 | 0.47 | 0.71 | A>A |
| chr18 | 52900000 | 53000000 | 0.34 | 0.60 | A>A |
| chr18 | 53000000 | 53100000 | 0.37 | 0.56 | A>A |
| chr18 | 53100000 | 53200000 | 0.30 | 0.56 | A>A |
| chr18 | 53200000 | 53300000 | 0.30 | 0.58 | A>A |
| chr18 | 53300000 | 53400000 | 0.22 | 0.50 | A>A |
| chr18 | 53400000 | 53500000 | 0.30 | 0.59 | A>A |
| chr18 | 53500000 | 53600000 | 0.31 | 0.56 | A>A |
| chr18 | 53600000 | 53700000 | 0.31 | 0.49 | A>A |

| | | | | | |
|---|---|---|---|---|---|
| chr18 | 53700000 | 53800000 | 0.28 | 0.49 | A>A |
| chr18 | 53800000 | 53900000 | 0.35 | 0.53 | A>A |
| chr18 | 53900000 | 54000000 | 0.36 | 0.54 | A>A |
| chr18 | 54000000 | 54100000 | 0.42 | 0.59 | A>A |
| chr18 | 54100000 | 54200000 | 0.33 | 0.56 | A>A |
| chr18 | 54200000 | 54300000 | 0.50 | 0.61 | A>A |
| chr18 | 54300000 | 54400000 | 0.64 | 0.69 | A>A |
| chr18 | 54400000 | 54500000 | 0.55 | 0.63 | A>A |
| chr18 | 54500000 | 54600000 | 0.58 | 0.64 | A>A |
| chr18 | 54600000 | 54700000 | 0.54 | 0.62 | A>A |
| chr18 | 54700000 | 54800000 | 0.44 | 0.55 | A>A |
| chr18 | 54800000 | 54900000 | 0.41 | 0.52 | A>A |
| chr18 | 54900000 | 55000000 | 0.47 | 0.50 | A>A |
| chr18 | 55000000 | 55100000 | 0.43 | 0.52 | A>A |
| chr18 | 55100000 | 55200000 | 0.63 | 0.57 | A>A |
| chr18 | 55200000 | 55300000 | 0.66 | 0.69 | A>A |
| chr18 | 55300000 | 55400000 | 0.84 | 0.72 | A>A |
| chr18 | 55400000 | 55500000 | 0.90 | 0.85 | A>A |
| chr18 | 55500000 | 55600000 | 0.75 | 0.78 | A>A |
| chr18 | 55600000 | 55700000 | 0.81 | 0.78 | A>A |
| chr18 | 55700000 | 55800000 | 0.70 | 0.74 | A>A |
| chr18 | 55800000 | 55900000 | 0.69 | 0.70 | A>A |
| chr18 | 55900000 | 56000000 | 0.75 | 0.77 | A>A |
| chr18 | 56000000 | 56100000 | 0.71 | 0.74 | A>A |
| chr18 | 56100000 | 56200000 | 0.70 | 0.72 | A>A |
| chr18 | 56200000 | 56300000 | 0.89 | 0.83 | A>A |
| chr18 | 56300000 | 56400000 | 0.77 | 0.81 | A>A |
| chr18 | 56400000 | 56500000 | 0.72 | 0.70 | A>A |
| chr18 | 56500000 | 56600000 | 0.92 | 0.89 | A>A |
| chr18 | 56600000 | 56700000 | 0.94 | 0.88 | A>A |
| chr18 | 56700000 | 56800000 | 0.72 | 0.69 | A>A |
| chr18 | 56800000 | 56900000 | 0.74 | 0.55 | A>A |
| chr18 | 56900000 | 57000000 | 0.66 | 0.56 | A>A |
| chr18 | 57000000 | 57100000 | 0.63 | 0.49 | A>A |
| chr18 | 57100000 | 57200000 | 0.25 | 0.14 | A>A |
| chr18 | 57200000 | 57300000 | 0.14 | 0.01 | A>A |
| chr18 | 57300000 | 57400000 | 0.31 | 0.07 | A>A |
| chr18 | 57400000 | 57500000 | 0.27 | 0.25 | A>A |
| chr18 | 57500000 | 57600000 | 0.47 | 0.44 | A>A |
| chr18 | 57600000 | 57700000 | 0.14 | 0.20 | A>A |
| chr18 | 57700000 | 57800000 | 0.12 | 0.19 | A>A |
| chr18 | 57800000 | 57900000 | -0.11 | 0.13 | A>A |
| chr18 | 57900000 | 58000000 | -0.30 | 0.10 | B>B |
| chr18 | 58000000 | 58100000 | -0.28 | -0.06 | B>B |
| chr18 | 58100000 | 58200000 | -0.47 | -0.36 | B>B |
| chr18 | 58200000 | 58300000 | -0.65 | -0.68 | B>B |
| chr18 | 58300000 | 58400000 | -0.99 | -1.16 | B>B |
| chr18 | 58400000 | 58500000 | -1.16 | -1.36 | B>B |
| chr18 | 58500000 | 58600000 | -1.32 | -1.40 | B>B |
| chr18 | 58600000 | 58700000 | -1.32 | -1.40 | B>B |
| chr18 | 58700000 | 58800000 | -1.23 | -1.40 | B>B |
| chr18 | 58800000 | 58900000 | -1.33 | -1.46 | B>B |
| chr18 | 58900000 | 59000000 | -1.02 | -1.46 | B>B |
| chr18 | 59000000 | 59100000 | -0.91 | -1.38 | B>B |
| chr18 | 59100000 | 59200000 | -0.63 | -1.22 | B>B |
| chr18 | 59200000 | 59300000 | -0.12 | -0.79 | A>B |

| | | | | | |
|---|---|---|---|---|---|
| chr18 | 59300000 | 59400000 | -0.36 | -1.16 | A>B |
| chr18 | 59400000 | 59500000 | -0.41 | -1.24 | A>B |
| chr18 | 59500000 | 59600000 | -0.17 | -0.94 | A>B |
| chr18 | 59600000 | 59700000 | -0.16 | -0.84 | A>B |
| chr18 | 59700000 | 59800000 | 0.15 | -0.24 | B>B |
| chr18 | 59800000 | 59900000 | 0.53 | 0.37 | A>A |
| chr18 | 59900000 | 60000000 | 0.78 | 0.66 | A>A |
| chr18 | 60000000 | 60100000 | 0.91 | 0.81 | A>A |
| chr18 | 60100000 | 60200000 | 1.06 | 0.87 | A>A |
| chr18 | 60200000 | 60300000 | 0.82 | 0.72 | A>A |
| chr18 | 60300000 | 60400000 | 0.75 | 0.76 | A>A |
| chr18 | 60400000 | 60500000 | 0.89 | 0.88 | A>A |
| chr18 | 60500000 | 60600000 | 0.83 | 0.82 | A>A |
| chr18 | 60600000 | 60700000 | 0.59 | 0.59 | A>A |
| chr18 | 60700000 | 60800000 | 0.65 | 0.43 | A>A |
| chr18 | 60800000 | 60900000 | 0.67 | 0.46 | A>A |
| chr18 | 60900000 | 61000000 | 0.70 | 0.48 | A>A |
| chr18 | 61000000 | 61100000 | 0.76 | 0.72 | A>A |
| chr18 | 61100000 | 61200000 | 0.17 | 0.54 | A>A |
| chr18 | 61200000 | 61300000 | -0.49 | 0.42 | B>A |
| chr18 | 61300000 | 61400000 | -0.87 | 0.42 | B>A |
| chr18 | 61400000 | 61500000 | -0.80 | 0.27 | B>A |
| chr18 | 61500000 | 61600000 | -0.44 | 0.05 | B>B |
| chr18 | 61600000 | 61700000 | -0.37 | -0.28 | B>B |
| chr18 | 61700000 | 61800000 | -1.03 | -1.09 | B>B |
| chr18 | 61800000 | 61900000 | -1.18 | -1.22 | B>B |
| chr18 | 61900000 | 62000000 | -1.25 | -1.34 | B>B |
| chr18 | 62000000 | 62100000 | -1.30 | -1.34 | B>B |
| chr18 | 62100000 | 62200000 | -1.23 | -1.29 | B>B |
| chr18 | 62200000 | 62300000 | -1.20 | -1.27 | B>B |
| chr18 | 62300000 | 62400000 | -1.16 | -1.24 | B>B |
| chr18 | 62400000 | 62500000 | -1.22 | -1.18 | B>B |
| chr18 | 62500000 | 62600000 | -1.14 | -1.22 | B>B |
| chr18 | 62600000 | 62700000 | -1.19 | -1.22 | B>B |
| chr18 | 62700000 | 62800000 | -1.06 | -1.23 | B>B |
| chr18 | 62800000 | 62900000 | -1.12 | -1.19 | B>B |
| chr18 | 62900000 | 63000000 | -1.12 | -1.19 | B>B |
| chr18 | 63000000 | 63100000 | -1.10 | -1.10 | B>B |
| chr18 | 63100000 | 63200000 | -1.07 | -1.13 | B>B |
| chr18 | 63200000 | 63300000 | -0.98 | -1.11 | B>B |
| chr18 | 63300000 | 63400000 | -0.90 | -1.09 | B>B |
| chr18 | 63400000 | 63500000 | -0.71 | -1.15 | B>B |
| chr18 | 63500000 | 63600000 | -0.93 | -1.10 | B>B |
| chr18 | 63600000 | 63700000 | -0.95 | -1.02 | B>B |
| chr18 | 63700000 | 63800000 | -0.99 | -1.11 | B>B |
| chr18 | 63800000 | 63900000 | -0.95 | -1.05 | B>B |
| chr18 | 63900000 | 64000000 | -0.94 | -0.98 | B>B |
| chr18 | 64000000 | 64100000 | -0.84 | -0.95 | B>B |
| chr18 | 64100000 | 64200000 | -0.40 | -0.64 | B>B |
| chr18 | 64200000 | 64300000 | -0.47 | -0.62 | B>B |
| chr18 | 64300000 | 64400000 | -0.82 | -0.76 | B>B |
| chr18 | 64400000 | 64500000 | -0.97 | -0.86 | B>B |
| chr18 | 64500000 | 64600000 | -1.15 | -0.90 | B>B |
| chr18 | 64600000 | 64700000 | -1.19 | -0.96 | B>B |
| chr18 | 64700000 | 64800000 | -1.14 | -0.94 | B>B |
| chr18 | 64800000 | 64900000 | -0.85 | -0.85 | B>B |

| | | | | | |
|---|---|---|---|---|---|
| chr18 | 64900000 | 65000000 | -0.86 | -0.90 | B>B |
| chr18 | 65000000 | 65100000 | -0.66 | -0.72 | B>B |
| chr18 | 65100000 | 65200000 | -0.62 | -0.75 | B>B |
| chr18 | 65200000 | 65300000 | -0.75 | -0.88 | B>B |
| chr18 | 65300000 | 65400000 | -0.92 | -0.96 | B>B |
| chr18 | 65400000 | 65500000 | -0.90 | -1.02 | B>B |
| chr18 | 65500000 | 65600000 | -0.95 | -1.11 | B>B |
| chr18 | 65600000 | 65700000 | -0.95 | -1.14 | B>B |
| chr18 | 65700000 | 65800000 | -1.08 | -1.17 | B>B |
| chr18 | 65800000 | 65900000 | -1.14 | -1.14 | B>B |
| chr18 | 65900000 | 66000000 | -1.17 | -1.15 | B>B |
| chr18 | 66000000 | 66100000 | -1.14 | -1.11 | B>B |
| chr18 | 66100000 | 66200000 | -1.12 | -1.04 | B>B |
| chr18 | 66200000 | 66300000 | -0.82 | -0.96 | B>B |
| chr18 | 66300000 | 66400000 | -0.30 | -0.48 | B>B |
| chr18 | 66400000 | 66500000 | -0.78 | -0.99 | B>B |
| chr18 | 66500000 | 66600000 | -0.74 | -1.06 | B>B |
| chr18 | 66600000 | 66700000 | -0.82 | -1.21 | B>B |
| chr18 | 66700000 | 66800000 | -0.84 | -1.21 | B>B |
| chr18 | 66800000 | 66900000 | -0.70 | -1.24 | B>B |
| chr18 | 66900000 | 67000000 | -0.77 | -1.16 | B>B |
| chr18 | 67000000 | 67100000 | -0.49 | -1.12 | A>B |
| chr18 | 67100000 | 67200000 | -0.29 | -1.11 | A>B |
| chr18 | 67200000 | 67300000 | -0.11 | -1.06 | A>B |
| chr18 | 67300000 | 67400000 | 0.05 | -1.07 | A>B |
| chr18 | 67400000 | 67500000 | 0.18 | -0.81 | A>B |
| chr18 | 67500000 | 67600000 | 0.21 | -0.14 | A>A |
| chr18 | 67600000 | 67700000 | 0.29 | 0.01 | A>A |
| chr18 | 67700000 | 67800000 | 0.43 | 0.04 | A>A |
| chr18 | 67800000 | 67900000 | 0.79 | 0.53 | A>A |
| chr18 | 67900000 | 68000000 | 1.15 | 0.96 | A>A |
| chr18 | 68000000 | 68100000 | 1.09 | 0.89 | A>A |
| chr18 | 68100000 | 68200000 | 0.22 | 0.10 | A>A |
| chr18 | 68200000 | 68300000 | -0.37 | -0.41 | B>B |
| chr18 | 68300000 | 68400000 | -0.62 | -0.61 | B>B |
| chr18 | 68400000 | 68500000 | -0.57 | -0.42 | B>B |
| chr18 | 68500000 | 68600000 | -0.45 | -0.33 | B>B |
| chr18 | 68600000 | 68700000 | -0.52 | -0.46 | B>B |
| chr18 | 68700000 | 68800000 | -0.72 | -0.60 | B>B |
| chr18 | 68800000 | 68900000 | -0.72 | -0.67 | B>B |
| chr18 | 68900000 | 69000000 | -0.91 | -0.58 | B>B |
| chr18 | 69000000 | 69100000 | -1.01 | -0.36 | B>A |
| chr18 | 69100000 | 69200000 | -1.09 | -0.23 | B>A |
| chr18 | 69200000 | 69300000 | -1.09 | -0.02 | B>A |
| chr18 | 69300000 | 69400000 | -1.12 | 0.13 | B>A |
| chr18 | 69400000 | 69500000 | -1.06 | 0.24 | B>A |
| chr18 | 69500000 | 69600000 | -0.98 | 0.41 | B>A |
| chr18 | 69600000 | 69700000 | -0.79 | 0.34 | B>A |
| chr18 | 69700000 | 69800000 | -0.38 | 0.44 | B>A |
| chr18 | 69800000 | 69900000 | -0.33 | 0.56 | B>A |
| chr18 | 69900000 | 70000000 | -0.23 | 0.51 | B>A |
| chr18 | 70000000 | 70100000 | -0.16 | 0.57 | B>A |
| chr18 | 70100000 | 70200000 | -0.08 | 0.39 | A>A |
| chr18 | 70200000 | 70300000 | -0.47 | 0.06 | B>A |
| chr18 | 70300000 | 70400000 | -0.54 | 0.02 | B>A |
| chr18 | 70400000 | 70500000 | -0.59 | 0.09 | B>A |

| | | | | | |
|---|---|---|---|---|---|
| chr18 | 70500000 | 70600000 | -0.86 | 0.08 | B>A |
| chr18 | 70600000 | 70700000 | -0.83 | 0.13 | B>A |
| chr18 | 70700000 | 70800000 | -0.52 | 0.31 | B>A |
| chr18 | 70800000 | 70900000 | 0.04 | 0.57 | B>A |
| chr18 | 70900000 | 71000000 | 0.27 | 0.84 | B>A |
| chr18 | 71000000 | 71100000 | -0.07 | 0.48 | B>A |
| chr18 | 71100000 | 71200000 | -0.21 | 0.28 | A>A |
| chr18 | 71200000 | 71300000 | -0.08 | 0.34 | A>A |
| chr18 | 71300000 | 71400000 | 0.16 | 0.38 | A>A |
| chr18 | 71400000 | 71500000 | 0.00 | 0.28 | A>A |
| chr18 | 71500000 | 71600000 | -0.14 | 0.12 | B>B |
| chr18 | 71600000 | 71700000 | -0.13 | -0.01 | B>B |
| chr18 | 71700000 | 71800000 | 0.05 | 0.11 | A>A |
| chr18 | 71800000 | 71900000 | 0.68 | 0.45 | A>A |
| chr18 | 71900000 | 72000000 | 0.97 | 0.68 | A>A |
| chr18 | 72000000 | 72100000 | 0.63 | 0.32 | A>A |
| chr18 | 72100000 | 72200000 | 0.66 | 0.37 | A>A |
| chr18 | 72200000 | 72300000 | 0.63 | 0.40 | A>A |
| chr18 | 72300000 | 72400000 | 0.45 | 0.25 | A>A |
| chr18 | 72400000 | 72500000 | 0.50 | 0.38 | A>A |
| chr18 | 72500000 | 72600000 | 0.49 | 0.42 | A>A |
| chr18 | 72600000 | 72700000 | 0.47 | 0.40 | A>A |
| chr18 | 72700000 | 72800000 | 0.36 | 0.36 | A>A |
| chr18 | 72800000 | 72900000 | 0.31 | 0.26 | A>A |
| chr18 | 72900000 | 73000000 | 0.43 | 0.41 | A>A |
| chr18 | 73000000 | 73100000 | 0.20 | 0.22 | A>A |
| chr18 | 73100000 | 73200000 | 0.20 | 0.17 | A>A |
| chr18 | 73200000 | 73300000 | 0.05 | -0.30 | B>B |
| chr18 | 73300000 | 73400000 | -0.24 | -0.55 | B>B |
| chr18 | 73400000 | 73500000 | -0.26 | -0.66 | B>B |
| chr18 | 73500000 | 73600000 | -0.30 | -0.65 | B>B |
| chr18 | 73600000 | 73700000 | -0.25 | -0.51 | B>B |
| chr18 | 73700000 | 73800000 | -0.41 | -0.65 | B>B |
| chr18 | 73800000 | 73900000 | -0.42 | -0.75 | B>B |
| chr18 | 73900000 | 74000000 | -0.20 | -0.54 | B>B |
| chr18 | 74000000 | 74100000 | 0.22 | -0.01 | A>A |
| chr18 | 74100000 | 74200000 | 0.80 | 0.60 | A>A |
| chr18 | 74200000 | 74300000 | 1.01 | 0.76 | A>A |
| chr18 | 74300000 | 74400000 | 0.89 | 0.64 | A>A |
| chr18 | 74400000 | 74500000 | 0.73 | 0.55 | A>A |
| chr18 | 74500000 | 74600000 | 0.92 | 0.94 | A>A |
| chr18 | 74600000 | 74700000 | 0.76 | 0.64 | A>A |
| chr18 | 74700000 | 74800000 | 0.71 | 0.50 | A>A |
| chr18 | 74800000 | 74900000 | 0.81 | 0.51 | A>A |
| chr18 | 74900000 | 75000000 | 0.44 | 0.08 | A>A |
| chr18 | 75000000 | 75100000 | 0.13 | -0.49 | A>B |
| chr18 | 75100000 | 75200000 | -0.08 | -0.76 | A>B |
| chr18 | 75200000 | 75300000 | -0.21 | -0.93 | A>B |
| chr18 | 75300000 | 75400000 | -0.16 | -0.89 | A>B |
| chr18 | 75400000 | 75500000 | -0.52 | -0.97 | B>B |
| chr18 | 75500000 | 75600000 | -0.78 | -1.19 | B>B |
| chr18 | 75600000 | 75700000 | -0.67 | -0.99 | B>B |
| chr18 | 75700000 | 75800000 | 0.00 | -1.22 | NO |
| chr18 | 75800000 | 75900000 | -0.81 | -1.15 | B>B |
| chr18 | 75900000 | 76000000 | -0.86 | -1.15 | B>B |
| chr18 | 76000000 | 76100000 | -0.94 | -1.08 | B>B |

| | | | | | |
|---|---|---|---|---|---|
| chr18 | 76100000 | 76200000 | -0.72 | -0.80 | B>B |
| chr18 | 76200000 | 76300000 | -0.64 | -0.50 | B>B |
| chr18 | 76300000 | 76400000 | -0.40 | -0.48 | B>B |
| chr18 | 76400000 | 76500000 | -0.17 | -0.40 | B>B |
| chr18 | 76500000 | 76600000 | -0.14 | -0.31 | B>B |
| chr18 | 76600000 | 76700000 | -0.10 | -0.23 | B>B |
| chr18 | 76700000 | 76800000 | 0.19 | 0.06 | A>A |
| chr18 | 76800000 | 76900000 | 0.65 | 0.49 | A>A |
| chr18 | 76900000 | 77000000 | 0.55 | 0.54 | A>A |
| chr18 | 77000000 | 77100000 | 0.58 | 0.58 | A>A |
| chr18 | 77100000 | 77200000 | 0.78 | 0.81 | A>A |
| chr18 | 77200000 | 77300000 | 0.85 | 0.86 | A>A |
| chr18 | 77300000 | 77400000 | 0.76 | 0.80 | A>A |
| chr18 | 77400000 | 77500000 | 0.87 | 0.97 | A>A |
| chr18 | 77500000 | 77600000 | 0.85 | 0.94 | A>A |
| chr18 | 77600000 | 77700000 | 0.87 | 1.03 | A>A |
| chr18 | 77700000 | 77800000 | 1.06 | 1.05 | A>A |
| chr18 | 77800000 | 77900000 | 1.11 | 1.00 | A>A |
| chr18 | 77900000 | 78000000 | 1.02 | 0.80 | A>A |
| chr18 | 78000000 | 78077248 | 0.00 | 0.00 | NO |
| chr19 | 0 | 100000 | 0.00 | 0.00 | NO |
| chr19 | 100000 | 200000 | 0.00 | 0.00 | NO |
| chr19 | 200000 | 300000 | 0.00 | 0.00 | NO |
| chr19 | 300000 | 400000 | 0.00 | 0.00 | NO |
| chr19 | 400000 | 500000 | 0.00 | 1.10 | NO |
| chr19 | 500000 | 600000 | 0.00 | 1.56 | NO |
| chr19 | 600000 | 700000 | 0.00 | 1.51 | NO |
| chr19 | 700000 | 800000 | 0.00 | 1.20 | NO |
| chr19 | 800000 | 900000 | 0.00 | 1.26 | NO |
| chr19 | 900000 | 1000000 | 0.00 | 1.18 | NO |
| chr19 | 1000000 | 1100000 | 0.00 | 0.97 | NO |
| chr19 | 1100000 | 1200000 | 0.00 | 0.00 | NO |
| chr19 | 1200000 | 1300000 | 0.00 | 1.02 | NO |
| chr19 | 1300000 | 1400000 | 1.15 | 1.41 | A>A |
| chr19 | 1400000 | 1500000 | 0.00 | 1.33 | NO |
| chr19 | 1500000 | 1600000 | 0.84 | 1.18 | A>A |
| chr19 | 1600000 | 1700000 | 1.04 | 1.12 | A>A |
| chr19 | 1700000 | 1800000 | 0.94 | 1.14 | A>A |
| chr19 | 1800000 | 1900000 | 0.85 | 1.06 | A>A |
| chr19 | 1900000 | 2000000 | 0.91 | 1.03 | A>A |
| chr19 | 2000000 | 2100000 | 0.00 | 1.01 | NO |
| chr19 | 2100000 | 2200000 | 0.74 | 0.96 | A>A |
| chr19 | 2200000 | 2300000 | 0.00 | 0.95 | NO |
| chr19 | 2300000 | 2400000 | 0.65 | 0.67 | A>A |
| chr19 | 2400000 | 2500000 | 0.75 | 0.70 | A>A |
| chr19 | 2500000 | 2600000 | 0.78 | 0.91 | A>A |
| chr19 | 2600000 | 2700000 | 0.47 | 0.59 | A>A |
| chr19 | 2700000 | 2800000 | 0.70 | 0.86 | A>A |
| chr19 | 2800000 | 2900000 | 0.00 | 0.85 | NO |
| chr19 | 2900000 | 3000000 | 0.00 | 0.73 | NO |
| chr19 | 3000000 | 3100000 | 0.78 | 1.00 | A>A |
| chr19 | 3100000 | 3200000 | 0.76 | 0.72 | A>A |
| chr19 | 3200000 | 3300000 | 0.56 | 0.70 | A>A |
| chr19 | 3300000 | 3400000 | 0.65 | 0.83 | A>A |
| chr19 | 3400000 | 3500000 | 0.64 | 0.78 | A>A |
| chr19 | 3500000 | 3600000 | 0.70 | 0.88 | A>A |

| | | | | | |
|---|---|---|---|---|---|
| chr19 | 3600000 | 3700000 | 0.69 | 0.83 | A>A |
| chr19 | 3700000 | 3800000 | 0.66 | 0.86 | A>A |
| chr19 | 3800000 | 3900000 | 0.00 | 0.70 | NO |
| chr19 | 3900000 | 4000000 | 0.69 | 0.75 | A>A |
| chr19 | 4000000 | 4100000 | 0.77 | 0.80 | A>A |
| chr19 | 4100000 | 4200000 | 0.77 | 0.85 | A>A |
| chr19 | 4200000 | 4300000 | 0.68 | 0.71 | A>A |
| chr19 | 4300000 | 4400000 | 0.86 | 0.92 | A>A |
| chr19 | 4400000 | 4500000 | 0.71 | 0.86 | A>A |
| chr19 | 4500000 | 4600000 | 0.00 | 0.83 | NO |
| chr19 | 4600000 | 4700000 | 0.76 | 0.78 | A>A |
| chr19 | 4700000 | 4800000 | 0.60 | 0.60 | A>A |
| chr19 | 4800000 | 4900000 | 0.00 | 0.49 | NO |
| chr19 | 4900000 | 5000000 | 0.69 | 0.49 | A>A |
| chr19 | 5000000 | 5100000 | 0.44 | 0.29 | A>A |
| chr19 | 5100000 | 5200000 | 0.56 | 0.35 | A>A |
| chr19 | 5200000 | 5300000 | 0.48 | 0.39 | A>A |
| chr19 | 5300000 | 5400000 | 0.46 | 0.30 | A>A |
| chr19 | 5400000 | 5500000 | 0.47 | 0.29 | A>A |
| chr19 | 5500000 | 5600000 | 0.45 | 0.41 | A>A |
| chr19 | 5600000 | 5700000 | 0.75 | 0.81 | A>A |
| chr19 | 5700000 | 5800000 | 0.82 | 0.69 | A>A |
| chr19 | 5800000 | 5900000 | 0.00 | 0.68 | NO |
| chr19 | 5900000 | 6000000 | 0.72 | 0.46 | A>A |
| chr19 | 6000000 | 6100000 | 0.80 | 0.43 | A>A |
| chr19 | 6100000 | 6200000 | 0.00 | 0.43 | NO |
| chr19 | 6200000 | 6300000 | 0.00 | 0.51 | NO |
| chr19 | 6300000 | 6400000 | 0.00 | 0.55 | NO |
| chr19 | 6400000 | 6500000 | 0.00 | 0.69 | NO |
| chr19 | 6500000 | 6600000 | 0.00 | 0.65 | NO |
| chr19 | 6600000 | 6700000 | 0.00 | 0.00 | NO |
| chr19 | 6700000 | 6800000 | 0.00 | 0.69 | NO |
| chr19 | 6800000 | 6900000 | 0.00 | 0.00 | NO |
| chr19 | 6900000 | 7000000 | 0.00 | 0.35 | NO |
| chr19 | 7000000 | 7100000 | 0.00 | 0.00 | NO |
| chr19 | 7100000 | 7200000 | 0.00 | 0.40 | NO |
| chr19 | 7200000 | 7300000 | 0.00 | 0.44 | NO |
| chr19 | 7300000 | 7400000 | 0.00 | 0.00 | NO |
| chr19 | 7400000 | 7500000 | 0.69 | 0.38 | A>A |
| chr19 | 7500000 | 7600000 | 1.08 | 0.72 | A>A |
| chr19 | 7600000 | 7700000 | 0.92 | 0.80 | A>A |
| chr19 | 7700000 | 7800000 | 0.00 | 0.69 | NO |
| chr19 | 7800000 | 7900000 | 0.00 | 0.00 | NO |
| chr19 | 7900000 | 8000000 | 1.11 | 0.77 | A>A |
| chr19 | 8000000 | 8100000 | 1.01 | 0.70 | A>A |
| chr19 | 8100000 | 8200000 | 0.00 | 0.69 | NO |
| chr19 | 8200000 | 8300000 | 0.00 | 0.64 | NO |
| chr19 | 8300000 | 8400000 | 0.00 | 0.00 | NO |
| chr19 | 8400000 | 8500000 | 0.00 | 0.00 | NO |
| chr19 | 8500000 | 8600000 | 0.00 | 0.90 | NO |
| chr19 | 8600000 | 8700000 | 0.00 | 0.00 | NO |
| chr19 | 8700000 | 8800000 | 0.00 | 0.00 | NO |
| chr19 | 8800000 | 8900000 | 0.00 | 0.00 | NO |
| chr19 | 8900000 | 9000000 | 0.00 | 0.00 | NO |
| chr19 | 9000000 | 9100000 | 0.00 | 0.00 | NO |
| chr19 | 9100000 | 9200000 | 0.00 | 0.00 | NO |

| | | | | | |
|---|---|---|---|---|---|
| chr19 | 9200000 | 9300000 | 0.00 | 0.45 | NO |
| chr19 | 9300000 | 9400000 | 0.00 | 0.00 | NO |
| chr19 | 9400000 | 9500000 | 0.00 | 0.31 | NO |
| chr19 | 9500000 | 9600000 | 0.00 | 0.00 | NO |
| chr19 | 9600000 | 9700000 | 0.00 | 0.00 | NO |
| chr19 | 9700000 | 9800000 | 0.00 | 0.00 | NO |
| chr19 | 9800000 | 9900000 | 0.00 | 0.00 | NO |
| chr19 | 9900000 | 10000000 | 0.47 | 0.69 | A>A |
| chr19 | 10000000 | 10100000 | 0.00 | 0.57 | NO |
| chr19 | 10100000 | 10200000 | 0.00 | 0.65 | NO |
| chr19 | 10200000 | 10300000 | 0.00 | 0.84 | NO |
| chr19 | 10300000 | 10400000 | 0.00 | 0.71 | NO |
| chr19 | 10400000 | 10500000 | 0.00 | 0.94 | NO |
| chr19 | 10500000 | 10600000 | 0.00 | 0.74 | NO |
| chr19 | 10600000 | 10700000 | 0.00 | 0.99 | NO |
| chr19 | 10700000 | 10800000 | 0.00 | 0.95 | NO |
| chr19 | 10800000 | 10900000 | 1.09 | 0.85 | A>A |
| chr19 | 10900000 | 11000000 | 1.05 | 0.91 | A>A |
| chr19 | 11000000 | 11100000 | 0.97 | 0.95 | A>A |
| chr19 | 11100000 | 11200000 | 0.00 | 0.92 | NO |
| chr19 | 11200000 | 11300000 | 0.88 | 0.99 | A>A |
| chr19 | 11300000 | 11400000 | 0.91 | 0.81 | A>A |
| chr19 | 11400000 | 11500000 | 0.71 | 1.08 | A>A |
| chr19 | 11500000 | 11600000 | 0.00 | 0.97 | NO |
| chr19 | 11600000 | 11700000 | 0.35 | 0.80 | A>A |
| chr19 | 11700000 | 11800000 | 0.00 | 0.00 | NO |
| chr19 | 11800000 | 11900000 | 0.00 | 0.00 | NO |
| chr19 | 11900000 | 12000000 | 0.00 | 0.00 | NO |
| chr19 | 12000000 | 12100000 | 0.00 | 0.00 | NO |
| chr19 | 12100000 | 12200000 | 0.00 | 0.00 | NO |
| chr19 | 12200000 | 12300000 | 0.00 | 0.00 | NO |
| chr19 | 12300000 | 12400000 | 0.00 | 0.00 | NO |
| chr19 | 12400000 | 12500000 | 0.00 | 0.79 | NO |
| chr19 | 12500000 | 12600000 | 0.00 | 0.00 | NO |
| chr19 | 12600000 | 12700000 | 0.00 | 0.00 | NO |
| chr19 | 12700000 | 12800000 | -0.22 | 1.19 | B>A |
| chr19 | 12800000 | 12900000 | 0.97 | 1.49 | B>A |
| chr19 | 12900000 | 13000000 | 0.93 | 1.37 | A>A |
| chr19 | 13000000 | 13100000 | 0.00 | 0.98 | NO |
| chr19 | 13100000 | 13200000 | 0.92 | 1.08 | A>A |
| chr19 | 13200000 | 13300000 | 0.96 | 0.79 | A>A |
| chr19 | 13300000 | 13400000 | 0.80 | 0.39 | A>A |
| chr19 | 13400000 | 13500000 | 0.50 | 0.10 | A>A |
| chr19 | 13500000 | 13600000 | 0.00 | 0.20 | NO |
| chr19 | 13600000 | 13700000 | 0.00 | 0.08 | NO |
| chr19 | 13700000 | 13800000 | 0.00 | 0.34 | NO |
| chr19 | 13800000 | 13900000 | 0.00 | 0.53 | NO |
| chr19 | 13900000 | 14000000 | 0.99 | 0.79 | A>A |
| chr19 | 14000000 | 14100000 | 1.05 | 1.02 | A>A |
| chr19 | 14100000 | 14200000 | 0.00 | 0.88 | NO |
| chr19 | 14200000 | 14300000 | 0.81 | 0.66 | A>A |
| chr19 | 14300000 | 14400000 | 0.00 | 0.00 | NO |
| chr19 | 14400000 | 14500000 | 0.00 | 0.71 | NO |
| chr19 | 14500000 | 14600000 | 0.98 | 0.93 | A>A |
| chr19 | 14600000 | 14700000 | 0.92 | 0.71 | A>A |
| chr19 | 14700000 | 14800000 | 0.00 | 0.10 | NO |

| | | | | | |
|---|---|---|---|---|---|
| chr19 | 14800000 | 14900000 | 0.00 | -0.01 | NO |
| chr19 | 14900000 | 15000000 | 0.00 | -0.11 | NO |
| chr19 | 15000000 | 15100000 | 0.00 | -0.16 | NO |
| chr19 | 15100000 | 15200000 | 0.00 | -0.15 | NO |
| chr19 | 15200000 | 15300000 | 0.71 | 0.21 | A>A |
| chr19 | 15300000 | 15400000 | 0.83 | 0.54 | A>A |
| chr19 | 15400000 | 15500000 | 0.00 | 0.69 | NO |
| chr19 | 15500000 | 15600000 | 0.85 | 0.57 | A>A |
| chr19 | 15600000 | 15700000 | 0.69 | 0.10 | A>B |
| chr19 | 15700000 | 15800000 | 0.00 | 0.05 | NO |
| chr19 | 15800000 | 15900000 | 0.00 | -0.06 | NO |
| chr19 | 15900000 | 16000000 | 0.00 | -0.06 | NO |
| chr19 | 16000000 | 16100000 | 0.00 | -0.04 | NO |
| chr19 | 16100000 | 16200000 | 0.69 | 0.25 | A>A |
| chr19 | 16200000 | 16300000 | 0.00 | 0.46 | NO |
| chr19 | 16300000 | 16400000 | 0.71 | 0.44 | A>A |
| chr19 | 16400000 | 16500000 | 0.89 | 0.62 | A>A |
| chr19 | 16500000 | 16600000 | 0.77 | 0.68 | A>A |
| chr19 | 16600000 | 16700000 | 0.94 | 0.88 | A>A |
| chr19 | 16700000 | 16800000 | 0.88 | 0.69 | A>A |
| chr19 | 16800000 | 16900000 | 0.60 | 0.42 | A>A |
| chr19 | 16900000 | 17000000 | 0.00 | 0.77 | NO |
| chr19 | 17000000 | 17100000 | 0.00 | 0.57 | NO |
| chr19 | 17100000 | 17200000 | 0.63 | 0.58 | A>A |
| chr19 | 17200000 | 17300000 | 0.82 | 0.92 | A>A |
| chr19 | 17300000 | 17400000 | 0.96 | 1.19 | A>A |
| chr19 | 17400000 | 17500000 | 1.01 | 1.21 | A>A |
| chr19 | 17500000 | 17600000 | 0.91 | 1.11 | A>A |
| chr19 | 17600000 | 17700000 | 0.94 | 1.23 | A>A |
| chr19 | 17700000 | 17800000 | 0.00 | 0.96 | NO |
| chr19 | 17800000 | 17900000 | 0.00 | 1.06 | NO |
| chr19 | 17900000 | 18000000 | 1.00 | 1.02 | A>A |
| chr19 | 18000000 | 18100000 | 0.00 | 1.21 | NO |
| chr19 | 18100000 | 18200000 | 0.00 | 1.25 | NO |
| chr19 | 18200000 | 18300000 | 1.28 | 1.41 | A>A |
| chr19 | 18300000 | 18400000 | 1.19 | 1.40 | A>A |
| chr19 | 18400000 | 18500000 | 0.00 | 1.48 | NO |
| chr19 | 18500000 | 18600000 | 0.98 | 1.28 | A>A |
| chr19 | 18600000 | 18700000 | 1.11 | 1.42 | A>A |
| chr19 | 18700000 | 18800000 | 0.97 | 1.17 | A>A |
| chr19 | 18800000 | 18900000 | 1.05 | 1.30 | A>A |
| chr19 | 18900000 | 19000000 | 0.83 | 1.05 | A>A |
| chr19 | 19000000 | 19100000 | 0.00 | 1.38 | NO |
| chr19 | 19100000 | 19200000 | 0.90 | 1.30 | A>A |
| chr19 | 19200000 | 19300000 | 0.76 | 1.18 | A>A |
| chr19 | 19300000 | 19400000 | 0.79 | 1.22 | A>A |
| chr19 | 19400000 | 19500000 | 0.67 | 1.20 | B>A |
| chr19 | 19500000 | 19600000 | 0.38 | 1.23 | B>A |
| chr19 | 19600000 | 19700000 | 0.09 | 1.32 | B>A |
| chr19 | 19700000 | 19800000 | -0.25 | 1.34 | B>A |
| chr19 | 19800000 | 19900000 | 0.00 | 0.00 | NO |
| chr19 | 19900000 | 20000000 | 0.00 | 0.00 | NO |
| chr19 | 20000000 | 20100000 | 0.00 | 0.00 | NO |
| chr19 | 20100000 | 20200000 | 0.00 | 0.00 | NO |
| chr19 | 20200000 | 20300000 | 0.00 | 0.00 | NO |
| chr19 | 20300000 | 20400000 | 0.00 | 0.00 | NO |

| | | | | | |
|---|---|---|---|---|---|
| chr19 | 20400000 | 20500000 | 0.00 | 0.00 | NO |
| chr19 | 20500000 | 20600000 | 0.00 | 0.00 | NO |
| chr19 | 20600000 | 20700000 | -1.19 | 0.13 | B>A |
| chr19 | 20700000 | 20800000 | 0.00 | 0.00 | NO |
| chr19 | 20800000 | 20900000 | 0.00 | 0.00 | NO |
| chr19 | 20900000 | 21000000 | 0.00 | 0.00 | NO |
| chr19 | 21000000 | 21100000 | 0.00 | 0.00 | NO |
| chr19 | 21100000 | 21200000 | 0.00 | 0.00 | NO |
| chr19 | 21200000 | 21300000 | 0.00 | 0.00 | NO |
| chr19 | 21300000 | 21400000 | 0.00 | 0.00 | NO |
| chr19 | 21400000 | 21500000 | -0.75 | -0.03 | B>A |
| chr19 | 21500000 | 21600000 | 0.00 | 0.00 | NO |
| chr19 | 21600000 | 21700000 | 0.00 | 0.00 | NO |
| chr19 | 21700000 | 21800000 | 0.00 | 0.00 | NO |
| chr19 | 21800000 | 21900000 | 0.00 | 0.00 | NO |
| chr19 | 21900000 | 22000000 | 0.00 | 0.00 | NO |
| chr19 | 22000000 | 22100000 | -1.21 | -0.36 | B>A |
| chr19 | 22100000 | 22200000 | -1.30 | -0.50 | B>A |
| chr19 | 22200000 | 22300000 | -1.26 | -0.38 | B>A |
| chr19 | 22300000 | 22400000 | -1.24 | -0.44 | B>A |
| chr19 | 22400000 | 22500000 | -1.11 | -0.48 | B>A |
| chr19 | 22500000 | 22600000 | 0.00 | -0.43 | NO |
| chr19 | 22600000 | 22700000 | -1.13 | -0.46 | B>A |
| chr19 | 22700000 | 22800000 | -1.09 | -0.42 | B>A |
| chr19 | 22800000 | 22900000 | -1.12 | -0.36 | B>A |
| chr19 | 22900000 | 23000000 | -1.09 | -0.28 | B>A |
| chr19 | 23000000 | 23100000 | -1.06 | -0.32 | B>A |
| chr19 | 23100000 | 23200000 | -0.98 | -0.26 | B>A |
| chr19 | 23200000 | 23300000 | -1.03 | -0.21 | B>A |
| chr19 | 23300000 | 23400000 | -1.00 | -0.12 | B>A |
| chr19 | 23400000 | 23500000 | -0.96 | -0.18 | B>A |
| chr19 | 23500000 | 23600000 | -0.95 | -0.16 | B>A |
| chr19 | 23600000 | 23700000 | -0.87 | -0.21 | B>A |
| chr19 | 23700000 | 23800000 | -0.91 | -0.12 | B>A |
| chr19 | 23800000 | 23900000 | -0.94 | -0.09 | B>A |
| chr19 | 23900000 | 24000000 | -0.93 | -0.12 | B>A |
| chr19 | 24000000 | 24100000 | -1.00 | -0.30 | B>A |
| chr19 | 24100000 | 24200000 | -0.94 | -0.23 | B>A |
| chr19 | 24200000 | 24300000 | -0.82 | -0.36 | B>B |
| chr19 | 24300000 | 24400000 | -0.97 | -0.53 | B>B |
| chr19 | 24400000 | 24500000 | 0.00 | 0.00 | NO |
| chr19 | 24500000 | 24600000 | -0.84 | -0.63 | B>B |
| chr19 | 24600000 | 24700000 | 0.00 | 0.00 | NO |
| chr19 | 24700000 | 24800000 | 0.00 | 0.00 | NO |
| chr19 | 24800000 | 24900000 | 0.00 | 0.00 | NO |
| chr19 | 24900000 | 25000000 | 0.00 | 0.00 | NO |
| chr19 | 25000000 | 25100000 | 0.00 | 0.00 | NO |
| chr19 | 25100000 | 25200000 | 0.00 | 0.00 | NO |
| chr19 | 25200000 | 25300000 | 0.00 | 0.00 | NO |
| chr19 | 25300000 | 25400000 | 0.00 | 0.00 | NO |
| chr19 | 25400000 | 25500000 | 0.00 | 0.00 | NO |
| chr19 | 25500000 | 25600000 | 0.00 | 0.00 | NO |
| chr19 | 25600000 | 25700000 | 0.00 | 0.00 | NO |
| chr19 | 25700000 | 25800000 | 0.00 | 0.00 | NO |
| chr19 | 25800000 | 25900000 | 0.00 | 0.00 | NO |
| chr19 | 25900000 | 26000000 | 0.00 | 0.00 | NO |

| | | | | | |
|---|---|---|---|---|---|
| chr19 | 26000000 | 26100000 | 0.00 | 0.00 | NO |
| chr19 | 26100000 | 26200000 | 0.00 | 0.00 | NO |
| chr19 | 26200000 | 26300000 | 0.00 | 0.00 | NO |
| chr19 | 26300000 | 26400000 | 0.00 | 0.00 | NO |
| chr19 | 26400000 | 26500000 | 0.00 | 0.00 | NO |
| chr19 | 26500000 | 26600000 | 0.00 | 0.00 | NO |
| chr19 | 26600000 | 26700000 | 0.00 | 0.00 | NO |
| chr19 | 26700000 | 26800000 | 0.00 | 0.00 | NO |
| chr19 | 26800000 | 26900000 | 0.00 | 0.00 | NO |
| chr19 | 26900000 | 27000000 | 0.00 | 0.00 | NO |
| chr19 | 27000000 | 27100000 | 0.00 | 0.00 | NO |
| chr19 | 27100000 | 27200000 | 0.00 | 0.00 | NO |
| chr19 | 27200000 | 27300000 | 0.00 | 0.00 | NO |
| chr19 | 27300000 | 27400000 | 0.00 | 0.00 | NO |
| chr19 | 27400000 | 27500000 | 0.00 | 0.00 | NO |
| chr19 | 27500000 | 27600000 | 0.00 | 0.00 | NO |
| chr19 | 27600000 | 27700000 | 0.00 | 0.00 | NO |
| chr19 | 27700000 | 27800000 | 0.00 | 0.00 | NO |
| chr19 | 27800000 | 27900000 | 0.00 | 0.00 | NO |
| chr19 | 27900000 | 28000000 | 0.00 | -1.02 | NO |
| chr19 | 28000000 | 28100000 | -1.26 | -1.10 | B>B |
| chr19 | 28100000 | 28200000 | -1.30 | -1.10 | B>B |
| chr19 | 28200000 | 28300000 | -1.11 | -1.01 | B>B |
| chr19 | 28300000 | 28400000 | -1.00 | -1.10 | B>B |
| chr19 | 28400000 | 28500000 | -0.99 | -1.08 | B>B |
| chr19 | 28500000 | 28600000 | -1.04 | -1.07 | B>B |
| chr19 | 28600000 | 28700000 | -1.05 | -1.04 | B>B |
| chr19 | 28700000 | 28800000 | -1.06 | -1.02 | B>B |
| chr19 | 28800000 | 28900000 | -0.95 | -1.02 | B>B |
| chr19 | 28900000 | 29000000 | -1.08 | -0.98 | B>B |
| chr19 | 29000000 | 29100000 | -1.09 | -0.99 | B>B |
| chr19 | 29100000 | 29200000 | -1.00 | -0.96 | B>B |
| chr19 | 29200000 | 29300000 | -0.96 | -0.97 | B>B |
| chr19 | 29300000 | 29400000 | -0.94 | -1.00 | B>B |
| chr19 | 29400000 | 29500000 | -0.92 | -0.93 | B>B |
| chr19 | 29500000 | 29600000 | -0.95 | -0.98 | B>B |
| chr19 | 29600000 | 29700000 | -0.74 | -0.92 | B>B |
| chr19 | 29700000 | 29800000 | -0.65 | -0.94 | B>B |
| chr19 | 29800000 | 29900000 | -0.64 | -1.00 | B>B |
| chr19 | 29900000 | 30000000 | -0.57 | -0.96 | B>B |
| chr19 | 30000000 | 30100000 | -0.46 | -0.87 | B>B |
| chr19 | 30100000 | 30200000 | -0.17 | -0.76 | A>B |
| chr19 | 30200000 | 30300000 | -0.01 | -0.68 | A>B |
| chr19 | 30300000 | 30400000 | 0.00 | -0.62 | A>B |
| chr19 | 30400000 | 30500000 | -0.02 | -0.61 | B>B |
| chr19 | 30500000 | 30600000 | -0.15 | -0.76 | A>B |
| chr19 | 30600000 | 30700000 | -0.24 | -0.85 | A>B |
| chr19 | 30700000 | 30800000 | -0.34 | -0.91 | B>B |
| chr19 | 30800000 | 30900000 | -0.34 | -0.98 | A>B |
| chr19 | 30900000 | 31000000 | -0.43 | -1.02 | A>B |
| chr19 | 31000000 | 31100000 | -0.56 | -1.07 | B>B |
| chr19 | 31100000 | 31200000 | -0.67 | -1.16 | B>B |
| chr19 | 31200000 | 31300000 | -0.96 | -1.18 | B>B |
| chr19 | 31300000 | 31400000 | -0.92 | -1.10 | B>B |
| chr19 | 31400000 | 31500000 | -0.72 | -1.00 | B>B |
| chr19 | 31500000 | 31600000 | -0.77 | -0.98 | B>B |

| | | | | | |
|---|---|---|---|---|---|
| chr19 | 31600000 | 31700000 | -0.71 | -0.97 | B>B |
| chr19 | 31700000 | 31800000 | -0.68 | -0.95 | B>B |
| chr19 | 31800000 | 31900000 | -0.62 | -0.95 | B>B |
| chr19 | 31900000 | 32000000 | -0.78 | -0.96 | B>B |
| chr19 | 32000000 | 32100000 | -0.81 | -0.98 | B>B |
| chr19 | 32100000 | 32200000 | -0.84 | -1.00 | B>B |
| chr19 | 32200000 | 32300000 | -0.87 | -1.03 | B>B |
| chr19 | 32300000 | 32400000 | -0.79 | -1.07 | B>B |
| chr19 | 32400000 | 32500000 | -0.75 | -1.08 | B>B |
| chr19 | 32500000 | 32600000 | -0.65 | -1.13 | B>B |
| chr19 | 32600000 | 32700000 | -0.56 | -1.15 | A>B |
| chr19 | 32700000 | 32800000 | -0.36 | -1.11 | A>B |
| chr19 | 32800000 | 32900000 | -0.07 | -0.84 | A>B |
| chr19 | 32900000 | 33000000 | -0.09 | -0.80 | A>B |
| chr19 | 33000000 | 33100000 | -0.10 | -0.82 | A>B |
| chr19 | 33100000 | 33200000 | 0.24 | -0.41 | A>B |
| chr19 | 33200000 | 33300000 | 0.19 | -0.47 | A>B |
| chr19 | 33300000 | 33400000 | 0.16 | -0.39 | B>B |
| chr19 | 33400000 | 33500000 | 0.30 | -0.26 | A>A |
| chr19 | 33500000 | 33600000 | 0.45 | 0.07 | A>A |
| chr19 | 33600000 | 33700000 | 0.35 | -0.11 | A>A |
| chr19 | 33700000 | 33800000 | 0.24 | -0.28 | B>B |
| chr19 | 33800000 | 33900000 | 0.24 | -0.37 | A>B |
| chr19 | 33900000 | 34000000 | 0.03 | -0.51 | B>B |
| chr19 | 34000000 | 34100000 | -0.02 | -0.65 | A>B |
| chr19 | 34100000 | 34200000 | -0.08 | -0.69 | A>B |
| chr19 | 34200000 | 34300000 | -0.03 | -0.58 | B>B |
| chr19 | 34300000 | 34400000 | -0.04 | -0.55 | B>B |
| chr19 | 34400000 | 34500000 | -0.14 | -0.64 | B>B |
| chr19 | 34500000 | 34600000 | -0.10 | -0.57 | B>B |
| chr19 | 34600000 | 34700000 | 0.18 | -0.25 | B>B |
| chr19 | 34700000 | 34800000 | 0.29 | 0.00 | A>A |
| chr19 | 34800000 | 34900000 | 0.37 | 0.19 | A>A |
| chr19 | 34900000 | 35000000 | 0.12 | 0.19 | A>A |
| chr19 | 35000000 | 35100000 | -0.52 | -0.07 | B>B |
| chr19 | 35100000 | 35200000 | -0.67 | -0.10 | B>A |
| chr19 | 35200000 | 35300000 | -0.83 | 0.05 | B>A |
| chr19 | 35300000 | 35400000 | -0.70 | 0.05 | B>A |
| chr19 | 35400000 | 35500000 | -0.22 | 0.36 | B>A |
| chr19 | 35500000 | 35600000 | 0.18 | 0.37 | A>A |
| chr19 | 35600000 | 35700000 | 0.21 | 0.43 | A>A |
| chr19 | 35700000 | 35800000 | 0.32 | 0.56 | A>A |
| chr19 | 35800000 | 35900000 | 0.40 | 0.48 | A>A |
| chr19 | 35900000 | 36000000 | 0.46 | 0.44 | A>A |
| chr19 | 36000000 | 36100000 | 0.58 | 0.56 | A>A |
| chr19 | 36100000 | 36200000 | 0.76 | 0.78 | A>A |
| chr19 | 36200000 | 36300000 | 0.91 | 0.91 | A>A |
| chr19 | 36300000 | 36400000 | 0.72 | 0.74 | A>A |
| chr19 | 36400000 | 36500000 | 0.58 | 0.53 | A>A |
| chr19 | 36500000 | 36600000 | 0.54 | 0.60 | A>A |
| chr19 | 36600000 | 36700000 | 0.07 | 0.51 | A>A |
| chr19 | 36700000 | 36800000 | -0.78 | 0.10 | B>A |
| chr19 | 36800000 | 36900000 | -1.16 | 0.11 | B>A |
| chr19 | 36900000 | 37000000 | -1.04 | 0.17 | B>A |
| chr19 | 37000000 | 37100000 | -1.18 | 0.18 | B>A |
| chr19 | 37100000 | 37200000 | -1.16 | 0.22 | B>A |

| | | | | | |
|---|---|---|---|---|---|
| chr19 | 37200000 | 37300000 | -1.43 | 0.27 | B>A |
| chr19 | 37300000 | 37400000 | -1.59 | 0.18 | B>A |
| chr19 | 37400000 | 37500000 | -1.71 | 0.01 | B>A |
| chr19 | 37500000 | 37600000 | -1.61 | 0.10 | B>A |
| chr19 | 37600000 | 37700000 | -1.43 | 0.10 | B>A |
| chr19 | 37700000 | 37800000 | -1.24 | 0.00 | B>A |
| chr19 | 37800000 | 37900000 | -1.12 | 0.07 | B>A |
| chr19 | 37900000 | 38000000 | -1.35 | 0.13 | B>A |
| chr19 | 38000000 | 38100000 | -1.49 | 0.23 | B>A |
| chr19 | 38100000 | 38200000 | -1.26 | 0.16 | B>A |
| chr19 | 38200000 | 38300000 | -0.99 | 0.13 | B>A |
| chr19 | 38300000 | 38400000 | -0.59 | 0.11 | B>A |
| chr19 | 38400000 | 38500000 | 0.05 | 0.04 | A>A |
| chr19 | 38500000 | 38600000 | -0.05 | 0.02 | B>B |
| chr19 | 38600000 | 38700000 | 0.08 | 0.17 | A>A |
| chr19 | 38700000 | 38800000 | 0.42 | 0.59 | A>A |
| chr19 | 38800000 | 38900000 | 0.47 | 0.55 | A>A |
| chr19 | 38900000 | 39000000 | 0.49 | 0.35 | A>A |
| chr19 | 39000000 | 39100000 | 0.49 | 0.44 | A>A |
| chr19 | 39100000 | 39200000 | 0.52 | 0.56 | A>A |
| chr19 | 39200000 | 39300000 | 0.52 | 0.55 | A>A |
| chr19 | 39300000 | 39400000 | 0.61 | 0.78 | A>A |
| chr19 | 39400000 | 39500000 | 0.52 | 0.52 | A>A |
| chr19 | 39500000 | 39600000 | 0.59 | 0.51 | A>A |
| chr19 | 39600000 | 39700000 | 0.51 | 0.53 | A>A |
| chr19 | 39700000 | 39800000 | 0.44 | 0.44 | A>A |
| chr19 | 39800000 | 39900000 | 0.66 | 0.65 | A>A |
| chr19 | 39900000 | 40000000 | 0.70 | 0.79 | A>A |
| chr19 | 40000000 | 40100000 | 0.30 | 0.28 | A>A |
| chr19 | 40100000 | 40200000 | 0.17 | 0.03 | A>A |
| chr19 | 40200000 | 40300000 | 0.22 | 0.08 | A>A |
| chr19 | 40300000 | 40400000 | 0.39 | 0.29 | A>A |
| chr19 | 40400000 | 40500000 | 0.49 | 0.50 | A>A |
| chr19 | 40500000 | 40600000 | 0.13 | 0.18 | A>A |
| chr19 | 40600000 | 40700000 | 0.33 | 0.27 | A>A |
| chr19 | 40700000 | 40800000 | 0.49 | 0.41 | A>A |
| chr19 | 40800000 | 40900000 | 0.55 | 0.52 | A>A |
| chr19 | 40900000 | 41000000 | 0.71 | 0.74 | A>A |
| chr19 | 41000000 | 41100000 | 0.60 | 0.68 | A>A |
| chr19 | 41100000 | 41200000 | 0.61 | 0.55 | A>A |
| chr19 | 41200000 | 41300000 | 0.64 | 0.62 | A>A |
| chr19 | 41300000 | 41400000 | 0.42 | 0.25 | A>A |
| chr19 | 41400000 | 41500000 | 0.41 | 0.17 | A>A |
| chr19 | 41500000 | 41600000 | 0.36 | 0.12 | A>A |
| chr19 | 41600000 | 41700000 | 0.40 | 0.24 | A>A |
| chr19 | 41700000 | 41800000 | 0.56 | 0.49 | A>A |
| chr19 | 41800000 | 41900000 | 0.51 | 0.51 | A>A |
| chr19 | 41900000 | 42000000 | 0.46 | 0.23 | A>A |
| chr19 | 42000000 | 42100000 | 0.24 | 0.11 | A>A |
| chr19 | 42100000 | 42200000 | 0.21 | 0.01 | A>A |
| chr19 | 42200000 | 42300000 | 0.28 | 0.21 | A>A |
| chr19 | 42300000 | 42400000 | 0.49 | 0.33 | A>A |
| chr19 | 42400000 | 42500000 | 0.54 | 0.37 | A>A |
| chr19 | 42500000 | 42600000 | 0.45 | 0.22 | A>A |
| chr19 | 42600000 | 42700000 | 0.48 | 0.29 | A>A |
| chr19 | 42700000 | 42800000 | 0.71 | 0.53 | A>A |

| | | | | | |
|---|---|---|---|---|---|
| chr19 | 42800000 | 42900000 | 0.58 | 0.32 | A>A |
| chr19 | 42900000 | 43000000 | 0.23 | -0.16 | A>A |
| chr19 | 43000000 | 43100000 | -0.02 | -0.59 | B>B |
| chr19 | 43100000 | 43200000 | -0.35 | -0.97 | A>B |
| chr19 | 43200000 | 43300000 | 0.00 | 0.00 | NO |
| chr19 | 43300000 | 43400000 | 0.00 | 0.00 | NO |
| chr19 | 43400000 | 43500000 | 0.00 | 0.00 | NO |
| chr19 | 43500000 | 43600000 | 0.00 | 0.00 | NO |
| chr19 | 43600000 | 43700000 | 0.00 | 0.00 | NO |
| chr19 | 43700000 | 43800000 | 0.00 | 0.00 | NO |
| chr19 | 43800000 | 43900000 | 0.00 | 0.07 | NO |
| chr19 | 43900000 | 44000000 | 0.32 | 0.46 | A>A |
| chr19 | 44000000 | 44100000 | 0.47 | 0.48 | A>A |
| chr19 | 44100000 | 44200000 | 0.44 | 0.63 | A>A |
| chr19 | 44200000 | 44300000 | 0.36 | 0.65 | A>A |
| chr19 | 44300000 | 44400000 | -0.32 | 0.28 | B>A |
| chr19 | 44400000 | 44500000 | -0.47 | 0.20 | B>A |
| chr19 | 44500000 | 44600000 | -0.31 | 0.32 | B>A |
| chr19 | 44600000 | 44700000 | -0.40 | 0.29 | B>A |
| chr19 | 44700000 | 44800000 | -0.52 | 0.19 | B>A |
| chr19 | 44800000 | 44900000 | -0.49 | 0.13 | B>A |
| chr19 | 44900000 | 45000000 | -0.51 | 0.32 | B>A |
| chr19 | 45000000 | 45100000 | -0.02 | 0.27 | A>A |
| chr19 | 45100000 | 45200000 | 0.41 | 0.55 | A>A |
| chr19 | 45200000 | 45300000 | 0.64 | 0.78 | A>A |
| chr19 | 45300000 | 45400000 | 0.58 | 0.69 | A>A |
| chr19 | 45400000 | 45500000 | 0.67 | 0.77 | A>A |
| chr19 | 45500000 | 45600000 | 0.71 | 0.79 | A>A |
| chr19 | 45600000 | 45700000 | 0.54 | 0.80 | A>A |
| chr19 | 45700000 | 45800000 | 0.83 | 0.90 | A>A |
| chr19 | 45800000 | 45900000 | 0.99 | 1.22 | A>A |
| chr19 | 45900000 | 46000000 | 0.73 | 0.91 | A>A |
| chr19 | 46000000 | 46100000 | 0.76 | 1.05 | A>A |
| chr19 | 46100000 | 46200000 | 0.71 | 0.96 | A>A |
| chr19 | 46200000 | 46300000 | 0.75 | 1.04 | A>A |
| chr19 | 46300000 | 46400000 | 0.74 | 0.91 | A>A |
| chr19 | 46400000 | 46500000 | 0.59 | 0.65 | A>A |
| chr19 | 46500000 | 46600000 | 0.43 | 0.36 | A>A |
| chr19 | 46600000 | 46700000 | 0.42 | 0.40 | A>A |
| chr19 | 46700000 | 46800000 | 0.41 | 0.30 | A>A |
| chr19 | 46800000 | 46900000 | 0.46 | 0.47 | A>A |
| chr19 | 46900000 | 47000000 | 0.40 | 0.41 | A>A |
| chr19 | 47000000 | 47100000 | 0.38 | 0.39 | A>A |
| chr19 | 47100000 | 47200000 | 0.54 | 0.59 | A>A |
| chr19 | 47200000 | 47300000 | 0.55 | 0.64 | A>A |
| chr19 | 47300000 | 47400000 | 0.46 | 0.48 | A>A |
| chr19 | 47400000 | 47500000 | 0.45 | 0.55 | A>A |
| chr19 | 47500000 | 47600000 | 0.48 | 0.51 | A>A |
| chr19 | 47600000 | 47700000 | 0.47 | 0.66 | A>A |
| chr19 | 47700000 | 47800000 | 0.51 | 0.72 | A>A |
| chr19 | 47800000 | 47900000 | 0.53 | 0.77 | A>A |
| chr19 | 47900000 | 48000000 | 0.44 | 0.46 | A>A |
| chr19 | 48000000 | 48100000 | 0.50 | 0.53 | A>A |
| chr19 | 48100000 | 48200000 | 0.50 | 0.53 | A>A |
| chr19 | 48200000 | 48300000 | 0.52 | 0.53 | A>A |
| chr19 | 48300000 | 48400000 | 0.51 | 0.43 | A>A |

| | | | | | |
|---|---|---|---|---|---|
| chr19 | 48400000 | 48500000 | 0.00 | 0.00 | NO |
| chr19 | 48500000 | 48600000 | 0.41 | 0.36 | A>A |
| chr19 | 48600000 | 48700000 | 0.50 | 0.57 | A>A |
| chr19 | 48700000 | 48800000 | 0.58 | 0.58 | A>A |
| chr19 | 48800000 | 48900000 | 0.63 | 0.75 | A>A |
| chr19 | 48900000 | 49000000 | 0.70 | 0.91 | A>A |
| chr19 | 49000000 | 49100000 | 0.65 | 0.89 | A>A |
| chr19 | 49100000 | 49200000 | 0.60 | 0.85 | A>A |
| chr19 | 49200000 | 49300000 | 0.60 | 0.70 | A>A |
| chr19 | 49300000 | 49400000 | 0.63 | 0.80 | A>A |
| chr19 | 49400000 | 49500000 | 0.70 | 0.80 | A>A |
| chr19 | 49500000 | 49600000 | 0.75 | 0.85 | A>A |
| chr19 | 49600000 | 49700000 | 0.74 | 0.83 | A>A |
| chr19 | 49700000 | 49800000 | 0.59 | 0.59 | A>A |
| chr19 | 49800000 | 49900000 | 0.72 | 0.79 | A>A |
| chr19 | 49900000 | 50000000 | 0.78 | 0.92 | A>A |
| chr19 | 50000000 | 50100000 | 0.73 | 1.01 | A>A |
| chr19 | 50100000 | 50200000 | 0.88 | 1.07 | A>A |
| chr19 | 50200000 | 50300000 | 0.83 | 0.90 | A>A |
| chr19 | 50300000 | 50400000 | 0.90 | 1.12 | A>A |
| chr19 | 50400000 | 50500000 | 0.70 | 0.80 | A>A |
| chr19 | 50500000 | 50600000 | 0.63 | 0.84 | A>A |
| chr19 | 50600000 | 50700000 | 0.55 | 0.42 | A>A |
| chr19 | 50700000 | 50800000 | 0.62 | 0.53 | A>A |
| chr19 | 50800000 | 50900000 | 0.61 | 0.53 | A>A |
| chr19 | 50900000 | 51000000 | 0.68 | 0.55 | A>A |
| chr19 | 51000000 | 51100000 | 0.61 | 0.33 | A>A |
| chr19 | 51100000 | 51200000 | 0.49 | 0.17 | A>A |
| chr19 | 51200000 | 51300000 | 0.48 | 0.19 | A>A |
| chr19 | 51300000 | 51400000 | 0.45 | -0.03 | A>A |
| chr19 | 51400000 | 51500000 | 0.37 | -0.09 | A>A |
| chr19 | 51500000 | 51600000 | 0.27 | -0.11 | A>A |
| chr19 | 51600000 | 51700000 | 0.02 | -0.23 | B>B |
| chr19 | 51700000 | 51800000 | -0.16 | -0.28 | B>B |
| chr19 | 51800000 | 51900000 | 0.04 | -0.02 | A>A |
| chr19 | 51900000 | 52000000 | -0.35 | -0.12 | B>B |
| chr19 | 52000000 | 52100000 | -0.47 | -0.12 | B>B |
| chr19 | 52100000 | 52200000 | -0.40 | -0.03 | B>B |
| chr19 | 52200000 | 52300000 | -0.60 | 0.24 | B>A |
| chr19 | 52300000 | 52400000 | -1.21 | 0.04 | B>A |
| chr19 | 52400000 | 52500000 | -1.21 | 0.32 | B>A |
| chr19 | 52500000 | 52600000 | -1.01 | 0.31 | B>A |
| chr19 | 52600000 | 52700000 | -1.01 | 0.31 | B>A |
| chr19 | 52700000 | 52800000 | -0.67 | 0.18 | B>A |
| chr19 | 52800000 | 52900000 | -1.15 | 0.27 | B>A |
| chr19 | 52900000 | 53000000 | -1.36 | 0.23 | B>A |
| chr19 | 53000000 | 53100000 | -1.47 | 0.24 | B>A |
| chr19 | 53100000 | 53200000 | -1.34 | 0.21 | B>A |
| chr19 | 53200000 | 53300000 | -1.33 | 0.24 | B>A |
| chr19 | 53300000 | 53400000 | -1.33 | 0.37 | B>A |
| chr19 | 53400000 | 53500000 | -1.38 | 0.36 | B>A |
| chr19 | 53500000 | 53600000 | -1.31 | 0.45 | B>A |
| chr19 | 53600000 | 53700000 | -1.25 | 0.21 | B>A |
| chr19 | 53700000 | 53800000 | -1.31 | 0.11 | B>A |
| chr19 | 53800000 | 53900000 | -1.14 | 0.21 | B>A |
| chr19 | 53900000 | 54000000 | -1.03 | 0.24 | B>A |

| | | | | | |
|---|---|---|---|---|---|
| chr19 | 54000000 | 54100000 | -0.87 | 0.30 | B>A |
| chr19 | 54100000 | 54200000 | -0.96 | 0.11 | B>A |
| chr19 | 54200000 | 54300000 | -0.71 | 0.08 | B>A |
| chr19 | 54300000 | 54400000 | 0.29 | 0.49 | A>A |
| chr19 | 54400000 | 54500000 | 0.43 | 0.56 | A>A |
| chr19 | 54500000 | 54600000 | 0.26 | 0.41 | A>A |
| chr19 | 54600000 | 54700000 | 0.58 | 0.69 | A>A |
| chr19 | 54700000 | 54800000 | 0.38 | 0.25 | A>A |
| chr19 | 54800000 | 54900000 | 0.15 | -0.01 | A>A |
| chr19 | 54900000 | 55000000 | 0.23 | 0.01 | A>A |
| chr19 | 55000000 | 55100000 | -0.17 | -0.63 | B>B |
| chr19 | 55100000 | 55200000 | -0.22 | -0.62 | B>B |
| chr19 | 55200000 | 55300000 | 0.00 | 0.00 | NO |
| chr19 | 55300000 | 55400000 | 0.00 | 0.00 | NO |
| chr19 | 55400000 | 55500000 | 0.07 | -0.49 | B>B |
| chr19 | 55500000 | 55600000 | 0.42 | 0.09 | A>A |
| chr19 | 55600000 | 55700000 | 0.69 | 0.52 | A>A |
| chr19 | 55700000 | 55800000 | 0.86 | 0.81 | A>A |
| chr19 | 55800000 | 55900000 | 1.05 | 0.85 | A>A |
| chr19 | 55900000 | 56000000 | 0.71 | 0.57 | A>A |
| chr19 | 56000000 | 56100000 | 0.76 | 0.69 | A>A |
| chr19 | 56100000 | 56200000 | 0.42 | 0.51 | A>A |
| chr19 | 56200000 | 56300000 | -0.41 | -0.40 | B>B |
| chr19 | 56300000 | 56400000 | -0.89 | -0.73 | B>B |
| chr19 | 56400000 | 56500000 | -1.05 | -0.74 | B>B |
| chr19 | 56500000 | 56600000 | -0.75 | -0.54 | B>B |
| chr19 | 56600000 | 56700000 | -0.32 | -0.15 | B>B |
| chr19 | 56700000 | 56800000 | -0.72 | -0.46 | B>B |
| chr19 | 56800000 | 56900000 | -0.83 | -0.20 | B>A |
| chr19 | 56900000 | 57000000 | -1.35 | -0.44 | B>A |
| chr19 | 57000000 | 57100000 | -1.22 | -0.46 | B>A |
| chr19 | 57100000 | 57200000 | -1.25 | -0.67 | B>A |
| chr19 | 57200000 | 57300000 | -1.36 | -0.80 | B>A |
| chr19 | 57300000 | 57400000 | -1.50 | -0.88 | B>A |
| chr19 | 57400000 | 57500000 | -1.46 | -0.86 | B>A |
| chr19 | 57500000 | 57600000 | -1.35 | -0.84 | B>A |
| chr19 | 57600000 | 57700000 | -1.10 | -0.57 | B>A |
| chr19 | 57700000 | 57800000 | -0.78 | -0.26 | B>A |
| chr19 | 57800000 | 57900000 | -0.59 | -0.05 | B>A |
| chr19 | 57900000 | 58000000 | -0.68 | 0.00 | B>A |
| chr19 | 58000000 | 58100000 | -0.71 | 0.08 | B>A |
| chr19 | 58100000 | 58200000 | -0.70 | 0.08 | B>A |
| chr19 | 58200000 | 58300000 | -0.66 | 0.06 | B>A |
| chr19 | 58300000 | 58400000 | -0.42 | 0.04 | B>B |
| chr19 | 58400000 | 58500000 | -0.81 | 0.31 | B>A |
| chr19 | 58500000 | 58600000 | -1.27 | 0.49 | B>A |
| chr19 | 58600000 | 58700000 | -1.06 | 0.43 | B>A |
| chr19 | 58700000 | 58800000 | -0.72 | 0.47 | B>A |
| chr19 | 58800000 | 58900000 | 0.15 | 0.73 | B>A |
| chr19 | 58900000 | 59000000 | 0.55 | 0.99 | A>A |
| chr19 | 59000000 | 59100000 | 0.84 | 0.82 | A>A |
| chr19 | 59100000 | 59128983 | 0.00 | 0.00 | NO |
| chr20 | 0 | 100000 | 0.00 | 0.00 | NO |
| chr20 | 100000 | 200000 | -0.29 | -0.43 | B>B |
| chr20 | 200000 | 300000 | 0.45 | 0.15 | A>A |
| chr20 | 300000 | 400000 | 0.92 | 0.76 | A>A |

| | | | | | |
|---|---|---|---|---|---|
| chr20 | 400000 | 500000 | 0.80 | 0.84 | A>A |
| chr20 | 500000 | 600000 | 0.83 | 0.66 | A>A |
| chr20 | 600000 | 700000 | 0.79 | 0.66 | A>A |
| chr20 | 700000 | 800000 | 0.43 | 0.40 | A>A |
| chr20 | 800000 | 900000 | 0.53 | 0.23 | A>A |
| chr20 | 900000 | 1000000 | 0.27 | 0.12 | A>A |
| chr20 | 1000000 | 1100000 | 0.19 | 0.05 | A>A |
| chr20 | 1100000 | 1200000 | 0.40 | 0.33 | A>A |
| chr20 | 1200000 | 1300000 | 0.44 | 0.48 | A>A |
| chr20 | 1300000 | 1400000 | 0.48 | 0.43 | A>A |
| chr20 | 1400000 | 1500000 | 0.31 | 0.13 | A>A |
| chr20 | 1500000 | 1600000 | 0.00 | 0.00 | NO |
| chr20 | 1600000 | 1700000 | -0.12 | -0.35 | B>B |
| chr20 | 1700000 | 1800000 | 0.17 | 0.01 | A>A |
| chr20 | 1800000 | 1900000 | 0.52 | 0.38 | A>A |
| chr20 | 1900000 | 2000000 | 0.29 | 0.17 | A>A |
| chr20 | 2000000 | 2100000 | 0.27 | 0.11 | A>A |
| chr20 | 2100000 | 2200000 | 0.33 | 0.27 | A>A |
| chr20 | 2200000 | 2300000 | 0.33 | 0.47 | A>A |
| chr20 | 2300000 | 2400000 | 0.43 | 0.36 | A>A |
| chr20 | 2400000 | 2500000 | 0.56 | 0.47 | A>A |
| chr20 | 2500000 | 2600000 | 0.76 | 0.61 | A>A |
| chr20 | 2600000 | 2700000 | 0.83 | 0.68 | A>A |
| chr20 | 2700000 | 2800000 | 0.89 | 0.80 | A>A |
| chr20 | 2800000 | 2900000 | 0.97 | 0.86 | A>A |
| chr20 | 2900000 | 3000000 | 0.77 | 0.83 | A>A |
| chr20 | 3000000 | 3100000 | 0.86 | 0.86 | A>A |
| chr20 | 3100000 | 3200000 | 1.01 | 0.99 | A>A |
| chr20 | 3200000 | 3300000 | 0.99 | 0.94 | A>A |
| chr20 | 3300000 | 3400000 | 0.85 | 0.82 | A>A |
| chr20 | 3400000 | 3500000 | 0.99 | 0.83 | A>A |
| chr20 | 3500000 | 3600000 | 0.91 | 0.88 | A>A |
| chr20 | 3600000 | 3700000 | 0.98 | 0.89 | A>A |
| chr20 | 3700000 | 3800000 | 1.17 | 1.01 | A>A |
| chr20 | 3800000 | 3900000 | 1.02 | 0.97 | A>A |
| chr20 | 3900000 | 4000000 | 0.92 | 0.92 | A>A |
| chr20 | 4000000 | 4100000 | 0.91 | 0.91 | A>A |
| chr20 | 4100000 | 4200000 | 0.85 | 0.83 | A>A |
| chr20 | 4200000 | 4300000 | 0.60 | 0.48 | A>A |
| chr20 | 4300000 | 4400000 | 0.62 | 0.46 | A>A |
| chr20 | 4400000 | 4500000 | 0.68 | 0.47 | A>A |
| chr20 | 4500000 | 4600000 | 0.57 | 0.55 | A>A |
| chr20 | 4600000 | 4700000 | 0.59 | 0.58 | A>A |
| chr20 | 4700000 | 4800000 | 0.62 | 0.66 | A>A |
| chr20 | 4800000 | 4900000 | 0.84 | 0.76 | A>A |
| chr20 | 4900000 | 5000000 | 0.75 | 0.74 | A>A |
| chr20 | 5000000 | 5100000 | 0.85 | 0.77 | A>A |
| chr20 | 5100000 | 5200000 | 0.61 | 0.56 | A>A |
| chr20 | 5200000 | 5300000 | 0.11 | 0.12 | A>A |
| chr20 | 5300000 | 5400000 | 0.02 | 0.23 | A>A |
| chr20 | 5400000 | 5500000 | 0.09 | 0.31 | A>A |
| chr20 | 5500000 | 5600000 | 0.47 | 0.62 | A>A |
| chr20 | 5600000 | 5700000 | 0.85 | 0.91 | A>A |
| chr20 | 5700000 | 5800000 | 0.68 | 0.75 | A>A |
| chr20 | 5800000 | 5900000 | 0.63 | 0.68 | A>A |
| chr20 | 5900000 | 6000000 | 0.42 | 0.54 | A>A |

| | | | | | |
|---|---|---|---|---|---|
| chr20 | 6000000 | 6100000 | 0.04 | 0.21 | A>A |
| chr20 | 6100000 | 6200000 | -0.90 | -0.81 | B>B |
| chr20 | 6200000 | 6300000 | -1.16 | -1.19 | B>B |
| chr20 | 6300000 | 6400000 | -1.31 | -1.46 | B>B |
| chr20 | 6400000 | 6500000 | -1.20 | -1.33 | B>B |
| chr20 | 6500000 | 6600000 | -1.12 | -1.07 | B>B |
| chr20 | 6600000 | 6700000 | -1.02 | -0.74 | B>B |
| chr20 | 6700000 | 6800000 | -0.92 | -0.66 | B>B |
| chr20 | 6800000 | 6900000 | -1.05 | -0.97 | B>B |
| chr20 | 6900000 | 7000000 | -1.13 | -1.03 | B>B |
| chr20 | 7000000 | 7100000 | -1.03 | -1.01 | B>B |
| chr20 | 7100000 | 7200000 | -1.10 | -1.02 | B>B |
| chr20 | 7200000 | 7300000 | -1.07 | -0.97 | B>B |
| chr20 | 7300000 | 7400000 | -1.02 | -0.77 | B>B |
| chr20 | 7400000 | 7500000 | -0.97 | -0.76 | B>B |
| chr20 | 7500000 | 7600000 | -0.87 | -0.75 | B>B |
| chr20 | 7600000 | 7700000 | -0.91 | -0.73 | B>B |
| chr20 | 7700000 | 7800000 | -0.90 | -0.69 | B>B |
| chr20 | 7800000 | 7900000 | -0.88 | -0.65 | B>B |
| chr20 | 7900000 | 8000000 | -0.71 | -0.37 | B>B |
| chr20 | 8000000 | 8100000 | -0.72 | -0.33 | B>B |
| chr20 | 8100000 | 8200000 | -0.91 | -0.40 | B>A |
| chr20 | 8200000 | 8300000 | -1.03 | -0.54 | B>B |
| chr20 | 8300000 | 8400000 | -0.98 | -0.61 | B>B |
| chr20 | 8400000 | 8500000 | -1.07 | -0.62 | B>B |
| chr20 | 8500000 | 8600000 | -0.89 | -0.47 | B>B |
| chr20 | 8600000 | 8700000 | -0.85 | -0.42 | B>B |
| chr20 | 8700000 | 8800000 | -0.43 | -0.16 | B>B |
| chr20 | 8800000 | 8900000 | -0.13 | 0.08 | B>B |
| chr20 | 8900000 | 9000000 | -0.03 | 0.18 | A>A |
| chr20 | 9000000 | 9100000 | 0.17 | 0.31 | A>A |
| chr20 | 9100000 | 9200000 | 0.19 | 0.32 | A>A |
| chr20 | 9200000 | 9300000 | 0.14 | 0.34 | A>A |
| chr20 | 9300000 | 9400000 | -0.18 | 0.09 | B>B |
| chr20 | 9400000 | 9500000 | -0.24 | 0.00 | B>B |
| chr20 | 9500000 | 9600000 | -0.22 | 0.08 | B>B |
| chr20 | 9600000 | 9700000 | -0.68 | -0.49 | B>B |
| chr20 | 9700000 | 9800000 | -0.91 | -0.73 | B>B |
| chr20 | 9800000 | 9900000 | -0.60 | -0.52 | B>B |
| chr20 | 9900000 | 10000000 | -0.16 | -0.03 | B>B |
| chr20 | 10000000 | 10100000 | 0.21 | 0.21 | A>A |
| chr20 | 10100000 | 10200000 | 0.27 | 0.29 | A>A |
| chr20 | 10200000 | 10300000 | 0.30 | 0.33 | A>A |
| chr20 | 10300000 | 10400000 | 0.30 | 0.32 | A>A |
| chr20 | 10400000 | 10500000 | 0.17 | 0.23 | A>A |
| chr20 | 10500000 | 10600000 | 0.19 | 0.18 | A>A |
| chr20 | 10600000 | 10700000 | 0.18 | 0.25 | A>A |
| chr20 | 10700000 | 10800000 | 0.40 | 0.31 | A>A |
| chr20 | 10800000 | 10900000 | 0.33 | 0.27 | A>A |
| chr20 | 10900000 | 11000000 | 0.32 | 0.28 | A>A |
| chr20 | 11000000 | 11100000 | 0.27 | 0.25 | A>A |
| chr20 | 11100000 | 11200000 | 0.17 | 0.24 | A>A |
| chr20 | 11200000 | 11300000 | 0.00 | 0.13 | A>A |
| chr20 | 11300000 | 11400000 | -0.08 | -0.03 | B>B |
| chr20 | 11400000 | 11500000 | -0.12 | -0.08 | B>B |
| chr20 | 11500000 | 11600000 | -0.46 | -0.43 | B>B |

| | | | | | |
|---|---|---|---|---|---|
| chr20 | 11600000 | 11700000 | -0.77 | -0.68 | B>B |
| chr20 | 11700000 | 11800000 | -0.89 | -0.93 | B>B |
| chr20 | 11800000 | 11900000 | -0.75 | -0.77 | B>B |
| chr20 | 11900000 | 12000000 | -0.86 | -0.84 | B>B |
| chr20 | 12000000 | 12100000 | -1.04 | -1.00 | B>B |
| chr20 | 12100000 | 12200000 | -1.09 | -1.03 | B>B |
| chr20 | 12200000 | 12300000 | -1.16 | -1.11 | B>B |
| chr20 | 12300000 | 12400000 | -1.21 | -1.15 | B>B |
| chr20 | 12400000 | 12500000 | -1.19 | -1.19 | B>B |
| chr20 | 12500000 | 12600000 | -1.30 | -1.24 | B>B |
| chr20 | 12600000 | 12700000 | -1.33 | -1.41 | B>B |
| chr20 | 12700000 | 12800000 | -1.16 | -1.30 | B>B |
| chr20 | 12800000 | 12900000 | -1.00 | -1.07 | B>B |
| chr20 | 12900000 | 13000000 | -0.49 | -0.32 | B>B |
| chr20 | 13000000 | 13100000 | -0.48 | -0.19 | B>B |
| chr20 | 13100000 | 13200000 | -0.50 | -0.49 | B>B |
| chr20 | 13200000 | 13300000 | -0.30 | -0.31 | B>B |
| chr20 | 13300000 | 13400000 | -0.18 | -0.18 | B>B |
| chr20 | 13400000 | 13500000 | -0.16 | -0.07 | B>B |
| chr20 | 13500000 | 13600000 | -0.07 | -0.03 | B>B |
| chr20 | 13600000 | 13700000 | 0.00 | 0.00 | B>B |
| chr20 | 13700000 | 13800000 | -0.13 | 0.00 | B>B |
| chr20 | 13800000 | 13900000 | -0.59 | -0.14 | B>B |
| chr20 | 13900000 | 14000000 | -0.69 | -0.04 | B>A |
| chr20 | 14000000 | 14100000 | -0.67 | -0.06 | B>A |
| chr20 | 14100000 | 14200000 | -0.50 | -0.01 | B>B |
| chr20 | 14200000 | 14300000 | -0.35 | 0.09 | B>B |
| chr20 | 14300000 | 14400000 | 0.00 | 0.22 | A>A |
| chr20 | 14400000 | 14500000 | 0.11 | 0.29 | A>A |
| chr20 | 14500000 | 14600000 | 0.09 | 0.30 | A>A |
| chr20 | 14600000 | 14700000 | 0.06 | 0.27 | A>A |
| chr20 | 14700000 | 14800000 | 0.15 | 0.28 | A>A |
| chr20 | 14800000 | 14900000 | 0.09 | 0.28 | A>A |
| chr20 | 14900000 | 15000000 | 0.04 | 0.26 | A>A |
| chr20 | 15000000 | 15100000 | 0.03 | 0.24 | A>A |
| chr20 | 15100000 | 15200000 | -0.01 | 0.19 | A>A |
| chr20 | 15200000 | 15300000 | -0.13 | 0.10 | B>B |
| chr20 | 15300000 | 15400000 | -0.27 | 0.01 | B>B |
| chr20 | 15400000 | 15500000 | -0.42 | -0.08 | B>B |
| chr20 | 15500000 | 15600000 | -0.58 | -0.18 | B>B |
| chr20 | 15600000 | 15700000 | -0.58 | -0.26 | B>B |
| chr20 | 15700000 | 15800000 | -0.75 | -0.53 | B>B |
| chr20 | 15800000 | 15900000 | -0.84 | -0.59 | B>B |
| chr20 | 15900000 | 16000000 | -0.80 | -0.65 | B>B |
| chr20 | 16000000 | 16100000 | -0.80 | -0.92 | B>B |
| chr20 | 16100000 | 16200000 | -0.83 | -0.91 | B>B |
| chr20 | 16200000 | 16300000 | -0.69 | -0.55 | B>B |
| chr20 | 16300000 | 16400000 | -0.46 | -0.31 | B>B |
| chr20 | 16400000 | 16500000 | -0.38 | -0.25 | B>B |
| chr20 | 16500000 | 16600000 | -0.27 | -0.21 | B>B |
| chr20 | 16600000 | 16700000 | -0.60 | -0.45 | B>B |
| chr20 | 16700000 | 16800000 | -0.67 | -0.58 | B>B |
| chr20 | 16800000 | 16900000 | -1.05 | -1.32 | B>B |
| chr20 | 16900000 | 17000000 | -1.22 | -1.50 | B>B |
| chr20 | 17000000 | 17100000 | -1.21 | -1.58 | B>B |
| chr20 | 17100000 | 17200000 | -1.08 | -1.55 | B>B |

| | | | | | |
|---|---|---|---|---|---|
| chr20 | 17200000 | 17300000 | -1.00 | -1.43 | B>B |
| chr20 | 17300000 | 17400000 | -0.96 | -1.30 | B>B |
| chr20 | 17400000 | 17500000 | -0.44 | -0.57 | B>B |
| chr20 | 17500000 | 17600000 | 0.29 | 0.20 | A>A |
| chr20 | 17600000 | 17700000 | 0.24 | 0.25 | A>A |
| chr20 | 17700000 | 17800000 | 0.27 | 0.31 | A>A |
| chr20 | 17800000 | 17900000 | 0.29 | 0.34 | A>A |
| chr20 | 17900000 | 18000000 | 0.33 | 0.29 | A>A |
| chr20 | 18000000 | 18100000 | 0.14 | 0.14 | A>A |
| chr20 | 18100000 | 18200000 | 0.08 | 0.21 | A>A |
| chr20 | 18200000 | 18300000 | 0.01 | 0.10 | A>A |
| chr20 | 18300000 | 18400000 | -0.16 | -0.09 | B>B |
| chr20 | 18400000 | 18500000 | 0.19 | 0.11 | A>A |
| chr20 | 18500000 | 18600000 | 0.09 | 0.00 | A>A |
| chr20 | 18600000 | 18700000 | -0.30 | -0.44 | B>B |
| chr20 | 18700000 | 18800000 | -0.53 | -0.68 | B>B |
| chr20 | 18800000 | 18900000 | -0.74 | -1.05 | B>B |
| chr20 | 18900000 | 19000000 | -0.80 | -1.21 | B>B |
| chr20 | 19000000 | 19100000 | -0.81 | -1.20 | B>B |
| chr20 | 19100000 | 19200000 | -0.75 | -1.15 | B>B |
| chr20 | 19200000 | 19300000 | -0.74 | -1.21 | B>B |
| chr20 | 19300000 | 19400000 | -0.91 | -1.37 | B>B |
| chr20 | 19400000 | 19500000 | -0.84 | -1.29 | B>B |
| chr20 | 19500000 | 19600000 | -0.63 | -1.04 | B>B |
| chr20 | 19600000 | 19700000 | -0.15 | -0.39 | B>B |
| chr20 | 19700000 | 19800000 | 0.28 | 0.09 | A>A |
| chr20 | 19800000 | 19900000 | 0.38 | 0.30 | A>A |
| chr20 | 19900000 | 20000000 | 0.26 | 0.28 | A>A |
| chr20 | 20000000 | 20100000 | -0.18 | -0.30 | B>B |
| chr20 | 20100000 | 20200000 | -0.66 | -0.80 | B>B |
| chr20 | 20200000 | 20300000 | -0.67 | -0.94 | B>B |
| chr20 | 20300000 | 20400000 | -0.46 | -0.52 | B>B |
| chr20 | 20400000 | 20500000 | -0.41 | -0.39 | B>B |
| chr20 | 20500000 | 20600000 | -0.38 | -0.38 | B>B |
| chr20 | 20600000 | 20700000 | -0.28 | -0.45 | B>B |
| chr20 | 20700000 | 20800000 | -0.37 | -0.76 | B>B |
| chr20 | 20800000 | 20900000 | -0.48 | -0.79 | B>B |
| chr20 | 20900000 | 21000000 | -0.69 | -1.04 | B>B |
| chr20 | 21000000 | 21100000 | -0.39 | -0.78 | B>B |
| chr20 | 21100000 | 21200000 | -0.25 | -0.32 | B>B |
| chr20 | 21200000 | 21300000 | -0.13 | -0.20 | B>B |
| chr20 | 21300000 | 21400000 | 0.08 | -0.08 | A>A |
| chr20 | 21400000 | 21500000 | -0.35 | -0.45 | B>B |
| chr20 | 21500000 | 21600000 | -0.55 | -0.74 | B>B |
| chr20 | 21600000 | 21700000 | -0.76 | -0.98 | B>B |
| chr20 | 21700000 | 21800000 | -0.79 | -1.17 | B>B |
| chr20 | 21800000 | 21900000 | -0.91 | -1.29 | B>B |
| chr20 | 21900000 | 22000000 | -0.93 | -1.33 | B>B |
| chr20 | 22000000 | 22100000 | -0.91 | -1.27 | B>B |
| chr20 | 22100000 | 22200000 | -0.85 | -1.19 | B>B |
| chr20 | 22200000 | 22300000 | -0.51 | -0.78 | B>B |
| chr20 | 22300000 | 22400000 | 0.22 | 0.06 | A>A |
| chr20 | 22400000 | 22500000 | 0.56 | 0.36 | A>A |
| chr20 | 22500000 | 22600000 | 0.40 | 0.18 | A>A |
| chr20 | 22600000 | 22700000 | 0.15 | 0.00 | A>A |
| chr20 | 22700000 | 22800000 | 0.38 | 0.32 | A>A |

| | | | | | |
|---|---|---|---|---|---|
| chr20 | 22800000 | 22900000 | 0.39 | 0.30 | A>A |
| chr20 | 22900000 | 23000000 | 0.37 | 0.36 | A>A |
| chr20 | 23000000 | 23100000 | 0.30 | 0.24 | A>A |
| chr20 | 23100000 | 23200000 | 0.25 | 0.20 | A>A |
| chr20 | 23200000 | 23300000 | 0.17 | 0.05 | A>A |
| chr20 | 23300000 | 23400000 | 0.24 | 0.18 | A>A |
| chr20 | 23400000 | 23500000 | -0.05 | -0.13 | B>B |
| chr20 | 23500000 | 23600000 | -0.14 | -0.17 | B>B |
| chr20 | 23600000 | 23700000 | -0.06 | -0.18 | B>B |
| chr20 | 23700000 | 23800000 | 0.00 | 0.00 | NO |
| chr20 | 23800000 | 23900000 | -0.31 | -0.72 | B>B |
| chr20 | 23900000 | 24000000 | -0.31 | -0.62 | B>B |
| chr20 | 24000000 | 24100000 | -0.32 | -1.08 | A>B |
| chr20 | 24100000 | 24200000 | -0.50 | -1.22 | A>B |
| chr20 | 24200000 | 24300000 | -0.59 | -1.14 | B>B |
| chr20 | 24300000 | 24400000 | -0.55 | -1.11 | B>B |
| chr20 | 24400000 | 24500000 | -0.40 | -0.90 | B>B |
| chr20 | 24500000 | 24600000 | -0.28 | -0.75 | B>B |
| chr20 | 24600000 | 24700000 | -0.10 | -0.53 | B>B |
| chr20 | 24700000 | 24800000 | -0.17 | -0.42 | B>B |
| chr20 | 24800000 | 24900000 | -0.23 | -0.55 | B>B |
| chr20 | 24900000 | 25000000 | 0.19 | 0.04 | A>A |
| chr20 | 25000000 | 25100000 | 0.58 | 0.43 | A>A |
| chr20 | 25100000 | 25200000 | 0.62 | 0.54 | A>A |
| chr20 | 25200000 | 25300000 | 0.64 | 0.62 | A>A |
| chr20 | 25300000 | 25400000 | 0.61 | 0.58 | A>A |
| chr20 | 25400000 | 25500000 | 0.45 | 0.54 | A>A |
| chr20 | 25500000 | 25600000 | 0.53 | 0.60 | A>A |
| chr20 | 25600000 | 25700000 | 0.34 | 0.39 | A>A |
| chr20 | 25700000 | 25800000 | 0.00 | 0.00 | NO |
| chr20 | 25800000 | 25900000 | 0.00 | 0.00 | NO |
| chr20 | 25900000 | 26000000 | 0.00 | 0.00 | NO |
| chr20 | 26000000 | 26100000 | 0.00 | 0.00 | NO |
| chr20 | 26100000 | 26200000 | 0.00 | 0.00 | NO |
| chr20 | 26200000 | 26300000 | 0.00 | 0.00 | NO |
| chr20 | 26300000 | 26400000 | 0.00 | 0.00 | NO |
| chr20 | 26400000 | 26500000 | 0.00 | 0.00 | NO |
| chr20 | 26500000 | 26600000 | 0.00 | 0.00 | NO |
| chr20 | 26600000 | 26700000 | 0.00 | 0.00 | NO |
| chr20 | 26700000 | 26800000 | 0.00 | 0.00 | NO |
| chr20 | 26800000 | 26900000 | 0.00 | 0.00 | NO |
| chr20 | 26900000 | 27000000 | 0.00 | 0.00 | NO |
| chr20 | 27000000 | 27100000 | 0.00 | 0.00 | NO |
| chr20 | 27100000 | 27200000 | 0.00 | 0.00 | NO |
| chr20 | 27200000 | 27300000 | 0.00 | 0.00 | NO |
| chr20 | 27300000 | 27400000 | 0.00 | 0.00 | NO |
| chr20 | 27400000 | 27500000 | 0.00 | 0.00 | NO |
| chr20 | 27500000 | 27600000 | 0.00 | 0.00 | NO |
| chr20 | 27600000 | 27700000 | 0.00 | 0.00 | NO |
| chr20 | 27700000 | 27800000 | 0.00 | 0.00 | NO |
| chr20 | 27800000 | 27900000 | 0.00 | 0.00 | NO |
| chr20 | 27900000 | 28000000 | 0.00 | 0.00 | NO |
| chr20 | 28000000 | 28100000 | 0.00 | 0.00 | NO |
| chr20 | 28100000 | 28200000 | 0.00 | 0.00 | NO |
| chr20 | 28200000 | 28300000 | 0.00 | 0.00 | NO |
| chr20 | 28300000 | 28400000 | 0.00 | 0.00 | NO |

| | | | | | |
|---|---|---|---|---|---|
| chr20 | 28400000 | 28500000 | 0.00 | 0.00 | NO |
| chr20 | 28500000 | 28600000 | 0.00 | 0.00 | NO |
| chr20 | 28600000 | 28700000 | 0.00 | 0.00 | NO |
| chr20 | 28700000 | 28800000 | 0.00 | 0.00 | NO |
| chr20 | 28800000 | 28900000 | 0.00 | 0.00 | NO |
| chr20 | 28900000 | 29000000 | 0.00 | 0.00 | NO |
| chr20 | 29000000 | 29100000 | 0.00 | 0.00 | NO |
| chr20 | 29100000 | 29200000 | 0.00 | 0.00 | NO |
| chr20 | 29200000 | 29300000 | 0.00 | 0.00 | NO |
| chr20 | 29300000 | 29400000 | 0.00 | 0.00 | NO |
| chr20 | 29400000 | 29500000 | 0.00 | 0.00 | NO |
| chr20 | 29500000 | 29600000 | 0.00 | 0.00 | NO |
| chr20 | 29600000 | 29700000 | 0.00 | 0.00 | NO |
| chr20 | 29700000 | 29800000 | 0.00 | 0.00 | NO |
| chr20 | 29800000 | 29900000 | 0.00 | 0.00 | NO |
| chr20 | 29900000 | 30000000 | -1.31 | -1.86 | B>B |
| chr20 | 30000000 | 30100000 | -0.32 | -0.93 | A>B |
| chr20 | 30100000 | 30200000 | 0.59 | 0.53 | A>A |
| chr20 | 30200000 | 30300000 | 0.60 | 0.74 | A>A |
| chr20 | 30300000 | 30400000 | 0.88 | 0.95 | A>A |
| chr20 | 30400000 | 30500000 | 0.92 | 0.89 | A>A |
| chr20 | 30500000 | 30600000 | 0.96 | 0.92 | A>A |
| chr20 | 30600000 | 30700000 | 0.89 | 0.87 | A>A |
| chr20 | 30700000 | 30800000 | 0.93 | 0.92 | A>A |
| chr20 | 30800000 | 30900000 | 0.92 | 0.89 | A>A |
| chr20 | 30900000 | 31000000 | 0.94 | 0.91 | A>A |
| chr20 | 31000000 | 31100000 | 0.88 | 0.88 | A>A |
| chr20 | 31100000 | 31200000 | 0.95 | 0.88 | A>A |
| chr20 | 31200000 | 31300000 | 0.82 | 0.76 | A>A |
| chr20 | 31300000 | 31400000 | 0.86 | 0.83 | A>A |
| chr20 | 31400000 | 31500000 | 0.73 | 0.70 | A>A |
| chr20 | 31500000 | 31600000 | 0.63 | 0.65 | A>A |
| chr20 | 31600000 | 31700000 | 0.62 | 0.62 | A>A |
| chr20 | 31700000 | 31800000 | 0.69 | 0.73 | A>A |
| chr20 | 31800000 | 31900000 | 0.65 | 0.67 | A>A |
| chr20 | 31900000 | 32000000 | 0.68 | 0.68 | A>A |
| chr20 | 32000000 | 32100000 | 0.64 | 0.70 | A>A |
| chr20 | 32100000 | 32200000 | 0.78 | 0.77 | A>A |
| chr20 | 32200000 | 32300000 | 0.71 | 0.72 | A>A |
| chr20 | 32300000 | 32400000 | 0.73 | 0.70 | A>A |
| chr20 | 32400000 | 32500000 | 0.69 | 0.66 | A>A |
| chr20 | 32500000 | 32600000 | 0.65 | 0.67 | A>A |
| chr20 | 32600000 | 32700000 | 0.65 | 0.62 | A>A |
| chr20 | 32700000 | 32800000 | 0.63 | 0.61 | A>A |
| chr20 | 32800000 | 32900000 | 0.63 | 0.58 | A>A |
| chr20 | 32900000 | 33000000 | 0.74 | 0.66 | A>A |
| chr20 | 33000000 | 33100000 | 0.67 | 0.62 | A>A |
| chr20 | 33100000 | 33200000 | 0.60 | 0.64 | A>A |
| chr20 | 33200000 | 33300000 | 0.71 | 0.66 | A>A |
| chr20 | 33300000 | 33400000 | 0.64 | 0.64 | A>A |
| chr20 | 33400000 | 33500000 | 0.66 | 0.63 | A>A |
| chr20 | 33500000 | 33600000 | 0.76 | 0.70 | A>A |
| chr20 | 33600000 | 33700000 | 0.69 | 0.65 | A>A |
| chr20 | 33700000 | 33800000 | 0.69 | 0.65 | A>A |
| chr20 | 33800000 | 33900000 | 0.67 | 0.70 | A>A |
| chr20 | 33900000 | 34000000 | 0.76 | 0.73 | A>A |

| | | | | | |
|---|---|---|---|---|---|
| chr20 | 34000000 | 34100000 | 0.68 | 0.67 | A>A |
| chr20 | 34100000 | 34200000 | 0.76 | 0.75 | A>A |
| chr20 | 34200000 | 34300000 | 0.73 | 0.69 | A>A |
| chr20 | 34300000 | 34400000 | 0.77 | 0.70 | A>A |
| chr20 | 34400000 | 34500000 | 0.70 | 0.69 | A>A |
| chr20 | 34500000 | 34600000 | 0.63 | 0.65 | A>A |
| chr20 | 34600000 | 34700000 | 0.84 | 0.81 | A>A |
| chr20 | 34700000 | 34800000 | 0.65 | 0.69 | A>A |
| chr20 | 34800000 | 34900000 | 0.56 | 0.64 | A>A |
| chr20 | 34900000 | 35000000 | 0.00 | 0.00 | NO |
| chr20 | 35000000 | 35100000 | 0.57 | 0.62 | A>A |
| chr20 | 35100000 | 35200000 | 0.71 | 0.72 | A>A |
| chr20 | 35200000 | 35300000 | 0.75 | 0.72 | A>A |
| chr20 | 35300000 | 35400000 | 0.77 | 0.72 | A>A |
| chr20 | 35400000 | 35500000 | 0.86 | 0.80 | A>A |
| chr20 | 35500000 | 35600000 | 0.72 | 0.72 | A>A |
| chr20 | 35600000 | 35700000 | 0.89 | 0.78 | A>A |
| chr20 | 35700000 | 35800000 | 0.87 | 0.83 | A>A |
| chr20 | 35800000 | 35900000 | 0.80 | 0.79 | A>A |
| chr20 | 35900000 | 36000000 | 1.00 | 0.91 | A>A |
| chr20 | 36000000 | 36100000 | 0.90 | 0.88 | A>A |
| chr20 | 36100000 | 36200000 | 0.75 | 0.70 | A>A |
| chr20 | 36200000 | 36300000 | 0.65 | 0.69 | A>A |
| chr20 | 36300000 | 36400000 | 0.80 | 0.80 | A>A |
| chr20 | 36400000 | 36500000 | 0.79 | 0.74 | A>A |
| chr20 | 36500000 | 36600000 | 0.68 | 0.78 | A>A |
| chr20 | 36600000 | 36700000 | 0.75 | 0.79 | A>A |
| chr20 | 36700000 | 36800000 | 0.93 | 0.96 | A>A |
| chr20 | 36800000 | 36900000 | 0.81 | 0.87 | A>A |
| chr20 | 36900000 | 37000000 | 0.77 | 0.74 | A>A |
| chr20 | 37000000 | 37100000 | 0.93 | 0.88 | A>A |
| chr20 | 37100000 | 37200000 | 0.96 | 0.89 | A>A |
| chr20 | 37200000 | 37300000 | 0.70 | 0.77 | A>A |
| chr20 | 37300000 | 37400000 | 0.78 | 0.64 | A>A |
| chr20 | 37400000 | 37500000 | 0.71 | 0.70 | A>A |
| chr20 | 37500000 | 37600000 | 0.68 | 0.67 | A>A |
| chr20 | 37600000 | 37700000 | 0.29 | 0.17 | A>A |
| chr20 | 37700000 | 37800000 | -0.44 | -0.77 | B>B |
| chr20 | 37800000 | 37900000 | -0.65 | -1.28 | A>B |
| chr20 | 37900000 | 38000000 | -0.91 | -1.43 | B>B |
| chr20 | 38000000 | 38100000 | -1.15 | -1.43 | B>B |
| chr20 | 38100000 | 38200000 | -1.12 | -1.43 | B>B |
| chr20 | 38200000 | 38300000 | -1.13 | -1.39 | B>B |
| chr20 | 38300000 | 38400000 | -1.11 | -1.37 | B>B |
| chr20 | 38400000 | 38500000 | -1.17 | -1.32 | B>B |
| chr20 | 38500000 | 38600000 | -1.10 | -1.31 | B>B |
| chr20 | 38600000 | 38700000 | -1.15 | -1.30 | B>B |
| chr20 | 38700000 | 38800000 | -1.01 | -1.26 | B>B |
| chr20 | 38800000 | 38900000 | -0.91 | -1.26 | B>B |
| chr20 | 38900000 | 39000000 | -0.88 | -1.27 | B>B |
| chr20 | 39000000 | 39100000 | -0.86 | -1.27 | B>B |
| chr20 | 39100000 | 39200000 | -0.73 | -1.17 | B>B |
| chr20 | 39200000 | 39300000 | -0.66 | -1.11 | B>B |
| chr20 | 39300000 | 39400000 | -0.52 | -0.93 | B>B |
| chr20 | 39400000 | 39500000 | -0.55 | -1.09 | B>B |
| chr20 | 39500000 | 39600000 | -0.31 | -0.67 | B>B |

| | | | | | |
|---|---|---|---|---|---|
| chr20 | 39600000 | 39700000 | 0.39 | 0.20 | A>A |
| chr20 | 39700000 | 39800000 | 0.29 | 0.20 | A>A |
| chr20 | 39800000 | 39900000 | 0.24 | 0.15 | A>A |
| chr20 | 39900000 | 40000000 | 0.31 | 0.21 | A>A |
| chr20 | 40000000 | 40100000 | 0.23 | 0.23 | A>A |
| chr20 | 40100000 | 40200000 | 0.22 | 0.20 | A>A |
| chr20 | 40200000 | 40300000 | 0.16 | 0.12 | A>A |
| chr20 | 40300000 | 40400000 | -0.37 | -0.67 | B>B |
| chr20 | 40400000 | 40500000 | -0.94 | -1.33 | B>B |
| chr20 | 40500000 | 40600000 | -0.98 | -1.43 | B>B |
| chr20 | 40600000 | 40700000 | -1.04 | -1.41 | B>B |
| chr20 | 40700000 | 40800000 | -1.02 | -1.41 | B>B |
| chr20 | 40800000 | 40900000 | -1.08 | -1.32 | B>B |
| chr20 | 40900000 | 41000000 | -1.07 | -1.34 | B>B |
| chr20 | 41000000 | 41100000 | -1.07 | -1.36 | B>B |
| chr20 | 41100000 | 41200000 | -1.09 | -1.36 | B>B |
| chr20 | 41200000 | 41300000 | -1.10 | -1.38 | B>B |
| chr20 | 41300000 | 41400000 | -1.15 | -1.36 | B>B |
| chr20 | 41400000 | 41500000 | -1.16 | -1.41 | B>B |
| chr20 | 41500000 | 41600000 | -1.16 | -1.42 | B>B |
| chr20 | 41600000 | 41700000 | -1.00 | -1.46 | B>B |
| chr20 | 41700000 | 41800000 | -0.92 | -1.38 | B>B |
| chr20 | 41800000 | 41900000 | -0.77 | -1.25 | B>B |
| chr20 | 41900000 | 42000000 | -0.68 | -1.27 | A>B |
| chr20 | 42000000 | 42100000 | -0.08 | -0.26 | B>B |
| chr20 | 42100000 | 42200000 | 0.62 | 0.59 | A>A |
| chr20 | 42200000 | 42300000 | 0.72 | 0.66 | A>A |
| chr20 | 42300000 | 42400000 | 0.57 | 0.54 | A>A |
| chr20 | 42400000 | 42500000 | 0.61 | 0.54 | A>A |
| chr20 | 42500000 | 42600000 | 0.76 | 0.69 | A>A |
| chr20 | 42600000 | 42700000 | 0.60 | 0.65 | A>A |
| chr20 | 42700000 | 42800000 | 0.56 | 0.61 | A>A |
| chr20 | 42800000 | 42900000 | 0.72 | 0.72 | A>A |
| chr20 | 42900000 | 43000000 | 0.87 | 0.82 | A>A |
| chr20 | 43000000 | 43100000 | 0.75 | 0.74 | A>A |
| chr20 | 43100000 | 43200000 | 0.88 | 0.78 | A>A |
| chr20 | 43200000 | 43300000 | 0.89 | 0.73 | A>A |
| chr20 | 43300000 | 43400000 | 1.02 | 0.77 | A>A |
| chr20 | 43400000 | 43500000 | 0.87 | 0.65 | A>A |
| chr20 | 43500000 | 43600000 | 0.82 | 0.67 | A>A |
| chr20 | 43600000 | 43700000 | 0.59 | 0.37 | A>A |
| chr20 | 43700000 | 43800000 | 0.17 | 0.03 | A>A |
| chr20 | 43800000 | 43900000 | 0.01 | -0.13 | B>B |
| chr20 | 43900000 | 44000000 | 0.39 | 0.29 | A>A |
| chr20 | 44000000 | 44100000 | 0.45 | 0.30 | A>A |
| chr20 | 44100000 | 44200000 | 0.03 | -0.14 | B>B |
| chr20 | 44200000 | 44300000 | -0.15 | -0.22 | B>B |
| chr20 | 44300000 | 44400000 | -0.03 | -0.18 | B>B |
| chr20 | 44400000 | 44500000 | 0.67 | 0.59 | A>A |
| chr20 | 44500000 | 44600000 | 0.92 | 0.77 | A>A |
| chr20 | 44600000 | 44700000 | 0.72 | 0.62 | A>A |
| chr20 | 44700000 | 44800000 | 0.65 | 0.57 | A>A |
| chr20 | 44800000 | 44900000 | 0.62 | 0.56 | A>A |
| chr20 | 44900000 | 45000000 | 0.59 | 0.55 | A>A |
| chr20 | 45000000 | 45100000 | 0.44 | 0.44 | A>A |
| chr20 | 45100000 | 45200000 | 0.23 | 0.38 | A>A |

| | | | | | |
|---|---|---|---|---|---|
| chr20 | 45200000 | 45300000 | 0.37 | 0.39 | A>A |
| chr20 | 45300000 | 45400000 | 0.45 | 0.48 | A>A |
| chr20 | 45400000 | 45500000 | 0.50 | 0.53 | A>A |
| chr20 | 45500000 | 45600000 | 0.53 | 0.60 | A>A |
| chr20 | 45600000 | 45700000 | 0.49 | 0.60 | A>A |
| chr20 | 45700000 | 45800000 | 0.47 | 0.51 | A>A |
| chr20 | 45800000 | 45900000 | 0.51 | 0.54 | A>A |
| chr20 | 45900000 | 46000000 | 0.72 | 0.71 | A>A |
| chr20 | 46000000 | 46100000 | 0.70 | 0.64 | A>A |
| chr20 | 46100000 | 46200000 | 0.68 | 0.67 | A>A |
| chr20 | 46200000 | 46300000 | 0.50 | 0.55 | A>A |
| chr20 | 46300000 | 46400000 | 0.42 | 0.46 | A>A |
| chr20 | 46400000 | 46500000 | 0.43 | 0.46 | A>A |
| chr20 | 46500000 | 46600000 | 0.42 | 0.50 | A>A |
| chr20 | 46600000 | 46700000 | 0.39 | 0.50 | A>A |
| chr20 | 46700000 | 46800000 | 0.37 | 0.45 | A>A |
| chr20 | 46800000 | 46900000 | 0.39 | 0.44 | A>A |
| chr20 | 46900000 | 47000000 | 0.47 | 0.44 | A>A |
| chr20 | 47000000 | 47100000 | 0.50 | 0.47 | A>A |
| chr20 | 47100000 | 47200000 | 0.51 | 0.47 | A>A |
| chr20 | 47200000 | 47300000 | 0.38 | 0.44 | A>A |
| chr20 | 47300000 | 47400000 | 0.62 | 0.61 | A>A |
| chr20 | 47400000 | 47500000 | 0.72 | 0.63 | A>A |
| chr20 | 47500000 | 47600000 | 0.65 | 0.56 | A>A |
| chr20 | 47600000 | 47700000 | 0.76 | 0.70 | A>A |
| chr20 | 47700000 | 47800000 | 0.75 | 0.73 | A>A |
| chr20 | 47800000 | 47900000 | 0.84 | 0.71 | A>A |
| chr20 | 47900000 | 48000000 | 0.67 | 0.66 | A>A |
| chr20 | 48000000 | 48100000 | 0.76 | 0.70 | A>A |
| chr20 | 48100000 | 48200000 | 0.65 | 0.59 | A>A |
| chr20 | 48200000 | 48300000 | 0.65 | 0.66 | A>A |
| chr20 | 48300000 | 48400000 | 0.81 | 0.79 | A>A |
| chr20 | 48400000 | 48500000 | 0.66 | 0.64 | A>A |
| chr20 | 48500000 | 48600000 | 0.68 | 0.68 | A>A |
| chr20 | 48600000 | 48700000 | 0.80 | 0.78 | A>A |
| chr20 | 48700000 | 48800000 | 0.73 | 0.74 | A>A |
| chr20 | 48800000 | 48900000 | 0.80 | 0.76 | A>A |
| chr20 | 48900000 | 49000000 | 0.84 | 0.86 | A>A |
| chr20 | 49000000 | 49100000 | 0.75 | 0.73 | A>A |
| chr20 | 49100000 | 49200000 | 0.72 | 0.74 | A>A |
| chr20 | 49200000 | 49300000 | 0.72 | 0.77 | A>A |
| chr20 | 49300000 | 49400000 | 0.84 | 0.82 | A>A |
| chr20 | 49400000 | 49500000 | 0.74 | 0.67 | A>A |
| chr20 | 49500000 | 49600000 | 0.68 | 0.69 | A>A |
| chr20 | 49600000 | 49700000 | 0.52 | 0.35 | A>A |
| chr20 | 49700000 | 49800000 | 0.52 | 0.21 | A>A |
| chr20 | 49800000 | 49900000 | 0.46 | 0.28 | A>A |
| chr20 | 49900000 | 50000000 | 0.44 | 0.38 | A>A |
| chr20 | 50000000 | 50100000 | 0.58 | 0.58 | A>A |
| chr20 | 50100000 | 50200000 | 0.66 | 0.59 | A>A |
| chr20 | 50200000 | 50300000 | 0.47 | 0.51 | A>A |
| chr20 | 50300000 | 50400000 | 0.67 | 0.62 | A>A |
| chr20 | 50400000 | 50500000 | 0.61 | 0.61 | A>A |
| chr20 | 50500000 | 50600000 | 0.62 | 0.69 | A>A |
| chr20 | 50600000 | 50700000 | 0.52 | 0.51 | A>A |
| chr20 | 50700000 | 50800000 | 0.19 | 0.19 | A>A |

| | | | | | |
|---|---|---|---|---|---|
| chr20 | 50800000 | 50900000 | -0.25 | -0.46 | B>B |
| chr20 | 50900000 | 51000000 | -0.70 | -1.04 | B>B |
| chr20 | 51000000 | 51100000 | -0.89 | -1.30 | B>B |
| chr20 | 51100000 | 51200000 | -1.06 | -1.40 | B>B |
| chr20 | 51200000 | 51300000 | -1.11 | -1.40 | B>B |
| chr20 | 51300000 | 51400000 | -1.02 | -1.35 | B>B |
| chr20 | 51400000 | 51500000 | -0.92 | -1.38 | B>B |
| chr20 | 51500000 | 51600000 | -0.77 | -1.24 | B>B |
| chr20 | 51600000 | 51700000 | -0.68 | -1.15 | B>B |
| chr20 | 51700000 | 51800000 | -0.60 | -1.12 | B>B |
| chr20 | 51800000 | 51900000 | -0.55 | -1.03 | B>B |
| chr20 | 51900000 | 52000000 | -0.33 | -0.97 | A>B |
| chr20 | 52000000 | 52100000 | -0.35 | -0.71 | B>B |
| chr20 | 52100000 | 52200000 | -0.09 | -0.24 | B>B |
| chr20 | 52200000 | 52300000 | 0.57 | 0.52 | A>A |
| chr20 | 52300000 | 52400000 | 0.64 | 0.52 | A>A |
| chr20 | 52400000 | 52500000 | 0.64 | 0.43 | A>A |
| chr20 | 52500000 | 52600000 | 0.14 | 0.03 | A>A |
| chr20 | 52600000 | 52700000 | -0.08 | -0.16 | B>B |
| chr20 | 52700000 | 52800000 | 0.57 | 0.42 | A>A |
| chr20 | 52800000 | 52900000 | -0.24 | -0.40 | B>B |
| chr20 | 52900000 | 53000000 | -0.50 | -0.69 | B>B |
| chr20 | 53000000 | 53100000 | -0.71 | -0.92 | B>B |
| chr20 | 53100000 | 53200000 | -0.76 | -1.02 | B>B |
| chr20 | 53200000 | 53300000 | -0.55 | -0.77 | B>B |
| chr20 | 53300000 | 53400000 | -0.90 | -1.18 | B>B |
| chr20 | 53400000 | 53500000 | -1.07 | -1.43 | B>B |
| chr20 | 53500000 | 53600000 | -1.08 | -1.38 | B>B |
| chr20 | 53600000 | 53700000 | -1.05 | -1.38 | B>B |
| chr20 | 53700000 | 53800000 | -1.05 | -1.36 | B>B |
| chr20 | 53800000 | 53900000 | -1.12 | -1.35 | B>B |
| chr20 | 53900000 | 54000000 | -1.09 | -1.35 | B>B |
| chr20 | 54000000 | 54100000 | -1.15 | -1.34 | B>B |
| chr20 | 54100000 | 54200000 | -1.08 | -1.33 | B>B |
| chr20 | 54200000 | 54300000 | -1.16 | -1.41 | B>B |
| chr20 | 54300000 | 54400000 | -1.08 | -1.47 | B>B |
| chr20 | 54400000 | 54500000 | -1.19 | -1.50 | B>B |
| chr20 | 54500000 | 54600000 | -1.24 | -1.53 | B>B |
| chr20 | 54600000 | 54700000 | -1.32 | -1.57 | B>B |
| chr20 | 54700000 | 54800000 | -1.15 | -1.40 | B>B |
| chr20 | 54800000 | 54900000 | -0.93 | -1.29 | B>B |
| chr20 | 54900000 | 55000000 | -0.08 | -0.31 | B>B |
| chr20 | 55000000 | 55100000 | 0.46 | 0.27 | A>A |
| chr20 | 55100000 | 55200000 | 0.54 | 0.46 | A>A |
| chr20 | 55200000 | 55300000 | 0.79 | 0.76 | A>A |
| chr20 | 55300000 | 55400000 | 0.86 | 0.80 | A>A |
| chr20 | 55400000 | 55500000 | 0.88 | 0.66 | A>A |
| chr20 | 55500000 | 55600000 | 0.70 | 0.54 | A>A |
| chr20 | 55600000 | 55700000 | 0.55 | 0.41 | A>A |
| chr20 | 55700000 | 55800000 | 0.62 | 0.47 | A>A |
| chr20 | 55800000 | 55900000 | 0.63 | 0.44 | A>A |
| chr20 | 55900000 | 56000000 | 0.76 | 0.66 | A>A |
| chr20 | 56000000 | 56100000 | 0.86 | 0.81 | A>A |
| chr20 | 56100000 | 56200000 | 0.58 | 0.61 | A>A |
| chr20 | 56200000 | 56300000 | 0.68 | 0.76 | A>A |
| chr20 | 56300000 | 56400000 | 0.83 | 0.88 | A>A |

| | | | | | |
|---|---|---|---|---|---|
| chr20 | 56400000 | 56500000 | 0.76 | 0.86 | A>A |
| chr20 | 56500000 | 56600000 | 0.66 | 0.84 | A>A |
| chr20 | 56600000 | 56700000 | 0.67 | 0.80 | A>A |
| chr20 | 56700000 | 56800000 | 0.55 | 0.64 | A>A |
| chr20 | 56800000 | 56900000 | 0.63 | 0.74 | A>A |
| chr20 | 56900000 | 57000000 | 0.75 | 0.79 | A>A |
| chr20 | 57000000 | 57100000 | 0.41 | 0.52 | A>A |
| chr20 | 57100000 | 57200000 | 0.40 | 0.61 | A>A |
| chr20 | 57200000 | 57300000 | 0.56 | 0.49 | A>A |
| chr20 | 57300000 | 57400000 | 0.37 | 0.08 | A>A |
| chr20 | 57400000 | 57500000 | 0.46 | 0.14 | A>A |
| chr20 | 57500000 | 57600000 | 0.75 | 0.32 | A>A |
| chr20 | 57600000 | 57700000 | 0.36 | -0.17 | A>A |
| chr20 | 57700000 | 57800000 | 0.13 | -0.52 | A>B |
| chr20 | 57800000 | 57900000 | 0.37 | -0.63 | A>B |
| chr20 | 57900000 | 58000000 | 0.35 | -0.50 | A>B |
| chr20 | 58000000 | 58100000 | 0.18 | -0.28 | B>B |
| chr20 | 58100000 | 58200000 | -0.12 | -0.33 | B>B |
| chr20 | 58200000 | 58300000 | -0.42 | -0.49 | B>B |
| chr20 | 58300000 | 58400000 | -0.45 | -0.13 | B>B |
| chr20 | 58400000 | 58500000 | -0.24 | 0.30 | B>A |
| chr20 | 58500000 | 58600000 | -0.14 | 0.09 | B>B |
| chr20 | 58600000 | 58700000 | -0.34 | -0.22 | B>B |
| chr20 | 58700000 | 58800000 | -0.47 | -0.61 | B>B |
| chr20 | 58800000 | 58900000 | -0.57 | -0.80 | B>B |
| chr20 | 58900000 | 59000000 | -0.82 | -1.31 | B>B |
| chr20 | 59000000 | 59100000 | -0.80 | -1.44 | A>B |
| chr20 | 59100000 | 59200000 | -0.85 | -1.48 | A>B |
| chr20 | 59200000 | 59300000 | -0.86 | -1.54 | A>B |
| chr20 | 59300000 | 59400000 | -0.76 | -1.39 | A>B |
| chr20 | 59400000 | 59500000 | -0.56 | -1.11 | B>B |
| chr20 | 59500000 | 59600000 | -0.31 | -1.06 | A>B |
| chr20 | 59600000 | 59700000 | -0.08 | -0.68 | A>B |
| chr20 | 59700000 | 59800000 | 0.08 | -0.40 | B>B |
| chr20 | 59800000 | 59900000 | 0.07 | -0.31 | B>B |
| chr20 | 59900000 | 60000000 | 0.00 | -0.27 | B>B |
| chr20 | 60000000 | 60100000 | 0.00 | -0.36 | B>B |
| chr20 | 60100000 | 60200000 | -0.06 | -0.23 | B>B |
| chr20 | 60200000 | 60300000 | -0.06 | -0.23 | B>B |
| chr20 | 60300000 | 60400000 | 0.08 | -0.15 | B>B |
| chr20 | 60400000 | 60500000 | 0.16 | 0.08 | A>A |
| chr20 | 60500000 | 60600000 | 0.66 | 0.75 | A>A |
| chr20 | 60600000 | 60700000 | 0.98 | 1.11 | A>A |
| chr20 | 60700000 | 60800000 | 0.98 | 1.15 | A>A |
| chr20 | 60800000 | 60900000 | 1.03 | 1.13 | A>A |
| chr20 | 60900000 | 61000000 | 1.09 | 1.10 | A>A |
| chr20 | 61000000 | 61100000 | 0.96 | 1.01 | A>A |
| chr20 | 61100000 | 61200000 | 0.00 | 0.00 | NO |
| chr20 | 61200000 | 61300000 | 0.00 | 0.00 | NO |
| chr20 | 61300000 | 61400000 | 0.94 | 0.96 | A>A |
| chr20 | 61400000 | 61500000 | 0.95 | 0.93 | A>A |
| chr20 | 61500000 | 61600000 | 1.05 | 0.97 | A>A |
| chr20 | 61600000 | 61700000 | 0.79 | 0.81 | A>A |
| chr20 | 61700000 | 61800000 | 0.84 | 0.85 | A>A |
| chr20 | 61800000 | 61900000 | 0.82 | 0.82 | A>A |
| chr20 | 61900000 | 62000000 | 0.79 | 0.77 | A>A |

| | | | | | |
|---|---|---|---|---|---|
| chr20 | 62000000 | 62100000 | 0.73 | 0.75 | A>A |
| chr20 | 62100000 | 62200000 | 0.66 | 0.71 | A>A |
| chr20 | 62200000 | 62300000 | 0.74 | 0.70 | A>A |
| chr20 | 62300000 | 62400000 | 0.59 | 0.66 | A>A |
| chr20 | 62400000 | 62500000 | 0.74 | 0.70 | A>A |
| chr20 | 62500000 | 62600000 | 0.73 | 0.71 | A>A |
| chr20 | 62600000 | 62700000 | 0.71 | 0.72 | A>A |
| chr20 | 62700000 | 62800000 | 0.77 | 0.75 | A>A |
| chr20 | 62800000 | 62900000 | 0.72 | 0.70 | A>A |
| chr20 | 62900000 | 63000000 | 0.00 | 0.00 | NO |
| chr20 | 63000000 | 63025520 | 0.00 | 0.00 | NO |
| chr21 | 0 | 100000 | 0.00 | 0.00 | NO |
| chr21 | 100000 | 200000 | 0.00 | 0.00 | NO |
| chr21 | 200000 | 300000 | 0.00 | 0.00 | NO |
| chr21 | 300000 | 400000 | 0.00 | 0.00 | NO |
| chr21 | 400000 | 500000 | 0.00 | 0.00 | NO |
| chr21 | 500000 | 600000 | 0.00 | 0.00 | NO |
| chr21 | 600000 | 700000 | 0.00 | 0.00 | NO |
| chr21 | 700000 | 800000 | 0.00 | 0.00 | NO |
| chr21 | 800000 | 900000 | 0.00 | 0.00 | NO |
| chr21 | 900000 | 1000000 | 0.00 | 0.00 | NO |
| chr21 | 1000000 | 1100000 | 0.00 | 0.00 | NO |
| chr21 | 1100000 | 1200000 | 0.00 | 0.00 | NO |
| chr21 | 1200000 | 1300000 | 0.00 | 0.00 | NO |
| chr21 | 1300000 | 1400000 | 0.00 | 0.00 | NO |
| chr21 | 1400000 | 1500000 | 0.00 | 0.00 | NO |
| chr21 | 1500000 | 1600000 | 0.00 | 0.00 | NO |
| chr21 | 1600000 | 1700000 | 0.00 | 0.00 | NO |
| chr21 | 1700000 | 1800000 | 0.00 | 0.00 | NO |
| chr21 | 1800000 | 1900000 | 0.00 | 0.00 | NO |
| chr21 | 1900000 | 2000000 | 0.00 | 0.00 | NO |
| chr21 | 2000000 | 2100000 | 0.00 | 0.00 | NO |
| chr21 | 2100000 | 2200000 | 0.00 | 0.00 | NO |
| chr21 | 2200000 | 2300000 | 0.00 | 0.00 | NO |
| chr21 | 2300000 | 2400000 | 0.00 | 0.00 | NO |
| chr21 | 2400000 | 2500000 | 0.00 | 0.00 | NO |
| chr21 | 2500000 | 2600000 | 0.00 | 0.00 | NO |
| chr21 | 2600000 | 2700000 | 0.00 | 0.00 | NO |
| chr21 | 2700000 | 2800000 | 0.00 | 0.00 | NO |
| chr21 | 2800000 | 2900000 | 0.00 | 0.00 | NO |
| chr21 | 2900000 | 3000000 | 0.00 | 0.00 | NO |
| chr21 | 3000000 | 3100000 | 0.00 | 0.00 | NO |
| chr21 | 3100000 | 3200000 | 0.00 | 0.00 | NO |
| chr21 | 3200000 | 3300000 | 0.00 | 0.00 | NO |
| chr21 | 3300000 | 3400000 | 0.00 | 0.00 | NO |
| chr21 | 3400000 | 3500000 | 0.00 | 0.00 | NO |
| chr21 | 3500000 | 3600000 | 0.00 | 0.00 | NO |
| chr21 | 3600000 | 3700000 | 0.00 | 0.00 | NO |
| chr21 | 3700000 | 3800000 | 0.00 | 0.00 | NO |
| chr21 | 3800000 | 3900000 | 0.00 | 0.00 | NO |
| chr21 | 3900000 | 4000000 | 0.00 | 0.00 | NO |
| chr21 | 4000000 | 4100000 | 0.00 | 0.00 | NO |
| chr21 | 4100000 | 4200000 | 0.00 | 0.00 | NO |
| chr21 | 4200000 | 4300000 | 0.00 | 0.00 | NO |
| chr21 | 4300000 | 4400000 | 0.00 | 0.00 | NO |
| chr21 | 4400000 | 4500000 | 0.00 | 0.00 | NO |

| | | | | | |
|---|---|---|---|---|---|
| chr21 | 4500000 | 4600000 | 0.00 | 0.00 | NO |
| chr21 | 4600000 | 4700000 | 0.00 | 0.00 | NO |
| chr21 | 4700000 | 4800000 | 0.00 | 0.00 | NO |
| chr21 | 4800000 | 4900000 | 0.00 | 0.00 | NO |
| chr21 | 4900000 | 5000000 | 0.00 | 0.00 | NO |
| chr21 | 5000000 | 5100000 | 0.00 | 0.00 | NO |
| chr21 | 5100000 | 5200000 | 0.00 | 0.00 | NO |
| chr21 | 5200000 | 5300000 | 0.00 | 0.00 | NO |
| chr21 | 5300000 | 5400000 | 0.00 | 0.00 | NO |
| chr21 | 5400000 | 5500000 | 0.00 | 0.00 | NO |
| chr21 | 5500000 | 5600000 | 0.00 | 0.00 | NO |
| chr21 | 5600000 | 5700000 | 0.00 | 0.00 | NO |
| chr21 | 5700000 | 5800000 | 0.00 | 0.00 | NO |
| chr21 | 5800000 | 5900000 | 0.00 | 0.00 | NO |
| chr21 | 5900000 | 6000000 | 0.00 | 0.00 | NO |
| chr21 | 6000000 | 6100000 | 0.00 | 0.00 | NO |
| chr21 | 6100000 | 6200000 | 0.00 | 0.00 | NO |
| chr21 | 6200000 | 6300000 | 0.00 | 0.00 | NO |
| chr21 | 6300000 | 6400000 | 0.00 | 0.00 | NO |
| chr21 | 6400000 | 6500000 | 0.00 | 0.00 | NO |
| chr21 | 6500000 | 6600000 | 0.00 | 0.00 | NO |
| chr21 | 6600000 | 6700000 | 0.00 | 0.00 | NO |
| chr21 | 6700000 | 6800000 | 0.00 | 0.00 | NO |
| chr21 | 6800000 | 6900000 | 0.00 | 0.00 | NO |
| chr21 | 6900000 | 7000000 | 0.00 | 0.00 | NO |
| chr21 | 7000000 | 7100000 | 0.00 | 0.00 | NO |
| chr21 | 7100000 | 7200000 | 0.00 | 0.00 | NO |
| chr21 | 7200000 | 7300000 | 0.00 | 0.00 | NO |
| chr21 | 7300000 | 7400000 | 0.00 | 0.00 | NO |
| chr21 | 7400000 | 7500000 | 0.00 | 0.00 | NO |
| chr21 | 7500000 | 7600000 | 0.00 | 0.00 | NO |
| chr21 | 7600000 | 7700000 | 0.00 | 0.00 | NO |
| chr21 | 7700000 | 7800000 | 0.00 | 0.00 | NO |
| chr21 | 7800000 | 7900000 | 0.00 | 0.00 | NO |
| chr21 | 7900000 | 8000000 | 0.00 | 0.00 | NO |
| chr21 | 8000000 | 8100000 | 0.00 | 0.00 | NO |
| chr21 | 8100000 | 8200000 | 0.00 | 0.00 | NO |
| chr21 | 8200000 | 8300000 | 0.00 | 0.00 | NO |
| chr21 | 8300000 | 8400000 | 0.00 | 0.00 | NO |
| chr21 | 8400000 | 8500000 | 0.00 | 0.00 | NO |
| chr21 | 8500000 | 8600000 | 0.00 | 0.00 | NO |
| chr21 | 8600000 | 8700000 | 0.00 | 0.00 | NO |
| chr21 | 8700000 | 8800000 | 0.00 | 0.00 | NO |
| chr21 | 8800000 | 8900000 | 0.00 | 0.00 | NO |
| chr21 | 8900000 | 9000000 | 0.00 | 0.00 | NO |
| chr21 | 9000000 | 9100000 | 0.00 | 0.00 | NO |
| chr21 | 9100000 | 9200000 | 0.00 | 0.00 | NO |
| chr21 | 9200000 | 9300000 | 0.00 | 0.00 | NO |
| chr21 | 9300000 | 9400000 | 0.00 | 0.00 | NO |
| chr21 | 9400000 | 9500000 | 0.00 | 0.00 | NO |
| chr21 | 9500000 | 9600000 | 0.00 | 0.00 | NO |
| chr21 | 9600000 | 9700000 | 0.00 | 0.00 | NO |
| chr21 | 9700000 | 9800000 | 0.00 | 0.00 | NO |
| chr21 | 9800000 | 9900000 | 0.00 | 0.00 | NO |
| chr21 | 9900000 | 10000000 | 0.00 | 0.00 | NO |
| chr21 | 10000000 | 10100000 | 0.00 | 0.00 | NO |

| | | | | | |
|---|---|---|---|---|---|
| chr21 | 10100000 | 10200000 | 0.00 | 0.00 | NO |
| chr21 | 10200000 | 10300000 | 0.00 | 0.00 | NO |
| chr21 | 10300000 | 10400000 | 0.00 | 0.00 | NO |
| chr21 | 10400000 | 10500000 | 0.00 | 0.00 | NO |
| chr21 | 10500000 | 10600000 | 0.00 | 0.00 | NO |
| chr21 | 10600000 | 10700000 | 0.00 | 0.00 | NO |
| chr21 | 10700000 | 10800000 | -0.91 | -0.85 | B>B |
| chr21 | 10800000 | 10900000 | 0.00 | 0.00 | NO |
| chr21 | 10900000 | 11000000 | 0.00 | 0.00 | NO |
| chr21 | 11000000 | 11100000 | -0.98 | -1.00 | B>B |
| chr21 | 11100000 | 11200000 | 0.00 | 0.00 | NO |
| chr21 | 11200000 | 11300000 | 0.00 | 0.00 | NO |
| chr21 | 11300000 | 11400000 | 0.00 | 0.00 | NO |
| chr21 | 11400000 | 11500000 | 0.00 | 0.00 | NO |
| chr21 | 11500000 | 11600000 | 0.00 | 0.00 | NO |
| chr21 | 11600000 | 11700000 | 0.00 | 0.00 | NO |
| chr21 | 11700000 | 11800000 | 0.00 | 0.00 | NO |
| chr21 | 11800000 | 11900000 | 0.00 | 0.00 | NO |
| chr21 | 11900000 | 12000000 | 0.00 | 0.00 | NO |
| chr21 | 12000000 | 12100000 | 0.00 | 0.00 | NO |
| chr21 | 12100000 | 12200000 | 0.00 | 0.00 | NO |
| chr21 | 12200000 | 12300000 | 0.00 | 0.00 | NO |
| chr21 | 12300000 | 12400000 | 0.00 | 0.00 | NO |
| chr21 | 12400000 | 12500000 | 0.00 | 0.00 | NO |
| chr21 | 12500000 | 12600000 | 0.00 | 0.00 | NO |
| chr21 | 12600000 | 12700000 | 0.00 | 0.00 | NO |
| chr21 | 12700000 | 12800000 | 0.00 | 0.00 | NO |
| chr21 | 12800000 | 12900000 | 0.00 | 0.00 | NO |
| chr21 | 12900000 | 13000000 | 0.00 | 0.00 | NO |
| chr21 | 13000000 | 13100000 | 0.00 | 0.00 | NO |
| chr21 | 13100000 | 13200000 | 0.00 | 0.00 | NO |
| chr21 | 13200000 | 13300000 | 0.00 | 0.00 | NO |
| chr21 | 13300000 | 13400000 | 0.00 | 0.00 | NO |
| chr21 | 13400000 | 13500000 | 0.00 | 0.00 | NO |
| chr21 | 13500000 | 13600000 | 0.00 | 0.00 | NO |
| chr21 | 13600000 | 13700000 | 0.00 | 0.00 | NO |
| chr21 | 13700000 | 13800000 | 0.00 | 0.00 | NO |
| chr21 | 13800000 | 13900000 | 0.00 | 0.00 | NO |
| chr21 | 13900000 | 14000000 | 0.00 | 0.00 | NO |
| chr21 | 14000000 | 14100000 | 0.00 | 0.00 | NO |
| chr21 | 14100000 | 14200000 | 0.00 | 0.00 | NO |
| chr21 | 14200000 | 14300000 | 0.00 | 0.00 | NO |
| chr21 | 14300000 | 14400000 | 0.00 | 0.00 | NO |
| chr21 | 14400000 | 14500000 | 0.00 | 0.00 | NO |
| chr21 | 14500000 | 14600000 | 0.00 | 0.00 | NO |
| chr21 | 14600000 | 14700000 | 0.00 | 0.00 | NO |
| chr21 | 14700000 | 14800000 | 0.00 | 0.00 | NO |
| chr21 | 14800000 | 14900000 | 0.00 | 0.00 | NO |
| chr21 | 14900000 | 15000000 | 0.00 | 0.00 | NO |
| chr21 | 15000000 | 15100000 | 0.00 | 0.00 | NO |
| chr21 | 15100000 | 15200000 | 0.00 | 0.00 | NO |
| chr21 | 15200000 | 15300000 | 0.00 | 0.00 | NO |
| chr21 | 15300000 | 15400000 | 0.00 | 0.00 | NO |
| chr21 | 15400000 | 15500000 | 0.00 | 0.00 | NO |
| chr21 | 15500000 | 15600000 | -0.24 | -0.52 | B>B |
| chr21 | 15600000 | 15700000 | 0.07 | -0.03 | A>A |

| | | | | | |
|---|---|---|---|---|---|
| chr21 | 15700000 | 15800000 | 0.55 | 0.29 | A>A |
| chr21 | 15800000 | 15900000 | 0.36 | 0.46 | A>A |
| chr21 | 15900000 | 16000000 | 0.46 | 0.41 | A>A |
| chr21 | 16000000 | 16100000 | 0.52 | 0.56 | A>A |
| chr21 | 16100000 | 16200000 | 0.66 | 0.50 | A>A |
| chr21 | 16200000 | 16300000 | 0.85 | 0.69 | A>A |
| chr21 | 16300000 | 16400000 | 0.99 | 0.80 | A>A |
| chr21 | 16400000 | 16500000 | 1.07 | 0.91 | A>A |
| chr21 | 16500000 | 16600000 | 1.19 | 0.99 | A>A |
| chr21 | 16600000 | 16700000 | 1.29 | 0.89 | A>A |
| chr21 | 16700000 | 16800000 | 1.24 | 1.01 | A>A |
| chr21 | 16800000 | 16900000 | 0.82 | 0.74 | A>A |
| chr21 | 16900000 | 17000000 | 0.72 | 0.61 | A>A |
| chr21 | 17000000 | 17100000 | 0.95 | 0.86 | A>A |
| chr21 | 17100000 | 17200000 | 0.99 | 0.82 | A>A |
| chr21 | 17200000 | 17300000 | 0.78 | 0.54 | A>A |
| chr21 | 17300000 | 17400000 | 0.51 | 0.15 | A>A |
| chr21 | 17400000 | 17500000 | 0.69 | 0.35 | A>A |
| chr21 | 17500000 | 17600000 | 0.97 | 0.67 | A>A |
| chr21 | 17600000 | 17700000 | 0.69 | 0.50 | A>A |
| chr21 | 17700000 | 17800000 | 0.57 | 0.37 | A>A |
| chr21 | 17800000 | 17900000 | 0.56 | 0.26 | A>A |
| chr21 | 17900000 | 18000000 | 0.68 | 0.43 | A>A |
| chr21 | 18000000 | 18100000 | 0.26 | 0.08 | A>A |
| chr21 | 18100000 | 18200000 | -0.02 | -0.06 | B>B |
| chr21 | 18200000 | 18300000 | -0.15 | -0.18 | B>B |
| chr21 | 18300000 | 18400000 | -0.30 | -0.23 | B>B |
| chr21 | 18400000 | 18500000 | -0.26 | -0.37 | B>B |
| chr21 | 18500000 | 18600000 | -0.46 | -0.53 | B>B |
| chr21 | 18600000 | 18700000 | -0.25 | -0.50 | B>B |
| chr21 | 18700000 | 18800000 | 0.27 | -0.14 | A>A |
| chr21 | 18800000 | 18900000 | 1.14 | 0.56 | A>A |
| chr21 | 18900000 | 19000000 | 0.67 | 0.10 | A>A |
| chr21 | 19000000 | 19100000 | -0.27 | -0.75 | B>B |
| chr21 | 19100000 | 19200000 | -0.18 | -0.54 | B>B |
| chr21 | 19200000 | 19300000 | -0.48 | -0.78 | B>B |
| chr21 | 19300000 | 19400000 | 0.00 | 0.00 | NO |
| chr21 | 19400000 | 19500000 | 0.00 | 0.00 | NO |
| chr21 | 19500000 | 19600000 | 0.00 | -1.56 | NO |
| chr21 | 19600000 | 19700000 | -1.27 | -1.48 | B>B |
| chr21 | 19700000 | 19800000 | -1.45 | -1.49 | B>B |
| chr21 | 19800000 | 19900000 | -1.33 | -1.49 | B>B |
| chr21 | 19900000 | 20000000 | -1.28 | -1.41 | B>B |
| chr21 | 20000000 | 20100000 | -1.22 | -1.35 | B>B |
| chr21 | 20100000 | 20200000 | -1.17 | -1.31 | B>B |
| chr21 | 20200000 | 20300000 | -1.14 | -1.28 | B>B |
| chr21 | 20300000 | 20400000 | -1.18 | -1.25 | B>B |
| chr21 | 20400000 | 20500000 | -1.05 | -1.22 | B>B |
| chr21 | 20500000 | 20600000 | -1.06 | -1.15 | B>B |
| chr21 | 20600000 | 20700000 | -1.01 | -1.12 | B>B |
| chr21 | 20700000 | 20800000 | -1.04 | -1.14 | B>B |
| chr21 | 20800000 | 20900000 | -1.02 | -1.11 | B>B |
| chr21 | 20900000 | 21000000 | -1.01 | -1.11 | B>B |
| chr21 | 21000000 | 21100000 | -1.01 | -1.07 | B>B |
| chr21 | 21100000 | 21200000 | -1.00 | -1.08 | B>B |
| chr21 | 21200000 | 21300000 | -0.98 | -1.06 | B>B |

| | | | | | |
|---|---|---|---|---|---|
| chr21 | 21300000 | 21400000 | -0.98 | -1.03 | B>B |
| chr21 | 21400000 | 21500000 | -0.91 | -1.01 | B>B |
| chr21 | 21500000 | 21600000 | -0.95 | -1.05 | B>B |
| chr21 | 21600000 | 21700000 | -0.92 | -1.01 | B>B |
| chr21 | 21700000 | 21800000 | -0.95 | -1.00 | B>B |
| chr21 | 21800000 | 21900000 | -0.95 | -1.04 | B>B |
| chr21 | 21900000 | 22000000 | -0.93 | -1.08 | B>B |
| chr21 | 22000000 | 22100000 | -0.97 | -1.05 | B>B |
| chr21 | 22100000 | 22200000 | -1.01 | -1.08 | B>B |
| chr21 | 22200000 | 22300000 | -1.01 | -1.12 | B>B |
| chr21 | 22300000 | 22400000 | -0.99 | -1.11 | B>B |
| chr21 | 22400000 | 22500000 | -1.00 | -1.07 | B>B |
| chr21 | 22500000 | 22600000 | -0.91 | -1.05 | B>B |
| chr21 | 22600000 | 22700000 | -0.94 | -0.98 | B>B |
| chr21 | 22700000 | 22800000 | -0.89 | -0.96 | B>B |
| chr21 | 22800000 | 22900000 | -0.90 | -0.99 | B>B |
| chr21 | 22900000 | 23000000 | -0.87 | -0.97 | B>B |
| chr21 | 23000000 | 23100000 | -0.84 | -0.94 | B>B |
| chr21 | 23100000 | 23200000 | -0.82 | -0.96 | B>B |
| chr21 | 23200000 | 23300000 | -0.84 | -0.92 | B>B |
| chr21 | 23300000 | 23400000 | -0.85 | -0.92 | B>B |
| chr21 | 23400000 | 23500000 | -0.85 | -0.93 | B>B |
| chr21 | 23500000 | 23600000 | -0.85 | -0.90 | B>B |
| chr21 | 23600000 | 23700000 | -0.87 | -0.96 | B>B |
| chr21 | 23700000 | 23800000 | -0.86 | -0.89 | B>B |
| chr21 | 23800000 | 23900000 | -0.84 | -0.95 | B>B |
| chr21 | 23900000 | 24000000 | -0.87 | -0.93 | B>B |
| chr21 | 24000000 | 24100000 | -0.90 | -0.99 | B>B |
| chr21 | 24100000 | 24200000 | -0.94 | -1.00 | B>B |
| chr21 | 24200000 | 24300000 | -0.92 | -1.06 | B>B |
| chr21 | 24300000 | 24400000 | -0.95 | -1.03 | B>B |
| chr21 | 24400000 | 24500000 | -0.93 | -1.00 | B>B |
| chr21 | 24500000 | 24600000 | -0.94 | -1.02 | B>B |
| chr21 | 24600000 | 24700000 | -0.94 | -1.01 | B>B |
| chr21 | 24700000 | 24800000 | -0.92 | -0.99 | B>B |
| chr21 | 24800000 | 24900000 | -0.93 | -0.97 | B>B |
| chr21 | 24900000 | 25000000 | -0.92 | -1.04 | B>B |
| chr21 | 25000000 | 25100000 | -0.94 | -1.05 | B>B |
| chr21 | 25100000 | 25200000 | -0.89 | -1.02 | B>B |
| chr21 | 25200000 | 25300000 | -0.86 | -1.01 | B>B |
| chr21 | 25300000 | 25400000 | -0.90 | -1.03 | B>B |
| chr21 | 25400000 | 25500000 | -0.92 | -1.00 | B>B |
| chr21 | 25500000 | 25600000 | -0.94 | -1.01 | B>B |
| chr21 | 25600000 | 25700000 | -0.92 | -0.98 | B>B |
| chr21 | 25700000 | 25800000 | -0.96 | -1.08 | B>B |
| chr21 | 25800000 | 25900000 | -0.97 | -1.06 | B>B |
| chr21 | 25900000 | 26000000 | -0.94 | -1.11 | B>B |
| chr21 | 26000000 | 26100000 | -1.03 | -1.08 | B>B |
| chr21 | 26100000 | 26200000 | -1.03 | -1.11 | B>B |
| chr21 | 26200000 | 26300000 | -1.03 | -1.14 | B>B |
| chr21 | 26300000 | 26400000 | -1.05 | -1.16 | B>B |
| chr21 | 26400000 | 26500000 | -1.09 | -1.25 | B>B |
| chr21 | 26500000 | 26600000 | -1.18 | -1.34 | B>B |
| chr21 | 26600000 | 26700000 | -1.17 | -1.33 | B>B |
| chr21 | 26700000 | 26800000 | -1.16 | -1.51 | B>B |
| chr21 | 26800000 | 26900000 | -0.92 | -1.44 | B>B |

| | | | | | |
|---|---|---|---|---|---|
| chr21 | 26900000 | 27000000 | -0.04 | -0.70 | A>B |
| chr21 | 27000000 | 27100000 | 0.25 | -0.25 | A>A |
| chr21 | 27100000 | 27200000 | 0.50 | 0.11 | A>A |
| chr21 | 27200000 | 27300000 | 0.39 | 0.24 | A>A |
| chr21 | 27300000 | 27400000 | 0.76 | 0.62 | A>A |
| chr21 | 27400000 | 27500000 | 0.87 | 0.63 | A>A |
| chr21 | 27500000 | 27600000 | 0.63 | 0.48 | A>A |
| chr21 | 27600000 | 27700000 | 0.24 | 0.36 | A>A |
| chr21 | 27700000 | 27800000 | 0.25 | 0.29 | A>A |
| chr21 | 27800000 | 27900000 | 0.07 | 0.28 | A>A |
| chr21 | 27900000 | 28000000 | -0.02 | 0.24 | A>A |
| chr21 | 28000000 | 28100000 | -0.23 | 0.05 | B>B |
| chr21 | 28100000 | 28200000 | -0.37 | -0.43 | B>B |
| chr21 | 28200000 | 28300000 | -0.59 | -0.79 | B>B |
| chr21 | 28300000 | 28400000 | -0.88 | -1.12 | B>B |
| chr21 | 28400000 | 28500000 | -0.93 | -1.14 | B>B |
| chr21 | 28500000 | 28600000 | -1.02 | -1.16 | B>B |
| chr21 | 28600000 | 28700000 | -1.07 | -1.08 | B>B |
| chr21 | 28700000 | 28800000 | -0.95 | -1.10 | B>B |
| chr21 | 28800000 | 28900000 | -1.03 | -1.11 | B>B |
| chr21 | 28900000 | 29000000 | -0.98 | -1.08 | B>B |
| chr21 | 29000000 | 29100000 | -1.00 | -1.06 | B>B |
| chr21 | 29100000 | 29200000 | -0.92 | -1.15 | B>B |
| chr21 | 29200000 | 29300000 | 0.00 | 0.00 | NO |
| chr21 | 29300000 | 29400000 | 0.00 | 0.00 | NO |
| chr21 | 29400000 | 29500000 | -0.99 | -1.14 | B>B |
| chr21 | 29500000 | 29600000 | -1.10 | -1.17 | B>B |
| chr21 | 29600000 | 29700000 | -1.04 | -1.18 | B>B |
| chr21 | 29700000 | 29800000 | -0.94 | -1.18 | B>B |
| chr21 | 29800000 | 29900000 | -0.79 | -1.22 | B>B |
| chr21 | 29900000 | 30000000 | -0.71 | -1.16 | B>B |
| chr21 | 30000000 | 30100000 | -0.61 | -1.09 | B>B |
| chr21 | 30100000 | 30200000 | -0.31 | -0.82 | B>B |
| chr21 | 30200000 | 30300000 | 0.43 | -0.21 | A>B |
| chr21 | 30300000 | 30400000 | 0.85 | 0.35 | A>A |
| chr21 | 30400000 | 30500000 | 0.79 | 0.53 | A>A |
| chr21 | 30500000 | 30600000 | 0.81 | 0.63 | A>A |
| chr21 | 30600000 | 30700000 | 0.87 | 0.65 | A>A |
| chr21 | 30700000 | 30800000 | 0.60 | 0.38 | A>A |
| chr21 | 30800000 | 30900000 | 0.23 | -0.08 | A>A |
| chr21 | 30900000 | 31000000 | -0.05 | -0.65 | A>B |
| chr21 | 31000000 | 31100000 | -0.08 | -0.93 | A>B |
| chr21 | 31100000 | 31200000 | -0.18 | -0.98 | A>B |
| chr21 | 31200000 | 31300000 | -0.42 | -1.01 | A>B |
| chr21 | 31300000 | 31400000 | -0.69 | -1.11 | B>B |
| chr21 | 31400000 | 31500000 | -0.76 | -1.03 | B>B |
| chr21 | 31500000 | 31600000 | -0.69 | -1.13 | B>B |
| chr21 | 31600000 | 31700000 | -0.83 | -1.16 | B>B |
| chr21 | 31700000 | 31800000 | -0.99 | -1.10 | B>B |
| chr21 | 31800000 | 31900000 | -1.17 | -1.19 | B>B |
| chr21 | 31900000 | 32000000 | -1.31 | -1.25 | B>B |
| chr21 | 32000000 | 32100000 | -1.33 | -1.35 | B>B |
| chr21 | 32100000 | 32200000 | -1.20 | -1.19 | B>B |
| chr21 | 32200000 | 32300000 | -1.12 | -1.08 | B>B |
| chr21 | 32300000 | 32400000 | -0.89 | -0.90 | B>B |
| chr21 | 32400000 | 32500000 | -0.23 | -0.34 | B>B |

| | | | | | |
|---|---|---|---|---|---|
| chr21 | 32500000 | 32600000 | 0.39 | 0.25 | A>A |
| chr21 | 32600000 | 32700000 | 0.65 | 0.57 | A>A |
| chr21 | 32700000 | 32800000 | 0.67 | 0.58 | A>A |
| chr21 | 32800000 | 32900000 | 0.73 | 0.52 | A>A |
| chr21 | 32900000 | 33000000 | 0.63 | 0.53 | A>A |
| chr21 | 33000000 | 33100000 | 0.82 | 0.58 | A>A |
| chr21 | 33100000 | 33200000 | 0.63 | 0.28 | A>A |
| chr21 | 33200000 | 33300000 | 0.64 | 0.20 | A>A |
| chr21 | 33300000 | 33400000 | 0.53 | -0.07 | A>B |
| chr21 | 33400000 | 33500000 | 0.59 | -0.15 | A>B |
| chr21 | 33500000 | 33600000 | 0.49 | -0.07 | A>A |
| chr21 | 33600000 | 33700000 | 0.63 | 0.44 | A>A |
| chr21 | 33700000 | 33800000 | 0.74 | 0.70 | A>A |
| chr21 | 33800000 | 33900000 | 0.85 | 0.74 | A>A |
| chr21 | 33900000 | 34000000 | 0.68 | 0.68 | A>A |
| chr21 | 34000000 | 34100000 | 0.72 | 0.64 | A>A |
| chr21 | 34100000 | 34200000 | 0.72 | 0.67 | A>A |
| chr21 | 34200000 | 34300000 | 0.50 | 0.45 | A>A |
| chr21 | 34300000 | 34400000 | 0.52 | 0.44 | A>A |
| chr21 | 34400000 | 34500000 | 0.62 | 0.47 | A>A |
| chr21 | 34500000 | 34600000 | 0.55 | 0.47 | A>A |
| chr21 | 34600000 | 34700000 | 0.70 | 0.69 | A>A |
| chr21 | 34700000 | 34800000 | 0.71 | 0.68 | A>A |
| chr21 | 34800000 | 34900000 | 0.65 | 0.68 | A>A |
| chr21 | 34900000 | 35000000 | 0.73 | 0.73 | A>A |
| chr21 | 35000000 | 35100000 | 0.71 | 0.72 | A>A |
| chr21 | 35100000 | 35200000 | 0.79 | 0.70 | A>A |
| chr21 | 35200000 | 35300000 | 0.63 | 0.61 | A>A |
| chr21 | 35300000 | 35400000 | 0.67 | 0.56 | A>A |
| chr21 | 35400000 | 35500000 | 0.58 | 0.57 | A>A |
| chr21 | 35500000 | 35600000 | 0.49 | 0.47 | A>A |
| chr21 | 35600000 | 35700000 | 0.42 | 0.38 | A>A |
| chr21 | 35700000 | 35800000 | 0.54 | 0.50 | A>A |
| chr21 | 35800000 | 35900000 | 0.58 | 0.46 | A>A |
| chr21 | 35900000 | 36000000 | 0.54 | 0.46 | A>A |
| chr21 | 36000000 | 36100000 | 0.38 | 0.41 | A>A |
| chr21 | 36100000 | 36200000 | 0.53 | 0.48 | A>A |
| chr21 | 36200000 | 36300000 | 0.56 | 0.52 | A>A |
| chr21 | 36300000 | 36400000 | 0.48 | 0.47 | A>A |
| chr21 | 36400000 | 36500000 | 0.33 | 0.43 | A>A |
| chr21 | 36500000 | 36600000 | 0.24 | 0.48 | A>A |
| chr21 | 36600000 | 36700000 | 0.34 | 0.47 | A>A |
| chr21 | 36700000 | 36800000 | 0.21 | 0.36 | A>A |
| chr21 | 36800000 | 36900000 | -0.01 | 0.22 | A>A |
| chr21 | 36900000 | 37000000 | -0.07 | 0.07 | B>B |
| chr21 | 37000000 | 37100000 | -0.04 | 0.06 | A>A |
| chr21 | 37100000 | 37200000 | 0.08 | 0.19 | A>A |
| chr21 | 37200000 | 37300000 | 0.25 | 0.24 | A>A |
| chr21 | 37300000 | 37400000 | 0.41 | 0.40 | A>A |
| chr21 | 37400000 | 37500000 | 0.66 | 0.58 | A>A |
| chr21 | 37500000 | 37600000 | 0.79 | 0.68 | A>A |
| chr21 | 37600000 | 37700000 | 0.76 | 0.67 | A>A |
| chr21 | 37700000 | 37800000 | 0.67 | 0.63 | A>A |
| chr21 | 37800000 | 37900000 | 0.63 | 0.62 | A>A |
| chr21 | 37900000 | 38000000 | 0.61 | 0.58 | A>A |
| chr21 | 38000000 | 38100000 | 0.57 | 0.57 | A>A |

| | | | | | |
|---|---|---|---|---|---|
| chr21 | 38100000 | 38200000 | 0.64 | 0.59 | A>A |
| chr21 | 38200000 | 38300000 | 0.60 | 0.60 | A>A |
| chr21 | 38300000 | 38400000 | 0.63 | 0.57 | A>A |
| chr21 | 38400000 | 38500000 | 0.70 | 0.61 | A>A |
| chr21 | 38500000 | 38600000 | 0.61 | 0.59 | A>A |
| chr21 | 38600000 | 38700000 | 0.64 | 0.59 | A>A |
| chr21 | 38700000 | 38800000 | 0.68 | 0.63 | A>A |
| chr21 | 38800000 | 38900000 | 0.64 | 0.60 | A>A |
| chr21 | 38900000 | 39000000 | 0.62 | 0.58 | A>A |
| chr21 | 39000000 | 39100000 | 0.44 | 0.50 | A>A |
| chr21 | 39100000 | 39200000 | 0.41 | 0.49 | A>A |
| chr21 | 39200000 | 39300000 | 0.41 | 0.52 | A>A |
| chr21 | 39300000 | 39400000 | 0.55 | 0.56 | A>A |
| chr21 | 39400000 | 39500000 | 0.47 | 0.52 | A>A |
| chr21 | 39500000 | 39600000 | 0.39 | 0.53 | A>A |
| chr21 | 39600000 | 39700000 | 0.55 | 0.56 | A>A |
| chr21 | 39700000 | 39800000 | 0.45 | 0.48 | A>A |
| chr21 | 39800000 | 39900000 | 0.39 | 0.50 | A>A |
| chr21 | 39900000 | 40000000 | 0.32 | 0.44 | A>A |
| chr21 | 40000000 | 40100000 | 0.35 | 0.47 | A>A |
| chr21 | 40100000 | 40200000 | 0.54 | 0.58 | A>A |
| chr21 | 40200000 | 40300000 | 0.57 | 0.58 | A>A |
| chr21 | 40300000 | 40400000 | 0.51 | 0.64 | A>A |
| chr21 | 40400000 | 40500000 | 0.48 | 0.53 | A>A |
| chr21 | 40500000 | 40600000 | 0.57 | 0.63 | A>A |
| chr21 | 40600000 | 40700000 | 0.88 | 0.74 | A>A |
| chr21 | 40700000 | 40800000 | 0.84 | 0.64 | A>A |
| chr21 | 40800000 | 40900000 | 0.50 | 0.37 | A>A |
| chr21 | 40900000 | 41000000 | 0.58 | 0.32 | A>A |
| chr21 | 41000000 | 41100000 | 0.50 | 0.42 | A>A |
| chr21 | 41100000 | 41200000 | -0.02 | -0.57 | B>B |
| chr21 | 41200000 | 41300000 | -0.18 | -1.37 | A>B |
| chr21 | 41300000 | 41400000 | -0.64 | -1.72 | A>B |
| chr21 | 41400000 | 41500000 | -0.52 | -1.46 | A>B |
| chr21 | 41500000 | 41600000 | -0.67 | -1.28 | A>B |
| chr21 | 41600000 | 41700000 | -0.99 | -1.81 | A>B |
| chr21 | 41700000 | 41800000 | -0.56 | -1.45 | A>B |
| chr21 | 41800000 | 41900000 | -0.16 | -0.71 | B>B |
| chr21 | 41900000 | 42000000 | -0.31 | -1.25 | A>B |
| chr21 | 42000000 | 42100000 | -0.50 | -1.64 | A>B |
| chr21 | 42100000 | 42200000 | -0.64 | -1.64 | A>B |
| chr21 | 42200000 | 42300000 | -0.43 | -1.72 | A>B |
| chr21 | 42300000 | 42400000 | -0.42 | -1.58 | A>B |
| chr21 | 42400000 | 42500000 | 0.13 | -0.70 | A>B |
| chr21 | 42500000 | 42600000 | 0.60 | 0.29 | A>A |
| chr21 | 42600000 | 42700000 | 0.60 | 0.34 | A>A |
| chr21 | 42700000 | 42800000 | 0.55 | 0.33 | A>A |
| chr21 | 42800000 | 42900000 | 0.60 | 0.50 | A>A |
| chr21 | 42900000 | 43000000 | 0.76 | 0.58 | A>A |
| chr21 | 43000000 | 43100000 | 0.70 | 0.66 | A>A |
| chr21 | 43100000 | 43200000 | 0.81 | 0.79 | A>A |
| chr21 | 43200000 | 43300000 | 0.62 | 0.62 | A>A |
| chr21 | 43300000 | 43400000 | 0.72 | 0.71 | A>A |
| chr21 | 43400000 | 43500000 | 0.67 | 0.69 | A>A |
| chr21 | 43500000 | 43600000 | 0.56 | 0.59 | A>A |
| chr21 | 43600000 | 43700000 | 0.66 | 0.67 | A>A |

| | | | | | |
|---|---|---|---|---|---|
| chr21 | 43700000 | 43800000 | 0.58 | 0.64 | A>A |
| chr21 | 43800000 | 43900000 | 0.47 | 0.64 | A>A |
| chr21 | 43900000 | 44000000 | 0.57 | 0.65 | A>A |
| chr21 | 44000000 | 44100000 | 0.53 | 0.60 | A>A |
| chr21 | 44100000 | 44200000 | 0.58 | 0.64 | A>A |
| chr21 | 44200000 | 44300000 | 0.51 | 0.59 | A>A |
| chr21 | 44300000 | 44400000 | 0.47 | 0.52 | A>A |
| chr21 | 44400000 | 44500000 | 0.61 | 0.55 | A>A |
| chr21 | 44500000 | 44600000 | 0.33 | 0.50 | A>A |
| chr21 | 44600000 | 44700000 | 0.00 | 0.00 | NO |
| chr21 | 44700000 | 44800000 | 0.37 | 0.44 | A>A |
| chr21 | 44800000 | 44900000 | 0.39 | 0.48 | A>A |
| chr21 | 44900000 | 45000000 | 0.46 | 0.48 | A>A |
| chr21 | 45000000 | 45100000 | 0.41 | 0.51 | A>A |
| chr21 | 45100000 | 45200000 | 0.54 | 0.58 | A>A |
| chr21 | 45200000 | 45300000 | 0.48 | 0.54 | A>A |
| chr21 | 45300000 | 45400000 | 0.56 | 0.54 | A>A |
| chr21 | 45400000 | 45500000 | 0.48 | 0.50 | A>A |
| chr21 | 45500000 | 45600000 | 0.45 | 0.53 | A>A |
| chr21 | 45600000 | 45700000 | 0.49 | 0.49 | A>A |
| chr21 | 45700000 | 45800000 | 0.49 | 0.49 | A>A |
| chr21 | 45800000 | 45900000 | 0.51 | 0.50 | A>A |
| chr21 | 45900000 | 46000000 | 0.38 | 0.49 | A>A |
| chr21 | 46000000 | 46100000 | 0.38 | 0.50 | A>A |
| chr21 | 46100000 | 46200000 | 0.35 | 0.50 | A>A |
| chr21 | 46200000 | 46300000 | 0.45 | 0.52 | A>A |
| chr21 | 46300000 | 46400000 | 0.45 | 0.54 | A>A |
| chr21 | 46400000 | 46500000 | 0.40 | 0.54 | A>A |
| chr21 | 46500000 | 46600000 | 0.41 | 0.53 | A>A |
| chr21 | 46600000 | 46700000 | 0.43 | 0.43 | A>A |
| chr21 | 46700000 | 46800000 | 0.43 | 0.52 | A>A |
| chr21 | 46800000 | 46900000 | 0.46 | 0.56 | A>A |
| chr21 | 46900000 | 47000000 | 0.38 | 0.46 | A>A |
| chr21 | 47000000 | 47100000 | 0.29 | 0.44 | A>A |
| chr21 | 47100000 | 47200000 | 0.31 | 0.41 | A>A |
| chr21 | 47200000 | 47300000 | 0.35 | 0.46 | A>A |
| chr21 | 47300000 | 47400000 | 0.43 | 0.47 | A>A |
| chr21 | 47400000 | 47500000 | 0.50 | 0.52 | A>A |
| chr21 | 47500000 | 47600000 | 0.45 | 0.54 | A>A |
| chr21 | 47600000 | 47700000 | 0.47 | 0.55 | A>A |
| chr21 | 47700000 | 47800000 | 0.55 | 0.58 | A>A |
| chr21 | 47800000 | 47900000 | 0.58 | 0.56 | A>A |
| chr21 | 47900000 | 48000000 | 0.57 | 0.55 | A>A |
| chr21 | 48000000 | 48100000 | 0.38 | 0.35 | A>A |
| chr21 | 48100000 | 48129895 | 0.00 | 0.00 | NO |
| chr22 | 0 | 100000 | 0.00 | 0.00 | NO |
| chr22 | 100000 | 200000 | 0.00 | 0.00 | NO |
| chr22 | 200000 | 300000 | 0.00 | 0.00 | NO |
| chr22 | 300000 | 400000 | 0.00 | 0.00 | NO |
| chr22 | 400000 | 500000 | 0.00 | 0.00 | NO |
| chr22 | 500000 | 600000 | 0.00 | 0.00 | NO |
| chr22 | 600000 | 700000 | 0.00 | 0.00 | NO |
| chr22 | 700000 | 800000 | 0.00 | 0.00 | NO |
| chr22 | 800000 | 900000 | 0.00 | 0.00 | NO |
| chr22 | 900000 | 1000000 | 0.00 | 0.00 | NO |
| chr22 | 1000000 | 1100000 | 0.00 | 0.00 | NO |

| | | | | | |
|---|---|---|---|---|---|
| chr22 | 1100000 | 1200000 | 0.00 | 0.00 | NO |
| chr22 | 1200000 | 1300000 | 0.00 | 0.00 | NO |
| chr22 | 1300000 | 1400000 | 0.00 | 0.00 | NO |
| chr22 | 1400000 | 1500000 | 0.00 | 0.00 | NO |
| chr22 | 1500000 | 1600000 | 0.00 | 0.00 | NO |
| chr22 | 1600000 | 1700000 | 0.00 | 0.00 | NO |
| chr22 | 1700000 | 1800000 | 0.00 | 0.00 | NO |
| chr22 | 1800000 | 1900000 | 0.00 | 0.00 | NO |
| chr22 | 1900000 | 2000000 | 0.00 | 0.00 | NO |
| chr22 | 2000000 | 2100000 | 0.00 | 0.00 | NO |
| chr22 | 2100000 | 2200000 | 0.00 | 0.00 | NO |
| chr22 | 2200000 | 2300000 | 0.00 | 0.00 | NO |
| chr22 | 2300000 | 2400000 | 0.00 | 0.00 | NO |
| chr22 | 2400000 | 2500000 | 0.00 | 0.00 | NO |
| chr22 | 2500000 | 2600000 | 0.00 | 0.00 | NO |
| chr22 | 2600000 | 2700000 | 0.00 | 0.00 | NO |
| chr22 | 2700000 | 2800000 | 0.00 | 0.00 | NO |
| chr22 | 2800000 | 2900000 | 0.00 | 0.00 | NO |
| chr22 | 2900000 | 3000000 | 0.00 | 0.00 | NO |
| chr22 | 3000000 | 3100000 | 0.00 | 0.00 | NO |
| chr22 | 3100000 | 3200000 | 0.00 | 0.00 | NO |
| chr22 | 3200000 | 3300000 | 0.00 | 0.00 | NO |
| chr22 | 3300000 | 3400000 | 0.00 | 0.00 | NO |
| chr22 | 3400000 | 3500000 | 0.00 | 0.00 | NO |
| chr22 | 3500000 | 3600000 | 0.00 | 0.00 | NO |
| chr22 | 3600000 | 3700000 | 0.00 | 0.00 | NO |
| chr22 | 3700000 | 3800000 | 0.00 | 0.00 | NO |
| chr22 | 3800000 | 3900000 | 0.00 | 0.00 | NO |
| chr22 | 3900000 | 4000000 | 0.00 | 0.00 | NO |
| chr22 | 4000000 | 4100000 | 0.00 | 0.00 | NO |
| chr22 | 4100000 | 4200000 | 0.00 | 0.00 | NO |
| chr22 | 4200000 | 4300000 | 0.00 | 0.00 | NO |
| chr22 | 4300000 | 4400000 | 0.00 | 0.00 | NO |
| chr22 | 4400000 | 4500000 | 0.00 | 0.00 | NO |
| chr22 | 4500000 | 4600000 | 0.00 | 0.00 | NO |
| chr22 | 4600000 | 4700000 | 0.00 | 0.00 | NO |
| chr22 | 4700000 | 4800000 | 0.00 | 0.00 | NO |
| chr22 | 4800000 | 4900000 | 0.00 | 0.00 | NO |
| chr22 | 4900000 | 5000000 | 0.00 | 0.00 | NO |
| chr22 | 5000000 | 5100000 | 0.00 | 0.00 | NO |
| chr22 | 5100000 | 5200000 | 0.00 | 0.00 | NO |
| chr22 | 5200000 | 5300000 | 0.00 | 0.00 | NO |
| chr22 | 5300000 | 5400000 | 0.00 | 0.00 | NO |
| chr22 | 5400000 | 5500000 | 0.00 | 0.00 | NO |
| chr22 | 5500000 | 5600000 | 0.00 | 0.00 | NO |
| chr22 | 5600000 | 5700000 | 0.00 | 0.00 | NO |
| chr22 | 5700000 | 5800000 | 0.00 | 0.00 | NO |
| chr22 | 5800000 | 5900000 | 0.00 | 0.00 | NO |
| chr22 | 5900000 | 6000000 | 0.00 | 0.00 | NO |
| chr22 | 6000000 | 6100000 | 0.00 | 0.00 | NO |
| chr22 | 6100000 | 6200000 | 0.00 | 0.00 | NO |
| chr22 | 6200000 | 6300000 | 0.00 | 0.00 | NO |
| chr22 | 6300000 | 6400000 | 0.00 | 0.00 | NO |
| chr22 | 6400000 | 6500000 | 0.00 | 0.00 | NO |
| chr22 | 6500000 | 6600000 | 0.00 | 0.00 | NO |
| chr22 | 6600000 | 6700000 | 0.00 | 0.00 | NO |

| | | | | | |
|---|---|---|---|---|---|
| chr22 | 6700000 | 6800000 | 0.00 | 0.00 | NO |
| chr22 | 6800000 | 6900000 | 0.00 | 0.00 | NO |
| chr22 | 6900000 | 7000000 | 0.00 | 0.00 | NO |
| chr22 | 7000000 | 7100000 | 0.00 | 0.00 | NO |
| chr22 | 7100000 | 7200000 | 0.00 | 0.00 | NO |
| chr22 | 7200000 | 7300000 | 0.00 | 0.00 | NO |
| chr22 | 7300000 | 7400000 | 0.00 | 0.00 | NO |
| chr22 | 7400000 | 7500000 | 0.00 | 0.00 | NO |
| chr22 | 7500000 | 7600000 | 0.00 | 0.00 | NO |
| chr22 | 7600000 | 7700000 | 0.00 | 0.00 | NO |
| chr22 | 7700000 | 7800000 | 0.00 | 0.00 | NO |
| chr22 | 7800000 | 7900000 | 0.00 | 0.00 | NO |
| chr22 | 7900000 | 8000000 | 0.00 | 0.00 | NO |
| chr22 | 8000000 | 8100000 | 0.00 | 0.00 | NO |
| chr22 | 8100000 | 8200000 | 0.00 | 0.00 | NO |
| chr22 | 8200000 | 8300000 | 0.00 | 0.00 | NO |
| chr22 | 8300000 | 8400000 | 0.00 | 0.00 | NO |
| chr22 | 8400000 | 8500000 | 0.00 | 0.00 | NO |
| chr22 | 8500000 | 8600000 | 0.00 | 0.00 | NO |
| chr22 | 8600000 | 8700000 | 0.00 | 0.00 | NO |
| chr22 | 8700000 | 8800000 | 0.00 | 0.00 | NO |
| chr22 | 8800000 | 8900000 | 0.00 | 0.00 | NO |
| chr22 | 8900000 | 9000000 | 0.00 | 0.00 | NO |
| chr22 | 9000000 | 9100000 | 0.00 | 0.00 | NO |
| chr22 | 9100000 | 9200000 | 0.00 | 0.00 | NO |
| chr22 | 9200000 | 9300000 | 0.00 | 0.00 | NO |
| chr22 | 9300000 | 9400000 | 0.00 | 0.00 | NO |
| chr22 | 9400000 | 9500000 | 0.00 | 0.00 | NO |
| chr22 | 9500000 | 9600000 | 0.00 | 0.00 | NO |
| chr22 | 9600000 | 9700000 | 0.00 | 0.00 | NO |
| chr22 | 9700000 | 9800000 | 0.00 | 0.00 | NO |
| chr22 | 9800000 | 9900000 | 0.00 | 0.00 | NO |
| chr22 | 9900000 | 10000000 | 0.00 | 0.00 | NO |
| chr22 | 10000000 | 10100000 | 0.00 | 0.00 | NO |
| chr22 | 10100000 | 10200000 | 0.00 | 0.00 | NO |
| chr22 | 10200000 | 10300000 | 0.00 | 0.00 | NO |
| chr22 | 10300000 | 10400000 | 0.00 | 0.00 | NO |
| chr22 | 10400000 | 10500000 | 0.00 | 0.00 | NO |
| chr22 | 10500000 | 10600000 | 0.00 | 0.00 | NO |
| chr22 | 10600000 | 10700000 | 0.00 | 0.00 | NO |
| chr22 | 10700000 | 10800000 | 0.00 | 0.00 | NO |
| chr22 | 10800000 | 10900000 | 0.00 | 0.00 | NO |
| chr22 | 10900000 | 11000000 | 0.00 | 0.00 | NO |
| chr22 | 11000000 | 11100000 | 0.00 | 0.00 | NO |
| chr22 | 11100000 | 11200000 | 0.00 | 0.00 | NO |
| chr22 | 11200000 | 11300000 | 0.00 | 0.00 | NO |
| chr22 | 11300000 | 11400000 | 0.00 | 0.00 | NO |
| chr22 | 11400000 | 11500000 | 0.00 | 0.00 | NO |
| chr22 | 11500000 | 11600000 | 0.00 | 0.00 | NO |
| chr22 | 11600000 | 11700000 | 0.00 | 0.00 | NO |
| chr22 | 11700000 | 11800000 | 0.00 | 0.00 | NO |
| chr22 | 11800000 | 11900000 | 0.00 | 0.00 | NO |
| chr22 | 11900000 | 12000000 | 0.00 | 0.00 | NO |
| chr22 | 12000000 | 12100000 | 0.00 | 0.00 | NO |
| chr22 | 12100000 | 12200000 | 0.00 | 0.00 | NO |
| chr22 | 12200000 | 12300000 | 0.00 | 0.00 | NO |

| | | | | | |
|---|---|---|---|---|---|
| chr22 | 12300000 | 12400000 | 0.00 | 0.00 | NO |
| chr22 | 12400000 | 12500000 | 0.00 | 0.00 | NO |
| chr22 | 12500000 | 12600000 | 0.00 | 0.00 | NO |
| chr22 | 12600000 | 12700000 | 0.00 | 0.00 | NO |
| chr22 | 12700000 | 12800000 | 0.00 | 0.00 | NO |
| chr22 | 12800000 | 12900000 | 0.00 | 0.00 | NO |
| chr22 | 12900000 | 13000000 | 0.00 | 0.00 | NO |
| chr22 | 13000000 | 13100000 | 0.00 | 0.00 | NO |
| chr22 | 13100000 | 13200000 | 0.00 | 0.00 | NO |
| chr22 | 13200000 | 13300000 | 0.00 | 0.00 | NO |
| chr22 | 13300000 | 13400000 | 0.00 | 0.00 | NO |
| chr22 | 13400000 | 13500000 | 0.00 | 0.00 | NO |
| chr22 | 13500000 | 13600000 | 0.00 | 0.00 | NO |
| chr22 | 13600000 | 13700000 | 0.00 | 0.00 | NO |
| chr22 | 13700000 | 13800000 | 0.00 | 0.00 | NO |
| chr22 | 13800000 | 13900000 | 0.00 | 0.00 | NO |
| chr22 | 13900000 | 14000000 | 0.00 | 0.00 | NO |
| chr22 | 14000000 | 14100000 | 0.00 | 0.00 | NO |
| chr22 | 14100000 | 14200000 | 0.00 | 0.00 | NO |
| chr22 | 14200000 | 14300000 | 0.00 | 0.00 | NO |
| chr22 | 14300000 | 14400000 | 0.00 | 0.00 | NO |
| chr22 | 14400000 | 14500000 | 0.00 | 0.00 | NO |
| chr22 | 14500000 | 14600000 | 0.00 | 0.00 | NO |
| chr22 | 14600000 | 14700000 | 0.00 | 0.00 | NO |
| chr22 | 14700000 | 14800000 | 0.00 | 0.00 | NO |
| chr22 | 14800000 | 14900000 | 0.00 | 0.00 | NO |
| chr22 | 14900000 | 15000000 | 0.00 | 0.00 | NO |
| chr22 | 15000000 | 15100000 | 0.00 | 0.00 | NO |
| chr22 | 15100000 | 15200000 | 0.00 | 0.00 | NO |
| chr22 | 15200000 | 15300000 | 0.00 | 0.00 | NO |
| chr22 | 15300000 | 15400000 | 0.00 | 0.00 | NO |
| chr22 | 15400000 | 15500000 | 0.00 | 0.00 | NO |
| chr22 | 15500000 | 15600000 | 0.00 | 0.00 | NO |
| chr22 | 15600000 | 15700000 | 0.00 | 0.00 | NO |
| chr22 | 15700000 | 15800000 | 0.00 | 0.00 | NO |
| chr22 | 15800000 | 15900000 | 0.00 | 0.00 | NO |
| chr22 | 15900000 | 16000000 | 0.00 | 0.00 | NO |
| chr22 | 16000000 | 16100000 | 0.00 | 0.00 | NO |
| chr22 | 16100000 | 16200000 | 0.00 | 0.00 | NO |
| chr22 | 16200000 | 16300000 | 0.00 | 0.00 | NO |
| chr22 | 16300000 | 16400000 | 0.00 | 0.00 | NO |
| chr22 | 16400000 | 16500000 | 0.00 | 0.00 | NO |
| chr22 | 16500000 | 16600000 | 0.00 | 0.00 | NO |
| chr22 | 16600000 | 16700000 | 0.00 | 0.00 | NO |
| chr22 | 16700000 | 16800000 | 0.00 | 0.00 | NO |
| chr22 | 16800000 | 16900000 | 0.00 | 0.00 | NO |
| chr22 | 16900000 | 17000000 | 0.00 | 0.00 | NO |
| chr22 | 17000000 | 17100000 | 0.00 | 0.00 | NO |
| chr22 | 17100000 | 17200000 | 0.00 | 0.00 | NO |
| chr22 | 17200000 | 17300000 | 0.00 | 0.00 | NO |
| chr22 | 17300000 | 17400000 | 0.00 | 0.00 | NO |
| chr22 | 17400000 | 17500000 | -0.96 | -1.71 | A>B |
| chr22 | 17500000 | 17600000 | -0.61 | -0.96 | B>B |
| chr22 | 17600000 | 17700000 | -0.39 | -0.86 | B>B |
| chr22 | 17700000 | 17800000 | -0.53 | -1.17 | A>B |
| chr22 | 17800000 | 17900000 | -0.46 | -1.17 | A>B |

| | | | | | |
|---|---|---|---|---|---|
| chr22 | 17900000 | 18000000 | -0.53 | -1.14 | A>B |
| chr22 | 18000000 | 18100000 | -0.16 | -0.40 | B>B |
| chr22 | 18100000 | 18200000 | 0.25 | 0.11 | A>A |
| chr22 | 18200000 | 18300000 | 0.14 | 0.32 | A>A |
| chr22 | 18300000 | 18400000 | 0.31 | 0.37 | A>A |
| chr22 | 18400000 | 18500000 | 0.46 | 0.49 | A>A |
| chr22 | 18500000 | 18600000 | 0.31 | 0.45 | A>A |
| chr22 | 18600000 | 18700000 | 0.00 | 0.00 | NO |
| chr22 | 18700000 | 18800000 | 0.00 | 0.00 | NO |
| chr22 | 18800000 | 18900000 | 0.00 | 0.00 | NO |
| chr22 | 18900000 | 19000000 | 0.00 | 0.81 | NO |
| chr22 | 19000000 | 19100000 | 0.77 | 0.86 | A>A |
| chr22 | 19100000 | 19200000 | 0.72 | 0.86 | A>A |
| chr22 | 19200000 | 19300000 | 0.60 | 0.71 | A>A |
| chr22 | 19300000 | 19400000 | 0.48 | 0.64 | A>A |
| chr22 | 19400000 | 19500000 | 0.74 | 0.69 | A>A |
| chr22 | 19500000 | 19600000 | 0.64 | 0.61 | A>A |
| chr22 | 19600000 | 19700000 | 0.54 | 0.63 | A>A |
| chr22 | 19700000 | 19800000 | 0.82 | 0.87 | A>A |
| chr22 | 19800000 | 19900000 | 0.95 | 0.86 | A>A |
| chr22 | 19900000 | 20000000 | 0.82 | 0.81 | A>A |
| chr22 | 20000000 | 20100000 | 1.00 | 1.01 | A>A |
| chr22 | 20100000 | 20200000 | 1.23 | 1.07 | A>A |
| chr22 | 20200000 | 20300000 | 1.03 | 1.01 | A>A |
| chr22 | 20300000 | 20400000 | 0.00 | 0.00 | NO |
| chr22 | 20400000 | 20500000 | 0.00 | 0.00 | NO |
| chr22 | 20500000 | 20600000 | 0.00 | 0.00 | NO |
| chr22 | 20600000 | 20700000 | 0.00 | 0.00 | NO |
| chr22 | 20700000 | 20800000 | 0.61 | 0.66 | A>A |
| chr22 | 20800000 | 20900000 | 0.93 | 0.91 | A>A |
| chr22 | 20900000 | 21000000 | 0.89 | 0.66 | A>A |
| chr22 | 21000000 | 21100000 | 0.00 | 0.00 | NO |
| chr22 | 21100000 | 21200000 | 0.91 | 0.78 | A>A |
| chr22 | 21200000 | 21300000 | 1.06 | 1.00 | A>A |
| chr22 | 21300000 | 21400000 | 1.23 | 1.09 | A>A |
| chr22 | 21400000 | 21500000 | 0.00 | 0.00 | NO |
| chr22 | 21500000 | 21600000 | 0.00 | 0.00 | NO |
| chr22 | 21600000 | 21700000 | 0.00 | 0.00 | NO |
| chr22 | 21700000 | 21800000 | 0.00 | 0.00 | NO |
| chr22 | 21800000 | 21900000 | 0.00 | 0.00 | NO |
| chr22 | 21900000 | 22000000 | 1.07 | 1.03 | A>A |
| chr22 | 22000000 | 22100000 | 0.92 | 0.90 | A>A |
| chr22 | 22100000 | 22200000 | 0.75 | 0.74 | A>A |
| chr22 | 22200000 | 22300000 | 0.63 | 0.52 | A>A |
| chr22 | 22300000 | 22400000 | 0.36 | 0.13 | A>A |
| chr22 | 22400000 | 22500000 | 0.12 | -0.10 | A>A |
| chr22 | 22500000 | 22600000 | -0.08 | -0.09 | B>B |
| chr22 | 22600000 | 22700000 | 0.00 | 0.00 | NO |
| chr22 | 22700000 | 22800000 | -0.07 | -0.07 | B>B |
| chr22 | 22800000 | 22900000 | 0.21 | 0.09 | A>A |
| chr22 | 22900000 | 23000000 | 0.10 | 0.04 | A>A |
| chr22 | 23000000 | 23100000 | 0.08 | 0.17 | A>A |
| chr22 | 23100000 | 23200000 | -0.07 | -0.19 | B>B |
| chr22 | 23200000 | 23300000 | -0.17 | -0.20 | B>B |
| chr22 | 23300000 | 23400000 | -0.10 | -0.22 | B>B |
| chr22 | 23400000 | 23500000 | -0.03 | 0.05 | A>A |

| | | | | | |
|---|---|---|---|---|---|
| chr22 | 23500000 | 23600000 | 0.32 | 0.40 | A>A |
| chr22 | 23600000 | 23700000 | 0.21 | 0.32 | A>A |
| chr22 | 23700000 | 23800000 | 0.33 | 0.37 | A>A |
| chr22 | 23800000 | 23900000 | 0.58 | 0.58 | A>A |
| chr22 | 23900000 | 24000000 | 0.28 | 0.44 | A>A |
| chr22 | 24000000 | 24100000 | 0.67 | 0.67 | A>A |
| chr22 | 24100000 | 24200000 | 0.79 | 0.72 | A>A |
| chr22 | 24200000 | 24300000 | 0.73 | 0.72 | A>A |
| chr22 | 24300000 | 24400000 | 0.00 | 0.00 | NO |
| chr22 | 24400000 | 24500000 | 0.65 | 0.71 | A>A |
| chr22 | 24500000 | 24600000 | 0.79 | 0.77 | A>A |
| chr22 | 24600000 | 24700000 | 0.00 | 0.00 | NO |
| chr22 | 24700000 | 24800000 | 0.55 | 0.54 | A>A |
| chr22 | 24800000 | 24900000 | 0.55 | 0.69 | A>A |
| chr22 | 24900000 | 25000000 | 0.71 | 0.68 | A>A |
| chr22 | 25000000 | 25100000 | 0.00 | 0.00 | NO |
| chr22 | 25100000 | 25200000 | 0.26 | 0.23 | A>A |
| chr22 | 25200000 | 25300000 | 0.47 | 0.41 | A>A |
| chr22 | 25300000 | 25400000 | 0.40 | 0.34 | A>A |
| chr22 | 25400000 | 25500000 | 0.00 | 0.12 | NO |
| chr22 | 25500000 | 25600000 | -0.01 | 0.03 | A>A |
| chr22 | 25600000 | 25700000 | 0.00 | -0.48 | NO |
| chr22 | 25700000 | 25800000 | -0.15 | -0.35 | B>B |
| chr22 | 25800000 | 25900000 | -0.24 | -0.47 | B>B |
| chr22 | 25900000 | 26000000 | -0.35 | -0.53 | B>B |
| chr22 | 26000000 | 26100000 | -0.33 | -0.60 | B>B |
| chr22 | 26100000 | 26200000 | -0.51 | -0.74 | B>B |
| chr22 | 26200000 | 26300000 | -0.83 | -1.22 | B>B |
| chr22 | 26300000 | 26400000 | -0.86 | -1.33 | B>B |
| chr22 | 26400000 | 26500000 | -0.83 | -1.37 | B>B |
| chr22 | 26500000 | 26600000 | -0.79 | -1.35 | B>B |
| chr22 | 26600000 | 26700000 | -0.84 | -1.35 | B>B |
| chr22 | 26700000 | 26800000 | -0.59 | -0.89 | B>B |
| chr22 | 26800000 | 26900000 | 0.04 | -0.32 | B>B |
| chr22 | 26900000 | 27000000 | 0.16 | -0.14 | A>A |
| chr22 | 27000000 | 27100000 | 0.11 | -0.02 | A>A |
| chr22 | 27100000 | 27200000 | 0.19 | -0.02 | A>A |
| chr22 | 27200000 | 27300000 | -0.06 | -0.05 | B>B |
| chr22 | 27300000 | 27400000 | -0.11 | -0.24 | B>B |
| chr22 | 27400000 | 27500000 | 0.01 | -0.20 | B>B |
| chr22 | 27500000 | 27600000 | 0.05 | -0.08 | B>B |
| chr22 | 27600000 | 27700000 | 0.06 | -0.05 | A>A |
| chr22 | 27700000 | 27800000 | -0.11 | -0.20 | B>B |
| chr22 | 27800000 | 27900000 | -0.04 | -0.04 | B>B |
| chr22 | 27900000 | 28000000 | -0.08 | -0.07 | B>B |
| chr22 | 28000000 | 28100000 | 0.03 | -0.03 | B>B |
| chr22 | 28100000 | 28200000 | 0.01 | -0.06 | B>B |
| chr22 | 28200000 | 28300000 | -0.02 | 0.03 | A>A |
| chr22 | 28300000 | 28400000 | 0.18 | 0.18 | A>A |
| chr22 | 28400000 | 28500000 | -0.20 | 0.02 | B>B |
| chr22 | 28500000 | 28600000 | -0.26 | -0.02 | B>B |
| chr22 | 28600000 | 28700000 | -0.18 | -0.04 | B>B |
| chr22 | 28700000 | 28800000 | -0.17 | -0.16 | B>B |
| chr22 | 28800000 | 28900000 | -0.13 | -0.01 | B>B |
| chr22 | 28900000 | 29000000 | -0.15 | -0.05 | B>B |
| chr22 | 29000000 | 29100000 | 0.28 | 0.18 | A>A |

| | | | | | |
|---|---|---|---|---|---|
| chr22 | 29100000 | 29200000 | 0.34 | 0.38 | A>A |
| chr22 | 29200000 | 29300000 | 0.11 | 0.29 | A>A |
| chr22 | 29300000 | 29400000 | 0.31 | 0.41 | A>A |
| chr22 | 29400000 | 29500000 | 0.11 | 0.33 | A>A |
| chr22 | 29500000 | 29600000 | 0.38 | 0.35 | A>A |
| chr22 | 29600000 | 29700000 | 0.69 | 0.63 | A>A |
| chr22 | 29700000 | 29800000 | 0.85 | 0.88 | A>A |
| chr22 | 29800000 | 29900000 | 0.82 | 0.74 | A>A |
| chr22 | 29900000 | 30000000 | 0.80 | 0.74 | A>A |
| chr22 | 30000000 | 30100000 | 0.56 | 0.55 | A>A |
| chr22 | 30100000 | 30200000 | 0.52 | 0.57 | A>A |
| chr22 | 30200000 | 30300000 | 0.56 | 0.59 | A>A |
| chr22 | 30300000 | 30400000 | 0.61 | 0.56 | A>A |
| chr22 | 30400000 | 30500000 | 0.25 | 0.39 | A>A |
| chr22 | 30500000 | 30600000 | 0.80 | 0.75 | A>A |
| chr22 | 30600000 | 30700000 | 1.08 | 0.86 | A>A |
| chr22 | 30700000 | 30800000 | 0.96 | 0.84 | A>A |
| chr22 | 30800000 | 30900000 | 0.81 | 0.80 | A>A |
| chr22 | 30900000 | 31000000 | 0.91 | 0.90 | A>A |
| chr22 | 31000000 | 31100000 | 0.76 | 0.78 | A>A |
| chr22 | 31100000 | 31200000 | 0.60 | 0.59 | A>A |
| chr22 | 31200000 | 31300000 | 0.83 | 0.74 | A>A |
| chr22 | 31300000 | 31400000 | 0.55 | 0.60 | A>A |
| chr22 | 31400000 | 31500000 | 0.62 | 0.64 | A>A |
| chr22 | 31500000 | 31600000 | 0.74 | 0.75 | A>A |
| chr22 | 31600000 | 31700000 | 0.83 | 0.73 | A>A |
| chr22 | 31700000 | 31800000 | 0.85 | 0.76 | A>A |
| chr22 | 31800000 | 31900000 | 0.84 | 0.77 | A>A |
| chr22 | 31900000 | 32000000 | 0.67 | 0.56 | A>A |
| chr22 | 32000000 | 32100000 | 0.50 | 0.41 | A>A |
| chr22 | 32100000 | 32200000 | 0.37 | 0.21 | A>A |
| chr22 | 32200000 | 32300000 | 0.27 | 0.11 | A>A |
| chr22 | 32300000 | 32400000 | 0.17 | -0.03 | A>A |
| chr22 | 32400000 | 32500000 | -0.09 | -0.37 | B>B |
| chr22 | 32500000 | 32600000 | -0.12 | -0.49 | B>B |
| chr22 | 32600000 | 32700000 | -0.15 | -0.27 | B>B |
| chr22 | 32700000 | 32800000 | -0.21 | -0.61 | B>B |
| chr22 | 32800000 | 32900000 | -0.16 | -0.60 | B>B |
| chr22 | 32900000 | 33000000 | -0.52 | -1.11 | A>B |
| chr22 | 33000000 | 33100000 | 0.00 | -1.16 | NO |
| chr22 | 33100000 | 33200000 | -0.51 | -1.14 | A>B |
| chr22 | 33200000 | 33300000 | -0.74 | -1.33 | A>B |
| chr22 | 33300000 | 33400000 | -0.86 | -1.29 | B>B |
| chr22 | 33400000 | 33500000 | -0.76 | -1.23 | B>B |
| chr22 | 33500000 | 33600000 | -0.84 | -1.24 | B>B |
| chr22 | 33600000 | 33700000 | -0.89 | -1.04 | B>B |
| chr22 | 33700000 | 33800000 | -0.37 | -0.54 | B>B |
| chr22 | 33800000 | 33900000 | -0.26 | -0.55 | B>B |
| chr22 | 33900000 | 34000000 | -0.39 | -0.51 | B>B |
| chr22 | 34000000 | 34100000 | -0.58 | -0.67 | B>B |
| chr22 | 34100000 | 34200000 | -0.54 | -0.70 | B>B |
| chr22 | 34200000 | 34300000 | -0.61 | -0.70 | B>B |
| chr22 | 34300000 | 34400000 | -0.84 | -1.10 | B>B |
| chr22 | 34400000 | 34500000 | -0.89 | -1.17 | B>B |
| chr22 | 34500000 | 34600000 | -0.93 | -1.07 | B>B |
| chr22 | 34600000 | 34700000 | -0.90 | -1.10 | B>B |

| | | | | | |
|---|---|---|---|---|---|
| chr22 | 34700000 | 34800000 | -0.93 | -0.97 | B>B |
| chr22 | 34800000 | 34900000 | -0.94 | -1.00 | B>B |
| chr22 | 34900000 | 35000000 | -0.84 | -0.97 | B>B |
| chr22 | 35000000 | 35100000 | -0.88 | -1.03 | B>B |
| chr22 | 35100000 | 35200000 | -0.85 | -1.11 | B>B |
| chr22 | 35200000 | 35300000 | -0.97 | -1.24 | B>B |
| chr22 | 35300000 | 35400000 | -1.00 | -1.43 | B>B |
| chr22 | 35400000 | 35500000 | -1.05 | -1.49 | B>B |
| chr22 | 35500000 | 35600000 | -0.89 | -1.30 | B>B |
| chr22 | 35600000 | 35700000 | -0.38 | -0.72 | B>B |
| chr22 | 35700000 | 35800000 | 0.23 | 0.26 | A>A |
| chr22 | 35800000 | 35900000 | 0.37 | 0.39 | A>A |
| chr22 | 35900000 | 36000000 | 0.31 | 0.21 | A>A |
| chr22 | 36000000 | 36100000 | 0.15 | 0.19 | A>A |
| chr22 | 36100000 | 36200000 | 0.10 | 0.15 | A>A |
| chr22 | 36200000 | 36300000 | 0.23 | 0.25 | A>A |
| chr22 | 36300000 | 36400000 | 0.19 | 0.22 | A>A |
| chr22 | 36400000 | 36500000 | 0.36 | 0.36 | A>A |
| chr22 | 36500000 | 36600000 | 0.34 | 0.33 | A>A |
| chr22 | 36600000 | 36700000 | 0.52 | 0.68 | A>A |
| chr22 | 36700000 | 36800000 | 0.78 | 0.88 | A>A |
| chr22 | 36800000 | 36900000 | 0.82 | 0.77 | A>A |
| chr22 | 36900000 | 37000000 | 0.54 | 0.39 | A>A |
| chr22 | 37000000 | 37100000 | 0.53 | 0.48 | A>A |
| chr22 | 37100000 | 37200000 | 0.40 | 0.33 | A>A |
| chr22 | 37200000 | 37300000 | 0.52 | 0.38 | A>A |
| chr22 | 37300000 | 37400000 | 0.52 | 0.28 | A>A |
| chr22 | 37400000 | 37500000 | 0.68 | 0.67 | A>A |
| chr22 | 37500000 | 37600000 | 0.70 | 0.66 | A>A |
| chr22 | 37600000 | 37700000 | 0.48 | 0.62 | A>A |
| chr22 | 37700000 | 37800000 | 0.82 | 0.80 | A>A |
| chr22 | 37800000 | 37900000 | 0.72 | 0.66 | A>A |
| chr22 | 37900000 | 38000000 | 0.72 | 0.80 | A>A |
| chr22 | 38000000 | 38100000 | 0.83 | 0.85 | A>A |
| chr22 | 38100000 | 38200000 | 0.87 | 0.89 | A>A |
| chr22 | 38200000 | 38300000 | 0.76 | 0.84 | A>A |
| chr22 | 38300000 | 38400000 | 0.72 | 0.71 | A>A |
| chr22 | 38400000 | 38500000 | 0.55 | 0.56 | A>A |
| chr22 | 38500000 | 38600000 | 0.57 | 0.60 | A>A |
| chr22 | 38600000 | 38700000 | 0.47 | 0.55 | A>A |
| chr22 | 38700000 | 38800000 | 0.56 | 0.46 | A>A |
| chr22 | 38800000 | 38900000 | 0.60 | 0.61 | A>A |
| chr22 | 38900000 | 39000000 | 0.62 | 0.61 | A>A |
| chr22 | 39000000 | 39100000 | 0.58 | 0.60 | A>A |
| chr22 | 39100000 | 39200000 | 0.68 | 0.62 | A>A |
| chr22 | 39200000 | 39300000 | 0.61 | 0.56 | A>A |
| chr22 | 39300000 | 39400000 | 0.59 | 0.54 | A>A |
| chr22 | 39400000 | 39500000 | 0.40 | 0.45 | A>A |
| chr22 | 39500000 | 39600000 | 0.45 | 0.42 | A>A |
| chr22 | 39600000 | 39700000 | 0.77 | 0.73 | A>A |
| chr22 | 39700000 | 39800000 | 0.52 | 0.54 | A>A |
| chr22 | 39800000 | 39900000 | 0.56 | 0.54 | A>A |
| chr22 | 39900000 | 40000000 | 0.08 | 0.11 | A>A |
| chr22 | 40000000 | 40100000 | 0.12 | 0.10 | A>A |
| chr22 | 40100000 | 40200000 | 0.11 | 0.03 | A>A |
| chr22 | 40200000 | 40300000 | -0.11 | -0.06 | B>B |

| | | | | | |
|---|---|---|---|---|---|
| chr22 | 40300000 | 40400000 | -0.01 | 0.07 | A>A |
| chr22 | 40400000 | 40500000 | -0.01 | 0.13 | A>A |
| chr22 | 40500000 | 40600000 | 0.26 | 0.20 | A>A |
| chr22 | 40600000 | 40700000 | 0.15 | 0.24 | A>A |
| chr22 | 40700000 | 40800000 | 0.28 | 0.29 | A>A |
| chr22 | 40800000 | 40900000 | 0.29 | 0.37 | A>A |
| chr22 | 40900000 | 41000000 | 0.27 | 0.42 | A>A |
| chr22 | 41000000 | 41100000 | 0.30 | 0.37 | A>A |
| chr22 | 41100000 | 41200000 | 0.30 | 0.45 | A>A |
| chr22 | 41200000 | 41300000 | 0.49 | 0.49 | A>A |
| chr22 | 41300000 | 41400000 | 0.45 | 0.40 | A>A |
| chr22 | 41400000 | 41500000 | 0.48 | 0.45 | A>A |
| chr22 | 41500000 | 41600000 | 0.47 | 0.50 | A>A |
| chr22 | 41600000 | 41700000 | 0.63 | 0.58 | A>A |
| chr22 | 41700000 | 41800000 | 0.58 | 0.62 | A>A |
| chr22 | 41800000 | 41900000 | 0.61 | 0.63 | A>A |
| chr22 | 41900000 | 42000000 | 0.57 | 0.65 | A>A |
| chr22 | 42000000 | 42100000 | 0.49 | 0.62 | A>A |
| chr22 | 42100000 | 42200000 | 0.54 | 0.47 | A>A |
| chr22 | 42200000 | 42300000 | 0.54 | 0.58 | A>A |
| chr22 | 42300000 | 42400000 | 0.41 | 0.54 | A>A |
| chr22 | 42400000 | 42500000 | 0.48 | 0.56 | A>A |
| chr22 | 42500000 | 42600000 | 0.29 | 0.33 | A>A |
| chr22 | 42600000 | 42700000 | 0.45 | 0.50 | A>A |
| chr22 | 42700000 | 42800000 | 0.50 | 0.56 | A>A |
| chr22 | 42800000 | 42900000 | 0.35 | 0.47 | A>A |
| chr22 | 42900000 | 43000000 | 0.76 | 0.81 | A>A |
| chr22 | 43000000 | 43100000 | 0.51 | 0.55 | A>A |
| chr22 | 43100000 | 43200000 | 0.51 | 0.62 | A>A |
| chr22 | 43200000 | 43300000 | 0.19 | 0.49 | A>A |
| chr22 | 43300000 | 43400000 | 0.40 | 0.52 | A>A |
| chr22 | 43400000 | 43500000 | 0.30 | 0.52 | A>A |
| chr22 | 43500000 | 43600000 | 0.54 | 0.54 | A>A |
| chr22 | 43600000 | 43700000 | 0.13 | 0.03 | A>A |
| chr22 | 43700000 | 43800000 | -0.02 | -0.01 | B>B |
| chr22 | 43800000 | 43900000 | -0.08 | -0.16 | B>B |
| chr22 | 43900000 | 44000000 | -0.17 | -0.23 | B>B |
| chr22 | 44000000 | 44100000 | -0.05 | -0.07 | B>B |
| chr22 | 44100000 | 44200000 | -0.17 | 0.01 | B>B |
| chr22 | 44200000 | 44300000 | -0.07 | 0.01 | B>B |
| chr22 | 44300000 | 44400000 | 0.17 | 0.26 | A>A |
| chr22 | 44400000 | 44500000 | 0.17 | 0.24 | A>A |
| chr22 | 44500000 | 44600000 | 0.06 | 0.02 | A>A |
| chr22 | 44600000 | 44700000 | 0.09 | -0.07 | A>A |
| chr22 | 44700000 | 44800000 | 0.13 | 0.14 | A>A |
| chr22 | 44800000 | 44900000 | 0.27 | 0.18 | A>A |
| chr22 | 44900000 | 45000000 | 0.19 | 0.14 | A>A |
| chr22 | 45000000 | 45100000 | 0.09 | 0.16 | A>A |
| chr22 | 45100000 | 45200000 | 0.32 | 0.26 | A>A |
| chr22 | 45200000 | 45300000 | 0.08 | 0.19 | A>A |
| chr22 | 45300000 | 45400000 | 0.17 | 0.07 | A>A |
| chr22 | 45400000 | 45500000 | 0.24 | 0.13 | A>A |
| chr22 | 45500000 | 45600000 | 0.35 | 0.31 | A>A |
| chr22 | 45600000 | 45700000 | 0.32 | 0.35 | A>A |
| chr22 | 45700000 | 45800000 | 0.28 | 0.23 | A>A |
| chr22 | 45800000 | 45900000 | 0.26 | 0.27 | A>A |

| | | | | | |
|---|---|---|---|---|---|
| chr22 | 45900000 | 46000000 | 0.10 | 0.16 | A>A |
| chr22 | 46000000 | 46100000 | 0.17 | 0.13 | A>A |
| chr22 | 46100000 | 46200000 | 0.19 | 0.13 | A>A |
| chr22 | 46200000 | 46300000 | -0.02 | 0.07 | A>A |
| chr22 | 46300000 | 46400000 | -0.01 | 0.15 | A>A |
| chr22 | 46400000 | 46500000 | 0.32 | 0.33 | A>A |
| chr22 | 46500000 | 46600000 | 0.25 | 0.36 | A>A |
| chr22 | 46600000 | 46700000 | 0.51 | 0.51 | A>A |
| chr22 | 46700000 | 46800000 | 0.57 | 0.54 | A>A |
| chr22 | 46800000 | 46900000 | 0.69 | 0.68 | A>A |
| chr22 | 46900000 | 47000000 | 0.49 | 0.54 | A>A |
| chr22 | 47000000 | 47100000 | 0.70 | 0.66 | A>A |
| chr22 | 47100000 | 47200000 | 0.43 | 0.50 | A>A |
| chr22 | 47200000 | 47300000 | -0.01 | 0.19 | A>A |
| chr22 | 47300000 | 47400000 | -0.14 | 0.08 | B>B |
| chr22 | 47400000 | 47500000 | -0.16 | -0.07 | B>B |
| chr22 | 47500000 | 47600000 | -0.44 | -0.14 | B>B |
| chr22 | 47600000 | 47700000 | -0.43 | -0.15 | B>B |
| chr22 | 47700000 | 47800000 | -0.32 | -0.07 | B>B |
| chr22 | 47800000 | 47900000 | -0.53 | -0.24 | B>B |
| chr22 | 47900000 | 48000000 | -0.55 | -0.26 | B>B |
| chr22 | 48000000 | 48100000 | -0.52 | -0.29 | B>B |
| chr22 | 48100000 | 48200000 | -0.49 | -0.28 | B>B |
| chr22 | 48200000 | 48300000 | -0.32 | -0.21 | B>B |
| chr22 | 48300000 | 48400000 | -0.25 | -0.14 | B>B |
| chr22 | 48400000 | 48500000 | -0.16 | -0.13 | B>B |
| chr22 | 48500000 | 48600000 | -0.03 | 0.07 | A>A |
| chr22 | 48600000 | 48700000 | 0.05 | 0.08 | A>A |
| chr22 | 48700000 | 48800000 | 0.08 | 0.24 | A>A |
| chr22 | 48800000 | 48900000 | 0.00 | 0.10 | A>A |
| chr22 | 48900000 | 49000000 | -0.52 | -0.22 | B>B |
| chr22 | 49000000 | 49100000 | -0.50 | -0.23 | B>B |
| chr22 | 49100000 | 49200000 | -0.51 | -0.33 | B>B |
| chr22 | 49200000 | 49300000 | -0.57 | -0.39 | B>B |
| chr22 | 49300000 | 49400000 | -0.64 | -0.43 | B>B |
| chr22 | 49400000 | 49500000 | -0.52 | -0.31 | B>B |
| chr22 | 49500000 | 49600000 | -0.41 | -0.39 | B>B |
| chr22 | 49600000 | 49700000 | -0.46 | -0.39 | B>B |
| chr22 | 49700000 | 49800000 | -0.54 | -0.17 | B>B |
| chr22 | 49800000 | 49900000 | -0.34 | -0.01 | B>B |
| chr22 | 49900000 | 50000000 | -0.31 | -0.23 | B>B |
| chr22 | 50000000 | 50100000 | -0.48 | -0.05 | B>B |
| chr22 | 50100000 | 50200000 | -0.04 | 0.25 | A>A |
| chr22 | 50200000 | 50300000 | 0.73 | 0.65 | A>A |
| chr22 | 50300000 | 50400000 | 1.09 | 0.92 | A>A |
| chr22 | 50400000 | 50500000 | 0.77 | 0.75 | A>A |
| chr22 | 50500000 | 50600000 | 0.76 | 0.71 | A>A |
| chr22 | 50600000 | 50700000 | 0.81 | 0.81 | A>A |
| chr22 | 50700000 | 50800000 | 1.02 | 0.98 | A>A |
| chr22 | 50800000 | 50900000 | 0.94 | 0.84 | A>A |
| chr22 | 50900000 | 51000000 | 1.04 | 0.84 | A>A |
| chr22 | 51000000 | 51100000 | 1.01 | 0.87 | A>A |
| chr22 | 51100000 | 51200000 | 1.02 | 0.65 | A>A |
| chr22 | 51200000 | 51300000 | 0.00 | 0.00 | NO |
| chr22 | 51300000 | 51304566 | 0.00 | 0.00 | NO |
| chrX | 0 | 100000 | 0.00 | 0.00 | NO |

| | | | | | |
|---|---|---|---|---|---|
| chrX | 100000 | 200000 | 0.00 | 0.00 | NO |
| chrX | 200000 | 300000 | 0.00 | 0.00 | NO |
| chrX | 300000 | 400000 | 0.00 | 0.00 | NO |
| chrX | 400000 | 500000 | 0.00 | 0.00 | NO |
| chrX | 500000 | 600000 | 0.00 | 0.00 | NO |
| chrX | 600000 | 700000 | 0.00 | 0.00 | NO |
| chrX | 700000 | 800000 | 0.00 | 0.00 | NO |
| chrX | 800000 | 900000 | 0.00 | 0.00 | NO |
| chrX | 900000 | 1000000 | 0.00 | 0.00 | NO |
| chrX | 1000000 | 1100000 | 0.00 | 0.00 | NO |
| chrX | 1100000 | 1200000 | 0.00 | 0.00 | NO |
| chrX | 1200000 | 1300000 | 0.00 | 0.00 | NO |
| chrX | 1300000 | 1400000 | 0.00 | 0.00 | NO |
| chrX | 1400000 | 1500000 | 0.00 | 0.00 | NO |
| chrX | 1500000 | 1600000 | 0.00 | 0.00 | NO |
| chrX | 1600000 | 1700000 | 0.00 | 0.00 | NO |
| chrX | 1700000 | 1800000 | 0.00 | 0.00 | NO |
| chrX | 1800000 | 1900000 | 0.00 | 0.00 | NO |
| chrX | 1900000 | 2000000 | 0.00 | 0.00 | NO |
| chrX | 2000000 | 2100000 | 0.00 | 0.00 | NO |
| chrX | 2100000 | 2200000 | 0.00 | 0.00 | NO |
| chrX | 2200000 | 2300000 | 0.00 | 0.00 | NO |
| chrX | 2300000 | 2400000 | 0.00 | 0.00 | NO |
| chrX | 2400000 | 2500000 | 0.00 | 0.00 | NO |
| chrX | 2500000 | 2600000 | 0.00 | 0.00 | NO |
| chrX | 2600000 | 2700000 | 0.00 | 0.00 | NO |
| chrX | 2700000 | 2800000 | 0.00 | 1.51 | NO |
| chrX | 2800000 | 2900000 | 1.40 | 1.23 | A>A |
| chrX | 2900000 | 3000000 | 1.14 | 0.79 | A>A |
| chrX | 3000000 | 3100000 | 0.48 | 0.32 | A>A |
| chrX | 3100000 | 3200000 | 0.61 | 0.35 | A>A |
| chrX | 3200000 | 3300000 | 0.44 | 0.06 | A>A |
| chrX | 3300000 | 3400000 | 0.37 | 0.18 | A>A |
| chrX | 3400000 | 3500000 | 0.61 | 0.33 | A>A |
| chrX | 3500000 | 3600000 | 0.94 | 0.53 | A>A |
| chrX | 3600000 | 3700000 | 1.10 | 0.58 | A>A |
| chrX | 3700000 | 3800000 | 0.00 | 0.00 | NO |
| chrX | 3800000 | 3900000 | 0.00 | 0.00 | NO |
| chrX | 3900000 | 4000000 | -0.39 | -0.55 | B>B |
| chrX | 4000000 | 4100000 | -0.64 | -0.69 | B>B |
| chrX | 4100000 | 4200000 | -0.72 | -0.57 | B>B |
| chrX | 4200000 | 4300000 | -0.71 | -0.61 | B>B |
| chrX | 4300000 | 4400000 | -0.42 | -0.45 | B>B |
| chrX | 4400000 | 4500000 | -0.15 | 0.03 | B>B |
| chrX | 4500000 | 4600000 | 0.29 | 0.57 | A>A |
| chrX | 4600000 | 4700000 | 0.47 | 0.54 | A>A |
| chrX | 4700000 | 4800000 | 0.64 | 0.64 | A>A |
| chrX | 4800000 | 4900000 | 0.52 | 0.61 | A>A |
| chrX | 4900000 | 5000000 | 0.23 | 0.54 | A>A |
| chrX | 5000000 | 5100000 | -0.09 | 0.12 | A>A |
| chrX | 5100000 | 5200000 | -0.38 | -0.05 | B>B |
| chrX | 5200000 | 5300000 | -0.47 | -0.53 | B>B |
| chrX | 5300000 | 5400000 | -0.61 | -0.52 | B>B |
| chrX | 5400000 | 5500000 | -0.68 | -0.63 | B>B |
| chrX | 5500000 | 5600000 | -0.69 | -0.63 | B>B |
| chrX | 5600000 | 5700000 | -0.66 | -0.56 | B>B |

| | | | | | |
|---|---|---|---|---|---|
| chrX | 5700000 | 5800000 | -0.65 | -0.61 | B>B |
| chrX | 5800000 | 5900000 | -0.68 | -0.59 | B>B |
| chrX | 5900000 | 6000000 | -0.61 | -0.58 | B>B |
| chrX | 6000000 | 6100000 | -0.65 | -0.57 | B>B |
| chrX | 6100000 | 6200000 | -0.51 | -0.40 | B>B |
| chrX | 6200000 | 6300000 | -0.55 | -0.57 | B>B |
| chrX | 6300000 | 6400000 | -0.67 | -0.57 | B>B |
| chrX | 6400000 | 6500000 | -0.56 | -0.57 | B>B |
| chrX | 6500000 | 6600000 | -0.59 | -0.54 | B>B |
| chrX | 6600000 | 6700000 | -0.50 | -0.48 | B>B |
| chrX | 6700000 | 6800000 | -0.52 | -0.55 | B>B |
| chrX | 6800000 | 6900000 | -0.45 | -0.50 | B>B |
| chrX | 6900000 | 7000000 | -0.02 | 0.01 | B>B |
| chrX | 7000000 | 7100000 | 0.18 | 0.33 | A>A |
| chrX | 7100000 | 7200000 | -0.24 | -0.25 | B>B |
| chrX | 7200000 | 7300000 | -0.42 | -0.45 | B>B |
| chrX | 7300000 | 7400000 | -0.49 | -0.53 | B>B |
| chrX | 7400000 | 7500000 | -0.56 | -0.54 | B>B |
| chrX | 7500000 | 7600000 | -0.54 | -0.54 | B>B |
| chrX | 7600000 | 7700000 | 0.00 | -0.61 | NO |
| chrX | 7700000 | 7800000 | -0.58 | -0.54 | B>B |
| chrX | 7800000 | 7900000 | -0.21 | -0.29 | B>B |
| chrX | 7900000 | 8000000 | -0.26 | -0.39 | B>B |
| chrX | 8000000 | 8100000 | -0.59 | -0.70 | B>B |
| chrX | 8100000 | 8200000 | -0.64 | -0.74 | B>B |
| chrX | 8200000 | 8300000 | -0.77 | -0.80 | B>B |
| chrX | 8300000 | 8400000 | -0.61 | -0.77 | B>B |
| chrX | 8400000 | 8500000 | -0.47 | -0.79 | B>B |
| chrX | 8500000 | 8600000 | -0.33 | -0.73 | B>B |
| chrX | 8600000 | 8700000 | -0.13 | -0.44 | B>B |
| chrX | 8700000 | 8800000 | 0.21 | 0.26 | A>A |
| chrX | 8800000 | 8900000 | 0.55 | 0.68 | A>A |
| chrX | 8900000 | 9000000 | 0.59 | 0.85 | A>A |
| chrX | 9000000 | 9100000 | 0.83 | 0.91 | A>A |
| chrX | 9100000 | 9200000 | 0.85 | 0.91 | A>A |
| chrX | 9200000 | 9300000 | 0.74 | 0.88 | A>A |
| chrX | 9300000 | 9400000 | 0.80 | 0.81 | A>A |
| chrX | 9400000 | 9500000 | 0.67 | 0.85 | A>A |
| chrX | 9500000 | 9600000 | 0.68 | 0.64 | A>A |
| chrX | 9600000 | 9700000 | 0.52 | 0.51 | A>A |
| chrX | 9700000 | 9800000 | 0.64 | 0.66 | A>A |
| chrX | 9800000 | 9900000 | 0.53 | 0.58 | A>A |
| chrX | 9900000 | 10000000 | 0.71 | 0.70 | A>A |
| chrX | 10000000 | 10100000 | 0.86 | 0.82 | A>A |
| chrX | 10100000 | 10200000 | 0.47 | 0.56 | A>A |
| chrX | 10200000 | 10300000 | 0.72 | 0.64 | A>A |
| chrX | 10300000 | 10400000 | 0.65 | 0.64 | A>A |
| chrX | 10400000 | 10500000 | 0.63 | 0.85 | A>A |
| chrX | 10500000 | 10600000 | 0.79 | 0.80 | A>A |
| chrX | 10600000 | 10700000 | 0.36 | 0.40 | A>A |
| chrX | 10700000 | 10800000 | 0.00 | 0.39 | NO |
| chrX | 10800000 | 10900000 | 0.22 | 0.31 | A>A |
| chrX | 10900000 | 11000000 | 0.13 | 0.28 | A>A |
| chrX | 11000000 | 11100000 | 0.20 | 0.16 | A>A |
| chrX | 11100000 | 11200000 | 0.11 | -0.06 | A>A |
| chrX | 11200000 | 11300000 | -0.37 | -0.72 | B>B |

| | | | | | |
|---|---|---|---|---|---|
| chrX | 11300000 | 11400000 | -0.34 | -0.91 | B>B |
| chrX | 11400000 | 11500000 | -0.25 | -0.79 | B>B |
| chrX | 11500000 | 11600000 | -0.16 | -0.48 | B>B |
| chrX | 11600000 | 11700000 | -0.14 | -0.68 | B>B |
| chrX | 11700000 | 11800000 | 0.11 | -0.25 | B>B |
| chrX | 11800000 | 11900000 | 0.35 | 0.38 | A>A |
| chrX | 11900000 | 12000000 | 0.40 | 0.46 | A>A |
| chrX | 12000000 | 12100000 | 0.41 | 0.49 | A>A |
| chrX | 12100000 | 12200000 | 0.39 | 0.56 | A>A |
| chrX | 12200000 | 12300000 | 0.42 | 0.58 | A>A |
| chrX | 12300000 | 12400000 | 0.45 | 0.34 | A>A |
| chrX | 12400000 | 12500000 | 0.27 | 0.08 | A>A |
| chrX | 12500000 | 12600000 | 0.01 | -0.05 | B>B |
| chrX | 12600000 | 12700000 | 0.15 | -0.18 | B>B |
| chrX | 12700000 | 12800000 | 0.23 | -0.06 | A>A |
| chrX | 12800000 | 12900000 | 0.55 | 0.22 | A>A |
| chrX | 12900000 | 13000000 | 0.55 | 0.33 | A>A |
| chrX | 13000000 | 13100000 | 1.08 | 0.85 | A>A |
| chrX | 13100000 | 13200000 | 0.90 | 0.82 | A>A |
| chrX | 13200000 | 13300000 | 1.00 | 0.73 | A>A |
| chrX | 13300000 | 13400000 | 0.87 | 0.89 | A>A |
| chrX | 13400000 | 13500000 | 0.78 | 0.68 | A>A |
| chrX | 13500000 | 13600000 | 0.45 | 0.22 | A>A |
| chrX | 13600000 | 13700000 | 0.67 | 0.37 | A>A |
| chrX | 13700000 | 13800000 | 0.53 | 0.32 | A>A |
| chrX | 13800000 | 13900000 | 0.38 | -0.22 | A>B |
| chrX | 13900000 | 14000000 | 0.28 | -0.36 | A>B |
| chrX | 14000000 | 14100000 | 0.12 | 0.01 | A>A |
| chrX | 14100000 | 14200000 | -0.26 | -0.41 | B>B |
| chrX | 14200000 | 14300000 | -0.28 | -0.64 | B>B |
| chrX | 14300000 | 14400000 | -0.38 | -0.64 | B>B |
| chrX | 14400000 | 14500000 | -0.47 | -0.62 | B>B |
| chrX | 14500000 | 14600000 | -0.35 | -0.57 | B>B |
| chrX | 14600000 | 14700000 | -0.22 | -0.59 | B>B |
| chrX | 14700000 | 14800000 | -0.19 | -0.55 | B>B |
| chrX | 14800000 | 14900000 | 0.23 | -0.13 | A>A |
| chrX | 14900000 | 15000000 | 0.28 | -0.07 | A>A |
| chrX | 15000000 | 15100000 | -0.09 | -0.44 | B>B |
| chrX | 15100000 | 15200000 | 0.09 | -0.27 | B>B |
| chrX | 15200000 | 15300000 | 0.36 | 0.03 | A>A |
| chrX | 15300000 | 15400000 | 0.70 | 0.57 | A>A |
| chrX | 15400000 | 15500000 | 0.68 | 0.48 | A>A |
| chrX | 15500000 | 15600000 | 0.65 | 0.43 | A>A |
| chrX | 15600000 | 15700000 | 0.58 | 0.53 | A>A |
| chrX | 15700000 | 15800000 | 0.71 | 0.65 | A>A |
| chrX | 15800000 | 15900000 | 0.60 | 0.49 | A>A |
| chrX | 15900000 | 16000000 | 0.38 | 0.25 | A>A |
| chrX | 16000000 | 16100000 | 0.30 | 0.29 | A>A |
| chrX | 16100000 | 16200000 | 0.31 | 0.27 | A>A |
| chrX | 16200000 | 16300000 | 0.34 | 0.17 | A>A |
| chrX | 16300000 | 16400000 | 0.02 | 0.16 | A>A |
| chrX | 16400000 | 16500000 | 0.31 | 0.17 | A>A |
| chrX | 16500000 | 16600000 | 0.49 | 0.24 | A>A |
| chrX | 16600000 | 16700000 | 0.59 | 0.46 | A>A |
| chrX | 16700000 | 16800000 | 0.87 | 0.85 | A>A |
| chrX | 16800000 | 16900000 | 0.94 | 0.83 | A>A |

| | | | | | |
|---|---|---|---|---|---|
| chrX | 16900000 | 17000000 | 1.03 | 0.84 | A>A |
| chrX | 17000000 | 17100000 | 0.78 | 0.72 | A>A |
| chrX | 17100000 | 17200000 | 0.41 | 0.51 | A>A |
| chrX | 17200000 | 17300000 | 0.49 | 0.46 | A>A |
| chrX | 17300000 | 17400000 | 0.60 | 0.61 | A>A |
| chrX | 17400000 | 17500000 | 0.87 | 0.79 | A>A |
| chrX | 17500000 | 17600000 | 0.58 | 0.78 | A>A |
| chrX | 17600000 | 17700000 | 0.66 | 0.83 | A>A |
| chrX | 17700000 | 17800000 | 0.47 | 0.46 | A>A |
| chrX | 17800000 | 17900000 | 0.36 | 0.30 | A>A |
| chrX | 17900000 | 18000000 | 0.28 | 0.17 | A>A |
| chrX | 18000000 | 18100000 | 0.25 | 0.14 | A>A |
| chrX | 18100000 | 18200000 | 0.27 | 0.15 | A>A |
| chrX | 18200000 | 18300000 | 0.48 | 0.36 | A>A |
| chrX | 18300000 | 18400000 | 0.72 | 0.56 | A>A |
| chrX | 18400000 | 18500000 | 0.70 | 0.67 | A>A |
| chrX | 18500000 | 18600000 | 0.79 | 0.82 | A>A |
| chrX | 18600000 | 18700000 | 0.88 | 0.80 | A>A |
| chrX | 18700000 | 18800000 | 0.87 | 0.90 | A>A |
| chrX | 18800000 | 18900000 | 0.91 | 0.85 | A>A |
| chrX | 18900000 | 19000000 | 0.68 | 0.56 | A>A |
| chrX | 19000000 | 19100000 | 0.69 | 0.66 | A>A |
| chrX | 19100000 | 19200000 | 0.70 | 0.63 | A>A |
| chrX | 19200000 | 19300000 | 0.68 | 0.54 | A>A |
| chrX | 19300000 | 19400000 | 0.49 | 0.49 | A>A |
| chrX | 19400000 | 19500000 | 0.75 | 0.89 | A>A |
| chrX | 19500000 | 19600000 | 0.79 | 0.85 | A>A |
| chrX | 19600000 | 19700000 | 0.92 | 0.86 | A>A |
| chrX | 19700000 | 19800000 | 0.82 | 1.05 | A>A |
| chrX | 19800000 | 19900000 | 0.83 | 0.85 | A>A |
| chrX | 19900000 | 20000000 | 0.69 | 0.86 | A>A |
| chrX | 20000000 | 20100000 | 0.82 | 0.91 | A>A |
| chrX | 20100000 | 20200000 | 0.76 | 0.80 | A>A |
| chrX | 20200000 | 20300000 | 0.81 | 0.81 | A>A |
| chrX | 20300000 | 20400000 | 0.97 | 0.91 | A>A |
| chrX | 20400000 | 20500000 | 0.78 | 0.70 | A>A |
| chrX | 20500000 | 20600000 | 0.27 | 0.47 | A>A |
| chrX | 20600000 | 20700000 | 0.44 | 0.41 | A>A |
| chrX | 20700000 | 20800000 | 0.17 | 0.48 | A>A |
| chrX | 20800000 | 20900000 | 0.11 | 0.43 | A>A |
| chrX | 20900000 | 21000000 | 0.17 | 0.40 | A>A |
| chrX | 21000000 | 21100000 | 0.15 | 0.38 | A>A |
| chrX | 21100000 | 21200000 | 0.25 | 0.52 | A>A |
| chrX | 21200000 | 21300000 | 0.21 | 0.53 | A>A |
| chrX | 21300000 | 21400000 | 0.11 | 0.32 | A>A |
| chrX | 21400000 | 21500000 | 0.24 | 0.40 | A>A |
| chrX | 21500000 | 21600000 | 0.17 | 0.30 | A>A |
| chrX | 21600000 | 21700000 | 0.30 | 0.26 | A>A |
| chrX | 21700000 | 21800000 | 0.38 | 0.28 | A>A |
| chrX | 21800000 | 21900000 | 0.50 | 0.64 | A>A |
| chrX | 21900000 | 22000000 | 0.67 | 0.50 | A>A |
| chrX | 22000000 | 22100000 | 0.39 | 0.27 | A>A |
| chrX | 22100000 | 22200000 | 0.32 | 0.36 | A>A |
| chrX | 22200000 | 22300000 | 0.11 | 0.11 | A>A |
| chrX | 22300000 | 22400000 | 0.03 | -0.11 | B>B |
| chrX | 22400000 | 22500000 | -0.22 | -0.36 | B>B |

| | | | | | |
|---|---|---|---|---|---|
| chrX | 22500000 | 22600000 | -0.24 | -0.49 | B>B |
| chrX | 22600000 | 22700000 | -0.33 | -0.51 | B>B |
| chrX | 22700000 | 22800000 | -0.10 | -0.36 | B>B |
| chrX | 22800000 | 22900000 | -0.16 | -0.37 | B>B |
| chrX | 22900000 | 23000000 | -0.09 | -0.39 | B>B |
| chrX | 23000000 | 23100000 | -0.08 | -0.42 | B>B |
| chrX | 23100000 | 23200000 | -0.10 | -0.50 | B>B |
| chrX | 23200000 | 23300000 | -0.10 | -0.36 | B>B |
| chrX | 23300000 | 23400000 | 0.17 | -0.18 | B>B |
| chrX | 23400000 | 23500000 | -0.06 | -0.18 | B>B |
| chrX | 23500000 | 23600000 | 0.00 | -0.06 | B>B |
| chrX | 23600000 | 23700000 | 0.37 | 0.20 | A>A |
| chrX | 23700000 | 23800000 | 0.94 | 0.87 | A>A |
| chrX | 23800000 | 23900000 | 1.27 | 1.06 | A>A |
| chrX | 23900000 | 24000000 | 0.99 | 0.80 | A>A |
| chrX | 24000000 | 24100000 | 0.95 | 0.97 | A>A |
| chrX | 24100000 | 24200000 | 1.27 | 0.99 | A>A |
| chrX | 24200000 | 24300000 | 0.98 | 0.68 | A>A |
| chrX | 24300000 | 24400000 | 0.79 | 0.48 | A>A |
| chrX | 24400000 | 24500000 | 0.86 | 0.38 | A>A |
| chrX | 24500000 | 24600000 | 0.84 | 0.28 | A>A |
| chrX | 24600000 | 24700000 | 0.72 | 0.17 | A>A |
| chrX | 24700000 | 24800000 | 0.85 | 0.33 | A>A |
| chrX | 24800000 | 24900000 | 0.87 | 0.57 | A>A |
| chrX | 24900000 | 25000000 | 0.93 | 0.71 | A>A |
| chrX | 25000000 | 25100000 | 0.60 | 0.33 | A>A |
| chrX | 25100000 | 25200000 | 0.27 | -0.02 | A>A |
| chrX | 25200000 | 25300000 | -0.11 | -0.57 | B>B |
| chrX | 25300000 | 25400000 | -0.35 | -0.81 | B>B |
| chrX | 25400000 | 25500000 | -0.55 | -0.75 | B>B |
| chrX | 25500000 | 25600000 | -0.62 | -0.73 | B>B |
| chrX | 25600000 | 25700000 | -0.71 | -0.78 | B>B |
| chrX | 25700000 | 25800000 | -0.72 | -0.83 | B>B |
| chrX | 25800000 | 25900000 | -0.69 | -0.84 | B>B |
| chrX | 25900000 | 26000000 | -0.68 | -0.86 | B>B |
| chrX | 26000000 | 26100000 | -0.78 | -0.84 | B>B |
| chrX | 26100000 | 26200000 | -0.82 | -0.89 | B>B |
| chrX | 26200000 | 26300000 | -0.70 | -0.85 | B>B |
| chrX | 26300000 | 26400000 | -0.78 | -0.89 | B>B |
| chrX | 26400000 | 26500000 | -0.83 | -0.90 | B>B |
| chrX | 26500000 | 26600000 | -0.79 | -0.83 | B>B |
| chrX | 26600000 | 26700000 | -0.55 | -0.57 | B>B |
| chrX | 26700000 | 26800000 | -0.80 | -0.72 | B>B |
| chrX | 26800000 | 26900000 | -0.72 | -0.82 | B>B |
| chrX | 26900000 | 27000000 | -0.85 | -0.75 | B>B |
| chrX | 27000000 | 27100000 | -0.84 | -0.89 | B>B |
| chrX | 27100000 | 27200000 | -0.74 | -0.73 | B>B |
| chrX | 27200000 | 27300000 | -0.57 | -0.58 | B>B |
| chrX | 27300000 | 27400000 | -0.60 | -0.02 | B>A |
| chrX | 27400000 | 27500000 | -0.48 | 0.32 | B>A |
| chrX | 27500000 | 27600000 | -0.35 | 0.45 | B>A |
| chrX | 27600000 | 27700000 | -0.21 | 0.46 | B>A |
| chrX | 27700000 | 27800000 | 0.05 | 0.70 | B>A |
| chrX | 27800000 | 27900000 | 0.10 | 0.75 | B>A |
| chrX | 27900000 | 28000000 | 0.20 | 0.71 | A>A |
| chrX | 28000000 | 28100000 | 0.13 | 0.78 | B>A |

| | | | | | |
|---|---|---|---|---|---|
| chrX | 28100000 | 28200000 | 0.03 | 0.71 | B>A |
| chrX | 28200000 | 28300000 | -0.18 | 0.75 | B>A |
| chrX | 28300000 | 28400000 | -0.15 | 0.79 | B>A |
| chrX | 28400000 | 28500000 | -0.25 | 0.79 | B>A |
| chrX | 28500000 | 28600000 | -0.13 | 0.59 | B>A |
| chrX | 28600000 | 28700000 | -0.12 | 0.53 | B>A |
| chrX | 28700000 | 28800000 | -0.22 | 0.50 | B>A |
| chrX | 28800000 | 28900000 | -0.03 | 0.46 | A>A |
| chrX | 28900000 | 29000000 | 0.01 | 0.47 | A>A |
| chrX | 29000000 | 29100000 | 0.02 | 0.36 | A>A |
| chrX | 29100000 | 29200000 | 0.14 | 0.18 | A>A |
| chrX | 29200000 | 29300000 | 0.44 | 0.23 | A>A |
| chrX | 29300000 | 29400000 | 0.53 | 0.32 | A>A |
| chrX | 29400000 | 29500000 | 0.61 | 0.38 | A>A |
| chrX | 29500000 | 29600000 | 0.70 | 0.40 | A>A |
| chrX | 29600000 | 29700000 | 0.63 | 0.44 | A>A |
| chrX | 29700000 | 29800000 | 0.72 | 0.31 | A>A |
| chrX | 29800000 | 29900000 | 0.65 | 0.30 | A>A |
| chrX | 29900000 | 30000000 | 0.33 | 0.02 | A>A |
| chrX | 30000000 | 30100000 | 0.29 | -0.10 | A>A |
| chrX | 30100000 | 30200000 | 0.11 | -0.08 | A>A |
| chrX | 30200000 | 30300000 | 0.37 | -0.08 | A>A |
| chrX | 30300000 | 30400000 | 0.47 | -0.10 | A>A |
| chrX | 30400000 | 30500000 | 0.39 | -0.38 | A>B |
| chrX | 30500000 | 30600000 | 0.22 | -0.17 | A>A |
| chrX | 30600000 | 30700000 | 0.40 | 0.06 | A>A |
| chrX | 30700000 | 30800000 | 0.17 | -0.19 | B>B |
| chrX | 30800000 | 30900000 | 0.27 | -0.09 | A>A |
| chrX | 30900000 | 31000000 | 0.18 | -0.53 | A>B |
| chrX | 31000000 | 31100000 | -0.29 | -0.73 | B>B |
| chrX | 31100000 | 31200000 | -0.52 | -0.81 | B>B |
| chrX | 31200000 | 31300000 | -0.44 | -0.78 | B>B |
| chrX | 31300000 | 31400000 | -0.47 | -0.76 | B>B |
| chrX | 31400000 | 31500000 | -0.57 | -0.88 | B>B |
| chrX | 31500000 | 31600000 | -0.69 | -0.84 | B>B |
| chrX | 31600000 | 31700000 | -0.59 | -0.76 | B>B |
| chrX | 31700000 | 31800000 | -0.61 | -0.83 | B>B |
| chrX | 31800000 | 31900000 | -0.77 | -0.86 | B>B |
| chrX | 31900000 | 32000000 | -0.78 | -0.86 | B>B |
| chrX | 32000000 | 32100000 | -0.88 | -0.91 | B>B |
| chrX | 32100000 | 32200000 | -0.77 | -0.89 | B>B |
| chrX | 32200000 | 32300000 | -0.73 | -0.88 | B>B |
| chrX | 32300000 | 32400000 | -0.80 | -0.91 | B>B |
| chrX | 32400000 | 32500000 | -0.82 | -0.93 | B>B |
| chrX | 32500000 | 32600000 | -0.75 | -0.81 | B>B |
| chrX | 32600000 | 32700000 | -0.74 | -0.91 | B>B |
| chrX | 32700000 | 32800000 | -0.90 | -0.91 | B>B |
| chrX | 32800000 | 32900000 | -0.78 | -0.85 | B>B |
| chrX | 32900000 | 33000000 | -0.82 | -0.86 | B>B |
| chrX | 33000000 | 33100000 | -0.78 | -0.92 | B>B |
| chrX | 33100000 | 33200000 | -0.76 | -0.87 | B>B |
| chrX | 33200000 | 33300000 | -0.86 | -0.93 | B>B |
| chrX | 33300000 | 33400000 | -0.72 | -0.87 | B>B |
| chrX | 33400000 | 33500000 | -0.67 | -0.73 | B>B |
| chrX | 33500000 | 33600000 | -0.74 | -0.83 | B>B |
| chrX | 33600000 | 33700000 | -0.75 | -0.85 | B>B |

| | | | | | |
|---|---|---|---|---|---|
| chrX | 33700000 | 33800000 | -0.67 | -0.86 | B>B |
| chrX | 33800000 | 33900000 | -0.83 | -0.81 | B>B |
| chrX | 33900000 | 34000000 | -0.76 | -0.83 | B>B |
| chrX | 34000000 | 34100000 | -0.76 | -0.90 | B>B |
| chrX | 34100000 | 34200000 | -0.70 | -0.83 | B>B |
| chrX | 34200000 | 34300000 | -0.71 | -0.74 | B>B |
| chrX | 34300000 | 34400000 | -0.50 | -0.59 | B>B |
| chrX | 34400000 | 34500000 | -0.44 | -0.49 | B>B |
| chrX | 34500000 | 34600000 | -0.19 | -0.06 | B>B |
| chrX | 34600000 | 34700000 | -0.01 | 0.25 | A>A |
| chrX | 34700000 | 34800000 | 0.14 | 0.50 | A>A |
| chrX | 34800000 | 34900000 | 0.22 | 0.53 | A>A |
| chrX | 34900000 | 35000000 | 0.25 | 0.52 | A>A |
| chrX | 35000000 | 35100000 | 0.40 | 0.59 | A>A |
| chrX | 35100000 | 35200000 | 0.31 | 0.73 | A>A |
| chrX | 35200000 | 35300000 | 0.41 | 0.69 | A>A |
| chrX | 35300000 | 35400000 | 0.48 | 0.67 | A>A |
| chrX | 35400000 | 35500000 | 0.44 | 0.72 | A>A |
| chrX | 35500000 | 35600000 | 0.54 | 0.74 | A>A |
| chrX | 35600000 | 35700000 | 0.58 | 0.72 | A>A |
| chrX | 35700000 | 35800000 | 0.38 | 0.66 | A>A |
| chrX | 35800000 | 35900000 | 0.28 | 0.67 | A>A |
| chrX | 35900000 | 36000000 | 0.18 | 0.50 | A>A |
| chrX | 36000000 | 36100000 | 0.02 | 0.64 | B>A |
| chrX | 36100000 | 36200000 | -0.13 | 0.61 | B>A |
| chrX | 36200000 | 36300000 | -0.09 | 0.63 | B>A |
| chrX | 36300000 | 36400000 | -0.12 | 0.69 | B>A |
| chrX | 36400000 | 36500000 | -0.11 | 0.72 | B>A |
| chrX | 36500000 | 36600000 | -0.27 | 0.58 | B>A |
| chrX | 36600000 | 36700000 | -0.23 | 0.49 | B>A |
| chrX | 36700000 | 36800000 | -0.30 | 0.28 | B>A |
| chrX | 36800000 | 36900000 | -0.33 | 0.27 | B>A |
| chrX | 36900000 | 37000000 | -0.18 | 0.12 | B>B |
| chrX | 37000000 | 37100000 | -0.22 | -0.09 | B>B |
| chrX | 37100000 | 37200000 | 0.00 | 0.00 | NO |
| chrX | 37200000 | 37300000 | 0.23 | 0.27 | A>A |
| chrX | 37300000 | 37400000 | 0.05 | 0.09 | A>A |
| chrX | 37400000 | 37500000 | -0.12 | -0.02 | B>B |
| chrX | 37500000 | 37600000 | -0.05 | 0.11 | A>A |
| chrX | 37600000 | 37700000 | 0.17 | 0.25 | A>A |
| chrX | 37700000 | 37800000 | 0.22 | 0.35 | A>A |
| chrX | 37800000 | 37900000 | 0.06 | 0.38 | A>A |
| chrX | 37900000 | 38000000 | -0.02 | 0.34 | A>A |
| chrX | 38000000 | 38100000 | 0.25 | 0.25 | A>A |
| chrX | 38100000 | 38200000 | 0.54 | 0.27 | A>A |
| chrX | 38200000 | 38300000 | 0.09 | -0.54 | A>B |
| chrX | 38300000 | 38400000 | -0.19 | -0.73 | B>B |
| chrX | 38400000 | 38500000 | 0.00 | -0.55 | B>B |
| chrX | 38500000 | 38600000 | 0.12 | -0.38 | B>B |
| chrX | 38600000 | 38700000 | 0.27 | -0.08 | A>A |
| chrX | 38700000 | 38800000 | 0.23 | -0.39 | A>B |
| chrX | 38800000 | 38900000 | 0.12 | -0.47 | A>B |
| chrX | 38900000 | 39000000 | 0.09 | -0.47 | B>B |
| chrX | 39000000 | 39100000 | 0.06 | -0.37 | B>B |
| chrX | 39100000 | 39200000 | 0.15 | -0.33 | B>B |
| chrX | 39200000 | 39300000 | 0.21 | -0.15 | A>A |

| | | | | | |
|---|---|---|---|---|---|
| chrX | 39300000 | 39400000 | 0.39 | 0.11 | A>A |
| chrX | 39400000 | 39500000 | 0.54 | 0.24 | A>A |
| chrX | 39500000 | 39600000 | 0.67 | 0.42 | A>A |
| chrX | 39600000 | 39700000 | 0.78 | 0.67 | A>A |
| chrX | 39700000 | 39800000 | 0.96 | 0.83 | A>A |
| chrX | 39800000 | 39900000 | 0.83 | 0.83 | A>A |
| chrX | 39900000 | 40000000 | 0.87 | 0.66 | A>A |
| chrX | 40000000 | 40100000 | 0.78 | 0.63 | A>A |
| chrX | 40100000 | 40200000 | 0.55 | 0.48 | A>A |
| chrX | 40200000 | 40300000 | 0.65 | 0.54 | A>A |
| chrX | 40300000 | 40400000 | 0.61 | 0.56 | A>A |
| chrX | 40400000 | 40500000 | 0.79 | 0.75 | A>A |
| chrX | 40500000 | 40600000 | 0.89 | 1.00 | A>A |
| chrX | 40600000 | 40700000 | 1.11 | 0.96 | A>A |
| chrX | 40700000 | 40800000 | 0.92 | 0.89 | A>A |
| chrX | 40800000 | 40900000 | 1.12 | 1.08 | A>A |
| chrX | 40900000 | 41000000 | 0.96 | 1.11 | A>A |
| chrX | 41000000 | 41100000 | 0.95 | 1.06 | A>A |
| chrX | 41100000 | 41200000 | 0.92 | 0.92 | A>A |
| chrX | 41200000 | 41300000 | 1.25 | 1.03 | A>A |
| chrX | 41300000 | 41400000 | 0.80 | 0.77 | A>A |
| chrX | 41400000 | 41500000 | 0.57 | 0.74 | A>A |
| chrX | 41500000 | 41600000 | 0.53 | 0.71 | A>A |
| chrX | 41600000 | 41700000 | 0.36 | 0.34 | A>A |
| chrX | 41700000 | 41800000 | 0.53 | 0.42 | A>A |
| chrX | 41800000 | 41900000 | 0.31 | 0.20 | A>A |
| chrX | 41900000 | 42000000 | 0.38 | 0.28 | A>A |
| chrX | 42000000 | 42100000 | 0.28 | 0.12 | A>A |
| chrX | 42100000 | 42200000 | 0.02 | -0.07 | B>B |
| chrX | 42200000 | 42300000 | 0.06 | -0.21 | B>B |
| chrX | 42300000 | 42400000 | -0.14 | -0.20 | B>B |
| chrX | 42400000 | 42500000 | -0.17 | -0.11 | B>B |
| chrX | 42500000 | 42600000 | -0.16 | 0.03 | B>B |
| chrX | 42600000 | 42700000 | -0.13 | 0.15 | A>A |
| chrX | 42700000 | 42800000 | -0.25 | -0.20 | B>B |
| chrX | 42800000 | 42900000 | -0.37 | -0.31 | B>B |
| chrX | 42900000 | 43000000 | -0.20 | -0.36 | B>B |
| chrX | 43000000 | 43100000 | -0.28 | -0.37 | B>B |
| chrX | 43100000 | 43200000 | -0.30 | -0.20 | B>B |
| chrX | 43200000 | 43300000 | -0.30 | -0.16 | B>B |
| chrX | 43300000 | 43400000 | -0.11 | -0.04 | B>B |
| chrX | 43400000 | 43500000 | 0.30 | 0.38 | A>A |
| chrX | 43500000 | 43600000 | 0.49 | 0.55 | A>A |
| chrX | 43600000 | 43700000 | 0.21 | 0.41 | A>A |
| chrX | 43700000 | 43800000 | 0.53 | 0.72 | A>A |
| chrX | 43800000 | 43900000 | 0.57 | 0.66 | A>A |
| chrX | 43900000 | 44000000 | 0.48 | 0.71 | A>A |
| chrX | 44000000 | 44100000 | 0.54 | 0.79 | A>A |
| chrX | 44100000 | 44200000 | 0.59 | 0.70 | A>A |
| chrX | 44200000 | 44300000 | 0.65 | 0.85 | A>A |
| chrX | 44300000 | 44400000 | 0.76 | 0.79 | A>A |
| chrX | 44400000 | 44500000 | 0.68 | 0.89 | A>A |
| chrX | 44500000 | 44600000 | 0.74 | 0.85 | A>A |
| chrX | 44600000 | 44700000 | 0.76 | 0.92 | A>A |
| chrX | 44700000 | 44800000 | 0.86 | 0.92 | A>A |
| chrX | 44800000 | 44900000 | 0.90 | 0.88 | A>A |

| | | | | | |
|---|---|---|---|---|---|
| chrX | 44900000 | 45000000 | 0.75 | 0.90 | A>A |
| chrX | 45000000 | 45100000 | 0.57 | 0.78 | A>A |
| chrX | 45100000 | 45200000 | 0.72 | 0.76 | A>A |
| chrX | 45200000 | 45300000 | 0.78 | 0.85 | A>A |
| chrX | 45300000 | 45400000 | 0.59 | 0.71 | A>A |
| chrX | 45400000 | 45500000 | 0.80 | 0.87 | A>A |
| chrX | 45500000 | 45600000 | 0.74 | 0.76 | A>A |
| chrX | 45600000 | 45700000 | 0.78 | 0.72 | A>A |
| chrX | 45700000 | 45800000 | 0.93 | 0.90 | A>A |
| chrX | 45800000 | 45900000 | 0.82 | 0.85 | A>A |
| chrX | 45900000 | 46000000 | 0.85 | 0.94 | A>A |
| chrX | 46000000 | 46100000 | 0.95 | 0.95 | A>A |
| chrX | 46100000 | 46200000 | 0.87 | 0.91 | A>A |
| chrX | 46200000 | 46300000 | 0.64 | 0.77 | A>A |
| chrX | 46300000 | 46400000 | 0.70 | 0.95 | A>A |
| chrX | 46400000 | 46500000 | 0.98 | 0.92 | A>A |
| chrX | 46500000 | 46600000 | 0.89 | 0.99 | A>A |
| chrX | 46600000 | 46700000 | 1.09 | 0.93 | A>A |
| chrX | 46700000 | 46800000 | 1.01 | 0.98 | A>A |
| chrX | 46800000 | 46900000 | 1.01 | 1.01 | A>A |
| chrX | 46900000 | 47000000 | 0.90 | 0.86 | A>A |
| chrX | 47000000 | 47100000 | 0.99 | 0.96 | A>A |
| chrX | 47100000 | 47200000 | 0.97 | 0.97 | A>A |
| chrX | 47200000 | 47300000 | 1.02 | 0.90 | A>A |
| chrX | 47300000 | 47400000 | 0.83 | 0.91 | A>A |
| chrX | 47400000 | 47500000 | 1.11 | 1.02 | A>A |
| chrX | 47500000 | 47600000 | 0.59 | 0.90 | A>A |
| chrX | 47600000 | 47700000 | 0.52 | 0.90 | A>A |
| chrX | 47700000 | 47800000 | 0.42 | 0.98 | B>A |
| chrX | 47800000 | 47900000 | 0.45 | 0.89 | A>A |
| chrX | 47900000 | 48000000 | 0.00 | 0.79 | NO |
| chrX | 48000000 | 48100000 | 0.00 | 0.00 | NO |
| chrX | 48100000 | 48200000 | 0.00 | 0.00 | NO |
| chrX | 48200000 | 48300000 | 0.00 | 0.00 | NO |
| chrX | 48300000 | 48400000 | 0.90 | 0.93 | A>A |
| chrX | 48400000 | 48500000 | 0.99 | 0.80 | A>A |
| chrX | 48500000 | 48600000 | 0.94 | 0.87 | A>A |
| chrX | 48600000 | 48700000 | 1.01 | 0.88 | A>A |
| chrX | 48700000 | 48800000 | 1.09 | 0.84 | A>A |
| chrX | 48800000 | 48900000 | 1.20 | 1.03 | A>A |
| chrX | 48900000 | 49000000 | 1.16 | 1.03 | A>A |
| chrX | 49000000 | 49100000 | 1.35 | 0.95 | A>A |
| chrX | 49100000 | 49200000 | 0.86 | 0.70 | A>A |
| chrX | 49200000 | 49300000 | 0.00 | 0.00 | NO |
| chrX | 49300000 | 49400000 | 0.00 | 0.00 | NO |
| chrX | 49400000 | 49500000 | 0.35 | -0.01 | A>A |
| chrX | 49500000 | 49600000 | 0.35 | 0.04 | A>A |
| chrX | 49600000 | 49700000 | 0.33 | 0.16 | A>A |
| chrX | 49700000 | 49800000 | 0.56 | 0.16 | A>A |
| chrX | 49800000 | 49900000 | 0.31 | -0.05 | A>A |
| chrX | 49900000 | 50000000 | 0.26 | -0.33 | A>B |
| chrX | 50000000 | 50100000 | 0.00 | -0.65 | NO |
| chrX | 50100000 | 50200000 | -0.15 | -0.78 | A>B |
| chrX | 50200000 | 50300000 | -0.21 | -0.72 | B>B |
| chrX | 50300000 | 50400000 | -0.21 | -0.70 | B>B |
| chrX | 50400000 | 50500000 | -0.25 | -0.78 | B>B |

| | | | | | |
|---|---|---|---|---|---|
| chrX | 50500000 | 50600000 | -0.14 | -0.82 | A>B |
| chrX | 50600000 | 50700000 | -0.21 | -0.85 | A>B |
| chrX | 50700000 | 50800000 | -0.34 | -0.87 | B>B |
| chrX | 50800000 | 50900000 | -0.42 | -0.84 | B>B |
| chrX | 50900000 | 51000000 | -0.52 | -0.89 | B>B |
| chrX | 51000000 | 51100000 | -0.44 | -0.79 | B>B |
| chrX | 51100000 | 51200000 | -0.13 | -0.48 | B>B |
| chrX | 51200000 | 51300000 | 0.03 | -0.23 | B>B |
| chrX | 51300000 | 51400000 | -0.13 | -0.35 | B>B |
| chrX | 51400000 | 51500000 | 0.00 | 0.00 | NO |
| chrX | 51500000 | 51600000 | 0.45 | 0.27 | A>A |
| chrX | 51600000 | 51700000 | 0.08 | 0.03 | A>A |
| chrX | 51700000 | 51800000 | 0.00 | 0.00 | NO |
| chrX | 51800000 | 51900000 | 0.00 | 0.00 | NO |
| chrX | 51900000 | 52000000 | 0.00 | 0.00 | NO |
| chrX | 52000000 | 52100000 | 0.15 | -0.04 | A>A |
| chrX | 52100000 | 52200000 | 0.00 | 0.00 | NO |
| chrX | 52200000 | 52300000 | 0.00 | 0.00 | NO |
| chrX | 52300000 | 52400000 | 0.00 | 0.00 | NO |
| chrX | 52400000 | 52500000 | 0.00 | 0.00 | NO |
| chrX | 52500000 | 52600000 | 0.00 | 0.00 | NO |
| chrX | 52600000 | 52700000 | 0.00 | 0.00 | NO |
| chrX | 52700000 | 52800000 | 0.00 | 0.00 | NO |
| chrX | 52800000 | 52900000 | 0.00 | 0.00 | NO |
| chrX | 52900000 | 53000000 | 0.00 | 0.00 | NO |
| chrX | 53000000 | 53100000 | 1.02 | 0.83 | A>A |
| chrX | 53100000 | 53200000 | 1.24 | 1.04 | A>A |
| chrX | 53200000 | 53300000 | 1.12 | 1.06 | A>A |
| chrX | 53300000 | 53400000 | 1.20 | 1.00 | A>A |
| chrX | 53400000 | 53500000 | 1.18 | 1.09 | A>A |
| chrX | 53500000 | 53600000 | 1.14 | 1.01 | A>A |
| chrX | 53600000 | 53700000 | 1.13 | 1.01 | A>A |
| chrX | 53700000 | 53800000 | 1.09 | 0.71 | A>A |
| chrX | 53800000 | 53900000 | 0.91 | 0.66 | A>A |
| chrX | 53900000 | 54000000 | 0.83 | 0.67 | A>A |
| chrX | 54000000 | 54100000 | 0.98 | 0.83 | A>A |
| chrX | 54100000 | 54200000 | 0.91 | 0.81 | A>A |
| chrX | 54200000 | 54300000 | 0.78 | 0.53 | A>A |
| chrX | 54300000 | 54400000 | 0.71 | 0.51 | A>A |
| chrX | 54400000 | 54500000 | 0.86 | 0.64 | A>A |
| chrX | 54500000 | 54600000 | 1.03 | 0.82 | A>A |
| chrX | 54600000 | 54700000 | 0.70 | 0.53 | A>A |
| chrX | 54700000 | 54800000 | 0.35 | 0.27 | A>A |
| chrX | 54800000 | 54900000 | 0.56 | 0.29 | A>A |
| chrX | 54900000 | 55000000 | 0.35 | 0.02 | A>A |
| chrX | 55000000 | 55100000 | 0.41 | 0.15 | A>A |
| chrX | 55100000 | 55200000 | 0.22 | 0.24 | A>A |
| chrX | 55200000 | 55300000 | 0.09 | -0.12 | B>B |
| chrX | 55300000 | 55400000 | -0.16 | -0.18 | B>B |
| chrX | 55400000 | 55500000 | 0.00 | 0.00 | NO |
| chrX | 55500000 | 55600000 | 0.00 | -0.17 | NO |
| chrX | 55600000 | 55700000 | -0.33 | -0.19 | B>B |
| chrX | 55700000 | 55800000 | 0.17 | 0.18 | A>A |
| chrX | 55800000 | 55900000 | -0.24 | 0.03 | B>B |
| chrX | 55900000 | 56000000 | -0.30 | 0.06 | B>B |
| chrX | 56000000 | 56100000 | -0.27 | 0.30 | B>A |

| | | | | | |
|---|---|---|---|---|---|
| chrX | 56100000 | 56200000 | -0.18 | 0.55 | B>A |
| chrX | 56200000 | 56300000 | -0.13 | 0.16 | A>A |
| chrX | 56300000 | 56400000 | 0.00 | 0.00 | NO |
| chrX | 56400000 | 56500000 | 0.00 | 0.00 | NO |
| chrX | 56500000 | 56600000 | 0.11 | -0.01 | A>A |
| chrX | 56600000 | 56700000 | -0.04 | 0.01 | B>B |
| chrX | 56700000 | 56800000 | 0.11 | 0.17 | A>A |
| chrX | 56800000 | 56900000 | 0.25 | 0.33 | A>A |
| chrX | 56900000 | 57000000 | 0.00 | 0.00 | NO |
| chrX | 57000000 | 57100000 | 0.20 | 0.31 | A>A |
| chrX | 57100000 | 57200000 | 0.31 | 0.44 | A>A |
| chrX | 57200000 | 57300000 | -0.04 | 0.26 | A>A |
| chrX | 57300000 | 57400000 | 0.02 | 0.52 | A>A |
| chrX | 57400000 | 57500000 | -0.23 | 0.48 | B>A |
| chrX | 57500000 | 57600000 | -0.26 | 0.08 | B>B |
| chrX | 57600000 | 57700000 | -0.02 | 0.16 | A>A |
| chrX | 57700000 | 57800000 | -0.23 | -0.09 | B>B |
| chrX | 57800000 | 57900000 | 0.00 | -0.21 | NO |
| chrX | 57900000 | 58000000 | -0.23 | -0.16 | B>B |
| chrX | 58000000 | 58100000 | -0.71 | -0.37 | B>B |
| chrX | 58100000 | 58200000 | -0.98 | -0.49 | B>B |
| chrX | 58200000 | 58300000 | -0.77 | -0.36 | B>B |
| chrX | 58300000 | 58400000 | -0.90 | 0.18 | B>A |
| chrX | 58400000 | 58500000 | 0.00 | 0.00 | NO |
| chrX | 58500000 | 58600000 | 0.00 | 0.00 | NO |
| chrX | 58600000 | 58700000 | 0.00 | 0.00 | NO |
| chrX | 58700000 | 58800000 | 0.00 | 0.00 | NO |
| chrX | 58800000 | 58900000 | 0.00 | 0.00 | NO |
| chrX | 58900000 | 59000000 | 0.00 | 0.00 | NO |
| chrX | 59000000 | 59100000 | 0.00 | 0.00 | NO |
| chrX | 59100000 | 59200000 | 0.00 | 0.00 | NO |
| chrX | 59200000 | 59300000 | 0.00 | 0.00 | NO |
| chrX | 59300000 | 59400000 | 0.00 | 0.00 | NO |
| chrX | 59400000 | 59500000 | 0.00 | 0.00 | NO |
| chrX | 59500000 | 59600000 | 0.00 | 0.00 | NO |
| chrX | 59600000 | 59700000 | 0.00 | 0.00 | NO |
| chrX | 59700000 | 59800000 | 0.00 | 0.00 | NO |
| chrX | 59800000 | 59900000 | 0.00 | 0.00 | NO |
| chrX | 59900000 | 60000000 | 0.00 | 0.00 | NO |
| chrX | 60000000 | 60100000 | 0.00 | 0.00 | NO |
| chrX | 60100000 | 60200000 | 0.00 | 0.00 | NO |
| chrX | 60200000 | 60300000 | 0.00 | 0.00 | NO |
| chrX | 60300000 | 60400000 | 0.00 | 0.00 | NO |
| chrX | 60400000 | 60500000 | 0.00 | 0.00 | NO |
| chrX | 60500000 | 60600000 | 0.00 | 0.00 | NO |
| chrX | 60600000 | 60700000 | 0.00 | 0.00 | NO |
| chrX | 60700000 | 60800000 | 0.00 | 0.00 | NO |
| chrX | 60800000 | 60900000 | 0.00 | 0.00 | NO |
| chrX | 60900000 | 61000000 | 0.00 | 0.00 | NO |
| chrX | 61000000 | 61100000 | 0.00 | 0.00 | NO |
| chrX | 61100000 | 61200000 | 0.00 | 0.00 | NO |
| chrX | 61200000 | 61300000 | 0.00 | 0.00 | NO |
| chrX | 61300000 | 61400000 | 0.00 | 0.00 | NO |
| chrX | 61400000 | 61500000 | 0.00 | 0.00 | NO |
| chrX | 61500000 | 61600000 | 0.00 | 0.00 | NO |
| chrX | 61600000 | 61700000 | 0.00 | 0.00 | NO |

| | | | | | |
|---|---|---|---|---|---|
| chrX | 61700000 | 61800000 | 0.00 | 0.00 | NO |
| chrX | 61800000 | 61900000 | 0.00 | 0.00 | NO |
| chrX | 61900000 | 62000000 | 0.00 | 0.40 | NO |
| chrX | 62000000 | 62100000 | -1.01 | 0.15 | B>A |
| chrX | 62100000 | 62200000 | 0.00 | -0.30 | NO |
| chrX | 62200000 | 62300000 | -1.04 | -0.36 | B>A |
| chrX | 62300000 | 62400000 | 0.00 | 0.00 | NO |
| chrX | 62400000 | 62500000 | 0.00 | 0.00 | NO |
| chrX | 62500000 | 62600000 | 0.05 | 0.47 | A>A |
| chrX | 62600000 | 62700000 | -0.03 | 0.53 | B>A |
| chrX | 62700000 | 62800000 | 0.12 | 0.57 | A>A |
| chrX | 62800000 | 62900000 | 0.01 | 0.45 | A>A |
| chrX | 62900000 | 63000000 | 0.13 | 0.07 | A>A |
| chrX | 63000000 | 63100000 | -0.30 | -0.53 | B>B |
| chrX | 63100000 | 63200000 | -0.55 | -0.84 | B>B |
| chrX | 63200000 | 63300000 | -0.72 | -0.92 | B>B |
| chrX | 63300000 | 63400000 | -0.46 | -0.87 | B>B |
| chrX | 63400000 | 63500000 | -0.30 | -0.59 | B>B |
| chrX | 63500000 | 63600000 | -0.47 | -0.70 | B>B |
| chrX | 63600000 | 63700000 | -0.52 | -0.67 | B>B |
| chrX | 63700000 | 63800000 | -0.77 | -0.91 | B>B |
| chrX | 63800000 | 63900000 | -0.81 | -0.94 | B>B |
| chrX | 63900000 | 64000000 | -0.90 | -1.00 | B>B |
| chrX | 64000000 | 64100000 | -0.76 | -0.93 | B>B |
| chrX | 64100000 | 64200000 | -0.63 | -0.72 | B>B |
| chrX | 64200000 | 64300000 | -0.59 | -0.80 | B>B |
| chrX | 64300000 | 64400000 | 0.00 | -0.94 | NO |
| chrX | 64400000 | 64500000 | -0.76 | -0.92 | B>B |
| chrX | 64500000 | 64600000 | -0.92 | -0.87 | B>B |
| chrX | 64600000 | 64700000 | -0.37 | -0.54 | B>B |
| chrX | 64700000 | 64800000 | 0.07 | 0.02 | A>A |
| chrX | 64800000 | 64900000 | 0.49 | 0.47 | A>A |
| chrX | 64900000 | 65000000 | 0.55 | 0.54 | A>A |
| chrX | 65000000 | 65100000 | 0.07 | 0.21 | A>A |
| chrX | 65100000 | 65200000 | -0.05 | 0.28 | A>A |
| chrX | 65200000 | 65300000 | -0.28 | -0.25 | B>B |
| chrX | 65300000 | 65400000 | -0.52 | -0.55 | B>B |
| chrX | 65400000 | 65500000 | -0.56 | -0.73 | B>B |
| chrX | 65500000 | 65600000 | -0.67 | -0.80 | B>B |
| chrX | 65600000 | 65700000 | -0.64 | -0.82 | B>B |
| chrX | 65700000 | 65800000 | -0.45 | -0.62 | B>B |
| chrX | 65800000 | 65900000 | -0.10 | -0.26 | B>B |
| chrX | 65900000 | 66000000 | -0.34 | -0.31 | B>B |
| chrX | 66000000 | 66100000 | -0.56 | -0.62 | B>B |
| chrX | 66100000 | 66200000 | -0.68 | -0.67 | B>B |
| chrX | 66200000 | 66300000 | -0.65 | -0.60 | B>B |
| chrX | 66300000 | 66400000 | -0.65 | -0.57 | B>B |
| chrX | 66400000 | 66500000 | -0.45 | -0.51 | B>B |
| chrX | 66500000 | 66600000 | -0.42 | -0.36 | B>B |
| chrX | 66600000 | 66700000 | -0.36 | -0.09 | B>B |
| chrX | 66700000 | 66800000 | -0.03 | 0.30 | A>A |
| chrX | 66800000 | 66900000 | -0.01 | 0.47 | A>A |
| chrX | 66900000 | 67000000 | -0.06 | 0.36 | A>A |
| chrX | 67000000 | 67100000 | -0.22 | 0.11 | B>B |
| chrX | 67100000 | 67200000 | -0.35 | -0.01 | B>B |
| chrX | 67200000 | 67300000 | -0.08 | 0.12 | A>A |

| | | | | | |
|---|---|---|---|---|---|
| chrX | 67300000 | 67400000 | 0.15 | 0.20 | A>A |
| chrX | 67400000 | 67500000 | 0.67 | 0.66 | A>A |
| chrX | 67500000 | 67600000 | 0.87 | 0.88 | A>A |
| chrX | 67600000 | 67700000 | 0.80 | 0.85 | A>A |
| chrX | 67700000 | 67800000 | 0.75 | 0.60 | A>A |
| chrX | 67800000 | 67900000 | 0.73 | 0.61 | A>A |
| chrX | 67900000 | 68000000 | 0.43 | 0.45 | A>A |
| chrX | 68000000 | 68100000 | 0.45 | 0.52 | A>A |
| chrX | 68100000 | 68200000 | 0.61 | 0.87 | A>A |
| chrX | 68200000 | 68300000 | 0.59 | 0.82 | A>A |
| chrX | 68300000 | 68400000 | 0.65 | 0.71 | A>A |
| chrX | 68400000 | 68500000 | 0.55 | 0.58 | A>A |
| chrX | 68500000 | 68600000 | 0.37 | 0.35 | A>A |
| chrX | 68600000 | 68700000 | 0.13 | 0.13 | A>A |
| chrX | 68700000 | 68800000 | 0.16 | 0.10 | A>A |
| chrX | 68800000 | 68900000 | 0.04 | -0.09 | B>B |
| chrX | 68900000 | 69000000 | -0.37 | -0.32 | B>B |
| chrX | 69000000 | 69100000 | -0.32 | -0.38 | B>B |
| chrX | 69100000 | 69200000 | -0.17 | -0.19 | B>B |
| chrX | 69200000 | 69300000 | 0.20 | 0.19 | A>A |
| chrX | 69300000 | 69400000 | 0.64 | 0.51 | A>A |
| chrX | 69400000 | 69500000 | 0.90 | 0.87 | A>A |
| chrX | 69500000 | 69600000 | 1.01 | 0.93 | A>A |
| chrX | 69600000 | 69700000 | 0.82 | 0.94 | A>A |
| chrX | 69700000 | 69800000 | 0.82 | 1.04 | A>A |
| chrX | 69800000 | 69900000 | 0.54 | 0.64 | A>A |
| chrX | 69900000 | 70000000 | 0.65 | 0.69 | A>A |
| chrX | 70000000 | 70100000 | 0.55 | 0.69 | A>A |
| chrX | 70100000 | 70200000 | 0.67 | 0.72 | A>A |
| chrX | 70200000 | 70300000 | 1.04 | 0.95 | A>A |
| chrX | 70300000 | 70400000 | 1.10 | 1.03 | A>A |
| chrX | 70400000 | 70500000 | 1.20 | 1.11 | A>A |
| chrX | 70500000 | 70600000 | 1.17 | 1.07 | A>A |
| chrX | 70600000 | 70700000 | 1.27 | 1.07 | A>A |
| chrX | 70700000 | 70800000 | 1.36 | 1.17 | A>A |
| chrX | 70800000 | 70900000 | 1.07 | 0.96 | A>A |
| chrX | 70900000 | 71000000 | 0.00 | 0.00 | NO |
| chrX | 71000000 | 71100000 | 0.00 | 0.00 | NO |
| chrX | 71100000 | 71200000 | 0.90 | 0.82 | A>A |
| chrX | 71200000 | 71300000 | 0.97 | 1.01 | A>A |
| chrX | 71300000 | 71400000 | 1.18 | 1.01 | A>A |
| chrX | 71400000 | 71500000 | 1.05 | 0.85 | A>A |
| chrX | 71500000 | 71600000 | 0.92 | 0.80 | A>A |
| chrX | 71600000 | 71700000 | 0.72 | 0.98 | A>A |
| chrX | 71700000 | 71800000 | 0.84 | 0.87 | A>A |
| chrX | 71800000 | 71900000 | 0.95 | 0.88 | A>A |
| chrX | 71900000 | 72000000 | 0.00 | 0.00 | NO |
| chrX | 72000000 | 72100000 | 0.00 | 0.00 | NO |
| chrX | 72100000 | 72200000 | 0.00 | 0.00 | NO |
| chrX | 72200000 | 72300000 | 0.00 | 0.00 | NO |
| chrX | 72300000 | 72400000 | 0.00 | -0.27 | NO |
| chrX | 72400000 | 72500000 | -0.24 | -0.18 | B>B |
| chrX | 72500000 | 72600000 | 0.00 | -0.30 | NO |
| chrX | 72600000 | 72700000 | -0.28 | -0.17 | B>B |
| chrX | 72700000 | 72800000 | 0.08 | 0.21 | A>A |
| chrX | 72800000 | 72900000 | 0.13 | 0.31 | A>A |

| | | | | | |
|---|---|---|---|---|---|
| chrX | 72900000 | 73000000 | -0.15 | -0.01 | B>B |
| chrX | 73000000 | 73100000 | 0.17 | 0.07 | A>A |
| chrX | 73100000 | 73200000 | 0.69 | 0.61 | A>A |
| chrX | 73200000 | 73300000 | 0.31 | 0.48 | A>A |
| chrX | 73300000 | 73400000 | 0.23 | 0.45 | A>A |
| chrX | 73400000 | 73500000 | 0.88 | 0.82 | A>A |
| chrX | 73500000 | 73600000 | 0.72 | 0.52 | A>A |
| chrX | 73600000 | 73700000 | 0.53 | 0.45 | A>A |
| chrX | 73700000 | 73800000 | 0.38 | 0.27 | A>A |
| chrX | 73800000 | 73900000 | 0.12 | -0.11 | A>A |
| chrX | 73900000 | 74000000 | -0.25 | -0.65 | B>B |
| chrX | 74000000 | 74100000 | -0.38 | -0.71 | B>B |
| chrX | 74100000 | 74200000 | -0.32 | -0.62 | B>B |
| chrX | 74200000 | 74300000 | -0.10 | -0.33 | B>B |
| chrX | 74300000 | 74400000 | 0.24 | 0.03 | A>A |
| chrX | 74400000 | 74500000 | -0.04 | -0.25 | B>B |
| chrX | 74500000 | 74600000 | -0.20 | -0.18 | B>B |
| chrX | 74600000 | 74700000 | -0.44 | -0.64 | B>B |
| chrX | 74700000 | 74800000 | -0.59 | -0.70 | B>B |
| chrX | 74800000 | 74900000 | -0.55 | -0.63 | B>B |
| chrX | 74900000 | 75000000 | -0.57 | -0.61 | B>B |
| chrX | 75000000 | 75100000 | -0.68 | -0.80 | B>B |
| chrX | 75100000 | 75200000 | -0.82 | -0.92 | B>B |
| chrX | 75200000 | 75300000 | -0.61 | -0.85 | B>B |
| chrX | 75300000 | 75400000 | -0.22 | -0.24 | B>B |
| chrX | 75400000 | 75500000 | -0.13 | -0.19 | B>B |
| chrX | 75500000 | 75600000 | -0.32 | -0.54 | B>B |
| chrX | 75600000 | 75700000 | -0.27 | -0.46 | B>B |
| chrX | 75700000 | 75800000 | 0.00 | -0.62 | NO |
| chrX | 75800000 | 75900000 | -0.54 | -0.79 | B>B |
| chrX | 75900000 | 76000000 | -0.65 | -0.95 | B>B |
| chrX | 76000000 | 76100000 | -0.49 | -0.81 | B>B |
| chrX | 76100000 | 76200000 | -0.38 | -0.79 | B>B |
| chrX | 76200000 | 76300000 | -0.25 | -0.65 | B>B |
| chrX | 76300000 | 76400000 | -0.30 | -0.67 | B>B |
| chrX | 76400000 | 76500000 | -0.23 | -0.57 | B>B |
| chrX | 76500000 | 76600000 | 0.00 | -0.27 | NO |
| chrX | 76600000 | 76700000 | 0.00 | 0.00 | NO |
| chrX | 76700000 | 76800000 | 0.47 | 0.35 | A>A |
| chrX | 76800000 | 76900000 | 0.69 | 0.64 | A>A |
| chrX | 76900000 | 77000000 | 0.83 | 0.86 | A>A |
| chrX | 77000000 | 77100000 | 0.98 | 0.89 | A>A |
| chrX | 77100000 | 77200000 | 0.93 | 0.85 | A>A |
| chrX | 77200000 | 77300000 | 0.86 | 0.82 | A>A |
| chrX | 77300000 | 77400000 | 0.65 | 0.52 | A>A |
| chrX | 77400000 | 77500000 | 0.06 | -0.33 | B>B |
| chrX | 77500000 | 77600000 | -0.17 | -0.66 | B>B |
| chrX | 77600000 | 77700000 | -0.44 | -0.87 | B>B |
| chrX | 77700000 | 77800000 | -0.39 | -0.72 | B>B |
| chrX | 77800000 | 77900000 | -0.27 | -0.39 | B>B |
| chrX | 77900000 | 78000000 | -0.16 | -0.37 | B>B |
| chrX | 78000000 | 78100000 | -0.43 | -0.72 | B>B |
| chrX | 78100000 | 78200000 | -0.56 | -0.79 | B>B |
| chrX | 78200000 | 78300000 | -0.73 | -0.83 | B>B |
| chrX | 78300000 | 78400000 | -0.64 | -0.67 | B>B |
| chrX | 78400000 | 78500000 | -0.79 | -0.76 | B>B |

| | | | | | |
|---|---|---|---|---|---|
| chrX | 78500000 | 78600000 | -0.69 | -0.63 | B>B |
| chrX | 78600000 | 78700000 | -0.60 | -0.42 | B>B |
| chrX | 78700000 | 78800000 | -0.73 | -0.64 | B>B |
| chrX | 78800000 | 78900000 | -0.77 | -0.90 | B>B |
| chrX | 78900000 | 79000000 | -0.90 | -0.87 | B>B |
| chrX | 79000000 | 79100000 | -0.82 | -0.93 | B>B |
| chrX | 79100000 | 79200000 | -0.89 | -0.84 | B>B |
| chrX | 79200000 | 79300000 | -0.68 | -0.64 | B>B |
| chrX | 79300000 | 79400000 | -0.62 | -0.41 | B>B |
| chrX | 79400000 | 79500000 | -0.25 | -0.17 | B>B |
| chrX | 79500000 | 79600000 | -0.01 | -0.11 | B>B |
| chrX | 79600000 | 79700000 | -0.14 | -0.40 | B>B |
| chrX | 79700000 | 79800000 | -0.20 | -0.37 | B>B |
| chrX | 79800000 | 79900000 | -0.20 | -0.36 | B>B |
| chrX | 79900000 | 80000000 | 0.42 | 0.14 | A>A |
| chrX | 80000000 | 80100000 | 0.39 | 0.20 | A>A |
| chrX | 80100000 | 80200000 | -0.40 | -0.37 | B>B |
| chrX | 80200000 | 80300000 | -0.58 | -0.61 | B>B |
| chrX | 80300000 | 80400000 | -0.34 | -0.36 | B>B |
| chrX | 80400000 | 80500000 | 0.02 | 0.07 | A>A |
| chrX | 80500000 | 80600000 | -0.01 | 0.16 | A>A |
| chrX | 80600000 | 80700000 | -0.39 | -0.22 | B>B |
| chrX | 80700000 | 80800000 | -0.69 | -0.28 | B>B |
| chrX | 80800000 | 80900000 | -0.80 | -0.34 | B>B |
| chrX | 80900000 | 81000000 | -0.74 | -0.29 | B>B |
| chrX | 81000000 | 81100000 | -0.75 | -0.04 | B>A |
| chrX | 81100000 | 81200000 | -0.70 | 0.11 | B>A |
| chrX | 81200000 | 81300000 | -0.68 | -0.10 | B>A |
| chrX | 81300000 | 81400000 | -0.65 | -0.08 | B>A |
| chrX | 81400000 | 81500000 | -0.63 | -0.03 | B>A |
| chrX | 81500000 | 81600000 | -0.47 | 0.21 | B>A |
| chrX | 81600000 | 81700000 | -0.30 | 0.36 | B>A |
| chrX | 81700000 | 81800000 | -0.22 | 0.40 | B>A |
| chrX | 81800000 | 81900000 | -0.14 | 0.43 | B>A |
| chrX | 81900000 | 82000000 | -0.14 | 0.63 | B>A |
| chrX | 82000000 | 82100000 | -0.10 | 0.72 | B>A |
| chrX | 82100000 | 82200000 | -0.20 | 0.65 | B>A |
| chrX | 82200000 | 82300000 | -0.21 | 0.39 | B>A |
| chrX | 82300000 | 82400000 | -0.41 | 0.40 | B>A |
| chrX | 82400000 | 82500000 | -0.43 | 0.37 | B>A |
| chrX | 82500000 | 82600000 | -0.64 | 0.21 | B>A |
| chrX | 82600000 | 82700000 | -0.63 | 0.13 | B>A |
| chrX | 82700000 | 82800000 | -0.66 | -0.12 | B>A |
| chrX | 82800000 | 82900000 | -0.65 | -0.32 | B>B |
| chrX | 82900000 | 83000000 | -0.78 | -0.58 | B>B |
| chrX | 83000000 | 83100000 | -0.79 | -0.75 | B>B |
| chrX | 83100000 | 83200000 | -0.65 | -0.77 | B>B |
| chrX | 83200000 | 83300000 | -0.53 | -0.67 | B>B |
| chrX | 83300000 | 83400000 | 0.03 | -0.03 | A>A |
| chrX | 83400000 | 83500000 | -0.10 | -0.07 | B>B |
| chrX | 83500000 | 83600000 | -0.37 | -0.36 | B>B |
| chrX | 83600000 | 83700000 | -0.46 | -0.47 | B>B |
| chrX | 83700000 | 83800000 | -0.19 | -0.47 | B>B |
| chrX | 83800000 | 83900000 | -0.54 | -0.70 | B>B |
| chrX | 83900000 | 84000000 | -0.55 | -0.76 | B>B |
| chrX | 84000000 | 84100000 | -0.56 | -0.84 | B>B |

| | | | | | |
|---|---|---|---|---|---|
| chrX | 84100000 | 84200000 | -0.41 | -0.68 | B>B |
| chrX | 84200000 | 84300000 | 0.03 | -0.29 | B>B |
| chrX | 84300000 | 84400000 | -0.09 | -0.25 | B>B |
| chrX | 84400000 | 84500000 | -0.09 | -0.33 | B>B |
| chrX | 84500000 | 84600000 | 0.17 | 0.14 | A>A |
| chrX | 84600000 | 84700000 | 0.25 | -0.04 | A>A |
| chrX | 84700000 | 84800000 | -0.24 | -0.45 | B>B |
| chrX | 84800000 | 84900000 | -0.59 | -0.68 | B>B |
| chrX | 84900000 | 85000000 | -0.68 | -0.81 | B>B |
| chrX | 85000000 | 85100000 | -0.50 | -0.75 | B>B |
| chrX | 85100000 | 85200000 | 0.17 | -0.02 | A>A |
| chrX | 85200000 | 85300000 | 0.46 | 0.34 | A>A |
| chrX | 85300000 | 85400000 | -0.20 | -0.48 | B>B |
| chrX | 85400000 | 85500000 | -0.59 | -0.82 | B>B |
| chrX | 85500000 | 85600000 | -0.78 | -0.91 | B>B |
| chrX | 85600000 | 85700000 | -0.84 | -0.98 | B>B |
| chrX | 85700000 | 85800000 | -0.88 | -0.94 | B>B |
| chrX | 85800000 | 85900000 | -0.83 | -0.96 | B>B |
| chrX | 85900000 | 86000000 | -0.88 | -1.02 | B>B |
| chrX | 86000000 | 86100000 | -0.80 | -1.01 | B>B |
| chrX | 86100000 | 86200000 | -0.79 | -0.95 | B>B |
| chrX | 86200000 | 86300000 | -0.79 | -0.95 | B>B |
| chrX | 86300000 | 86400000 | -0.88 | -0.94 | B>B |
| chrX | 86400000 | 86500000 | -0.84 | -0.91 | B>B |
| chrX | 86500000 | 86600000 | -0.60 | -0.52 | B>B |
| chrX | 86600000 | 86700000 | -0.29 | 0.15 | B>B |
| chrX | 86700000 | 86800000 | 0.03 | 0.37 | A>A |
| chrX | 86800000 | 86900000 | -0.06 | 0.21 | A>A |
| chrX | 86900000 | 87000000 | -0.53 | -0.48 | B>B |
| chrX | 87000000 | 87100000 | -0.78 | -0.89 | B>B |
| chrX | 87100000 | 87200000 | -0.94 | -0.94 | B>B |
| chrX | 87200000 | 87300000 | -0.87 | -1.03 | B>B |
| chrX | 87300000 | 87400000 | -0.88 | -1.00 | B>B |
| chrX | 87400000 | 87500000 | -0.90 | -1.04 | B>B |
| chrX | 87500000 | 87600000 | -0.86 | -0.99 | B>B |
| chrX | 87600000 | 87700000 | -0.94 | -1.03 | B>B |
| chrX | 87700000 | 87800000 | -0.93 | -1.04 | B>B |
| chrX | 87800000 | 87900000 | -0.90 | -0.99 | B>B |
| chrX | 87900000 | 88000000 | -1.02 | -0.97 | B>B |
| chrX | 88000000 | 88100000 | -1.02 | -1.05 | B>B |
| chrX | 88100000 | 88200000 | -1.01 | -0.99 | B>B |
| chrX | 88200000 | 88300000 | -1.06 | -0.97 | B>B |
| chrX | 88300000 | 88400000 | -1.03 | -1.05 | B>B |
| chrX | 88400000 | 88500000 | 0.00 | -0.90 | NO |
| chrX | 88500000 | 88600000 | 0.00 | 0.00 | NO |
| chrX | 88600000 | 88700000 | 0.00 | 0.00 | NO |
| chrX | 88700000 | 88800000 | 0.00 | 0.00 | NO |
| chrX | 88800000 | 88900000 | 0.00 | 0.00 | NO |
| chrX | 88900000 | 89000000 | 0.00 | 0.00 | NO |
| chrX | 89000000 | 89100000 | 0.00 | 0.00 | NO |
| chrX | 89100000 | 89200000 | 0.00 | 0.00 | NO |
| chrX | 89200000 | 89300000 | 0.00 | 0.00 | NO |
| chrX | 89300000 | 89400000 | 0.00 | 0.00 | NO |
| chrX | 89400000 | 89500000 | 0.00 | 0.00 | NO |
| chrX | 89500000 | 89600000 | 0.00 | 0.00 | NO |
| chrX | 89600000 | 89700000 | 0.00 | 0.00 | NO |

| | | | | | |
|---|---|---|---|---|---|
| chrX | 89700000 | 89800000 | 0.00 | 0.00 | NO |
| chrX | 89800000 | 89900000 | 0.00 | 0.00 | NO |
| chrX | 89900000 | 90000000 | 0.00 | 0.00 | NO |
| chrX | 90000000 | 90100000 | 0.00 | 0.00 | NO |
| chrX | 90100000 | 90200000 | 0.00 | 0.00 | NO |
| chrX | 90200000 | 90300000 | 0.00 | 0.00 | NO |
| chrX | 90300000 | 90400000 | -1.16 | -1.18 | B>B |
| chrX | 90400000 | 90500000 | 0.00 | 0.00 | NO |
| chrX | 90500000 | 90600000 | -1.24 | -1.19 | B>B |
| chrX | 90600000 | 90700000 | -1.18 | -1.18 | B>B |
| chrX | 90700000 | 90800000 | 0.00 | 0.00 | NO |
| chrX | 90800000 | 90900000 | 0.00 | 0.00 | NO |
| chrX | 90900000 | 91000000 | 0.00 | 0.00 | NO |
| chrX | 91000000 | 91100000 | 0.00 | 0.00 | NO |
| chrX | 91100000 | 91200000 | 0.00 | 0.00 | NO |
| chrX | 91200000 | 91300000 | 0.00 | 0.00 | NO |
| chrX | 91300000 | 91400000 | 0.00 | 0.00 | NO |
| chrX | 91400000 | 91500000 | 0.00 | -0.58 | NO |
| chrX | 91500000 | 91600000 | 0.00 | 0.00 | NO |
| chrX | 91600000 | 91700000 | 0.00 | 0.00 | NO |
| chrX | 91700000 | 91800000 | 0.00 | 0.00 | NO |
| chrX | 91800000 | 91900000 | 0.00 | 0.00 | NO |
| chrX | 91900000 | 92000000 | 0.00 | 0.00 | NO |
| chrX | 92000000 | 92100000 | 0.00 | 0.00 | NO |
| chrX | 92100000 | 92200000 | 0.00 | 0.00 | NO |
| chrX | 92200000 | 92300000 | 0.00 | 0.00 | NO |
| chrX | 92300000 | 92400000 | 0.00 | 0.00 | NO |
| chrX | 92400000 | 92500000 | -1.03 | -1.09 | B>B |
| chrX | 92500000 | 92600000 | -1.04 | -1.11 | B>B |
| chrX | 92600000 | 92700000 | -1.03 | -1.03 | B>B |
| chrX | 92700000 | 92800000 | -0.93 | -1.03 | B>B |
| chrX | 92800000 | 92900000 | -0.79 | -1.00 | B>B |
| chrX | 92900000 | 93000000 | -0.60 | -0.89 | B>B |
| chrX | 93000000 | 93100000 | -0.82 | -0.96 | B>B |
| chrX | 93100000 | 93200000 | -0.84 | -0.85 | B>B |
| chrX | 93200000 | 93300000 | -0.77 | -0.90 | B>B |
| chrX | 93300000 | 93400000 | -0.89 | -0.81 | B>B |
| chrX | 93400000 | 93500000 | -0.93 | -0.82 | B>B |
| chrX | 93500000 | 93600000 | -0.95 | -0.62 | B>B |
| chrX | 93600000 | 93700000 | -0.89 | -0.34 | B>A |
| chrX | 93700000 | 93800000 | -0.93 | -0.23 | B>A |
| chrX | 93800000 | 93900000 | -0.94 | -0.30 | B>A |
| chrX | 93900000 | 94000000 | -0.87 | -0.70 | B>B |
| chrX | 94000000 | 94100000 | -0.86 | -0.83 | B>B |
| chrX | 94100000 | 94200000 | -0.82 | -0.96 | B>B |
| chrX | 94200000 | 94300000 | -0.90 | -0.99 | B>B |
| chrX | 94300000 | 94400000 | -0.91 | -0.96 | B>B |
| chrX | 94400000 | 94500000 | -0.83 | -0.97 | B>B |
| chrX | 94500000 | 94600000 | -0.84 | -0.89 | B>B |
| chrX | 94600000 | 94700000 | -0.85 | -0.91 | B>B |
| chrX | 94700000 | 94800000 | -0.92 | -0.96 | B>B |
| chrX | 94800000 | 94900000 | -0.90 | -0.94 | B>B |
| chrX | 94900000 | 95000000 | -0.80 | -0.93 | B>B |
| chrX | 95000000 | 95100000 | -0.72 | -0.88 | B>B |
| chrX | 95100000 | 95200000 | -0.71 | -0.88 | B>B |
| chrX | 95200000 | 95300000 | -0.75 | -0.80 | B>B |

| | | | | | |
|---|---|---|---|---|---|
| chrX | 95300000 | 95400000 | -0.64 | -0.77 | B>B |
| chrX | 95400000 | 95500000 | -0.53 | -0.64 | B>B |
| chrX | 95500000 | 95600000 | -0.28 | -0.32 | B>B |
| chrX | 95600000 | 95700000 | 0.24 | 0.34 | A>A |
| chrX | 95700000 | 95800000 | 0.40 | 0.52 | A>A |
| chrX | 95800000 | 95900000 | 0.31 | 0.32 | A>A |
| chrX | 95900000 | 96000000 | 0.49 | 0.37 | A>A |
| chrX | 96000000 | 96100000 | 0.43 | 0.26 | A>A |
| chrX | 96100000 | 96200000 | 0.49 | 0.41 | A>A |
| chrX | 96200000 | 96300000 | 0.39 | 0.32 | A>A |
| chrX | 96300000 | 96400000 | 0.59 | 0.58 | A>A |
| chrX | 96400000 | 96500000 | 0.64 | 0.64 | A>A |
| chrX | 96500000 | 96600000 | 0.55 | 0.55 | A>A |
| chrX | 96600000 | 96700000 | 0.59 | 0.53 | A>A |
| chrX | 96700000 | 96800000 | 0.45 | 0.44 | A>A |
| chrX | 96800000 | 96900000 | 0.24 | 0.37 | A>A |
| chrX | 96900000 | 97000000 | -0.01 | -0.03 | B>B |
| chrX | 97000000 | 97100000 | -0.23 | -0.45 | B>B |
| chrX | 97100000 | 97200000 | -0.46 | -0.68 | B>B |
| chrX | 97200000 | 97300000 | -0.64 | -0.79 | B>B |
| chrX | 97300000 | 97400000 | -0.67 | -0.77 | B>B |
| chrX | 97400000 | 97500000 | -0.74 | -0.87 | B>B |
| chrX | 97500000 | 97600000 | -0.72 | -0.78 | B>B |
| chrX | 97600000 | 97700000 | -0.78 | -0.80 | B>B |
| chrX | 97700000 | 97800000 | -0.72 | -0.82 | B>B |
| chrX | 97800000 | 97900000 | -0.83 | -0.84 | B>B |
| chrX | 97900000 | 98000000 | -0.80 | -0.85 | B>B |
| chrX | 98000000 | 98100000 | -0.83 | -0.88 | B>B |
| chrX | 98100000 | 98200000 | -0.87 | -0.83 | B>B |
| chrX | 98200000 | 98300000 | -0.93 | -0.92 | B>B |
| chrX | 98300000 | 98400000 | -0.88 | -0.91 | B>B |
| chrX | 98400000 | 98500000 | -0.87 | -0.88 | B>B |
| chrX | 98500000 | 98600000 | -0.88 | -0.84 | B>B |
| chrX | 98600000 | 98700000 | -0.90 | -0.93 | B>B |
| chrX | 98700000 | 98800000 | -0.96 | -0.89 | B>B |
| chrX | 98800000 | 98900000 | -0.87 | -0.97 | B>B |
| chrX | 98900000 | 99000000 | -0.86 | -1.01 | B>B |
| chrX | 99000000 | 99100000 | -0.78 | -0.97 | B>B |
| chrX | 99100000 | 99200000 | -0.85 | -0.97 | B>B |
| chrX | 99200000 | 99300000 | -0.76 | -0.92 | B>B |
| chrX | 99300000 | 99400000 | -0.80 | -0.98 | B>B |
| chrX | 99400000 | 99500000 | -0.68 | -0.85 | B>B |
| chrX | 99500000 | 99600000 | -0.49 | -0.84 | B>B |
| chrX | 99600000 | 99700000 | -0.19 | -0.47 | B>B |
| chrX | 99700000 | 99800000 | 0.10 | -0.17 | B>B |
| chrX | 99800000 | 99900000 | 0.55 | 0.27 | A>A |
| chrX | 99900000 | 100000000 | 0.88 | 0.79 | A>A |
| chrX | 1E+08 | 100100000 | 0.96 | 0.83 | A>A |
| chrX | 1E+08 | 100200000 | 1.14 | 1.09 | A>A |
| chrX | 1E+08 | 100300000 | 1.35 | 1.20 | A>A |
| chrX | 1E+08 | 100400000 | 1.31 | 1.16 | A>A |
| chrX | 1E+08 | 100500000 | 1.08 | 0.89 | A>A |
| chrX | 1.01E+08 | 100600000 | 0.99 | 0.80 | A>A |
| chrX | 1.01E+08 | 100700000 | 0.96 | 0.76 | A>A |
| chrX | 1.01E+08 | 100800000 | 0.47 | 0.07 | A>A |
| chrX | 1.01E+08 | 100900000 | 0.36 | 0.17 | A>A |

| | | | | | |
|---|---|---|---|---|---|
| chrX | 1.01E+08 | 101000000 | 0.18 | 0.01 | A>A |
| chrX | 1.01E+08 | 101100000 | 0.10 | -0.06 | A>A |
| chrX | 1.01E+08 | 101200000 | -0.15 | -0.08 | B>B |
| chrX | 1.01E+08 | 101300000 | -0.34 | -0.28 | B>B |
| chrX | 1.01E+08 | 101400000 | -0.18 | -0.30 | B>B |
| chrX | 1.01E+08 | 101500000 | 0.00 | 0.00 | NO |
| chrX | 1.02E+08 | 101600000 | 0.00 | 0.00 | NO |
| chrX | 1.02E+08 | 101700000 | 0.00 | 0.00 | NO |
| chrX | 1.02E+08 | 101800000 | 0.00 | 0.00 | NO |
| chrX | 1.02E+08 | 101900000 | 0.64 | 0.47 | A>A |
| chrX | 1.02E+08 | 102000000 | 0.85 | 0.86 | A>A |
| chrX | 1.02E+08 | 102100000 | 0.80 | 0.78 | A>A |
| chrX | 1.02E+08 | 102200000 | 0.51 | 0.44 | A>A |
| chrX | 1.02E+08 | 102300000 | 0.34 | 0.21 | A>A |
| chrX | 1.02E+08 | 102400000 | 0.46 | 0.16 | A>A |
| chrX | 1.02E+08 | 102500000 | 0.60 | 0.44 | A>A |
| chrX | 1.03E+08 | 102600000 | 0.93 | 0.71 | A>A |
| chrX | 1.03E+08 | 102700000 | 1.31 | 0.97 | A>A |
| chrX | 1.03E+08 | 102800000 | 0.71 | 0.57 | A>A |
| chrX | 1.03E+08 | 102900000 | 0.97 | 0.89 | A>A |
| chrX | 1.03E+08 | 103000000 | 1.04 | 0.70 | A>A |
| chrX | 1.03E+08 | 103100000 | 0.70 | 0.41 | A>A |
| chrX | 1.03E+08 | 103200000 | 0.46 | 0.35 | A>A |
| chrX | 1.03E+08 | 103300000 | 0.53 | 0.26 | A>A |
| chrX | 1.03E+08 | 103400000 | 0.59 | 0.46 | A>A |
| chrX | 1.03E+08 | 103500000 | 0.80 | 0.53 | A>A |
| chrX | 1.04E+08 | 103600000 | 0.44 | 0.26 | A>A |
| chrX | 1.04E+08 | 103700000 | 0.24 | 0.22 | A>A |
| chrX | 1.04E+08 | 103800000 | -0.04 | -0.23 | B>B |
| chrX | 1.04E+08 | 103900000 | -0.27 | -0.57 | B>B |
| chrX | 1.04E+08 | 104000000 | -0.40 | -0.91 | B>B |
| chrX | 1.04E+08 | 104100000 | -0.56 | -1.00 | B>B |
| chrX | 1.04E+08 | 104200000 | -0.48 | -0.81 | B>B |
| chrX | 1.04E+08 | 104300000 | -0.69 | -1.03 | B>B |
| chrX | 1.04E+08 | 104400000 | -0.72 | -1.04 | B>B |
| chrX | 1.04E+08 | 104500000 | -0.63 | -0.96 | B>B |
| chrX | 1.05E+08 | 104600000 | -0.79 | -1.02 | B>B |
| chrX | 1.05E+08 | 104700000 | -0.81 | -1.10 | B>B |
| chrX | 1.05E+08 | 104800000 | -0.78 | -1.13 | B>B |
| chrX | 1.05E+08 | 104900000 | -0.75 | -1.07 | B>B |
| chrX | 1.05E+08 | 105000000 | -0.70 | -1.15 | B>B |
| chrX | 1.05E+08 | 105100000 | -0.63 | -1.06 | B>B |
| chrX | 1.05E+08 | 105200000 | -0.58 | -1.01 | B>B |
| chrX | 1.05E+08 | 105300000 | -0.58 | -1.03 | B>B |
| chrX | 1.05E+08 | 105400000 | -0.55 | -0.93 | B>B |
| chrX | 1.05E+08 | 105500000 | -0.38 | -0.80 | B>B |
| chrX | 1.06E+08 | 105600000 | -0.33 | -0.79 | B>B |
| chrX | 1.06E+08 | 105700000 | -0.10 | -0.43 | B>B |
| chrX | 1.06E+08 | 105800000 | 0.00 | -0.26 | B>B |
| chrX | 1.06E+08 | 105900000 | 0.05 | -0.14 | B>B |
| chrX | 1.06E+08 | 106000000 | 0.24 | 0.02 | A>A |
| chrX | 1.06E+08 | 106100000 | 0.52 | 0.32 | A>A |
| chrX | 1.06E+08 | 106200000 | 1.02 | 0.86 | A>A |
| chrX | 1.06E+08 | 106300000 | 0.91 | 0.79 | A>A |
| chrX | 1.06E+08 | 106400000 | 1.07 | 1.08 | A>A |
| chrX | 1.06E+08 | 106500000 | 1.09 | 0.99 | A>A |

| | | | | | |
|---|---|---|---|---|---|
| chrX | 1.07E+08 | 106600000 | 0.74 | 0.66 | A>A |
| chrX | 1.07E+08 | 106700000 | 0.86 | 0.70 | A>A |
| chrX | 1.07E+08 | 106800000 | 0.81 | 0.69 | A>A |
| chrX | 1.07E+08 | 106900000 | 1.06 | 0.88 | A>A |
| chrX | 1.07E+08 | 107000000 | 1.16 | 1.17 | A>A |
| chrX | 1.07E+08 | 107100000 | 0.90 | 0.99 | A>A |
| chrX | 1.07E+08 | 107200000 | 0.92 | 0.78 | A>A |
| chrX | 1.07E+08 | 107300000 | 1.03 | 0.83 | A>A |
| chrX | 1.07E+08 | 107400000 | 0.82 | 0.78 | A>A |
| chrX | 1.07E+08 | 107500000 | 0.71 | 0.61 | A>A |
| chrX | 1.08E+08 | 107600000 | 0.89 | 0.85 | A>A |
| chrX | 1.08E+08 | 107700000 | 1.13 | 1.08 | A>A |
| chrX | 1.08E+08 | 107800000 | 0.94 | 0.88 | A>A |
| chrX | 1.08E+08 | 107900000 | 0.78 | 0.59 | A>A |
| chrX | 1.08E+08 | 108000000 | 0.35 | 0.32 | A>A |
| chrX | 1.08E+08 | 108100000 | 0.05 | -0.33 | B>B |
| chrX | 1.08E+08 | 108200000 | -0.12 | -0.57 | B>B |
| chrX | 1.08E+08 | 108300000 | -0.29 | -0.66 | B>B |
| chrX | 1.08E+08 | 108400000 | -0.25 | -0.60 | B>B |
| chrX | 1.08E+08 | 108500000 | -0.24 | -0.57 | B>B |
| chrX | 1.09E+08 | 108600000 | -0.12 | -0.48 | B>B |
| chrX | 1.09E+08 | 108700000 | 0.07 | -0.21 | B>B |
| chrX | 1.09E+08 | 108800000 | 0.33 | 0.19 | A>A |
| chrX | 1.09E+08 | 108900000 | 0.50 | 0.26 | A>A |
| chrX | 1.09E+08 | 109000000 | 0.77 | 0.74 | A>A |
| chrX | 1.09E+08 | 109100000 | 0.98 | 0.93 | A>A |
| chrX | 1.09E+08 | 109200000 | 0.74 | 0.56 | A>A |
| chrX | 1.09E+08 | 109300000 | 1.13 | 0.95 | A>A |
| chrX | 1.09E+08 | 109400000 | 0.88 | 0.78 | A>A |
| chrX | 1.09E+08 | 109500000 | 0.85 | 0.84 | A>A |
| chrX | 1.1E+08 | 109600000 | 0.95 | 0.98 | A>A |
| chrX | 1.1E+08 | 109700000 | 0.46 | 0.28 | A>A |
| chrX | 1.1E+08 | 109800000 | -0.05 | -0.40 | B>B |
| chrX | 1.1E+08 | 109900000 | -0.08 | -0.54 | B>B |
| chrX | 1.1E+08 | 110000000 | -0.08 | -0.55 | B>B |
| chrX | 1.1E+08 | 110100000 | -0.04 | -0.40 | B>B |
| chrX | 1.1E+08 | 110200000 | -0.11 | -0.55 | B>B |
| chrX | 1.1E+08 | 110300000 | -0.07 | -0.58 | B>B |
| chrX | 1.1E+08 | 110400000 | -0.08 | -0.51 | B>B |
| chrX | 1.1E+08 | 110500000 | -0.21 | -0.66 | B>B |
| chrX | 1.11E+08 | 110600000 | -0.15 | -0.73 | B>B |
| chrX | 1.11E+08 | 110700000 | -0.19 | -0.76 | B>B |
| chrX | 1.11E+08 | 110800000 | -0.13 | -0.68 | B>B |
| chrX | 1.11E+08 | 110900000 | 0.15 | -0.49 | A>B |
| chrX | 1.11E+08 | 111000000 | 0.48 | 0.11 | A>A |
| chrX | 1.11E+08 | 111100000 | 0.10 | -0.40 | B>B |
| chrX | 1.11E+08 | 111200000 | -0.02 | -0.69 | A>B |
| chrX | 1.11E+08 | 111300000 | -0.20 | -0.85 | A>B |
| chrX | 1.11E+08 | 111400000 | -0.35 | -0.83 | B>B |
| chrX | 1.11E+08 | 111500000 | -0.50 | -0.91 | B>B |
| chrX | 1.12E+08 | 111600000 | -0.57 | -0.98 | B>B |
| chrX | 1.12E+08 | 111700000 | -0.61 | -0.92 | B>B |
| chrX | 1.12E+08 | 111800000 | -0.62 | -0.83 | B>B |
| chrX | 1.12E+08 | 111900000 | -0.38 | -0.75 | B>B |
| chrX | 1.12E+08 | 112000000 | -0.30 | -0.69 | B>B |
| chrX | 1.12E+08 | 112100000 | -0.16 | -0.60 | B>B |

| | | | | | |
|---|---|---|---|---|---|
| chrX | 1.12E+08 | 112200000 | -0.33 | -0.73 | B>B |
| chrX | 1.12E+08 | 112300000 | -0.55 | -0.86 | B>B |
| chrX | 1.12E+08 | 112400000 | -0.58 | -0.90 | B>B |
| chrX | 1.12E+08 | 112500000 | -0.62 | -0.91 | B>B |
| chrX | 1.13E+08 | 112600000 | -0.62 | -0.94 | B>B |
| chrX | 1.13E+08 | 112700000 | -0.67 | -0.95 | B>B |
| chrX | 1.13E+08 | 112800000 | -0.55 | -0.89 | B>B |
| chrX | 1.13E+08 | 112900000 | -0.57 | -0.95 | B>B |
| chrX | 1.13E+08 | 113000000 | -0.57 | -0.83 | B>B |
| chrX | 1.13E+08 | 113100000 | -0.54 | -0.74 | B>B |
| chrX | 1.13E+08 | 113200000 | -0.44 | -0.56 | B>B |
| chrX | 1.13E+08 | 113300000 | -0.38 | -0.53 | B>B |
| chrX | 1.13E+08 | 113400000 | -0.35 | -0.46 | B>B |
| chrX | 1.13E+08 | 113500000 | -0.41 | -0.49 | B>B |
| chrX | 1.14E+08 | 113600000 | 0.00 | -0.30 | NO |
| chrX | 1.14E+08 | 113700000 | -0.41 | -0.51 | B>B |
| chrX | 1.14E+08 | 113800000 | -0.48 | -0.50 | B>B |
| chrX | 1.14E+08 | 113900000 | -0.34 | -0.41 | B>B |
| chrX | 1.14E+08 | 114000000 | -0.41 | -0.47 | B>B |
| chrX | 1.14E+08 | 114100000 | -0.36 | -0.47 | B>B |
| chrX | 1.14E+08 | 114200000 | -0.29 | -0.43 | B>B |
| chrX | 1.14E+08 | 114300000 | -0.14 | -0.29 | B>B |
| chrX | 1.14E+08 | 114400000 | -0.08 | -0.12 | B>B |
| chrX | 1.14E+08 | 114500000 | 0.25 | 0.38 | A>A |
| chrX | 1.15E+08 | 114600000 | 0.67 | 0.64 | A>A |
| chrX | 1.15E+08 | 114700000 | 1.15 | 1.24 | A>A |
| chrX | 1.15E+08 | 114800000 | 1.24 | 1.17 | A>A |
| chrX | 1.15E+08 | 114900000 | 0.85 | 1.03 | A>A |
| chrX | 1.15E+08 | 115000000 | 0.00 | 0.00 | NO |
| chrX | 1.15E+08 | 115100000 | -0.34 | 0.70 | B>A |
| chrX | 1.15E+08 | 115200000 | -0.54 | 0.67 | B>A |
| chrX | 1.15E+08 | 115300000 | -0.77 | 0.65 | B>A |
| chrX | 1.15E+08 | 115400000 | -0.84 | 0.91 | B>A |
| chrX | 1.15E+08 | 115500000 | -0.94 | 1.19 | B>A |
| chrX | 1.16E+08 | 115600000 | -0.92 | 1.16 | B>A |
| chrX | 1.16E+08 | 115700000 | -0.79 | 0.67 | B>A |
| chrX | 1.16E+08 | 115800000 | -0.88 | 0.72 | B>A |
| chrX | 1.16E+08 | 115900000 | -0.90 | 0.77 | B>A |
| chrX | 1.16E+08 | 116000000 | -0.87 | 0.84 | B>A |
| chrX | 1.16E+08 | 116100000 | -0.92 | 0.70 | B>A |
| chrX | 1.16E+08 | 116200000 | -0.99 | 0.42 | B>A |
| chrX | 1.16E+08 | 116300000 | -0.93 | 0.39 | B>A |
| chrX | 1.16E+08 | 116400000 | -0.96 | 0.28 | B>A |
| chrX | 1.16E+08 | 116500000 | -0.84 | 0.10 | B>A |
| chrX | 1.17E+08 | 116600000 | -0.89 | 0.15 | B>A |
| chrX | 1.17E+08 | 116700000 | -0.94 | 0.19 | B>A |
| chrX | 1.17E+08 | 116800000 | -0.86 | 0.14 | B>A |
| chrX | 1.17E+08 | 116900000 | -0.81 | 0.22 | B>A |
| chrX | 1.17E+08 | 117000000 | -0.70 | 0.21 | B>A |
| chrX | 1.17E+08 | 117100000 | -0.62 | 0.56 | B>A |
| chrX | 1.17E+08 | 117200000 | -0.43 | 0.68 | B>A |
| chrX | 1.17E+08 | 117300000 | -0.11 | 0.84 | B>A |
| chrX | 1.17E+08 | 117400000 | -0.05 | 0.80 | B>A |
| chrX | 1.17E+08 | 117500000 | 0.63 | 1.07 | A>A |
| chrX | 1.18E+08 | 117600000 | 0.95 | 1.13 | A>A |
| chrX | 1.18E+08 | 117700000 | 1.19 | 1.15 | A>A |

| | | | | | |
|---|---|---|---|---|---|
| chrX | 1.18E+08 | 117800000 | 1.11 | 1.04 | A>A |
| chrX | 1.18E+08 | 117900000 | 1.45 | 1.33 | A>A |
| chrX | 1.18E+08 | 118000000 | 1.04 | 0.93 | A>A |
| chrX | 1.18E+08 | 118100000 | 1.21 | 1.07 | A>A |
| chrX | 1.18E+08 | 118200000 | 1.10 | 1.04 | A>A |
| chrX | 1.18E+08 | 118300000 | 1.22 | 0.99 | A>A |
| chrX | 1.18E+08 | 118400000 | 1.26 | 1.05 | A>A |
| chrX | 1.18E+08 | 118500000 | 1.28 | 1.06 | A>A |
| chrX | 1.19E+08 | 118600000 | 1.45 | 1.13 | A>A |
| chrX | 1.19E+08 | 118700000 | 1.33 | 1.13 | A>A |
| chrX | 1.19E+08 | 118800000 | 1.45 | 1.31 | A>A |
| chrX | 1.19E+08 | 118900000 | 1.46 | 1.07 | A>A |
| chrX | 1.19E+08 | 119000000 | 1.56 | 1.15 | A>A |
| chrX | 1.19E+08 | 119100000 | 1.40 | 1.11 | A>A |
| chrX | 1.19E+08 | 119200000 | 1.15 | 0.99 | A>A |
| chrX | 1.19E+08 | 119300000 | 0.00 | 0.92 | NO |
| chrX | 1.19E+08 | 119400000 | 1.11 | 0.95 | A>A |
| chrX | 1.19E+08 | 119500000 | 1.04 | 0.82 | A>A |
| chrX | 1.2E+08 | 119600000 | 0.86 | 0.86 | A>A |
| chrX | 1.2E+08 | 119700000 | 0.90 | 1.02 | A>A |
| chrX | 1.2E+08 | 119800000 | 1.19 | 1.14 | A>A |
| chrX | 1.2E+08 | 119900000 | 0.61 | 0.81 | A>A |
| chrX | 1.2E+08 | 120000000 | 0.09 | 0.45 | A>A |
| chrX | 1.2E+08 | 120100000 | 0.00 | 0.00 | NO |
| chrX | 1.2E+08 | 120200000 | -0.02 | 0.48 | A>A |
| chrX | 1.2E+08 | 120300000 | -0.16 | 0.63 | B>A |
| chrX | 1.2E+08 | 120400000 | -0.23 | 0.55 | B>A |
| chrX | 1.2E+08 | 120500000 | -0.39 | 0.49 | B>A |
| chrX | 1.21E+08 | 120600000 | -0.57 | 0.54 | B>A |
| chrX | 1.21E+08 | 120700000 | -0.69 | 0.57 | B>A |
| chrX | 1.21E+08 | 120800000 | -0.71 | 0.58 | B>A |
| chrX | 1.21E+08 | 120900000 | -0.80 | 0.67 | B>A |
| chrX | 1.21E+08 | 121000000 | -0.65 | 0.82 | B>A |
| chrX | 1.21E+08 | 121100000 | -0.71 | 0.79 | B>A |
| chrX | 1.21E+08 | 121200000 | -0.78 | 0.75 | B>A |
| chrX | 1.21E+08 | 121300000 | -0.80 | 0.63 | B>A |
| chrX | 1.21E+08 | 121400000 | -0.76 | 0.71 | B>A |
| chrX | 1.21E+08 | 121500000 | -0.80 | 0.55 | B>A |
| chrX | 1.22E+08 | 121600000 | -0.70 | 0.00 | NO |
| chrX | 1.22E+08 | 121700000 | -0.79 | 0.00 | NO |
| chrX | 1.22E+08 | 121800000 | -0.76 | 0.00 | NO |
| chrX | 1.22E+08 | 121900000 | -0.70 | 0.00 | NO |
| chrX | 1.22E+08 | 122000000 | -0.56 | 0.00 | NO |
| chrX | 1.22E+08 | 122100000 | -0.44 | 0.00 | NO |
| chrX | 1.22E+08 | 122200000 | -0.28 | 0.00 | NO |
| chrX | 1.22E+08 | 122300000 | -0.09 | 0.00 | NO |
| chrX | 1.22E+08 | 122400000 | 0.07 | 0.48 | A>A |
| chrX | 1.22E+08 | 122500000 | 0.05 | 0.56 | B>A |
| chrX | 1.23E+08 | 122600000 | 0.13 | 0.55 | A>A |
| chrX | 1.23E+08 | 122700000 | 0.48 | 0.73 | A>A |
| chrX | 1.23E+08 | 122800000 | 0.94 | 1.11 | A>A |
| chrX | 1.23E+08 | 122900000 | 1.15 | 1.21 | A>A |
| chrX | 1.23E+08 | 123000000 | 1.14 | 1.22 | A>A |
| chrX | 1.23E+08 | 123100000 | 1.07 | 1.10 | A>A |
| chrX | 1.23E+08 | 123200000 | 1.05 | 1.06 | A>A |
| chrX | 1.23E+08 | 123300000 | 0.75 | 0.76 | A>A |

| | | | | | |
|---|---|---|---|---|---|
| chrX | 1.23E+08 | 123400000 | 0.31 | 0.22 | A>A |
| chrX | 1.23E+08 | 123500000 | 0.05 | -0.11 | B>B |
| chrX | 1.24E+08 | 123600000 | -0.14 | -0.37 | B>B |
| chrX | 1.24E+08 | 123700000 | -0.24 | -0.73 | B>B |
| chrX | 1.24E+08 | 123800000 | -0.36 | -0.84 | B>B |
| chrX | 1.24E+08 | 123900000 | -0.35 | -0.91 | B>B |
| chrX | 1.24E+08 | 124000000 | -0.45 | -0.81 | B>B |
| chrX | 1.24E+08 | 124100000 | -0.49 | -0.80 | B>B |
| chrX | 1.24E+08 | 124200000 | -0.45 | -0.77 | B>B |
| chrX | 1.24E+08 | 124300000 | -0.39 | -0.64 | B>B |
| chrX | 1.24E+08 | 124400000 | -0.26 | -0.31 | B>B |
| chrX | 1.24E+08 | 124500000 | -0.27 | -0.21 | B>B |
| chrX | 1.25E+08 | 124600000 | -0.47 | -0.56 | B>B |
| chrX | 1.25E+08 | 124700000 | -0.66 | -0.69 | B>B |
| chrX | 1.25E+08 | 124800000 | -0.75 | -0.72 | B>B |
| chrX | 1.25E+08 | 124900000 | -0.80 | -0.70 | B>B |
| chrX | 1.25E+08 | 125000000 | -0.71 | -0.74 | B>B |
| chrX | 1.25E+08 | 125100000 | -0.75 | -0.72 | B>B |
| chrX | 1.25E+08 | 125200000 | -0.77 | -0.62 | B>B |
| chrX | 1.25E+08 | 125300000 | -0.62 | -0.55 | B>B |
| chrX | 1.25E+08 | 125400000 | -0.71 | -0.42 | B>B |
| chrX | 1.25E+08 | 125500000 | -0.77 | -0.26 | B>B |
| chrX | 1.26E+08 | 125600000 | -0.70 | -0.06 | B>A |
| chrX | 1.26E+08 | 125700000 | -0.79 | 0.13 | B>A |
| chrX | 1.26E+08 | 125800000 | -0.75 | 0.09 | B>A |
| chrX | 1.26E+08 | 125900000 | -0.73 | -0.22 | B>B |
| chrX | 1.26E+08 | 126000000 | -0.74 | -0.57 | B>B |
| chrX | 1.26E+08 | 126100000 | -0.80 | -0.65 | B>B |
| chrX | 1.26E+08 | 126200000 | -0.74 | -0.73 | B>B |
| chrX | 1.26E+08 | 126300000 | -0.82 | -0.74 | B>B |
| chrX | 1.26E+08 | 126400000 | -0.71 | -0.75 | B>B |
| chrX | 1.26E+08 | 126500000 | -0.76 | -0.74 | B>B |
| chrX | 1.27E+08 | 126600000 | -0.74 | -0.77 | B>B |
| chrX | 1.27E+08 | 126700000 | -0.75 | -0.79 | B>B |
| chrX | 1.27E+08 | 126800000 | -0.72 | -0.85 | B>B |
| chrX | 1.27E+08 | 126900000 | -0.76 | -0.76 | B>B |
| chrX | 1.27E+08 | 127000000 | -0.75 | -0.80 | B>B |
| chrX | 1.27E+08 | 127100000 | -0.74 | -0.74 | B>B |
| chrX | 1.27E+08 | 127200000 | -0.79 | -0.76 | B>B |
| chrX | 1.27E+08 | 127300000 | -0.67 | -0.79 | B>B |
| chrX | 1.27E+08 | 127400000 | -0.76 | -0.76 | B>B |
| chrX | 1.27E+08 | 127500000 | -0.68 | -0.81 | B>B |
| chrX | 1.28E+08 | 127600000 | -0.65 | -0.73 | B>B |
| chrX | 1.28E+08 | 127700000 | -0.68 | -0.64 | B>B |
| chrX | 1.28E+08 | 127800000 | -0.60 | -0.62 | B>B |
| chrX | 1.28E+08 | 127900000 | -0.56 | -0.52 | B>B |
| chrX | 1.28E+08 | 128000000 | -0.58 | -0.59 | B>B |
| chrX | 1.28E+08 | 128100000 | -0.58 | -0.72 | B>B |
| chrX | 1.28E+08 | 128200000 | -0.43 | -0.63 | B>B |
| chrX | 1.28E+08 | 128300000 | -0.34 | -0.62 | B>B |
| chrX | 1.28E+08 | 128400000 | -0.29 | -0.59 | B>B |
| chrX | 1.28E+08 | 128500000 | -0.20 | -0.20 | B>B |
| chrX | 1.29E+08 | 128600000 | 0.35 | 0.20 | A>A |
| chrX | 1.29E+08 | 128700000 | 1.03 | 0.92 | A>A |
| chrX | 1.29E+08 | 128800000 | 1.03 | 1.04 | A>A |
| chrX | 1.29E+08 | 128900000 | 1.28 | 1.14 | A>A |

| | | | | | |
|---|---|---|---|---|---|
| chrX | 1.29E+08 | 129000000 | 1.13 | 1.04 | A>A |
| chrX | 1.29E+08 | 129100000 | 1.29 | 1.14 | A>A |
| chrX | 1.29E+08 | 129200000 | 1.26 | 1.11 | A>A |
| chrX | 1.29E+08 | 129300000 | 1.16 | 1.18 | A>A |
| chrX | 1.29E+08 | 129400000 | 1.10 | 1.08 | A>A |
| chrX | 1.29E+08 | 129500000 | 1.04 | 0.89 | A>A |
| chrX | 1.3E+08 | 129600000 | 0.85 | 0.68 | A>A |
| chrX | 1.3E+08 | 129700000 | 0.66 | 0.52 | A>A |
| chrX | 1.3E+08 | 129800000 | 0.78 | 0.68 | A>A |
| chrX | 1.3E+08 | 129900000 | 0.81 | 0.82 | A>A |
| chrX | 1.3E+08 | 130000000 | 0.80 | 0.79 | A>A |
| chrX | 1.3E+08 | 130100000 | 0.52 | 0.60 | A>A |
| chrX | 1.3E+08 | 130200000 | 0.29 | 0.32 | A>A |
| chrX | 1.3E+08 | 130300000 | 0.35 | 0.34 | A>A |
| chrX | 1.3E+08 | 130400000 | 0.25 | 0.36 | A>A |
| chrX | 1.3E+08 | 130500000 | 0.37 | 0.57 | A>A |
| chrX | 1.31E+08 | 130600000 | 0.40 | 0.70 | A>A |
| chrX | 1.31E+08 | 130700000 | 0.50 | 0.81 | A>A |
| chrX | 1.31E+08 | 130800000 | 0.31 | 0.52 | A>A |
| chrX | 1.31E+08 | 130900000 | 0.12 | 0.28 | A>A |
| chrX | 1.31E+08 | 131000000 | 0.12 | 0.06 | A>A |
| chrX | 1.31E+08 | 131100000 | 0.17 | -0.08 | A>A |
| chrX | 1.31E+08 | 131200000 | 0.31 | 0.12 | A>A |
| chrX | 1.31E+08 | 131300000 | 0.24 | 0.11 | A>A |
| chrX | 1.31E+08 | 131400000 | 0.24 | 0.05 | A>A |
| chrX | 1.31E+08 | 131500000 | -0.07 | -0.41 | B>B |
| chrX | 1.32E+08 | 131600000 | 0.06 | -0.46 | B>B |
| chrX | 1.32E+08 | 131700000 | 0.02 | -0.37 | B>B |
| chrX | 1.32E+08 | 131800000 | -0.02 | -0.42 | B>B |
| chrX | 1.32E+08 | 131900000 | -0.07 | -0.63 | B>B |
| chrX | 1.32E+08 | 132000000 | 0.04 | -0.78 | A>B |
| chrX | 1.32E+08 | 132100000 | 0.00 | -0.73 | A>B |
| chrX | 1.32E+08 | 132200000 | -0.11 | -0.69 | B>B |
| chrX | 1.32E+08 | 132300000 | -0.01 | -0.89 | A>B |
| chrX | 1.32E+08 | 132400000 | 0.00 | -0.72 | A>B |
| chrX | 1.32E+08 | 132500000 | 0.04 | -0.66 | A>B |
| chrX | 1.33E+08 | 132600000 | 0.28 | -0.44 | A>B |
| chrX | 1.33E+08 | 132700000 | 0.20 | -0.44 | A>B |
| chrX | 1.33E+08 | 132800000 | -0.01 | -0.34 | B>B |
| chrX | 1.33E+08 | 132900000 | 0.11 | -0.08 | A>A |
| chrX | 1.33E+08 | 133000000 | 0.10 | -0.19 | B>B |
| chrX | 1.33E+08 | 133100000 | 0.12 | -0.29 | B>B |
| chrX | 1.33E+08 | 133200000 | 0.07 | -0.35 | B>B |
| chrX | 1.33E+08 | 133300000 | 0.23 | -0.19 | A>A |
| chrX | 1.33E+08 | 133400000 | 0.42 | 0.10 | A>A |
| chrX | 1.33E+08 | 133500000 | 0.52 | 0.16 | A>A |
| chrX | 1.34E+08 | 133600000 | 0.89 | 0.76 | A>A |
| chrX | 1.34E+08 | 133700000 | 0.96 | 0.94 | A>A |
| chrX | 1.34E+08 | 133800000 | 1.47 | 1.22 | A>A |
| chrX | 1.34E+08 | 133900000 | 1.23 | 1.13 | A>A |
| chrX | 1.34E+08 | 134000000 | 1.09 | 0.98 | A>A |
| chrX | 1.34E+08 | 134100000 | 1.20 | 1.07 | A>A |
| chrX | 1.34E+08 | 134200000 | 1.00 | 1.03 | A>A |
| chrX | 1.34E+08 | 134300000 | 0.00 | 0.00 | NO |
| chrX | 1.34E+08 | 134400000 | 0.00 | 0.00 | NO |
| chrX | 1.34E+08 | 134500000 | 0.45 | 0.71 | A>A |

| | | | | | |
|---|---|---|---|---|---|
| chrX | 1.35E+08 | 134600000 | 0.52 | 0.80 | A>A |
| chrX | 1.35E+08 | 134700000 | 0.60 | 0.55 | A>A |
| chrX | 1.35E+08 | 134800000 | 0.36 | 0.43 | A>A |
| chrX | 1.35E+08 | 134900000 | 0.00 | 0.00 | NO |
| chrX | 1.35E+08 | 135000000 | 0.00 | 0.00 | NO |
| chrX | 1.35E+08 | 135100000 | 0.75 | 0.82 | A>A |
| chrX | 1.35E+08 | 135200000 | 0.84 | 1.03 | A>A |
| chrX | 1.35E+08 | 135300000 | 1.04 | 1.06 | A>A |
| chrX | 1.35E+08 | 135400000 | 0.74 | 0.61 | A>A |
| chrX | 1.35E+08 | 135500000 | 0.47 | 0.28 | A>A |
| chrX | 1.36E+08 | 135600000 | 0.60 | 0.45 | A>A |
| chrX | 1.36E+08 | 135700000 | 0.51 | 0.35 | A>A |
| chrX | 1.36E+08 | 135800000 | 0.42 | 0.31 | A>A |
| chrX | 1.36E+08 | 135900000 | 0.82 | 0.46 | A>A |
| chrX | 1.36E+08 | 136000000 | 1.05 | 0.79 | A>A |
| chrX | 1.36E+08 | 136100000 | 0.50 | 0.24 | A>A |
| chrX | 1.36E+08 | 136200000 | 0.27 | -0.21 | A>A |
| chrX | 1.36E+08 | 136300000 | -0.11 | -0.64 | B>B |
| chrX | 1.36E+08 | 136400000 | -0.19 | -0.84 | A>B |
| chrX | 1.36E+08 | 136500000 | -0.27 | -0.81 | B>B |
| chrX | 1.37E+08 | 136600000 | -0.21 | -0.80 | A>B |
| chrX | 1.37E+08 | 136700000 | -0.18 | -0.82 | A>B |
| chrX | 1.37E+08 | 136800000 | -0.36 | -0.87 | B>B |
| chrX | 1.37E+08 | 136900000 | -0.57 | -0.90 | B>B |
| chrX | 1.37E+08 | 137000000 | -0.72 | -0.95 | B>B |
| chrX | 1.37E+08 | 137100000 | -0.74 | -0.89 | B>B |
| chrX | 1.37E+08 | 137200000 | -0.70 | -0.88 | B>B |
| chrX | 1.37E+08 | 137300000 | -0.77 | -0.86 | B>B |
| chrX | 1.37E+08 | 137400000 | -0.88 | -0.89 | B>B |
| chrX | 1.37E+08 | 137500000 | -0.93 | -0.94 | B>B |
| chrX | 1.38E+08 | 137600000 | -1.07 | -0.99 | B>B |
| chrX | 1.38E+08 | 137700000 | -0.99 | -0.98 | B>B |
| chrX | 1.38E+08 | 137800000 | -0.73 | -0.87 | B>B |
| chrX | 1.38E+08 | 137900000 | -0.63 | -0.88 | B>B |
| chrX | 1.38E+08 | 138000000 | -0.75 | -0.89 | B>B |
| chrX | 1.38E+08 | 138100000 | -0.82 | -0.88 | B>B |
| chrX | 1.38E+08 | 138200000 | -0.75 | -0.88 | B>B |
| chrX | 1.38E+08 | 138300000 | -0.48 | -0.77 | B>B |
| chrX | 1.38E+08 | 138400000 | -0.68 | -0.82 | B>B |
| chrX | 1.38E+08 | 138500000 | -0.71 | -0.81 | B>B |
| chrX | 1.39E+08 | 138600000 | -0.67 | -0.75 | B>B |
| chrX | 1.39E+08 | 138700000 | -0.37 | -0.75 | B>B |
| chrX | 1.39E+08 | 138800000 | -0.11 | -0.52 | B>B |
| chrX | 1.39E+08 | 138900000 | 0.37 | 0.01 | A>A |
| chrX | 1.39E+08 | 139000000 | 0.65 | 0.48 | A>A |
| chrX | 1.39E+08 | 139100000 | 0.21 | -0.15 | A>A |
| chrX | 1.39E+08 | 139200000 | -0.09 | -0.51 | B>B |
| chrX | 1.39E+08 | 139300000 | -0.32 | -0.80 | B>B |
| chrX | 1.39E+08 | 139400000 | -0.68 | -0.98 | B>B |
| chrX | 1.39E+08 | 139500000 | -0.57 | -1.02 | B>B |
| chrX | 1.4E+08 | 139600000 | -0.32 | -0.85 | B>B |
| chrX | 1.4E+08 | 139700000 | -0.30 | -0.79 | B>B |
| chrX | 1.4E+08 | 139800000 | -0.06 | -0.61 | B>B |
| chrX | 1.4E+08 | 139900000 | 0.02 | -0.18 | B>B |
| chrX | 1.4E+08 | 140000000 | 0.58 | 0.33 | A>A |
| chrX | 1.4E+08 | 140100000 | 0.89 | 0.66 | A>A |

| | | | | | |
|---|---|---|---|---|---|
| chrX | 1.4E+08 | 140200000 | 0.00 | 0.00 | NO |
| chrX | 1.4E+08 | 140300000 | 0.83 | 0.59 | A>A |
| chrX | 1.4E+08 | 140400000 | 0.00 | 0.62 | NO |
| chrX | 1.4E+08 | 140500000 | 0.82 | 0.61 | A>A |
| chrX | 1.41E+08 | 140600000 | 0.80 | 0.80 | A>A |
| chrX | 1.41E+08 | 140700000 | 0.00 | 0.00 | NO |
| chrX | 1.41E+08 | 140800000 | 0.61 | 0.51 | A>A |
| chrX | 1.41E+08 | 140900000 | 0.70 | 0.54 | A>A |
| chrX | 1.41E+08 | 141000000 | 0.58 | 0.50 | A>A |
| chrX | 1.41E+08 | 141100000 | 0.52 | 0.28 | A>A |
| chrX | 1.41E+08 | 141200000 | 0.33 | 0.13 | A>A |
| chrX | 1.41E+08 | 141300000 | 0.10 | -0.26 | B>B |
| chrX | 1.41E+08 | 141400000 | -0.06 | -0.37 | B>B |
| chrX | 1.41E+08 | 141500000 | -0.19 | -0.50 | B>B |
| chrX | 1.42E+08 | 141600000 | -0.35 | -0.60 | B>B |
| chrX | 1.42E+08 | 141700000 | -0.40 | -0.60 | B>B |
| chrX | 1.42E+08 | 141800000 | -0.48 | -0.66 | B>B |
| chrX | 1.42E+08 | 141900000 | -0.61 | -0.71 | B>B |
| chrX | 1.42E+08 | 142000000 | -0.62 | -0.79 | B>B |
| chrX | 1.42E+08 | 142100000 | -0.71 | -0.81 | B>B |
| chrX | 1.42E+08 | 142200000 | -0.76 | -0.74 | B>B |
| chrX | 1.42E+08 | 142300000 | -0.74 | -0.81 | B>B |
| chrX | 1.42E+08 | 142400000 | -0.79 | -0.79 | B>B |
| chrX | 1.42E+08 | 142500000 | -0.74 | -0.83 | B>B |
| chrX | 1.43E+08 | 142600000 | -0.81 | -0.82 | B>B |
| chrX | 1.43E+08 | 142700000 | -0.76 | -0.75 | B>B |
| chrX | 1.43E+08 | 142800000 | -0.73 | -0.63 | B>B |
| chrX | 1.43E+08 | 142900000 | -0.86 | -0.57 | B>B |
| chrX | 1.43E+08 | 143000000 | -0.86 | -0.48 | B>B |
| chrX | 1.43E+08 | 143100000 | -0.88 | -0.43 | B>B |
| chrX | 1.43E+08 | 143200000 | -0.90 | -0.55 | B>B |
| chrX | 1.43E+08 | 143300000 | 0.00 | 0.00 | NO |
| chrX | 1.43E+08 | 143400000 | -0.91 | -0.74 | B>B |
| chrX | 1.43E+08 | 143500000 | -0.93 | -0.84 | B>B |
| chrX | 1.44E+08 | 143600000 | 0.00 | -0.83 | NO |
| chrX | 1.44E+08 | 143700000 | -0.89 | -0.90 | B>B |
| chrX | 1.44E+08 | 143800000 | -0.91 | -0.89 | B>B |
| chrX | 1.44E+08 | 143900000 | -1.07 | -0.89 | B>B |
| chrX | 1.44E+08 | 144000000 | -0.98 | -0.88 | B>B |
| chrX | 1.44E+08 | 144100000 | -1.02 | -0.99 | B>B |
| chrX | 1.44E+08 | 144200000 | -0.93 | -0.91 | B>B |
| chrX | 1.44E+08 | 144300000 | -0.83 | -0.82 | B>B |
| chrX | 1.44E+08 | 144400000 | -1.01 | -0.85 | B>B |
| chrX | 1.44E+08 | 144500000 | -1.06 | -0.92 | B>B |
| chrX | 1.45E+08 | 144600000 | -1.02 | -0.95 | B>B |
| chrX | 1.45E+08 | 144700000 | -1.10 | -0.98 | B>B |
| chrX | 1.45E+08 | 144800000 | -1.15 | -1.02 | B>B |
| chrX | 1.45E+08 | 144900000 | -0.91 | -0.97 | B>B |
| chrX | 1.45E+08 | 145000000 | -0.96 | -0.96 | B>B |
| chrX | 1.45E+08 | 145100000 | -1.05 | -1.01 | B>B |
| chrX | 1.45E+08 | 145200000 | -0.93 | -0.91 | B>B |
| chrX | 1.45E+08 | 145300000 | -1.01 | -1.02 | B>B |
| chrX | 1.45E+08 | 145400000 | -1.10 | -1.06 | B>B |
| chrX | 1.45E+08 | 145500000 | -1.00 | -1.03 | B>B |
| chrX | 1.46E+08 | 145600000 | -1.07 | -1.06 | B>B |
| chrX | 1.46E+08 | 145700000 | -1.12 | -1.09 | B>B |

| | | | | | |
|---|---|---|---|---|---|
| chrX | 1.46E+08 | 145800000 | -1.11 | -1.05 | B>B |
| chrX | 1.46E+08 | 145900000 | -1.08 | -1.04 | B>B |
| chrX | 1.46E+08 | 146000000 | -1.11 | -1.11 | B>B |
| chrX | 1.46E+08 | 146100000 | -1.14 | -1.06 | B>B |
| chrX | 1.46E+08 | 146200000 | -1.11 | -1.08 | B>B |
| chrX | 1.46E+08 | 146300000 | -1.02 | -1.10 | B>B |
| chrX | 1.46E+08 | 146400000 | -1.01 | -1.02 | B>B |
| chrX | 1.46E+08 | 146500000 | -1.15 | -1.06 | B>B |
| chrX | 1.47E+08 | 146600000 | -1.07 | -1.07 | B>B |
| chrX | 1.47E+08 | 146700000 | -1.01 | -0.94 | B>B |
| chrX | 1.47E+08 | 146800000 | -1.00 | -0.99 | B>B |
| chrX | 1.47E+08 | 146900000 | -0.86 | -0.92 | B>B |
| chrX | 1.47E+08 | 147000000 | -0.33 | -0.49 | B>B |
| chrX | 1.47E+08 | 147100000 | 0.00 | -0.27 | B>B |
| chrX | 1.47E+08 | 147200000 | -0.55 | -0.74 | B>B |
| chrX | 1.47E+08 | 147300000 | -0.86 | -0.94 | B>B |
| chrX | 1.47E+08 | 147400000 | -0.95 | -1.06 | B>B |
| chrX | 1.47E+08 | 147500000 | -0.88 | -0.93 | B>B |
| chrX | 1.48E+08 | 147600000 | -0.74 | -0.88 | B>B |
| chrX | 1.48E+08 | 147700000 | -0.83 | -0.97 | B>B |
| chrX | 1.48E+08 | 147800000 | -0.96 | -0.91 | B>B |
| chrX | 1.48E+08 | 147900000 | -0.99 | -1.02 | B>B |
| chrX | 1.48E+08 | 148000000 | -0.80 | -0.98 | B>B |
| chrX | 1.48E+08 | 148100000 | -0.81 | -1.01 | B>B |
| chrX | 1.48E+08 | 148200000 | -0.87 | -0.80 | B>B |
| chrX | 1.48E+08 | 148300000 | -0.86 | -0.89 | B>B |
| chrX | 1.48E+08 | 148400000 | -0.50 | -0.79 | B>B |
| chrX | 1.48E+08 | 148500000 | -0.36 | -0.87 | B>B |
| chrX | 1.49E+08 | 148600000 | 0.27 | -0.08 | A>A |
| chrX | 1.49E+08 | 148700000 | 0.00 | 0.00 | NO |
| chrX | 1.49E+08 | 148800000 | 1.05 | 0.29 | A>B |
| chrX | 1.49E+08 | 148900000 | 0.00 | 0.00 | NO |
| chrX | 1.49E+08 | 149000000 | 0.00 | 0.00 | NO |
| chrX | 1.49E+08 | 149100000 | 0.00 | 0.00 | NO |
| chrX | 1.49E+08 | 149200000 | 1.04 | 1.06 | A>A |
| chrX | 1.49E+08 | 149300000 | 1.29 | 1.33 | A>A |
| chrX | 1.49E+08 | 149400000 | 1.13 | 1.23 | A>A |
| chrX | 1.49E+08 | 149500000 | 1.22 | 1.30 | A>A |
| chrX | 1.5E+08 | 149600000 | 1.21 | 1.19 | A>A |
| chrX | 1.5E+08 | 149700000 | 1.13 | 1.12 | A>A |
| chrX | 1.5E+08 | 149800000 | 1.18 | 1.12 | A>A |
| chrX | 1.5E+08 | 149900000 | 1.02 | 0.97 | A>A |
| chrX | 1.5E+08 | 150000000 | 1.07 | 1.05 | A>A |
| chrX | 1.5E+08 | 150100000 | 1.42 | 1.50 | A>A |
| chrX | 1.5E+08 | 150200000 | 1.57 | 1.13 | A>A |
| chrX | 1.5E+08 | 150300000 | 0.94 | 0.55 | A>A |
| chrX | 1.5E+08 | 150400000 | 0.30 | 0.00 | A>A |
| chrX | 1.5E+08 | 150500000 | -0.04 | -0.35 | B>B |
| chrX | 1.51E+08 | 150600000 | 0.40 | 0.18 | A>A |
| chrX | 1.51E+08 | 150700000 | 0.29 | -0.27 | A>A |
| chrX | 1.51E+08 | 150800000 | 0.20 | -0.27 | B>B |
| chrX | 1.51E+08 | 150900000 | 0.15 | -0.17 | B>B |
| chrX | 1.51E+08 | 151000000 | 0.28 | 0.20 | A>A |
| chrX | 1.51E+08 | 151100000 | 0.56 | 0.61 | A>A |
| chrX | 1.51E+08 | 151200000 | 0.69 | 0.75 | A>A |
| chrX | 1.51E+08 | 151300000 | 0.50 | 0.65 | A>A |

| | | | | | |
|---|---|---|---|---|---|
| chrX | 1.51E+08 | 151400000 | 0.13 | 0.40 | A>A |
| chrX | 1.51E+08 | 151500000 | -0.19 | 0.19 | A>A |
| chrX | 1.52E+08 | 151600000 | -0.22 | -0.01 | B>B |
| chrX | 1.52E+08 | 151700000 | -0.28 | -0.05 | B>B |
| chrX | 1.52E+08 | 151800000 | -0.09 | -0.01 | B>B |
| chrX | 1.52E+08 | 151900000 | 0.00 | 0.23 | NO |
| chrX | 1.52E+08 | 152000000 | 0.00 | 0.32 | NO |
| chrX | 1.52E+08 | 152100000 | 1.40 | 1.05 | A>A |
| chrX | 1.52E+08 | 152200000 | 1.44 | 1.06 | A>A |
| chrX | 1.52E+08 | 152300000 | 0.00 | 0.00 | NO |
| chrX | 1.52E+08 | 152400000 | 0.00 | 0.00 | NO |
| chrX | 1.52E+08 | 152500000 | 0.00 | 0.80 | NO |
| chrX | 1.53E+08 | 152600000 | 0.00 | 0.76 | NO |
| chrX | 1.53E+08 | 152700000 | 1.53 | 1.16 | A>A |
| chrX | 1.53E+08 | 152800000 | 1.63 | 1.39 | A>A |
| chrX | 1.53E+08 | 152900000 | 1.82 | 1.49 | A>A |
| chrX | 1.53E+08 | 153000000 | 1.72 | 1.59 | A>A |
| chrX | 1.53E+08 | 153100000 | 1.61 | 1.50 | A>A |
| chrX | 1.53E+08 | 153200000 | 1.72 | 1.67 | A>A |
| chrX | 1.53E+08 | 153300000 | 1.69 | 1.46 | A>A |
| chrX | 1.53E+08 | 153400000 | 1.67 | 1.59 | A>A |
| chrX | 1.53E+08 | 153500000 | 0.00 | 0.00 | NO |
| chrX | 1.54E+08 | 153600000 | 1.60 | 1.43 | A>A |
| chrX | 1.54E+08 | 153700000 | 2.02 | 1.68 | A>A |
| chrX | 1.54E+08 | 153800000 | 1.74 | 1.44 | A>A |
| chrX | 1.54E+08 | 153900000 | 0.00 | 0.00 | NO |
| chrX | 1.54E+08 | 154000000 | 1.83 | 1.34 | A>A |
| chrX | 1.54E+08 | 154100000 | 1.95 | 1.61 | A>A |
| chrX | 1.54E+08 | 154200000 | 1.43 | 1.29 | A>A |
| chrX | 1.54E+08 | 154300000 | 1.84 | 1.52 | A>A |
| chrX | 1.54E+08 | 154400000 | 1.45 | 1.28 | A>A |
| chrX | 1.54E+08 | 154500000 | 1.00 | 1.20 | A>A |
| chrX | 1.55E+08 | 154600000 | 0.00 | 0.00 | NO |
| chrX | 1.55E+08 | 154700000 | 0.00 | 0.00 | NO |
| chrX | 1.55E+08 | 154800000 | 0.00 | 0.00 | NO |
| chrX | 1.55E+08 | 154900000 | 0.56 | 1.17 | B>A |
| chrX | 1.55E+08 | 155000000 | 0.00 | 0.00 | NO |
| chrX | 1.55E+08 | 155100000 | 0.00 | 0.00 | NO |
| chrX | 1.55E+08 | 155200000 | 0.00 | 0.00 | NO |
| chrX | 1.55E+08 | 155270560 | 0.00 | 0.00 | NO |
